# Phylogenetic Tree-based Pipeline for Uncovering Mutational Patterns during Influenza Virus Evolution

**DOI:** 10.1101/708420

**Authors:** Fransiskus Xaverius Ivan, Akhila Deshpande, Chun Wei Lim, Xinrui Zhou, Jie Zheng, Chee Keong Kwoh

## Abstract

Various computational and statistical approaches have been proposed to uncover the mutational patterns of rapidly evolving influenza viral genes. Nonetheless, the approaches mainly rely on sequence alignments which could potentially lead to spurious mutations obtained by comparing sequences from different clades that coexist during particular periods of time. To address this issue, we propose a phylogenetic tree-based pipeline that takes into account the evolutionary structure in the sequence data. Assuming that the sequences evolve progressively under a strict molecular clock, considering a competitive model that is based on a certain Markov model, and using a resampling approach to obtain robust estimates, we could capture statistically significant single-mutations and co-mutations during the sequence evolution. Moreover, by considering the results obtained from analyses that consider all paths and the longest path in the resampled trees, we can categorize the mutational sites and suggest their relevance. Here we applied the pipeline to investigate the 50 years of evolution of the HA sequences of influenza A/H3N2 viruses. In addition to confirming previous knowledge on the A/H3N2 HA evolution, we also demonstrate the use of the pipeline to classify mutational sites according to whether they are able to enhance antigenic drift, compensate other mutations that enhance antigenic drift, or both.

## Introduction

Seasonal influenza viruses, especially the influenza A viruses, have exhibited frequent mutations with a rapid evolutionary rate. The hemagglutinin (HA) of influenza has the highest mutational rate among all influenza viral proteins [1]. Besides, the HA is considered as a major culprit for the antigenicity of influenza and a primary target for the influenza vaccine [2, 3]. Cumulative mutations can lead to the antigenic drift of influenza, enable the viruses to mismatch the influenza vaccine, escape the human immune system, and even raise an epidemic [4]. Therefore, it is crucial to surveil and predict the mutations of influenza. The knowledge of mutational patterns can improve our understanding about the mechanism of antigenic drift.

Discovering the dependencies among mutations is a non-trivial and active area of bioinformatics. Non-independent mutations of amino acids may co-occur, or occur chronologically, generally sharing a common constraint or protein function domain [5]. The directed mutagenesis experiments are a classical type of method to identify functional dependencies between amino acid sites [6]. However, the complexity of possible the experiments limits the capacity of research. Subsequently, various statistical and computational models have been proposed as complementary tools to evaluate the correlation between amino acid sites [7], annotate protein functional domains [8], reveal possible amino acid interactions, and predict the interactions between motifs or proteins [9, 10].

As to the influenza viruses, many computational methods detecting the antigenic mutations have been proposed. For example, Smith *et al*. pioneered the mapping of antigenic evolution and genetic evolution, revealing that the influenza viruses undergo continuous genetic evolution pressure, while the antigenic evolution is more punctuated with 11 antigenic clusters of influenza A/H3N2 being detected. The comparison between genetic and antigenic evolution indicated that some mutations bear a disproportionately large effect on the antigenicity of influenza [11]. Shih *et al*. analyzed the frequency changes of all HA1 amino acids, showing that the positive selection on HA1 is ongoing most of the time. However, the antigenic drift of influenza is punctuated which can be changed by a single substitution at antigenic sites of HA1, or in most cases, by simultaneous multiple fixations [12]. Koel *et al*. extended the works by investigating the antigenic clusters and all observed substitutions. It was found that seven cluster-transition substitutions were responsible for the antigenic cluster transitions, all of which located at or around the receptor-binding sites of HA [13]. Recently, Quan *et al*. developed a computational model RECDS (recognition of cluster-transition determining sites) using a gradient boosting classifier to rank the importance of all HA sites, and evaluate the contribution of an HA amino acid site to the antigenic evolutionary history of influenza viruses [14]. The RECDS is a feature-based (both sequence-based features and structure-based features) computational model under the assumption that features dominating antigenicity are highly conserved. Statistical models on positive selection sites are mainly based on the ratio of nonsynonymous to synonymous mutations (dN/dS ratio) [15]. Tusche *et al*. integrated the dN/dS as a measure of selection, the ancestral information inferred from phylogenetic trees, and spatial proximity of sites to identify regions under selective pressure [16].

However, these methods do not pay attention to the substitution dependency on the HA. Information theory based strategies are the most extensively used to measure the covariance between mutations [17]. For example, Baker et al. developed a web-based tool CoeViz for calculating and visualizing covariance metrics (mutual information, chi-square statistics, Pearson correlation, and joint Shannon entropy) [18]. Xia et al. constructed a site transition network based on the pairwise mutual information between amino acids of the HA sequences [19]. The network incorporating correlation information between residues improved the prediction of site mutations with an accuracy of 70%. Besides, Elma *et al*. considered the information of HA evolution. A mass-based protein phylogenetic model was proposed to identify functional comutations [20]. Alternatively, machine learning approaches are also applied to detect comutation patterns. For example, Chen et al. applied association rule mining to explore co-occurring mutations on H3 [21]. Du et al. proposed a feature-based Naïve Bayesian network to predict antigenic clusters [22].

However, those methods mainly depend on the protein sequences, lacking the chronological and 3D structural information. In this study, we proposed a pipeline for uncovering not only single-mutations under positive selection pressure, but also co-mutations of influenza viral protein sequences. Besides, we analyzed the co-mutations of hemagglutinin sequences of human influenza A/H3N2 to evaluate the effectiveness and robustness of the proposed pipeline. The detected mutation sites are highly overlapped with those reported to be under positive selection pressure, especially interfaces exposed to the antigenic binding. The proposed pipeline is promising to be applied to analyzing the molecular evolution of all influenza proteins.

## Methods

The flowchart of the proposed pipeline for uncovering significant single-mutations and comutations in particular influenza protein sequences is presented in **Fig. 1**. The overall pipeline composed of five major procedures, i.e., (*i*) Sub-pipeline 1 that retrieved, clustered and aligned a subset of sequence data from local influenza genome datasets, (*ii*) Sub-pipeline 2 that identified and removed outliers detected following the linear regression of root-to-tip distances in inferred neighbor joining (NJ) tree against isolation dates, (*iii*) Sub-pipeline 3 that extracted substitution model parameters from a maximum likelihood (ML) tree reconstructed from aligned sequences and used them to simulate sequence evolution, (*iv*) Sub-pipeline 4 that reconstructed resampled trees from aligned sequence data (either real or simulated one), and (*v*) Sub-pipeline 5 that calculated supports for single-mutations and co-mutations detected in the resampled trees. The evolutionary parameters, i.e., the rate of substitution and the date of origin, were required for co-mutation detection and remaining analyses (interpretation), and could be robustly estimated from the root-to-tip regressions of the resampled trees. At the final stage, the distributions of supports for the single-mutations/co-mutations from simulated sequence data were used to set a threshold for claiming significant single-mutations/co-mutations from the real sequence data. The details of the local influenza genome datasets and steps in each major procedure in the pipeline are described shortly, while the use of the pipeline for analyzing the evolutionary patterns of the hemagglutinin (HA) sequences of human influenza A/H3N2 viruses are presented in the Results and Discussions.

**Fig. 1.**
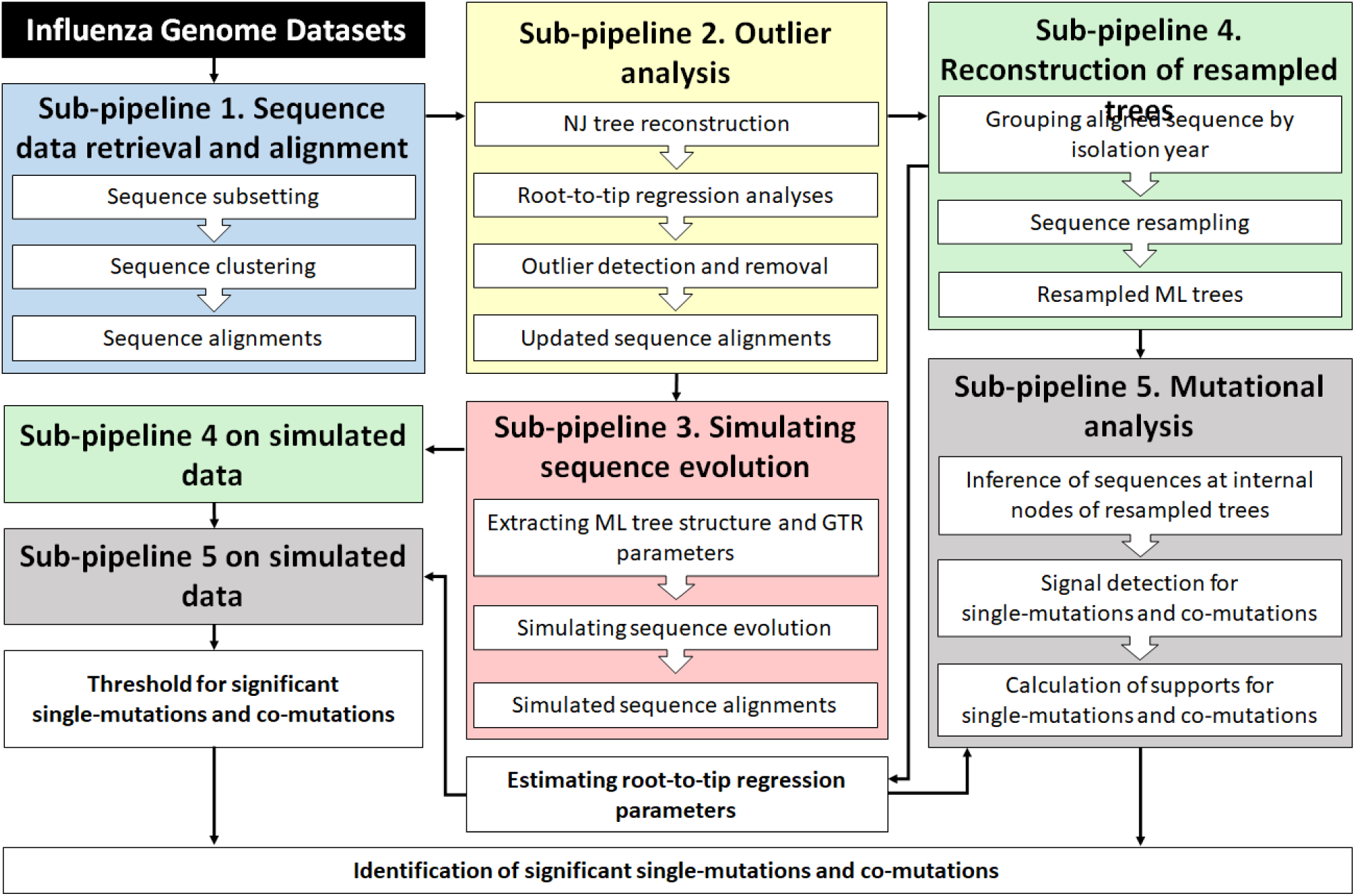
Phylogenetic tree-based pipelines for uncovering significant single-mutations and co-mutations in evolving influenza viral proteins.

### Influenza genome datasets

Local datasets consisting of influenza virus genomes, transcriptomes, proteomes, and their metadata (including the virus types, virus subtypes, virus names, and date of isolation) were created for this work. The records were retrieved from NCBI Influenza Virus Resource [23] or GISAID database [24]. Only records of influenza viruses whose genome was complete, associated coding/protein sequences could be identified and were not too short, and information of the host, location, and date of collection (for the date, if only the day was missing, then it was set to the 15th of the month; if the day and the month were missing, then it was set to 30 June of the year) was available, were included in the datasets. The records were cleaned and reformatted into one tab-delimited text file of metadata and eight tab-delimited text files of sequence data that correspond to each of the eight segments of influenza virus genome.

### Sub-pipeline 1 – Sequence data retrieval and alignment

This pipeline was used to subset the full nucleotide sequences, coding sequences and protein sequences of a particular gene of a specific influenza A subtype or influenza B lineage from the local datasets described previously, and each of the dataset was stored into a fasta file. The description of each fasta sequence record included the sequence ID, gene name, genome ID, name of the corresponding influenza virus strain, country and the date of virus isolation. The subsetted, non-redundant coding sequences were then aligned to codon position. For fast alignment and considering the sequences were highly similar, the protein sequences were first clustered using the CD-HIT tool [25] to obtain clusters of sequences whose percent identity to a representative sequence was above a certain threshold (we used athreshold of 98%). Clusters containing protein sequences of different length were split according to their length. Subsequently, the representatives of CD-HIT clusters were aligned with the muscle package [26] and the protein alignment was then used to guide the alignment of the corresponding coding sequences to codon position. The alignment of each of the rest of the coding sequences to the alignment of the representatives was done according to the alignment of its corresponding representative. The results of the alignment were visualized with MEGA7 software [27] for inspection.

### Sub-pipeline 2 – Outlier analysis

For outlier detection, we assumed that the sequence evolution follows a strict molecular clock, i.e., all branches in the phylogenetic tree evolve at the same rate. To evaluate this assumption, the genetic distances based on Jukes-Cantor (JC) substitution model [28] were calculated from the aligned coding sequences and used to construct an NJ tree [29]. Assuming the sequences evolve progressively, the phylogenetic tree was rooted using one of the earliest coding sequence as an outgroup. The root-to-tip regression analysis was then used to explore the association between genetic distances of the samples from the tree root and sampling dates. Denoting these two variables as *d*_*r,i*_ and *t*_*i*_, respectively, where *r* represents the tree root and *i* represents the samples or tree tips, the regression model can be written as: *E*[*d*_*r,i*_] = *μ*(*t*_*i*_ − *t*_*r*_). The gradient (*μ*) and *x*-intercept (*t*_*r*_) provide estimates for the substitution rate and the time of the tree root (the date of origin), respectively. Given the nature of the sequence data that is heterochronous (collected at different time points), a strong linear correlation between *d*_*r,i*_ and *t*_*i*_ suggests a high level of strict clock-like signals. Due to the non-independency of the individual data points, the root-to-tip linear regression is not appropriate for statistical hypothesis [30]. Nonetheless, the regression approach is reasonably used for identifying outliers. Here we identified a data point as an outlier if the absolute value of its residual from the regression line was larger than five times interquartile range.

### Sub-pipeline 3 – Simulating sequence evolution

After outliers were removed, a new phylogenetic tree was reconstructed using a more complex substitution model and algorithm. In particular, we reconstructed an ML tree using *GTR* + *G* + *I* substitution model implemented in phangorn package [31]. The *GTR* substitution model [32] is a type of continuous-time Markov model that is most general neutral, independent, finite-sites and time-reversible model. Its parameters consist of four equilibrium base frequency parameters (*π*_*A*_: the frequency of base A, *π*_*G*_: the frequency of base G, *π*_*C*_: the frequency of base C, and *π*_*T*_: the frequency of base T) and six substitution rate parameters (*α*: the substitution rate parameter for A → G and G → A, *β*: the substitution rate parameter for A → C and C → A, *γ*: the substitution rate parameter for A → T and T → A, *δ*: the substitution rate parameter for G → C and C → G, *ε*: the substitution rate parameter for G → T and T → G, and *η*: the substitution rate parameter for C → T and T → C). These parameters form the equilibrium base frequency vector Π = (*π*_*A*_, *π*_*G*_, *π*_*C*_, *π*_*T*_) and the rate matrix

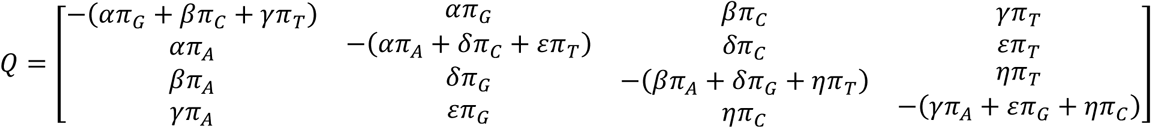

for the continuous-time Markov model. When considering the *GTR* + *G* + *I* model, a discrete Gamma distribution (+*G*) is used to take into account the rate heterogeneity among sites and a fixed fraction of sites is assumed to be evolutionary invariable (+*I*). These add two parameters for the Gamma distribution, i.e., the number of rate categories and the shape parameters, and another parameter for the proportion of invariant sites into the model. The estimated *GTR* + *G* + *I* substitution model parameters, the structure of the ML tree (that included the length of their branches) and the sequence at the tree root (inferred using Fitch algorithm [33]), were then used to simulate sequence evolution with Pyvolve [34]. Note that Pyvolve does not model insertion/deletion, hence any gap in the root sequence were removed. Sequences produced at the tip of the tree as the results of simulation were used to create a new sequence dataset that is referred to as simulated sequence dataset.

### Sub-pipeline 4 – Reconstruction of resampled trees

To reconstruct resampled phylogenetic trees from real or simulated sequence dataset, the aligned sequences were first grouped according to their sampling year. Before grouping, one of the earliest sequence was first singled out and it will always be included for sampled tree reconstruction. The grouping was done year by year, i.e., starting from the earliest year to the latest year, and the earlier sequences were grouped into a single year group if the total number of the sequences was more than a certain threshold (here we used a threshold of 20). After sequence grouping, we repeatedly and randomly sampled a fixed number of aligned sequences from each year group and added the earliest sequence to the sample. An ML phylogenetic tree was then reconstructed for each sample using a *GTR* + *G* + *I* substitution model implemented in phangorn package. The resampled ML trees were rooted using the earliest sequence as an outgroup and then used to calculate bootstrap estimates for the substitution rate and the date of sequence origin, i.e., by averaging the estimates obtained from each tree using the root-to-tip regression approach.

### Sub-pipeline 5 – Mutational analysis with resampled trees

Mutational analysis was done using resampled phylogenetic trees from each of the real and simulated sequence data. Each edge length or distance between two adjacent nodes in the trees was associated with the evolutionary distance, i.e., the number of nucleotide substitutions per site estimated based on the chosen substitution model. The distance between any two nodes (of interest, between ancestor and predecessor) in the tree was calculated by summing the length of edges in the path connecting the two nodes. The coding and protein sequences at each internal node of each tree were inferred using the Fitch’s algorithm [33]. Amino acid mutations were detected at each node (except for the root) by comparing its protein sequence to its parent’s protein sequence. Each amino acid mutation was in the form AA1-*p*-AA2, representing a mutation of a given amino acid AA1 in the parent node to another amino acid AA2 in the child node at a given site *p* in the sequence. Finally, the distance of each node to the root of the tree was also recorded.

The resampled phylogenetic trees were used to calculate support values that indicate the signal strength of single-mutational and co-mutational events during sequence evolution. To calculate supports for single-mutations, each of single-mutations observed in the trees was mapped to a list of real numbers representing the distances of the nodes where the mutation observed to their corresponding root. Then, for each mutation, we smoothed the distribution of its distance data with a Gaussian kernel density estimate [35], followed by the detection of the peaks that were defined as local maxima centered in any interval for the distance. Assuming *h*_1_, *h*_2_, …, *h*_*k*_ are the heights of the detected peaks for mutation *m* at distance to the root *d*_1_, *d*_2_, …, *d*_*k*_, then the strength of the signal for *m* at distance *d*_*i*_, denoted by *S*(*m, d*_*i*_), was calculated as follows: 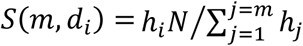, where *N* is the number of observations for the mutation of interest. The formula is indicative of the portion of observations that support the observed mutation. In addition to calculating supports for single-mutations found in any node in the resampled trees, we also calculated supports for single-mutations that were only observed in the longest paths of the resampled trees. The version of mutational analysis that considers any path in the resampled trees is termed as all path analysis, while the one that considers the longest path is termed as the longest path analysis.

The supports for co-mutations were calculated in similar way. Here, we considered co-mutations as any possible pair of single-mutations (the order of the single-mutations does not matter) observed at a single node or from two different nodes that had ancestor-predecessor relationship and distance below a certain threshold (which ought to be influenced by the estimated substitution rate). Each co-mutation was mapped to the distance of the ancestral node to the root of its resampled tree. Note that if the co-mutation was observed at a single node, then the node was considered as both ancestor and predecessor associated with the co-mutation. Algorithmically, the co-mutation list and the map can be created while walking from an initial node (any node other than the root), in the direction to the associated root, up to the node whose distance to the initial node is below a certain threshold. The rest of the procedure is as described previously for calculating the supports for single mutations. The complete procedure for calculating supports for co-mutations is formalized in **Algorithm 1**.

**Algorithm 1.**
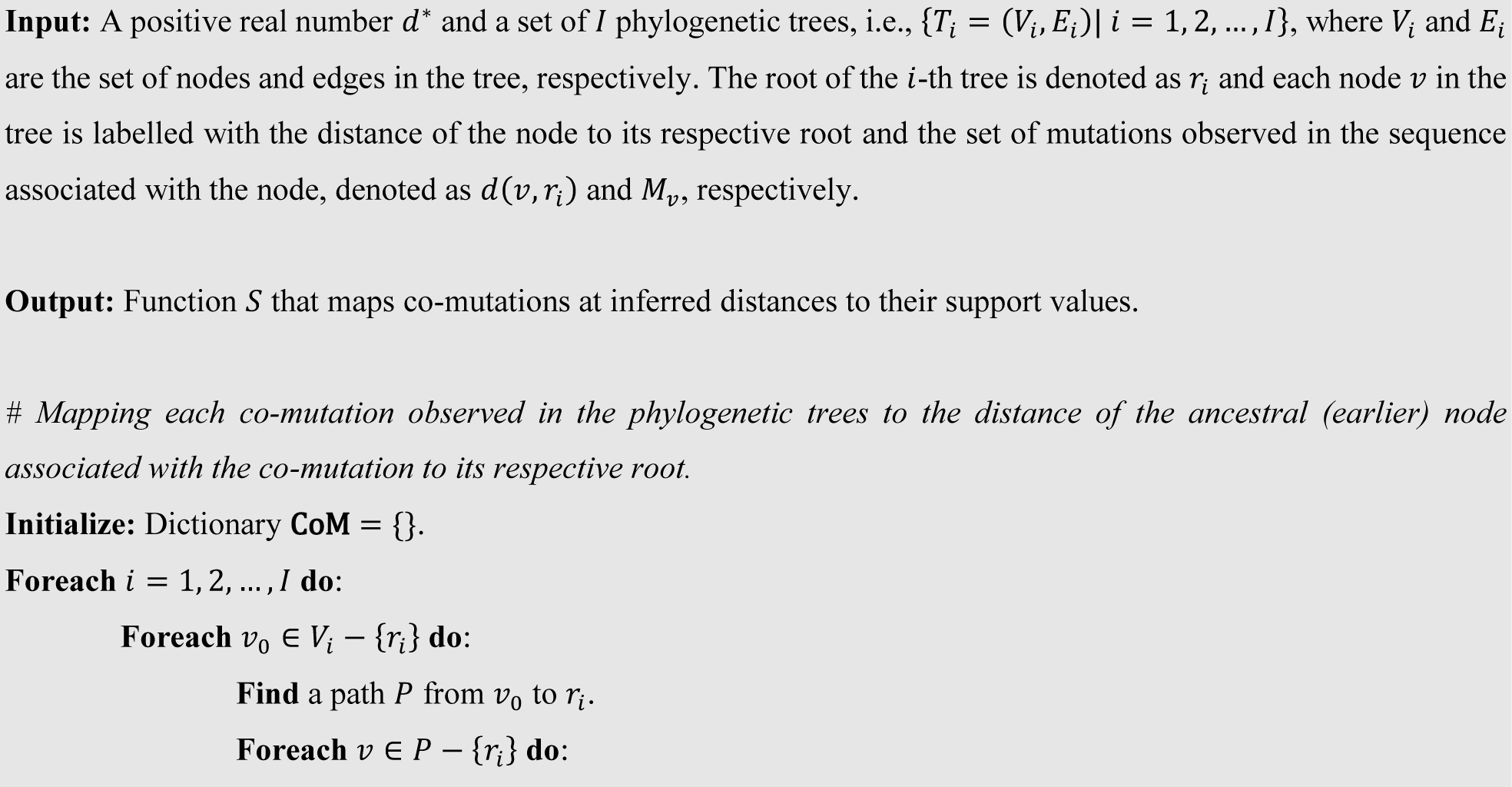
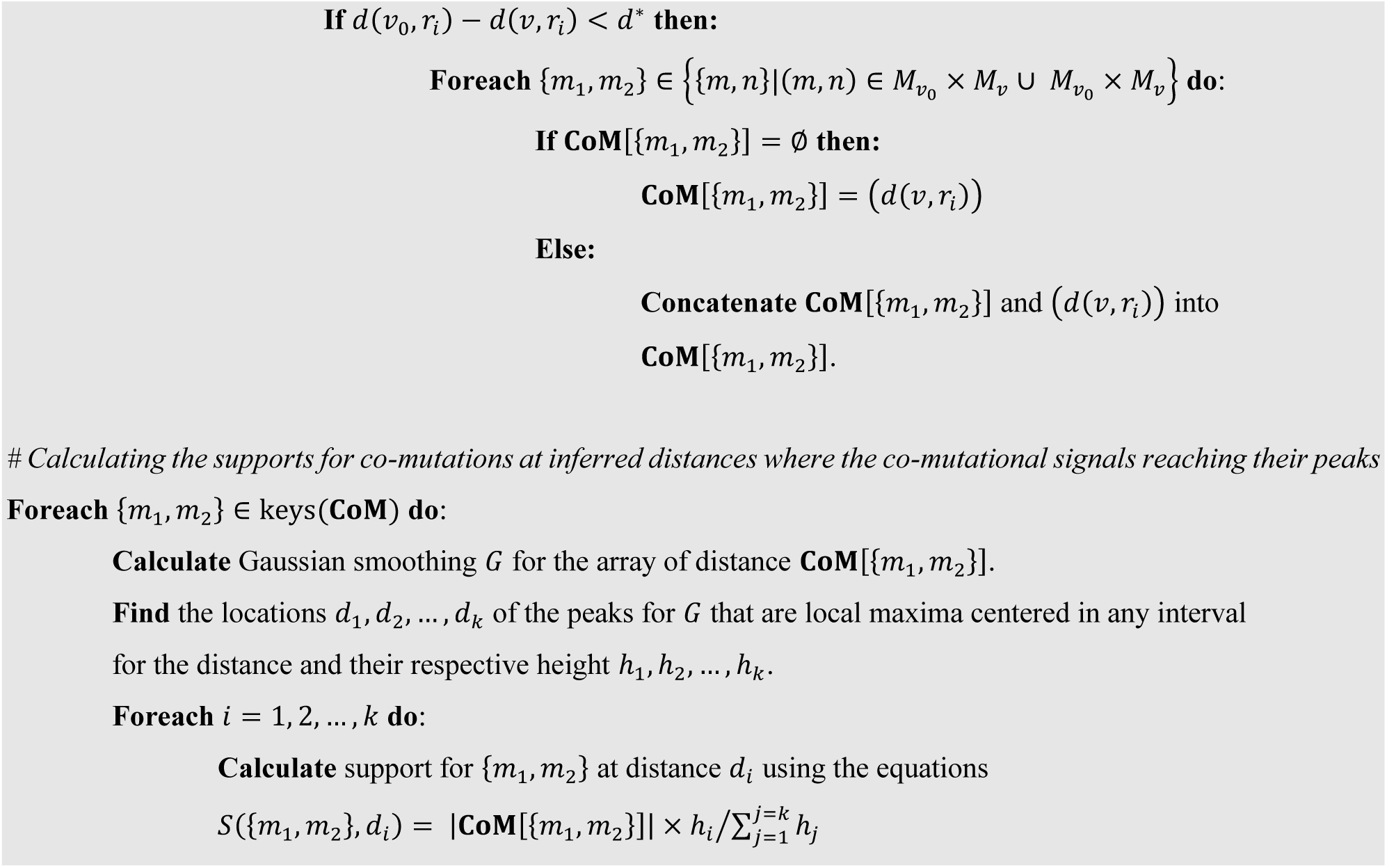

Pseudocode for calculating supports for co-mutations from sampled phylogenetic trees.

### Identification of significant single-mutations and co-mutations

The simulated sequence data generated by Pyvolve were under the assumptions of continuous-time Markov model (Markov process), which include the neutrality and site independence. Hence, we could evaluate whether the real sequence data followed the two assumptions by comparing the distribution of relevant statistics calculated from the real sequence data with that calculated from the simulated sequence data. Here we compared the distributions of supports for single-mutations and co-mutations (as described in the previous section) to evaluate the neutrality and site-independency, respectively. Given the site neutrality assumption was rejected, then a certain quantile (e.g., 95% quantile) of the distribution of supports for simulated single-mutations could be used as a threshold for identifying significant single-mutations in the real data. In similar fashion, if the site independence assumption was rejected, then a certain quantile of the distribution of supports for simulated co-mutations could be used as a threshold for identifying significant co-mutations in the real data.

### Other analyses

To optimize the pipeline and assess the robustness of the output, we calculated the overlap coefficient and Kendall rank correlation between the list of top single-mutations/co-mutations output by two different runs of the pipeline. For this, the supports for each unique single-mutation/co-mutation from each run were first summed and then the single-mutations/co-mutations were sorted in descending order according to their aggregated support. Given the top N single-mutations/co-mutations *X*_*N*_*=(X*_1_, *X*_2_, …, *X*_*N*_) from the first run and *Y*_*N*_ = (*Y*_1_, *Y*_2_, …, *Y*_*N*_) from the second run, the overlap coefficient was calculated using the following formula: overlap(*X*_*N*_, *Y*_*N*_) = | *X*_*N*_ ∩ *Y*_*N*_|/*N*.

For calculating the Kendall rank correlation, we first determined the union of *X*_*N*_ and *Y*_*N*_, i.e., *X*_*N*_ ∪ *Y*_*N*_. Then, we assigned a ranking for each single-mutation/co-mutation in *X*_*N*_ ∪ *Y*_*N*_ according to the first run ordering as well as the second run ordering. For all single-mutations/co-mutations that were in *X*_*N*_ ∪ *Y*_*N*_ but not in *X*_*N*_, the first run assigned their ranking to *N* + 1. In the same way, for all single-mutations/co-mutations that were in *X*_*N*_ ∪ *Y*_*N*_ but not in *Y*_*N*_, the second run assigned their ranking to *N* + 1. The two ranking assignments for single-mutations/co-mutations in *X*_*N*_ ∪ *Y*_*N*_, each of them was sorted in descending order, were then used to calculate the Kendall rank correlation: τ = ((number of concordant pairs) − (number of discordant pairs))/*(L*(*L* − 1)/2), where *L* = |*X*_*N*_ ∪ *Y*_*N*_| and assuming *(q*_1_, *q*_2_, …, *q*_*L*_) and (*r*_1_, *r*_2_, …, *r*_*L*_) be the sorted ranks by the first run and the second run, respectively, pairs of observations *(q*_*i*_, *r*_*i*_) and *(q*_*j*_, *r*_*j*_), where *i* < *j*, are said to be concordant if *q*_*i*_ > *q*_*j*_ and *r*_*i*_ > *r*_*j*_ and discordant if *q*_*i*_ > *q*_*j*_ and *r*_*i*_ < *r*_*j*_, or if *q*_*i*_ < *q*_*j*_ and *r*_*i*_ > *r*_*j*_ (if *q*_*i*_ = *q*_*j*_ and *r*_*i*_ = *r*_*j*_, the pair is neither concordant not discordant).

Finally, for the interpretation of the single-mutations and co-mutations output by the pipeline, each amino acid site was mapped to H3 numbering and epitope regions (epitope A, B, C, D and E). The mapping of the sites to epitope regions was based on the mapping provided in [36].

## Results and Discussions

### HA sequences of influenza A/H3N2 and outlier analysis

We explored the use of the pipeline to uncover significant single-mutations and co-mutations during the evolution of the A/H3N2 HA. For this, the pipeline first subsetted 7,727 non-redundant from 14,301 A/H3N2 HA sequences available in the local influenza genome datasets (the sequence metadata are provided in **Table S1**; the acknowledgement table for sequences obtained from GISAID is provided in **Table S2**). An NJ tree was then reconstructed from the aligned sequences and used for checking the assumption of the constant rate of evolution of the HA sequences. As shown in **Fig. 2A**, the assumption was strongly supported by multiple R-squared value of the root-to-tip regression that was >0.95. Then, using a multiplier for standard deviation of 5 for outlier detection, we identified 58 outliers that were mainly dominated by the sequences collected in the middle of 2012 from a number of regions in North America, including Indiana, Iowa, Michigan Minnesota, Pennsylvania and Ohio. In the phylogenetic tree, the outliers appeared as the tips on the long branch emerging from an internal node at a particular distance to root (**Fig. 2C**; the outliers emerge at distance of 0.11).

**Fig. 2.**
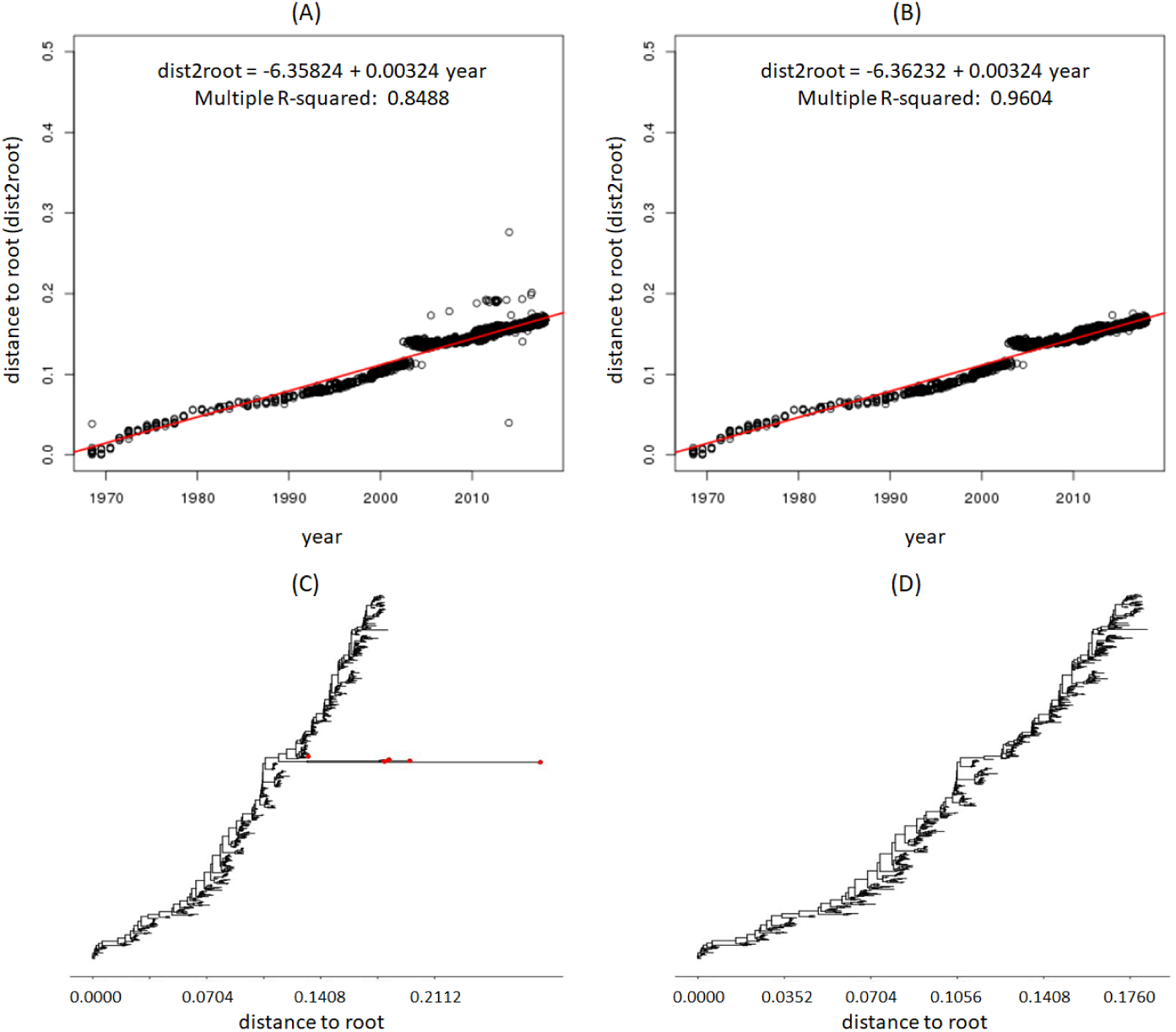
Root-to-tip regressions and phylogenetic trees of the HA sequences of influenza A/H3N2 before and after removing outliers. Outliers are any data point more than five times standard deviation from the average distances of all data points to the regression line. (A) Regression of root-to-tip genetic distance against sampling time for all HA sequences before outlier removal. (B) Regression of root-to-tip genetic distance against sampling time after outlier removal. (C) Neighbor joining tree of a subset of sequences that contain some outliers. The tips corresponding to the outliers are in red. (D) Neighbor joining tree of the same subset of sequences but the outliers are not included.

For further analysis, the outliers were removed. The removal of the outliers improved the regression model (**Fig. 2B**), but it did not remove some obvious gap around year 2003-2005 in the scatter plot. Following some investigation, the gap could be linked to the reassortment event and genome-wide selective sweep during the period that replaced the HA of the major circulating influenza A/H3N2 lineage (clade A) with the HA of a minor co-circulating H3N2 lineage (clade B) [37]. The existence of this phenomenon highlights the importance of the phylogenetic tree-based mutational analysis we proposed – sequence alignment-based approaches may lead to misleading list of mutations when analyzing sequence data arise from such phenomenon.

Despite the gap, we could still safely assume that the substitution rate of the HA of influenza A/H3N2 was constant during the period of sequence data collection due to high R-squared value and its improvement after removing outliers. Indeed, previous studies such as by [38] supported this assumption. Additionally, the assumption that the HA sequences evolve progressively was supported by the ladder-like structure of the phylogenetic tree of HA sequences that excluded the outliers (**Fig. 2D**). Biologically, the ladder-like phylogeny of the HA sequences has been regarded as the consequence of strong directional selection, driven by host immunity [39].

### Estimation of evolutionary parameters and simulation of sequence evolution

Following the outlier analysis, we reconstructed an ML tree under *GTR* + *G* + *I* substitution model using the alignment of all sequence data except the outliers. For *GTR* + *G* + *I* substitution model, the estimated discrete gamma model parameters were 4 for the number of rate categories and 1.1003 for the shape parameters; the estimated proportion of invariant sites was 0.2500; the estimated equilibrium base frequency parameters were 0.4061, 0.1688, 0.1842 and 0.2409 for nucleotide A, C, G and T, respectively; and the estimated rate matrix as follows:

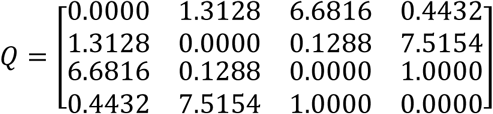

The estimated *GTR* + *G* + *I* substitution model parameters, along with the structure of the ML tree (that included the length of their branches) and the inferred sequence at the tree root, were used to generated simulated HA sequence dataset under *GTR* + *G* + *I* substitution model (see Methods).

In addition, we also estimated the substitution rate and the date of origin of the HA of influenza A/H3N2 sequence. We initially estimated these parameters from the root-to-tip regression that corresponds to the ML tree above, which gave the substitution rate of 0.004618 substitution per year and the date of origin of 1967.34). However, in the downstream analyses, the estimated parameters did not provide reasonable estimated years for the inferred mutations. Thus, we took a different approach, i.e., a bootstrap approach, that averaged the estimated regression parameters calculated from each of the 1000 resampled ML trees reconstructed in the next stage of the pipeline. Using this approach, the estimates for the substitution rate and the date of origin were 0.004369 substitution per site per year and 1967.90, respectively. These parameter values were proven to be better for mutational analyses. In addition to estimating the years of inferred mutations, the estimate for the substitution rate was also used to calculated the threshold distance between ancestor and predecessor in the resampled phylogenetic trees (the *d*^∗^ in **Algorithm 1**) for the identification of co-mutations. In particular, we set the expected number of substitutions per site in one year, i.e., 0.004369 substitutions, as the value for *d*^∗^. One reason for using such *d*^∗^ is due to the fact that influenza epidemics occur yearly and vaccines are updated almost every year by WHO; thus, significant mutational patterns should be observed within 1 year.

### Parameter optimization for mutational analyses using resampled phylogenetic trees and setting the threshold for identifying significant single-mutations and co-mutations

Two parameters associated with the reconstruction of resampled trees in **Sub-pipeline 4** were considered to significantly affect the output of the mutational analysis by **Sub-pipeline 5** and thus optimized. The first one was the number of sequences randomly selected from each isolation year group (or in other words, sample size per year group), which effectively determine the size of the resampled phylogenetic trees (i.e., the number of taxa in the resampled phylogenetic trees). The second one was the number of the resampled trees or the number of the resampling iterations. The values to be explored for the first parameter were 5, 10, 15 and 20, while for the second parameter were 300, 700, 1000, 1500 and 2000. The optimal parameters were determined by investigating the robustness of the output, i.e., comparing the top 500 single-mutations/co-mutations (after summing the supports for each unique single-mutation/co-mutation and sorting the single-mutations/co-mutations in descending order according to their aggregated support) that were output by the pipeline using different combination of these two parameters. In particular, we varied one parameter while fixing another, and calculated the overlap coefficient and Kendall rank correlation between two ranking groups output by the runs whose parameters being varied were consecutive.

As shown in **Fig. 3**, the overlap coefficients between two ranking groups were very high (>.95 and close to 1) for single-mutations regardless we varied the size of the trees or the number of resampled phylogenetic trees. On the other hand, the Kendall rank correlations between two ranking groups stayed high when the moving parameter was the number of resampled trees. However, the correlation got lower when the moving parameter was the sample size per year group; it reached <0.80 when we compared the sample size of 5 and 10. For co-mutations (**Fig. 4**), we once again observed that when the moving parameter was the number of resampled trees, the values for both overlap coefficients and Kendall rank correlations were in general still high (>0.90), except when comparing the number of resampled trees of 300 vs 700 (but still >0.85). But when the moving parameter was the sample size per year group, apparently the overlap coefficients and Kendall rank correlations were higher when we compared the sample size of 10 and 15. Overall, we may conclude that changing the number of resampled trees when it is already >700 does not affect the output of the pipeline significantly, and that the sample size per year group between 10 and 15 provides a more consistent result. The same conclusion could be drawn when we lowered the number of top single-/co-mutations to 100 (data not shown).

**Fig. 3.**
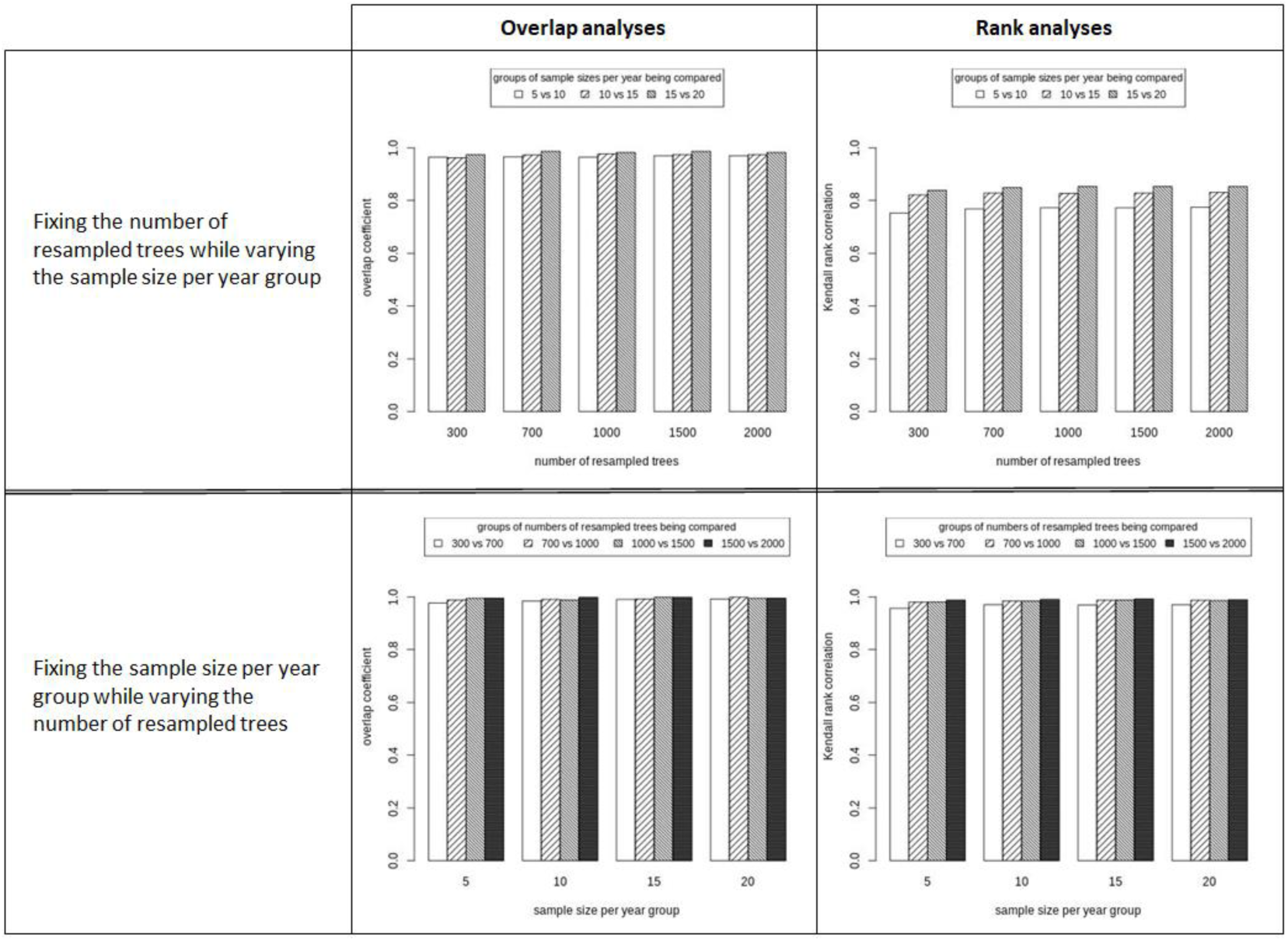
The robustness of single-mutations output by the proposed pipeline when varying the number of resampled trees and sample size per year group.

**Fig. 4.**
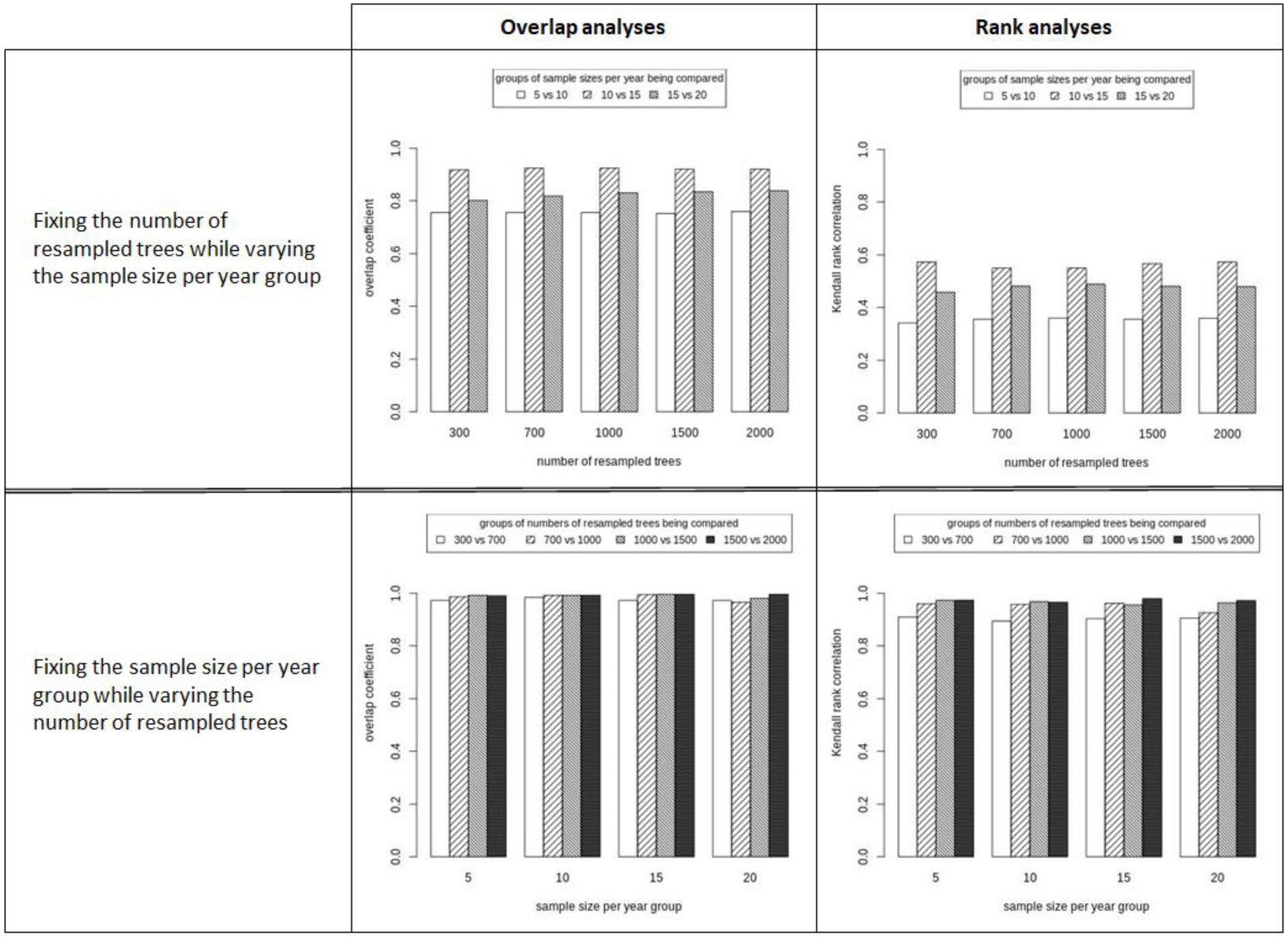
The robustness of co-mutations output by the proposed pipeline when varying the number of resampled trees and sample size per year group.

For further analyses throughout the paper, we fixed the number of resampled trees to 1,000 and the sample size per year group to 15. To demonstrate that these parameters provided robust output, the overall pipeline was run 10 times independently. In similar fashion to previous, the overlap coefficients and Kendall rank correlations between top 500 single-mutations/co-mutations output by two different runs (note that in total, there were 45 pairs of runs) were calculated to assess the robustness of the pipeline. But here, the overlap coefficients and Kendall rank correlations were also calculated for lists of single-mutations/co-mutations that were associated with the simulated sequence datasets in addition to the real one. As it can be seen in **Fig. 5A** and **5B**, the overlap coefficients and Kendall rank correlation between two ranking groups in the case of both real and simulated sequence datasets were very high (>0.90) for single-mutations. For co-mutations, the overlap coefficients were also still high for both datasets (>0.90); however, the Kendall rank correlations dropped to about 0.85 and 0.72 for real and simulated datasets, respectively. Of interest, the overlap coefficients and Kendall rank correlation for real dataset were generally higher than those for simulated dataset. This result indicates that top single-mutations/co-mutations were highly maintained in the analyses of real dataset, and thus some of them must be at the top not by chance.

**Fig. 5.**
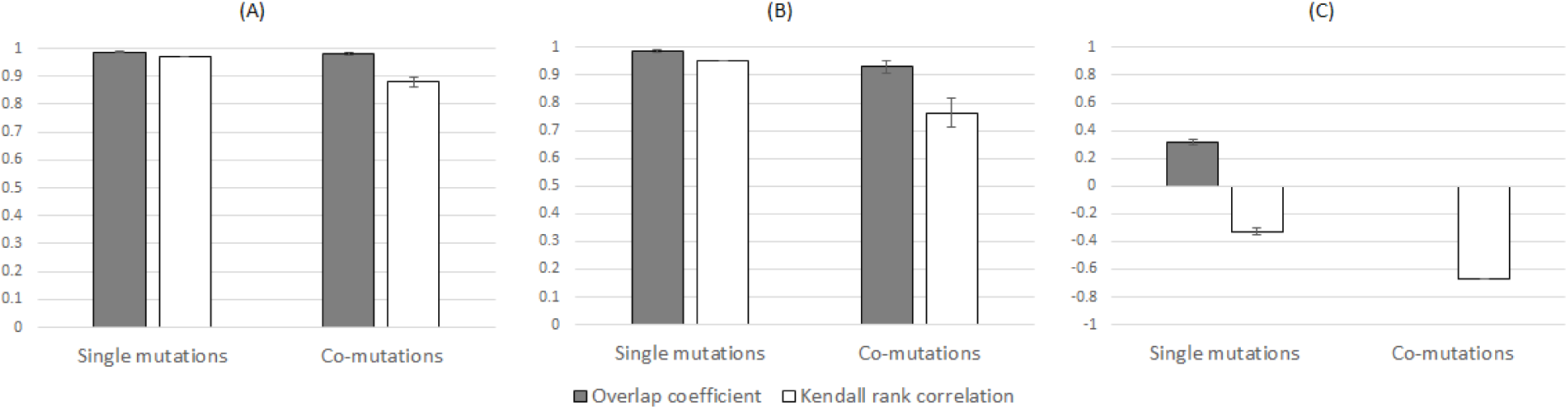
Evaluating the robustness of the proposed analysis pipeline on HA proteins sequences of influenza A/H3N2. Averages of overlap coefficients and Kendall rank correlations for all possible pairwise comparisons between the top lists of single-mutations and co-mutations output by 10 different runs on **(A)** real dataset, **(B)** the same simulated dataset and **(C)** different simulated datasets. The overlap coefficients and Kendall rank correlations were calculated based on top 500 single-mutations or co-mutations of each run.

In addition to inspecting the overlap coefficient and Kendall rank correlation, we also evaluated the robustness of the output by examining the QQ plots that compare distributions of supports for single-mutations/co-mutations from two different runs. If two support distributions are similar, then the points in the QQ-plots will be mainly scattered on the line *y* = *x*. As exemplified in **Fig. 6A, 6B, 6E** and **6F**, the Q-Q plots indeed suggest that different pipeline runs on the same dataset (real or simulated one) output distributions of supports for single-mutations/co-mutations that were highly similar. Thus, the pipeline was robust in term of producing lists of single-mutations/co-mutations that have particular support distributions.

**Fig. 6.**
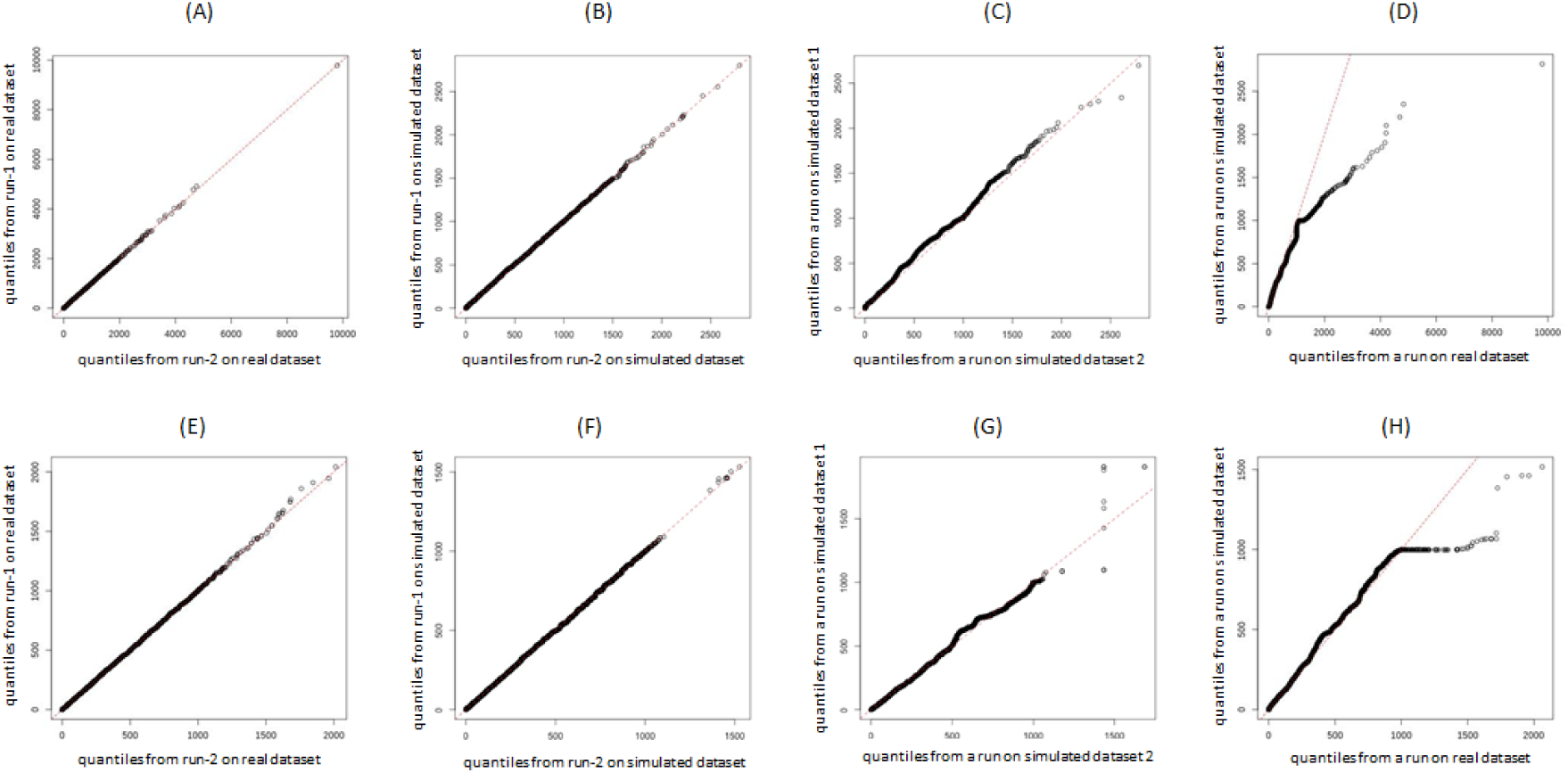
Q-Q plots that compare two distributions of supports for single-mutations and co-mutations output by two different runs on the real dataset (**A** and **E**, respectively), two different runs on the same simulated dataset (**B** and **F**, respectively), two different runs on different simulated datasets (**C** and **G**, respectively), and a run on real dataset versus a run on simulated dataset (**D** and **H**, respectively).

Next, we compared the lists of single-mutations and co-mutations output by **Sub-pipeline 4** and **Sub-pipeline 5** on different simulated sequence datasets. Expectedly, since different simulations likely produce different mutations, we observed low overlap coefficients (which was even 0 for co-mutation case) and negative Kendall rank correlations (that indicated disagreement) between top 500 single-mutations/co-mutations from different datasets (**Fig. 5C**). But mechanistically, different simulations were expected to produce similar distributions of supports for single-mutations/co-mutations. Indeed, despite data points that deviates from the line *y* = *x* in the right tail, this was confirmed by the corresponding QQ-plots (**Fig. 6C** and **6G**).

The deviation from the line *y* = *x* in the right tail was more obvious when we compared the distributions of supports generated from real and simulated datasets (exemplified in **Fig. 6D** and **6H** for single-mutations and co-mutations, respectively). Overall, this once again indicates that the real dataset contained more extremes (i.e., single-mutations/co-mutations with a high support value) than simulated datasets, which appeared not by chance. Thus, as described in the Methods, we may use the support data given by simulated dataset for the identification of statistically significant single-mutations and co-mutations during the real sequence evolution. For this purpose, we set the 95% quantile of the support distributions for single-mutations from simulated dataset as a threshold for the significance of single-mutations from real dataset for both all path and the longest path analysis, and the 99% quantile for the co-mutation case. The 95% quantile for single-mutations gave a threshold of 999.85 and 1000 for all path and the longest path analyses, respectively, and the 99% quantile for co-mutations gave a threshold of 994. As it can be observed in **Fig. 6D** and **6H**, the threshold for all path’s single-mutation and co-mutation analyses were close to the beginning of the deviating points. The appropriateness of choosing higher quantile as a threshold for significant co-mutations was due to higher coverage of co-mutations whose pairs of single-mutations were both significant (94.5% coverage when using 99% quantile, compared to 62.9% coverage when using 95% quantile).

### Patterns of significant single-mutations during the evolution of the HA of influenza A/H3N2 viruses

In all path analysis, 346 significant single-mutations during the evolution of the HA of human influenza A/H3N2 were identified. The majority of the mutations, i.e., 73.2% of the total significant single-mutations observed in the trees occurred in the epitope regions of the HA protein. In more details, the number of single-mutations observed in epitope A, B, C, D and E were 60, 60, 38, 63, and 32 respectively. Nonetheless, a significant number of single-mutations (93 mutations) was also observed in the non-epitope regions. In the longest path analysis, we identified 117 significant single-mutations whose majority (77.8%) occurred in the epitope regions, i.e., 24, 24, 10, 18 and 15 significant single-mutations observed in epitope A, B, C, D and E, respectively. The number of significant single-mutations observed in the non-epitope regions for the longest path analysis was 26. Almost all significant single-mutations in the longest path analysis were also observed in all path analysis, i.e., 111 out of 117.

Sites 144 and 145 in epitope A had the most frequent significant single-mutation occurrences in all path analysis, which were 8 and 11 times, respectively (**Table 1**). Interestingly, the mutations at sites 144 and 145 occurred obvious co-occurrences despite their very close proximity in the HA structure (**Fig.7A**). Sites 45 in epitope C and 193 in epitope B followed the list with the number of significant single-mutation occurrences of 7 times. Nonetheless, only 4 mutations at site 144, 3 mutations at site 145, 1 mutation at site 193 and none at site 45 were identified in the longest path analysis (**Fig.7B**). On the other hand, 5 significant single-mutations at site 189 in epitope B were all observed in the longest path analysis, and this made site 189 as the top site that had the most frequent significant single-mutation occurrences in the longest path (**Fig. 7A** and **7B**). The five significant amino acid substitutions occurring at this position were all different: Q to K (estimated year of occurrence in 1975), K to R (in 1985), R to S (in 1991), S to N (in 2003) and N to K (in 2010) (**Fig. 8**), which may indicate the key role of site 189 as a major driver for the evolution of the HA of influenza A/H3N2 viruses.

**Fig. 7.**
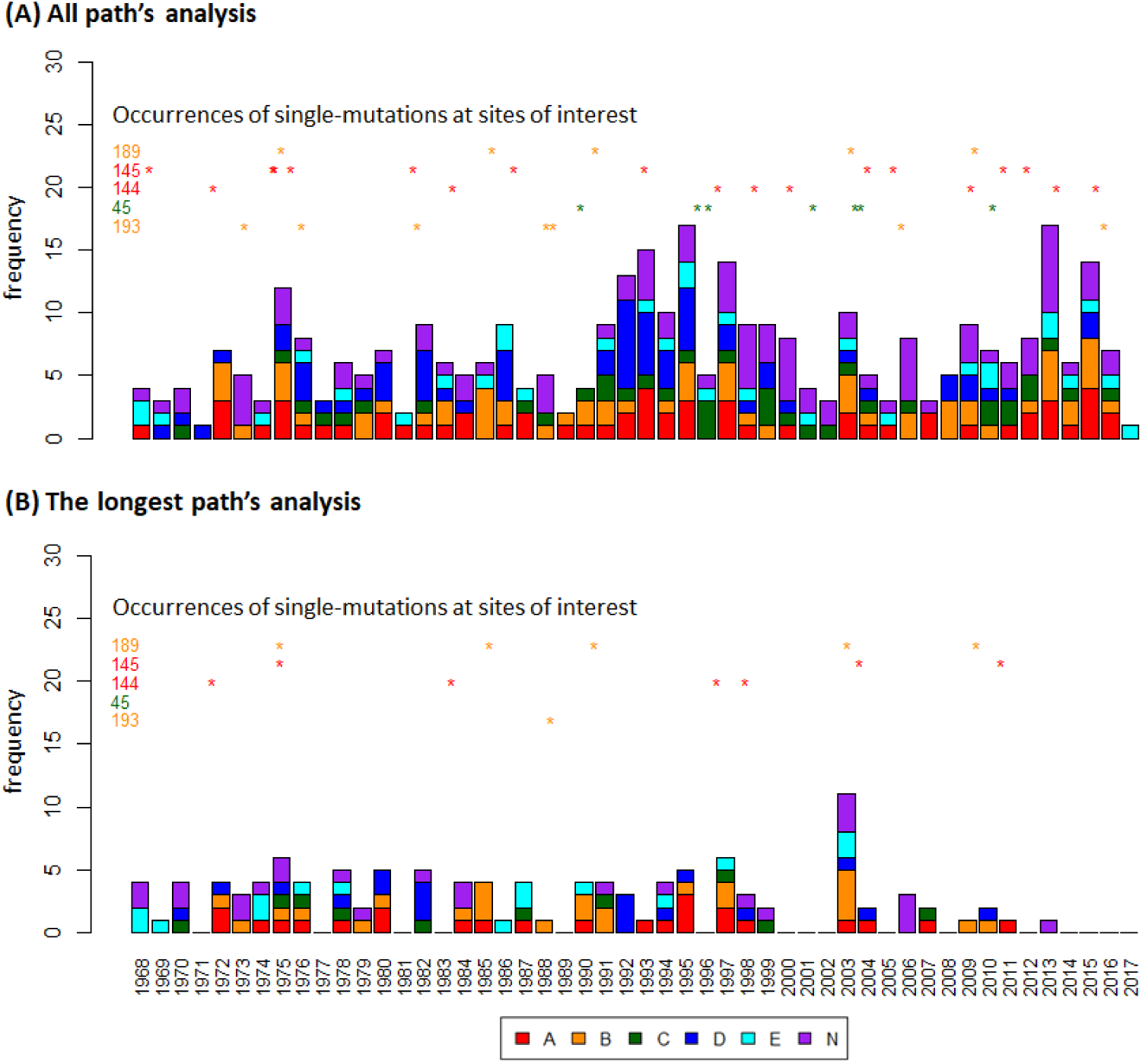
The yearly frequency of significant single-mutations during the evolution of the HA of influenza A/H3N2 detected in (A) all path analysis and (B) the longest path analysis. The occurrences of significant single-mutations at sites of interest are indicated by stars in the corresponding rows. The contribution of each of epitope regions (A, B, C, D and E) and non-epitope region (N) to the total yearly frequency are indicated by color (red for epitope A, orange for epitope B, green for epitope C, blue for epitope D, cyan for epitope E and purple for N).

**Table 1.** The most frequent single-mutations during the evolution of the HA of influenza A/H3N2 observed in all path analysis and the longest path analysis.

| Location in the resampled trees | Number of occurrences | HA sites |  |  |  |  |  |
| --- | --- | --- | --- | --- | --- | --- | --- |
|  |  | Epitope A | Epitope B | Epitope C | Epitope D | Epitope E | Non-epitope |
| All path analysis | 11 | 145 |  |  |  |  |  |
|  | 8 | 144 |  |  |  |  |  |
|  | 7 |  | 193 | 45 |  |  |  |
|  | 6 | 124 | 138 | 53 | 173 | 62 |  |
|  | 5 | 137, 142 | 159, 189 |  | 219, 226, 229 | 92 | 3, 347 |
| The longest path analysis | 5 |  | 189 |  |  |  |  |
|  | 4 | 144 |  |  |  |  |  |
|  | 3 | 124, 133, 145 | 155, 156 | 50 | 172, 226 | 83 |  |

The distribution of the significant single-mutation occurrences in all path analysis over the years from 1968 to 2018 is shown in **Fig. 7A**, and the distribution of the occurrences in the longest path analysis is shown in **Fig. 7B**. In **Fig. 7A**, we can observe the fluctuation of the number of significant single-mutation occurrences and a trend in which more mutations tend to be higher in some ranges of years (e.g., 1991-1995 and 1997-1999) and less in other ranges of years (e.g., 1987-1989, 2000-2002 and 2004-2008). In **Fig. 7B**, a relatively consistent pattern in the number of significant single-mutations in the longest path can be observed before around year 2000, where >3 and ≤3 significant single-mutations were alternatively observed across the years. But after 1998, the number was generally ≤3 (often 0) over the years except in 2003, when the number spiked to 11. Considering significant single-mutations occurred over the years in all path analysis, the absence of significant single-mutations in the longest path analysis is very likely an indication of the presence of multiple competing lineages. The absence in the period 2000-2002 could be linked to the presence of multiple competing lineages of clades A, B and C as reported in [40], while the absence in the recent periods is due to the divergence of clade 3c that began in early 2011 [41]. Furthermore, the fluctuation in the number of significant single-mutations in both all path and the longest path analyses is relevant with the previous report in [42], which confirmed alternating periods of stasis (neutral evolution without apparent substantial antigenic change) and rapid fitness change in the evolution of the HA sequence of influenza A/H3N2.

To further validate our results, we investigated the overlap between the sites associated with significant single-mutations in our list and the sites that have been reported to be under selection pressure in other two studies. First we compared our results against the results by Bush et al. [43] that were based on analysis of sequences collected between 1983 and 1997, only sites associated with significant single-mutations that occurred in the period were considered. As a result, we found that the majority of sites under positive selection pressure in the report were also in our list, i.e., 23 out of 30 sites. The sites that were captured included sites 121, 124, 133, 135, 137, 138, 142, 145, 156, 158, 159, 186, 193, 194, 196, 197, 201, 219, 226, 246, 262, 275 and 276; while the sites that were not captured included sites 80, 128, 182, 190, 220, 310 and 312. In contrast, we recovered only few sites under negative selection pressure in the report, i.e., 3 out of 18 sites. Indeed, these observations were expected since the significant single-mutations we captured were the ones that ought to be fixed in the following generation of HA sequences of the viruses. The coverage of sites under positive selection pressure was further confirmed when comparing our list with the result in [44], which interestingly had a moderate overlap with the result in Bush et al. (only 13 sites in the overlap; 22 sites in [44] are not in [43], and 17 sites in [43] are not in [44]). In particular, our list of significant single-mutations in the period before 2012 (to match with the collection dates of sequences in [44]) covered almost all of the sites in the patches under positive selection pressure uncovered in the study, which include sites 47, 48, 50. 53, 62, 92, 94, 137, 140, 142, 144, 145, 156-159, 172-175, 186, 188, 189, 192, 193, 196-199, 220, 229, 275 and 276 (sites 91 and 171 were not covered).

**Fig. 8.**
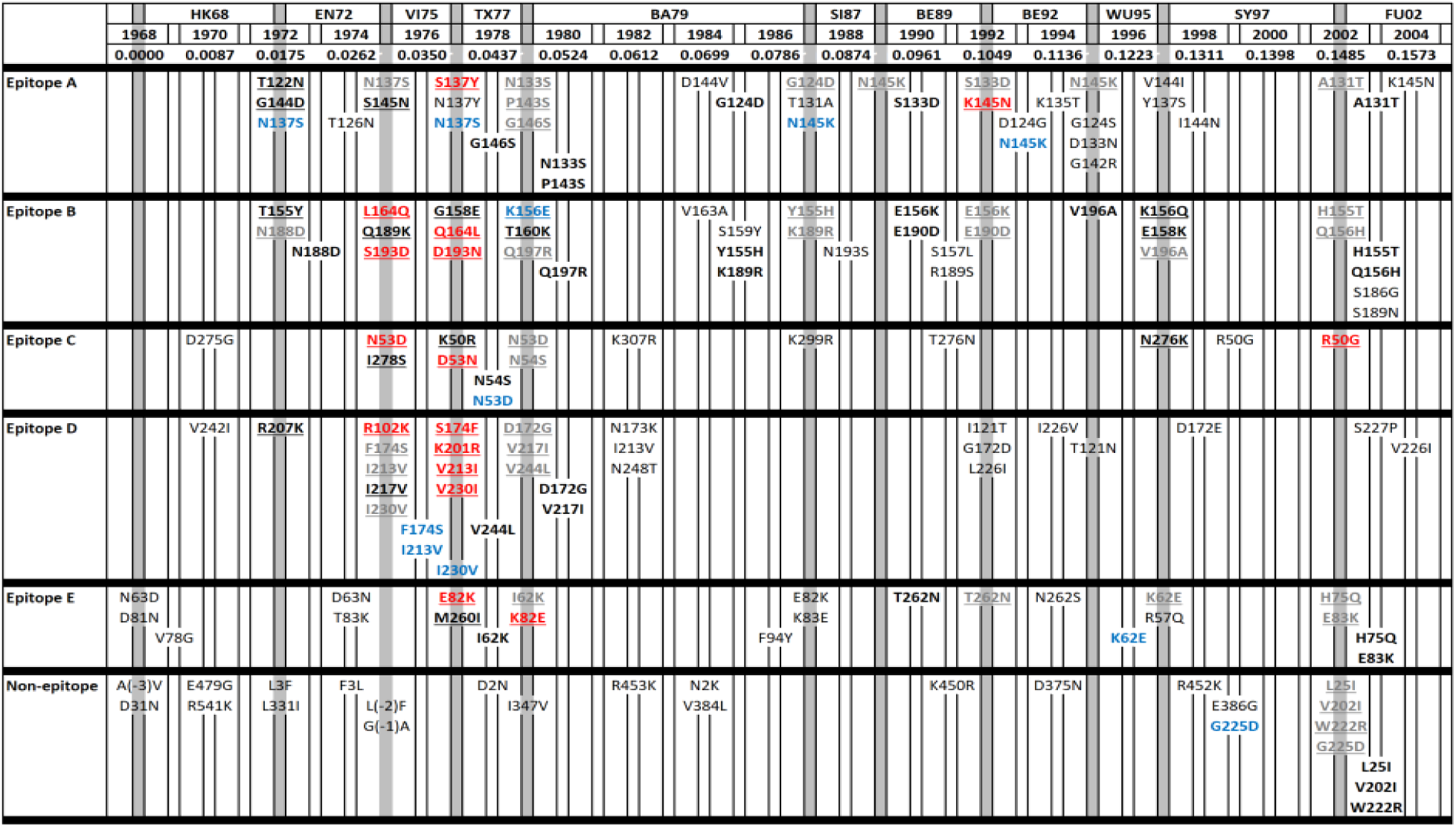
Overlap between significant single-mutations and mutations playing a role in antigenic cluster transitions of influenza A/H3N2 (as reported in [11]). Significant single-mutations obtained from the longest path analysis are in black; they are bolded if reported in [11] and underlined if their occurrence is in very close proximity to the year of the new antigenic cluster emergence. Significant-single mutations only obtained from all path analysis and reported in [11] are in bold blue; one of them is underlined to indicate that its occurrence was in very close proximity to the year of the new antigenic cluster emergence. Mutations reported in [11] that are not found in our analysis is in bold red and underlined. Mutations reported in [11] that are found in our analysis but their occurrence are not in very close proximity are in bold grey and underlined. (HK: Hong Kong, EN: England, VI: Victoria, TX: Texas, BA: Bangkok, SI: Sichuan, BE: Beijing, WU: Wuhan, SY: Sydney, FU: Fujian).

**Fig. 9.**
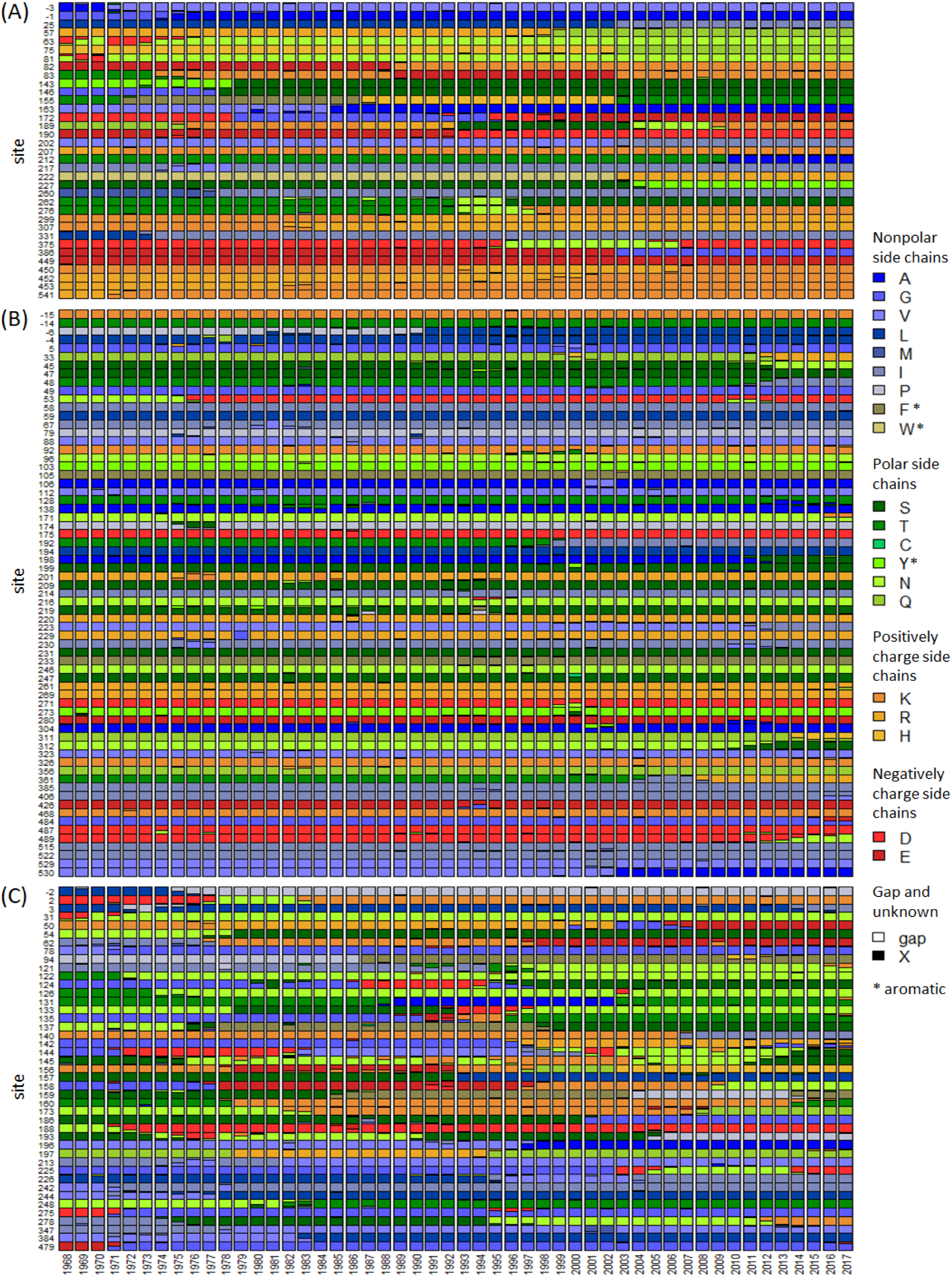
Yearly frequency of amino acid residues during 50 years of evolution of the HA of influenza A/H3N2 viruses for: (A) sites only found in the longest path analysis, (B) sites only found in all path analysis, and (C) sites found in both all path and the longest path analyses.

Next, we also revealed that the majority of single-mutations in the relevant period were associated with antigenic cluster transitions as reported in [11]. As shown in **Fig. 8**, out of 67 single-mutations (4 of them in non-epitope region) in the report, 51 of them were recovered in our analysis: 40 at the longest path (in black and bolded; 15 of them are underlined to indicate that their occurrence was in very close proximity to the year of the new antigenic cluster emergence) and 11 at non-longest paths (in blue and bolded; 1 of them is underlined to indicate that its occurrence was in very close proximity to the year of the new antigenic cluster emergence). In the table, we also showed additional 65 single-mutations that were not in the report.

Additionally, we also noted that our analysis recovered almost all mutations at the 7 sites near the receptor binding site (i.e., 145, 155, 156, 158, 159, 189 and 193) that had been experimentally shown to be responsible for antigenic cluster transitions during influenza A/H3N2 virus evolution [13]. These include T155Y during transition from HK68 to EN72; Q189K during transition from EN72 to VI75; G158E during transition from VI75 to TX77; K156E during transition from TX77 to BA79; Y155H, S159Y and K189R during transition from BA79 to SI87; N145K and N193S during transition from SI87 to BE89; S133D and E156K during transition from SI87 to BE92; N145K during transition from BE92 to WU95; K135T, K156Q and E158K during transition from WU95 to SY97; and Q156H during transition from SY97 to FU02. Only mutation D193N during transition from VI75 to TX77 was not recovered. Moreover, the significant single-mutations found in this study also recovered the top 15 cluster-transition determining sites recently reported in [45], which included sites 122, 133, 135, 144, 145, 155, 156, 158, 189, 190, 193, 197, 262, 276 and 278.

Lastly, we present the frequency patterns of amino acid residues during 50 years of evolution of the HA of influenza A/H3N2 viruses at each site associated with significant single-mutations found in our analyses. Given the set of significant single-mutations from all path analysis denoted by *A* and the set of significant single-mutations from the longest path analysis denoted by *B*, we grouped the sites into three categories: (1) sites appeared in *B* but not in *A* − *B*, (2) sites appeared in *A* − *B* but not in *B*, and (3) sites appeared in *B* and *A* − *B*. **Fig. 9A** reveals that the hallmark of mutational pattern at sites in the first group was the numerous replacements of a dominant amino acid residue with another dominant amino acid residue, and each dominant amino acids generally could dominate for a long period of time. On the other hand, **Fig. 9B** reveals that sites in the second group often presented temporary appearance of competing amino acid residues. Even though the competing amino acid residue once became the majority, it failed to dominate for a long term. Finally, **Fig. 9C** reveals that sites in the third group presented more dynamics in their mutational patterns, which combined the characteristics mentioned earlier. Practically, with regards to the notions in [11], sites in the first group may play more roles in the enhancement of antigenic drift or shaping the evolution of the HA; sites in the second group may play more roles in compensatory mutations for retaining higher fitness and associated with clades emerged during specific epidemic seasons; and sites found in the third group could both enhance antigenic drift as well as compensate other mutations that enhanced antigenic drift.

### Patterns of significant co-mutations during the evolution of the HA of influenza A/H3N2 viruses

Using a threshold distance between ancestor and predecessor in the resampled phylogenetic trees (the *d*^∗^ in **Algorithm 1**) of 0.004369 substitution per site for co-mutation detection and the 99% quantile of support distribution for co-mutations from the simulated data as a threshold for significance, and only considered co-mutations consisting of a pair of significant single-mutations, we identified 343 significant co-mutations output by the pipeline. However, when considering site pairs of the observed co-mutations, no site pair was observed more than twice during influenza A/H3N2 virus evolution. In fact, we only identified 8 site pairs that occurred twice, including 3-144, 62-144, 62-158, 121-142, 144-158, 155-189, 159-225 and 226-262; the rests occurred only once. Nonetheless, when considering the co-mutational networks, some sites had higher degree or number of co-mutational incidents with other sites. The site with the highest degree was 144, with a degree of 20. Sites 145 and 189 with a degree of 13 followed the top list. Sites 124 and 226 had a degree of 12; sites 92 and 156 had a degree of 11; and the rest had a degree of 10 or less.

When considering the epitopes, we found that the co-mutations mainly involved sites in non-epitope region (N) and epitope A, B and D. The frequencies for co-mutations involving epitopes A and B and involving N and epitope B were the highest, i.e., 33 times. The frequencies for co-mutations involving N and epitope D, N and epitope A, N and N, epitopes A and D, and epitopes B and D were 28, 27, 27, 23, and 22, respectively; the rests were 20 or less. Next, epitope region with the highest degree was epitope A (104), followed by B (80), D (54), E (26) and C (13). The degree of N was higher than the degree of epitopes C, D and E, i.e., 66. This observation suggests the importance of mutations in non-epitope region that may play a role in maintaining the integrity of the HA.

Next, we explored the temporal patterns of the significant co-mutations found in this study. For this, we grouped the significant co-mutations by the estimated years of their occurrences by using year group 1968-1972, 1973-1977, 1978-1982, and so on until 2013-2017 (as a note, there was no co-mutation observed in 2018). The networks of co-mutational site pairs observed in each year group and their transitions are shown in **Fig. 10**. The yearly frequency of co-mutations for each year group is also shown on the left or right of the corresponding network. As an initial observation, we can see that the number of co-mutations over the years were continuously up and down. For some years, the number of co-mutations was even very low (less than 5 and even 0), while for some other years, the number was quite high (>10). Then, we can also observe that for some transitions between year groups, the overlap between the sites were relatively small. Only one site was shared by year groups 1973-1977 and 1978-1982, 1983-1987 and 1988-1992, and 1998-2002 and 2003-2007; two sites were shared by year groups 2003-2007 and 2008-2012; and three sites were shared by year groups 1968-1972 and 1973-1977. Larger overlaps were observed between year groups 1978-1982 and 1983-1987 (4 overlapping sites), 1988-1992 and 1993-1997 (9 sites), 1993-1997 and 1998-2002 (4 sites), and 2008-2012 and 2013-2017 (6 sites). In addition, we can also observe a number of cliques in the co-mutational networks. Of particular interest, we can see that sites with higher degree, i.e., 124, 144, 145, 189 and 226, were usually part of the cliques.

**Fig. 10.**
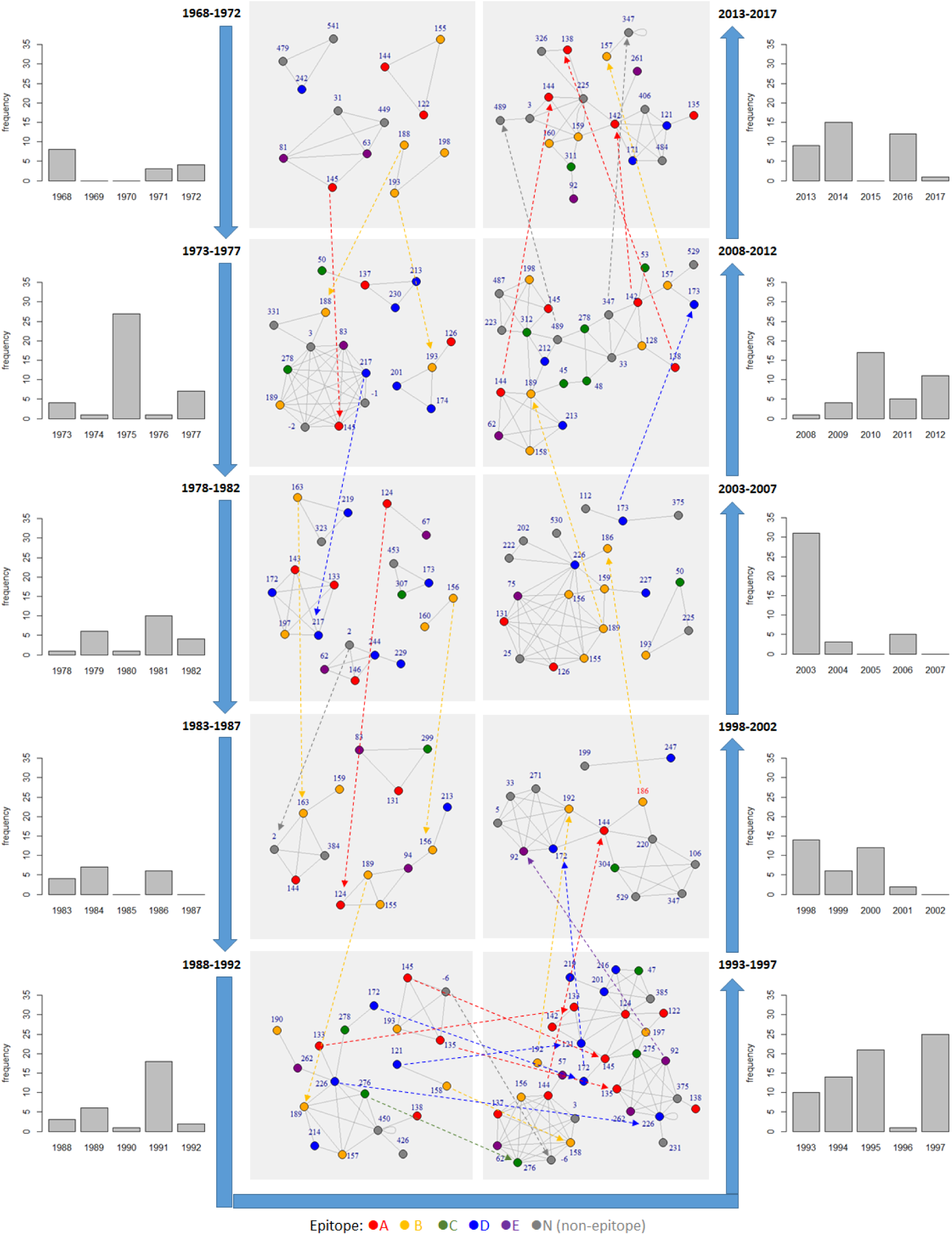
Networks of site pairs that significantly co-mutated every lustrum (a period of 5 years) during 50 years of evolution of the HA of influenza A/H3N2 viruses. The networks considered all significant co-mutations associated with significant single-mutations in all path analysis.

When considering co-mutations whose pair consisting of significant single-mutations in the longest path analysis, site pairs 137-158 and 155-189 co-mutated twice. Interestingly, sites 83 had the highest degree (13), followed by sites 144 (10), 131 (8), 137 (7), 156 (7) and 189 (7). The co-mutations involving epitopes A and B stayed at the top (16 times), and epitope A still had the highest degree (49). Finally, the corresponding temporal patterns of the significant co-mutations (**Fig. 11**) also revealed the presence of a number of cliques. The lack and absence of the co-mutational networks in the last 2 periods corresponds to the lack and absence of significant single-mutations in the longest path explained previously.

**Fig. 11.**
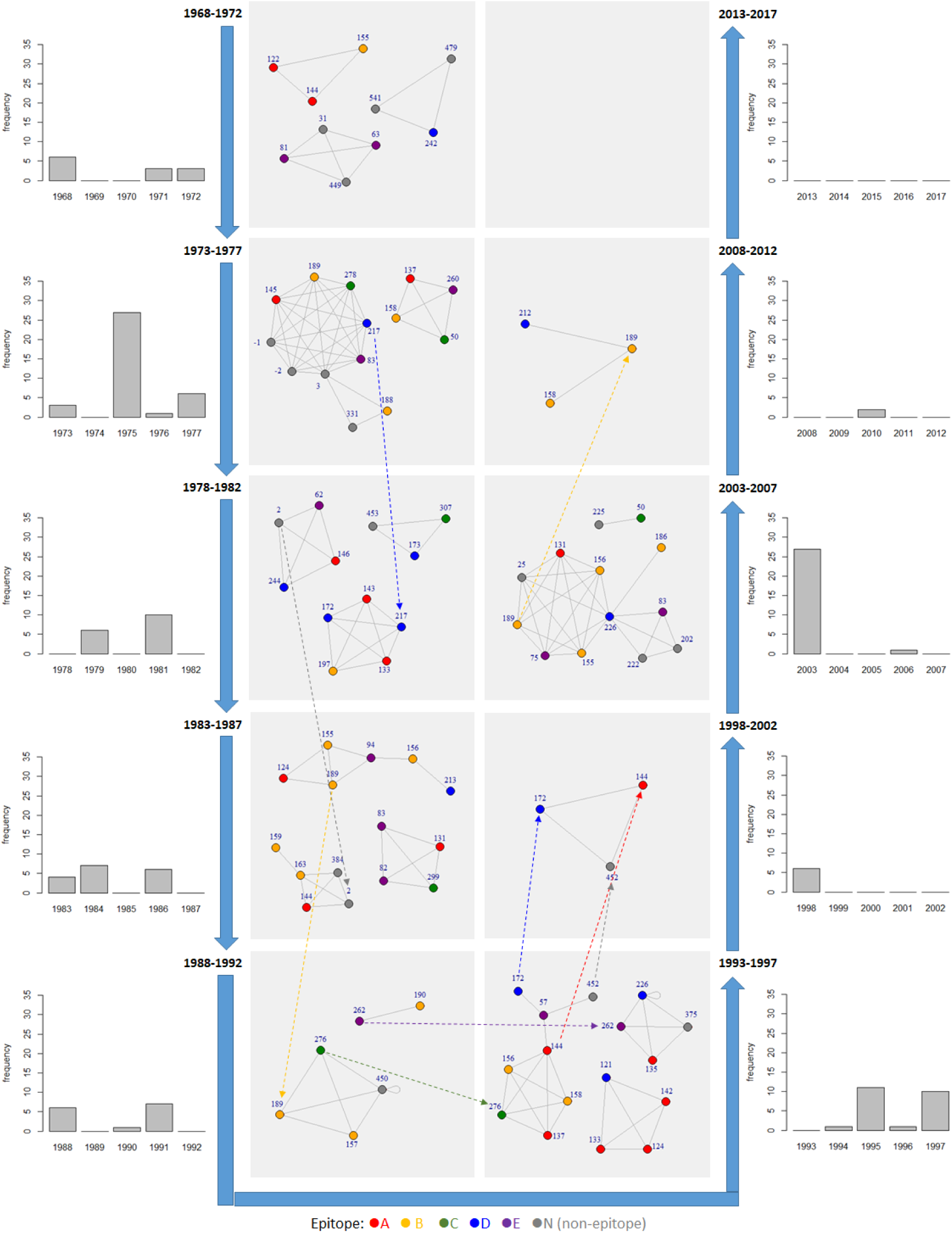
Networks of site pairs that significantly co-mutated every lustrum (a period of 5 years) during 50 years of evolution of the HA of influenza A/H3N2 viruses. The networks only considered significant co-mutations associated with significant single-mutations in the longest path analysis.

Overall, consistent with previous report by [42] and [46], our observation suggests that during the evolution of influenza A/H3N2, the increased fitness of the HA was occasionally contributed by simultaneous multi-site co-mutations. Here we argue that the events were likely driven by mutations at a number of influential sites frequently observed as part of cliques in **Fig. 10** and **11**, including sites 83, 144, 145, and 189. Furthermore, we also noted that a new configuration of amino acids at these sites seemed to drive mutations at different sites that were not explored in the previous years.

## Conclusion

In this study, we present a novel phylogenetic tree-based pipeline for analyzing mutational patterns during the evolution of influenza virus sequences. We demonstrated the use of the pipeline to investigate the single-mutational and co-mutational patterns of the HA sequences of influenza A/H3N2 viruses. In addition to known biologically significant mutations in HA and related patterns, our approach allowed the identification of three groups of sites based the outcomes of all path and the longest path analyses on the resampled phylogenetic trees. Sites in each group were shown to exhibit specific characteristics of mutational pattern, which could be linked to their roles in antigenic drift: enhancing antigenic drift, compensating other mutations that enhance antigenic drift, or both. This classification may potentially be useful for evaluating candidate vaccines targeting the HA.

## Supplementary Material

Supplementary data are available at Molecular Biology and Evolution online. The codes for the proposed pipeline are available at DR-NTU (Data) https://doi.org/10.21979/N9/PDYCUD.

## Acknowledgement

This project was supported by AcRF Tier 2 grant MOE2014-T2-2-023, Ministry of Education, Singapore and A*STAR-NTU-SUTD AI Partnership Grant.

## Author Contributions

FXI conceived and designed the overall pipeline. FXI, AD and CWL contributed to the writing of python/R/shell codes. FXI wrote the article; XZ helped the writing of the introduction and discussions. JZ and CK reviewed the article.

## Notes

https://doi.org/10.21979/N9/PDYCUD

