## Supplementary material for "Phylogenetic Tree-based Pipeline for Uncovering Mutational Patterns during Influenza Virus Evolution": Table S1

**Table S1.** GenBank and GISAID accessions and metadata for sequences used in this study.

| Sequence ID | Influenza strain | Collection date | Database source |
|---|---|---|---|
| DQ415320 | A/TW/3446/02(H3N2) | 2002 | NCBI |
| DQ415322 | A/TW/229/03(H3N2) | 2003 | NCBI |
| DQ415319 | A/TW/872/02(H3N2) | 2002 | NCBI |
| DQ415321 | A/TW/3565/02(H3N2) | 2002 | NCBI |
| DQ415323 | A/TW/3286/03(H3N2) | 2003 | NCBI |
| CY034100 | A/Panama/2007/1999(H3N2) | 1999 | NCBI |
| CY034116 | A/Wisconsin/67/2005(H3N2) | 2005-08 | NCBI |
| DQ415324 | A/TW/36/04(H3N2) | 2004 | NCBI |
| DQ415325 | A/Taiwan/220/2004(H3N2) | 2004 | NCBI |
| DQ415326 | A/TW/875/04(H3N2) | 2004 | NCBI |
| CY036903 | A/Shandong/9/1993(H3N2) | 1993 | NCBI |
| CY033606 | A/Beijing/32/1992(H3N2) | 1992 | NCBI |
| CY026163 | A/Colorado/UR06-0535/2007(H3N2) | 7/3/2007 | NCBI |
| CY025341 | A/Florida/UR06-0597/2007(H3N2) | 14/3/2007 | NCBI |
| CY025349 | A/Colorado/UR06-0206/2007(H3N2) | 6/2/2007 | NCBI |
| CY025277 | A/Illinois/UR06-0567/2007(H3N2) | 9/3/2007 | NCBI |
| CY025018 | A/Auckland/583/2000(H3N2) | 12/2/2000 | NCBI |
| DQ508849 | A/Leningrad/360/1986(H3N2) | 1986 | NCBI |
| DQ508825 | A/Bangkok/1/1979(H3N2) | 1979 | NCBI |
| CY025050 | A/Auckland/610/2002(H3N2) | 29/7/2002 | NCBI |
| DQ469962 | A/Ontario/RV1273/2005(H3N2) | 2005 | NCBI |
| CY025042 | A/Auckland/598/2000(H3N2) | 9/10/2000 | NCBI |
| CY025421 | A/California/UR06-0347/2007(H3N2) | 19/2/2007 | NCBI |
| CY025485 | A/NewYork/UR06-0515/2007(H3N2) | 5/3/2007 | NCBI |
| CY026275 | A/Texas/UR06-0566/2007(H3N2) | 9/3/2007 | NCBI |
| CY026555 | A/Texas/UR06-0356/2007(H3N2) | 20/2/2007 | NCBI |
| CY025611 | A/Colorado/UR06-0557/2007(H3N2) | 8/3/2007 | NCBI |
| CY025643 | A/NewYork/UR06-0040/2007(H3N2) | 17/1/2007 | NCBI |
| CY027579 | A/California/UR06-0463/2007(H3N2) | 28/2/2007 | NCBI |
| CY025731 | A/NewYork/UR06-0529/2007(H3N2) | 6/3/2007 | NCBI |
| CY025747 | A/Texas/UR06-0480/2007(H3N2) | 2/3/2007 | NCBI |
| CY028740 | A/Virginia/UR06-0489/2007(H3N2) | 1/3/2007 | NCBI |
| CY025835 | A/NewYork/UR06-0373/2007(H3N2) | 21/2/2007 | NCBI |

| | | | |
|---|---|---|---|
| CY026251 | A/Colorado/UR06-0558/2007(H3N2) | 7/3/2007 | NCBI |
| CY026747 | A/Illinois/UR06-0528/2007(H3N2) | 6/3/2007 | NCBI |
| CY025907 | A/Kentucky/UR06-0370/2007(H3N2) | 21/2/2007 | NCBI |
| CY025931 | A/Colorado/UR06-0453/2007(H3N2) | 27/2/2007 | NCBI |
| CY026843 | A/Oregon/UR06-0289/2007(H3N2) | 13/2/2007 | NCBI |
| CY027075 | A/Oregon/UR06-0273/2007(H3N2) | 13/2/2007 | NCBI |
| CY027107 | A/Oregon/UR06-0389/2007(H3N2) | 19/2/2007 | NCBI |
| CY027795 | A/Oregon/UR06-0200/2007(H3N2) | 6/2/2007 | NCBI |
| CY026035 | A/Alabama/UR06-0482/2007(H3N2) | 1/3/2007 | NCBI |
| CY025477 | A/Illinois/UR06-0334/2007(H3N2) | 19/2/2007 | NCBI |
| CY030053 | A/Colorado/UR06-0452/2007(H3N2) | 28/2/2007 | NCBI |
| CY027571 | A/Colorado/UR06-0534/2007(H3N2) | 7/3/2007 | NCBI |
| CY026603 | A/Washington/UR06-0225/2007(H3N2) | 8/2/2007 | NCBI |
| CY025739 | A/California/UR06-0118/2007(H3N2) | 29/1/2007 | NCBI |
| CY027587 | A/Illinois/UR06-0546/2007(H3N2) | 6/3/2007 | NCBI |
| CY026707 | A/Colorado/UR06-0454/2007(H3N2) | 26/2/2007 | NCBI |
| CY026259 | A/Illinois/UR06-0555/2007(H3N2) | 7/3/2007 | NCBI |
| CY025867 | A/Vermont/UR06-0483/2007(H3N2) | 2/3/2007 | NCBI |
| CY026827 | A/Oregon/UR06-0450/2007(H3N2) | 26/2/2007 | NCBI |
| CY026883 | A/Kentucky/UR06-0158/2007(H3N2) | 5/2/2007 | NCBI |
| CY027715 | A/Illinois/UR06-0402/2007(H3N2) | 24/2/2007 | NCBI |
| CY027499 | A/Ohio/UR06-0410/2007(H3N2) | 23/2/2007 | NCBI |
| CY026307 | A/Virginia/UR06-0021/2007(H3N2) | 8/1/2007 | NCBI |
| CY027867 | A/Virginia/UR06-0580/2007(H3N2) | 12/3/2007 | NCBI |
| CY027547 | A/Colorado/UR06-0024/2007(H3N2) | 8/1/2007 | NCBI |
| CY025413 | A/Vermont/UR06-0524/2007(H3N2) | 5/3/2007 | NCBI |
| CY033457 | A/Vermont/UR06-0329/2007(H3N2) | 16/2/2007 | NCBI |
| CY028299 | A/Ohio/UR06-0256/2007(H3N2) | 12/2/2007 | NCBI |
| CY025603 | A/Vermont/UR06-0471/2007(H3N2) | 28/2/2007 | NCBI |
| CY025755 | A/Florida/UR06-0150/2007(H3N2) | 2/2/2007 | NCBI |
| CY025875 | A/Vermont/UR06-0470/2007(H3N2) | 27/2/2007 | NCBI |
| CY025883 | A/Vermont/UR06-0486/2007(H3N2) | 1/3/2007 | NCBI |
| CY025899 | A/Illinois/UR06-0436/2007(H3N2) | 27/2/2007 | NCBI |
| CY026923 | A/Ohio/UR06-0494/2007(H3N2) | 2/3/2007 | NCBI |
| CY027683 | A/Oregon/UR06-0202/2007(H3N2) | 6/2/2007 | NCBI |
| CY028371 | A/Oregon/UR06-0221/2007(H3N2) | 8/2/2007 | NCBI |
| CY026027 | A/Illinois/UR06-0030/2007(H3N2) | 15/1/2007 | NCBI |

| | | | |
|---|---|---|---|
| EU268227 | A/Wyoming/03/2003(H3N2) | 2003 | NCBI |
| CY034108 | A/Wyoming/03/2003(H3N2) | 2003 | NCBI |
| CY027539 | A/Colorado/UR06-022/2007(H3N2) | 8/1/2007 | NCBI |
| CY030205 | A/Virginia/UR06-0605/2007(H3N2) | 13/3/2007 | NCBI |
| CY034958 | A/HongKong/1-1-MA-12C/1968(H3N2) | 1968 | NCBI |
| CY033988 | A/HongKong/1-1-MA-12D/1968(H3N2) | 1968 | NCBI |
| CY034966 | A/HongKong/1-1-MA-12E/1968(H3N2) | 1968 | NCBI |
| CY033025 | A/HongKong/1-5/1968(H3N2) | 1968 | NCBI |
| CY033996 | A/HongKong/1-8/1968(H3N2) | 1968 | NCBI |
| CY033057 | A/HongKong/1-10/1968(H3N2) | 1968 | NCBI |
| CY033121 | A/HongKong/1-4-MA21-3/1968(H3N2) | 1968 | NCBI |
| CY034012 | A/HongKong/1-6-MA21-2/1968(H3N2) | 1968 | NCBI |
| CY034020 | A/HongKong/1-8-MA21-1/1968(H3N2) | 1968 | NCBI |
| CY033545 | A/HongKong/1-11-MA21-2/1968(H3N2) | 1968 | NCBI |
| CY034036 | A/HongKong/1-12-MA21-3/1968(H3N2) | 1968 | NCBI |
| CY032501 | A/Managua/18/2007(H3N2) | 21/6/2007 | NCBI |
| CY032429 | A/Managua/20/2007(H3N2) | 22/6/2007 | NCBI |
| CY032485 | A/Managua/22/2007(H3N2) | 20/6/2007 | NCBI |
| CY032437 | A/Managua/25/2007(H3N2) | 27/6/2007 | NCBI |
| CY032477 | A/Managua/29/2007(H3N2) | 22/6/2007 | NCBI |
| CY036807 | A/X-117(PuertoRico/8/1934-Beijing/32/1992)(H3N2) | 1992 | NCBI |
| CY033638 | A/NewYork/55/2004(H3N2) | 2004 | NCBI |
| CY034414 | A/NewYork/UR06-0607/2007(H3N2) | 14/3/2007 | NCBI |
| EU399751 | A/Ontario/1252/2007(H3N2) | 2007 | NCBI |
| CY034950 | A/HongKong/1-1-MA-12B/1968(H3N2) | 1968 | NCBI |
| CY034004 | A/HongKong/1-11/1968(H3N2) | 1968 | NCBI |
| CY033113 | A/HongKong/1-4-MA21-2/1968(H3N2) | 1968 | NCBI |
| CY033137 | A/HongKong/1-8-MA21-2/1968(H3N2) | 1968 | NCBI |
| CY034982 | A/HongKong/1-10-MA21-2/1968(H3N2) | 1968 | NCBI |
| CY033537 | A/HongKong/1-11-MA21-1/1968(H3N2) | 1968 | NCBI |
| CY035102 | A/Managua/21/2007(H3N2) | 21/6/2007 | NCBI |
| CY035086 | A/Managua/3/2007(H3N2) | 7/6/2007 | NCBI |
| CY032549 | A/Managua/31/2007(H3N2) | 18/6/2007 | NCBI |
| CY032541 | A/Managua/7/2007(H3N2) | 5/6/2007 | NCBI |
| CY035110 | A/NymcX-157B(PuertoRico/8/1934-NewYork/55/2004)(H3 | 2004 | NCBI |
| CY037311 | A/Kuwait/AF03/2008(H3N2) | 2/2/2008 | NCBI |
| CY037319 | A/Korea/AF05/2008(H3N2) | 19/2/2008 | NCBI |

| | | | |
|---|---|---|---|
| CY035214 | A/Siena/10/1990(H3N2) | 18/1/1990 | NCBI |
| CY038503 | A/Siena/3/1995(H3N2) | 27/1/1995 | NCBI |
| CY039399 | A/Taiwan/70002/2007(H3N2) | 2/1/2007 | NCBI |
| CY032961 | A/Yucatan/844/2003(H3N2) | 19/3/2003 | NCBI |
| CY034942 | A/HongKong/1-1-MA-12A/1968(H3N2) | 1968 | NCBI |
| CY033033 | A/HongKong/1-6/1968(H3N2) | 1968 | NCBI |
| CY033049 | A/HongKong/1-9/1968(H3N2) | 1968 | NCBI |
| CY033129 | A/HongKong/1-5-MA21-2/1968(H3N2) | 1968 | NCBI |
| CY034028 | A/HongKong/1-8-MA21-3/1968(H3N2) | 1968 | NCBI |
| CY033513 | A/HongKong/1-9-MA21-1/1968(H3N2) | 1968 | NCBI |
| CY033521 | A/HongKong/1-9-MA21-2/1968(H3N2) | 1968 | NCBI |
| CY033529 | A/HongKong/1-9-MA21-3/1968(H3N2) | 1968 | NCBI |
| CY034084 | A/Managua/17/2007(H3N2) | 21/6/2007 | NCBI |
| CY035094 | A/Managua/26/2007(H3N2) | 23/6/2007 | NCBI |
| CY032445 | A/Managua/27/2007(H3N2) | 24/6/2007 | NCBI |
| CY032533 | A/Managua/28/2007(H3N2) | 11/6/2007 | NCBI |
| CY034092 | A/Managua/5/2007(H3N2) | 13/6/2007 | NCBI |
| CY033646 | A/NymcX-161(PuertoRico/8/1934-Wisconsin/67/2005)(H3 | 2005 | NCBI |
| CY033630 | A/NymcX-149C6(PuertoRico/8/1934-Wyoming/03/2003)(H | 2003 | NCBI |
| CY031555 | A/Kentucky/UR06-0571/2007(H3N2) | 12/3/2007 | NCBI |
| CY031563 | A/California/UR06-0588/2007(H3N2) | 12/3/2007 | NCBI |
| CY037303 | A/Qatar/AF02/2008(H3N2) | 29/1/2008 | NCBI |
| CY035198 | A/Siena/3/1988(H3N2) | 5/2/1988 | NCBI |
| CY036847 | A/Siena/1/1997(H3N2) | 6/1/1997 | NCBI |
| CY035046 | A/Pennsylvania/PIT06/2008(H3N2) | 17/1/2008 | NCBI |
| CY035054 | A/Pennsylvania/PIT08/2008(H3N2) | 15/1/2008 | NCBI |
| CY035062 | A/Pennsylvania/PIT22/2008(H3N2) | 23/1/2008 | NCBI |
| CY025501 | A/Vermont/UR06-0484/2007(H3N2) | 2/3/2007 | NCBI |
| CY026195 | A/California/UR06-0589/2007(H3N2) | 13/3/2007 | NCBI |
| CY027563 | A/Texas/UR06-0358/2007(H3N2) | 20/2/2007 | NCBI |
| CY025707 | A/Illinois/UR06-0478/2007(H3N2) | 1/3/2007 | NCBI |
| CY025715 | A/NewYork/UR06-0437/2007(H3N2) | 27/2/2007 | NCBI |
| CY025723 | A/Colorado/UR06-0531/2007(H3N2) | 6/3/2007 | NCBI |
| CY027195 | A/Vermont/UR06-0469/2007(H3N2) | 1/3/2007 | NCBI |
| CY026667 | A/Colorado/UR06-0279/2007(H3N2) | 14/2/2007 | NCBI |
| CY025843 | A/NewYork/UR06-0510/2007(H3N2) | 5/3/2007 | NCBI |
| CY025851 | A/Texas/UR06-0418/2007(H3N2) | 26/2/2007 | NCBI |

| | | | |
|---|---|---|---|
| CY025859 | A/Alabama/UR06-0545/2007(H3N2) | 7/3/2007 | NCBI |
| CY026771 | A/Washington/UR06-0252/2007(H3N2) | 12/2/2007 | NCBI |
| CY026787 | A/California/UR06-0565/2007(H3N2) | 7/3/2007 | NCBI |
| CY026019 | A/Illinois/UR06-0036/2007(H3N2) | 15/1/2007 | NCBI |
| CY027123 | A/Oregon/UR06-0272/2007(H3N2) | 13/2/2007 | NCBI |
| CY026147 | A/Colorado/UR06-0023/2007(H3N2) | 8/1/2007 | NCBI |
| CY035022 | A/Brisbane/10/2007(H3N2) | 6/2/2007 | NCBI |
| CY035206 | A/Siena/4/1990(H3N2) | 8/1/1990 | NCBI |
| CY039407 | A/Taiwan/70113/2007(H3N2) | 15/1/2007 | NCBI |
| CY040066 | A/Taiwan/2361/2007(H3N2) | 12/11/2007 | NCBI |
| CY035038 | A/Pennsylvania/PIT02/2008(H3N2) | 23/1/2008 | NCBI |
| CY035182 | A/Pennsylvania/PIT31/2008(H3N2) | 8/2/2008 | NCBI |
| CY036935 | A/Pennsylvania/PIT42/2008(H3N2) | 20/2/2008 | NCBI |
| CY038543 | A/Managua/3243.01/2007(H3N2) | 24/6/2007 | NCBI |
| CY036983 | A/Florida/UR07-0101/2008(H3N2) | 26/2/2008 | NCBI |
| CY037623 | A/Florida/UR07-0146/2008(H3N2) | 10/3/2008 | NCBI |
| CY036959 | A/Ohio/UR07-0043/2008(H3N2) | 19/2/2008 | NCBI |
| CY037551 | A/Ohio/UR07-0058/2008(H3N2) | 19/2/2008 | NCBI |
| CY038607 | A/HongKong/HKU19/2004(H3N2) | 1/6/2004 | NCBI |
| CY040314 | A/HongKong/HKU38/2004(H3N2) | 23/9/2004 | NCBI |
| CY038679 | A/HongKong/HKU42/2005(H3N2) | 6/1/2005 | NCBI |
| CY038711 | A/HongKong/HKU67/2005(H3N2) | 23/5/2005 | NCBI |
| CY038727 | A/HongKong/HKU69/2005(H3N2) | 27/5/2005 | NCBI |
| CY037607 | A/NewYork/UR07-0111/2008(H3N2) | 28/2/2008 | NCBI |
| CY037519 | A/Ohio/UR07-0031/2008(H3N2) | 18/2/2008 | NCBI |
| CY037599 | A/Kansas/UR07-0110/2008(H3N2) | 28/2/2008 | NCBI |
| CY037831 | A/Kentucky/UR07-0124/2008(H3N2) | 4/3/2008 | NCBI |
| CY037791 | A/Kentucky/UR07-0028/2008(H3N2) | 15/2/2008 | NCBI |
| CY037823 | A/Kentucky/UR07-0085/2008(H3N2) | 25/2/2008 | NCBI |
| CY041466 | A/Florida/UR07-0095/2008(H3N2) | 26/2/2008 | NCBI |
| CY039415 | A/Mississippi/UR07-0024/2008(H3N2) | 14/2/2008 | NCBI |
| CY041474 | A/Florida/UR07-0154/2008(H3N2) | 12/3/2008 | NCBI |
| CY040082 | A/Taiwan/9/2004(H3N2) | 2/1/2004 | NCBI |
| CY039431 | A/Kentucky/UR07-0116/2008(H3N2) | 3/3/2008 | NCBI |
| CY035174 | A/Pennsylvania/PIT25/2008(H3N2) | 11/2/2008 | NCBI |
| CY035190 | A/Pennsylvania/PIT43/2008(H3N2) | 20/2/2008 | NCBI |
| CY036927 | A/Pennsylvania/PIT04/2008(H3N2) | 23/1/2008 | NCBI |

| | | | |
|---|---|---|---|
| CY037631 | A/NewYork/UR07-0153/2008(H3N2) | 12/3/2008 | NCBI |
| CY037511 | A/Mississippi/UR07-0025/2008(H3N2) | 14/2/2008 | NCBI |
| CY036967 | A/NewYork/UR07-0080/2008(H3N2) | 25/2/2008 | NCBI |
| CY037727 | A/Florida/UR07-0102/2008(H3N2) | 26/2/2008 | NCBI |
| CY038567 | A/HongKong/HKU2/2004(H3N2) | 13/2/2004 | NCBI |
| CY038583 | A/HongKong/HKU11/2004(H3N2) | 25/3/2004 | NCBI |
| CY038647 | A/HongKong/HKU34/2004(H3N2) | 18/8/2004 | NCBI |
| CY038735 | A/HongKong/HKU70/2005(H3N2) | 30/5/2005 | NCBI |
| CY037647 | A/Kentucky/UR07-0164/2008(H3N2) | 20/3/2008 | NCBI |
| CY037847 | A/Kansas/UR07-0136/2008(H3N2) | 5/3/2008 | NCBI |
| CY037567 | A/Kentucky/UR07-0081/2008(H3N2) | 25/2/2008 | NCBI |
| CY037479 | A/Ohio/UR07-0005/2008(H3N2) | 12/2/2008 | NCBI |
| CY037591 | A/Kentucky/UR07-0108/2008(H3N2) | 27/2/2008 | NCBI |
| CY037839 | A/Kansas/UR07-0129/2008(H3N2) | 5/3/2008 | NCBI |
| CY040098 | A/Taiwan/70120/2008(H3N2) | 15/1/2008 | NCBI |
| CY039423 | A/Kentucky/UR07-0038/2008(H3N2) | 18/2/2008 | NCBI |
| CY038511 | A/Managua/3616.01/2007(H3N2) | 7/6/2007 | NCBI |
| CY038519 | A/Managua/112.01/2007(H3N2) | 7/6/2007 | NCBI |
| CY038527 | A/Managua/4127.01/2007(H3N2) | 5/7/2007 | NCBI |
| CY037487 | A/Kansas/UR07-0007/2008(H3N2) | 12/2/2008 | NCBI |
| CY039439 | A/Managua/4348.01/2007(H3N2) | 5/6/2007 | NCBI |
| CY039487 | A/HongKong/HKU1/2004(H3N2) | 10/2/2004 | NCBI |
| CY038591 | A/HongKong/HKU12/2004(H3N2) | 29/3/2004 | NCBI |
| CY038631 | A/HongKong/HKU29/2004(H3N2) | 21/7/2004 | NCBI |
| CY038639 | A/HongKong/HKU31/2004(H3N2) | 30/7/2004 | NCBI |
| CY039495 | A/HongKong/HKU32/2004(H3N2) | 2/8/2004 | NCBI |
| CY039503 | A/HongKong/HKU50/2005(H3N2) | 11/4/2005 | NCBI |
| CY038695 | A/HongKong/HKU62/2005(H3N2) | 13/5/2005 | NCBI |
| CY038703 | A/HongKong/HKU65/2005(H3N2) | 17/5/2005 | NCBI |
| CY040338 | A/HongKong/HKU75/2005(H3N2) | 17/6/2005 | NCBI |
| CY040346 | A/HongKong/HKU76/2005(H3N2) | 21/6/2005 | NCBI |
| CY040354 | A/HongKong/HKU77/2005(H3N2) | 29/6/2005 | NCBI |
| CY038751 | A/HongKong/HKU79/2005(H3N2) | 16/7/2005 | NCBI |
| CY037711 | A/NewYork/UR07-0093/2008(H3N2) | 25/2/2008 | NCBI |
| CY037703 | A/Kentucky/UR07-0072/2008(H3N2) | 21/2/2008 | NCBI |
| CY037495 | A/Ohio/UR07-0016/2008(H3N2) | 14/2/2008 | NCBI |
| CY037671 | A/Kentucky/UR07-0015/2008(H3N2) | 14/2/2008 | NCBI |

| | | | |
|---|---|---|---|
| CY028443 | A/Kentucky/UR06-0044/2007(H3N2) | 17/1/2007 | NCBI |
| CY028475 | A/Texas/UR06-0603/2007(H3N2) | 14/3/2007 | NCBI |
| CY030197 | A/Illinois/UR06-0600/2007(H3N2) | 14/3/2007 | NCBI |
| CY032969 | A/Yucatan/ME6057/2003(H3N2) | 20/3/2003 | NCBI |
| CY033505 | A/HongKong/1-1-MA-12/1968(H3N2) | 1968 | NCBI |
| CY034974 | A/HongKong/1-1-MA-20D/1968(H3N2) | 1968 | NCBI |
| CY033081 | A/HongKong/1-1-MA21-2/1968(H3N2) | 1968 | NCBI |
| CY033089 | A/HongKong/1-1-MA21-3/1968(H3N2) | 1968 | NCBI |
| CY033097 | A/HongKong/1-2-MA21-1/1968(H3N2) | 1968 | NCBI |
| CY033105 | A/HongKong/1-2-MA21-2/1968(H3N2) | 1968 | NCBI |
| CY033553 | A/HongKong/1-11-MA21-3/1968(H3N2) | 1968 | NCBI |
| CY033561 | A/HongKong/1-12-MA21-1/1968(H3N2) | 1968 | NCBI |
| CY033569 | A/HongKong/1-12-MA21-2/1968(H3N2) | 1968 | NCBI |
| CY032525 | A/Managua/14/2007(H3N2) | 21/6/2007 | NCBI |
| CY032461 | A/Managua/15/2007(H3N2) | 21/6/2007 | NCBI |
| CY032469 | A/Managua/16/2007(H3N2) | 21/6/2007 | NCBI |
| CY032453 | A/Managua/32/2007(H3N2) | 27/6/2007 | NCBI |
| CY032517 | A/Managua/33/2007(H3N2) | 12/6/2007 | NCBI |
| CY032493 | A/Managua/34/2007(H3N2) | 23/6/2007 | NCBI |
| CY035134 | A/NymcX-171A(PuertoRico/8/1934-Brisbane/10/2007)(H3 | 2007 | NCBI |
| CY035006 | A/NymcX-157A(PuertoRico/8/1934-NewYork/55/2004)(H3 | 2004 | NCBI |
| CY035014 | A/NymcX-171(PuertoRico/8/1934-Brisbane/10/2007)(H3N | 2007 | NCBI |
| CY035142 | A/NymcX-171B(PuertoRico/8/1934-Brisbane/10/2007)(H3 | 2007 | NCBI |
| CY034406 | A/Vermont/UR06-0512/2007(H3N2) | 5/3/2007 | NCBI |
| CY036831 | A/Siena/1/1991(H3N2) | 17/12/1991 | NCBI |
| CY035222 | A/Siena/5/1991(H3N2) | 21/12/1991 | NCBI |
| CY035230 | A/Siena/6/1991(H3N2) | 24/12/1991 | NCBI |
| CY037343 | A/Genoa/1/2002(H3N2) | 2/12/2002 | NCBI |
| CY037351 | A/Genoa/14/2003(H3N2) | 24/10/2003 | NCBI |
| CY040050 | A/Taiwan/70307/2007(H3N2) | 3/2/2007 | NCBI |
| CY035150 | A/Pennsylvania/PIT05/2008(H3N2) | 22/1/2008 | NCBI |
| CY035158 | A/Pennsylvania/PIT12/2008(H3N2) | 10/1/2008 | NCBI |
| CY035166 | A/Pennsylvania/PIT20/2008(H3N2) | 25/1/2008 | NCBI |
| CY037359 | A/Managua/4902.01/2007(H3N2) | 3/8/2007 | NCBI |
| CY038495 | A/Pennsylvania/PIT39/2008(H3N2) | 21/2/2008 | NCBI |
| CY038559 | A/Managua/1867.01/2007(H3N2) | 23/6/2007 | NCBI |
| CY037559 | A/Kentucky/UR07-0068/2008(H3N2) | 21/2/2008 | NCBI |

| | | | |
|---|---|---|---|
| CY036975 | A/Ohio/UR07-0089/2008(H3N2) | 26/2/2008 | NCBI |
| CY036991 | A/Kansas/UR07-0135/2008(H3N2) | 5/3/2008 | NCBI |
| CY036999 | A/Ohio/UR07-0140/2008(H3N2) | 6/3/2008 | NCBI |
| CY037583 | A/Kentucky/UR07-0107/2008(H3N2) | 27/2/2008 | NCBI |
| CY037535 | A/Kentucky/UR07-0041/2008(H3N2) | 18/2/2008 | NCBI |
| CY040298 | A/HongKong/HKU18/2004(H3N2) | 28/5/2004 | NCBI |
| CY038615 | A/HongKong/HKU20/2004(H3N2) | 15/6/2004 | NCBI |
| CY038623 | A/HongKong/HKU22/2004(H3N2) | 29/6/2004 | NCBI |
| CY040306 | A/HongKong/HKU35/2004(H3N2) | 24/8/2004 | NCBI |
| CY038663 | A/HongKong/HKU37/2004(H3N2) | 17/9/2004 | NCBI |
| CY038671 | A/HongKong/HKU39/2004(H3N2) | 18/10/2004 | NCBI |
| CY040322 | A/HongKong/HKU52/2005(H3N2) | 16/4/2005 | NCBI |
| CY038743 | A/HongKong/HKU71/2005(H3N2) | 31/5/2005 | NCBI |
| CY037743 | A/NewYork/UR07-0133/2008(H3N2) | 6/3/2008 | NCBI |
| CY037719 | A/Mississippi/UR07-0100/2008(H3N2) | 25/2/2008 | NCBI |
| CY037543 | A/California/UR07-0053/2008(H3N2) | 19/2/2008 | NCBI |
| CY037527 | A/Kentucky/UR07-0037/2008(H3N2) | 18/2/2008 | NCBI |
| CY037799 | A/Ohio/UR07-0035/2008(H3N2) | 18/2/2008 | NCBI |
| CY037807 | A/Kansas/UR07-0047/2008(H3N2) | 19/2/2008 | NCBI |
| CY037815 | A/Kentucky/UR07-0049/2008(H3N2) | 19/2/2008 | NCBI |
| CY041458 | A/Ohio/UR07-0013/2008(H3N2) | 13/2/2008 | NCBI |
| CY043744 | A/HongKong/HKU28/2004(H3N2) | 17/7/2004 | NCBI |
| CY043768 | A/HongKong/HKU78/2005(H3N2) | 9/7/2005 | NCBI |
| CY043728 | A/Siena/3/1991(H3N2) | 19/12/1991 | NCBI |
| CY043760 | A/HongKong/HKU72/2005(H3N2) | 1/6/2005 | NCBI |
| CY043752 | A/HongKong/HKU49/2005(H3N2) | 4/4/2005 | NCBI |
| CY044397 | A/HongKong/HKU73/2005(H3N2) | 2/6/2005 | NCBI |
| CY044293 | A/HongKong/1-10-MA21-3/1968(H3N2) | 1968 | NCBI |
| CY044333 | A/Texas/AF01/2007(H3N2) | 7/1/2008 | NCBI |
| CY044357 | A/Pennsylvania/PIT34/2008(H3N2) | 6/2/2008 | NCBI |
| CY044381 | A/NewYork/UR07-0160/2008(H3N2) | 17/3/2008 | NCBI |
| CY044277 | A/HongKong/1-5-MA21-3/1968(H3N2) | 1968 | NCBI |
| CY044285 | A/HongKong/1-6-MA21-1/1968(H3N2) | 1968 | NCBI |
| CY044301 | A/X-31B(PuertoRico/8/1934-Aichi/2/1968)(H3N2) | 1968 | NCBI |
| CY044476 | A/Boston/10/2008(H3N2) | 3/3/2008 | NCBI |
| CY044492 | A/Boston/14/2007(H3N2) | 15/12/2007 | NCBI |
| CY044500 | A/Boston/19/2008(H3N2) | 28/1/2008 | NCBI |

| | | | |
|---|---|---|---|
| CY044628 | A/Boston/45/2008(H3N2) | 19/2/2008 | NCBI |
| CY044636 | A/Boston/46/2008(H3N2) | 19/2/2008 | NCBI |
| CY044644 | A/Boston/47/2008(H3N2) | 20/2/2008 | NCBI |
| CY044652 | A/Boston/48/2008(H3N2) | 20/2/2008 | NCBI |
| CY044780 | A/Boston/68/2008(H3N2) | 1/3/2008 | NCBI |
| CY044788 | A/Boston/71/2008(H3N2) | 2/3/2008 | NCBI |
| CY044820 | A/Boston/88/2008(H3N2) | 16/4/2008 | NCBI |
| CY044445 | A/Boston/3/2008(H3N2) | 24/2/2008 | NCBI |
| CY044461 | A/Boston/5/2008(H3N2) | 26/2/2008 | NCBI |
| CY044508 | A/Boston/20/2008(H3N2) | 28/1/2008 | NCBI |
| CY044596 | A/Boston/40/2008(H3N2) | 15/2/2008 | NCBI |
| CY044612 | A/Boston/43/2008(H3N2) | 19/2/2008 | NCBI |
| CY044676 | A/Boston/51/2008(H3N2) | 21/2/2008 | NCBI |
| CY044740 | A/Boston/61/2008(H3N2) | 26/2/2008 | NCBI |
| CY044756 | A/Boston/63/2008(H3N2) | 26/2/2008 | NCBI |
| CY044772 | A/Boston/66/2008(H3N2) | 27/2/2008 | NCBI |
| CY044828 | A/Boston/92/2008(H3N2) | 2008 | NCBI |
| CY044852 | A/Boston/97/2008(H3N2) | 14/3/2008 | NCBI |
| CY044429 | A/Boston/1/2008(H3N2) | 13/2/2008 | NCBI |
| CY044453 | A/Boston/4/2008(H3N2) | 25/2/2008 | NCBI |
| CY044540 | A/Boston/29/2008(H3N2) | 6/2/2008 | NCBI |
| CY044604 | A/Boston/42/2008(H3N2) | 18/2/2008 | NCBI |
| CY044620 | A/Boston/44/2008(H3N2) | 19/2/2008 | NCBI |
| CY044668 | A/Boston/50/2008(H3N2) | 21/2/2008 | NCBI |
| CY044692 | A/Boston/55/2008(H3N2) | 24/2/2008 | NCBI |
| CY044748 | A/Boston/62/2008(H3N2) | 26/2/2008 | NCBI |
| CY044844 | A/Boston/96/2008(H3N2) | 20/2/2008 | NCBI |
| CY044548 | A/Boston/31/2008(H3N2) | 7/2/2008 | NCBI |
| CY044572 | A/Boston/36/2008(H3N2) | 13/2/2008 | NCBI |
| CY044580 | A/Boston/38/2008(H3N2) | 14/2/2008 | NCBI |
| CY044588 | A/Boston/39/2008(H3N2) | 14/2/2008 | NCBI |
| CY044708 | A/Boston/57/2008(H3N2) | 24/2/2008 | NCBI |
| CY044716 | A/Boston/58/2008(H3N2) | 24/2/2008 | NCBI |
| CY044724 | A/Boston/59/2008(H3N2) | 24/2/2008 | NCBI |
| CY044732 | A/Boston/60/2008(H3N2) | 25/2/2008 | NCBI |
| CY044796 | A/Boston/78/2008(H3N2) | 11/3/2008 | NCBI |
| CY044804 | A/Boston/79/2008(H3N2) | 12/3/2008 | NCBI |

| | | | |
|---|---|---|---|
| CY044812 | A/Boston/81/2008(H3N2) | 13/3/2008 | NCBI |
| CY044469 | A/Boston/9/2008(H3N2) | 27/2/2008 | NCBI |
| CY045716 | A/HongKong/1-1-MA-20C/1968(H3N2) | 1968 | NCBI |
| CY045828 | A/X-119(PuertoRico/8/1934-Harbin/15/1992)(H3N2) | 1992 | NCBI |
| CY045836 | A/NymcX-161B(PuertoRico/8/1934-Wisconsin/67/2005)(H | 2005 | NCBI |
| CY045724 | A/HongKong/1-5-MA21-1/1968(H3N2) | 1968 | NCBI |
| CY045732 | A/HongKong/1-6-MA21-3/1968(H3N2) | 1968 | NCBI |
| CY050532 | A/NewYork/3079/2009(H3N2) | 27/4/2009 | NCBI |
| CY050540 | A/NewYork/3087/2009(H3N2) | 30/4/2009 | NCBI |
| CY050556 | A/NewYork/3103/2009(H3N2) | 28/4/2009 | NCBI |
| CY050508 | A/NewYork/3064/2009(H3N2) | 28/4/2009 | NCBI |
| CY050572 | A/NewYork/3127/2009(H3N2) | 28/4/2009 | NCBI |
| CY050500 | A/NewYork/3062/2009(H3N2) | 28/4/2009 | NCBI |
| CY050564 | A/NewYork/3104/2009(H3N2) | 29/4/2009 | NCBI |
| CY050580 | A/NewYork/3135/2009(H3N2) | 2/5/2009 | NCBI |
| CY050452 | A/NewYork/1670/2009(H3N2) | 27/4/2009 | NCBI |
| CY050468 | A/NewYork/1685/2009(H3N2) | 26/4/2009 | NCBI |
| CY050492 | A/NewYork/3060/2009(H3N2) | 27/4/2009 | NCBI |
| CY050460 | A/NewYork/1671/2009(H3N2) | 27/4/2009 | NCBI |
| CY050668 | A/NewYork/3164/2009(H3N2) | 1/5/2009 | NCBI |
| CY050676 | A/NewYork/3272/2009(H3N2) | 29/4/2009 | NCBI |
| CY050684 | A/NewYork/3273/2009(H3N2) | 29/4/2009 | NCBI |
| CY050788 | A/NewYork/3491/2009(H3N2) | 11/5/2009 | NCBI |
| CY050796 | A/NewYork/3496/2009(H3N2) | 13/5/2009 | NCBI |
| CY050588 | A/NewYork/3006/2009(H3N2) | 28/4/2009 | NCBI |
| CY050716 | A/NewYork/3297/2009(H3N2) | 27/4/2009 | NCBI |
| CY050732 | A/NewYork/3309/2009(H3N2) | 5/5/2009 | NCBI |
| CY050820 | A/NewYork/3738/2009(H3N2) | 20/5/2009 | NCBI |
| CY050836 | A/NewYork/3487/2009(H3N2) | 11/5/2009 | NCBI |
| CY050636 | A/NewYork/3146/2009(H3N2) | 1/5/2009 | NCBI |
| CY050652 | A/NewYork/3148/2009(H3N2) | 2009 | NCBI |
| CY050708 | A/NewYork/3280/2009(H3N2) | 3/5/2009 | NCBI |
| CY050724 | A/NewYork/3143/2009(H3N2) | 30/4/2009 | NCBI |
| CY050828 | A/NewYork/3752/2009(H3N2) | 1/6/2009 | NCBI |
| CY050596 | A/NewYork/3011/2009(H3N2) | 27/4/2009 | NCBI |
| CY050604 | A/NewYork/3018/2009(H3N2) | 27/4/2009 | NCBI |
| CY050620 | A/NewYork/3137/2009(H3N2) | 2/5/2009 | NCBI |

| | | | |
|---|---|---|---|
| CY050628 | A/NewYork/3139/2009(H3N2) | 29/4/2009 | NCBI |
| FJ912976 | A/Thailand/CU356/2008(H3N2) | 2008 | NCBI |
| FJ912984 | A/Thailand/CU370/2008(H3N2) | 2008 | NCBI |
| FJ912992 | A/Thailand/CU379/2008(H3N2) | 2008 | NCBI |
| GQ983548 | A/Thailand/CU-B106/2009(H3N2) | 22/6/2009 | NCBI |
| GQ902793 | A/Thailand/CU-B4/2009(H3N2) | 13/6/2009 | NCBI |
| CY055091 | A/NewYork/4165/2009(H3N2) | 12/6/2009 | NCBI |
| CY055099 | A/NewYork/3743/2009(H3N2) | 20/5/2009 | NCBI |
| CY049748 | A/Tokyo/Ut-Sk-1/2007(H3N2) | 2007 | NCBI |
| GU271974 | A/Thailand/CU-B1672/2009(H3N2) | 30/10/2009 | NCBI |
| GU271982 | A/Thailand/CU-B1697/2009(H3N2) | 10/11/2009 | NCBI |
| GU271990 | A/Thailand/CU-H16/2009(H3N2) | 17/6/2009 | NCBI |
| GU907114 | A/Nanjing/1/2009(H3N2) | 10/4/2009 | NCBI |
| EU283414 | A/Hiroshima/52/2005(H3N2) | 2005 | NCBI |
| EU625363 | A/Thailand/CU-1101/2008(H3N2) | 2008 | NCBI |
| EU625364 | A/Thailand/CU-1102/2008(H3N2) | 2008 | NCBI |
| EU625365 | A/Thailand/CU-1103/2008(H3N2) | 2008 | NCBI |
| EU021274 | A/Thailand/CU228/2006(H3N2) | 2006 | NCBI |
| EU021282 | A/Thailand/CU231/2006(H3N2) | 2006 | NCBI |
| EU021272 | A/Thailand/CU280/2007(H3N2) | 2007 | NCBI |
| EU021276 | A/Thailand/CU282/2007(H3N2) | 2007 | NCBI |
| EU021268 | A/Thailand/CU46/2006(H3N2) | 2006 | NCBI |
| CY058780 | A/NewYork/3527/2009(H3N2) | 18/5/2009 | NCBI |
| CY058756 | A/NewYork/3316/2009(H3N2) | 1/5/2009 | NCBI |
| CY058764 | A/NewYork/3360/2009(H3N2) | 6/5/2009 | NCBI |
| CY058772 | A/NewYork/3494/2009(H3N2) | 13/5/2009 | NCBI |
| CY058796 | A/NewYork/3626/2009(H3N2) | 19/5/2009 | NCBI |
| CY058804 | A/NewYork/4255/2009(H3N2) | 24/5/2009 | NCBI |
| CY061898 | A/Australia/19/2009(H3N2) | 21/7/2009 | NCBI |
| CY061890 | A/Australia/2/2009(H3N2) | 16/7/2009 | NCBI |
| CY064815 | A/California/VRDL136/2009(H3N2) | 26/4/2009 | NCBI |
| CY064887 | A/California/VRDL158/2009(H3N2) | 27/4/2009 | NCBI |
| CY064847 | A/California/VRDL143/2009(H3N2) | 27/4/2009 | NCBI |
| CY064855 | A/California/VRDL144/2009(H3N2) | 27/4/2009 | NCBI |
| CY064863 | A/California/VRDL145/2009(H3N2) | 27/4/2009 | NCBI |
| CY064879 | A/California/VRDL147/2009(H3N2) | 28/4/2009 | NCBI |
| CY064823 | A/California/VRDL138/2009(H3N2) | 27/4/2009 | NCBI |

| | | | |
|---|---|---|---|
| CY065976 | A/Philippines/2-MA/1982(H3N2) | 1/6/1982 | NCBI |
| CY065968 | A/Philippines/2/1982(H3N2) | 1/6/1982 | NCBI |
| CY066511 | A/Australia/18/2009(H3N2) | 21/7/2009 | NCBI |
| CY066519 | A/California/VRDL156/2009(H3N2) | 28/4/2009 | NCBI |
| CY067197 | A/California/VRDL241/2009(H3N2) | 29/4/2009 | NCBI |
| CY067205 | A/California/VRDL291/2009(H3N2) | 1/5/2009 | NCBI |
| CY067253 | A/California/VRDL316/2009(H3N2) | 4/5/2009 | NCBI |
| CY067213 | A/California/VRDL296/2009(H3N2) | 30/4/2009 | NCBI |
| CY067221 | A/California/VRDL302/2009(H3N2) | 28/4/2009 | NCBI |
| CY067229 | A/California/VRDL304/2009(H3N2) | 7/5/2009 | NCBI |
| CY067237 | A/California/VRDL307/2009(H3N2) | 2/5/2009 | NCBI |
| CY067245 | A/California/VRDL314/2009(H3N2) | 1/5/2009 | NCBI |
| CY067921 | A/California/VRDL176/2009(H3N2) | 27/4/2009 | NCBI |
| CY067937 | A/California/VRDL286/2009(H3N2) | 30/4/2009 | NCBI |
| CY067945 | A/California/VRDL287/2009(H3N2) | 4/5/2009 | NCBI |
| CY067953 | A/California/VRDL288/2009(H3N2) | 30/4/2009 | NCBI |
| CY067929 | A/California/VRDL177/2009(H3N2) | 28/4/2009 | NCBI |
| CY067985 | A/California/VRDL342/2009(H3N2) | 20/5/2009 | NCBI |
| CY068001 | A/California/VRDL344/2009(H3N2) | 19/5/2009 | NCBI |
| CY068041 | A/California/VRDL354/2009(H3N2) | 26/5/2009 | NCBI |
| CY067969 | A/California/VRDL339/2009(H3N2) | 4/5/2009 | NCBI |
| CY067977 | A/California/VRDL340/2009(H3N2) | 4/5/2009 | NCBI |
| CY067993 | A/California/VRDL343/2009(H3N2) | 11/5/2009 | NCBI |
| CY068009 | A/California/VRDL345/2009(H3N2) | 26/5/2009 | NCBI |
| CY068049 | A/California/VRDL355/2009(H3N2) | 11/5/2009 | NCBI |
| CY067961 | A/California/VRDL328/2009(H3N2) | 9/5/2009 | NCBI |
| CY068017 | A/California/VRDL349/2009(H3N2) | 26/5/2009 | NCBI |
| CY068025 | A/California/VRDL351/2009(H3N2) | 23/5/2009 | NCBI |
| CY068033 | A/California/VRDL352/2009(H3N2) | 11/5/2009 | NCBI |
| CY068081 | A/California/VRDL362/2009(H3N2) | 11/8/2009 | NCBI |
| CY068089 | A/California/VRDL367/2009(H3N2) | 25/4/2009 | NCBI |
| CY068129 | A/California/VRDL396/2009(H3N2) | 28/4/2009 | NCBI |
| CY068057 | A/California/VRDL357/2009(H3N2) | 5/6/2009 | NCBI |
| CY068065 | A/California/VRDL360/2009(H3N2) | 3/6/2009 | NCBI |
| CY068073 | A/California/VRDL361/2009(H3N2) | 23/6/2009 | NCBI |
| CY068097 | A/California/VRDL368/2009(H3N2) | 27/4/2009 | NCBI |
| CY068105 | A/California/VRDL373/2009(H3N2) | 27/4/2009 | NCBI |

| | | | |
|---|---|---|---|
| CY068113 | A/California/VRDL374/2009(H3N2) | 25/4/2009 | NCBI |
| CY068121 | A/California/VRDL375/2009(H3N2) | 27/4/2009 | NCBI |
| CY068153 | A/California/VRDL161/2009(H3N2) | 28/4/2009 | NCBI |
| CY068161 | A/California/VRDL164/2009(H3N2) | 28/4/2009 | NCBI |
| CY068217 | A/California/VRDL173/2009(H3N2) | 28/4/2009 | NCBI |
| CY068257 | A/California/VRDL230/2009(H3N2) | 29/4/2009 | NCBI |
| CY068273 | A/California/VRDL233/2009(H3N2) | 29/4/2009 | NCBI |
| CY068305 | A/California/VRDL250/2009(H3N2) | 29/4/2009 | NCBI |
| CY068361 | A/California/VRDL265/2009(H3N2) | 1/4/2009 | NCBI |
| CY068169 | A/California/VRDL165/2009(H3N2) | 29/4/2009 | NCBI |
| CY068209 | A/California/VRDL172/2009(H3N2) | 28/4/2009 | NCBI |
| CY068225 | A/California/VRDL188/2009(H3N2) | 29/4/2009 | NCBI |
| CY068249 | A/California/VRDL216/2009(H3N2) | 28/4/2009 | NCBI |
| CY068265 | A/California/VRDL231/2009(H3N2) | 30/4/2009 | NCBI |
| CY068297 | A/California/VRDL248/2009(H3N2) | 27/4/2009 | NCBI |
| CY068313 | A/California/VRDL253/2009(H3N2) | 30/4/2009 | NCBI |
| CY068353 | A/California/VRDL263/2009(H3N2) | 30/4/2009 | NCBI |
| CY068177 | A/California/VRDL166/2009(H3N2) | 28/4/2009 | NCBI |
| CY068185 | A/California/VRDL168/2009(H3N2) | 29/4/2009 | NCBI |
| CY068193 | A/California/VRDL169/2009(H3N2) | 28/4/2009 | NCBI |
| CY068201 | A/California/VRDL170/2009(H3N2) | 28/4/2009 | NCBI |
| CY068281 | A/California/VRDL243/2009(H3N2) | 29/4/2009 | NCBI |
| CY068289 | A/California/VRDL247/2009(H3N2) | 29/4/2009 | NCBI |
| CY068137 | A/California/VRDL159/2009(H3N2) | 28/4/2009 | NCBI |
| CY068145 | A/California/VRDL160/2009(H3N2) | 28/4/2009 | NCBI |
| CY068233 | A/California/VRDL211/2009(H3N2) | 28/4/2009 | NCBI |
| CY068241 | A/California/VRDL212/2009(H3N2) | 29/4/2009 | NCBI |
| CY068321 | A/California/VRDL259/2009(H3N2) | 29/4/2009 | NCBI |
| CY068329 | A/California/VRDL260/2009(H3N2) | 1/5/2009 | NCBI |
| CY068337 | A/California/VRDL261/2009(H3N2) | 2/5/2009 | NCBI |
| CY068409 | A/California/VRDL277/2009(H3N2) | 30/4/2009 | NCBI |
| CY068425 | A/California/VRDL279/2009(H3N2) | 30/4/2009 | NCBI |
| CY068449 | A/California/VRDL294/2009(H3N2) | 30/4/2009 | NCBI |
| CY068473 | A/California/VRDL309/2009(H3N2) | 2/5/2009 | NCBI |
| CY068489 | A/California/VRDL312/2009(H3N2) | 1/5/2009 | NCBI |
| CY068529 | A/California/VRDL319/2009(H3N2) | 6/5/2009 | NCBI |
| CY068545 | A/California/VRDL322/2009(H3N2) | 4/5/2009 | NCBI |

| | | | |
|---|---|---|---|
| CY068585 | A/California/VRDL332/2009(H3N2) | 11/5/2009 | NCBI |
| CY068601 | A/California/VRDL334/2009(H3N2) | 18/5/2009 | NCBI |
| CY068377 | A/California/VRDL268/2009(H3N2) | 30/4/2009 | NCBI |
| CY068385 | A/California/VRDL271/2009(H3N2) | 30/4/2009 | NCBI |
| CY068393 | A/California/VRDL272/2009(H3N2) | 30/4/2009 | NCBI |
| CY068457 | A/California/VRDL305/2009(H3N2) | 1/5/2009 | NCBI |
| CY068465 | A/California/VRDL308/2009(H3N2) | 29/4/2009 | NCBI |
| CY068561 | A/California/VRDL327/2009(H3N2) | 11/5/2009 | NCBI |
| CY068569 | A/California/VRDL329/2009(H3N2) | 9/5/2009 | NCBI |
| CY068401 | A/California/VRDL274/2009(H3N2) | 30/4/2009 | NCBI |
| CY068417 | A/California/VRDL278/2009(H3N2) | 30/4/2009 | NCBI |
| CY068433 | A/California/VRDL285/2009(H3N2) | 30/4/2009 | NCBI |
| CY068441 | A/California/VRDL293/2009(H3N2) | 30/4/2009 | NCBI |
| CY068481 | A/California/VRDL310/2009(H3N2) | 30/4/2009 | NCBI |
| CY068497 | A/California/VRDL313/2009(H3N2) | 1/5/2009 | NCBI |
| CY068521 | A/California/VRDL318/2009(H3N2) | 6/5/2009 | NCBI |
| CY068537 | A/California/VRDL321/2009(H3N2) | 4/5/2009 | NCBI |
| CY068553 | A/California/VRDL325/2009(H3N2) | 3/5/2009 | NCBI |
| CY068577 | A/California/VRDL331/2009(H3N2) | 11/5/2009 | NCBI |
| CY068593 | A/California/VRDL333/2009(H3N2) | 6/5/2009 | NCBI |
| CY068633 | A/California/VRDL394/2009(H3N2) | 28/4/2009 | NCBI |
| CY068505 | A/California/VRDL315/2009(H3N2) | 1/5/2009 | NCBI |
| CY068513 | A/California/VRDL317/2009(H3N2) | 6/5/2009 | NCBI |
| CY068609 | A/California/VRDL335/2009(H3N2) | 3/5/2009 | NCBI |
| CY068617 | A/California/VRDL336/2009(H3N2) | 5/5/2009 | NCBI |
| CY068625 | A/California/VRDL338/2009(H3N2) | 18/5/2009 | NCBI |
| CY068694 | A/California/VRDL183/2009(H3N2) | 27/4/2009 | NCBI |
| CY068710 | A/California/VRDL185/2009(H3N2) | 29/4/2009 | NCBI |
| CY068678 | A/California/VRDL178/2009(H3N2) | 28/4/2009 | NCBI |
| CY068686 | A/California/VRDL181/2009(H3N2) | 28/4/2009 | NCBI |
| CY068702 | A/California/VRDL184/2009(H3N2) | 28/4/2009 | NCBI |
| CY068718 | A/California/VRDL187/2009(H3N2) | 28/4/2009 | NCBI |
| CY068742 | A/California/VRDL199/2009(H3N2) | 28/4/2009 | NCBI |
| CY068758 | A/California/VRDL203/2009(H3N2) | 28/4/2009 | NCBI |
| CY068774 | A/California/VRDL205/2009(H3N2) | 29/4/2009 | NCBI |
| CY068822 | A/California/VRDL381/2009(H3N2) | 27/4/2009 | NCBI |
| CY068838 | A/California/VRDL384/2009(H3N2) | 25/4/2009 | NCBI |

| | | | |
|---|---|---|---|
| CY068862 | A/California/VRDL391/2009(H3N2) | 28/4/2009 | NCBI |
| CY068878 | A/California/VRDL393/2009(H3N2) | 29/4/2009 | NCBI |
| CY068726 | A/California/VRDL196/2009(H3N2) | 29/4/2009 | NCBI |
| CY068734 | A/California/VRDL198/2009(H3N2) | 28/4/2009 | NCBI |
| CY068814 | A/California/VRDL269/2009(H3N2) | 30/4/2009 | NCBI |
| CY068782 | A/California/VRDL207/2009(H3N2) | 27/4/2009 | NCBI |
| CY068790 | A/California/VRDL208/2009(H3N2) | 29/4/2009 | NCBI |
| CY068798 | A/California/VRDL210/2009(H3N2) | 28/4/2009 | NCBI |
| CY068846 | A/California/VRDL387/2009(H3N2) | 27/4/2009 | NCBI |
| CY068854 | A/California/VRDL390/2009(H3N2) | 26/4/2009 | NCBI |
| CY068750 | A/California/VRDL201/2009(H3N2) | 29/4/2009 | NCBI |
| CY068766 | A/California/VRDL204/2009(H3N2) | 28/4/2009 | NCBI |
| CY068806 | A/California/VRDL246/2009(H3N2) | 29/4/2009 | NCBI |
| CY068830 | A/California/VRDL383/2009(H3N2) | 26/4/2009 | NCBI |
| CY068870 | A/California/VRDL392/2009(H3N2) | 28/4/2009 | NCBI |
| CY069317 | A/Japan/WRAIR1037P/2009(H3N2) | 9/1/2009 | NCBI |
| CY069325 | A/Korea/WRAIR1038P/2009(H3N2) | 9/4/2009 | NCBI |
| CY069429 | A/Japan/WRAIR1140P/2009(H3N2) | 9/2/2009 | NCBI |
| CY070919 | A/California/VRDL330/2009(H3N2) | 9/5/2009 | NCBI |
| CY070967 | A/NewYork/20342/2010(H3N2) | 16/6/2010 | NCBI |
| CY070943 | A/Managua/1807.03/2010(H3N2) | 7/6/2010 | NCBI |
| CY070959 | A/Managua/2867.01/2010(H3N2) | 10/6/2010 | NCBI |
| CY070951 | A/Managua/1181.03/2010(H3N2) | 3/6/2010 | NCBI |
| CY070927 | A/Managua/5806.01/2010(H3N2) | 8/6/2010 | NCBI |
| CY070935 | A/Managua/4131.04/2010(H3N2) | 6/6/2010 | NCBI |
| HM641169 | A/HongKong/1-6-MA21-1/1968(H3N2) | 1968 | NCBI |
| HM641171 | A/HongKong/1-5-MA21-1/1968(H3N2) | 1968 | NCBI |
| HM641170 | A/HongKong/1-5-MA21-3/1968(H3N2) | 1968 | NCBI |
| HM641174 | A/HongKong/1-2-MA21-2/1968(H3N2) | 1968 | NCBI |
| HM641173 | A/HongKong/1-2-MA21-3/1968(H3N2) | 1968 | NCBI |
| HM641172 | A/HongKong/1-4-MA21-1/1968(H3N2) | 1968 | NCBI |
| HM641176 | A/HongKong/1-10-MA21-2/1968(H3N2) | 1968 | NCBI |
| HM641177 | A/HongKong/1-10-MA21-3/1968(H3N2) | 1968 | NCBI |
| HM641168 | A/HongKong/1-6-MA21-2/1968(H3N2) | 1968 | NCBI |
| HM641178 | A/HongKong/MA/1968(H3N2) | 1968 | NCBI |
| CY072214 | A/NewYork/20343/2010(H3N2) | 16/6/2010 | NCBI |
| CY072190 | A/California/VRDL347/2009(H3N2) | 23/5/2009 | NCBI |

| | | | |
|---|---|---|---|
| CY072198 | A/California/VRDL385/2009(H3N2) | 26/4/2009 | NCBI |
| CY072206 | A/California/VRDL386/2009(H3N2) | 26/4/2009 | NCBI |
| CY073757 | A/California/VRDL363/2009(H3N2) | 2009 | NCBI |
| CY073869 | A/Managua/1422.03/2010(H3N2) | 4/6/2010 | NCBI |
| CY074723 | A/Managua/905.02/2010(H3N2) | 2/6/2010 | NCBI |
| CY074739 | A/Managua/2912.04/2010(H3N2) | 13/6/2010 | NCBI |
| CY074779 | A/Managua/1535.03/2010(H3N2) | 8/6/2010 | NCBI |
| CY074787 | A/Managua/90.01/2010(H3N2) | 12/6/2010 | NCBI |
| CY074715 | A/Managua/5871.02/2010(H3N2) | 8/6/2010 | NCBI |
| CY074731 | A/Managua/194.01/2010(H3N2) | 8/6/2010 | NCBI |
| CY074795 | A/Managua/1784.03/2010(H3N2) | 3/6/2010 | NCBI |
| CY074747 | A/Managua/1167.03/2010(H3N2) | 7/6/2010 | NCBI |
| CY074755 | A/Managua/80.1/2010(H3N2) | 7/6/2010 | NCBI |
| CY074763 | A/Managua/1856.03/2010(H3N2) | 9/6/2010 | NCBI |
| CY074771 | A/Managua/3244.02/2010(H3N2) | 6/6/2010 | NCBI |
| CY074675 | A/Managua/395.02/2010(H3N2) | 3/6/2010 | NCBI |
| CY074683 | A/Managua/5906.01/2010(H3N2) | 4/6/2010 | NCBI |
| CY074691 | A/Managua/4763.03/2010(H3N2) | 6/6/2010 | NCBI |
| CY074699 | A/Managua/3192.01/2010(H3N2) | 5/6/2010 | NCBI |
| CY074707 | A/Managua/4009.02/2010(H3N2) | 14/6/2010 | NCBI |
| CY074803 | A/Managua/3715.02/2010(H3N2) | 7/6/2010 | NCBI |
| CY074811 | A/Managua/5674.05/2010(H3N2) | 5/6/2010 | NCBI |
| CY074819 | A/Managua/4218.02/2010(H3N2) | 8/6/2010 | NCBI |
| CY074843 | A/Managua/1166.03/2010(H3N2) | 6/6/2010 | NCBI |
| CY074899 | A/Managua/4203.02/2010(H3N2) | 7/6/2010 | NCBI |
| CY074859 | A/Managua/506.02/2010(H3N2) | 5/6/2010 | NCBI |
| CY074867 | A/Managua/4134.04/2010(H3N2) | 7/6/2010 | NCBI |
| CY074875 | A/Managua/3527.02/2010(H3N2) | 5/6/2010 | NCBI |
| CY074883 | A/Managua/5863.02/2010(H3N2) | 5/6/2010 | NCBI |
| CY074835 | A/Managua/410.02/2010(H3N2) | 8/6/2010 | NCBI |
| CY074851 | A/Managua/5543.01/2010(H3N2) | 4/6/2010 | NCBI |
| CY074891 | A/Managua/3570.04/2010(H3N2) | 6/6/2010 | NCBI |
| CY074827 | A/Managua/3433.02/2010(H3N2) | 4/6/2010 | NCBI |
| CY074923 | A/Managua/4456.02/2010(H3N2) | 6/6/2010 | NCBI |
| CY074931 | A/Managua/305-10/2010(H3N2) | 19/3/2010 | NCBI |
| CY074907 | A/Managua/5259.02/2010(H3N2) | 8/6/2010 | NCBI |
| CY074915 | A/Managua/2614.02/2010(H3N2) | 7/6/2010 | NCBI |

| | | | |
|---|---|---|---|
| CY074942 | A/Thailand/CU-H1071/2009(H3N2) | 10/12/2009 | NCBI |
| CY074958 | A/Thailand/CU-H1443/2010(H3N2) | 4/2/2010 | NCBI |
| CY074950 | A/Thailand/CU-H1285/2010(H3N2) | 21/1/2010 | NCBI |
| CY077425 | A/California/VRDL356/2009(H3N2) | 2/6/2009 | NCBI |
| HQ664924 | A/Nanjing/1654/2010(H3N2) | 10/9/2010 | NCBI |
| HQ664914 | A/Nanjing/1655/2010(H3N2) | 10/9/2010 | NCBI |
| HQ664931 | A/Nanjing/1663/2010(H3N2) | 10/9/2010 | NCBI |
| HQ166052 | A/Netherlands/213/2003(H3N2) | 20/4/2003 | NCBI |
| CY080483 | A/NewYork/3644/2009(H3N2) | 23/5/2009 | NCBI |
| CY080499 | A/HongKong/1-2-MA21-3/1968(H3N2) | 1968 | NCBI |
| CY080459 | A/NewYork/3421/2009(H3N2) | 4/5/2009 | NCBI |
| CY080491 | A/NewYork/3932/2009(H3N2) | 4/6/2009 | NCBI |
| CY080467 | A/NewYork/3470/2009(H3N2) | 16/5/2009 | NCBI |
| CY080475 | A/NewYork/3519/2009(H3N2) | 15/5/2009 | NCBI |
| CY080563 | A/Australia/55/2009(H3N2) | 30/7/2009 | NCBI |
| CY080515 | A/HongKong/1-4-MA21-1/1968(H3N2) | 1968 | NCBI |
| CY080547 | A/Australia/30/2009(H3N2) | 23/7/2009 | NCBI |
| CY080523 | A/HongKong/1-10-MA21-1/1968(H3N2) | 1968 | NCBI |
| CY080539 | A/Australia/22/2009(H3N2) | 21/7/2009 | NCBI |
| CY080555 | A/Australia/34/2009(H3N2) | 24/7/2009 | NCBI |
| CY081470 | A/Cambodia/NHRCC00002/2009(H3N2) | 14/10/2009 | NCBI |
| CY081462 | A/Cambodia/NHRCC00001/2009(H3N2) | 14/10/2009 | NCBI |
| CY081478 | A/Cambodia/NHRCC00006/2009(H3N2) | 14/10/2009 | NCBI |
| CY084334 | A/NewYork/6630/2009(H3N2) | 13/11/2009 | NCBI |
| CY084385 | A/NewYork/3318/2009(H3N2) | 28/4/2009 | NCBI |
| CY084401 | A/NewYork/3904/2009(H3N2) | 4/6/2009 | NCBI |
| CY084393 | A/NewYork/3417/2009(H3N2) | 6/5/2009 | NCBI |
| CY084409 | A/NewYork/3918/2009(H3N2) | 5/6/2009 | NCBI |
| CY084417 | A/NewYork/4362/2009(H3N2) | 23/6/2009 | NCBI |
| CY088774 | A/Managua/511.03/2010(H3N2) | 2/6/2010 | NCBI |
| CY088782 | A/Managua/5664.02/2010(H3N2) | 3/6/2010 | NCBI |
| CY088790 | A/Managua/5427.01/2010(H3N2) | 2/6/2010 | NCBI |
| CY088907 | A/Managua/3039.01/2010(H3N2) | 1/6/2010 | NCBI |
| CY088915 | A/Managua/2145.01/2010(H3N2) | 2/6/2010 | NCBI |
| CY088923 | A/Managua/5801.02/2010(H3N2) | 2/6/2010 | NCBI |
| CY088931 | A/Managua/4747.04/2010(H3N2) | 2/6/2010 | NCBI |
| CY088939 | A/Managua/38.01/2010(H3N2) | 2/6/2010 | NCBI |

| | | | |
|---|---|---|---|
| CY088995 | A/Managua/1685.01/2010(H3N2) | 10/6/2010 | NCBI |
| CY089003 | A/Managua/4076.04/2010(H3N2) | 10/6/2010 | NCBI |
| CY089011 | A/Managua/4604.04/2010(H3N2) | 11/6/2010 | NCBI |
| CY089019 | A/Managua/5870.02/2010(H3N2) | 11/6/2010 | NCBI |
| CY088883 | A/Managua/4355.03/2010(H3N2) | 30/5/2010 | NCBI |
| CY088899 | A/Managua/1155.01/2010(H3N2) | 1/6/2010 | NCBI |
| CY088963 | A/Managua/5898.02/2010(H3N2) | 6/6/2010 | NCBI |
| CY088979 | A/Managua/3408.02/2010(H3N2) | 7/6/2010 | NCBI |
| CY088875 | A/Managua/5099.03/2010(H3N2) | 28/5/2010 | NCBI |
| CY088891 | A/Managua/1260.02/2010(H3N2) | 1/6/2010 | NCBI |
| CY088947 | A/Managua/5444.02/2010(H3N2) | 1/6/2010 | NCBI |
| CY088971 | A/Managua/3209.01/2010(H3N2) | 7/6/2010 | NCBI |
| CY088987 | A/Managua/3659.02/2010(H3N2) | 9/6/2010 | NCBI |
| CY089027 | A/Managua/4855.03/2010(H3N2) | 3/6/2010 | NCBI |
| CY088843 | A/Managua/3345.02/2010(H3N2) | 17/5/2010 | NCBI |
| CY088851 | A/Managua/2654.01/2010(H3N2) | 21/5/2010 | NCBI |
| CY088859 | A/Managua/5271.04/2010(H3N2) | 28/5/2010 | NCBI |
| CY088867 | A/Managua/3704.06/2010(H3N2) | 28/5/2010 | NCBI |
| CY088955 | A/Managua/3622.02/2010(H3N2) | 3/6/2010 | NCBI |
| CY089540 | A/California/VRDL359/2009(H3N2) | 8/6/2009 | NCBI |
| CY089629 | A/NewYork/3750/2009(H3N2) | 21/5/2009 | NCBI |
| CY089733 | A/Boston/40/2009(H3N2) | 27/2/2009 | NCBI |
| CY089741 | A/Boston/73/2009(H3N2) | 1/4/2009 | NCBI |
| CY089749 | A/Boston/75/2009(H3N2) | 16/1/2009 | NCBI |
| CY089773 | A/Boston/99/2009(H3N2) | 27/4/2009 | NCBI |
| CY089765 | A/Boston/97/2009(H3N2) | 11/5/2009 | NCBI |
| CY090877 | A/Sydney/DD2-02/2010(H3N2) | 4/9/2010 | NCBI |
| CY090885 | A/Georgia/NHRC0001/1999(H3N2) | 12/10/1999 | NCBI |
| CY090901 | A/SouthCarolina/NHRC0001/1999(H3N2) | 29/11/1999 | NCBI |
| CY090949 | A/SouthCarolina/NHRC0002/2003(H3N2) | 13/11/2003 | NCBI |
| CY090965 | A/NewJersey/NHRC0001/2003(H3N2) | 26/11/2003 | NCBI |
| CY090869 | A/Sydney/DD2-01/2010(H3N2) | 13/9/2010 | NCBI |
| CY090893 | A/Illinois/NHRC0001/1999(H3N2) | 1/11/1999 | NCBI |
| CY090909 | A/Missouri/NHRC0001/2000(H3N2) | 7/2/2000 | NCBI |
| CY090941 | A/California/NHRC0001/2003(H3N2) | 13/11/2003 | NCBI |
| CY090957 | A/Texas/NHRC0001/2003(H3N2) | 20/11/2003 | NCBI |
| CY090917 | A/Illinois/NHRC0001/2002(H3N2) | 13/2/2002 | NCBI |

| | | | |
|---|---|---|---|
| CY090925 | A/SouthCarolina/NHRC0001/2003(H3N2) | 21/10/2003 | NCBI |
| CY090933 | A/Missouri/NHRC0001/2003(H3N2) | 6/11/2003 | NCBI |
| CY090973 | A/Georgia/NHRC0001/2004(H3N2) | 20/1/2004 | NCBI |
| CY090981 | A/Missouri/NHRC0001/2005(H3N2) | 12/1/2005 | NCBI |
| CY090989 | A/SouthCarolina/NHRC0001/2005(H3N2) | 11/2/2005 | NCBI |
| CY090997 | A/SouthCarolina/NHRC0002/2005(H3N2) | 10/3/2005 | NCBI |
| CY091037 | A/Texas/NHRC0001/2006(H3N2) | 13/2/2006 | NCBI |
| CY091045 | A/Illinois/NHRC0001/2006(H3N2) | 15/2/2006 | NCBI |
| CY091053 | A/California/NHRC0001/2006(H3N2) | 21/2/2006 | NCBI |
| CY091061 | A/SouthCarolina/NHRC0001/2006(H3N2) | 28/3/2006 | NCBI |
| CY091005 | A/Illinois/NHRC0001/2005(H3N2) | 6/4/2005 | NCBI |
| CY091021 | A/Texas/NHRC0002/2005(H3N2) | 19/12/2005 | NCBI |
| CY091013 | A/California/NHRC0001/2005(H3N2) | 30/11/2005 | NCBI |
| CY091029 | A/Illinois/NHRC0002/2005(H3N2) | 23/12/2005 | NCBI |
| CY091069 | A/Texas/NHRC0002/2006(H3N2) | 4/4/2006 | NCBI |
| CY091077 | A/Missouri/NHRC0001/2006(H3N2) | 5/4/2006 | NCBI |
| CY091093 | A/Missouri/NHRC0001/2002(H3N2) | 9/1/2002 | NCBI |
| CY091133 | A/Texas/NHRC0001/2005(H3N2) | 16/2/2005 | NCBI |
| CY091085 | A/Illinois/NHRC0002/2006(H3N2) | 6/4/2006 | NCBI |
| CY091141 | A/Georgia/NHRC0001/2006(H3N2) | 9/1/2006 | NCBI |
| CY091101 | A/Texas/NHRC0001/2002(H3N2) | 22/1/2002 | NCBI |
| CY091109 | A/Georgia/NHRC0001/2002(H3N2) | 2/2/2002 | NCBI |
| CY091117 | A/Missouri/NHRC0002/2004(H3N2) | 9/1/2004 | NCBI |
| CY091125 | A/Georgia/NHRC0001/2005(H3N2) | 25/1/2005 | NCBI |
| CY091149 | A/Singapore/NHRC0001/2003(H3N2) | 28/9/2003 | NCBI |
| CY091157 | A/Singapore/NHRC0005/2003(H3N2) | 3/10/2003 | NCBI |
| CY091165 | A/Singapore/NHRC0011/2003(H3N2) | 11/10/2003 | NCBI |
| CY091173 | A/California/NHRC0003/2003(H3N2) | 17/12/2003 | NCBI |
| CY091181 | A/Australia/NHRC0001/2003(H3N2) | 1/7/2003 | NCBI |
| CY091285 | A/Australia/NHRC0006/2003(H3N2) | 7/7/2003 | NCBI |
| CY091293 | A/Australia/NHRC0003/2003(H3N2) | 1/7/2003 | NCBI |
| CY091301 | A/Australia/NHRC0002/2003(H3N2) | 1/7/2003 | NCBI |
| CY091309 | A/HongKong/NHRC0001/2005(H3N2) | 15/6/2005 | NCBI |
| CY091405 | A/Australia/NHRC0003/2005(H3N2) | 15/4/2005 | NCBI |
| CY091413 | A/Australia/NHRC0002/2005(H3N2) | 15/4/2005 | NCBI |
| CY091421 | A/Australia/NHRC0001/2005(H3N2) | 13/4/2005 | NCBI |
| CY091429 | A/California/NHRC0002/2005(H3N2) | 7/2/2005 | NCBI |

| | | | |
|---|---|---|---|
| CY091541 | A/California/NHRC0008/2005(H3N2) | 5/12/2005 | NCBI |
| CY091549 | A/California/NHRC0009/2005(H3N2) | 7/2/2005 | NCBI |
| CY091189 | A/Australia/NHRC0004/2003(H3N2) | 2/7/2003 | NCBI |
| CY091205 | A/Australia/NHRC0010/2003(H3N2) | 10/7/2003 | NCBI |
| CY091261 | A/Singapore/NHRC0007/2003(H3N2) | 8/10/2003 | NCBI |
| CY091277 | A/Australia/NHRC0008/2003(H3N2) | 8/7/2003 | NCBI |
| CY091325 | A/Australia/NHRC0014/2005(H3N2) | 17/6/2005 | NCBI |
| CY091341 | A/Australia/NHRC0012/2005(H3N2) | 13/6/2005 | NCBI |
| CY091373 | A/Australia/NHRC0007/2005(H3N2) | 13/6/2005 | NCBI |
| CY091389 | A/Australia/NHRC0005/2005(H3N2) | 16/4/2005 | NCBI |
| CY091445 | A/Malaysia/NHRC0002/2004(H3N2) | 25/9/2004 | NCBI |
| CY091461 | A/California/NHRC0006/2003(H3N2) | 10/12/2003 | NCBI |
| CY091517 | A/California/NHRC0005/2005(H3N2) | 29/11/2005 | NCBI |
| CY091533 | A/California/NHRC0007/2005(H3N2) | 15/11/2005 | NCBI |
| CY091221 | A/California/NHRC0002/2003(H3N2) | 16/12/2003 | NCBI |
| CY091229 | A/Singapore/NHRC0012/2003(H3N2) | 13/10/2003 | NCBI |
| CY091237 | A/Singapore/NHRC0010/2003(H3N2) | 11/10/2003 | NCBI |
| CY091245 | A/Singapore/NHRC0009/2003(H3N2) | 9/10/2003 | NCBI |
| CY091349 | A/Australia/NHRC0011/2005(H3N2) | 13/6/2005 | NCBI |
| CY091357 | A/Australia/NHRC0009/2005(H3N2) | 13/6/2005 | NCBI |
| CY091365 | A/Australia/NHRC0008/2005(H3N2) | 13/6/2005 | NCBI |
| CY091469 | A/Singapore/NHRC0006/2003(H3N2) | 7/10/2003 | NCBI |
| CY091477 | A/Singapore/NHRC0004/2003(H3N2) | 1/10/2003 | NCBI |
| CY091485 | A/Singapore/NHRC0003/2003(H3N2) | 30/9/2003 | NCBI |
| CY091493 | A/Singapore/NHRC0002/2003(H3N2) | 29/9/2003 | NCBI |
| CY091501 | A/California/NHRC0003/2005(H3N2) | 25/10/2005 | NCBI |
| CY091565 | A/SouthCarolina/NHRC0001/2011(H3N2) | 28/1/2011 | NCBI |
| CY091573 | A/Illinois/NHRC0002/2011(H3N2) | 19/1/2011 | NCBI |
| CY091581 | A/California/NHRC0001/2011(H3N2) | 14/1/2011 | NCBI |
| CY091197 | A/Australia/NHRC0007/2003(H3N2) | 7/7/2003 | NCBI |
| CY091213 | A/California/NHRC0004/2003(H3N2) | 19/12/2003 | NCBI |
| CY091253 | A/Singapore/NHRC0008/2003(H3N2) | 8/10/2003 | NCBI |
| CY091269 | A/Australia/NHRC0009/2003(H3N2) | 9/7/2003 | NCBI |
| CY091317 | A/Australia/NHRC0015/2005(H3N2) | 19/6/2005 | NCBI |
| CY091333 | A/Australia/NHRC0013/2005(H3N2) | 15/6/2005 | NCBI |
| CY091381 | A/Australia/NHRC0006/2005(H3N2) | 13/6/2005 | NCBI |
| CY091397 | A/Australia/NHRC0004/2005(H3N2) | 15/4/2005 | NCBI |

| | | | |
|---|---|---|---|
| CY091437 | A/Malaysia/NHRC0003/2004(H3N2) | 26/9/2004 | NCBI |
| CY091453 | A/Malaysia/NHRC0001/2004(H3N2) | 15/9/2004 | NCBI |
| CY091509 | A/California/NHRC0004/2005(H3N2) | 2/11/2005 | NCBI |
| CY091525 | A/California/NHRC0006/2005(H3N2) | 2/12/2005 | NCBI |
| CY091557 | A/Texas/NHRC0001/2011(H3N2) | 25/1/2011 | NCBI |
| CY092217 | A/NewJersey/NHRC0001/2005(H3N2) | 10/2/2005 | NCBI |
| CY092201 | A/Illinois/NHRC0001/2003(H3N2) | 28/10/2003 | NCBI |
| CY092249 | A/California/NHRC0002/2007(H3N2) | 30/4/2007 | NCBI |
| CY092257 | A/California/NHRC0010/2005(H3N2) | 8/2/2005 | NCBI |
| CY092281 | A/California/NHRC0002/2011(H3N2) | 16/1/2011 | NCBI |
| CY092225 | A/Australia/NHRC0005/2003(H3N2) | 8/7/2003 | NCBI |
| CY092233 | A/Australia/NHRC0010/2005(H3N2) | 13/6/2005 | NCBI |
| CY092241 | A/California/NHRC0001/2007(H3N2) | 19/3/2007 | NCBI |
| CY092209 | A/Georgia/NHRC0001/2003(H3N2) | 5/12/2003 | NCBI |
| CY092265 | A/California/NHRC0003/2011(H3N2) | 28/1/2011 | NCBI |
| CY092273 | A/California/NHRC0004/2011(H3N2) | 28/1/2011 | NCBI |
| CY092297 | A/Managua/58-11/2011(H3N2) | 25/1/2011 | NCBI |
| CY092289 | A/Managua/38-11/2011(H3N2) | 28/1/2011 | NCBI |
| CY092305 | A/Managua/2492.04/2010(H3N2) | 2/7/2010 | NCBI |
| CY092313 | A/Managua/3424.01/2010(H3N2) | 30/6/2010 | NCBI |
| CY092353 | A/California/VRDL197/2009(H3N2) | 29/4/2009 | NCBI |
| CY092377 | A/California/VRDL257/2009(H3N2) | 1/5/2009 | NCBI |
| CY092361 | A/California/VRDL242/2009(H3N2) | 29/4/2009 | NCBI |
| CY092369 | A/California/VRDL255/2009(H3N2) | 29/4/2009 | NCBI |
| CY092329 | A/California/VRDL182/2009(H3N2) | 28/4/2009 | NCBI |
| CY093117 | A/Managua/5744.03/2010(H3N2) | 7/6/2010 | NCBI |
| CY093359 | A/Arizona/WRAIR1562P/2009(H3N2) | 1/3/2009 | NCBI |
| CY093351 | A/Kuwait/WRAIR1561P/2009(H3N2) | 1/1/2009 | NCBI |
| CY093319 | A/Kyrgyzstan/WRAIR1256P/2008(H3N2) | 1/12/2008 | NCBI |
| CY093327 | A/Nevada/WRAIR1258P/2009(H3N2) | 1/2/2009 | NCBI |
| CY093255 | A/Florida/WRAIR1241P/2009(H3N2) | 1/5/2009 | NCBI |
| CY093335 | A/Guam/WRAIR1557P/2009(H3N2) | 1/2/2009 | NCBI |
| CY093271 | A/Iraq/WRAIR1244P/2009(H3N2) | 1/2/2009 | NCBI |
| CY093535 | A/MexicoCity/WRAIR3579N/2010(H3N2) | 9/12/2010 | NCBI |
| CY093543 | A/MexicoCity/WRAIR3579T/2010(H3N2) | 9/12/2010 | NCBI |
| CY093263 | A/Alabama/WRAIR1242P/2009(H3N2) | 1/2/2009 | NCBI |
| CY093224 | A/DistrictofColumbia/WRAIR0300/2010(H3N2) | 17/11/2010 | NCBI |

| | | | |
|---|---|---|---|
| CY093303 | A/Iraq/WRAIR1252P/2009(H3N2) | 1/2/2009 | NCBI |
| CY093311 | A/Kyrgyzstan/WRAIR1255P/2008(H3N2) | 1/12/2008 | NCBI |
| CY093287 | A/NewJersey/WRAIR1248P/2009(H3N2) | 1/1/2009 | NCBI |
| CY093240 | A/SouthKorea/WRAIR1171P/2009(H3N2) | 1/1/2009 | NCBI |
| CY093343 | A/Texas/WRAIR1558P/2009(H3N2) | 1/2/2009 | NCBI |
| CY093415 | A/Bogota/WRAIR3457T/2010(H3N2) | 1/12/2010 | NCBI |
| CY093279 | A/Colorado/WRAIR1246P/2009(H3N2) | 1/5/2009 | NCBI |
| CY093447 | A/SanSalvador/WRAIR3537T/2010(H3N2) | 11/8/2010 | NCBI |
| CY093423 | A/SantoDomingo/WRAIR3514T/2010(H3N2) | 29/6/2010 | NCBI |
| CY093431 | A/SantoDomingo/WRAIR3516N/2010(H3N2) | 29/6/2010 | NCBI |
| CY093439 | A/SantoDomingo/WRAIR3516T/2010(H3N2) | 29/6/2010 | NCBI |
| CY096843 | A/Canterbury/200/2004(H3N2) | 16/9/2004 | NCBI |
| CY096859 | A/Memphis/3/1988(H3N2) | 1988 | NCBI |
| CY098081 | A/Managua/1645.03/2010(H3N2) | 9/6/2010 | NCBI |
| CY098065 | A/Managua/5828.02/2010(H3N2) | 3/6/2010 | NCBI |
| CY098073 | A/Managua/3587.02/2010(H3N2) | 3/6/2010 | NCBI |
| CY025541 | A/Vermont/UR06-0448/2007(H3N2) | 26/2/2007 | NCBI |
| CY038783 | A/Mississippi/UR07-0003/2008(H3N2) | 11/2/2008 | NCBI |
| CY038791 | A/California/UR07-0017/2008(H3N2) | 13/2/2008 | NCBI |
| CY038799 | A/Florida/UR07-0021/2008(H3N2) | 14/2/2008 | NCBI |
| CY038815 | A/California/UR07-0067/2008(H3N2) | 20/2/2008 | NCBI |
| CY038831 | A/Ohio/UR07-0073/2008(H3N2) | 21/2/2008 | NCBI |
| CY038847 | A/Kansas/UR07-0096/2008(H3N2) | 26/2/2008 | NCBI |
| CY038855 | A/Kentucky/UR07-0148/2008(H3N2) | 10/3/2008 | NCBI |
| CY038863 | A/Kentucky/UR07-0149/2008(H3N2) | 10/3/2008 | NCBI |
| CY038871 | A/Florida/UR07-0150/2008(H3N2) | 10/3/2008 | NCBI |
| CY038911 | A/Managua/419.01/2007(H3N2) | 28/6/2007 | NCBI |
| CY038935 | A/HongKong/HKU3/2004(H3N2) | 19/2/2004 | NCBI |
| CY038943 | A/HongKong/HKU4/2004(H3N2) | 1/3/2004 | NCBI |
| CY038951 | A/HongKong/HKU6/2004(H3N2) | 6/3/2004 | NCBI |
| CY038959 | A/HongKong/HKU7/2004(H3N2) | 10/3/2004 | NCBI |
| CY038975 | A/HongKong/HKU10/2004(H3N2) | 22/3/2004 | NCBI |
| CY038983 | A/HongKong/HKU14/2004(H3N2) | 16/4/2004 | NCBI |
| CY038991 | A/HongKong/HKU15/2004(H3N2) | 23/4/2004 | NCBI |
| CY039007 | A/HongKong/HKU21/2004(H3N2) | 19/6/2004 | NCBI |
| CY039015 | A/HongKong/HKU23/2004(H3N2) | 2/7/2004 | NCBI |
| CY039023 | A/HongKong/HKU25/2004(H3N2) | 7/7/2004 | NCBI |

| | | | |
|---|---|---|---|
| CY039031 | A/HongKong/HKU30/2004(H3N2) | 22/7/2004 | NCBI |
| CY039039 | A/HongKong/HKU33/2004(H3N2) | 9/8/2004 | NCBI |
| CY039047 | A/HongKong/HKU40/2004(H3N2) | 23/11/2004 | NCBI |
| CY039055 | A/HongKong/HKU45/2005(H3N2) | 7/3/2005 | NCBI |
| CY039063 | A/HongKong/HKU47/2005(H3N2) | 19/3/2005 | NCBI |
| CY039095 | A/Ohio/UR07-0034/2008(H3N2) | 18/2/2008 | NCBI |
| CY039111 | A/Ohio/UR07-0045/2008(H3N2) | 19/2/2008 | NCBI |
| CY039159 | A/HongKong/HKU5/2004(H3N2) | 4/3/2004 | NCBI |
| CY039167 | A/HongKong/HKU24/2004(H3N2) | 5/7/2004 | NCBI |
| CY039175 | A/HongKong/HKU44/2005(H3N2) | 28/2/2005 | NCBI |
| CY039183 | A/HongKong/HKU48/2005(H3N2) | 1/4/2005 | NCBI |
| CY039207 | A/HongKong/HKU58/2005(H3N2) | 5/5/2005 | NCBI |
| CY039215 | A/HongKong/HKU59/2005(H3N2) | 7/5/2005 | NCBI |
| CY039223 | A/HongKong/HKU61/2005(H3N2) | 11/5/2005 | NCBI |
| CY039231 | A/HongKong/HKU63/2005(H3N2) | 14/5/2005 | NCBI |
| CY039239 | A/HongKong/HKU64/2005(H3N2) | 16/5/2005 | NCBI |
| CY039247 | A/HongKong/HKU66/2005(H3N2) | 21/5/2005 | NCBI |
| JQ070800 | A/Indiana/08/2011(H3N2) | 27/7/2011 | NCBI |
| JQ290172 | A/Iowa/08/2011(H3N2) | 14/11/2011 | NCBI |
| JQ290180 | A/Iowa/07/2011(H3N2) | 14/11/2011 | NCBI |
| JQ290188 | A/Iowa/09/2011(H3N2) | 14/11/2011 | NCBI |
| CY104076 | A/HaNoi/ARI189/2003(H3N2) | 2003 | NCBI |
| CY104084 | A/TayNguyen/TN152/2003(H3N2) | 18/9/2003 | NCBI |
| CY104092 | A/TayNguyen/TN170/2003(H3N2) | 29/10/2003 | NCBI |
| CY104100 | A/TayNguyen/TN171/2003(H3N2) | 19/10/2003 | NCBI |
| CY104108 | A/TayNguyen/TN83/2003(H3N2) | 2003 | NCBI |
| CY104116 | A/TayNguyen/TN85/2003(H3N2) | 2003 | NCBI |
| CY104124 | A/TayNguyen/TN92/2003(H3N2) | 2003 | NCBI |
| CY104132 | A/BacGiang/BG003/2004(H3N2) | 13/2/2004 | NCBI |
| CY104140 | A/HaNoi/HN30110/2004(H3N2) | 30/1/2004 | NCBI |
| CY104148 | A/NamDinh/HN30119/2004(H3N2) | 31/1/2004 | NCBI |
| CY104156 | A/HaNoi/HN30127/2004(H3N2) | 1/2/2004 | NCBI |
| CY104164 | A/HaNoi/HN30136/2004(H3N2) | 4/2/2004 | NCBI |
| CY104172 | A/HaNoi/HN30108/2004(H3N2) | 2004 | NCBI |
| CY104180 | A/Hanoi/HN30109/2004(H3N2) | 2004 | NCBI |
| CY104188 | A/HungYen/HY028/2004(H3N2) | 2004 | NCBI |
| CY104196 | A/TayNguyen/TN46/2004(H3N2) | 2004 | NCBI |

| | | | |
|---|---|---|---|
| CY104204 | A/TayNguyen/TN49/2004(H3N2) | 2004 | NCBI |
| CY104212 | A/HaNoi/ISBM16/2005(H3N2) | 24/3/2005 | NCBI |
| CY104220 | A/TayNguyen/TN340/2005(H3N2) | 15/7/2005 | NCBI |
| CY104228 | A/TayNguyen/TN346/2005(H3N2) | 1/8/2005 | NCBI |
| CY104236 | A/TayNguyen/TN360/2005(H3N2) | 11/8/2005 | NCBI |
| CY104252 | A/TayNguyen/TN381/2005(H3N2) | 30/8/2005 | NCBI |
| CY104260 | A/HaNoi/Q372/2005(H3N2) | 30/6/2005 | NCBI |
| CY104268 | A/HaNoi/Q427/2005(H3N2) | 22/7/2005 | NCBI |
| CY104316 | A/HaNoi/BM061/2007(H3N2) | 22/3/2007 | NCBI |
| CY104324 | A/HaNoi/BT17/2007(H3N2) | 22/1/2007 | NCBI |
| CY104332 | A/HaNoi/BT262/2007(H3N2) | 3/8/2007 | NCBI |
| CY104340 | A/HaNoi/GS0740/2007(H3N2) | 27/7/2007 | NCBI |
| CY104348 | A/Hue/H020/2007(H3N2) | 1/10/2007 | NCBI |
| CY104356 | A/HoaBinh/HB378/2007(H3N2) | 3/10/2007 | NCBI |
| CY104372 | A/HungYen/HY3/2007(H3N2) | 2007 | NCBI |
| CY104380 | A/LangSon/LS156/2007(H3N2) | 29/5/2007 | NCBI |
| CY104388 | A/HaNoi/N015/2007(H3N2) | 10/1/2007 | NCBI |
| CY104396 | A/HaNoi/N027/2007(H3N2) | 17/1/2007 | NCBI |
| CY104404 | A/HaNoi/N064/2007(H3N2) | 9/2/2007 | NCBI |
| CY104412 | A/HaNoi/N078/2007(H3N2) | 10/3/2007 | NCBI |
| CY104420 | A/HaNoi/Q011/2007(H3N2) | 4/1/2007 | NCBI |
| CY104428 | A/HaNoi/Q039/2007(H3N2) | 15/1/2007 | NCBI |
| CY104436 | A/HaNoi/Q097/2007(H3N2) | 2/2/2007 | NCBI |
| CY104444 | A/HaNoi/Q118/2007(H3N2) | 12/2/2007 | NCBI |
| CY104452 | A/HaNoi/Q163/2007(H3N2) | 12/3/2007 | NCBI |
| CY104460 | A/HaNoi/Q430/2007(H3N2) | 18/6/2007 | NCBI |
| CY104468 | A/HaNoi/Q502/2007(H3N2) | 12/7/2007 | NCBI |
| CY104484 | A/HaNoi/Q664/2007(H3N2) | 7/9/2007 | NCBI |
| CY104492 | A/HaNoi/Q672/2007(H3N2) | 11/9/2007 | NCBI |
| CY104500 | A/HaNoi/Q748/2007(H3N2) | 8/10/2007 | NCBI |
| CY104508 | A/HaNoi/TX050/2007(H3N2) | 23/1/2007 | NCBI |
| CY104516 | A/HaNoi/TX058/2007(H3N2) | 24/1/2007 | NCBI |
| CY104524 | A/HaNoi/TX320/2007(H3N2) | 31/7/2007 | NCBI |
| CY104532 | A/HaNoi/TX322/2007(H3N2) | 2/8/2007 | NCBI |
| CY104548 | A/BacNinh/N52/2007(H3N2) | 2/1/2007 | NCBI |
| CY104622 | A/DaNang/DN446/2008(H3N2) | 21/11/2008 | NCBI |
| CY104630 | A/KhanhHoa/KH466/2008(H3N2) | 3/12/2008 | NCBI |

| | | | |
|---|---|---|---|
| CY104646 | A/KhanhHoa/KH494/2008(H3N2) | 22/12/2008 | NCBI |
| CY104678 | A/TayNguyen/HCM955/2007(H3N2) | 9/7/2007 | NCBI |
| CY105190 | A/HaNoi/1766/2001(H3N2) | 2001 | NCBI |
| CY105206 | A/Hanoi/BM766/2003(H3N2) | 16/6/2003 | NCBI |
| CY105214 | A/Hanoi/BM767/2003(H3N2) | 16/6/2003 | NCBI |
| CY105238 | A/TayNguyen/TN160/2003(H3N2) | 25/9/2003 | NCBI |
| CY105246 | A/TayNguyen/TN186/2003(H3N2) | 18/11/2003 | NCBI |
| CY105254 | A/HaNoi/HN695/2003(H3N2) | 14/8/2003 | NCBI |
| CY105262 | A/TayNguyen/TN905/2003(H3N2) | 2003 | NCBI |
| CY105270 | A/HaNoi/662/2004(H3N2) | 2004 | NCBI |
| CY105278 | A/HaNoi/BM769/2004(H3N2) | 2004 | NCBI |
| CY105286 | A/HaNoi/HN30130/2004(H3N2) | 2/2/2004 | NCBI |
| CY105294 | A/Hanoi/HN30135/2004(H3N2) | 4/2/2004 | NCBI |
| CY105302 | A/HaNoi/HN30137/2004(H3N2) | 4/2/2004 | NCBI |
| CY105310 | A/HaNoi/HN30147/2004(H3N2) | 5/2/2004 | NCBI |
| CY105318 | A/Hanoi/HN30240/2004(H3N2) | 10/3/2004 | NCBI |
| CY105326 | A/Hanoi/HN3069/2004(H3N2) | 18/1/2004 | NCBI |
| CY105334 | A/Hanoi/HN3094/2004(H3N2) | 28/1/2004 | NCBI |
| CY105342 | A/HaNoi/BM544/2004(H3N2) | 2004 | NCBI |
| CY105358 | A/TayNguyen/TN335/2005(H3N2) | 14/6/2005 | NCBI |
| CY105366 | A/TayNguyen/TN343/2005(H3N2) | 19/7/2005 | NCBI |
| CY105374 | A/TayNguyen/TN348/2005(H3N2) | 1/8/2005 | NCBI |
| CY105382 | A/TayNguyen/TN362/2005(H3N2) | 18/8/2005 | NCBI |
| CY105390 | A/TayNguyen/TN363/2005(H3N2) | 18/8/2005 | NCBI |
| CY105398 | A/TayNguyen/TN367/2005(H3N2) | 18/8/2005 | NCBI |
| CY105406 | A/TayNguyen/TN368/2005(H3N2) | 20/8/2005 | NCBI |
| CY105414 | A/TayNguyen/TN380/2005(H3N2) | 30/8/2005 | NCBI |
| CY105422 | A/TayNguyen/TN388/2005(H3N2) | 26/9/2005 | NCBI |
| CY105430 | A/TayNguyen/TN393/2005(H3N2) | 3/10/2005 | NCBI |
| CY105438 | A/TayNguyen/TN403/2005(H3N2) | 10/10/2005 | NCBI |
| CY105446 | A/TayNguyen/TN405/2005(H3N2) | 13/10/2005 | NCBI |
| CY105454 | A/TayNguyen/TN406/2005(H3N2) | 13/10/2005 | NCBI |
| CY105462 | A/TayNguyen/TN410/2005(H3N2) | 17/10/2005 | NCBI |
| CY105470 | A/TayNguyen/TN332/2005(H3N2) | 14/6/2005 | NCBI |
| CY105478 | A/HaNoi/BM171/2007(H3N2) | 11/7/2007 | NCBI |
| CY105486 | A/HaNoi/BM238/2007(H3N2) | 12/9/2007 | NCBI |
| CY105494 | A/HaNoi/BM38/2007(H3N2) | 9/2/2007 | NCBI |

| | | | |
|---|---|---|---|
| CY105502 | A/HaNoi/BT251/2007(H3N2) | 26/7/2007 | NCBI |
| CY105510 | A/HaNoi/GS0737/2007(H3N2) | 25/7/2007 | NCBI |
| CY105518 | A/HaNoi/GS0738/2007(H3N2) | 27/7/2007 | NCBI |
| CY105526 | A/Hue/H511/2007(H3N2) | 30/11/2007 | NCBI |
| CY105534 | A/Hue/H515/2007(H3N2) | 4/12/2007 | NCBI |
| CY105542 | A/Hue/H519/2007(H3N2) | 6/12/2007 | NCBI |
| CY105550 | A/Hue/H521/2007(H3N2) | 7/12/2007 | NCBI |
| CY105558 | A/HungYen/HY6/2007(H3N2) | 2007 | NCBI |
| CY105566 | A/LangSon/LS218/2007(H3N2) | 24/7/2007 | NCBI |
| CY105574 | A/LangSon/LS281/2007(H3N2) | 13/9/2007 | NCBI |
| CY105582 | A/HaNoi/N035/2007(H3N2) | 22/1/2007 | NCBI |
| CY105590 | A/HaNoi/N062/2007(H3N2) | 8/2/2007 | NCBI |
| CY105598 | A/HaNoi/N073/2007(H3N2) | 6/3/2007 | NCBI |
| CY105606 | A/HaNoi/N080/2007(H3N2) | 10/3/2007 | NCBI |
| CY105614 | A/HaNoi/N210/2007(H3N2) | 21/6/2007 | NCBI |
| CY105622 | A/HaNoi/N317/2007(H3N2) | 17/10/2007 | NCBI |
| CY105630 | A/HaNoi/N71/2007(H3N2) | 2/3/2007 | NCBI |
| CY105638 | A/HaNoi/Q031/2007(H3N2) | 11/1/2007 | NCBI |
| CY105646 | A/HaNoi/Q071/2007(H3N2) | 25/1/2007 | NCBI |
| CY105654 | A/HaNoi/Q094/2007(H3N2) | 2/2/2007 | NCBI |
| CY105662 | A/HaNoi/Q099/2007(H3N2) | 5/2/2007 | NCBI |
| CY105670 | A/HaNoi/Q148/2007(H3N2) | 6/3/2007 | NCBI |
| CY105678 | A/HaNam/Q187/2007(H3N2) | 20/3/2007 | NCBI |
| CY105686 | A/HaNoi/Q220/2007(H3N2) | 30/3/2007 | NCBI |
| CY105694 | A/HaNoi/Q464/2007(H3N2) | 29/6/2007 | NCBI |
| CY105702 | A/HaNoi/Q492/2007(H3N2) | 10/7/2007 | NCBI |
| CY105710 | A/HaNoi/Q499/2007(H3N2) | 12/7/2007 | NCBI |
| CY105718 | A/HaNoi/Q545/2007(H3N2) | 27/7/2007 | NCBI |
| CY105726 | A/HaNoi/Q578/2007(H3N2) | 8/8/2007 | NCBI |
| CY105734 | A/HaNoi/Q638/2007(H3N2) | 29/8/2007 | NCBI |
| CY105742 | A/HaNoi/Q656/2007(H3N2) | 5/9/2007 | NCBI |
| CY105750 | A/HaNoi/Q769/2007(H3N2) | 15/10/2007 | NCBI |
| CY105758 | A/TayNguyen/TN341/2007(H3N2) | 25/6/2007 | NCBI |
| CY105766 | A/TayNguyen/TN367/2007(H3N2) | 4/7/2007 | NCBI |
| CY105774 | A/HaNoi/TX019/2007(H3N2) | 10/1/2007 | NCBI |
| CY105782 | A/HaNoi/TX022/2007(H3N2) | 12/1/2007 | NCBI |
| CY105790 | A/HaNoi/TX023/2007(H3N2) | 12/1/2007 | NCBI |

| | | | |
|---|---|---|---|
| CY105798 | A/HaNoi/TX067/2007(H3N2) | 28/1/2007 | NCBI |
| CY105806 | A/HaNoi/TX072/2007(H3N2) | 29/1/2007 | NCBI |
| CY105814 | A/HaNoi/Q924/2007(H3N2) | 5/12/2007 | NCBI |
| CY105822 | A/DaNang/DN453/2008(H3N2) | 26/11/2008 | NCBI |
| CY105830 | A/Hue/H05/2008(H3N2) | 4/1/2008 | NCBI |
| CY105838 | A/Hue/H207/2008(H3N2) | 9/6/2008 | NCBI |
| CY105846 | A/Hue/H39/2008(H3N2) | 28/1/2008 | NCBI |
| CY105854 | A/KhanhHoa/KH363/2008(H3N2) | 23/9/2008 | NCBI |
| CY105862 | A/KhanhHoa/KH474/2008(H3N2) | 10/12/2008 | NCBI |
| CY105870 | A/KhanhHoa/KH475/2008(H3N2) | 10/12/2008 | NCBI |
| CY105878 | A/TayNguyen/HCM52/2006(H3N2) | 10/1/2006 | NCBI |
| CY105886 | A/HaNoi/311/2005(H3N2) | 2005 | NCBI |
| CY106576 | A/HongKong/H090-662-V10/2009(H3N2) | 30/6/2009 | NCBI |
| CY106584 | A/HongKong/H090-669-V10/2009(H3N2) | 4/7/2009 | NCBI |
| CY106592 | A/HongKong/H090-671-V10/2009(H3N2) | 6/7/2009 | NCBI |
| CY106600 | A/HongKong/H090-672-V10/2009(H3N2) | 6/7/2009 | NCBI |
| CY106608 | A/HongKong/H090-674-V10/2009(H3N2) | 7/7/2009 | NCBI |
| CY106616 | A/HongKong/H090-688-V10/2009(H3N2) | 10/7/2009 | NCBI |
| CY106624 | A/HongKong/H090-689-V10/2009(H3N2) | 9/7/2009 | NCBI |
| CY106632 | A/HongKong/H090-689-V20/2009(H3N2) | 12/7/2009 | NCBI |
| CY106640 | A/HongKong/H090-689-V22/2009(H3N2) | 12/7/2009 | NCBI |
| CY106648 | A/HongKong/H090-692-V10/2009(H3N2) | 10/7/2009 | NCBI |
| CY106656 | A/HongKong/H090-692-V12/2009(H3N2) | 10/7/2009 | NCBI |
| CY106664 | A/HongKong/H090-695-V10/2009(H3N2) | 10/7/2009 | NCBI |
| CY106672 | A/HongKong/H090-703-V10/2009(H3N2) | 17/7/2009 | NCBI |
| CY106680 | A/HongKong/H090-707-V10/2009(H3N2) | 14/7/2009 | NCBI |
| CY106688 | A/HongKong/H090-710-V10/2009(H3N2) | 14/7/2009 | NCBI |
| CY106696 | A/HongKong/H090-714-V10/2009(H3N2) | 15/7/2009 | NCBI |
| CY106704 | A/HongKong/H090-720-V10/2009(H3N2) | 18/7/2009 | NCBI |
| CY106712 | A/HongKong/H090-720-V20/2009(H3N2) | 21/7/2009 | NCBI |
| CY106720 | A/HongKong/H090-720-V21/2009(H3N2) | 21/7/2009 | NCBI |
| CY106728 | A/HongKong/H090-720-V31/2009(H3N2) | 24/7/2009 | NCBI |
| CY106736 | A/HongKong/H090-724-V10/2009(H3N2) | 20/7/2009 | NCBI |
| CY106744 | A/HongKong/H090-724-V20/2009(H3N2) | 22/7/2009 | NCBI |
| CY106752 | A/HongKong/H090-725-V10/2009(H3N2) | 20/7/2009 | NCBI |
| CY106760 | A/HongKong/H090-726-V10/2009(H3N2) | 20/7/2009 | NCBI |
| CY106768 | A/HongKong/H090-727-V10/2009(H3N2) | 21/7/2009 | NCBI |

| | | | |
|---|---|---|---|
| CY106776 | A/HongKong/H090-729-V10/2009(H3N2) | 21/7/2009 | NCBI |
| CY106784 | A/HongKong/H090-733-V10/2009(H3N2) | 22/7/2009 | NCBI |
| CY106792 | A/HongKong/H090-734-V10/2009(H3N2) | 22/7/2009 | NCBI |
| CY106800 | A/HongKong/H090-734-V20/2009(H3N2) | 24/7/2009 | NCBI |
| CY106808 | A/HongKong/H090-734-V32/2009(H3N2) | 27/7/2009 | NCBI |
| CY106816 | A/HongKong/H090-736-V10/2009(H3N2) | 23/7/2009 | NCBI |
| CY106824 | A/HongKong/H090-737-V10/2009(H3N2) | 23/7/2009 | NCBI |
| CY106832 | A/HongKong/H090-739-V10/2009(H3N2) | 23/7/2009 | NCBI |
| CY106840 | A/HongKong/H090-739-V22/2009(H3N2) | 27/7/2009 | NCBI |
| CY106848 | A/HongKong/H090-739-V23/2009(H3N2) | 27/7/2009 | NCBI |
| CY106856 | A/HongKong/H090-739-V33/2009(H3N2) | 29/7/2009 | NCBI |
| CY106864 | A/HongKong/H090-741-V10/2009(H3N2) | 24/7/2009 | NCBI |
| CY106872 | A/HongKong/H090-747-V10/2009(H3N2) | 28/7/2009 | NCBI |
| CY106880 | A/HongKong/H090-747-V20/2009(H3N2) | 30/7/2009 | NCBI |
| CY106888 | A/HongKong/H090-747-V22/2009(H3N2) | 30/7/2009 | NCBI |
| CY106896 | A/HongKong/H090-750-V10/2009(H3N2) | 29/7/2009 | NCBI |
| CY106904 | A/HongKong/H090-752-V10/2009(H3N2) | 29/7/2009 | NCBI |
| CY106912 | A/HongKong/H090-755-V20/2009(H3N2) | 1/8/2009 | NCBI |
| CY106920 | A/HongKong/H090-755-V31/2009(H3N2) | 5/8/2009 | NCBI |
| CY106928 | A/HongKong/H090-756-V10/2009(H3N2) | 30/7/2009 | NCBI |
| CY106936 | A/HongKong/H090-763-V10/2009(H3N2) | 31/7/2009 | NCBI |
| CY106944 | A/HongKong/H090-763-V23/2009(H3N2) | 3/8/2009 | NCBI |
| CY106952 | A/HongKong/H090-764-V10/2009(H3N2) | 31/7/2009 | NCBI |
| CY106960 | A/HongKong/H090-768-V10/2009(H3N2) | 2/8/2009 | NCBI |
| CY106968 | A/HongKong/H090-769-V10/2009(H3N2) | 3/8/2009 | NCBI |
| CY106984 | A/HongKong/H090-781-V10/2009(H3N2) | 6/8/2009 | NCBI |
| CY106992 | A/HongKong/H090-783-V10/2009(H3N2) | 7/8/2009 | NCBI |
| CY110759 | A/Kunming/1-Va10/2005(H3N2) | 2005 | NCBI |
| CY087976 | A/England/430/2003(H3N2) | 24/10/2003 | NCBI |
| CY087982 | A/England/431/2003(H3N2) | 2003-10 | NCBI |
| CY087990 | A/Scotland/52/2003(H3N2) | 23/10/2003 | NCBI |
| CY087998 | A/England/446/2003(H3N2) | 27/10/2003 | NCBI |
| CY088006 | A/Scotland/50/2003(H3N2) | 2003-11 | NCBI |
| CY088014 | A/England/419/2003(H3N2) | 3/11/2003 | NCBI |
| CY088022 | A/England/442/2003(H3N2) | 3/11/2003 | NCBI |
| CY088038 | A/England/438/2003(H3N2) | 12/11/2003 | NCBI |
| CY088046 | A/Scotland/61/2003(H3N2) | 26/10/2003 | NCBI |

| | | | |
|---|---|---|---|
| CY088054 | A/England/445/2003(H3N2) | 3/11/2003 | NCBI |
| CY088062 | A/England/740/2003(H3N2) | 2003-11 | NCBI |
| CY088070 | A/England/642/2003(H3N2) | 17/11/2003 | NCBI |
| CY088086 | A/England/460/2003(H3N2) | 11/11/2003 | NCBI |
| CY088094 | A/Scotland/71/2003(H3N2) | 4/11/2003 | NCBI |
| CY088102 | A/Scotland/72/2003(H3N2) | 28/10/2003 | NCBI |
| CY088110 | A/England/475/2003(H3N2) | 3/11/2003 | NCBI |
| CY088118 | A/England/467/2003(H3N2) | 6/11/2003 | NCBI |
| CY088126 | A/Scotland/76/2003(H3N2) | 3/11/2003 | NCBI |
| CY088134 | A/Scotland/78/2003(H3N2) | 7/11/2003 | NCBI |
| CY088150 | A/Scotland/81/2003(H3N2) | 12/11/2003 | NCBI |
| CY088174 | A/England/539/2003(H3N2) | 2003-11 | NCBI |
| CY088182 | A/England/479/2003(H3N2) | 19/11/2003 | NCBI |
| CY088190 | A/England/944/2003(H3N2) | 17/11/2003 | NCBI |
| CY088206 | A/England/518/2003(H3N2) | 5/11/2003 | NCBI |
| CY088214 | A/England/787/2003(H3N2) | 2003-11 | NCBI |
| CY088222 | A/England/520/2003(H3N2) | 20/11/2003 | NCBI |
| CY088230 | A/England/531/2003(H3N2) | 20/11/2003 | NCBI |
| CY088246 | A/England/789/2003(H3N2) | 2003-11 | NCBI |
| CY088254 | A/England/525/2003(H3N2) | 27/11/2003 | NCBI |
| CY088270 | A/England/585/2003(H3N2) | 25/11/2003 | NCBI |
| CY088278 | A/England/487/2003(H3N2) | 24/11/2003 | NCBI |
| CY088318 | A/England/485/2003(H3N2) | 5/12/2003 | NCBI |
| CY088326 | A/England/508/2003(H3N2) | 27/11/2003 | NCBI |
| CY088334 | A/England/568/2003(H3N2) | 26/11/2003 | NCBI |
| CY088342 | A/England/957/2003(H3N2) | 28/11/2003 | NCBI |
| CY088350 | A/England/567/2003(H3N2) | 4/12/2003 | NCBI |
| CY088358 | A/England/805/2003(H3N2) | 2003-11 | NCBI |
| CY088366 | A/England/558/2003(H3N2) | 5/12/2003 | NCBI |
| CY088374 | A/England/534/2003(H3N2) | 26/11/2003 | NCBI |
| CY088390 | A/England/522/2003(H3N2) | 12/12/2003 | NCBI |
| CY088398 | A/England/909/2003(H3N2) | 9/12/2003 | NCBI |
| CY088406 | A/England/547/2003(H3N2) | 18/12/2003 | NCBI |
| CY088414 | A/England/552/2003(H3N2) | 18/12/2003 | NCBI |
| CY088422 | A/England/587/2003(H3N2) | 6/12/2003 | NCBI |
| CY088430 | A/England/917/2003(H3N2) | 2/12/2003 | NCBI |
| CY088449 | A/England/592/2003(H3N2) | 16/12/2003 | NCBI |

| | | | |
|---|---|---|---|
| CY088457 | A/England/593/2003(H3N2) | 11/12/2003 | NCBI |
| CY088465 | A/England/602/2003(H3N2) | 14/12/2003 | NCBI |
| CY088475 | A/England/4024313/2003(H3N2) | 2003-12 | NCBI |
| CY088483 | A/Fujian/411/2002(H3N2) | 2002 | NCBI |
| CY111126 | A/Boston/DOA01/2011(H3N2) | 3/1/2011 | NCBI |
| CY111134 | A/Boston/DOA02/2011(H3N2) | 4/1/2011 | NCBI |
| CY111142 | A/Boston/DOA03/2011(H3N2) | 3/1/2011 | NCBI |
| CY111150 | A/Boston/DOA04/2011(H3N2) | 5/1/2011 | NCBI |
| CY111158 | A/Boston/DOA05/2011(H3N2) | 11/1/2011 | NCBI |
| CY111166 | A/Boston/DOA06/2011(H3N2) | 12/1/2011 | NCBI |
| CY111174 | A/Boston/DOA07/2011(H3N2) | 13/1/2011 | NCBI |
| CY111182 | A/Boston/DOA09/2011(H3N2) | 13/1/2011 | NCBI |
| CY111190 | A/Boston/DOA17/2011(H3N2) | 21/1/2011 | NCBI |
| CY111198 | A/Boston/DOA15/2011(H3N2) | 18/1/2011 | NCBI |
| CY111214 | A/Boston/DOA13/2011(H3N2) | 19/1/2011 | NCBI |
| CY111222 | A/Boston/DOA10/2011(H3N2) | 13/1/2011 | NCBI |
| CY111230 | A/Boston/DOA11/2011(H3N2) | 13/1/2011 | NCBI |
| CY111238 | A/Boston/DOA12/2011(H3N2) | 15/1/2011 | NCBI |
| CY111246 | A/Boston/DOA20/2011(H3N2) | 28/1/2011 | NCBI |
| CY111270 | A/Boston/DOA32/2011(H3N2) | 7/2/2011 | NCBI |
| CY111302 | A/Boston/DOA48/2011(H3N2) | 16/2/2011 | NCBI |
| CY111310 | A/Boston/DOA51/2011(H3N2) | 18/2/2011 | NCBI |
| CY111318 | A/Boston/DOA52/2011(H3N2) | 18/2/2011 | NCBI |
| CY111326 | A/Boston/DOA55/2011(H3N2) | 21/2/2011 | NCBI |
| CY111334 | A/Boston/DOA57/2011(H3N2) | 23/2/2011 | NCBI |
| CY111342 | A/Boston/DOA61/2011(H3N2) | 24/2/2011 | NCBI |
| CY111350 | A/Boston/DOA66/2011(H3N2) | 26/2/2011 | NCBI |
| CY111358 | A/Boston/DOA67/2011(H3N2) | 28/2/2011 | NCBI |
| CY111366 | A/Boston/DOA69/2011(H3N2) | 2/3/2011 | NCBI |
| CY111382 | A/Boston/DOA71/2011(H3N2) | 5/3/2011 | NCBI |
| CY111390 | A/Boston/DOA64/2011(H3N2) | 26/2/2011 | NCBI |
| CY111406 | A/Boston/DOA77/2011(H3N2) | 13/3/2011 | NCBI |
| CY111414 | A/Boston/DOA78/2011(H3N2) | 14/3/2011 | NCBI |
| CY111422 | A/Boston/DOA79/2011(H3N2) | 12/3/2011 | NCBI |
| CY111430 | A/Boston/DOA80/2011(H3N2) | 16/3/2011 | NCBI |
| CY111438 | A/Boston/DOA81/2011(H3N2) | 17/3/2011 | NCBI |
| CY111446 | A/Boston/DOA82/2011(H3N2) | 17/3/2011 | NCBI |

| | | | |
|---|---|---|---|
| CY111454 | A/Boston/DOA85/2011(H3N2) | 23/3/2011 | NCBI |
| CY111462 | A/Boston/DOA86/2011(H3N2) | 29/3/2011 | NCBI |
| CY111470 | A/Boston/DOA87/2011(H3N2) | 25/4/2011 | NCBI |
| CY111478 | A/Boston/DOA75/2011(H3N2) | 11/3/2011 | NCBI |
| CY111522 | A/WesternAustralia/6/2000(H3N2) | 4/10/2000 | NCBI |
| CY111546 | A/Boston/DOA33/2011(H3N2) | 8/2/2011 | NCBI |
| CY112233 | A/Bilthoven/15793/1968(H3N2) | 1968 | NCBI |
| CY112241 | A/Bilthoven/16398/1968(H3N2) | 1968 | NCBI |
| CY112249 | A/HongKong/1/1968(H3N2) | 1968 | NCBI |
| CY112257 | A/Bilthoven/808/1969(H3N2) | 1969 | NCBI |
| CY112265 | A/Bilthoven/93/1970(H3N2) | 1970 | NCBI |
| CY112273 | A/Bilthoven/2668/1970(H3N2) | 1970 | NCBI |
| CY112289 | A/HongKong/107/1971(H3N2) | 1971 | NCBI |
| CY112297 | A/Bilthoven/6022/1972(H3N2) | 1972 | NCBI |
| CY112305 | A/Bilthoven/21793/1972(H3N2) | 1972 | NCBI |
| CY112321 | A/Bilthoven/748/1973(H3N2) | 1973 | NCBI |
| CY112329 | A/Bilthoven/5146/1974(H3N2) | 1974 | NCBI |
| CY112337 | A/Bilthoven/4273/1975(H3N2) | 1975 | NCBI |
| CY112353 | A/Rotterdam/577/1980(H3N2) | 1980 | NCBI |
| CY112361 | A/Bilthoven/4791/1981(H3N2) | 1981 | NCBI |
| CY112396 | A/Sichuan/2/1987(H3N2) | 1987 | NCBI |
| CY112404 | A/Beijing/352/1989(H3N2) | 1989 | NCBI |
| CY112420 | A/Geneva/5007/1989(H3N2) | 1989 | NCBI |
| CY112428 | A/HongKong/1/1989(H3N2) | 1989 | NCBI |
| CY112436 | A/Singapore/53/1989(H3N2) | 1989 | NCBI |
| CY112444 | A/Geneva/6447/1991(H3N2) | 27/11/1991 | NCBI |
| CY112452 | A/Stockholm/20/1991(H3N2) | 17/12/1991 | NCBI |
| CY112460 | A/Amsterdam/4112/1992(H3N2) | 1992 | NCBI |
| CY112476 | A/Finland/218/1992(H3N2) | 24/1/1992 | NCBI |
| CY112484 | A/Finland/220/1992(H3N2) | 24/1/1992 | NCBI |
| CY112492 | A/Finland/247/1992(H3N2) | 1992 | NCBI |
| CY112500 | A/Paris/490/1991(H3N2) | 1/12/1991 | NCBI |
| CY112508 | A/Rotterdam/100540/1992(H3N2) | 26/1/1992 | NCBI |
| CY112516 | A/Sendai/C273/1992(H3N2) | 1992 | NCBI |
| CY112524 | A/SouthAustralia/8/1992(H3N2) | 5/3/1992 | NCBI |
| CY112532 | A/Stockholm/8/1992(H3N2) | 2/3/1992 | NCBI |
| CY112540 | A/Victoria/33/1992(H3N2) | 29/6/1992 | NCBI |

| | | | |
|---|---|---|---|
| CY112548 | A/Lyon/1803/1993(H3N2) | 27/11/1993 | NCBI |
| CY112556 | A/Lyon/22686/1993(H3N2) | 2/12/1993 | NCBI |
| CY112573 | A/Madrid/G130/1993(H3N2) | 1993 | NCBI |
| CY112581 | A/Netherlands/3/1992(H3N2) | 22/11/1992 | NCBI |
| CY112589 | A/Netherlands/17/1993(H3N2) | 1/1/1993 | NCBI |
| CY112597 | A/Netherlands/115/1993(H3N2) | 18/2/1993 | NCBI |
| CY112605 | A/Netherlands/179/1993(H3N2) | 26/2/1993 | NCBI |
| CY112613 | A/Netherlands/241/1993(H3N2) | 13/10/1993 | NCBI |
| CY112621 | A/Netherlands/371/1993(H3N2) | 2/12/1993 | NCBI |
| CY112637 | A/Oslo/2219/1993(H3N2) | 14/11/1993 | NCBI |
| CY112645 | A/Oslo/244/1997(H3N2) | 1997 | NCBI |
| CY112653 | A/Shiga/6/1993(H3N2) | 1993 | NCBI |
| CY112661 | A/Singapore/19/1993(H3N2) | 21/12/1993 | NCBI |
| CY112669 | A/Stockholm/20/1993(H3N2) | 3/11/1993 | NCBI |
| CY112677 | A/Victoria/104/1993(H3N2) | 3/9/1993 | NCBI |
| CY112685 | A/Wellington/59/1993(H3N2) | 1993 | NCBI |
| CY112693 | A/Yamagata/62/1993(H3N2) | 1993 | NCBI |
| CY112701 | A/England/7/1994(H3N2) | 1994 | NCBI |
| CY112709 | A/HongKong/1/1994(H3N2) | 1/6/1994 | NCBI |
| CY112717 | A/HongKong/2/1994(H3N2) | 1/6/1994 | NCBI |
| CY112725 | A/HongKong/55/1994(H3N2) | 26/7/1994 | NCBI |
| CY112733 | A/Johannesburg/33/1994(H3N2) | 1994 | NCBI |
| CY112741 | A/Johannesburg/47/1994(H3N2) | 1994 | NCBI |
| CY112749 | A/SouthAustralia/15/1994(H3N2) | 3/6/1994 | NCBI |
| CY112757 | A/SouthAustralia/25/1994(H3N2) | 9/6/1994 | NCBI |
| CY112765 | A/HongKong/32/1995(H3N2) | 1/3/1995 | NCBI |
| CY112773 | A/HongKong/38/1995(H3N2) | 1/8/1995 | NCBI |
| CY112781 | A/HongKong/55/1995(H3N2) | 12/10/1995 | NCBI |
| CY112789 | A/Nanchang/933/1995(H3N2) | 1995 | NCBI |
| CY112797 | A/Netherlands/271/1995(H3N2) | 2/12/1995 | NCBI |
| CY112813 | A/Victoria/75/1995(H3N2) | 27/7/1995 | NCBI |
| CY112821 | A/Wuhan/359/1995(H3N2) | 1995 | NCBI |
| CY112829 | A/Singapore/1/1996(H3N2) | 12/2/1996 | NCBI |
| CY112837 | A/HongKong/1/1997(H3N2) | 1997 | NCBI |
| CY112845 | A/HongKong/280/1997(H3N2) | 20/6/1997 | NCBI |
| CY112853 | A/Johannesburg/10/1997(H3N2) | 1/7/1997 | NCBI |
| CY112861 | A/Nice/491/1996(H3N2) | 30/12/1996 | NCBI |

| | | | |
|---|---|---|---|
| CY112869 | A/Oslo/21/1997(H3N2) | 1997 | NCBI |
| CY112877 | A/Oslo/2352/1993(H3N2) | 13/11/1993 | NCBI |
| CY112885 | A/Sydney/5/1997(H3N2) | 1997 | NCBI |
| CY112893 | A/Netherlands/414/1998(H3N2) | 12/5/1998 | NCBI |
| CY112901 | A/Netherlands/427/1998(H3N2) | 16/11/1998 | NCBI |
| CY112909 | A/Moscow/10/1999(H3N2) | 1999 | NCBI |
| CY112917 | A/Panama/2007/1999(H3N2) | 1999 | NCBI |
| CY112925 | A/Netherlands/118/2001(H3N2) | 14/3/2001 | NCBI |
| CY112933 | A/Fujian/411/2002(H3N2) | 2002 | NCBI |
| CY112941 | A/Netherlands/1/2002(H3N2) | 7/1/2002 | NCBI |
| CY112949 | A/Netherlands/20/2003(H3N2) | 18/2/2003 | NCBI |
| CY112957 | A/Netherlands/22/2003(H3N2) | 23/2/2003 | NCBI |
| CY112965 | A/Netherlands/88/2003(H3N2) | 5/3/2003 | NCBI |
| CY112981 | A/Netherlands/217/2003(H3N2) | 20/4/2003 | NCBI |
| CY112989 | A/Netherlands/132/2004(H3N2) | 4/12/2004 | NCBI |
| CY112997 | A/Netherlands/548/2005(H3N2) | 26/12/2005 | NCBI |
| CY113013 | A/Netherlands/377/2008(H3N2) | 27/9/2008 | NCBI |
| CY113021 | A/Netherlands/69/2009(H3N2) | 12/1/2009 | NCBI |
| CY113037 | A/Netherlands/761/2009(H3N2) | 25/6/2009 | NCBI |
| CY113045 | A/Bilthoven/908/1969(H3N2) | 1969 | NCBI |
| CY113053 | A/Bilthoven/17938/1969(H3N2) | 1969 | NCBI |
| CY113061 | A/Bilthoven/21438/1971(H3N2) | 1971 | NCBI |
| CY113069 | A/Bilthoven/21801/1971(H3N2) | 1971 | NCBI |
| CY113077 | A/Bilthoven/23290/1972(H3N2) | 1972 | NCBI |
| CY113085 | A/England/42/1972(H3N2) | 1972 | NCBI |
| CY113093 | A/Bilthoven/552/1973(H3N2) | 1973 | NCBI |
| CY113101 | A/Bilthoven/3517/1973(H3N2) | 1973 | NCBI |
| CY113109 | A/PortChalmers/1/1973(H3N2) | 1973 | NCBI |
| CY113117 | A/Bilthoven/5930/1974(H3N2) | 1974 | NCBI |
| CY113125 | A/Bilthoven/5931/1974(H3N2) | 1974 | NCBI |
| CY113133 | A/Bilthoven/7398/1974(H3N2) | 1974 | NCBI |
| CY113141 | A/Bilthoven/9459/1974(H3N2) | 1974 | NCBI |
| CY113149 | A/Bilthoven/334/1975(H3N2) | 1975 | NCBI |
| CY113157 | A/Bilthoven/1843/1975(H3N2) | 1975 | NCBI |
| CY113165 | A/Bilthoven/2600/1975(H3N2) | 1975 | NCBI |
| CY113173 | A/Bilthoven/2813/1975(H3N2) | 1975 | NCBI |
| CY113181 | A/Victoria/3/1975(H3N2) | 1975 | NCBI |

| | | | |
|---|---|---|---|
| CY113189 | A/Bilthoven/628/1976(H3N2) | 1976 | NCBI |
| CY113197 | A/Bilthoven/1761/1976(H3N2) | 1976 | NCBI |
| CY113205 | A/Bilthoven/5029/1976(H3N2) | 1976 | NCBI |
| CY113213 | A/Bilthoven/5657/1976(H3N2) | 1976 | NCBI |
| CY113221 | A/Bilthoven/6545/1976(H3N2) | 1976 | NCBI |
| CY113229 | A/Amsterdam/1609/1977(H3N2) | 1977 | NCBI |
| CY113237 | A/Bilthoven/3895/1977(H3N2) | 1977 | NCBI |
| CY113245 | A/Rotterdam/5828/1977(H3N2) | 1977 | NCBI |
| CY113253 | A/Rotterdam/8179/1977(H3N2) | 1977 | NCBI |
| CY113261 | A/Texas/1/1977(H3N2) | 1977 | NCBI |
| CY113269 | A/Netherlands/209/1980(H3N2) | 1/12/1980 | NCBI |
| CY113277 | A/Lyon/2380/1981(H3N2) | 1981 | NCBI |
| CY113285 | A/Bilthoven/10684/1982(H3N2) | 1982 | NCBI |
| CY113293 | A/Netherlands/241/1982(H3N2) | 1/12/1982 | NCBI |
| CY113301 | A/Philippines/2/1982(H3N2) | 1982 | NCBI |
| CY113309 | A/Oslo/13676/1983(H3N2) | 1983 | NCBI |
| CY113317 | A/Guildford/V728/1985(H3N2) | 1985 | NCBI |
| CY113325 | A/Netherlands/330/1985(H3N2) | 1/1/1985 | NCBI |
| CY113333 | A/Netherlands/333/1985(H3N2) | 1/2/1985 | NCBI |
| CY113341 | A/Stockholm/10/1985(H3N2) | 1985 | NCBI |
| CY113349 | A/Wellington/4/1985(H3N2) | 1985 | NCBI |
| CY113357 | A/Colorado/2/1986(H3N2) | 1986 | NCBI |
| CY113365 | A/Shanghai/11/1987(H3N2) | 1987 | NCBI |
| CY113373 | A/Atlanta/3572/1988(H3N2) | 1988 | NCBI |
| CY113381 | A/England/427/1988(H3N2) | 1988 | NCBI |
| CY113389 | A/Netherlands/450/1988(H3N2) | 1/3/1988 | NCBI |
| CY113397 | A/Stockholm/12/1988(H3N2) | 1988 | NCBI |
| CY113405 | A/Atlanta/211/1989(H3N2) | 1989 | NCBI |
| CY113413 | A/England/138/1989(H3N2) | 1989 | NCBI |
| CY113421 | A/Netherlands/620/1989(H3N2) | 27/11/1989 | NCBI |
| CY113429 | A/Netherlands/650/1989(H3N2) | 13/12/1989 | NCBI |
| CY113437 | A/Netherlands/738/1989(H3N2) | 17/12/1989 | NCBI |
| CY113445 | A/Singapore/3/1993(H3N2) | 6/1/1993 | NCBI |
| CY113453 | A/Singapore/34/1989(H3N2) | 1989 | NCBI |
| CY113461 | A/Singapore/35/1989(H3N2) | 1989 | NCBI |
| CY113469 | A/Singapore/40/1989(H3N2) | 1989 | NCBI |
| CY113477 | A/Victoria/1/1989(H3N2) | 1989 | NCBI |

| | | | |
|---|---|---|---|
| CY113485 | A/Wellington/5/1989(H3N2) | 1989 | NCBI |
| CY113493 | A/Waikato/1/1989(H3N2) | 1989 | NCBI |
| CY113501 | A/Memphis/2/1990(H3N2) | 1990 | NCBI |
| CY113509 | A/Memphis/5/1990(H3N2) | 1990 | NCBI |
| CY113517 | A/Shanghai/24/1990(H3N2) | 1990 | NCBI |
| CY113525 | A/Suita/1/1990(H3N2) | 1990 | NCBI |
| CY113533 | A/Victoria/2/1990(H3N2) | 1990 | NCBI |
| CY113541 | A/Wellington/3/1990(H3N2) | 1990 | NCBI |
| CY113549 | A/Canberra/1/1991(H3N2) | 1991 | NCBI |
| CY113557 | A/England/260/1991(H3N2) | 15/9/1991 | NCBI |
| CY113565 | A/Geneva/5366/1991(H3N2) | 1991 | NCBI |
| CY113573 | A/Lyon/1149/1991(H3N2) | 9/12/1991 | NCBI |
| CY113581 | A/Lyon/1182/1991(H3N2) | 12/12/1991 | NCBI |
| CY113589 | A/Lyon/1189/1991(H3N2) | 12/12/1991 | NCBI |
| CY113597 | A/Lyon/1276/1991(H3N2) | 12/12/1991 | NCBI |
| CY113605 | A/Lyon/1337/1991(H3N2) | 16/12/1991 | NCBI |
| CY113613 | A/Lyon/1373/1991(H3N2) | 16/12/1991 | NCBI |
| CY113621 | A/Lyon/1594/1991(H3N2) | 26/12/1991 | NCBI |
| CY113629 | A/Lyon/23672/1991(H3N2) | 17/12/1991 | NCBI |
| CY113637 | A/Lyon/24103/1991(H3N2) | 24/12/1991 | NCBI |
| CY113645 | A/Lyon/24222/1991(H3N2) | 26/12/1991 | NCBI |
| CY113653 | A/Madrid/G12/1991(H3N2) | 1991 | NCBI |
| CY113661 | A/Netherlands/816/1991(H3N2) | 19/12/1991 | NCBI |
| CY113669 | A/Netherlands/891/1991(H3N2) | 4/12/1991 | NCBI |
| CY113677 | A/Beijing/32/1992(H3N2) | 1992 | NCBI |
| CY113685 | A/Geneva/5113/1992(H3N2) | 15/1/1992 | NCBI |
| CY113693 | A/Houston/56798/1992(H3N2) | 3/4/1992 | NCBI |
| CY113701 | A/Houston/56829/1992(H3N2) | 3/4/1992 | NCBI |
| CY113709 | A/Houston/56941/1992(H3N2) | 3/4/1992 | NCBI |
| CY113717 | A/Madrid/G58/1992(H3N2) | 14/1/1992 | NCBI |
| CY113725 | A/Netherlands/819/1992(H3N2) | 9/1/1992 | NCBI |
| CY113733 | A/Netherlands/823/1992(H3N2) | 3/2/1992 | NCBI |
| CY113741 | A/Netherlands/935/1992(H3N2) | 10/1/1992 | NCBI |
| CY113749 | A/Netherlands/938/1991(H3N2) | 19/12/1991 | NCBI |
| CY113757 | A/Nijmegen/3126/1992(H3N2) | 9/1/1992 | NCBI |
| CY113765 | A/Nijmegen/3129/1992(H3N2) | 7/2/1992 | NCBI |
| CY113773 | A/Oviedo/31/1991(H3N2) | 28/12/1991 | NCBI |

| | | | |
|---|---|---|---|
| CY113781 | A/Paris/320/1991(H3N2) | 1/12/1991 | NCBI |
| CY113789 | A/Paris/325/1991(H3N2) | 1/12/1991 | NCBI |
| CY113797 | A/Paris/407/1991(H3N2) | 1/12/1991 | NCBI |
| CY113805 | A/Paris/417/1991(H3N2) | 1/12/1991 | NCBI |
| CY113813 | A/Paris/424/1991(H3N2) | 1/12/1991 | NCBI |
| CY113821 | A/Paris/457/1991(H3N2) | 1/12/1991 | NCBI |
| CY113829 | A/Paris/467/1991(H3N2) | 1/12/1991 | NCBI |
| CY113837 | A/Paris/512/1991(H3N2) | 1/12/1991 | NCBI |
| CY113845 | A/Paris/548/1991(H3N2) | 1/12/1991 | NCBI |
| CY113853 | A/Paris/564/1991(H3N2) | 1/12/1991 | NCBI |
| CY113861 | A/Paris/583/1992(H3N2) | 1/1/1992 | NCBI |
| CY113869 | A/Paris/597/1991(H3N2) | 1/12/1991 | NCBI |
| CY113877 | A/Paris/614/1992(H3N2) | 1/1/1992 | NCBI |
| CY113885 | A/SouthAustralia/23/1992(H3N2) | 31/3/1992 | NCBI |
| CY113893 | A/SouthAustralia/27/1992(H3N2) | 6/4/1992 | NCBI |
| CY113901 | A/Stockholm/7/1992(H3N2) | 14/2/1992 | NCBI |
| CY113909 | A/Stockholm/12/1992(H3N2) | 1/11/1992 | NCBI |
| CY113917 | A/Stockholm/13/1992(H3N2) | 1/12/1992 | NCBI |
| CY113925 | A/Tilburg/5957/1991(H3N2) | 4/12/1991 | NCBI |
| CY113933 | A/Umea/1982/1992(H3N2) | 1/10/1992 | NCBI |
| CY113941 | A/Umea/2000/1992(H3N2) | 1/11/1992 | NCBI |
| CY113949 | A/Victoria/68/1992(H3N2) | 17/7/1992 | NCBI |
| CY113957 | A/Akita/4/1993(H3N2) | 1993 | NCBI |
| CY113965 | A/Enschede/5458/1993(H3N2) | 1993 | NCBI |
| CY113973 | A/Guangdong/25/1993(H3N2) | 1993 | NCBI |
| CY113981 | A/Lyon/672/1993(H3N2) | 22/2/1993 | NCBI |
| CY113989 | A/Lyon/1815/1993(H3N2) | 30/11/1993 | NCBI |
| CY113997 | A/Lyon/23602/1993(H3N2) | 14/12/1993 | NCBI |
| CY114005 | A/Madrid/G122/1993(H3N2) | 1993 | NCBI |
| CY114013 | A/Madrid/G101/1993(H3N2) | 1993 | NCBI |
| CY114021 | A/Madrid/G109/1993(H3N2) | 1993 | NCBI |
| CY114029 | A/Madrid/G116/1993(H3N2) | 1993 | NCBI |
| CY114037 | A/Madrid/G252/1993(H3N2) | 1993 | NCBI |
| CY114045 | A/Netherlands/101/1993(H3N2) | 15/2/1993 | NCBI |
| CY114053 | A/Netherlands/126/1993(H3N2) | 23/2/1993 | NCBI |
| CY114061 | A/Netherlands/165/1993(H3N2) | 5/3/1993 | NCBI |
| CY114069 | A/Netherlands/357/1993(H3N2) | 6/12/1993 | NCBI |

| | | | |
|---|---|---|---|
| CY114077 | A/Netherlands/372/1993(H3N2) | 1/12/1993 | NCBI |
| CY114085 | A/Netherlands/398/1993(H3N2) | 27/11/1993 | NCBI |
| CY114093 | A/Paris/287/1993(H3N2) | 1993 | NCBI |
| CY114101 | A/Scotland/142/1993(H3N2) | 1993 | NCBI |
| CY114109 | A/Scotland/160/1993(H3N2) | 1993 | NCBI |
| CY114117 | A/Shandong/9/1993(H3N2) | 1993 | NCBI |
| CY114125 | A/Singapore/36/1989(H3N2) | 1989 | NCBI |
| CY114133 | A/Yamagata/56/1993(H3N2) | 1993 | NCBI |
| CY114141 | A/Yamagata/61/1993(H3N2) | 1993 | NCBI |
| CY114149 | A/HongKong/56/1994(H3N2) | 2/8/1994 | NCBI |
| CY114157 | A/Netherlands/18/1994(H3N2) | 5/1/1994 | NCBI |
| CY114165 | A/Finland/338/1995(H3N2) | 1995 | NCBI |
| CY114173 | A/Finland/339/1995(H3N2) | 1995 | NCBI |
| CY114181 | A/Finland/381/1995(H3N2) | 1995 | NCBI |
| CY114189 | A/Geneva/A9509/1995(H3N2) | 1/2/1995 | NCBI |
| CY114197 | A/HongKong/3/1995(H3N2) | 24/1/1995 | NCBI |
| CY114205 | A/HongKong/49/1995(H3N2) | 1/9/1995 | NCBI |
| CY114213 | A/Lyon/2279/1995(H3N2) | 20/11/1995 | NCBI |
| CY114221 | A/Netherlands/1/1995(H3N2) | 11/1/1995 | NCBI |
| CY114229 | A/Geneva/3958/1996(H3N2) | 1996 | NCBI |
| CY114237 | A/HongKong/20/1996(H3N2) | 24/2/1996 | NCBI |
| CY114245 | A/HongKong/42/1996(H3N2) | 4/3/1996 | NCBI |
| CY114253 | A/HongKong/357/1996(H3N2) | 19/7/1996 | NCBI |
| CY114261 | A/HongKong/358/1996(H3N2) | 19/7/1996 | NCBI |
| CY114269 | A/HongKong/434/1996(H3N2) | 2/10/1996 | NCBI |
| CY114277 | A/Lyon/1781/1996(H3N2) | 26/10/1996 | NCBI |
| CY114285 | A/Netherlands/91/1996(H3N2) | 30/1/1996 | NCBI |
| CY114293 | A/Auckland/10/1997(H3N2) | 1/8/1997 | NCBI |
| CY114301 | A/Netherlands/300/1997(H3N2) | 23/12/1997 | NCBI |
| CY114309 | A/Netherlands/462/1998(H3N2) | 16/12/1998 | NCBI |
| CY114317 | A/Netherlands/3/2000(H3N2) | 6/1/2000 | NCBI |
| CY114325 | A/Netherlands/124/2001(H3N2) | 20/3/2001 | NCBI |
| CY114333 | A/Netherlands/126/2001(H3N2) | 21/3/2001 | NCBI |
| CY114341 | A/Netherlands/120/2002(H3N2) | 26/2/2002 | NCBI |
| CY114349 | A/Finland/170/2003(H3N2) | 2003 | NCBI |
| CY114357 | A/Netherlands/222/2003(H3N2) | 14/4/2003 | NCBI |
| CY114365 | A/Netherlands/312/2003(H3N2) | 25/9/2003 | NCBI |

| | | | |
|---|---|---|---|
| CY114373 | A/California/7/2004(H3N2) | 2004 | NCBI |
| CY114381 | A/Wisconsin/67/2005(H3N2) | 2005 | NCBI |
| CY114389 | A/Netherlands/42/2006(H3N2) | 25/2/2006 | NCBI |
| CY114397 | A/Netherlands/363/2006(H3N2) | 24/12/2006 | NCBI |
| CY114405 | A/Netherlands/69/2007(H3N2) | 19/2/2007 | NCBI |
| CY114413 | A/Netherlands/348/2007(H3N2) | 6/11/2007 | NCBI |
| CY114421 | A/Netherlands/063/2011(H3N2) | 5/1/2011 | NCBI |
| CY114429 | A/Bangkok/1/1979(H3N2) | 1979 | NCBI |
| CY114437 | A/Netherlands/233/1982(H3N2) | 1/4/1982 | NCBI |
| CY114445 | A/Victoria/1/1988(H3N2) | 1988 | NCBI |
| CY114453 | A/Oklahoma/5/1988(H3N2) | 1988 | NCBI |
| CY114461 | A/Beijing/353/1989(H3N2) | 1989 | NCBI |
| CY114469 | A/UnitedKingdom/261/1991(H3N2) | 1991 | NCBI |
| CY114477 | A/Brisbane/8/1996(H3N2) | 1996 | NCBI |
| CY114485 | A/Netherlands/5/1998(H3N2) | 14/1/1998 | NCBI |
| CY114493 | A/Netherlands/301/1999(H3N2) | 1/10/1999 | NCBI |
| CY114501 | A/Netherlands/009/2010(H3N2) | 21/4/2010 | NCBI |
| CY114509 | A/Netherlands/034/2010(H3N2) | 23/12/2010 | NCBI |
| CY115464 | A/HongKong/H090-662-V10/2009(H3N2) | 30/6/2009 | NCBI |
| CY115472 | A/HongKong/H090-669-V10/2009(H3N2) | 4/7/2009 | NCBI |
| CY115480 | A/HongKong/H090-671-V10/2009(H3N2) | 6/7/2009 | NCBI |
| CY115488 | A/HongKong/H090-672-V10/2009(H3N2) | 6/7/2009 | NCBI |
| CY115496 | A/HongKong/H090-674-V10/2009(H3N2) | 7/7/2009 | NCBI |
| CY115504 | A/HongKong/H090-688-V10/2009(H3N2) | 10/7/2009 | NCBI |
| CY115512 | A/HongKong/H090-689-V10/2009(H3N2) | 9/7/2009 | NCBI |
| CY115520 | A/HongKong/H090-689-V20/2009(H3N2) | 12/7/2009 | NCBI |
| CY115528 | A/HongKong/H090-689-V22/2009(H3N2) | 12/7/2009 | NCBI |
| CY115536 | A/HongKong/H090-692-V10/2009(H3N2) | 10/7/2009 | NCBI |
| CY115544 | A/HongKong/H090-695-V10/2009(H3N2) | 10/7/2009 | NCBI |
| CY115552 | A/HongKong/H090-703-V10/2009(H3N2) | 17/7/2009 | NCBI |
| CY115560 | A/HongKong/H090-707-V10/2009(H3N2) | 14/7/2009 | NCBI |
| CY115568 | A/HongKong/H090-710-V10/2009(H3N2) | 14/7/2009 | NCBI |
| CY115576 | A/HongKong/H090-714-V10/2009(H3N2) | 15/7/2009 | NCBI |
| CY115584 | A/HongKong/H090-720-V10/2009(H3N2) | 18/7/2009 | NCBI |
| CY115592 | A/HongKong/H090-720-V20/2009(H3N2) | 21/7/2009 | NCBI |
| CY115600 | A/HongKong/H090-720-V21/2009(H3N2) | 21/7/2009 | NCBI |
| CY115608 | A/HongKong/H090-720-V31/2009(H3N2) | 24/7/2009 | NCBI |

| | | | |
|---|---|---|---|
| CY115616 | A/HongKong/H090-724-V10/2009(H3N2) | 20/7/2009 | NCBI |
| CY115624 | A/HongKong/H090-724-V20/2009(H3N2) | 22/7/2009 | NCBI |
| CY115632 | A/HongKong/H090-725-V10/2009(H3N2) | 20/7/2009 | NCBI |
| CY115640 | A/HongKong/H090-726-V10/2009(H3N2) | 20/7/2009 | NCBI |
| CY115648 | A/HongKong/H090-729-V10/2009(H3N2) | 21/7/2009 | NCBI |
| CY115656 | A/HongKong/H090-733-V10/2009(H3N2) | 22/7/2009 | NCBI |
| CY115664 | A/HongKong/H090-734-V10/2009(H3N2) | 22/7/2009 | NCBI |
| CY115672 | A/HongKong/H090-734-V20/2009(H3N2) | 24/7/2009 | NCBI |
| CY115680 | A/HongKong/H090-734-V32/2009(H3N2) | 27/7/2009 | NCBI |
| CY115688 | A/HongKong/H090-736-V10/2009(H3N2) | 23/7/2009 | NCBI |
| CY115696 | A/HongKong/H090-737-V10/2009(H3N2) | 23/7/2009 | NCBI |
| CY115704 | A/HongKong/H090-739-V10/2009(H3N2) | 23/7/2009 | NCBI |
| CY115712 | A/HongKong/H090-739-V22/2009(H3N2) | 27/7/2009 | NCBI |
| CY115720 | A/HongKong/H090-739-V23/2009(H3N2) | 27/7/2009 | NCBI |
| CY115728 | A/HongKong/H090-739-V33/2009(H3N2) | 29/7/2009 | NCBI |
| CY115736 | A/HongKong/H090-741-V10/2009(H3N2) | 24/7/2009 | NCBI |
| CY115744 | A/HongKong/H090-747-V10/2009(H3N2) | 28/7/2009 | NCBI |
| CY115752 | A/HongKong/H090-747-V22/2009(H3N2) | 30/7/2009 | NCBI |
| CY115760 | A/HongKong/H090-750-V10/2009(H3N2) | 29/7/2009 | NCBI |
| CY115768 | A/HongKong/H090-752-V10/2009(H3N2) | 29/7/2009 | NCBI |
| CY115776 | A/HongKong/H090-755-V31/2009(H3N2) | 5/8/2009 | NCBI |
| CY115784 | A/HongKong/H090-756-V10/2009(H3N2) | 30/7/2009 | NCBI |
| CY115792 | A/HongKong/H090-763-V10/2009(H3N2) | 31/7/2009 | NCBI |
| CY115800 | A/HongKong/H090-763-V23/2009(H3N2) | 3/8/2009 | NCBI |
| CY115808 | A/HongKong/H090-764-V10/2009(H3N2) | 31/7/2009 | NCBI |
| CY115816 | A/HongKong/H090-769-V10/2009(H3N2) | 3/8/2009 | NCBI |
| CY115824 | A/HongKong/H090-770-V10/2009(H3N2) | 3/8/2009 | NCBI |
| CY116699 | A/Boston/DOA16/2011(H3N2) | 19/1/2011 | NCBI |
| CY116707 | A/Boston/DOA24/2011(H3N2) | 1/2/2011 | NCBI |
| CY116715 | A/Boston/DOA68/2011(H3N2) | 1/3/2011 | NCBI |
| CY117581 | A/Boston/DOA42/2011(H3N2) | 14/2/2011 | NCBI |
| CY117589 | A/Boston/DOA74/2011(H3N2) | 9/3/2011 | NCBI |
| CY117698 | A/Malaysia/10081/1996(H3N2) | 24/5/1996 | NCBI |
| CY117706 | A/Malaysia/10135/1996(H3N2) | 30/5/1996 | NCBI |
| CY117728 | A/Malaysia/10879/1999(H3N2) | 2/2/1999 | NCBI |
| CY117736 | A/Malaysia/10970/1999(H3N2) | 11/2/1999 | NCBI |
| CY117752 | A/Malaysia/13241/1997(H3N2) | 24/7/1997 | NCBI |

| | | | |
|---|---|---|---|
| CY117760 | A/Malaysia/13328/1999(H3N2) | 23/12/1999 | NCBI |
| CY117768 | A/Malaysia/13329/1999(H3N2) | 23/12/1999 | NCBI |
| CY117776 | A/Malaysia/14590/2000(H3N2) | 26/6/2000 | NCBI |
| CY117784 | A/Malaysia/14712/1997(H3N2) | 20/12/1997 | NCBI |
| CY117800 | A/Malaysia/13381/1999(H3N2) | 30/12/1999 | NCBI |
| CY117980 | A/Malaysia/26227/2003(H3N2) | 7/11/2003 | NCBI |
| CY117988 | A/Malaysia/29997/2004(H3N2) | 20/10/2004 | NCBI |
| CY118004 | A/Malaysia/33817/2006(H3N2) | 24/2/2006 | NCBI |
| CY118028 | A/Malaysia/1666665/2006(H3N2) | 22/10/2006 | NCBI |
| CY118051 | A/Malaysia/1768409/2007(H3N2) | 20/4/2007 | NCBI |
| CY118067 | A/Malaysia/1777545/2007(H3N2) | 6/5/2007 | NCBI |
| CY118075 | A/Malaysia/1779027/2007(H3N2) | 10/5/2007 | NCBI |
| CY118115 | A/Malaysia/1912341/2007(H3N2) | 18/12/2007 | NCBI |
| CY118123 | A/Malaysia/1918618/2007(H3N2) | 31/12/2007 | NCBI |
| CY118131 | A/Malaysia/1959476/2008(H3N2) | 10/3/2008 | NCBI |
| CY118151 | A/Malaysia/1997829/2008(H3N2) | 12/5/2008 | NCBI |
| CY118175 | A/Malaysia/2075556/2008(H3N2) | 15/9/2008 | NCBI |
| CY118418 | A/Malaysia/07145/1995(H3N2) | 3/7/1995 | NCBI |
| CY118426 | A/Malaysia/07831/1995(H3N2) | 27/9/1995 | NCBI |
| CY118442 | A/Malaysia/10111/1996(H3N2) | 19/6/1996 | NCBI |
| CY118450 | A/Malaysia/10360/1996(H3N2) | 2/7/1996 | NCBI |
| CY118458 | A/Malaysia/10370/1996(H3N2) | 3/7/1996 | NCBI |
| CY118466 | A/Malaysia/10807/1999(H3N2) | 23/1/1999 | NCBI |
| CY118474 | A/Malaysia/10717/1999(H3N2) | 6/1/1999 | NCBI |
| CY118482 | A/Malaysia/10877/1999(H3N2) | 2/2/1999 | NCBI |
| CY118490 | A/Malaysia/10969/1999(H3N2) | 11/2/1999 | NCBI |
| CY118498 | A/Malaysia/10997/1999(H3N2) | 18/2/1999 | NCBI |
| CY118506 | A/Malaysia/11473/1996(H3N2) | 24/12/1996 | NCBI |
| CY118514 | A/Malaysia/12132/1999(H3N2) | 16/7/1999 | NCBI |
| CY118522 | A/Malaysia/12550/1999(H3N2) | 13/9/1999 | NCBI |
| CY118530 | A/Malaysia/12726/1997(H3N2) | 2/6/1997 | NCBI |
| CY118538 | A/Malaysia/12766/1999(H3N2) | 9/10/1999 | NCBI |
| CY118546 | A/Malaysia/12904/1999(H3N2) | 28/10/1999 | NCBI |
| CY118554 | A/Malaysia/12942/1999(H3N2) | 2/11/1999 | NCBI |
| CY118562 | A/Malaysia/12974/1999(H3N2) | 5/11/1999 | NCBI |
| CY118570 | A/Malaysia/12990/1999(H3N2) | 13/11/1999 | NCBI |
| CY118578 | A/Malaysia/13002/1999(H3N2) | 11/11/1999 | NCBI |

| | | | |
|---|---|---|---|
| CY118586 | A/Malaysia/13094/1999(H3N2) | 20/11/1999 | NCBI |
| CY118594 | A/Malaysia/13246/1999(H3N2) | 13/12/1999 | NCBI |
| CY118602 | A/Malaysia/17230/1998(H3N2) | 25/5/1998 | NCBI |
| CY118610 | A/Malaysia/17332/1998(H3N2) | 19/6/1998 | NCBI |
| CY118618 | A/Malaysia/17392/1998(H3N2) | 15/6/1998 | NCBI |
| CY118626 | A/Malaysia/17998/2001(H3N2) | 30/1/2001 | NCBI |
| CY118634 | A/Malaysia/10759/1999(H3N2) | 22/1/1999 | NCBI |
| CY118642 | A/Malaysia/18121/2001(H3N2) | 26/2/2001 | NCBI |
| CY118650 | A/Malaysia/18128/2001(H3N2) | 15/2/2001 | NCBI |
| CY118658 | A/Malaysia/23606/2002(H3N2) | 18/11/2002 | NCBI |
| CY118666 | A/Malaysia/25849/2003(H3N2) | 22/9/2003 | NCBI |
| CY118674 | A/Malaysia/25920/2003(H3N2) | 30/9/2003 | NCBI |
| CY118682 | A/Malaysia/25957/2003(H3N2) | 5/10/2003 | NCBI |
| CY118690 | A/Malaysia/26003/2003(H3N2) | 10/10/2003 | NCBI |
| CY118698 | A/Malaysia/26430/2003(H3N2) | 8/12/2003 | NCBI |
| CY118706 | A/Malaysia/26468/2003(H3N2) | 11/12/2003 | NCBI |
| CY118714 | A/Malaysia/26571/2003(H3N2) | 22/12/2003 | NCBI |
| CY118722 | A/Malaysia/26682/2004(H3N2) | 5/1/2004 | NCBI |
| CY118730 | A/Malaysia/26822/2004(H3N2) | 19/1/2004 | NCBI |
| CY118738 | A/Malaysia/26989/2004(H3N2) | 7/2/2004 | NCBI |
| CY118746 | A/Malaysia/28114/2004(H3N2) | 24/5/2004 | NCBI |
| CY118754 | A/Malaysia/28991/2004(H3N2) | 23/6/2004 | NCBI |
| CY118762 | A/Malaysia/29197/2004(H3N2) | 20/7/2004 | NCBI |
| CY118770 | A/Malaysia/29242/2004(H3N2) | 26/7/2004 | NCBI |
| CY118778 | A/Malaysia/29255/2004(H3N2) | 27/7/2004 | NCBI |
| CY118786 | A/Malaysia/29350/2004(H3N2) | 4/8/2004 | NCBI |
| CY118794 | A/Malaysia/29361/2004(H3N2) | 4/8/2004 | NCBI |
| CY118802 | A/Malaysia/29711/2004(H3N2) | 14/9/2004 | NCBI |
| CY118810 | A/Malaysia/29890/2004(H3N2) | 6/10/2004 | NCBI |
| CY118818 | A/Malaysia/29930/2004(H3N2) | 12/10/2004 | NCBI |
| CY118826 | A/Malaysia/31944/2005(H3N2) | 7/7/2005 | NCBI |
| CY118834 | A/Malaysia/31998/2005(H3N2) | 15/7/2005 | NCBI |
| CY118842 | A/Malaysia/32171/2005(H3N2) | 4/8/2005 | NCBI |
| CY118850 | A/Malaysia/32133/2005(H3N2) | 8/8/2005 | NCBI |
| CY118858 | A/Malaysia/32226/2005(H3N2) | 10/8/2005 | NCBI |
| CY118866 | A/Malaysia/33386/2005(H3N2) | 30/12/2005 | NCBI |
| CY118874 | A/Malaysia/33464/2006(H3N2) | 11/1/2006 | NCBI |

| | | | |
|---|---|---|---|
| CY118882 | A/Malaysia/33827/2006(H3N2) | 11/1/2006 | NCBI |
| CY118890 | A/Malaysia/33561/2006(H3N2) | 23/1/2006 | NCBI |
| CY118898 | A/Malaysia/34015/2006(H3N2) | 16/3/2006 | NCBI |
| CY118906 | A/Malaysia/1657167/2006(H3N2) | 12/10/2006 | NCBI |
| CY118914 | A/Malaysia/1665090/2006(H3N2) | 19/10/2006 | NCBI |
| CY118922 | A/Malaysia/1674395/2006(H3N2) | 15/11/2006 | NCBI |
| CY118930 | A/Malaysia/1676558/2006(H3N2) | 17/11/2006 | NCBI |
| CY118938 | A/Malaysia/1681079/2006(H3N2) | 24/11/2006 | NCBI |
| CY118946 | A/Malaysia/1755590/2007(H3N2) | 30/3/2007 | NCBI |
| CY118954 | A/Malaysia/1756502/2007(H3N2) | 2/4/2007 | NCBI |
| CY118962 | A/Malaysia/1756937/2007(H3N2) | 2/4/2007 | NCBI |
| CY118970 | A/Malaysia/1767091/2007(H3N2) | 1/4/2007 | NCBI |
| CY118978 | A/Malaysia/1768035/2007(H3N2) | 20/4/2007 | NCBI |
| CY118986 | A/Malaysia/1773914/2007(H3N2) | 1/5/2007 | NCBI |
| CY118994 | A/Malaysia/1779036/2007(H3N2) | 10/5/2007 | NCBI |
| CY119002 | A/Malaysia/1781328/2007(H3N2) | 14/5/2007 | NCBI |
| CY119010 | A/Malaysia/1912346/2007(H3N2) | 18/12/2007 | NCBI |
| CY119018 | A/Malaysia/1922209/2008(H3N2) | 7/1/2008 | NCBI |
| CY119026 | A/Malaysia/1959864/2008(H3N2) | 10/3/2008 | NCBI |
| CY119034 | A/Malaysia/1963888/2008(H3N2) | 17/3/2008 | NCBI |
| CY119042 | A/Malaysia/1965896/2008(H3N2) | 19/3/2008 | NCBI |
| CY119050 | A/Malaysia/1770921/2007(H3N2) | 25/4/2007 | NCBI |
| CY119058 | A/Malaysia/2205714/2009(H3N2) | 16/4/2009 | NCBI |
| CY119082 | A/Malaysia/12106/1999(H3N2) | 14/7/1999 | NCBI |
| CY121024 | A/Beijing/353/1989(H3N2) | 1989 | NCBI |
| CY121117 | A/Aichi/2/1968(H3N2) | 1968 | NCBI |
| CY121125 | A/NewYork/55/2004(H3N2) | 2004 | NCBI |
| CY121157 | A/England/42/1972(H3N2) | 1972 | NCBI |
| CY121181 | A/PortChalmers/1/1973(H3N2) | 1973 | NCBI |
| CY121197 | A/Victoria/3/1975(H3N2) | 1975 | NCBI |
| CY121213 | A/Iowa/119/2010(H3N2) | 2010 | NCBI |
| CY121245 | A/Philippines/2/1982(H3N2) | 1982 | NCBI |
| CY121277 | A/Leningrad/360/1986(H3N2) | 1986 | NCBI |
| CY121293 | A/Sichuan/2/1987(H3N2) | 1987 | NCBI |
| CY121309 | A/Shanghai/11/1987(H3N2) | 1987 | NCBI |
| CY121325 | A/Harbin/15/1992(H3N2) | 1992 | NCBI |
| CY121341 | A/Johannesburg/33/1994(H3N2) | 1994 | NCBI |

| | | | |
|---|---|---|---|
| CY121357 | A/SouthAfrica/1147/1996(H3N2) | 1996 | NCBI |
| CY121373 | A/Moscow/10/1999(H3N2) | 1999 | NCBI |
| CY121408 | A/UlanUde/01/2000(H3N2) | 2000 | NCBI |
| CY121424 | A/California/32/1999(H3N2) | 1999 | NCBI |
| CY121448 | A/Fujian/445/2003(H3N2) | 2003 | NCBI |
| CY121480 | A/Wellington/01/2004(H3N2) | 2004 | NCBI |
| CY121496 | A/Perth/10/2010(H3N2) | 2010 | NCBI |
| CY121512 | A/Nepal/921/2006(H3N2) | 2006 | NCBI |
| CY121528 | A/Wisconsin/03/2007(H3N2) | 2007 | NCBI |
| CY121568 | A/BRISBANE/9/2006(H3N2) | 2006 | NCBI |
| CY121632 | A/Uruguay/716/2007(H3N2) | 2007 | NCBI |
| CY121736 | A/HongKong/26560/2009(H3N2) | 2009 | NCBI |
| CY121792 | A/Brisbane/11/2010(H3N2) | 2010 | NCBI |
| CY121800 | A/RhodeIsland/01/2010(H3N2) | 2010 | NCBI |
| CY125717 | A/Boston/DOA27/2011(H3N2) | 21/2/2011 | NCBI |
| JX437828 | A/Singapore/H2009.332C/2009(H3N2) | 1/5/2009 | NCBI |
| JX437829 | A/Singapore/H2009.334C/2009(H3N2) | 2/5/2009 | NCBI |
| JX437830 | A/Singapore/H2009.471C/2009(H3N2) | 21/6/2009 | NCBI |
| JX437831 | A/Singapore/H2010.321C/2010(H3N2) | 27/4/2010 | NCBI |
| JX437832 | A/Singapore/H2010.370C/2010(H3N2) | 15/5/2010 | NCBI |
| JX437833 | A/Singapore/H2010.471C/2010(H3N2) | 21/6/2010 | NCBI |
| JX437834 | A/Singapore/H2010.559C/2010(H3N2) | 23/7/2010 | NCBI |
| JX437835 | A/Singapore/H2010.564C/2010(H3N2) | 25/7/2010 | NCBI |
| JX437836 | A/Singapore/H2010.666C/2010(H3N2) | 31/8/2010 | NCBI |
| JX437837 | A/Singapore/H2011.140C/2011(H3N2) | 20/2/2011 | NCBI |
| JX437838 | A/Singapore/H2011.460cC/2011(H3N2) | 17/6/2011 | NCBI |
| JX437839 | A/Singapore/H2011.504C/2011(H3N2) | 3/7/2011 | NCBI |
| JX437840 | A/Singapore/H2011.704C/2011(H3N2) | 14/9/2011 | NCBI |
| JX437841 | A/Singapore/H2011.808bC/2011(H3N2) | 22/10/2011 | NCBI |
| JX437842 | A/Singapore/C2011.545C/2011(H3N2) | 18/7/2011 | NCBI |
| JX437708 | A/Singapore/C2009.458V/2009(H3N2) | 16/6/2009 | NCBI |
| JX437709 | A/Singapore/C2009.518V/2009(H3N2) | 8/7/2009 | NCBI |
| JX437710 | A/Singapore/C2009.715V/2009(H3N2) | 18/9/2009 | NCBI |
| JX437711 | A/Singapore/C2009.803bV/2009(H3N2) | 20/10/2009 | NCBI |
| JX437712 | A/Singapore/C2010.036V/2010(H3N2) | 13/1/2010 | NCBI |
| JX437713 | A/Singapore/C2010.304V/2010(H3N2) | 21/4/2010 | NCBI |
| JX437714 | A/Singapore/C2010.362V/2010(H3N2) | 12/5/2010 | NCBI |

| | | | |
|---|---|---|---|
| JX437715 | A/Singapore/C2010.471aV/2010(H3N2) | 21/6/2010 | NCBI |
| JX437716 | A/Singapore/C2010.682V/2010(H3N2) | 6/6/2010 | NCBI |
| JX437717 | A/Singapore/C2010.803V/2010(H3N2) | 20/10/2010 | NCBI |
| JX437718 | A/Singapore/C2010.937V/2010(H3N2) | 8/12/2010 | NCBI |
| JX437719 | A/Singapore/C2011.027V/2011(H3N2) | 10/1/2011 | NCBI |
| JX437720 | A/Singapore/C2011.362V/2011(H3N2) | 12/5/2011 | NCBI |
| JX437721 | A/Singapore/C2011.507V/2011(H3N2) | 4/7/2011 | NCBI |
| JX437722 | A/Singapore/C2011.584V/2011(H3N2) | 1/8/2011 | NCBI |
| CY125791 | A/Boston/DOA94/2012(H3N2) | 19/4/2012 | NCBI |
| CY100546 | A/Mexico/InDRE2112/2005(H3N2) | 11/1/2005 | NCBI |
| CY100570 | A/Mexico/InDRE2227/2005(H3N2) | 11/9/2005 | NCBI |
| CY100586 | A/Mexico/InDRE2601/2003(H3N2) | 2003 | NCBI |
| CY100602 | A/Mexico/InDRE29/2006(H3N2) | 16/1/2006 | NCBI |
| CY100618 | A/Mexico/InDRE835/2003(H3N2) | 4/7/2003 | NCBI |
| CY100626 | A/Mexico/InDRE940/2003(H3N2) | 2003 | NCBI |
| CY134481 | A/Boston/DOA29/2011(H3N2) | 27/1/2011 | NCBI |
| CY134505 | A/Boston/DOA36/2011(H3N2) | 10/2/2011 | NCBI |
| CY134513 | A/Boston/DOA38/2011(H3N2) | 12/2/2011 | NCBI |
| CY134529 | A/Boston/DOA41/2011(H3N2) | 13/2/2011 | NCBI |
| CY134545 | A/Boston/DOA45/2011(H3N2) | 15/2/2011 | NCBI |
| CY134561 | A/Boston/DOA47/2011(H3N2) | 14/2/2011 | NCBI |
| CY134577 | A/Boston/DOA50/2011(H3N2) | 18/2/2011 | NCBI |
| CY134593 | A/Boston/DOA54/2011(H3N2) | 22/2/2011 | NCBI |
| CY134601 | A/Boston/DOA60/2011(H3N2) | 24/2/2011 | NCBI |
| CY134609 | A/Boston/DOA63/2011(H3N2) | 25/2/2011 | NCBI |
| CY134748 | A/Texas/JMM_3/2012(H3N2) | 11/11/2012 | NCBI |
| CY134756 | A/Texas/JMM_4/2012(H3N2) | 12/11/2012 | NCBI |
| CY134764 | A/Texas/JMM_5/2012(H3N2) | 14/11/2012 | NCBI |
| CY134772 | A/Texas/JMM_8/2012(H3N2) | 20/11/2012 | NCBI |
| CY134780 | A/Texas/JMM_9/2012(H3N2) | 20/11/2012 | NCBI |
| CY134788 | A/Texas/JMM_10/2012(H3N2) | 20/11/2012 | NCBI |
| CY134796 | A/Texas/JMM_11/2012(H3N2) | 20/11/2012 | NCBI |
| CY134804 | A/Texas/JMM_12/2012(H3N2) | 20/11/2012 | NCBI |
| CY134812 | A/Texas/JMM_13/2012(H3N2) | 20/11/2012 | NCBI |
| CY134820 | A/Texas/JMM_14/2012(H3N2) | 21/11/2012 | NCBI |
| CY134828 | A/Texas/JMM_15/2012(H3N2) | 22/11/2012 | NCBI |
| CY134836 | A/Texas/JMM_16/2012(H3N2) | 24/11/2012 | NCBI |

| | | | |
|---|---|---|---|
| CY134844 | A/Texas/JMM_17/2012(H3N2) | 24/11/2012 | NCBI |
| CY134852 | A/Texas/JMM_18/2012(H3N2) | 24/11/2012 | NCBI |
| CY134860 | A/Texas/JMM_19/2012(H3N2) | 25/11/2012 | NCBI |
| CY134868 | A/Texas/JMM_20/2012(H3N2) | 25/11/2012 | NCBI |
| CY134876 | A/Texas/JMM_21/2012(H3N2) | 25/11/2012 | NCBI |
| CY134884 | A/Texas/JMM_22/2012(H3N2) | 25/11/2012 | NCBI |
| CY134892 | A/Texas/JMM_23/2012(H3N2) | 25/11/2012 | NCBI |
| CY134900 | A/Texas/JMM_24/2012(H3N2) | 26/11/2012 | NCBI |
| CY134908 | A/Texas/JMM_25/2012(H3N2) | 26/11/2012 | NCBI |
| CY134916 | A/Texas/JMM_26/2012(H3N2) | 26/11/2012 | NCBI |
| CY134924 | A/Texas/JMM_27/2012(H3N2) | 28/11/2012 | NCBI |
| CY134932 | A/Texas/JMM_29/2012(H3N2) | 28/11/2012 | NCBI |
| CY134940 | A/Texas/JMM_30/2012(H3N2) | 28/11/2012 | NCBI |
| CY134948 | A/Texas/JMM_31/2012(H3N2) | 28/11/2012 | NCBI |
| CY134956 | A/Texas/JMM_32/2012(H3N2) | 28/11/2012 | NCBI |
| CY134964 | A/Texas/JMM_33/2012(H3N2) | 29/11/2012 | NCBI |
| CY134972 | A/Texas/JMM_34/2012(H3N2) | 29/11/2012 | NCBI |
| CY134980 | A/Texas/JMM_35/2012(H3N2) | 29/11/2012 | NCBI |
| CY134988 | A/Texas/JMM_36/2012(H3N2) | 29/11/2012 | NCBI |
| CY134996 | A/Texas/JMM_37/2012(H3N2) | 30/11/2012 | NCBI |
| CY135004 | A/Texas/JMM_38/2012(H3N2) | 30/11/2012 | NCBI |
| CY135012 | A/Texas/JMM_39/2012(H3N2) | 30/11/2012 | NCBI |
| CY135020 | A/Texas/JMM_40/2012(H3N2) | 1/12/2012 | NCBI |
| CY135028 | A/Texas/JMM_41/2012(H3N2) | 1/12/2012 | NCBI |
| CY135036 | A/Texas/JMM_43/2012(H3N2) | 2/12/2012 | NCBI |
| CY135044 | A/Texas/JMM_44/2012(H3N2) | 2/12/2012 | NCBI |
| CY135052 | A/Texas/JMM_45/2012(H3N2) | 3/12/2012 | NCBI |
| CY135060 | A/Texas/JMM_46/2012(H3N2) | 4/12/2012 | NCBI |
| CY135068 | A/Texas/JMM_47/2012(H3N2) | 4/12/2012 | NCBI |
| CY135076 | A/Texas/JMM_48/2012(H3N2) | 4/12/2012 | NCBI |
| CY135084 | A/Texas/JMM_49/2012(H3N2) | 6/12/2012 | NCBI |
| CY135092 | A/Texas/JMM_50/2012(H3N2) | 6/12/2012 | NCBI |
| CY135100 | A/Texas/JMM_51/2012(H3N2) | 6/12/2012 | NCBI |
| CY135124 | A/Texas/JMM_54/2012(H3N2) | 6/12/2012 | NCBI |
| CY135132 | A/Texas/JMM_56/2012(H3N2) | 7/12/2012 | NCBI |
| CY135140 | A/Texas/JMM_57/2012(H3N2) | 7/12/2012 | NCBI |
| CY135148 | A/Texas/JMM_58/2012(H3N2) | 7/12/2012 | NCBI |

| | | | |
|---|---|---|---|
| CY135156 | A/Texas/JMM_59/2012(H3N2) | 8/12/2012 | NCBI |
| CY135164 | A/Texas/JMM_60/2012(H3N2) | 8/12/2012 | NCBI |
| CY134732 | A/Texas/JMM_1/2012(H3N2) | 3/11/2012 | NCBI |
| CY134740 | A/Texas/JMM_2/2012(H3N2) | 4/11/2012 | NCBI |
| KC882439 | A/Iowa/19/2010(H3N2) | 30/12/2010 | NCBI |
| KC882860 | A/Iowa/19/2010(H3N2) | 30/12/2010 | NCBI |
| KC882479 | A/Texas/24/2010(H3N2) | 31/12/2010 | NCBI |
| KC882563 | A/NewJersey/01/2011(H3N2) | 11/1/2011 | NCBI |
| KC882639 | A/Texas/06/2011(H3N2) | 5/2/2011 | NCBI |
| KC882908 | A/California/16/2011(H3N2) | 16/1/2011 | NCBI |
| KC883049 | A/Wisconsin/22/2011(H3N2) | 15/4/2011 | NCBI |
| KC883070 | A/Illinois/03/2011(H3N2) | 22/1/2011 | NCBI |
| KC883233 | A/Massachusetts/06/2010(H3N2) | 31/12/2010 | NCBI |
| KC883402 | A/Alaska/15/2011(H3N2) | 1/3/2011 | NCBI |
| KC892437 | A/California/17/2012(H3N2) | 25/1/2012 | NCBI |
| KC892845 | A/Nevada/12/2012(H3N2) | 30/3/2012 | NCBI |
| KC892444 | A/Washington/16/2012(H3N2) | 2/3/2012 | NCBI |
| KC892829 | A/California/22/2012(H3N2) | 20/2/2012 | NCBI |
| KC892822 | A/NewMexico/07/2012(H3N2) | 16/2/2012 | NCBI |
| KC892838 | A/Michigan/03/2012(H3N2) | 6/3/2012 | NCBI |
| KF014197 | A/Singapore/C2009.496/2009(H3N2) | 30/6/2009 | NCBI |
| KF014199 | A/Singapore/C2009.485a/2009(H3N2) | 26/6/2009 | NCBI |
| KF014200 | A/Singapore/C2009.485b/2009(H3N2) | 26/6/2009 | NCBI |
| KF014201 | A/Singapore/C2009.515/2009(H3N2) | 7/7/2009 | NCBI |
| KF014202 | A/Singapore/C2009.863/2009(H3N2) | 11/11/2009 | NCBI |
| KF014205 | A/Singapore/C2011.173/2011(H3N2) | 4/3/2011 | NCBI |
| KF014206 | A/Singapore/C2011.244/2011(H3N2) | 30/3/2011 | NCBI |
| KF014207 | A/Singapore/C2011.301/2011(H3N2) | 20/4/2011 | NCBI |
| KF014209 | A/Singapore/C2011.384/2011(H3N2) | 20/5/2011 | NCBI |
| KF014210 | A/Singapore/C2011.411/2011(H3N2) | 30/5/2011 | NCBI |
| KF014211 | A/Singapore/C2011.422/2011(H3N2) | 3/6/2011 | NCBI |
| KF014212 | A/Singapore/C2011.452/2011(H3N2) | 14/6/2011 | NCBI |
| KF014213 | A/Singapore/C2011.458/2011(H3N2) | 16/6/2011 | NCBI |
| KF014214 | A/Singapore/C2011.471/2011(H3N2) | 21/6/2011 | NCBI |
| KF014215 | A/Singapore/C2011.477/2011(H3N2) | 23/6/2011 | NCBI |
| KF014216 | A/Singapore/C2011.493/2011(H3N2) | 29/6/2011 | NCBI |
| KF014217 | A/Singapore/C2011.496/2011(H3N2) | 30/6/2011 | NCBI |

| | | | |
|---|---|---|---|
| KF014219 | A/Singapore/C2011.512/2011(H3N2) | 6/7/2011 | NCBI |
| KF014220 | A/Singapore/C2011.518/2011(H3N2) | 8/7/2011 | NCBI |
| KF014221 | A/Singapore/C2011.564/2011(H3N2) | 25/7/2011 | NCBI |
| KF014222 | A/Singapore/C2011.573/2011(H3N2) | 28/7/2011 | NCBI |
| KF014224 | A/Singapore/C2011.614/2011(H3N2) | 12/8/2011 | NCBI |
| KF014225 | A/Singapore/C2011.641/2011(H3N2) | 22/8/2011 | NCBI |
| KF014226 | A/Singapore/C2011.647/2011(H3N2) | 24/8/2011 | NCBI |
| KF014227 | A/Singapore/C2011.803/2011(H3N2) | 20/10/2011 | NCBI |
| KF014228 | A/Singapore/C2011.825/2011(H3N2) | 28/10/2011 | NCBI |
| KF014229 | A/Singapore/H2009.389b/2009(H3N2) | 22/5/2009 | NCBI |
| KF014230 | A/Singapore/H2009.485/2009(H3N2) | 26/6/2009 | NCBI |
| KF014231 | A/Singapore/H2009.510/2009(H3N2) | 5/7/2009 | NCBI |
| KF014232 | A/Singapore/H2009.518/2009(H3N2) | 8/7/2009 | NCBI |
| KF014233 | A/Singapore/H2009.679/2009(H3N2) | 5/9/2009 | NCBI |
| KF014234 | A/Singapore/H2010.211/2010(H3N2) | 18/3/2010 | NCBI |
| KF014235 | A/Singapore/H2010.301/2010(H3N2) | 20/4/2010 | NCBI |
| KF014236 | A/Singapore/H2010.384/2010(H3N2) | 20/5/2010 | NCBI |
| KF014237 | A/Singapore/H2010.389/2010(H3N2) | 22/5/2010 | NCBI |
| KF014238 | A/Singapore/H2010.619/2010(H3N2) | 14/8/2010 | NCBI |
| KF014239 | A/Singapore/H2010.797/2010(H3N2) | 18/10/2010 | NCBI |
| KF014240 | A/Singapore/H2010.822/2010(H3N2) | 27/10/2010 | NCBI |
| KF014241 | A/Singapore/H2011.447/2011(H3N2) | 12/6/2011 | NCBI |
| KF014242 | A/Singapore/H2011.460a/2011(H3N2) | 17/6/2011 | NCBI |
| KF014243 | A/Singapore/H2011.463/2011(H3N2) | 18/6/2011 | NCBI |
| KF014244 | A/Singapore/H2011.471a/2011(H3N2) | 21/6/2011 | NCBI |
| KF014245 | A/Singapore/H2011.471b/2011(H3N2) | 21/6/2011 | NCBI |
| KF014246 | A/Singapore/H2011.471c/2011(H3N2) | 21/6/2011 | NCBI |
| KF014247 | A/Singapore/H2011.474/2011(H3N2) | 22/6/2011 | NCBI |
| KF014248 | A/Singapore/H2011.479/2011(H3N2) | 24/6/2011 | NCBI |
| KF014249 | A/Singapore/H2011.482/2011(H3N2) | 25/6/2011 | NCBI |
| KF014250 | A/Singapore/H2011.496/2011(H3N2) | 30/6/2011 | NCBI |
| KF014251 | A/Singapore/H2011.499a/2011(H3N2) | 1/7/2011 | NCBI |
| KF014252 | A/Singapore/H2011.507/2011(H3N2) | 4/7/2011 | NCBI |
| KF014253 | A/Singapore/H2011.515/2011(H3N2) | 7/7/2011 | NCBI |
| KF014254 | A/Singapore/H2011.518/2011(H3N2) | 8/7/2011 | NCBI |
| KF014255 | A/Singapore/H2011.570/2011(H3N2) | 27/7/2011 | NCBI |
| KF014256 | A/Singapore/H2011.751/2011(H3N2) | 1/10/2011 | NCBI |

| | | | |
|---|---|---|---|
| KF014257 | A/Singapore/H2011.797/2011(H3N2) | 18/10/2011 | NCBI |
| KF014258 | A/Singapore/H2011.808a/2011(H3N2) | 22/10/2011 | NCBI |
| CY146809 | A/Victoria/JY2/1968(H3N2) | 1968 | NCBI |
| CY147430 | A/PortChalmers/JY2/1973(H3N2) | 1973 | NCBI |
| CY147438 | A/HongKong/JY2/1968(H3N2) | 1968 | NCBI |
| CY147446 | A/NorthernTerritories/60-JY2/1968(H3N2) | 1968 | NCBI |
| CY147579 | A/Mexico/24003/2008(H3N2) | 1/11/2008 | NCBI |
| CY147619 | A/Mexico/24008/2009(H3N2) | 1/4/2009 | NCBI |
| CY147627 | A/Mexico/24009/2008(H3N2) | 1/11/2008 | NCBI |
| CY147635 | A/Mexico/24010/2008(H3N2) | 1/11/2008 | NCBI |
| CY147643 | A/Mexico/24011/2008(H3N2) | 1/11/2008 | NCBI |
| CY147651 | A/Mexico/24012/2008(H3N2) | 2/12/2008 | NCBI |
| CY147659 | A/Mexico/24013/2008(H3N2) | 1/12/2008 | NCBI |
| CY147667 | A/Mexico/24014/2008(H3N2) | 1/12/2008 | NCBI |
| CY147723 | A/Mexico/24025/2009(H3N2) | 27/4/2009 | NCBI |
| KF432083 | A/Singapore/H2013.060/2013(H3N2) | 22/1/2013 | NCBI |
| CY148292 | A/Boston/DOA2-095/2012(H3N2) | 12/9/2012 | NCBI |
| CY148300 | A/Boston/DOA2-097/2012(H3N2) | 12/11/2012 | NCBI |
| CY148308 | A/Boston/DOA2-098/2012(H3N2) | 12/12/2012 | NCBI |
| CY148324 | A/Boston/DOA2-100/2012(H3N2) | 13/12/2012 | NCBI |
| CY148332 | A/Boston/DOA2-101/2012(H3N2) | 14/12/2012 | NCBI |
| CY148340 | A/Boston/DOA2-102/2012(H3N2) | 15/12/2012 | NCBI |
| CY148348 | A/Boston/DOA2-103/2012(H3N2) | 17/12/2012 | NCBI |
| CY148356 | A/Boston/DOA2-104/2012(H3N2) | 22/12/2012 | NCBI |
| CY148364 | A/Boston/DOA2-105/2012(H3N2) | 21/12/2012 | NCBI |
| CY148372 | A/Boston/DOA2-106/2012(H3N2) | 14/11/2012 | NCBI |
| CY148380 | A/Boston/DOA2-107/2012(H3N2) | 1/11/2012 | NCBI |
| CY148388 | A/Boston/DOA2-108/2012(H3N2) | 11/10/2012 | NCBI |
| CY148396 | A/Boston/DOA2-109/2012(H3N2) | 13/9/2012 | NCBI |
| CY148404 | A/Boston/DOA2-110/2012(H3N2) | 7/12/2012 | NCBI |
| CY148412 | A/Boston/DOA2-111/2012(H3N2) | 2/12/2012 | NCBI |
| CY148420 | A/Boston/DOA2-112/2012(H3N2) | 4/12/2012 | NCBI |
| CY148428 | A/Boston/DOA2-113/2012(H3N2) | 2/12/2012 | NCBI |
| CY148436 | A/Boston/DOA2-114/2012(H3N2) | 3/12/2012 | NCBI |
| CY148444 | A/Boston/DOA2-115/2012(H3N2) | 5/12/2012 | NCBI |
| CY148452 | A/Boston/DOA2-116/2012(H3N2) | 3/12/2012 | NCBI |
| CY148460 | A/Boston/DOA2-117/2012(H3N2) | 7/12/2012 | NCBI |

| | | | |
|---|---|---|---|
| CY148468 | A/Boston/DOA2-119/2012(H3N2) | 24/12/2012 | NCBI |
| CY148476 | A/Boston/DOA2-120/2012(H3N2) | 24/12/2012 | NCBI |
| CY148484 | A/Boston/DOA2-121/2012(H3N2) | 24/12/2012 | NCBI |
| CY148492 | A/Boston/DOA2-122/2012(H3N2) | 26/12/2012 | NCBI |
| CY148500 | A/Boston/DOA2-123/2012(H3N2) | 26/12/2012 | NCBI |
| CY148508 | A/Boston/DOA2-124/2012(H3N2) | 26/12/2012 | NCBI |
| CY148516 | A/Boston/DOA2-125/2012(H3N2) | 26/12/2012 | NCBI |
| CY148524 | A/Boston/DOA2-126/2012(H3N2) | 16/12/2012 | NCBI |
| CY148532 | A/Boston/DOA2-127/2012(H3N2) | 17/12/2012 | NCBI |
| CY148540 | A/Boston/DOA2-128/2012(H3N2) | 18/12/2012 | NCBI |
| CY148548 | A/Boston/DOA2-129/2012(H3N2) | 18/12/2012 | NCBI |
| CY148556 | A/Boston/DOA2-130/2012(H3N2) | 18/12/2012 | NCBI |
| CY148564 | A/Boston/DOA2-131/2012(H3N2) | 19/12/2012 | NCBI |
| CY148572 | A/Boston/DOA2-132/2012(H3N2) | 19/12/2012 | NCBI |
| CY148580 | A/Boston/DOA2-133/2012(H3N2) | 20/12/2012 | NCBI |
| CY148588 | A/Boston/DOA2-134/2012(H3N2) | 20/12/2012 | NCBI |
| CY148596 | A/Boston/DOA2-135/2012(H3N2) | 20/12/2012 | NCBI |
| CY148604 | A/Boston/DOA2-136/2012(H3N2) | 20/12/2012 | NCBI |
| CY148612 | A/Boston/DOA2-137/2012(H3N2) | 21/12/2012 | NCBI |
| CY148620 | A/Boston/DOA2-138/2012(H3N2) | 20/12/2012 | NCBI |
| CY148628 | A/Boston/DOA2-140/2012(H3N2) | 10/12/2012 | NCBI |
| CY148644 | A/Boston/DOA2-142/2012(H3N2) | 21/12/2012 | NCBI |
| CY148652 | A/Boston/DOA2-143/2012(H3N2) | 18/12/2012 | NCBI |
| CY148660 | A/Boston/DOA2-144/2013(H3N2) | 16/1/2013 | NCBI |
| CY148668 | A/Boston/DOA2-145/2013(H3N2) | 20/1/2013 | NCBI |
| CY148676 | A/Boston/DOA2-146/2013(H3N2) | 20/1/2013 | NCBI |
| CY148684 | A/Boston/DOA2-147/2012(H3N2) | 30/12/2012 | NCBI |
| CY148692 | A/Boston/DOA2-148/2013(H3N2) | 8/1/2013 | NCBI |
| CY148700 | A/Boston/DOA2-149/2013(H3N2) | 9/1/2013 | NCBI |
| CY148708 | A/Boston/DOA2-150/2013(H3N2) | 5/1/2013 | NCBI |
| CY148716 | A/Boston/DOA2-151/2013(H3N2) | 10/1/2013 | NCBI |
| CY148724 | A/Boston/DOA2-153/2013(H3N2) | 18/1/2013 | NCBI |
| CY148732 | A/Boston/DOA2-154/2013(H3N2) | 16/1/2013 | NCBI |
| CY148740 | A/Boston/DOA2-155/2012(H3N2) | 29/12/2012 | NCBI |
| CY148748 | A/Boston/DOA2-156/2013(H3N2) | 13/1/2013 | NCBI |
| CY148756 | A/Boston/DOA2-157/2013(H3N2) | 9/1/2013 | NCBI |
| CY148764 | A/Boston/DOA2-158/2013(H3N2) | 3/1/2013 | NCBI |

| | | | |
|---|---|---|---|
| CY148772 | A/Boston/DOA2-159/2013(H3N2) | 10/1/2013 | NCBI |
| CY148780 | A/Boston/DOA2-161/2013(H3N2) | 2/1/2013 | NCBI |
| CY148788 | A/Boston/DOA2-162/2012(H3N2) | 10/1/2012 | NCBI |
| CY148796 | A/Boston/DOA2-163/2013(H3N2) | 11/1/2013 | NCBI |
| CY148804 | A/Boston/DOA2-166/2013(H3N2) | 4/1/2013 | NCBI |
| CY148812 | A/Boston/DOA2-167/2012(H3N2) | 31/12/2012 | NCBI |
| CY148820 | A/Boston/DOA2-168/2012(H3N2) | 18/12/2012 | NCBI |
| CY148828 | A/Boston/DOA2-169/2012(H3N2) | 31/12/2012 | NCBI |
| CY148836 | A/Boston/DOA2-170/2013(H3N2) | 12/1/2013 | NCBI |
| CY148844 | A/Boston/DOA2-171/2012(H3N2) | 29/12/2012 | NCBI |
| CY148852 | A/Boston/DOA2-172/2013(H3N2) | 17/1/2013 | NCBI |
| CY148860 | A/Boston/DOA2-173/2013(H3N2) | 6/1/2013 | NCBI |
| CY148868 | A/Boston/DOA2-174/2013(H3N2) | 4/1/2013 | NCBI |
| CY148876 | A/Boston/DOA2-175/2013(H3N2) | 6/1/2013 | NCBI |
| CY148884 | A/Boston/DOA2-176/2013(H3N2) | 2/1/2013 | NCBI |
| CY148892 | A/Boston/DOA2-177/2013(H3N2) | 7/1/2013 | NCBI |
| CY148900 | A/Boston/DOA2-178/2013(H3N2) | 8/1/2013 | NCBI |
| CY148908 | A/Boston/DOA2-179/2013(H3N2) | 15/1/2013 | NCBI |
| CY148916 | A/Boston/DOA2-180/2013(H3N2) | 13/1/2013 | NCBI |
| CY148924 | A/Boston/DOA2-181/2013(H3N2) | 18/1/2013 | NCBI |
| CY148932 | A/Boston/DOA2-182/2013(H3N2) | 8/1/2013 | NCBI |
| CY148940 | A/Boston/DOA2-183/2013(H3N2) | 9/1/2013 | NCBI |
| CY148948 | A/Boston/DOA2-184/2013(H3N2) | 7/1/2013 | NCBI |
| CY148956 | A/Boston/DOA2-185/2013(H3N2) | 6/1/2013 | NCBI |
| CY148964 | A/Boston/DOA2-186/2013(H3N2) | 11/1/2013 | NCBI |
| CY148972 | A/Boston/DOA2-187/2013(H3N2) | 16/1/2013 | NCBI |
| CY148980 | A/Boston/DOA2-189/2013(H3N2) | 14/1/2013 | NCBI |
| CY148996 | A/Boston/DOA2-191/2013(H3N2) | 2/1/2013 | NCBI |
| CY149004 | A/Boston/DOA2-192/2013(H3N2) | 1/1/2013 | NCBI |
| CY149012 | A/Boston/DOA2-194/2013(H3N2) | 17/1/2013 | NCBI |
| CY149020 | A/Boston/DOA2-196/2013(H3N2) | 23/1/2013 | NCBI |
| CY149028 | A/Boston/DOA2-199/2013(H3N2) | 17/1/2013 | NCBI |
| CY149036 | A/Boston/DOA2-201/2013(H3N2) | 15/1/2013 | NCBI |
| CY149044 | A/Boston/DOA2-202/2013(H3N2) | 11/1/2013 | NCBI |
| CY149052 | A/Boston/DOA2-203/2012(H3N2) | 29/12/2012 | NCBI |
| CY149060 | A/Boston/DOA2-206/2013(H3N2) | 18/1/2013 | NCBI |
| CY149068 | A/Boston/DOA2-207/2013(H3N2) | 3/1/2013 | NCBI |

| | | | |
|---|---|---|---|
| CY149076 | A/Boston/DOA2-209/2012(H3N2) | 28/12/2012 | NCBI |
| CY149084 | A/Boston/DOA2-212/2013(H3N2) | 9/1/2013 | NCBI |
| CY149092 | A/Boston/DOA2-213/2013(H3N2) | 22/1/2013 | NCBI |
| CY149100 | A/Boston/DOA2-214/2013(H3N2) | 7/1/2013 | NCBI |
| CY149108 | A/Boston/DOA2-215/2013(H3N2) | 15/1/2013 | NCBI |
| CY149116 | A/Boston/DOA2-217/2013(H3N2) | 15/1/2013 | NCBI |
| CY149124 | A/Boston/DOA2-218/2013(H3N2) | 15/1/2013 | NCBI |
| CY149132 | A/Boston/DOA2-219/2012(H3N2) | 29/12/2012 | NCBI |
| CY149140 | A/Boston/DOA2-220/2013(H3N2) | 7/1/2013 | NCBI |
| CY149148 | A/Boston/DOA2-221/2013(H3N2) | 23/1/2013 | NCBI |
| CY149156 | A/Boston/DOA2-222/2012(H3N2) | 28/12/2012 | NCBI |
| CY149164 | A/Boston/DOA2-223/2012(H3N2) | 30/12/2012 | NCBI |
| CY149172 | A/Boston/DOA2-224/2013(H3N2) | 1/1/2013 | NCBI |
| CY149180 | A/Boston/DOA2-225/2013(H3N2) | 22/1/2013 | NCBI |
| CY149188 | A/Boston/DOA2-226/2013(H3N2) | 5/1/2013 | NCBI |
| CY149196 | A/Boston/DOA2-227/2013(H3N2) | 11/1/2013 | NCBI |
| CY149204 | A/Boston/DOA2-228/2013(H3N2) | 5/1/2013 | NCBI |
| CY149212 | A/Boston/DOA2-229/2013(H3N2) | 11/1/2013 | NCBI |
| CY149220 | A/Boston/DOA2-230/2013(H3N2) | 17/1/2013 | NCBI |
| CY149228 | A/Boston/DOA2-232/2013(H3N2) | 10/1/2013 | NCBI |
| CY149236 | A/Boston/DOA2-233/2013(H3N2) | 3/1/2013 | NCBI |
| CY149244 | A/Boston/DOA2-235/2013(H3N2) | 7/1/2013 | NCBI |
| CY149252 | A/Boston/DOA2-236/2013(H3N2) | 10/1/2013 | NCBI |
| CY149260 | A/Boston/DOA2-238/2013(H3N2) | 4/1/2013 | NCBI |
| CY149268 | A/Boston/DOA2-239/2013(H3N2) | 9/1/2013 | NCBI |
| CY149276 | A/Boston/DOA2-240/2013(H3N2) | 8/1/2013 | NCBI |
| CY149284 | A/Boston/DOA2-241/2013(H3N2) | 4/1/2013 | NCBI |
| CY149292 | A/Boston/DOA2-242/2013(H3N2) | 4/1/2013 | NCBI |
| CY149300 | A/Boston/DOA2-243/2013(H3N2) | 10/1/2013 | NCBI |
| CY149308 | A/Boston/DOA2-244/2013(H3N2) | 28/1/2013 | NCBI |
| CY149316 | A/Boston/DOA2-245/2013(H3N2) | 25/1/2013 | NCBI |
| CY149324 | A/Boston/DOA2-246/2013(H3N2) | 26/1/2013 | NCBI |
| CY148636 | A/Boston/DOA2-141/2012(H3N2) | 21/12/2012 | NCBI |
| KF789787 | A/Michigan/32/2012(H3N2) | 18/11/2012 | NCBI |
| KF790407 | A/Alaska/02/2013(H3N2) | 20/2/2013 | NCBI |
| KF789782 | A/Michigan/33/2012(H3N2) | 3/12/2012 | NCBI |
| KF789847 | A/Washington/07/2013(H3N2) | 7/1/2013 | NCBI |

| | | | |
|---|---|---|---|
| CY116589 | A/Netherlands/178/1995(H3N2) | 1995 | NCBI |
| KC296480 | A/Bilthoven/16190/1968(H3N2) | 1968 | NCBI |
| KC296467 | A/Bilthoven/2271/1976(H3N2) | 1976 | NCBI |
| CY160288 | A/Peru/PER001/2010(H3N2) | 25/10/2010 | NCBI |
| CY160296 | A/Peru/PER002/2010(H3N2) | 19/11/2010 | NCBI |
| CY160304 | A/Peru/PER003/2010(H3N2) | 30/10/2010 | NCBI |
| CY160312 | A/Peru/PER004/2010(H3N2) | 20/11/2010 | NCBI |
| CY160320 | A/Peru/PER005/2010(H3N2) | 30/10/2010 | NCBI |
| CY160328 | A/Peru/PER006/2011(H3N2) | 25/8/2011 | NCBI |
| CY160336 | A/Peru/PER007/2010(H3N2) | 8/11/2010 | NCBI |
| CY160344 | A/Peru/PER008/2010(H3N2) | 22/10/2010 | NCBI |
| CY160352 | A/Peru/PER009/2010(H3N2) | 26/10/2010 | NCBI |
| CY160360 | A/Peru/PER010/2010(H3N2) | 2/11/2010 | NCBI |
| CY160368 | A/Peru/PER011/2010(H3N2) | 13/11/2010 | NCBI |
| CY160376 | A/Peru/PER012/2010(H3N2) | 13/12/2010 | NCBI |
| CY160384 | A/Peru/PER013/2010(H3N2) | 24/9/2010 | NCBI |
| CY160392 | A/Peru/PER014/2011(H3N2) | 4/8/2011 | NCBI |
| CY160400 | A/Peru/PER015/2010(H3N2) | 26/11/2010 | NCBI |
| CY160408 | A/Peru/PER016/2011(H3N2) | 17/10/2011 | NCBI |
| CY160416 | A/Peru/PER017/2011(H3N2) | 10/9/2011 | NCBI |
| CY160424 | A/Peru/PER018/2012(H3N2) | 7/3/2012 | NCBI |
| CY160432 | A/Peru/PER019/2010(H3N2) | 23/11/2010 | NCBI |
| CY160440 | A/Peru/PER020/2011(H3N2) | 23/10/2011 | NCBI |
| CY160448 | A/Peru/PER021/2010(H3N2) | 28/10/2010 | NCBI |
| CY160456 | A/Peru/PER022/2010(H3N2) | 24/11/2010 | NCBI |
| CY160464 | A/Peru/PER023/2010(H3N2) | 6/12/2010 | NCBI |
| CY160472 | A/Peru/PER024/2010(H3N2) | 22/10/2010 | NCBI |
| CY160480 | A/Peru/PER025/2010(H3N2) | 20/11/2010 | NCBI |
| CY160488 | A/Peru/PER026/2010(H3N2) | 6/10/2010 | NCBI |
| CY160496 | A/Peru/PER027/2011(H3N2) | 23/8/2011 | NCBI |
| CY160504 | A/Peru/PER028/2011(H3N2) | 27/8/2011 | NCBI |
| CY160512 | A/Peru/PER029/2010(H3N2) | 22/10/2010 | NCBI |
| CY160520 | A/Peru/PER030/2010(H3N2) | 20/11/2010 | NCBI |
| CY160528 | A/Peru/PER031/2012(H3N2) | 27/7/2012 | NCBI |
| CY160536 | A/Peru/PER032/2010(H3N2) | 28/10/2010 | NCBI |
| CY160544 | A/Peru/PER033/2010(H3N2) | 16/11/2010 | NCBI |
| CY160552 | A/Peru/PER034/2011(H3N2) | 12/10/2011 | NCBI |

| | | | |
|---|---|---|---|
| CY160560 | A/Peru/PER035/2010(H3N2) | 20/10/2010 | NCBI |
| CY160568 | A/Peru/PER036/2011(H3N2) | 29/8/2011 | NCBI |
| CY160576 | A/Peru/PER037/2012(H3N2) | 23/5/2012 | NCBI |
| CY160584 | A/Peru/PER038/2010(H3N2) | 11/10/2010 | NCBI |
| CY160592 | A/Peru/PER039/2011(H3N2) | 2/9/2011 | NCBI |
| CY160600 | A/Peru/PER040/2010(H3N2) | 18/10/2010 | NCBI |
| CY160608 | A/Peru/PER041/2011(H3N2) | 25/8/2011 | NCBI |
| CY160616 | A/Peru/PER042/2012(H3N2) | 2/5/2012 | NCBI |
| CY160624 | A/Peru/PER043/2011(H3N2) | 26/8/2011 | NCBI |
| CY160632 | A/Peru/PER044/2011(H3N2) | 3/9/2011 | NCBI |
| CY160648 | A/Peru/PER046/2010(H3N2) | 3/11/2010 | NCBI |
| CY160656 | A/Peru/PER047/2011(H3N2) | 6/9/2011 | NCBI |
| CY160664 | A/Peru/PER048/2010(H3N2) | 30/9/2010 | NCBI |
| CY160672 | A/Peru/PER049/2011(H3N2) | 20/9/2011 | NCBI |
| CY160680 | A/Peru/PER050/2010(H3N2) | 6/10/2010 | NCBI |
| CY160688 | A/Peru/PER051/2012(H3N2) | 5/3/2012 | NCBI |
| CY160696 | A/Peru/PER052/2010(H3N2) | 23/11/2010 | NCBI |
| CY160704 | A/Peru/PER053/2010(H3N2) | 17/9/2010 | NCBI |
| CY160712 | A/Peru/PER054/2010(H3N2) | 15/11/2010 | NCBI |
| CY160720 | A/Peru/PER055/2011(H3N2) | 7/1/2011 | NCBI |
| CY160728 | A/Peru/PER056/2011(H3N2) | 23/8/2011 | NCBI |
| CY160736 | A/Peru/PER057/2010(H3N2) | 24/11/2010 | NCBI |
| CY160744 | A/Peru/PER058/2010(H3N2) | 13/10/2010 | NCBI |
| CY160752 | A/Peru/PER059/2010(H3N2) | 23/10/2010 | NCBI |
| CY160760 | A/Peru/PER060/2010(H3N2) | 29/10/2010 | NCBI |
| CY160768 | A/Peru/PER061/2011(H3N2) | 2/9/2011 | NCBI |
| CY160776 | A/Peru/PER062/2010(H3N2) | 4/11/2010 | NCBI |
| CY160784 | A/Peru/PER063/2012(H3N2) | 2/5/2012 | NCBI |
| CY160792 | A/Peru/PER064/2010(H3N2) | 26/10/2010 | NCBI |
| CY160800 | A/Peru/PER065/2010(H3N2) | 4/11/2010 | NCBI |
| CY160808 | A/Peru/PER066/2011(H3N2) | 15/11/2011 | NCBI |
| CY160816 | A/Peru/PER067/2011(H3N2) | 31/10/2011 | NCBI |
| CY160824 | A/Peru/PER068/2010(H3N2) | 13/12/2010 | NCBI |
| CY160832 | A/Peru/PER069/2011(H3N2) | 13/9/2011 | NCBI |
| CY160840 | A/Peru/PER070/2010(H3N2) | 17/11/2010 | NCBI |
| CY160848 | A/Peru/PER071/2010(H3N2) | 16/10/2010 | NCBI |
| CY160856 | A/Peru/PER072/2012(H3N2) | 27/2/2012 | NCBI |

| | | | |
|---|---|---|---|
| CY160864 | A/Peru/PER073/2010(H3N2) | 17/11/2010 | NCBI |
| CY160872 | A/Peru/PER074/2010(H3N2) | 27/10/2010 | NCBI |
| CY160880 | A/Peru/PER075/2010(H3N2) | 21/10/2010 | NCBI |
| CY160888 | A/Peru/PER076/2010(H3N2) | 18/10/2010 | NCBI |
| CY160896 | A/Peru/PER077/2010(H3N2) | 16/11/2010 | NCBI |
| CY160904 | A/Peru/PER078/2012(H3N2) | 4/4/2012 | NCBI |
| CY160912 | A/Peru/PER079/2010(H3N2) | 27/10/2010 | NCBI |
| CY160920 | A/Peru/PER080/2010(H3N2) | 22/10/2010 | NCBI |
| CY160928 | A/Peru/PER081/2012(H3N2) | 3/2/2012 | NCBI |
| CY160936 | A/Peru/PER082/2010(H3N2) | 30/10/2010 | NCBI |
| CY160944 | A/Peru/PER083/2010(H3N2) | 16/9/2010 | NCBI |
| CY160952 | A/Peru/PER084/2010(H3N2) | 23/10/2010 | NCBI |
| CY160960 | A/Peru/PER085/2011(H3N2) | 26/10/2011 | NCBI |
| CY160968 | A/Peru/PER086/2011(H3N2) | 12/8/2011 | NCBI |
| CY160976 | A/Peru/PER087/2011(H3N2) | 13/9/2011 | NCBI |
| CY160984 | A/Peru/PER088/2010(H3N2) | 13/10/2010 | NCBI |
| CY160992 | A/Peru/PER089/2010(H3N2) | 26/10/2010 | NCBI |
| CY161000 | A/Peru/PER090/2010(H3N2) | 14/10/2010 | NCBI |
| CY161008 | A/Peru/PER092/2011(H3N2) | 23/11/2011 | NCBI |
| CY161016 | A/Peru/PER093/2010(H3N2) | 16/11/2010 | NCBI |
| CY161024 | A/Peru/PER094/2010(H3N2) | 15/10/2010 | NCBI |
| CY161032 | A/Peru/PER095/2010(H3N2) | 23/11/2010 | NCBI |
| CY161040 | A/Peru/PER096/2012(H3N2) | 30/4/2012 | NCBI |
| CY161048 | A/Peru/PER097/2011(H3N2) | 13/9/2011 | NCBI |
| CY161056 | A/Peru/PER098/2011(H3N2) | 15/9/2011 | NCBI |
| CY161064 | A/Peru/PER099/2012(H3N2) | 30/1/2012 | NCBI |
| CY161072 | A/Peru/PER100/2012(H3N2) | 11/5/2012 | NCBI |
| CY161080 | A/Peru/PER101/2010(H3N2) | 16/10/2010 | NCBI |
| CY161088 | A/Peru/PER102/2012(H3N2) | 9/3/2012 | NCBI |
| CY161096 | A/Peru/PER103/2010(H3N2) | 19/11/2010 | NCBI |
| CY161104 | A/Peru/PER104/2010(H3N2) | 4/10/2010 | NCBI |
| CY161112 | A/Peru/PER105/2012(H3N2) | 4/4/2012 | NCBI |
| CY161120 | A/Peru/PER106/2011(H3N2) | 6/9/2011 | NCBI |
| CY161128 | A/Peru/PER107/2010(H3N2) | 21/9/2010 | NCBI |
| CY161136 | A/Peru/PER108/2012(H3N2) | 16/4/2012 | NCBI |
| CY161144 | A/Peru/PER109/2010(H3N2) | 10/11/2010 | NCBI |
| CY161152 | A/Peru/PER110/2010(H3N2) | 14/12/2010 | NCBI |

| | | | |
|---|---|---|---|
| CY161160 | A/Peru/PER111/2012(H3N2) | 1/6/2012 | NCBI |
| CY161168 | A/Peru/PER112/2010(H3N2) | 8/11/2010 | NCBI |
| CY161176 | A/Peru/PER113/2012(H3N2) | 28/3/2012 | NCBI |
| CY161184 | A/Peru/PER114/2010(H3N2) | 16/11/2010 | NCBI |
| CY161192 | A/Peru/PER115/2011(H3N2) | 28/9/2011 | NCBI |
| CY161200 | A/Peru/PER116/2010(H3N2) | 22/10/2010 | NCBI |
| CY161208 | A/Peru/PER117/2012(H3N2) | 1/1/2012 | NCBI |
| CY161216 | A/Peru/PER118/2011(H3N2) | 24/10/2011 | NCBI |
| CY161224 | A/Peru/PER119/2010(H3N2) | 12/11/2010 | NCBI |
| CY161232 | A/Peru/PER120/2010(H3N2) | 29/11/2010 | NCBI |
| CY161240 | A/Peru/PER121/2011(H3N2) | 21/10/2011 | NCBI |
| CY161248 | A/Peru/PER122/2011(H3N2) | 17/8/2011 | NCBI |
| CY161256 | A/Peru/PER123/2010(H3N2) | 17/11/2010 | NCBI |
| CY161264 | A/Peru/PER124/2012(H3N2) | 4/6/2012 | NCBI |
| CY161272 | A/Peru/PER125/2010(H3N2) | 26/10/2010 | NCBI |
| CY161280 | A/Peru/PER127/2010(H3N2) | 8/11/2010 | NCBI |
| CY161288 | A/Peru/PER128/2010(H3N2) | 22/10/2010 | NCBI |
| CY161296 | A/Peru/PER129/2010(H3N2) | 28/10/2010 | NCBI |
| CY161304 | A/Peru/PER130/2011(H3N2) | 9/9/2011 | NCBI |
| CY161312 | A/Peru/PER131/2010(H3N2) | 10/11/2010 | NCBI |
| CY161320 | A/Peru/PER132/2011(H3N2) | 7/9/2011 | NCBI |
| CY161328 | A/Peru/PER133/2010(H3N2) | 4/10/2010 | NCBI |
| CY161336 | A/Peru/PER134/2011(H3N2) | 16/11/2011 | NCBI |
| CY161344 | A/Peru/PER135/2010(H3N2) | 22/10/2010 | NCBI |
| CY161352 | A/Peru/PER136/2011(H3N2) | 31/10/2011 | NCBI |
| CY161360 | A/Peru/PER137/2011(H3N2) | 2/8/2011 | NCBI |
| CY161368 | A/Peru/PER138/2010(H3N2) | 10/11/2010 | NCBI |
| CY161376 | A/Peru/PER139/2010(H3N2) | 11/10/2010 | NCBI |
| CY161384 | A/Peru/PER140/2010(H3N2) | 13/11/2010 | NCBI |
| CY161392 | A/Peru/PER141/2010(H3N2) | 11/11/2010 | NCBI |
| CY161400 | A/Peru/PER142/2011(H3N2) | 7/10/2011 | NCBI |
| CY161408 | A/Peru/PER143/2010(H3N2) | 21/10/2010 | NCBI |
| CY161416 | A/Peru/PER144/2012(H3N2) | 1/2/2012 | NCBI |
| CY161424 | A/Peru/PER145/2010(H3N2) | 18/10/2010 | NCBI |
| CY161432 | A/Peru/PER146/2010(H3N2) | 17/11/2010 | NCBI |
| CY161440 | A/Peru/PER147/2010(H3N2) | 8/12/2010 | NCBI |
| CY161448 | A/Peru/PER148/2010(H3N2) | 27/10/2010 | NCBI |

| | | | |
|---|---|---|---|
| CY161456 | A/Peru/PER149/2011(H3N2) | 7/9/2011 | NCBI |
| CY161464 | A/Peru/PER150/2010(H3N2) | 3/12/2010 | NCBI |
| CY161472 | A/Peru/PER151/2011(H3N2) | 20/1/2011 | NCBI |
| CY161480 | A/Peru/PER152/2011(H3N2) | 13/9/2011 | NCBI |
| CY161488 | A/Peru/PER153/2012(H3N2) | 11/4/2012 | NCBI |
| CY161496 | A/Peru/PER154/2010(H3N2) | 18/12/2010 | NCBI |
| CY161504 | A/Peru/PER155/2011(H3N2) | 19/10/2011 | NCBI |
| CY161512 | A/Peru/PER156/2012(H3N2) | 6/6/2012 | NCBI |
| CY161520 | A/Peru/PER157/2010(H3N2) | 13/11/2010 | NCBI |
| CY161528 | A/Peru/PER158/2011(H3N2) | 6/9/2011 | NCBI |
| CY161536 | A/Peru/PER159/2011(H3N2) | 20/9/2011 | NCBI |
| CY161544 | A/Peru/PER160/2010(H3N2) | 25/10/2010 | NCBI |
| CY161552 | A/Peru/PER161/2011(H3N2) | 6/9/2011 | NCBI |
| CY161560 | A/Peru/PER162/2010(H3N2) | 30/9/2010 | NCBI |
| CY161568 | A/Peru/PER163/2011(H3N2) | 8/9/2011 | NCBI |
| CY161576 | A/Peru/PER164/2010(H3N2) | 12/11/2010 | NCBI |
| CY161584 | A/Peru/PER165/2010(H3N2) | 9/11/2010 | NCBI |
| CY161592 | A/Peru/PER166/2011(H3N2) | 26/8/2011 | NCBI |
| CY161600 | A/Peru/PER167/2012(H3N2) | 4/5/2012 | NCBI |
| CY161608 | A/Peru/PER168/2010(H3N2) | 3/12/2010 | NCBI |
| CY161616 | A/Peru/PER169/2010(H3N2) | 11/10/2010 | NCBI |
| CY161624 | A/Peru/PER170/2010(H3N2) | 4/11/2010 | NCBI |
| CY161632 | A/Peru/PER171/2010(H3N2) | 19/10/2010 | NCBI |
| CY161640 | A/Peru/PER172/2010(H3N2) | 3/11/2010 | NCBI |
| CY161648 | A/Peru/PER173/2010(H3N2) | 15/12/2010 | NCBI |
| CY161656 | A/Peru/PER174/2010(H3N2) | 10/11/2010 | NCBI |
| CY161664 | A/Peru/PER175/2012(H3N2) | 18/5/2012 | NCBI |
| CY161672 | A/Peru/PER176/2011(H3N2) | 6/9/2011 | NCBI |
| CY161680 | A/Peru/PER177/2010(H3N2) | 3/12/2010 | NCBI |
| CY161688 | A/Peru/PER178/2010(H3N2) | 27/11/2010 | NCBI |
| CY161696 | A/Peru/PER179/2011(H3N2) | 3/9/2011 | NCBI |
| CY161704 | A/Peru/PER180/2010(H3N2) | 4/11/2010 | NCBI |
| CY161712 | A/Peru/PER181/2010(H3N2) | 27/10/2010 | NCBI |
| CY161720 | A/Peru/PER182/2010(H3N2) | 29/9/2010 | NCBI |
| CY161728 | A/Peru/PER183/2010(H3N2) | 11/11/2010 | NCBI |
| CY161736 | A/Peru/PER184/2010(H3N2) | 14/10/2010 | NCBI |
| CY161744 | A/Peru/PER185/2011(H3N2) | 5/10/2011 | NCBI |

| | | | |
|---|---|---|---|
| CY161752 | A/Peru/PER186/2012(H3N2) | 31/5/2012 | NCBI |
| CY161760 | A/Peru/PER187/2011(H3N2) | 7/11/2011 | NCBI |
| CY161768 | A/Peru/PER188/2010(H3N2) | 13/10/2010 | NCBI |
| CY161776 | A/Peru/PER189/2011(H3N2) | 25/1/2011 | NCBI |
| CY161784 | A/Peru/PER190/2010(H3N2) | 7/12/2010 | NCBI |
| CY161792 | A/Peru/PER191/2010(H3N2) | 15/11/2010 | NCBI |
| CY161800 | A/Peru/PER192/2010(H3N2) | 23/10/2010 | NCBI |
| CY161808 | A/Peru/PER193/2010(H3N2) | 27/11/2010 | NCBI |
| CY161816 | A/Peru/PER194/2011(H3N2) | 1/9/2011 | NCBI |
| CY161824 | A/Peru/PER195/2010(H3N2) | 10/9/2010 | NCBI |
| CY161832 | A/Peru/PER196/2010(H3N2) | 18/10/2010 | NCBI |
| CY161840 | A/Peru/PER197/2011(H3N2) | 14/11/2011 | NCBI |
| CY161848 | A/Peru/PER198/2010(H3N2) | 29/11/2010 | NCBI |
| CY161856 | A/Peru/PER199/2011(H3N2) | 29/8/2011 | NCBI |
| CY161864 | A/Peru/PER200/2012(H3N2) | 4/6/2012 | NCBI |
| CY161872 | A/Peru/PER201/2010(H3N2) | 15/11/2010 | NCBI |
| CY161880 | A/Peru/PER202/2010(H3N2) | 8/11/2010 | NCBI |
| CY161888 | A/Peru/PER203/2011(H3N2) | 14/9/2011 | NCBI |
| CY161896 | A/Peru/PER204/2010(H3N2) | 20/12/2010 | NCBI |
| CY161904 | A/Peru/PER205/2011(H3N2) | 12/9/2011 | NCBI |
| CY161912 | A/Peru/PER206/2011(H3N2) | 7/9/2011 | NCBI |
| CY161920 | A/Peru/PER207/2010(H3N2) | 27/10/2010 | NCBI |
| CY161928 | A/Peru/PER208/2010(H3N2) | 27/10/2010 | NCBI |
| CY161936 | A/Peru/PER209/2012(H3N2) | 27/2/2012 | NCBI |
| CY161944 | A/Peru/PER210/2010(H3N2) | 13/9/2010 | NCBI |
| CY161952 | A/Peru/PER211/2010(H3N2) | 22/10/2010 | NCBI |
| CY161960 | A/Peru/PER212/2010(H3N2) | 2/11/2010 | NCBI |
| CY161968 | A/Peru/PER213/2010(H3N2) | 18/10/2010 | NCBI |
| CY161976 | A/Peru/PER214/2010(H3N2) | 4/11/2010 | NCBI |
| CY161984 | A/Peru/PER215/2010(H3N2) | 20/12/2010 | NCBI |
| CY161992 | A/Peru/PER216/2012(H3N2) | 1/6/2012 | NCBI |
| CY162000 | A/Peru/PER217/2012(H3N2) | 8/2/2012 | NCBI |
| CY162008 | A/Peru/PER218/2010(H3N2) | 15/10/2010 | NCBI |
| CY162016 | A/Peru/PER219/2012(H3N2) | 7/5/2012 | NCBI |
| CY162024 | A/Peru/PER220/2010(H3N2) | 2/11/2010 | NCBI |
| CY162032 | A/Peru/PER221/2012(H3N2) | 2/5/2012 | NCBI |
| CY162040 | A/Peru/PER222/2012(H3N2) | 9/4/2012 | NCBI |

| | | | |
|---|---|---|---|
| CY162048 | A/Peru/PER223/2010(H3N2) | 9/12/2010 | NCBI |
| CY162056 | A/Peru/PER224/2010(H3N2) | 2/11/2010 | NCBI |
| CY162064 | A/Peru/PER225/2010(H3N2) | 27/10/2010 | NCBI |
| CY162072 | A/Peru/PER226/2011(H3N2) | 7/10/2011 | NCBI |
| CY162080 | A/Peru/PER227/2011(H3N2) | 31/8/2011 | NCBI |
| CY162088 | A/Peru/PER228/2010(H3N2) | 2/11/2010 | NCBI |
| CY162096 | A/Peru/PER229/2010(H3N2) | 6/11/2010 | NCBI |
| CY162104 | A/Peru/PER230/2010(H3N2) | 26/10/2010 | NCBI |
| CY162112 | A/Peru/PER231/2011(H3N2) | 12/9/2011 | NCBI |
| CY162120 | A/Peru/PER232/2012(H3N2) | 18/5/2012 | NCBI |
| CY162128 | A/Peru/PER233/2010(H3N2) | 13/10/2010 | NCBI |
| CY162136 | A/Peru/PER234/2010(H3N2) | 3/12/2010 | NCBI |
| CY162144 | A/Peru/PER235/2011(H3N2) | 27/5/2011 | NCBI |
| CY162152 | A/Peru/PER236/2012(H3N2) | 17/3/2012 | NCBI |
| CY162160 | A/Peru/PER237/2010(H3N2) | 3/12/2010 | NCBI |
| CY162168 | A/Peru/PER238/2011(H3N2) | 22/8/2011 | NCBI |
| CY162184 | A/Peru/PER241/2010(H3N2) | 6/11/2010 | NCBI |
| CY162192 | A/Peru/PER242/2010(H3N2) | 2/11/2010 | NCBI |
| CY162200 | A/Peru/PER243/2011(H3N2) | 31/8/2011 | NCBI |
| CY162208 | A/Peru/PER244/2011(H3N2) | 26/8/2011 | NCBI |
| CY162216 | A/Peru/PER245/2010(H3N2) | 15/10/2010 | NCBI |
| CY162224 | A/Peru/PER246/2010(H3N2) | 26/11/2010 | NCBI |
| CY162232 | A/Peru/PER247/2011(H3N2) | 26/8/2011 | NCBI |
| CY162240 | A/Peru/PER248/2010(H3N2) | 20/10/2010 | NCBI |
| CY162248 | A/Peru/PER250/2010(H3N2) | 4/11/2010 | NCBI |
| CY162256 | A/Peru/PER251/2010(H3N2) | 12/10/2010 | NCBI |
| CY162264 | A/Peru/PER252/2010(H3N2) | 21/10/2010 | NCBI |
| CY162272 | A/Peru/PER253/2010(H3N2) | 16/11/2010 | NCBI |
| CY162280 | A/Peru/PER254/2010(H3N2) | 8/11/2010 | NCBI |
| CY162288 | A/Peru/PER255/2011(H3N2) | 6/9/2011 | NCBI |
| CY162296 | A/Peru/PER256/2010(H3N2) | 22/11/2010 | NCBI |
| CY162304 | A/Peru/PER257/2010(H3N2) | 20/10/2010 | NCBI |
| CY162312 | A/Peru/PER258/2010(H3N2) | 6/11/2010 | NCBI |
| CY162320 | A/Peru/PER259/2010(H3N2) | 15/10/2010 | NCBI |
| CY162328 | A/Peru/PER260/2012(H3N2) | 4/6/2012 | NCBI |
| CY162336 | A/Peru/PER262/2012(H3N2) | 19/4/2012 | NCBI |
| CY162344 | A/Peru/PER263/2010(H3N2) | 9/11/2010 | NCBI |

| | | | |
|---|---|---|---|
| CY162352 | A/Peru/PER264/2011(H3N2) | 19/9/2011 | NCBI |
| CY162360 | A/Peru/PER265/2011(H3N2) | 14/9/2011 | NCBI |
| CY162368 | A/Peru/PER266/2010(H3N2) | 28/10/2010 | NCBI |
| CY162376 | A/Peru/PER267/2010(H3N2) | 8/11/2010 | NCBI |
| CY162384 | A/Peru/PER268/2011(H3N2) | 12/9/2011 | NCBI |
| CY162392 | A/Peru/PER269/2011(H3N2) | 15/9/2011 | NCBI |
| CY162400 | A/Peru/PER270/2010(H3N2) | 4/11/2010 | NCBI |
| CY162408 | A/Peru/PER271/2011(H3N2) | 13/9/2011 | NCBI |
| CY162416 | A/Peru/PER272/2010(H3N2) | 16/10/2010 | NCBI |
| CY162424 | A/Peru/PER273/2011(H3N2) | 19/8/2011 | NCBI |
| CY162432 | A/Peru/PER274/2010(H3N2) | 18/10/2010 | NCBI |
| CY162440 | A/Peru/PER275/2011(H3N2) | 21/9/2011 | NCBI |
| CY162448 | A/Peru/PER276/2010(H3N2) | 4/11/2010 | NCBI |
| CY162456 | A/Peru/PER277/2010(H3N2) | 14/10/2010 | NCBI |
| CY162464 | A/Peru/PER278/2010(H3N2) | 4/11/2010 | NCBI |
| CY162472 | A/Peru/PER279/2010(H3N2) | 20/9/2010 | NCBI |
| CY162480 | A/Peru/PER280/2012(H3N2) | 18/5/2012 | NCBI |
| CY162488 | A/Peru/PER281/2011(H3N2) | 23/8/2011 | NCBI |
| CY162496 | A/Peru/PER282/2011(H3N2) | 30/9/2011 | NCBI |
| CY162504 | A/Peru/PER283/2011(H3N2) | 7/10/2011 | NCBI |
| CY162512 | A/Peru/PER284/2011(H3N2) | 23/10/2011 | NCBI |
| CY162520 | A/Peru/PER285/2010(H3N2) | 30/11/2010 | NCBI |
| CY162528 | A/Peru/PER286/2010(H3N2) | 28/10/2010 | NCBI |
| CY162536 | A/Peru/PER287/2011(H3N2) | 9/9/2011 | NCBI |
| CY162544 | A/Peru/PER288/2010(H3N2) | 27/10/2010 | NCBI |
| CY162552 | A/Peru/PER289/2011(H3N2) | 2/9/2011 | NCBI |
| CY162560 | A/Peru/PER290/2011(H3N2) | 29/8/2011 | NCBI |
| CY162568 | A/Peru/PER291/2010(H3N2) | 22/10/2010 | NCBI |
| CY162576 | A/Peru/PER292/2010(H3N2) | 20/11/2010 | NCBI |
| CY162584 | A/Peru/PER294/2010(H3N2) | 20/10/2010 | NCBI |
| CY162592 | A/Peru/PER295/2010(H3N2) | 20/12/2010 | NCBI |
| CY162600 | A/Peru/PER297/2010(H3N2) | 26/10/2010 | NCBI |
| CY162608 | A/Peru/PER298/2011(H3N2) | 7/9/2011 | NCBI |
| CY162616 | A/Peru/PER299/2010(H3N2) | 20/11/2010 | NCBI |
| CY162624 | A/Peru/PER300/2010(H3N2) | 7/10/2010 | NCBI |
| CY162632 | A/Peru/PER301/2012(H3N2) | 2/4/2012 | NCBI |
| CY162640 | A/Peru/PER302/2011(H3N2) | 30/5/2011 | NCBI |

| | | | |
|---|---|---|---|
| CY162648 | A/Peru/PER303/2010(H3N2) | 30/9/2010 | NCBI |
| CY162656 | A/Peru/PER304/2010(H3N2) | 29/11/2010 | NCBI |
| CY162664 | A/Peru/PER305/2012(H3N2) | 7/5/2012 | NCBI |
| CY162672 | A/Peru/PER306/2010(H3N2) | 5/11/2010 | NCBI |
| CY162680 | A/Peru/PER307/2011(H3N2) | 17/10/2011 | NCBI |
| CY162688 | A/Peru/PER308/2010(H3N2) | 17/9/2010 | NCBI |
| CY162696 | A/Peru/PER309/2012(H3N2) | 9/5/2012 | NCBI |
| CY162704 | A/Peru/PER310/2010(H3N2) | 3/12/2010 | NCBI |
| CY162712 | A/Peru/PER311/2010(H3N2) | 17/11/2010 | NCBI |
| CY162720 | A/Peru/PER312/2010(H3N2) | 22/10/2010 | NCBI |
| CY162728 | A/Peru/PER313/2012(H3N2) | 1/1/2012 | NCBI |
| CY162736 | A/Peru/PER314/2012(H3N2) | 18/4/2012 | NCBI |
| CY162744 | A/Peru/PER315/2010(H3N2) | 21/10/2010 | NCBI |
| CY162752 | A/Peru/PER316/2012(H3N2) | 10/4/2012 | NCBI |
| CY162760 | A/Peru/PER317/2010(H3N2) | 25/10/2010 | NCBI |
| CY162768 | A/Peru/PER318/2010(H3N2) | 18/10/2010 | NCBI |
| CY162776 | A/Peru/PER319/2010(H3N2) | 27/10/2010 | NCBI |
| CY162784 | A/Peru/PER320/2010(H3N2) | 9/11/2010 | NCBI |
| CY162792 | A/Peru/PER321/2010(H3N2) | 30/9/2010 | NCBI |
| CY162800 | A/Peru/PER322/2011(H3N2) | 10/9/2011 | NCBI |
| CY162808 | A/Peru/PER323/2010(H3N2) | 19/10/2010 | NCBI |
| CY162816 | A/Peru/PER324/2011(H3N2) | 20/10/2011 | NCBI |
| CY162824 | A/Peru/PER325/2010(H3N2) | 2/11/2010 | NCBI |
| CY162832 | A/Peru/PER326/2012(H3N2) | 17/4/2012 | NCBI |
| CY162840 | A/Peru/PER327/2010(H3N2) | 23/9/2010 | NCBI |
| CY162848 | A/Peru/PER328/2011(H3N2) | 30/6/2011 | NCBI |
| CY162856 | A/Peru/PER329/2010(H3N2) | 19/10/2010 | NCBI |
| CY162864 | A/Peru/PER330/2010(H3N2) | 4/11/2010 | NCBI |
| CY162872 | A/Peru/PER331/2012(H3N2) | 21/5/2012 | NCBI |
| CY162880 | A/Peru/PER332/2010(H3N2) | 8/11/2010 | NCBI |
| CY162888 | A/Peru/PER333/2011(H3N2) | 30/9/2011 | NCBI |
| CY162896 | A/Peru/PER334/2011(H3N2) | 26/10/2011 | NCBI |
| CY162904 | A/Peru/PER335/2010(H3N2) | 11/11/2010 | NCBI |
| CY162912 | A/Peru/PER336/2010(H3N2) | 29/11/2010 | NCBI |
| CY162920 | A/Peru/PER337/2011(H3N2) | 20/8/2011 | NCBI |
| CY162928 | A/Peru/PER338/2010(H3N2) | 28/10/2010 | NCBI |
| CY162936 | A/Peru/PER339/2010(H3N2) | 2/11/2010 | NCBI |

| | | | |
|---|---|---|---|
| CY162944 | A/Peru/PER340/2010(H3N2) | 3/11/2010 | NCBI |
| CY162952 | A/Peru/PER341/2011(H3N2) | 25/10/2011 | NCBI |
| CY162960 | A/Peru/PER342/2010(H3N2) | 20/10/2010 | NCBI |
| CY162968 | A/Peru/PER343/2011(H3N2) | 3/11/2011 | NCBI |
| CY162976 | A/Peru/PER344/2010(H3N2) | 26/10/2010 | NCBI |
| CY162984 | A/Peru/PER345/2011(H3N2) | 22/9/2011 | NCBI |
| CY162992 | A/Peru/PER346/2011(H3N2) | 19/11/2011 | NCBI |
| CY163000 | A/Peru/PER347/2010(H3N2) | 13/10/2010 | NCBI |
| CY163008 | A/Peru/PER349/2010(H3N2) | 27/10/2010 | NCBI |
| CY163016 | A/Peru/PER350/2010(H3N2) | 8/11/2010 | NCBI |
| CY163024 | A/Peru/PER351/2010(H3N2) | 22/10/2010 | NCBI |
| CY163032 | A/Peru/PER352/2011(H3N2) | 20/9/2011 | NCBI |
| CY163040 | A/Peru/PER353/2012(H3N2) | 30/4/2012 | NCBI |
| CY163048 | A/Peru/PER354/2012(H3N2) | 21/5/2012 | NCBI |
| CY163056 | A/Peru/PER355/2011(H3N2) | 6/9/2011 | NCBI |
| CY163064 | A/Peru/PER356/2012(H3N2) | 4/2/2012 | NCBI |
| CY163072 | A/Peru/PER357/2010(H3N2) | 26/10/2010 | NCBI |
| CY163080 | A/Peru/PER358/2012(H3N2) | 8/2/2012 | NCBI |
| CY163088 | A/Peru/PER359/2012(H3N2) | 28/3/2012 | NCBI |
| CY163096 | A/Peru/PER360/2010(H3N2) | 21/10/2010 | NCBI |
| CY163104 | A/Peru/PER361/2010(H3N2) | 20/11/2010 | NCBI |
| CY163112 | A/Peru/PER362/2011(H3N2) | 23/2/2011 | NCBI |
| CY163120 | A/Peru/PER363/2012(H3N2) | 7/5/2012 | NCBI |
| CY163128 | A/Peru/PER364/2011(H3N2) | 22/9/2011 | NCBI |
| CY163136 | A/Peru/PER365/2010(H3N2) | 6/11/2010 | NCBI |
| CY163144 | A/Peru/PER367/2010(H3N2) | 16/11/2010 | NCBI |
| CY163152 | A/Peru/PER369/2010(H3N2) | 20/11/2010 | NCBI |
| CY163160 | A/Peru/PER370/2010(H3N2) | 12/11/2010 | NCBI |
| CY163168 | A/Peru/PER371/2012(H3N2) | 5/7/2012 | NCBI |
| CY163176 | A/Peru/PER372/2010(H3N2) | 14/10/2010 | NCBI |
| CY163184 | A/Peru/PER373/2010(H3N2) | 15/11/2010 | NCBI |
| CY163192 | A/Peru/PER374/2011(H3N2) | 13/1/2011 | NCBI |
| CY163200 | A/Peru/PER375/2010(H3N2) | 23/11/2010 | NCBI |
| CY163208 | A/Peru/PER376/2010(H3N2) | 15/11/2010 | NCBI |
| CY163216 | A/Peru/PER377/2010(H3N2) | 28/10/2010 | NCBI |
| CY163224 | A/Peru/PER378/2011(H3N2) | 16/9/2011 | NCBI |
| CY163232 | A/Peru/PER379/2010(H3N2) | 27/9/2010 | NCBI |

| | | | |
|---|---|---|---|
| CY163240 | A/Peru/PER380/2010(H3N2) | 13/10/2010 | NCBI |
| CY163248 | A/Peru/PER381/2010(H3N2) | 21/10/2010 | NCBI |
| CY163256 | A/Peru/PER382/2010(H3N2) | 26/10/2010 | NCBI |
| CY163264 | A/Peru/PER383/2010(H3N2) | 6/12/2010 | NCBI |
| CY163272 | A/Peru/PER384/2012(H3N2) | 9/4/2012 | NCBI |
| CY163280 | A/Peru/PER385/2011(H3N2) | 14/9/2011 | NCBI |
| CY163288 | A/Peru/PER386/2010(H3N2) | 28/10/2010 | NCBI |
| CY163296 | A/Peru/PER387/2010(H3N2) | 5/11/2010 | NCBI |
| CY163304 | A/Peru/PER388/2011(H3N2) | 14/9/2011 | NCBI |
| CY163312 | A/Peru/PER389/2012(H3N2) | 2/4/2012 | NCBI |
| CY163320 | A/Peru/PER390/2010(H3N2) | 11/10/2010 | NCBI |
| CY163328 | A/Peru/PER391/2010(H3N2) | 17/11/2010 | NCBI |
| CY163336 | A/Peru/PER392/2010(H3N2) | 29/10/2010 | NCBI |
| CY163344 | A/Peru/PER393/2010(H3N2) | 28/10/2010 | NCBI |
| CY163352 | A/Peru/PER394/2011(H3N2) | 7/9/2011 | NCBI |
| CY163360 | A/Peru/PER395/2011(H3N2) | 24/10/2011 | NCBI |
| CY163368 | A/Peru/PER396/2011(H3N2) | 7/9/2011 | NCBI |
| CY163376 | A/Peru/PER397/2011(H3N2) | 10/9/2011 | NCBI |
| CY163384 | A/Peru/PER398/2010(H3N2) | 11/10/2010 | NCBI |
| CY163392 | A/Peru/PER399/2012(H3N2) | 3/4/2012 | NCBI |
| CY163400 | A/Peru/PER400/2012(H3N2) | 7/3/2012 | NCBI |
| KF034879 | A/Suzhou/1275/2013(H3N2) | 4/4/2013 | NCBI |
| KJ473722 | A/Guangdong/ST798/2008(H3N2) | 2008 | NCBI |
| CY167484 | A/Tennessee/F2011/2011(H3N2) | 7/1/2011 | NCBI |
| CY167500 | A/Tennessee/F2018/2011(H3N2) | 21/1/2011 | NCBI |
| CY167508 | A/Tennessee/F2018A/2011(H3N2) | 24/1/2011 | NCBI |
| CY167516 | A/Tennessee/F2019/2011(H3N2) | 21/1/2011 | NCBI |
| CY167524 | A/Tennessee/F2019A/2011(H3N2) | 24/1/2011 | NCBI |
| CY167532 | A/Tennessee/F2021/2011(H3N2) | 28/1/2011 | NCBI |
| CY167540 | A/Tennessee/F2021B/2011(H3N2) | 4/2/2011 | NCBI |
| CY167564 | A/Tennessee/F2031A/2011(H3N2) | 7/2/2011 | NCBI |
| CY167572 | A/Tennessee/F2034/2011(H3N2) | 7/2/2011 | NCBI |
| CY167596 | A/Tennessee/F2048/2011(H3N2) | 16/2/2011 | NCBI |
| CY167604 | A/Tennessee/F2048A/2011(H3N2) | 18/2/2011 | NCBI |
| CY167612 | A/Tennessee/F2053/2011(H3N2) | 22/2/2011 | NCBI |
| CY167620 | A/Tennessee/F2054A/2011(H3N2) | 25/2/2011 | NCBI |
| CY167628 | A/Tennessee/F2057c54A/2011(H3N2) | 7/3/2011 | NCBI |

| | | | |
|---|---|---|---|
| CY167644 | A/Tennessee/F2065/2011(H3N2) | 2/3/2011 | NCBI |
| CY167652 | A/Tennessee/F2065B/2011(H3N2) | 7/3/2011 | NCBI |
| CY167668 | A/Tennessee/F2078/2011(H3N2) | 15/3/2011 | NCBI |
| CY167676 | A/Tennessee/F2078A/2011(H3N2) | 18/3/2011 | NCBI |
| CY167684 | A/Tennessee/F2079/2011(H3N2) | 15/3/2011 | NCBI |
| CY167692 | A/Tennessee/F2079A/2011(H3N2) | 18/3/2011 | NCBI |
| CY167700 | A/Tennessee/F2079B/2011(H3N2) | 22/3/2011 | NCBI |
| CY167708 | A/Tennessee/F2080c7879/2011(H3N2) | 15/3/2011 | NCBI |
| CY167732 | A/Tennessee/F2084c83/2011(H3N2) | 16/3/2011 | NCBI |
| CY167740 | A/Tennessee/F2084c83A/2011(H3N2) | 23/3/2011 | NCBI |
| CY167764 | A/Tennessee/F2090B/2011(H3N2) | 7/4/2011 | NCBI |
| CY172183 | A/NewYork/1000/2005(H3N2) | 30/8/2005 | NCBI |
| CY172191 | A/NewYork/1001/2005(H3N2) | 10/11/2005 | NCBI |
| CY172199 | A/NewYork/1002/2005(H3N2) | 27/12/2005 | NCBI |
| CY172207 | A/NewYork/1005/2006(H3N2) | 4/1/2006 | NCBI |
| CY172215 | A/NewYork/1006/2006(H3N2) | 8/1/2006 | NCBI |
| CY172223 | A/NewYork/1008/2006(H3N2) | 11/1/2006 | NCBI |
| CY172231 | A/NewYork/1009/2006(H3N2) | 12/1/2006 | NCBI |
| CY172239 | A/NewYork/1010/2006(H3N2) | 13/1/2006 | NCBI |
| CY172247 | A/NewYork/1011/2006(H3N2) | 24/1/2006 | NCBI |
| CY172255 | A/NewYork/1012/2006(H3N2) | 24/1/2006 | NCBI |
| CY172263 | A/NewYork/1013/2006(H3N2) | 6/2/2006 | NCBI |
| CY172271 | A/NewYork/1014/2006(H3N2) | 7/2/2006 | NCBI |
| CY172279 | A/NewYork/1015/2006(H3N2) | 7/2/2006 | NCBI |
| CY172287 | A/NewYork/1016/2006(H3N2) | 14/2/2006 | NCBI |
| CY172295 | A/NewYork/1017/2006(H3N2) | 14/2/2006 | NCBI |
| CY172303 | A/NewYork/1018/2006(H3N2) | 15/2/2006 | NCBI |
| CY172311 | A/NewYork/1019/2006(H3N2) | 20/4/2006 | NCBI |
| CY172319 | A/NewYork/1020/2006(H3N2) | 15/2/2006 | NCBI |
| CY172327 | A/NewYork/1021/2006(H3N2) | 15/2/2006 | NCBI |
| CY172335 | A/NewYork/1022/2006(H3N2) | 16/2/2006 | NCBI |
| CY172343 | A/NewYork/1023/2006(H3N2) | 16/2/2006 | NCBI |
| CY172351 | A/NewYork/1024/2006(H3N2) | 17/2/2006 | NCBI |
| CY172359 | A/NewYork/1025/2006(H3N2) | 17/2/2006 | NCBI |
| CY172367 | A/NewYork/1026/2006(H3N2) | 17/2/2006 | NCBI |
| CY172375 | A/NewYork/1027/2006(H3N2) | 17/2/2006 | NCBI |
| CY172383 | A/NewYork/1028/2006(H3N2) | 20/2/2006 | NCBI |

| | | | |
|---|---|---|---|
| CY172391 | A/NewYork/1029/2006(H3N2) | 24/2/2006 | NCBI |
| CY172399 | A/NewYork/1030/2006(H3N2) | 27/2/2006 | NCBI |
| CY172407 | A/NewYork/1031/2006(H3N2) | 28/2/2006 | NCBI |
| CY172415 | A/NewYork/1032/2006(H3N2) | 6/3/2006 | NCBI |
| CY172423 | A/NewYork/1033/2006(H3N2) | 7/3/2006 | NCBI |
| CY172431 | A/NewYork/1034/2006(H3N2) | 7/3/2006 | NCBI |
| CY172439 | A/NewYork/1035/2006(H3N2) | 7/3/2006 | NCBI |
| CY172447 | A/NewYork/1036/2006(H3N2) | 8/3/2006 | NCBI |
| CY172455 | A/NewYork/1037/2006(H3N2) | 9/3/2006 | NCBI |
| CY172463 | A/NewYork/1039/2006(H3N2) | 12/3/2006 | NCBI |
| CY172471 | A/NewYork/1040/2006(H3N2) | 12/3/2006 | NCBI |
| CY172479 | A/NewYork/1041/2006(H3N2) | 13/3/2006 | NCBI |
| CY172487 | A/NewYork/1042/2006(H3N2) | 13/3/2006 | NCBI |
| CY172495 | A/NewYork/1043/2006(H3N2) | 14/3/2006 | NCBI |
| CY172503 | A/NewYork/1044/2006(H3N2) | 16/3/2006 | NCBI |
| CY172511 | A/NewYork/1045/2006(H3N2) | 20/3/2006 | NCBI |
| CY172519 | A/NewYork/1046/2006(H3N2) | 22/3/2006 | NCBI |
| CY172527 | A/NewYork/1048/2006(H3N2) | 31/3/2006 | NCBI |
| CY172735 | A/NewYork/1077/2007(H3N2) | 10/2/2007 | NCBI |
| CY172743 | A/NewYork/1078/2007(H3N2) | 12/2/2007 | NCBI |
| CY172751 | A/NewYork/1079/2007(H3N2) | 11/2/2007 | NCBI |
| CY172759 | A/NewYork/1080/2007(H3N2) | 16/2/2007 | NCBI |
| CY172767 | A/NewYork/1082/2007(H3N2) | 28/2/2007 | NCBI |
| CY172775 | A/NewYork/1083/2007(H3N2) | 2/3/2007 | NCBI |
| CY172783 | A/NewYork/1084/2007(H3N2) | 5/3/2007 | NCBI |
| CY172791 | A/NewYork/1085/2007(H3N2) | 7/3/2007 | NCBI |
| CY172799 | A/NewYork/1086/2007(H3N2) | 8/3/2007 | NCBI |
| CY172807 | A/NewYork/1087/2007(H3N2) | 15/3/2007 | NCBI |
| CY172815 | A/NewYork/1088/2007(H3N2) | 16/3/2007 | NCBI |
| CY172823 | A/NewYork/1089/2007(H3N2) | 23/3/2007 | NCBI |
| CY172831 | A/NewYork/1090/2007(H3N2) | 26/3/2007 | NCBI |
| CY172839 | A/NewYork/1093/2007(H3N2) | 11/4/2007 | NCBI |
| CY172847 | A/NewYork/1095/2007(H3N2) | 13/4/2007 | NCBI |
| CY172855 | A/NewYork/1096/2007(H3N2) | 13/4/2007 | NCBI |
| CY172863 | A/NewYork/1097/2007(H3N2) | 15/4/2007 | NCBI |
| CY172871 | A/NewYork/1098/2007(H3N2) | 16/4/2007 | NCBI |
| CY172879 | A/NewYork/1099/2007(H3N2) | 24/4/2007 | NCBI |

| | | | |
|---|---|---|---|
| CY172887 | A/NewYork/1100/2007(H3N2) | 6/6/2007 | NCBI |
| CY172895 | A/NewYork/1102/2007(H3N2) | 6/2/2007 | NCBI |
| CY172903 | A/NewYork/1103/2007(H3N2) | 26/3/2007 | NCBI |
| CY173087 | A/NewYork/1128/2008(H3N2) | 18/1/2008 | NCBI |
| CY173095 | A/NewYork/1129/2008(H3N2) | 24/1/2008 | NCBI |
| CY173103 | A/NewYork/1130/2008(H3N2) | 24/1/2008 | NCBI |
| CY173111 | A/NewYork/1131/2008(H3N2) | 30/1/2008 | NCBI |
| CY173119 | A/NewYork/1133/2008(H3N2) | 29/1/2008 | NCBI |
| CY173127 | A/NewYork/1134/2008(H3N2) | 28/1/2008 | NCBI |
| CY173135 | A/NewYork/1135/2008(H3N2) | 4/2/2008 | NCBI |
| CY173143 | A/NewYork/1136/2008(H3N2) | 5/2/2008 | NCBI |
| CY173151 | A/NewYork/1138/2008(H3N2) | 6/2/2008 | NCBI |
| CY173159 | A/NewYork/1139/2008(H3N2) | 6/2/2008 | NCBI |
| CY173167 | A/NewYork/1140/2008(H3N2) | 10/2/2008 | NCBI |
| CY173175 | A/NewYork/1141/2008(H3N2) | 13/2/2008 | NCBI |
| CY173183 | A/NewYork/1142/2008(H3N2) | 18/2/2008 | NCBI |
| CY173191 | A/NewYork/1143/2008(H3N2) | 18/2/2008 | NCBI |
| CY173199 | A/NewYork/1144/2008(H3N2) | 19/2/2008 | NCBI |
| CY173207 | A/NewYork/1145/2008(H3N2) | 19/2/2008 | NCBI |
| CY173215 | A/NewYork/1146/2008(H3N2) | 19/2/2008 | NCBI |
| CY173223 | A/NewYork/1147/2008(H3N2) | 27/2/2008 | NCBI |
| CY173231 | A/NewYork/1148/2008(H3N2) | 3/3/2008 | NCBI |
| CY173239 | A/NewYork/1150/2008(H3N2) | 3/3/2008 | NCBI |
| CY173247 | A/NewYork/1151/2008(H3N2) | 6/3/2008 | NCBI |
| CY173255 | A/NewYork/1152/2008(H3N2) | 10/3/2008 | NCBI |
| CY173263 | A/NewYork/1153/2008(H3N2) | 12/3/2008 | NCBI |
| CY173271 | A/NewYork/1154/2008(H3N2) | 11/3/2008 | NCBI |
| CY173279 | A/NewYork/1155/2008(H3N2) | 27/3/2008 | NCBI |
| CY173287 | A/NewYork/1156/2008(H3N2) | 28/3/2008 | NCBI |
| CY173295 | A/NewYork/1157/2008(H3N2) | 21/4/2008 | NCBI |
| CY173511 | A/NewYork/1188/2009(H3N2) | 26/1/2009 | NCBI |
| CY173519 | A/NewYork/1189/2009(H3N2) | 16/2/2009 | NCBI |
| CY173527 | A/NewYork/1190/2009(H3N2) | 22/2/2009 | NCBI |
| CY173535 | A/NewYork/1191/2009(H3N2) | 25/2/2009 | NCBI |
| CY173543 | A/NewYork/1192/2009(H3N2) | 1/3/2009 | NCBI |
| CY173551 | A/NewYork/1193/2009(H3N2) | 16/3/2009 | NCBI |
| CY173559 | A/NewYork/1194/2009(H3N2) | 16/3/2009 | NCBI |

| | | | |
|---|---|---|---|
| CY173567 | A/NewYork/1195/2009(H3N2) | 9/2/2009 | NCBI |
| CY173575 | A/NewYork/1196/2009(H3N2) | 19/2/2009 | NCBI |
| CY173583 | A/NewYork/1197/2009(H3N2) | 24/2/2009 | NCBI |
| CY173591 | A/NewYork/1198/2009(H3N2) | 2/3/2009 | NCBI |
| CY173599 | A/NewYork/1003/2005(H3N2) | 20/12/2005 | NCBI |
| CY167887 | A/Boston/YGA_00002/2012(H3N2) | 26/10/2012 | NCBI |
| CY167895 | A/Boston/YGA_00003/2012(H3N2) | 27/11/2012 | NCBI |
| CY167903 | A/Boston/YGA_00004/2012(H3N2) | 5/12/2012 | NCBI |
| CY167911 | A/Boston/YGA_00006/2012(H3N2) | 14/12/2012 | NCBI |
| CY167919 | A/Boston/YGA_00007/2012(H3N2) | 18/12/2012 | NCBI |
| CY167927 | A/Boston/YGA_00008/2012(H3N2) | 19/12/2012 | NCBI |
| CY167935 | A/Boston/YGA_00009/2012(H3N2) | 20/12/2012 | NCBI |
| CY167943 | A/Boston/YGA_00010/2012(H3N2) | 20/12/2012 | NCBI |
| CY167951 | A/Boston/YGA_00011/2012(H3N2) | 27/12/2012 | NCBI |
| CY167959 | A/Boston/YGA_00012/2012(H3N2) | 27/12/2012 | NCBI |
| CY167967 | A/Boston/YGA_00014/2012(H3N2) | 28/12/2012 | NCBI |
| CY167975 | A/Boston/YGA_00015/2012(H3N2) | 28/12/2012 | NCBI |
| CY167983 | A/Boston/YGA_00016/2012(H3N2) | 28/12/2012 | NCBI |
| CY167991 | A/Boston/YGA_00017/2012(H3N2) | 29/12/2012 | NCBI |
| CY167999 | A/Boston/YGA_00019/2012(H3N2) | 31/12/2012 | NCBI |
| CY168007 | A/Boston/YGA_00020/2012(H3N2) | 28/12/2012 | NCBI |
| CY168015 | A/Boston/YGA_00021/2012(H3N2) | 31/12/2012 | NCBI |
| CY168023 | A/Boston/YGA_00022/2013(H3N2) | 2/1/2013 | NCBI |
| CY168031 | A/Boston/YGA_00024/2013(H3N2) | 3/1/2013 | NCBI |
| CY168039 | A/Boston/YGA_00025/2013(H3N2) | 3/1/2013 | NCBI |
| CY168047 | A/Boston/YGA_00026/2013(H3N2) | 3/1/2013 | NCBI |
| CY168055 | A/Boston/YGA_00027/2013(H3N2) | 3/1/2013 | NCBI |
| CY168063 | A/Boston/YGA_00028/2013(H3N2) | 3/1/2013 | NCBI |
| CY168071 | A/Boston/YGA_00030/2013(H3N2) | 4/1/2013 | NCBI |
| CY168079 | A/Boston/YGA_00031/2013(H3N2) | 9/1/2013 | NCBI |
| CY168087 | A/Boston/YGA_00035/2013(H3N2) | 10/1/2013 | NCBI |
| CY168103 | A/Boston/YGA_00038/2013(H3N2) | 11/1/2013 | NCBI |
| CY168111 | A/Boston/YGA_00039/2013(H3N2) | 11/1/2013 | NCBI |
| CY168119 | A/Boston/YGA_00040/2013(H3N2) | 11/1/2013 | NCBI |
| CY168127 | A/Boston/YGA_00041/2013(H3N2) | 11/1/2013 | NCBI |
| CY168135 | A/Boston/YGA_00042/2013(H3N2) | 12/1/2013 | NCBI |
| CY168143 | A/Boston/YGA_00043/2013(H3N2) | 12/1/2013 | NCBI |

| | | | |
|---|---|---|---|
| CY168151 | A/Boston/YGA_00044/2013(H3N2) | 13/1/2013 | NCBI |
| CY168159 | A/Boston/YGA_00045/2013(H3N2) | 13/1/2013 | NCBI |
| CY168167 | A/Boston/YGA_00046/2013(H3N2) | 14/1/2013 | NCBI |
| CY168175 | A/Boston/YGA_00047/2013(H3N2) | 15/1/2013 | NCBI |
| CY168183 | A/Boston/YGA_00048/2013(H3N2) | 15/1/2013 | NCBI |
| CY168191 | A/Boston/YGA_00049/2013(H3N2) | 15/1/2013 | NCBI |
| CY168199 | A/Boston/YGA_00050/2013(H3N2) | 18/1/2013 | NCBI |
| CY168207 | A/Boston/YGA_00051/2013(H3N2) | 18/1/2013 | NCBI |
| CY168215 | A/Boston/YGA_00052/2013(H3N2) | 18/1/2013 | NCBI |
| CY168223 | A/Boston/YGA_00053/2013(H3N2) | 21/1/2013 | NCBI |
| CY168231 | A/Boston/YGA_00054/2013(H3N2) | 24/1/2013 | NCBI |
| CY168239 | A/Boston/YGA_00055/2013(H3N2) | 1/2/2013 | NCBI |
| CY168247 | A/Boston/YGA_00056/2013(H3N2) | 1/2/2013 | NCBI |
| CY168255 | A/Boston/YGA_00059/2013(H3N2) | 5/2/2013 | NCBI |
| CY168263 | A/Boston/YGA_00060/2013(H3N2) | 6/2/2013 | NCBI |
| CY168271 | A/Boston/YGA_00061/2013(H3N2) | 6/2/2013 | NCBI |
| CY168279 | A/Boston/YGA_00062/2013(H3N2) | 11/2/2013 | NCBI |
| CY168287 | A/Boston/YGA_00064/2013(H3N2) | 15/2/2013 | NCBI |
| CY168295 | A/Boston/YGA_00066/2013(H3N2) | 21/2/2013 | NCBI |
| CY168303 | A/Boston/YGA_00067/2013(H3N2) | 26/2/2013 | NCBI |
| CY168311 | A/Boston/YGA_00068/2013(H3N2) | 28/2/2013 | NCBI |
| CY168319 | A/Boston/YGA_00070/2013(H3N2) | 13/1/2013 | NCBI |
| CY168327 | A/Boston/YGA_00071/2013(H3N2) | 13/1/2013 | NCBI |
| CY168335 | A/Boston/YGA_00074/2013(H3N2) | 18/1/2013 | NCBI |
| CY168343 | A/Boston/YGA_00076/2013(H3N2) | 19/1/2013 | NCBI |
| CY168351 | A/Boston/YGA_00077/2013(H3N2) | 19/1/2013 | NCBI |
| CY168359 | A/Boston/YGA_00078/2013(H3N2) | 20/1/2013 | NCBI |
| CY168367 | A/Boston/YGA_00079/2013(H3N2) | 21/1/2013 | NCBI |
| CY168375 | A/Boston/YGA_00080/2013(H3N2) | 22/1/2013 | NCBI |
| CY168383 | A/Boston/YGA_00081/2013(H3N2) | 23/1/2013 | NCBI |
| CY168391 | A/Boston/YGA_00082/2013(H3N2) | 23/1/2013 | NCBI |
| CY168399 | A/Boston/YGA_00083/2013(H3N2) | 23/1/2013 | NCBI |
| CY168407 | A/Boston/YGA_00084/2013(H3N2) | 25/1/2013 | NCBI |
| CY168415 | A/Boston/YGA_00086/2013(H3N2) | 29/1/2013 | NCBI |
| CY168431 | A/Boston/YGA_00088/2013(H3N2) | 4/2/2013 | NCBI |
| CY168439 | A/Boston/YGA_00089/2012(H3N2) | 19/12/2012 | NCBI |
| CY168447 | A/Boston/YGA_00090/2012(H3N2) | 20/12/2012 | NCBI |

| | | | |
|---|---|---|---|
| CY168455 | A/Boston/YGA_00091/2013(H3N2) | 4/2/2013 | NCBI |
| CY168463 | A/Boston/YGA_00092/2013(H3N2) | 1/2/2013 | NCBI |
| CY168471 | A/Boston/YGA_00093/2012(H3N2) | 20/12/2012 | NCBI |
| CY168479 | A/Boston/YGA_00094/2013(H3N2) | 31/1/2013 | NCBI |
| CY168487 | A/Boston/YGA_00095/2013(H3N2) | 2/2/2013 | NCBI |
| CY168495 | A/Boston/YGA_00096/2013(H3N2) | 28/1/2013 | NCBI |
| CY168503 | A/Boston/YGA_00097/2013(H3N2) | 23/3/2013 | NCBI |
| CY168511 | A/Boston/YGA_00098/2013(H3N2) | 1/4/2013 | NCBI |
| CY168519 | A/Boston/YGA_00099/2013(H3N2) | 23/4/2013 | NCBI |
| CY168527 | A/Boston/YGA_01001/2012(H3N2) | 19/12/2012 | NCBI |
| CY168543 | A/Boston/YGA_01003/2012(H3N2) | 19/12/2012 | NCBI |
| CY168551 | A/Boston/YGA_01004/2012(H3N2) | 20/12/2012 | NCBI |
| CY168559 | A/Boston/YGA_01005/2012(H3N2) | 20/12/2012 | NCBI |
| CY168567 | A/Boston/YGA_01006/2012(H3N2) | 20/12/2012 | NCBI |
| CY168575 | A/Boston/YGA_01008/2012(H3N2) | 20/12/2012 | NCBI |
| CY168583 | A/Boston/YGA_01009/2012(H3N2) | 20/12/2012 | NCBI |
| CY168591 | A/Boston/YGA_01010/2012(H3N2) | 20/12/2012 | NCBI |
| CY168599 | A/Boston/YGA_01011/2012(H3N2) | 21/12/2012 | NCBI |
| CY168607 | A/Boston/YGA_01012/2012(H3N2) | 21/12/2012 | NCBI |
| CY168615 | A/Boston/YGA_01013/2012(H3N2) | 21/12/2012 | NCBI |
| CY168623 | A/Boston/YGA_01014/2012(H3N2) | 21/12/2012 | NCBI |
| CY168631 | A/Boston/YGA_01015/2012(H3N2) | 21/12/2012 | NCBI |
| CY168639 | A/Boston/YGA_01016/2012(H3N2) | 22/12/2012 | NCBI |
| CY168647 | A/Boston/YGA_01017/2012(H3N2) | 21/12/2012 | NCBI |
| CY168655 | A/Boston/YGA_01018/2012(H3N2) | 22/12/2012 | NCBI |
| CY168663 | A/Boston/YGA_01020/2012(H3N2) | 22/12/2012 | NCBI |
| CY168671 | A/Boston/YGA_01022/2012(H3N2) | 22/12/2012 | NCBI |
| CY168679 | A/Boston/YGA_01023/2012(H3N2) | 22/12/2012 | NCBI |
| CY168687 | A/Boston/YGA_01024/2012(H3N2) | 23/12/2012 | NCBI |
| CY168695 | A/Boston/YGA_01025/2012(H3N2) | 22/12/2012 | NCBI |
| CY168703 | A/Boston/YGA_01026/2012(H3N2) | 23/12/2012 | NCBI |
| CY168711 | A/Boston/YGA_01027/2012(H3N2) | 23/12/2012 | NCBI |
| CY168719 | A/Boston/YGA_01028/2012(H3N2) | 23/12/2012 | NCBI |
| CY168727 | A/Boston/YGA_01029/2012(H3N2) | 23/12/2012 | NCBI |
| CY168735 | A/Boston/YGA_01030/2012(H3N2) | 23/12/2012 | NCBI |
| CY168743 | A/Boston/YGA_01031/2012(H3N2) | 23/12/2012 | NCBI |
| CY168751 | A/Boston/YGA_01033/2012(H3N2) | 24/12/2012 | NCBI |

| | | | |
|---|---|---|---|
| CY168759 | A/Boston/YGA_01035/2012(H3N2) | 24/12/2012 | NCBI |
| CY168767 | A/Boston/YGA_01036/2012(H3N2) | 24/12/2012 | NCBI |
| CY168775 | A/Boston/YGA_01037/2012(H3N2) | 24/12/2012 | NCBI |
| CY168783 | A/Boston/YGA_01038/2012(H3N2) | 24/12/2012 | NCBI |
| CY168791 | A/Boston/YGA_01039/2012(H3N2) | 24/12/2012 | NCBI |
| CY168799 | A/Boston/YGA_01040/2012(H3N2) | 24/12/2012 | NCBI |
| CY168815 | A/Boston/YGA_01042/2012(H3N2) | 25/12/2012 | NCBI |
| CY168823 | A/Boston/YGA_01043/2012(H3N2) | 25/12/2012 | NCBI |
| CY168831 | A/Boston/YGA_01044/2012(H3N2) | 25/12/2012 | NCBI |
| CY168839 | A/Boston/YGA_01045/2012(H3N2) | 25/12/2012 | NCBI |
| CY168847 | A/Boston/YGA_01046/2012(H3N2) | 25/12/2012 | NCBI |
| CY168855 | A/Boston/YGA_01047/2012(H3N2) | 25/12/2012 | NCBI |
| CY168863 | A/Boston/YGA_01048/2012(H3N2) | 25/12/2012 | NCBI |
| CY168871 | A/Boston/YGA_01049/2012(H3N2) | 26/12/2012 | NCBI |
| CY168887 | A/Boston/YGA_01051/2012(H3N2) | 25/12/2012 | NCBI |
| CY168895 | A/Boston/YGA_01052/2012(H3N2) | 26/12/2012 | NCBI |
| CY168903 | A/Boston/YGA_01053/2012(H3N2) | 26/12/2012 | NCBI |
| CY168911 | A/Boston/YGA_01054/2012(H3N2) | 26/12/2012 | NCBI |
| CY168919 | A/Boston/YGA_01055/2012(H3N2) | 26/12/2012 | NCBI |
| CY168927 | A/Boston/YGA_01056/2012(H3N2) | 26/12/2012 | NCBI |
| CY168935 | A/Boston/YGA_01058/2012(H3N2) | 26/12/2012 | NCBI |
| CY168943 | A/Boston/YGA_01059/2012(H3N2) | 26/12/2012 | NCBI |
| CY168951 | A/Boston/YGA_01060/2012(H3N2) | 26/12/2012 | NCBI |
| CY168959 | A/Boston/YGA_01061/2012(H3N2) | 26/12/2012 | NCBI |
| CY168967 | A/Boston/YGA_01062/2012(H3N2) | 27/12/2012 | NCBI |
| CY168975 | A/Boston/YGA_01063/2012(H3N2) | 27/12/2012 | NCBI |
| CY168983 | A/Boston/YGA_01064/2012(H3N2) | 27/12/2012 | NCBI |
| CY168991 | A/Boston/YGA_01065/2012(H3N2) | 27/12/2012 | NCBI |
| CY168999 | A/Boston/YGA_01066/2012(H3N2) | 27/12/2012 | NCBI |
| CY169007 | A/Boston/YGA_01067/2012(H3N2) | 27/12/2012 | NCBI |
| CY169015 | A/Boston/YGA_01068/2012(H3N2) | 27/12/2012 | NCBI |
| CY169023 | A/Boston/YGA_01070/2012(H3N2) | 27/12/2012 | NCBI |
| CY169031 | A/Boston/YGA_01071/2012(H3N2) | 27/12/2012 | NCBI |
| CY169039 | A/Boston/YGA_01072/2012(H3N2) | 27/12/2012 | NCBI |
| CY169047 | A/Boston/YGA_01073/2012(H3N2) | 28/12/2012 | NCBI |
| CY169055 | A/Boston/YGA_01074/2012(H3N2) | 28/12/2012 | NCBI |
| CY169063 | A/Boston/YGA_01075/2012(H3N2) | 28/12/2012 | NCBI |

| | | | |
|---|---|---|---|
| CY169071 | A/Boston/YGA_01076/2012(H3N2) | 28/12/2012 | NCBI |
| CY169079 | A/Boston/YGA_01077/2012(H3N2) | 28/12/2012 | NCBI |
| CY169087 | A/Boston/YGA_01078/2012(H3N2) | 28/12/2012 | NCBI |
| CY169095 | A/Boston/YGA_01079/2012(H3N2) | 28/12/2012 | NCBI |
| CY169103 | A/Boston/YGA_01081/2012(H3N2) | 28/12/2012 | NCBI |
| CY169111 | A/Boston/YGA_01082/2012(H3N2) | 28/12/2012 | NCBI |
| CY169119 | A/Boston/YGA_01083/2012(H3N2) | 28/12/2012 | NCBI |
| CY169127 | A/Boston/YGA_01084/2012(H3N2) | 28/12/2012 | NCBI |
| CY169135 | A/Boston/YGA_01085/2012(H3N2) | 28/12/2012 | NCBI |
| CY169143 | A/Boston/YGA_01086/2012(H3N2) | 28/12/2012 | NCBI |
| CY169151 | A/Boston/YGA_01088/2012(H3N2) | 29/12/2012 | NCBI |
| CY169159 | A/Boston/YGA_01089/2012(H3N2) | 29/12/2012 | NCBI |
| CY169167 | A/Boston/YGA_01090/2012(H3N2) | 29/12/2012 | NCBI |
| CY169175 | A/Boston/YGA_01091/2012(H3N2) | 29/12/2012 | NCBI |
| CY169183 | A/Boston/YGA_01092/2012(H3N2) | 29/12/2012 | NCBI |
| CY169191 | A/Boston/YGA_01093/2012(H3N2) | 29/12/2012 | NCBI |
| CY169199 | A/Boston/YGA_01095/2012(H3N2) | 30/12/2012 | NCBI |
| CY169207 | A/Boston/YGA_01096/2012(H3N2) | 30/12/2012 | NCBI |
| CY169215 | A/Boston/YGA_01097/2012(H3N2) | 30/12/2012 | NCBI |
| CY169223 | A/Boston/YGA_01098/2012(H3N2) | 30/12/2012 | NCBI |
| CY169231 | A/Boston/YGA_01099/2012(H3N2) | 31/12/2012 | NCBI |
| CY169239 | A/Boston/YGA_01100/2012(H3N2) | 31/12/2012 | NCBI |
| CY169247 | A/Boston/YGA_01101/2012(H3N2) | 31/12/2012 | NCBI |
| CY169255 | A/Boston/YGA_01102/2012(H3N2) | 31/12/2012 | NCBI |
| CY169263 | A/Boston/YGA_01103/2012(H3N2) | 31/12/2012 | NCBI |
| CY169271 | A/Boston/YGA_01104/2012(H3N2) | 31/12/2012 | NCBI |
| CY169279 | A/Boston/YGA_01105/2012(H3N2) | 31/12/2012 | NCBI |
| CY169287 | A/Boston/YGA_01106/2012(H3N2) | 31/12/2012 | NCBI |
| CY169295 | A/Boston/YGA_01107/2012(H3N2) | 31/12/2012 | NCBI |
| CY169303 | A/Boston/YGA_01108/2012(H3N2) | 31/12/2012 | NCBI |
| CY169311 | A/Boston/YGA_01109/2012(H3N2) | 31/12/2012 | NCBI |
| CY169319 | A/Boston/YGA_01110/2012(H3N2) | 31/12/2012 | NCBI |
| CY169327 | A/Boston/YGA_01111/2013(H3N2) | 1/1/2013 | NCBI |
| CY169335 | A/Boston/YGA_01112/2013(H3N2) | 1/1/2013 | NCBI |
| CY169343 | A/Boston/YGA_01113/2013(H3N2) | 1/1/2013 | NCBI |
| CY169351 | A/Boston/YGA_01114/2013(H3N2) | 1/1/2013 | NCBI |
| CY169359 | A/Boston/YGA_01115/2013(H3N2) | 1/1/2013 | NCBI |

| | | | |
|---|---|---|---|
| CY169367 | A/Boston/YGA_01118/2013(H3N2) | 2/1/2013 | NCBI |
| CY169375 | A/Boston/YGA_01120/2013(H3N2) | 2/1/2013 | NCBI |
| CY169383 | A/Boston/YGA_01121/2013(H3N2) | 2/1/2013 | NCBI |
| CY169391 | A/Boston/YGA_01122/2013(H3N2) | 2/1/2013 | NCBI |
| CY169399 | A/Boston/YGA_01123/2013(H3N2) | 2/1/2013 | NCBI |
| CY169407 | A/Boston/YGA_01124/2013(H3N2) | 2/1/2013 | NCBI |
| CY169415 | A/Boston/YGA_01125/2013(H3N2) | 2/1/2013 | NCBI |
| CY169423 | A/Boston/YGA_01126/2013(H3N2) | 2/1/2013 | NCBI |
| CY169431 | A/Boston/YGA_01127/2013(H3N2) | 2/1/2013 | NCBI |
| CY169439 | A/Boston/YGA_01128/2013(H3N2) | 2/1/2013 | NCBI |
| CY169447 | A/Boston/YGA_01129/2013(H3N2) | 3/1/2013 | NCBI |
| CY169455 | A/Boston/YGA_01130/2013(H3N2) | 2/1/2013 | NCBI |
| CY169463 | A/Boston/YGA_01131/2013(H3N2) | 3/1/2013 | NCBI |
| CY169471 | A/Boston/YGA_01132/2013(H3N2) | 3/1/2013 | NCBI |
| CY169479 | A/Boston/YGA_01133/2013(H3N2) | 3/1/2013 | NCBI |
| CY169487 | A/Boston/YGA_01134/2013(H3N2) | 3/1/2013 | NCBI |
| CY169495 | A/Boston/YGA_01135/2013(H3N2) | 3/1/2013 | NCBI |
| CY169503 | A/Boston/YGA_01136/2013(H3N2) | 3/1/2013 | NCBI |
| CY169511 | A/Boston/YGA_01137/2013(H3N2) | 3/1/2013 | NCBI |
| CY169519 | A/Boston/YGA_01139/2013(H3N2) | 3/1/2013 | NCBI |
| CY169527 | A/Boston/YGA_01140/2013(H3N2) | 3/1/2013 | NCBI |
| CY169535 | A/Boston/YGA_01141/2013(H3N2) | 4/1/2013 | NCBI |
| CY169543 | A/Boston/YGA_01142/2013(H3N2) | 4/1/2013 | NCBI |
| CY169551 | A/Boston/YGA_01143/2013(H3N2) | 4/1/2013 | NCBI |
| CY169559 | A/Boston/YGA_01144/2013(H3N2) | 4/1/2013 | NCBI |
| CY169567 | A/Boston/YGA_01145/2013(H3N2) | 4/1/2013 | NCBI |
| CY169575 | A/Boston/YGA_01146/2013(H3N2) | 5/1/2013 | NCBI |
| CY169583 | A/Boston/YGA_01147/2013(H3N2) | 5/1/2013 | NCBI |
| CY169591 | A/Boston/YGA_01148/2013(H3N2) | 5/1/2013 | NCBI |
| CY169599 | A/Boston/YGA_01149/2013(H3N2) | 5/1/2013 | NCBI |
| CY169607 | A/Boston/YGA_01150/2013(H3N2) | 5/1/2013 | NCBI |
| CY169615 | A/Boston/YGA_01151/2013(H3N2) | 5/1/2013 | NCBI |
| CY169623 | A/Boston/YGA_01152/2013(H3N2) | 5/1/2013 | NCBI |
| CY169631 | A/Boston/YGA_01153/2013(H3N2) | 5/1/2013 | NCBI |
| CY169639 | A/Boston/YGA_01154/2013(H3N2) | 6/1/2013 | NCBI |
| CY169647 | A/Boston/YGA_01155/2013(H3N2) | 6/1/2013 | NCBI |
| CY169655 | A/Boston/YGA_01156/2013(H3N2) | 5/1/2013 | NCBI |

| | | | |
|---|---|---|---|
| CY169663 | A/Boston/YGA_01157/2013(H3N2) | 6/1/2013 | NCBI |
| CY169671 | A/Boston/YGA_01158/2013(H3N2) | 6/1/2013 | NCBI |
| CY169679 | A/Boston/YGA_01159/2013(H3N2) | 6/1/2013 | NCBI |
| CY169687 | A/Boston/YGA_01160/2013(H3N2) | 7/1/2013 | NCBI |
| CY169695 | A/Boston/YGA_01161/2013(H3N2) | 7/1/2013 | NCBI |
| CY169703 | A/Boston/YGA_01162/2013(H3N2) | 7/1/2013 | NCBI |
| CY169711 | A/Boston/YGA_01163/2013(H3N2) | 10/1/2013 | NCBI |
| CY169719 | A/Boston/YGA_01164/2013(H3N2) | 10/1/2013 | NCBI |
| CY169727 | A/Boston/YGA_01165/2013(H3N2) | 10/1/2013 | NCBI |
| CY169735 | A/Boston/YGA_01166/2013(H3N2) | 10/1/2013 | NCBI |
| CY169743 | A/Boston/YGA_01167/2013(H3N2) | 10/1/2013 | NCBI |
| CY169751 | A/Boston/YGA_01168/2013(H3N2) | 11/1/2013 | NCBI |
| CY169759 | A/Boston/YGA_01169/2013(H3N2) | 13/1/2013 | NCBI |
| CY169767 | A/Boston/YGA_01170/2013(H3N2) | 13/1/2013 | NCBI |
| CY169775 | A/Boston/YGA_01171/2013(H3N2) | 14/1/2013 | NCBI |
| CY169783 | A/Boston/YGA_01172/2013(H3N2) | 14/1/2013 | NCBI |
| CY169791 | A/Boston/YGA_01173/2013(H3N2) | 17/1/2013 | NCBI |
| CY169799 | A/Boston/YGA_01174/2013(H3N2) | 17/1/2013 | NCBI |
| CY169807 | A/Boston/YGA_01175/2013(H3N2) | 18/1/2013 | NCBI |
| CY169815 | A/Boston/YGA_01176/2013(H3N2) | 18/1/2013 | NCBI |
| CY169823 | A/Boston/YGA_01180/2013(H3N2) | 20/1/2013 | NCBI |
| CY169831 | A/Boston/YGA_01181/2013(H3N2) | 20/1/2013 | NCBI |
| CY169839 | A/Boston/YGA_01182/2013(H3N2) | 21/1/2013 | NCBI |
| CY169847 | A/Boston/YGA_01183/2013(H3N2) | 21/1/2013 | NCBI |
| CY169855 | A/Boston/YGA_01184/2013(H3N2) | 21/1/2013 | NCBI |
| CY169871 | A/Boston/YGA_01186/2013(H3N2) | 21/1/2013 | NCBI |
| CY169879 | A/Boston/YGA_01189/2013(H3N2) | 23/1/2013 | NCBI |
| CY169887 | A/Boston/YGA_01190/2013(H3N2) | 23/1/2013 | NCBI |
| CY169895 | A/Boston/YGA_01191/2013(H3N2) | 23/1/2013 | NCBI |
| CY169903 | A/Boston/YGA_01192/2013(H3N2) | 23/1/2013 | NCBI |
| CY169911 | A/Boston/YGA_01193/2013(H3N2) | 24/1/2013 | NCBI |
| CY169919 | A/Boston/YGA_01194/2013(H3N2) | 24/1/2013 | NCBI |
| CY169927 | A/Boston/YGA_01196/2013(H3N2) | 24/1/2013 | NCBI |
| CY169935 | A/Boston/YGA_01197/2013(H3N2) | 28/1/2013 | NCBI |
| CY169943 | A/Boston/YGA_01198/2013(H3N2) | 28/1/2013 | NCBI |
| CY169951 | A/Boston/YGA_01199/2013(H3N2) | 29/1/2013 | NCBI |
| CY169959 | A/Boston/YGA_01200/2013(H3N2) | 1/2/2013 | NCBI |

| | | | |
|---|---|---|---|
| CY169967 | A/Boston/YGA_01201/2013(H3N2) | 2/2/2013 | NCBI |
| CY169983 | A/Boston/YGA_01203/2013(H3N2) | 5/2/2013 | NCBI |
| CY169991 | A/Boston/YGA_01204/2013(H3N2) | 5/2/2013 | NCBI |
| CY169999 | A/Boston/YGA_01205/2013(H3N2) | 5/2/2013 | NCBI |
| CY170007 | A/Boston/YGA_01206/2013(H3N2) | 7/2/2013 | NCBI |
| CY170015 | A/Boston/YGA_01207/2013(H3N2) | 7/2/2013 | NCBI |
| CY170023 | A/Boston/YGA_01208/2013(H3N2) | 10/2/2013 | NCBI |
| CY170031 | A/Boston/YGA_01209/2013(H3N2) | 11/2/2013 | NCBI |
| CY170039 | A/Boston/YGA_01210/2013(H3N2) | 13/2/2013 | NCBI |
| CY170047 | A/Boston/YGA_01212/2013(H3N2) | 20/2/2013 | NCBI |
| CY170055 | A/Boston/YGA_01213/2013(H3N2) | 21/2/2013 | NCBI |
| CY170063 | A/Boston/YGA_01214/2013(H3N2) | 7/3/2013 | NCBI |
| CY170071 | A/Boston/YGA_01216/2013(H3N2) | 14/3/2013 | NCBI |
| CY170103 | A/Boston/YGA_02001/2012(H3N2) | 6/12/2012 | NCBI |
| CY170111 | A/Boston/YGA_02002/2012(H3N2) | 12/12/2012 | NCBI |
| CY170119 | A/Boston/YGA_02003/2012(H3N2) | 17/12/2012 | NCBI |
| CY170127 | A/Boston/YGA_02004/2012(H3N2) | 17/12/2012 | NCBI |
| CY170135 | A/Boston/YGA_02005/2012(H3N2) | 17/12/2012 | NCBI |
| CY170143 | A/Boston/YGA_02006/2012(H3N2) | 18/12/2012 | NCBI |
| CY170151 | A/Boston/YGA_02008/2012(H3N2) | 21/12/2012 | NCBI |
| CY170159 | A/Boston/YGA_02009/2012(H3N2) | 21/12/2012 | NCBI |
| CY170167 | A/Boston/YGA_02010/2012(H3N2) | 24/12/2012 | NCBI |
| CY170175 | A/Boston/YGA_02012/2012(H3N2) | 27/12/2012 | NCBI |
| CY170183 | A/Boston/YGA_02013/2012(H3N2) | 27/12/2012 | NCBI |
| CY170191 | A/Boston/YGA_02014/2012(H3N2) | 31/12/2012 | NCBI |
| CY170199 | A/Boston/YGA_02015/2012(H3N2) | 31/12/2012 | NCBI |
| CY170207 | A/Boston/YGA_02017/2012(H3N2) | 31/12/2012 | NCBI |
| CY170215 | A/Boston/YGA_02018/2013(H3N2) | 2/1/2013 | NCBI |
| CY170223 | A/Boston/YGA_02019/2013(H3N2) | 4/1/2013 | NCBI |
| CY170231 | A/Boston/YGA_02021/2013(H3N2) | 4/1/2013 | NCBI |
| CY170239 | A/Boston/YGA_02022/2013(H3N2) | 7/1/2013 | NCBI |
| CY170247 | A/Boston/YGA_02023/2013(H3N2) | 7/1/2013 | NCBI |
| CY170263 | A/Boston/YGA_02025/2013(H3N2) | 8/1/2013 | NCBI |
| CY170271 | A/Boston/YGA_02026/2013(H3N2) | 9/1/2013 | NCBI |
| CY170279 | A/Boston/YGA_02027/2013(H3N2) | 10/1/2013 | NCBI |
| CY170287 | A/Boston/YGA_02031/2013(H3N2) | 14/1/2013 | NCBI |
| CY170295 | A/Boston/YGA_02032/2013(H3N2) | 14/1/2013 | NCBI |

| | | | |
|---|---|---|---|
| CY170303 | A/Boston/YGA_02033/2013(H3N2) | 15/1/2013 | NCBI |
| CY170311 | A/Boston/YGA_02034/2013(H3N2) | 16/1/2013 | NCBI |
| CY170319 | A/Boston/YGA_02035/2013(H3N2) | 16/1/2013 | NCBI |
| CY170327 | A/Boston/YGA_02036/2013(H3N2) | 17/1/2013 | NCBI |
| CY170335 | A/Boston/YGA_02037/2013(H3N2) | 18/1/2013 | NCBI |
| CY170343 | A/Boston/YGA_02039/2013(H3N2) | 19/1/2013 | NCBI |
| CY170351 | A/Boston/YGA_02040/2013(H3N2) | 20/1/2013 | NCBI |
| CY170359 | A/Boston/YGA_02041/2013(H3N2) | 21/1/2013 | NCBI |
| CY170367 | A/Boston/YGA_02042/2013(H3N2) | 22/1/2013 | NCBI |
| CY170375 | A/Boston/YGA_02043/2013(H3N2) | 24/1/2013 | NCBI |
| CY170391 | A/Boston/YGA_02047/2013(H3N2) | 27/1/2013 | NCBI |
| CY170399 | A/Boston/YGA_02049/2013(H3N2) | 31/1/2013 | NCBI |
| CY170407 | A/Boston/YGA_02051/2013(H3N2) | 1/2/2013 | NCBI |
| CY170415 | A/Boston/YGA_02053/2013(H3N2) | 12/2/2013 | NCBI |
| CY170423 | A/Boston/YGA_02054/2013(H3N2) | 12/2/2013 | NCBI |
| CY170431 | A/Boston/YGA_02055/2013(H3N2) | 12/2/2013 | NCBI |
| CY170439 | A/Boston/YGA_02058/2013(H3N2) | 20/2/2013 | NCBI |
| CY170447 | A/SantaClara/YGA_03001/2013(H3N2) | 2/1/2013 | NCBI |
| CY170455 | A/SantaClara/YGA_03002/2013(H3N2) | 23/1/2013 | NCBI |
| CY170463 | A/SantaClara/YGA_03003/2013(H3N2) | 7/1/2013 | NCBI |
| CY170471 | A/SantaClara/YGA_03004/2013(H3N2) | 30/1/2013 | NCBI |
| CY170479 | A/SantaClara/YGA_03005/2013(H3N2) | 22/1/2013 | NCBI |
| CY170487 | A/SantaClara/YGA_03006/2013(H3N2) | 15/1/2013 | NCBI |
| CY170495 | A/SantaClara/YGA_03007/2013(H3N2) | 29/1/2013 | NCBI |
| CY170503 | A/SantaClara/YGA_03008/2013(H3N2) | 23/1/2013 | NCBI |
| CY170511 | A/SantaClara/YGA_03009/2013(H3N2) | 23/1/2013 | NCBI |
| CY170519 | A/SantaClara/YGA_03010/2013(H3N2) | 13/1/2013 | NCBI |
| CY170527 | A/SantaClara/YGA_03011/2013(H3N2) | 8/1/2013 | NCBI |
| CY170535 | A/SantaClara/YGA_03012/2013(H3N2) | 21/1/2013 | NCBI |
| CY170543 | A/SantaClara/YGA_03013/2013(H3N2) | 2/1/2013 | NCBI |
| CY170551 | A/SantaClara/YGA_03014/2013(H3N2) | 9/1/2013 | NCBI |
| CY170559 | A/SantaClara/YGA_03015/2013(H3N2) | 23/1/2013 | NCBI |
| CY170567 | A/SantaClara/YGA_03016/2013(H3N2) | 17/1/2013 | NCBI |
| CY170575 | A/SantaClara/YGA_03017/2013(H3N2) | 9/1/2013 | NCBI |
| CY170583 | A/SantaClara/YGA_03018/2013(H3N2) | 16/1/2013 | NCBI |
| CY170591 | A/SantaClara/YGA_03019/2013(H3N2) | 16/1/2013 | NCBI |
| CY170599 | A/SantaClara/YGA_03020/2013(H3N2) | 20/1/2013 | NCBI |

| | | | |
|---|---|---|---|
| CY170607 | A/SantaClara/YGA_03021/2013(H3N2) | 2/1/2013 | NCBI |
| CY170615 | A/SantaClara/YGA_03022/2013(H3N2) | 29/1/2013 | NCBI |
| CY170623 | A/SantaClara/YGA_03023/2013(H3N2) | 29/1/2013 | NCBI |
| CY170631 | A/SantaClara/YGA_03024/2013(H3N2) | 22/1/2013 | NCBI |
| CY170639 | A/SantaClara/YGA_03025/2013(H3N2) | 8/1/2013 | NCBI |
| CY170647 | A/SantaClara/YGA_03026/2013(H3N2) | 7/1/2013 | NCBI |
| CY170655 | A/SantaClara/YGA_03027/2013(H3N2) | 9/1/2013 | NCBI |
| CY170663 | A/SantaClara/YGA_03028/2013(H3N2) | 25/1/2013 | NCBI |
| CY170687 | A/SantaClara/YGA_03032/2013(H3N2) | 7/1/2013 | NCBI |
| CY170703 | A/SantaClara/YGA_03034/2013(H3N2) | 17/1/2013 | NCBI |
| CY170711 | A/SantaClara/YGA_03035/2013(H3N2) | 17/1/2013 | NCBI |
| CY170719 | A/SantaClara/YGA_03036/2013(H3N2) | 30/1/2013 | NCBI |
| CY170727 | A/SantaClara/YGA_03037/2013(H3N2) | 31/1/2013 | NCBI |
| CY170735 | A/SantaClara/YGA_03038/2013(H3N2) | 12/1/2013 | NCBI |
| CY170743 | A/SantaClara/YGA_03039/2013(H3N2) | 9/1/2013 | NCBI |
| CY170751 | A/SantaClara/YGA_03040/2013(H3N2) | 13/1/2013 | NCBI |
| CY170759 | A/SantaClara/YGA_03041/2013(H3N2) | 29/1/2013 | NCBI |
| CY170767 | A/SantaClara/YGA_03042/2013(H3N2) | 3/1/2013 | NCBI |
| CY170775 | A/SantaClara/YGA_03044/2013(H3N2) | 24/1/2013 | NCBI |
| CY170783 | A/SantaClara/YGA_03045/2013(H3N2) | 2/2/2013 | NCBI |
| CY170791 | A/SantaClara/YGA_03046/2013(H3N2) | 2/2/2013 | NCBI |
| CY170799 | A/SantaClara/YGA_03047/2013(H3N2) | 2/2/2013 | NCBI |
| CY170807 | A/SantaClara/YGA_03048/2013(H3N2) | 18/1/2013 | NCBI |
| CY170815 | A/SantaClara/YGA_03049/2012(H3N2) | 23/12/2012 | NCBI |
| CY170823 | A/SantaClara/YGA_03050/2013(H3N2) | 24/1/2013 | NCBI |
| CY170831 | A/SantaClara/YGA_03052/2013(H3N2) | 25/1/2013 | NCBI |
| CY170847 | A/SantaClara/YGA_03054/2013(H3N2) | 15/1/2013 | NCBI |
| CY170855 | A/SantaClara/YGA_03055/2013(H3N2) | 14/1/2013 | NCBI |
| CY170863 | A/SantaClara/YGA_03056/2013(H3N2) | 24/1/2013 | NCBI |
| CY170871 | A/SantaClara/YGA_03057/2012(H3N2) | 24/12/2012 | NCBI |
| CY170879 | A/SantaClara/YGA_03059/2013(H3N2) | 27/1/2013 | NCBI |
| CY170887 | A/SantaClara/YGA_03060/2013(H3N2) | 6/1/2013 | NCBI |
| CY170895 | A/SantaClara/YGA_03061/2013(H3N2) | 4/2/2013 | NCBI |
| CY170903 | A/SantaClara/YGA_03062/2012(H3N2) | 23/12/2012 | NCBI |
| CY170911 | A/SantaClara/YGA_03063/2013(H3N2) | 21/1/2013 | NCBI |
| CY170919 | A/SantaClara/YGA_03064/2013(H3N2) | 25/1/2013 | NCBI |
| CY170935 | A/SantaClara/YGA_03066/2013(H3N2) | 27/1/2013 | NCBI |

| | | | |
|---|---|---|---|
| CY170943 | A/SantaClara/YGA_03067/2012(H3N2) | 28/12/2012 | NCBI |
| CY170951 | A/SantaClara/YGA_03068/2012(H3N2) | 5/12/2012 | NCBI |
| CY170959 | A/SantaClara/YGA_03069/2013(H3N2) | 10/1/2013 | NCBI |
| CY170967 | A/SantaClara/YGA_03070/2013(H3N2) | 3/2/2013 | NCBI |
| CY167868 | A/SantaClara/YGA_03071/2013(H3N2) | 1/2/2013 | NCBI |
| CY170975 | A/SantaClara/YGA_03072/2013(H3N2) | 1/2/2013 | NCBI |
| CY170983 | A/SantaClara/YGA_03073/2013(H3N2) | 7/1/2013 | NCBI |
| CY170991 | A/SantaClara/YGA_03074/2013(H3N2) | 8/1/2013 | NCBI |
| CY170999 | A/SantaClara/YGA_03075/2013(H3N2) | 4/2/2013 | NCBI |
| CY171007 | A/SantaClara/YGA_03076/2013(H3N2) | 3/2/2013 | NCBI |
| CY171015 | A/SantaClara/YGA_03077/2013(H3N2) | 9/1/2013 | NCBI |
| CY171023 | A/SantaClara/YGA_03078/2013(H3N2) | 22/1/2013 | NCBI |
| CY171031 | A/SantaClara/YGA_03079/2013(H3N2) | 24/1/2013 | NCBI |
| CY171039 | A/SantaClara/YGA_03080/2013(H3N2) | 14/1/2013 | NCBI |
| CY171047 | A/Chicago/YGA_04001/2012(H3N2) | 5/11/2012 | NCBI |
| CY171055 | A/Chicago/YGA_04002/2012(H3N2) | 12/11/2012 | NCBI |
| CY171063 | A/Chicago/YGA_04006/2012(H3N2) | 27/11/2012 | NCBI |
| CY171071 | A/Chicago/YGA_04007/2012(H3N2) | 27/11/2012 | NCBI |
| CY171079 | A/Chicago/YGA_04008/2012(H3N2) | 29/11/2012 | NCBI |
| CY171087 | A/Chicago/YGA_04009/2012(H3N2) | 30/11/2012 | NCBI |
| CY171095 | A/Chicago/YGA_04010/2012(H3N2) | 2/12/2012 | NCBI |
| CY171103 | A/Chicago/YGA_04011/2012(H3N2) | 2/12/2012 | NCBI |
| CY171111 | A/Chicago/YGA_04012/2012(H3N2) | 3/12/2012 | NCBI |
| CY171119 | A/Chicago/YGA_04013/2012(H3N2) | 4/12/2012 | NCBI |
| CY171127 | A/Chicago/YGA_04014/2012(H3N2) | 5/12/2012 | NCBI |
| CY171135 | A/Chicago/YGA_04015/2012(H3N2) | 6/12/2012 | NCBI |
| CY171143 | A/Chicago/YGA_04017/2012(H3N2) | 7/12/2012 | NCBI |
| CY171151 | A/Chicago/YGA_04018/2012(H3N2) | 7/12/2012 | NCBI |
| CY171167 | A/Chicago/YGA_04020/2012(H3N2) | 7/12/2012 | NCBI |
| CY171175 | A/Chicago/YGA_04035/2012(H3N2) | 13/12/2012 | NCBI |
| CY171183 | A/Chicago/YGA_04036/2012(H3N2) | 13/12/2012 | NCBI |
| CY171191 | A/Chicago/YGA_04037/2012(H3N2) | 13/12/2012 | NCBI |
| CY171199 | A/Chicago/YGA_04038/2012(H3N2) | 14/12/2012 | NCBI |
| CY171207 | A/Chicago/YGA_04039/2012(H3N2) | 15/12/2012 | NCBI |
| CY171215 | A/Chicago/YGA_04046/2012(H3N2) | 17/12/2012 | NCBI |
| CY171223 | A/Chicago/YGA_04047/2012(H3N2) | 17/12/2012 | NCBI |
| CY171231 | A/Chicago/YGA_04048/2012(H3N2) | 17/12/2012 | NCBI |

| | | | |
|---|---|---|---|
| CY171239 | A/Chicago/YGA_04053/2012(H3N2) | 18/12/2012 | NCBI |
| CY171247 | A/Chicago/YGA_04054/2012(H3N2) | 18/12/2012 | NCBI |
| CY171255 | A/Chicago/YGA_04055/2012(H3N2) | 18/12/2012 | NCBI |
| CY171263 | A/Chicago/YGA_04056/2012(H3N2) | 18/12/2012 | NCBI |
| CY171271 | A/Chicago/YGA_04064/2012(H3N2) | 18/12/2012 | NCBI |
| CY171279 | A/Chicago/YGA_04065/2012(H3N2) | 19/12/2012 | NCBI |
| CY171287 | A/Chicago/YGA_04066/2012(H3N2) | 19/12/2012 | NCBI |
| CY171295 | A/Chicago/YGA_04081/2012(H3N2) | 24/12/2012 | NCBI |
| CY171303 | A/Chicago/YGA_04084/2012(H3N2) | 24/12/2012 | NCBI |
| CY171311 | A/Chicago/YGA_04085/2012(H3N2) | 25/12/2012 | NCBI |
| CY171319 | A/Chicago/YGA_04086/2012(H3N2) | 25/12/2012 | NCBI |
| CY171327 | A/Chicago/YGA_04089/2012(H3N2) | 26/12/2012 | NCBI |
| CY171335 | A/Chicago/YGA_04090/2012(H3N2) | 26/12/2012 | NCBI |
| CY171343 | A/Chicago/YGA_04092/2012(H3N2) | 26/12/2012 | NCBI |
| CY171351 | A/Chicago/YGA_04094/2012(H3N2) | 26/12/2012 | NCBI |
| CY171359 | A/Chicago/YGA_04095/2012(H3N2) | 26/12/2012 | NCBI |
| CY171367 | A/Chicago/YGA_04096/2012(H3N2) | 26/12/2012 | NCBI |
| CY171375 | A/Chicago/YGA_04097/2012(H3N2) | 26/12/2012 | NCBI |
| CY171383 | A/Chicago/YGA_04098/2012(H3N2) | 27/12/2012 | NCBI |
| CY171391 | A/Chicago/YGA_04099/2012(H3N2) | 27/12/2012 | NCBI |
| CY171399 | A/Chicago/YGA_04102/2012(H3N2) | 27/12/2012 | NCBI |
| CY171407 | A/Chicago/YGA_04103/2012(H3N2) | 27/12/2012 | NCBI |
| CY171415 | A/Chicago/YGA_04104/2012(H3N2) | 27/12/2012 | NCBI |
| CY171423 | A/Chicago/YGA_04105/2012(H3N2) | 27/12/2012 | NCBI |
| CY171431 | A/Chicago/YGA_04106/2012(H3N2) | 28/12/2012 | NCBI |
| CY171439 | A/Chicago/YGA_04107/2012(H3N2) | 28/12/2012 | NCBI |
| CY171447 | A/Chicago/YGA_04110/2012(H3N2) | 28/12/2012 | NCBI |
| CY171455 | A/Chicago/YGA_04111/2012(H3N2) | 28/12/2012 | NCBI |
| CY171463 | A/Chicago/YGA_04112/2012(H3N2) | 28/12/2012 | NCBI |
| CY171471 | A/Chicago/YGA_04113/2012(H3N2) | 28/12/2012 | NCBI |
| CY171479 | A/Chicago/YGA_04114/2012(H3N2) | 28/12/2012 | NCBI |
| CY171487 | A/Chicago/YGA_04115/2012(H3N2) | 28/12/2012 | NCBI |
| CY171495 | A/Chicago/YGA_04117/2012(H3N2) | 29/12/2012 | NCBI |
| CY171503 | A/Chicago/YGA_04118/2012(H3N2) | 29/12/2012 | NCBI |
| CY171511 | A/Chicago/YGA_04119/2012(H3N2) | 29/12/2012 | NCBI |
| CY171519 | A/Chicago/YGA_04120/2012(H3N2) | 30/12/2012 | NCBI |
| CY171527 | A/Chicago/YGA_04121/2012(H3N2) | 30/12/2012 | NCBI |

| | | | |
|---|---|---|---|
| CY171535 | A/Chicago/YGA_04122/2012(H3N2) | 30/12/2012 | NCBI |
| CY171551 | A/Chicago/YGA_04124/2012(H3N2) | 31/12/2012 | NCBI |
| CY171559 | A/Chicago/YGA_04126/2012(H3N2) | 31/12/2012 | NCBI |
| CY171567 | A/Chicago/YGA_04127/2013(H3N2) | 1/1/2013 | NCBI |
| CY171575 | A/Chicago/YGA_04128/2013(H3N2) | 1/1/2013 | NCBI |
| CY171583 | A/Chicago/YGA_04129/2013(H3N2) | 1/1/2013 | NCBI |
| CY171591 | A/Chicago/YGA_04130/2013(H3N2) | 1/1/2013 | NCBI |
| CY171599 | A/Chicago/YGA_04131/2013(H3N2) | 1/1/2013 | NCBI |
| CY171607 | A/Chicago/YGA_04132/2013(H3N2) | 1/1/2013 | NCBI |
| CY171615 | A/Chicago/YGA_04133/2013(H3N2) | 2/1/2013 | NCBI |
| CY171623 | A/Chicago/YGA_04134/2013(H3N2) | 2/1/2013 | NCBI |
| CY171631 | A/Chicago/YGA_04135/2013(H3N2) | 3/1/2013 | NCBI |
| CY171639 | A/Chicago/YGA_04136/2013(H3N2) | 3/1/2013 | NCBI |
| CY171647 | A/Chicago/YGA_04142/2013(H3N2) | 4/1/2013 | NCBI |
| CY171655 | A/Chicago/YGA_04149/2013(H3N2) | 7/1/2013 | NCBI |
| CY171663 | A/Chicago/YGA_04150/2013(H3N2) | 7/1/2013 | NCBI |
| CY171671 | A/Chicago/YGA_04164/2013(H3N2) | 9/1/2013 | NCBI |
| CY171679 | A/Chicago/YGA_04165/2013(H3N2) | 9/1/2013 | NCBI |
| CY171687 | A/Chicago/YGA_04183/2013(H3N2) | 14/1/2013 | NCBI |
| CY171695 | A/Chicago/YGA_04184/2013(H3N2) | 14/1/2013 | NCBI |
| CY171703 | A/Chicago/YGA_04187/2013(H3N2) | 16/1/2013 | NCBI |
| CY171711 | A/Chicago/YGA_04188/2013(H3N2) | 16/1/2013 | NCBI |
| CY171719 | A/Chicago/YGA_04193/2013(H3N2) | 19/1/2013 | NCBI |
| CY171727 | A/Chicago/YGA_04198/2013(H3N2) | 23/1/2013 | NCBI |
| CY171735 | A/Chicago/YGA_04200/2013(H3N2) | 25/1/2013 | NCBI |
| KJ623709 | A/Pavia/07/2014(H3N2) | 2014-01 | NCBI |
| CY181585 | A/Nicaragua/AGA2-01/2012(H3N2) | 15/9/2012 | NCBI |
| CY181593 | A/Nicaragua/AGA2-02/2012(H3N2) | 16/9/2012 | NCBI |
| CY181601 | A/Nicaragua/AGA2-03/2012(H3N2) | 19/9/2012 | NCBI |
| CY181609 | A/Nicaragua/AGA2-04/2012(H3N2) | 19/9/2012 | NCBI |
| CY181617 | A/Nicaragua/AGA2-05/2012(H3N2) | 20/9/2012 | NCBI |
| CY181625 | A/Nicaragua/AGA2-06/2012(H3N2) | 18/9/2012 | NCBI |
| CY181633 | A/Nicaragua/AGA2-07/2012(H3N2) | 20/9/2012 | NCBI |
| CY181641 | A/Nicaragua/AGA2-08/2012(H3N2) | 27/9/2012 | NCBI |
| CY181649 | A/Nicaragua/AGA2-09/2012(H3N2) | 31/8/2012 | NCBI |
| CY181657 | A/Nicaragua/AGA2-11/2012(H3N2) | 5/9/2012 | NCBI |
| CY181665 | A/Nicaragua/AGA2-12/2012(H3N2) | 29/9/2012 | NCBI |

| | | | |
|---|---|---|---|
| CY181673 | A/Nicaragua/AGA2-13/2012(H3N2) | 30/9/2012 | NCBI |
| CY181681 | A/Nicaragua/AGA2-14/2012(H3N2) | 1/10/2012 | NCBI |
| CY181689 | A/Nicaragua/AGA2-17/2012(H3N2) | 3/10/2012 | NCBI |
| CY181697 | A/Nicaragua/AGA2-18/2012(H3N2) | 7/9/2012 | NCBI |
| CY181705 | A/Nicaragua/AGA2-19/2012(H3N2) | 8/10/2012 | NCBI |
| CY181713 | A/Nicaragua/AGA2-20/2012(H3N2) | 11/10/2012 | NCBI |
| CY181721 | A/Nicaragua/AGA2-21/2012(H3N2) | 10/10/2012 | NCBI |
| CY181729 | A/Nicaragua/AGA2-23/2011(H3N2) | 4/10/2011 | NCBI |
| CY181745 | A/Nicaragua/AGA2-25/2011(H3N2) | 6/10/2011 | NCBI |
| CY181785 | A/Nicaragua/AGA2-30/2011(H3N2) | 10/10/2011 | NCBI |
| CY181793 | A/Nicaragua/AGA2-32/2011(H3N2) | 12/10/2011 | NCBI |
| CY181841 | A/Nicaragua/AGA2-40/2011(H3N2) | 17/10/2011 | NCBI |
| CY181905 | A/Nicaragua/AGA2-48/2011(H3N2) | 24/10/2011 | NCBI |
| CY181913 | A/Nicaragua/AGA2-49/2011(H3N2) | 24/10/2011 | NCBI |
| CY182017 | A/Nicaragua/AGA2-64/2011(H3N2) | 3/11/2011 | NCBI |
| CY182025 | A/Nicaragua/AGA2-65/2011(H3N2) | 6/11/2011 | NCBI |
| CY182041 | A/Nicaragua/AGA2-67/2011(H3N2) | 7/11/2011 | NCBI |
| CY182073 | A/Nicaragua/AGA2-72/2012(H3N2) | 9/7/2012 | NCBI |
| CY182089 | A/Nicaragua/AGA2-76/2012(H3N2) | 26/7/2012 | NCBI |
| CY182129 | A/Nicaragua/AGA2-81/2012(H3N2) | 10/8/2012 | NCBI |
| CY182145 | A/Nicaragua/AGA2-84/2012(H3N2) | 18/8/2012 | NCBI |
| CY182153 | A/Nicaragua/AGA2-85/2012(H3N2) | 17/8/2012 | NCBI |
| CY182161 | A/Nicaragua/AGA2-86/2012(H3N2) | 20/8/2012 | NCBI |
| CY182169 | A/Nicaragua/AGA2-88/2012(H3N2) | 30/6/2012 | NCBI |
| CY182177 | A/Nicaragua/AGA2-89/2012(H3N2) | 11/11/2012 | NCBI |
| CY182185 | A/Nicaragua/AGA2-90/2012(H3N2) | 12/11/2012 | NCBI |
| CY182193 | A/Nicaragua/AGA2-91/2012(H3N2) | 15/10/2012 | NCBI |
| CY182201 | A/Nicaragua/AGA2-92/2012(H3N2) | 12/10/2012 | NCBI |
| CY182209 | A/Nicaragua/AGA2-93/2012(H3N2) | 17/10/2012 | NCBI |
| CY182217 | A/Nicaragua/AGA2-94/2012(H3N2) | 17/10/2012 | NCBI |
| CY182225 | A/Nicaragua/AGA2-95/2012(H3N2) | 20/10/2012 | NCBI |
| CY182233 | A/Nicaragua/AGA2-96/2012(H3N2) | 23/10/2012 | NCBI |
| CY182241 | A/Nicaragua/AGA2-97/2012(H3N2) | 22/10/2012 | NCBI |
| CY182249 | A/Nicaragua/AGA2-100/2012(H3N2) | 26/10/2012 | NCBI |
| CY182257 | A/Nicaragua/AGA2-101/2012(H3N2) | 30/10/2012 | NCBI |
| CY182265 | A/Nicaragua/AGA2-102/2012(H3N2) | 30/10/2012 | NCBI |
| CY182273 | A/Nicaragua/AGA2-103/2012(H3N2) | 6/11/2012 | NCBI |

| | | | |
|---|---|---|---|
| CY182281 | A/Nicaragua/AGA2-104/2012(H3N2) | 3/11/2012 | NCBI |
| CY182289 | A/Nicaragua/AGA2-105/2012(H3N2) | 12/10/2012 | NCBI |
| CY182297 | A/Nicaragua/AGA2-106/2012(H3N2) | 12/10/2012 | NCBI |
| CY182305 | A/Nicaragua/AGA2-107/2012(H3N2) | 11/10/2012 | NCBI |
| CY182313 | A/Nicaragua/AGA2-108/2012(H3N2) | 11/10/2012 | NCBI |
| CY182321 | A/Nicaragua/AGA2-109/2012(H3N2) | 19/11/2012 | NCBI |
| CY182329 | A/Nicaragua/AGA2-110/2012(H3N2) | 6/12/2012 | NCBI |
| CY182337 | A/Nicaragua/AGA2-112/2013(H3N2) | 11/2/2013 | NCBI |
| CY182345 | A/Nicaragua/AGA2-113/2013(H3N2) | 17/2/2013 | NCBI |
| CY182353 | A/Nicaragua/AGA2-114/2013(H3N2) | 21/2/2013 | NCBI |
| CY182689 | A/Houston/JMM_61/2012(H3N2) | 9/12/2012 | NCBI |
| CY182697 | A/Houston/JMM_62/2012(H3N2) | 9/12/2012 | NCBI |
| CY182705 | A/Houston/JMM_63/2012(H3N2) | 10/12/2012 | NCBI |
| CY182721 | A/Houston/JMM_65/2012(H3N2) | 12/12/2012 | NCBI |
| CY182729 | A/Houston/JMM_66/2012(H3N2) | 12/12/2012 | NCBI |
| CY182737 | A/Houston/JMM_67/2012(H3N2) | 13/12/2012 | NCBI |
| CY182745 | A/Houston/JMM_68/2012(H3N2) | 13/12/2012 | NCBI |
| CY182753 | A/Houston/JMM_69/2012(H3N2) | 14/12/2012 | NCBI |
| CY182761 | A/Houston/JMM_70/2012(H3N2) | 14/12/2012 | NCBI |
| CY182769 | A/Houston/JMM_71/2012(H3N2) | 15/12/2012 | NCBI |
| CY182777 | A/Houston/JMM_72/2012(H3N2) | 15/12/2012 | NCBI |
| CY182785 | A/Houston/JMM_73/2012(H3N2) | 15/12/2012 | NCBI |
| CY182793 | A/Houston/JMM_74/2012(H3N2) | 15/12/2012 | NCBI |
| CY182801 | A/Houston/JMM_75/2012(H3N2) | 16/12/2012 | NCBI |
| CY182809 | A/Houston/JMM_76/2012(H3N2) | 16/12/2012 | NCBI |
| CY182817 | A/Houston/JMM_77/2012(H3N2) | 16/12/2012 | NCBI |
| CY182825 | A/Houston/JMM_78/2012(H3N2) | 17/12/2012 | NCBI |
| CY182833 | A/Houston/JMM_79/2012(H3N2) | 17/12/2012 | NCBI |
| CY182841 | A/Houston/JMM_80/2012(H3N2) | 18/12/2012 | NCBI |
| CY182849 | A/Houston/JMM_81/2012(H3N2) | 18/12/2012 | NCBI |
| CY182857 | A/Houston/JMM_82/2012(H3N2) | 18/12/2012 | NCBI |
| CY182865 | A/Houston/JMM_84/2012(H3N2) | 19/12/2012 | NCBI |
| CY182873 | A/Houston/JMM_85/2012(H3N2) | 19/12/2012 | NCBI |
| CY182881 | A/Houston/JMM_86/2012(H3N2) | 20/12/2012 | NCBI |
| CY182889 | A/Houston/JMM_87/2012(H3N2) | 20/12/2012 | NCBI |
| CY182897 | A/Houston/JMM_88/2012(H3N2) | 21/12/2012 | NCBI |
| CY182905 | A/Houston/JMM_90/2012(H3N2) | 22/12/2012 | NCBI |

| | | | |
|---|---|---|---|
| CY182913 | A/Houston/JMM_91/2012(H3N2) | 22/12/2012 | NCBI |
| CY182921 | A/Houston/JMM_92/2012(H3N2) | 23/12/2012 | NCBI |
| CY182937 | A/Houston/JMM_94/2012(H3N2) | 28/12/2012 | NCBI |
| CY182945 | A/Houston/JMM_95/2012(H3N2) | 28/12/2012 | NCBI |
| CY182953 | A/Houston/JMM_96/2012(H3N2) | 29/12/2012 | NCBI |
| CY182961 | A/Houston/JMM_97/2012(H3N2) | 29/12/2012 | NCBI |
| CY182969 | A/Houston/JMM_98/2012(H3N2) | 29/12/2012 | NCBI |
| CY182977 | A/Houston/JMM_99/2012(H3N2) | 29/12/2012 | NCBI |
| CY182985 | A/Houston/JMM_101/2013(H3N2) | 1/1/2013 | NCBI |
| CY182993 | A/Houston/JMM_102/2013(H3N2) | 1/1/2013 | NCBI |
| CY183001 | A/Houston/JMM_103/2013(H3N2) | 2/1/2013 | NCBI |
| CY183009 | A/Houston/JMM_104/2013(H3N2) | 2/1/2013 | NCBI |
| CY183017 | A/Houston/JMM_105/2013(H3N2) | 2/1/2013 | NCBI |
| CY183025 | A/Houston/JMM_106/2013(H3N2) | 2/1/2013 | NCBI |
| CY183033 | A/Houston/JMM_107/2013(H3N2) | 3/1/2013 | NCBI |
| CY183041 | A/Houston/JMM_108/2013(H3N2) | 3/1/2013 | NCBI |
| CY183049 | A/Houston/JMM_109/2013(H3N2) | 3/1/2013 | NCBI |
| CY183057 | A/Houston/JMM_110/2013(H3N2) | 3/1/2013 | NCBI |
| CY183065 | A/Houston/JMM_111/2013(H3N2) | 4/1/2013 | NCBI |
| CY183073 | A/Houston/JMM_112/2013(H3N2) | 4/1/2013 | NCBI |
| CY183081 | A/Houston/JMM_113/2013(H3N2) | 4/1/2013 | NCBI |
| CY183089 | A/Houston/JMM_115/2013(H3N2) | 5/1/2013 | NCBI |
| CY183097 | A/Houston/JMM_116/2013(H3N2) | 5/1/2013 | NCBI |
| CY183105 | A/Houston/JMM_117/2013(H3N2) | 5/1/2013 | NCBI |
| CY183113 | A/Houston/JMM_118/2013(H3N2) | 5/1/2013 | NCBI |
| CY183121 | A/Houston/JMM_119/2013(H3N2) | 7/1/2013 | NCBI |
| CY183129 | A/Houston/JMM_120/2013(H3N2) | 8/1/2013 | NCBI |
| CY183137 | A/Houston/JMM_121/2013(H3N2) | 8/1/2013 | NCBI |
| CY183153 | A/Houston/JMM_123/2013(H3N2) | 8/1/2013 | NCBI |
| CY183161 | A/Houston/JMM_124/2013(H3N2) | 8/1/2013 | NCBI |
| CY183169 | A/Houston/JMM_125/2013(H3N2) | 8/1/2013 | NCBI |
| CY183177 | A/Houston/JMM_126/2013(H3N2) | 8/1/2013 | NCBI |
| CY183185 | A/Houston/JMM_127/2013(H3N2) | 9/1/2013 | NCBI |
| CY183193 | A/Houston/JMM_128/2013(H3N2) | 9/1/2013 | NCBI |
| CY183201 | A/Houston/JMM_129/2013(H3N2) | 9/1/2013 | NCBI |
| CY183209 | A/Houston/JMM_130/2013(H3N2) | 9/1/2013 | NCBI |
| CY183225 | A/Houston/JMM_132/2013(H3N2) | 10/1/2013 | NCBI |

| | | | |
|---|---|---|---|
| CY183233 | A/Houston/JMM_134/2013(H3N2) | 11/1/2013 | NCBI |
| CY183241 | A/Houston/JMM_135/2013(H3N2) | 11/1/2013 | NCBI |
| CY183249 | A/Houston/JMM_136/2013(H3N2) | 11/1/2013 | NCBI |
| CY183257 | A/Houston/JMM_137/2013(H3N2) | 11/1/2013 | NCBI |
| CY183265 | A/Houston/JMM_139/2013(H3N2) | 12/1/2013 | NCBI |
| CY183273 | A/Houston/JMM_140/2013(H3N2) | 12/1/2013 | NCBI |
| CY183281 | A/Houston/JMM_141/2013(H3N2) | 12/1/2013 | NCBI |
| CY183289 | A/Houston/JMM_142/2013(H3N2) | 12/1/2013 | NCBI |
| CY183297 | A/Houston/JMM_144/2013(H3N2) | 13/1/2013 | NCBI |
| CY183305 | A/Houston/JMM_145/2013(H3N2) | 14/1/2013 | NCBI |
| CY183313 | A/Houston/JMM_146/2013(H3N2) | 19/1/2013 | NCBI |
| CY183321 | A/Houston/JMM_147/2013(H3N2) | 19/1/2013 | NCBI |
| CY186019 | A/Houston/JMM_149/2013(H3N2) | 22/1/2013 | NCBI |
| CY186027 | A/Houston/JMM_150/2013(H3N2) | 25/1/2013 | NCBI |
| CY186035 | A/Houston/JMM_151/2013(H3N2) | 26/1/2013 | NCBI |
| CY186051 | A/Houston/JMM_153/2013(H3N2) | 22/1/2013 | NCBI |
| CY186059 | A/Houston/JMM_154/2013(H3N2) | 22/1/2013 | NCBI |
| CY186075 | A/Houston/JMM_156/2013(H3N2) | 26/1/2013 | NCBI |
| CY186083 | A/Houston/JMM_157/2013(H3N2) | 30/1/2013 | NCBI |
| CY186091 | A/Houston/JMM_158/2013(H3N2) | 1/2/2013 | NCBI |
| CY186107 | A/Houston/JMM_160/2013(H3N2) | 15/1/2013 | NCBI |
| CY186115 | A/Houston/JMM_161/2013(H3N2) | 23/1/2013 | NCBI |
| CY186123 | A/Houston/JMM_162/2013(H3N2) | 27/1/2013 | NCBI |
| CY186139 | A/Houston/JMM_164/2013(H3N2) | 20/1/2013 | NCBI |
| CY186147 | A/Houston/JMM_166/2013(H3N2) | 20/1/2013 | NCBI |
| CY186155 | A/Houston/JMM_167/2013(H3N2) | 14/1/2013 | NCBI |
| CY186163 | A/Houston/JMM_168/2013(H3N2) | 14/1/2013 | NCBI |
| CY186171 | A/Houston/JMM_169/2013(H3N2) | 30/1/2013 | NCBI |
| CY186179 | A/Houston/JMM_170/2013(H3N2) | 22/1/2013 | NCBI |
| CY186195 | A/Houston/JMM_172/2013(H3N2) | 21/1/2013 | NCBI |
| CY186203 | A/Houston/JMM_174/2013(H3N2) | 20/1/2013 | NCBI |
| CY186211 | A/Houston/JMM_175/2013(H3N2) | 14/1/2013 | NCBI |
| CY186219 | A/Houston/JMM_176/2013(H3N2) | 17/1/2013 | NCBI |
| CY186227 | A/Houston/JMM_177/2013(H3N2) | 15/1/2013 | NCBI |
| CY186235 | A/Houston/JMM_178/2013(H3N2) | 15/1/2013 | NCBI |
| CY186243 | A/Houston/JMM_179/2013(H3N2) | 5/2/2013 | NCBI |
| CY186251 | A/Houston/JMM_180/2013(H3N2) | 8/2/2013 | NCBI |

| | | | |
|---|---|---|---|
| KJ577149 | A/Thailand/VIROAF1/2012(H3N2) | 5/10/2012 | NCBI |
| KJ577157 | A/Thailand/VIROAF2/2012(H3N2) | 30/7/2012 | NCBI |
| KJ577189 | A/Thailand/VIROAF6/2012(H3N2) | 13/9/2012 | NCBI |
| KJ855475 | A/Mexico/TLA2227/2005(H3N2) | 11/9/2005 | NCBI |
| KJ855467 | A/Mexico/JAL25216/2009(H3N2) | 22/5/2009 | NCBI |
| KJ855459 | A/Mexico/JAL25206/2009(H3N2) | 22/5/2009 | NCBI |
| KJ855451 | A/Mexico/DIF2662/2003(H3N2) | 15/12/2003 | NCBI |
| KJ855443 | A/Mexico/MEX2640/2005(H3N2) | 22/11/2005 | NCBI |
| KJ855435 | A/Mexico/DIF2601/2005(H3N2) | 18/11/2005 | NCBI |
| KJ855427 | A/Mexico/QUE2270/2005(H3N2) | 11/8/2005 | NCBI |
| KJ855419 | A/Mexico/DIF2246/2005(H3N2) | 11/9/2005 | NCBI |
| KJ855411 | A/Mexico/DIF2160/2005(H3N2) | 11/1/2005 | NCBI |
| KJ855403 | A/Mexico/DIF2112/2005(H3N2) | 11/1/2005 | NCBI |
| KJ855395 | A/Mexico/NAY2090/2007(H3N2) | 14/4/2007 | NCBI |
| KJ855387 | A/Mexico/DIF940/2003(H3N2) | 23/4/2003 | NCBI |
| KJ855379 | A/Mexico/DIF835/2003(H3N2) | 4/7/2003 | NCBI |
| KJ855371 | A/Mexico/DIF756/2003(H3N2) | 4/2/2003 | NCBI |
| KJ855363 | A/Mexico/DIF29/2006(H3N2) | 16/1/2006 | NCBI |
| KJ855355 | A/Mexico/VER58/2012(H3N2) | 17/1/2012 | NCBI |
| KJ855347 | A/Mexico/VER59/2011(H3N2) | 9/12/2011 | NCBI |
| KJ855339 | A/Mexico/VER60/2011(H3N2) | 24/12/2011 | NCBI |
| KJ855331 | A/Mexico/VER40/2011(H3N2) | 1/12/2011 | NCBI |
| KM064110 | A/Utah/07/2013(H3N2) | 5/7/2013 | NCBI |
| KM064453 | A/Massachusetts/11/2013(H3N2) | 10/7/2013 | NCBI |
| KM064435 | A/Arkansas/13/2013(H3N2) | 15/8/2013 | NCBI |
| KJ942592 | A/Indiana/10/2011(H3N2) | 2011 | NCBI |
| KJ942608 | A/Hawaii/22/2012(H3N2) | 2012 | NCBI |
| KJ942616 | A/Texas/50/2012(H3N2) | 2012 | NCBI |
| KJ942624 | A/Minnesota/11/2010(H3N2) | 2010 | NCBI |
| KC306165 | A/Victoria/361/2011(H3N2) | 2011 | NCBI |
| KJ942680 | A/Victoria/361/2011(H3N2) | 2011 | NCBI |
| KJ942688 | A/Indiana/21/2012(H3N2) | 2012 | NCBI |
| KJ942704 | A/Ohio/02/2012(H3N2) | 2012 | NCBI |
| KJ942792 | A/Brisbane/299/2011(H3N2) | 2011 | NCBI |
| KJ942911 | A/Santiago/p5d2/2012(H3N2) | 22/6/2012 | NCBI |
| KJ942919 | A/Santiago/op5d1/2012(H3N2) | 28/6/2012 | NCBI |
| KJ942935 | A/Santiago/p36d1/2013(H3N2) | 30/8/2013 | NCBI |

| | | | |
|---|---|---|---|
| KJ942943 | A/Santiago/p36d0/2013(H3N2) | 29/8/2013 | NCBI |
| KJ942951 | A/Santiago/p19d2/2012(H3N2) | 6/7/2012 | NCBI |
| KJ942959 | A/Santiago/p37d1/2012(H3N2) | 31/10/2012 | NCBI |
| KJ942975 | A/Santiago/p19d1/2012(H3N2) | 5/7/2012 | NCBI |
| KJ942991 | A/Santiago/p37d0/2012(H3N2) | 30/10/2012 | NCBI |
| KJ943018 | A/Santiago/op3d1/2012(H3N2) | 24/7/2012 | NCBI |
| KJ943026 | A/Santiago/p34d0/2013(H3N2) | 7/8/2013 | NCBI |
| KJ943058 | A/Santiago/p27d0/2013(H3N2) | 12/7/2013 | NCBI |
| KJ943066 | A/Santiago/p19d0/2012(H3N2) | 29/6/2012 | NCBI |
| KJ943074 | A/Santiago/p36d2/2013(H3N2) | 31/8/2013 | NCBI |
| KJ943093 | A/Santiago/p38-2012d1/2012(H3N2) | 27/12/2012 | NCBI |
| KJ943101 | A/Santiago/op20d1/2012(H3N2) | 8/11/2012 | NCBI |
| KJ943109 | A/Santiago/p5d3/2012(H3N2) | 23/6/2012 | NCBI |
| KJ943138 | A/Santiago/p5d0/2012(H3N2) | 20/6/2012 | NCBI |
| KJ943146 | A/Santiago/p5d1/2012(H3N2) | 21/6/2012 | NCBI |
| KJ943154 | A/Santiago/p37d3/2012(H3N2) | 2/11/2012 | NCBI |
| KJ943162 | A/Santiago/p27d3/2013(H3N2) | 15/7/2013 | NCBI |
| KJ943183 | A/Santiago/op7d1/2012(H3N2) | 28/6/2012 | NCBI |
| KJ943191 | A/Santiago/p34d1/2013(H3N2) | 8/8/2013 | NCBI |
| KJ943216 | A/Santiago/p36d5/2013(H3N2) | 2/9/2013 | NCBI |
| KJ943240 | A/Santiago/p38-2012d3/2012(H3N2) | 29/12/2012 | NCBI |
| KJ943248 | A/Santiago/op17d0/2012(H3N2) | 1/6/2012 | NCBI |
| KJ943557 | A/Santiago/p24d2/2012(H3N2) | 21/7/2012 | NCBI |
| KJ943565 | A/Santiago/p2d2/2012(H3N2) | 14/6/2012 | NCBI |
| KJ943573 | A/Santiago/p3d0/2012(H3N2) | 14/6/2012 | NCBI |
| KJ943581 | A/Santiago/p9d3/2012(H3N2) | 25/6/2012 | NCBI |
| KJ943589 | A/Santiago/p9d1/2012(H3N2) | 23/6/2012 | NCBI |
| KJ943597 | A/Santiago/p6d1/2012(H3N2) | 21/6/2012 | NCBI |
| KJ943609 | A/Santiago/p1d2/2012(H3N2) | 6/6/2012 | NCBI |
| KJ943618 | A/Santiago/p24d1/2012(H3N2) | 20/7/2012 | NCBI |
| KJ943626 | A/Santiago/p18d00/2012(H3N2) | 5/7/2012 | NCBI |
| KJ943634 | A/Santiago/p4d3/2012(H3N2) | 20/6/2012 | NCBI |
| KJ943642 | A/Santiago/p9d0/2012(H3N2) | 22/6/2012 | NCBI |
| KJ943664 | A/Santiago/p9d2/2012(H3N2) | 24/6/2012 | NCBI |
| KJ943678 | A/Santiago/p20d0/2012(H3N2) | 7/7/2012 | NCBI |
| KJ943686 | A/Santiago/p1d1/2012(H3N2) | 5/6/2012 | NCBI |
| KJ943699 | A/Santiago/p9d5/2012(H3N2) | 26/6/2012 | NCBI |

| | | | |
|---|---|---|---|
| KJ943712 | A/Santiago/p4d2/2012(H3N2) | 19/6/2012 | NCBI |
| KJ943720 | A/Santiago/p18d0/2012(H3N2) | 6/7/2012 | NCBI |
| KJ943729 | A/Santiago/p4d1/2012(H3N2) | 18/6/2012 | NCBI |
| KJ943737 | A/Santiago/p6d2/2012(H3N2) | 22/6/2012 | NCBI |
| KJ943751 | A/Santiago/p25d2/2012(H3N2) | 21/7/2012 | NCBI |
| KJ943759 | A/Santiago/p6d3/2012(H3N2) | 23/6/2012 | NCBI |
| KJ943767 | A/Santiago/p2d1/2012(H3N2) | 13/6/2012 | NCBI |
| KJ943775 | A/Santiago/p6d5/2012(H3N2) | 25/6/2012 | NCBI |
| KJ943783 | A/Santiago/p4d0/2012(H3N2) | 16/6/2012 | NCBI |
| KM069479 | A/Singapore/S2009.329/2009(H3N2) | 30/4/2009 | NCBI |
| KM069480 | A/Singapore/S2009.332b/2009(H3N2) | 1/5/2009 | NCBI |
| KM069481 | A/Singapore/S2009.359/2009(H3N2) | 11/5/2009 | NCBI |
| KM069482 | A/Singapore/S2009.362b/2009(H3N2) | 12/5/2009 | NCBI |
| KM069483 | A/Singapore/S2009.362c/2009(H3N2) | 12/5/2009 | NCBI |
| KM069484 | A/Singapore/S2009.364c/2009(H3N2) | 13/5/2009 | NCBI |
| KM069485 | A/Singapore/S2009.367a/2009(H3N2) | 14/5/2009 | NCBI |
| KM069486 | A/Singapore/S2009.378a/2009(H3N2) | 18/5/2009 | NCBI |
| KM069488 | A/Singapore/S2009.436/2009(H3N2) | 8/6/2009 | NCBI |
| KM069489 | A/Singapore/C2009.589/2009(H3N2) | 3/8/2009 | NCBI |
| KM069490 | A/Singapore/S2009.655/2009(H3N2) | 27/8/2009 | NCBI |
| KM069491 | A/Singapore/C2009.666/2009(H3N2) | 31/8/2009 | NCBI |
| KM069492 | A/Singapore/C2009.677/2009(H3N2) | 4/9/2009 | NCBI |
| KM069493 | A/Singapore/C2009.693/2009(H3N2) | 10/9/2009 | NCBI |
| KM069494 | A/Singapore/C2009.707/2009(H3N2) | 15/9/2009 | NCBI |
| KM069495 | A/Singapore/C2009.784/2009(H3N2) | 13/10/2009 | NCBI |
| KM069496 | A/Singapore/C2009.803a/2009(H3N2) | 20/10/2009 | NCBI |
| KM069497 | A/Singapore/C2009.934/2009(H3N2) | 7/12/2009 | NCBI |
| KM069498 | A/Singapore/C2010.049/2010(H3N2) | 18/1/2010 | NCBI |
| KM069499 | A/Singapore/C2010.241/2010(H3N2) | 29/3/2010 | NCBI |
| KM069500 | A/Singapore/S2010.304b/2010(H3N2) | 21/4/2010 | NCBI |
| KM069501 | A/Singapore/C2010.307/2010(H3N2) | 22/4/2010 | NCBI |
| KM069502 | A/Singapore/C2010.310/2010(H3N2) | 23/4/2010 | NCBI |
| KM069503 | A/Singapore/H2010.310/2010(H3N2) | 23/4/2010 | NCBI |
| KM069504 | A/Singapore/H2010.323/2010(H3N2) | 28/4/2010 | NCBI |
| KM069505 | A/Singapore/S2010.345b/2010(H3N2) | 6/5/2010 | NCBI |
| KM069506 | A/Singapore/S2010.345c/2010(H3N2) | 6/5/2010 | NCBI |
| KM069507 | A/Singapore/S2010.348/2010(H3N2) | 7/5/2010 | NCBI |

| | | | |
|---|---|---|---|
| KM069508 | A/Singapore/S2010.359a/2010(H3N2) | 11/5/2010 | NCBI |
| KM069509 | A/Singapore/S2010.362b/2010(H3N2) | 12/5/2010 | NCBI |
| KM069511 | A/Singapore/C2010.384/2010(H3N2) | 20/5/2010 | NCBI |
| KM069512 | A/Singapore/H2010.400/2010(H3N2) | 26/5/2010 | NCBI |
| KM069513 | A/Singapore/C2010.416/2010(H3N2) | 1/6/2010 | NCBI |
| KM069514 | A/Singapore/S2010.416a/2010(H3N2) | 1/6/2010 | NCBI |
| KM069516 | A/Singapore/H2010.422/2010(H3N2) | 3/6/2010 | NCBI |
| KM069517 | A/Singapore/C2010.441/2010(H3N2) | 10/6/2010 | NCBI |
| KM069518 | A/Singapore/C2010.471b/2010(H3N2) | 21/6/2010 | NCBI |
| KM069521 | A/Singapore/H2010.570/2010(H3N2) | 27/7/2010 | NCBI |
| KM069522 | A/Singapore/S2010.570/2010(H3N2) | 27/7/2010 | NCBI |
| KM069523 | A/Singapore/H2010.586/2010(H3N2) | 2/8/2010 | NCBI |
| KM069524 | A/Singapore/S2010.611a/2010(H3N2) | 11/8/2010 | NCBI |
| KM069526 | A/Singapore/S2010.614/2010(H3N2) | 12/8/2010 | NCBI |
| KM069527 | A/Singapore/C2010.625/2010(H3N2) | 16/8/2010 | NCBI |
| KM069528 | A/Singapore/S2010.627a/2010(H3N2) | 17/8/2010 | NCBI |
| KM069529 | A/Singapore/S2010.627b/2010(H3N2) | 17/8/2010 | NCBI |
| KM069530 | A/Singapore/S2010.655b/2010(H3N2) | 27/8/2010 | NCBI |
| KM069531 | A/Singapore/C2010.663/2010(H3N2) | 30/8/2010 | NCBI |
| KM069532 | A/Singapore/S2010.682/2010(H3N2) | 6/9/2010 | NCBI |
| KM069533 | A/Singapore/C2010.721/2010(H3N2) | 20/9/2010 | NCBI |
| KM069534 | A/Singapore/C2010.742/2010(H3N2) | 28/9/2010 | NCBI |
| KM069536 | A/Singapore/C2010.784b/2010(H3N2) | 13/10/2010 | NCBI |
| KM069539 | A/Singapore/S2011.090/2011(H3N2) | 2/2/2011 | NCBI |
| KM069540 | A/Singapore/H2011.521/2011(H3N2) | 9/7/2011 | NCBI |
| KM069541 | A/Singapore/S2011.575/2011(H3N2) | 29/7/2011 | NCBI |
| KM069542 | A/Singapore/S2011.611b/2011(H3N2) | 11/8/2011 | NCBI |
| KM069543 | A/Singapore/H2011.625/2011(H3N2) | 16/8/2011 | NCBI |
| KM069544 | A/Singapore/S2011.701a/2011(H3N2) | 13/9/2011 | NCBI |
| KM069545 | A/Singapore/S2011.737/2011(H3N2) | 26/9/2011 | NCBI |
| KM069546 | A/Singapore/S2011.740/2011(H3N2) | 27/9/2011 | NCBI |
| KM069547 | A/Singapore/H2011.748/2011(H3N2) | 30/9/2011 | NCBI |
| KM069548 | A/Singapore/H2011.795/2011(H3N2) | 17/10/2011 | NCBI |
| KM069549 | A/Singapore/H2011.814/2011(H3N2) | 24/10/2011 | NCBI |
| KM069550 | A/Singapore/C2011.860/2011(H3N2) | 10/11/2011 | NCBI |
| KM069551 | A/Singapore/C2011.863/2011(H3N2) | 11/11/2011 | NCBI |
| KM069552 | A/Singapore/H2011.871/2011(H3N2) | 14/11/2011 | NCBI |

| | | | |
|---|---|---|---|
| KM069553 | A/Singapore/H2011.904/2011(H3N2) | 26/11/2011 | NCBI |
| KM069554 | A/Singapore/H2011.907/2011(H3N2) | 27/11/2011 | NCBI |
| KM069555 | A/Singapore/H2011.923/2011(H3N2) | 3/12/2011 | NCBI |
| KM069556 | A/Singapore/C2011.992/2011(H3N2) | 28/12/2011 | NCBI |
| KM069557 | A/Singapore/C2012.011/2012(H3N2) | 4/1/2012 | NCBI |
| KM069558 | A/Singapore/S2012.014/2012(H3N2) | 5/1/2012 | NCBI |
| KM069559 | A/Singapore/S2012.181/2012(H3N2) | 6/3/2012 | NCBI |
| KM069560 | A/Singapore/C2012.197/2012(H3N2) | 12/3/2012 | NCBI |
| KM069561 | A/Singapore/C2012.208/2012(H3N2) | 16/3/2012 | NCBI |
| KM069562 | A/Singapore/C2012.247/2012(H3N2) | 30/3/2012 | NCBI |
| KM069563 | A/Singapore/C2012.274/2012(H3N2) | 9/4/2012 | NCBI |
| KM069564 | A/Singapore/H2012.301/2012(H3N2) | 19/4/2012 | NCBI |
| KM069565 | A/Singapore/H2012.375/2012(H3N2) | 16/5/2012 | NCBI |
| KM069566 | A/Singapore/C2012.389/2012(H3N2) | 21/5/2012 | NCBI |
| KM069567 | A/Singapore/S2012.389/2012(H3N2) | 21/5/2012 | NCBI |
| KM069568 | A/Singapore/H2012.490/2012(H3N2) | 27/6/2012 | NCBI |
| KM069569 | A/Singapore/H2012.562a/2012(H3N2) | 23/7/2012 | NCBI |
| KM069570 | A/Singapore/H2012.567/2012(H3N2) | 25/7/2012 | NCBI |
| KM069571 | A/Singapore/C2012.669/2012(H3N2) | 1/9/2012 | NCBI |
| KM069572 | A/Singapore/H2012.705/2012(H3N2) | 14/9/2012 | NCBI |
| KM069574 | A/Singapore/C2012.801/2012(H3N2) | 19/10/2012 | NCBI |
| KM069575 | A/Singapore/H2012.828/2012(H3N2) | 29/10/2012 | NCBI |
| KM069576 | A/Singapore/S2012.833/2012(H3N2) | 31/10/2012 | NCBI |
| KM069577 | A/Singapore/H2012.855/2012(H3N2) | 8/11/2012 | NCBI |
| KM069578 | A/Singapore/H2012.923/2012(H3N2) | 3/12/2012 | NCBI |
| KM069579 | A/Singapore/H2012.934/2012(H3N2) | 7/12/2012 | NCBI |
| KM069580 | A/Singapore/H2012.956/2012(H3N2) | 15/12/2012 | NCBI |
| KM069583 | A/Singapore/H2013.384/2013(H3N2) | 20/5/2013 | NCBI |
| KM069584 | A/Singapore/H2013.411/2013(H3N2) | 30/5/2013 | NCBI |
| KM069585 | A/Singapore/H2013.422a/2013(H3N2) | 3/6/2013 | NCBI |
| KM069586 | A/Singapore/H2013.422b/2013(H3N2) | 3/6/2013 | NCBI |
| KM069587 | A/Singapore/H2013.422c/2013(H3N2) | 3/6/2013 | NCBI |
| KM069589 | A/Singapore/H2013.430a/2013(H3N2) | 6/6/2013 | NCBI |
| KM069590 | A/Singapore/H2013.430b/2013(H3N2) | 6/6/2013 | NCBI |
| KM069591 | A/Singapore/H2013.447/2013(H3N2) | 12/6/2013 | NCBI |
| KM069592 | A/Singapore/H2013.463/2013(H3N2) | 18/6/2013 | NCBI |
| KM069593 | A/Singapore/H2013.529/2013(H3N2) | 12/7/2013 | NCBI |

| | | | |
|---|---|---|---|
| KM069594 | A/Singapore/H2013.545/2013(H3N2) | 18/7/2013 | NCBI |
| KM069595 | A/Singapore/H2013.696/2013(H3N2) | 11/9/2013 | NCBI |
| KM069596 | A/Singapore/H2013.718a/2013(H3N2) | 19/9/2013 | NCBI |
| KM069598 | A/Singapore/H2013.721c/2013(H3N2) | 20/9/2013 | NCBI |
| KM069599 | A/Singapore/H2013.721e/2013(H3N2) | 20/9/2013 | NCBI |
| KM069600 | A/Singapore/H2013.751/2013(H3N2) | 1/10/2013 | NCBI |
| KM069601 | A/Singapore/H2013.778/2013(H3N2) | 11/10/2013 | NCBI |
| KF612194 | A/SantaCruz/2821/2012(H3N2) | 15/6/2012 | NCBI |
| KF612193 | A/SantaCruz/174/2011(H3N2) | 5/9/2011 | NCBI |
| KF612192 | A/SantaCruz/1391/2012(H3N2) | 7/8/2012 | NCBI |
| CY187873 | A/Nicaragua/4430_02/2013(H3N2) | 16/7/2013 | NCBI |
| CY187881 | A/Nicaragua/6596_07/2013(H3N2) | 2/8/2013 | NCBI |
| CY187889 | A/Nicaragua/7477_04/2013(H3N2) | 13/8/2013 | NCBI |
| CY187905 | A/Nicaragua/3442_07/2013(H3N2) | 7/8/2013 | NCBI |
| CY187913 | A/Nicaragua/6676_09/2013(H3N2) | 5/7/2013 | NCBI |
| CY187921 | A/Nicaragua/6201_05/2013(H3N2) | 31/7/2013 | NCBI |
| CY187929 | A/Nicaragua/7525_05/2013(H3N2) | 8/7/2013 | NCBI |
| CY187937 | A/Nicaragua/5300_01/2013(H3N2) | 8/7/2013 | NCBI |
| CY187945 | A/Nicaragua/6426_03/2013(H3N2) | 28/7/2013 | NCBI |
| CY187953 | A/Nicaragua/7699_01/2013(H3N2) | 6/8/2013 | NCBI |
| CY187961 | A/Nicaragua/6909_03/2013(H3N2) | 21/7/2013 | NCBI |
| CY187969 | A/Nicaragua/6965_03/2013(H3N2) | 12/7/2013 | NCBI |
| CY187977 | A/Nicaragua/6496_12/2013(H3N2) | 10/7/2013 | NCBI |
| CY187985 | A/Nicaragua/6823_15/2013(H3N2) | 19/7/2013 | NCBI |
| CY187993 | A/Nicaragua/7622_01/2013(H3N2) | 22/8/2013 | NCBI |
| CY188017 | A/Nicaragua/4871_07/2013(H3N2) | 3/8/2013 | NCBI |
| CY188025 | A/Nicaragua/6958_04/2013(H3N2) | 25/7/2013 | NCBI |
| CY188033 | A/Nicaragua/6889_07/2013(H3N2) | 12/7/2013 | NCBI |
| CY188041 | A/Nicaragua/6104_06/2013(H3N2) | 20/7/2013 | NCBI |
| CY188057 | A/Nicaragua/5805_08/2013(H3N2) | 21/7/2013 | NCBI |
| CY188081 | A/Nicaragua/4860_12/2013(H3N2) | 15/10/2013 | NCBI |
| CY188089 | A/Nicaragua/5349_05/2013(H3N2) | 27/7/2013 | NCBI |
| CY188097 | A/Nicaragua/7228_01/2013(H3N2) | 18/7/2013 | NCBI |
| CY188113 | A/Nicaragua/6118_03/2013(H3N2) | 26/7/2013 | NCBI |
| CY188121 | A/Nicaragua/4236_02/2013(H3N2) | 31/7/2013 | NCBI |
| CY188129 | A/Nicaragua/6890_09/2013(H3N2) | 6/8/2013 | NCBI |
| CY188137 | A/Nicaragua/6480_07/2013(H3N2) | 18/7/2013 | NCBI |

| | | | |
|---|---|---|---|
| CY188145 | A/Nicaragua/5026_03/2013(H3N2) | 10/10/2013 | NCBI |
| CY188177 | A/Nicaragua/7607_03/2013(H3N2) | 5/7/2013 | NCBI |
| CY188193 | A/Nicaragua/6228_04/2013(H3N2) | 14/7/2013 | NCBI |
| CY188201 | A/Nicaragua/6414_07/2013(H3N2) | 2/8/2013 | NCBI |
| CY188209 | A/Nicaragua/6966_06/2013(H3N2) | 28/7/2013 | NCBI |
| CY188217 | A/Nicaragua/6322_04/2013(H3N2) | 26/7/2013 | NCBI |
| CY188225 | A/Nicaragua/7628_01/2013(H3N2) | 12/8/2013 | NCBI |
| CY188233 | A/Nicaragua/7828_01/2013(H3N2) | 20/9/2013 | NCBI |
| CY188241 | A/Nicaragua/7568_04/2013(H3N2) | 20/7/2013 | NCBI |
| CY188249 | A/Nicaragua/6163_03/2013(H3N2) | 9/8/2013 | NCBI |
| CY188265 | A/Nicaragua/7660_03/2013(H3N2) | 28/8/2013 | NCBI |
| CY188273 | A/Nicaragua/7655_01/2013(H3N2) | 29/7/2013 | NCBI |
| CY188297 | A/Nicaragua/3360_03/2013(H3N2) | 8/8/2013 | NCBI |
| CY188305 | A/Nicaragua/6397_14/2013(H3N2) | 13/7/2013 | NCBI |
| CY188321 | A/Nicaragua/6971_03/2013(H3N2) | 25/7/2013 | NCBI |
| CY188329 | A/Nicaragua/7637_03/2013(H3N2) | 28/7/2013 | NCBI |
| CY188345 | A/Nicaragua/5605_09/2013(H3N2) | 23/7/2013 | NCBI |
| CY188369 | A/Nicaragua/6840_02/2013(H3N2) | 29/6/2013 | NCBI |
| CY188377 | A/Nicaragua/6249_03/2013(H3N2) | 24/6/2013 | NCBI |
| CY188401 | A/Nicaragua/6854_05/2013(H3N2) | 4/7/2013 | NCBI |
| CY188433 | A/Nicaragua/6162_03/2013(H3N2) | 8/3/2013 | NCBI |
| CY188457 | A/Nicaragua/5968_01/2013(H3N2) | 30/6/2013 | NCBI |
| CY188489 | A/Nicaragua/6898_05/2013(H3N2) | 30/6/2013 | NCBI |
| CY188497 | A/Nicaragua/6962_04/2013(H3N2) | 4/6/2013 | NCBI |
| CY188521 | A/Nicaragua/6952_06/2013(H3N2) | 2/7/2013 | NCBI |
| CY188545 | A/Nicaragua/7541_02/2013(H3N2) | 25/6/2013 | NCBI |
| CY188553 | A/Nicaragua/6865_15/2013(H3N2) | 2/7/2013 | NCBI |
| CY188577 | A/Nicaragua/6678_01/2013(H3N2) | 4/7/2013 | NCBI |
| CY188585 | A/Nicaragua/6180_11/2013(H3N2) | 28/6/2013 | NCBI |
| CY188593 | A/Nicaragua/7482_04/2013(H3N2) | 1/7/2013 | NCBI |
| CY188601 | A/Nicaragua/726_01/2007(H3N2) | 27/6/2007 | NCBI |
| CY188609 | A/Nicaragua/5674_05/2010(H3N2) | 5/6/2010 | NCBI |
| CY188617 | A/Nicaragua/5969_01/2010(H3N2) | 7/6/2010 | NCBI |
| CY188633 | A/Nicaragua/7462_02/2012(H3N2) | 15/9/2012 | NCBI |
| CY188641 | A/Nicaragua/4855_04/2013(H3N2) | 5/7/2013 | NCBI |
| CY188649 | A/Nicaragua/5898_06/2013(H3N2) | 8/7/2013 | NCBI |
| CY188657 | A/Nicaragua/421_02/2013(H3N2) | 21/7/2013 | NCBI |

| | | | |
|---|---|---|---|
| CY188665 | A/Nicaragua/5754_07/2013(H3N2) | 22/7/2013 | NCBI |
| CY188681 | A/Nicaragua/6464_25/2013(H3N2) | 27/6/2013 | NCBI |
| KF533055 | A/Singapore/139N/2007(H3N2) | 10/8/2007 | NCBI |
| KF533050 | A/Singapore/25Z/2007(H3N2) | 10/5/2007 | NCBI |
| KF533054 | A/Singapore/87Z/2007(H3N2) | 18/5/2007 | NCBI |
| KF533053 | A/Singapore/78L/2007(H3N2) | 18/5/2007 | NCBI |
| KF533052 | A/Singapore/68Q/2007(H3N2) | 16/5/2007 | NCBI |
| KF533051 | A/Singapore/64K/2007(H3N2) | 16/5/2007 | NCBI |
| KP223135 | A/Singapore/238T/2007(H3N2) | 17/9/2007 | NCBI |
| KP223134 | A/Singapore/105L/2007(H3N2) | 25/5/2007 | NCBI |
| KP223133 | A/Singapore/103C/2007(H3N2) | 25/5/2007 | NCBI |
| KP223132 | A/Singapore/53C/2007(H3N2) | 14/5/2007 | NCBI |
| CY189543 | A/Nicaragua/30014_01_TR2/2012(H3N2) | 30/8/2012 | NCBI |
| CY189551 | A/Nicaragua/930_01_TR1/2012(H3N2) | 28/8/2012 | NCBI |
| CY189567 | A/Nicaragua/5892_01_TR1/2013(H3N2) | 2/7/2013 | NCBI |
| CY189575 | A/Nicaragua/30143_01_tr3/2013(H3N2) | 5/7/2013 | NCBI |
| CY189583 | A/Nicaragua/7545_01_TR3/2013(H3N2) | 5/7/2013 | NCBI |
| CY189591 | A/Nicaragua/30185_01_TR1/2013(H3N2) | 5/7/2013 | NCBI |
| CY189599 | A/Nicaragua/30190_01_TR1/2013(H3N2) | 8/7/2013 | NCBI |
| CY189607 | A/Nicaragua/30193_01_TR1/2013(H3N2) | 8/7/2013 | NCBI |
| CY189615 | A/Nicaragua/6019_01_TR1/2013(H3N2) | 8/7/2013 | NCBI |
| CY189623 | A/Nicaragua/30208_01_tr1/2013(H3N2) | 9/7/2013 | NCBI |
| CY189631 | A/Nicaragua/3026_01_TR1/2013(H3N2) | 9/7/2013 | NCBI |
| CY189639 | A/Nicaragua/5465_01_TR3/2013(H3N2) | 10/7/2013 | NCBI |
| CY189655 | A/Nicaragua/30184_01_TR2/2013(H3N2) | 10/7/2013 | NCBI |
| CY189687 | A/Nicaragua/6777_01_TR1/2013(H3N2) | 14/7/2013 | NCBI |
| CY189695 | A/Nicaragua/7693_01_TR3/2013(H3N2) | 15/7/2013 | NCBI |
| CY189703 | A/Nicaragua/7457_02/2013(H3N2) | 9/7/2013 | NCBI |
| CY189751 | A/Nicaragua/30071_01_TR1/2013(H3N2) | 14/6/2013 | NCBI |
| CY189767 | A/Nicaragua/30072_01_TR1/2013(H3N2) | 14/6/2013 | NCBI |
| LC032873 | A/Kyoto/13K025/2014(H3N2) | 5/2/2014 | NCBI |
| LC032889 | A/Kyoto/13K054/2014(H3N2) | 13/2/2014 | NCBI |
| LC032993 | A/Hokkaido/13H009/2014(H3N2) | 14/1/2014 | NCBI |
| LC033001 | A/Hokkaido/13H011/2014(H3N2) | 22/1/2014 | NCBI |
| LC033057 | A/Niigata/13NU003/2014(H3N2) | 30/1/2014 | NCBI |
| LC033113 | A/Niigata/13F173/2014(H3N2) | 17/2/2014 | NCBI |
| LC033121 | A/Niigata/13F071/2014(H3N2) | 5/2/2014 | NCBI |

| | | | |
|---|---|---|---|
| LC033129 | A/Niigata/13F024/2014(H3N2) | 31/1/2014 | NCBI |
| LC033137 | A/Niigata/13F335/2014(H3N2) | 3/3/2014 | NCBI |
| LC033145 | A/Niigata/13F416/2014(H3N2) | 10/3/2014 | NCBI |
| LC033305 | A/Nagasaki/13N004/2014(H3N2) | 2/1/2014 | NCBI |
| LC033313 | A/Nagasaki/13N007/2014(H3N2) | 13/1/2014 | NCBI |
| LC033321 | A/Nagasaki/13N020/2014(H3N2) | 23/1/2014 | NCBI |
| LC033329 | A/Nagasaki/13N046/2014(H3N2) | 30/1/2014 | NCBI |
| LC033337 | A/Nagasaki/13N049/2014(H3N2) | 3/2/2014 | NCBI |
| KP765772 | A/HuNan/01/2014(H3N2) | 1/1/2014 | NCBI |
| KJ609214 | A/Brisbane/10/2007(H3N2) | 6/2/2007 | NCBI |
| KJ609206 | A/Perth/16/2009(H3N2) | 7/4/2009 | NCBI |
| KR057501 | A/Wisconsin/27/2014(H3N2) | 5/8/2014 | NCBI |
| KR057515 | A/Maryland/18/2014(H3N2) | 2/5/2014 | NCBI |
| KR057523 | A/Maryland/19/2014(H3N2) | 1/5/2014 | NCBI |
| KR057531 | A/Texas/34/2014(H3N2) | 10/6/2014 | NCBI |
| KR057542 | A/Hawaii/44/2014(H3N2) | 18/7/2014 | NCBI |
| KR057550 | A/Hawaii/46/2014(H3N2) | 20/7/2014 | NCBI |
| KR057558 | A/Hawaii/47/2014(H3N2) | 18/7/2014 | NCBI |
| KR057566 | A/Hawaii/45/2014(H3N2) | 26/7/2014 | NCBI |
| KR057592 | A/Hawaii/35/2014(H3N2) | 10/8/2014 | NCBI |
| KR057608 | A/Hawaii/36/2014(H3N2) | 8/8/2014 | NCBI |
| KR057616 | A/Hawaii/37/2014(H3N2) | 13/8/2014 | NCBI |
| KR057624 | A/Hawaii/38/2014(H3N2) | 11/8/2014 | NCBI |
| KR057632 | A/Hawaii/39/2014(H3N2) | 13/8/2014 | NCBI |
| KR057643 | A/Michigan/08/2014(H3N2) | 14/8/2014 | NCBI |
| KR057651 | A/Texas/39/2014(H3N2) | 20/8/2014 | NCBI |
| KR057659 | A/NewHampshire/18/2014(H3N2) | 28/8/2014 | NCBI |
| KR057670 | A/California/15/2014(H3N2) | 25/8/2014 | NCBI |
| KR057678 | A/California/16/2014(H3N2) | 25/8/2014 | NCBI |
| KR057686 | A/California/17/2014(H3N2) | 11/8/2014 | NCBI |
| KR057694 | A/California/18/2014(H3N2) | 8/8/2014 | NCBI |
| CY068369 | A/California/VRDL266/2009(H3N2) | 30/4/2009 | NCBI |
| KR137646 | A/Bethesda/NIH12/2011(H3N2) | 15/2/2011 | NCBI |
| KJ848674 | A/Thailand/VIROAF1CS/2012(H3N2) | 5/10/2012 | NCBI |
| KM438082 | A/Quebec/26-220405/2005(H3N2) | 22/4/2005 | NCBI |
| KM438087 | A/Quebec/26-281106/2006(H3N2) | 28/11/2006 | NCBI |
| KM438088 | A/Quebec/26-191206/2006(H3N2) | 19/12/2006 | NCBI |

| | | | |
|---|---|---|---|
| KM438089 | A/Quebec/26-151105/2005(H3N2) | 15/11/2005 | NCBI |
| KM438091 | A/Quebec/26-031205/2005(H3N2) | 3/12/2005 | NCBI |
| KM438092 | A/Quebec/26-310106/2006(H3N2) | 31/1/2006 | NCBI |
| KM438093 | A/Quebec/26-270106/2006(H3N2) | 27/1/2006 | NCBI |
| KT836001 | A/SouthDakota/23/2015(H3N2) | 29/1/2015 | NCBI |
| KT835727 | A/Maryland/02/2015(H3N2) | 24/1/2015 | NCBI |
| KT835729 | A/Nebraska/05/2015(H3N2) | 31/1/2015 | NCBI |
| KT835967 | A/California/19/2015(H3N2) | 27/1/2015 | NCBI |
| KT835726 | A/Wisconsin/30/2015(H3N2) | 26/1/2015 | NCBI |
| KT835887 | A/Iowa/12/2015(H3N2) | 31/1/2015 | NCBI |
| KT835784 | A/Kansas/02/2015(H3N2) | 30/1/2015 | NCBI |
| KT835793 | A/Kansas/09/2015(H3N2) | 30/1/2015 | NCBI |
| KT835710 | A/SouthDakota/25/2015(H3N2) | 29/1/2015 | NCBI |
| KT835840 | A/Florida/20/2015(H3N2) | 30/1/2015 | NCBI |
| KT835832 | A/WestVirginia/02/2015(H3N2) | 28/1/2015 | NCBI |
| KT835718 | A/Missouri/05/2015(H3N2) | 28/1/2015 | NCBI |
| KT835723 | A/Tennessee/10/2015(H3N2) | 27/1/2015 | NCBI |
| KT836350 | A/Wisconsin/05/2015(H3N2) | 27/1/2015 | NCBI |
| KT835968 | A/Washington/04/2015(H3N2) | 30/1/2015 | NCBI |
| KT835734 | A/Colorado/09/2015(H3N2) | 22/1/2015 | NCBI |
| KT836011 | A/Iowa/17/2015(H3N2) | 29/1/2015 | NCBI |
| KT836106 | A/Delaware/05/2015(H3N2) | 30/1/2015 | NCBI |
| KT836364 | A/Maryland/01/2015(H3N2) | 22/1/2015 | NCBI |
| KT835742 | A/Montana/09/2015(H3N2) | 23/1/2015 | NCBI |
| KT836052 | A/Alabama/02/2015(H3N2) | 26/1/2015 | NCBI |
| KT835789 | A/NewJersey/02/2015(H3N2) | 26/1/2015 | NCBI |
| KT835753 | A/Wyoming/14/2015(H3N2) | 22/1/2015 | NCBI |
| KT835917 | A/Wisconsin/27/2015(H3N2) | 26/1/2015 | NCBI |
| KT835761 | A/Nevada/32/2015(H3N2) | 23/1/2015 | NCBI |
| KT835769 | A/Nevada/33/2015(H3N2) | 24/1/2015 | NCBI |
| KT835777 | A/Oklahoma/15/2015(H3N2) | 30/1/2015 | NCBI |
| KT836358 | A/NewHampshire/07/2015(H3N2) | 28/1/2015 | NCBI |
| KT835948 | A/Washington/30/2015(H3N2) | 30/1/2015 | NCBI |
| KT835935 | A/NewHampshire/06/2015(H3N2) | 29/1/2015 | NCBI |
| KT835788 | A/Nevada/23/2015(H3N2) | 26/1/2015 | NCBI |
| KT835922 | A/Arizona/01/2015(H3N2) | 30/1/2015 | NCBI |
| KT835797 | A/Nevada/12/2015(H3N2) | 30/1/2015 | NCBI |

| | | | |
|---|---|---|---|
| KT835801 | A/Indiana/06/2015(H3N2) | 21/1/2015 | NCBI |
| KT835821 | A/Delaware/06/2015(H3N2) | 28/1/2015 | NCBI |
| KT835995 | A/Missouri/12/2015(H3N2) | 31/1/2015 | NCBI |
| KT835818 | A/SouthDakota/22/2015(H3N2) | 29/1/2015 | NCBI |
| KT835972 | A/Delaware/09/2015(H3N2) | 24/1/2015 | NCBI |
| KT835830 | A/Utah/06/2015(H3N2) | 29/1/2015 | NCBI |
| KT836043 | A/Idaho/05/2015(H3N2) | 28/1/2015 | NCBI |
| KT835862 | A/Washington/29/2015(H3N2) | 28/1/2015 | NCBI |
| KT835929 | A/Washington/01/2015(H3N2) | 21/1/2015 | NCBI |
| KT835946 | A/Massachusetts/03/2015(H3N2) | 20/1/2015 | NCBI |
| KT835899 | A/NorthCarolina/02/2015(H3N2) | 28/1/2015 | NCBI |
| KT835859 | A/Arkansas/04/2015(H3N2) | 21/1/2015 | NCBI |
| KT836317 | A/Texas/31/2015(H3N2) | 27/1/2015 | NCBI |
| KT836206 | A/Massachusetts/02/2015(H3N2) | 20/1/2015 | NCBI |
| KT835907 | A/Alabama/04/2015(H3N2) | 22/1/2015 | NCBI |
| KT835927 | A/Washington/18/2015(H3N2) | 21/1/2015 | NCBI |
| KT835975 | A/NewYork/03/2015(H3N2) | 23/1/2015 | NCBI |
| KT836144 | A/Texas/30/2015(H3N2) | 26/1/2015 | NCBI |
| KT835947 | A/Florida/17/2015(H3N2) | 26/1/2015 | NCBI |
| KT835914 | A/Pennsylvania/04/2015(H3N2) | 27/1/2015 | NCBI |
| KT836135 | A/NewYork/12/2015(H3N2) | 22/1/2015 | NCBI |
| KT836204 | A/NewYork/04/2015(H3N2) | 22/1/2015 | NCBI |
| KT836326 | A/Texas/27/2015(H3N2) | 28/1/2015 | NCBI |
| KT836090 | A/Wisconsin/19/2015(H3N2) | 22/1/2015 | NCBI |
| KT835952 | A/Iowa/13/2015(H3N2) | 30/1/2015 | NCBI |
| KT836050 | A/NewJersey/12/2015(H3N2) | 29/1/2015 | NCBI |
| KT836066 | A/NewYork/18/2015(H3N2) | 23/1/2015 | NCBI |
| KT836045 | A/California/20/2015(H3N2) | 20/1/2015 | NCBI |
| KT836064 | A/WestVirginia/07/2015(H3N2) | 25/1/2015 | NCBI |
| KT835979 | A/Massachusetts/12/2015(H3N2) | 26/1/2015 | NCBI |
| KT835987 | A/Maine/24/2015(H3N2) | 29/1/2015 | NCBI |
| KT835997 | A/NorthCarolina/25/2015(H3N2) | 25/1/2015 | NCBI |
| KT836034 | A/NorthCarolina/24/2015(H3N2) | 27/1/2015 | NCBI |
| KT836059 | A/Texas/56/2015(H3N2) | 31/1/2015 | NCBI |
| KT836216 | A/NewMexico/07/2015(H3N2) | 20/1/2015 | NCBI |
| KT836288 | A/Florida/21/2015(H3N2) | 21/1/2015 | NCBI |
| KT836104 | A/Washington/06/2015(H3N2) | 27/1/2015 | NCBI |

| | | | |
|---|---|---|---|
| KT836091 | A/WestVirginia/06/2015(H3N2) | 22/1/2015 | NCBI |
| KT836273 | A/Texas/24/2015(H3N2) | 21/1/2015 | NCBI |
| KT836179 | A/Texas/32/2015(H3N2) | 26/1/2015 | NCBI |
| KT836321 | A/NewJersey/09/2015(H3N2) | 26/1/2015 | NCBI |
| KT836094 | A/Kentucky/04/2015(H3N2) | 22/1/2015 | NCBI |
| KT836178 | A/Florida/24/2015(H3N2) | 20/1/2015 | NCBI |
| KT836333 | A/Idaho/04/2015(H3N2) | 26/1/2015 | NCBI |
| KT836089 | A/Nevada/15/2015(H3N2) | 23/1/2015 | NCBI |
| KT836319 | A/Missouri/09/2015(H3N2) | 24/1/2015 | NCBI |
| KT836218 | A/Texas/34/2015(H3N2) | 26/1/2015 | NCBI |
| KT836271 | A/Ohio/03/2015(H3N2) | 21/1/2015 | NCBI |
| KT836145 | A/Arkansas/05/2015(H3N2) | 20/1/2015 | NCBI |
| KT836134 | A/Texas/29/2015(H3N2) | 24/1/2015 | NCBI |
| KT836376 | A/NewYork/02/2015(H3N2) | 21/1/2015 | NCBI |
| KT836308 | A/Texas/33/2015(H3N2) | 27/1/2015 | NCBI |
| KT836187 | A/NewHampshire/05/2015(H3N2) | 22/1/2015 | NCBI |
| KT836258 | A/NorthCarolina/03/2015(H3N2) | 20/1/2015 | NCBI |
| KT836147 | A/Texas/23/2015(H3N2) | 21/1/2015 | NCBI |
| KT836192 | A/Texas/28/2015(H3N2) | 21/1/2015 | NCBI |
| KT836155 | A/NewJersey/10/2015(H3N2) | 26/1/2015 | NCBI |
| KT836399 | A/Virginia/10/2015(H3N2) | 20/1/2015 | NCBI |
| KT836386 | A/Missouri/11/2015(H3N2) | 24/1/2015 | NCBI |
| KT836940 | A/Maryland/26/2014(H3N2) | 16/9/2014 | NCBI |
| KT836943 | A/Florida/35/2014(H3N2) | 23/9/2014 | NCBI |
| KT836949 | A/Texas/40/2014(H3N2) | 22/9/2014 | NCBI |
| KT837305 | A/NewHampshire/19/2014(H3N2) | 28/9/2014 | NCBI |
| KT837123 | A/Alaska/40/2014(H3N2) | 8/9/2014 | NCBI |
| KT837036 | A/Minnesota/22/2014(H3N2) | 20/9/2014 | NCBI |
| KT837037 | A/Pennsylvania/40/2014(H3N2) | 13/9/2014 | NCBI |
| KT837031 | A/Michigan/11/2014(H3N2) | 13/9/2014 | NCBI |
| KT837011 | A/Michigan/15/2014(H3N2) | 24/9/2014 | NCBI |
| KT836991 | A/Michigan/12/2014(H3N2) | 16/9/2014 | NCBI |
| KT836982 | A/Minnesota/21/2014(H3N2) | 17/9/2014 | NCBI |
| KT836988 | A/Michigan/13/2014(H3N2) | 18/9/2014 | NCBI |
| KT836998 | A/Maryland/27/2014(H3N2) | 17/9/2014 | NCBI |
| KT836992 | A/Michigan/14/2014(H3N2) | 19/9/2014 | NCBI |
| KT837042 | A/NewMexico/27/2014(H3N2) | 18/9/2014 | NCBI |

| | | | |
|---|---|---|---|
| KT837091 | A/Wisconsin/26/2014(H3N2) | 25/9/2014 | NCBI |
| KT837064 | A/Florida/34/2014(H3N2) | 30/9/2014 | NCBI |
| KT837083 | A/Pennsylvania/45/2014(H3N2) | 25/9/2014 | NCBI |
| KT837059 | A/Pennsylvania/44/2014(H3N2) | 27/9/2014 | NCBI |
| KT837060 | A/NorthCarolina/20/2014(H3N2) | 8/9/2014 | NCBI |
| KT837066 | A/Wisconsin/25/2014(H3N2) | 25/9/2014 | NCBI |
| KT837099 | A/Michigan/09/2014(H3N2) | 1/9/2014 | NCBI |
| KT837151 | A/California/19/2014(H3N2) | 4/9/2014 | NCBI |
| KT837104 | A/Michigan/10/2014(H3N2) | 10/9/2014 | NCBI |
| KT837132 | A/Alaska/52/2014(H3N2) | 3/9/2014 | NCBI |
| KT837267 | A/Wisconsin/34/2014(H3N2) | 30/9/2014 | NCBI |
| KT837259 | A/Arkansas/01/2014(H3N2) | 17/9/2014 | NCBI |
| KT837201 | A/Hawaii/40/2014(H3N2) | 7/9/2014 | NCBI |
| KT837202 | A/Ohio/05/2014(H3N2) | 30/9/2014 | NCBI |
| KT837234 | A/NorthCarolina/23/2014(H3N2) | 9/9/2014 | NCBI |
| KT837251 | A/NewYork/19/2014(H3N2) | 23/9/2014 | NCBI |
| KT837176 | A/Arkansas/02/2014(H3N2) | 17/9/2014 | NCBI |
| KT837256 | A/NewYork/20/2014(H3N2) | 25/9/2014 | NCBI |
| KT837174 | A/NewJersey/22/2014(H3N2) | 16/9/2014 | NCBI |
| KT837178 | A/Arkansas/04/2014(H3N2) | 22/9/2014 | NCBI |
| KT837180 | A/NorthCarolina/24/2014(H3N2) | 24/9/2014 | NCBI |
| KT837206 | A/NewYork/18/2014(H3N2) | 18/9/2014 | NCBI |
| KT837273 | A/Arkansas/05/2014(H3N2) | 24/9/2014 | NCBI |
| KT837286 | A/Arkansas/06/2014(H3N2) | 24/9/2014 | NCBI |
| KT837299 | A/Pennsylvania/43/2014(H3N2) | 24/9/2014 | NCBI |
| KT837303 | A/Pennsylvania/42/2014(H3N2) | 19/9/2014 | NCBI |
| KT837311 | A/Arkansas/07/2014(H3N2) | 29/9/2014 | NCBI |
| KT838345 | A/Alabama/05/2014(H3N2) | 20/10/2014 | NCBI |
| KT837326 | A/Connecticut/21/2014(H3N2) | 14/10/2014 | NCBI |
| KT837447 | A/Louisiana/07/2014(H3N2) | 10/10/2014 | NCBI |
| KT837461 | A/Texas/43/2014(H3N2) | 15/10/2014 | NCBI |
| KT838182 | A/Florida/45/2014(H3N2) | 12/10/2014 | NCBI |
| KT837494 | A/Louisiana/09/2014(H3N2) | 11/10/2014 | NCBI |
| KT837410 | A/Louisiana/06/2014(H3N2) | 7/10/2014 | NCBI |
| KT837434 | A/Indiana/10/2014(H3N2) | 25/10/2014 | NCBI |
| KT837436 | A/Maine/05/2014(H3N2) | 14/10/2014 | NCBI |
| KT837347 | A/Colorado/16/2014(H3N2) | 3/10/2014 | NCBI |

| | | | |
|---|---|---|---|
| KT837415 | A/Louisiana/14/2014(H3N2) | 22/10/2014 | NCBI |
| KT837342 | A/DistrictOfColumbia/03/2014(H3N2) | 30/10/2014 | NCBI |
| KT837448 | A/California/53/2014(H3N2) | 26/10/2014 | NCBI |
| KT838321 | A/Louisiana/11/2014(H3N2) | 21/10/2014 | NCBI |
| KT837490 | A/Oregon/05/2014(H3N2) | 9/10/2014 | NCBI |
| KT838304 | A/Connecticut/19/2014(H3N2) | 3/10/2014 | NCBI |
| KT837376 | A/Oregon/06/2014(H3N2) | 9/10/2014 | NCBI |
| KT838036 | A/Louisiana/08/2014(H3N2) | 11/10/2014 | NCBI |
| KT838329 | A/California/52/2014(H3N2) | 24/10/2014 | NCBI |
| KT838314 | A/California/51/2014(H3N2) | 19/10/2014 | NCBI |
| KT837367 | A/NewHampshire/21/2014(H3N2) | 17/10/2014 | NCBI |
| KT837484 | A/Oregon/09/2014(H3N2) | 5/10/2014 | NCBI |
| KT837474 | A/Texas/42/2014(H3N2) | 22/10/2014 | NCBI |
| KT838029 | A/NorthCarolina/21/2014(H3N2) | 6/10/2014 | NCBI |
| KT837861 | A/NorthCarolina/21/2014(H3N2) | 6/10/2014 | NCBI |
| KT837514 | A/California/50/2014(H3N2) | 18/10/2014 | NCBI |
| KT837773 | A/Florida/38/2014(H3N2) | 7/10/2014 | NCBI |
| KT837887 | A/Florida/37/2014(H3N2) | 7/10/2014 | NCBI |
| KT837740 | A/Minnesota/31/2014(H3N2) | 28/10/2014 | NCBI |
| KT837392 | A/Texas/37/2014(H3N2) | 1/10/2014 | NCBI |
| KT837413 | A/Arkansas/13/2014(H3N2) | 28/10/2014 | NCBI |
| KT837411 | A/Texas/66/2014(H3N2) | 30/10/2014 | NCBI |
| KT837713 | A/Delaware/22/2014(H3N2) | 21/10/2014 | NCBI |
| KT837827 | A/Florida/39/2014(H3N2) | 7/10/2014 | NCBI |
| KT837762 | A/Iowa/14/2014(H3N2) | 12/10/2014 | NCBI |
| KT837722 | A/Alaska/47/2014(H3N2) | 5/10/2014 | NCBI |
| KT837598 | A/Washington/18/2014(H3N2) | 6/10/2014 | NCBI |
| KT837619 | A/Nevada/20/2014(H3N2) | 27/10/2014 | NCBI |
| KT838288 | A/NorthDakota/06/2014(H3N2) | 1/10/2014 | NCBI |
| KT838131 | A/Hawaii/48/2014(H3N2) | 18/10/2014 | NCBI |
| KT838138 | A/Arkansas/10/2014(H3N2) | 20/10/2014 | NCBI |
| KT837568 | A/Ohio/03/2014(H3N2) | 29/10/2014 | NCBI |
| KT837683 | A/Iowa/15/2014(H3N2) | 20/10/2014 | NCBI |
| KT837647 | A/Montana/14/2014(H3N2) | 28/10/2014 | NCBI |
| KT837602 | A/NorthDakota/09/2014(H3N2) | 24/10/2014 | NCBI |
| KT837533 | A/Montana/15/2014(H3N2) | 30/10/2014 | NCBI |
| KT837674 | A/Mississippi/08/2014(H3N2) | 23/10/2014 | NCBI |

| | | | |
|---|---|---|---|
| KT837612 | A/Texas/48/2014(H3N2) | 27/10/2014 | NCBI |
| KT837626 | A/Ohio/04/2014(H3N2) | 29/10/2014 | NCBI |
| KT837538 | A/Mississippi/09/2014(H3N2) | 29/10/2014 | NCBI |
| KT837666 | A/Kansas/08/2014(H3N2) | 29/10/2014 | NCBI |
| KT837574 | A/Louisiana/13/2014(H3N2) | 21/10/2014 | NCBI |
| KT837960 | A/NewYork/28/2014(H3N2) | 14/10/2014 | NCBI |
| KT837668 | A/Maryland/33/2014(H3N2) | 29/10/2014 | NCBI |
| KT837611 | A/Iowa/16/2014(H3N2) | 21/10/2014 | NCBI |
| KT837659 | A/Alabama/07/2014(H3N2) | 23/10/2014 | NCBI |
| KT837560 | A/Iowa/13/2014(H3N2) | 9/10/2014 | NCBI |
| KT837669 | A/Texas/46/2014(H3N2) | 30/10/2014 | NCBI |
| KT837642 | A/Louisiana/12/2014(H3N2) | 21/10/2014 | NCBI |
| KT837566 | A/Kansas/07/2014(H3N2) | 27/10/2014 | NCBI |
| KT837951 | A/Arkansas/11/2014(H3N2) | 20/10/2014 | NCBI |
| KT837971 | A/Florida/43/2014(H3N2) | 6/10/2014 | NCBI |
| KT837787 | A/Florida/40/2014(H3N2) | 7/10/2014 | NCBI |
| KT837755 | A/Maryland/34/2014(H3N2) | 23/10/2014 | NCBI |
| KT837754 | A/Alaska/46/2014(H3N2) | 4/10/2014 | NCBI |
| KT837826 | A/California/27/2014(H3N2) | 10/10/2014 | NCBI |
| KT837837 | A/Mississippi/03/2014(H3N2) | 15/10/2014 | NCBI |
| KT837930 | A/Hawaii/49/2014(H3N2) | 18/10/2014 | NCBI |
| KT837715 | A/Texas/38/2014(H3N2) | 6/10/2014 | NCBI |
| KT838161 | A/Washington/20/2014(H3N2) | 10/10/2014 | NCBI |
| KT837770 | A/Colorado/15/2014(H3N2) | 4/10/2014 | NCBI |
| KT837807 | A/Mississippi/04/2014(H3N2) | 16/10/2014 | NCBI |
| KT837817 | A/Mississippi/05/2014(H3N2) | 14/10/2014 | NCBI |
| KT837843 | A/Wyoming/08/2014(H3N2) | 9/10/2014 | NCBI |
| KT837812 | A/Mississippi/06/2014(H3N2) | 14/10/2014 | NCBI |
| KT838114 | A/Virginia/16/2014(H3N2) | 14/10/2014 | NCBI |
| KT837779 | A/Maine/04/2014(H3N2) | 9/10/2014 | NCBI |
| KT837816 | A/Kentucky/10/2014(H3N2) | 4/10/2014 | NCBI |
| KT837800 | A/Florida/36/2014(H3N2) | 3/10/2014 | NCBI |
| KT837808 | A/NewYork/26/2014(H3N2) | 14/10/2014 | NCBI |
| KT837838 | A/Wyoming/09/2014(H3N2) | 9/10/2014 | NCBI |
| KT838136 | A/Washington/19/2014(H3N2) | 22/10/2014 | NCBI |
| KT838116 | A/Wisconsin/44/2014(H3N2) | 23/10/2014 | NCBI |
| KT838098 | A/Iowa/17/2014(H3N2) | 25/10/2014 | NCBI |

| | | | |
|---|---|---|---|
| KT838001 | A/NewYork/24/2014(H3N2) | 10/10/2014 | NCBI |
| KT838242 | A/Colorado/21/2014(H3N2) | 21/10/2014 | NCBI |
| KT838061 | A/Wisconsin/46/2014(H3N2) | 28/10/2014 | NCBI |
| KT838088 | A/Illinois/04/2014(H3N2) | 27/10/2014 | NCBI |
| KT838047 | A/Maine/06/2014(H3N2) | 18/10/2014 | NCBI |
| KT837993 | A/Arkansas/12/2014(H3N2) | 21/10/2014 | NCBI |
| KT837933 | A/Maryland/32/2014(H3N2) | 15/10/2014 | NCBI |
| KT838163 | A/Colorado/19/2014(H3N2) | 18/10/2014 | NCBI |
| KT837999 | A/Maryland/28/2014(H3N2) | 14/10/2014 | NCBI |
| KT837997 | A/NewYork/35/2014(H3N2) | 21/10/2014 | NCBI |
| KT838041 | A/NewYork/36/2014(H3N2) | 23/10/2014 | NCBI |
| KT838015 | A/NewYork/34/2014(H3N2) | 17/10/2014 | NCBI |
| KT838076 | A/NewYork/31/2014(H3N2) | 17/10/2014 | NCBI |
| KT838269 | A/Texas/41/2014(H3N2) | 20/10/2014 | NCBI |
| KT838287 | A/Mississippi/07/2014(H3N2) | 6/10/2014 | NCBI |
| KT838145 | A/Montana/11/2014(H3N2) | 17/10/2014 | NCBI |
| KT838222 | A/Arkansas/08/2014(H3N2) | 1/10/2014 | NCBI |
| KT838150 | A/Minnesota/28/2014(H3N2) | 18/10/2014 | NCBI |
| KT838244 | A/Delaware/19/2014(H3N2) | 11/10/2014 | NCBI |
| KT838203 | A/NewMexico/28/2014(H3N2) | 15/10/2014 | NCBI |
| KT839265 | A/Michigan/19/2014(H3N2) | 14/11/2014 | NCBI |
| KT839366 | A/Louisiana/28/2014(H3N2) | 24/11/2014 | NCBI |
| KT838396 | A/Michigan/18/2014(H3N2) | 10/11/2014 | NCBI |
| KT839386 | A/Illinois/07/2014(H3N2) | 10/11/2014 | NCBI |
| KT838362 | A/RhodeIsland/21/2014(H3N2) | 30/11/2014 | NCBI |
| KT838381 | A/Texas/55/2014(H3N2) | 18/11/2014 | NCBI |
| KT839231 | A/Louisiana/20/2014(H3N2) | 19/11/2014 | NCBI |
| KT838844 | A/Louisiana/20/2014(H3N2) | 19/11/2014 | NCBI |
| KT838367 | A/Texas/69/2014(H3N2) | 24/11/2014 | NCBI |
| KT838376 | A/Texas/54/2014(H3N2) | 17/11/2014 | NCBI |
| KT839381 | A/Alabama/08/2014(H3N2) | 5/11/2014 | NCBI |
| KT838394 | A/Oklahoma/09/2014(H3N2) | 24/11/2014 | NCBI |
| KT838878 | A/Missouri/11/2014(H3N2) | 18/11/2014 | NCBI |
| KT839286 | A/Pennsylvania/48/2014(H3N2) | 28/11/2014 | NCBI |
| KT838698 | A/Alabama/09/2014(H3N2) | 5/11/2014 | NCBI |
| KT839293 | A/Alabama/09/2014(H3N2) | 5/11/2014 | NCBI |
| KT839275 | A/NewHampshire/28/2014(H3N2) | 28/11/2014 | NCBI |

| | | | |
|---|---|---|---|
| KT838385 | A/Pennsylvania/47/2014(H3N2) | 28/11/2014 | NCBI |
| KT839331 | A/Texas/68/2014(H3N2) | 24/11/2014 | NCBI |
| KT838590 | A/Utah/17/2014(H3N2) | 6/11/2014 | NCBI |
| KT839304 | A/Utah/17/2014(H3N2) | 6/11/2014 | NCBI |
| KT838399 | A/Texas/70/2014(H3N2) | 24/11/2014 | NCBI |
| KT839351 | A/NewJersey/29/2014(H3N2) | 20/11/2014 | NCBI |
| KT839330 | A/NewMexico/29/2014(H3N2) | 18/11/2014 | NCBI |
| KT839362 | A/NorthCarolina/31/2014(H3N2) | 13/11/2014 | NCBI |
| KT838865 | A/NorthCarolina/31/2014(H3N2) | 13/11/2014 | NCBI |
| KT839272 | A/Idaho/07/2014(H3N2) | 25/11/2014 | NCBI |
| KT839376 | A/Michigan/17/2014(H3N2) | 10/11/2014 | NCBI |
| KT838412 | A/NewMexico/30/2014(H3N2) | 28/11/2014 | NCBI |
| KT838415 | A/Minnesota/36/2014(H3N2) | 23/11/2014 | NCBI |
| KT839143 | A/Colorado/39/2014(H3N2) | 29/11/2014 | NCBI |
| KT838416 | A/Mississippi/15/2014(H3N2) | 25/11/2014 | NCBI |
| KT838424 | A/Indiana/21/2014(H3N2) | 27/11/2014 | NCBI |
| KT839216 | A/Mississippi/17/2014(H3N2) | 13/11/2014 | NCBI |
| KT838468 | A/Ohio/07/2014(H3N2) | 19/11/2014 | NCBI |
| KT838423 | A/Iowa/20/2014(H3N2) | 14/11/2014 | NCBI |
| KT838504 | A/Ohio/08/2014(H3N2) | 22/11/2014 | NCBI |
| KT839110 | A/Montana/20/2014(H3N2) | 11/11/2014 | NCBI |
| KT838507 | A/Hawaii/60/2014(H3N2) | 3/11/2014 | NCBI |
| KT838500 | A/Maryland/40/2014(H3N2) | 25/11/2014 | NCBI |
| KT838474 | A/Minnesota/41/2014(H3N2) | 23/11/2014 | NCBI |
| KT838441 | A/Minnesota/35/2014(H3N2) | 29/11/2014 | NCBI |
| KT838483 | A/Oklahoma/10/2014(H3N2) | 24/11/2014 | NCBI |
| KT838452 | A/Wyoming/17/2014(H3N2) | 24/11/2014 | NCBI |
| KT839204 | A/Texas/52/2014(H3N2) | 5/11/2014 | NCBI |
| KT838445 | A/Nevada/18/2014(H3N2) | 10/11/2014 | NCBI |
| KT838838 | A/NorthCarolina/28/2014(H3N2) | 4/11/2014 | NCBI |
| KT838916 | A/Delaware/25/2014(H3N2) | 20/11/2014 | NCBI |
| KT838853 | A/Delaware/25/2014(H3N2) | 20/11/2014 | NCBI |
| KT838475 | A/Massachusetts/18/2014(H3N2) | 30/11/2014 | NCBI |
| KT838809 | A/Illinois/05/2014(H3N2) | 3/11/2014 | NCBI |
| KT839229 | A/Illinois/05/2014(H3N2) | 3/11/2014 | NCBI |
| KT839223 | A/Ohio/15/2014(H3N2) | 11/11/2014 | NCBI |
| KT838489 | A/Colorado/28/2014(H3N2) | 20/11/2014 | NCBI |

| | | | |
|---|---|---|---|
| KT839200 | A/Georgia/15/2014(H3N2) | 13/11/2014 | NCBI |
| KT839159 | A/Minnesota/34/2014(H3N2) | 12/11/2014 | NCBI |
| KT838749 | A/NorthCarolina/33/2014(H3N2) | 13/11/2014 | NCBI |
| KT838647 | A/Mississippi/10/2014(H3N2) | 3/11/2014 | NCBI |
| KT838664 | A/Georgia/13/2014(H3N2) | 14/11/2014 | NCBI |
| KT838622 | A/Louisiana/17/2014(H3N2) | 18/11/2014 | NCBI |
| KT838856 | A/Louisiana/17/2014(H3N2) | 18/11/2014 | NCBI |
| KT838666 | A/Nevada/17/2014(H3N2) | 10/11/2014 | NCBI |
| KT838536 | A/Indiana/15/2014(H3N2) | 10/11/2014 | NCBI |
| KT838707 | A/Utah/22/2014(H3N2) | 8/11/2014 | NCBI |
| KT838579 | A/Utah/20/2014(H3N2) | 5/11/2014 | NCBI |
| KT838545 | A/Virginia/26/2014(H3N2) | 25/11/2014 | NCBI |
| KT838547 | A/Mississippi/12/2014(H3N2) | 5/11/2014 | NCBI |
| KT839047 | A/Utah/19/2014(H3N2) | 7/11/2014 | NCBI |
| KT839037 | A/NorthCarolina/32/2014(H3N2) | 13/11/2014 | NCBI |
| KT838551 | A/Mississippi/11/2014(H3N2) | 4/11/2014 | NCBI |
| KT838557 | A/Indiana/11/2014(H3N2) | 5/11/2014 | NCBI |
| KT838576 | A/Indiana/16/2014(H3N2) | 10/11/2014 | NCBI |
| KT838600 | A/Utah/18/2014(H3N2) | 2/11/2014 | NCBI |
| KT838706 | A/Indiana/13/2014(H3N2) | 7/11/2014 | NCBI |
| KT838651 | A/Missouri/09/2014(H3N2) | 3/11/2014 | NCBI |
| KT838594 | A/Idaho/04/2014(H3N2) | 5/11/2014 | NCBI |
| KT838599 | A/Utah/21/2014(H3N2) | 6/11/2014 | NCBI |
| KT838806 | A/Louisiana/22/2014(H3N2) | 19/11/2014 | NCBI |
| KT838816 | A/Massachusetts/20/2014(H3N2) | 18/11/2014 | NCBI |
| KT838793 | A/Oregon/13/2014(H3N2) | 18/11/2014 | NCBI |
| KT838670 | A/Indiana/12/2014(H3N2) | 5/11/2014 | NCBI |
| KT838753 | A/Louisiana/21/2014(H3N2) | 19/11/2014 | NCBI |
| KT838783 | A/Oregon/11/2014(H3N2) | 21/11/2014 | NCBI |
| KT838732 | A/Michigan/26/2014(H3N2) | 25/11/2014 | NCBI |
| KT838769 | A/Minnesota/37/2014(H3N2) | 16/11/2014 | NCBI |
| KT839040 | A/Washington/33/2014(H3N2) | 21/11/2014 | NCBI |
| KT838690 | A/Hawaii/59/2014(H3N2) | 13/11/2014 | NCBI |
| KT838731 | A/Indiana/17/2014(H3N2) | 11/11/2014 | NCBI |
| KT839225 | A/Wisconsin/64/2014(H3N2) | 22/11/2014 | NCBI |
| KT838788 | A/Washington/36/2014(H3N2) | 21/11/2014 | NCBI |
| KT838931 | A/Wyoming/15/2014(H3N2) | 18/11/2014 | NCBI |

| | | | |
|---|---|---|---|
| KT838694 | A/Wyoming/15/2014(H3N2) | 18/11/2014 | NCBI |
| KT838757 | A/Wyoming/11/2014(H3N2) | 7/11/2014 | NCBI |
| KT838759 | A/Louisiana/19/2014(H3N2) | 18/11/2014 | NCBI |
| KT838765 | A/Louisiana/18/2014(H3N2) | 18/11/2014 | NCBI |
| KT838766 | A/NorthCarolina/35/2014(H3N2) | 14/11/2014 | NCBI |
| KT839021 | A/Colorado/25/2014(H3N2) | 9/11/2014 | NCBI |
| KT838833 | A/NewHampshire/23/2014(H3N2) | 19/11/2014 | NCBI |
| KT838930 | A/NewHampshire/23/2014(H3N2) | 19/11/2014 | NCBI |
| KT839070 | A/Ohio/14/2014(H3N2) | 9/11/2014 | NCBI |
| KT838939 | A/Florida/58/2014(H3N2) | 8/11/2014 | NCBI |
| KT839017 | A/NorthCarolina/34/2014(H3N2) | 13/11/2014 | NCBI |
| KT839011 | A/NorthCarolina/29/2014(H3N2) | 8/11/2014 | NCBI |
| KT839005 | A/Georgia/06/2014(H3N2) | 20/11/2014 | NCBI |
| KT838984 | A/Georgia/04/2014(H3N2) | 20/11/2014 | NCBI |
| KT838991 | A/Kansas/10/2014(H3N2) | 19/11/2014 | NCBI |
| KT838921 | A/Texas/57/2014(H3N2) | 25/11/2014 | NCBI |
| KT839006 | A/NewJersey/26/2014(H3N2) | 17/11/2014 | NCBI |
| KT838940 | A/Missouri/10/2014(H3N2) | 17/11/2014 | NCBI |
| KT839090 | A/NewJersey/27/2014(H3N2) | 19/11/2014 | NCBI |
| KT839000 | A/Georgia/07/2014(H3N2) | 20/11/2014 | NCBI |
| KT838934 | A/Georgia/05/2014(H3N2) | 20/11/2014 | NCBI |
| KT838997 | A/Delaware/24/2014(H3N2) | 11/11/2014 | NCBI |
| KT838949 | A/Massachusetts/14/2014(H3N2) | 19/11/2014 | NCBI |
| KT839002 | A/Wyoming/14/2014(H3N2) | 14/11/2014 | NCBI |
| KT839370 | A/Texas/56/2014(H3N2) | 25/11/2014 | NCBI |
| KT839108 | A/Georgia/18/2014(H3N2) | 29/11/2014 | NCBI |
| KT839061 | A/Wisconsin/49/2014(H3N2) | 9/11/2014 | NCBI |
| KT839195 | A/Texas/62/2014(H3N2) | 3/11/2014 | NCBI |
| KT839117 | A/Indiana/19/2014(H3N2) | 12/11/2014 | NCBI |
| KT839166 | A/Colorado/24/2014(H3N2) | 7/11/2014 | NCBI |
| KT839124 | A/Wisconsin/48/2014(H3N2) | 8/11/2014 | NCBI |
| KT839152 | A/Georgia/23/2014(H3N2) | 26/11/2014 | NCBI |
| KT839084 | A/Georgia/19/2014(H3N2) | 14/11/2014 | NCBI |
| KT839116 | A/Georgia/16/2014(H3N2) | 27/11/2014 | NCBI |
| KT839151 | A/Virginia/36/2014(H3N2) | 25/11/2014 | NCBI |
| KT839323 | A/Pennsylvania/49/2014(H3N2) | 28/11/2014 | NCBI |
| KT839341 | A/Kentucky/14/2014(H3N2) | 26/11/2014 | NCBI |

| | | | |
|---|---|---|---|
| KT840260 | A/Tennessee/24/2014(H3N2) | 2/12/2014 | NCBI |
| KT839487 | A/Pennsylvania/51/2014(H3N2) | 2/12/2014 | NCBI |
| KT839850 | A/Louisiana/38/2014(H3N2) | 6/12/2014 | NCBI |
| KT840257 | A/Kansas/12/2014(H3N2) | 8/12/2014 | NCBI |
| KT839725 | A/NewHampshire/29/2014(H3N2) | 4/12/2014 | NCBI |
| KT840264 | A/RhodeIsland/22/2014(H3N2) | 8/12/2014 | NCBI |
| KT839403 | A/Missouri/13/2014(H3N2) | 1/12/2014 | NCBI |
| KT840276 | A/Nevada/21/2014(H3N2) | 2/12/2014 | NCBI |
| KT839470 | A/Virginia/37/2014(H3N2) | 5/12/2014 | NCBI |
| KT840216 | A/Nevada/22/2014(H3N2) | 4/12/2014 | NCBI |
| KT840235 | A/Wyoming/18/2014(H3N2) | 9/12/2014 | NCBI |
| KT839648 | A/Wyoming/20/2014(H3N2) | 9/12/2014 | NCBI |
| KT840214 | A/Washington/35/2014(H3N2) | 1/12/2014 | NCBI |
| KT840247 | A/Georgia/24/2014(H3N2) | 1/12/2014 | NCBI |
| KT840270 | A/NewHampshire/32/2014(H3N2) | 8/12/2014 | NCBI |
| KT839420 | A/Georgia/28/2014(H3N2) | 3/12/2014 | NCBI |
| KT839437 | A/Wisconsin/57/2014(H3N2) | 9/12/2014 | NCBI |
| KT839543 | A/Wisconsin/59/2014(H3N2) | 9/12/2014 | NCBI |
| KT839551 | A/Wisconsin/55/2014(H3N2) | 8/12/2014 | NCBI |
| KT839424 | A/NorthDakota/10/2014(H3N2) | 4/12/2014 | NCBI |
| KT839471 | A/NorthDakota/11/2014(H3N2) | 2/12/2014 | NCBI |
| KT839427 | A/Washington/37/2014(H3N2) | 8/12/2014 | NCBI |
| KT839522 | A/Delaware/31/2014(H3N2) | 7/12/2014 | NCBI |
| KT839429 | A/Florida/65/2014(H3N2) | 5/12/2014 | NCBI |
| KT839447 | A/Wisconsin/54/2014(H3N2) | 8/12/2014 | NCBI |
| KT839463 | A/Wisconsin/56/2014(H3N2) | 9/12/2014 | NCBI |
| KT839615 | A/Wisconsin/60/2014(H3N2) | 10/12/2014 | NCBI |
| KT839526 | A/Wisconsin/52/2014(H3N2) | 2/12/2014 | NCBI |
| KT839558 | A/Massachusetts/16/2014(H3N2) | 3/12/2014 | NCBI |
| KT840272 | A/Missouri/14/2014(H3N2) | 2/12/2014 | NCBI |
| KT839524 | A/Florida/66/2014(H3N2) | 8/12/2014 | NCBI |
| KT840277 | A/Georgia/30/2014(H3N2) | 8/12/2014 | NCBI |
| KT839549 | A/Mississippi/16/2014(H3N2) | 8/12/2014 | NCBI |
| KT839501 | A/Wisconsin/58/2014(H3N2) | 9/12/2014 | NCBI |
| KT839568 | A/Georgia/29/2014(H3N2) | 6/12/2014 | NCBI |
| KT839565 | A/Louisiana/39/2014(H3N2) | 10/12/2014 | NCBI |
| KT839585 | A/NorthCarolina/41/2014(H3N2) | 3/12/2014 | NCBI |

| | | | |
|---|---|---|---|
| KT839499 | A/Massachusetts/17/2014(H3N2) | 2/12/2014 | NCBI |
| KT839547 | A/Kentucky/15/2014(H3N2) | 2/12/2014 | NCBI |
| KT839530 | A/Wisconsin/53/2014(H3N2) | 2/12/2014 | NCBI |
| KT840083 | A/Pennsylvania/52/2014(H3N2) | 2/12/2014 | NCBI |
| KT839932 | A/Kentucky/30/2014(H3N2) | 1/12/2014 | NCBI |
| KT839646 | A/Missouri/17/2014(H3N2) | 4/12/2014 | NCBI |
| KT839576 | A/Pennsylvania/65/2014(H3N2) | 9/12/2014 | NCBI |
| KT840109 | A/Illinois/12/2014(H3N2) | 3/12/2014 | NCBI |
| KT840183 | A/Indiana/24/2014(H3N2) | 1/12/2014 | NCBI |
| KT839602 | A/Colorado/38/2014(H3N2) | 3/12/2014 | NCBI |
| KT839593 | A/Minnesota/38/2014(H3N2) | 7/12/2014 | NCBI |
| KT839722 | A/Tennessee/23/2014(H3N2) | 4/12/2014 | NCBI |
| KT839596 | A/Maryland/44/2014(H3N2) | 10/12/2014 | NCBI |
| KT839622 | A/Delaware/38/2014(H3N2) | 2/12/2014 | NCBI |
| KT839666 | A/Georgia/21/2014(H3N2) | 2/12/2014 | NCBI |
| KT839631 | A/Virginia/42/2014(H3N2) | 9/12/2014 | NCBI |
| KT839694 | A/Tennessee/25/2014(H3N2) | 8/12/2014 | NCBI |
| KT840177 | A/Illinois/11/2014(H3N2) | 2/12/2014 | NCBI |
| KT839755 | A/Indiana/26/2014(H3N2) | 8/12/2014 | NCBI |
| KT839652 | A/NorthCarolina/40/2014(H3N2) | 4/12/2014 | NCBI |
| KT839852 | A/Alabama/12/2014(H3N2) | 9/12/2014 | NCBI |
| KT839990 | A/Alabama/10/2014(H3N2) | 4/12/2014 | NCBI |
| KT839770 | A/Illinois/09/2014(H3N2) | 1/12/2014 | NCBI |
| KT839664 | A/Minnesota/39/2014(H3N2) | 4/12/2014 | NCBI |
| KT839669 | A/Florida/76/2014(H3N2) | 1/12/2014 | NCBI |
| KT839714 | A/Massachusetts/19/2014(H3N2) | 2/12/2014 | NCBI |
| KT839702 | A/California/40/2014(H3N2) | 10/12/2014 | NCBI |
| KT839729 | A/Nebraska/15/2014(H3N2) | 5/12/2014 | NCBI |
| KT840130 | A/Texas/89/2014(H3N2) | 7/12/2014 | NCBI |
| KT839894 | A/California/44/2014(H3N2) | 4/12/2014 | NCBI |
| KT839984 | A/Texas/75/2014(H3N2) | 2/12/2014 | NCBI |
| KT840006 | A/Pennsylvania/67/2014(H3N2) | 4/12/2014 | NCBI |
| KT840114 | A/Arkansas/14/2014(H3N2) | 8/12/2014 | NCBI |
| KT839951 | A/Utah/34/2014(H3N2) | 5/12/2014 | NCBI |
| KT839989 | A/Ohio/09/2014(H3N2) | 4/12/2014 | NCBI |
| KT839906 | A/Alabama/11/2014(H3N2) | 8/12/2014 | NCBI |
| KT840093 | A/Utah/35/2014(H3N2) | 7/12/2014 | NCBI |

| | | | |
|---|---|---|---|
| KT839892 | A/Tennessee/18/2014(H3N2) | 9/12/2014 | NCBI |
| KT839861 | A/Arkansas/15/2014(H3N2) | 9/12/2014 | NCBI |
| KT840120 | A/Texas/80/2014(H3N2) | 2/12/2014 | NCBI |
| KT839927 | A/Mississippi/20/2014(H3N2) | 9/12/2014 | NCBI |
| KT839933 | A/Virginia/29/2014(H3N2) | 9/12/2014 | NCBI |
| KT839815 | A/Tennessee/17/2014(H3N2) | 8/12/2014 | NCBI |
| KT840205 | A/Florida/85/2014(H3N2) | 3/12/2014 | NCBI |
| KT840034 | A/Virginia/32/2014(H3N2) | 10/12/2014 | NCBI |
| KT839873 | A/Nevada/24/2014(H3N2) | 4/12/2014 | NCBI |
| KT839912 | A/Mississippi/19/2014(H3N2) | 8/12/2014 | NCBI |
| KT839842 | A/Utah/33/2014(H3N2) | 2/12/2014 | NCBI |
| KT840140 | A/Texas/74/2014(H3N2) | 8/12/2014 | NCBI |
| KT839974 | A/Tennessee/16/2014(H3N2) | 8/12/2014 | NCBI |
| KT840132 | A/Tennessee/15/2014(H3N2) | 9/12/2014 | NCBI |
| KT839947 | A/California/42/2014(H3N2) | 4/12/2014 | NCBI |
| KT839994 | A/Montana/26/2014(H3N2) | 8/12/2014 | NCBI |
| KT840077 | A/Montana/27/2014(H3N2) | 9/12/2014 | NCBI |
| KT839845 | A/Indiana/25/2014(H3N2) | 5/12/2014 | NCBI |
| KT839955 | A/Texas/71/2014(H3N2) | 8/12/2014 | NCBI |
| KT839966 | A/NorthDakota/14/2014(H3N2) | 3/12/2014 | NCBI |
| KT840095 | A/Virginia/33/2014(H3N2) | 8/12/2014 | NCBI |
| KT840064 | A/Ohio/10/2014(H3N2) | 9/12/2014 | NCBI |
| KT840023 | A/Wyoming/19/2014(H3N2) | 10/12/2014 | NCBI |
| KT839868 | A/Utah/27/2014(H3N2) | 5/12/2014 | NCBI |
| KT839893 | A/Delaware/35/2014(H3N2) | 7/12/2014 | NCBI |
| KT840081 | A/Missouri/16/2014(H3N2) | 10/12/2014 | NCBI |
| KT839922 | A/NewHampshire/30/2014(H3N2) | 1/12/2014 | NCBI |
| KT840133 | A/Texas/81/2014(H3N2) | 10/12/2014 | NCBI |
| KT839946 | A/Texas/72/2014(H3N2) | 8/12/2014 | NCBI |
| KT839972 | A/Texas/73/2014(H3N2) | 8/12/2014 | NCBI |
| KT840226 | A/Pennsylvania/56/2014(H3N2) | 1/12/2014 | NCBI |
| KT840176 | A/Missouri/18/2014(H3N2) | 10/12/2014 | NCBI |
| KT840180 | A/Georgia/20/2014(H3N2) | 3/12/2014 | NCBI |
| KT840253 | A/Texas/88/2014(H3N2) | 5/12/2014 | NCBI |
| KT840841 | A/Georgia/38/2014(H3N2) | 17/12/2014 | NCBI |
| KT840287 | A/Delaware/42/2014(H3N2) | 22/12/2014 | NCBI |
| KT841091 | A/DistrictOfColumbia/05/2014(H3N2) | 19/12/2014 | NCBI |

| | | | |
|---|---|---|---|
| KT840935 | A/Florida/89/2014(H3N2) | 18/12/2014 | NCBI |
| KT840513 | A/Colorado/36/2014(H3N2) | 17/12/2014 | NCBI |
| KT841061 | A/Idaho/10/2014(H3N2) | 18/12/2014 | NCBI |
| KT840438 | A/RhodeIsland/23/2014(H3N2) | 19/12/2014 | NCBI |
| KT840368 | A/WestVirginia/06/2014(H3N2) | 14/12/2014 | NCBI |
| KT840663 | A/Wisconsin/68/2014(H3N2) | 16/12/2014 | NCBI |
| KT841087 | A/Texas/93/2014(H3N2) | 16/12/2014 | NCBI |
| KT840328 | A/Texas/96/2014(H3N2) | 26/12/2014 | NCBI |
| KT840394 | A/DistrictOfColumbia/08/2014(H3N2) | 27/12/2014 | NCBI |
| KT840955 | A/NewYork/40/2014(H3N2) | 29/12/2014 | NCBI |
| KT840640 | A/Idaho/08/2014(H3N2) | 16/12/2014 | NCBI |
| KT841085 | A/California/43/2014(H3N2) | 23/12/2014 | NCBI |
| KT840321 | A/Delaware/39/2014(H3N2) | 29/12/2014 | NCBI |
| KT840718 | A/Illinois/14/2014(H3N2) | 18/12/2014 | NCBI |
| KT840476 | A/Kentucky/24/2014(H3N2) | 22/12/2014 | NCBI |
| KT840519 | A/Texas/106/2014(H3N2) | 20/12/2014 | NCBI |
| KT840434 | A/Kentucky/25/2014(H3N2) | 22/12/2014 | NCBI |
| KT840421 | A/Michigan/31/2014(H3N2) | 22/12/2014 | NCBI |
| KT840649 | A/Idaho/09/2014(H3N2) | 18/12/2014 | NCBI |
| KT840322 | A/DistrictOfColumbia/10/2014(H3N2) | 28/12/2014 | NCBI |
| KT840725 | A/Missouri/21/2014(H3N2) | 17/12/2014 | NCBI |
| KT841079 | A/Hawaii/63/2014(H3N2) | 15/12/2014 | NCBI |
| KT840998 | A/Hawaii/63/2014(H3N2) | 15/12/2014 | NCBI |
| KT840479 | A/Washington/60/2014(H3N2) | 23/12/2014 | NCBI |
| KT841040 | A/Iowa/31/2014(H3N2) | 25/12/2014 | NCBI |
| KT840330 | A/Georgia/45/2014(H3N2) | 15/12/2014 | NCBI |
| KT840399 | A/Vermont/22/2014(H3N2) | 20/12/2014 | NCBI |
| KT840395 | A/Michigan/32/2014(H3N2) | 26/12/2014 | NCBI |
| KT840487 | A/Florida/92/2014(H3N2) | 22/12/2014 | NCBI |
| KT840968 | A/Texas/94/2014(H3N2) | 23/12/2014 | NCBI |
| KT841080 | A/NewYork/39/2014(H3N2) | 31/12/2014 | NCBI |
| KT840341 | A/Washington/53/2014(H3N2) | 26/12/2014 | NCBI |
| KT840344 | A/Missouri/20/2014(H3N2) | 26/12/2014 | NCBI |
| KT840941 | A/Maryland/53/2014(H3N2) | 17/12/2014 | NCBI |
| KT840468 | A/Kentucky/27/2014(H3N2) | 23/12/2014 | NCBI |
| KT840352 | A/Wisconsin/72/2014(H3N2) | 19/12/2014 | NCBI |
| KT840494 | A/Michigan/27/2014(H3N2) | 12/12/2014 | NCBI |

| | | | |
|---|---|---|---|
| KT840361 | A/Texas/107/2014(H3N2) | 15/12/2014 | NCBI |
| KT841025 | A/Wisconsin/75/2014(H3N2) | 22/12/2014 | NCBI |
| KT840415 | A/Texas/95/2014(H3N2) | 26/12/2014 | NCBI |
| KT840532 | A/Colorado/35/2014(H3N2) | 12/12/2014 | NCBI |
| KT840514 | A/Michigan/30/2014(H3N2) | 16/12/2014 | NCBI |
| KT840542 | A/Texas/105/2014(H3N2) | 26/12/2014 | NCBI |
| KT840475 | A/Michigan/28/2014(H3N2) | 29/12/2014 | NCBI |
| KT840445 | A/Washington/57/2014(H3N2) | 29/12/2014 | NCBI |
| KT840669 | A/Pennsylvania/64/2014(H3N2) | 11/12/2014 | NCBI |
| KT840717 | A/Montana/30/2014(H3N2) | 18/12/2014 | NCBI |
| KT840515 | A/DistrictOfColumbia/04/2014(H3N2) | 26/12/2014 | NCBI |
| KT840668 | A/Maryland/45/2014(H3N2) | 12/12/2014 | NCBI |
| KT840639 | A/Nevada/30/2014(H3N2) | 17/12/2014 | NCBI |
| KT840507 | A/Pennsylvania/63/2014(H3N2) | 13/12/2014 | NCBI |
| KT840509 | A/Washington/48/2014(H3N2) | 19/12/2014 | NCBI |
| KT840525 | A/Missouri/19/2014(H3N2) | 23/12/2014 | NCBI |
| KT840638 | A/Washington/47/2014(H3N2) | 17/12/2014 | NCBI |
| KT840538 | A/Tennessee/20/2014(H3N2) | 15/12/2014 | NCBI |
| KT841013 | A/Hawaii/65/2014(H3N2) | 24/12/2014 | NCBI |
| KT840730 | A/Washington/46/2014(H3N2) | 17/12/2014 | NCBI |
| KT840688 | A/Montana/28/2014(H3N2) | 12/12/2014 | NCBI |
| KT840964 | A/Florida/80/2014(H3N2) | 18/12/2014 | NCBI |
| KT840735 | A/Massachusetts/25/2014(H3N2) | 11/12/2014 | NCBI |
| KT840873 | A/Virginia/34/2014(H3N2) | 11/12/2014 | NCBI |
| KT840664 | A/Massachusetts/27/2014(H3N2) | 21/12/2014 | NCBI |
| KT840611 | A/Kentucky/21/2014(H3N2) | 17/12/2014 | NCBI |
| KT840695 | A/Massachusetts/22/2014(H3N2) | 17/12/2014 | NCBI |
| KT840982 | A/Maryland/50/2014(H3N2) | 17/12/2014 | NCBI |
| KT840727 | A/Virginia/30/2014(H3N2) | 11/12/2014 | NCBI |
| KT840791 | A/Florida/73/2014(H3N2) | 11/12/2014 | NCBI |
| KT840748 | A/Vermont/10/2014(H3N2) | 13/12/2014 | NCBI |
| KT840736 | A/Massachusetts/23/2014(H3N2) | 14/12/2014 | NCBI |
| KT840636 | A/Wisconsin/69/2014(H3N2) | 17/12/2014 | NCBI |
| KT840701 | A/Nebraska/14/2014(H3N2) | 11/12/2014 | NCBI |
| KT840689 | A/Virginia/31/2014(H3N2) | 11/12/2014 | NCBI |
| KT840680 | A/Pennsylvania/60/2014(H3N2) | 22/12/2014 | NCBI |
| KT840733 | A/Vermont/09/2014(H3N2) | 14/12/2014 | NCBI |

| | | | |
|---|---|---|---|
| KT840844 | A/Texas/79/2014(H3N2) | 15/12/2014 | NCBI |
| KT840772 | A/Virginia/41/2014(H3N2) | 28/12/2014 | NCBI |
| KT840987 | A/Delaware/43/2014(H3N2) | 17/12/2014 | NCBI |
| KT840760 | A/Idaho/12/2014(H3N2) | 16/12/2014 | NCBI |
| KT840809 | A/Nevada/27/2014(H3N2) | 22/12/2014 | NCBI |
| KT840812 | A/Washington/39/2014(H3N2) | 23/12/2014 | NCBI |
| KT840805 | A/Maine/08/2014(H3N2) | 30/12/2014 | NCBI |
| KT840816 | A/Texas/82/2014(H3N2) | 11/12/2014 | NCBI |
| KT840904 | A/Vermont/11/2014(H3N2) | 18/12/2014 | NCBI |
| KT840926 | A/Delaware/37/2014(H3N2) | 16/12/2014 | NCBI |
| KT840874 | A/Vermont/12/2014(H3N2) | 11/12/2014 | NCBI |
| KT840883 | A/Maine/07/2014(H3N2) | 29/12/2014 | NCBI |
| KT840846 | A/Nevada/26/2014(H3N2) | 12/12/2014 | NCBI |
| KT840823 | A/Delaware/36/2014(H3N2) | 16/12/2014 | NCBI |
| KT840924 | A/Virginia/35/2014(H3N2) | 22/12/2014 | NCBI |
| KT840851 | A/Nevada/25/2014(H3N2) | 15/12/2014 | NCBI |
| KT840890 | A/Washington/52/2014(H3N2) | 15/12/2014 | NCBI |
| KT840984 | A/Pennsylvania/54/2014(H3N2) | 13/12/2014 | NCBI |
| KT840959 | A/Nevada/29/2014(H3N2) | 23/12/2014 | NCBI |
| KT841005 | A/Ohio/12/2014(H3N2) | 27/12/2014 | NCBI |
| KT841007 | A/Georgia/39/2014(H3N2) | 16/12/2014 | NCBI |
| KT841064 | A/Texas/84/2014(H3N2) | 11/12/2014 | NCBI |
| KT841056 | A/Texas/85/2014(H3N2) | 11/12/2014 | NCBI |
| KT841052 | A/NewYork/38/2014(H3N2) | 31/12/2014 | NCBI |
| KT841159 | A/NewMexico/01/2015(H3N2) | 3/1/2015 | NCBI |
| KT841458 | A/NewYork/09/2015(H3N2) | 8/1/2015 | NCBI |
| KT841218 | A/NorthCarolina/05/2015(H3N2) | 8/1/2015 | NCBI |
| KT841101 | A/Georgia/03/2015(H3N2) | 6/1/2015 | NCBI |
| KT841721 | A/NewYork/05/2015(H3N2) | 6/1/2015 | NCBI |
| KT841099 | A/Pennsylvania/09/2015(H3N2) | 7/1/2015 | NCBI |
| KT841911 | A/Colorado/06/2015(H3N2) | 8/1/2015 | NCBI |
| KT841110 | A/SouthDakota/01/2015(H3N2) | 5/1/2015 | NCBI |
| KT841103 | A/Georgia/07/2015(H3N2) | 2/1/2015 | NCBI |
| KT841569 | A/California/02/2015(H3N2) | 6/1/2015 | NCBI |
| KT841324 | A/Pennsylvania/10/2015(H3N2) | 8/1/2015 | NCBI |
| KT841106 | A/Pennsylvania/10/2015(H3N2) | 8/1/2015 | NCBI |
| KT841147 | A/NorthDakota/02/2015(H3N2) | 1/1/2015 | NCBI |

| | | | |
|---|---|---|---|
| KT841206 | A/Mississippi/01/2015(H3N2) | 4/1/2015 | NCBI |
| KT841198 | A/Michigan/02/2015(H3N2) | 5/1/2015 | NCBI |
| KT841754 | A/Wisconsin/01/2015(H3N2) | 2/1/2015 | NCBI |
| KT841258 | A/Michigan/03/2015(H3N2) | 2/1/2015 | NCBI |
| KT841988 | A/Mississippi/02/2015(H3N2) | 5/1/2015 | NCBI |
| KT841134 | A/Iowa/04/2015(H3N2) | 6/1/2015 | NCBI |
| KT841888 | A/Idaho/10/2015(H3N2) | 5/1/2015 | NCBI |
| KT841283 | A/Texas/04/2015(H3N2) | 7/1/2015 | NCBI |
| KT841229 | A/Alabama/01/2015(H3N2) | 6/1/2015 | NCBI |
| KT841640 | A/Florida/12/2015(H3N2) | 7/1/2015 | NCBI |
| KT841260 | A/Pennsylvania/06/2015(H3N2) | 7/1/2015 | NCBI |
| KT841175 | A/Wisconsin/03/2015(H3N2) | 2/1/2015 | NCBI |
| KT841124 | A/NorthCarolina/06/2015(H3N2) | 7/1/2015 | NCBI |
| KT841253 | A/California/09/2015(H3N2) | 6/1/2015 | NCBI |
| KT841437 | A/Texas/05/2015(H3N2) | 4/1/2015 | NCBI |
| KT841133 | A/Minnesota/02/2015(H3N2) | 7/1/2015 | NCBI |
| KT841709 | A/Michigan/05/2015(H3N2) | 8/1/2015 | NCBI |
| KT841797 | A/California/04/2015(H3N2) | 7/1/2015 | NCBI |
| KT841338 | A/Maine/04/2015(H3N2) | 2/1/2015 | NCBI |
| KT841343 | A/Virginia/06/2015(H3N2) | 3/1/2015 | NCBI |
| KT841144 | A/Idaho/11/2015(H3N2) | 2/1/2015 | NCBI |
| KT841223 | A/Virginia/02/2015(H3N2) | 4/1/2015 | NCBI |
| KT841376 | A/Virginia/03/2015(H3N2) | 4/1/2015 | NCBI |
| KT841305 | A/Delaware/10/2015(H3N2) | 1/1/2015 | NCBI |
| KT841226 | A/Michigan/12/2015(H3N2) | 8/1/2015 | NCBI |
| KT841242 | A/Washington/02/2015(H3N2) | 2/1/2015 | NCBI |
| KT841630 | A/Minnesota/01/2015(H3N2) | 4/1/2015 | NCBI |
| KT841306 | A/California/06/2015(H3N2) | 1/1/2015 | NCBI |
| KT841445 | A/California/03/2015(H3N2) | 5/1/2015 | NCBI |
| KT841528 | A/Indiana/01/2015(H3N2) | 8/1/2015 | NCBI |
| KT841320 | A/California/08/2015(H3N2) | 5/1/2015 | NCBI |
| KT841348 | A/Pennsylvania/08/2015(H3N2) | 7/1/2015 | NCBI |
| KT841869 | A/Florida/16/2015(H3N2) | 6/1/2015 | NCBI |
| KT841318 | A/Idaho/08/2015(H3N2) | 5/1/2015 | NCBI |
| KT841863 | A/Oregon/02/2015(H3N2) | 2/1/2015 | NCBI |
| KT841651 | A/NorthDakota/03/2015(H3N2) | 1/1/2015 | NCBI |
| KT841775 | A/Florida/02/2015(H3N2) | 2/1/2015 | NCBI |

| | | | |
|---|---|---|---|
| KT841261 | A/Pennsylvania/01/2015(H3N2) | 2/1/2015 | NCBI |
| KT841441 | A/Florida/04/2015(H3N2) | 7/1/2015 | NCBI |
| KT841566 | A/Pennsylvania/11/2015(H3N2) | 7/1/2015 | NCBI |
| KT841327 | A/Texas/02/2015(H3N2) | 6/1/2015 | NCBI |
| KT841423 | A/NewYork/08/2015(H3N2) | 7/1/2015 | NCBI |
| KT841357 | A/RhodeIsland/01/2015(H3N2) | 1/1/2015 | NCBI |
| KT841859 | A/Florida/07/2015(H3N2) | 5/1/2015 | NCBI |
| KT841455 | A/Georgia/05/2015(H3N2) | 2/1/2015 | NCBI |
| KT841853 | A/Maine/03/2015(H3N2) | 1/1/2015 | NCBI |
| KT841323 | A/SouthDakota/20/2015(H3N2) | 6/1/2015 | NCBI |
| KT841780 | A/Montana/01/2015(H3N2) | 6/1/2015 | NCBI |
| KT841495 | A/Mississippi/03/2015(H3N2) | 5/1/2015 | NCBI |
| KT841362 | A/Delaware/03/2015(H3N2) | 6/1/2015 | NCBI |
| KT841686 | A/Colorado/01/2015(H3N2) | 2/1/2015 | NCBI |
| KT841417 | A/Tennessee/06/2015(H3N2) | 6/1/2015 | NCBI |
| KT841405 | A/Tennessee/05/2015(H3N2) | 6/1/2015 | NCBI |
| KT841984 | A/Delaware/02/2015(H3N2) | 2/1/2015 | NCBI |
| KT841752 | A/Colorado/04/2015(H3N2) | 5/1/2015 | NCBI |
| KT841714 | A/Florida/01/2015(H3N2) | 2/1/2015 | NCBI |
| KT841729 | A/Florida/06/2015(H3N2) | 3/1/2015 | NCBI |
| KT841418 | A/Nevada/06/2015(H3N2) | 8/1/2015 | NCBI |
| KT841843 | A/Florida/09/2015(H3N2) | 5/1/2015 | NCBI |
| KT841430 | A/SouthCarolina/01/2015(H3N2) | 3/1/2015 | NCBI |
| KT841654 | A/Florida/11/2015(H3N2) | 6/1/2015 | NCBI |
| KT841668 | A/Oregon/01/2015(H3N2) | 1/1/2015 | NCBI |
| KT841521 | A/Maine/01/2015(H3N2) | 1/1/2015 | NCBI |
| KT841463 | A/RhodeIsland/05/2015(H3N2) | 7/1/2015 | NCBI |
| KT841860 | A/Delaware/01/2015(H3N2) | 2/1/2015 | NCBI |
| KT841894 | A/Florida/03/2015(H3N2) | 2/1/2015 | NCBI |
| KT841450 | A/Georgia/06/2015(H3N2) | 2/1/2015 | NCBI |
| KT841453 | A/Wyoming/01/2015(H3N2) | 6/1/2015 | NCBI |
| KT841693 | A/NewYork/07/2015(H3N2) | 7/1/2015 | NCBI |
| KT841688 | A/Nebraska/01/2015(H3N2) | 5/1/2015 | NCBI |
| KT841606 | A/Maine/05/2015(H3N2) | 2/1/2015 | NCBI |
| KT841511 | A/NewYork/10/2015(H3N2) | 8/1/2015 | NCBI |
| KT841760 | A/NorthCarolina/08/2015(H3N2) | 6/1/2015 | NCBI |
| KT841680 | A/RhodeIsland/03/2015(H3N2) | 6/1/2015 | NCBI |

| | | | |
|---|---|---|---|
| KT841782 | A/RhodeIsland/02/2015(H3N2) | 5/1/2015 | NCBI |
| KT841673 | A/RhodeIsland/04/2015(H3N2) | 7/1/2015 | NCBI |
| KT841637 | A/NewYork/06/2015(H3N2) | 6/1/2015 | NCBI |
| KT841612 | A/Michigan/04/2015(H3N2) | 6/1/2015 | NCBI |
| KT841905 | A/Missouri/02/2015(H3N2) | 4/1/2015 | NCBI |
| KT841587 | A/RhodeIsland/06/2015(H3N2) | 6/1/2015 | NCBI |
| KT841644 | A/Nevada/04/2015(H3N2) | 7/1/2015 | NCBI |
| KT841505 | A/Oklahoma/02/2015(H3N2) | 3/1/2015 | NCBI |
| KT841523 | A/NewYork/11/2015(H3N2) | 5/1/2015 | NCBI |
| KT841705 | A/Nebraska/02/2015(H3N2) | 5/1/2015 | NCBI |
| KT841538 | A/Missouri/03/2015(H3N2) | 8/1/2015 | NCBI |
| KT841906 | A/California/01/2015(H3N2) | 4/1/2015 | NCBI |
| KT841560 | A/Oklahoma/01/2015(H3N2) | 5/1/2015 | NCBI |
| KT841878 | A/Missouri/01/2015(H3N2) | 3/1/2015 | NCBI |
| KT841834 | A/WestVirginia/01/2015(H3N2) | 2/1/2015 | NCBI |
| KT841779 | A/Utah/02/2015(H3N2) | 8/1/2015 | NCBI |
| KT841800 | A/Florida/10/2015(H3N2) | 2/1/2015 | NCBI |
| KT841820 | A/Utah/01/2015(H3N2) | 2/1/2015 | NCBI |
| KT841684 | A/Kentucky/01/2015(H3N2) | 1/1/2015 | NCBI |
| KT841880 | A/Colorado/05/2015(H3N2) | 7/1/2015 | NCBI |
| KT841842 | A/Wisconsin/02/2015(H3N2) | 6/1/2015 | NCBI |
| KT841829 | A/Arkansas/01/2015(H3N2) | 8/1/2015 | NCBI |
| KT841970 | A/California/44/2015(H3N2) | 4/1/2015 | NCBI |
| KT841898 | A/Kentucky/02/2015(H3N2) | 6/1/2015 | NCBI |
| KT841897 | A/Nevada/03/2015(H3N2) | 8/1/2015 | NCBI |
| KT841992 | A/Alabama/07/2015(H3N2) | 11/2/2015 | NCBI |
| KT842157 | A/Wisconsin/22/2015(H3N2) | 27/2/2015 | NCBI |
| KT842730 | A/Kentucky/07/2015(H3N2) | 24/2/2015 | NCBI |
| KT842679 | A/NorthDakota/05/2015(H3N2) | 3/2/2015 | NCBI |
| KT842055 | A/NewHampshire/15/2015(H3N2) | 25/2/2015 | NCBI |
| KT842000 | A/Alabama/09/2015(H3N2) | 14/2/2015 | NCBI |
| KT841998 | A/SouthDakota/30/2015(H3N2) | 13/2/2015 | NCBI |
| KT842007 | A/Nevada/09/2015(H3N2) | 5/2/2015 | NCBI |
| KT842694 | A/Michigan/17/2015(H3N2) | 17/2/2015 | NCBI |
| KT842195 | A/Kansas/01/2015(H3N2) | 1/2/2015 | NCBI |
| KT842010 | A/NewYork/13/2015(H3N2) | 10/2/2015 | NCBI |
| KT842052 | A/Utah/07/2015(H3N2) | 10/2/2015 | NCBI |

| | | | |
|---|---|---|---|
| KT842117 | A/NewYork/14/2015(H3N2) | 5/2/2015 | NCBI |
| KT842064 | A/Texas/38/2015(H3N2) | 27/2/2015 | NCBI |
| KT842789 | A/NewYork/16/2015(H3N2) | 9/2/2015 | NCBI |
| KT842066 | A/SouthDakota/14/2015(H3N2) | 27/2/2015 | NCBI |
| KT842017 | A/NewYork/20/2015(H3N2) | 25/2/2015 | NCBI |
| KT842025 | A/Florida/18/2015(H3N2) | 2/2/2015 | NCBI |
| KT842582 | A/Texas/39/2015(H3N2) | 18/2/2015 | NCBI |
| KT842738 | A/SouthDakota/29/2015(H3N2) | 24/2/2015 | NCBI |
| KT842116 | A/Ohio/04/2015(H3N2) | 6/2/2015 | NCBI |
| KT842675 | A/Kentucky/08/2015(H3N2) | 24/2/2015 | NCBI |
| KT842028 | A/Massachusetts/09/2015(H3N2) | 25/2/2015 | NCBI |
| KT842513 | A/Kansas/08/2015(H3N2) | 19/2/2015 | NCBI |
| KT842119 | A/Pennsylvania/03/2015(H3N2) | 5/2/2015 | NCBI |
| KT842727 | A/Pennsylvania/05/2015(H3N2) | 6/2/2015 | NCBI |
| KT842755 | A/Nevada/10/2015(H3N2) | 14/2/2015 | NCBI |
| KT842740 | A/Florida/19/2015(H3N2) | 1/2/2015 | NCBI |
| KT842136 | A/NorthCarolina/23/2015(H3N2) | 9/2/2015 | NCBI |
| KT842320 | A/Missouri/06/2015(H3N2) | 9/2/2015 | NCBI |
| KT842711 | A/NewHampshire/14/2015(H3N2) | 27/2/2015 | NCBI |
| KT842134 | A/Nevada/19/2015(H3N2) | 18/2/2015 | NCBI |
| KT842239 | A/Indiana/13/2015(H3N2) | 17/2/2015 | NCBI |
| KT842475 | A/NewJersey/04/2015(H3N2) | 1/2/2015 | NCBI |
| KT842783 | A/Hawaii/02/2015(H3N2) | 3/2/2015 | NCBI |
| KT842065 | A/Maine/10/2015(H3N2) | 18/2/2015 | NCBI |
| KT842663 | A/Maryland/03/2015(H3N2) | 16/2/2015 | NCBI |
| KT842144 | A/Maine/09/2015(H3N2) | 13/2/2015 | NCBI |
| KT842112 | A/Texas/09/2015(H3N2) | 3/2/2015 | NCBI |
| KT842766 | A/SouthDakota/05/2015(H3N2) | 3/2/2015 | NCBI |
| KT842672 | A/Wisconsin/23/2015(H3N2) | 26/2/2015 | NCBI |
| KT842686 | A/NewYork/15/2015(H3N2) | 5/2/2015 | NCBI |
| KT842085 | A/NorthCarolina/11/2015(H3N2) | 9/2/2015 | NCBI |
| KT842400 | A/Massachusetts/15/2015(H3N2) | 19/2/2015 | NCBI |
| KT842092 | A/Idaho/09/2015(H3N2) | 4/2/2015 | NCBI |
| KT842581 | A/RhodeIsland/08/2015(H3N2) | 4/2/2015 | NCBI |
| KT842346 | A/Nevada/17/2015(H3N2) | 18/2/2015 | NCBI |
| KT842225 | A/Wisconsin/14/2015(H3N2) | 10/2/2015 | NCBI |
| KT842133 | A/Washington/05/2015(H3N2) | 6/2/2015 | NCBI |

| | | | |
|---|---|---|---|
| KT842508 | A/Texas/16/2015(H3N2) | 4/2/2015 | NCBI |
| KT842403 | A/Delaware/18/2015(H3N2) | 23/2/2015 | NCBI |
| KT842559 | A/SouthCarolina/07/2015(H3N2) | 5/2/2015 | NCBI |
| KT842159 | A/Florida/31/2015(H3N2) | 5/2/2015 | NCBI |
| KT842158 | A/Hawaii/07/2015(H3N2) | 5/2/2015 | NCBI |
| KT842287 | A/Idaho/06/2015(H3N2) | 9/2/2015 | NCBI |
| KT842149 | A/Kentucky/11/2015(H3N2) | 17/2/2015 | NCBI |
| KT842253 | A/Virginia/04/2015(H3N2) | 4/2/2015 | NCBI |
| KT842171 | A/NewHampshire/12/2015(H3N2) | 1/2/2015 | NCBI |
| KT842521 | A/Wisconsin/20/2015(H3N2) | 24/2/2015 | NCBI |
| KT842705 | A/Wisconsin/13/2015(H3N2) | 22/2/2015 | NCBI |
| KT842186 | A/California/73/2015(H3N2) | 20/2/2015 | NCBI |
| KT842409 | A/Arkansas/06/2015(H3N2) | 17/2/2015 | NCBI |
| KT842393 | A/Delaware/13/2015(H3N2) | 24/2/2015 | NCBI |
| KT842474 | A/NewMexico/06/2015(H3N2) | 27/2/2015 | NCBI |
| KT842214 | A/California/47/2015(H3N2) | 19/2/2015 | NCBI |
| KT842209 | A/Wisconsin/15/2015(H3N2) | 3/2/2015 | NCBI |
| KT842382 | A/NewHampshire/18/2015(H3N2) | 11/2/2015 | NCBI |
| KT842395 | A/NewJersey/11/2015(H3N2) | 21/2/2015 | NCBI |
| KT842311 | A/NewMexico/10/2015(H3N2) | 7/2/2015 | NCBI |
| KT842215 | A/Texas/18/2015(H3N2) | 8/2/2015 | NCBI |
| KT842230 | A/Wisconsin/08/2015(H3N2) | 6/2/2015 | NCBI |
| KT842500 | A/Oregon/04/2015(H3N2) | 14/2/2015 | NCBI |
| KT842506 | A/Nevada/13/2015(H3N2) | 11/2/2015 | NCBI |
| KT842292 | A/Oklahoma/06/2015(H3N2) | 11/2/2015 | NCBI |
| KT842223 | A/Texas/11/2015(H3N2) | 16/2/2015 | NCBI |
| KT842425 | A/Washington/12/2015(H3N2) | 18/2/2015 | NCBI |
| KT842463 | A/Alabama/08/2015(H3N2) | 14/2/2015 | NCBI |
| KT842246 | A/Nevada/18/2015(H3N2) | 18/2/2015 | NCBI |
| KT842257 | A/NewYork/19/2015(H3N2) | 17/2/2015 | NCBI |
| KT842252 | A/Oklahoma/05/2015(H3N2) | 11/2/2015 | NCBI |
| KT842300 | A/Tennessee/11/2015(H3N2) | 3/2/2015 | NCBI |
| KT842261 | A/Washington/13/2015(H3N2) | 25/2/2015 | NCBI |
| KT842404 | A/Washington/16/2015(H3N2) | 13/2/2015 | NCBI |
| KT842297 | A/California/12/2015(H3N2) | 15/2/2015 | NCBI |
| KT842363 | A/Iowa/09/2015(H3N2) | 3/2/2015 | NCBI |
| KT842280 | A/Minnesota/23/2015(H3N2) | 19/2/2015 | NCBI |

| | | | |
|---|---|---|---|
| KT842328 | A/Utah/11/2015(H3N2) | 20/2/2015 | NCBI |
| KT842430 | A/Wisconsin/09/2015(H3N2) | 18/2/2015 | NCBI |
| KT842465 | A/Wisconsin/07/2015(H3N2) | 13/2/2015 | NCBI |
| KT842498 | A/NewHampshire/13/2015(H3N2) | 18/2/2015 | NCBI |
| KT842339 | A/Mississippi/10/2015(H3N2) | 23/2/2015 | NCBI |
| KT842345 | A/NewJersey/05/2015(H3N2) | 23/2/2015 | NCBI |
| KT842350 | A/Michigan/27/2015(H3N2) | 24/2/2015 | NCBI |
| KT842786 | A/Tennessee/07/2015(H3N2) | 2/2/2015 | NCBI |
| KT842575 | A/California/16/2015(H3N2) | 12/2/2015 | NCBI |
| KT842646 | A/Tennessee/09/2015(H3N2) | 2/2/2015 | NCBI |
| KT842670 | A/Tennessee/08/2015(H3N2) | 3/2/2015 | NCBI |
| KT842604 | A/Hawaii/12/2015(H3N2) | 10/2/2015 | NCBI |
| KT842613 | A/Hawaii/14/2015(H3N2) | 17/2/2015 | NCBI |
| KT842621 | A/Hawaii/15/2015(H3N2) | 15/2/2015 | NCBI |
| KT842629 | A/Hawaii/23/2015(H3N2) | 25/2/2015 | NCBI |
| KT842798 | A/Ohio/10/2015(H3N2) | 21/3/2015 | NCBI |
| KT843283 | A/SouthDakota/34/2015(H3N2) | 11/3/2015 | NCBI |
| KT843241 | A/Idaho/12/2015(H3N2) | 13/3/2015 | NCBI |
| KT843317 | A/NewJersey/14/2015(H3N2) | 12/3/2015 | NCBI |
| KT843328 | A/NewJersey/15/2015(H3N2) | 2/3/2015 | NCBI |
| KT843199 | A/Wisconsin/21/2015(H3N2) | 9/3/2015 | NCBI |
| KT842804 | A/Kentucky/06/2015(H3N2) | 22/3/2015 | NCBI |
| KT843212 | A/NorthCarolina/15/2015(H3N2) | 17/3/2015 | NCBI |
| KT842877 | A/Nevada/21/2015(H3N2) | 2/3/2015 | NCBI |
| KT842874 | A/Washington/21/2015(H3N2) | 6/3/2015 | NCBI |
| KT842873 | A/Maine/18/2015(H3N2) | 3/3/2015 | NCBI |
| KT842810 | A/SouthDakota/15/2015(H3N2) | 5/3/2015 | NCBI |
| KT842811 | A/NewHampshire/16/2015(H3N2) | 3/3/2015 | NCBI |
| KT842812 | A/NorthDakota/06/2015(H3N2) | 8/3/2015 | NCBI |
| KT842892 | A/Florida/27/2015(H3N2) | 5/3/2015 | NCBI |
| KT843271 | A/Montana/07/2015(H3N2) | 7/3/2015 | NCBI |
| KT842830 | A/NewJersey/13/2015(H3N2) | 21/3/2015 | NCBI |
| KT843246 | A/SouthDakota/35/2015(H3N2) | 14/3/2015 | NCBI |
| KT843024 | A/Ohio/11/2015(H3N2) | 20/3/2015 | NCBI |
| KT842822 | A/Delaware/22/2015(H3N2) | 8/3/2015 | NCBI |
| KT843150 | A/Indiana/10/2015(H3N2) | 3/3/2015 | NCBI |
| KT842831 | A/Wisconsin/24/2015(H3N2) | 9/3/2015 | NCBI |

| | | | |
|---|---|---|---|
| KT843101 | A/Texas/19/2015(H3N2) | 2/3/2015 | NCBI |
| KT843266 | A/NewMexico/12/2015(H3N2) | 10/3/2015 | NCBI |
| KT842849 | A/RhodeIsland/12/2015(H3N2) | 30/3/2015 | NCBI |
| KT842834 | A/Hawaii/08/2015(H3N2) | 5/3/2015 | NCBI |
| KT842858 | A/Maine/17/2015(H3N2) | 2/3/2015 | NCBI |
| KT842876 | A/Texas/43/2015(H3N2) | 28/3/2015 | NCBI |
| KT843209 | A/Texas/36/2015(H3N2) | 8/3/2015 | NCBI |
| KT842854 | A/Maine/20/2015(H3N2) | 11/3/2015 | NCBI |
| KT843208 | A/NorthCarolina/16/2015(H3N2) | 15/3/2015 | NCBI |
| KT842862 | A/SouthDakota/11/2015(H3N2) | 31/3/2015 | NCBI |
| KT843247 | A/Vermont/05/2015(H3N2) | 13/3/2015 | NCBI |
| KT843303 | A/Florida/26/2015(H3N2) | 17/3/2015 | NCBI |
| KT843340 | A/Texas/37/2015(H3N2) | 3/3/2015 | NCBI |
| KT843126 | A/Kansas/04/2015(H3N2) | 3/3/2015 | NCBI |
| KT843293 | A/Oregon/05/2015(H3N2) | 8/3/2015 | NCBI |
| KT843113 | A/Wisconsin/16/2015(H3N2) | 4/3/2015 | NCBI |
| KT842921 | A/Hawaii/27/2015(H3N2) | 28/3/2015 | NCBI |
| KT843013 | A/Washington/33/2015(H3N2) | 12/3/2015 | NCBI |
| KT842976 | A/Washington/35/2015(H3N2) | 18/3/2015 | NCBI |
| KT843000 | A/Delaware/35/2015(H3N2) | 2/3/2015 | NCBI |
| KT842933 | A/Maryland/12/2015(H3N2) | 2/3/2015 | NCBI |
| KT843014 | A/Idaho/17/2015(H3N2) | 14/3/2015 | NCBI |
| KT843181 | A/Idaho/17/2015(H3N2) | 14/3/2015 | NCBI |
| KT843178 | A/NewMexico/18/2015(H3N2) | 18/3/2015 | NCBI |
| KT843145 | A/RhodeIsland/13/2015(H3N2) | 2/3/2015 | NCBI |
| KT842948 | A/Massachusetts/16/2015(H3N2) | 5/3/2015 | NCBI |
| KT843006 | A/California/81/2015(H3N2) | 7/3/2015 | NCBI |
| KT843184 | A/Texas/61/2015(H3N2) | 8/3/2015 | NCBI |
| KT842999 | A/California/78/2015(H3N2) | 3/3/2015 | NCBI |
| KT843007 | A/Washington/34/2015(H3N2) | 16/3/2015 | NCBI |
| KT842985 | A/Iowa/14/2015(H3N2) | 12/3/2015 | NCBI |
| KT843278 | A/Idaho/18/2015(H3N2) | 27/3/2015 | NCBI |
| KT843098 | A/NewJersey/06/2015(H3N2) | 2/3/2015 | NCBI |
| KT843042 | A/Iowa/11/2015(H3N2) | 5/3/2015 | NCBI |
| KT843046 | A/California/24/2015(H3N2) | 3/3/2015 | NCBI |
| KT843031 | A/Oregon/06/2015(H3N2) | 4/3/2015 | NCBI |
| KT843055 | A/NewJersey/16/2015(H3N2) | 3/3/2015 | NCBI |

| | | | |
|---|---|---|---|
| KT843119 | A/Vermont/06/2015(H3N2) | 8/3/2015 | NCBI |
| KT843096 | A/Maine/16/2015(H3N2) | 2/3/2015 | NCBI |
| KT843063 | A/Massachusetts/18/2015(H3N2) | 29/3/2015 | NCBI |
| KT843071 | A/Nebraska/16/2015(H3N2) | 23/3/2015 | NCBI |
| KT843079 | A/Maryland/13/2015(H3N2) | 4/3/2015 | NCBI |
| KT843086 | A/RhodeIsland/15/2015(H3N2) | 12/3/2015 | NCBI |
| KT843091 | A/Florida/43/2015(H3N2) | 31/3/2015 | NCBI |
| KT843159 | A/Washington/31/2015(H3N2) | 15/3/2015 | NCBI |
| KT843172 | A/Hawaii/24/2015(H3N2) | 4/3/2015 | NCBI |
| KT843478 | A/Minnesota/28/2015(H3N2) | 21/4/2015 | NCBI |
| KT843427 | A/Arizona/07/2015(H3N2) | 14/4/2015 | NCBI |
| KT843346 | A/SouthDakota/12/2015(H3N2) | 4/4/2015 | NCBI |
| KT843367 | A/NewJersey/49/2015(H3N2) | 14/4/2015 | NCBI |
| KT843528 | A/Hawaii/33/2015(H3N2) | 19/4/2015 | NCBI |
| KT843384 | A/Wisconsin/66/2015(H3N2) | 21/4/2015 | NCBI |
| KT843385 | A/Florida/49/2015(H3N2) | 26/4/2015 | NCBI |
| KT843359 | A/Hawaii/28/2015(H3N2) | 4/4/2015 | NCBI |
| KT843447 | A/Idaho/23/2015(H3N2) | 14/4/2015 | NCBI |
| KT843423 | A/Nevada/36/2015(H3N2) | 6/4/2015 | NCBI |
| KT843414 | A/NewMexico/19/2015(H3N2) | 1/4/2015 | NCBI |
| KT843433 | A/Alaska/10/2015(H3N2) | 12/4/2015 | NCBI |
| KT843370 | A/Minnesota/27/2015(H3N2) | 12/4/2015 | NCBI |
| KT843424 | A/California/82/2015(H3N2) | 28/4/2015 | NCBI |
| KT843499 | A/RhodeIsland/24/2015(H3N2) | 3/4/2015 | NCBI |
| KT843492 | A/Indiana/14/2015(H3N2) | 8/4/2015 | NCBI |
| KT843469 | A/Minnesota/25/2015(H3N2) | 1/4/2015 | NCBI |
| KT843387 | A/Minnesota/26/2015(H3N2) | 2/4/2015 | NCBI |
| KT843470 | A/Alaska/09/2015(H3N2) | 18/4/2015 | NCBI |
| KT843394 | A/Wisconsin/36/2015(H3N2) | 4/4/2015 | NCBI |
| KT843489 | A/Massachusetts/19/2015(H3N2) | 13/4/2015 | NCBI |
| KT843483 | A/Florida/45/2015(H3N2) | 18/4/2015 | NCBI |
| KT843434 | A/Hawaii/37/2015(H3N2) | 14/4/2015 | NCBI |
| KT843532 | A/Florida/46/2015(H3N2) | 18/4/2015 | NCBI |
| KT843519 | A/NewJersey/48/2015(H3N2) | 14/4/2015 | NCBI |
| KT844394 | A/Massachusetts/23/2015(H3N2) | 5/8/2015 | NCBI |
| KT843543 | A/Hawaii/44/2015(H3N2) | 9/7/2015 | NCBI |
| KT844383 | A/Minnesota/30/2015(H3N2) | 18/5/2015 | NCBI |

| | | | |
|---|---|---|---|
| KT844175 | A/Alaska/142/2015(H3N2) | 10/8/2015 | NCBI |
| KT843557 | A/Massachusetts/22/2015(H3N2) | 7/8/2015 | NCBI |
| KT844382 | A/Washington/37/2015(H3N2) | 6/5/2015 | NCBI |
| KT843941 | A/Alaska/66/2015(H3N2) | 24/6/2015 | NCBI |
| KT843552 | A/Florida/64/2015(H3N2) | 17/8/2015 | NCBI |
| KT843553 | A/Alaska/65/2015(H3N2) | 26/6/2015 | NCBI |
| KT844183 | A/Alaska/143/2015(H3N2) | 6/8/2015 | NCBI |
| KT844384 | A/Florida/50/2015(H3N2) | 3/6/2015 | NCBI |
| KT844372 | A/Virginia/32/2015(H3N2) | 17/6/2015 | NCBI |
| KT843576 | A/NewMexico/22/2015(H3N2) | 27/7/2015 | NCBI |
| KT843597 | A/Ohio/20/2015(H3N2) | 23/7/2015 | NCBI |
| KT843584 | A/Iowa/26/2015(H3N2) | 30/7/2015 | NCBI |
| KT843592 | A/Iowa/34/2015(H3N2) | 30/7/2015 | NCBI |
| KT844191 | A/Alaska/144/2015(H3N2) | 3/8/2015 | NCBI |
| KT843606 | A/Iowa/20/2015(H3N2) | 25/7/2015 | NCBI |
| KT843603 | A/Alaska/68/2015(H3N2) | 25/6/2015 | NCBI |
| KT844045 | A/Ohio/21/2015(H3N2) | 29/7/2015 | NCBI |
| KT843613 | A/Alaska/69/2015(H3N2) | 26/6/2015 | NCBI |
| KT843673 | A/Oregon/18/2015(H3N2) | 18/7/2015 | NCBI |
| KT843621 | A/Alaska/14/2015(H3N2) | 3/6/2015 | NCBI |
| KT843963 | A/NorthCarolina/26/2015(H3N2) | 30/6/2015 | NCBI |
| KT844265 | A/Alaska/17/2015(H3N2) | 1/6/2015 | NCBI |
| KT843923 | A/NorthCarolina/29/2015(H3N2) | 17/8/2015 | NCBI |
| KT844215 | A/Alaska/147/2015(H3N2) | 24/7/2015 | NCBI |
| KT843656 | A/Iowa/30/2015(H3N2) | 30/7/2015 | NCBI |
| KT843650 | A/Hawaii/47/2015(H3N2) | 5/7/2015 | NCBI |
| KT844061 | A/Arizona/10/2015(H3N2) | 1/8/2015 | NCBI |
| KT843655 | A/Alaska/26/2015(H3N2) | 14/6/2015 | NCBI |
| KT843651 | A/Florida/52/2015(H3N2) | 22/6/2015 | NCBI |
| KT843667 | A/Hawaii/46/2015(H3N2) | 7/7/2015 | NCBI |
| KT843682 | A/Ohio/23/2015(H3N2) | 3/8/2015 | NCBI |
| KT843690 | A/Ohio/24/2015(H3N2) | 3/8/2015 | NCBI |
| KT843699 | A/Iowa/37/2015(H3N2) | 30/7/2015 | NCBI |
| KT843707 | A/Iowa/38/2015(H3N2) | 30/7/2015 | NCBI |
| KT843715 | A/NewYork/57/2015(H3N2) | 21/7/2015 | NCBI |
| KT843723 | A/NewYork/58/2015(H3N2) | 3/8/2015 | NCBI |
| KT843731 | A/NewYork/59/2015(H3N2) | 29/7/2015 | NCBI |

| | | | |
|---|---|---|---|
| KT843739 | A/NewYork/60/2015(H3N2) | 1/8/2015 | NCBI |
| KT843747 | A/NewYork/62/2015(H3N2) | 4/8/2015 | NCBI |
| KT843755 | A/NewYork/63/2015(H3N2) | 6/8/2015 | NCBI |
| KT843763 | A/NewYork/64/2015(H3N2) | 10/8/2015 | NCBI |
| KT843951 | A/Wisconsin/74/2015(H3N2) | 1/9/2015 | NCBI |
| KT844044 | A/Iowa/24/2015(H3N2) | 30/7/2015 | NCBI |
| KT843776 | A/Alaska/29/2015(H3N2) | 18/6/2015 | NCBI |
| KT843797 | A/Hawaii/45/2015(H3N2) | 8/6/2015 | NCBI |
| KT843784 | A/Alaska/30/2015(H3N2) | 19/6/2015 | NCBI |
| KT844038 | A/Alaska/31/2015(H3N2) | 20/6/2015 | NCBI |
| KT843801 | A/Alaska/33/2015(H3N2) | 19/6/2015 | NCBI |
| KT843809 | A/Alaska/34/2015(H3N2) | 19/6/2015 | NCBI |
| KT843817 | A/Alaska/35/2015(H3N2) | 17/6/2015 | NCBI |
| KT843825 | A/Alaska/36/2015(H3N2) | 16/6/2015 | NCBI |
| KT843833 | A/Alaska/37/2015(H3N2) | 21/6/2015 | NCBI |
| KT843841 | A/Alaska/38/2015(H3N2) | 21/6/2015 | NCBI |
| KT844331 | A/Alaska/13/2015(H3N2) | 5/6/2015 | NCBI |
| KT843979 | A/Florida/55/2015(H3N2) | 4/7/2015 | NCBI |
| KT844339 | A/Alaska/18/2015(H3N2) | 1/6/2015 | NCBI |
| KT843857 | A/Florida/59/2015(H3N2) | 9/8/2015 | NCBI |
| KT843865 | A/Florida/61/2015(H3N2) | 19/8/2015 | NCBI |
| KT843873 | A/Florida/62/2015(H3N2) | 18/8/2015 | NCBI |
| KT843881 | A/Florida/63/2015(H3N2) | 18/8/2015 | NCBI |
| KT843899 | A/Ohio/26/2015(H3N2) | 17/8/2015 | NCBI |
| KT843907 | A/Montana/16/2015(H3N2) | 8/8/2015 | NCBI |
| KT843915 | A/Montana/17/2015(H3N2) | 13/8/2015 | NCBI |
| KT843955 | A/Michigan/38/2015(H3N2) | 12/6/2015 | NCBI |
| KT843931 | A/NorthCarolina/28/2015(H3N2) | 20/8/2015 | NCBI |
| KT843938 | A/Wisconsin/69/2015(H3N2) | 1/7/2015 | NCBI |
| KT843986 | A/Illinois/12/2015(H3N2) | 30/6/2015 | NCBI |
| KT843973 | A/Wisconsin/75/2015(H3N2) | 1/9/2015 | NCBI |
| KT843987 | A/Michigan/37/2015(H3N2) | 27/6/2015 | NCBI |
| KT843991 | A/Iowa/19/2015(H3N2) | 22/7/2015 | NCBI |
| KT843995 | A/NewJersey/50/2015(H3N2) | 5/7/2015 | NCBI |
| KT844007 | A/Iowa/25/2015(H3N2) | 30/7/2015 | NCBI |
| KT844015 | A/Iowa/28/2015(H3N2) | 30/7/2015 | NCBI |
| KT844051 | A/Washington/36/2015(H3N2) | 10/5/2015 | NCBI |

| | | | |
|---|---|---|---|
| KT844239 | A/Alaska/150/2015(H3N2) | 19/7/2015 | NCBI |
| KT844069 | A/Arizona/11/2015(H3N2) | 14/7/2015 | NCBI |
| KT844335 | A/Hawaii/43/2015(H3N2) | 8/7/2015 | NCBI |
| KT844085 | A/Texas/65/2015(H3N2) | 6/8/2015 | NCBI |
| KT844093 | A/Iowa/22/2015(H3N2) | 30/7/2015 | NCBI |
| KT844101 | A/Iowa/23/2015(H3N2) | 30/7/2015 | NCBI |
| KT844109 | A/Iowa/29/2015(H3N2) | 30/7/2015 | NCBI |
| KT844117 | A/Iowa/31/2015(H3N2) | 30/7/2015 | NCBI |
| KT844127 | A/Alaska/135/2015(H3N2) | 2/8/2015 | NCBI |
| KT844135 | A/Alaska/137/2015(H3N2) | 2/8/2015 | NCBI |
| KT844143 | A/Alaska/138/2015(H3N2) | 3/8/2015 | NCBI |
| KT844151 | A/Alaska/139/2015(H3N2) | 4/8/2015 | NCBI |
| KT844159 | A/Alaska/140/2015(H3N2) | 11/8/2015 | NCBI |
| KT844167 | A/Alaska/141/2015(H3N2) | 5/8/2015 | NCBI |
| KT844199 | A/Alaska/145/2015(H3N2) | 24/7/2015 | NCBI |
| KT844207 | A/Alaska/146/2015(H3N2) | 23/7/2015 | NCBI |
| KT844244 | A/Florida/57/2015(H3N2) | 26/7/2015 | NCBI |
| KT844223 | A/Alaska/148/2015(H3N2) | 26/7/2015 | NCBI |
| KT844231 | A/Alaska/149/2015(H3N2) | 7/7/2015 | NCBI |
| KT844252 | A/Florida/58/2015(H3N2) | 25/7/2015 | NCBI |
| KT844260 | A/Washington/39/2015(H3N2) | 14/7/2015 | NCBI |
| KT844280 | A/Hawaii/34/2015(H3N2) | 21/5/2015 | NCBI |
| KT844288 | A/Hawaii/35/2015(H3N2) | 26/5/2015 | NCBI |
| KT844296 | A/Hawaii/36/2015(H3N2) | 11/6/2015 | NCBI |
| KT844324 | A/Minnesota/29/2015(H3N2) | 16/5/2015 | NCBI |
| KT844307 | A/Oregon/16/2015(H3N2) | 15/7/2015 | NCBI |
| KT844316 | A/Florida/51/2015(H3N2) | 9/6/2015 | NCBI |
| KT844347 | A/Alaska/21/2015(H3N2) | 8/6/2015 | NCBI |
| KT844356 | A/Alaska/25/2015(H3N2) | 17/6/2015 | NCBI |
| KT844366 | A/Arizona/09/2015(H3N2) | 21/6/2015 | NCBI |
| KT880082 | A/Wisconsin/20/2015(H3N2) | 24/2/2015 | NCBI |
| KM654789 | A/Connecticut/Flu056/2012(H3N2) | 30/12/2012 | NCBI |
| KM654790 | A/Connecticut/Flu066/2012(H3N2) | 30/12/2012 | NCBI |
| KM654793 | A/Connecticut/Flu086/2012(H3N2) | 31/12/2012 | NCBI |
| KM654811 | A/Connecticut/Flu055/2012(H3N2) | 27/12/2012 | NCBI |
| KM654816 | A/Connecticut/Flu090/2013(H3N2) | 1/1/2013 | NCBI |
| KM654828 | A/Connecticut/Flu166/2013(H3N2) | 26/1/2013 | NCBI |

| | | | |
|---|---|---|---|
| KU198899 | A/Moscow/CRIE-141/2015(H3N2) | 27/1/2015 | NCBI |
| KP335870 | A/Thailand/CU-B4929/2011(H3N2) | 18/7/2011 | NCBI |
| KP335872 | A/Thailand/CU-B5105/2011(H3N2) | 3/8/2011 | NCBI |
| KP335877 | A/Thailand/CU-B5614/2011(H3N2) | 5/9/2011 | NCBI |
| KP335879 | A/Thailand/CU-B5697/2011(H3N2) | 16/9/2011 | NCBI |
| KP335882 | A/Thailand/CU-B5848/2011(H3N2) | 17/10/2011 | NCBI |
| KP335883 | A/Thailand/CU-B5873/2011(H3N2) | 24/10/2011 | NCBI |
| KP335886 | A/Thailand/CU-B5909/2011(H3N2) | 3/11/2011 | NCBI |
| KP335887 | A/Thailand/CU-B5928/2011(H3N2) | 15/11/2011 | NCBI |
| KP335888 | A/Thailand/CU-C2417/2011(H3N2) | 8/12/2011 | NCBI |
| KP335873 | A/Thailand/CU-H3020/2011(H3N2) | 15/8/2011 | NCBI |
| KP335890 | A/Thailand/CU-B6091/2012(H3N2) | 23/2/2012 | NCBI |
| KP335891 | A/Thailand/CU-B6251/2012(H3N2) | 6/6/2012 | NCBI |
| KP335892 | A/Thailand/CU-B6274/2012(H3N2) | 22/6/2012 | NCBI |
| KP335893 | A/Thailand/CU-B6309/2012(H3N2) | 2/7/2012 | NCBI |
| KP335896 | A/Thailand/CU-B6936/2012(H3N2) | 13/9/2012 | NCBI |
| KP335902 | A/Thailand/CU-B7235/2012(H3N2) | 26/11/2012 | NCBI |
| KP335904 | A/Thailand/CU-B7367/2012(H3N2) | 22/12/2012 | NCBI |
| KP335889 | A/Thailand/CU-H3141/2012(H3N2) | 9/1/2012 | NCBI |
| KP335895 | A/Thailand/CU-H3368/2012(H3N2) | 31/8/2012 | NCBI |
| KP335899 | A/Thailand/CU-H3453/2012(H3N2) | 16/10/2012 | NCBI |
| KP335907 | A/Thailand/CU-B7536/2013(H3N2) | 28/1/2013 | NCBI |
| KP335908 | A/Thailand/CU-B7585/2013(H3N2) | 2/2/2013 | NCBI |
| KP335912 | A/Thailand/CU-B7765/2013(H3N2) | 18/3/2013 | NCBI |
| KP335918 | A/Thailand/CU-A114/2013(H3N2) | 4/6/2013 | NCBI |
| KP335920 | A/Thailand/CU-A134/2013(H3N2) | 20/6/2013 | NCBI |
| KP335922 | A/Thailand/CU-A166/2013(H3N2) | 10/7/2013 | NCBI |
| KP335925 | A/Thailand/CU-A182/2013(H3N2) | 18/7/2013 | NCBI |
| KP335928 | A/Thailand/CU-A305/2013(H3N2) | 29/8/2013 | NCBI |
| KP335939 | A/Thailand/CU-A459/2013(H3N2) | 29/10/2013 | NCBI |
| KP335914 | A/Thailand/CU-B7885/2013(H3N2) | 22/4/2013 | NCBI |
| KP335926 | A/Thailand/CU-B8121/2013(H3N2) | 2/8/2013 | NCBI |
| KP335927 | A/Thailand/CU-B8127/2013(H3N2) | 5/8/2013 | NCBI |
| KP335933 | A/Thailand/CU-B8364/2013(H3N2) | 23/9/2013 | NCBI |
| KP335934 | A/Thailand/CU-B8518/2013(H3N2) | 2/10/2013 | NCBI |
| KP335940 | A/Thailand/CU-B8736/2013(H3N2) | 14/11/2013 | NCBI |
| KP335943 | A/Thailand/CU-B8772/2013(H3N2) | 11/12/2013 | NCBI |

| | | | |
|---|---|---|---|
| KP335930 | A/Thailand/CU-C4087/2013(H3N2) | 4/9/2013 | NCBI |
| KP335945 | A/Thailand/CU-C4364/2013(H3N2) | 18/12/2013 | NCBI |
| KP335936 | A/Thailand/CU-H3567/2013(H3N2) | 10/10/2013 | NCBI |
| KP335947 | A/Thailand/CU-B8849/2014(H3N2) | 7/1/2014 | NCBI |
| KP335959 | A/Thailand/CU-B10282/2014(H3N2) | 16/6/2014 | NCBI |
| KP335961 | A/Thailand/CU-B10345/2014(H3N2) | 9/7/2014 | NCBI |
| KP335964 | A/Thailand/CU-B10509/2014(H3N2) | 4/8/2014 | NCBI |
| KP335965 | A/Thailand/CU-B10521/2014(H3N2) | 5/8/2014 | NCBI |
| KP335971 | A/Thailand/CU-B10828/2014(H3N2) | 10/9/2014 | NCBI |
| KP335948 | A/Thailand/CU-C4406/2014(H3N2) | 7/1/2014 | NCBI |
| KP335952 | A/Thailand/CU-C4546/2014(H3N2) | 12/2/2014 | NCBI |
| KP335956 | A/Thailand/CU-CB166/2014(H3N2) | 1/4/2014 | NCBI |
| KP335950 | A/Thailand/CU-H3595/2014(H3N2) | 4/2/2014 | NCBI |
| KP335955 | A/Thailand/CU-H3626/2014(H3N2) | 7/3/2014 | NCBI |
| KP335960 | A/Thailand/CU-H3649/2014(H3N2) | 9/7/2014 | NCBI |
| KP335973 | A/Thailand/CU-B10975/2014(H3N2) | 6/10/2014 | NCBI |
| KP335975 | A/Thailand/CU-B11065/2014(H3N2) | 14/10/2014 | NCBI |
| KP335977 | A/Thailand/CU-B11202/2014(H3N2) | 1/11/2014 | NCBI |
| KP335978 | A/Thailand/CU-H3680/2014(H3N2) | 13/11/2014 | NCBI |
| KP638230 | A/Thailand/SS07134/2008(H3N2) | 21/10/2008 | NCBI |
| KP638222 | A/Thailand/SS07028/2007(H3N2) | 26/9/2007 | NCBI |
| KP638214 | A/Thailand/SR08183/2011(H3N2) | 17/6/2011 | NCBI |
| KP638206 | A/Thailand/SP11679/2012(H3N2) | 13/7/2012 | NCBI |
| KP638198 | A/Thailand/MR11632/2012(H3N2) | 8/8/2012 | NCBI |
| KP638190 | A/Thailand/MR10547/2011(H3N2) | 22/8/2011 | NCBI |
| KP638182 | A/Thailand/MR10462/2011(H3N2) | 4/2/2011 | NCBI |
| KP638174 | A/Thailand/MR08398/2010(H3N2) | 1/9/2010 | NCBI |
| KP638166 | A/Thailand/MR07161/2008(H3N2) | 30/7/2008 | NCBI |
| KP638158 | A/Thailand/MR07159/2008(H3N2) | 29/7/2008 | NCBI |
| KP638150 | A/Thailand/SP07127/2008(H3N2) | 23/6/2008 | NCBI |
| KP638142 | A/Thailand/SP07043/2007(H3N2) | 10/9/2007 | NCBI |
| KP638134 | A/Thailand/SN11893/2013(H3N2) | 23/1/2013 | NCBI |
| KP638126 | A/Thailand/SN11792/2012(H3N2) | 30/7/2012 | NCBI |
| KP638118 | A/Thailand/SN11650/2011(H3N2) | 19/9/2011 | NCBI |
| KP638110 | A/Thailand/SN10452/2010(H3N2) | 7/10/2010 | NCBI |
| KP638102 | A/Thailand/SN07141/2008(H3N2) | 23/7/2008 | NCBI |
| KP638094 | A/Thailand/SN07100/2007(H3N2) | 21/12/2007 | NCBI |

| | | | |
|---|---|---|---|
| KP638086 | A/Thailand/PY12558/2013(H3N2) | 11/2/2013 | NCBI |
| KP638078 | A/Thailand/PY10316/2011(H3N2) | 9/8/2011 | NCBI |
| KP638070 | A/Thailand/PY08156/2010(H3N2) | 17/8/2010 | NCBI |
| KP638062 | A/Thailand/PY08064/2009(H3N2) | 12/10/2009 | NCBI |
| KP638054 | A/Thailand/MR07094/2007(H3N2) | 14/11/2007 | NCBI |
| KP638046 | A/Thailand/MR07011/2007(H3N2) | 28/3/2007 | NCBI |
| KP638038 | A/Thailand/KS10537/2013(H3N2) | 19/2/2013 | NCBI |
| KP638030 | A/Thailand/KS10532/2013(H3N2) | 8/2/2013 | NCBI |
| KP638022 | A/Thailand/KS10525/2012(H3N2) | 7/12/2012 | NCBI |
| KP638014 | A/Thailand/KS10521/2012(H3N2) | 16/11/2012 | NCBI |
| KP638006 | A/Thailand/KS10428/2011(H3N2) | 13/9/2011 | NCBI |
| KP637998 | A/Thailand/KS10406/2011(H3N2) | 25/7/2011 | NCBI |
| KP637990 | A/Thailand/KS08398/2010(H3N2) | 14/10/2010 | NCBI |
| KP637982 | A/Thailand/KS08384/2010(H3N2) | 17/8/2010 | NCBI |
| KP637974 | A/Thailand/KS08309/2009(H3N2) | 29/10/2009 | NCBI |
| KP637966 | A/Thailand/KS08239/2009(H3N2) | 6/3/2009 | NCBI |
| KP637958 | A/Thailand/KS08204/2008(H3N2) | 13/10/2008 | NCBI |
| KP637950 | A/Thailand/KS07193/2008(H3N2) | 12/6/2008 | NCBI |
| KP637942 | A/Thailand/KS07111/2007(H3N2) | 27/11/2007 | NCBI |
| KP637934 | A/Thailand/KS07007/2007(H3N2) | 4/5/2007 | NCBI |
| KP637926 | A/Thailand/IY07007/2007(H3N2) | 3/5/2007 | NCBI |
| KP637918 | A/Thailand/IY07006/2007(H3N2) | 3/5/2007 | NCBI |
| KU529244 | A/Singapore/H2013.238/2013(H3N2) | 28/3/2013 | NCBI |
| KU529245 | A/Singapore/H2013.403/2013(H3N2) | 27/5/2013 | NCBI |
| KU529246 | A/Singapore/H2013.422d/2013(H3N2) | 3/6/2013 | NCBI |
| KU529247 | A/Singapore/H2013.721a/2013(H3N2) | 20/9/2013 | NCBI |
| KU529248 | A/Singapore/H2013.721d/2013(H3N2) | 20/9/2013 | NCBI |
| KU529249 | A/Singapore/H2013.721f/2013(H3N2) | 20/9/2013 | NCBI |
| KU529250 | A/Singapore/H2013.729/2013(H3N2) | 23/9/2013 | NCBI |
| KU589428 | A/Alaska/46/2015(H3N2) | 21/6/2015 | NCBI |
| KU591314 | A/Missouri/42/2015(H3N2) | 4/10/2015 | NCBI |
| KU589438 | A/Oregon/17/2015(H3N2) | 1/7/2015 | NCBI |
| KU589447 | A/Florida/60/2015(H3N2) | 9/8/2015 | NCBI |
| KU589456 | A/Tennessee/18/2015(H3N2) | 18/8/2015 | NCBI |
| KU589472 | A/Tennessee/20/2015(H3N2) | 19/8/2015 | NCBI |
| KU591376 | A/Pennsylvania/40/2015(H3N2) | 22/10/2015 | NCBI |
| KU589485 | A/NewHampshire/38/2015(H3N2) | 2/9/2015 | NCBI |

| | | | |
|---|---|---|---|
| KU589493 | A/NewYork/65/2015(H3N2) | 25/8/2015 | NCBI |
| KU589502 | A/NewMexico/26/2015(H3N2) | 13/9/2015 | NCBI |
| KU589510 | A/Alaska/179/2015(H3N2) | 16/8/2015 | NCBI |
| KU589521 | A/Wisconsin/76/2015(H3N2) | 1/9/2015 | NCBI |
| KU589529 | A/Michigan/42/2015(H3N2) | 18/8/2015 | NCBI |
| KU589534 | A/Alaska/188/2015(H3N2) | 15/8/2015 | NCBI |
| KU589541 | A/Alaska/189/2015(H3N2) | 13/8/2015 | NCBI |
| KU589549 | A/Alaska/191/2015(H3N2) | 18/8/2015 | NCBI |
| KU589556 | A/Alaska/192/2015(H3N2) | 16/8/2015 | NCBI |
| KU589564 | A/Alaska/193/2015(H3N2) | 17/8/2015 | NCBI |
| KU589936 | A/Michigan/53/2015(H3N2) | 12/9/2015 | NCBI |
| KU589580 | A/Michigan/54/2015(H3N2) | 19/9/2015 | NCBI |
| KU589952 | A/Michigan/55/2015(H3N2) | 18/9/2015 | NCBI |
| KU590413 | A/Idaho/25/2015(H3N2) | 18/9/2015 | NCBI |
| KU590617 | A/Vermont/08/2015(H3N2) | 19/8/2015 | NCBI |
| KU591292 | A/Florida/80/2015(H3N2) | 28/10/2015 | NCBI |
| KU589623 | A/Alaska/67/2015(H3N2) | 1/7/2015 | NCBI |
| KU589631 | A/Ohio/18/2015(H3N2) | 23/7/2015 | NCBI |
| KU589639 | A/Oregon/11/2015(H3N2) | 30/6/2015 | NCBI |
| KU591895 | A/Michigan/67/2015(H3N2) | 10/10/2015 | NCBI |
| KU589648 | A/Nebraska/17/2015(H3N2) | 15/7/2015 | NCBI |
| KU589656 | A/Utah/15/2015(H3N2) | 13/7/2015 | NCBI |
| KU589667 | A/Alaska/49/2015(H3N2) | 15/6/2015 | NCBI |
| KU589675 | A/Alaska/50/2015(H3N2) | 16/6/2015 | NCBI |
| KU590101 | A/Alaska/151/2015(H3N2) | 21/7/2015 | NCBI |
| KU590109 | A/Alaska/152/2015(H3N2) | 20/7/2015 | NCBI |
| KU589699 | A/Alaska/154/2015(H3N2) | 24/7/2015 | NCBI |
| KU590132 | A/Alaska/155/2015(H3N2) | 8/8/2015 | NCBI |
| KU590140 | A/Alaska/156/2015(H3N2) | 3/8/2015 | NCBI |
| KU590696 | A/Ohio/27/2015(H3N2) | 17/9/2015 | NCBI |
| KU590704 | A/Ohio/28/2015(H3N2) | 17/9/2015 | NCBI |
| KU590720 | A/Massachusetts/26/2015(H3N2) | 24/9/2015 | NCBI |
| KU590249 | A/NewMexico/23/2015(H3N2) | 12/9/2015 | NCBI |
| KU590256 | A/NewMexico/25/2015(H3N2) | 10/9/2015 | NCBI |
| KU590264 | A/Iowa/40/2015(H3N2) | 4/9/2015 | NCBI |
| KU590272 | A/Iowa/41/2015(H3N2) | 10/9/2015 | NCBI |
| KU589778 | A/Iowa/42/2015(H3N2) | 10/9/2015 | NCBI |

| | | | |
|---|---|---|---|
| KU590287 | A/Iowa/43/2015(H3N2) | 14/9/2015 | NCBI |
| KU590736 | A/Wyoming/18/2015(H3N2) | 22/9/2015 | NCBI |
| KU589801 | A/Massachusetts/25/2015(H3N2) | 10/9/2015 | NCBI |
| KU589809 | A/Indiana/17/2015(H3N2) | 31/8/2015 | NCBI |
| KU589817 | A/Indiana/19/2015(H3N2) | 1/9/2015 | NCBI |
| KU589824 | A/Hawaii/34/2015(H3N2) | 21/5/2015 | NCBI |
| KU591415 | A/Indiana/20/2015(H3N2) | 1/9/2015 | NCBI |
| KU589832 | A/Alaska/48/2015(H3N2) | 21/6/2015 | NCBI |
| KU589840 | A/NewYork/55/2015(H3N2) | 10/7/2015 | NCBI |
| KU591433 | A/Indiana/22/2015(H3N2) | 1/9/2015 | NCBI |
| KU589851 | A/Indiana/25/2015(H3N2) | 3/9/2015 | NCBI |
| KU589859 | A/NewHampshire/39/2015(H3N2) | 9/9/2015 | NCBI |
| KU589867 | A/NewHampshire/40/2015(H3N2) | 14/9/2015 | NCBI |
| KU590091 | A/Montana/23/2015(H3N2) | 17/9/2015 | NCBI |
| KU589875 | A/Minnesota/41/2015(H3N2) | 14/9/2015 | NCBI |
| KU589880 | A/Montana/25/2015(H3N2) | 19/9/2015 | NCBI |
| KU589883 | A/Florida/69/2015(H3N2) | 15/9/2015 | NCBI |
| KU591456 | A/Montana/26/2015(H3N2) | 17/9/2015 | NCBI |
| KU589891 | A/Florida/70/2015(H3N2) | 15/9/2015 | NCBI |
| KU589899 | A/Florida/71/2015(H3N2) | 15/9/2015 | NCBI |
| KU590728 | A/Kentucky/15/2015(H3N2) | 28/9/2015 | NCBI |
| KU589912 | A/Wisconsin/83/2015(H3N2) | 24/9/2015 | NCBI |
| KU591475 | A/Florida/72/2015(H3N2) | 15/9/2015 | NCBI |
| KU589920 | A/Wisconsin/84/2015(H3N2) | 1/10/2015 | NCBI |
| KU591492 | A/Wisconsin/81/2015(H3N2) | 18/9/2015 | NCBI |
| KU589928 | A/Michigan/52/2015(H3N2) | 14/9/2015 | NCBI |
| KU589931 | A/Wisconsin/82/2015(H3N2) | 17/9/2015 | NCBI |
| KU591508 | A/Tennessee/21/2015(H3N2) | 15/9/2015 | NCBI |
| KU589960 | A/Montana/28/2015(H3N2) | 24/9/2015 | NCBI |
| KU589968 | A/Montana/29/2015(H3N2) | 26/9/2015 | NCBI |
| KU590305 | A/Alaska/157/2015(H3N2) | 13/8/2015 | NCBI |
| KU589982 | A/Alaska/158/2015(H3N2) | 14/8/2015 | NCBI |
| KU589990 | A/Alaska/159/2015(H3N2) | 17/8/2015 | NCBI |
| KU589998 | A/Alaska/162/2015(H3N2) | 19/8/2015 | NCBI |
| KU590015 | A/Alaska/160/2015(H3N2) | 18/8/2015 | NCBI |
| KU590023 | A/Oregon/19/2015(H3N2) | 8/10/2015 | NCBI |
| KU590486 | A/Oregon/20/2015(H3N2) | 9/10/2015 | NCBI |

| | | | |
|---|---|---|---|
| KU590031 | A/Indiana/18/2015(H3N2) | 1/9/2015 | NCBI |
| KU590039 | A/Indiana/26/2015(H3N2) | 3/9/2015 | NCBI |
| KU590047 | A/Indiana/27/2015(H3N2) | 3/9/2015 | NCBI |
| KU590055 | A/Montana/18/2015(H3N2) | 11/9/2015 | NCBI |
| KU590117 | A/Alaska/153/2015(H3N2) | 3/8/2015 | NCBI |
| KU590147 | A/Indiana/24/2015(H3N2) | 3/9/2015 | NCBI |
| KU591483 | A/Wisconsin/80/2015(H3N2) | 14/9/2015 | NCBI |
| KU590321 | A/Alaska/208/2015(H3N2) | 2/9/2015 | NCBI |
| KU590342 | A/Florida/74/2015(H3N2) | 3/10/2015 | NCBI |
| KU590350 | A/Florida/75/2015(H3N2) | 1/10/2015 | NCBI |
| KU590365 | A/NorthCarolina/32/2015(H3N2) | 28/9/2015 | NCBI |
| KU590372 | A/Maine/39/2015(H3N2) | 7/10/2015 | NCBI |
| KU590380 | A/Maine/40/2015(H3N2) | 9/10/2015 | NCBI |
| KU590388 | A/Maine/41/2015(H3N2) | 12/10/2015 | NCBI |
| KU590396 | A/Missouri/41/2015(H3N2) | 4/10/2015 | NCBI |
| KU590398 | A/Minnesota/42/2015(H3N2) | 14/9/2015 | NCBI |
| KU590405 | A/Minnesota/43/2015(H3N2) | 17/9/2015 | NCBI |
| KU591425 | A/Indiana/21/2015(H3N2) | 1/9/2015 | NCBI |
| KU590494 | A/Oregon/21/2015(H3N2) | 9/10/2015 | NCBI |
| KU590502 | A/Oregon/22/2015(H3N2) | 9/10/2015 | NCBI |
| KU590510 | A/Oregon/23/2015(H3N2) | 12/10/2015 | NCBI |
| KU590518 | A/Oregon/24/2015(H3N2) | 12/10/2015 | NCBI |
| KU590526 | A/Alaska/253/2015(H3N2) | 2/10/2015 | NCBI |
| KU590534 | A/Alaska/254/2015(H3N2) | 5/10/2015 | NCBI |
| KU590542 | A/Alaska/255/2015(H3N2) | 6/10/2015 | NCBI |
| KU590550 | A/Alaska/256/2015(H3N2) | 5/10/2015 | NCBI |
| KU590563 | A/Washington/40/2015(H3N2) | 1/9/2015 | NCBI |
| KU590571 | A/Washington/41/2015(H3N2) | 31/8/2015 | NCBI |
| KU590579 | A/Washington/42/2015(H3N2) | 31/8/2015 | NCBI |
| KU590587 | A/Washington/43/2015(H3N2) | 31/8/2015 | NCBI |
| KU590595 | A/Washington/44/2015(H3N2) | 7/9/2015 | NCBI |
| KU590633 | A/Pennsylvania/36/2015(H3N2) | 2/10/2015 | NCBI |
| KU590606 | A/NewHampshire/41/2015(H3N2) | 10/10/2015 | NCBI |
| KU590625 | A/Vermont/09/2015(H3N2) | 30/9/2015 | NCBI |
| KU590855 | A/Texas/75/2015(H3N2) | 8/12/2015 | NCBI |
| KU590643 | A/Pennsylvania/50/2015(H3N2) | 23/11/2015 | NCBI |
| KU590652 | A/Alaska/40/2015(H3N2) | 19/6/2015 | NCBI |

| | | | |
|---|---|---|---|
| KU590660 | A/Alaska/44/2015(H3N2) | 21/6/2015 | NCBI |
| KU590668 | A/Alaska/45/2015(H3N2) | 23/6/2015 | NCBI |
| KU590676 | A/Oregon/14/2015(H3N2) | 29/6/2015 | NCBI |
| KU591379 | A/Alaska/183/2015(H3N2) | 16/8/2015 | NCBI |
| KU590687 | A/Tennessee/19/2015(H3N2) | 18/8/2015 | NCBI |
| KU590712 | A/Ohio/29/2015(H3N2) | 17/9/2015 | NCBI |
| KU591259 | A/Washington/45/2015(H3N2) | 16/6/2015 | NCBI |
| KU591821 | A/Alaska/222/2015(H3N2) | 8/9/2015 | NCBI |
| KU590770 | A/Alaska/225/2015(H3N2) | 11/9/2015 | NCBI |
| KU591853 | A/Alaska/239/2015(H3N2) | 16/9/2015 | NCBI |
| KU590787 | A/Hawaii/62/2015(H3N2) | 28/9/2015 | NCBI |
| KU590796 | A/Alaska/242/2015(H3N2) | 13/9/2015 | NCBI |
| KU590804 | A/Nebraska/19/2015(H3N2) | 23/10/2015 | NCBI |
| KU590812 | A/Florida/86/2015(H3N2) | 2/11/2015 | NCBI |
| KU590820 | A/California/94/2015(H3N2) | 20/10/2015 | NCBI |
| KU591700 | A/California/98/2015(H3N2) | 19/10/2015 | NCBI |
| KU590836 | A/Texas/71/2015(H3N2) | 11/11/2015 | NCBI |
| KU590845 | A/Washington/62/2015(H3N2) | 3/12/2015 | NCBI |
| KU591318 | A/Wisconsin/86/2015(H3N2) | 19/10/2015 | NCBI |
| KU590861 | A/Alaska/54/2015(H3N2) | 20/6/2015 | NCBI |
| KU590868 | A/Alaska/55/2015(H3N2) | 18/6/2015 | NCBI |
| KU590876 | A/Alaska/56/2015(H3N2) | 18/6/2015 | NCBI |
| KU590884 | A/Alaska/57/2015(H3N2) | 18/6/2015 | NCBI |
| KU590892 | A/Alaska/61/2015(H3N2) | 18/6/2015 | NCBI |
| KU590903 | A/Hawaii/58/2015(H3N2) | 15/9/2015 | NCBI |
| KU590911 | A/Hawaii/60/2015(H3N2) | 26/9/2015 | NCBI |
| KU590917 | A/Hawaii/61/2015(H3N2) | 28/9/2015 | NCBI |
| KU590924 | A/Texas/69/2015(H3N2) | 6/10/2015 | NCBI |
| KU590931 | A/Texas/70/2015(H3N2) | 6/10/2015 | NCBI |
| KU590942 | A/Alaska/116/2015(H3N2) | 6/7/2015 | NCBI |
| KU590951 | A/Mississippi/13/2015(H3N2) | 13/8/2015 | NCBI |
| KU590968 | A/NorthCarolina/33/2015(H3N2) | 16/10/2015 | NCBI |
| KU590976 | A/Delaware/41/2015(H3N2) | 14/10/2015 | NCBI |
| KU590983 | A/Arkansas/08/2015(H3N2) | 12/10/2015 | NCBI |
| KU590991 | A/Arkansas/09/2015(H3N2) | 12/10/2015 | NCBI |
| KU590999 | A/Maine/42/2015(H3N2) | 15/10/2015 | NCBI |
| KU591007 | A/Maine/43/2015(H3N2) | 19/10/2015 | NCBI |

| | | | |
|---|---|---|---|
| KU591015 | A/California/83/2015(H3N2) | 10/9/2015 | NCBI |
| KU591023 | A/California/85/2015(H3N2) | 9/9/2015 | NCBI |
| KU591031 | A/California/86/2015(H3N2) | 5/9/2015 | NCBI |
| KU591039 | A/California/87/2015(H3N2) | 1/10/2015 | NCBI |
| KU591047 | A/California/89/2015(H3N2) | 24/9/2015 | NCBI |
| KU591055 | A/California/91/2015(H3N2) | 1/10/2015 | NCBI |
| KU591063 | A/Wisconsin/85/2015(H3N2) | 16/10/2015 | NCBI |
| KU591071 | A/SouthCarolina/09/2015(H3N2) | 2015 | NCBI |
| KU591079 | A/Pennsylvania/42/2015(H3N2) | 19/10/2015 | NCBI |
| KU591087 | A/Florida/81/2015(H3N2) | 5/10/2015 | NCBI |
| KU591095 | A/Iowa/50/2015(H3N2) | 14/10/2015 | NCBI |
| KU591103 | A/WestVirginia/23/2015(H3N2) | 8/10/2015 | NCBI |
| KU591111 | A/Massachusetts/28/2015(H3N2) | 10/10/2015 | NCBI |
| KU591123 | A/Washington/49/2015(H3N2) | 4/10/2015 | NCBI |
| KU591131 | A/Washington/50/2015(H3N2) | 11/10/2015 | NCBI |
| KU591140 | A/Hawaii/65/2015(H3N2) | 4/10/2015 | NCBI |
| KU591147 | A/Hawaii/66/2015(H3N2) | 3/10/2015 | NCBI |
| KU591155 | A/Hawaii/67/2015(H3N2) | 5/10/2015 | NCBI |
| KU591163 | A/Hawaii/68/2015(H3N2) | 5/10/2015 | NCBI |
| KU591442 | A/Colorado/19/2015(H3N2) | 15/11/2015 | NCBI |
| KU591175 | A/Colorado/21/2015(H3N2) | 19/11/2015 | NCBI |
| KU591184 | A/Colorado/24/2015(H3N2) | 17/11/2015 | NCBI |
| KU591192 | A/Maryland/17/2015(H3N2) | 8/10/2015 | NCBI |
| KU591208 | A/Alaska/51/2015(H3N2) | 15/6/2015 | NCBI |
| KU591217 | A/Alaska/58/2015(H3N2) | 17/6/2015 | NCBI |
| KU591225 | A/Alaska/60/2015(H3N2) | 15/6/2015 | NCBI |
| KU591235 | A/Indiana/28/2015(H3N2) | 28/9/2015 | NCBI |
| KU591246 | A/Oklahoma/20/2015(H3N2) | 9/10/2015 | NCBI |
| KU592166 | A/Hawaii/74/2015(H3N2) | 9/10/2015 | NCBI |
| KU592179 | A/SouthDakota/37/2015(H3N2) | 11/8/2015 | NCBI |
| KU591268 | A/Alaska/257/2015(H3N2) | 11/10/2015 | NCBI |
| KU591276 | A/Alaska/258/2015(H3N2) | 12/10/2015 | NCBI |
| KU591284 | A/Alaska/259/2015(H3N2) | 20/10/2015 | NCBI |
| KU591300 | A/Michigan/62/2015(H3N2) | 29/9/2015 | NCBI |
| KU591308 | A/Colorado/16/2015(H3N2) | 21/10/2015 | NCBI |
| KU591326 | A/Washington/55/2015(H3N2) | 7/10/2015 | NCBI |
| KU591334 | A/Washington/56/2015(H3N2) | 20/10/2015 | NCBI |

| | | | |
|---|---|---|---|
| KU591350 | A/Maryland/18/2015(H3N2) | 29/10/2015 | NCBI |
| KU591358 | A/Maryland/19/2015(H3N2) | 1/11/2015 | NCBI |
| KU591366 | A/Louisiana/07/2015(H3N2) | 11/11/2015 | NCBI |
| KU591392 | A/Washington/51/2015(H3N2) | 9/10/2015 | NCBI |
| KU591405 | A/Alaska/185/2015(H3N2) | 20/8/2015 | NCBI |
| KU591399 | A/Massachusetts/14/2014(H3N2) | 19/11/2014 | NCBI |
| KU591464 | A/Montana/27/2015(H3N2) | 20/9/2015 | NCBI |
| KU591519 | A/NewYork/71/2015(H3N2) | 3/11/2015 | NCBI |
| KU591527 | A/Florida/84/2015(H3N2) | 6/11/2015 | NCBI |
| KU591535 | A/Oregon/25/2015(H3N2) | 19/10/2015 | NCBI |
| KU591543 | A/Oregon/27/2015(H3N2) | 23/10/2015 | NCBI |
| KU591552 | A/Alaska/212/2015(H3N2) | 30/8/2015 | NCBI |
| KU591560 | A/Alaska/215/2015(H3N2) | 4/9/2015 | NCBI |
| KU591908 | A/Hawaii/69/2015(H3N2) | 4/10/2015 | NCBI |
| KU591566 | A/Hawaii/70/2015(H3N2) | 5/10/2015 | NCBI |
| KU591574 | A/Colorado/15/2015(H3N2) | 11/10/2015 | NCBI |
| KU591582 | A/Alaska/260/2015(H3N2) | 22/10/2015 | NCBI |
| KU591590 | A/Alaska/262/2015(H3N2) | 25/10/2015 | NCBI |
| KU591598 | A/WestVirginia/24/2015(H3N2) | 19/10/2015 | NCBI |
| KU592150 | A/Hawaii/72/2015(H3N2) | 8/10/2015 | NCBI |
| KU591930 | A/Florida/82/2015(H3N2) | 26/10/2015 | NCBI |
| KU591622 | A/Wisconsin/87/2015(H3N2) | 29/10/2015 | NCBI |
| KU591630 | A/Louisiana/06/2015(H3N2) | 2/11/2015 | NCBI |
| KU591638 | A/Florida/85/2015(H3N2) | 2/11/2015 | NCBI |
| KU591653 | A/Florida/87/2015(H3N2) | 8/11/2015 | NCBI |
| KU591946 | A/SouthCarolina/10/2015(H3N2) | 6/10/2015 | NCBI |
| KU591669 | A/Wyoming/19/2015(H3N2) | 3/11/2015 | NCBI |
| KU591677 | A/California/93/2015(H3N2) | 20/10/2015 | NCBI |
| KU591692 | A/California/95/2015(H3N2) | 20/10/2015 | NCBI |
| KU591708 | A/Washington/58/2015(H3N2) | 9/11/2015 | NCBI |
| KU591735 | A/California/84/2015(H3N2) | 30/8/2015 | NCBI |
| KU591743 | A/Georgia/12/2015(H3N2) | 2015 | NCBI |
| KU591751 | A/Georgia/13/2015(H3N2) | 9/10/2015 | NCBI |
| KU591759 | A/Florida/76/2015(H3N2) | 20/10/2015 | NCBI |
| KU591767 | A/Florida/77/2015(H3N2) | 24/10/2015 | NCBI |
| KU591789 | A/Iowa/45/2015(H3N2) | 22/10/2015 | NCBI |
| KU591797 | A/Alaska/216/2015(H3N2) | 6/9/2015 | NCBI |

| | | | |
|---|---|---|---|
| KU591805 | A/Alaska/220/2015(H3N2) | 4/9/2015 | NCBI |
| KU591813 | A/Alaska/221/2015(H3N2) | 3/9/2015 | NCBI |
| KU591829 | A/Alaska/227/2015(H3N2) | 11/9/2015 | NCBI |
| KU591837 | A/Alaska/231/2015(H3N2) | 15/9/2015 | NCBI |
| KU591845 | A/Alaska/237/2015(H3N2) | 12/9/2015 | NCBI |
| KU591861 | A/Alaska/244/2015(H3N2) | 14/9/2015 | NCBI |
| KU591876 | A/Alaska/246/2015(H3N2) | 13/9/2015 | NCBI |
| KU591900 | A/Michigan/58/2015(H3N2) | 4/10/2015 | NCBI |
| KU591922 | A/Hawaii/75/2015(H3N2) | 17/10/2015 | NCBI |
| KU591938 | A/Minnesota/47/2015(H3N2) | 16/11/2015 | NCBI |
| KU591954 | A/SouthCarolina/11/2015(H3N2) | 27/10/2015 | NCBI |
| KU591970 | A/Washington/57/2015(H3N2) | 7/11/2015 | NCBI |
| KU591984 | A/Alaska/218/2015(H3N2) | 7/9/2015 | NCBI |
| KU591992 | A/Alaska/219/2015(H3N2) | 5/9/2015 | NCBI |
| KU592000 | A/Alaska/228/2015(H3N2) | 12/9/2015 | NCBI |
| KU592008 | A/Alaska/229/2015(H3N2) | 13/9/2015 | NCBI |
| KU592016 | A/Alaska/230/2015(H3N2) | 14/9/2015 | NCBI |
| KU592024 | A/Alaska/232/2015(H3N2) | 9/9/2015 | NCBI |
| KU592032 | A/Alaska/235/2015(H3N2) | 16/9/2015 | NCBI |
| KU592040 | A/Alaska/236/2015(H3N2) | 16/9/2015 | NCBI |
| KU592048 | A/Alaska/238/2015(H3N2) | 16/9/2015 | NCBI |
| KU592056 | A/Alaska/240/2015(H3N2) | 15/9/2015 | NCBI |
| KU592064 | A/Alaska/241/2015(H3N2) | 16/9/2015 | NCBI |
| KU592072 | A/Alaska/243/2015(H3N2) | 14/9/2015 | NCBI |
| KU592080 | A/Alaska/247/2015(H3N2) | 17/9/2015 | NCBI |
| KU592088 | A/Alaska/249/2015(H3N2) | 17/9/2015 | NCBI |
| KU592112 | A/Hawaii/71/2015(H3N2) | 9/10/2015 | NCBI |
| KU592120 | A/Alaska/261/2015(H3N2) | 26/10/2015 | NCBI |
| KU592142 | A/Washington/54/2015(H3N2) | 17/10/2015 | NCBI |
| KU592158 | A/Hawaii/73/2015(H3N2) | 12/10/2015 | NCBI |
| KU592171 | A/SouthDakota/36/2015(H3N2) | 1/8/2015 | NCBI |
| KU592187 | A/SouthDakota/39/2015(H3N2) | 13/10/2015 | NCBI |
| KU592744 | A/Pennsylvania/09/2015(H3N2) | 7/1/2015 | NCBI |
| KU592760 | A/Maryland/26/2014(H3N2) | 16/9/2014 | NCBI |
| KU592823 | A/Michigan/15/2014(H3N2) | 24/9/2014 | NCBI |
| KU592736 | A/Louisiana/20/2014(H3N2) | 19/11/2014 | NCBI |
| KU592839 | A/Louisiana/39/2013(H3N2) | 29/12/2013 | NCBI |

| | | | |
|---|---|---|---|
| KU592779 | A/NorthCarolina/13/2014(H3N2) | 15/4/2014 | NCBI |
| KU592771 | A/Wisconsin/66/2015(H3N2) | 21/4/2015 | NCBI |
| KU299817 | A/Cambodia/V1221301/2011(H3N2) | 14/10/2011 | NCBI |
| KU299809 | A/Cambodia/V1028352/2011(H3N2) | 18/10/2011 | NCBI |
| KU299801 | A/Cambodia/V1028340/2011(H3N2) | 18/10/2011 | NCBI |
| KU299793 | A/Cambodia/V1012345/2011(H3N2) | 4/10/2011 | NCBI |
| KU888853 | A/DC/1/2014(H3N2) | 30/12/2014 | NCBI |
| KU933501 | A/Montana/28/2015(H3N2) | 24/9/2015 | NCBI |
| KX133410 | A/Gainesville/01/2016(H3N2) | 18/3/2016 | NCBI |
| KX003350 | A/Texas/64/2015(H3N2) | 1/7/2015 | NCBI |
| KU591203 | A/Hawaii/43/2015(H3N2) | 8/7/2015 | NCBI |
| KX003360 | A/SouthDakota/38/2015(H3N2) | 9/9/2015 | NCBI |
| KX003370 | A/Michigan/60/2015(H3N2) | 5/10/2015 | NCBI |
| KU592195 | A/Oregon/25/2015(H3N2) | 19/10/2015 | NCBI |
| KX003382 | A/Arizona/23/2015(H3N2) | 26/10/2015 | NCBI |
| KX003390 | A/Arizona/24/2015(H3N2) | 2/11/2015 | NCBI |
| KX003398 | A/Arizona/25/2015(H3N2) | 2/11/2015 | NCBI |
| KX003406 | A/Arizona/27/2015(H3N2) | 18/10/2015 | NCBI |
| KX003413 | A/Oklahoma/21/2015(H3N2) | 18/11/2015 | NCBI |
| KX003421 | A/Florida/94/2015(H3N2) | 16/11/2015 | NCBI |
| KX003429 | A/Nevada/37/2015(H3N2) | 13/11/2015 | NCBI |
| KX003445 | A/Indiana/29/2015(H3N2) | 27/11/2015 | NCBI |
| KX003453 | A/Washington/60/2015(H3N2) | 3/11/2015 | NCBI |
| KX003458 | A/Washington/61/2015(H3N2) | 1/11/2015 | NCBI |
| KX003463 | A/Texas/73/2015(H3N2) | 2/12/2015 | NCBI |
| KX003471 | A/Nevada/38/2015(H3N2) | 14/12/2015 | NCBI |
| KX003479 | A/Nevada/39/2015(H3N2) | 15/12/2015 | NCBI |
| KX003487 | A/SouthDakota/40/2015(H3N2) | 14/11/2015 | NCBI |
| KX003495 | A/Alaska/45/2015(H3N2) | 23/6/2015 | NCBI |
| KX003502 | A/Alaska/47/2015(H3N2) | 23/6/2015 | NCBI |
| KU590075 | A/Alaska/49/2015(H3N2) | 15/6/2015 | NCBI |
| KU590083 | A/Alaska/50/2015(H3N2) | 16/6/2015 | NCBI |
| KU590097 | A/Alaska/59/2015(H3N2) | 17/6/2015 | NCBI |
| KX003515 | A/Alaska/62/2015(H3N2) | 21/6/2015 | NCBI |
| KX003523 | A/Alaska/63/2015(H3N2) | 24/6/2015 | NCBI |
| KX003531 | A/Alaska/64/2015(H3N2) | 24/6/2015 | NCBI |
| KX003539 | A/Alaska/75/2015(H3N2) | 23/6/2015 | NCBI |

| | | | |
|---|---|---|---|
| KX003547 | A/Alaska/77/2015(H3N2) | 25/6/2015 | NCBI |
| KX003555 | A/Alaska/79/2015(H3N2) | 24/6/2015 | NCBI |
| KX003563 | A/Alaska/80/2015(H3N2) | 30/6/2015 | NCBI |
| KX003571 | A/Alaska/83/2015(H3N2) | 6/7/2015 | NCBI |
| KX003579 | A/Alaska/86/2015(H3N2) | 6/7/2015 | NCBI |
| KX003587 | A/Alaska/93/2015(H3N2) | 1/7/2015 | NCBI |
| KX003595 | A/Alaska/96/2015(H3N2) | 30/6/2015 | NCBI |
| KX003603 | A/Alaska/97/2015(H3N2) | 2/7/2015 | NCBI |
| KX003611 | A/Alaska/98/2015(H3N2) | 3/7/2015 | NCBI |
| KX003619 | A/Alaska/99/2015(H3N2) | 3/7/2015 | NCBI |
| KX003627 | A/Alaska/100/2015(H3N2) | 3/7/2015 | NCBI |
| KX003635 | A/Alaska/101/2015(H3N2) | 4/7/2015 | NCBI |
| KX003643 | A/Alaska/102/2015(H3N2) | 28/6/2015 | NCBI |
| KX003651 | A/Alaska/103/2015(H3N2) | 4/7/2015 | NCBI |
| KX003659 | A/Alaska/104/2015(H3N2) | 4/7/2015 | NCBI |
| KX003667 | A/Alaska/105/2015(H3N2) | 4/7/2015 | NCBI |
| KX003674 | A/Alaska/106/2015(H3N2) | 27/6/2015 | NCBI |
| KX003680 | A/Alaska/107/2015(H3N2) | 3/7/2015 | NCBI |
| KX003686 | A/Alaska/108/2015(H3N2) | 6/7/2015 | NCBI |
| KX003693 | A/Alaska/109/2015(H3N2) | 7/7/2015 | NCBI |
| KU590854 | A/Alaska/111/2015(H3N2) | 9/7/2015 | NCBI |
| KX003707 | A/Alaska/112/2015(H3N2) | 4/7/2015 | NCBI |
| KX003715 | A/Alaska/113/2015(H3N2) | 10/7/2015 | NCBI |
| KX003723 | A/Alaska/114/2015(H3N2) | 6/7/2015 | NCBI |
| KX003731 | A/Alaska/115/2015(H3N2) | 4/7/2015 | NCBI |
| KX003739 | A/NewYork/64/2015(H3N2) | 10/8/2015 | NCBI |
| KX003747 | A/Alaska/117/2015(H3N2) | 7/7/2015 | NCBI |
| KX003755 | A/Alaska/118/2015(H3N2) | 15/7/2015 | NCBI |
| KX003763 | A/Alaska/119/2015(H3N2) | 8/7/2015 | NCBI |
| KX003779 | A/Alaska/120/2015(H3N2) | 10/7/2015 | NCBI |
| KX003787 | A/Alaska/121/2015(H3N2) | 9/7/2015 | NCBI |
| KX003795 | A/Alaska/122/2015(H3N2) | 5/7/2015 | NCBI |
| KX003803 | A/Alaska/123/2015(H3N2) | 15/7/2015 | NCBI |
| KX003811 | A/Alaska/124/2015(H3N2) | 14/7/2015 | NCBI |
| KX003819 | A/Alaska/126/2015(H3N2) | 17/7/2015 | NCBI |
| KX003827 | A/Alaska/127/2015(H3N2) | 17/7/2015 | NCBI |
| KX003835 | A/Alaska/128/2015(H3N2) | 16/7/2015 | NCBI |

| | | | |
|---|---|---|---|
| KX003843 | A/Alaska/129/2015(H3N2) | 18/7/2015 | NCBI |
| KX003851 | A/Alaska/131/2015(H3N2) | 23/7/2015 | NCBI |
| KX003859 | A/Alaska/132/2015(H3N2) | 18/7/2015 | NCBI |
| KX003867 | A/Alaska/133/2015(H3N2) | 25/7/2015 | NCBI |
| KX003875 | A/Alaska/134/2015(H3N2) | 26/7/2015 | NCBI |
| KX003883 | A/Alaska/159/2015(H3N2) | 17/8/2015 | NCBI |
| KX003891 | A/Nebraska/18/2015(H3N2) | 4/9/2015 | NCBI |
| KX003898 | A/Minnesota/40/2015(H3N2) | 1/9/2015 | NCBI |
| KX003906 | A/Alabama/10/2015(H3N2) | 8/9/2015 | NCBI |
| KX003914 | A/Indiana/20/2015(H3N2) | 1/9/2015 | NCBI |
| KX003922 | A/Pennsylvania/32/2015(H3N2) | 16/9/2015 | NCBI |
| KU590063 | A/Montana/19/2015(H3N2) | 13/9/2015 | NCBI |
| KX003929 | A/Montana/20/2015(H3N2) | 15/9/2015 | NCBI |
| KX003935 | A/Montana/21/2015(H3N2) | 14/9/2015 | NCBI |
| KX003941 | A/Montana/24/2015(H3N2) | 18/9/2015 | NCBI |
| KX003950 | A/NewJersey/52/2015(H3N2) | 1/9/2015 | NCBI |
| KX003958 | A/Alaska/206/2015(H3N2) | 7/9/2015 | NCBI |
| KX003966 | A/Alaska/207/2015(H3N2) | 7/9/2015 | NCBI |
| KX003974 | A/Alaska/209/2015(H3N2) | 7/9/2015 | NCBI |
| KX003982 | A/Alaska/210/2015(H3N2) | 7/9/2015 | NCBI |
| KX003990 | A/Alaska/213/2015(H3N2) | 30/8/2015 | NCBI |
| KX003997 | A/Alaska/214/2015(H3N2) | 31/8/2015 | NCBI |
| KX004005 | A/Washington/45/2015(H3N2) | 16/6/2015 | NCBI |
| KX004013 | A/Washington/46/2015(H3N2) | 15/9/2015 | NCBI |
| KX004021 | A/Washington/47/2015(H3N2) | 19/9/2015 | NCBI |
| KU591381 | A/Washington/48/2015(H3N2) | 19/9/2015 | NCBI |
| KU592165 | A/Montana/30/2015(H3N2) | 6/10/2015 | NCBI |
| KU589847 | A/Montana/31/2015(H3N2) | 7/10/2015 | NCBI |
| KX417083 | A/Michigan/50/2016(H3N2) | 16/2/2016 | NCBI |
| KX415624 | A/Nevada/22/2016(H3N2) | 4/3/2016 | NCBI |
| KX004045 | A/Montana/32/2015(H3N2) | 12/10/2015 | NCBI |
| KX411927 | A/Arizona/15/2015(H3N2) | 20/9/2015 | NCBI |
| KX411935 | A/Arizona/16/2015(H3N2) | 28/9/2015 | NCBI |
| KX411943 | A/Arizona/17/2015(H3N2) | 28/9/2015 | NCBI |
| KX411950 | A/Arizona/18/2015(H3N2) | 30/9/2015 | NCBI |
| KX411958 | A/Arizona/19/2015(H3N2) | 29/9/2015 | NCBI |
| KU590454 | A/Arizona/20/2015(H3N2) | 4/10/2015 | NCBI |

| | | | |
|---|---|---|---|
| KX411966 | A/Arizona/21/2015(H3N2) | 2/10/2015 | NCBI |
| KU590008 | A/Arizona/22/2015(H3N2) | 7/10/2015 | NCBI |
| KX411981 | A/Maine/43/2015(H3N2) | 19/10/2015 | NCBI |
| KX411989 | A/Wisconsin/86/2015(H3N2) | 19/10/2015 | NCBI |
| KX411996 | A/NewYork/67/2015(H3N2) | 6/9/2015 | NCBI |
| KX412004 | A/NewYork/69/2015(H3N2) | 12/10/2015 | NCBI |
| KX412012 | A/Florida/76/2015(H3N2) | 20/10/2015 | NCBI |
| KX412020 | A/Florida/77/2015(H3N2) | 24/10/2015 | NCBI |
| KU591774 | A/Florida/78/2015(H3N2) | 22/10/2015 | NCBI |
| KX412029 | A/Florida/79/2015(H3N2) | 26/10/2015 | NCBI |
| KU591781 | A/Iowa/44/2015(H3N2) | 21/10/2015 | NCBI |
| KX412045 | A/Iowa/45/2015(H3N2) | 22/10/2015 | NCBI |
| KX412061 | A/Iowa/46/2015(H3N2) | 21/10/2015 | NCBI |
| KX412069 | A/Iowa/47/2015(H3N2) | 22/10/2015 | NCBI |
| KX412077 | A/Iowa/48/2015(H3N2) | 24/10/2015 | NCBI |
| KX412085 | A/Iowa/49/2015(H3N2) | 24/10/2015 | NCBI |
| KX412093 | A/Alaska/163/2015(H3N2) | 4/8/2015 | NCBI |
| KX412101 | A/Alaska/164/2015(H3N2) | 3/8/2015 | NCBI |
| KX412109 | A/Alaska/165/2015(H3N2) | 19/8/2015 | NCBI |
| KX412117 | A/Alaska/166/2015(H3N2) | 15/8/2015 | NCBI |
| KX412125 | A/Alaska/167/2015(H3N2) | 15/8/2015 | NCBI |
| KX412133 | A/Alaska/168/2015(H3N2) | 15/8/2015 | NCBI |
| KX412141 | A/Alaska/170/2015(H3N2) | 21/8/2015 | NCBI |
| KX412149 | A/Alaska/171/2015(H3N2) | 25/8/2015 | NCBI |
| KU591548 | A/Alaska/172/2015(H3N2) | 26/8/2015 | NCBI |
| KX412163 | A/Alaska/173/2015(H3N2) | 29/8/2015 | NCBI |
| KX412171 | A/Alaska/174/2015(H3N2) | 30/8/2015 | NCBI |
| KX412179 | A/Alaska/176/2015(H3N2) | 19/8/2015 | NCBI |
| KX412187 | A/Alaska/177/2015(H3N2) | 21/8/2015 | NCBI |
| KX412195 | A/Alaska/178/2015(H3N2) | 18/8/2015 | NCBI |
| KX412200 | A/Alaska/184/2015(H3N2) | 20/8/2015 | NCBI |
| KX412204 | A/Alaska/187/2015(H3N2) | 19/8/2015 | NCBI |
| KX412209 | A/Alaska/194/2015(H3N2) | 17/8/2015 | NCBI |
| KX412217 | A/Alaska/195/2015(H3N2) | 24/8/2015 | NCBI |
| KX412225 | A/Alaska/197/2015(H3N2) | 29/8/2015 | NCBI |
| KX412233 | A/Alaska/198/2015(H3N2) | 29/8/2015 | NCBI |
| KX412241 | A/Alaska/199/2015(H3N2) | 29/8/2015 | NCBI |

| | | | |
|---|---|---|---|
| KX412249 | A/Alaska/201/2015(H3N2) | 7/9/2015 | NCBI |
| KX412257 | A/Alaska/202/2015(H3N2) | 7/9/2015 | NCBI |
| KX412265 | A/Alaska/204/2015(H3N2) | 5/9/2015 | NCBI |
| KX412273 | A/Alaska/205/2015(H3N2) | 7/9/2015 | NCBI |
| KU591976 | A/Alaska/217/2015(H3N2) | 7/9/2015 | NCBI |
| KX412284 | A/Alaska/224/2015(H3N2) | 12/9/2015 | NCBI |
| KX412292 | A/Alaska/226/2015(H3N2) | 10/9/2015 | NCBI |
| KX412300 | A/Alaska/233/2015(H3N2) | 16/9/2015 | NCBI |
| KX412308 | A/Alaska/234/2015(H3N2) | 16/9/2015 | NCBI |
| KU591868 | A/Alaska/245/2015(H3N2) | 10/9/2015 | NCBI |
| KX412317 | A/Alaska/248/2015(H3N2) | 19/9/2015 | NCBI |
| KX412324 | A/Alaska/250/2015(H3N2) | 20/9/2015 | NCBI |
| KX412331 | A/Alaska/251/2015(H3N2) | 18/9/2015 | NCBI |
| KX412338 | A/Alaska/252/2015(H3N2) | 21/9/2015 | NCBI |
| KU591892 | A/Hawaii/63/2015(H3N2) | 28/9/2015 | NCBI |
| KX412347 | A/Hawaii/64/2015(H3N2) | 28/9/2015 | NCBI |
| KX412363 | A/Pennsylvania/33/2015(H3N2) | 21/9/2015 | NCBI |
| KX412371 | A/Pennsylvania/34/2015(H3N2) | 28/9/2015 | NCBI |
| KX412379 | A/Pennsylvania/35/2015(H3N2) | 30/9/2015 | NCBI |
| KX412387 | A/Delaware/43/2015(H3N2) | 23/10/2015 | NCBI |
| KX412395 | A/Massachusetts/29/2015(H3N2) | 14/10/2015 | NCBI |
| KX412403 | A/Washington/52/2015(H3N2) | 16/10/2015 | NCBI |
| KX412411 | A/Washington/59/2015(H3N2) | 21/10/2015 | NCBI |
| KU592127 | A/Alaska/262/2015(H3N2) | 25/10/2015 | NCBI |
| KU592134 | A/Washington/53/2015(H3N2) | 17/10/2015 | NCBI |
| KX412421 | A/Louisiana/06/2015(H3N2) | 2/11/2015 | NCBI |
| KX412429 | A/Colorado/21/2015(H3N2) | 19/11/2015 | NCBI |
| KX412437 | A/Colorado/23/2015(H3N2) | 22/11/2015 | NCBI |
| KX412445 | A/Colorado/24/2015(H3N2) | 17/11/2015 | NCBI |
| KU591684 | A/California/94/2015(H3N2) | 20/10/2015 | NCBI |
| KX412454 | A/California/96/2015(H3N2) | 21/10/2015 | NCBI |
| KX412462 | A/California/97/2015(H3N2) | 24/10/2015 | NCBI |
| KX412470 | A/Illinois/19/2015(H3N2) | 12/10/2015 | NCBI |
| KX412478 | A/Louisiana/07/2015(H3N2) | 11/11/2015 | NCBI |
| KX412486 | A/Texas/72/2015(H3N2) | 8/11/2015 | NCBI |
| KX412502 | A/Indiana/31/2015(H3N2) | 14/12/2015 | NCBI |
| KX412510 | A/Delaware/44/2015(H3N2) | 2/12/2015 | NCBI |

| | | | |
|---|---|---|---|
| KX412518 | A/Texas/74/2015(H3N2) | 21/12/2015 | NCBI |
| KX412534 | A/Washington/62/2015(H3N2) | 3/12/2015 | NCBI |
| KX412542 | A/Washington/64/2015(H3N2) | 15/12/2015 | NCBI |
| KX412550 | A/Washington/65/2015(H3N2) | 15/12/2015 | NCBI |
| KX412558 | A/Delaware/46/2015(H3N2) | 21/12/2015 | NCBI |
| KX412566 | A/Delaware/47/2015(H3N2) | 26/12/2015 | NCBI |
| KX412574 | A/Delaware/48/2015(H3N2) | 29/12/2015 | NCBI |
| KX412582 | A/Idaho/27/2015(H3N2) | 30/11/2015 | NCBI |
| KX412590 | A/Idaho/28/2015(H3N2) | 1/12/2015 | NCBI |
| KX412598 | A/Idaho/29/2015(H3N2) | 4/12/2015 | NCBI |
| KX412606 | A/Idaho/30/2015(H3N2) | 5/12/2015 | NCBI |
| KX412614 | A/Idaho/31/2015(H3N2) | 8/12/2015 | NCBI |
| KX412622 | A/Idaho/32/2015(H3N2) | 8/12/2015 | NCBI |
| KX412630 | A/NewMexico/30/2015(H3N2) | 13/12/2015 | NCBI |
| KX412646 | A/Pennsylvania/52/2015(H3N2) | 26/12/2015 | NCBI |
| KX412654 | A/Connecticut/11/2015(H3N2) | 20/12/2015 | NCBI |
| KX412662 | A/Alabama/02/2016(H3N2) | 27/1/2016 | NCBI |
| KX412670 | A/Alabama/03/2016(H3N2) | 27/1/2016 | NCBI |
| KX412678 | A/Alabama/04/2016(H3N2) | 31/1/2016 | NCBI |
| KX412686 | A/Alabama/05/2016(H3N2) | 31/1/2016 | NCBI |
| KX412694 | A/Illinois/27/2015(H3N2) | 16/12/2015 | NCBI |
| KX412702 | A/Washington/69/2015(H3N2) | 28/12/2015 | NCBI |
| KX412718 | A/NorthCarolina/35/2015(H3N2) | 30/12/2015 | NCBI |
| KX412726 | A/Nevada/01/2016(H3N2) | 3/1/2016 | NCBI |
| KX412734 | A/NorthCarolina/02/2016(H3N2) | 12/1/2016 | NCBI |
| KX412742 | A/NorthCarolina/03/2016(H3N2) | 12/1/2016 | NCBI |
| KX412750 | A/NorthCarolina/04/2016(H3N2) | 13/1/2016 | NCBI |
| KX412766 | A/NorthCarolina/05/2016(H3N2) | 13/1/2016 | NCBI |
| KX412774 | A/NorthCarolina/06/2016(H3N2) | 12/1/2016 | NCBI |
| KX412782 | A/NewYork/76/2015(H3N2) | 1/12/2015 | NCBI |
| KX412790 | A/NewYork/77/2015(H3N2) | 7/12/2015 | NCBI |
| KX412798 | A/Georgia/03/2016(H3N2) | 12/1/2016 | NCBI |
| KX412814 | A/NewJersey/55/2015(H3N2) | 13/12/2015 | NCBI |
| KX412822 | A/Nevada/03/2016(H3N2) | 8/1/2016 | NCBI |
| KX412830 | A/Texas/01/2016(H3N2) | 2/1/2016 | NCBI |
| KX412838 | A/Alabama/06/2016(H3N2) | 20/1/2016 | NCBI |
| KX412846 | A/Pennsylvania/03/2016(H3N2) | 23/1/2016 | NCBI |

| | | | |
|---|---|---|---|
| KX412862 | A/Texas/06/2016(H3N2) | 20/1/2016 | NCBI |
| KX412878 | A/Texas/08/2016(H3N2) | 25/1/2016 | NCBI |
| KX412886 | A/NewMexico/32/2015(H3N2) | 16/12/2015 | NCBI |
| KX412894 | A/SouthCarolina/05/2016(H3N2) | 25/1/2016 | NCBI |
| KX412902 | A/Missouri/02/2016(H3N2) | 26/1/2016 | NCBI |
| KX412910 | A/Texas/14/2016(H3N2) | 3/2/2016 | NCBI |
| KX412918 | A/Georgia/16/2016(H3N2) | 2/2/2016 | NCBI |
| KX412926 | A/Georgia/18/2016(H3N2) | 10/2/2016 | NCBI |
| KX412934 | A/Georgia/19/2016(H3N2) | 15/2/2016 | NCBI |
| KX412942 | A/Michigan/30/2016(H3N2) | 22/2/2016 | NCBI |
| KX412950 | A/Michigan/31/2016(H3N2) | 23/2/2016 | NCBI |
| KX412958 | A/Washington/72/2015(H3N2) | 22/9/2015 | NCBI |
| KX412966 | A/Washington/73/2015(H3N2) | 18/9/2015 | NCBI |
| KX412974 | A/Washington/74/2015(H3N2) | 22/9/2015 | NCBI |
| KX412982 | A/Washington/75/2015(H3N2) | 24/9/2015 | NCBI |
| KX412990 | A/Washington/76/2015(H3N2) | 21/10/2015 | NCBI |
| KX412998 | A/Alaska/269/2015(H3N2) | 16/11/2015 | NCBI |
| KX413006 | A/Idaho/44/2015(H3N2) | 30/12/2015 | NCBI |
| KX413014 | A/Louisiana/08/2015(H3N2) | 9/12/2015 | NCBI |
| KX413022 | A/Louisiana/11/2015(H3N2) | 19/12/2015 | NCBI |
| KX413030 | A/Kansas/02/2016(H3N2) | 22/1/2016 | NCBI |
| KX413038 | A/Kansas/03/2016(H3N2) | 14/1/2016 | NCBI |
| KX413046 | A/NewJersey/56/2015(H3N2) | 9/10/2015 | NCBI |
| KX413054 | A/California/133/2015(H3N2) | 29/12/2015 | NCBI |
| KX413062 | A/Texas/15/2016(H3N2) | 3/1/2016 | NCBI |
| KX413070 | A/Maine/03/2016(H3N2) | 13/2/2016 | NCBI |
| KX413078 | A/Maine/04/2016(H3N2) | 16/2/2016 | NCBI |
| KX413086 | A/Mississippi/02/2016(H3N2) | 4/2/2016 | NCBI |
| KX413094 | A/Arizona/32/2016(H3N2) | 8/1/2016 | NCBI |
| KX413102 | A/Arizona/37/2016(H3N2) | 7/2/2016 | NCBI |
| KX413110 | A/Virginia/26/2016(H3N2) | 27/3/2016 | NCBI |
| KX413118 | A/Virginia/27/2016(H3N2) | 26/3/2016 | NCBI |
| KX413126 | A/Virginia/32/2016(H3N2) | 4/4/2016 | NCBI |
| KX413134 | A/Texas/65/2016(H3N2) | 28/3/2016 | NCBI |
| KX413142 | A/Texas/66/2016(H3N2) | 21/1/2016 | NCBI |
| KX413150 | A/Texas/67/2016(H3N2) | 15/1/2016 | NCBI |
| KX413158 | A/Texas/71/2016(H3N2) | 2/2/2016 | NCBI |

| | | | |
|---|---|---|---|
| KX413166 | A/Texas/74/2016(H3N2) | 26/2/2016 | NCBI |
| KX413174 | A/Texas/76/2016(H3N2) | 21/2/2016 | NCBI |
| KX413182 | A/Texas/77/2016(H3N2) | 11/2/2016 | NCBI |
| KX413190 | A/Texas/78/2016(H3N2) | 21/2/2016 | NCBI |
| KX413198 | A/Texas/79/2016(H3N2) | 21/2/2016 | NCBI |
| KX413206 | A/Texas/80/2016(H3N2) | 18/2/2016 | NCBI |
| KX413214 | A/Texas/82/2016(H3N2) | 22/2/2016 | NCBI |
| KX413222 | A/Texas/83/2016(H3N2) | 26/2/2016 | NCBI |
| KX413238 | A/Texas/85/2016(H3N2) | 28/2/2016 | NCBI |
| KX413246 | A/Texas/86/2016(H3N2) | 25/2/2016 | NCBI |
| KX413262 | A/Texas/88/2016(H3N2) | 24/2/2016 | NCBI |
| KX413270 | A/Virginia/37/2016(H3N2) | 20/4/2016 | NCBI |
| KX413278 | A/Georgia/37/2016(H3N2) | 22/2/2016 | NCBI |
| KX413286 | A/Georgia/40/2016(H3N2) | 6/3/2016 | NCBI |
| KX413294 | A/Georgia/45/2016(H3N2) | 6/3/2016 | NCBI |
| KU591715 | A/Michigan/66/2015(H3N2) | 15/10/2015 | NCBI |
| KX413311 | A/Michigan/67/2015(H3N2) | 10/10/2015 | NCBI |
| KU589661 | A/Iowa/54/2015(H3N2) | 1/11/2015 | NCBI |
| KX413331 | A/Hawaii/76/2015(H3N2) | 18/10/2015 | NCBI |
| KX413338 | A/Hawaii/77/2015(H3N2) | 2/11/2015 | NCBI |
| KX413346 | A/California/100/2015(H3N2) | 4/11/2015 | NCBI |
| KX413354 | A/California/101/2015(H3N2) | 4/11/2015 | NCBI |
| KX413362 | A/California/106/2015(H3N2) | 3/11/2015 | NCBI |
| KX413369 | A/California/107/2015(H3N2) | 18/10/2015 | NCBI |
| KX413377 | A/California/108/2015(H3N2) | 12/11/2015 | NCBI |
| KX413385 | A/Pennsylvania/44/2015(H3N2) | 7/11/2015 | NCBI |
| KX413401 | A/Pennsylvania/45/2015(H3N2) | 12/11/2015 | NCBI |
| KX413417 | A/Florida/89/2015(H3N2) | 14/11/2015 | NCBI |
| KX413425 | A/Florida/90/2015(H3N2) | 16/11/2015 | NCBI |
| KX413433 | A/California/110/2015(H3N2) | 19/11/2015 | NCBI |
| KX413449 | A/California/111/2015(H3N2) | 20/11/2015 | NCBI |
| KX413457 | A/Alaska/264/2015(H3N2) | 11/11/2015 | NCBI |
| KX413465 | A/Pennsylvania/50/2015(H3N2) | 23/11/2015 | NCBI |
| KX414161 | A/Arizona/29/2015(H3N2) | 27/11/2015 | NCBI |
| KX413481 | A/Colorado/27/2015(H3N2) | 8/12/2015 | NCBI |
| KX413489 | A/Florida/91/2015(H3N2) | 1/12/2015 | NCBI |
| KX413497 | A/Florida/93/2015(H3N2) | 4/12/2015 | NCBI |

| | | | |
|---|---|---|---|
| KX413505 | A/California/112/2015(H3N2) | 18/11/2015 | NCBI |
| KX413513 | A/California/113/2015(H3N2) | 28/11/2015 | NCBI |
| KX413521 | A/Alaska/266/2015(H3N2) | 26/11/2015 | NCBI |
| KX413529 | A/Alaska/267/2015(H3N2) | 27/11/2015 | NCBI |
| KX413537 | A/Washington/66/2015(H3N2) | 1/12/2015 | NCBI |
| KX413553 | A/Minnesota/51/2015(H3N2) | 14/12/2015 | NCBI |
| KX413561 | A/Minnesota/52/2015(H3N2) | 13/12/2015 | NCBI |
| KX413569 | A/Washington/68/2015(H3N2) | 8/12/2015 | NCBI |
| KX413577 | A/Hawaii/78/2015(H3N2) | 5/12/2015 | NCBI |
| KX413601 | A/Hawaii/79/2015(H3N2) | 11/12/2015 | NCBI |
| KX413609 | A/Florida/96/2015(H3N2) | 7/12/2015 | NCBI |
| KX413617 | A/Montana/36/2015(H3N2) | 10/12/2015 | NCBI |
| KX413633 | A/California/117/2015(H3N2) | 5/12/2015 | NCBI |
| KX413641 | A/California/118/2015(H3N2) | 15/12/2015 | NCBI |
| KX413649 | A/Florida/97/2015(H3N2) | 14/12/2015 | NCBI |
| KX413657 | A/Florida/98/2015(H3N2) | 11/12/2015 | NCBI |
| KX413665 | A/Colorado/28/2015(H3N2) | 12/12/2015 | NCBI |
| KX413673 | A/Colorado/29/2015(H3N2) | 17/12/2015 | NCBI |
| KX413689 | A/Minnesota/54/2015(H3N2) | 20/12/2015 | NCBI |
| KX413697 | A/Hawaii/80/2015(H3N2) | 11/12/2015 | NCBI |
| KX413705 | A/Hawaii/81/2015(H3N2) | 13/12/2015 | NCBI |
| KX413713 | A/Montana/39/2015(H3N2) | 24/12/2015 | NCBI |
| KX413721 | A/California/119/2015(H3N2) | 28/12/2015 | NCBI |
| KX413729 | A/Michigan/77/2015(H3N2) | 15/11/2015 | NCBI |
| KX413737 | A/Michigan/78/2015(H3N2) | 14/12/2015 | NCBI |
| KX413753 | A/Arizona/41/2015(H3N2) | 6/12/2015 | NCBI |
| KX413761 | A/Arizona/42/2015(H3N2) | 9/12/2015 | NCBI |
| KX413769 | A/Arizona/43/2015(H3N2) | 11/12/2015 | NCBI |
| KX414137 | A/Arizona/44/2015(H3N2) | 21/12/2015 | NCBI |
| KX413785 | A/Delaware/49/2015(H3N2) | 29/12/2015 | NCBI |
| KX413793 | A/Delaware/01/2016(H3N2) | 10/1/2016 | NCBI |
| KX413801 | A/Florida/104/2015(H3N2) | 23/12/2015 | NCBI |
| KX413809 | A/Florida/01/2016(H3N2) | 7/1/2016 | NCBI |
| KX413817 | A/NewYork/03/2016(H3N2) | 4/1/2016 | NCBI |
| KX413825 | A/California/02/2016(H3N2) | 7/1/2016 | NCBI |
| KX413833 | A/California/121/2015(H3N2) | 30/12/2015 | NCBI |
| KX413841 | A/California/122/2015(H3N2) | 29/12/2015 | NCBI |

| | | | |
|---|---|---|---|
| KX413849 | A/California/123/2015(H3N2) | 28/12/2015 | NCBI |
| KX413857 | A/California/124/2015(H3N2) | 11/12/2015 | NCBI |
| KX413865 | A/California/126/2015(H3N2) | 3/12/2015 | NCBI |
| KX413873 | A/California/127/2015(H3N2) | 19/12/2015 | NCBI |
| KX413889 | A/California/130/2015(H3N2) | 29/12/2015 | NCBI |
| KX413897 | A/Utah/03/2016(H3N2) | 6/1/2016 | NCBI |
| KX413905 | A/Hawaii/03/2016(H3N2) | 8/1/2016 | NCBI |
| KX413913 | A/Hawaii/04/2016(H3N2) | 12/1/2016 | NCBI |
| KX413921 | A/Oregon/01/2016(H3N2) | 8/1/2016 | NCBI |
| KX413929 | A/Colorado/38/2015(H3N2) | 26/12/2015 | NCBI |
| KX413937 | A/Colorado/39/2015(H3N2) | 27/12/2015 | NCBI |
| KX413945 | A/Colorado/37/2015(H3N2) | 30/12/2015 | NCBI |
| KX413953 | A/Colorado/40/2015(H3N2) | 14/12/2015 | NCBI |
| KX413961 | A/Colorado/03/2016(H3N2) | 6/1/2016 | NCBI |
| KX413969 | A/NewYork/07/2016(H3N2) | 11/1/2016 | NCBI |
| KX413977 | A/California/03/2016(H3N2) | 6/1/2016 | NCBI |
| KX413985 | A/California/131/2015(H3N2) | 30/12/2015 | NCBI |
| KX413993 | A/California/04/2016(H3N2) | 11/1/2016 | NCBI |
| KX414001 | A/California/05/2016(H3N2) | 2/1/2016 | NCBI |
| KX414017 | A/Washington/04/2016(H3N2) | 8/1/2016 | NCBI |
| KX414033 | A/Washington/05/2016(H3N2) | 12/1/2016 | NCBI |
| KX414041 | A/Washington/77/2015(H3N2) | 27/12/2015 | NCBI |
| KX414049 | A/Washington/08/2016(H3N2) | 1/1/2016 | NCBI |
| KX414057 | A/Pennsylvania/08/2016(H3N2) | 8/1/2016 | NCBI |
| KX414065 | A/Texas/11/2016(H3N2) | 16/1/2016 | NCBI |
| KX414073 | A/Texas/12/2016(H3N2) | 18/1/2016 | NCBI |
| KX414081 | A/Florida/06/2016(H3N2) | 10/1/2016 | NCBI |
| KX414089 | A/Florida/07/2016(H3N2) | 22/1/2016 | NCBI |
| KX414097 | A/Indiana/10/2016(H3N2) | 19/1/2016 | NCBI |
| KX414121 | A/Indiana/11/2016(H3N2) | 17/1/2016 | NCBI |
| KX414129 | A/NewYork/13/2016(H3N2) | 16/1/2016 | NCBI |
| KX414145 | A/Arizona/52/2015(H3N2) | 22/12/2015 | NCBI |
| KX414153 | A/Arizona/53/2015(H3N2) | 29/12/2015 | NCBI |
| KX414169 | A/Arizona/58/2015(H3N2) | 21/12/2015 | NCBI |
| KX414177 | A/Arizona/59/2015(H3N2) | 28/12/2015 | NCBI |
| KX414185 | A/Arizona/11/2016(H3N2) | 13/1/2016 | NCBI |
| KX414193 | A/Utah/07/2016(H3N2) | 15/1/2016 | NCBI |

| | | | |
|---|---|---|---|
| KX414201 | A/California/11/2016(H3N2) | 6/1/2016 | NCBI |
| KX414209 | A/Hawaii/05/2016(H3N2) | 5/1/2016 | NCBI |
| KX414217 | A/Hawaii/07/2016(H3N2) | 12/1/2016 | NCBI |
| KX414225 | A/Colorado/07/2016(H3N2) | 13/1/2016 | NCBI |
| KX414233 | A/Colorado/08/2016(H3N2) | 21/1/2016 | NCBI |
| KX414241 | A/Minnesota/09/2016(H3N2) | 20/1/2016 | NCBI |
| KX414257 | A/Washington/17/2016(H3N2) | 19/1/2016 | NCBI |
| KX414273 | A/Washington/18/2016(H3N2) | 20/1/2016 | NCBI |
| KX414281 | A/Arkansas/02/2016(H3N2) | 28/1/2016 | NCBI |
| KX414289 | A/Arkansas/03/2016(H3N2) | 1/2/2016 | NCBI |
| KX414297 | A/Montana/09/2016(H3N2) | 12/1/2016 | NCBI |
| KX414305 | A/Montana/13/2016(H3N2) | 30/1/2016 | NCBI |
| KX414313 | A/California/13/2016(H3N2) | 14/1/2016 | NCBI |
| KX414321 | A/California/20/2016(H3N2) | 26/1/2016 | NCBI |
| KX414329 | A/California/21/2016(H3N2) | 20/1/2016 | NCBI |
| KX414337 | A/California/24/2016(H3N2) | 3/1/2016 | NCBI |
| KX414345 | A/California/25/2016(H3N2) | 16/1/2016 | NCBI |
| KX414361 | A/Texas/22/2016(H3N2) | 1/2/2016 | NCBI |
| KX414353 | A/Texas/22/2016(H3N2) | 1/2/2016 | NCBI |
| KX414369 | A/Indiana/16/2016(H3N2) | 22/1/2016 | NCBI |
| KX414377 | A/Indiana/17/2016(H3N2) | 25/1/2016 | NCBI |
| KX414385 | A/Indiana/18/2016(H3N2) | 25/1/2016 | NCBI |
| KX414393 | A/Indiana/18/2016(H3N2) | 25/1/2016 | NCBI |
| KX414401 | A/Indiana/19/2016(H3N2) | 2/2/2016 | NCBI |
| KX414409 | A/Vermont/02/2016(H3N2) | 30/1/2016 | NCBI |
| KX414417 | A/Florida/52/2016(H3N2) | 24/1/2016 | NCBI |
| KX414425 | A/Florida/08/2016(H3N2) | 28/1/2016 | NCBI |
| KX414433 | A/Florida/09/2016(H3N2) | 26/1/2016 | NCBI |
| KX414441 | A/Florida/10/2016(H3N2) | 26/1/2016 | NCBI |
| KX414449 | A/Utah/12/2016(H3N2) | 30/1/2016 | NCBI |
| KX414457 | A/NewYork/14/2016(H3N2) | 29/1/2016 | NCBI |
| KX414465 | A/NewYork/79/2015(H3N2) | 14/12/2015 | NCBI |
| KX414473 | A/Idaho/05/2016(H3N2) | 26/1/2016 | NCBI |
| KX414481 | A/Idaho/05/2016(H3N2) | 26/1/2016 | NCBI |
| KX414489 | A/Arizona/22/2016(H3N2) | 23/1/2016 | NCBI |
| KX414497 | A/Arizona/23/2016(H3N2) | 20/1/2016 | NCBI |
| KX414505 | A/Arizona/21/2016(H3N2) | 24/1/2016 | NCBI |

| | | | |
|---|---|---|---|
| KX414513 | A/Arizona/24/2016(H3N2) | 20/1/2016 | NCBI |
| KX414521 | A/Arizona/25/2016(H3N2) | 20/1/2016 | NCBI |
| KX414529 | A/Arizona/26/2016(H3N2) | 20/1/2016 | NCBI |
| KX414537 | A/Arizona/27/2016(H3N2) | 18/1/2016 | NCBI |
| KX414545 | A/Wisconsin/22/2016(H3N2) | 31/1/2016 | NCBI |
| KX414553 | A/Washington/19/2016(H3N2) | 3/2/2016 | NCBI |
| KX414561 | A/Washington/19/2016(H3N2) | 3/2/2016 | NCBI |
| KX414569 | A/Michigan/24/2016(H3N2) | 1/2/2016 | NCBI |
| KX414577 | A/Washington/26/2016(H3N2) | 19/1/2016 | NCBI |
| KX414585 | A/Washington/26/2016(H3N2) | 19/1/2016 | NCBI |
| KX414593 | A/Colorado/11/2016(H3N2) | 28/1/2016 | NCBI |
| KX414601 | A/Colorado/12/2016(H3N2) | 28/1/2016 | NCBI |
| KX414617 | A/Minnesota/61/2015(H3N2) | 27/12/2015 | NCBI |
| KX414609 | A/Minnesota/61/2015(H3N2) | 27/12/2015 | NCBI |
| KX414625 | A/Florida/13/2016(H3N2) | 1/2/2016 | NCBI |
| KX414633 | A/Florida/13/2016(H3N2) | 1/2/2016 | NCBI |
| KX414641 | A/Florida/14/2016(H3N2) | 31/1/2016 | NCBI |
| KX414649 | A/Florida/15/2016(H3N2) | 22/1/2016 | NCBI |
| KX414657 | A/Florida/17/2016(H3N2) | 10/2/2016 | NCBI |
| KX414665 | A/Florida/18/2016(H3N2) | 9/2/2016 | NCBI |
| KX414673 | A/Oregon/02/2016(H3N2) | 14/1/2016 | NCBI |
| KX414681 | A/Oregon/04/2016(H3N2) | 2/2/2016 | NCBI |
| KX414689 | A/California/35/2016(H3N2) | 7/2/2016 | NCBI |
| KX414697 | A/California/36/2016(H3N2) | 6/2/2016 | NCBI |
| KX414705 | A/California/37/2016(H3N2) | 27/1/2016 | NCBI |
| KX414713 | A/Tennessee/02/2016(H3N2) | 5/2/2016 | NCBI |
| KX414721 | A/NewYork/18/2016(H3N2) | 5/2/2016 | NCBI |
| KX414729 | A/NewYork/19/2016(H3N2) | 9/2/2016 | NCBI |
| KX414737 | A/Montana/14/2016(H3N2) | 2/2/2016 | NCBI |
| KX414745 | A/NewMexico/05/2016(H3N2) | 27/1/2016 | NCBI |
| KX414753 | A/NewMexico/06/2016(H3N2) | 28/1/2016 | NCBI |
| KX414761 | A/NewMexico/06/2016(H3N2) | 28/1/2016 | NCBI |
| KX414769 | A/Michigan/29/2016(H3N2) | 5/2/2016 | NCBI |
| KX414777 | A/Texas/23/2016(H3N2) | 24/1/2016 | NCBI |
| KX414785 | A/Texas/26/2016(H3N2) | 5/2/2016 | NCBI |
| KX414793 | A/Arizona/39/2016(H3N2) | 5/2/2016 | NCBI |
| KX414801 | A/Utah/17/2016(H3N2) | 9/2/2016 | NCBI |

| | | | |
|---|---|---|---|
| KX414809 | A/Utah/14/2016(H3N2) | 9/2/2016 | NCBI |
| KX414817 | A/California/40/2016(H3N2) | 15/2/2016 | NCBI |
| KX414825 | A/California/41/2016(H3N2) | 15/2/2016 | NCBI |
| KX414833 | A/California/138/2015(H3N2) | 4/12/2015 | NCBI |
| KX414848 | A/California/135/2015(H3N2) | 8/12/2015 | NCBI |
| KX414856 | A/California/136/2015(H3N2) | 10/12/2015 | NCBI |
| KX414864 | A/California/43/2016(H3N2) | 11/1/2016 | NCBI |
| KX414880 | A/California/38/2016(H3N2) | 20/1/2016 | NCBI |
| KX414872 | A/California/38/2016(H3N2) | 20/1/2016 | NCBI |
| KX414888 | A/Texas/29/2016(H3N2) | 9/2/2016 | NCBI |
| KX414896 | A/Texas/30/2016(H3N2) | 16/2/2016 | NCBI |
| KX414904 | A/Maryland/05/2016(H3N2) | 5/2/2016 | NCBI |
| KX414912 | A/Alaska/06/2016(H3N2) | 23/1/2016 | NCBI |
| KX414920 | A/Alaska/05/2016(H3N2) | 31/1/2016 | NCBI |
| KX414928 | A/NewJersey/06/2016(H3N2) | 18/1/2016 | NCBI |
| KX414936 | A/Florida/20/2016(H3N2) | 11/2/2016 | NCBI |
| KX414944 | A/Florida/22/2016(H3N2) | 19/2/2016 | NCBI |
| KX414952 | A/Florida/53/2016(H3N2) | 15/2/2016 | NCBI |
| KX414960 | A/Florida/25/2016(H3N2) | 18/2/2016 | NCBI |
| KX414968 | A/Wisconsin/27/2016(H3N2) | 18/2/2016 | NCBI |
| KX414976 | A/Wisconsin/29/2016(H3N2) | 19/2/2016 | NCBI |
| KX414992 | A/Texas/33/2016(H3N2) | 20/2/2016 | NCBI |
| KX414984 | A/Texas/33/2016(H3N2) | 20/2/2016 | NCBI |
| KX415000 | A/Texas/34/2016(H3N2) | 22/2/2016 | NCBI |
| KX415008 | A/Indiana/24/2016(H3N2) | 14/2/2016 | NCBI |
| KX415016 | A/Indiana/25/2016(H3N2) | 16/2/2016 | NCBI |
| KX415024 | A/Nebraska/10/2016(H3N2) | 17/2/2016 | NCBI |
| KX415032 | A/Nebraska/11/2016(H3N2) | 13/2/2016 | NCBI |
| KX415040 | A/Colorado/15/2016(H3N2) | 18/2/2016 | NCBI |
| KX415048 | A/Washington/29/2016(H3N2) | 18/2/2016 | NCBI |
| KX415056 | A/NorthDakota/18/2016(H3N2) | 18/2/2016 | NCBI |
| KX415064 | A/Minnesota/13/2016(H3N2) | 10/2/2016 | NCBI |
| KX415072 | A/NewHampshire/03/2016(H3N2) | 15/2/2016 | NCBI |
| KX415080 | A/NewYork/20/2016(H3N2) | 12/2/2016 | NCBI |
| KX415088 | A/Washington/35/2016(H3N2) | 5/2/2016 | NCBI |
| KX415096 | A/Washington/36/2016(H3N2) | 12/2/2016 | NCBI |
| KX415104 | A/Oregon/05/2016(H3N2) | 16/2/2016 | NCBI |

| | | | |
|---|---|---|---|
| KX415112 | A/California/46/2016(H3N2) | 2/2/2016 | NCBI |
| KX415120 | A/California/47/2016(H3N2) | 4/2/2016 | NCBI |
| KX415128 | A/California/48/2016(H3N2) | 30/1/2016 | NCBI |
| KX415136 | A/California/49/2016(H3N2) | 1/2/2016 | NCBI |
| KX415144 | A/California/54/2016(H3N2) | 3/2/2016 | NCBI |
| KX415152 | A/California/58/2016(H3N2) | 3/2/2016 | NCBI |
| KX415160 | A/California/60/2016(H3N2) | 9/2/2016 | NCBI |
| KX415168 | A/NorthCarolina/13/2016(H3N2) | 22/2/2016 | NCBI |
| KX415176 | A/NorthCarolina/13/2016(H3N2) | 22/2/2016 | NCBI |
| KX415192 | A/RhodeIsland/11/2016(H3N2) | 17/2/2016 | NCBI |
| KX415200 | A/California/137/2015(H3N2) | 31/12/2015 | NCBI |
| KX415208 | A/California/62/2016(H3N2) | 12/1/2016 | NCBI |
| KX415216 | A/California/63/2016(H3N2) | 15/1/2016 | NCBI |
| KX415224 | A/California/64/2016(H3N2) | 21/1/2016 | NCBI |
| KX415232 | A/California/65/2016(H3N2) | 28/2/2016 | NCBI |
| KX415240 | A/Connecticut/04/2016(H3N2) | 15/2/2016 | NCBI |
| KX415248 | A/Connecticut/07/2016(H3N2) | 4/2/2016 | NCBI |
| KX415256 | A/Hawaii/13/2016(H3N2) | 13/2/2016 | NCBI |
| KX415264 | A/Hawaii/14/2016(H3N2) | 18/2/2016 | NCBI |
| KX415272 | A/Hawaii/15/2016(H3N2) | 18/2/2016 | NCBI |
| KX415280 | A/Hawaii/16/2016(H3N2) | 15/2/2016 | NCBI |
| KX415288 | A/Kentucky/07/2016(H3N2) | 24/2/2016 | NCBI |
| KX415296 | A/Kentucky/07/2016(H3N2) | 24/2/2016 | NCBI |
| KX415320 | A/NewJersey/07/2016(H3N2) | 25/2/2016 | NCBI |
| KX415328 | A/Virginia/25/2016(H3N2) | 1/3/2016 | NCBI |
| KX415336 | A/NewYork/29/2016(H3N2) | 21/2/2016 | NCBI |
| KX415344 | A/Idaho/11/2016(H3N2) | 20/2/2016 | NCBI |
| KX415352 | A/Utah/21/2016(H3N2) | 1/3/2016 | NCBI |
| KX415360 | A/Utah/18/2016(H3N2) | 28/2/2016 | NCBI |
| KX415368 | A/Montana/25/2016(H3N2) | 29/2/2016 | NCBI |
| KX415376 | A/NewMexico/09/2016(H3N2) | 21/2/2016 | NCBI |
| KX415384 | A/NewMexico/10/2016(H3N2) | 19/2/2016 | NCBI |
| KX415392 | A/Wisconsin/32/2016(H3N2) | 26/2/2016 | NCBI |
| KX415400 | A/Wisconsin/33/2016(H3N2) | 29/2/2016 | NCBI |
| KX415416 | A/Indiana/28/2016(H3N2) | 23/2/2016 | NCBI |
| KX415432 | A/Florida/28/2016(H3N2) | 2/3/2016 | NCBI |
| KX415440 | A/Florida/30/2016(H3N2) | 4/3/2016 | NCBI |

| | | | |
|---|---|---|---|
| KX415448 | A/Florida/31/2016(H3N2) | 26/2/2016 | NCBI |
| KX415464 | A/Arizona/44/2016(H3N2) | 16/2/2016 | NCBI |
| KX415472 | A/Arizona/41/2016(H3N2) | 13/2/2016 | NCBI |
| KX415480 | A/Arizona/41/2016(H3N2) | 13/2/2016 | NCBI |
| KX415488 | A/Louisiana/04/2016(H3N2) | 17/2/2016 | NCBI |
| KX415496 | A/Louisiana/07/2016(H3N2) | 24/2/2016 | NCBI |
| KX415504 | A/Colorado/18/2016(H3N2) | 1/3/2016 | NCBI |
| KX415512 | A/Colorado/19/2016(H3N2) | 1/3/2016 | NCBI |
| KX415520 | A/Texas/48/2016(H3N2) | 8/3/2016 | NCBI |
| KX415528 | A/Texas/49/2016(H3N2) | 10/3/2016 | NCBI |
| KX415536 | A/Michigan/44/2016(H3N2) | 17/2/2016 | NCBI |
| KX415544 | A/Michigan/47/2016(H3N2) | 7/3/2016 | NCBI |
| KX415552 | A/Michigan/47/2016(H3N2) | 7/3/2016 | NCBI |
| KX415560 | A/NewHampshire/07/2016(H3N2) | 11/2/2016 | NCBI |
| KX415568 | A/NewHampshire/09/2016(H3N2) | 9/3/2016 | NCBI |
| KX415576 | A/Minnesota/28/2016(H3N2) | 28/2/2016 | NCBI |
| KX415584 | A/Minnesota/30/2016(H3N2) | 5/3/2016 | NCBI |
| KX415592 | A/Connecticut/13/2016(H3N2) | 28/2/2016 | NCBI |
| KX415600 | A/Connecticut/13/2016(H3N2) | 28/2/2016 | NCBI |
| KX415608 | A/Connecticut/14/2016(H3N2) | 28/2/2016 | NCBI |
| KX415616 | A/Nevada/24/2016(H3N2) | 28/2/2016 | NCBI |
| KX415632 | A/NorthCarolina/14/2016(H3N2) | 10/3/2016 | NCBI |
| KX415650 | A/Pennsylvania/20/2016(H3N2) | 29/2/2016 | NCBI |
| KX415640 | A/Pennsylvania/20/2016(H3N2) | 29/2/2016 | NCBI |
| KX415664 | A/NorthDakota/19/2016(H3N2) | 5/3/2016 | NCBI |
| KX415656 | A/NorthDakota/19/2016(H3N2) | 5/3/2016 | NCBI |
| KX415672 | A/NorthDakota/20/2016(H3N2) | 6/3/2016 | NCBI |
| KX415680 | A/Kentucky/11/2016(H3N2) | 8/3/2016 | NCBI |
| KX415688 | A/Kansas/16/2016(H3N2) | 7/3/2016 | NCBI |
| KX415696 | A/Kansas/17/2016(H3N2) | 14/3/2016 | NCBI |
| KX415704 | A/Alabama/09/2016(H3N2) | 11/2/2016 | NCBI |
| KX415712 | A/Texas/52/2016(H3N2) | 18/2/2016 | NCBI |
| KX415720 | A/Texas/52/2016(H3N2) | 18/2/2016 | NCBI |
| KX415728 | A/Texas/54/2016(H3N2) | 28/2/2016 | NCBI |
| KX415737 | A/Texas/54/2016(H3N2) | 28/2/2016 | NCBI |
| KX415753 | A/Wisconsin/36/2016(H3N2) | 11/3/2016 | NCBI |
| KX415744 | A/Wisconsin/36/2016(H3N2) | 11/3/2016 | NCBI |

| | | | |
|---|---|---|---|
| KX415760 | A/Wisconsin/37/2016(H3N2) | 11/3/2016 | NCBI |
| KX415768 | A/Hawaii/17/2016(H3N2) | 26/2/2016 | NCBI |
| KX415776 | A/Hawaii/19/2016(H3N2) | 23/2/2016 | NCBI |
| KX415784 | A/California/69/2016(H3N2) | 24/2/2016 | NCBI |
| KX415792 | A/California/69/2016(H3N2) | 24/2/2016 | NCBI |
| KX415800 | A/California/70/2016(H3N2) | 8/2/2016 | NCBI |
| KX415808 | A/NewMexico/13/2016(H3N2) | 25/2/2016 | NCBI |
| KX415816 | A/Texas/56/2016(H3N2) | 15/2/2016 | NCBI |
| KX415824 | A/Texas/57/2016(H3N2) | 22/2/2016 | NCBI |
| KX415832 | A/Texas/58/2016(H3N2) | 10/2/2016 | NCBI |
| KX415848 | A/Texas/60/2016(H3N2) | 8/3/2016 | NCBI |
| KX415840 | A/Texas/60/2016(H3N2) | 8/3/2016 | NCBI |
| KX415856 | A/Texas/62/2016(H3N2) | 11/3/2016 | NCBI |
| KX415864 | A/Mississippi/11/2016(H3N2) | 5/3/2016 | NCBI |
| KX415872 | A/Mississippi/11/2016(H3N2) | 5/3/2016 | NCBI |
| KX415880 | A/Indiana/30/2016(H3N2) | 1/3/2016 | NCBI |
| KX415888 | A/Indiana/30/2016(H3N2) | 1/3/2016 | NCBI |
| KX415896 | A/Indiana/31/2016(H3N2) | 7/3/2016 | NCBI |
| KX415912 | A/California/74/2016(H3N2) | 28/2/2016 | NCBI |
| KX415904 | A/California/74/2016(H3N2) | 28/2/2016 | NCBI |
| KX415920 | A/California/75/2016(H3N2) | 20/1/2016 | NCBI |
| KX415928 | A/California/79/2016(H3N2) | 15/2/2016 | NCBI |
| KX415936 | A/California/80/2016(H3N2) | 15/2/2016 | NCBI |
| KX415944 | A/California/82/2016(H3N2) | 16/2/2016 | NCBI |
| KX415952 | A/California/88/2016(H3N2) | 25/2/2016 | NCBI |
| KX415960 | A/California/92/2016(H3N2) | 3/3/2016 | NCBI |
| KX415968 | A/California/94/2016(H3N2) | 29/2/2016 | NCBI |
| KX415976 | A/NewMexico/22/2016(H3N2) | 8/2/2016 | NCBI |
| KX415984 | A/Utah/23/2016(H3N2) | 8/3/2016 | NCBI |
| KX415992 | A/Idaho/14/2016(H3N2) | 4/3/2016 | NCBI |
| KX416000 | A/Idaho/16/2016(H3N2) | 7/3/2016 | NCBI |
| KX416008 | A/Nevada/20/2016(H3N2) | 10/3/2016 | NCBI |
| KX416016 | A/Nevada/21/2016(H3N2) | 9/3/2016 | NCBI |
| KX416024 | A/Florida/35/2016(H3N2) | 7/3/2016 | NCBI |
| KX416032 | A/Florida/38/2016(H3N2) | 13/3/2016 | NCBI |
| KX416040 | A/Florida/42/2016(H3N2) | 16/3/2016 | NCBI |
| KX416048 | A/Florida/43/2016(H3N2) | 18/3/2016 | NCBI |

| | | | |
|---|---|---|---|
| KX416056 | A/Virginia/39/2016(H3N2) | 10/3/2016 | NCBI |
| KX416064 | A/Wyoming/07/2016(H3N2) | 10/3/2016 | NCBI |
| KX416072 | A/Wyoming/08/2016(H3N2) | 14/3/2016 | NCBI |
| KX416080 | A/Pennsylvania/37/2016(H3N2) | 18/3/2016 | NCBI |
| KX416088 | A/Pennsylvania/41/2016(H3N2) | 10/3/2016 | NCBI |
| KX416096 | A/Colorado/22/2016(H3N2) | 6/3/2016 | NCBI |
| KX416104 | A/Colorado/23/2016(H3N2) | 8/3/2016 | NCBI |
| KX416112 | A/Washington/46/2016(H3N2) | 9/3/2016 | NCBI |
| KX416120 | A/Washington/47/2016(H3N2) | 14/3/2016 | NCBI |
| KX416128 | A/NewYork/43/2016(H3N2) | 29/2/2016 | NCBI |
| KX416136 | A/NewHampshire/11/2016(H3N2) | 18/3/2016 | NCBI |
| KX416144 | A/NewHampshire/13/2016(H3N2) | 18/3/2016 | NCBI |
| KX416152 | A/Louisiana/09/2016(H3N2) | 7/3/2016 | NCBI |
| KX416160 | A/Michigan/59/2016(H3N2) | 22/2/2016 | NCBI |
| KX416176 | A/Texas/118/2016(H3N2) | 16/3/2016 | NCBI |
| KX416184 | A/Texas/119/2016(H3N2) | 24/3/2016 | NCBI |
| KX416192 | A/NorthCarolina/35/2016(H3N2) | 21/3/2016 | NCBI |
| KX416200 | A/NorthCarolina/36/2016(H3N2) | 24/3/2016 | NCBI |
| KX416208 | A/Delaware/15/2016(H3N2) | 20/3/2016 | NCBI |
| KX416216 | A/Delaware/16/2016(H3N2) | 28/3/2016 | NCBI |
| KX416224 | A/Kentucky/14/2016(H3N2) | 8/3/2016 | NCBI |
| KX416232 | A/Kentucky/15/2016(H3N2) | 9/3/2016 | NCBI |
| KX416240 | A/Oregon/10/2016(H3N2) | 15/3/2016 | NCBI |
| KX416248 | A/Oregon/12/2016(H3N2) | 18/3/2016 | NCBI |
| KX416256 | A/Texas/120/2016(H3N2) | 20/3/2016 | NCBI |
| KX416264 | A/Virginia/42/2016(H3N2) | 21/3/2016 | NCBI |
| KX416272 | A/Virginia/43/2016(H3N2) | 20/3/2016 | NCBI |
| KX416280 | A/NewMexico/29/2016(H3N2) | 20/3/2016 | NCBI |
| KX416288 | A/NewMexico/30/2016(H3N2) | 20/3/2016 | NCBI |
| KX416296 | A/NewJersey/15/2016(H3N2) | 27/3/2016 | NCBI |
| KX416304 | A/Wisconsin/52/2016(H3N2) | 16/3/2016 | NCBI |
| KX416312 | A/Wisconsin/53/2016(H3N2) | 21/3/2016 | NCBI |
| KX416320 | A/Texas/126/2016(H3N2) | 21/3/2016 | NCBI |
| KX416328 | A/Maryland/11/2016(H3N2) | 24/2/2016 | NCBI |
| KX416336 | A/Iowa/02/2016(H3N2) | 3/3/2016 | NCBI |
| KX416344 | A/Iowa/03/2016(H3N2) | 23/2/2016 | NCBI |
| KX416352 | A/Utah/28/2016(H3N2) | 24/3/2016 | NCBI |

| | | | |
|---|---|---|---|
| KX416360 | A/Utah/29/2016(H3N2) | 23/3/2016 | NCBI |
| KX416368 | A/Michigan/64/2016(H3N2) | 3/3/2016 | NCBI |
| KX416376 | A/Michigan/65/2016(H3N2) | 4/3/2016 | NCBI |
| KX416384 | A/Missouri/10/2016(H3N2) | 23/3/2016 | NCBI |
| KX416392 | A/Indiana/34/2016(H3N2) | 20/3/2016 | NCBI |
| KX416400 | A/Indiana/35/2016(H3N2) | 21/3/2016 | NCBI |
| KX416408 | A/Minnesota/43/2016(H3N2) | 28/3/2016 | NCBI |
| KX416416 | A/Florida/44/2016(H3N2) | 30/3/2016 | NCBI |
| KX416424 | A/Florida/47/2016(H3N2) | 28/3/2016 | NCBI |
| KX416432 | A/Hawaii/28/2016(H3N2) | 29/2/2016 | NCBI |
| KX416440 | A/Hawaii/29/2016(H3N2) | 1/3/2016 | NCBI |
| KX416456 | A/Florida/50/2016(H3N2) | 26/3/2016 | NCBI |
| KX416448 | A/Florida/50/2016(H3N2) | 26/3/2016 | NCBI |
| KX416464 | A/Florida/51/2016(H3N2) | 29/3/2016 | NCBI |
| KX416472 | A/Idaho/17/2016(H3N2) | 15/3/2016 | NCBI |
| KX416480 | A/Idaho/20/2016(H3N2) | 2016 | NCBI |
| KX416488 | A/Wyoming/10/2016(H3N2) | 21/3/2016 | NCBI |
| KX416496 | A/Wyoming/14/2016(H3N2) | 22/3/2016 | NCBI |
| KX416504 | A/Wyoming/15/2016(H3N2) | 23/3/2016 | NCBI |
| KX416512 | A/Colorado/26/2016(H3N2) | 15/3/2016 | NCBI |
| KX416520 | A/Colorado/27/2016(H3N2) | 27/3/2016 | NCBI |
| KX416528 | A/NorthDakota/23/2016(H3N2) | 15/3/2016 | NCBI |
| KX416536 | A/NorthDakota/26/2016(H3N2) | 22/3/2016 | NCBI |
| KX416544 | A/Arizona/49/2016(H3N2) | 6/3/2016 | NCBI |
| KX416552 | A/Arizona/50/2016(H3N2) | 5/3/2016 | NCBI |
| KX416560 | A/Massachusetts/28/2016(H3N2) | 28/3/2016 | NCBI |
| KX416568 | A/Massachusetts/28/2016(H3N2) | 28/3/2016 | NCBI |
| KX416576 | A/NewJersey/17/2016(H3N2) | 6/4/2016 | NCBI |
| KX416584 | A/NewJersey/18/2016(H3N2) | 2/4/2016 | NCBI |
| KX416592 | A/NewJersey/18/2016(H3N2) | 2/4/2016 | NCBI |
| KX416600 | A/Iowa/09/2016(H3N2) | 23/3/2016 | NCBI |
| KX416608 | A/Iowa/10/2016(H3N2) | 3/4/2016 | NCBI |
| KX416616 | A/NewHampshire/15/2016(H3N2) | 10/3/2016 | NCBI |
| KX416624 | A/NewHampshire/16/2016(H3N2) | 30/3/2016 | NCBI |
| KX416632 | A/Illinois/07/2016(H3N2) | 1/2/2016 | NCBI |
| KX416640 | A/Illinois/08/2016(H3N2) | 8/2/2016 | NCBI |
| KX416648 | A/Illinois/09/2016(H3N2) | 9/2/2016 | NCBI |

| | | | |
|---|---|---|---|
| KX416656 | A/Illinois/10/2016(H3N2) | 10/2/2016 | NCBI |
| KX416664 | A/Illinois/11/2016(H3N2) | 1/2/2016 | NCBI |
| KX416672 | A/Illinois/12/2016(H3N2) | 30/1/2016 | NCBI |
| KX416680 | A/California/96/2016(H3N2) | 15/3/2016 | NCBI |
| KX416688 | A/California/98/2016(H3N2) | 20/3/2016 | NCBI |
| KX416696 | A/Nevada/30/2016(H3N2) | 3/4/2016 | NCBI |
| KX416704 | A/Maryland/15/2016(H3N2) | 11/3/2016 | NCBI |
| KX416712 | A/Kansas/22/2016(H3N2) | 2/4/2016 | NCBI |
| KX416720 | A/Kansas/25/2016(H3N2) | 8/4/2016 | NCBI |
| KX416728 | A/Pennsylvania/65/2016(H3N2) | 4/4/2016 | NCBI |
| KX416736 | A/NewYork/56/2016(H3N2) | 2/3/2016 | NCBI |
| KX416744 | A/NewYork/57/2016(H3N2) | 7/3/2016 | NCBI |
| KX416752 | A/NewYork/58/2016(H3N2) | 20/3/2016 | NCBI |
| KX416760 | A/Michigan/66/2016(H3N2) | 15/3/2016 | NCBI |
| KX416768 | A/Michigan/70/2016(H3N2) | 4/4/2016 | NCBI |
| KX416776 | A/NewMexico/33/2016(H3N2) | 27/3/2016 | NCBI |
| KX416784 | A/NewMexico/34/2016(H3N2) | 29/3/2016 | NCBI |
| KX416792 | A/Louisiana/12/2016(H3N2) | 29/3/2016 | NCBI |
| KX416800 | A/Louisiana/13/2016(H3N2) | 29/3/2016 | NCBI |
| KX416808 | A/Hawaii/32/2016(H3N2) | 2/3/2016 | NCBI |
| KX416816 | A/Hawaii/33/2016(H3N2) | 5/3/2016 | NCBI |
| KX416824 | A/Washington/52/2016(H3N2) | 11/4/2016 | NCBI |
| KX416832 | A/Washington/55/2016(H3N2) | 12/4/2016 | NCBI |
| KX416840 | A/Utah/32/2016(H3N2) | 7/4/2016 | NCBI |
| KX416848 | A/Utah/33/2016(H3N2) | 7/4/2016 | NCBI |
| KX416856 | A/Indiana/40/2016(H3N2) | 4/4/2016 | NCBI |
| KX416864 | A/Florida/55/2016(H3N2) | 5/4/2016 | NCBI |
| KX416872 | A/Vermont/16/2016(H3N2) | 15/4/2016 | NCBI |
| KX416880 | A/Kentucky/21/2016(H3N2) | 7/4/2016 | NCBI |
| KX416888 | A/Kentucky/20/2016(H3N2) | 4/4/2016 | NCBI |
| KX416896 | A/SouthDakota/19/2016(H3N2) | 28/2/2016 | NCBI |
| KX416904 | A/SouthDakota/20/2016(H3N2) | 28/2/2016 | NCBI |
| KX416912 | A/SouthDakota/26/2016(H3N2) | 15/4/2016 | NCBI |
| KX416920 | A/Louisiana/17/2016(H3N2) | 18/3/2016 | NCBI |
| KX416928 | A/NorthCarolina/44/2016(H3N2) | 12/4/2016 | NCBI |
| KX416936 | A/NorthCarolina/46/2016(H3N2) | 19/4/2016 | NCBI |
| KX416944 | A/Texas/133/2016(H3N2) | 13/4/2016 | NCBI |

| | | | |
|---|---|---|---|
| KX416952 | A/Texas/134/2016(H3N2) | 20/4/2016 | NCBI |
| KX416960 | A/Iowa/12/2016(H3N2) | 28/3/2016 | NCBI |
| KX416968 | A/Texas/130/2016(H3N2) | 2/4/2016 | NCBI |
| KX416976 | A/Texas/131/2016(H3N2) | 7/4/2016 | NCBI |
| KX416984 | A/Wisconsin/61/2016(H3N2) | 15/4/2016 | NCBI |
| KX416992 | A/Wisconsin/62/2016(H3N2) | 18/4/2016 | NCBI |
| KX417000 | A/Minnesota/59/2016(H3N2) | 24/4/2016 | NCBI |
| KX417008 | A/Texas/136/2016(H3N2) | 22/4/2016 | NCBI |
| KX417016 | A/Iowa/15/2016(H3N2) | 20/4/2016 | NCBI |
| KX417024 | A/Wisconsin/67/2016(H3N2) | 3/5/2016 | NCBI |
| KX417032 | A/Missouri/13/2016(H3N2) | 27/4/2016 | NCBI |
| KX571022 | A/Brisbane/E1/2007(H3N2) | 12/2/2007 | NCBI |
| KT888780 | A/Iquitos/CFI1471/2011(H3N2) | 3/11/2011 | NCBI |
| KT888781 | A/Iquitos/CFI1510/2011(H3N2) | 30/11/2011 | NCBI |
| KT888782 | A/Iquitos/CFI1576/2012(H3N2) | 5/1/2012 | NCBI |
| KT888783 | A/Tegucigalpa/FHT00191/2011(H3N2) | 4/10/2011 | NCBI |
| KT888784 | A/Iquitos/FPI002788/2011(H3N2) | 1/12/2011 | NCBI |
| KT888785 | A/PuertoMaldonado/FPM00365/2011(H3N2) | 17/10/2011 | NCBI |
| KT888786 | A/Piura/FPP00919/2011(H3N2) | 25/10/2011 | NCBI |
| KT888787 | A/Piura/FPP01106/2011(H3N2) | 29/12/2011 | NCBI |
| KT888788 | A/Pucallpa/FPU00363/2011(H3N2) | 19/12/2011 | NCBI |
| KT888789 | A/Pucallpa/FPU00370/2011(H3N2) | 27/12/2011 | NCBI |
| KT888790 | A/Masaya/INI00222/2011(H3N2) | 13/10/2011 | NCBI |
| KT888791 | A/Masaya/INI00224/2011(H3N2) | 14/10/2011 | NCBI |
| KT888792 | A/Managua/INI00283/2011(H3N2) | 23/10/2011 | NCBI |
| KT888793 | A/Trujillo/IPE00693/2011(H3N2) | 7/10/2011 | NCBI |
| KT888794 | A/Puno/IPE00792/2011(H3N2) | 25/11/2011 | NCBI |
| KT888795 | A/Callao/IPE00830/2012(H3N2) | 10/1/2012 | NCBI |
| KT888796 | A/Callao/IPE00833/2012(H3N2) | 11/1/2012 | NCBI |
| KT889202 | A/San_Diego/BRD11439/2011(H3N2) | 2/11/2011 | NCBI |
| KT889204 | A/California/BRD11499N/2012(H3N2) | 30/1/2012 | NCBI |
| KT889205 | A/California/BRD11503N/2012(H3N2) | 30/1/2012 | NCBI |
| KT889207 | A/San_Diego/FDX11070/2011(H3N2) | 15/12/2011 | NCBI |
| KT889209 | A/California/FDX21048/2012(H3N2) | 2012 | NCBI |
| KT889210 | A/Illinois/FDX50085/2012(H3N2) | 2012 | NCBI |
| KT889211 | A/Illinois/FDX50179/2012(H3N2) | 2012 | NCBI |
| KT889212 | A/California/SAR40343N/2012(H3N2) | 2012 | NCBI |

| | | | |
|---|---|---|---|
| KT889213 | A/Yokosuka/SE680/2012(H3N2) | 24/1/2012 | NCBI |
| KT889214 | A/Yokosuka/SE863/2012(H3N2) | 3/2/2012 | NCBI |
| KT889230 | A/Moscow/1514A00504307T_CP2/2011(H3N2) | 7/2/2011 | NCBI |
| KT889231 | A/Moscow/1514A00504942N/2012(H3N2) | 19/1/2012 | NCBI |
| KT889232 | A/Moscow/1514A00504950T/2012(H3N2) | 11/1/2012 | NCBI |
| KT889233 | A/Moscow/1514A00505149T/2012(H3N2) | 12/4/2012 | NCBI |
| KT889234 | A/Moscow/1514A00505153N/2012(H3N2) | 23/3/2012 | NCBI |
| KT889235 | A/Moscow/1514A00505162N/2012(H3N2) | 13/4/2012 | NCBI |
| KT889236 | A/MexicoCity/1514A00904139N_CP2/2010(H3N2) | 29/12/2010 | NCBI |
| KT889237 | A/MexicoCity/1514A00905313N/2013(H3N2) | 12/3/2013 | NCBI |
| KT889238 | A/MexicoCity/1514A00905315N/2013(H3N2) | 11/3/2013 | NCBI |
| KT889239 | A/MexicoCity/1514A00905321T/2013(H3N2) | 26/2/2013 | NCBI |
| KT889240 | A/MexicoCity/1514A00905324N/2013(H3N2) | 4/1/2013 | NCBI |
| KT889241 | A/Amman/1514A01004462T/2012(H3N2) | 11/1/2012 | NCBI |
| KT889242 | A/Amman/1514A01004465T/2011(H3N2) | 5/12/2011 | NCBI |
| KT889243 | A/Kabul/1514A01305424T/2013(H3N2) | 16/1/2013 | NCBI |
| KT889244 | A/Kabul/1514A01305426N/2013(H3N2) | 13/1/2013 | NCBI |
| KT889245 | A/Kabul/1514A01305429N/2013(H3N2) | 10/1/2013 | NCBI |
| KT889246 | A/Kabul/1514A01305430N/2013(H3N2) | 7/1/2013 | NCBI |
| KT889247 | A/Kabul/1514A01305435T/2013(H3N2) | 10/1/2013 | NCBI |
| KT889248 | A/Kabul/1514A01305436N/2013(H3N2) | 15/1/2013 | NCBI |
| KT889249 | A/HongKong/1514A01704826T/2011(H3N2) | 15/11/2011 | NCBI |
| KT889250 | A/Taipei/1514A02605215N/2013(H3N2) | 7/5/2013 | NCBI |
| KT889251 | A/Taipei/1514A02605215T/2013(H3N2) | 7/5/2013 | NCBI |
| KT889252 | A/TelAviv/1514A02705506T/2013(H3N2) | 31/1/2013 | NCBI |
| KT889253 | A/Prague/1514A06404885N/2012(H3N2) | 12/4/2012 | NCBI |
| KT889254 | A/Pristina/1514A06505401N/2013(H3N2) | 6/2/2013 | NCBI |
| KT889255 | A/Sarajevo/1514A06604486N/2012(H3N2) | 11/1/2012 | NCBI |
| KT889256 | A/AddisAbaba/1514A07305892N/2013(H3N2) | 26/9/2013 | NCBI |
| KT889257 | A/Budapest/1514A07804502N/2011(H3N2) | 12/12/2011 | NCBI |
| KT889258 | A/Budapest/1514A07805419T/2013(H3N2) | 5/2/2013 | NCBI |
| KT889259 | A/Budapest/1514A07805832T/2013(H3N2) | 6/2/2013 | NCBI |
| KT889260 | A/Thailand/1552AN015808_CP2/2010(H3N2) | 1/11/2010 | NCBI |
| KT889261 | A/Kuwait/F072/2014(H3N2) | 2014-01 | NCBI |
| KT889262 | A/Kuwait/F074/2014(H3N2) | 2014-01 | NCBI |
| KT889263 | A/Kuwait/F075/2014(H3N2) | 2014-01 | NCBI |
| KT889264 | A/Kuwait/F078/2014(H3N2) | 2014-01 | NCBI |

| | | | |
|---|---|---|---|
| KT889265 | A/Kuwait/F083/2014(H3N2) | 2014-01 | NCBI |
| KT889266 | A/Kuwait/F089/2014(H3N2) | 2014-01 | NCBI |
| KT889267 | A/Kuwait/F093/2014(H3N2) | 2014-01 | NCBI |
| KT889268 | A/Kuwait/F099/2014(H3N2) | 2014-01 | NCBI |
| KT889269 | A/Kuwait/F102/2014(H3N2) | 2014-01 | NCBI |
| KT889270 | A/Kuwait/F123/2014(H3N2) | 2014-02 | NCBI |
| KT889271 | A/Kuwait/F124/2014(H3N2) | 2014-02 | NCBI |
| KT889272 | A/Kuwait/F142/2014(H3N2) | 2014-02 | NCBI |
| KT889273 | A/Kuwait/F146/2014(H3N2) | 2014-02 | NCBI |
| KT889274 | A/Kuwait/F147/2014(H3N2) | 2014-02 | NCBI |
| KT889275 | A/Kuwait/F167/2014(H3N2) | 2014-03 | NCBI |
| KT889129 | A/Japan/17/2012(H3N2) | 31/1/2012 | NCBI |
| KT889130 | A/Korea/18/2012(H3N2) | 24/1/2012 | NCBI |
| KT889131 | A/Illinois/19/2012(H3N2) | 15/2/2012 | NCBI |
| KT889132 | A/Nebraska/20/2012(H3N2) | 8/2/2012 | NCBI |
| KT889133 | A/Alaska/21usafsam/2012(H3N2) | 29/1/2012 | NCBI |
| KT889134 | A/Japan/22/2011(H3N2) | 20/12/2011 | NCBI |
| KT889135 | A/NorthCarolina/23/2012(H3N2) | 28/2/2012 | NCBI |
| KT889136 | A/NorthCarolina/24usafsam/2012(H3N2) | 16/2/2012 | NCBI |
| KT889137 | A/Nevada/25usafsam/2012(H3N2) | 5/3/2012 | NCBI |
| KT889138 | A/Japan/26/2012(H3N2) | 5/3/2012 | NCBI |
| KT889139 | A/Alaska/27/2012(H3N2) | 6/3/2012 | NCBI |
| KT889140 | A/NewYork/28/2012(H3N2) | 6/3/2012 | NCBI |
| KT889141 | A/Washington/29/2012(H3N2) | 17/2/2012 | NCBI |
| KT889142 | A/Florida/30/2012(H3N2) | 2/3/2012 | NCBI |
| KT889143 | A/Oklahoma/31/2012(H3N2) | 1/3/2012 | NCBI |
| KT889144 | A/Nebraska/32/2012(H3N2) | 29/2/2012 | NCBI |
| KT889145 | A/Illinois/34/2012(H3N2) | 5/3/2012 | NCBI |
| KT889146 | A/Korea/37/2012(H3N2) | 19/1/2012 | NCBI |
| KT889147 | A/Colorado/38/2012(H3N2) | 13/2/2012 | NCBI |
| KT889148 | A/Washington/39/2012(H3N2) | 14/2/2012 | NCBI |
| KT889149 | A/Kyrgyzstan/40/2012(H3N2) | 12/1/2012 | NCBI |
| KT889150 | A/Kyrgyzstan/41/2012(H3N2) | 12/1/2012 | NCBI |
| KT889151 | A/Japan/42/2012(H3N2) | 31/1/2012 | NCBI |
| KT889152 | A/Japan/43/2012(H3N2) | 5/2/2012 | NCBI |
| KT889153 | A/Colorado/44/2012(H3N2) | 9/1/2012 | NCBI |
| KT889154 | A/Korea/45/2012(H3N2) | 13/1/2012 | NCBI |

| | | | |
|---|---|---|---|
| KT889155 | A/Korea/47/2012(H3N2) | 20/1/2012 | NCBI |
| KT889156 | A/Nebraska/48/2012(H3N2) | 9/1/2012 | NCBI |
| KT889158 | A/Texas/55121/2012(H3N2) | 2012 | NCBI |
| KT888462 | A/Kuwait/F179-9291/2013(H3N2) | 2013-11 | NCBI |
| KT888463 | A/Kuwait/F181-3/2014(H3N2) | 2014-02 | NCBI |
| KT888602 | A/Philippines/FLU_AFPA_00013/2011(H3N2) | 10/10/2011 | NCBI |
| KT888603 | A/Philippines/FLU_AFPA_00063/2012(H3N2) | 1/1/2012 | NCBI |
| KT888604 | A/Bhutan/FLU_BTA_00010/2011(H3N2) | 27/9/2011 | NCBI |
| KT888605 | A/Bhutan/FLU_BTD_SV0023_CP2/2009(H3N2) | 15/1/2009 | NCBI |
| KT888606 | A/Bhutan/FLU_BTI_00001/2011(H3N2) | 27/9/2011 | NCBI |
| KT888607 | A/Philippines/FLU_PHB_0332/2011(H3N2) | 22/11/2011 | NCBI |
| KT888608 | A/Philippines/FLU_PHB_0334/2011(H3N2) | 29/11/2011 | NCBI |
| KT888609 | A/Philippines/FLU_PHB_0341/2011(H3N2) | 6/12/2011 | NCBI |
| KT888610 | A/Philippines/FLU_PHC_0105_CP2/2009(H3N2) | 31/3/2009 | NCBI |
| KT888611 | A/Bulacao/FLU_PHC_0275/2011(H3N2) | 15/11/2011 | NCBI |
| KT888612 | A/Philippines/FLU_PHC_0284/2011(H3N2) | 29/11/2011 | NCBI |
| KT888613 | A/Bulacao/FLU_PHC_0286/2011(H3N2) | 29/11/2011 | NCBI |
| KT888614 | A/Bulacao/FLU_PHC_0291/2011(H3N2) | 6/12/2011 | NCBI |
| KT888615 | A/Philippines/FLU_PHC_0304/2011(H3N2) | 20/12/2011 | NCBI |
| KT888616 | A/Philippines/FLU_PHD_0142_CP2/2009(H3N2) | 27/7/2009 | NCBI |
| KT888617 | A/Bangkok/FLU_PMKA0055_CP2/2009(H3N2) | 9/11/2009 | NCBI |
| KT888618 | A/Bangkok/FLU_PMKA0967_CP3/2010(H3N2) | 26/10/2010 | NCBI |
| KT888619 | A/Bangkok/FLU_PMKA1169/2011(H3N2) | 25/3/2011 | NCBI |
| KT888620 | A/Bangkok/FLU_PMKA1400/2011(H3N2) | 28/7/2011 | NCBI |
| KT888621 | A/Bangkok/FLU_PMKA1455/2011(H3N2) | 5/8/2011 | NCBI |
| KT888622 | A/Bangkok/FLU_PMKA1464/2011(H3N2) | 8/8/2011 | NCBI |
| KT888623 | A/Bangkok/FLU_PMKA1475/2011(H3N2) | 9/8/2011 | NCBI |
| KT888624 | A/Bangkok/FLU_PMKA1480/2011(H3N2) | 10/8/2011 | NCBI |
| KT888625 | A/Thailand/FLU_PMKA1493/2011(H3N2) | 11/8/2011 | NCBI |
| KT888626 | A/Bangkok/FLU_PMKA1561/2011(H3N2) | 25/8/2011 | NCBI |
| KT888627 | A/Bangkok/FLU_PMKA1573/2011(H3N2) | 29/8/2011 | NCBI |
| KT888628 | A/Thailand/FLU_PMKA1633/2011(H3N2) | 14/9/2011 | NCBI |
| KT888629 | A/Thailand/FLU_PMKA1835/2012(H3N2) | 20/2/2012 | NCBI |
| KT888630 | A/Bangkok/FLU_PMKB0204/2011(H3N2) | 18/8/2011 | NCBI |
| KT888631 | A/Thailand/FLU_PMKD0388/2011(H3N2) | 14/9/2011 | NCBI |
| KT888632 | A/Myanmar/FLUEBMSV00004/2011(H3N2) | 21/9/2011 | NCBI |
| KT888633 | A/India/FLUEINSV0056/2011(H3N2) | 12/9/2011 | NCBI |

| | | | |
|---|---|---|---|
| KT888634 | A/Jakarta/FLUEJKSV0024/2012(H3N2) | 12/1/2012 | NCBI |
| KT888635 | A/Jakarta/FLUEJKSV0025/2012(H3N2) | 12/1/2012 | NCBI |
| KT888636 | A/Indonesia/FLUEJKSV0026/2012(H3N2) | 13/1/2012 | NCBI |
| KT888637 | A/Jakarta/FLUEJKSV0031/2012(H3N2) | 1/2/2012 | NCBI |
| KT888638 | A/Jakarta/FLUEJKSV0033/2012(H3N2) | 13/2/2012 | NCBI |
| KT888639 | A/Thailand/FLUETBSV0120/2012(H3N2) | 14/2/2012 | NCBI |
| KT888640 | A/Kathmandu/FLUNP09A_387_CP3/2009(H3N2) | 28/7/2009 | NCBI |
| KT888641 | A/Kathmandu/FLUNP09M_299_CP2/2009(H3N2) | 10/7/2009 | NCBI |
| KT888642 | A/Kathmandu/FLUNP09N_001_CP2/2009(H3N2) | 20/5/2009 | NCBI |
| KT888643 | A/Thailand/SV0466_11/2011(H3N2) | 19/8/2011 | NCBI |
| KT888644 | A/Bangkok/SV0471_11/2011(H3N2) | 25/8/2011 | NCBI |
| KX885891 | A/Texas/88/2016(H3N2) | 24/2/2016 | NCBI |
| KX885880 | A/Idaho/33/2016(H3N2) | 7/6/2016 | NCBI |
| KX885896 | A/Nebraska/19/2015(H3N2) | 23/10/2015 | NCBI |
| CY207299 | A/NewYork/WC-LVD-14-064/2014(H3N2) | 4/11/2014 | NCBI |
| CY207307 | A/NewYork/WC-LVD-14-066/2014(H3N2) | 29/12/2014 | NCBI |
| CY207315 | A/NewYork/WC-LVD-14-067/2014(H3N2) | 24/11/2014 | NCBI |
| CY207323 | A/NewYork/WC-LVD-14-069/2014(H3N2) | 2/12/2014 | NCBI |
| CY207331 | A/NewYork/WC-LVD-14-070/2014(H3N2) | 2/12/2014 | NCBI |
| CY207339 | A/NewYork/WC-LVD-14-071/2014(H3N2) | 8/12/2014 | NCBI |
| CY207347 | A/NewYork/WC-LVD-14-072/2014(H3N2) | 10/12/2014 | NCBI |
| CY207355 | A/NewYork/WC-LVD-14-073/2014(H3N2) | 3/12/2014 | NCBI |
| CY207363 | A/NewYork/WC-LVD-14-074/2014(H3N2) | 17/12/2014 | NCBI |
| CY207371 | A/NewYork/WC-LVD-14-075/2014(H3N2) | 21/12/2014 | NCBI |
| CY207379 | A/NewYork/WC-LVD-14-076/2014(H3N2) | 21/12/2014 | NCBI |
| CY207387 | A/NewYork/WC-LVD-14-077/2014(H3N2) | 22/12/2014 | NCBI |
| CY207395 | A/NewYork/WC-LVD-14-079/2014(H3N2) | 22/12/2014 | NCBI |
| CY207403 | A/NewYork/WC-LVD-14-080/2014(H3N2) | 25/12/2014 | NCBI |
| CY207411 | A/NewYork/WC-LVD-14-081/2014(H3N2) | 28/12/2014 | NCBI |
| CY207419 | A/NewYork/WC-LVD-15-001/2015(H3N2) | 7/1/2015 | NCBI |
| CY207427 | A/NewYork/WC-LVD-15-002/2015(H3N2) | 4/1/2015 | NCBI |
| CY207435 | A/NewYork/WC-LVD-15-003/2015(H3N2) | 4/1/2015 | NCBI |
| CY207443 | A/NewYork/WC-LVD-15-004/2015(H3N2) | 4/1/2015 | NCBI |
| CY207451 | A/NewYork/WC-LVD-14-082/2014(H3N2) | 22/12/2014 | NCBI |
| CY207459 | A/NewYork/WC-LVD-14-083/2014(H3N2) | 23/12/2014 | NCBI |
| CY207467 | A/NewYork/WC-LVD-14-084/2014(H3N2) | 30/12/2014 | NCBI |
| CY207475 | A/NewYork/WC-LVD-14-085/2014(H3N2) | 30/12/2014 | NCBI |

| | | | |
|---|---|---|---|
| CY207483 | A/NewYork/WC-LVD-14-086/2014(H3N2) | 24/12/2014 | NCBI |
| CY207491 | A/NewYork/WC-LVD-15-005/2015(H3N2) | 15/1/2015 | NCBI |
| CY207499 | A/NewYork/WC-LVD-15-006/2015(H3N2) | 19/1/2015 | NCBI |
| CY207507 | A/NewYork/WC-LVD-14-087/2014(H3N2) | 27/12/2014 | NCBI |
| CY207515 | A/NewYork/WC-LVD-15-007/2015(H3N2) | 5/1/2015 | NCBI |
| CY207523 | A/NewYork/WC-LVD-15-008/2015(H3N2) | 14/1/2015 | NCBI |
| CY207531 | A/NewYork/WC-LVD-15-009/2015(H3N2) | 8/1/2015 | NCBI |
| CY207539 | A/NewYork/WC-LVD-15-011/2015(H3N2) | 14/1/2015 | NCBI |
| CY207547 | A/NewYork/WC-LVD-15-012/2015(H3N2) | 14/1/2015 | NCBI |
| CY207555 | A/NewYork/WC-LVD-14-088/2014(H3N2) | 29/12/2014 | NCBI |
| CY207563 | A/NewYork/WC-LVD-14-089/2014(H3N2) | 28/12/2014 | NCBI |
| CY207571 | A/NewYork/WC-LVD-14-090/2014(H3N2) | 13/12/2014 | NCBI |
| CY207579 | A/NewYork/WC-LVD-15-013/2015(H3N2) | 20/1/2015 | NCBI |
| CY207587 | A/NewYork/WC-LVD-14-091/2014(H3N2) | 26/12/2014 | NCBI |
| CY207595 | A/NewYork/WC-LVD-15-014/2015(H3N2) | 19/1/2015 | NCBI |
| CY207603 | A/NewYork/WC-LVD-15-015/2015(H3N2) | 16/1/2015 | NCBI |
| CY207611 | A/NewYork/WC-LVD-14-092/2014(H3N2) | 14/12/2014 | NCBI |
| CY207619 | A/NewYork/WC-LVD-14-093/2014(H3N2) | 21/12/2014 | NCBI |
| CY207627 | A/NewYork/WC-LVD-14-094/2014(H3N2) | 26/12/2014 | NCBI |
| CY207635 | A/NewYork/WC-LVD-15-016/2015(H3N2) | 28/1/2015 | NCBI |
| CY207643 | A/NewYork/WC-LVD-15-017/2015(H3N2) | 28/1/2015 | NCBI |
| CY207651 | A/NewYork/WC-LVD-15-018/2015(H3N2) | 14/1/2015 | NCBI |
| CY207659 | A/NewYork/WC-LVD-15-020/2015(H3N2) | 3/2/2015 | NCBI |
| CY207667 | A/NewYork/WC-LVD-15-021/2015(H3N2) | 2/2/2015 | NCBI |
| CY207675 | A/NewYork/WC-LVD-15-022/2015(H3N2) | 25/1/2015 | NCBI |
| CY207683 | A/NewYork/WC-LVD-15-024/2015(H3N2) | 30/1/2015 | NCBI |
| CY207691 | A/NewYork/WC-LVD-15-025/2015(H3N2) | 4/2/2015 | NCBI |
| CY207699 | A/NewYork/WC-LVD-15-026/2015(H3N2) | 3/2/2015 | NCBI |
| CY207707 | A/NewYork/WC-LVD-15-027/2015(H3N2) | 23/2/2015 | NCBI |
| CY207715 | A/NewYork/WC-LVD-15-028/2015(H3N2) | 28/1/2015 | NCBI |
| CY207723 | A/NewYork/WC-LVD-15-029/2015(H3N2) | 10/2/2015 | NCBI |
| CY207731 | A/NewYork/WC-LVD-15-031/2015(H3N2) | 11/2/2015 | NCBI |
| CY207739 | A/NewYork/WC-LVD-15-032/2015(H3N2) | 18/2/2015 | NCBI |
| CY207747 | A/NewYork/WC-LVD-15-033/2015(H3N2) | 17/2/2015 | NCBI |
| CY207755 | A/NewYork/WC-LVD-15-034/2015(H3N2) | 16/1/2015 | NCBI |
| CY207763 | A/NewYork/WC-LVD-15-035/2015(H3N2) | 27/1/2015 | NCBI |
| CY207771 | A/NewYork/WC-LVD-15-037/2015(H3N2) | 3/3/2015 | NCBI |

| | | | |
|---|---|---|---|
| CY207779 | A/NewYork/WC-LVD-15-038/2015(H3N2) | 4/3/2015 | NCBI |
| CY207787 | A/NewYork/WC-LVD-15-039/2015(H3N2) | 8/3/2015 | NCBI |
| CY207795 | A/NewYork/WC-LVD-15-040/2015(H3N2) | 7/3/2015 | NCBI |
| CY207803 | A/NewYork/WC-LVD-15-041/2015(H3N2) | 18/3/2015 | NCBI |
| CY207811 | A/NewYork/WC-LVD-15-042/2015(H3N2) | 14/3/2015 | NCBI |
| CY207819 | A/NewYork/WC-LVD-15-043/2015(H3N2) | 21/2/2015 | NCBI |
| CY207827 | A/NewYork/WC-LVD-15-044/2015(H3N2) | 21/3/2015 | NCBI |
| CY207835 | A/NewYork/WC-LVD-15-045/2015(H3N2) | 3/8/2015 | NCBI |
| CY207843 | A/NewYork/WC-LVD-15-046/2015(H3N2) | 28/3/2015 | NCBI |
| CY207851 | A/NewYork/WC-LVD-15-047/2015(H3N2) | 15/4/2015 | NCBI |
| CY207859 | A/NewYork/WC-LVD-14-095/2014(H3N2) | 18/12/2014 | NCBI |
| CY207867 | A/NewYork/WC-LVD-14-096/2014(H3N2) | 12/12/2014 | NCBI |
| CY207875 | A/NewYork/WC-LVD-14-097/2014(H3N2) | 10/11/2014 | NCBI |
| CY207883 | A/NewYork/WC-LVD-14-098/2014(H3N2) | 8/12/2014 | NCBI |
| CY207891 | A/NewYork/WC-LVD-15-048/2015(H3N2) | 30/1/2015 | NCBI |
| CY207899 | A/NewYork/WC-LVD-15-049/2015(H3N2) | 22/1/2015 | NCBI |
| CY207907 | A/NewYork/WC-LVD-14-099/2014(H3N2) | 23/12/2014 | NCBI |
| CY207915 | A/NewYork/WC-LVD-14-100/2014(H3N2) | 29/11/2014 | NCBI |
| CY207923 | A/NewYork/WC-LVD-15-051/2015(H3N2) | 12/1/2015 | NCBI |
| CY207931 | A/NewYork/WC-LVD-14-102/0214(H3N2) | 18/12/2014 | NCBI |
| CY207939 | A/NewYork/WC-LVD-15-052/2015(H3N2) | 2/2/2015 | NCBI |
| CY207947 | A/NewYork/WC-LVD-15-053/2015(H3N2) | 23/1/2015 | NCBI |
| KX916100 | A/NorthCarolina/02/2016(H3N2) | 12/1/2016 | NCBI |
| KX916108 | A/Kansas/03/2016(H3N2) | 14/1/2016 | NCBI |
| KX916116 | A/Arizona/32/2016(H3N2) | 8/1/2016 | NCBI |
| KX916124 | A/Virginia/27/2016(H3N2) | 26/3/2016 | NCBI |
| KX916132 | A/Virginia/32/2016(H3N2) | 4/4/2016 | NCBI |
| KX916212 | A/Idaho/33/2016(H3N2) | 7/6/2016 | NCBI |
| KX916244 | A/Virginia/37/2016(H3N2) | 20/4/2016 | NCBI |
| KX917004 | A/California/43/2016(H3N2) | 11/1/2016 | NCBI |
| KX917012 | A/Indiana/25/2016(H3N2) | 16/2/2016 | NCBI |
| KX917108 | A/NorthCarolina/14/2016(H3N2) | 10/3/2016 | NCBI |
| KX917116 | A/NorthDakota/19/2016(H3N2) | 5/3/2016 | NCBI |
| KX917124 | A/NorthDakota/20/2016(H3N2) | 6/3/2016 | NCBI |
| KX917156 | A/Utah/23/2016(H3N2) | 8/3/2016 | NCBI |
| KX917172 | A/Idaho/16/2016(H3N2) | 7/3/2016 | NCBI |
| KX917180 | A/Nevada/20/2016(H3N2) | 10/3/2016 | NCBI |

| | | | |
|---|---|---|---|
| KX917188 | A/Virginia/39/2016(H3N2) | 10/3/2016 | NCBI |
| KX917204 | A/Pennsylvania/41/2016(H3N2) | 10/3/2016 | NCBI |
| KX917212 | A/Colorado/22/2016(H3N2) | 6/3/2016 | NCBI |
| KX917236 | A/NewHampshire/11/2016(H3N2) | 18/3/2016 | NCBI |
| KX917244 | A/NorthCarolina/35/2016(H3N2) | 21/3/2016 | NCBI |
| KX917252 | A/Delaware/15/2016(H3N2) | 20/3/2016 | NCBI |
| KX917268 | A/NewMexico/30/2016(H3N2) | 20/3/2016 | NCBI |
| KX917292 | A/Texas/126/2016(H3N2) | 21/3/2016 | NCBI |
| KX917300 | A/Utah/28/2016(H3N2) | 24/3/2016 | NCBI |
| KX917308 | A/Indiana/35/2016(H3N2) | 21/3/2016 | NCBI |
| KX917316 | A/Florida/44/2016(H3N2) | 30/3/2016 | NCBI |
| KX917332 | A/Wyoming/15/2016(H3N2) | 23/3/2016 | NCBI |
| KX917364 | A/Maryland/15/2016(H3N2) | 11/3/2016 | NCBI |
| KX917372 | A/NewYork/56/2016(H3N2) | 2/3/2016 | NCBI |
| KX917380 | A/NewYork/58/2016(H3N2) | 20/3/2016 | NCBI |
| KX917444 | A/Indiana/40/2016(H3N2) | 4/4/2016 | NCBI |
| KX917540 | A/NorthCarolina/44/2016(H3N2) | 12/4/2016 | NCBI |
| KX917636 | A/Texas/133/2016(H3N2) | 13/4/2016 | NCBI |
| KX917644 | A/Texas/134/2016(H3N2) | 20/4/2016 | NCBI |
| KX917676 | A/Texas/130/2016(H3N2) | 2/4/2016 | NCBI |
| KX917884 | A/Texas/136/2016(H3N2) | 22/4/2016 | NCBI |
| KX918236 | A/Wisconsin/67/2016(H3N2) | 3/5/2016 | NCBI |
| KX916084 | A/Ohio/31/2015(H3N2) | 2/12/2015 | NCBI |
| KX916092 | A/Connecticut/09/2015(H3N2) | 26/12/2015 | NCBI |
| KX916148 | A/Pennsylvania/73/2016(H3N2) | 4/5/2016 | NCBI |
| KX916156 | A/Pennsylvania/74/2016(H3N2) | 23/5/2016 | NCBI |
| KX916164 | A/Hawaii/48/2016(H3N2) | 13/4/2016 | NCBI |
| KX916180 | A/Hawaii/49/2016(H3N2) | 18/4/2016 | NCBI |
| KX916188 | A/Utah/42/2016(H3N2) | 26/5/2016 | NCBI |
| KX916196 | A/Utah/41/2016(H3N2) | 24/5/2016 | NCBI |
| KX916220 | A/Idaho/34/2016(H3N2) | 20/6/2016 | NCBI |
| KX916228 | A/Texas/175/2016(H3N2) | 22/6/2016 | NCBI |
| KX916236 | A/Texas/176/2016(H3N2) | 22/6/2016 | NCBI |
| KX916252 | A/Georgia/23/2016(H3N2) | 18/2/2016 | NCBI |
| KX916260 | A/Georgia/27/2016(H3N2) | 24/2/2016 | NCBI |
| KX916268 | A/Georgia/46/2016(H3N2) | 20/3/2016 | NCBI |
| KX916276 | A/Georgia/47/2016(H3N2) | 29/3/2016 | NCBI |

| | | | |
|---|---|---|---|
| KX916284 | A/Georgia/48/2016(H3N2) | 4/4/2016 | NCBI |
| KX916292 | A/Georgia/49/2016(H3N2) | 4/4/2016 | NCBI |
| KX916300 | A/Georgia/52/2016(H3N2) | 17/4/2016 | NCBI |
| KX916308 | A/Hawaii/56/2016(H3N2) | 11/6/2016 | NCBI |
| KX916316 | A/Hawaii/58/2016(H3N2) | 24/6/2016 | NCBI |
| KX916324 | A/California/120/2016(H3N2) | 27/6/2016 | NCBI |
| KX916332 | A/NewYork/86/2016(H3N2) | 28/7/2016 | NCBI |
| KX916340 | A/Ohio/34/2016(H3N2) | 17/8/2016 | NCBI |
| KX916348 | A/Hawaii/63/2016(H3N2) | 31/7/2016 | NCBI |
| KX916356 | A/Hawaii/64/2016(H3N2) | 3/8/2016 | NCBI |
| KX916364 | A/Iowa/23/2016(H3N2) | 19/8/2016 | NCBI |
| KX916372 | A/Montana/44/2016(H3N2) | 19/8/2016 | NCBI |
| KX916380 | A/Montana/45/2016(H3N2) | 22/8/2016 | NCBI |
| KX916388 | A/Wisconsin/77/2016(H3N2) | 12/8/2016 | NCBI |
| KX916396 | A/Texas/180/2016(H3N2) | 28/8/2016 | NCBI |
| KX916404 | A/Georgia/57/2016(H3N2) | 24/5/2016 | NCBI |
| KX916412 | A/Texas/171/2016(H3N2) | 16/5/2016 | NCBI |
| KX916420 | A/Texas/172/2016(H3N2) | 17/5/2016 | NCBI |
| KX916428 | A/Texas/174/2016(H3N2) | 23/5/2016 | NCBI |
| KX916436 | A/Texas/170/2016(H3N2) | 8/5/2016 | NCBI |
| KX916444 | A/Pennsylvania/82/2016(H3N2) | 14/4/2016 | NCBI |
| KX916460 | A/California/118/2016(H3N2) | 30/5/2016 | NCBI |
| KX916468 | A/Minnesota/71/2016(H3N2) | 23/6/2016 | NCBI |
| KX916476 | A/Nebraska/20/2016(H3N2) | 27/6/2016 | NCBI |
| KX916484 | A/Hawaii/54/2016(H3N2) | 21/5/2016 | NCBI |
| KX916492 | A/Hawaii/55/2016(H3N2) | 7/6/2016 | NCBI |
| KX916500 | A/Texas/177/2016(H3N2) | 23/6/2016 | NCBI |
| KX916508 | A/Georgia/58/2016(H3N2) | 5/7/2016 | NCBI |
| KX916524 | A/SouthCarolina/15/2016(H3N2) | 11/5/2016 | NCBI |
| KX916532 | A/SouthCarolina/16/2016(H3N2) | 11/5/2016 | NCBI |
| KX916540 | A/Georgia/53/2016(H3N2) | 17/4/2016 | NCBI |
| KX916548 | A/Georgia/56/2016(H3N2) | 30/4/2016 | NCBI |
| KX916556 | A/Georgia/54/2016(H3N2) | 18/4/2016 | NCBI |
| KX916564 | A/Vermont/25/2016(H3N2) | 18/4/2016 | NCBI |
| KX916572 | A/Vermont/23/2016(H3N2) | 14/4/2016 | NCBI |
| KX916580 | A/Vermont/22/2016(H3N2) | 13/4/2016 | NCBI |
| KX916588 | A/Vermont/24/2016(H3N2) | 17/4/2016 | NCBI |

| | | | |
|---|---|---|---|
| KX916596 | A/Vermont/26/2016(H3N2) | 21/4/2016 | NCBI |
| KX916604 | A/Vermont/27/2016(H3N2) | 5/5/2016 | NCBI |
| KX916612 | A/Vermont/28/2016(H3N2) | 5/5/2016 | NCBI |
| KX916620 | A/SouthCarolina/21/2016(H3N2) | 25/5/2016 | NCBI |
| KX916636 | A/SouthCarolina/17/2016(H3N2) | 23/5/2016 | NCBI |
| KX916644 | A/SouthCarolina/19/2016(H3N2) | 24/5/2016 | NCBI |
| KX916652 | A/SouthCarolina/22/2016(H3N2) | 25/5/2016 | NCBI |
| KX916660 | A/SouthCarolina/23/2016(H3N2) | 25/5/2016 | NCBI |
| KX916668 | A/SouthCarolina/20/2016(H3N2) | 24/5/2016 | NCBI |
| KX916676 | A/SouthCarolina/24/2016(H3N2) | 25/5/2016 | NCBI |
| KX916684 | A/SouthCarolina/25/2016(H3N2) | 25/5/2016 | NCBI |
| KX916692 | A/Texas/158/2016(H3N2) | 23/3/2016 | NCBI |
| KX916700 | A/Texas/166/2016(H3N2) | 11/4/2016 | NCBI |
| KX916708 | A/Arizona/66/2016(H3N2) | 12/5/2016 | NCBI |
| KX916716 | A/Arizona/65/2016(H3N2) | 9/5/2016 | NCBI |
| KX916724 | A/Idaho/31/2016(H3N2) | 13/5/2016 | NCBI |
| KX916732 | A/Idaho/32/2016(H3N2) | 15/5/2016 | NCBI |
| KX916740 | A/Hawaii/42/2016(H3N2) | 20/3/2016 | NCBI |
| KX916748 | A/Hawaii/46/2016(H3N2) | 31/3/2016 | NCBI |
| KX916756 | A/Washington/75/2016(H3N2) | 13/5/2016 | NCBI |
| KX916772 | A/Washington/74/2016(H3N2) | 22/4/2016 | NCBI |
| KX916780 | A/Alaska/23/2016(H3N2) | 15/6/2016 | NCBI |
| KX916788 | A/Alaska/22/2016(H3N2) | 6/6/2016 | NCBI |
| KX916796 | A/Florida/68/2016(H3N2) | 13/6/2016 | NCBI |
| KX916804 | A/Missouri/16/2016(H3N2) | 9/7/2016 | NCBI |
| KX916812 | A/Wisconsin/72/2016(H3N2) | 4/7/2016 | NCBI |
| KX916820 | A/Florida/70/2016(H3N2) | 15/7/2016 | NCBI |
| KX916828 | A/Arizona/67/2016(H3N2) | 4/5/2016 | NCBI |
| KX916836 | A/Hawaii/59/2016(H3N2) | 11/7/2016 | NCBI |
| KX916844 | A/Hawaii/60/2016(H3N2) | 11/7/2016 | NCBI |
| KX916852 | A/Hawaii/62/2016(H3N2) | 23/7/2016 | NCBI |
| KX916860 | A/Hawaii/61/2016(H3N2) | 19/7/2016 | NCBI |
| KX916868 | A/Ohio/30/2016(H3N2) | 10/8/2016 | NCBI |
| KX916876 | A/Texas/179/2016(H3N2) | 15/8/2016 | NCBI |
| KX916884 | A/Texas/178/2016(H3N2) | 4/8/2016 | NCBI |
| KX916892 | A/Nebraska/21/2016(H3N2) | 11/8/2016 | NCBI |
| KX916900 | A/Kentucky/29/2016(H3N2) | 25/8/2016 | NCBI |

| | | | |
|---|---|---|---|
| KX916908 | A/Michigan/97/2016(H3N2) | 17/8/2016 | NCBI |
| KX916916 | A/Michigan/98/2016(H3N2) | 21/8/2016 | NCBI |
| KX916924 | A/Wisconsin/74/2016(H3N2) | 8/8/2016 | NCBI |
| KX916932 | A/NewYork/83/2016(H3N2) | 13/7/2016 | NCBI |
| KX916940 | A/NewYork/81/2016(H3N2) | 12/7/2016 | NCBI |
| KX916948 | A/NewYork/82/2016(H3N2) | 12/7/2016 | NCBI |
| KX916956 | A/NewYork/84/2016(H3N2) | 13/7/2016 | NCBI |
| KX916964 | A/NewYork/85/2016(H3N2) | 13/7/2016 | NCBI |
| KX916972 | A/Wisconsin/73/2016(H3N2) | 14/7/2016 | NCBI |
| KX916980 | A/Utah/43/2016(H3N2) | 10/7/2016 | NCBI |
| KX916988 | A/Vermont/01/2016(H3N2) | 13/1/2016 | NCBI |
| KX916996 | A/California/12/2016(H3N2) | 12/1/2016 | NCBI |
| KX917020 | A/Delaware/10/2016(H3N2) | 1/3/2016 | NCBI |
| KX917028 | A/Delaware/11/2016(H3N2) | 2/3/2016 | NCBI |
| KX917036 | A/California/61/2016(H3N2) | 9/1/2016 | NCBI |
| KX917044 | A/Connecticut/05/2016(H3N2) | 14/2/2016 | NCBI |
| KX917052 | A/Connecticut/06/2016(H3N2) | 7/2/2016 | NCBI |
| KX917060 | A/Connecticut/12/2016(H3N2) | 16/2/2016 | NCBI |
| KX949247 | A/Kentucky/06/2016(H3N2) | 19/2/2016 | NCBI |
| KX917076 | A/Virginia/24/2016(H3N2) | 1/3/2016 | NCBI |
| KX917084 | A/NewYork/25/2016(H3N2) | 8/2/2016 | NCBI |
| KX917092 | A/NewYork/26/2016(H3N2) | 5/2/2016 | NCBI |
| KX917100 | A/NewYork/27/2016(H3N2) | 11/2/2016 | NCBI |
| KX917132 | A/Kentucky/10/2016(H3N2) | 17/2/2016 | NCBI |
| KX917140 | A/SouthCarolina/08/2016(H3N2) | 22/2/2016 | NCBI |
| KX917148 | A/Texas/59/2016(H3N2) | 26/2/2016 | NCBI |
| KX917196 | A/Pennsylvania/40/2016(H3N2) | 10/3/2016 | NCBI |
| KX917220 | A/Washington/41/2016(H3N2) | 4/3/2016 | NCBI |
| KX917228 | A/Massachusetts/26/2016(H3N2) | 14/3/2016 | NCBI |
| KX917260 | A/Texas/121/2016(H3N2) | 20/3/2016 | NCBI |
| KX917276 | A/SouthCarolina/12/2016(H3N2) | 5/3/2016 | NCBI |
| KX917284 | A/Texas/123/2016(H3N2) | 21/3/2016 | NCBI |
| KX917340 | A/Washington/51/2016(H3N2) | 30/3/2016 | NCBI |
| KX917348 | A/NorthCarolina/40/2016(H3N2) | 30/3/2016 | NCBI |
| KX917356 | A/NorthCarolina/41/2016(H3N2) | 5/4/2016 | NCBI |
| KX917388 | A/NewYork/60/2016(H3N2) | 31/3/2016 | NCBI |
| KX917396 | A/Michigan/69/2016(H3N2) | 31/3/2016 | NCBI |

| | | | |
|---|---|---|---|
| KX917412 | A/NewYork/62/2016(H3N2) | 11/4/2016 | NCBI |
| KX917420 | A/Texas/129/2016(H3N2) | 11/4/2016 | NCBI |
| KX917436 | A/Mississippi/16/2016(H3N2) | 18/4/2016 | NCBI |
| KX917452 | A/Florida/57/2016(H3N2) | 6/4/2016 | NCBI |
| KX917460 | A/Vermont/17/2016(H3N2) | 16/4/2016 | NCBI |
| KX917468 | A/Washington/56/2016(H3N2) | 21/3/2016 | NCBI |
| KX917476 | A/Washington/57/2016(H3N2) | 21/3/2016 | NCBI |
| KX917484 | A/Minnesota/65/2016(H3N2) | 15/4/2016 | NCBI |
| KX917492 | A/Colorado/28/2016(H3N2) | 30/3/2016 | NCBI |
| KX917500 | A/Colorado/32/2016(H3N2) | 9/4/2016 | NCBI |
| KX917508 | A/SouthDakota/21/2016(H3N2) | 20/3/2016 | NCBI |
| KX917516 | A/SouthDakota/23/2016(H3N2) | 29/3/2016 | NCBI |
| KX917524 | A/SouthDakota/24/2016(H3N2) | 31/3/2016 | NCBI |
| KX917532 | A/Louisiana/18/2016(H3N2) | 21/3/2016 | NCBI |
| KX917548 | A/Arizona/56/2016(H3N2) | 13/3/2016 | NCBI |
| KX917556 | A/Arizona/57/2016(H3N2) | 13/3/2016 | NCBI |
| KX917564 | A/Nevada/34/2016(H3N2) | 20/4/2016 | NCBI |
| KX917572 | A/Nevada/34/2016(H3N2) | 20/4/2016 | NCBI |
| KX917580 | A/Washington/60/2016(H3N2) | 2/4/2016 | NCBI |
| KX917588 | A/Oklahoma/09/2016(H3N2) | 29/3/2016 | NCBI |
| KX917596 | A/Idaho/21/2016(H3N2) | 8/4/2016 | NCBI |
| KX917604 | A/Idaho/24/2016(H3N2) | 16/4/2016 | NCBI |
| KX917612 | A/Washington/63/2016(H3N2) | 6/4/2016 | NCBI |
| KX917620 | A/NewJersey/20/2016(H3N2) | 18/4/2016 | NCBI |
| KX917628 | A/NewJersey/22/2016(H3N2) | 20/4/2016 | NCBI |
| KX917652 | A/NewMexico/36/2016(H3N2) | 4/4/2016 | NCBI |
| KX917660 | A/Hawaii/38/2016(H3N2) | 11/3/2016 | NCBI |
| KX917668 | A/Hawaii/39/2016(H3N2) | 24/3/2016 | NCBI |
| KX917684 | A/Montana/43/2016(H3N2) | 8/4/2016 | NCBI |
| KX917692 | A/Pennsylvania/64/2016(H3N2) | 2/4/2016 | NCBI |
| KX917700 | A/Pennsylvania/62/2016(H3N2) | 31/3/2016 | NCBI |
| KX917708 | A/Florida/58/2016(H3N2) | 10/4/2016 | NCBI |
| KX917716 | A/Florida/59/2016(H3N2) | 10/4/2016 | NCBI |
| KX917724 | A/Wyoming/19/2016(H3N2) | 4/4/2016 | NCBI |
| KX917732 | A/Wyoming/22/2016(H3N2) | 17/4/2016 | NCBI |
| KX917748 | A/Ohio/20/2016(H3N2) | 25/4/2016 | NCBI |
| KX917740 | A/Ohio/20/2016(H3N2) | 25/4/2016 | NCBI |

| | | | |
|---|---|---|---|
| KX917756 | A/Virginia/55/2016(H3N2) | 20/4/2016 | NCBI |
| KX917764 | A/Virginia/52/2016(H3N2) | 13/4/2016 | NCBI |
| KX917772 | A/RhodeIsland/21/2016(H3N2) | 17/4/2016 | NCBI |
| KX917780 | A/RhodeIsland/20/2016(H3N2) | 13/4/2016 | NCBI |
| KX917788 | A/Utah/34/2016(H3N2) | 22/4/2016 | NCBI |
| KX917796 | A/Utah/36/2016(H3N2) | 23/4/2016 | NCBI |
| KX917804 | A/Colorado/31/2016(H3N2) | 7/4/2016 | NCBI |
| KX917812 | A/Colorado/33/2016(H3N2) | 9/4/2016 | NCBI |
| KX917820 | A/Colorado/36/2016(H3N2) | 22/4/2016 | NCBI |
| KX917828 | A/Colorado/37/2016(H3N2) | 22/4/2016 | NCBI |
| KX917836 | A/Colorado/38/2016(H3N2) | 1/5/2016 | NCBI |
| KX917844 | A/Minnesota/63/2016(H3N2) | 7/4/2016 | NCBI |
| KX917852 | A/Minnesota/69/2016(H3N2) | 11/4/2016 | NCBI |
| KX917860 | A/Alaska/19/2016(H3N2) | 25/4/2016 | NCBI |
| KX917868 | A/Arizona/60/2016(H3N2) | 20/4/2016 | NCBI |
| KX917876 | A/Arizona/59/2016(H3N2) | 18/4/2016 | NCBI |
| KX917892 | A/Kentucky/22/2016(H3N2) | 14/4/2016 | NCBI |
| KX917900 | A/Kentucky/25/2016(H3N2) | 28/4/2016 | NCBI |
| KX917908 | A/NewJersey/24/2016(H3N2) | 4/5/2016 | NCBI |
| KX917916 | A/NewJersey/23/2016(H3N2) | 2/5/2016 | NCBI |
| KX917924 | A/Idaho/25/2016(H3N2) | 18/4/2016 | NCBI |
| KX917932 | A/Idaho/27/2016(H3N2) | 27/4/2016 | NCBI |
| KX917940 | A/Wyoming/26/2016(H3N2) | 25/4/2016 | NCBI |
| KX917948 | A/Washington/72/2016(H3N2) | 2/5/2016 | NCBI |
| KX917956 | A/Washington/65/2016(H3N2) | 7/4/2016 | NCBI |
| KX917964 | A/Washington/66/2016(H3N2) | 8/4/2016 | NCBI |
| KX917972 | A/Washington/71/2016(H3N2) | 24/4/2016 | NCBI |
| KX917980 | A/Nevada/35/2016(H3N2) | 27/4/2016 | NCBI |
| KX917988 | A/Nevada/35/2016(H3N2) | 27/4/2016 | NCBI |
| KX917996 | A/Vermont/21/2016(H3N2) | 28/4/2016 | NCBI |
| KX918004 | A/Vermont/20/2016(H3N2) | 27/4/2016 | NCBI |
| KX918012 | A/Nevada/36/2016(H3N2) | 29/4/2016 | NCBI |
| KX918020 | A/NewMexico/44/2016(H3N2) | 12/4/2016 | NCBI |
| KX918028 | A/California/111/2016(H3N2) | 20/4/2016 | NCBI |
| KX918036 | A/California/110/2016(H3N2) | 17/4/2016 | NCBI |
| KX918044 | A/NewYork/80/2016(H3N2) | 29/4/2016 | NCBI |
| KX918052 | A/NewYork/77/2016(H3N2) | 26/4/2016 | NCBI |

| | | | |
|---|---|---|---|
| KX918060 | A/Ohio/23/2016(H3N2) | 4/5/2016 | NCBI |
| KX918068 | A/Ohio/24/2016(H3N2) | 8/5/2016 | NCBI |
| KX918076 | A/Ohio/24/2016(H3N2) | 8/5/2016 | NCBI |
| KX918084 | A/Ohio/25/2016(H3N2) | 11/5/2016 | NCBI |
| KX918092 | A/California/107/2016(H3N2) | 10/4/2016 | NCBI |
| KX918108 | A/California/105/2016(H3N2) | 7/4/2016 | NCBI |
| KX918100 | A/California/105/2016(H3N2) | 7/4/2016 | NCBI |
| KX918116 | A/California/101/2016(H3N2) | 25/3/2016 | NCBI |
| KX918124 | A/Utah/39/2016(H3N2) | 9/5/2016 | NCBI |
| KX918132 | A/Oregon/15/2016(H3N2) | 23/4/2016 | NCBI |
| KX918140 | A/Oregon/16/2016(H3N2) | 29/4/2016 | NCBI |
| KX918148 | A/Connecticut/30/2016(H3N2) | 1/4/2016 | NCBI |
| KX918156 | A/Connecticut/30/2016(H3N2) | 1/4/2016 | NCBI |
| KX918164 | A/Connecticut/27/2016(H3N2) | 28/3/2016 | NCBI |
| KX918172 | A/Connecticut/27/2016(H3N2) | 28/3/2016 | NCBI |
| KX918180 | A/Connecticut/26/2016(H3N2) | 27/3/2016 | NCBI |
| KX918188 | A/Connecticut/20/2016(H3N2) | 16/3/2016 | NCBI |
| KX918196 | A/Connecticut/25/2016(H3N2) | 26/3/2016 | NCBI |
| KX918204 | A/Connecticut/19/2016(H3N2) | 11/3/2016 | NCBI |
| KX918212 | A/Connecticut/18/2016(H3N2) | 8/3/2016 | NCBI |
| KX918220 | A/Florida/64/2016(H3N2) | 5/5/2016 | NCBI |
| KX918228 | A/Wisconsin/65/2016(H3N2) | 25/4/2016 | NCBI |
| KX918244 | A/Pennsylvania/72/2016(H3N2) | 3/5/2016 | NCBI |
| KX918252 | A/Pennsylvania/70/2016(H3N2) | 2/5/2016 | NCBI |
| KX918260 | A/Louisiana/21/2016(H3N2) | 14/4/2016 | NCBI |
| KX918268 | A/Louisiana/23/2016(H3N2) | 22/4/2016 | NCBI |
| KX918276 | A/Louisiana/23/2016(H3N2) | 22/4/2016 | NCBI |
| KX918284 | A/Kentucky/26/2016(H3N2) | 25/4/2016 | NCBI |
| KX918292 | A/Hawaii/47/2016(H3N2) | 1/4/2016 | NCBI |
| KX918300 | A/Iowa/18/2016(H3N2) | 11/5/2016 | NCBI |
| KX918308 | A/Iowa/20/2016(H3N2) | 13/5/2016 | NCBI |
| KX918316 | A/NewYork/78/2016(H3N2) | 26/4/2016 | NCBI |
| KX918324 | A/NewYork/78/2016(H3N2) | 26/4/2016 | NCBI |
| KX918340 | A/NewYork/68/2016(H3N2) | 6/4/2016 | NCBI |
| KX918332 | A/NewYork/68/2016(H3N2) | 6/4/2016 | NCBI |
| KX918348 | A/Florida/67/2016(H3N2) | 11/5/2016 | NCBI |
| KX918356 | A/Vermont/29/2016(H3N2) | 13/5/2016 | NCBI |

| | | | |
|---|---|---|---|
| KX918364 | A/Vermont/30/2016(H3N2) | 18/5/2016 | NCBI |
| KX918372 | A/Mississippi/14/2016(H3N2) | 18/5/2016 | NCBI |
| KX918380 | A/Mississippi/15/2016(H3N2) | 18/5/2016 | NCBI |
| KX918388 | A/Wisconsin/69/2016(H3N2) | 11/5/2016 | NCBI |
| KX918396 | A/Wisconsin/69/2016(H3N2) | 11/5/2016 | NCBI |
| KX918404 | A/Texas/173/2016(H3N2) | 21/5/2016 | NCBI |
| KX918412 | A/Texas/173/2016(H3N2) | 21/5/2016 | NCBI |
| KX949259 | A/Florida/93/2015(H3N2) | 4/12/2015 | NCBI |
| KX949170 | A/Florida/52/2016(H3N2) | 24/1/2016 | NCBI |
| KX949183 | A/NewYork/18/2016(H3N2) | 5/2/2016 | NCBI |
| KX949152 | A/NewYork/21/2016(H3N2) | 13/2/2016 | NCBI |
| KX949158 | A/NewYork/19/2016(H3N2) | 9/2/2016 | NCBI |
| KX949243 | A/Oklahoma/02/2016(H3N2) | 14/1/2016 | NCBI |
| KX949302 | A/Florida/22/2016(H3N2) | 19/2/2016 | NCBI |
| KX949330 | A/Florida/17/2016(H3N2) | 10/2/2016 | NCBI |
| KX949338 | A/Florida/28/2016(H3N2) | 2/3/2016 | NCBI |
| KX949346 | A/NorthDakota/26/2016(H3N2) | 22/3/2016 | NCBI |
| KY003432 | A/NorthCarolina/51/2016(H3N2) | 28/8/2016 | NCBI |
| KY003548 | A/Texas/183/2016(H3N2) | 31/8/2016 | NCBI |
| KY003503 | A/Texas/189/2016(H3N2) | 6/9/2016 | NCBI |
| KY003522 | A/California/122/2016(H3N2) | 12/7/2016 | NCBI |
| KY003421 | A/Iowa/25/2016(H3N2) | 9/9/2016 | NCBI |
| KY003479 | A/Montana/46/2016(H3N2) | 9/9/2016 | NCBI |
| KY003487 | A/Hawaii/67/2016(H3N2) | 4/9/2016 | NCBI |
| KY003384 | A/RhodeIsland/29/2016(H3N2) | 13/9/2016 | NCBI |
| KY003363 | A/RhodeIsland/28/2016(H3N2) | 13/9/2016 | NCBI |
| KY003478 | A/RhodeIsland/26/2016(H3N2) | 13/9/2016 | NCBI |
| KY003405 | A/RhodeIsland/27/2016(H3N2) | 13/9/2016 | NCBI |
| KY003485 | A/Maine/21/2016(H3N2) | 12/9/2016 | NCBI |
| KY003371 | A/RhodeIsland/25/2016(H3N2) | 13/9/2016 | NCBI |
| KY003480 | A/NewHampshire/20/2016(H3N2) | 7/9/2016 | NCBI |
| KY003415 | A/Alaska/25/2016(H3N2) | 10/9/2016 | NCBI |
| KY003462 | A/NorthDakota/27/2016(H3N2) | 15/9/2016 | NCBI |
| KY003458 | A/Alaska/24/2016(H3N2) | 8/9/2016 | NCBI |
| KY003578 | A/Texas/182/2016(H3N2) | 31/8/2016 | NCBI |
| KY003379 | A/NorthCarolina/52/2016(H3N2) | 8/9/2016 | NCBI |
| KY003406 | A/Iowa/24/2016(H3N2) | 2/9/2016 | NCBI |

| | | | |
|---|---|---|---|
| KY003496 | A/Hawaii/68/2016(H3N2) | 7/9/2016 | NCBI |
| KY003413 | A/Pennsylvania/86/2016(H3N2) | 14/9/2016 | NCBI |
| KY003504 | A/Texas/185/2016(H3N2) | 1/9/2016 | NCBI |
| KY003566 | A/Texas/187/2016(H3N2) | 6/9/2016 | NCBI |
| KY003575 | A/Texas/184/2016(H3N2) | 31/8/2016 | NCBI |
| KY003526 | A/Missouri/18/2016(H3N2) | 11/7/2016 | NCBI |
| KY003555 | A/Michigan/81/2016(H3N2) | 12/7/2016 | NCBI |
| KY003550 | A/Ohio/31/2016(H3N2) | 14/8/2016 | NCBI |
| KY003528 | A/Nebraska/22/2016(H3N2) | 6/9/2016 | NCBI |
| KY003564 | A/California/121/2016(H3N2) | 9/7/2016 | NCBI |
| KY045405 | A/NewYork/89/2016(H3N2) | 8/8/2016 | NCBI |
| KY045466 | A/NewYork/95/2016(H3N2) | 29/8/2016 | NCBI |
| KY045445 | A/NewYork/91/2016(H3N2) | 23/8/2016 | NCBI |
| KY045408 | A/NewYork/93/2016(H3N2) | 24/8/2016 | NCBI |
| KY045457 | A/NewYork/96/2016(H3N2) | 1/9/2016 | NCBI |
| KY045455 | A/NewYork/92/2016(H3N2) | 23/8/2016 | NCBI |
| KY045433 | A/NewYork/90/2016(H3N2) | 9/8/2016 | NCBI |
| KY045454 | A/NewYork/94/2016(H3N2) | 27/8/2016 | NCBI |
| KY045443 | A/NewYork/88/2016(H3N2) | 27/7/2016 | NCBI |
| KX879573 | A/Alchi/2/1968(H3N2) | 1968 | NCBI |
| KY049988 | A/Qinghai/178/2010(H3N2) | 15/3/2010 | NCBI |
| KY115814 | A/Texas/181/2016(H3N2) | 28/8/2016 | NCBI |
| KY115822 | A/Ohio/37/2016(H3N2) | 7/9/2016 | NCBI |
| KY115838 | A/NorthCarolina/54/2016(H3N2) | 20/9/2016 | NCBI |
| KY115846 | A/NorthCarolina/53/2016(H3N2) | 8/9/2016 | NCBI |
| KY115862 | A/Illinois/45/2016(H3N2) | 20/8/2016 | NCBI |
| KY115870 | A/Illinois/46/2016(H3N2) | 14/9/2016 | NCBI |
| KY115878 | A/Michigan/100/2016(H3N2) | 31/8/2016 | NCBI |
| KY115886 | A/Michigan/101/2016(H3N2) | 6/9/2016 | NCBI |
| KY115894 | A/Michigan/103/2016(H3N2) | 11/9/2016 | NCBI |
| KY115902 | A/Michigan/104/2016(H3N2) | 17/9/2016 | NCBI |
| KY115918 | A/Nebraska/24/2016(H3N2) | 19/9/2016 | NCBI |
| KY115926 | A/Nebraska/23/2016(H3N2) | 6/9/2016 | NCBI |
| KY115942 | A/Wisconsin/83/2016(H3N2) | 24/9/2016 | NCBI |
| KY115950 | A/Missouri/15/2016(H3N2) | 2016 | NCBI |
| KY115958 | A/Michigan/78/2016(H3N2) | 21/5/2016 | NCBI |
| KY115966 | A/Nevada/37/2016(H3N2) | 29/7/2016 | NCBI |

| | | | |
|---|---|---|---|
| KY115974 | A/Michigan/99/2016(H3N2) | 24/8/2016 | NCBI |
| KY115982 | A/Florida/74/2016(H3N2) | 7/9/2016 | NCBI |
| KY115990 | A/Florida/75/2016(H3N2) | 7/9/2016 | NCBI |
| KY115998 | A/Wisconsin/78/2016(H3N2) | 7/9/2016 | NCBI |
| KY116006 | A/NewYork/88/2016(H3N2) | 27/7/2016 | NCBI |
| KY116014 | A/NewYork/96/2016(H3N2) | 1/9/2016 | NCBI |
| KY116022 | A/NewYork/89/2016(H3N2) | 8/8/2016 | NCBI |
| KY116030 | A/NewYork/90/2016(H3N2) | 9/8/2016 | NCBI |
| KY116038 | A/NewYork/91/2016(H3N2) | 23/8/2016 | NCBI |
| KY116046 | A/NewYork/92/2016(H3N2) | 23/8/2016 | NCBI |
| KY116054 | A/NewYork/93/2016(H3N2) | 24/8/2016 | NCBI |
| KY116062 | A/NewYork/94/2016(H3N2) | 27/8/2016 | NCBI |
| KY116070 | A/NewYork/95/2016(H3N2) | 29/8/2016 | NCBI |
| KY116078 | A/Indiana/17/2016(H3N2) | 25/1/2016 | NCBI |
| KY116086 | A/Alaska/14/2016(H3N2) | 2016 | NCBI |
| KY116094 | A/Florida/66/2016(H3N2) | 2016 | NCBI |
| KY116102 | A/Maine/22/2016(H3N2) | 17/9/2016 | NCBI |
| KY116118 | A/Alaska/26/2016(H3N2) | 20/9/2016 | NCBI |
| KY116126 | A/Iowa/26/2016(H3N2) | 16/9/2016 | NCBI |
| KY116134 | A/Iowa/27/2016(H3N2) | 26/9/2016 | NCBI |
| KY116142 | A/NewMexico/48/2016(H3N2) | 5/9/2016 | NCBI |
| KY116158 | A/NewYork/99/2016(H3N2) | 23/9/2016 | NCBI |
| KY116166 | A/NewYork/98/2016(H3N2) | 21/9/2016 | NCBI |
| KY116182 | A/Wisconsin/85/2016(H3N2) | 2/10/2016 | NCBI |
| KY116198 | A/Wisconsin/84/2016(H3N2) | 29/9/2016 | NCBI |
| KY116206 | A/Pennsylvania/87/2016(H3N2) | 21/9/2016 | NCBI |
| KY116222 | A/Pennsylvania/88/2016(H3N2) | 27/9/2016 | NCBI |
| KY116238 | A/Pennsylvania/89/2016(H3N2) | 27/9/2016 | NCBI |
| KY116246 | A/Florida/76/2016(H3N2) | 19/9/2016 | NCBI |
| KY116262 | A/Hawaii/73/2016(H3N2) | 16/9/2016 | NCBI |
| KY116270 | A/Hawaii/74/2016(H3N2) | 16/9/2016 | NCBI |
| KY116278 | A/Hawaii/71/2016(H3N2) | 13/9/2016 | NCBI |
| KY116286 | A/Hawaii/77/2016(H3N2) | 22/9/2016 | NCBI |
| KY116294 | A/Hawaii/78/2016(H3N2) | 22/9/2016 | NCBI |
| KY116302 | A/Hawaii/76/2016(H3N2) | 21/9/2016 | NCBI |
| KY116310 | A/Hawaii/75/2016(H3N2) | 19/9/2016 | NCBI |
| KY116318 | A/Hawaii/81/2016(H3N2) | 24/9/2016 | NCBI |

| | | | |
|---|---|---|---|
| KY116326 | A/Hawaii/79/2016(H3N2) | 22/9/2016 | NCBI |
| KY116334 | A/Hawaii/80/2016(H3N2) | 23/9/2016 | NCBI |
| KY116342 | A/Minnesota/72/2016(H3N2) | 2/10/2016 | NCBI |
| KY116358 | A/Minnesota/73/2016(H3N2) | 4/10/2016 | NCBI |
| KY116366 | A/Wyoming/29/2016(H3N2) | 3/10/2016 | NCBI |
| KY116374 | A/California/124/2016(H3N2) | 10/8/2016 | NCBI |
| KY116382 | A/California/125/2016(H3N2) | 10/8/2016 | NCBI |
| KY116390 | A/California/126/2016(H3N2) | 10/8/2016 | NCBI |
| KY116398 | A/California/128/2016(H3N2) | 27/9/2016 | NCBI |
| KY116406 | A/California/127/2016(H3N2) | 24/9/2016 | NCBI |
| KY116414 | A/California/129/2016(H3N2) | 5/10/2016 | NCBI |
| KY116422 | A/California/130/2016(H3N2) | 10/10/2016 | NCBI |
| KY116430 | A/Nevada/38/2016(H3N2) | 22/9/2016 | NCBI |
| KY116438 | A/Washington/77/2016(H3N2) | 3/10/2016 | NCBI |
| KY116446 | A/Washington/78/2016(H3N2) | 5/10/2016 | NCBI |
| KY116462 | A/Florida/79/2016(H3N2) | 3/10/2016 | NCBI |
| KY116470 | A/Florida/78/2016(H3N2) | 30/9/2016 | NCBI |
| KY116478 | A/Wyoming/30/2016(H3N2) | 7/10/2016 | NCBI |
| KY116486 | A/Arizona/70/2016(H3N2) | 27/9/2016 | NCBI |
| KY116494 | A/Arizona/71/2016(H3N2) | 30/9/2016 | NCBI |
| KY116502 | A/Idaho/37/2016(H3N2) | 7/10/2016 | NCBI |
| KY116510 | A/Illinois/47/2016(H3N2) | 7/10/2016 | NCBI |
| KY116518 | A/Michigan/105/2016(H3N2) | 22/9/2016 | NCBI |
| KY116526 | A/Michigan/106/2016(H3N2) | 28/9/2016 | NCBI |
| KY116534 | A/NewHampshire/21/2016(H3N2) | 4/10/2016 | NCBI |
| KY116542 | A/Iowa/29/2016(H3N2) | 6/10/2016 | NCBI |
| KY116550 | A/Iowa/28/2016(H3N2) | 1/10/2016 | NCBI |
| KY116558 | A/Massachusetts/36/2016(H3N2) | 4/10/2016 | NCBI |
| KY116566 | A/Massachusetts/37/2016(H3N2) | 13/10/2016 | NCBI |
| KY116574 | A/NewYork/100/2016(H3N2) | 26/9/2016 | NCBI |
| KY116582 | A/California/135/2016(H3N2) | 9/10/2016 | NCBI |
| KY116590 | A/California/132/2016(H3N2) | 2/10/2016 | NCBI |
| KY116598 | A/California/131/2016(H3N2) | 30/9/2016 | NCBI |
| KY116606 | A/California/133/2016(H3N2) | 3/10/2016 | NCBI |
| KY116614 | A/California/134/2016(H3N2) | 3/10/2016 | NCBI |
| KY116622 | A/Wisconsin/86/2016(H3N2) | 14/10/2016 | NCBI |
| KY116630 | A/Connecticut/34/2016(H3N2) | 4/10/2016 | NCBI |

| | | | |
|---|---|---|---|
| KY116638 | A/Florida/80/2016(H3N2) | 1/10/2016 | NCBI |
| KY116646 | A/Minnesota/74/2016(H3N2) | 5/10/2016 | NCBI |
| KY116654 | A/Minnesota/75/2016(H3N2) | 10/10/2016 | NCBI |
| KY116662 | A/Ohio/38/2016(H3N2) | 16/10/2016 | NCBI |
| KY116670 | A/Maine/23/2016(H3N2) | 23/10/2016 | NCBI |
| KY116918 | A/Michigan/15/2014(H3N2) | 24/9/2014 | NCBI |
| KY116959 | A/Alaska/232/2015(H3N2) | 9/9/2015 | NCBI |
| KY116942 | A/Montana/28/2015(H3N2) | 24/9/2015 | NCBI |
| KY044140 | A/Idaho/33/2016(H3N2) | 7/6/2016 | NCBI |
| KY116951 | A/Alaska/240/2015(H3N2) | 15/9/2015 | NCBI |
| KY116967 | A/RhodeIsland/09/2012(H3N2) | 28/11/2012 | NCBI |
| KY116975 | A/WestVirginia/20/2012(H3N2) | 20/11/2012 | NCBI |
| KY116991 | A/Virginia/51/2014(H3N2) | 29/12/2014 | NCBI |
| KY117007 | A/Alaska/140/2015(H3N2) | 11/8/2015 | NCBI |
| KY289029 | A/Hawaii/75/2016(H3N2) | 19/9/2016 | NCBI |
| KY289650 | A/NewYork/100/2016(H3N2) | 26/9/2016 | NCBI |
| KY289038 | A/California/140/2016(H3N2) | 17/10/2016 | NCBI |
| KY289027 | A/Michigan/105/2016(H3N2) | 22/9/2016 | NCBI |
| KY289039 | A/Idaho/37/2016(H3N2) | 7/10/2016 | NCBI |
| KY289314 | A/NewMexico/48/2016(H3N2) | 5/9/2016 | NCBI |
| KY289033 | A/Hawaii/81/2016(H3N2) | 24/9/2016 | NCBI |
| KY289101 | A/NorthCarolina/58/2016(H3N2) | 27/10/2016 | NCBI |
| KY289044 | A/Florida/87/2016(H3N2) | 27/10/2016 | NCBI |
| KY289122 | A/Florida/89/2016(H3N2) | 29/10/2016 | NCBI |
| KY289051 | A/Delaware/27/2016(H3N2) | 26/10/2016 | NCBI |
| KY289141 | A/NewHampshire/22/2016(H3N2) | 20/10/2016 | NCBI |
| KY289151 | A/WestVirginia/06/2016(H3N2) | 25/10/2016 | NCBI |
| KY289073 | A/Tennessee/28/2016(H3N2) | 28/9/2016 | NCBI |
| KY289132 | A/Vermont/32/2016(H3N2) | 8/10/2016 | NCBI |
| KY289146 | A/Connecticut/35/2016(H3N2) | 27/10/2016 | NCBI |
| KY289131 | A/WestVirginia/07/2016(H3N2) | 25/10/2016 | NCBI |
| KY289170 | A/Florida/84/2016(H3N2) | 15/10/2016 | NCBI |
| KY289066 | A/Pennsylvania/92/2016(H3N2) | 29/10/2016 | NCBI |
| KY289082 | A/NewHampshire/23/2016(H3N2) | 28/10/2016 | NCBI |
| KY289103 | A/Virginia/61/2016(H3N2) | 24/10/2016 | NCBI |
| KY289092 | A/Tennessee/29/2016(H3N2) | 25/10/2016 | NCBI |
| KY289115 | A/Florida/81/2016(H3N2) | 3/10/2016 | NCBI |

| | | | |
|---|---|---|---|
| KY289165 | A/NorthCarolina/57/2016(H3N2) | 17/10/2016 | NCBI |
| KY289196 | A/Florida/86/2016(H3N2) | 17/10/2016 | NCBI |
| KY289149 | A/NewHampshire/21/2016(H3N2) | 4/10/2016 | NCBI |
| KY289121 | A/Massachusetts/37/2016(H3N2) | 13/10/2016 | NCBI |
| KY289206 | A/Massachusetts/36/2016(H3N2) | 4/10/2016 | NCBI |
| KY289148 | A/Wisconsin/86/2016(H3N2) | 14/10/2016 | NCBI |
| KY289213 | A/Michigan/106/2016(H3N2) | 28/9/2016 | NCBI |
| KY289252 | A/Wisconsin/84/2016(H3N2) | 29/9/2016 | NCBI |
| KY289289 | A/Illinois/47/2016(H3N2) | 7/10/2016 | NCBI |
| KY289312 | A/Washington/78/2016(H3N2) | 5/10/2016 | NCBI |
| KY289237 | A/California/130/2016(H3N2) | 10/10/2016 | NCBI |
| KY289308 | A/Minnesota/72/2016(H3N2) | 2/10/2016 | NCBI |
| KY289263 | A/Iowa/26/2016(H3N2) | 16/9/2016 | NCBI |
| KY289297 | A/Connecticut/34/2016(H3N2) | 4/10/2016 | NCBI |
| KY289257 | A/Florida/80/2016(H3N2) | 1/10/2016 | NCBI |
| KY289276 | A/Florida/78/2016(H3N2) | 30/9/2016 | NCBI |
| KY289274 | A/Iowa/27/2016(H3N2) | 26/9/2016 | NCBI |
| KY289355 | A/California/151/2016(H3N2) | 2/11/2016 | NCBI |
| KY289374 | A/California/145/2016(H3N2) | 17/10/2016 | NCBI |
| KY289437 | A/Utah/45/2016(H3N2) | 26/10/2016 | NCBI |
| KY289427 | A/California/149/2016(H3N2) | 17/10/2016 | NCBI |
| KY289361 | A/Idaho/38/2016(H3N2) | 13/10/2016 | NCBI |
| KY289460 | A/Washington/81/2016(H3N2) | 12/10/2016 | NCBI |
| KY289402 | A/Oregon/19/2016(H3N2) | 19/10/2016 | NCBI |
| KY289326 | A/California/146/2016(H3N2) | 17/10/2016 | NCBI |
| KY289329 | A/California/147/2016(H3N2) | 17/10/2016 | NCBI |
| KY289439 | A/Washington/82/2016(H3N2) | 16/10/2016 | NCBI |
| KY289332 | A/Montana/53/2016(H3N2) | 21/10/2016 | NCBI |
| KY289497 | A/Texas/190/2016(H3N2) | 21/10/2016 | NCBI |
| KY289473 | A/Montana/52/2016(H3N2) | 20/10/2016 | NCBI |
| KY289347 | A/California/143/2016(H3N2) | 26/10/2016 | NCBI |
| KY289348 | A/Utah/44/2016(H3N2) | 21/10/2016 | NCBI |
| KY289350 | A/Alabama/13/2016(H3N2) | 19/10/2016 | NCBI |
| KY289475 | A/Minnesota/76/2016(H3N2) | 25/10/2016 | NCBI |
| KY289416 | A/California/148/2016(H3N2) | 17/10/2016 | NCBI |
| KY289390 | A/Oregon/18/2016(H3N2) | 18/10/2016 | NCBI |
| KY289450 | A/California/144/2016(H3N2) | 27/10/2016 | NCBI |

| | | | |
|---|---|---|---|
| KY289384 | A/California/150/2016(H3N2) | 2/11/2016 | NCBI |
| KY289394 | A/Alabama/14/2016(H3N2) | 19/10/2016 | NCBI |
| KY289457 | A/Minnesota/77/2016(H3N2) | 27/10/2016 | NCBI |
| KY289529 | A/Wisconsin/87/2016(H3N2) | 21/10/2016 | NCBI |
| KY289593 | A/California/141/2016(H3N2) | 20/10/2016 | NCBI |
| KY289590 | A/Hawaii/82/2016(H3N2) | 27/9/2016 | NCBI |
| KY289584 | A/Iowa/30/2016(H3N2) | 12/10/2016 | NCBI |
| KY289539 | A/Nevada/39/2016(H3N2) | 12/10/2016 | NCBI |
| KY289544 | A/Michigan/107/2016(H3N2) | 21/10/2016 | NCBI |
| KY289595 | A/Nevada/40/2016(H3N2) | 13/10/2016 | NCBI |
| KY289619 | A/Nevada/41/2016(H3N2) | 17/10/2016 | NCBI |
| KY289591 | A/California/142/2016(H3N2) | 20/10/2016 | NCBI |
| KY289583 | A/California/137/2016(H3N2) | 2/10/2016 | NCBI |
| KY289627 | A/Washington/80/2016(H3N2) | 19/10/2016 | NCBI |
| KY289579 | A/Washington/79/2016(H3N2) | 14/10/2016 | NCBI |
| KY289599 | A/Iowa/31/2016(H3N2) | 18/10/2016 | NCBI |
| KY289614 | A/Hawaii/84/2016(H3N2) | 29/9/2016 | NCBI |
| KY289567 | A/Wisconsin/89/2016(H3N2) | 30/10/2016 | NCBI |
| CY208728 | A/Nebraska/22/2016(H3N2) | 6/9/2016 | NCBI |
| CY208736 | A/NewHampshire/20/2016(H3N2) | 7/9/2016 | NCBI |
| CY208744 | A/Iowa/24/2016(H3N2) | 2/9/2016 | NCBI |
| CY208752 | A/Iowa/25/2016(H3N2) | 9/9/2016 | NCBI |
| CY208760 | A/Hawaii/67/2016(H3N2) | 4/9/2016 | NCBI |
| CY208768 | A/Hawaii/68/2016(H3N2) | 7/9/2016 | NCBI |
| CY208776 | A/Montana/46/2016(H3N2) | 9/9/2016 | NCBI |
| CY208784 | A/Illinois/46/2016(H3N2) | 14/9/2016 | NCBI |
| CY208792 | A/Michigan/101/2016(H3N2) | 6/9/2016 | NCBI |
| CY208800 | A/Michigan/103/2016(H3N2) | 11/9/2016 | NCBI |
| CY208808 | A/Nebraska/23/2016(H3N2) | 6/9/2016 | NCBI |
| CY208816 | A/Texas/183/2016(H3N2) | 31/8/2016 | NCBI |
| CY208824 | A/Texas/185/2016(H3N2) | 1/9/2016 | NCBI |
| CY208832 | A/Florida/74/2016(H3N2) | 7/9/2016 | NCBI |
| CY208840 | A/Florida/75/2016(H3N2) | 7/9/2016 | NCBI |
| CY208848 | A/Texas/182/2016(H3N2) | 31/8/2016 | NCBI |
| CY208856 | A/NorthCarolina/51/2016(H3N2) | 28/8/2016 | NCBI |
| CY208864 | A/NorthCarolina/52/2016(H3N2) | 8/9/2016 | NCBI |
| CY208872 | A/RhodeIsland/25/2016(H3N2) | 13/9/2016 | NCBI |

| | | | |
|---|---|---|---|
| CY208880 | A/RhodeIsland/26/2016(H3N2) | 13/9/2016 | NCBI |
| CY208888 | A/RhodeIsland/27/2016(H3N2) | 13/9/2016 | NCBI |
| CY208896 | A/RhodeIsland/28/2016(H3N2) | 13/9/2016 | NCBI |
| CY208904 | A/RhodeIsland/29/2016(H3N2) | 13/9/2016 | NCBI |
| CY208912 | A/Maine/21/2016(H3N2) | 12/9/2016 | NCBI |
| CY208920 | A/Alaska/25/2016(H3N2) | 10/9/2016 | NCBI |
| CY208928 | A/Alaska/24/2016(H3N2) | 8/9/2016 | NCBI |
| CY208936 | A/Pennsylvania/86/2016(H3N2) | 14/9/2016 | NCBI |
| CY208944 | A/Hawaii/71/2016(H3N2) | 13/9/2016 | NCBI |
| CY208952 | A/Minnesota/75/2016(H3N2) | 10/10/2016 | NCBI |
| CY208960 | A/Florida/82/2016(H3N2) | 3/10/2016 | NCBI |
| CY208968 | A/Florida/83/2016(H3N2) | 14/10/2016 | NCBI |
| CY208976 | A/NorthCarolina/56/2016(H3N2) | 26/9/2016 | NCBI |
| CY208984 | A/NorthCarolina/57/2016(H3N2) | 17/10/2016 | NCBI |
| CY208992 | A/Iowa/31/2016(H3N2) | 18/10/2016 | NCBI |
| CY209000 | A/Delaware/28/2016(H3N2) | 27/10/2016 | NCBI |
| CY209008 | A/Florida/87/2016(H3N2) | 27/10/2016 | NCBI |
| CY209016 | A/NewHampshire/22/2016(H3N2) | 20/10/2016 | NCBI |
| CY209024 | A/Wisconsin/89/2016(H3N2) | 30/10/2016 | NCBI |
| CY209032 | A/Alabama/13/2016(H3N2) | 19/10/2016 | NCBI |
| CY209040 | A/Virginia/61/2016(H3N2) | 24/10/2016 | NCBI |
| CY209048 | A/Pennsylvania/92/2016(H3N2) | 29/10/2016 | NCBI |
| CY209056 | A/WestVirginia/06/2016(H3N2) | 25/10/2016 | NCBI |
| CY209064 | A/Florida/88/2016(H3N2) | 28/10/2016 | NCBI |
| CY209072 | A/Tennessee/29/2016(H3N2) | 25/10/2016 | NCBI |
| CY209080 | A/NorthCarolina/58/2016(H3N2) | 27/10/2016 | NCBI |
| CY209088 | A/Colorado/40/2016(H3N2) | 29/10/2016 | NCBI |
| CY209096 | A/Colorado/41/2016(H3N2) | 5/11/2016 | NCBI |
| CY209104 | A/Iowa/33/2016(H3N2) | 27/10/2016 | NCBI |
| CY209112 | A/Iowa/34/2016(H3N2) | 30/10/2016 | NCBI |
| CY209120 | A/Alaska/28/2016(H3N2) | 31/10/2016 | NCBI |
| CY209128 | A/Alaska/29/2016(H3N2) | 31/10/2016 | NCBI |
| CY209136 | A/NewMexico/49/2016(H3N2) | 27/10/2016 | NCBI |
| CY209144 | A/NewMexico/50/2016(H3N2) | 27/10/2016 | NCBI |
| CY209152 | A/Hawaii/87/2016(H3N2) | 13/10/2016 | NCBI |
| CY209160 | A/Hawaii/89/2016(H3N2) | 21/10/2016 | NCBI |
| CY209168 | A/Hawaii/90/2016(H3N2) | 21/10/2016 | NCBI |

| | | | |
|---|---|---|---|
| CY209176 | A/Hawaii/91/2016(H3N2) | 25/10/2016 | NCBI |
| CY209184 | A/SouthDakota/28/2016(H3N2) | 28/9/2016 | NCBI |
| CY209192 | A/SouthDakota/27/2016(H3N2) | 26/9/2016 | NCBI |
| CY209200 | A/SouthDakota/29/2016(H3N2) | 17/10/2016 | NCBI |
| CY209208 | A/SouthDakota/32/2016(H3N2) | 4/11/2016 | NCBI |
| CY209216 | A/Michigan/109/2016(H3N2) | 1/11/2016 | NCBI |
| CY209224 | A/Michigan/110/2016(H3N2) | 6/11/2016 | NCBI |
| CY209232 | A/Wisconsin/90/2016(H3N2) | 9/11/2016 | NCBI |
| CY209240 | A/Wisconsin/91/2016(H3N2) | 14/11/2016 | NCBI |
| CY209248 | A/Nevada/43/2016(H3N2) | 27/10/2016 | NCBI |
| CY209256 | A/Nevada/44/2016(H3N2) | 1/11/2016 | NCBI |
| CY209264 | A/Nevada/45/2016(H3N2) | 3/11/2016 | NCBI |
| CY209272 | A/Nevada/46/2016(H3N2) | 8/11/2016 | NCBI |
| CY209280 | A/Oregon/20/2016(H3N2) | 1/11/2016 | NCBI |
| CY209288 | A/Oregon/21/2016(H3N2) | 2/11/2016 | NCBI |
| CY209296 | A/Washington/83/2016(H3N2) | 25/10/2016 | NCBI |
| CY209304 | A/Washington/84/2016(H3N2) | 25/10/2016 | NCBI |
| CY209312 | A/Washington/85/2016(H3N2) | 31/10/2016 | NCBI |
| CY209320 | A/Washington/86/2016(H3N2) | 9/11/2016 | NCBI |
| CY209328 | A/Utah/46/2016(H3N2) | 11/11/2016 | NCBI |
| CY209336 | A/Utah/47/2016(H3N2) | 11/11/2016 | NCBI |
| CY209344 | A/Oklahoma/14/2016(H3N2) | 9/11/2016 | NCBI |
| CY209352 | A/Illinois/48/2016(H3N2) | 13/11/2016 | NCBI |
| CY209360 | A/Minnesota/78/2016(H3N2) | 3/11/2016 | NCBI |
| CY209368 | A/Minnesota/79/2016(H3N2) | 12/11/2016 | NCBI |
| CY209376 | A/Montana/56/2016(H3N2) | 27/10/2016 | NCBI |
| CY209384 | A/Montana/58/2016(H3N2) | 5/11/2016 | NCBI |
| CY209392 | A/California/154/2016(H3N2) | 8/10/2016 | NCBI |
| CY209400 | A/California/155/2016(H3N2) | 10/10/2016 | NCBI |
| CY209408 | A/California/156/2016(H3N2) | 15/10/2016 | NCBI |
| CY209416 | A/California/157/2016(H3N2) | 17/10/2016 | NCBI |
| CY209424 | A/California/158/2016(H3N2) | 10/11/2016 | NCBI |
| KY321927 | A/HongKong/01/1968(H3N2) | 1968 | NCBI |
| CY210171 | A/Louisiana/24/2016(H3N2) | 25/10/2016 | NCBI |
| CY210179 | A/Louisiana/25/2016(H3N2) | 2/11/2016 | NCBI |
| CY210187 | A/Louisiana/27/2016(H3N2) | 15/11/2016 | NCBI |
| CY210195 | A/Louisiana/28/2016(H3N2) | 15/11/2016 | NCBI |

| | | | |
|---|---|---|---|
| CY210252 | A/Delaware/28/2016(H3N2) | 27/10/2016 | NCBI |
| CY210260 | A/Pennsylvania/90/2016(H3N2) | 18/10/2016 | NCBI |
| CY210268 | A/Massachusetts/38/2016(H3N2) | 22/10/2016 | NCBI |
| CY210290 | A/NewYork/102/2016(H3N2) | 18/10/2016 | NCBI |
| CY210298 | A/NewYork/103/2016(H3N2) | 26/10/2016 | NCBI |
| CY210306 | A/Connecticut/37/2016(H3N2) | 5/11/2016 | NCBI |
| CY210314 | A/Connecticut/36/2016(H3N2) | 30/10/2016 | NCBI |
| CY210322 | A/Ohio/39/2016(H3N2) | 22/10/2016 | NCBI |
| CY210337 | A/Florida/91/2016(H3N2) | 24/10/2016 | NCBI |
| CY210345 | A/Florida/90/2016(H3N2) | 22/10/2016 | NCBI |
| CY210353 | A/Pennsylvania/93/2016(H3N2) | 9/11/2016 | NCBI |
| CY210361 | A/Pennsylvania/95/2016(H3N2) | 15/11/2016 | NCBI |
| CY210369 | A/Pennsylvania/96/2016(H3N2) | 15/11/2016 | NCBI |
| CY210377 | A/Delaware/32/2016(H3N2) | 15/11/2016 | NCBI |
| CY210393 | A/WestVirginia/08/2016(H3N2) | 31/10/2016 | NCBI |
| CY210401 | A/NewYork/104/2016(H3N2) | 25/10/2016 | NCBI |
| CY210409 | A/NewYork/105/2016(H3N2) | 1/11/2016 | NCBI |
| CY210424 | A/Delaware/34/2016(H3N2) | 17/11/2016 | NCBI |
| CY210432 | A/Alaska/30/2016(H3N2) | 10/11/2016 | NCBI |
| CY210440 | A/Alaska/31/2016(H3N2) | 20/11/2016 | NCBI |
| CY210448 | A/Wyoming/32/2016(H3N2) | 9/11/2016 | NCBI |
| CY210456 | A/Colorado/42/2016(H3N2) | 5/11/2016 | NCBI |
| CY210464 | A/Colorado/43/2016(H3N2) | 13/11/2016 | NCBI |
| CY210472 | A/Maryland/20/2016(H3N2) | 4/11/2016 | NCBI |
| CY210480 | A/Maryland/21/2016(H3N2) | 17/11/2016 | NCBI |
| CY210488 | A/NorthCarolina/59/2016(H3N2) | 15/11/2016 | NCBI |
| CY210496 | A/NorthCarolina/60/2016(H3N2) | 16/11/2016 | NCBI |
| CY210504 | A/Michigan/112/2016(H3N2) | 14/11/2016 | NCBI |
| CY210512 | A/Michigan/111/2016(H3N2) | 4/11/2016 | NCBI |
| CY210520 | A/Hawaii/94/2016(H3N2) | 3/11/2016 | NCBI |
| CY210528 | A/Hawaii/95/2016(H3N2) | 7/11/2016 | NCBI |
| CY210536 | A/NewMexico/51/2016(H3N2) | 18/11/2016 | NCBI |
| CY210544 | A/NewHampshire/24/2016(H3N2) | 18/11/2016 | NCBI |
| CY210559 | A/Florida/94/2016(H3N2) | 7/11/2016 | NCBI |
| CY210567 | A/Florida/92/2016(H3N2) | 5/11/2016 | NCBI |
| CY210575 | A/Florida/95/2016(H3N2) | 11/11/2016 | NCBI |
| CY210583 | A/Florida/96/2016(H3N2) | 15/11/2016 | NCBI |

| | | | |
|---|---|---|---|
| CY210591 | A/Utah/48/2016(H3N2) | 20/11/2016 | NCBI |
| CY210599 | A/Utah/49/2016(H3N2) | 25/11/2016 | NCBI |
| CY210607 | A/Idaho/41/2016(H3N2) | 22/11/2016 | NCBI |
| CY210615 | A/Idaho/40/2016(H3N2) | 18/11/2016 | NCBI |
| CY210623 | A/Nebraska/25/2016(H3N2) | 16/11/2016 | NCBI |
| CY210631 | A/Missouri/19/2016(H3N2) | 21/11/2016 | NCBI |
| CY210639 | A/California/163/2016(H3N2) | 19/11/2016 | NCBI |
| CY210647 | A/California/162/2016(H3N2) | 15/11/2016 | NCBI |
| CY210655 | A/California/160/2016(H3N2) | 10/11/2016 | NCBI |
| CY210663 | A/California/161/2016(H3N2) | 11/11/2016 | NCBI |
| CY210671 | A/Wisconsin/92/2016(H3N2) | 25/11/2016 | NCBI |
| CY210679 | A/Wisconsin/93/2016(H3N2) | 26/11/2016 | NCBI |
| CY210687 | A/Alabama/16/2016(H3N2) | 27/11/2016 | NCBI |
| CY210695 | A/Alabama/15/2016(H3N2) | 16/11/2016 | NCBI |
| CY210703 | A/Pennsylvania/97/2016(H3N2) | 11/11/2016 | NCBI |
| CY210711 | A/Pennsylvania/98/2016(H3N2) | 19/11/2016 | NCBI |
| CY210719 | A/Maryland/23/2016(H3N2) | 26/11/2016 | NCBI |
| CY210727 | A/Maryland/24/2016(H3N2) | 29/11/2016 | NCBI |
| CY210735 | A/Georgia/62/2016(H3N2) | 4/11/2016 | NCBI |
| CY210743 | A/Georgia/61/2016(H3N2) | 7/10/2016 | NCBI |
| CY210751 | A/Georgia/59/2016(H3N2) | 2/10/2016 | NCBI |
| CY210759 | A/Ohio/40/2016(H3N2) | 17/11/2016 | NCBI |
| CY210767 | A/Ohio/41/2016(H3N2) | 17/11/2016 | NCBI |
| CY210775 | A/NewYork/106/2016(H3N2) | 15/11/2016 | NCBI |
| CY210783 | A/NewYork/107/2016(H3N2) | 26/11/2016 | NCBI |
| CY210791 | A/Massachusetts/41/2016(H3N2) | 29/11/2016 | NCBI |
| CY210799 | A/Virginia/62/2016(H3N2) | 30/11/2016 | NCBI |
| CY210807 | A/Virginia/63/2016(H3N2) | 30/11/2016 | NCBI |
| CY210815 | A/Vermont/33/2016(H3N2) | 19/11/2016 | NCBI |
| CY210823 | A/Vermont/34/2016(H3N2) | 1/12/2016 | NCBI |
| CY211474 | A/Louisiana/26/2016(H3N2) | 14/11/2016 | NCBI |
| CY211482 | A/Louisiana/31/2016(H3N2) | 26/11/2016 | NCBI |
| CY211490 | A/Louisiana/30/2016(H3N2) | 23/11/2016 | NCBI |
| CY211498 | A/Michigan/113/2016(H3N2) | 17/11/2016 | NCBI |
| CY211506 | A/Michigan/114/2016(H3N2) | 26/11/2016 | NCBI |
| CY211514 | A/Alaska/26/2016(H3N2) | 20/9/2016 | NCBI |
| CY211522 | A/Alaska/27/2016(H3N2) | 25/9/2016 | NCBI |

| | | | |
|---|---|---|---|
| CY211530 | A/Hawaii/73/2016(H3N2) | 16/9/2016 | NCBI |
| CY211538 | A/Hawaii/74/2016(H3N2) | 16/9/2016 | NCBI |
| CY211546 | A/Hawaii/77/2016(H3N2) | 22/9/2016 | NCBI |
| CY211554 | A/Hawaii/78/2016(H3N2) | 22/9/2016 | NCBI |
| CY211562 | A/Hawaii/76/2016(H3N2) | 21/9/2016 | NCBI |
| CY211570 | A/Wyoming/29/2016(H3N2) | 3/10/2016 | NCBI |
| CY211578 | A/California/126/2016(H3N2) | 10/8/2016 | NCBI |
| CY211586 | A/California/129/2016(H3N2) | 5/10/2016 | NCBI |
| CY211594 | A/Washington/77/2016(H3N2) | 3/10/2016 | NCBI |
| CY211602 | A/Arizona/70/2016(H3N2) | 27/9/2016 | NCBI |
| CY211610 | A/Idaho/35/2016(H3N2) | 27/9/2016 | NCBI |
| CY211618 | A/California/135/2016(H3N2) | 9/10/2016 | NCBI |
| CY211626 | A/California/139/2016(H3N2) | 16/10/2016 | NCBI |
| CY211634 | A/California/141/2016(H3N2) | 20/10/2016 | NCBI |
| CY211642 | A/Oregon/19/2016(H3N2) | 19/10/2016 | NCBI |
| CY211650 | A/Oregon/18/2016(H3N2) | 18/10/2016 | NCBI |
| CY211658 | A/Vermont/32/2016(H3N2) | 8/10/2016 | NCBI |
| CY211666 | A/Florida/86/2016(H3N2) | 17/10/2016 | NCBI |
| CY211674 | A/Pennsylvania/90/2016(H3N2) | 18/10/2016 | NCBI |
| CY211689 | A/NewYork/103/2016(H3N2) | 26/10/2016 | NCBI |
| CY211708 | A/NewJersey/26/2016(H3N2) | 17/11/2016 | NCBI |
| CY211716 | A/Mississippi/17/2016(H3N2) | 23/11/2016 | NCBI |
| CY211731 | A/Illinois/49/2016(H3N2) | 15/11/2016 | NCBI |
| CY211739 | A/Iowa/36/2016(H3N2) | 15/11/2016 | NCBI |
| CY211747 | A/Iowa/35/2016(H3N2) | 14/11/2016 | NCBI |
| CY211755 | A/Minnesota/80/2016(H3N2) | 12/11/2016 | NCBI |
| CY211763 | A/Indiana/43/2016(H3N2) | 29/11/2016 | NCBI |
| CY211771 | A/Kentucky/30/2016(H3N2) | 5/12/2016 | NCBI |
| CY211779 | A/Texas/191/2016(H3N2) | 3/12/2016 | NCBI |
| CY211787 | A/Massachusetts/40/2016(H3N2) | 18/11/2016 | NCBI |
| CY211795 | A/Delaware/37/2016(H3N2) | 4/12/2016 | NCBI |
| CY211810 | A/Montana/62/2016(H3N2) | 22/11/2016 | NCBI |
| CY211818 | A/Montana/64/2016(H3N2) | 3/12/2016 | NCBI |
| CY211826 | A/California/165/2016(H3N2) | 17/11/2016 | NCBI |
| CY211834 | A/California/166/2016(H3N2) | 17/11/2016 | NCBI |
| CY211842 | A/California/164/2016(H3N2) | 2016 | NCBI |
| CY211850 | A/California/168/2016(H3N2) | 25/11/2016 | NCBI |

| | | | |
|---|---|---|---|
| CY211858 | A/California/167/2016(H3N2) | 24/11/2016 | NCBI |
| CY211866 | A/California/169/2016(H3N2) | 25/11/2016 | NCBI |
| CY211874 | A/NewHampshire/25/2016(H3N2) | 27/11/2016 | NCBI |
| CY211882 | A/Tennessee/32/2016(H3N2) | 5/12/2016 | NCBI |
| CY211890 | A/Tennessee/33/2016(H3N2) | 6/12/2016 | NCBI |
| CY211898 | A/Arizona/78/2016(H3N2) | 24/11/2016 | NCBI |
| CY211906 | A/Arizona/77/2016(H3N2) | 22/11/2016 | NCBI |
| CY211914 | A/Hawaii/96/2016(H3N2) | 15/11/2016 | NCBI |
| CY211922 | A/Hawaii/97/2016(H3N2) | 21/11/2016 | NCBI |
| CY211930 | A/Oregon/22/2016(H3N2) | 26/11/2016 | NCBI |
| CY211938 | A/Oregon/23/2016(H3N2) | 26/11/2016 | NCBI |
| CY211946 | A/Iowa/37/2016(H3N2) | 24/11/2016 | NCBI |
| CY211954 | A/Iowa/38/2016(H3N2) | 4/12/2016 | NCBI |
| CY211962 | A/WestVirginia/10/2016(H3N2) | 25/11/2016 | NCBI |
| CY211970 | A/WestVirginia/11/2016(H3N2) | 7/12/2016 | NCBI |
| CY211978 | A/NorthCarolina/62/2016(H3N2) | 2/12/2016 | NCBI |
| CY211986 | A/NorthCarolina/63/2016(H3N2) | 8/12/2016 | NCBI |
| CY211994 | A/Florida/100/2016(H3N2) | 27/11/2016 | NCBI |
| CY212002 | A/Florida/101/2016(H3N2) | 29/11/2016 | NCBI |
| CY212010 | A/Florida/98/2016(H3N2) | 20/11/2016 | NCBI |
| CY212018 | A/Illinois/50/2016(H3N2) | 10/12/2016 | NCBI |
| CY212033 | A/Wisconsin/98/2016(H3N2) | 6/12/2016 | NCBI |
| CY212041 | A/Maine/24/2016(H3N2) | 8/12/2016 | NCBI |
| CY212049 | A/Maine/25/2016(H3N2) | 14/12/2016 | NCBI |
| CY212057 | A/Florida/103/2016(H3N2) | 8/12/2016 | NCBI |
| CY212065 | A/Florida/102/2016(H3N2) | 1/12/2016 | NCBI |
| CY212073 | A/Oklahoma/17/2016(H3N2) | 30/11/2016 | NCBI |
| CY212081 | A/Oklahoma/18/2016(H3N2) | 2/12/2016 | NCBI |
| CY212338 | A/Delaware/35/2016(H3N2) | 20/11/2016 | NCBI |
| CY212346 | A/Missouri/19/2016(H3N2) | 21/11/2016 | NCBI |
| CY212354 | A/Mississippi/17/2016(H3N2) | 23/11/2016 | NCBI |
| CY212362 | A/Pennsylvania/97/2016(H3N2) | 11/11/2016 | NCBI |
| CY212370 | A/NorthDakota/28/2016(H3N2) | 15/12/2016 | NCBI |
| CY212378 | A/Minnesota/82/2016(H3N2) | 7/12/2016 | NCBI |
| CY212386 | A/Minnesota/83/2016(H3N2) | 11/12/2016 | NCBI |
| CY212394 | A/Maryland/25/2016(H3N2) | 21/10/2016 | NCBI |
| CY212402 | A/Maryland/27/2016(H3N2) | 2/12/2016 | NCBI |

| | | | |
|---|---|---|---|
| CY212410 | A/Maryland/26/2016(H3N2) | 28/11/2016 | NCBI |
| CY212418 | A/Massachusetts/42/2016(H3N2) | 7/12/2016 | NCBI |
| CY212426 | A/Massachusetts/43/2016(H3N2) | 16/12/2016 | NCBI |
| CY212434 | A/Virginia/64/2016(H3N2) | 6/12/2016 | NCBI |
| CY212442 | A/Virginia/66/2016(H3N2) | 9/12/2016 | NCBI |
| CY212450 | A/Virginia/65/2016(H3N2) | 6/12/2016 | NCBI |
| CY212458 | A/Virginia/67/2016(H3N2) | 13/12/2016 | NCBI |
| CY212466 | A/Virginia/68/2016(H3N2) | 14/12/2016 | NCBI |
| CY212474 | A/Alaska/32/2016(H3N2) | 5/12/2016 | NCBI |
| CY212482 | A/Alaska/33/2016(H3N2) | 5/12/2016 | NCBI |
| CY212490 | A/California/179/2016(H3N2) | 3/12/2016 | NCBI |
| CY212498 | A/California/180/2016(H3N2) | 9/12/2016 | NCBI |
| CY212506 | A/California/181/2016(H3N2) | 11/12/2016 | NCBI |
| CY212514 | A/California/177/2016(H3N2) | 28/11/2016 | NCBI |
| CY212522 | A/California/178/2016(H3N2) | 28/11/2016 | NCBI |
| CY212530 | A/California/176/2016(H3N2) | 26/11/2016 | NCBI |
| CY212538 | A/California/175/2016(H3N2) | 5/11/2016 | NCBI |
| CY212546 | A/Colorado/44/2016(H3N2) | 25/11/2016 | NCBI |
| CY212554 | A/Colorado/45/2016(H3N2) | 25/11/2016 | NCBI |
| CY212562 | A/Idaho/42/2016(H3N2) | 2/12/2016 | NCBI |
| CY212570 | A/Idaho/43/2016(H3N2) | 7/12/2016 | NCBI |
| CY212578 | A/NewMexico/54/2016(H3N2) | 26/11/2016 | NCBI |
| CY212586 | A/Nevada/49/2016(H3N2) | 21/11/2016 | NCBI |
| CY212594 | A/Nevada/50/2016(H3N2) | 29/11/2016 | NCBI |
| CY212602 | A/Nevada/51/2016(H3N2) | 29/11/2016 | NCBI |
| CY212610 | A/Nevada/53/2016(H3N2) | 6/12/2016 | NCBI |
| CY212618 | A/Nevada/52/2016(H3N2) | 5/12/2016 | NCBI |
| CY212626 | A/Nevada/54/2016(H3N2) | 8/12/2016 | NCBI |
| CY212634 | A/Nevada/55/2016(H3N2) | 8/12/2016 | NCBI |
| CY212642 | A/Utah/51/2016(H3N2) | 8/12/2016 | NCBI |
| CY212650 | A/Utah/50/2016(H3N2) | 6/12/2016 | NCBI |
| CY212658 | A/Wyoming/33/2016(H3N2) | 4/12/2016 | NCBI |
| CY212666 | A/Wyoming/34/2016(H3N2) | 7/12/2016 | NCBI |
| CY212674 | A/NewYork/109/2016(H3N2) | 30/11/2016 | NCBI |
| CY212682 | A/NewYork/110/2016(H3N2) | 6/12/2016 | NCBI |
| CY212690 | A/NewYork/111/2016(H3N2) | 14/12/2016 | NCBI |
| CY212698 | A/Illinois/51/2016(H3N2) | 13/12/2016 | NCBI |

| | | | |
|---|---|---|---|
| CY212706 | A/Illinois/52/2016(H3N2) | 13/12/2016 | NCBI |
| CY212714 | A/Pennsylvania/101/2016(H3N2) | 9/12/2016 | NCBI |
| CY212722 | A/Pennsylvania/102/2016(H3N2) | 9/12/2016 | NCBI |
| CY212730 | A/Pennsylvania/99/2016(H3N2) | 8/12/2016 | NCBI |
| CY212738 | A/Pennsylvania/100/2016(H3N2) | 8/12/2016 | NCBI |
| CY212746 | A/Nebraska/26/2016(H3N2) | 6/12/2016 | NCBI |
| CY212754 | A/Nebraska/27/2016(H3N2) | 13/12/2016 | NCBI |
| CY212762 | A/Missouri/20/2016(H3N2) | 11/12/2016 | NCBI |
| CY212770 | A/Michigan/117/2016(H3N2) | 7/12/2016 | NCBI |
| CY212778 | A/Michigan/118/2016(H3N2) | 8/12/2016 | NCBI |
| CY212786 | A/Alabama/17/2016(H3N2) | 14/12/2016 | NCBI |
| CY212794 | A/Alabama/18/2016(H3N2) | 14/12/2016 | NCBI |
| CY212802 | A/Vermont/35/2016(H3N2) | 5/12/2016 | NCBI |
| CY212810 | A/Vermont/36/2016(H3N2) | 13/12/2016 | NCBI |
| CY212818 | A/Texas/193/2016(H3N2) | 7/12/2016 | NCBI |
| CY212826 | A/Texas/195/2016(H3N2) | 19/12/2016 | NCBI |
| CY212834 | A/Indiana/44/2016(H3N2) | 2/12/2016 | NCBI |
| CY212842 | A/Indiana/45/2016(H3N2) | 14/12/2016 | NCBI |
| CY212850 | A/Arizona/79/2016(H3N2) | 10/12/2016 | NCBI |
| CY212858 | A/Arizona/80/2016(H3N2) | 12/12/2016 | NCBI |
| CY212866 | A/Hawaii/98/2016(H3N2) | 28/11/2016 | NCBI |
| CY212874 | A/Hawaii/99/2016(H3N2) | 3/12/2016 | NCBI |
| CY212882 | A/California/186/2016(H3N2) | 14/12/2016 | NCBI |
| CY212890 | A/California/184/2016(H3N2) | 13/12/2016 | NCBI |
| CY212898 | A/California/187/2016(H3N2) | 14/12/2016 | NCBI |
| CY212906 | A/California/185/2016(H3N2) | 13/12/2016 | NCBI |
| CY212914 | A/California/188/2016(H3N2) | 15/12/2016 | NCBI |
| CY212922 | A/California/189/2016(H3N2) | 15/12/2016 | NCBI |
| CY212930 | A/Montana/66/2016(H3N2) | 9/12/2016 | NCBI |
| CY212938 | A/Montana/67/2016(H3N2) | 9/12/2016 | NCBI |
| CY212946 | A/Washington/100/2016(H3N2) | 13/12/2016 | NCBI |
| CY212954 | A/Washington/101/2016(H3N2) | 15/12/2016 | NCBI |
| CY212962 | A/California/183/2016(H3N2) | 6/12/2016 | NCBI |
| CY212970 | A/California/182/2016(H3N2) | 2/12/2016 | NCBI |
| CY212978 | A/California/190/2016(H3N2) | 15/12/2016 | NCBI |
| CY212986 | A/California/191/2016(H3N2) | 15/12/2016 | NCBI |
| CY212994 | A/Delaware/39/2016(H3N2) | 19/12/2016 | NCBI |

| | | | |
|---|---|---|---|
| CY213002 | A/Delaware/40/2016(H3N2) | 19/12/2016 | NCBI |
| CY213010 | A/NewYork/112/2016(H3N2) | 1/12/2016 | NCBI |
| CY213018 | A/NewYork/113/2016(H3N2) | 3/12/2016 | NCBI |
| CY213026 | A/Kentucky/31/2016(H3N2) | 20/12/2016 | NCBI |
| CY213034 | A/NewHampshire/27/2016(H3N2) | 12/12/2016 | NCBI |
| CY213042 | A/NewHampshire/28/2016(H3N2) | 17/12/2016 | NCBI |
| CY213050 | A/Florida/104/2016(H3N2) | 1/12/2016 | NCBI |
| CY213058 | A/Florida/105/2016(H3N2) | 8/12/2016 | NCBI |
| CY213066 | A/Louisiana/32/2016(H3N2) | 6/12/2016 | NCBI |
| CY213074 | A/Louisiana/33/2016(H3N2) | 15/12/2016 | NCBI |
| CY213082 | A/Iowa/40/2016(H3N2) | 13/12/2016 | NCBI |
| CY213090 | A/Iowa/42/2016(H3N2) | 20/12/2016 | NCBI |
| CY213098 | A/Illinois/53/2016(H3N2) | 16/12/2016 | NCBI |
| CY213106 | A/Illinois/54/2016(H3N2) | 16/12/2016 | NCBI |
| CY213114 | A/Tennessee/34/2016(H3N2) | 12/12/2016 | NCBI |
| CY213122 | A/Wisconsin/102/2016(H3N2) | 19/12/2016 | NCBI |
| CY213130 | A/Minnesota/85/2016(H3N2) | 19/12/2016 | NCBI |
| CY213138 | A/Texas/197/2016(H3N2) | 25/12/2016 | NCBI |
| CY213146 | A/Texas/201/2016(H3N2) | 29/12/2016 | NCBI |
| CY213154 | A/Alaska/35/2016(H3N2) | 13/12/2016 | NCBI |
| CY213162 | A/Washington/106/2016(H3N2) | 18/12/2016 | NCBI |
| CY213170 | A/Washington/103/2016(H3N2) | 4/11/2016 | NCBI |
| CY213178 | A/Washington/102/2016(H3N2) | 31/10/2016 | NCBI |
| CY213186 | A/Washington/105/2016(H3N2) | 19/11/2016 | NCBI |
| CY213194 | A/Washington/104/2016(H3N2) | 15/11/2016 | NCBI |
| CY213202 | A/Washington/108/2016(H3N2) | 19/12/2016 | NCBI |
| CY213210 | A/Utah/53/2016(H3N2) | 22/12/2016 | NCBI |
| CY213218 | A/Utah/52/2016(H3N2) | 20/12/2016 | NCBI |
| CY213226 | A/NewMexico/55/2016(H3N2) | 14/12/2016 | NCBI |
| CY213234 | A/NewMexico/56/2016(H3N2) | 15/12/2016 | NCBI |
| CY213242 | A/California/192/2016(H3N2) | 14/12/2016 | NCBI |
| CY213250 | A/Texas/196/2016(H3N2) | 22/12/2016 | NCBI |
| CY213258 | A/Texas/198/2016(H3N2) | 25/12/2016 | NCBI |
| CY213266 | A/Mississippi/19/2016(H3N2) | 6/12/2016 | NCBI |
| CY213274 | A/Mississippi/20/2016(H3N2) | 13/12/2016 | NCBI |
| CY213282 | A/Texas/202/2016(H3N2) | 29/12/2016 | NCBI |
| CY213290 | A/Missouri/21/2016(H3N2) | 19/12/2016 | NCBI |

| | | | |
|---|---|---|---|
| CY213298 | A/NorthDakota/30/2016(H3N2) | 28/12/2016 | NCBI |
| CY213306 | A/NorthDakota/31/2016(H3N2) | 29/12/2016 | NCBI |
| CY213314 | A/Michigan/120/2016(H3N2) | 25/12/2016 | NCBI |
| CY213322 | A/Michigan/119/2016(H3N2) | 23/12/2016 | NCBI |
| CY213778 | A/Louisiana/26/2016(H3N2) | 14/11/2016 | NCBI |
| CY213786 | A/Louisiana/29/2016(H3N2) | 15/11/2016 | NCBI |
| CY213794 | A/Delaware/36/2016(H3N2) | 30/11/2016 | NCBI |
| CY213802 | A/California/133/2016(H3N2) | 3/10/2016 | NCBI |
| CY213810 | A/Ohio/38/2016(H3N2) | 16/10/2016 | NCBI |
| CY213818 | A/Nevada/39/2016(H3N2) | 12/10/2016 | NCBI |
| CY213826 | A/Alabama/14/2016(H3N2) | 19/10/2016 | NCBI |
| CY213834 | A/Texas/190/2016(H3N2) | 21/10/2016 | NCBI |
| CY213842 | A/Washington/82/2016(H3N2) | 16/10/2016 | NCBI |
| CY213850 | A/Minnesota/76/2016(H3N2) | 25/10/2016 | NCBI |
| CY213858 | A/California/146/2016(H3N2) | 17/10/2016 | NCBI |
| CY213866 | A/Ohio/39/2016(H3N2) | 22/10/2016 | NCBI |
| CY213874 | A/SouthDakota/32/2016(H3N2) | 4/11/2016 | NCBI |
| CY213882 | A/Michigan/110/2016(H3N2) | 6/11/2016 | NCBI |
| CY213890 | A/Wisconsin/90/2016(H3N2) | 9/11/2016 | NCBI |
| CY213898 | A/Wisconsin/91/2016(H3N2) | 14/11/2016 | NCBI |
| CY213906 | A/Oklahoma/14/2016(H3N2) | 9/11/2016 | NCBI |
| CY213914 | A/Pennsylvania/95/2016(H3N2) | 15/11/2016 | NCBI |
| CY213922 | A/WestVirginia/08/2016(H3N2) | 31/10/2016 | NCBI |
| CY213930 | A/NewYork/105/2016(H3N2) | 1/11/2016 | NCBI |
| CY213938 | A/Colorado/44/2016(H3N2) | 25/11/2016 | NCBI |
| CY213946 | A/Wyoming/34/2016(H3N2) | 7/12/2016 | NCBI |
| CY213954 | A/Washington/107/2016(H3N2) | 18/12/2016 | NCBI |
| CY213962 | A/Illinois/60/2016(H3N2) | 25/12/2016 | NCBI |
| CY213970 | A/Iowa/43/2016(H3N2) | 9/12/2016 | NCBI |
| CY213978 | A/Iowa/45/2016(H3N2) | 30/12/2016 | NCBI |
| CY213986 | A/Louisiana/34/2016(H3N2) | 19/12/2016 | NCBI |
| CY213994 | A/Louisiana/35/2016(H3N2) | 27/12/2016 | NCBI |
| CY214002 | A/Wisconsin/02/2017(H3N2) | 3/1/2017 | NCBI |
| CY214010 | A/Wisconsin/03/2017(H3N2) | 3/1/2017 | NCBI |
| CY214018 | A/Indiana/46/2016(H3N2) | 22/12/2016 | NCBI |
| CY214026 | A/Indiana/47/2016(H3N2) | 29/12/2016 | NCBI |
| CY214034 | A/Texas/01/2017(H3N2) | 2/1/2017 | NCBI |

| | | | |
|---|---|---|---|
| CY214042 | A/Illinois/55/2016(H3N2) | 14/12/2016 | NCBI |
| CY214050 | A/Illinois/56/2016(H3N2) | 15/12/2016 | NCBI |
| CY214058 | A/Illinois/61/2016(H3N2) | 26/12/2016 | NCBI |
| CY214066 | A/Illinois/57/2016(H3N2) | 16/12/2016 | NCBI |
| CY214074 | A/Illinois/01/2017(H3N2) | 3/1/2017 | NCBI |
| CY214082 | A/Illinois/63/2016(H3N2) | 30/12/2016 | NCBI |
| CY214090 | A/Illinois/62/2016(H3N2) | 29/12/2016 | NCBI |
| CY214098 | A/California/195/2016(H3N2) | 23/12/2016 | NCBI |
| CY214106 | A/California/199/2016(H3N2) | 25/12/2016 | NCBI |
| CY214114 | A/California/196/2016(H3N2) | 23/12/2016 | NCBI |
| CY214122 | A/California/197/2016(H3N2) | 23/12/2016 | NCBI |
| CY214130 | A/California/200/2016(H3N2) | 25/12/2016 | NCBI |
| CY214138 | A/California/201/2016(H3N2) | 25/12/2016 | NCBI |
| CY214146 | A/California/198/2016(H3N2) | 23/12/2016 | NCBI |
| CY214154 | A/California/202/2016(H3N2) | 25/12/2016 | NCBI |
| CY214162 | A/California/203/2016(H3N2) | 25/12/2016 | NCBI |
| CY214170 | A/California/204/2016(H3N2) | 25/12/2016 | NCBI |
| CY214178 | A/Colorado/47/2016(H3N2) | 30/11/2016 | NCBI |
| CY214186 | A/Colorado/48/2016(H3N2) | 5/12/2016 | NCBI |
| CY214194 | A/Montana/68/2016(H3N2) | 20/12/2016 | NCBI |
| CY214202 | A/Montana/69/2016(H3N2) | 23/12/2016 | NCBI |
| CY214210 | A/Hawaii/103/2016(H3N2) | 7/12/2016 | NCBI |
| CY214218 | A/Hawaii/102/2016(H3N2) | 6/12/2016 | NCBI |
| CY214226 | A/Kansas/28/2016(H3N2) | 21/12/2016 | NCBI |
| CY214234 | A/Kansas/29/2016(H3N2) | 24/12/2016 | NCBI |
| CY214242 | A/Kansas/01/2017(H3N2) | 1/1/2017 | NCBI |
| CY214250 | A/Kansas/30/2016(H3N2) | 28/12/2016 | NCBI |
| CY214346 | A/Pennsylvania/01/2017(H3N2) | 10/1/2017 | NCBI |
| CY214354 | A/Vermont/37/2016(H3N2) | 23/12/2016 | NCBI |
| CY214362 | A/Vermont/38/2016(H3N2) | 27/12/2016 | NCBI |
| CY214370 | A/Delaware/38/2016(H3N2) | 4/12/2016 | NCBI |
| CY214378 | A/NewHampshire/26/2016(H3N2) | 4/12/2016 | NCBI |
| CY214386 | A/Tennessee/35/2016(H3N2) | 13/12/2016 | NCBI |
| CY214394 | A/Kentucky/32/2016(H3N2) | 26/12/2016 | NCBI |
| CY214402 | A/Kentucky/33/2016(H3N2) | 26/12/2016 | NCBI |
| CY214410 | A/RhodeIsland/30/2016(H3N2) | 28/12/2016 | NCBI |
| CY214418 | A/RhodeIsland/31/2016(H3N2) | 29/12/2016 | NCBI |

| | | | |
|---|---|---|---|
| CY214426 | A/Pennsylvania/105/2016(H3N2) | 17/12/2016 | NCBI |
| CY214434 | A/Pennsylvania/106/2016(H3N2) | 21/12/2016 | NCBI |
| CY214442 | A/Pennsylvania/104/2016(H3N2) | 15/12/2016 | NCBI |
| CY214450 | A/Minnesota/85/2016(H3N2) | 19/12/2016 | NCBI |
| CY214458 | A/Massachusetts/44/2016(H3N2) | 18/12/2016 | NCBI |
| CY214466 | A/Massachusetts/45/2016(H3N2) | 18/12/2016 | NCBI |
| CY214474 | A/Maryland/30/2016(H3N2) | 12/12/2016 | NCBI |
| CY214482 | A/Maryland/31/2016(H3N2) | 19/12/2016 | NCBI |
| CY214490 | A/Florida/106/2016(H3N2) | 26/12/2016 | NCBI |
| CY214498 | A/Florida/107/2016(H3N2) | 26/12/2016 | NCBI |
| CY214506 | A/NorthCarolina/64/2016(H3N2) | 15/12/2016 | NCBI |
| CY214514 | A/NorthCarolina/65/2016(H3N2) | 19/12/2016 | NCBI |
| CY214522 | A/Georgia/64/2016(H3N2) | 24/12/2016 | NCBI |
| CY214530 | A/Georgia/65/2016(H3N2) | 29/12/2016 | NCBI |
| CY214538 | A/Delaware/02/2017(H3N2) | 4/1/2017 | NCBI |
| CY214546 | A/WestVirginia/12/2016(H3N2) | 7/12/2016 | NCBI |
| CY214554 | A/WestVirginia/13/2016(H3N2) | 13/12/2016 | NCBI |
| CY214562 | A/WestVirginia/14/2016(H3N2) | 16/12/2016 | NCBI |
| CY214570 | A/WestVirginia/01/2017(H3N2) | 3/1/2017 | NCBI |
| CY214578 | A/NewJersey/27/2016(H3N2) | 20/12/2016 | NCBI |
| CY214586 | A/NewJersey/28/2016(H3N2) | 30/12/2016 | NCBI |
| CY214594 | A/NewJersey/01/2017(H3N2) | 1/1/2017 | NCBI |
| CY214602 | A/Florida/108/2016(H3N2) | 16/12/2016 | NCBI |
| CY214610 | A/Florida/109/2016(H3N2) | 16/12/2016 | NCBI |
| CY214618 | A/Kentucky/02/2017(H3N2) | 4/1/2017 | NCBI |
| CY214626 | A/Kentucky/01/2017(H3N2) | 3/1/2017 | NCBI |
| CY214634 | A/Tennessee/37/2016(H3N2) | 27/12/2016 | NCBI |
| CY214642 | A/Tennessee/01/2017(H3N2) | 1/1/2017 | NCBI |
| CY214650 | A/Washington/110/2016(H3N2) | 27/12/2016 | NCBI |
| CY214658 | A/Washington/111/2016(H3N2) | 27/12/2016 | NCBI |
| CY214666 | A/NewMexico/57/2016(H3N2) | 25/12/2016 | NCBI |
| CY214674 | A/NewMexico/58/2016(H3N2) | 26/12/2016 | NCBI |
| CY214682 | A/NewHampshire/01/2017(H3N2) | 2/1/2017 | NCBI |
| CY214690 | A/NewHampshire/02/2017(H3N2) | 3/1/2017 | NCBI |
| CY214698 | A/Maine/02/2017(H3N2) | 6/1/2017 | NCBI |
| CY214706 | A/Maine/01/2017(H3N2) | 4/1/2017 | NCBI |
| CY214714 | A/Kentucky/05/2017(H3N2) | 9/1/2017 | NCBI |

| | | | |
|---|---|---|---|
| CY214722 | A/NewYork/114/2016(H3N2) | 29/12/2016 | NCBI |
| CY214730 | A/NewYork/01/2017(H3N2) | 2/1/2017 | NCBI |
| CY214738 | A/Alaska/36/2016(H3N2) | 27/12/2016 | NCBI |
| CY214746 | A/Alaska/37/2016(H3N2) | 30/12/2016 | NCBI |
| CY214754 | A/Oregon/01/2017(H3N2) | 2/1/2017 | NCBI |
| CY214762 | A/Oregon/02/2017(H3N2) | 2/1/2017 | NCBI |
| CY214770 | A/Utah/02/2017(H3N2) | 5/1/2017 | NCBI |
| CY214778 | A/Utah/01/2017(H3N2) | 4/1/2017 | NCBI |
| CY214786 | A/Wyoming/01/2017(H3N2) | 3/1/2017 | NCBI |
| CY214794 | A/Wyoming/02/2017(H3N2) | 5/1/2017 | NCBI |
| CY214802 | A/Connecticut/38/2016(H3N2) | 30/11/2016 | NCBI |
| CY214810 | A/Connecticut/39/2016(H3N2) | 4/12/2016 | NCBI |
| CY214818 | A/Texas/10/2017(H3N2) | 10/1/2017 | NCBI |
| CY214826 | A/Texas/07/2017(H3N2) | 5/1/2017 | NCBI |
| CY214834 | A/Maryland/01/2017(H3N2) | 4/1/2017 | NCBI |
| CY214842 | A/Maryland/03/2017(H3N2) | 6/1/2017 | NCBI |
| CY214850 | A/Minnesota/02/2017(H3N2) | 7/1/2017 | NCBI |
| CY214858 | A/Minnesota/03/2017(H3N2) | 9/1/2017 | NCBI |
| CY214866 | A/California/213/2016(H3N2) | 30/12/2016 | NCBI |
| CY214874 | A/California/212/2016(H3N2) | 29/12/2016 | NCBI |
| CY214882 | A/California/214/2016(H3N2) | 31/12/2016 | NCBI |
| CY214890 | A/California/206/2016(H3N2) | 24/12/2016 | NCBI |
| CY214898 | A/California/209/2016(H3N2) | 26/12/2016 | NCBI |
| CY214906 | A/California/207/2016(H3N2) | 24/12/2016 | NCBI |
| CY214914 | A/California/211/2016(H3N2) | 28/12/2016 | NCBI |
| CY214922 | A/California/208/2016(H3N2) | 25/12/2016 | NCBI |
| CY214930 | A/California/210/2016(H3N2) | 27/12/2016 | NCBI |
| CY214938 | A/Virginia/04/2017(H3N2) | 3/1/2017 | NCBI |
| CY214946 | A/Virginia/03/2017(H3N2) | 2/1/2017 | NCBI |
| CY214954 | A/Virginia/01/2017(H3N2) | 1/1/2017 | NCBI |
| CY214962 | A/Virginia/02/2017(H3N2) | 1/1/2017 | NCBI |
| CY214970 | A/Hawaii/104/2016(H3N2) | 23/12/2016 | NCBI |
| CY214978 | A/Hawaii/105/2016(H3N2) | 30/12/2016 | NCBI |
| CY214986 | A/Colorado/01/2017(H3N2) | 5/1/2017 | NCBI |
| CY214994 | A/Colorado/02/2017(H3N2) | 5/1/2017 | NCBI |
| CY215002 | A/Massachusetts/01/2017(H3N2) | 4/1/2017 | NCBI |
| CY215010 | A/Massachusetts/02/2017(H3N2) | 7/1/2017 | NCBI |

| | | | |
|---|---|---|---|
| CY215018 | A/Missouri/02/2017(H3N2) | 10/1/2017 | NCBI |
| CY215026 | A/Missouri/01/2017(H3N2) | 7/1/2017 | NCBI |
| CY215034 | A/Texas/05/2017(H3N2) | 3/1/2017 | NCBI |
| CY215042 | A/Washington/04/2017(H3N2) | 4/1/2017 | NCBI |
| CY215050 | A/Washington/05/2017(H3N2) | 4/1/2017 | NCBI |
| CY215058 | A/Montana/70/2016(H3N2) | 29/12/2016 | NCBI |
| CY215066 | A/Montana/01/2017(H3N2) | 1/1/2017 | NCBI |
| CY215074 | A/Nevada/58/2016(H3N2) | 19/12/2016 | NCBI |
| CY215082 | A/Nevada/59/2016(H3N2) | 22/12/2016 | NCBI |
| CY215090 | A/Nevada/56/2016(H3N2) | 8/12/2016 | NCBI |
| CY215098 | A/Nevada/57/2016(H3N2) | 10/12/2016 | NCBI |
| CY215106 | A/Florida/01/2017(H3N2) | 4/1/2017 | NCBI |
| CY215114 | A/Florida/02/2017(H3N2) | 11/1/2017 | NCBI |
| CY215122 | A/Michigan/122/2016(H3N2) | 6/12/2016 | NCBI |
| CY215130 | A/Michigan/124/2016(H3N2) | 23/12/2016 | NCBI |
| CY215138 | A/Michigan/125/2016(H3N2) | 23/12/2016 | NCBI |
| CY215146 | A/Oklahoma/02/2017(H3N2) | 2/1/2017 | NCBI |
| CY215154 | A/Oklahoma/03/2017(H3N2) | 2/1/2017 | NCBI |
| CY215162 | A/Mississippi/02/2017(H3N2) | 12/1/2017 | NCBI |
| CY215170 | A/Mississippi/01/2017(H3N2) | 11/1/2017 | NCBI |
| CY215665 | A/Pennsylvania/103/2016(H3N2) | 21/12/2016 | NCBI |
| CY215673 | A/Louisiana/25/2016(H3N2) | 2/11/2016 | NCBI |
| CY215681 | A/Louisiana/27/2016(H3N2) | 15/11/2016 | NCBI |
| CY215689 | A/Louisiana/30/2016(H3N2) | 23/11/2016 | NCBI |
| CY215697 | A/Texas/192/2016(H3N2) | 21/12/2016 | NCBI |
| CY215705 | A/Hawaii/87/2016(H3N2) | 13/10/2016 | NCBI |
| CY215713 | A/NewHampshire/24/2016(H3N2) | 18/11/2016 | NCBI |
| CY215721 | A/Florida/94/2016(H3N2) | 7/11/2016 | NCBI |
| CY215729 | A/Nebraska/25/2016(H3N2) | 16/11/2016 | NCBI |
| CY215737 | A/Wisconsin/93/2016(H3N2) | 26/11/2016 | NCBI |
| CY215745 | A/Illinois/49/2016(H3N2) | 15/11/2016 | NCBI |
| CY215753 | A/Iowa/36/2016(H3N2) | 15/11/2016 | NCBI |
| CY215761 | A/Minnesota/80/2016(H3N2) | 12/11/2016 | NCBI |
| CY215769 | A/Maryland/24/2016(H3N2) | 29/11/2016 | NCBI |
| CY215777 | A/Massachusetts/41/2016(H3N2) | 29/11/2016 | NCBI |
| CY215785 | A/Vermont/34/2016(H3N2) | 1/12/2016 | NCBI |
| CY215793 | A/Montana/64/2016(H3N2) | 3/12/2016 | NCBI |

| | | | |
|---|---|---|---|
| CY215801 | A/California/168/2016(H3N2) | 25/11/2016 | NCBI |
| CY215809 | A/Arizona/77/2016(H3N2) | 22/11/2016 | NCBI |
| CY215817 | A/Hawaii/96/2016(H3N2) | 15/11/2016 | NCBI |
| CY215825 | A/Iowa/37/2016(H3N2) | 24/11/2016 | NCBI |
| CY215833 | A/Illinois/50/2016(H3N2) | 10/12/2016 | NCBI |
| CY215841 | A/Wisconsin/97/2016(H3N2) | 6/12/2016 | NCBI |
| CY215849 | A/Colorado/49/2016(H3N2) | 26/12/2016 | NCBI |
| CY215857 | A/Hawaii/106/2016(H3N2) | 12/12/2016 | NCBI |
| CY215865 | A/Idaho/45/2016(H3N2) | 2016 | NCBI |
| CY215873 | A/Idaho/46/2016(H3N2) | 24/12/2016 | NCBI |
| CY215881 | A/Idaho/02/2017(H3N2) | 7/1/2017 | NCBI |
| CY215889 | A/Illinois/08/2017(H3N2) | 15/1/2017 | NCBI |
| CY215897 | A/Illinois/11/2017(H3N2) | 17/1/2017 | NCBI |
| CY215905 | A/NorthCarolina/03/2017(H3N2) | 18/1/2017 | NCBI |
| CY215913 | A/NorthCarolina/01/2017(H3N2) | 8/1/2017 | NCBI |
| CY215921 | A/Connecticut/43/2016(H3N2) | 28/12/2016 | NCBI |
| CY215929 | A/Connecticut/44/2016(H3N2) | 29/12/2016 | NCBI |
| CY215937 | A/Iowa/01/2017(H3N2) | 17/1/2017 | NCBI |
| CY215945 | A/Iowa/02/2017(H3N2) | 17/1/2017 | NCBI |
| CY215953 | A/Florida/03/2017(H3N2) | 7/1/2017 | NCBI |
| CY215961 | A/Delaware/04/2017(H3N2) | 17/1/2017 | NCBI |
| CY215969 | A/Delaware/05/2017(H3N2) | 17/1/2017 | NCBI |
| CY215977 | A/Illinois/07/2017(H3N2) | 12/1/2017 | NCBI |
| CY215985 | A/Illinois/09/2017(H3N2) | 16/1/2017 | NCBI |
| CY215993 | A/Illinois/06/2017(H3N2) | 10/1/2017 | NCBI |
| CY216001 | A/Illinois/10/2017(H3N2) | 16/1/2017 | NCBI |
| CY216009 | A/Illinois/04/2017(H3N2) | 7/1/2017 | NCBI |
| CY216017 | A/Illinois/05/2017(H3N2) | 8/1/2017 | NCBI |
| CY216025 | A/Georgia/03/2017(H3N2) | 9/1/2017 | NCBI |
| CY216033 | A/Georgia/04/2017(H3N2) | 9/1/2017 | NCBI |
| CY216041 | A/Tennessee/02/2017(H3N2) | 14/1/2017 | NCBI |
| CY216049 | A/Tennessee/04/2017(H3N2) | 15/1/2017 | NCBI |
| CY216057 | A/Alaska/01/2017(H3N2) | 8/1/2017 | NCBI |
| CY216065 | A/Alaska/02/2017(H3N2) | 11/1/2017 | NCBI |
| CY216073 | A/NewMexico/02/2017(H3N2) | 10/1/2017 | NCBI |
| CY216081 | A/NewMexico/01/2017(H3N2) | 9/1/2017 | NCBI |
| CY216089 | A/Nevada/01/2017(H3N2) | 1/1/2017 | NCBI |

| | | | |
|---|---|---|---|
| CY216097 | A/Nevada/02/2017(H3N2) | 1/1/2017 | NCBI |
| CY216105 | A/Pennsylvania/118/2016(H3N2) | 27/12/2016 | NCBI |
| CY216113 | A/Pennsylvania/04/2017(H3N2) | 9/1/2017 | NCBI |
| CY216121 | A/Pennsylvania/119/2016(H3N2) | 30/12/2016 | NCBI |
| CY216129 | A/Kentucky/09/2017(H3N2) | 17/1/2017 | NCBI |
| CY216137 | A/Kentucky/07/2017(H3N2) | 16/1/2017 | NCBI |
| CY216145 | A/NewHampshire/04/2017(H3N2) | 20/1/2017 | NCBI |
| CY216153 | A/NewHampshire/03/2017(H3N2) | 18/1/2017 | NCBI |
| CY216161 | A/California/02/2017(H3N2) | 3/1/2017 | NCBI |
| CY216169 | A/NorthDakota/02/2017(H3N2) | 14/1/2017 | NCBI |
| CY216177 | A/Ohio/02/2017(H3N2) | 10/1/2017 | NCBI |
| CY216185 | A/NorthDakota/03/2017(H3N2) | 16/1/2017 | NCBI |
| CY216193 | A/Louisiana/03/2017(H3N2) | 11/1/2017 | NCBI |
| CY216201 | A/Louisiana/02/2017(H3N2) | 10/1/2017 | NCBI |
| CY216209 | A/SouthDakota/04/2017(H3N2) | 15/1/2017 | NCBI |
| CY216217 | A/SouthDakota/05/2017(H3N2) | 15/1/2017 | NCBI |
| CY216225 | A/SouthDakota/02/2017(H3N2) | 5/1/2017 | NCBI |
| CY216233 | A/SouthDakota/03/2017(H3N2) | 5/1/2017 | NCBI |
| CY216241 | A/SouthDakota/35/2016(H3N2) | 20/12/2016 | NCBI |
| CY216249 | A/SouthDakota/36/2016(H3N2) | 21/12/2016 | NCBI |
| CY216257 | A/SouthDakota/33/2016(H3N2) | 11/12/2016 | NCBI |
| CY216265 | A/SouthDakota/34/2016(H3N2) | 13/12/2016 | NCBI |
| CY216273 | A/NewJersey/03/2017(H3N2) | 7/1/2017 | NCBI |
| CY216281 | A/NewJersey/04/2017(H3N2) | 8/1/2017 | NCBI |
| CY216289 | A/NewJersey/05/2017(H3N2) | 15/1/2017 | NCBI |
| CY216297 | A/NewJersey/02/2017(H3N2) | 5/1/2017 | NCBI |
| CY216305 | A/NewJersey/06/2017(H3N2) | 18/1/2017 | NCBI |
| CY216313 | A/Maine/04/2017(H3N2) | 22/1/2017 | NCBI |
| CY216321 | A/Maine/03/2017(H3N2) | 20/1/2017 | NCBI |
| CY216329 | A/Maine/07/2017(H3N2) | 23/1/2017 | NCBI |
| CY216337 | A/Maine/05/2017(H3N2) | 22/1/2017 | NCBI |
| CY216345 | A/Maine/06/2017(H3N2) | 22/1/2017 | NCBI |
| CY216353 | A/Utah/05/2017(H3N2) | 19/1/2017 | NCBI |
| CY216361 | A/Utah/03/2017(H3N2) | 15/1/2017 | NCBI |
| CY216369 | A/Oregon/03/2017(H3N2) | 9/1/2017 | NCBI |
| CY216377 | A/Oregon/04/2017(H3N2) | 16/1/2017 | NCBI |
| CY216385 | A/Wyoming/03/2017(H3N2) | 12/1/2017 | NCBI |

| | | | |
|---|---|---|---|
| CY216393 | A/Wyoming/04/2017(H3N2) | 16/1/2017 | NCBI |
| CY216401 | A/Nebraska/01/2017(H3N2) | 9/1/2017 | NCBI |
| CY216409 | A/Nebraska/02/2017(H3N2) | 10/1/2017 | NCBI |
| CY216417 | A/California/03/2017(H3N2) | 5/1/2017 | NCBI |
| CY216425 | A/California/08/2017(H3N2) | 10/1/2017 | NCBI |
| CY216433 | A/California/09/2017(H3N2) | 10/1/2017 | NCBI |
| CY216441 | A/Wisconsin/15/2017(H3N2) | 20/1/2017 | NCBI |
| CY216449 | A/Wisconsin/14/2017(H3N2) | 15/1/2017 | NCBI |
| CY216457 | A/Vermont/02/2017(H3N2) | 17/1/2017 | NCBI |
| CY216465 | A/Vermont/01/2017(H3N2) | 15/1/2017 | NCBI |
| CY216473 | A/Connecticut/03/2017(H3N2) | 9/1/2017 | NCBI |
| CY216481 | A/Connecticut/04/2017(H3N2) | 11/1/2017 | NCBI |
| CY216489 | A/RhodeIsland/01/2017(H3N2) | 17/1/2017 | NCBI |
| CY216497 | A/RhodeIsland/02/2017(H3N2) | 18/1/2017 | NCBI |
| CY216505 | A/Arkansas/05/2017(H3N2) | 18/1/2017 | NCBI |
| CY216513 | A/Arkansas/03/2017(H3N2) | 17/1/2017 | NCBI |
| CY216521 | A/Arkansas/04/2017(H3N2) | 17/1/2017 | NCBI |
| CY216529 | A/Arkansas/02/2017(H3N2) | 10/1/2017 | NCBI |
| CY216537 | A/Arkansas/01/2017(H3N2) | 9/1/2017 | NCBI |
| CY216545 | A/Arkansas/15/2016(H3N2) | 27/12/2016 | NCBI |
| CY216553 | A/Arkansas/14/2016(H3N2) | 16/12/2016 | NCBI |
| CY216561 | A/SouthCarolina/04/2017(H3N2) | 17/1/2017 | NCBI |
| CY216569 | A/SouthCarolina/02/2017(H3N2) | 4/1/2017 | NCBI |
| CY216577 | A/Texas/19/2017(H3N2) | 18/1/2017 | NCBI |
| CY216585 | A/Texas/20/2017(H3N2) | 18/1/2017 | NCBI |
| CY216593 | A/Indiana/05/2017(H3N2) | 11/1/2017 | NCBI |
| CY216601 | A/Indiana/06/2017(H3N2) | 13/1/2017 | NCBI |
| CY216609 | A/Alabama/02/2017(H3N2) | 5/1/2017 | NCBI |
| CY216617 | A/Alabama/01/2017(H3N2) | 4/1/2017 | NCBI |
| CY216625 | A/Alabama/03/2017(H3N2) | 19/1/2017 | NCBI |
| CY216633 | A/Alabama/04/2017(H3N2) | 23/1/2017 | NCBI |
| CY216641 | A/Maryland/04/2017(H3N2) | 10/1/2017 | NCBI |
| CY216649 | A/Maryland/05/2017(H3N2) | 12/1/2017 | NCBI |
| CY216657 | A/Virginia/07/2017(H3N2) | 18/1/2017 | NCBI |
| CY216665 | A/Virginia/08/2017(H3N2) | 22/1/2017 | NCBI |
| CY216673 | A/WestVirginia/15/2016(H3N2) | 27/12/2016 | NCBI |
| CY216681 | A/WestVirginia/02/2017(H3N2) | 14/1/2017 | NCBI |

| | | | |
|---|---|---|---|
| CY216689 | A/Minnesota/05/2017(H3N2) | 19/1/2017 | NCBI |
| CY216697 | A/Minnesota/07/2017(H3N2) | 24/1/2017 | NCBI |
| CY216705 | A/Florida/05/2017(H3N2) | 20/1/2017 | NCBI |
| CY216713 | A/Massachusetts/03/2017(H3N2) | 27/1/2017 | NCBI |
| CY216721 | A/Oklahoma/05/2017(H3N2) | 23/1/2017 | NCBI |
| CY216729 | A/Oklahoma/06/2017(H3N2) | 25/1/2017 | NCBI |
| CY216737 | A/Texas/23/2017(H3N2) | 26/1/2017 | NCBI |
| CY216745 | A/Texas/24/2017(H3N2) | 26/1/2017 | NCBI |
| CY216753 | A/NewYork/03/2017(H3N2) | 24/1/2017 | NCBI |
| CY216761 | A/NewYork/02/2017(H3N2) | 22/1/2017 | NCBI |
| CY216769 | A/Missouri/03/2017(H3N2) | 5/1/2017 | NCBI |
| CY216777 | A/Mississippi/03/2017(H3N2) | 25/1/2017 | NCBI |
| CY216785 | A/Mississippi/04/2017(H3N2) | 27/1/2017 | NCBI |
| CY216793 | A/NorthDakota/04/2017(H3N2) | 27/1/2017 | NCBI |
| CY216801 | A/NorthDakota/05/2017(H3N2) | 29/1/2017 | NCBI |
| KY509835 | A/Chungbuk/107/2016(H3N2) | 18/1/2016 | NCBI |
| KY509836 | A/Gyeonggi/458/2016(H3N2) | 22/2/2016 | NCBI |
| KY509837 | A/Seoul/1595/2015(H3N2) | 14/9/2015 | NCBI |
| KY487530 | A/Columbia/P0044/2014(H3N2) | 21/12/2014 | NCBI |
| KY487322 | A/Columbia/P0045/2014(H3N2) | 21/12/2014 | NCBI |
| KY487641 | A/Bethesda/P0055/2015(H3N2) | 8/1/2015 | NCBI |
| KY487154 | A/Columbia/P0079/2014(H3N2) | 25/12/2014 | NCBI |
| CY217882 | A/Maryland/23/2016(H3N2) | 26/11/2016 | NCBI |
| CY218929 | A/Delaware/03/2017(H3N2) | 8/1/2017 | NCBI |
| CY217890 | A/Michigan/115/2016(H3N2) | 17/11/2016 | NCBI |
| CY217906 | A/Michigan/116/2016(H3N2) | 12/12/2016 | NCBI |
| CY217914 | A/Washington/87/2016(H3N2) | 29/11/2016 | NCBI |
| CY217922 | A/Washington/90/2016(H3N2) | 5/12/2016 | NCBI |
| CY217930 | A/Washington/91/2016(H3N2) | 5/12/2016 | NCBI |
| CY217938 | A/Washington/92/2016(H3N2) | 5/12/2016 | NCBI |
| CY217946 | A/Washington/93/2016(H3N2) | 6/12/2016 | NCBI |
| CY217954 | A/Wisconsin/95/2016(H3N2) | 29/11/2016 | NCBI |
| CY217962 | A/Wisconsin/96/2016(H3N2) | 13/12/2016 | NCBI |
| CY217970 | A/Washington/94/2016(H3N2) | 12/12/2016 | NCBI |
| CY217978 | A/Washington/95/2016(H3N2) | 13/12/2016 | NCBI |
| CY217986 | A/Washington/97/2016(H3N2) | 13/12/2016 | NCBI |
| CY217994 | A/Washington/98/2016(H3N2) | 27/12/2016 | NCBI |

| | | | |
|---|---|---|---|
| CY218002 | A/Washington/99/2016(H3N2) | 27/12/2016 | NCBI |
| CY218030 | A/Michigan/01/2017(H3N2) | 3/1/2017 | NCBI |
| CY218038 | A/Michigan/02/2017(H3N2) | 3/1/2017 | NCBI |
| CY218046 | A/Michigan/03/2017(H3N2) | 3/1/2017 | NCBI |
| CY218054 | A/Michigan/04/2017(H3N2) | 4/1/2017 | NCBI |
| CY218062 | A/Michigan/05/2017(H3N2) | 4/1/2017 | NCBI |
| CY218086 | A/Washington/02/2017(H3N2) | 10/1/2017 | NCBI |
| CY218094 | A/Washington/03/2017(H3N2) | 10/1/2017 | NCBI |
| CY218110 | A/Wisconsin/06/2017(H3N2) | 13/1/2017 | NCBI |
| CY218118 | A/Wisconsin/07/2017(H3N2) | 13/1/2017 | NCBI |
| CY218126 | A/Wisconsin/08/2017(H3N2) | 13/1/2017 | NCBI |
| CY218142 | A/Pennsylvania/107/2016(H3N2) | 9/12/2016 | NCBI |
| CY218150 | A/Pennsylvania/111/2016(H3N2) | 14/12/2016 | NCBI |
| CY218166 | A/Pennsylvania/113/2016(H3N2) | 15/12/2016 | NCBI |
| CY218174 | A/Pennsylvania/114/2016(H3N2) | 15/12/2016 | NCBI |
| CY218182 | A/Pennsylvania/116/2016(H3N2) | 21/12/2016 | NCBI |
| CY218198 | A/Pennsylvania/117/2016(H3N2) | 28/12/2016 | NCBI |
| CY218206 | A/Pennsylvania/02/2017(H3N2) | 3/1/2017 | NCBI |
| CY218222 | A/Pennsylvania/03/2017(H3N2) | 10/1/2017 | NCBI |
| CY218952 | A/Minnesota/82/2016(H3N2) | 7/12/2016 | NCBI |
| CY218960 | A/NewYork/111/2016(H3N2) | 14/12/2016 | NCBI |
| CY218982 | A/Colorado/49/2016(H3N2) | 26/12/2016 | NCBI |
| CY218997 | A/Hawaii/106/2016(H3N2) | 12/12/2016 | NCBI |
| CY219005 | A/Mississippi/20/2016(H3N2) | 13/12/2016 | NCBI |
| CY219047 | A/Kentucky/05/2017(H3N2) | 9/1/2017 | NCBI |
| CY219055 | A/NewYork/114/2016(H3N2) | 29/12/2016 | NCBI |
| CY219076 | A/Kansas/29/2016(H3N2) | 24/12/2016 | NCBI |
| CY219098 | A/Virginia/05/2017(H3N2) | 5/1/2017 | NCBI |
| CY219110 | A/Missouri/01/2017(H3N2) | 7/1/2017 | NCBI |
| CY219138 | A/Washington/12/2017(H3N2) | 4/1/2017 | NCBI |
| CY219146 | A/Washington/11/2017(H3N2) | 3/1/2017 | NCBI |
| CY219154 | A/Colorado/03/2017(H3N2) | 18/1/2017 | NCBI |
| CY219162 | A/Colorado/04/2017(H3N2) | 24/1/2017 | NCBI |
| CY219170 | A/Arizona/01/2017(H3N2) | 2/1/2017 | NCBI |
| CY219178 | A/Arizona/81/2016(H3N2) | 27/12/2016 | NCBI |
| CY219186 | A/Montana/03/2017(H3N2) | 17/1/2017 | NCBI |
| CY219194 | A/Montana/05/2017(H3N2) | 23/1/2017 | NCBI |

| | | | |
|---|---|---|---|
| CY219209 | A/Idaho/04/2017(H3N2) | 17/1/2017 | NCBI |
| CY219217 | A/Idaho/06/2017(H3N2) | 19/1/2017 | NCBI |
| CY219244 | A/California/11/2017(H3N2) | 20/1/2017 | NCBI |
| CY219252 | A/California/12/2017(H3N2) | 20/1/2017 | NCBI |
| CY219260 | A/Nevada/03/2017(H3N2) | 11/1/2017 | NCBI |
| CY219268 | A/Nevada/04/2017(H3N2) | 14/1/2017 | NCBI |
| CY219276 | A/Nevada/05/2017(H3N2) | 17/1/2017 | NCBI |
| CY220031 | A/Oregon/19/2016(H3N2) | 19/10/2016 | NCBI |
| CY220039 | A/Maryland/23/2016(H3N2) | 26/11/2016 | NCBI |
| CY220047 | A/Georgia/01/2017(H3N2) | 10/1/2017 | NCBI |
| CY220063 | A/Texas/11/2017(H3N2) | 22/1/2017 | NCBI |
| CY220071 | A/Texas/12/2017(H3N2) | 24/1/2017 | NCBI |
| CY220079 | A/Texas/13/2017(H3N2) | 26/1/2017 | NCBI |
| CY220087 | A/Texas/14/2017(H3N2) | 27/1/2017 | NCBI |
| CY220095 | A/Delaware/06/2017(H3N2) | 26/1/2017 | NCBI |
| CY220103 | A/Delaware/01/2017(H3N2) | 3/1/2017 | NCBI |
| CY220111 | A/Delaware/07/2017(H3N2) | 2/2/2017 | NCBI |
| CY220119 | A/Delaware/08/2017(H3N2) | 2/2/2017 | NCBI |
| CY220127 | A/Connecticut/06/2017(H3N2) | 18/1/2017 | NCBI |
| CY220135 | A/Texas/32/2017(H3N2) | 31/1/2017 | NCBI |
| CY220143 | A/Texas/33/2017(H3N2) | 31/1/2017 | NCBI |
| CY220151 | A/Michigan/19/2017(H3N2) | 11/1/2017 | NCBI |
| CY220159 | A/Michigan/20/2017(H3N2) | 15/1/2017 | NCBI |
| CY220167 | A/Iowa/03/2017(H3N2) | 12/1/2017 | NCBI |
| CY220175 | A/Iowa/05/2017(H3N2) | 23/1/2017 | NCBI |
| CY220183 | A/NewJersey/07/2017(H3N2) | 30/1/2017 | NCBI |
| CY220191 | A/NewJersey/08/2017(H3N2) | 30/1/2017 | NCBI |
| CY220199 | A/NewJersey/09/2017(H3N2) | 30/1/2017 | NCBI |
| CY220207 | A/NewJersey/10/2017(H3N2) | 30/1/2017 | NCBI |
| CY220215 | A/NewJersey/11/2017(H3N2) | 2/2/2017 | NCBI |
| CY220223 | A/Oregon/06/2017(H3N2) | 24/1/2017 | NCBI |
| CY220231 | A/Oregon/05/2017(H3N2) | 14/1/2017 | NCBI |
| CY220239 | A/Alaska/04/2017(H3N2) | 24/1/2017 | NCBI |
| CY220247 | A/Alaska/03/2017(H3N2) | 23/1/2017 | NCBI |
| CY220255 | A/California/18/2017(H3N2) | 28/1/2017 | NCBI |
| CY220263 | A/California/20/2017(H3N2) | 29/1/2017 | NCBI |
| CY220271 | A/California/21/2017(H3N2) | 30/1/2017 | NCBI |

| | | | |
|---|---|---|---|
| CY220279 | A/Indiana/08/2017(H3N2) | 23/1/2017 | NCBI |
| CY220287 | A/Indiana/11/2017(H3N2) | 31/1/2017 | NCBI |
| CY220295 | A/Indiana/07/2017(H3N2) | 15/1/2017 | NCBI |
| CY220303 | A/Texas/35/2017(H3N2) | 1/2/2017 | NCBI |
| CY220311 | A/Texas/36/2017(H3N2) | 4/2/2017 | NCBI |
| CY220319 | A/NewHampshire/06/2017(H3N2) | 31/1/2017 | NCBI |
| CY220327 | A/NewHampshire/07/2017(H3N2) | 5/2/2017 | NCBI |
| CY220335 | A/Ohio/04/2017(H3N2) | 1/2/2017 | NCBI |
| CY220343 | A/Maine/08/2017(H3N2) | 31/1/2017 | NCBI |
| CY220351 | A/Maine/09/2017(H3N2) | 5/2/2017 | NCBI |
| CY220359 | A/Florida/07/2017(H3N2) | 16/1/2017 | NCBI |
| CY220367 | A/Florida/08/2017(H3N2) | 24/1/2017 | NCBI |
| CY220375 | A/Kentucky/11/2017(H3N2) | 27/1/2017 | NCBI |
| CY220383 | A/Kentucky/12/2017(H3N2) | 27/1/2017 | NCBI |
| CY220391 | A/California/15/2017(H3N2) | 3/1/2017 | NCBI |
| CY220399 | A/California/22/2017(H3N2) | 1/2/2017 | NCBI |
| CY220407 | A/NewMexico/04/2017(H3N2) | 22/1/2017 | NCBI |
| CY220415 | A/NewMexico/05/2017(H3N2) | 23/1/2017 | NCBI |
| CY220423 | A/Arizona/04/2017(H3N2) | 30/1/2017 | NCBI |
| CY220431 | A/Arizona/05/2017(H3N2) | 31/1/2017 | NCBI |
| CY220439 | A/Wyoming/05/2017(H3N2) | 24/1/2017 | NCBI |
| CY220447 | A/Wyoming/06/2017(H3N2) | 26/1/2017 | NCBI |
| CY220455 | A/Utah/08/2017(H3N2) | 4/2/2017 | NCBI |
| CY220463 | A/Utah/06/2017(H3N2) | 31/1/2017 | NCBI |
| CY220471 | A/Vermont/03/2017(H3N2) | 25/1/2017 | NCBI |
| CY220479 | A/Vermont/04/2017(H3N2) | 1/2/2017 | NCBI |
| CY220487 | A/Illinois/12/2017(H3N2) | 31/1/2017 | NCBI |
| CY220495 | A/Illinois/13/2017(H3N2) | 1/2/2017 | NCBI |
| CY220503 | A/Wisconsin/19/2017(H3N2) | 23/1/2017 | NCBI |
| CY220511 | A/Wisconsin/20/2017(H3N2) | 23/1/2017 | NCBI |
| CY220519 | A/NorthCarolina/05/2017(H3N2) | 30/1/2017 | NCBI |
| CY220527 | A/NorthCarolina/04/2017(H3N2) | 26/1/2017 | NCBI |
| CY220535 | A/Louisiana/04/2017(H3N2) | 24/1/2017 | NCBI |
| CY220543 | A/Louisiana/05/2017(H3N2) | 30/1/2017 | NCBI |
| CY220551 | A/Alabama/06/2017(H3N2) | 7/2/2017 | NCBI |
| CY220559 | A/Alabama/07/2017(H3N2) | 7/2/2017 | NCBI |
| CY220567 | A/Kansas/04/2017(H3N2) | 4/2/2017 | NCBI |

| | | | |
|---|---|---|---|
| CY220575 | A/Kansas/03/2017(H3N2) | 3/2/2017 | NCBI |
| CY220583 | A/Colorado/05/2017(H3N2) | 31/1/2017 | NCBI |
| CY220591 | A/Colorado/06/2017(H3N2) | 31/1/2017 | NCBI |
| CY220599 | A/Washington/16/2017(H3N2) | 25/1/2017 | NCBI |
| CY220607 | A/Washington/17/2017(H3N2) | 27/1/2017 | NCBI |
| CY220615 | A/Hawaii/107/2016(H3N2) | 28/12/2016 | NCBI |
| CY220623 | A/Hawaii/02/2017(H3N2) | 15/1/2017 | NCBI |
| CY220631 | A/Hawaii/03/2017(H3N2) | 17/1/2017 | NCBI |
| CY220639 | A/Montana/07/2017(H3N2) | 29/1/2017 | NCBI |
| CY220647 | A/Montana/06/2017(H3N2) | 26/1/2017 | NCBI |
| CY220655 | A/Minnesota/08/2017(H3N2) | 27/1/2017 | NCBI |
| CY220663 | A/Minnesota/10/2017(H3N2) | 6/2/2017 | NCBI |
| CY220671 | A/California/26/2017(H3N2) | 31/1/2017 | NCBI |
| CY220679 | A/California/27/2017(H3N2) | 2/2/2017 | NCBI |
| CY220687 | A/California/25/2017(H3N2) | 28/1/2017 | NCBI |
| CY220695 | A/California/28/2017(H3N2) | 3/2/2017 | NCBI |
| CY220703 | A/California/24/2017(H3N2) | 9/1/2017 | NCBI |
| CY220711 | A/NorthDakota/06/2017(H3N2) | 7/2/2017 | NCBI |
| CY220719 | A/NorthDakota/07/2017(H3N2) | 8/2/2017 | NCBI |
| CY220727 | A/Texas/40/2017(H3N2) | 13/2/2017 | NCBI |
| CY220735 | A/Texas/38/2017(H3N2) | 1/2/2017 | NCBI |
| CY220743 | A/Michigan/22/2017(H3N2) | 8/2/2017 | NCBI |
| CY220751 | A/Arizona/08/2017(H3N2) | 5/2/2017 | NCBI |
| CY220759 | A/Missouri/05/2017(H3N2) | 26/1/2017 | NCBI |
| CY220767 | A/Missouri/06/2017(H3N2) | 27/1/2017 | NCBI |
| KY487182 | A/Columbia/P0064/2014(H3N2) | 27/12/2014 | NCBI |
| KY615552 | A/Columbia/P0047/2014(H3N2) | 27/12/2014 | NCBI |
| KY487462 | A/Linkou/0061/2016(H3N2) | 18/1/2016 | NCBI |
| KY487741 | A/Bethesda/P0061/2015(H3N2) | 15/1/2015 | NCBI |
| KY487654 | A/Columbia/P0112/2014(H3N2) | 27/12/2014 | NCBI |
| KY487272 | A/Columbia/P0041/2014(H3N2) | 21/12/2014 | NCBI |
| KY615379 | A/Bethesda/P0040/2015(H3N2) | 3/1/2015 | NCBI |
| KY615543 | A/Keelung/0026/2016(H3N2) | 15/1/2016 | NCBI |
| KY487580 | A/Bethesda/P0044/2015(H3N2) | 10/1/2015 | NCBI |
| KY615391 | A/Baltimore/0148/2016(H3N2) | 9/4/2016 | NCBI |
| KY487317 | A/Bethesda/P0018/2015(H3N2) | 13/1/2015 | NCBI |
| KY487284 | A/Bethesda/P0053/2015(H3N2) | 11/1/2015 | NCBI |

| | | | |
|---|---|---|---|
| KY615469 | A/Bethesda/P0045/2015(H3N2) | 13/1/2015 | NCBI |
| KY615405 | A/Columbia/P0067/2014(H3N2) | 26/12/2014 | NCBI |
| KY487069 | A/Columbia/P0038/2014(H3N2) | 21/12/2014 | NCBI |
| KY487559 | A/Baltimore/P0040/2015(H3N2) | 8/1/2015 | NCBI |
| KY487421 | A/Bethesda/P0060/2015(H3N2) | 19/1/2015 | NCBI |
| KY487630 | A/Bethesda/P0004/2015(H3N2) | 3/1/2015 | NCBI |
| KY615529 | A/Bethesda/P0051/2015(H3N2) | 15/1/2015 | NCBI |
| KY487157 | A/Columbia/P0035/2014(H3N2) | 19/12/2014 | NCBI |
| KY487132 | A/Taipei/0011/2015(H3N2) | 24/12/2015 | NCBI |
| KY615506 | A/Columbia/P0075/2014(H3N2) | 27/12/2014 | NCBI |
| KY487207 | A/Columbia/P0087/2015(H3N2) | 22/1/2015 | NCBI |
| KY487270 | A/Keelung/0023/2016(H3N2) | 7/1/2016 | NCBI |
| KY487356 | A/Columbia/P0090/2015(H3N2) | 22/1/2015 | NCBI |
| KY615520 | A/Columbia/P0072/2014(H3N2) | 26/12/2014 | NCBI |
| KY487318 | A/Linkou/0081/2016(H3N2) | 26/1/2016 | NCBI |
| KY487176 | A/Baltimore/P0008/2014(H3N2) | 14/12/2014 | NCBI |
| KY615514 | A/Bethesda/P0054/2015(H3N2) | 13/1/2015 | NCBI |
| KY615528 | A/Taipei/0020/2016(H3N2) | 5/1/2016 | NCBI |
| KY487765 | A/Columbia/P0033/2014(H3N2) | 19/12/2014 | NCBI |
| CY221758 | A/Texas/15/2017(H3N2) | 20/1/2017 | NCBI |
| CY221766 | A/Texas/16/2017(H3N2) | 23/1/2017 | NCBI |
| CY221774 | A/Florida/97/2016(H3N2) | 15/11/2016 | NCBI |
| CY221782 | A/Texas/203/2016(H3N2) | 22/12/2016 | NCBI |
| CY221790 | A/Illinois/58/2016(H3N2) | 17/12/2016 | NCBI |
| CY221798 | A/Massachusetts/02/2017(H3N2) | 7/1/2017 | NCBI |
| CY221806 | A/NewHampshire/07/2017(H3N2) | 5/2/2017 | NCBI |
| CY221814 | A/Washington/15/2017(H3N2) | 24/1/2017 | NCBI |
| CY221822 | A/Wisconsin/23/2017(H3N2) | 12/2/2017 | NCBI |
| CY221830 | A/Louisiana/08/2017(H3N2) | 7/2/2017 | NCBI |
| CY221838 | A/Louisiana/09/2017(H3N2) | 15/2/2017 | NCBI |
| CY221846 | A/Texas/45/2017(H3N2) | 13/2/2017 | NCBI |
| CY221854 | A/Texas/46/2017(H3N2) | 13/2/2017 | NCBI |
| CY221862 | A/Indiana/13/2017(H3N2) | 3/2/2017 | NCBI |
| CY221870 | A/Indiana/14/2017(H3N2) | 3/2/2017 | NCBI |
| CY221878 | A/Iowa/06/2017(H3N2) | 1/2/2017 | NCBI |
| CY221886 | A/Iowa/08/2017(H3N2) | 14/2/2017 | NCBI |
| CY221894 | A/Nebraska/04/2017(H3N2) | 2/2/2017 | NCBI |

| | | | |
|---|---|---|---|
| CY221902 | A/Nebraska/03/2017(H3N2) | 19/1/2017 | NCBI |
| CY221910 | A/Texas/50/2017(H3N2) | 16/2/2017 | NCBI |
| CY221918 | A/Texas/52/2017(H3N2) | 20/2/2017 | NCBI |
| CY221926 | A/Texas/47/2017(H3N2) | 13/2/2017 | NCBI |
| CY221934 | A/Texas/48/2017(H3N2) | 13/2/2017 | NCBI |
| CY221942 | A/Delaware/09/2017(H3N2) | 15/2/2017 | NCBI |
| CY221957 | A/NewMexico/06/2017(H3N2) | 7/2/2017 | NCBI |
| CY221965 | A/NewMexico/07/2017(H3N2) | 8/2/2017 | NCBI |
| CY221973 | A/California/29/2017(H3N2) | 29/1/2017 | NCBI |
| CY221981 | A/Wyoming/07/2017(H3N2) | 9/2/2017 | NCBI |
| CY221989 | A/Wyoming/08/2017(H3N2) | 13/2/2017 | NCBI |
| CY221997 | A/Utah/09/2017(H3N2) | 15/2/2017 | NCBI |
| CY222005 | A/Idaho/09/2017(H3N2) | 24/1/2017 | NCBI |
| CY222013 | A/Idaho/10/2017(H3N2) | 1/2/2017 | NCBI |
| CY222021 | A/Washington/20/2017(H3N2) | 8/2/2017 | NCBI |
| CY222029 | A/Washington/23/2017(H3N2) | 12/2/2017 | NCBI |
| CY222037 | A/Nevada/06/2017(H3N2) | 6/2/2017 | NCBI |
| CY222045 | A/Nevada/07/2017(H3N2) | 8/2/2017 | NCBI |
| CY222053 | A/Alaska/05/2017(H3N2) | 9/2/2017 | NCBI |
| CY222061 | A/Alaska/06/2017(H3N2) | 9/2/2017 | NCBI |
| KY681553 | A/SaudiArabia/01/2013(H3N2) | 13/10/2013 | NCBI |
| KY681556 | A/SaudiArabia/04/2013(H3N2) | 15/10/2013 | NCBI |
| KY681557 | A/SaudiArabia/05/2013(H3N2) | 15/10/2013 | NCBI |
| KY681558 | A/SaudiArabia/06/2014(H3N2) | 2/10/2014 | NCBI |
| KY681559 | A/SaudiArabia/07/2014(H3N2) | 4/10/2014 | NCBI |
| KY681560 | A/SaudiArabia/08/2014(H3N2) | 4/10/2014 | NCBI |
| KY681564 | A/SaudiArabia/15/2015(H3N2) | 22/9/2015 | NCBI |
| KY681565 | A/SaudiArabia/16/2015(H3N2) | 22/9/2015 | NCBI |
| KY681566 | A/SaudiArabia/18/2015(H3N2) | 22/9/2015 | NCBI |
| KY681567 | A/SaudiArabia/19/2015(H3N2) | 22/9/2015 | NCBI |
| KY681568 | A/SaudiArabia/20/2015(H3N2) | 22/9/2015 | NCBI |
| KY681571 | A/SaudiArabia/24/2015(H3N2) | 25/9/2015 | NCBI |
| KY681572 | A/SaudiArabia/25/2015(H3N2) | 25/9/2015 | NCBI |
| KY681573 | A/SaudiArabia/26/2015(H3N2) | 25/9/2015 | NCBI |
| KY681574 | A/SaudiArabia/27/2015(H3N2) | 25/9/2015 | NCBI |
| KY681575 | A/SaudiArabia/28/2015(H3N2) | 25/9/2015 | NCBI |
| KY681576 | A/SaudiArabia/29/2015(H3N2) | 25/9/2015 | NCBI |

| | | | |
|---|---|---|---|
| KY681577 | A/SaudiArabia/31/2015(H3N2) | 25/9/2015 | NCBI |
| CY222533 | A/Texas/12/2017(H3N2) | 24/1/2017 | NCBI |
| CY222541 | A/Texas/53/2017(H3N2) | 22/2/2017 | NCBI |
| CY222549 | A/SouthCarolina/06/2017(H3N2) | 23/2/2017 | NCBI |
| CY222557 | A/SouthCarolina/07/2017(H3N2) | 23/2/2017 | NCBI |
| CY222565 | A/Florida/03/2017(H3N2) | 7/1/2017 | NCBI |
| CY222573 | A/Alaska/02/2017(H3N2) | 11/1/2017 | NCBI |
| CY222581 | A/NewMexico/02/2017(H3N2) | 10/1/2017 | NCBI |
| CY222589 | A/Pennsylvania/118/2016(H3N2) | 27/12/2016 | NCBI |
| CY222597 | A/California/02/2017(H3N2) | 3/1/2017 | NCBI |
| CY222605 | A/SouthDakota/36/2016(H3N2) | 21/12/2016 | NCBI |
| CY222613 | A/NewJersey/05/2017(H3N2) | 15/1/2017 | NCBI |
| CY222621 | A/Maine/07/2017(H3N2) | 23/1/2017 | NCBI |
| CY222629 | A/Wyoming/03/2017(H3N2) | 12/1/2017 | NCBI |
| CY222637 | A/Wyoming/04/2017(H3N2) | 16/1/2017 | NCBI |
| CY222645 | A/Vermont/02/2017(H3N2) | 17/1/2017 | NCBI |
| CY222653 | A/Arkansas/04/2017(H3N2) | 17/1/2017 | NCBI |
| CY222661 | A/Indiana/05/2017(H3N2) | 11/1/2017 | NCBI |
| CY222669 | A/Virginia/08/2017(H3N2) | 22/1/2017 | NCBI |
| CY222677 | A/WestVirginia/15/2016(H3N2) | 27/12/2016 | NCBI |
| CY222685 | A/Montana/05/2017(H3N2) | 23/1/2017 | NCBI |
| CY222693 | A/Idaho/06/2017(H3N2) | 19/1/2017 | NCBI |
| CY222701 | A/NewYork/03/2017(H3N2) | 24/1/2017 | NCBI |
| CY222717 | A/California/11/2017(H3N2) | 20/1/2017 | NCBI |
| CY222725 | A/Mississippi/03/2017(H3N2) | 25/1/2017 | NCBI |
| CY222733 | A/Nevada/05/2017(H3N2) | 17/1/2017 | NCBI |
| CY222741 | A/Iowa/05/2017(H3N2) | 23/1/2017 | NCBI |
| CY222749 | A/Florida/07/2017(H3N2) | 16/1/2017 | NCBI |
| CY222757 | A/Kentucky/12/2017(H3N2) | 27/1/2017 | NCBI |
| CY222765 | A/NorthCarolina/05/2017(H3N2) | 30/1/2017 | NCBI |
| CY222773 | A/Louisiana/05/2017(H3N2) | 30/1/2017 | NCBI |
| CY222781 | A/Maryland/10/2017(H3N2) | 31/1/2017 | NCBI |
| CY222789 | A/Maryland/07/2017(H3N2) | 2017 | NCBI |
| CY222797 | A/NewYork/06/2017(H3N2) | 5/2/2017 | NCBI |
| CY222805 | A/NewYork/07/2017(H3N2) | 7/2/2017 | NCBI |
| CY222813 | A/Texas/40/2017(H3N2) | 13/2/2017 | NCBI |
| CY222821 | A/Massachusetts/04/2017(H3N2) | 1/2/2017 | NCBI |

| | | | |
|---|---|---|---|
| CY222829 | A/Massachusetts/05/2017(H3N2) | 1/2/2017 | NCBI |
| CY222837 | A/Pennsylvania/06/2017(H3N2) | 8/2/2017 | NCBI |
| CY222845 | A/Pennsylvania/07/2017(H3N2) | 8/2/2017 | NCBI |
| CY222853 | A/Arkansas/11/2017(H3N2) | 30/1/2017 | NCBI |
| CY222861 | A/Arkansas/12/2017(H3N2) | 31/1/2017 | NCBI |
| CY222869 | A/Arkansas/13/2017(H3N2) | 1/2/2017 | NCBI |
| CY222877 | A/Texas/54/2017(H3N2) | 16/2/2017 | NCBI |
| CY222885 | A/Texas/55/2017(H3N2) | 16/2/2017 | NCBI |
| CY222893 | A/Minnesota/13/2017(H3N2) | 17/2/2017 | NCBI |
| CY222901 | A/Minnesota/12/2017(H3N2) | 16/2/2017 | NCBI |
| CY222909 | A/Mississippi/06/2017(H3N2) | 16/2/2017 | NCBI |
| CY222917 | A/Mississippi/08/2017(H3N2) | 23/2/2017 | NCBI |
| CY222925 | A/Missouri/09/2017(H3N2) | 21/2/2017 | NCBI |
| CY222933 | A/Missouri/10/2017(H3N2) | 22/2/2017 | NCBI |
| CY222941 | A/Hawaii/05/2017(H3N2) | 23/1/2017 | NCBI |
| CY222949 | A/Hawaii/07/2017(H3N2) | 27/1/2017 | NCBI |
| CY222957 | A/Montana/09/2017(H3N2) | 11/2/2017 | NCBI |
| CY222965 | A/Montana/72/2016(H3N2) | 7/2/2016 | NCBI |
| CY222973 | A/Colorado/07/2017(H3N2) | 31/1/2017 | NCBI |
| CY222981 | A/Colorado/08/2017(H3N2) | 3/2/2017 | NCBI |
| CY222989 | A/Oregon/08/2017(H3N2) | 14/2/2017 | NCBI |
| CY222997 | A/Oregon/07/2017(H3N2) | 6/2/2017 | NCBI |
| CY223005 | A/Michigan/24/2017(H3N2) | 14/2/2017 | NCBI |
| CY223013 | A/Kansas/06/2017(H3N2) | 10/2/2017 | NCBI |
| CY223021 | A/Kansas/07/2017(H3N2) | 13/2/2017 | NCBI |
| CY223029 | A/Washington/24/2017(H3N2) | 30/1/2017 | NCBI |
| CY223037 | A/Washington/25/2017(H3N2) | 21/2/2017 | NCBI |
| CY223045 | A/Alabama/08/2017(H3N2) | 20/2/2017 | NCBI |
| CY223053 | A/Alabama/09/2017(H3N2) | 24/2/2017 | NCBI |
| CY223061 | A/NorthDakota/09/2017(H3N2) | 27/2/2017 | NCBI |
| CY223069 | A/NorthDakota/11/2017(H3N2) | 28/2/2017 | NCBI |
| CY223077 | A/Texas/60/2017(H3N2) | 27/2/2017 | NCBI |
| CY223085 | A/Texas/61/2017(H3N2) | 27/2/2017 | NCBI |
| CY223093 | A/Illinois/15/2017(H3N2) | 7/2/2017 | NCBI |
| CY223101 | A/Illinois/17/2017(H3N2) | 20/2/2017 | NCBI |
| CY223109 | A/Illinois/16/2017(H3N2) | 19/2/2017 | NCBI |
| CY223117 | A/Illinois/18/2017(H3N2) | 21/2/2017 | NCBI |

| | | | |
|---|---|---|---|
| CY223125 | A/Illinois/19/2017(H3N2) | 22/2/2017 | NCBI |
| CY223133 | A/Illinois/21/2017(H3N2) | 26/2/2017 | NCBI |
| CY223141 | A/Illinois/20/2017(H3N2) | 24/2/2017 | NCBI |
| CY223149 | A/Indiana/15/2017(H3N2) | 17/2/2017 | NCBI |
| CY223157 | A/Indiana/17/2017(H3N2) | 20/2/2017 | NCBI |
| CY223165 | A/Alaska/08/2017(H3N2) | 13/2/2017 | NCBI |
| CY223173 | A/Alaska/09/2017(H3N2) | 21/2/2017 | NCBI |
| CY223181 | A/Wyoming/11/2017(H3N2) | 2/3/2017 | NCBI |
| CY223189 | A/Wyoming/10/2017(H3N2) | 28/2/2017 | NCBI |
| CY223197 | A/Arizona/09/2017(H3N2) | 17/2/2017 | NCBI |
| CY223205 | A/Arizona/11/2017(H3N2) | 19/2/2017 | NCBI |
| CY223213 | A/NewMexico/09/2017(H3N2) | 22/2/2017 | NCBI |
| CY223221 | A/NewMexico/11/2017(H3N2) | 28/2/2017 | NCBI |
| CY223229 | A/Utah/13/2017(H3N2) | 27/2/2017 | NCBI |
| CY223237 | A/Utah/12/2017(H3N2) | 25/2/2017 | NCBI |
| CY223253 | A/Oklahoma/07/2017(H3N2) | 6/2/2017 | NCBI |
| CY223261 | A/Oklahoma/11/2017(H3N2) | 22/2/2017 | NCBI |
| CY223269 | A/Iowa/11/2017(H3N2) | 20/2/2017 | NCBI |
| CY223277 | A/Iowa/12/2017(H3N2) | 22/2/2017 | NCBI |
| CY223285 | A/Louisiana/10/2017(H3N2) | 25/1/2017 | NCBI |
| CY223293 | A/Texas/62/2017(H3N2) | 1/3/2017 | NCBI |
| CY223301 | A/Wisconsin/30/2017(H3N2) | 24/2/2017 | NCBI |
| CY223309 | A/Wisconsin/31/2017(H3N2) | 1/3/2017 | NCBI |
| CY223317 | A/Nebraska/06/2017(H3N2) | 27/2/2017 | NCBI |
| CY223677 | A/California/23/2017(H3N2) | 9/1/2017 | NCBI |
| CY221950 | A/Virginia/69/2016(H3N2) | 26/12/2016 | NCBI |
| CY223686 | A/Michigan/20/2017(H3N2) | 15/1/2017 | NCBI |
| CY223694 | A/Maryland/10/2017(H3N2) | 31/1/2017 | NCBI |
| CY223702 | A/Maryland/07/2017(H3N2) | 2017 | NCBI |
| CY223710 | A/Ohio/05/2017(H3N2) | 7/2/2017 | NCBI |
| CY223718 | A/NewYork/06/2017(H3N2) | 5/2/2017 | NCBI |
| CY223726 | A/NewYork/07/2017(H3N2) | 7/2/2017 | NCBI |
| CY223734 | A/Florida/10/2017(H3N2) | 8/2/2017 | NCBI |
| CY223742 | A/Florida/11/2017(H3N2) | 13/2/2017 | NCBI |
| CY223750 | A/Massachusetts/04/2017(H3N2) | 1/2/2017 | NCBI |
| CY223758 | A/Massachusetts/05/2017(H3N2) | 1/2/2017 | NCBI |
| CY223766 | A/Pennsylvania/06/2017(H3N2) | 8/2/2017 | NCBI |

| | | | |
|---|---|---|---|
| CY223774 | A/Pennsylvania/05/2017(H3N2) | 7/2/2017 | NCBI |
| CY223782 | A/Pennsylvania/07/2017(H3N2) | 8/2/2017 | NCBI |
| CY223790 | A/Tennessee/16/2017(H3N2) | 12/2/2017 | NCBI |
| CY223798 | A/Tennessee/15/2017(H3N2) | 9/2/2017 | NCBI |
| CY223806 | A/Connecticut/09/2017(H3N2) | 31/1/2017 | NCBI |
| CY223814 | A/RhodeIsland/03/2017(H3N2) | 24/1/2017 | NCBI |
| CY223830 | A/RhodeIsland/04/2017(H3N2) | 30/1/2017 | NCBI |
| CY223838 | A/RhodeIsland/07/2017(H3N2) | 13/2/2017 | NCBI |
| CY223846 | A/RhodeIsland/06/2017(H3N2) | 7/2/2017 | NCBI |
| CY223854 | A/Maine/10/2017(H3N2) | 14/2/2017 | NCBI |
| CY223870 | A/Maine/11/2017(H3N2) | 15/2/2017 | NCBI |
| CY223878 | A/Delaware/12/2017(H3N2) | 13/2/2017 | NCBI |
| CY223886 | A/Delaware/13/2017(H3N2) | 13/2/2017 | NCBI |
| CY223894 | A/NewHampshire/08/2017(H3N2) | 8/2/2017 | NCBI |
| CY223902 | A/NewHampshire/09/2017(H3N2) | 14/2/2017 | NCBI |
| CY223910 | A/Florida/13/2017(H3N2) | 11/2/2017 | NCBI |
| CY223918 | A/Florida/12/2017(H3N2) | 10/2/2017 | NCBI |
| CY223926 | A/Georgia/05/2017(H3N2) | 15/2/2017 | NCBI |
| CY223934 | A/Georgia/06/2017(H3N2) | 15/2/2017 | NCBI |
| CY223942 | A/Kentucky/13/2017(H3N2) | 2/2/2017 | NCBI |
| CY223950 | A/Kentucky/15/2017(H3N2) | 9/2/2017 | NCBI |
| CY223958 | A/NewJersey/13/2017(H3N2) | 14/2/2017 | NCBI |
| CY223966 | A/NewJersey/14/2017(H3N2) | 14/2/2017 | NCBI |
| CY223974 | A/NewJersey/15/2017(H3N2) | 14/2/2017 | NCBI |
| CY223982 | A/Vermont/06/2017(H3N2) | 8/2/2017 | NCBI |
| CY223990 | A/Virginia/11/2017(H3N2) | 14/2/2017 | NCBI |
| CY223998 | A/Virginia/09/2017(H3N2) | 13/2/2017 | NCBI |
| CY224006 | A/Virginia/10/2017(H3N2) | 13/2/2017 | NCBI |
| CY224014 | A/NorthCarolina/09/2017(H3N2) | 3/2/2017 | NCBI |
| CY224022 | A/Maryland/14/2017(H3N2) | 16/2/2017 | NCBI |
| CY224030 | A/WestVirginia/06/2017(H3N2) | 1/2/2017 | NCBI |
| CY224038 | A/WestVirginia/07/2017(H3N2) | 4/2/2017 | NCBI |
| CY224046 | A/NewYork/11/2017(H3N2) | 11/2/2017 | NCBI |
| CY224054 | A/Pennsylvania/15/2017(H3N2) | 15/2/2017 | NCBI |
| CY224062 | A/Florida/15/2017(H3N2) | 14/2/2017 | NCBI |
| CY224070 | A/Louisiana/13/2017(H3N2) | 22/2/2017 | NCBI |
| CY224078 | A/Hawaii/09/2017(H3N2) | 19/2/2017 | NCBI |

| | | | |
|---|---|---|---|
| CY224086 | A/Hawaii/11/2017(H3N2) | 26/2/2017 | NCBI |
| CY224094 | A/Oregon/10/2017(H3N2) | 23/2/2017 | NCBI |
| CY224102 | A/Oregon/09/2017(H3N2) | 24/1/2017 | NCBI |
| CY224110 | A/Idaho/12/2017(H3N2) | 27/2/2017 | NCBI |
| CY224118 | A/Idaho/14/2017(H3N2) | 6/3/2017 | NCBI |
| CY224126 | A/Montana/10/2017(H3N2) | 26/2/2017 | NCBI |
| CY224134 | A/Montana/11/2017(H3N2) | 8/3/2017 | NCBI |
| CY224142 | A/California/38/2017(H3N2) | 8/3/2017 | NCBI |
| CY224150 | A/California/35/2017(H3N2) | 17/2/2017 | NCBI |
| CY224158 | A/California/36/2017(H3N2) | 18/2/2017 | NCBI |
| KY697310 | A/Gainesville/09/2016(H3N2) | 6/9/2016 | NCBI |
| KY697318 | A/Gainesville/10/2016(H3N2) | 6/9/2016 | NCBI |
| KY697326 | A/Gainesville/13/2016(H3N2) | 20/12/2016 | NCBI |
| KY697334 | A/Gainesville/14/2016(H3N2) | 20/12/2016 | NCBI |
| KY697350 | A/Gainesville/02/2017(H3N2) | 18/1/2017 | NCBI |
| KY697358 | A/Gainesville/10/2017(H3N2) | 26/1/2017 | NCBI |
| KY697366 | A/Gainesville/11/2017(H3N2) | 18/2/2017 | NCBI |
| KY697374 | A/Gainesville/12/2017(H3N2) | 18/2/2017 | NCBI |
| CY226454 | A/Washington/96/2016(H3N2) | 13/12/2016 | NCBI |
| CY218023 | A/Michigan/121/2016(H3N2) | 27/12/2016 | NCBI |
| CY226471 | A/Wisconsin/04/2017(H3N2) | 9/1/2017 | NCBI |
| CY226479 | A/Wisconsin/05/2017(H3N2) | 11/1/2017 | NCBI |
| CY226487 | A/Pennsylvania/108/2016(H3N2) | 12/12/2016 | NCBI |
| CY226495 | A/Pennsylvania/110/2016(H3N2) | 12/12/2016 | NCBI |
| CY226503 | A/Pennsylvania/112/2016(H3N2) | 15/12/2016 | NCBI |
| CY226511 | A/Pennsylvania/115/2016(H3N2) | 15/12/2016 | NCBI |
| CY226519 | A/Pennsylvania/117/2016(H3N2) | 28/12/2016 | NCBI |
| CY226527 | A/Wisconsin/10/2017(H3N2) | 18/1/2017 | NCBI |
| CY226535 | A/Wisconsin/11/2017(H3N2) | 24/1/2017 | NCBI |
| CY226543 | A/Washington/07/2017(H3N2) | 20/1/2017 | NCBI |
| CY226551 | A/Washington/09/2017(H3N2) | 24/1/2017 | NCBI |
| CY226559 | A/Tennessee/38/2016(H3N2) | 15/12/2016 | NCBI |
| CY226567 | A/Tennessee/39/2016(H3N2) | 16/12/2016 | NCBI |
| CY226575 | A/Tennessee/40/2016(H3N2) | 27/12/2016 | NCBI |
| CY226583 | A/Tennessee/09/2017(H3N2) | 10/1/2017 | NCBI |
| CY226591 | A/Tennessee/10/2017(H3N2) | 13/1/2017 | NCBI |
| CY226607 | A/Tennessee/12/2017(H3N2) | 24/1/2017 | NCBI |

| | | | |
|---|---|---|---|
| CY226615 | A/Tennessee/41/2016(H3N2) | 27/12/2016 | NCBI |
| CY226623 | A/Tennessee/07/2017(H3N2) | 9/1/2017 | NCBI |
| CY226647 | A/Tennessee/06/2017(H3N2) | 4/1/2017 | NCBI |
| CY226655 | A/Tennessee/13/2017(H3N2) | 26/1/2017 | NCBI |
| CY226671 | A/Michigan/16/2017(H3N2) | 27/1/2017 | NCBI |
| CY226695 | A/Michigan/18/2017(H3N2) | 7/2/2017 | NCBI |
| CY226703 | A/Michigan/14/2017(H3N2) | 3/1/2017 | NCBI |
| CY226711 | A/Michigan/17/2017(H3N2) | 2/2/2017 | NCBI |
| CY224830 | A/California/23/2017(H3N2) | 9/1/2017 | NCBI |
| CY226719 | A/Michigan/07/2017(H3N2) | 10/1/2017 | NCBI |
| CY226727 | A/Michigan/10/2017(H3N2) | 16/1/2017 | NCBI |
| CY226735 | A/Michigan/12/2017(H3N2) | 23/1/2017 | NCBI |
| CY226743 | A/Michigan/08/2017(H3N2) | 11/1/2017 | NCBI |
| CY226751 | A/Michigan/09/2017(H3N2) | 11/1/2017 | NCBI |
| CY226759 | A/Michigan/11/2017(H3N2) | 19/1/2017 | NCBI |
| CY226767 | A/Michigan/13/2017(H3N2) | 23/1/2017 | NCBI |
| CY224838 | A/Delaware/15/2017(H3N2) | 24/2/2017 | NCBI |
| CY226775 | A/Texas/204/2016(H3N2) | 19/12/2016 | NCBI |
| CY226783 | A/Texas/211/2016(H3N2) | 22/12/2016 | NCBI |
| CY226791 | A/Texas/209/2016(H3N2) | 21/12/2016 | NCBI |
| CY226799 | A/Texas/214/2016(H3N2) | 29/12/2016 | NCBI |
| CY226807 | A/Texas/25/2017(H3N2) | 3/1/2017 | NCBI |
| CY226815 | A/Texas/26/2017(H3N2) | 3/1/2017 | NCBI |
| CY226823 | A/Texas/27/2017(H3N2) | 8/1/2017 | NCBI |
| CY226831 | A/Texas/28/2017(H3N2) | 14/1/2017 | NCBI |
| CY226839 | A/Texas/29/2017(H3N2) | 17/1/2017 | NCBI |
| CY226847 | A/Washington/13/2017(H3N2) | 7/2/2017 | NCBI |
| CY226863 | A/Washington/14/2017(H3N2) | 9/2/2017 | NCBI |
| CY226895 | A/Wisconsin/17/2017(H3N2) | 13/2/2017 | NCBI |
| CY226926 | A/Texas/205/2016(H3N2) | 19/12/2016 | NCBI |
| CY226934 | A/Texas/206/2016(H3N2) | 19/12/2016 | NCBI |
| CY226942 | A/Texas/207/2016(H3N2) | 19/12/2016 | NCBI |
| CY226958 | A/Texas/208/2016(H3N2) | 19/12/2016 | NCBI |
| CY226966 | A/Texas/213/2016(H3N2) | 22/12/2016 | NCBI |
| CY226990 | A/Texas/215/2016(H3N2) | 29/12/2016 | NCBI |
| CY226998 | A/Wisconsin/26/2017(H3N2) | 22/2/2017 | NCBI |
| CY227006 | A/Wisconsin/27/2017(H3N2) | 22/2/2017 | NCBI |

| | | | |
|---|---|---|---|
| CY227030 | A/Washington/18/2017(H3N2) | 15/2/2017 | NCBI |
| CY227038 | A/Washington/19/2017(H3N2) | 16/2/2017 | NCBI |
| CY224846 | A/SouthCarolina/06/2017(H3N2) | 23/2/2017 | NCBI |
| CY224854 | A/Washington/05/2017(H3N2) | 4/1/2017 | NCBI |
| CY224862 | A/Nevada/58/2016(H3N2) | 19/12/2016 | NCBI |
| CY224870 | A/Montana/07/2017(H3N2) | 29/1/2017 | NCBI |
| CY224878 | A/Michigan/21/2017(H3N2) | 7/2/2017 | NCBI |
| CY224886 | A/Arizona/06/2017(H3N2) | 31/1/2017 | NCBI |
| CY224894 | A/Florida/10/2017(H3N2) | 8/2/2017 | NCBI |
| CY224902 | A/Florida/11/2017(H3N2) | 13/2/2017 | NCBI |
| CY224910 | A/Pennsylvania/05/2017(H3N2) | 7/2/2017 | NCBI |
| CY224918 | A/Tennessee/16/2017(H3N2) | 12/2/2017 | NCBI |
| CY224926 | A/Tennessee/15/2017(H3N2) | 9/2/2017 | NCBI |
| CY224942 | A/Connecticut/08/2017(H3N2) | 31/1/2017 | NCBI |
| CY224950 | A/Connecticut/09/2017(H3N2) | 31/1/2017 | NCBI |
| CY224958 | A/RhodeIsland/03/2017(H3N2) | 24/1/2017 | NCBI |
| CY224966 | A/RhodeIsland/07/2017(H3N2) | 13/2/2017 | NCBI |
| CY224974 | A/RhodeIsland/06/2017(H3N2) | 7/2/2017 | NCBI |
| CY224982 | A/Maine/10/2017(H3N2) | 14/2/2017 | NCBI |
| CY224990 | A/Delaware/12/2017(H3N2) | 13/2/2017 | NCBI |
| CY224998 | A/Delaware/13/2017(H3N2) | 13/2/2017 | NCBI |
| CY225014 | A/Delaware/10/2017(H3N2) | 9/2/2017 | NCBI |
| CY225030 | A/Delaware/11/2017(H3N2) | 9/2/2017 | NCBI |
| CY225038 | A/NewHampshire/08/2017(H3N2) | 8/2/2017 | NCBI |
| CY225046 | A/NewHampshire/09/2017(H3N2) | 14/2/2017 | NCBI |
| CY225054 | A/Florida/13/2017(H3N2) | 11/2/2017 | NCBI |
| CY225062 | A/Florida/12/2017(H3N2) | 10/2/2017 | NCBI |
| CY225070 | A/Georgia/05/2017(H3N2) | 15/2/2017 | NCBI |
| CY225078 | A/Georgia/06/2017(H3N2) | 15/2/2017 | NCBI |
| CY225086 | A/Kentucky/13/2017(H3N2) | 2/2/2017 | NCBI |
| CY225094 | A/Kentucky/15/2017(H3N2) | 9/2/2017 | NCBI |
| CY225102 | A/Nevada/06/2017(H3N2) | 6/2/2017 | NCBI |
| CY225110 | A/NewJersey/13/2017(H3N2) | 14/2/2017 | NCBI |
| CY225118 | A/NewJersey/14/2017(H3N2) | 14/2/2017 | NCBI |
| CY225126 | A/NewJersey/15/2017(H3N2) | 14/2/2017 | NCBI |
| CY225134 | A/Virginia/11/2017(H3N2) | 14/2/2017 | NCBI |
| CY225142 | A/Virginia/09/2017(H3N2) | 13/2/2017 | NCBI |

| | | | |
|---|---|---|---|
| CY225150 | A/Virginia/10/2017(H3N2) | 13/2/2017 | NCBI |
| CY225158 | A/Texas/48/2017(H3N2) | 13/2/2017 | NCBI |
| CY225166 | A/NorthCarolina/09/2017(H3N2) | 3/2/2017 | NCBI |
| CY225174 | A/Arkansas/13/2017(H3N2) | 1/2/2017 | NCBI |
| CY225182 | A/Maryland/14/2017(H3N2) | 16/2/2017 | NCBI |
| CY225190 | A/Missouri/09/2017(H3N2) | 21/2/2017 | NCBI |
| CY225198 | A/WestVirginia/06/2017(H3N2) | 1/2/2017 | NCBI |
| CY225206 | A/WestVirginia/07/2017(H3N2) | 4/2/2017 | NCBI |
| CY225214 | A/NewYork/11/2017(H3N2) | 11/2/2017 | NCBI |
| CY225222 | A/Pennsylvania/15/2017(H3N2) | 15/2/2017 | NCBI |
| CY225230 | A/Illinois/22/2017(H3N2) | 6/3/2017 | NCBI |
| CY225238 | A/Illinois/23/2017(H3N2) | 6/3/2017 | NCBI |
| CY225246 | A/Minnesota/17/2017(H3N2) | 5/3/2017 | NCBI |
| CY225254 | A/Minnesota/16/2017(H3N2) | 4/3/2017 | NCBI |
| CY225262 | A/Texas/65/2017(H3N2) | 7/3/2017 | NCBI |
| CY225270 | A/SouthDakota/15/2017(H3N2) | 27/2/2017 | NCBI |
| CY225278 | A/SouthDakota/16/2017(H3N2) | 1/3/2017 | NCBI |
| CY225286 | A/SouthDakota/09/2017(H3N2) | 8/2/2017 | NCBI |
| CY225294 | A/SouthDakota/07/2017(H3N2) | 2/2/2017 | NCBI |
| CY225302 | A/SouthDakota/11/2017(H3N2) | 13/2/2017 | NCBI |
| CY225310 | A/SouthDakota/12/2017(H3N2) | 13/2/2017 | NCBI |
| CY225318 | A/Missouri/17/2017(H3N2) | 22/2/2017 | NCBI |
| CY225326 | A/Missouri/18/2017(H3N2) | 23/2/2017 | NCBI |
| CY225334 | A/Washington/30/2017(H3N2) | 3/3/2017 | NCBI |
| CY225342 | A/Washington/29/2017(H3N2) | 27/2/2017 | NCBI |
| CY225350 | A/Alaska/10/2017(H3N2) | 7/3/2017 | NCBI |
| CY225358 | A/Wyoming/12/2017(H3N2) | 6/3/2017 | NCBI |
| CY225366 | A/Wyoming/13/2017(H3N2) | 9/3/2017 | NCBI |
| CY225374 | A/Colorado/09/2017(H3N2) | 3/3/2017 | NCBI |
| CY225382 | A/Colorado/10/2017(H3N2) | 16/3/2017 | NCBI |
| CY225390 | A/California/41/2017(H3N2) | 7/2/2017 | NCBI |
| CY225398 | A/Utah/18/2017(H3N2) | 16/3/2017 | NCBI |
| CY225406 | A/Utah/17/2017(H3N2) | 12/3/2017 | NCBI |
| CY225414 | A/NewMexico/14/2017(H3N2) | 9/3/2017 | NCBI |
| CY225422 | A/NewMexico/15/2017(H3N2) | 13/3/2017 | NCBI |
| CY227062 | A/Alaska/29/2016(H3N2) | 31/10/2016 | NCBI |
| CY227070 | A/California/163/2016(H3N2) | 19/11/2016 | NCBI |

| | | | |
|---|---|---|---|
| CY227078 | A/Minnesota/83/2016(H3N2) | 11/12/2016 | NCBI |
| CY227086 | A/Nevada/54/2016(H3N2) | 8/12/2016 | NCBI |
| CY227094 | A/Utah/50/2016(H3N2) | 6/12/2016 | NCBI |
| CY227102 | A/Missouri/20/2016(H3N2) | 11/12/2016 | NCBI |
| CY227110 | A/NorthCarolina/12/2017(H3N2) | 24/1/2017 | NCBI |
| CY227118 | A/Maryland/11/2017(H3N2) | 28/1/2017 | NCBI |
| CY227126 | A/Maryland/12/2017(H3N2) | 3/2/2017 | NCBI |
| CY227134 | A/Maryland/15/2017(H3N2) | 16/2/2017 | NCBI |
| CY227142 | A/Texas/54/2017(H3N2) | 16/2/2017 | NCBI |
| CY227150 | A/WestVirginia/03/2017(H3N2) | 23/1/2017 | NCBI |
| CY227158 | A/WestVirginia/04/2017(H3N2) | 26/1/2017 | NCBI |
| CY227166 | A/Massachusetts/08/2017(H3N2) | 24/2/2017 | NCBI |
| CY227174 | A/Massachusetts/07/2017(H3N2) | 20/2/2017 | NCBI |
| CY227182 | A/NewYork/10/2017(H3N2) | 9/2/2017 | NCBI |
| CY227190 | A/Pennsylvania/16/2017(H3N2) | 20/2/2017 | NCBI |
| CY227198 | A/Pennsylvania/17/2017(H3N2) | 20/2/2017 | NCBI |
| CY227206 | A/Florida/14/2017(H3N2) | 14/2/2017 | NCBI |
| CY227214 | A/Connecticut/10/2017(H3N2) | 10/2/2017 | NCBI |
| CY227222 | A/Connecticut/11/2017(H3N2) | 10/2/2017 | NCBI |
| CY227230 | A/Delaware/16/2017(H3N2) | 28/2/2017 | NCBI |
| CY227238 | A/Delaware/17/2017(H3N2) | 1/3/2017 | NCBI |
| CY227246 | A/NewYork/14/2017(H3N2) | 28/2/2017 | NCBI |
| CY227254 | A/NewYork/13/2017(H3N2) | 23/2/2017 | NCBI |
| CY227262 | A/NewYork/12/2017(H3N2) | 21/2/2017 | NCBI |
| CY227278 | A/NewHampshire/12/2017(H3N2) | 22/2/2017 | NCBI |
| CY227286 | A/NewHampshire/13/2017(H3N2) | 23/2/2017 | NCBI |
| CY227294 | A/NewJersey/16/2017(H3N2) | 26/2/2017 | NCBI |
| CY227302 | A/NewJersey/17/2017(H3N2) | 27/2/2017 | NCBI |
| CY227318 | A/Kentucky/17/2017(H3N2) | 23/2/2017 | NCBI |
| CY227326 | A/Kentucky/18/2017(H3N2) | 1/3/2017 | NCBI |
| CY227334 | A/Texas/60/2017(H3N2) | 27/2/2017 | NCBI |
| CY227342 | A/Illinois/17/2017(H3N2) | 20/2/2017 | NCBI |
| CY227350 | A/Illinois/18/2017(H3N2) | 21/2/2017 | NCBI |
| CY227358 | A/Illinois/19/2017(H3N2) | 22/2/2017 | NCBI |
| CY227366 | A/Illinois/20/2017(H3N2) | 24/2/2017 | NCBI |
| CY227374 | A/Indiana/15/2017(H3N2) | 17/2/2017 | NCBI |
| CY227382 | A/Oklahoma/11/2017(H3N2) | 22/2/2017 | NCBI |

| | | | |
|---|---|---|---|
| CY227390 | A/Wisconsin/30/2017(H3N2) | 24/2/2017 | NCBI |
| CY227398 | A/Florida/17/2017(H3N2) | 19/2/2017 | NCBI |
| CY227406 | A/Florida/16/2017(H3N2) | 3/2/2017 | NCBI |
| CY227414 | A/NorthCarolina/16/2017(H3N2) | 23/2/2017 | NCBI |
| CY227422 | A/NorthCarolina/14/2017(H3N2) | 12/2/2017 | NCBI |
| CY227430 | A/Florida/18/2017(H3N2) | 4/3/2017 | NCBI |
| CY227438 | A/Florida/19/2017(H3N2) | 8/3/2017 | NCBI |
| CY227446 | A/Vermont/08/2017(H3N2) | 2/3/2017 | NCBI |
| CY227454 | A/Maine/12/2017(H3N2) | 8/3/2017 | NCBI |
| CY227462 | A/Maryland/21/2017(H3N2) | 24/2/2017 | NCBI |
| CY227470 | A/Maryland/23/2017(H3N2) | 2/3/2017 | NCBI |
| CY227478 | A/Delaware/20/2017(H3N2) | 13/3/2017 | NCBI |
| CY227486 | A/Delaware/21/2017(H3N2) | 13/3/2017 | NCBI |
| CY227494 | A/Texas/67/2017(H3N2) | 28/2/2017 | NCBI |
| CY227502 | A/Texas/68/2017(H3N2) | 1/3/2017 | NCBI |
| CY227510 | A/Mississippi/09/2017(H3N2) | 9/3/2017 | NCBI |
| CY227518 | A/Pennsylvania/26/2017(H3N2) | 6/3/2017 | NCBI |
| CY227526 | A/Pennsylvania/27/2017(H3N2) | 6/3/2017 | NCBI |
| CY227534 | A/Maine/14/2017(H3N2) | 13/3/2017 | NCBI |
| CY227542 | A/Maine/15/2017(H3N2) | 16/3/2017 | NCBI |
| CY227550 | A/NewJersey/19/2017(H3N2) | 5/3/2017 | NCBI |
| CY227558 | A/NewJersey/20/2017(H3N2) | 8/3/2017 | NCBI |
| CY227566 | A/NewJersey/21/2017(H3N2) | 10/3/2017 | NCBI |
| CY227574 | A/Connecticut/13/2017(H3N2) | 5/3/2017 | NCBI |
| CY227582 | A/Indiana/20/2017(H3N2) | 9/3/2017 | NCBI |
| CY227590 | A/Indiana/18/2017(H3N2) | 6/3/2017 | NCBI |
| CY227598 | A/Iowa/14/2017(H3N2) | 13/3/2017 | NCBI |
| CY227606 | A/Iowa/15/2017(H3N2) | 13/3/2017 | NCBI |
| CY227614 | A/NorthCarolina/18/2017(H3N2) | 14/2/2017 | NCBI |
| CY227622 | A/NorthCarolina/20/2017(H3N2) | 7/3/2017 | NCBI |
| CY227630 | A/Tennessee/28/2017(H3N2) | 14/3/2017 | NCBI |
| CY227638 | A/Tennessee/29/2017(H3N2) | 14/3/2017 | NCBI |
| CY227646 | A/NewHampshire/15/2017(H3N2) | 14/3/2017 | NCBI |
| CY227654 | A/NewHampshire/14/2017(H3N2) | 9/3/2017 | NCBI |
| CY227662 | A/Florida/20/2017(H3N2) | 2/3/2017 | NCBI |
| CY227670 | A/Florida/21/2017(H3N2) | 8/3/2017 | NCBI |
| CY227678 | A/Louisiana/19/2017(H3N2) | 8/3/2017 | NCBI |

| | | | |
|---|---|---|---|
| CY227686 | A/Louisiana/17/2017(H3N2) | 5/3/2017 | NCBI |
| CY227694 | A/Arkansas/14/2017(H3N2) | 22/2/2017 | NCBI |
| CY227702 | A/Arkansas/15/2017(H3N2) | 22/2/2017 | NCBI |
| CY227710 | A/Arkansas/16/2017(H3N2) | 27/2/2017 | NCBI |
| CY227718 | A/Wisconsin/37/2017(H3N2) | 13/3/2017 | NCBI |
| CY227726 | A/RhodeIsland/10/2017(H3N2) | 23/2/2017 | NCBI |
| CY227734 | A/Michigan/34/2017(H3N2) | 18/2/2017 | NCBI |
| CY227742 | A/Michigan/35/2017(H3N2) | 25/2/2017 | NCBI |
| CY227750 | A/Texas/71/2017(H3N2) | 18/3/2017 | NCBI |
| CY227758 | A/Texas/70/2017(H3N2) | 17/3/2017 | NCBI |
| CY227766 | A/Illinois/24/2017(H3N2) | 13/3/2017 | NCBI |
| CY227774 | A/Illinois/25/2017(H3N2) | 14/3/2017 | NCBI |
| CY227782 | A/Texas/73/2017(H3N2) | 21/3/2017 | NCBI |
| CY227790 | A/Texas/74/2017(H3N2) | 21/3/2017 | NCBI |
| CY227798 | A/NorthDakota/13/2017(H3N2) | 16/3/2017 | NCBI |
| CY227806 | A/NorthDakota/14/2017(H3N2) | 17/3/2017 | NCBI |
| CY227814 | A/Minnesota/20/2017(H3N2) | 15/3/2017 | NCBI |
| CY227822 | A/Minnesota/21/2017(H3N2) | 21/3/2017 | NCBI |
| CY227830 | A/Texas/72/2017(H3N2) | 8/3/2017 | NCBI |
| CY227838 | A/Texas/75/2017(H3N2) | 25/3/2017 | NCBI |
| CY227846 | A/Oklahoma/13/2017(H3N2) | 2/3/2017 | NCBI |
| CY227854 | A/Oklahoma/14/2017(H3N2) | 4/3/2017 | NCBI |
| KY935523 | A/environment/Indiana/16TOSU3553/2016(H3N2) | 1/6/2016 | NCBI |
| KY935491 | A/environment/Indiana/16TOSU3767/2016(H3N2) | 1/7/2016 | NCBI |
| KY935535 | A/environment/Indiana/16TOSU3854/2016(H3N2) | 1/7/2016 | NCBI |
| CY230005 | A/Michigan/33/2017(H3N2) | 13/3/2017 | NCBI |
| CY230021 | A/Michigan/26/2017(H3N2) | 18/2/2017 | NCBI |
| CY230045 | A/Michigan/30/2017(H3N2) | 27/2/2017 | NCBI |
| CY230069 | A/Michigan/31/2017(H3N2) | 7/3/2017 | NCBI |
| CY230085 | A/Michigan/25/2017(H3N2) | 13/2/2017 | NCBI |
| CY230093 | A/Michigan/27/2017(H3N2) | 24/2/2017 | NCBI |
| CY230101 | A/Michigan/28/2017(H3N2) | 26/2/2017 | NCBI |
| CY230117 | A/Michigan/32/2017(H3N2) | 9/3/2017 | NCBI |
| CY230133 | A/Michigan/116/2016(H3N2) | 12/12/2016 | NCBI |
| CY230141 | A/Washington/94/2016(H3N2) | 12/12/2016 | NCBI |
| CY230149 | A/Pennsylvania/110/2016(H3N2) | 12/12/2016 | NCBI |
| CY230157 | A/Pennsylvania/115/2016(H3N2) | 15/12/2016 | NCBI |

| | | | |
|---|---|---|---|
| CY230165 | A/Pennsylvania/116/2016(H3N2) | 21/12/2016 | NCBI |
| CY230173 | A/Tennessee/40/2016(H3N2) | 27/12/2016 | NCBI |
| CY230189 | A/Washington/27/2017(H3N2) | 1/3/2017 | NCBI |
| CY230197 | A/Washington/28/2017(H3N2) | 6/3/2017 | NCBI |
| CY230213 | A/Wisconsin/28/2017(H3N2) | 7/3/2017 | NCBI |
| CY230229 | A/Wisconsin/29/2017(H3N2) | 7/3/2017 | NCBI |
| CY230253 | A/Tennessee/18/2017(H3N2) | 7/2/2017 | NCBI |
| CY230277 | A/Tennessee/26/2017(H3N2) | 10/3/2017 | NCBI |
| CY230285 | A/Tennessee/21/2017(H3N2) | 20/2/2017 | NCBI |
| CY230293 | A/Tennessee/17/2017(H3N2) | 6/2/2017 | NCBI |
| CY230301 | A/Tennessee/23/2017(H3N2) | 27/2/2017 | NCBI |
| CY227862 | A/Texas/53/2017(H3N2) | 22/2/2017 | NCBI |
| CY230317 | A/Washington/31/2017(H3N2) | 10/3/2017 | NCBI |
| CY230325 | A/Washington/32/2017(H3N2) | 15/3/2017 | NCBI |
| CY230349 | A/Wisconsin/34/2017(H3N2) | 21/3/2017 | NCBI |
| CY230357 | A/Wisconsin/35/2017(H3N2) | 21/3/2017 | NCBI |
| CY230373 | A/Texas/26/2017(H3N2) | 3/1/2017 | NCBI |
| CY230381 | A/Wisconsin/16/2017(H3N2) | 13/2/2017 | NCBI |
| CY230390 | A/Washington/19/2017(H3N2) | 16/2/2017 | NCBI |
| CY227870 | A/Michigan/37/2017(H3N2) | 22/2/2017 | NCBI |
| CY227878 | A/Louisiana/15/2017(H3N2) | 16/3/2017 | NCBI |
| CY227894 | A/Delaware/18/2017(H3N2) | 11/3/2017 | NCBI |
| CY227902 | A/Delaware/19/2017(H3N2) | 21/3/2017 | NCBI |
| CY227910 | A/Montana/13/2017(H3N2) | 15/3/2017 | NCBI |
| CY227918 | A/Montana/12/2017(H3N2) | 5/3/2017 | NCBI |
| CY227926 | A/SouthCarolina/09/2017(H3N2) | 21/3/2017 | NCBI |
| CY227934 | A/SouthCarolina/11/2017(H3N2) | 23/3/2017 | NCBI |
| CY227942 | A/Maine/23/2016(H3N2) | 23/10/2016 | NCBI |
| CY227950 | A/Hawaii/90/2016(H3N2) | 21/10/2016 | NCBI |
| CY227958 | A/SouthDakota/29/2016(H3N2) | 17/10/2016 | NCBI |
| CY227966 | A/Washington/83/2016(H3N2) | 25/10/2016 | NCBI |
| CY227982 | A/Hawaii/95/2016(H3N2) | 7/11/2016 | NCBI |
| CY227990 | A/California/162/2016(H3N2) | 15/11/2016 | NCBI |
| CY227998 | A/Delaware/37/2016(H3N2) | 4/12/2016 | NCBI |
| CY228006 | A/Florida/102/2016(H3N2) | 1/12/2016 | NCBI |
| CY228014 | A/Nevada/55/2016(H3N2) | 8/12/2016 | NCBI |
| CY228022 | A/Illinois/52/2016(H3N2) | 13/12/2016 | NCBI |

| | | | |
|---|---|---|---|
| CY228030 | A/Nebraska/26/2016(H3N2) | 6/12/2016 | NCBI |
| CY228038 | A/Nebraska/27/2016(H3N2) | 13/12/2016 | NCBI |
| CY228046 | A/Alabama/18/2016(H3N2) | 14/12/2016 | NCBI |
| CY228054 | A/Arizona/80/2016(H3N2) | 12/12/2016 | NCBI |
| CY228062 | A/Washington/101/2016(H3N2) | 15/12/2016 | NCBI |
| CY228070 | A/Iowa/42/2016(H3N2) | 20/12/2016 | NCBI |
| CY228078 | A/Texas/197/2016(H3N2) | 25/12/2016 | NCBI |
| CY228086 | A/Alaska/35/2016(H3N2) | 13/12/2016 | NCBI |
| CY228094 | A/Utah/53/2016(H3N2) | 22/12/2016 | NCBI |
| CY228102 | A/Utah/52/2016(H3N2) | 20/12/2016 | NCBI |
| CY228117 | A/Maryland/30/2016(H3N2) | 12/12/2016 | NCBI |
| CY228125 | A/Texas/198/2016(H3N2) | 25/12/2016 | NCBI |
| CY228133 | A/Maine/02/2017(H3N2) | 6/1/2017 | NCBI |
| CY228141 | A/Maine/01/2017(H3N2) | 4/1/2017 | NCBI |
| CY228149 | A/Illinois/56/2016(H3N2) | 15/12/2016 | NCBI |
| CY228157 | A/Illinois/01/2017(H3N2) | 3/1/2017 | NCBI |
| CY228165 | A/California/199/2016(H3N2) | 25/12/2016 | NCBI |
| CY228173 | A/Texas/07/2017(H3N2) | 5/1/2017 | NCBI |
| CY228181 | A/Maryland/03/2017(H3N2) | 6/1/2017 | NCBI |
| CY228189 | A/California/214/2016(H3N2) | 31/12/2016 | NCBI |
| CY228197 | A/Virginia/03/2017(H3N2) | 2/1/2017 | NCBI |
| CY228205 | A/Massachusetts/03/2017(H3N2) | 27/1/2017 | NCBI |
| CY228213 | A/Iowa/03/2017(H3N2) | 12/1/2017 | NCBI |
| CY228221 | A/Wisconsin/20/2017(H3N2) | 23/1/2017 | NCBI |
| CY228229 | A/Michigan/22/2017(H3N2) | 8/2/2017 | NCBI |
| CY228237 | A/Texas/46/2017(H3N2) | 13/2/2017 | NCBI |
| CY228245 | A/Hawaii/07/2017(H3N2) | 27/1/2017 | NCBI |
| CY228253 | A/Montana/09/2017(H3N2) | 11/2/2017 | NCBI |
| CY228261 | A/Oregon/08/2017(H3N2) | 14/2/2017 | NCBI |
| CY228269 | A/Illinois/15/2017(H3N2) | 7/2/2017 | NCBI |
| CY228277 | A/NewMexico/11/2017(H3N2) | 28/2/2017 | NCBI |
| CY228285 | A/Illinois/22/2017(H3N2) | 6/3/2017 | NCBI |
| CY228293 | A/Illinois/23/2017(H3N2) | 6/3/2017 | NCBI |
| CY228301 | A/Hawaii/11/2017(H3N2) | 26/2/2017 | NCBI |
| CY228309 | A/SouthDakota/15/2017(H3N2) | 27/2/2017 | NCBI |
| CY228317 | A/SouthDakota/11/2017(H3N2) | 13/2/2017 | NCBI |
| CY228325 | A/Missouri/17/2017(H3N2) | 22/2/2017 | NCBI |

| | | | |
|---|---|---|---|
| CY228333 | A/Vermont/07/2017(H3N2) | 17/2/2017 | NCBI |
| CY228341 | A/Tennessee/28/2017(H3N2) | 14/3/2017 | NCBI |
| CY228349 | A/Louisiana/17/2017(H3N2) | 5/3/2017 | NCBI |
| CY228357 | A/Kentucky/23/2017(H3N2) | 16/3/2017 | NCBI |
| CY228365 | A/Kentucky/22/2017(H3N2) | 9/3/2017 | NCBI |
| CY228373 | A/Maryland/25/2017(H3N2) | 27/2/2017 | NCBI |
| CY228381 | A/Maryland/28/2017(H3N2) | 11/3/2017 | NCBI |
| CY228389 | A/WestVirginia/09/2017(H3N2) | 13/2/2017 | NCBI |
| CY228397 | A/WestVirginia/10/2017(H3N2) | 21/2/2017 | NCBI |
| CY228405 | A/WestVirginia/12/2017(H3N2) | 14/3/2017 | NCBI |
| CY228413 | A/Florida/22/2017(H3N2) | 20/3/2017 | NCBI |
| CY228429 | A/Florida/23/2017(H3N2) | 23/3/2017 | NCBI |
| CY228445 | A/NewYork/17/2017(H3N2) | 6/2/2017 | NCBI |
| CY228453 | A/NewYork/18/2017(H3N2) | 23/2/2017 | NCBI |
| CY228461 | A/Nevada/08/2017(H3N2) | 2/3/2017 | NCBI |
| CY228469 | A/Massachusetts/10/2017(H3N2) | 9/3/2017 | NCBI |
| CY228485 | A/Virginia/22/2017(H3N2) | 24/3/2017 | NCBI |
| CY228493 | A/Virginia/20/2017(H3N2) | 17/3/2017 | NCBI |
| CY228509 | A/Vermont/09/2017(H3N2) | 13/3/2017 | NCBI |
| CY228517 | A/Vermont/10/2017(H3N2) | 13/3/2017 | NCBI |
| CY228533 | A/Mississippi/12/2017(H3N2) | 20/3/2017 | NCBI |
| CY228541 | A/Mississippi/13/2017(H3N2) | 23/3/2017 | NCBI |
| CY228549 | A/Tennessee/31/2017(H3N2) | 28/3/2017 | NCBI |
| CY228565 | A/Delaware/23/2017(H3N2) | 29/3/2017 | NCBI |
| CY228581 | A/Delaware/22/2017(H3N2) | 23/3/2017 | NCBI |
| CY228589 | A/Washington/33/2017(H3N2) | 19/3/2017 | NCBI |
| CY228597 | A/Washington/34/2017(H3N2) | 19/3/2017 | NCBI |
| CY228605 | A/Colorado/11/2017(H3N2) | 21/3/2017 | NCBI |
| CY228613 | A/Illinois/28/2017(H3N2) | 23/3/2017 | NCBI |
| CY228621 | A/Illinois/27/2017(H3N2) | 19/3/2017 | NCBI |
| CY228629 | A/Florida/26/2017(H3N2) | 26/3/2017 | NCBI |
| CY228637 | A/Florida/25/2017(H3N2) | 22/3/2017 | NCBI |
| CY228645 | A/Connecticut/15/2017(H3N2) | 20/3/2017 | NCBI |
| CY228653 | A/Maine/16/2017(H3N2) | 29/3/2017 | NCBI |
| CY228661 | A/Maine/17/2017(H3N2) | 30/3/2017 | NCBI |
| CY228677 | A/Hawaii/12/2017(H3N2) | 28/2/2017 | NCBI |
| CY228685 | A/Hawaii/15/2017(H3N2) | 15/3/2017 | NCBI |

| | | | |
|---|---|---|---|
| CY228693 | A/Wisconsin/41/2017(H3N2) | 26/3/2017 | NCBI |
| CY228701 | A/Wisconsin/39/2017(H3N2) | 19/3/2017 | NCBI |
| CY228709 | A/Michigan/39/2017(H3N2) | 16/3/2017 | NCBI |
| CY228717 | A/Michigan/40/2017(H3N2) | 17/3/2017 | NCBI |
| CY228725 | A/Arkansas/17/2017(H3N2) | 19/3/2017 | NCBI |
| CY228733 | A/Arkansas/18/2017(H3N2) | 19/3/2017 | NCBI |
| CY228741 | A/Arkansas/19/2017(H3N2) | 27/3/2017 | NCBI |
| CY228749 | A/Iowa/17/2017(H3N2) | 21/3/2017 | NCBI |
| CY228757 | A/Oklahoma/19/2017(H3N2) | 13/3/2017 | NCBI |
| CY228765 | A/Oklahoma/20/2017(H3N2) | 14/3/2017 | NCBI |
| CY228773 | A/NewHampshire/16/2017(H3N2) | 25/3/2017 | NCBI |
| CY228781 | A/NewHampshire/17/2017(H3N2) | 28/3/2017 | NCBI |
| CY228797 | A/NewJersey/22/2017(H3N2) | 20/3/2017 | NCBI |
| CY228805 | A/NewJersey/23/2017(H3N2) | 21/3/2017 | NCBI |
| CY228813 | A/Alaska/11/2017(H3N2) | 21/3/2017 | NCBI |
| CY228821 | A/Wyoming/14/2017(H3N2) | 16/3/2017 | NCBI |
| CY228829 | A/Wyoming/15/2017(H3N2) | 18/3/2017 | NCBI |
| CY228837 | A/NewMexico/16/2017(H3N2) | 26/3/2017 | NCBI |
| CY228845 | A/NewMexico/17/2017(H3N2) | 26/3/2017 | NCBI |
| CY228853 | A/Utah/20/2017(H3N2) | 30/3/2017 | NCBI |
| CY228861 | A/Utah/21/2017(H3N2) | 30/3/2017 | NCBI |
| CY228869 | A/California/50/2017(H3N2) | 1/2/2017 | NCBI |
| CY228877 | A/California/52/2017(H3N2) | 2/2/2017 | NCBI |
| CY228885 | A/Idaho/19/2017(H3N2) | 6/3/2017 | NCBI |
| CY228893 | A/Idaho/20/2017(H3N2) | 9/3/2017 | NCBI |
| CY228901 | A/Minnesota/23/2017(H3N2) | 28/3/2017 | NCBI |
| CY228909 | A/Minnesota/24/2017(H3N2) | 28/3/2017 | NCBI |
| CY228917 | A/Illinois/29/2017(H3N2) | 2/4/2017 | NCBI |
| CY228925 | A/Michigan/50/2017(H3N2) | 23/3/2017 | NCBI |
| CY228933 | A/Michigan/51/2017(H3N2) | 30/3/2017 | NCBI |
| KY949688 | A/Baltimore/0221/2017(H3N2) | 3/1/2017 | NCBI |
| KY950007 | A/Baltimore/0223/2017(H3N2) | 5/1/2017 | NCBI |
| KY949992 | A/Baltimore/0227/2017(H3N2) | 7/1/2017 | NCBI |
| KY950141 | A/Baltimore/0230/2017(H3N2) | 9/1/2017 | NCBI |
| KY950201 | A/Baltimore/0233/2017(H3N2) | 10/1/2017 | NCBI |
| KY949842 | A/Baltimore/0237/2017(H3N2) | 18/1/2017 | NCBI |
| KY949986 | A/Baltimore/0238/2017(H3N2) | 20/1/2017 | NCBI |

| | | | |
|---|---|---|---|
| KY949711 | A/Baltimore/0239/2017(H3N2) | 20/1/2017 | NCBI |
| KY949788 | A/Baltimore/0244/2017(H3N2) | 24/1/2017 | NCBI |
| KY949989 | A/Baltimore/0247/2017(H3N2) | 25/1/2017 | NCBI |
| KY949928 | A/Baltimore/0252/2017(H3N2) | 27/1/2017 | NCBI |
| KY950053 | A/Baltimore/0257/2017(H3N2) | 30/1/2017 | NCBI |
| KY949661 | A/Baltimore/0261/2017(H3N2) | 31/1/2017 | NCBI |
| KY949955 | A/Baltimore/0262/2017(H3N2) | 31/1/2017 | NCBI |
| KY950099 | A/Baltimore/0281/2017(H3N2) | 6/2/2017 | NCBI |
| KY949907 | A/Baltimore/0284/2017(H3N2) | 6/2/2017 | NCBI |
| KY950167 | A/Baltimore/0287/2017(H3N2) | 7/2/2017 | NCBI |
| KY950120 | A/Baltimore/0289/2017(H3N2) | 8/2/2017 | NCBI |
| KY949900 | A/Baltimore/0293/2017(H3N2) | 9/2/2017 | NCBI |
| KY949654 | A/Baltimore/0294/2017(H3N2) | 9/2/2017 | NCBI |
| KY949978 | A/Baltimore/0295/2017(H3N2) | 9/2/2017 | NCBI |
| KY949739 | A/Baltimore/0297/2017(H3N2) | 9/2/2017 | NCBI |
| KY950166 | A/Baltimore/0300/2017(H3N2) | 10/2/2017 | NCBI |
| KY949820 | A/Baltimore/0301/2017(H3N2) | 12/2/2017 | NCBI |
| KY949985 | A/Baltimore/0303/2017(H3N2) | 13/2/2017 | NCBI |
| KY950048 | A/Baltimore/0306/2017(H3N2) | 14/2/2017 | NCBI |
| KY950005 | A/Baltimore/0307/2017(H3N2) | 14/2/2017 | NCBI |
| KY949821 | A/Baltimore/0311/2017(H3N2) | 14/2/2017 | NCBI |
| KY950164 | A/Keelung/0081/2016(H3N2) | 29/11/2016 | NCBI |
| KY949869 | A/Keelung/0082/2016(H3N2) | 30/11/2016 | NCBI |
| KY949873 | A/Keelung/0089/2016(H3N2) | 8/12/2016 | NCBI |
| KY950075 | A/Keelung/0091/2016(H3N2) | 9/12/2016 | NCBI |
| KY949735 | A/Keelung/0095/2016(H3N2) | 14/12/2016 | NCBI |
| KY949885 | A/Keelung/0097/2016(H3N2) | 20/12/2016 | NCBI |
| KY949899 | A/Keelung/0099/2016(H3N2) | 21/12/2016 | NCBI |
| KY949642 | A/Linkou/0153/2016(H3N2) | 10/11/2016 | NCBI |
| KY949757 | A/Linkou/0164/2016(H3N2) | 25/11/2016 | NCBI |
| KY950107 | A/Linkou/0166/2016(H3N2) | 28/11/2016 | NCBI |
| KY949886 | A/Linkou/0168/2016(H3N2) | 1/12/2016 | NCBI |
| KY950205 | A/Linkou/0169/2016(H3N2) | 2/12/2016 | NCBI |
| KY949854 | A/Linkou/0173/2016(H3N2) | 8/12/2016 | NCBI |
| KY950074 | A/Linkou/0175/2016(H3N2) | 12/12/2016 | NCBI |
| KY949649 | A/Linkou/0181/2016(H3N2) | 21/12/2016 | NCBI |
| KY950159 | A/Linkou/0185/2016(H3N2) | 24/12/2016 | NCBI |

| | | | |
|---|---|---|---|
| KY949883 | A/Linkou/0186/2016(H3N2) | 26/12/2016 | NCBI |
| KY949839 | A/Linkou/0187/2016(H3N2) | 27/12/2016 | NCBI |
| KY950192 | A/Linkou/0191/2016(H3N2) | 30/12/2016 | NCBI |
| KY949789 | A/Taipei/0088/2016(H3N2) | 6/12/2016 | NCBI |
| KY949902 | A/Taipei/0092/2016(H3N2) | 19/12/2016 | NCBI |
| KY949971 | A/Taipei/0095/2016(H3N2) | 22/12/2016 | NCBI |
| CY230433 | A/Michigan/37/2017(H3N2) | 22/2/2017 | NCBI |
| CY230441 | A/Delaware/19/2017(H3N2) | 21/3/2017 | NCBI |
| CY230449 | A/Montana/13/2017(H3N2) | 15/3/2017 | NCBI |
| CY230457 | A/Montana/12/2017(H3N2) | 5/3/2017 | NCBI |
| CY228110 | A/California/192/2016(H3N2) | 14/12/2016 | NCBI |
| CY230980 | A/NewJersey/12/2017(H3N2) | 13/2/2017 | NCBI |
| CY230988 | A/NewJersey/12/2017(H3N2) | 13/2/2017 | NCBI |
| CY230466 | A/California/36/2017(H3N2) | 18/2/2017 | NCBI |
| CY230474 | A/Maine/15/2017(H3N2) | 16/3/2017 | NCBI |
| CY230482 | A/Wyoming/13/2017(H3N2) | 9/3/2017 | NCBI |
| CY230490 | A/NorthCarolina/20/2017(H3N2) | 7/3/2017 | NCBI |
| CY230498 | A/NewHampshire/15/2017(H3N2) | 14/3/2017 | NCBI |
| CY230506 | A/Louisiana/19/2017(H3N2) | 8/3/2017 | NCBI |
| CY230514 | A/Colorado/09/2017(H3N2) | 3/3/2017 | NCBI |
| CY230522 | A/NewMexico/15/2017(H3N2) | 13/3/2017 | NCBI |
| CY230530 | A/Arizona/13/2017(H3N2) | 22/2/2017 | NCBI |
| CY230538 | A/Arizona/14/2017(H3N2) | 25/2/2017 | NCBI |
| CY230546 | A/Maryland/28/2017(H3N2) | 11/3/2017 | NCBI |
| CY230554 | A/WestVirginia/09/2017(H3N2) | 13/2/2017 | NCBI |
| CY230562 | A/WestVirginia/12/2017(H3N2) | 14/3/2017 | NCBI |
| CY230570 | A/WestVirginia/11/2017(H3N2) | 6/3/2017 | NCBI |
| CY231125 | A/WestVirginia/11/2017(H3N2) | 6/3/2017 | NCBI |
| CY230578 | A/Texas/73/2017(H3N2) | 21/3/2017 | NCBI |
| CY230586 | A/NewYork/17/2017(H3N2) | 6/2/2017 | NCBI |
| CY230594 | A/Oregon/11/2017(H3N2) | 10/3/2017 | NCBI |
| CY230602 | A/Oregon/12/2017(H3N2) | 14/3/2017 | NCBI |
| CY230610 | A/Nevada/08/2017(H3N2) | 2/3/2017 | NCBI |
| CY230618 | A/Nevada/09/2017(H3N2) | 6/3/2017 | NCBI |
| CY230626 | A/Virginia/22/2017(H3N2) | 24/3/2017 | NCBI |
| CY230634 | A/Oklahoma/13/2017(H3N2) | 2/3/2017 | NCBI |
| CY230642 | A/Arizona/19/2017(H3N2) | 21/3/2017 | NCBI |

| | | | |
|---|---|---|---|
| CY230650 | A/Arizona/20/2017(H3N2) | 27/3/2017 | NCBI |
| CY230658 | A/Hawaii/16/2017(H3N2) | 9/3/2017 | NCBI |
| CY230666 | A/Hawaii/17/2017(H3N2) | 20/3/2017 | NCBI |
| CY230674 | A/Alabama/17/2017(H3N2) | 30/3/2017 | NCBI |
| CY230682 | A/Alabama/16/2017(H3N2) | 29/3/2017 | NCBI |
| CY230690 | A/Alabama/15/2017(H3N2) | 20/3/2017 | NCBI |
| CY230698 | A/Alabama/13/2017(H3N2) | 13/3/2017 | NCBI |
| CY230706 | A/Alabama/12/2017(H3N2) | 9/3/2017 | NCBI |
| CY230714 | A/Alabama/10/2017(H3N2) | 6/3/2017 | NCBI |
| CY230722 | A/Alabama/11/2017(H3N2) | 7/3/2017 | NCBI |
| CY230730 | A/NewMexico/18/2017(H3N2) | 4/4/2017 | NCBI |
| CY230738 | A/Alaska/12/2017(H3N2) | 30/3/2017 | NCBI |
| CY230746 | A/Alaska/14/2017(H3N2) | 4/4/2017 | NCBI |
| CY230754 | A/Louisiana/22/2017(H3N2) | 19/3/2017 | NCBI |
| CY230762 | A/Louisiana/23/2017(H3N2) | 20/3/2017 | NCBI |
| CY230770 | A/Louisiana/25/2017(H3N2) | 11/4/2017 | NCBI |
| CY230778 | A/Louisiana/26/2017(H3N2) | 11/4/2017 | NCBI |
| CY230786 | A/Iowa/19/2017(H3N2) | 3/4/2017 | NCBI |
| CY230794 | A/SouthDakota/19/2017(H3N2) | 29/3/2017 | NCBI |
| CY230802 | A/Wisconsin/46/2017(H3N2) | 2/4/2017 | NCBI |
| CY230810 | A/Wisconsin/47/2017(H3N2) | 3/4/2017 | NCBI |
| CY230818 | A/Utah/24/2017(H3N2) | 14/4/2017 | NCBI |
| CY230826 | A/Utah/23/2017(H3N2) | 13/4/2017 | NCBI |
| CY230834 | A/Idaho/21/2017(H3N2) | 20/3/2017 | NCBI |
| CY230842 | A/Idaho/23/2017(H3N2) | 31/3/2017 | NCBI |
| CY230850 | A/Oregon/13/2017(H3N2) | 25/2/2017 | NCBI |
| CY230858 | A/Oregon/14/2017(H3N2) | 28/2/2017 | NCBI |
| CY230866 | A/Wyoming/16/2017(H3N2) | 6/4/2017 | NCBI |
| CY230874 | A/Wyoming/17/2017(H3N2) | 11/4/2017 | NCBI |
| CY230882 | A/Indiana/21/2017(H3N2) | 5/4/2017 | NCBI |
| CY230890 | A/Indiana/22/2017(H3N2) | 10/4/2017 | NCBI |
| KY935475 | A/PassoFundo/LACENRS-1854/2015(H3N2) | 20/7/2015 | NCBI |
| KY935467 | A/PortoAlegre/LACENRS-1155/2015(H3N2) | 3/6/2015 | NCBI |
| KY925323 | A/Alegrete/LACENRS-1624/2014(H3N2) | 18/7/2014 | NCBI |
| KY925762 | A/Arvorezinha/LACENRS-2055/2013(H3N2) | 3/7/2013 | NCBI |
| KY925449 | A/Bage/LACENRS-1200/2014(H3N2) | 23/6/2014 | NCBI |
| KY926322 | A/Bage/LACENRS-2050/2013(H3N2) | 4/7/2013 | NCBI |

| | | | |
|---|---|---|---|
| KY925459 | A/Bage/LACENRS-294/2011(H3N2) | 16/6/2011 | NCBI |
| KY926167 | A/BomRetirodoSul/LACENRS-3406/2013(H3N2) | 22/8/2013 | NCBI |
| KY925783 | A/CambaradoSul/LACENRS-1939/2015(H3N2) | 29/7/2015 | NCBI |
| KY925136 | A/CambaradoSul/LACENRS-2049/2015(H3N2) | 10/8/2015 | NCBI |
| KY925954 | A/Canela/LACENRS-102/2011(H3N2) | 20/5/2011 | NCBI |
| KY925720 | A/Canoas/LACENRS-1212/2013(H3N2) | 8/6/2013 | NCBI |
| KY925421 | A/Canoas/LACENRS-1283/2015(H3N2) | 15/6/2015 | NCBI |
| KY926059 | A/Canoas/LACENRS-1317/2014(H3N2) | 7/7/2014 | NCBI |
| KY925589 | A/Canoas/LACENRS-1660/2014(H3N2) | 22/7/2014 | NCBI |
| KY925125 | A/Canoas/LACENRS-1793/2015(H3N2) | 17/7/2015 | NCBI |
| KY925599 | A/Canoas/LACENRS-773/2015(H3N2) | 6/5/2015 | NCBI |
| KY926096 | A/CruzAlta/LACENRS-2045/2014(H3N2) | 20/8/2014 | NCBI |
| KY926275 | A/CruzAlta/LACENRS-2623/2014(H3N2) | 19/10/2014 | NCBI |
| KY925426 | A/Farroupilha/LACENRS-1297/2014(H3N2) | 2/7/2014 | NCBI |
| KY926340 | A/FredericoWestphalen/LACENRS-204/2012(H3N2) | 13/6/2012 | NCBI |
| KY925138 | A/FredericoWestphalen/LACENRS-2130/2012(H3N2) | 26/7/2012 | NCBI |
| KY925015 | A/NovoHamburgo/LACENRS-201/2011(H3N2) | 10/6/2011 | NCBI |
| KY925394 | A/PaimFilho/LACENRS-32/2011(H3N2) | 6/4/2011 | NCBI |
| KY925800 | A/PassoFundo/LACENRS-1533/2013(H3N2) | 19/6/2013 | NCBI |
| KY925287 | A/Pelotas/LACENRS-1787/2015(H3N2) | 16/7/2015 | NCBI |
| KY925917 | A/PortoAlegre/LACENRS-1013/2016(H3N2) | 22/4/2016 | NCBI |
| KY926296 | A/PortoAlegre/LACENRS-1030/2015(H3N2) | 25/5/2015 | NCBI |
| KY925109 | A/PortoAlegre/LACENRS-1075/2015(H3N2) | 29/5/2015 | NCBI |
| KY925690 | A/PortoAlegre/LACENRS-1112/2015(H3N2) | 2/6/2015 | NCBI |
| KY925566 | A/PortoAlegre/LACENRS-1351/2014(H3N2) | 2/7/2014 | NCBI |
| KY925258 | A/PortoAlegre/LACENRS-1406/2014(H3N2) | 11/7/2014 | NCBI |
| KY926137 | A/PortoAlegre/LACENRS-1447/2014(H3N2) | 15/7/2014 | NCBI |
| KY926344 | A/PortoAlegre/LACENRS-1738/2015(H3N2) | 14/7/2015 | NCBI |
| KY925261 | A/PortoAlegre/LACENRS-1887/2015(H3N2) | 25/7/2015 | NCBI |
| KY925215 | A/PortoAlegre/LACENRS-1969/2015(H3N2) | 30/7/2015 | NCBI |
| KY924932 | A/PortoAlegre/LACENRS-2021/2014(H3N2) | 18/8/2014 | NCBI |
| KY925870 | A/PortoAlegre/LACENRS-2232/2014(H3N2) | 8/9/2014 | NCBI |
| KY925119 | A/PortoAlegre/LACENRS-2376/2014(H3N2) | 22/9/2014 | NCBI |
| KY926033 | A/PortoAlegre/LACENRS-275/2013(H3N2) | 28/3/2013 | NCBI |
| KY925024 | A/PortoAlegre/LACENRS-557/2011(H3N2) | 30/6/2011 | NCBI |
| KY926181 | A/PortoAlegre/LACENRS-820/2015(H3N2) | 11/5/2015 | NCBI |
| KY926256 | A/PortoAlegre/LACENRS-822/2011(H3N2) | 14/7/2011 | NCBI |

| | | | |
|---|---|---|---|
| KY925865 | A/PortoAlegre/LACENRS-846/2015(H3N2) | 9/5/2015 | NCBI |
| KY925818 | A/PortoAlegre/LACENRS-921/2015(H3N2) | 18/5/2015 | NCBI |
| KY925101 | A/Quarai/LACENRS-3508/2013(H3N2) | 28/8/2013 | NCBI |
| KY925163 | A/RioGrande/LACENRS-3171/2013(H3N2) | 12/8/2013 | NCBI |
| KY926393 | A/SantaCruzdoSul/LACENRS-1625/2014(H3N2) | 21/7/2014 | NCBI |
| KY925679 | A/SantaCruzdoSul/LACENRS-913/2011(H3N2) | 11/7/2011 | NCBI |
| KY926064 | A/SantaMaria/LACENRS-231/2011(H3N2) | 14/6/2011 | NCBI |
| KY925827 | A/SantoAntoniodaPatrulha/LACENRS-60/2011(H3N2) | 3/5/2011 | NCBI |
| KY925124 | A/SaoBorja/LACENRS-2679/2013(H3N2) | 23/7/2013 | NCBI |
| KY925368 | A/SapucaiadoSul/LACENRS-779/2011(H3N2) | 12/7/2011 | NCBI |
| KY925943 | A/Soledade/LACENRS-820/2011(H3N2) | 9/7/2011 | NCBI |
| KY926398 | A/Uruguaiana/LACENRS-368/2011(H3N2) | 20/6/2011 | NCBI |
| KY925772 | A/Viamao/LACENRS-733/2014(H3N2) | 28/5/2014 | NCBI |
| KY926260 | A/Viamao/LACENRS-974/2015(H3N2) | 21/5/2015 | NCBI |
| KY926222 | A/Ilopolis/LACENRS-1322/2014(H3N2) | 7/7/2014 | NCBI |
| KY925945 | A/Ilopolis/LACENRS-3175/2013(H3N2) | 12/8/2013 | NCBI |
| CY233259 | A/Tennessee/33/2017(H3N2) | 20/3/2017 | NCBI |
| CY233290 | A/Tennessee/32/2017(H3N2) | 9/3/2017 | NCBI |
| CY233306 | A/Tennessee/35/2017(H3N2) | 23/3/2017 | NCBI |
| CY233322 | A/Washington/31/2017(H3N2) | 10/3/2017 | NCBI |
| CY233330 | A/Washington/32/2017(H3N2) | 15/3/2017 | NCBI |
| CY233338 | A/Wisconsin/35/2017(H3N2) | 21/3/2017 | NCBI |
| CY233346 | A/Wisconsin/44/2017(H3N2) | 13/4/2017 | NCBI |
| CY233384 | A/Michigan/44/2017(H3N2) | 22/3/2017 | NCBI |
| CY233400 | A/Michigan/47/2017(H3N2) | 30/3/2017 | NCBI |
| CY233408 | A/Michigan/48/2017(H3N2) | 3/4/2017 | NCBI |
| CY233416 | A/Michigan/43/2017(H3N2) | 13/3/2017 | NCBI |
| CY233432 | A/Michigan/45/2017(H3N2) | 22/3/2017 | NCBI |
| CY233448 | A/Michigan/46/2017(H3N2) | 27/3/2017 | NCBI |
| CY232272 | A/Nevada/09/2017(H3N2) | 6/3/2017 | NCBI |
| CY232280 | A/Texas/72/2017(H3N2) | 8/3/2017 | NCBI |
| CY232288 | A/Texas/75/2017(H3N2) | 25/3/2017 | NCBI |
| CY232296 | A/Maryland/29/2017(H3N2) | 9/3/2017 | NCBI |
| CY232316 | A/Kentucky/25/2017(H3N2) | 30/3/2017 | NCBI |
| CY232324 | A/Virginia/25/2017(H3N2) | 30/3/2017 | NCBI |
| CY232332 | A/Virginia/26/2017(H3N2) | 30/3/2017 | NCBI |
| CY232340 | A/NorthCarolina/25/2017(H3N2) | 23/3/2017 | NCBI |

| | | | |
|---|---|---|---|
| CY232348 | A/NorthCarolina/22/2017(H3N2) | 8/3/2017 | NCBI |
| CY232356 | A/Ohio/10/2017(H3N2) | 4/4/2017 | NCBI |
| CY232364 | A/Massachusetts/11/2017(H3N2) | 30/3/2017 | NCBI |
| CY232372 | A/Georgia/07/2017(H3N2) | 16/3/2017 | NCBI |
| CY232380 | A/Georgia/08/2017(H3N2) | 20/3/2017 | NCBI |
| CY232388 | A/NewYork/19/2017(H3N2) | 30/1/2017 | NCBI |
| CY232396 | A/NewYork/20/2017(H3N2) | 29/3/2017 | NCBI |
| CY232404 | A/Florida/27/2017(H3N2) | 30/3/2017 | NCBI |
| CY232412 | A/Florida/28/2017(H3N2) | 4/4/2017 | NCBI |
| CY232428 | A/Delaware/25/2017(H3N2) | 4/4/2017 | NCBI |
| CY232436 | A/Delaware/24/2017(H3N2) | 3/4/2017 | NCBI |
| CY232444 | A/WestVirginia/13/2017(H3N2) | 13/3/2017 | NCBI |
| CY232452 | A/WestVirginia/15/2017(H3N2) | 28/3/2017 | NCBI |
| CY232460 | A/WestVirginia/16/2017(H3N2) | 1/4/2017 | NCBI |
| CY232468 | A/Florida/32/2017(H3N2) | 31/3/2017 | NCBI |
| CY232476 | A/Florida/34/2017(H3N2) | 9/4/2017 | NCBI |
| CY232484 | A/RhodeIsland/12/2017(H3N2) | 26/3/2017 | NCBI |
| CY232492 | A/RhodeIsland/13/2017(H3N2) | 10/4/2017 | NCBI |
| CY232500 | A/Vermont/11/2017(H3N2) | 1/4/2017 | NCBI |
| CY232508 | A/Vermont/12/2017(H3N2) | 1/4/2017 | NCBI |
| CY232516 | A/NewHampshire/18/2017(H3N2) | 6/4/2017 | NCBI |
| CY232524 | A/NewHampshire/19/2017(H3N2) | 8/4/2017 | NCBI |
| CY232532 | A/Connecticut/18/2017(H3N2) | 28/3/2017 | NCBI |
| CY232540 | A/Connecticut/19/2017(H3N2) | 31/3/2017 | NCBI |
| CY232554 | A/Maine/19/2017(H3N2) | 13/4/2017 | NCBI |
| CY232562 | A/Tennessee/38/2017(H3N2) | 11/4/2017 | NCBI |
| CY232570 | A/NorthCarolina/27/2017(H3N2) | 5/4/2017 | NCBI |
| CY232578 | A/NorthCarolina/26/2017(H3N2) | 3/4/2017 | NCBI |
| CY232586 | A/Minnesota/26/2017(H3N2) | 12/4/2017 | NCBI |
| CY232594 | A/Arkansas/20/2017(H3N2) | 2/4/2017 | NCBI |
| CY232602 | A/Michigan/52/2017(H3N2) | 6/4/2017 | NCBI |
| CY232610 | A/Michigan/53/2017(H3N2) | 16/4/2017 | NCBI |
| CY232618 | A/Louisiana/27/2017(H3N2) | 21/4/2017 | NCBI |
| CY232626 | A/Louisiana/28/2017(H3N2) | 21/4/2017 | NCBI |
| CY232634 | A/Louisiana/29/2017(H3N2) | 21/4/2017 | NCBI |
| CY234176 | A/Florida/30/2017(H3N2) | 7/2/2017 | NCBI |
| CY234184 | A/Florida/31/2017(H3N2) | 25/2/2017 | NCBI |

| | | | |
|---|---|---|---|
| CY234192 | A/Arizona/18/2017(H3N2) | 13/3/2017 | NCBI |
| CY234200 | A/SouthCarolina/12/2017(H3N2) | 10/4/2017 | NCBI |
| CY234208 | A/SouthCarolina/12/2017(H3N2) | 10/4/2017 | NCBI |
| CY234224 | A/SouthCarolina/13/2017(H3N2) | 12/4/2017 | NCBI |
| CY234216 | A/SouthCarolina/13/2017(H3N2) | 12/4/2017 | NCBI |
| CY234232 | A/Louisiana/02/2017(H3N2) | 10/1/2017 | NCBI |
| CY234240 | A/NewJersey/16/2017(H3N2) | 26/2/2017 | NCBI |
| CY234248 | A/California/38/2017(H3N2) | 8/3/2017 | NCBI |
| CY234256 | A/Pennsylvania/27/2017(H3N2) | 6/3/2017 | NCBI |
| CY234264 | A/Texas/70/2017(H3N2) | 17/3/2017 | NCBI |
| CY234272 | A/Maryland/25/2017(H3N2) | 27/2/2017 | NCBI |
| CY234280 | A/Oregon/11/2017(H3N2) | 10/3/2017 | NCBI |
| CY234288 | A/Tennessee/30/2017(H3N2) | 28/3/2017 | NCBI |
| CY234296 | A/Illinois/27/2017(H3N2) | 19/3/2017 | NCBI |
| CY234304 | A/Florida/26/2017(H3N2) | 26/3/2017 | NCBI |
| CY234312 | A/Wisconsin/39/2017(H3N2) | 19/3/2017 | NCBI |
| CY234320 | A/Michigan/39/2017(H3N2) | 16/3/2017 | NCBI |
| CY234328 | A/Arkansas/17/2017(H3N2) | 19/3/2017 | NCBI |
| CY231146 | A/Iowa/16/2017(H3N2) | 21/3/2017 | NCBI |
| CY234336 | A/Iowa/16/2017(H3N2) | 21/3/2017 | NCBI |
| CY234344 | A/Alaska/11/2017(H3N2) | 21/3/2017 | NCBI |
| CY234352 | A/NewMexico/16/2017(H3N2) | 26/3/2017 | NCBI |
| CY234360 | A/Utah/21/2017(H3N2) | 30/3/2017 | NCBI |
| CY234368 | A/Maryland/29/2017(H3N2) | 9/3/2017 | NCBI |
| CY232303 | A/Maryland/30/2017(H3N2) | 11/3/2017 | NCBI |
| CY234376 | A/Maryland/30/2017(H3N2) | 11/3/2017 | NCBI |
| CY234384 | A/Georgia/08/2017(H3N2) | 20/3/2017 | NCBI |
| CY234392 | A/Illinois/29/2017(H3N2) | 2/4/2017 | NCBI |
| CY234400 | A/Michigan/51/2017(H3N2) | 30/3/2017 | NCBI |
| CY234408 | A/Washington/37/2017(H3N2) | 20/3/2017 | NCBI |
| CY234416 | A/Washington/36/2017(H3N2) | 19/3/2017 | NCBI |
| CY234424 | A/RhodeIsland/12/2017(H3N2) | 26/3/2017 | NCBI |
| CY234432 | A/Louisiana/26/2017(H3N2) | 11/4/2017 | NCBI |
| CY234440 | A/Connecticut/18/2017(H3N2) | 28/3/2017 | NCBI |
| CY232547 | A/Maine/18/2017(H3N2) | 13/4/2017 | NCBI |
| CY234450 | A/Tennessee/38/2017(H3N2) | 11/4/2017 | NCBI |
| CY234458 | A/Indiana/22/2017(H3N2) | 10/4/2017 | NCBI |

| | | | |
|---|---|---|---|
| CY234466 | A/Maryland/32/2017(H3N2) | 10/4/2017 | NCBI |
| CY234474 | A/Maryland/33/2017(H3N2) | 15/4/2017 | NCBI |
| CY234482 | A/Hawaii/18/2017(H3N2) | 24/3/2017 | NCBI |
| CY234490 | A/Massachusetts/12/2017(H3N2) | 7/4/2017 | NCBI |
| CY234498 | A/Massachusetts/13/2017(H3N2) | 17/4/2017 | NCBI |
| CY234506 | A/Arizona/25/2017(H3N2) | 16/4/2017 | NCBI |
| CY234514 | A/NewYork/22/2017(H3N2) | 5/4/2017 | NCBI |
| CY234522 | A/NewYork/21/2017(H3N2) | 28/3/2017 | NCBI |
| CY234530 | A/Oklahoma/24/2017(H3N2) | 30/3/2017 | NCBI |
| CY234538 | A/Oklahoma/24/2017(H3N2) | 30/3/2017 | NCBI |
| CY234546 | A/NewHampshire/20/2017(H3N2) | 23/4/2017 | NCBI |
| CY234554 | A/Florida/35/2017(H3N2) | 19/4/2017 | NCBI |
| CY234562 | A/Florida/36/2017(H3N2) | 19/4/2017 | NCBI |
| CY234570 | A/Ohio/11/2017(H3N2) | 17/4/2017 | NCBI |
| CY234578 | A/Connecticut/21/2017(H3N2) | 17/4/2017 | NCBI |
| CY234586 | A/Connecticut/20/2017(H3N2) | 14/4/2017 | NCBI |
| CY234594 | A/Connecticut/20/2017(H3N2) | 14/4/2017 | NCBI |
| CY234602 | A/Iowa/20/2017(H3N2) | 11/4/2017 | NCBI |
| CY234610 | A/Iowa/21/2017(H3N2) | 21/4/2017 | NCBI |
| CY234618 | A/Wisconsin/48/2017(H3N2) | 18/4/2017 | NCBI |
| CY234626 | A/Wisconsin/49/2017(H3N2) | 20/4/2017 | NCBI |
| CY234634 | A/Indiana/23/2017(H3N2) | 13/4/2017 | NCBI |
| CY234642 | A/Indiana/24/2017(H3N2) | 15/4/2017 | NCBI |
| CY234650 | A/Maryland/34/2017(H3N2) | 24/4/2017 | NCBI |
| CY234658 | A/Maryland/35/2017(H3N2) | 26/4/2017 | NCBI |
| CY234666 | A/Mississippi/15/2017(H3N2) | 17/4/2017 | NCBI |
| CY234674 | A/Mississippi/16/2017(H3N2) | 21/4/2017 | NCBI |
| CY234682 | A/Minnesota/29/2017(H3N2) | 29/4/2017 | NCBI |
| CY234690 | A/Minnesota/28/2017(H3N2) | 24/4/2017 | NCBI |
| CY234698 | A/Michigan/57/2017(H3N2) | 17/4/2017 | NCBI |
| CY234706 | A/Michigan/58/2017(H3N2) | 1/5/2017 | NCBI |
| CY234714 | A/NewYork/28/2017(H3N2) | 19/4/2017 | NCBI |
| CY234722 | A/NewYork/27/2017(H3N2) | 9/4/2017 | NCBI |
| CY234730 | A/Texas/106/2017(H3N2) | 25/4/2017 | NCBI |
| CY234738 | A/Texas/107/2017(H3N2) | 26/4/2017 | NCBI |
| CY234746 | A/Alabama/21/2017(H3N2) | 9/5/2017 | NCBI |
| CY234754 | A/Alabama/18/2017(H3N2) | 4/5/2017 | NCBI |

| | | | |
|---|---|---|---|
| CY234762 | A/Alabama/19/2017(H3N2) | 4/5/2017 | NCBI |
| CY234770 | A/Alabama/20/2017(H3N2) | 4/5/2017 | NCBI |
| CY234778 | A/Connecticut/22/2017(H3N2) | 28/4/2017 | NCBI |
| CY234786 | A/Connecticut/23/2017(H3N2) | 1/5/2017 | NCBI |
| CY234794 | A/Louisiana/42/2017(H3N2) | 2/5/2017 | NCBI |
| CY234802 | A/Louisiana/41/2017(H3N2) | 26/4/2017 | NCBI |
| CY234810 | A/Wisconsin/312/2017(H3N2) | 1/5/2017 | NCBI |
| CY234818 | A/Wisconsin/313/2017(H3N2) | 8/5/2017 | NCBI |
| CY234826 | A/NorthCarolina/29/2017(H3N2) | 27/3/2017 | NCBI |
| CY234834 | A/Florida/38/2017(H3N2) | 30/4/2017 | NCBI |
| CY234842 | A/Florida/37/2017(H3N2) | 27/4/2017 | NCBI |
| CY234850 | A/Virginia/27/2017(H3N2) | 7/4/2017 | NCBI |
| CY234858 | A/Virginia/28/2017(H3N2) | 19/4/2017 | NCBI |
| CY234866 | A/WestVirginia/18/2017(H3N2) | 30/4/2017 | NCBI |
| CY234874 | A/Maryland/36/2017(H3N2) | 29/4/2017 | NCBI |
| CY235368 | A/NewJersey/24/2017(H3N2) | 24/2/2017 | NCBI |
| CY238242 | A/Tennessee/32/2017(H3N2) | 9/3/2017 | NCBI |
| CY238250 | A/Texas/82/2017(H3N2) | 1/2/2017 | NCBI |
| CY238258 | A/Texas/84/2017(H3N2) | 7/2/2017 | NCBI |
| CY238274 | A/Texas/87/2017(H3N2) | 12/2/2017 | NCBI |
| CY238282 | A/Texas/91/2017(H3N2) | 24/2/2017 | NCBI |
| CY238290 | A/Texas/92/2017(H3N2) | 28/2/2017 | NCBI |
| CY238298 | A/Texas/95/2017(H3N2) | 6/3/2017 | NCBI |
| CY238354 | A/Texas/97/2017(H3N2) | 14/3/2017 | NCBI |
| CY238362 | A/Texas/99/2017(H3N2) | 21/3/2017 | NCBI |
| CY238386 | A/Texas/76/2017(H3N2) | 9/1/2017 | NCBI |
| CY238394 | A/Texas/77/2017(H3N2) | 11/1/2017 | NCBI |
| CY238425 | A/Texas/85/2017(H3N2) | 4/2/2017 | NCBI |
| CY238433 | A/Texas/88/2017(H3N2) | 8/2/2017 | NCBI |
| CY238441 | A/Texas/90/2017(H3N2) | 14/2/2017 | NCBI |
| CY238465 | A/Texas/93/2017(H3N2) | 21/2/2017 | NCBI |
| CY238473 | A/Texas/94/2017(H3N2) | 27/2/2017 | NCBI |
| CY238497 | A/Texas/96/2017(H3N2) | 6/3/2017 | NCBI |
| CY238505 | A/Texas/98/2017(H3N2) | 12/3/2017 | NCBI |
| CY238561 | A/Texas/102/2017(H3N2) | 29/3/2017 | NCBI |
| CY238569 | A/Texas/104/2017(H3N2) | 31/3/2017 | NCBI |
| CY238585 | A/Michigan/55/2017(H3N2) | 11/4/2017 | NCBI |

| | | | |
|---|---|---|---|
| CY235376 | A/Louisiana/31/2017(H3N2) | 14/5/2017 | NCBI |
| CY235384 | A/Louisiana/32/2017(H3N2) | 14/5/2017 | NCBI |
| CY235392 | A/Louisiana/33/2017(H3N2) | 14/5/2017 | NCBI |
| CY238593 | A/Michigan/54/2017(H3N2) | 10/4/2017 | NCBI |
| CY238601 | A/Michigan/56/2017(H3N2) | 11/4/2017 | NCBI |
| CY238639 | A/Tennessee/41/2017(H3N2) | 27/4/2017 | NCBI |
| CY238662 | A/Tennessee/40/2017(H3N2) | 17/4/2017 | NCBI |
| CY235400 | A/SouthCarolina/14/2017(H3N2) | 27/4/2017 | NCBI |
| CY235408 | A/Louisiana/40/2017(H3N2) | 17/5/2017 | NCBI |
| CY235416 | A/Louisiana/35/2017(H3N2) | 13/5/2017 | NCBI |
| CY235424 | A/Louisiana/36/2017(H3N2) | 13/5/2017 | NCBI |
| CY235432 | A/Louisiana/38/2017(H3N2) | 14/5/2017 | NCBI |
| CY235440 | A/Arizona/29/2017(H3N2) | 15/5/2017 | NCBI |
| CY238678 | A/Wisconsin/43/2017(H3N2) | 13/4/2017 | NCBI |
| CY238686 | A/Wisconsin/44/2017(H3N2) | 13/4/2017 | NCBI |
| CY233370 | A/Michigan/42/2017(H3N2) | 13/3/2017 | NCBI |
| CY238701 | A/Michigan/48/2017(H3N2) | 3/4/2017 | NCBI |
| CY238714 | A/Michigan/46/2017(H3N2) | 27/3/2017 | NCBI |
| CY238722 | A/Michigan/49/2017(H3N2) | 3/4/2017 | NCBI |
| CY235456 | A/Wisconsin/98/2016(H3N2) | 6/12/2016 | NCBI |
| CY235464 | A/NewMexico/09/2017(H3N2) | 22/2/2017 | NCBI |
| CY235472 | A/Connecticut/12/2017(H3N2) | 4/3/2017 | NCBI |
| CY235480 | A/Hawaii/16/2017(H3N2) | 9/3/2017 | NCBI |
| CY235488 | A/WestVirginia/14/2017(H3N2) | 15/3/2017 | NCBI |
| CY235496 | A/Alaska/13/2017(H3N2) | 3/4/2017 | NCBI |
| CY235512 | A/Utah/23/2017(H3N2) | 13/4/2017 | NCBI |
| CY235520 | A/Idaho/23/2017(H3N2) | 31/3/2017 | NCBI |
| CY235528 | A/Maryland/33/2017(H3N2) | 15/4/2017 | NCBI |
| CY235536 | A/Nevada/10/2017(H3N2) | 22/3/2017 | NCBI |
| CY235552 | A/Nevada/11/2017(H3N2) | 25/3/2017 | NCBI |
| CY235568 | A/Hawaii/19/2017(H3N2) | 27/3/2017 | NCBI |
| CY235584 | A/Hawaii/18/2017(H3N2) | 24/3/2017 | NCBI |
| CY235592 | A/Arizona/24/2017(H3N2) | 9/4/2017 | NCBI |
| CY235608 | A/Arizona/25/2017(H3N2) | 16/4/2017 | NCBI |
| CY235616 | A/Washington/40/2017(H3N2) | 17/4/2017 | NCBI |
| CY235632 | A/Alaska/16/2017(H3N2) | 22/4/2017 | NCBI |
| CY235648 | A/Utah/26/2017(H3N2) | 24/4/2017 | NCBI |

| | | | |
|---|---|---|---|
| CY235664 | A/Utah/27/2017(H3N2) | 25/4/2017 | NCBI |
| CY235680 | A/Nevada/12/2017(H3N2) | 25/4/2017 | NCBI |
| CY235696 | A/Arizona/28/2017(H3N2) | 29/4/2017 | NCBI |
| CY235712 | A/Arizona/27/2017(H3N2) | 25/4/2017 | NCBI |
| CY235720 | A/Hawaii/21/2017(H3N2) | 6/4/2017 | NCBI |
| CY235736 | A/Hawaii/23/2017(H3N2) | 11/4/2017 | NCBI |
| CY235744 | A/Idaho/24/2017(H3N2) | 7/4/2017 | NCBI |
| CY235752 | A/Idaho/25/2017(H3N2) | 28/4/2017 | NCBI |
| CY235768 | A/Washington/289/2017(H3N2) | 4/4/2017 | NCBI |
| CY235776 | A/Washington/291/2017(H3N2) | 26/4/2017 | NCBI |
| CY235784 | A/Utah/29/2017(H3N2) | 11/5/2017 | NCBI |
| CY235792 | A/Utah/28/2017(H3N2) | 7/5/2017 | NCBI |
| CY235800 | A/Louisiana/41/2017(H3N2) | 26/4/2017 | NCBI |
| CY235808 | A/Minnesota/30/2017(H3N2) | 8/5/2017 | NCBI |
| CY235816 | A/Minnesota/31/2017(H3N2) | 13/5/2017 | NCBI |
| CY235824 | A/Hawaii/26/2017(H3N2) | 8/5/2017 | NCBI |
| CY235832 | A/SouthDakota/22/2017(H3N2) | 13/3/2017 | NCBI |
| CY235840 | A/SouthDakota/21/2017(H3N2) | 12/3/2017 | NCBI |
| CY235848 | A/SouthDakota/25/2017(H3N2) | 18/4/2017 | NCBI |
| CY235856 | A/SouthDakota/23/2017(H3N2) | 21/3/2017 | NCBI |
| CY235864 | A/SouthDakota/24/2017(H3N2) | 29/3/2017 | NCBI |
| CY235872 | A/Florida/39/2017(H3N2) | 10/5/2017 | NCBI |
| CY235880 | A/Florida/40/2017(H3N2) | 19/5/2017 | NCBI |
| CY235888 | A/NewYork/31/2017(H3N2) | 25/4/2017 | NCBI |
| CY235896 | A/NewYork/30/2017(H3N2) | 20/4/2017 | NCBI |
| CY235904 | A/Iowa/22/2017(H3N2) | 10/5/2017 | NCBI |
| CY235912 | A/Florida/41/2017(H3N2) | 22/5/2017 | NCBI |
| CY235920 | A/NorthDakota/16/2017(H3N2) | 14/5/2017 | NCBI |
| CY235928 | A/NorthDakota/17/2017(H3N2) | 14/5/2017 | NCBI |
| CY235936 | A/NorthDakota/18/2017(H3N2) | 22/5/2017 | NCBI |
| CY235944 | A/Wisconsin/314/2017(H3N2) | 15/5/2017 | NCBI |
| CY235952 | A/Louisiana/46/2017(H3N2) | 31/5/2017 | NCBI |
| CY235960 | A/Louisiana/47/2017(H3N2) | 2/6/2017 | NCBI |
| CY235968 | A/Minnesota/32/2017(H3N2) | 12/5/2017 | NCBI |
| CY235976 | A/Minnesota/33/2017(H3N2) | 15/5/2017 | NCBI |
| CY235984 | A/Maryland/37/2017(H3N2) | 11/5/2017 | NCBI |
| CY235992 | A/Hawaii/28/2017(H3N2) | 11/5/2017 | NCBI |

| | | | |
|---|---|---|---|
| CY236000 | A/California/58/2017(H3N2) | 31/5/2017 | NCBI |
| CY238771 | A/Washington/27/2017(H3N2) | 1/3/2017 | NCBI |
| CY238779 | A/Wisconsin/29/2017(H3N2) | 7/3/2017 | NCBI |
| CY238787 | A/Tennessee/37/2017(H3N2) | 13/4/2017 | NCBI |
| CY238739 | A/Texas/68/2017(H3N2) | 1/3/2017 | NCBI |
| CY238755 | A/Florida/23/2017(H3N2) | 23/3/2017 | NCBI |
| CY238795 | A/Louisiana/39/2017(H3N2) | 16/5/2017 | NCBI |
| CY238803 | A/NorthCarolina/62/2016(H3N2) | 2/12/2016 | NCBI |
| CY238811 | A/Washington/16/2017(H3N2) | 25/1/2017 | NCBI |
| CY238819 | A/NewYork/27/2017(H3N2) | 9/4/2017 | NCBI |
| CY238827 | A/Hawaii/27/2017(H3N2) | 16/5/2017 | NCBI |
| CY238835 | A/SouthDakota/22/2017(H3N2) | 13/3/2017 | NCBI |
| CY238843 | A/Wisconsin/314/2017(H3N2) | 15/5/2017 | NCBI |
| CY238851 | A/Washington/292/2017(H3N2) | 8/5/2017 | NCBI |
| CY238859 | A/Utah/31/2017(H3N2) | 7/6/2017 | NCBI |
| CY238867 | A/Utah/30/2017(H3N2) | 2/6/2017 | NCBI |
| CY238875 | A/Texas/274/2017(H3N2) | 30/5/2017 | NCBI |
| CY238883 | A/Arizona/32/2017(H3N2) | 1/5/2017 | NCBI |
| CY238891 | A/Arizona/35/2017(H3N2) | 21/5/2017 | NCBI |
| CY238899 | A/Arizona/37/2017(H3N2) | 21/5/2017 | NCBI |
| CY238907 | A/Florida/42/2017(H3N2) | 7/6/2017 | NCBI |
| CY238915 | A/Arizona/38/2017(H3N2) | 13/6/2017 | NCBI |
| CY238923 | A/Arizona/39/2017(H3N2) | 16/6/2017 | NCBI |
| CY238931 | A/Michigan/262/2017(H3N2) | 22/5/2017 | NCBI |
| CY238939 | A/Michigan/263/2017(H3N2) | 24/5/2017 | NCBI |
| CY238947 | A/Michigan/264/2017(H3N2) | 9/6/2017 | NCBI |
| KU322006 | A/Milano/112/2011(H3N2) | 2011 | NCBI |
| KU322007 | A/Milano/13/2014(H3N2) | 2014 | NCBI |
| KU322008 | A/Milano/22/2014(H3N2) | 2014 | NCBI |
| KU322009 | A/Milano/27/2013(H3N2) | 2013 | NCBI |
| KU322010 | A/Milano/28/2013(H3N2) | 2013 | NCBI |
| KU322011 | A/Milano/40/2012(H3N2) | 2012 | NCBI |
| KU322012 | A/Milano/111/2014(H3N2) | 2014 | NCBI |
| KU322013 | A/Milano/217/2011(H3N2) | 2011 | NCBI |
| KU322015 | A/Milano/118/2015(H3N2) | 2015 | NCBI |
| KU322016 | A/Milano/212/2015(H3N2) | 2015 | NCBI |
| KU322017 | A/Milano/270/2015(H3N2) | 2015 | NCBI |

| | | | |
|---|---|---|---|
| KU322018 | A/Milano/278/2015(H3N2) | 2015 | NCBI |
| KU322318 | A/Parma/3/2012(H3N2) | 2012 | NCBI |
| KU322277 | A/Parma/173/2011(H3N2) | 2011 | NCBI |
| KU322278 | A/Parma/539/2015(H3N2) | 2015 | NCBI |
| KU322279 | A/Parma/471/2015(H3N2) | 2015 | NCBI |
| KU322280 | A/Parma/287/2015(H3N2) | 2015 | NCBI |
| KU322281 | A/Parma/278/2015(H3N2) | 2015 | NCBI |
| KU322282 | A/Parma/157/2015(H3N2) | 2015 | NCBI |
| KU322285 | A/Parma/306/2015(H3N2) | 2015 | NCBI |
| KU322286 | A/Parma/151/2015(H3N2) | 2015 | NCBI |
| KU322293 | A/Parma/83/2014(H3N2) | 2014 | NCBI |
| KU322294 | A/Parma/82/2014(H3N2) | 2014 | NCBI |
| KU322295 | A/Parma/78/2014(H3N2) | 2014 | NCBI |
| KU322296 | A/Parma/77/2014(H3N2) | 2014 | NCBI |
| KU322297 | A/Parma/63/2014(H3N2) | 2014 | NCBI |
| KU322299 | A/Parma/40/2014(H3N2) | 2014 | NCBI |
| KU322309 | A/Parma/171/2011(H3N2) | 2011 | NCBI |
| KU322310 | A/Parma/172/2011(H3N2) | 2011 | NCBI |
| KU322311 | A/Parma/22/2012(H3N2) | 2012 | NCBI |
| KU322312 | A/Parma/21/2012(H3N2) | 2012 | NCBI |
| KU322313 | A/Parma/58/2012(H3N2) | 2012 | NCBI |
| KU322314 | A/Parma/152/2012(H3N2) | 2012 | NCBI |
| KU322315 | A/Parma/153/2012(H3N2) | 2012 | NCBI |
| KU322316 | A/Parma/156/2012(H3N2) | 2012 | NCBI |
| CY239649 | A/Tennessee/36/2017(H3N2) | 27/3/2017 | NCBI |
| CY238410 | A/Texas/80/2017(H3N2) | 25/1/2017 | NCBI |
| CY239657 | A/Texas/80/2017(H3N2) | 25/1/2017 | NCBI |
| CY239665 | A/Louisiana/37/2017(H3N2) | 13/5/2017 | NCBI |
| CY239673 | A/Louisiana/43/2017(H3N2) | 18/5/2017 | NCBI |
| CY239681 | A/Arizona/29/2017(H3N2) | 15/5/2017 | NCBI |
| CY239689 | A/Tennessee/33/2016(H3N2) | 6/12/2016 | NCBI |
| CY239697 | A/California/179/2016(H3N2) | 3/12/2016 | NCBI |
| CY239705 | A/Wisconsin/19/2017(H3N2) | 23/1/2017 | NCBI |
| CY239713 | A/Alaska/08/2017(H3N2) | 13/2/2017 | NCBI |
| CY239721 | A/Wyoming/15/2017(H3N2) | 18/3/2017 | NCBI |
| CY239736 | A/Alabama/16/2017(H3N2) | 29/3/2017 | NCBI |
| CY239744 | A/Alabama/13/2017(H3N2) | 13/3/2017 | NCBI |

| | | | |
|---|---|---|---|
| CY239752 | A/Florida/34/2017(H3N2) | 9/4/2017 | NCBI |
| CY239760 | A/Vermont/12/2017(H3N2) | 1/4/2017 | NCBI |
| CY239768 | A/Maine/19/2017(H3N2) | 13/4/2017 | NCBI |
| CY239776 | A/SouthDakota/19/2017(H3N2) | 29/3/2017 | NCBI |
| CY239784 | A/Utah/24/2017(H3N2) | 14/4/2017 | NCBI |
| CY239792 | A/NorthCarolina/26/2017(H3N2) | 3/4/2017 | NCBI |
| CY239800 | A/Louisiana/27/2017(H3N2) | 21/4/2017 | NCBI |
| CY239808 | A/Iowa/20/2017(H3N2) | 11/4/2017 | NCBI |
| CY239816 | A/Wisconsin/48/2017(H3N2) | 18/4/2017 | NCBI |
| CY239824 | A/Minnesota/28/2017(H3N2) | 24/4/2017 | NCBI |
| CY239832 | A/Michigan/58/2017(H3N2) | 1/5/2017 | NCBI |
| CY239840 | A/NorthDakota/18/2017(H3N2) | 22/5/2017 | NCBI |
| CY239848 | A/Minnesota/33/2017(H3N2) | 15/5/2017 | NCBI |
| CY236042 | A/Maryland/39/2017(H3N2) | 15/5/2017 | NCBI |
| CY239856 | A/Maryland/39/2017(H3N2) | 15/5/2017 | NCBI |
| CY239864 | A/Hawaii/28/2017(H3N2) | 11/5/2017 | NCBI |
| CY239872 | A/WestVirginia/21/2017(H3N2) | 8/6/2017 | NCBI |
| CY239880 | A/WestVirginia/20/2017(H3N2) | 6/6/2017 | NCBI |
| CY239888 | A/Florida/44/2017(H3N2) | 23/6/2017 | NCBI |
| CY239896 | A/Florida/43/2017(H3N2) | 20/6/2017 | NCBI |
| CY239904 | A/Alaska/17/2017(H3N2) | 23/6/2017 | NCBI |
| CY239912 | A/Texas/275/2017(H3N2) | 29/6/2017 | NCBI |
| CY239920 | A/Arizona/42/2017(H3N2) | 18/6/2017 | NCBI |
| CY239928 | A/Arizona/43/2017(H3N2) | 22/6/2017 | NCBI |
| CY239936 | A/Florida/45/2017(H3N2) | 25/6/2017 | NCBI |
| CY239944 | A/Florida/46/2017(H3N2) | 5/7/2017 | NCBI |
| CY239952 | A/Hawaii/31/2017(H3N2) | 8/6/2017 | NCBI |
| CY239960 | A/Hawaii/32/2017(H3N2) | 15/6/2017 | NCBI |
| CY239968 | A/Wisconsin/315/2017(H3N2) | 23/6/2017 | NCBI |
| CY239976 | A/Wisconsin/316/2017(H3N2) | 7/7/2017 | NCBI |
| CY239984 | A/Michigan/267/2017(H3N2) | 17/6/2017 | NCBI |
| CY239992 | A/Michigan/268/2017(H3N2) | 19/6/2017 | NCBI |
| CY240000 | A/NewJersey/28/2017(H3N2) | 14/7/2017 | NCBI |
| CY240008 | A/Texas/277/2017(H3N2) | 26/4/2017 | NCBI |
| CY240016 | A/Michigan/270/2017(H3N2) | 1/7/2017 | NCBI |
| CY240031 | A/NewJersey/26/2017(H3N2) | 21/6/2017 | NCBI |
| CY240039 | A/NewJersey/27/2017(H3N2) | 25/6/2017 | NCBI |

| | | | |
|---|---|---|---|
| CY240047 | A/Connecticut/24/2017(H3N2) | 13/6/2017 | NCBI |
| CY240055 | A/Connecticut/25/2017(H3N2) | 24/6/2017 | NCBI |
| CY240063 | A/Alaska/22/2017(H3N2) | 10/7/2017 | NCBI |
| CY240071 | A/Alaska/23/2017(H3N2) | 10/7/2017 | NCBI |
| CY240079 | A/Alaska/21/2017(H3N2) | 16/6/2017 | NCBI |
| CY240087 | A/Alaska/19/2017(H3N2) | 6/5/2017 | NCBI |
| CY240095 | A/Texas/279/2017(H3N2) | 29/6/2017 | NCBI |
| CY240103 | A/Texas/278/2017(H3N2) | 27/6/2017 | NCBI |
| CY240403 | A/Nicaragua/5929_02/2010(H3N2) | 2/6/2010 | NCBI |
| CY240411 | A/Nicaragua/5864_01/2010(H3N2) | 2/6/2010 | NCBI |
| CY240419 | A/Nicaragua/6006_02/2010(H3N2) | 6/6/2010 | NCBI |
| CY240427 | A/Nicaragua/5991_01/2010(H3N2) | 30/5/2010 | NCBI |
| CY240435 | A/Nicaragua/5907_02/2010(H3N2) | 2/6/2010 | NCBI |
| CY240443 | A/Nicaragua/5913_01/2010(H3N2) | 4/6/2010 | NCBI |
| CY240451 | A/Nicaragua/5891_01/2010(H3N2) | 6/6/2010 | NCBI |
| CY240459 | A/Nicaragua/5908_03/2010(H3N2) | 4/6/2010 | NCBI |
| CY240467 | A/Nicaragua/5872_01/2010(H3N2) | 7/6/2010 | NCBI |
| CY240475 | A/Nicaragua/6127_06/2014(H3N2) | 10/10/2014 | NCBI |
| CY240483 | A/Nicaragua/6585_04/2014(H3N2) | 6/9/2014 | NCBI |
| CY240491 | A/Nicaragua/5916_01/2010(H3N2) | 14/6/2010 | NCBI |
| CY240499 | A/Nicaragua/5867_01/2010(H3N2) | 30/5/2010 | NCBI |
| CY240507 | A/Nicaragua/5710_01/2010(H3N2) | 6/6/2010 | NCBI |
| CY240515 | A/Nicaragua/6530_19/2014(H3N2) | 10/11/2014 | NCBI |
| CY240523 | A/Nicaragua/6200_06/2014(H3N2) | 11/10/2014 | NCBI |
| CY240531 | A/Nicaragua/6024_04/2014(H3N2) | 24/9/2014 | NCBI |
| CY240539 | A/Nicaragua/6275_07/2014(H3N2) | 24/9/2014 | NCBI |
| CY240547 | A/Nicaragua/6302_05/2014(H3N2) | 27/10/2014 | NCBI |
| CY240555 | A/Nicaragua/6597_07/2014(H3N2) | 3/10/2014 | NCBI |
| CY240563 | A/Nicaragua/5920_08/2014(H3N2) | 3/10/2014 | NCBI |
| CY240571 | A/Nicaragua/7945_05/2014(H3N2) | 9/10/2014 | NCBI |
| CY240579 | A/Nicaragua/6219_07/2014(H3N2) | 12/10/2014 | NCBI |
| CY240587 | A/Nicaragua/6445_03/2014(H3N2) | 13/10/2014 | NCBI |
| CY240595 | A/Nicaragua/4537_01/2007(H3N2) | 11/6/2007 | NCBI |
| CY240603 | A/Nicaragua/4367_01/2007(H3N2) | 20/6/2007 | NCBI |
| CY240611 | A/Nicaragua/5627_04/2014(H3N2) | 22/10/2014 | NCBI |
| CY240619 | A/Nicaragua/4877_07/2014(H3N2) | 4/10/2014 | NCBI |
| CY240627 | A/Nicaragua/7487_12/2014(H3N2) | 14/10/2014 | NCBI |

| | | | |
|---|---|---|---|
| CY240635 | A/Nicaragua/4130_01/2007(H3N2) | 14/6/2007 | NCBI |
| CY240643 | A/Nicaragua/7596_05/2014(H3N2) | 26/10/2014 | NCBI |
| CY240651 | A/Nicaragua/4509_01/2007(H3N2) | 21/6/2007 | NCBI |
| CY240659 | A/Nicaragua/6826_06/2014(H3N2) | 15/10/2014 | NCBI |
| CY240667 | A/Nicaragua/7460_02/2014(H3N2) | 14/10/2014 | NCBI |
| CY240675 | A/Nicaragua/5359_04/2014(H3N2) | 3/10/2014 | NCBI |
| CY240683 | A/Nicaragua/5861_04/2014(H3N2) | 2/10/2014 | NCBI |
| CY240691 | A/Nicaragua/4305_07/2014(H3N2) | 2/10/2014 | NCBI |
| CY240699 | A/Nicaragua/5298_04/2014(H3N2) | 6/10/2014 | NCBI |
| CY240707 | A/Nicaragua/8020_05/2014(H3N2) | 7/10/2014 | NCBI |
| CY240715 | A/Nicaragua/6866_14/2014(H3N2) | 8/10/2014 | NCBI |
| CY240723 | A/Nicaragua/9073_02/2016(H3N2) | 14/12/2016 | NCBI |
| CY240731 | A/Nicaragua/9516_01/2016(H3N2) | 12/12/2016 | NCBI |
| CY240739 | A/Nicaragua/7718_04/2016(H3N2) | 14/12/2016 | NCBI |
| CY240747 | A/Nicaragua/6893_05/2016(H3N2) | 2/12/2016 | NCBI |
| CY240755 | A/Nicaragua/8774_10/2016(H3N2) | 22/12/2016 | NCBI |
| CY240763 | A/Nicaragua/7473_10/2016(H3N2) | 22/12/2016 | NCBI |
| CY242030 | A/California/59/2017(H3N2) | 26/6/2017 | NCBI |
| CY242038 | A/California/60/2017(H3N2) | 1/7/2017 | NCBI |
| CY242046 | A/Wisconsin/317/2017(H3N2) | 21/7/2017 | NCBI |
| CY242054 | A/WestVirginia/22/2017(H3N2) | 29/6/2017 | NCBI |
| CY242062 | A/Arizona/45/2017(H3N2) | 11/7/2017 | NCBI |
| CY242070 | A/Arizona/46/2017(H3N2) | 17/7/2017 | NCBI |
| CY242078 | A/Florida/47/2017(H3N2) | 9/7/2017 | NCBI |
| CY242086 | A/Florida/48/2017(H3N2) | 9/7/2017 | NCBI |
| CY242102 | A/Florida/29/2017(H3N2) | 7/1/2017 | NCBI |
| CY242110 | A/Wisconsin/19/2017(H3N2) | 23/1/2017 | NCBI |
| CY242126 | A/Texas/71/2017(H3N2) | 18/3/2017 | NCBI |
| CY242142 | A/Virginia/27/2017(H3N2) | 7/4/2017 | NCBI |
| CY242150 | A/Arizona/35/2017(H3N2) | 21/5/2017 | NCBI |
| CY242158 | A/Arizona/39/2017(H3N2) | 16/6/2017 | NCBI |
| CY242166 | A/Michigan/263/2017(H3N2) | 24/5/2017 | NCBI |
| CY242174 | A/Michigan/264/2017(H3N2) | 9/6/2017 | NCBI |
| CY242182 | A/Florida/44/2017(H3N2) | 23/6/2017 | NCBI |
| CY242190 | A/Florida/43/2017(H3N2) | 20/6/2017 | NCBI |
| CY242198 | A/Arizona/43/2017(H3N2) | 22/6/2017 | NCBI |
| CY242206 | A/NewJersey/28/2017(H3N2) | 14/7/2017 | NCBI |

| | | | |
|---|---|---|---|
| CY242214 | A/Connecticut/24/2017(H3N2) | 13/6/2017 | NCBI |
| CY242222 | A/Alaska/22/2017(H3N2) | 10/7/2017 | NCBI |
| CY242230 | A/Texas/278/2017(H3N2) | 27/6/2017 | NCBI |
| CY242238 | A/Washington/294/2017(H3N2) | 7/7/2017 | NCBI |
| CY242246 | A/Hawaii/34/2017(H3N2) | 5/7/2017 | NCBI |
| CY242254 | A/Hawaii/35/2017(H3N2) | 14/7/2017 | NCBI |
| CY242262 | A/Wyoming/18/2017(H3N2) | 18/7/2017 | NCBI |
| CY242270 | A/Michigan/273/2017(H3N2) | 20/7/2017 | NCBI |
| CY242278 | A/Michigan/275/2017(H3N2) | 20/7/2017 | NCBI |
| CY242286 | A/Ohio/31/2017(H3N2) | 25/7/2017 | NCBI |
| CY242294 | A/Ohio/33/2017(H3N2) | 31/7/2017 | NCBI |
| CY242302 | A/Ohio/32/2017(H3N2) | 29/7/2017 | NCBI |
| CY242310 | A/Alaska/24/2017(H3N2) | 18/7/2017 | NCBI |
| CY242318 | A/Alaska/25/2017(H3N2) | 21/7/2017 | NCBI |
| CY242326 | A/Texas/280/2017(H3N2) | 8/7/2017 | NCBI |
| CY242334 | A/Connecticut/27/2017(H3N2) | 24/7/2017 | NCBI |
| CY242342 | A/Connecticut/26/2017(H3N2) | 23/7/2017 | NCBI |
| CY242350 | A/Florida/50/2017(H3N2) | 26/7/2017 | NCBI |
| CY242358 | A/Florida/51/2017(H3N2) | 28/7/2017 | NCBI |
| CY242366 | A/California/63/2017(H3N2) | 26/7/2017 | NCBI |
| CY242374 | A/California/64/2017(H3N2) | 26/7/2017 | NCBI |
| CY242382 | A/California/65/2017(H3N2) | 26/7/2017 | NCBI |
| CY242390 | A/Arizona/47/2017(H3N2) | 30/7/2017 | NCBI |
| CY242398 | A/Wisconsin/318/2017(H3N2) | 24/7/2017 | NCBI |
| CY242406 | A/Wisconsin/319/2017(H3N2) | 29/7/2017 | NCBI |
| CY242414 | A/Florida/52/2017(H3N2) | 4/8/2017 | NCBI |
| CY242422 | A/Florida/53/2017(H3N2) | 6/8/2017 | NCBI |
| CY243423 | A/Missouri/24/2016(H3N2) | 18/1/2016 | NCBI |
| CY243431 | A/Washington/106/2016(H3N2) | 18/12/2016 | NCBI |
| CY243439 | A/Delaware/32/2016(H3N2) | 15/11/2016 | NCBI |
| CY243447 | A/Idaho/37/2016(H3N2) | 7/10/2016 | NCBI |
| CY243455 | A/Michigan/274/2017(H3N2) | 20/7/2017 | NCBI |
| CY243463 | A/Nebraska/07/2017(H3N2) | 31/7/2017 | NCBI |
| CY243471 | A/SouthDakota/26/2017(H3N2) | 4/8/2017 | NCBI |
| CY243479 | A/Hawaii/37/2017(H3N2) | 25/7/2017 | NCBI |
| CY243487 | A/Hawaii/38/2017(H3N2) | 25/7/2017 | NCBI |
| CY243495 | A/NewYork/32/2017(H3N2) | 28/6/2017 | NCBI |

| | | | |
|---|---|---|---|
| CY243503 | A/Pennsylvania/257/2017(H3N2) | 8/8/2017 | NCBI |
| CY243511 | A/NewJersey/30/2017(H3N2) | 13/8/2017 | NCBI |
| CY243519 | A/Alaska/26/2017(H3N2) | 2/6/2017 | NCBI |
| CY243527 | A/Alaska/28/2017(H3N2) | 5/6/2017 | NCBI |
| CY243535 | A/Alaska/27/2017(H3N2) | 3/6/2017 | NCBI |
| CY243543 | A/Alaska/29/2017(H3N2) | 8/6/2017 | NCBI |
| CY243551 | A/Alaska/30/2017(H3N2) | 18/6/2017 | NCBI |
| CY243559 | A/Alaska/31/2017(H3N2) | 26/6/2017 | NCBI |
| CY243567 | A/Alaska/32/2017(H3N2) | 30/6/2017 | NCBI |
| CY243575 | A/Alaska/33/2017(H3N2) | 30/6/2017 | NCBI |
| CY243583 | A/Alaska/34/2017(H3N2) | 1/7/2017 | NCBI |
| CY243591 | A/Alaska/37/2017(H3N2) | 11/7/2017 | NCBI |
| CY243599 | A/Alaska/35/2017(H3N2) | 7/7/2017 | NCBI |
| CY243607 | A/Alaska/36/2017(H3N2) | 9/7/2017 | NCBI |
| CY243615 | A/Alaska/38/2017(H3N2) | 14/7/2017 | NCBI |
| CY243623 | A/Alaska/40/2017(H3N2) | 16/7/2017 | NCBI |
| CY243631 | A/Alaska/42/2017(H3N2) | 24/7/2017 | NCBI |
| CY243639 | A/Alaska/41/2017(H3N2) | 21/7/2017 | NCBI |
| CY243647 | A/Alaska/43/2017(H3N2) | 29/7/2017 | NCBI |
| CY243655 | A/Alaska/45/2017(H3N2) | 30/7/2017 | NCBI |
| CY243670 | A/Alaska/44/2017(H3N2) | 29/7/2017 | NCBI |
| CY243678 | A/Alaska/46/2017(H3N2) | 31/7/2017 | NCBI |
| CY243686 | A/Iowa/23/2017(H3N2) | 3/8/2017 | NCBI |
| CY243694 | A/Iowa/24/2017(H3N2) | 3/8/2017 | NCBI |
| CY243702 | A/Iowa/25/2017(H3N2) | 10/8/2017 | NCBI |
| CY243710 | A/Florida/55/2017(H3N2) | 31/7/2017 | NCBI |
| CY243718 | A/Florida/57/2017(H3N2) | 6/8/2017 | NCBI |
| CY243726 | A/Wisconsin/320/2017(H3N2) | 5/8/2017 | NCBI |
| CY243734 | A/Wisconsin/321/2017(H3N2) | 14/8/2017 | NCBI |
| CY243742 | A/Florida/58/2017(H3N2) | 14/8/2017 | NCBI |
| CY243750 | A/Florida/59/2017(H3N2) | 17/8/2017 | NCBI |
| CY243758 | A/Missouri/28/2017(H3N2) | 14/8/2017 | NCBI |
| CY243862 | A/Michigan/55/2017(H3N2) | 11/4/2017 | NCBI |
| CY243838 | A/California/28/2017(H3N2) | 3/2/2017 | NCBI |
| MF736999 | A/Karachay-Cherkess/CRIE-301/2017(H3N2) | 28/2/2017 | NCBI |
| MF736972 | A/Smolensk/CRIE-44/2016(H3N2) | 7/12/2016 | NCBI |
| MF736983 | A/Vologda/CRIE-172/2016(H3N2) | 17/12/2016 | NCBI |

| | | | |
|---|---|---|---|
| CY244473 | A/California/59/2017(H3N2) | 26/6/2017 | NCBI |
| CY244481 | A/California/60/2017(H3N2) | 1/7/2017 | NCBI |
| CY244489 | A/Florida/48/2017(H3N2) | 9/7/2017 | NCBI |
| CY244497 | A/Michigan/273/2017(H3N2) | 20/7/2017 | NCBI |
| CY244505 | A/Michigan/275/2017(H3N2) | 20/7/2017 | NCBI |
| CY244513 | A/Alaska/24/2017(H3N2) | 18/7/2017 | NCBI |
| CY244521 | A/Alaska/25/2017(H3N2) | 21/7/2017 | NCBI |
| CY244529 | A/Alaska/47/2017(H3N2) | 3/8/2017 | NCBI |
| CY244537 | A/Texas/282/2017(H3N2) | 21/8/2017 | NCBI |
| CY244545 | A/Texas/283/2017(H3N2) | 22/8/2017 | NCBI |
| CY244553 | A/Arizona/48/2017(H3N2) | 31/7/2017 | NCBI |
| CY244561 | A/Massachusetts/14/2017(H3N2) | 13/7/2017 | NCBI |
| CY244569 | A/Maryland/40/2017(H3N2) | 17/8/2017 | NCBI |
| CY244577 | A/Montana/14/2017(H3N2) | 24/8/2017 | NCBI |
| CY244585 | A/NorthDakota/20/2017(H3N2) | 6/8/2017 | NCBI |
| CY244593 | A/NorthDakota/21/2017(H3N2) | 8/8/2017 | NCBI |
| CY244601 | A/California/68/2017(H3N2) | 11/8/2017 | NCBI |
| CY244609 | A/California/67/2017(H3N2) | 5/8/2017 | NCBI |
| CY244617 | A/Alaska/49/2017(H3N2) | 6/8/2017 | NCBI |
| CY244625 | A/Alaska/50/2017(H3N2) | 6/8/2017 | NCBI |
| CY244633 | A/Alaska/51/2017(H3N2) | 7/8/2017 | NCBI |
| CY244641 | A/Pennsylvania/258/2017(H3N2) | 15/8/2017 | NCBI |
| CY244649 | A/Pennsylvania/259/2017(H3N2) | 17/8/2017 | NCBI |
| CY244657 | A/Pennsylvania/262/2017(H3N2) | 24/8/2017 | NCBI |
| CY244665 | A/Pennsylvania/261/2017(H3N2) | 23/8/2017 | NCBI |
| CY244673 | A/Pennsylvania/260/2017(H3N2) | 22/8/2017 | NCBI |
| CY244681 | A/Ohio/36/2017(H3N2) | 21/8/2017 | NCBI |
| CY244689 | A/Ohio/37/2017(H3N2) | 23/8/2017 | NCBI |
| CY244697 | A/Hawaii/42/2017(H3N2) | 6/8/2017 | NCBI |
| CY244705 | A/Hawaii/41/2017(H3N2) | 4/8/2017 | NCBI |
| CY244713 | A/NorthCarolina/31/2017(H3N2) | 28/8/2017 | NCBI |
| CY244721 | A/Michigan/276/2017(H3N2) | 11/8/2017 | NCBI |
| CY244729 | A/Michigan/278/2017(H3N2) | 15/8/2017 | NCBI |
| CY244737 | A/Louisiana/48/2017(H3N2) | 21/8/2017 | NCBI |
| CY244745 | A/Louisiana/49/2017(H3N2) | 21/8/2017 | NCBI |
| CY244753 | A/Louisiana/50/2017(H3N2) | 21/8/2017 | NCBI |
| CY244761 | A/Mississippi/17/2017(H3N2) | 23/8/2017 | NCBI |

| | | | |
|---|---|---|---|
| CY244774 | A/Mississippi/19/2017(H3N2) | 25/8/2017 | NCBI |
| CY244878 | A/Washington/106/2016(H3N2) | 18/12/2016 | NCBI |
| CY244886 | A/Texas/71/2017(H3N2) | 18/3/2017 | NCBI |
| CY244894 | A/Virginia/30/2017(H3N2) | 30/8/2017 | NCBI |
| CY244902 | A/Arkansas/18/2017(H3N2) | 19/3/2017 | NCBI |
| CY244910 | A/Connecticut/26/2017(H3N2) | 23/7/2017 | NCBI |
| CY244918 | A/Florida/52/2017(H3N2) | 4/8/2017 | NCBI |
| CY244926 | A/Hawaii/37/2017(H3N2) | 25/7/2017 | NCBI |
| CY244934 | A/Pennsylvania/257/2017(H3N2) | 8/8/2017 | NCBI |
| CY244942 | A/Alaska/28/2017(H3N2) | 5/6/2017 | NCBI |
| CY244950 | A/Alaska/30/2017(H3N2) | 18/6/2017 | NCBI |
| CY244958 | A/Alaska/31/2017(H3N2) | 26/6/2017 | NCBI |
| CY244966 | A/Alaska/33/2017(H3N2) | 30/6/2017 | NCBI |
| CY244974 | A/Alaska/35/2017(H3N2) | 7/7/2017 | NCBI |
| CY244982 | A/Alaska/36/2017(H3N2) | 9/7/2017 | NCBI |
| CY244990 | A/Alaska/47/2017(H3N2) | 3/8/2017 | NCBI |
| CY244998 | A/Alaska/44/2017(H3N2) | 29/7/2017 | NCBI |
| CY245006 | A/Pennsylvania/259/2017(H3N2) | 17/8/2017 | NCBI |
| CY245014 | A/Michigan/278/2017(H3N2) | 15/8/2017 | NCBI |
| CY245022 | A/Louisiana/49/2017(H3N2) | 21/8/2017 | NCBI |
| CY245030 | A/Louisiana/50/2017(H3N2) | 21/8/2017 | NCBI |
| CY245038 | A/Idaho/26/2017(H3N2) | 16/7/2017 | NCBI |
| CY245046 | A/Idaho/27/2017(H3N2) | 27/8/2017 | NCBI |
| CY245054 | A/NewYork/33/2017(H3N2) | 2/8/2017 | NCBI |
| CY245062 | A/Florida/60/2017(H3N2) | 19/8/2017 | NCBI |
| CY245070 | A/Wisconsin/323/2017(H3N2) | 22/8/2017 | NCBI |
| CY245078 | A/Wisconsin/322/2017(H3N2) | 20/8/2017 | NCBI |
| CY245086 | A/Washington/295/2017(H3N2) | 20/8/2017 | NCBI |
| CY245094 | A/NewHampshire/21/2017(H3N2) | 2/9/2017 | NCBI |
| CY245102 | A/Montana/15/2017(H3N2) | 1/9/2017 | NCBI |
| CY245110 | A/Maryland/41/2017(H3N2) | 28/8/2017 | NCBI |
| CY245118 | A/Delaware/29/2017(H3N2) | 6/9/2017 | NCBI |
| CY245126 | A/Alaska/55/2017(H3N2) | 30/8/2017 | NCBI |
| CY245134 | A/Alaska/57/2017(H3N2) | 1/9/2017 | NCBI |
| CY245142 | A/Alaska/56/2017(H3N2) | 31/8/2017 | NCBI |
| CY245150 | A/Alaska/54/2017(H3N2) | 28/8/2017 | NCBI |
| CY245158 | A/Arkansas/21/2017(H3N2) | 31/8/2017 | NCBI |

| | | | |
|---|---|---|---|
| CY245166 | A/California/72/2017(H3N2) | 23/8/2017 | NCBI |
| CY245174 | A/California/71/2017(H3N2) | 17/8/2017 | NCBI |
| CY245182 | A/Ohio/39/2017(H3N2) | 5/9/2017 | NCBI |
| CY245190 | A/Ohio/40/2017(H3N2) | 6/9/2017 | NCBI |
| CY245198 | A/Arizona/49/2017(H3N2) | 16/8/2017 | NCBI |
| CY245206 | A/Arizona/50/2017(H3N2) | 22/8/2017 | NCBI |
| CY245214 | A/Vermont/13/2017(H3N2) | 3/9/2017 | NCBI |
| CY245222 | A/Vermont/14/2017(H3N2) | 5/9/2017 | NCBI |
| CY245230 | A/Michigan/283/2017(H3N2) | 30/8/2017 | NCBI |
| CY245238 | A/Michigan/282/2017(H3N2) | 29/8/2017 | NCBI |
| CY245246 | A/Nebraska/09/2017(H3N2) | 7/9/2017 | NCBI |
| CY245254 | A/Texas/284/2017(H3N2) | 8/9/2017 | NCBI |
| CY245262 | A/Texas/285/2017(H3N2) | 8/9/2017 | NCBI |
| CY245270 | A/Florida/63/2017(H3N2) | 6/9/2017 | NCBI |
| CY245278 | A/Florida/64/2017(H3N2) | 6/9/2017 | NCBI |
| CY245286 | A/Mississippi/20/2017(H3N2) | 3/9/2017 | NCBI |
| CY245294 | A/Mississippi/21/2017(H3N2) | 3/9/2017 | NCBI |
| CY245302 | A/Wisconsin/324/2017(H3N2) | 5/9/2017 | NCBI |
| CY245310 | A/Wisconsin/326/2017(H3N2) | 7/9/2017 | NCBI |
| CY245318 | A/Michigan/285/2017(H3N2) | 11/9/2017 | NCBI |
| CY245326 | A/Michigan/286/2017(H3N2) | 11/9/2017 | NCBI |
| CY245334 | A/Michigan/287/2017(H3N2) | 12/9/2017 | NCBI |
| CY245342 | A/Michigan/288/2017(H3N2) | 16/9/2017 | NCBI |
| CY245350 | A/California/76/2017(H3N2) | 1/8/2017 | NCBI |
| CY245358 | A/California/75/2017(H3N2) | 30/7/2017 | NCBI |
| CY245366 | A/California/77/2017(H3N2) | 2/8/2017 | NCBI |
| CY245374 | A/California/78/2017(H3N2) | 5/8/2017 | NCBI |
| CY245382 | A/Connecticut/28/2017(H3N2) | 10/8/2017 | NCBI |
| CY245390 | A/Connecticut/29/2017(H3N2) | 30/8/2017 | NCBI |
| CY245398 | A/Connecticut/30/2017(H3N2) | 9/9/2017 | NCBI |
| CY245406 | A/Colorado/15/2017(H3N2) | 6/9/2017 | NCBI |
| CY245414 | A/Colorado/14/2017(H3N2) | 1/9/2017 | NCBI |
| CY245422 | A/Colorado/12/2017(H3N2) | 7/8/2017 | NCBI |
| CY245430 | A/Colorado/13/2017(H3N2) | 12/8/2017 | NCBI |
| CY245438 | A/Iowa/26/2017(H3N2) | 24/8/2017 | NCBI |
| CY245446 | A/Iowa/27/2017(H3N2) | 2/9/2017 | NCBI |
| CY245454 | A/Hawaii/44/2017(H3N2) | 13/8/2017 | NCBI |

| | | | |
|---|---|---|---|
| CY245462 | A/Hawaii/45/2017(H3N2) | 22/8/2017 | NCBI |
| CY245470 | A/Washington/297/2017(H3N2) | 16/9/2017 | NCBI |
| CY246966 | A/Florida/61/2017(H3N2) | 29/8/2017 | NCBI |
| CY246974 | A/SouthCarolina/16/2017(H3N2) | 4/9/2017 | NCBI |
| CY246982 | A/Maryland/47/2017(H3N2) | 11/9/2017 | NCBI |
| CY246990 | A/Maryland/48/2017(H3N2) | 25/9/2017 | NCBI |
| CY249835 | A/Maryland/53/2017(H3N2) | 28/9/2017 | NCBI |
| CY246998 | A/Hawaii/42/2017(H3N2) | 6/8/2017 | NCBI |
| CY247006 | A/Idaho/26/2017(H3N2) | 16/7/2017 | NCBI |
| CY247014 | A/Wisconsin/323/2017(H3N2) | 22/8/2017 | NCBI |
| CY247022 | A/Alaska/55/2017(H3N2) | 30/8/2017 | NCBI |
| CY247030 | A/Alaska/57/2017(H3N2) | 1/9/2017 | NCBI |
| CY247038 | A/Arizona/50/2017(H3N2) | 22/8/2017 | NCBI |
| CY247046 | A/Vermont/14/2017(H3N2) | 5/9/2017 | NCBI |
| CY247054 | A/Texas/285/2017(H3N2) | 8/9/2017 | NCBI |
| CY247062 | A/Nebraska/10/2017(H3N2) | 17/9/2017 | NCBI |
| CY247070 | A/Montana/16/2017(H3N2) | 8/9/2017 | NCBI |
| CY247078 | A/Montana/17/2017(H3N2) | 9/9/2017 | NCBI |
| CY247086 | A/Alaska/60/2017(H3N2) | 22/8/2017 | NCBI |
| CY247094 | A/Alaska/61/2017(H3N2) | 5/9/2017 | NCBI |
| CY247102 | A/Alaska/62/2017(H3N2) | 7/9/2017 | NCBI |
| CY247110 | A/Georgia/14/2017(H3N2) | 15/9/2017 | NCBI |
| CY247118 | A/Georgia/13/2017(H3N2) | 7/9/2017 | NCBI |
| CY247126 | A/Georgia/12/2017(H3N2) | 23/8/2017 | NCBI |
| CY247134 | A/Georgia/11/2017(H3N2) | 24/7/2017 | NCBI |
| CY247142 | A/Georgia/10/2017(H3N2) | 13/6/2017 | NCBI |
| CY247150 | A/Georgia/09/2017(H3N2) | 23/5/2017 | NCBI |
| CY247158 | A/Georgia/15/2017(H3N2) | 16/9/2017 | NCBI |
| CY247166 | A/Louisiana/52/2017(H3N2) | 25/7/2017 | NCBI |
| CY247174 | A/Louisiana/51/2017(H3N2) | 9/7/2017 | NCBI |
| CY247182 | A/Delaware/30/2017(H3N2) | 17/9/2017 | NCBI |
| CY247197 | A/Ohio/42/2017(H3N2) | 19/9/2017 | NCBI |
| CY247205 | A/SouthDakota/27/2017(H3N2) | 13/9/2017 | NCBI |
| CY247213 | A/SouthDakota/28/2017(H3N2) | 13/9/2017 | NCBI |
| CY247221 | A/NewJersey/31/2017(H3N2) | 19/9/2017 | NCBI |
| CY249843 | A/Virginia/31/2017(H3N2) | 5/9/2017 | NCBI |
| CY249851 | A/Florida/65/2017(H3N2) | 18/9/2017 | NCBI |

| | | | |
|---|---|---|---|
| CY249859 | A/Florida/68/2017(H3N2) | 24/9/2017 | NCBI |
| CY249867 | A/Hawaii/49/2017(H3N2) | 12/9/2017 | NCBI |
| CY249875 | A/Hawaii/50/2017(H3N2) | 23/9/2017 | NCBI |
| CY249883 | A/NewMexico/20/2017(H3N2) | 13/9/2017 | NCBI |
| CY249891 | A/NewMexico/21/2017(H3N2) | 19/9/2017 | NCBI |
| CY249899 | A/Wyoming/20/2017(H3N2) | 22/9/2017 | NCBI |
| CY249907 | A/NorthCarolina/32/2017(H3N2) | 17/9/2017 | NCBI |
| CY249915 | A/NorthCarolina/34/2017(H3N2) | 27/9/2017 | NCBI |
| CY249923 | A/NorthCarolina/33/2017(H3N2) | 25/9/2017 | NCBI |
| CY249931 | A/Wisconsin/328/2017(H3N2) | 25/9/2017 | NCBI |
| CY249939 | A/Wisconsin/327/2017(H3N2) | 22/9/2017 | NCBI |
| CY249947 | A/Arizona/51/2017(H3N2) | 11/9/2017 | NCBI |
| CY249955 | A/California/79/2017(H3N2) | 27/9/2017 | NCBI |
| CY249963 | A/Tennessee/57/2017(H3N2) | 8/9/2017 | NCBI |
| CY249971 | A/Tennessee/58/2017(H3N2) | 20/9/2017 | NCBI |
| CY249979 | A/NewHampshire/22/2017(H3N2) | 27/9/2017 | NCBI |
| CY249987 | A/Pennsylvania/263/2017(H3N2) | 1/10/2017 | NCBI |
| CY249995 | A/Michigan/289/2017(H3N2) | 21/9/2017 | NCBI |
| CY250003 | A/Michigan/292/2017(H3N2) | 25/9/2017 | NCBI |
| CY250011 | A/Missouri/29/2017(H3N2) | 24/9/2017 | NCBI |
| CY250019 | A/Missouri/30/2017(H3N2) | 26/9/2017 | NCBI |
| CY250027 | A/Alaska/63/2017(H3N2) | 25/9/2017 | NCBI |
| CY250035 | A/Alaska/64/2017(H3N2) | 25/9/2017 | NCBI |
| CY250043 | A/Indiana/25/2017(H3N2) | 27/9/2017 | NCBI |
| CY250051 | A/Indiana/26/2017(H3N2) | 30/9/2017 | NCBI |
| CY250059 | A/Minnesota/36/2017(H3N2) | 27/9/2017 | NCBI |
| CY250067 | A/Minnesota/37/2017(H3N2) | 28/9/2017 | NCBI |
| CY250075 | A/Texas/287/2017(H3N2) | 28/9/2017 | NCBI |
| CY250083 | A/Texas/286/2017(H3N2) | 24/9/2017 | NCBI |
| CY250091 | A/Montana/19/2017(H3N2) | 23/9/2017 | NCBI |
| CY250099 | A/Montana/18/2017(H3N2) | 22/9/2017 | NCBI |
| CY250107 | A/Louisiana/53/2017(H3N2) | 4/10/2017 | NCBI |
| CY250115 | A/Louisiana/54/2017(H3N2) | 9/10/2017 | NCBI |
| CY250324 | A/Texas/71/2017(H3N2) | 18/3/2017 | NCBI |
| CY250340 | A/Iowa/23/2017(H3N2) | 3/8/2017 | NCBI |
| CY250348 | A/Missouri/28/2017(H3N2) | 14/8/2017 | NCBI |
| CY250356 | A/Texas/283/2017(H3N2) | 22/8/2017 | NCBI |

| | | | |
|---|---|---|---|
| CY250364 | A/NewYork/33/2017(H3N2) | 2/8/2017 | NCBI |
| CY250372 | A/NewHampshire/21/2017(H3N2) | 2/9/2017 | NCBI |
| CY250380 | A/Florida/63/2017(H3N2) | 6/9/2017 | NCBI |
| CY250388 | A/Iowa/26/2017(H3N2) | 24/8/2017 | NCBI |
| CY250396 | A/Iowa/27/2017(H3N2) | 2/9/2017 | NCBI |
| CY250404 | A/Washington/296/2017(H3N2) | 16/9/2017 | NCBI |
| CY250515 | A/Arkansas/23/2017(H3N2) | 9/10/2017 | NCBI |
| CY250523 | A/Arkansas/24/2017(H3N2) | 12/10/2017 | NCBI |
| CY250531 | A/Arkansas/25/2017(H3N2) | 12/10/2017 | NCBI |
| CY250539 | A/Arkansas/26/2017(H3N2) | 12/10/2017 | NCBI |
| CY250547 | A/Illinois/31/2017(H3N2) | 10/9/2017 | NCBI |
| CY250555 | A/Illinois/32/2017(H3N2) | 4/10/2017 | NCBI |
| CY250563 | A/Michigan/295/2017(H3N2) | 7/10/2017 | NCBI |
| CY250571 | A/Iowa/28/2017(H3N2) | 5/10/2017 | NCBI |
| CY250579 | A/Iowa/29/2017(H3N2) | 6/10/2017 | NCBI |
| CY250587 | A/Wisconsin/331/2017(H3N2) | 6/10/2017 | NCBI |
| CY250595 | A/Wisconsin/330/2017(H3N2) | 29/9/2017 | NCBI |
| CY250603 | A/Texas/288/2017(H3N2) | 13/10/2017 | NCBI |
| CY250611 | A/Texas/289/2017(H3N2) | 13/10/2017 | NCBI |
| CY250619 | A/Alabama/25/2017(H3N2) | 19/10/2017 | NCBI |
| CY250627 | A/Alabama/24/2017(H3N2) | 17/10/2017 | NCBI |
| CY250635 | A/Louisiana/56/2017(H3N2) | 19/10/2017 | NCBI |
| CY250643 | A/Louisiana/55/2017(H3N2) | 18/10/2017 | NCBI |
| CY250651 | A/Texas/290/2017(H3N2) | 18/10/2017 | NCBI |
| CY250659 | A/Texas/291/2017(H3N2) | 19/10/2017 | NCBI |
| CY250667 | A/Indiana/28/2017(H3N2) | 6/10/2017 | NCBI |
| CY250675 | A/Missouri/31/2017(H3N2) | 19/10/2017 | NCBI |
| CY250683 | A/Missouri/32/2017(H3N2) | 24/10/2017 | NCBI |
| CY250691 | A/Arkansas/27/2017(H3N2) | 20/10/2017 | NCBI |
| CY250699 | A/Arkansas/29/2017(H3N2) | 23/10/2017 | NCBI |
| CY250817 | A/Victoria/5025/2014(H3N2) | 1/9/2014 | NCBI |
| CY250825 | A/SouthAustralia/1/2012(H3N2) | 7/2/2012 | NCBI |
| CY250833 | A/Brisbane/1/2012(H3N2) | 6/1/2012 | NCBI |
| CY250841 | A/Brisbane/6/2012(H3N2) | 2/2/2012 | NCBI |
| CY250849 | A/Brisbane/11/2012(H3N2) | 13/2/2012 | NCBI |
| CY250857 | A/Brisbane/14/2012(H3N2) | 6/2/2012 | NCBI |
| CY250865 | A/Brisbane/21/2012(H3N2) | 4/3/2012 | NCBI |

| | | | |
|---|---|---|---|
| CY250873 | A/Brisbane/27/2012(H3N2) | 19/3/2012 | NCBI |
| CY250881 | A/Brisbane/26/2012(H3N2) | 8/3/2012 | NCBI |
| CY250889 | A/Victoria/508/2012(H3N2) | 5/5/2012 | NCBI |
| CY250897 | A/Brisbane/38/2012(H3N2) | 18/4/2012 | NCBI |
| CY250905 | A/Victoria/3/2012(H3N2) | 7/5/2012 | NCBI |
| CY250913 | A/Sydney/204/2012(H3N2) | 14/5/2012 | NCBI |
| CY250921 | A/Sydney/200/2012(H3N2) | 9/1/2012 | NCBI |
| CY250929 | A/Victoria/7/2012(H3N2) | 16/5/2012 | NCBI |
| CY250937 | A/Sydney/21/2012(H3N2) | 18/5/2012 | NCBI |
| CY250945 | A/SouthAustralia/36/2012(H3N2) | 28/5/2012 | NCBI |
| CY250953 | A/SouthAustralia/39/2012(H3N2) | 1/6/2012 | NCBI |
| CY250961 | A/Tasmania/7/2012(H3N2) | 19/6/2012 | NCBI |
| CY250969 | A/Victoria/27/2012(H3N2) | 12/6/2012 | NCBI |
| CY250977 | A/SouthAustralia/54/2012(H3N2) | 26/6/2012 | NCBI |
| CY250985 | A/Canberra/5/2012(H3N2) | 6/6/2012 | NCBI |
| CY250993 | A/Victoria/43/2012(H3N2) | 21/6/2012 | NCBI |
| CY251001 | A/SouthAustralia/66/2012(H3N2) | 24/6/2012 | NCBI |
| CY251009 | A/Victoria/535/2012(H3N2) | 18/6/2012 | NCBI |
| CY251017 | A/Victoria/50/2012(H3N2) | 25/6/2012 | NCBI |
| CY251025 | A/Victoria/78/2012(H3N2) | 3/7/2012 | NCBI |
| CY251033 | A/Victoria/84/2012(H3N2) | 4/7/2012 | NCBI |
| CY251041 | A/SouthAustralia/108/2012(H3N2) | 5/7/2012 | NCBI |
| CY251049 | A/SouthAustralia/109/2012(H3N2) | 5/7/2012 | NCBI |
| CY251057 | A/SouthAustralia/111/2012(H3N2) | 6/7/2012 | NCBI |
| CY251065 | A/SouthAustralia/114/2012(H3N2) | 6/7/2012 | NCBI |
| CY251073 | A/Victoria/87/2012(H3N2) | 4/7/2012 | NCBI |
| CY251081 | A/Victoria/346/2012(H3N2) | 12/7/2012 | NCBI |
| CY251103 | A/Sydney/1001/2012(H3N2) | 16/7/2012 | NCBI |
| CY251111 | A/Darwin/1001/2012(H3N2) | 16/7/2012 | NCBI |
| CY251119 | A/Canberra/1001/2012(H3N2) | 16/7/2012 | NCBI |
| CY251127 | A/Victoria/1002/2012(H3N2) | 16/7/2012 | NCBI |
| CY251135 | A/Sydney/1002/2012(H3N2) | 17/7/2012 | NCBI |
| CY251143 | A/Victoria/113/2012(H3N2) | 10/7/2012 | NCBI |
| CY251151 | A/Victoria/120/2012(H3N2) | 16/7/2012 | NCBI |
| CY251159 | A/Brisbane/61/2012(H3N2) | 6/6/2012 | NCBI |
| CY251167 | A/Canberra/33/2012(H3N2) | 7/7/2012 | NCBI |
| CY251175 | A/Canberra/35/2012(H3N2) | 8/7/2012 | NCBI |

| | | | |
|---|---|---|---|
| CY251183 | A/Victoria/148/2012(H3N2) | 25/7/2012 | NCBI |
| CY251191 | A/Victoria/641/2012(H3N2) | 18/7/2012 | NCBI |
| CY251199 | A/Victoria/692/2012(H3N2) | 1/8/2012 | NCBI |
| CY251207 | A/Victoria/142/2012(H3N2) | 23/7/2012 | NCBI |
| CY251215 | A/Victoria/147/2012(H3N2) | 24/7/2012 | NCBI |
| CY251223 | A/Victoria/448/2012(H3N2) | 6/8/2012 | NCBI |
| CY251231 | A/Victoria/723/2012(H3N2) | 8/8/2012 | NCBI |
| CY251239 | A/Perth/229/2012(H3N2) | 8/8/2012 | NCBI |
| CY251247 | A/Sydney/205/2012(H3N2) | 28/5/2012 | NCBI |
| CY251255 | A/Sydney/233/2012(H3N2) | 25/7/2012 | NCBI |
| CY251263 | A/Victoria/184/2012(H3N2) | 14/8/2012 | NCBI |
| CY251271 | A/Tasmania/60/2012(H3N2) | 20/8/2012 | NCBI |
| CY251279 | A/SouthAustralia/133/2012(H3N2) | 20/8/2012 | NCBI |
| CY251287 | A/Perth/254/2012(H3N2) | 10/7/2012 | NCBI |
| CY251295 | A/Victoria/187/2012(H3N2) | 16/8/2012 | NCBI |
| CY251303 | A/Sydney/195/2012(H3N2) | 6/8/2012 | NCBI |
| CY251311 | A/Victoria/173/2012(H3N2) | 6/8/2012 | NCBI |
| CY251319 | A/Victoria/175/2012(H3N2) | 4/8/2012 | NCBI |
| CY251327 | A/SouthAustralia/148/2012(H3N2) | 10/9/2012 | NCBI |
| CY251335 | A/Sydney/245/2012(H3N2) | 4/9/2012 | NCBI |
| CY251343 | A/SouthAustralia/157/2012(H3N2) | 27/9/2012 | NCBI |
| CY251351 | A/Victoria/918/2012(H3N2) | 4/10/2012 | NCBI |
| CY251359 | A/Perth/153/2012(H3N2) | 2/7/2012 | NCBI |
| CY251367 | A/SouthAustralia/160/2012(H3N2) | 7/10/2012 | NCBI |
| CY251375 | A/Victoria/1003/2012(H3N2) | 13/8/2012 | NCBI |
| CY251383 | A/Townsville/1004/2012(H3N2) | 13/8/2012 | NCBI |
| CY251391 | A/Victoria/213/2012(H3N2) | 28/8/2012 | NCBI |
| CY251399 | A/Victoria/1002A/2012(H3N2) | 19/7/2012 | NCBI |
| CY251407 | A/Victoria/1003A/2012(H3N2) | 20/7/2012 | NCBI |
| CY251415 | A/Perth/296/2012(H3N2) | 11/12/2012 | NCBI |
| CY251423 | A/SouthAustralia/2/2013(H3N2) | 9/1/2013 | NCBI |
| CY251431 | A/Newcastle/1/2013(H3N2) | 14/1/2013 | NCBI |
| CY251439 | A/Victoria/710/2013(H3N2) | 3/2/2013 | NCBI |
| CY251447 | A/Victoria/700/2013(H3N2) | 1/2/2013 | NCBI |
| CY251455 | A/Brisbane/1/2013(H3N2) | 3/1/2013 | NCBI |
| CY251463 | A/Brisbane/16/2013(H3N2) | 13/2/2013 | NCBI |
| CY251471 | A/Brisbane/21/2013(H3N2) | 27/2/2013 | NCBI |

| | | | |
|---|---|---|---|
| CY251479 | A/SouthAustralia/8/2013(H3N2) | 6/3/2013 | NCBI |
| CY251487 | A/Victoria/505/2013(H3N2) | 17/6/2013 | NCBI |
| CY251495 | A/Victoria/504/2013(H3N2) | 17/6/2013 | NCBI |
| CY251503 | A/Victoria/506/2013(H3N2) | 16/6/2013 | NCBI |
| CY251511 | A/SouthAustralia/29/2013(H3N2) | 30/6/2013 | NCBI |
| CY251519 | A/Sydney/28/2013(H3N2) | 19/6/2013 | NCBI |
| CY251527 | A/Perth/508/2013(H3N2) | 20/7/2013 | NCBI |
| CY251535 | A/Victoria/528/2013(H3N2) | 19/7/2013 | NCBI |
| CY251543 | A/Perth/510/2013(H3N2) | 27/7/2013 | NCBI |
| CY251551 | A/Perth/164/2013(H3N2) | 29/8/2013 | NCBI |
| CY251559 | A/SouthAustralia/79/2013(H3N2) | 23/9/2013 | NCBI |
| CY251567 | A/Brisbane/141/2013(H3N2) | 18/8/2013 | NCBI |
| CY251575 | A/Victoria/640/2013(H3N2) | 14/10/2013 | NCBI |
| CY251583 | A/Sydney/1014/2013(H3N2) | 24/10/2013 | NCBI |
| CY251591 | A/Sydney/1015/2013(H3N2) | 24/10/2013 | NCBI |
| CY251599 | A/Perth/247/2013(H3N2) | 7/10/2013 | NCBI |
| CY251607 | A/Canberra/113/2013(H3N2) | 11/10/2013 | NCBI |
| CY251615 | A/Brisbane/193/2013(H3N2) | 14/10/2013 | NCBI |
| CY251623 | A/Canberra/127/2013(H3N2) | 21/11/2013 | NCBI |
| CY251631 | A/Brisbane/198/2013(H3N2) | 6/11/2013 | NCBI |
| CY251639 | A/Brisbane/204/2013(H3N2) | 14/11/2013 | NCBI |
| CY251647 | A/Tasmania/3/2014(H3N2) | 6/1/2014 | NCBI |
| CY251655 | A/Perth/2/2014(H3N2) | 1/1/2014 | NCBI |
| CY251663 | A/Sydney/8/2014(H3N2) | 11/1/2014 | NCBI |
| CY251671 | A/Sydney/20/2014(H3N2) | 26/1/2014 | NCBI |
| CY251679 | A/Brisbane/43/2014(H3N2) | 17/2/2014 | NCBI |
| CY251687 | A/Sydney/26/2014(H3N2) | 1/3/2014 | NCBI |
| CY251695 | A/Tasmania/11/2014(H3N2) | 16/3/2014 | NCBI |
| CY251703 | A/Canberra/2/2014(H3N2) | 23/1/2014 | NCBI |
| CY251711 | A/Canberra/3/2014(H3N2) | 27/1/2014 | NCBI |
| CY251719 | A/Victoria/806/2014(H3N2) | 28/4/2014 | NCBI |
| CY251727 | A/Newcastle/20/2014(H3N2) | 7/2/2014 | NCBI |
| CY251735 | A/Victoria/503/2014(H3N2) | 5/5/2014 | NCBI |
| CY251743 | A/Sydney/30/2014(H3N2) | 2/3/2014 | NCBI |
| CY251751 | A/Perth/522/2014(H3N2) | 11/3/2014 | NCBI |
| CY251759 | A/Sydney/37/2014(H3N2) | 15/2/2014 | NCBI |
| CY251767 | A/Perth/526/2014(H3N2) | 17/3/2014 | NCBI |

| | | | |
|---|---|---|---|
| CY251775 | A/Brisbane/100/2014(H3N2) | 7/4/2014 | NCBI |
| CY251783 | A/Sydney/71/2014(H3N2) | 9/2/2014 | NCBI |
| CY251791 | A/SouthAustralia/24/2014(H3N2) | 8/6/2014 | NCBI |
| CY251799 | A/Tasmania/15/2014(H3N2) | 4/6/2014 | NCBI |
| CY251807 | A/Newcastle/22/2014(H3N2) | 16/6/2014 | NCBI |
| CY251815 | A/Newcastle/25/2014(H3N2) | 4/5/2014 | NCBI |
| CY251823 | A/SouthAustralia/40/2014(H3N2) | 20/6/2014 | NCBI |
| CY251831 | A/SouthAustralia/55/2014(H3N2) | 29/6/2014 | NCBI |
| CY251839 | A/SouthAustralia/1002/2014(H3N2) | 25/6/2014 | NCBI |
| CY251847 | A/Victoria/24/2014(H3N2) | 24/6/2014 | NCBI |
| CY251855 | A/Victoria/27/2014(H3N2) | 25/6/2014 | NCBI |
| CY251863 | A/Sydney/81/2014(H3N2) | 28/5/2014 | NCBI |
| CY251871 | A/SouthAustralia/1005/2014(H3N2) | 3/7/2014 | NCBI |
| CY251879 | A/Sydney/84/2014(H3N2) | 22/5/2014 | NCBI |
| CY251887 | A/Sydney/85/2014(H3N2) | 3/6/2014 | NCBI |
| CY251895 | A/SouthAustralia/1007/2014(H3N2) | 9/7/2014 | NCBI |
| CY251903 | A/Victoria/528/2014(H3N2) | 17/7/2014 | NCBI |
| CY251911 | A/Canberra/31/2014(H3N2) | 13/7/2014 | NCBI |
| CY251919 | A/Sydney/103/2014(H3N2) | 24/6/2014 | NCBI |
| CY251927 | A/SouthAustralia/91/2014(H3N2) | 18/7/2014 | NCBI |
| CY251935 | A/Victoria/69/2014(H3N2) | 16/7/2014 | NCBI |
| CY251943 | A/Sydney/1007/2014(H3N2) | 30/7/2014 | NCBI |
| CY251951 | A/Sydney/1005/2014(H3N2) | 28/7/2014 | NCBI |
| CY251959 | A/Canberra/82/2014(H3N2) | 31/7/2014 | NCBI |
| CY251967 | A/Newcastle/1016/2014(H3N2) | 6/8/2014 | NCBI |
| CY251975 | A/Victoria/265/2014(H3N2) | 11/8/2014 | NCBI |
| CY251983 | A/Sydney/1004/2014(H3N2) | 25/7/2014 | NCBI |
| CY251991 | A/Newcastle/63/2014(H3N2) | 6/8/2014 | NCBI |
| CY251999 | A/Tasmania/37/2014(H3N2) | 6/8/2014 | NCBI |
| CY252007 | A/Perth/551/2014(H3N2) | 8/5/2014 | NCBI |
| CY252015 | A/Brisbane/262/2014(H3N2) | 1/8/2014 | NCBI |
| CY252023 | A/Sydney/156/2014(H3N2) | 16/7/2014 | NCBI |
| CY252031 | A/Victoria/1012/2014(H3N2) | 12/9/2014 | NCBI |
| CY252039 | A/SouthAustralia/1032/2014(H3N2) | 8/9/2014 | NCBI |
| CY252047 | A/Victoria/5003/2014(H3N2) | 25/8/2014 | NCBI |
| CY252055 | A/Victoria/5006/2014(H3N2) | 25/8/2014 | NCBI |
| CY252063 | A/Victoria/5018/2014(H3N2) | 29/8/2014 | NCBI |

| | | | |
|---|---|---|---|
| CY252071 | A/Victoria/5054/2014(H3N2) | 16/9/2014 | NCBI |
| CY252079 | A/Victoria/5060/2014(H3N2) | 19/9/2014 | NCBI |
| CY252087 | A/Tasmania/122/2014(H3N2) | 5/9/2014 | NCBI |
| CY252095 | A/Victoria/5062/2014(H3N2) | 23/9/2014 | NCBI |
| CY252103 | A/Victoria/5064/2014(H3N2) | 22/9/2014 | NCBI |
| CY252111 | A/Victoria/5065/2014(H3N2) | 24/9/2014 | NCBI |
| CY252119 | A/Victoria/5070/2014(H3N2) | 1/10/2014 | NCBI |
| CY252127 | A/Victoria/5071/2014(H3N2) | 3/10/2014 | NCBI |
| CY252135 | A/Victoria/673/2014(H3N2) | 7/9/2014 | NCBI |
| CY252143 | A/Brisbane/341/2014(H3N2) | 3/10/2014 | NCBI |
| CY252151 | A/Victoria/5075/2014(H3N2) | 30/10/2014 | NCBI |
| CY252159 | A/Victoria/5076/2014(H3N2) | 27/10/2014 | NCBI |
| CY252167 | A/Sydney/1065/2014(H3N2) | 1/9/2014 | NCBI |
| CY252175 | A/Victoria/115/2014(H3N2) | 12/8/2014 | NCBI |
| CY252183 | A/Brisbane/368/2014(H3N2) | 21/11/2014 | NCBI |
| CY252191 | A/Brisbane/359/2014(H3N2) | 3/11/2014 | NCBI |
| CY252199 | A/Sydney/327/2014(H3N2) | 3/12/2014 | NCBI |
| CY252207 | A/Perth/1055/2014(H3N2) | 6/11/2014 | NCBI |
| CY252215 | A/Victoria/6001/2015(H3N2) | 4/1/2015 | NCBI |
| CY252223 | A/SouthAustralia/1/2015(H3N2) | 7/2/2015 | NCBI |
| CY252231 | A/SouthAustralia/9/2015(H3N2) | 12/2/2015 | NCBI |
| CY252239 | A/SouthAustralia/11/2015(H3N2) | 13/2/2015 | NCBI |
| CY252247 | A/Victoria/901/2015(H3N2) | 7/2/2015 | NCBI |
| CY252255 | A/Sydney/6/2015(H3N2) | 5/1/2015 | NCBI |
| CY252263 | A/Sydney/5/2015(H3N2) | 25/1/2015 | NCBI |
| CY252271 | A/Tasmania/2/2015(H3N2) | 23/2/2015 | NCBI |
| CY252279 | A/Canberra/8/2015(H3N2) | 27/1/2015 | NCBI |
| CY252287 | A/Sydney/4/2015(H3N2) | 5/1/2015 | NCBI |
| CY252295 | A/SouthAustralia/12/2015(H3N2) | 19/2/2015 | NCBI |
| CY252303 | A/SouthAustralia/16/2015(H3N2) | 17/2/2015 | NCBI |
| CY252311 | A/Sydney/10/2015(H3N2) | 20/2/2015 | NCBI |
| CY252319 | A/SouthAustralia/21/2015(H3N2) | 13/3/2015 | NCBI |
| CY252327 | A/Sydney/18/2015(H3N2) | 11/3/2015 | NCBI |
| CY252335 | A/Sydney/500/2015(H3N2) | 9/1/2015 | NCBI |
| CY252343 | A/Brisbane/20/2015(H3N2) | 2/2/2015 | NCBI |
| CY252351 | A/Brisbane/22/2015(H3N2) | 31/1/2015 | NCBI |
| CY252359 | A/Sydney/529/2014(H3N2) | 20/11/2014 | NCBI |

| | | | |
|---|---|---|---|
| CY252367 | A/Brisbane/12/2015(H3N2) | 27/1/2015 | NCBI |
| CY252375 | A/SouthAustralia/36/2015(H3N2) | 23/3/2015 | NCBI |
| CY252383 | A/SouthAustralia/37/2015(H3N2) | 26/3/2015 | NCBI |
| CY252391 | A/Sydney/530/2014(H3N2) | 4/12/2014 | NCBI |
| CY252399 | A/Townsville/1001/2015(H3N2) | 23/3/2015 | NCBI |
| CY252407 | A/SouthAustralia/33/2015(H3N2) | 23/3/2015 | NCBI |
| CY252415 | A/Sydney/12/2015(H3N2) | 11/3/2015 | NCBI |
| CY252423 | A/SouthAustralia/34/2015(H3N2) | 23/3/2015 | NCBI |
| CY252431 | A/Townsville/1002/2015(H3N2) | 10/4/2015 | NCBI |
| CY252439 | A/Brisbane/40/2015(H3N2) | 14/3/2015 | NCBI |
| CY252447 | A/Townsville/8/2015(H3N2) | 16/3/2015 | NCBI |
| CY252455 | A/Brisbane/43/2015(H3N2) | 21/3/2015 | NCBI |
| CY252463 | A/Townsville/10/2015(H3N2) | 22/3/2015 | NCBI |
| CY252471 | A/Brisbane/47/2015(H3N2) | 18/3/2015 | NCBI |
| CY252479 | A/SouthAustralia/49/2015(H3N2) | 6/4/2015 | NCBI |
| CY252487 | A/SouthAustralia/41/2015(H3N2) | 8/4/2015 | NCBI |
| CY252495 | A/Townsville/11/2015(H3N2) | 23/3/2015 | NCBI |
| CY252503 | A/Sydney/20/2015(H3N2) | 24/3/2015 | NCBI |
| CY252511 | A/Sydney/29/2015(H3N2) | 9/4/2015 | NCBI |
| CY252519 | A/Victoria/509/2015(H3N2) | 24/4/2015 | NCBI |
| CY252527 | A/Townsville/12/2015(H3N2) | 25/3/2015 | NCBI |
| CY252535 | A/SouthAustralia/57/2015(H3N2) | 20/4/2015 | NCBI |
| CY252543 | A/Canberra/12/2015(H3N2) | 12/2/2015 | NCBI |
| CY252551 | A/SouthAustralia/52/2015(H3N2) | 22/4/2015 | NCBI |
| CY252559 | A/Canberra/13/2015(H3N2) | 23/3/2015 | NCBI |
| CY252567 | A/SouthAustralia/60/2015(H3N2) | 3/5/2015 | NCBI |
| CY252575 | A/SouthAustralia/61/2015(H3N2) | 9/5/2015 | NCBI |
| CY252583 | A/SouthAustralia/62/2015(H3N2) | 10/5/2015 | NCBI |
| CY252591 | A/Brisbane/71/2015(H3N2) | 31/3/2015 | NCBI |
| CY252599 | A/Brisbane/82/2015(H3N2) | 22/4/2015 | NCBI |
| CY252607 | A/Brisbane/88/2015(H3N2) | 5/5/2015 | NCBI |
| CY252615 | A/Brisbane/79/2015(H3N2) | 5/4/2015 | NCBI |
| CY252623 | A/Brisbane/80/2015(H3N2) | 22/4/2015 | NCBI |
| CY252631 | A/SouthAustralia/64/2015(H3N2) | 22/5/2015 | NCBI |
| CY252639 | A/Newcastle/12/2015(H3N2) | 6/5/2015 | NCBI |
| CY252647 | A/Perth/15/2015(H3N2) | 20/2/2015 | NCBI |
| CY252655 | A/Perth/26/2015(H3N2) | 21/3/2015 | NCBI |

| | | | |
|---|---|---|---|
| CY252663 | A/SouthAustralia/70/2015(H3N2) | 2/6/2015 | NCBI |
| CY252671 | A/Victoria/511/2015(H3N2) | 16/5/2015 | NCBI |
| CY252679 | A/Perth/39/2015(H3N2) | 12/5/2015 | NCBI |
| CY252687 | A/Brisbane/105/2015(H3N2) | 12/5/2015 | NCBI |
| CY252695 | A/Perth/31/2015(H3N2) | 13/4/2015 | NCBI |
| CY252703 | A/Perth/33/2015(H3N2) | 7/4/2015 | NCBI |
| CY252711 | A/Perth/36/2015(H3N2) | 3/5/2015 | NCBI |
| CY252719 | A/Perth/40/2015(H3N2) | 12/5/2015 | NCBI |
| CY252727 | A/Perth/43/2015(H3N2) | 21/5/2015 | NCBI |
| CY252735 | A/Victoria/1000/2015(H3N2) | 17/6/2015 | NCBI |
| CY252743 | A/Victoria/803/2015(H3N2) | 17/6/2015 | NCBI |
| CY252751 | A/Brisbane/102/2015(H3N2) | 7/5/2015 | NCBI |
| CY252759 | A/Tasmania/7/2015(H3N2) | 20/4/2015 | NCBI |
| CY252767 | A/Brisbane/1000/2015(H3N2) | 5/6/2015 | NCBI |
| CY252775 | A/SouthAustralia/1002/2015(H3N2) | 9/6/2015 | NCBI |
| CY252783 | A/Sydney/1005/2015(H3N2) | 19/6/2015 | NCBI |
| CY252791 | A/Sydney/1006/2015(H3N2) | 18/6/2015 | NCBI |
| CY252799 | A/Brisbane/107/2015(H3N2) | 25/5/2015 | NCBI |
| CY252807 | A/SouthAustralia/1001/2015(H3N2) | 15/5/2015 | NCBI |
| CY252815 | A/Victoria/4/2015(H3N2) | 24/4/2015 | NCBI |
| CY252823 | A/Victoria/7/2015(H3N2) | 15/5/2015 | NCBI |
| CY252831 | A/Victoria/11/2015(H3N2) | 20/6/2015 | NCBI |
| CY252839 | A/Brisbane/115/2015(H3N2) | 9/6/2015 | NCBI |
| CY252847 | A/Brisbane/119/2015(H3N2) | 10/6/2015 | NCBI |
| CY252855 | A/Brisbane/123/2015(H3N2) | 13/6/2015 | NCBI |
| CY252863 | A/Canberra/20/2015(H3N2) | 6/6/2015 | NCBI |
| CY252871 | A/Victoria/5/2015(H3N2) | 29/4/2015 | NCBI |
| CY252879 | A/Victoria/9/2015(H3N2) | 21/5/2015 | NCBI |
| CY252887 | A/Tasmania/25/2015(H3N2) | 26/6/2015 | NCBI |
| CY252895 | A/Sydney/44/2015(H3N2) | 9/6/2015 | NCBI |
| CY252903 | A/SouthAustralia/79/2015(H3N2) | 25/6/2015 | NCBI |
| CY252922 | A/Newcastle/1001/2015(H3N2) | 30/6/2015 | NCBI |
| CY252930 | A/SouthAustralia/1005/2015(H3N2) | 2/7/2015 | NCBI |
| CY252938 | A/Newcastle/17/2015(H3N2) | 24/6/2015 | NCBI |
| CY252946 | A/Tasmania/28/2015(H3N2) | 8/7/2015 | NCBI |
| CY252954 | A/Victoria/515/2015(H3N2) | 28/6/2015 | NCBI |
| CY252962 | A/Newcastle/14/2015(H3N2) | 21/6/2015 | NCBI |

| | | | |
|---|---|---|---|
| CY252970 | A/Victoria/516/2015(H3N2) | 6/7/2015 | NCBI |
| CY252978 | A/Victoria/519/2015(H3N2) | 27/6/2015 | NCBI |
| CY252986 | A/Victoria/522/2015(H3N2) | 30/6/2015 | NCBI |
| CY252994 | A/Victoria/523/2015(H3N2) | 1/7/2015 | NCBI |
| CY253002 | A/Victoria/527/2015(H3N2) | 25/6/2015 | NCBI |
| CY253010 | A/Victoria/528/2015(H3N2) | 1/7/2015 | NCBI |
| CY253018 | A/Victoria/535/2015(H3N2) | 11/7/2015 | NCBI |
| CY253026 | A/Victoria/538/2015(H3N2) | 9/7/2015 | NCBI |
| CY253034 | A/Victoria/536/2015(H3N2) | 11/7/2015 | NCBI |
| CY253042 | A/Tasmania/1005/2015(H3N2) | 30/6/2015 | NCBI |
| CY253050 | A/Newcastle/1002/2015(H3N2) | 1/7/2015 | NCBI |
| CY253058 | A/Victoria/2502/2015(H3N2) | 8/7/2015 | NCBI |
| CY253066 | A/Victoria/16/2015(H3N2) | 9/7/2015 | NCBI |
| CY253074 | A/Brisbane/127/2015(H3N2) | 24/6/2015 | NCBI |
| CY253082 | A/Brisbane/128/2015(H3N2) | 16/6/2015 | NCBI |
| CY253090 | A/Brisbane/130/2015(H3N2) | 6/7/2015 | NCBI |
| CY253098 | A/Brisbane/132/2015(H3N2) | 29/6/2015 | NCBI |
| CY253106 | A/Townsville/16/2015(H3N2) | 19/6/2015 | NCBI |
| CY253114 | A/Brisbane/134/2015(H3N2) | 21/6/2015 | NCBI |
| CY253122 | A/Victoria/820/2015(H3N2) | 23/7/2015 | NCBI |
| CY253130 | A/Brisbane/140/2015(H3N2) | 6/7/2015 | NCBI |
| CY253138 | A/Brisbane/141/2015(H3N2) | 26/6/2015 | NCBI |
| CY253146 | A/Sydney/57/2015(H3N2) | 19/6/2015 | NCBI |
| CY253154 | A/Victoria/837/2015(H3N2) | 3/8/2015 | NCBI |
| CY253162 | A/Victoria/861/2015(H3N2) | 10/8/2015 | NCBI |
| CY253170 | A/Sydney/54/2015(H3N2) | 17/6/2015 | NCBI |
| CY253178 | A/Victoria/835/2015(H3N2) | 1/8/2015 | NCBI |
| CY253186 | A/Victoria/852/2015(H3N2) | 11/8/2015 | NCBI |
| CY253194 | A/Victoria/853/2015(H3N2) | 7/8/2015 | NCBI |
| CY253202 | A/Victoria/862/2015(H3N2) | 10/8/2015 | NCBI |
| CY253210 | A/Sydney/52/2015(H3N2) | 17/6/2015 | NCBI |
| CY253218 | A/Sydney/53/2015(H3N2) | 17/6/2015 | NCBI |
| CY253226 | A/Perth/61/2015(H3N2) | 10/7/2015 | NCBI |
| CY253234 | A/Tasmania/1012/2015(H3N2) | 4/8/2015 | NCBI |
| CY253242 | A/Sydney/1030/2015(H3N2) | 5/8/2015 | NCBI |
| CY253250 | A/SouthAustralia/1014/2015(H3N2) | 2/8/2015 | NCBI |
| CY253258 | A/Perth/50/2015(H3N2) | 18/6/2015 | NCBI |

| | | | |
|---|---|---|---|
| CY253266 | A/Perth/51/2015(H3N2) | 23/6/2015 | NCBI |
| CY253274 | A/Perth/53/2015(H3N2) | 29/6/2015 | NCBI |
| CY253282 | A/Perth/59/2015(H3N2) | 7/7/2015 | NCBI |
| CY253290 | A/Perth/62/2015(H3N2) | 12/7/2015 | NCBI |
| CY253298 | A/Brisbane/1005/2015(H3N2) | 10/8/2015 | NCBI |
| CY253306 | A/Sydney/1028/2015(H3N2) | 9/8/2015 | NCBI |
| CY253314 | A/Brisbane/1002/2015(H3N2) | 15/7/2015 | NCBI |
| CY253322 | A/Sydney/1029/2015(H3N2) | 4/8/2015 | NCBI |
| CY253330 | A/Victoria/1001/2015(H3N2) | 2/7/2015 | NCBI |
| CY253338 | A/Sydney/1010/2015(H3N2) | 14/7/2015 | NCBI |
| CY253346 | A/Brisbane/1003/2015(H3N2) | 28/7/2015 | NCBI |
| CY253354 | A/Brisbane/1004/2015(H3N2) | 24/7/2015 | NCBI |
| CY253362 | A/Victoria/17/2015(H3N2) | 8/7/2015 | NCBI |
| CY253370 | A/Victoria/19/2015(H3N2) | 17/7/2015 | NCBI |
| CY253378 | A/Victoria/21/2015(H3N2) | 2/7/2015 | NCBI |
| CY253386 | A/Perth/69/2015(H3N2) | 16/7/2015 | NCBI |
| CY253394 | A/Perth/89/2015(H3N2) | 3/8/2015 | NCBI |
| CY253402 | A/Perth/94/2015(H3N2) | 5/8/2015 | NCBI |
| CY253410 | A/Perth/97/2015(H3N2) | 6/8/2015 | NCBI |
| CY253418 | A/Perth/113/2015(H3N2) | 4/7/2015 | NCBI |
| CY253426 | A/Perth/114/2015(H3N2) | 7/7/2015 | NCBI |
| CY253434 | A/Perth/116/2015(H3N2) | 15/7/2015 | NCBI |
| CY253442 | A/Tasmania/1014/2015(H3N2) | 20/8/2015 | NCBI |
| CY253450 | A/Sydney/1011/2015(H3N2) | 20/8/2015 | NCBI |
| CY253458 | A/Tasmania/1015/2015(H3N2) | 17/8/2015 | NCBI |
| CY253466 | A/SouthAustralia/1018/2015(H3N2) | 17/8/2015 | NCBI |
| CY253474 | A/Tasmania/1016/2015(H3N2) | 17/8/2015 | NCBI |
| CY253482 | A/Sydney/1020/2015(H3N2) | 21/8/2015 | NCBI |
| CY253490 | A/Sydney/1023/2015(H3N2) | 19/8/2015 | NCBI |
| CY253498 | A/SouthAustralia/1021/2015(H3N2) | 25/8/2015 | NCBI |
| CY253506 | A/Tasmania/1021/2015(H3N2) | 20/8/2015 | NCBI |
| CY253514 | A/Sydney/1024/2015(H3N2) | 20/8/2015 | NCBI |
| CY253522 | A/SouthAustralia/1023/2015(H3N2) | 24/8/2015 | NCBI |
| CY253530 | A/Victoria/3070/2015(H3N2) | 7/9/2015 | NCBI |
| CY253538 | A/Victoria/36/2015(H3N2) | 3/8/2015 | NCBI |
| CY253546 | A/Victoria/37/2015(H3N2) | 5/8/2015 | NCBI |
| CY253554 | A/Victoria/48/2015(H3N2) | 10/8/2015 | NCBI |

| | | | |
|---|---|---|---|
| CY253562 | A/Victoria/23/2015(H3N2) | 23/7/2015 | NCBI |
| CY253570 | A/Victoria/24/2015(H3N2) | 24/7/2015 | NCBI |
| CY253578 | A/Perth/118/2015(H3N2) | 22/7/2015 | NCBI |
| CY253586 | A/Victoria/3067/2015(H3N2) | 7/9/2015 | NCBI |
| CY253594 | A/Sydney/1033/2015(H3N2) | 7/9/2015 | NCBI |
| CY253602 | A/Sydney/1034/2015(H3N2) | 7/9/2015 | NCBI |
| CY253610 | A/SouthAustralia/129/2015(H3N2) | 12/9/2015 | NCBI |
| CY253618 | A/SouthAustralia/132/2015(H3N2) | 10/9/2015 | NCBI |
| CY253626 | A/Victoria/43/2015(H3N2) | 7/8/2015 | NCBI |
| CY253634 | A/Perth/65/2015(H3N2) | 13/7/2015 | NCBI |
| CY253642 | A/SouthAustralia/1017/2015(H3N2) | 17/8/2015 | NCBI |
| CY253650 | A/Brisbane/1009/2015(H3N2) | 22/8/2015 | NCBI |
| CY253658 | A/Victoria/3072/2015(H3N2) | 7/9/2015 | NCBI |
| CY253666 | A/Victoria/3079/2015(H3N2) | 10/9/2015 | NCBI |
| CY253674 | A/Sydney/1013/2015(H3N2) | 12/8/2015 | NCBI |
| CY253682 | A/Sydney/1019/2015(H3N2) | 14/8/2015 | NCBI |
| CY253690 | A/Victoria/3071/2015(H3N2) | 7/9/2015 | NCBI |
| CY253698 | A/Victoria/3073/2015(H3N2) | 7/9/2015 | NCBI |
| CY253706 | A/Tasmania/1007/2015(H3N2) | 6/7/2015 | NCBI |
| CY253714 | A/Perth/92/2015(H3N2) | 3/8/2015 | NCBI |
| CY253722 | A/Victoria/3015/2015(H3N2) | 1/9/2015 | NCBI |
| CY253730 | A/Victoria/3036/2015(H3N2) | 24/8/2015 | NCBI |
| CY253738 | A/Victoria/3047/2015(H3N2) | 21/8/2015 | NCBI |
| CY253746 | A/SouthAustralia/119/2015(H3N2) | 21/8/2015 | NCBI |
| CY253754 | A/Sydney/1022/2015(H3N2) | 20/8/2015 | NCBI |
| CY253762 | A/Newcastle/1003/2015(H3N2) | 14/7/2015 | NCBI |
| CY253770 | A/SouthAustralia/1006/2015(H3N2) | 10/7/2015 | NCBI |
| CY253778 | A/Victoria/1006/2015(H3N2) | 14/7/2015 | NCBI |
| CY253786 | A/Perth/76/2015(H3N2) | 29/7/2015 | NCBI |
| CY253794 | A/Victoria/3054/2015(H3N2) | 3/9/2015 | NCBI |
| CY253802 | A/Victoria/3009/2015(H3N2) | 1/9/2015 | NCBI |
| CY253810 | A/Victoria/3011/2015(H3N2) | 1/9/2015 | NCBI |
| CY253818 | A/Sydney/98/2015(H3N2) | 1/9/2015 | NCBI |
| CY253826 | A/Victoria/3085/2015(H3N2) | 17/9/2015 | NCBI |
| CY253834 | A/Brisbane/171/2015(H3N2) | 24/8/2015 | NCBI |
| CY253842 | A/Brisbane/174/2015(H3N2) | 7/9/2015 | NCBI |
| CY253850 | A/SouthAustralia/133/2015(H3N2) | 25/9/2015 | NCBI |

| | | | |
|---|---|---|---|
| CY253858 | A/Brisbane/172/2015(H3N2) | 3/9/2015 | NCBI |
| CY253866 | A/Townsville/29/2015(H3N2) | 9/9/2015 | NCBI |
| CY253874 | A/Victoria/629/2015(H3N2) | 13/9/2015 | NCBI |
| CY253882 | A/SouthAustralia/135/2015(H3N2) | 24/9/2015 | NCBI |
| CY253890 | A/Victoria/640/2015(H3N2) | 24/9/2015 | NCBI |
| CY253898 | A/Victoria/3100/2015(H3N2) | 15/9/2015 | NCBI |
| CY253906 | A/Tasmania/1022/2015(H3N2) | 28/8/2015 | NCBI |
| CY253914 | A/SouthAustralia/139/2015(H3N2) | 3/9/2015 | NCBI |
| CY253922 | A/Sydney/1043/2015(H3N2) | 14/9/2015 | NCBI |
| CY253930 | A/Sydney/1036/2015(H3N2) | 27/8/2015 | NCBI |
| CY253938 | A/Sydney/1042/2015(H3N2) | 9/9/2015 | NCBI |
| CY253946 | A/Victoria/55/2015(H3N2) | 24/8/2015 | NCBI |
| CY253954 | A/Victoria/64/2015(H3N2) | 4/9/2015 | NCBI |
| CY253962 | A/Victoria/63/2015(H3N2) | 3/9/2015 | NCBI |
| CY253970 | A/SouthAustralia/145/2015(H3N2) | 9/10/2015 | NCBI |
| CY253978 | A/Victoria/1014/2015(H3N2) | 30/9/2015 | NCBI |
| CY253986 | A/Darwin/1000/2015(H3N2) | 8/10/2015 | NCBI |
| CY253994 | A/Victoria/51/2015(H3N2) | 19/8/2015 | NCBI |
| CY254002 | A/SouthAustralia/146/2015(H3N2) | 7/10/2015 | NCBI |
| CY254010 | A/SouthAustralia/162/2015(H3N2) | 19/10/2015 | NCBI |
| CY254018 | A/Victoria/50/2015(H3N2) | 18/8/2015 | NCBI |
| CY254026 | A/SouthAustralia/163/2015(H3N2) | 24/10/2015 | NCBI |
| CY254034 | A/SouthAustralia/1036/2015(H3N2) | 12/10/2015 | NCBI |
| CY254042 | A/SouthAustralia/167/2015(H3N2) | 18/9/2015 | NCBI |
| CY254050 | A/Townsville/32/2015(H3N2) | 21/9/2015 | NCBI |
| CY254058 | A/SouthAustralia/1038/2015(H3N2) | 15/9/2015 | NCBI |
| CY254066 | A/SouthAustralia/166/2015(H3N2) | 17/9/2015 | NCBI |
| CY254074 | A/Victoria/52/2015(H3N2) | 21/8/2015 | NCBI |
| CY254082 | A/Victoria/56/2015(H3N2) | 24/8/2015 | NCBI |
| CY254090 | A/Brisbane/179/2015(H3N2) | 23/9/2015 | NCBI |
| CY254098 | A/Sydney/223/2015(H3N2) | 11/9/2015 | NCBI |
| CY254106 | A/Sydney/248/2015(H3N2) | 2/9/2015 | NCBI |
| CY254114 | A/Victoria/53/2015(H3N2) | 24/8/2015 | NCBI |
| CY254122 | A/Victoria/2118/2015(H3N2) | 1/9/2015 | NCBI |
| CY254130 | A/Sydney/224/2015(H3N2) | 4/9/2015 | NCBI |
| CY254138 | A/Sydney/225/2015(H3N2) | 5/9/2015 | NCBI |
| CY254146 | A/SouthAustralia/172/2015(H3N2) | 31/10/2015 | NCBI |

| | | | |
|---|---|---|---|
| CY254154 | A/Sydney/242/2015(H3N2) | 13/9/2015 | NCBI |
| CY254162 | A/Sydney/241/2015(H3N2) | 26/10/2015 | NCBI |
| CY254170 | A/Brisbane/183/2015(H3N2) | 13/10/2015 | NCBI |
| CY254178 | A/Brisbane/184/2015(H3N2) | 24/10/2015 | NCBI |
| CY254186 | A/Brisbane/188/2015(H3N2) | 14/10/2015 | NCBI |
| CY254194 | A/Townsville/37/2015(H3N2) | 18/10/2015 | NCBI |
| CY254202 | A/Brisbane/197/2015(H3N2) | 9/11/2015 | NCBI |
| CY254210 | A/Brisbane/195/2015(H3N2) | 2/11/2015 | NCBI |
| CY254218 | A/Victoria/455/2015(H3N2) | 15/12/2015 | NCBI |
| CY254226 | A/Sydney/9/2016(H3N2) | 8/1/2016 | NCBI |
| CY254234 | A/Sydney/29/2016(H3N2) | 10/1/2016 | NCBI |
| CY254242 | A/SouthAustralia/3/2016(H3N2) | 3/1/2016 | NCBI |
| CY254265 | A/Tasmania/3/2016(H3N2) | 11/2/2016 | NCBI |
| CY254273 | A/Brisbane/30/2016(H3N2) | 27/1/2016 | NCBI |
| CY254281 | A/Sydney/80/2016(H3N2) | 3/3/2016 | NCBI |
| CY254289 | A/Victoria/2015/2016(H3N2) | 15/3/2016 | NCBI |
| CY254297 | A/SouthAustralia/18/2016(H3N2) | 22/3/2016 | NCBI |
| CY254305 | A/Brisbane/66/2016(H3N2) | 8/3/2016 | NCBI |
| CY254313 | A/Brisbane/68/2016(H3N2) | 14/3/2016 | NCBI |
| CY254321 | A/Newcastle/30/2016(H3N2) | 29/3/2016 | NCBI |
| CY254329 | A/Canberra/7/2016(H3N2) | 10/4/2016 | NCBI |
| CY254337 | A/Canberra/5/2016(H3N2) | 19/2/2016 | NCBI |
| CY254345 | A/SouthAustralia/31/2016(H3N2) | 10/6/2016 | NCBI |
| CY254353 | A/SouthAustralia/29/2016(H3N2) | 31/5/2016 | NCBI |
| CY254361 | A/Newcastle/1000/2016(H3N2) | 18/5/2016 | NCBI |
| CY254369 | A/SouthAustralia/27/2016(H3N2) | 9/5/2016 | NCBI |
| CY254377 | A/Sydney/101/2016(H3N2) | 4/6/2016 | NCBI |
| CY254385 | A/Victoria/12/2016(H3N2) | 18/6/2016 | NCBI |
| CY254393 | A/SouthAustralia/38/2016(H3N2) | 3/7/2016 | NCBI |
| CY254401 | A/SouthAustralia/39/2016(H3N2) | 3/7/2016 | NCBI |
| CY254409 | A/SouthAustralia/41/2016(H3N2) | 2/7/2016 | NCBI |
| CY254417 | A/SouthAustralia/34/2016(H3N2) | 24/6/2016 | NCBI |
| CY254425 | A/SouthAustralia/35/2016(H3N2) | 20/6/2016 | NCBI |
| CY254433 | A/SouthAustralia/51/2016(H3N2) | 6/7/2016 | NCBI |
| CY254441 | A/Victoria/2045/2016(H3N2) | 16/7/2016 | NCBI |
| CY254449 | A/SouthAustralia/54/2016(H3N2) | 12/7/2016 | NCBI |
| CY254457 | A/SouthAustralia/55/2016(H3N2) | 11/7/2016 | NCBI |

| | | | |
|---|---|---|---|
| CY254465 | A/Sydney/1014/2016(H3N2) | 11/7/2016 | NCBI |
| CY254473 | A/Tasmania/1001/2016(H3N2) | 20/7/2016 | NCBI |
| CY254481 | A/Sydney/1018/2016(H3N2) | 20/7/2016 | NCBI |
| CY254489 | A/Sydney/129/2016(H3N2) | 30/6/2016 | NCBI |
| CY254497 | A/Newcastle/58/2016(H3N2) | 10/7/2016 | NCBI |
| CY254505 | A/Newcastle/64/2016(H3N2) | 7/7/2016 | NCBI |
| CY254513 | A/Victoria/508/2016(H3N2) | 23/7/2016 | NCBI |
| CY254521 | A/SouthAustralia/60/2016(H3N2) | 13/7/2016 | NCBI |
| CY254529 | A/Brisbane/1001/2016(H3N2) | 13/7/2016 | NCBI |
| CY254537 | A/Brisbane/122/2016(H3N2) | 9/6/2016 | NCBI |
| CY254545 | A/Sydney/142/2016(H3N2) | 30/6/2016 | NCBI |
| CY254553 | A/SouthAustralia/62/2016(H3N2) | 20/7/2016 | NCBI |
| CY254561 | A/Canberra/13/2016(H3N2) | 22/7/2016 | NCBI |
| CY254569 | A/Victoria/512/2016(H3N2) | 13/7/2016 | NCBI |
| CY254577 | A/Sydney/171/2016(H3N2) | 29/7/2016 | NCBI |
| CY254593 | A/Victoria/542/2016(H3N2) | 1/8/2016 | NCBI |
| CY254601 | A/Victoria/513/2016(H3N2) | 20/7/2016 | NCBI |
| CY254609 | A/Sydney/1013/2016(H3N2) | 8/7/2016 | NCBI |
| CY254617 | A/Sydney/115/2016(H3N2) | 1/7/2016 | NCBI |
| CY254625 | A/Sydney/140/2016(H3N2) | 10/7/2016 | NCBI |
| CY254633 | A/Sydney/143/2016(H3N2) | 2/7/2016 | NCBI |
| CY254641 | A/Canberra/11/2016(H3N2) | 20/7/2016 | NCBI |
| CY254649 | A/Perth/22/2016(H3N2) | 15/4/2016 | NCBI |
| CY254657 | A/Newcastle/78/2016(H3N2) | 25/7/2016 | NCBI |
| CY254665 | A/Victoria/42/2016(H3N2) | 20/7/2016 | NCBI |
| CY254673 | A/Newcastle/68/2016(H3N2) | 30/7/2016 | NCBI |
| CY254681 | A/Victoria/2049/2016(H3N2) | 24/7/2016 | NCBI |
| CY254689 | A/Victoria/546/2016(H3N2) | 31/7/2016 | NCBI |
| CY254697 | A/Victoria/808/2016(H3N2) | 13/8/2016 | NCBI |
| CY254705 | A/Victoria/548/2016(H3N2) | 15/8/2016 | NCBI |
| CY254713 | A/Canberra/1001/2016(H3N2) | 25/7/2016 | NCBI |
| CY254721 | A/Tasmania/21/2016(H3N2) | 12/8/2016 | NCBI |
| CY254729 | A/Brisbane/158/2016(H3N2) | 4/8/2016 | NCBI |
| CY254737 | A/Brisbane/165/2016(H3N2) | 31/7/2016 | NCBI |
| CY254745 | A/Victoria/580/2016(H3N2) | 9/8/2016 | NCBI |
| CY254753 | A/Brisbane/1004/2016(H3N2) | 25/7/2016 | NCBI |
| CY254761 | A/Victoria/52/2016(H3N2) | 30/7/2016 | NCBI |

| | | | |
|---|---|---|---|
| CY254769 | A/Victoria/70/2016(H3N2) | 12/8/2016 | NCBI |
| CY254777 | A/Newcastle/102/2016(H3N2) | 5/8/2016 | NCBI |
| CY254785 | A/Victoria/2111/2016(H3N2) | 18/8/2016 | NCBI |
| CY254793 | A/Victoria/2074/2016(H3N2) | 9/8/2016 | NCBI |
| CY254801 | A/Victoria/554/2016(H3N2) | 11/8/2016 | NCBI |
| CY254809 | A/Victoria/561/2016(H3N2) | 10/8/2016 | NCBI |
| CY254817 | A/Victoria/566/2016(H3N2) | 8/8/2016 | NCBI |
| CY254825 | A/Victoria/576/2016(H3N2) | 5/8/2016 | NCBI |
| CY254833 | A/Victoria/909/2016(H3N2) | 11/8/2016 | NCBI |
| CY254841 | A/Newcastle/91/2016(H3N2) | 4/8/2016 | NCBI |
| CY254849 | A/Victoria/63/2016(H3N2) | 8/8/2016 | NCBI |
| CY254857 | A/Victoria/2073/2016(H3N2) | 9/8/2016 | NCBI |
| CY254865 | A/Victoria/2097/2016(H3N2) | 16/8/2016 | NCBI |
| CY254873 | A/Victoria/2102/2016(H3N2) | 16/8/2016 | NCBI |
| CY254881 | A/Perth/78/2016(H3N2) | 24/7/2016 | NCBI |
| CY254889 | A/Brisbane/1014/2016(H3N2) | 16/8/2016 | NCBI |
| CY254897 | A/Victoria/582/2016(H3N2) | 17/8/2016 | NCBI |
| CY254905 | A/Victoria/589/2016(H3N2) | 21/8/2016 | NCBI |
| CY254913 | A/Sydney/266/2016(H3N2) | 13/8/2016 | NCBI |
| CY254921 | A/Victoria/2079/2016(H3N2) | 11/8/2016 | NCBI |
| CY254929 | A/Tasmania/24/2016(H3N2) | 12/8/2016 | NCBI |
| CY254937 | A/Brisbane/1008/2016(H3N2) | 9/8/2016 | NCBI |
| CY254945 | A/Brisbane/1009/2016(H3N2) | 10/8/2016 | NCBI |
| CY254953 | A/Sydney/1059/2016(H3N2) | 16/8/2016 | NCBI |
| CY254961 | A/Sydney/1065/2016(H3N2) | 15/8/2016 | NCBI |
| CY254969 | A/SouthAustralia/90/2016(H3N2) | 16/8/2016 | NCBI |
| CY254977 | A/Victoria/600/2016(H3N2) | 23/8/2016 | NCBI |
| CY254985 | A/Darwin/1002/2016(H3N2) | 15/8/2016 | NCBI |
| CY254993 | A/Victoria/71/2016(H3N2) | 22/8/2016 | NCBI |
| CY255001 | A/Newcastle/134/2016(H3N2) | 23/8/2016 | NCBI |
| CY255009 | A/Victoria/72/2016(H3N2) | 29/8/2016 | NCBI |
| CY255017 | A/Brisbane/200/2016(H3N2) | 29/8/2016 | NCBI |
| CY255025 | A/Sydney/1092/2016(H3N2) | 2/9/2016 | NCBI |
| CY255033 | A/SouthAustralia/106/2016(H3N2) | 9/9/2016 | NCBI |
| CY255041 | A/SouthAustralia/108/2016(H3N2) | 9/9/2016 | NCBI |
| CY255049 | A/SouthAustralia/118/2016(H3N2) | 12/9/2016 | NCBI |
| CY255057 | A/SouthAustralia/114/2016(H3N2) | 15/9/2016 | NCBI |

| | | | |
|---|---|---|---|
| CY255065 | A/SouthAustralia/115/2016(H3N2) | 15/9/2016 | NCBI |
| CY255073 | A/Victoria/2516/2016(H3N2) | 29/8/2016 | NCBI |
| CY255081 | A/Sydney/1094/2016(H3N2) | 8/9/2016 | NCBI |
| CY255089 | A/Victoria/1012/2016(H3N2) | 16/9/2016 | NCBI |
| CY255097 | A/SouthAustralia/1007/2016(H3N2) | 12/9/2016 | NCBI |
| CY255105 | A/Victoria/1014/2016(H3N2) | 12/9/2016 | NCBI |
| CY255113 | A/SouthAustralia/107/2016(H3N2) | 10/9/2016 | NCBI |
| CY255121 | A/SouthAustralia/112/2016(H3N2) | 8/9/2016 | NCBI |
| CY255129 | A/SouthAustralia/117/2016(H3N2) | 13/9/2016 | NCBI |
| CY255137 | A/Sydney/297/2016(H3N2) | 9/9/2016 | NCBI |
| CY255145 | A/Tasmania/141/2016(H3N2) | 26/8/2016 | NCBI |
| CY255161 | A/SouthAustralia/125/2016(H3N2) | 18/9/2016 | NCBI |
| CY255169 | A/Newcastle/187/2016(H3N2) | 10/9/2016 | NCBI |
| CY255177 | A/Newcastle/190/2016(H3N2) | 16/9/2016 | NCBI |
| CY255185 | A/Brisbane/224/2016(H3N2) | 25/9/2016 | NCBI |
| CY255193 | A/Brisbane/225/2016(H3N2) | 25/8/2016 | NCBI |
| CY255201 | A/Brisbane/231/2016(H3N2) | 7/9/2016 | NCBI |
| CY255209 | A/Darwin/1008/2016(H3N2) | 22/9/2016 | NCBI |
| CY255217 | A/Newcastle/180/2016(H3N2) | 14/9/2016 | NCBI |
| CY255225 | A/Tasmania/92/2016(H3N2) | 6/9/2016 | NCBI |
| CY255233 | A/Tasmania/97/2016(H3N2) | 19/9/2016 | NCBI |
| CY255241 | A/Victoria/109/2016(H3N2) | 2/9/2016 | NCBI |
| CY255249 | A/Victoria/123/2016(H3N2) | 10/9/2016 | NCBI |
| CY255257 | A/Newcastle/201/2016(H3N2) | 10/10/2016 | NCBI |
| CY255265 | A/Victoria/624/2016(H3N2) | 16/9/2016 | NCBI |
| CY255273 | A/Canberra/80/2016(H3N2) | 18/9/2016 | NCBI |
| CY255281 | A/Canberra/93/2016(H3N2) | 4/10/2016 | NCBI |
| CY255289 | A/Victoria/648/2016(H3N2) | 2016 | NCBI |
| CY255297 | A/Victoria/657/2016(H3N2) | 2016 | NCBI |
| CY255305 | A/Townsville/1003/2016(H3N2) | 13/10/2016 | NCBI |
| CY255313 | A/Brisbane/273/2016(H3N2) | 17/10/2016 | NCBI |
| CY255321 | A/Brisbane/285/2016(H3N2) | 20/10/2016 | NCBI |
| CY255329 | A/Brisbane/287/2016(H3N2) | 24/10/2016 | NCBI |
| CY255337 | A/Brisbane/291/2016(H3N2) | 3/11/2016 | NCBI |
| CY255345 | A/SouthAustralia/137/2016(H3N2) | 26/10/2016 | NCBI |
| CY255353 | A/SouthAustralia/138/2016(H3N2) | 27/10/2016 | NCBI |
| CY255361 | A/SouthAustralia/141/2016(H3N2) | 9/11/2016 | NCBI |

| | | | |
|---|---|---|---|
| CY255369 | A/SouthAustralia/143/2016(H3N2) | 12/11/2016 | NCBI |
| CY255377 | A/SouthAustralia/147/2016(H3N2) | 11/11/2016 | NCBI |
| CY255385 | A/SouthAustralia/148/2016(H3N2) | 11/11/2016 | NCBI |
| CY255393 | A/Victoria/668/2016(H3N2) | 1/11/2016 | NCBI |
| CY255401 | A/Brisbane/280/2016(H3N2) | 27/10/2016 | NCBI |
| CY255409 | A/Brisbane/290/2016(H3N2) | 15/10/2016 | NCBI |
| CY255417 | A/Brisbane/296/2016(H3N2) | 7/11/2016 | NCBI |
| CY255425 | A/Brisbane/298/2016(H3N2) | 29/10/2016 | NCBI |
| CY255433 | A/Brisbane/299/2016(H3N2) | 29/10/2016 | NCBI |
| CY255441 | A/Brisbane/300/2016(H3N2) | 30/10/2016 | NCBI |
| CY255449 | A/Brisbane/302/2016(H3N2) | 31/10/2016 | NCBI |
| CY255457 | A/Brisbane/303/2016(H3N2) | 23/10/2016 | NCBI |
| CY255465 | A/Victoria/660/2016(H3N2) | 26/10/2016 | NCBI |
| CY255473 | A/Victoria/663/2016(H3N2) | 9/11/2016 | NCBI |
| CY255481 | A/Victoria/670/2016(H3N2) | 11/11/2016 | NCBI |
| CY255489 | A/Victoria/146/2016(H3N2) | 6/10/2016 | NCBI |
| CY255497 | A/Victoria/150/2016(H3N2) | 17/10/2016 | NCBI |
| CY255505 | A/Tasmania/213/2016(H3N2) | 13/11/2016 | NCBI |
| CY255513 | A/SouthAustralia/172/2016(H3N2) | 14/12/2016 | NCBI |
| CY255521 | A/SouthAustralia/160/2016(H3N2) | 19/11/2016 | NCBI |
| CY255529 | A/SouthAustralia/164/2016(H3N2) | 6/12/2016 | NCBI |
| CY255537 | A/SouthAustralia/173/2016(H3N2) | 13/12/2016 | NCBI |
| CY255545 | A/Townsville/51/2016(H3N2) | 13/12/2016 | NCBI |
| CY255553 | A/Brisbane/324/2016(H3N2) | 31/12/2016 | NCBI |
| CY255561 | A/Brisbane/1/2017(H3N2) | 2/1/2017 | NCBI |
| CY255569 | A/Brisbane/3/2017(H3N2) | 4/1/2017 | NCBI |
| CY255577 | A/Perth/78A/2016(H3N2) | 29/7/2016 | NCBI |
| CY254585 | A/Victoria/539/2016(H3N2) | 26/7/2016 | NCBI |
| CY255588 | A/Oregon/19/2016(H3N2) | 19/10/2016 | NCBI |
| CY255596 | A/SouthCarolina/04/2017(H3N2) | 17/1/2017 | NCBI |
| CY255612 | A/Ohio/39/2017(H3N2) | 5/9/2017 | NCBI |
| CY255620 | A/Vermont/13/2017(H3N2) | 3/9/2017 | NCBI |
| CY255628 | A/Mississippi/20/2017(H3N2) | 3/9/2017 | NCBI |
| CY255636 | A/Mississippi/21/2017(H3N2) | 3/9/2017 | NCBI |
| CY255644 | A/Colorado/14/2017(H3N2) | 1/9/2017 | NCBI |
| CY255749 | A/Louisiana/53/2017(H3N2) | 4/10/2017 | NCBI |
| CY255757 | A/Louisiana/54/2017(H3N2) | 9/10/2017 | NCBI |

| | | | |
|---|---|---|---|
| CY255765 | A/Arkansas/22/2017(H3N2) | 8/10/2017 | NCBI |
| CY255773 | A/Arkansas/23/2017(H3N2) | 9/10/2017 | NCBI |
| CY255781 | A/Arkansas/25/2017(H3N2) | 12/10/2017 | NCBI |
| CY255789 | A/NewJersey/33/2017(H3N2) | 6/10/2017 | NCBI |
| CY255797 | A/Washington/300/2017(H3N2) | 11/9/2017 | NCBI |
| CY255805 | A/Washington/298/2017(H3N2) | 17/8/2017 | NCBI |
| CY255813 | A/Florida/72/2017(H3N2) | 6/10/2017 | NCBI |
| CY255821 | A/Florida/70/2017(H3N2) | 4/10/2017 | NCBI |
| CY255829 | A/Nevada/14/2017(H3N2) | 3/10/2017 | NCBI |
| CY255837 | A/Nevada/15/2017(H3N2) | 6/10/2017 | NCBI |
| CY255845 | A/Nevada/16/2017(H3N2) | 7/10/2017 | NCBI |
| CY255853 | A/Nevada/17/2017(H3N2) | 12/10/2017 | NCBI |
| CY255861 | A/Utah/33/2017(H3N2) | 12/10/2017 | NCBI |
| CY255869 | A/Maryland/81/2017(H3N2) | 8/10/2017 | NCBI |
| CY255877 | A/Maryland/83/2017(H3N2) | 9/10/2017 | NCBI |
| CY255885 | A/NewHampshire/23/2017(H3N2) | 12/10/2017 | NCBI |
| CY255893 | A/California/81/2017(H3N2) | 4/10/2017 | NCBI |
| CY255901 | A/California/82/2017(H3N2) | 4/10/2017 | NCBI |
| CY255909 | A/Idaho/30/2017(H3N2) | 12/9/2017 | NCBI |
| CY255917 | A/Idaho/29/2017(H3N2) | 6/9/2017 | NCBI |
| CY255925 | A/Idaho/28/2017(H3N2) | 26/8/2017 | NCBI |
| CY255933 | A/Idaho/31/2017(H3N2) | 3/10/2017 | NCBI |
| CY255941 | A/Idaho/32/2017(H3N2) | 3/10/2017 | NCBI |
| CY255949 | A/Pennsylvania/264/2017(H3N2) | 10/10/2017 | NCBI |
| CY255957 | A/Florida/71/2017(H3N2) | 4/10/2017 | NCBI |
| CY255965 | A/Florida/73/2017(H3N2) | 15/10/2017 | NCBI |
| CY255973 | A/SouthCarolina/17/2017(H3N2) | 18/10/2017 | NCBI |
| CY255981 | A/Montana/24/2017(H3N2) | 17/10/2017 | NCBI |
| CY255989 | A/Montana/23/2017(H3N2) | 15/10/2017 | NCBI |
| CY255997 | A/Montana/20/2017(H3N2) | 9/10/2017 | NCBI |
| CY256005 | A/Montana/21/2017(H3N2) | 10/10/2017 | NCBI |
| CY256013 | A/Montana/22/2017(H3N2) | 11/10/2017 | NCBI |
| CY256021 | A/Montana/25/2017(H3N2) | 18/10/2017 | NCBI |
| CY256029 | A/NewYork/35/2017(H3N2) | 13/10/2017 | NCBI |
| CY256037 | A/Arizona/53/2017(H3N2) | 2/10/2017 | NCBI |
| CY256045 | A/Arizona/54/2017(H3N2) | 9/10/2017 | NCBI |
| CY256053 | A/Arizona/52/2017(H3N2) | 25/9/2017 | NCBI |

| | | | |
|---|---|---|---|
| CY256061 | A/Oregon/15/2017(H3N2) | 28/9/2017 | NCBI |
| CY256069 | A/California/89/2017(H3N2) | 6/10/2017 | NCBI |
| CY256077 | A/California/88/2017(H3N2) | 5/10/2017 | NCBI |
| CY256085 | A/California/87/2017(H3N2) | 4/10/2017 | NCBI |
| CY256093 | A/California/90/2017(H3N2) | 13/10/2017 | NCBI |
| CY256101 | A/California/86/2017(H3N2) | 3/10/2017 | NCBI |
| CY256109 | A/California/85/2017(H3N2) | 2/10/2017 | NCBI |
| CY256117 | A/California/91/2017(H3N2) | 16/10/2017 | NCBI |
| CY256125 | A/California/92/2017(H3N2) | 17/10/2017 | NCBI |
| CY256133 | A/California/83/2017(H3N2) | 1/10/2017 | NCBI |
| CY256141 | A/California/84/2017(H3N2) | 1/10/2017 | NCBI |
| CY256149 | A/Washington/307/2017(H3N2) | 17/10/2017 | NCBI |
| CY256157 | A/Washington/306/2017(H3N2) | 14/10/2017 | NCBI |
| CY256165 | A/Washington/303/2017(H3N2) | 28/9/2017 | NCBI |
| CY256173 | A/Washington/302/2017(H3N2) | 25/9/2017 | NCBI |
| CY256181 | A/Washington/304/2017(H3N2) | 30/9/2017 | NCBI |
| CY256189 | A/Hawaii/51/2017(H3N2) | 2/10/2017 | NCBI |
| CY256197 | A/Hawaii/52/2017(H3N2) | 6/10/2017 | NCBI |
| CY256205 | A/Idaho/34/2017(H3N2) | 19/10/2017 | NCBI |
| CY256213 | A/Idaho/33/2017(H3N2) | 13/10/2017 | NCBI |
| CY256221 | A/NewMexico/23/2017(H3N2) | 17/10/2017 | NCBI |
| CY256229 | A/NewMexico/24/2017(H3N2) | 17/10/2017 | NCBI |
| CY256237 | A/Alaska/65/2017(H3N2) | 12/10/2017 | NCBI |
| CY256245 | A/Wyoming/22/2017(H3N2) | 22/10/2017 | NCBI |
| CY256253 | A/Wyoming/21/2017(H3N2) | 5/10/2017 | NCBI |
| CY256261 | A/California/102/2017(H3N2) | 21/10/2017 | NCBI |
| CY256269 | A/California/96/2017(H3N2) | 18/10/2017 | NCBI |
| CY256277 | A/California/97/2017(H3N2) | 20/10/2017 | NCBI |
| CY256285 | A/California/103/2017(H3N2) | 22/10/2017 | NCBI |
| CY256293 | A/California/104/2017(H3N2) | 22/10/2017 | NCBI |
| CY256301 | A/California/98/2017(H3N2) | 20/10/2017 | NCBI |
| CY256309 | A/California/100/2017(H3N2) | 20/10/2017 | NCBI |
| CY256317 | A/California/101/2017(H3N2) | 20/10/2017 | NCBI |
| CY256325 | A/California/95/2017(H3N2) | 14/10/2017 | NCBI |
| CY256333 | A/California/94/2017(H3N2) | 9/10/2017 | NCBI |
| CY256341 | A/California/93/2017(H3N2) | 6/10/2017 | NCBI |
| CY256349 | A/California/106/2017(H3N2) | 26/10/2017 | NCBI |

| | | | |
|---|---|---|---|
| CY256357 | A/California/107/2017(H3N2) | 26/10/2017 | NCBI |
| CY256365 | A/California/105/2017(H3N2) | 23/10/2017 | NCBI |
| CY256946 | A/Nicaragua/7473_10/2016(H3N2) | 22/12/2016 | NCBI |
| CY256954 | A/Nicaragua/8774_10/2016(H3N2) | 22/12/2016 | NCBI |
| CY256962 | A/Nicaragua/8165_10/2016(H3N2) | 12/12/2016 | NCBI |
| CY256970 | A/Nicaragua/8628_03/2016(H3N2) | 11/11/2016 | NCBI |
| CY256978 | A/Nicaragua/5893_05/2016(H3N2) | 12/11/2016 | NCBI |
| CY256986 | A/Nicaragua/6031_14/2016(H3N2) | 13/11/2016 | NCBI |
| CY256994 | A/Nicaragua/6271_08/2016(H3N2) | 12/11/2016 | NCBI |
| CY257002 | A/Nicaragua/7310_17/2016(H3N2) | 13/11/2016 | NCBI |
| CY257010 | A/Nicaragua/7547_21/2016(H3N2) | 13/11/2016 | NCBI |
| CY257018 | A/Nicaragua/7943_11/2016(H3N2) | 12/11/2016 | NCBI |
| CY257026 | A/Nicaragua/9083_05/2016(H3N2) | 12/11/2016 | NCBI |
| CY257034 | A/Nicaragua/9108_03/2016(H3N2) | 12/11/2016 | NCBI |
| CY257042 | A/Nicaragua/9479_02/2016(H3N2) | 13/11/2016 | NCBI |
| CY257050 | A/Nicaragua/4714_06/2016(H3N2) | 13/11/2016 | NCBI |
| CY257058 | A/Nicaragua/5595_13/2016(H3N2) | 14/11/2016 | NCBI |
| CY257066 | A/Nicaragua/6490_16/2016(H3N2) | 14/11/2016 | NCBI |
| CY257074 | A/Nicaragua/7400_01/2016(H3N2) | 13/11/2016 | NCBI |
| CY257082 | A/Nicaragua/8154_11/2016(H3N2) | 14/11/2016 | NCBI |
| CY257090 | A/Nicaragua/9159_03/2016(H3N2) | 12/11/2016 | NCBI |
| CY257098 | A/Nicaragua/5660_06/2016(H3N2) | 15/11/2016 | NCBI |
| CY257106 | A/Nicaragua/8605_19/2016(H3N2) | 31/10/2016 | NCBI |
| CY257114 | A/Nicaragua/9094_05/2016(H3N2) | 2/11/2016 | NCBI |
| CY257122 | A/Nicaragua/6823_22/2016(H3N2) | 3/11/2016 | NCBI |
| CY257130 | A/Nicaragua/8100_07/2016(H3N2) | 3/11/2016 | NCBI |
| CY257138 | A/Nicaragua/6600_12/2016(H3N2) | 4/11/2016 | NCBI |
| CY257146 | A/Nicaragua/5960_05/2016(H3N2) | 3/11/2016 | NCBI |
| CY257154 | A/Nicaragua/4340_06/2016(H3N2) | 4/11/2016 | NCBI |
| CY257162 | A/Nicaragua/4515_11/2016(H3N2) | 4/11/2016 | NCBI |
| CY257170 | A/Nicaragua/7777_10/2016(H3N2) | 5/11/2016 | NCBI |
| CY257178 | A/Nicaragua/8004_09/2016(H3N2) | 5/11/2016 | NCBI |
| CY257186 | A/Nicaragua/8023_04/2016(H3N2) | 6/11/2016 | NCBI |
| CY257194 | A/Nicaragua/8156_07/2016(H3N2) | 6/11/2016 | NCBI |
| CY257202 | A/Nicaragua/7243_09/2016(H3N2) | 6/11/2016 | NCBI |
| CY257210 | A/Nicaragua/7964_05/2016(H3N2) | 6/11/2016 | NCBI |
| CY257218 | A/Nicaragua/9156_03/2016(H3N2) | 7/11/2016 | NCBI |

| | | | |
|---|---|---|---|
| CY257226 | A/Nicaragua/9503_01/2016(H3N2) | 7/11/2016 | NCBI |
| CY257234 | A/Nicaragua/7748_02/2016(H3N2) | 7/11/2016 | NCBI |
| CY257242 | A/Nicaragua/8031_10/2016(H3N2) | 29/11/2016 | NCBI |
| CY257250 | A/Nicaragua/8694_11/2016(H3N2) | 29/11/2016 | NCBI |
| CY257258 | A/Nicaragua/0645_06/2016(H3N2) | 9/12/2016 | NCBI |
| CY257266 | A/Nicaragua/8396_01/2016(H3N2) | 21/11/2016 | NCBI |
| CY257274 | A/Nicaragua/8647_08/2016(H3N2) | 20/11/2016 | NCBI |
| CY257282 | A/Nicaragua/6838_08/2016(H3N2) | 23/11/2016 | NCBI |
| CY257290 | A/Nicaragua/7965_13/2016(H3N2) | 23/11/2016 | NCBI |
| CY257298 | A/Nicaragua/5978_06/2016(H3N2) | 23/11/2016 | NCBI |
| CY257306 | A/Nicaragua/8744_07/2016(H3N2) | 27/11/2016 | NCBI |
| CY257314 | A/Nicaragua/6678_06/2016(H3N2) | 29/11/2016 | NCBI |
| CY257322 | A/Nicaragua/8771_02/2016(H3N2) | 25/10/2016 | NCBI |
| CY257330 | A/Nicaragua/4444_05/2016(H3N2) | 27/10/2016 | NCBI |
| CY257338 | A/Nicaragua/9290_01/2016(H3N2) | 27/10/2016 | NCBI |
| CY257346 | A/Nicaragua/5270_04/2016(H3N2) | 28/10/2016 | NCBI |
| CY257354 | A/Nicaragua/7056_03/2016(H3N2) | 28/10/2016 | NCBI |
| CY257362 | A/Nicaragua/8711_03/2016(H3N2) | 28/10/2016 | NCBI |
| CY257370 | A/Nicaragua/9372_03/2016(H3N2) | 29/10/2016 | NCBI |
| CY257378 | A/Nicaragua/8493_03/2016(H3N2) | 30/10/2016 | NCBI |
| CY257386 | A/Nicaragua/6006_09/2016(H3N2) | 31/10/2016 | NCBI |
| CY257394 | A/Nicaragua/6252_04/2016(H3N2) | 31/10/2016 | NCBI |
| CY257402 | A/Nicaragua/6842_18/2016(H3N2) | 1/11/2016 | NCBI |
| CY257410 | A/Nicaragua/8337_09/2016(H3N2) | 31/10/2016 | NCBI |
| CY257418 | A/Nicaragua/5307_03/2016(H3N2) | 23/10/2016 | NCBI |
| CY257426 | A/Nicaragua/6487_12/2016(H3N2) | 22/10/2016 | NCBI |
| CY257434 | A/Nicaragua/8368_02/2016(H3N2) | 24/10/2016 | NCBI |
| CY257442 | A/Nicaragua/4993_07/2014(H3N2) | 5/10/2014 | NCBI |
| CY257453 | A/Montana/15/2017(H3N2) | 1/9/2017 | NCBI |
| CY257461 | A/Georgia/15/2017(H3N2) | 16/9/2017 | NCBI |
| MG279222 | A/Singapore/NHRC_RGN0208/2017(H3N2) | 3/7/2017 | NCBI |
| MG279230 | A/Singapore/NHRC_RGN0210/2017(H3N2) | 4/7/2017 | NCBI |
| MG279238 | A/Singapore/NHRC_RGN0212/2017(H3N2) | 4/7/2017 | NCBI |
| MG279246 | A/SouthCarolina/NHRC_75653/2017(H3N2) | 23/6/2017 | NCBI |
| MG279254 | A/SouthCarolina/NHRC_75690/2017(H3N2) | 1/9/2017 | NCBI |
| MG279262 | A/California/NHRC_M2921/2017(H3N2) | 20/6/2017 | NCBI |
| MG279270 | A/California/NHRC_MAP1043A/2017(H3N2) | 2/8/2017 | NCBI |

| | | | |
|---|---|---|---|
| MG279278 | A/California/NHRC_MAP4061A/2017(H3N2) | 6/7/2017 | NCBI |
| MG279294 | A/California/NHRC_CG5905/2017(H3N2) | 4/4/2017 | NCBI |
| MG279322 | A/California/NHRC_28273/2017(H3N2) | 2/10/2017 | NCBI |
| CY257549 | A/Hawaii/49/2017(H3N2) | 12/9/2017 | NCBI |
| CY257557 | A/Hawaii/50/2017(H3N2) | 23/9/2017 | NCBI |
| CY257565 | A/NewMexico/20/2017(H3N2) | 13/9/2017 | NCBI |
| CY257573 | A/NewMexico/21/2017(H3N2) | 19/9/2017 | NCBI |
| CY257581 | A/Wyoming/20/2017(H3N2) | 22/9/2017 | NCBI |
| CY257589 | A/NorthCarolina/32/2017(H3N2) | 17/9/2017 | NCBI |
| CY257597 | A/California/79/2017(H3N2) | 27/9/2017 | NCBI |
| CY257605 | A/Tennessee/58/2017(H3N2) | 20/9/2017 | NCBI |
| CY257613 | A/Missouri/29/2017(H3N2) | 24/9/2017 | NCBI |
| CY257621 | A/Alaska/63/2017(H3N2) | 25/9/2017 | NCBI |
| CY257629 | A/Alaska/64/2017(H3N2) | 25/9/2017 | NCBI |
| CY257637 | A/Indiana/25/2017(H3N2) | 27/9/2017 | NCBI |
| CY257645 | A/Minnesota/36/2017(H3N2) | 27/9/2017 | NCBI |
| CY257653 | A/Minnesota/37/2017(H3N2) | 28/9/2017 | NCBI |
| CY257661 | A/Texas/287/2017(H3N2) | 28/9/2017 | NCBI |
| CY257669 | A/Texas/286/2017(H3N2) | 24/9/2017 | NCBI |
| CY257677 | A/Montana/19/2017(H3N2) | 23/9/2017 | NCBI |
| CY257685 | A/Montana/18/2017(H3N2) | 22/9/2017 | NCBI |
| CY257693 | A/Arkansas/24/2017(H3N2) | 12/10/2017 | NCBI |
| CY257701 | A/Arkansas/26/2017(H3N2) | 12/10/2017 | NCBI |
| CY257709 | A/NewJersey/33/2017(H3N2) | 6/10/2017 | NCBI |
| CY257717 | A/Washington/300/2017(H3N2) | 11/9/2017 | NCBI |
| CY257725 | A/Florida/72/2017(H3N2) | 6/10/2017 | NCBI |
| CY257733 | A/Florida/70/2017(H3N2) | 4/10/2017 | NCBI |
| CY257741 | A/Nevada/14/2017(H3N2) | 3/10/2017 | NCBI |
| CY257749 | A/Nevada/15/2017(H3N2) | 6/10/2017 | NCBI |
| CY257757 | A/Nevada/17/2017(H3N2) | 12/10/2017 | NCBI |
| CY257765 | A/Maryland/82/2017(H3N2) | 9/10/2017 | NCBI |
| CY257773 | A/Maryland/81/2017(H3N2) | 8/10/2017 | NCBI |
| CY257781 | A/Maryland/83/2017(H3N2) | 9/10/2017 | NCBI |
| CY257789 | A/NewHampshire/23/2017(H3N2) | 12/10/2017 | NCBI |
| CY257797 | A/California/81/2017(H3N2) | 4/10/2017 | NCBI |
| CY257805 | A/California/82/2017(H3N2) | 4/10/2017 | NCBI |
| CY257813 | A/Idaho/30/2017(H3N2) | 12/9/2017 | NCBI |

| | | | |
|---|---|---|---|
| CY257821 | A/Idaho/29/2017(H3N2) | 6/9/2017 | NCBI |
| CY257829 | A/Idaho/31/2017(H3N2) | 3/10/2017 | NCBI |
| CY257837 | A/Idaho/32/2017(H3N2) | 3/10/2017 | NCBI |
| CY257845 | A/Illinois/31/2017(H3N2) | 10/9/2017 | NCBI |
| CY257853 | A/Illinois/32/2017(H3N2) | 4/10/2017 | NCBI |
| CY257861 | A/Michigan/295/2017(H3N2) | 7/10/2017 | NCBI |
| CY257869 | A/Iowa/28/2017(H3N2) | 5/10/2017 | NCBI |
| CY257877 | A/Iowa/29/2017(H3N2) | 6/10/2017 | NCBI |
| CY257885 | A/Wisconsin/331/2017(H3N2) | 6/10/2017 | NCBI |
| CY257893 | A/Wisconsin/330/2017(H3N2) | 29/9/2017 | NCBI |
| CY257901 | A/Pennsylvania/264/2017(H3N2) | 10/10/2017 | NCBI |
| CY257909 | A/Florida/71/2017(H3N2) | 4/10/2017 | NCBI |
| CY257917 | A/Florida/73/2017(H3N2) | 15/10/2017 | NCBI |
| CY257925 | A/Texas/288/2017(H3N2) | 13/10/2017 | NCBI |
| CY257933 | A/Alabama/24/2017(H3N2) | 17/10/2017 | NCBI |
| CY257941 | A/Louisiana/56/2017(H3N2) | 19/10/2017 | NCBI |
| CY257949 | A/Arkansas/27/2017(H3N2) | 20/10/2017 | NCBI |
| CY257957 | A/Arkansas/29/2017(H3N2) | 23/10/2017 | NCBI |
| CY257965 | A/Maryland/84/2017(H3N2) | 8/10/2017 | NCBI |
| CY257973 | A/Maryland/86/2017(H3N2) | 22/10/2017 | NCBI |
| CY257981 | A/Michigan/296/2017(H3N2) | 1/10/2017 | NCBI |
| CY257989 | A/Michigan/298/2017(H3N2) | 13/10/2017 | NCBI |
| CY257997 | A/Massachusetts/16/2017(H3N2) | 12/10/2017 | NCBI |
| CY258005 | A/Massachusetts/17/2017(H3N2) | 16/10/2017 | NCBI |
| CY258013 | A/Delaware/34/2017(H3N2) | 16/10/2017 | NCBI |
| CY258021 | A/Delaware/35/2017(H3N2) | 20/10/2017 | NCBI |
| CY258029 | A/Illinois/33/2017(H3N2) | 19/10/2017 | NCBI |
| CY258037 | A/Iowa/31/2017(H3N2) | 15/10/2017 | NCBI |
| CY258045 | A/Iowa/32/2017(H3N2) | 19/10/2017 | NCBI |
| CY258060 | A/Wisconsin/333/2017(H3N2) | 24/10/2017 | NCBI |
| CY258068 | A/Florida/74/2017(H3N2) | 22/10/2017 | NCBI |
| CY258076 | A/Florida/77/2017(H3N2) | 25/10/2017 | NCBI |
| CY258084 | A/NorthCarolina/36/2017(H3N2) | 23/10/2017 | NCBI |
| CY258092 | A/Ohio/43/2017(H3N2) | 10/10/2017 | NCBI |
| CY258100 | A/Ohio/46/2017(H3N2) | 15/10/2017 | NCBI |
| CY258108 | A/Ohio/44/2017(H3N2) | 10/10/2017 | NCBI |
| CY258116 | A/Texas/293/2017(H3N2) | 22/10/2017 | NCBI |

| | | | |
|---|---|---|---|
| CY258124 | A/Nebraska/12/2017(H3N2) | 3/10/2017 | NCBI |
| CY258132 | A/Nebraska/13/2017(H3N2) | 11/10/2017 | NCBI |
| CY258140 | A/Nebraska/14/2017(H3N2) | 20/10/2017 | NCBI |
| CY258148 | A/Oklahoma/25/2017(H3N2) | 4/10/2017 | NCBI |
| CY258156 | A/Oklahoma/26/2017(H3N2) | 6/10/2017 | NCBI |
| CY258164 | A/SouthDakota/29/2017(H3N2) | 9/10/2017 | NCBI |
| CY258172 | A/SouthDakota/30/2017(H3N2) | 21/10/2017 | NCBI |
| CY258180 | A/NewYork/36/2017(H3N2) | 21/10/2017 | NCBI |
| CY258188 | A/NewYork/37/2017(H3N2) | 26/10/2017 | NCBI |
| CY258196 | A/Colorado/17/2017(H3N2) | 29/10/2017 | NCBI |
| CY258204 | A/Texas/296/2017(H3N2) | 30/10/2017 | NCBI |
| CY258212 | A/Texas/297/2017(H3N2) | 30/10/2017 | NCBI |
| CY258220 | A/Nevada/18/2017(H3N2) | 14/10/2017 | NCBI |
| CY258228 | A/Nevada/20/2017(H3N2) | 16/10/2017 | NCBI |
| CY258236 | A/Virginia/32/2017(H3N2) | 21/10/2017 | NCBI |
| CY258244 | A/Virginia/33/2017(H3N2) | 25/10/2017 | NCBI |
| CY258252 | A/Indiana/29/2017(H3N2) | 1/11/2017 | NCBI |
| CY258260 | A/SouthCarolina/18/2017(H3N2) | 3/11/2017 | NCBI |
| CY258268 | A/SouthCarolina/20/2017(H3N2) | 3/11/2017 | NCBI |
| CY258276 | A/Pennsylvania/265/2017(H3N2) | 1/11/2017 | NCBI |
| CY258284 | A/Florida/78/2017(H3N2) | 27/10/2017 | NCBI |
| CY258292 | A/Florida/80/2017(H3N2) | 2/11/2017 | NCBI |
| CY258300 | A/Kansas/08/2017(H3N2) | 2/11/2017 | NCBI |
| CY258308 | A/Louisiana/57/2017(H3N2) | 28/10/2017 | NCBI |
| CY258316 | A/Louisiana/58/2017(H3N2) | 28/10/2017 | NCBI |
| CY258324 | A/Minnesota/39/2017(H3N2) | 31/10/2017 | NCBI |
| CY258332 | A/Minnesota/38/2017(H3N2) | 26/10/2017 | NCBI |
| CY258340 | A/NewYork/39/2017(H3N2) | 30/10/2017 | NCBI |
| CY258348 | A/NewYork/38/2017(H3N2) | 24/10/2017 | NCBI |
| CY258356 | A/California/114/2017(H3N2) | 26/10/2017 | NCBI |
| CY258364 | A/California/112/2017(H3N2) | 25/10/2017 | NCBI |
| CY258372 | A/California/113/2017(H3N2) | 25/10/2017 | NCBI |
| CY258380 | A/California/108/2017(H3N2) | 4/10/2017 | NCBI |
| CY258388 | A/California/109/2017(H3N2) | 5/10/2017 | NCBI |
| CY258396 | A/California/110/2017(H3N2) | 5/10/2017 | NCBI |
| CY258404 | A/Oregon/18/2017(H3N2) | 21/10/2017 | NCBI |
| CY258412 | A/Montana/26/2017(H3N2) | 18/10/2017 | NCBI |

| | | | |
|---|---|---|---|
| CY258420 | A/Montana/28/2017(H3N2) | 23/10/2017 | NCBI |
| CY258428 | A/Montana/29/2017(H3N2) | 24/10/2017 | NCBI |
| CY258436 | A/Montana/27/2017(H3N2) | 19/10/2017 | NCBI |
| CY258444 | A/Montana/30/2017(H3N2) | 30/10/2017 | NCBI |
| CY258452 | A/Montana/31/2017(H3N2) | 1/11/2017 | NCBI |
| CY258460 | A/Montana/32/2017(H3N2) | 4/11/2017 | NCBI |
| CY258959 | A/NewYork/A-WC-LVD-16-001/2016(H3N2) | 21/10/2016 | NCBI |
| CY258967 | A/NewYork/A-WC-LVD-16-002/2016(H3N2) | 26/10/2016 | NCBI |
| CY258975 | A/NewYork/A-WC-LVD-16-003/2016(H3N2) | 14/11/2016 | NCBI |
| CY258983 | A/NewYork/A-WC-LVD-16-004/2016(H3N2) | 12/12/2016 | NCBI |
| CY258991 | A/NewYork/A-WC-LVD-16-005/2016(H3N2) | 23/12/2016 | NCBI |
| CY258999 | A/NewYork/A-WC-LVD-16-006/2016(H3N2) | 27/12/2016 | NCBI |
| CY259007 | A/NewYork/A-WC-LVD-16-007/2016(H3N2) | 30/12/2016 | NCBI |
| CY259015 | A/NewYork/A-WC-LVD-17-001/2017(H3N2) | 4/1/2017 | NCBI |
| CY259023 | A/NewYork/A-WC-LVD-17-002/2017(H3N2) | 8/1/2017 | NCBI |
| CY259031 | A/NewYork/A-WC-LVD-17-003/2017(H3N2) | 13/1/2017 | NCBI |
| CY259039 | A/NewYork/A-WC-LVD-17-005/2017(H3N2) | 5/1/2017 | NCBI |
| CY259047 | A/NewYork/A-WC-LVD-17-007/2017(H3N2) | 19/1/2017 | NCBI |
| CY259055 | A/NewYork/A-WC-LVD-17-008/2017(H3N2) | 17/1/2017 | NCBI |
| CY259063 | A/NewYork/A-WC-LVD-17-009/2017(H3N2) | 19/1/2017 | NCBI |
| CY259071 | A/NewYork/A-WC-LVD-17-010/2017(H3N2) | 24/1/2017 | NCBI |
| CY259079 | A/NewYork/A-WC-LVD-17-011/2017(H3N2) | 22/1/2017 | NCBI |
| CY259087 | A/NewYork/A-WC-LVD-17-012/2017(H3N2) | 23/1/2017 | NCBI |
| CY259095 | A/NewYork/A-WC-LVD-17-013/2017(H3N2) | 23/1/2017 | NCBI |
| CY259103 | A/NewYork/A-WC-LVD-17-014/2017(H3N2) | 23/1/2017 | NCBI |
| CY259111 | A/NewYork/A-WC-LVD-17-015/2017(H3N2) | 23/1/2017 | NCBI |
| CY259119 | A/NewYork/A-WC-LVD-17-016/2017(H3N2) | 25/1/2017 | NCBI |
| CY259127 | A/NewYork/A-WC-LVD-17-017/2017(H3N2) | 20/1/2017 | NCBI |
| CY259135 | A/NewYork/A-WC-LVD-17-018/2017(H3N2) | 30/1/2017 | NCBI |
| CY259143 | A/NewYork/A-WC-LVD-17-019/2017(H3N2) | 31/1/2017 | NCBI |
| CY259151 | A/NewYork/A-WC-LVD-17-021/2017(H3N2) | 3/1/2017 | NCBI |
| CY259159 | A/NewYork/A-WC-LVD-17-022/2017(H3N2) | 31/1/2017 | NCBI |
| CY259167 | A/NewYork/A-WC-LVD-17-024/2017(H3N2) | 2/2/2017 | NCBI |
| CY259175 | A/NewYork/A-WC-LVD-17-025/2017(H3N2) | 2/2/2017 | NCBI |
| CY259183 | A/NewYork/A-WC-LVD-17-026/2017(H3N2) | 29/1/2017 | NCBI |
| CY259191 | A/NewYork/A-WC-LVD-17-027/2017(H3N2) | 7/2/2017 | NCBI |
| CY259199 | A/NewYork/A-WC-LVD-17-028/2017(H3N2) | 7/2/2017 | NCBI |

| | | | |
|---|---|---|---|
| CY259207 | A/NewYork/A-WC-LVD-17-029/2017(H3N2) | 8/2/2017 | NCBI |
| CY259215 | A/NewYork/A-WC-LVD-17-030/2017(H3N2) | 11/2/2017 | NCBI |
| CY259223 | A/NewYork/A-WC-LVD-17-031/2017(H3N2) | 7/2/2017 | NCBI |
| CY259231 | A/NewYork/A-WC-LVD-17-032/2017(H3N2) | 12/2/2017 | NCBI |
| CY259247 | A/NewYork/A-WC-LVD-17-034/2017(H3N2) | 27/2/2017 | NCBI |
| CY259255 | A/NewYork/A-WC-LVD-17-035/2017(H3N2) | 25/2/2017 | NCBI |
| CY259263 | A/NewYork/A-WC-LVD-17-036/2017(H3N2) | 5/3/2017 | NCBI |
| CY259271 | A/NewYork/A-WC-LVD-17-038/2017(H3N2) | 13/3/2017 | NCBI |
| CY259279 | A/NewYork/A-WC-LVD-17-039/2017(H3N2) | 8/3/2017 | NCBI |
| CY259287 | A/NewYork/A-WC-LVD-17-040/2017(H3N2) | 12/3/2017 | NCBI |
| CY259295 | A/NewYork/A-WC-LVD-17-041/2017(H3N2) | 17/2/2017 | NCBI |
| CY259303 | A/NewYork/A-WC-LVD-17-042/2017(H3N2) | 2/4/2017 | NCBI |
| CY259311 | A/NewYork/A-WC-LVD-17-043/2017(H3N2) | 30/3/2017 | NCBI |
| CY259319 | A/NewYork/A-WC-LVD-17-044/2017(H3N2) | 8/4/2017 | NCBI |
| CY259327 | A/NewYork/A-WC-LVD-17-045/2017(H3N2) | 8/4/2017 | NCBI |
| CY259335 | A/NewYork/A-WC-LVD-17-046/2017(H3N2) | 6/4/2017 | NCBI |
| CY259343 | A/NewYork/A-WC-LVD-17-047/2017(H3N2) | 25/4/2017 | NCBI |
| CY259351 | A/NewYork/A-WC-LVD-17-048/2017(H3N2) | 21/2/2017 | NCBI |
| CY259359 | A/NewYork/A-WC-LVD-17-049/2017(H3N2) | 23/2/2017 | NCBI |
| CY259367 | A/NewYork/A-WC-LVD-17-050/2017(H3N2) | 14/2/2017 | NCBI |
| CY259375 | A/NewYork/A-WC-LVD-17-051/2017(H3N2) | 9/1/2017 | NCBI |
| CY259383 | A/NewYork/A-WC-LVD-17-052/2017(H3N2) | 28/2/2017 | NCBI |
| CY259391 | A/NewYork/A-WC-LVD-17-053/2017(H3N2) | 24/3/2017 | NCBI |
| CY259455 | A/NewYork/A-WC-LVD-16-015/2016(H3N2) | 14/4/2016 | NCBI |
| CY259471 | A/NewYork/A-WC-LVD-16-017/2016(H3N2) | 16/3/2016 | NCBI |
| CY259487 | A/NewYork/A-WC-LVD-16-020/2016(H3N2) | 2/3/2016 | NCBI |
| CY259527 | A/NewYork/A-WC-LVD-16-025/2016(H3N2) | 20/3/2016 | NCBI |
| CY259575 | A/NewYork/A-WC-LVD-15-054/2015(H3N2) | 3/11/2015 | NCBI |
| CY259639 | A/NewYork/A-WC-LVD-16-038/2016(H3N2) | 2/2/2016 | NCBI |
| CY259775 | A/NewYork/A-WC-LVD-16-056/2016(H3N2) | 21/2/2016 | NCBI |
| CY259823 | A/NewYork/A-WC-LVD-16-064/2016(H3N2) | 17/2/2016 | NCBI |
| CY259831 | A/NewYork/A-WC-LVD-16-065/2016(H3N2) | 8/3/2016 | NCBI |
| CY259874 | A/Texas/289/2017(H3N2) | 13/10/2017 | NCBI |
| CY259882 | A/Alabama/25/2017(H3N2) | 19/10/2017 | NCBI |
| CY259890 | A/Louisiana/55/2017(H3N2) | 18/10/2017 | NCBI |
| CY259898 | A/Texas/290/2017(H3N2) | 18/10/2017 | NCBI |
| CY259906 | A/Texas/291/2017(H3N2) | 19/10/2017 | NCBI |

| | | | |
|---|---|---|---|
| CY259914 | A/Indiana/28/2017(H3N2) | 6/10/2017 | NCBI |
| CY259922 | A/Missouri/31/2017(H3N2) | 19/10/2017 | NCBI |
| CY259930 | A/Missouri/32/2017(H3N2) | 24/10/2017 | NCBI |
| CY259938 | A/Illinois/33/2017(H3N2) | 19/10/2017 | NCBI |
| CY259946 | A/Iowa/31/2017(H3N2) | 15/10/2017 | NCBI |
| CY259954 | A/Iowa/32/2017(H3N2) | 19/10/2017 | NCBI |
| CY259962 | A/Wisconsin/333/2017(H3N2) | 24/10/2017 | NCBI |
| CY259970 | A/Texas/293/2017(H3N2) | 22/10/2017 | NCBI |
| CY259978 | A/NewMexico/24/2017(H3N2) | 17/10/2017 | NCBI |
| CY259986 | A/California/96/2017(H3N2) | 18/10/2017 | NCBI |
| CY259994 | A/Iowa/36/2017(H3N2) | 28/10/2017 | NCBI |
| CY260002 | A/Iowa/37/2017(H3N2) | 2/11/2017 | NCBI |
| CY260010 | A/Wisconsin/334/2017(H3N2) | 2/11/2017 | NCBI |
| CY260018 | A/Wisconsin/335/2017(H3N2) | 7/11/2017 | NCBI |
| CY260026 | A/SouthDakota/31/2017(H3N2) | 31/10/2017 | NCBI |
| CY260034 | A/SouthDakota/32/2017(H3N2) | 31/10/2017 | NCBI |
| CY260042 | A/Michigan/299/2017(H3N2) | 3/11/2017 | NCBI |
| CY260050 | A/Michigan/300/2017(H3N2) | 5/11/2017 | NCBI |
| CY260058 | A/Texas/298/2017(H3N2) | 6/11/2017 | NCBI |
| CY260066 | A/Texas/299/2017(H3N2) | 7/11/2017 | NCBI |
| CY260074 | A/Mississippi/30/2017(H3N2) | 7/11/2017 | NCBI |
| CY260082 | A/Mississippi/31/2017(H3N2) | 8/11/2017 | NCBI |
| CY260090 | A/Illinois/34/2017(H3N2) | 3/11/2017 | NCBI |
| CY260098 | A/Illinois/35/2017(H3N2) | 7/11/2017 | NCBI |
| CY260106 | A/Arkansas/30/2017(H3N2) | 28/10/2017 | NCBI |
| CY260114 | A/Arkansas/31/2017(H3N2) | 31/10/2017 | NCBI |
| CY260122 | A/Arkansas/33/2017(H3N2) | 6/11/2017 | NCBI |
| CY260130 | A/Arkansas/34/2017(H3N2) | 6/11/2017 | NCBI |
| CY260138 | A/Oklahoma/28/2017(H3N2) | 29/10/2017 | NCBI |
| CY260146 | A/Oklahoma/29/2017(H3N2) | 1/11/2017 | NCBI |
| CY260265 | A/Nevada/25/2017(H3N2) | 6/11/2017 | NCBI |
| CY260273 | A/Nevada/24/2017(H3N2) | 3/11/2017 | NCBI |
| CY260309 | A/Utah/33/2017(H3N2) | 12/10/2017 | NCBI |
| CY260324 | A/SouthCarolina/17/2017(H3N2) | 18/10/2017 | NCBI |
| CY260332 | A/Montana/24/2017(H3N2) | 17/10/2017 | NCBI |
| CY260340 | A/Montana/22/2017(H3N2) | 11/10/2017 | NCBI |
| CY260348 | A/Montana/25/2017(H3N2) | 18/10/2017 | NCBI |

| | | | |
|---|---|---|---|
| CY260356 | A/California/86/2017(H3N2) | 3/10/2017 | NCBI |
| CY260364 | A/California/92/2017(H3N2) | 17/10/2017 | NCBI |
| CY260405 | A/Nebraska/14/2017(H3N2) | 20/10/2017 | NCBI |
| CY260413 | A/Wyoming/22/2017(H3N2) | 22/10/2017 | NCBI |
| CY260421 | A/California/104/2017(H3N2) | 22/10/2017 | NCBI |
| CY260449 | A/California/107/2017(H3N2) | 26/10/2017 | NCBI |
| CY260457 | A/Texas/296/2017(H3N2) | 30/10/2017 | NCBI |
| CY260465 | A/Virginia/33/2017(H3N2) | 25/10/2017 | NCBI |
| CY260480 | A/Kansas/08/2017(H3N2) | 2/11/2017 | NCBI |
| CY260488 | A/Minnesota/39/2017(H3N2) | 31/10/2017 | NCBI |
| CY260496 | A/NewYork/39/2017(H3N2) | 30/10/2017 | NCBI |
| CY260504 | A/Delaware/38/2017(H3N2) | 1/11/2017 | NCBI |
| CY260519 | A/Maryland/89/2017(H3N2) | 3/11/2017 | NCBI |
| CY260527 | A/Ohio/47/2017(H3N2) | 3/11/2017 | NCBI |
| CY260535 | A/Iowa/37/2017(H3N2) | 2/11/2017 | NCBI |
| CY260543 | A/Washington/308/2017(H3N2) | 18/10/2017 | NCBI |
| CY260551 | A/Wyoming/24/2017(H3N2) | 28/10/2017 | NCBI |
| CY260559 | A/Wyoming/25/2017(H3N2) | 6/11/2017 | NCBI |
| CY260567 | A/Texas/299/2017(H3N2) | 7/11/2017 | NCBI |
| CY260575 | A/Mississippi/30/2017(H3N2) | 7/11/2017 | NCBI |
| CY260583 | A/Connecticut/32/2017(H3N2) | 28/10/2017 | NCBI |
| CY260591 | A/Connecticut/33/2017(H3N2) | 30/10/2017 | NCBI |
| CY260606 | A/Florida/82/2017(H3N2) | 30/10/2017 | NCBI |
| CY260614 | A/WestVirginia/23/2017(H3N2) | 26/10/2017 | NCBI |
| CY260622 | A/WestVirginia/24/2017(H3N2) | 29/10/2017 | NCBI |
| CY260630 | A/Idaho/35/2017(H3N2) | 28/10/2017 | NCBI |
| CY260638 | A/Idaho/36/2017(H3N2) | 5/11/2017 | NCBI |
| CY260646 | A/Nevada/21/2017(H3N2) | 2/11/2017 | NCBI |
| CY260654 | A/Nevada/23/2017(H3N2) | 4/11/2017 | NCBI |
| CY260662 | A/Colorado/18/2017(H3N2) | 2/11/2017 | NCBI |
| CY260670 | A/Colorado/19/2017(H3N2) | 8/11/2017 | NCBI |
| CY260678 | A/NewMexico/25/2017(H3N2) | 4/11/2017 | NCBI |
| CY260686 | A/NewMexico/26/2017(H3N2) | 6/11/2017 | NCBI |
| CY260694 | A/Utah/34/2017(H3N2) | 4/11/2017 | NCBI |
| CY260702 | A/Utah/35/2017(H3N2) | 8/11/2017 | NCBI |
| CY260710 | A/Arizona/59/2017(H3N2) | 19/10/2017 | NCBI |
| CY260718 | A/Arizona/62/2017(H3N2) | 30/10/2017 | NCBI |

| | | | |
|---|---|---|---|
| CY260726 | A/NorthCarolina/39/2017(H3N2) | 28/10/2017 | NCBI |
| CY260739 | A/NewHampshire/24/2017(H3N2) | 1/11/2017 | NCBI |
| CY260747 | A/Kansas/10/2017(H3N2) | 19/11/2017 | NCBI |
| CY260755 | A/Kansas/09/2017(H3N2) | 13/11/2017 | NCBI |
| CY260763 | A/Minnesota/40/2017(H3N2) | 5/11/2017 | NCBI |
| CY260771 | A/Minnesota/41/2017(H3N2) | 6/11/2017 | NCBI |
| CY260785 | A/Texas/302/2017(H3N2) | 15/11/2017 | NCBI |
| CY260793 | A/Texas/301/2017(H3N2) | 8/11/2017 | NCBI |
| CY260801 | A/Missouri/33/2017(H3N2) | 30/10/2017 | NCBI |
| CY260809 | A/Missouri/34/2017(H3N2) | 5/11/2017 | NCBI |
| CY260817 | A/NewYork/41/2017(H3N2) | 8/11/2017 | NCBI |
| CY260825 | A/NewYork/40/2017(H3N2) | 31/10/2017 | NCBI |
| CY260833 | A/California/118/2017(H3N2) | 5/11/2017 | NCBI |
| CY260841 | A/California/125/2017(H3N2) | 15/11/2017 | NCBI |
| CY260849 | A/California/120/2017(H3N2) | 6/11/2017 | NCBI |
| CY260857 | A/Alaska/69/2017(H3N2) | 9/11/2017 | NCBI |
| CY260865 | A/Missouri/36/2017(H3N2) | 2/11/2017 | NCBI |
| CY260873 | A/Missouri/35/2017(H3N2) | 1/11/2017 | NCBI |
| CY260881 | A/Missouri/39/2017(H3N2) | 11/11/2017 | NCBI |
| CY260889 | A/Missouri/37/2017(H3N2) | 8/11/2017 | NCBI |
| CY260897 | A/Missouri/38/2017(H3N2) | 10/11/2017 | NCBI |
| CY260905 | A/Missouri/40/2017(H3N2) | 15/11/2017 | NCBI |
| CY260913 | A/Missouri/41/2017(H3N2) | 16/11/2017 | NCBI |
| CY260921 | A/NorthDakota/23/2017(H3N2) | 10/11/2017 | NCBI |
| CY260929 | A/NorthDakota/26/2017(H3N2) | 21/11/2017 | NCBI |
| CY260937 | A/NorthDakota/24/2017(H3N2) | 19/11/2017 | NCBI |
| CY260945 | A/NorthDakota/25/2017(H3N2) | 20/11/2017 | NCBI |
| CY260953 | A/Maryland/90/2017(H3N2) | 12/11/2017 | NCBI |
| CY260961 | A/Maryland/91/2017(H3N2) | 17/11/2017 | NCBI |
| CY260969 | A/Delaware/39/2017(H3N2) | 13/11/2017 | NCBI |
| CY260977 | A/Delaware/40/2017(H3N2) | 14/11/2017 | NCBI |
| CY260985 | A/NewHampshire/25/2017(H3N2) | 17/11/2017 | NCBI |
| CY260993 | A/NewHampshire/26/2017(H3N2) | 21/11/2017 | NCBI |
| CY261001 | A/Montana/34/2017(H3N2) | 11/11/2017 | NCBI |
| CY261009 | A/Montana/33/2017(H3N2) | 9/11/2017 | NCBI |
| CY261017 | A/Montana/35/2017(H3N2) | 14/11/2017 | NCBI |
| CY261025 | A/Connecticut/36/2017(H3N2) | 19/11/2017 | NCBI |

| | | | |
|---|---|---|---|
| CY261033 | A/Connecticut/34/2017(H3N2) | 15/11/2017 | NCBI |
| CY261041 | A/Florida/91/2017(H3N2) | 12/11/2017 | NCBI |
| CY261049 | A/Florida/92/2017(H3N2) | 12/11/2017 | NCBI |
| CY261057 | A/Massachusetts/19/2017(H3N2) | 15/11/2017 | NCBI |
| CY261065 | A/Massachusetts/18/2017(H3N2) | 16/10/2017 | NCBI |
| CY261073 | A/Iowa/39/2017(H3N2) | 14/11/2017 | NCBI |
| CY261081 | A/Iowa/40/2017(H3N2) | 15/11/2017 | NCBI |
| CY261089 | A/Arkansas/35/2017(H3N2) | 12/11/2017 | NCBI |
| CY261097 | A/Arkansas/36/2017(H3N2) | 13/11/2017 | NCBI |
| CY261105 | A/Arkansas/37/2017(H3N2) | 13/11/2017 | NCBI |
| CY261113 | A/Arkansas/38/2017(H3N2) | 15/11/2017 | NCBI |
| CY261121 | A/Arkansas/39/2017(H3N2) | 16/11/2017 | NCBI |
| CY261129 | A/Arkansas/40/2017(H3N2) | 19/11/2017 | NCBI |
| CY261137 | A/Texas/304/2017(H3N2) | 17/11/2017 | NCBI |
| CY261145 | A/Texas/303/2017(H3N2) | 16/11/2017 | NCBI |
| CY261153 | A/Idaho/37/2017(H3N2) | 14/11/2017 | NCBI |
| CY261161 | A/Idaho/38/2017(H3N2) | 17/11/2017 | NCBI |
| CY261169 | A/NewMexico/27/2017(H3N2) | 13/11/2017 | NCBI |
| CY261177 | A/NewMexico/28/2017(H3N2) | 18/11/2017 | NCBI |
| CY261185 | A/Colorado/20/2017(H3N2) | 7/11/2017 | NCBI |
| CY261193 | A/Colorado/21/2017(H3N2) | 17/11/2017 | NCBI |
| CY261201 | A/Hawaii/53/2017(H3N2) | 19/10/2017 | NCBI |
| CY261209 | A/Hawaii/55/2017(H3N2) | 22/10/2017 | NCBI |
| CY261217 | A/Wisconsin/336/2017(H3N2) | 16/11/2017 | NCBI |
| CY261225 | A/Wisconsin/337/2017(H3N2) | 20/11/2017 | NCBI |
| CY261240 | A/Ohio/49/2017(H3N2) | 14/11/2017 | NCBI |
| CY261248 | A/Ohio/50/2017(H3N2) | 14/11/2017 | NCBI |
| CY261256 | A/NewYork/42/2017(H3N2) | 25/11/2017 | NCBI |
| CY261264 | A/NewYork/43/2017(H3N2) | 26/11/2017 | NCBI |
| CY261272 | A/Pennsylvania/268/2017(H3N2) | 21/11/2017 | NCBI |
| CY261280 | A/Pennsylvania/267/2017(H3N2) | 19/11/2017 | NCBI |
| CY261288 | A/Illinois/36/2017(H3N2) | 13/11/2017 | NCBI |
| CY261296 | A/Illinois/38/2017(H3N2) | 20/11/2017 | NCBI |
| CY261304 | A/Oregon/23/2017(H3N2) | 12/11/2017 | NCBI |
| CY261312 | A/Oregon/24/2017(H3N2) | 17/11/2017 | NCBI |
| CY261320 | A/Wyoming/26/2017(H3N2) | 15/11/2017 | NCBI |
| CY261328 | A/Wyoming/27/2017(H3N2) | 22/11/2017 | NCBI |

| | | | |
|---|---|---|---|
| CY261343 | A/Utah/38/2017(H3N2) | 22/11/2017 | NCBI |
| CY261351 | A/California/127/2017(H3N2) | 30/10/2017 | NCBI |
| CY261359 | A/California/126/2017(H3N2) | 28/10/2017 | NCBI |
| CY261367 | A/California/128/2017(H3N2) | 30/10/2017 | NCBI |
| CY261375 | A/California/129/2017(H3N2) | 2/11/2017 | NCBI |
| CY261383 | A/California/130/2017(H3N2) | 19/11/2017 | NCBI |
| CY261391 | A/California/131/2017(H3N2) | 23/11/2017 | NCBI |
| CY261399 | A/California/133/2017(H3N2) | 26/11/2017 | NCBI |
| CY261407 | A/California/132/2017(H3N2) | 25/11/2017 | NCBI |
| CY261415 | A/Florida/95/2017(H3N2) | 18/11/2017 | NCBI |
| CY261423 | A/Florida/96/2017(H3N2) | 19/11/2017 | NCBI |
| CY261431 | A/Oklahoma/33/2017(H3N2) | 20/11/2017 | NCBI |
| CY261439 | A/SouthDakota/34/2017(H3N2) | 19/11/2017 | NCBI |
| CY261447 | A/SouthDakota/35/2017(H3N2) | 21/11/2017 | NCBI |
| CY261455 | A/SouthDakota/33/2017(H3N2) | 17/11/2017 | NCBI |
| CY261463 | A/SouthDakota/36/2017(H3N2) | 24/11/2017 | NCBI |
| CY261471 | A/NorthDakota/27/2017(H3N2) | 22/11/2017 | NCBI |
| CY261479 | A/NorthDakota/30/2017(H3N2) | 27/11/2017 | NCBI |
| CY261487 | A/NorthDakota/29/2017(H3N2) | 26/11/2017 | NCBI |
| CY261495 | A/Nebraska/15/2017(H3N2) | 19/11/2017 | NCBI |
| CY261503 | A/Nebraska/16/2017(H3N2) | 23/11/2017 | NCBI |
| CY261511 | A/Mississippi/32/2017(H3N2) | 10/11/2017 | NCBI |
| CY261519 | A/Mississippi/34/2017(H3N2) | 27/11/2017 | NCBI |
| CY261527 | A/Louisiana/61/2017(H3N2) | 8/11/2017 | NCBI |
| CY261535 | A/Louisiana/63/2017(H3N2) | 14/11/2017 | NCBI |
| CY261543 | A/Nevada/26/2017(H3N2) | 22/11/2017 | NCBI |
| CY261551 | A/Montana/36/2017(H3N2) | 15/11/2017 | NCBI |
| CY261559 | A/Montana/37/2017(H3N2) | 25/11/2017 | NCBI |
| CY261567 | A/Montana/39/2017(H3N2) | 28/11/2017 | NCBI |
| CY261575 | A/Montana/38/2017(H3N2) | 26/11/2017 | NCBI |
| CY261583 | A/Minnesota/42/2017(H3N2) | 19/11/2017 | NCBI |
| CY261591 | A/Minnesota/43/2017(H3N2) | 27/11/2017 | NCBI |
| CY261599 | A/Missouri/42/2017(H3N2) | 25/11/2017 | NCBI |
| CY261607 | A/Missouri/43/2017(H3N2) | 25/11/2017 | NCBI |
| CY261615 | A/Indiana/30/2017(H3N2) | 27/11/2017 | NCBI |
| CY261623 | A/Indiana/31/2017(H3N2) | 28/11/2017 | NCBI |
| CY261631 | A/Alaska/71/2017(H3N2) | 24/11/2017 | NCBI |

| | | | |
|---|---|---|---|
| CY261639 | A/Alaska/72/2017(H3N2) | 24/11/2017 | NCBI |
| CY261647 | A/Alaska/73/2017(H3N2) | 27/11/2017 | NCBI |
| CY261655 | A/Alaska/74/2017(H3N2) | 29/11/2017 | NCBI |
| CY261663 | A/California/136/2017(H3N2) | 22/11/2017 | NCBI |
| CY261671 | A/Texas/305/2017(H3N2) | 27/11/2017 | NCBI |
| CY261679 | A/Texas/306/2017(H3N2) | 27/11/2017 | NCBI |
| CY263136 | A/Kentucky/27/2017(H3N2) | 27/11/2017 | NCBI |
| CY263144 | A/Kentucky/28/2017(H3N2) | 29/11/2017 | NCBI |
| CY263152 | A/NewYork/45/2017(H3N2) | 27/11/2017 | NCBI |
| CY263160 | A/NewYork/47/2017(H3N2) | 30/11/2017 | NCBI |
| CY263168 | A/Illinois/41/2017(H3N2) | 4/12/2017 | NCBI |
| CY263176 | A/Virginia/34/2017(H3N2) | 24/11/2017 | NCBI |
| CY263184 | A/Virginia/35/2017(H3N2) | 24/11/2017 | NCBI |
| CY263192 | A/NorthCarolina/41/2017(H3N2) | 30/11/2017 | NCBI |
| CY263200 | A/NorthCarolina/42/2017(H3N2) | 4/12/2017 | NCBI |
| CY264052 | A/Arizona/67/2016(H3N2) | 4/5/2016 | NCBI |
| CY264633 | A/RhodeIsland/09/2012(H3N2) | 28/11/2012 | NCBI |
| CY264641 | A/WestVirginia/20/2012(H3N2) | 20/11/2012 | NCBI |
| CY264649 | A/Iowa/03/2014(H3N2) | 6/2/2014 | NCBI |
| CY264657 | A/NewMexico/10/2014(H3N2) | 5/4/2014 | NCBI |
| CY264665 | A/Massachusetts/33/2014(H3N2) | 20/2/2014 | NCBI |
| CY264673 | A/Pennsylvania/28/2014(H3N2) | 15/4/2014 | NCBI |
| CY264681 | A/Utah/05/2014(H3N2) | 23/3/2014 | NCBI |
| CY264689 | A/Colorado/11/2014(H3N2) | 13/4/2014 | NCBI |
| CY264697 | A/Texas/22/2014(H3N2) | 28/4/2014 | NCBI |
| CY264705 | A/Maryland/23/2014(H3N2) | 16/5/2014 | NCBI |
| CY264713 | A/Michigan/08/2017(H3N2) | 11/1/2017 | NCBI |
| CY264076 | A/Missouri/44/2017(H3N2) | 2017 | NCBI |
| CY264084 | A/Arizona/63/2017(H3N2) | 2/11/2017 | NCBI |
| CY264092 | A/Arizona/66/2017(H3N2) | 7/11/2017 | NCBI |
| CY264100 | A/Nevada/28/2017(H3N2) | 14/11/2017 | NCBI |
| CY264115 | A/Delaware/42/2017(H3N2) | 4/12/2017 | NCBI |
| CY264123 | A/Arkansas/41/2017(H3N2) | 1/12/2017 | NCBI |
| CY264131 | A/Arkansas/43/2017(H3N2) | 3/12/2017 | NCBI |
| CY264139 | A/Washington/314/2017(H3N2) | 17/11/2017 | NCBI |
| CY264147 | A/Colorado/22/2017(H3N2) | 22/11/2017 | NCBI |
| CY264155 | A/Colorado/23/2017(H3N2) | 25/11/2017 | NCBI |

| | | | |
|---|---|---|---|
| CY264163 | A/SouthDakota/37/2017(H3N2) | 29/11/2017 | NCBI |
| CY264171 | A/SouthDakota/38/2017(H3N2) | 30/11/2017 | NCBI |
| CY264179 | A/Alabama/27/2017(H3N2) | 12/11/2017 | NCBI |
| CY264187 | A/Alabama/30/2017(H3N2) | 3/12/2017 | NCBI |
| CY264195 | A/Alabama/31/2017(H3N2) | 3/12/2017 | NCBI |
| CY264203 | A/Alabama/29/2017(H3N2) | 1/12/2017 | NCBI |
| CY264211 | A/Kentucky/29/2017(H3N2) | 5/12/2017 | NCBI |
| CY264219 | A/Kentucky/30/2017(H3N2) | 7/12/2017 | NCBI |
| CY264227 | A/Kentucky/32/2017(H3N2) | 8/12/2017 | NCBI |
| CY264235 | A/Kentucky/31/2017(H3N2) | 7/12/2017 | NCBI |
| CY264243 | A/Connecticut/37/2017(H3N2) | 24/11/2017 | NCBI |
| CY264251 | A/WestVirginia/27/2017(H3N2) | 3/12/2017 | NCBI |
| CY264259 | A/WestVirginia/25/2017(H3N2) | 1/12/2017 | NCBI |
| CY264267 | A/RhodeIsland/15/2017(H3N2) | 28/11/2017 | NCBI |
| CY264275 | A/Massachusetts/20/2017(H3N2) | 26/11/2017 | NCBI |
| CY264283 | A/NewMexico/29/2017(H3N2) | 30/11/2017 | NCBI |
| CY264291 | A/NewMexico/30/2017(H3N2) | 2/12/2017 | NCBI |
| CY264299 | A/Hawaii/56/2017(H3N2) | 8/11/2017 | NCBI |
| CY264307 | A/Hawaii/57/2017(H3N2) | 24/11/2017 | NCBI |
| CY264315 | A/Hawaii/58/2017(H3N2) | 25/11/2017 | NCBI |
| CY264323 | A/Hawaii/60/2017(H3N2) | 26/11/2017 | NCBI |
| CY264337 | A/NewYork/49/2017(H3N2) | 30/11/2017 | NCBI |
| CY264345 | A/Wisconsin/339/2017(H3N2) | 9/12/2017 | NCBI |
| CY264353 | A/Wisconsin/340/2017(H3N2) | 9/12/2017 | NCBI |
| CY264361 | A/Florida/97/2017(H3N2) | 29/11/2017 | NCBI |
| CY264369 | A/Florida/98/2017(H3N2) | 1/12/2017 | NCBI |
| CY264377 | A/NewHampshire/27/2017(H3N2) | 21/11/2017 | NCBI |
| CY264385 | A/NewHampshire/29/2017(H3N2) | 27/11/2017 | NCBI |
| CY264393 | A/Maryland/92/2017(H3N2) | 21/11/2017 | NCBI |
| CY264401 | A/Maryland/93/2017(H3N2) | 24/11/2017 | NCBI |
| CY264409 | A/Tennessee/59/2017(H3N2) | 24/11/2017 | NCBI |
| CY264417 | A/California/138/2017(H3N2) | 2/12/2017 | NCBI |
| CY264425 | A/California/139/2017(H3N2) | 3/12/2017 | NCBI |
| CY264433 | A/Tennessee/62/2017(H3N2) | 3/12/2017 | NCBI |
| CY264441 | A/Iowa/44/2017(H3N2) | 27/11/2017 | NCBI |
| CY264449 | A/Iowa/45/2017(H3N2) | 3/12/2017 | NCBI |
| CY264457 | A/Utah/42/2017(H3N2) | 8/12/2017 | NCBI |

| | | | |
|---|---|---|---|
| CY264465 | A/Utah/41/2017(H3N2) | 7/12/2017 | NCBI |
| CY264473 | A/Arizona/68/2017(H3N2) | 20/11/2017 | NCBI |
| CY264481 | A/Arizona/70/2017(H3N2) | 24/11/2017 | NCBI |
| CY264489 | A/Idaho/39/2017(H3N2) | 1/12/2017 | NCBI |
| CY264497 | A/Idaho/40/2017(H3N2) | 3/12/2017 | NCBI |
| CY264505 | A/Wyoming/29/2017(H3N2) | 5/12/2017 | NCBI |
| CY264513 | A/Wyoming/31/2017(H3N2) | 8/12/2017 | NCBI |
| CY264521 | A/Illinois/42/2017(H3N2) | 4/12/2017 | NCBI |
| CY264529 | A/Texas/309/2017(H3N2) | 26/11/2017 | NCBI |
| CY264537 | A/Texas/308/2017(H3N2) | 20/11/2017 | NCBI |
| CY264545 | A/SouthCarolina/21/2017(H3N2) | 30/11/2017 | NCBI |
| CY264553 | A/Pennsylvania/269/2017(H3N2) | 2/12/2017 | NCBI |
| CY264561 | A/Pennsylvania/270/2017(H3N2) | 5/12/2017 | NCBI |
| CY264569 | A/Pennsylvania/271/2017(H3N2) | 6/12/2017 | NCBI |
| CY264577 | A/Oregon/26/2017(H3N2) | 21/11/2017 | NCBI |
| CY264585 | A/Oregon/27/2017(H3N2) | 28/11/2017 | NCBI |
| CY264743 | A/Wisconsin/327/2017(H3N2) | 22/9/2017 | NCBI |
| CY266026 | A/Illinois/40/2017(H3N2) | 8/10/2017 | NCBI |
| CY266042 | A/Illinois/39/2017(H3N2) | 4/10/2017 | NCBI |
| CY266058 | A/Florida/84/2017(H3N2) | 23/10/2017 | NCBI |
| CY266074 | A/Florida/85/2017(H3N2) | 24/10/2017 | NCBI |
| CY266090 | A/Florida/87/2017(H3N2) | 25/10/2017 | NCBI |
| CY266106 | A/Florida/86/2017(H3N2) | 24/10/2017 | NCBI |
| CY266122 | A/NorthCarolina/40/2017(H3N2) | 7/10/2017 | NCBI |
| CY266138 | A/Florida/88/2017(H3N2) | 8/11/2017 | NCBI |
| CY266154 | A/Florida/89/2017(H3N2) | 9/11/2017 | NCBI |
| CY266170 | A/Florida/90/2017(H3N2) | 9/11/2017 | NCBI |
| CY266186 | A/Florida/93/2017(H3N2) | 13/11/2017 | NCBI |
| CY264766 | A/Nebraska/17/2017(H3N2) | 4/12/2017 | NCBI |
| CY264774 | A/Nebraska/18/2017(H3N2) | 7/12/2017 | NCBI |
| CY264782 | A/Arkansas/44/2017(H3N2) | 10/12/2017 | NCBI |
| CY264790 | A/Kansas/11/2017(H3N2) | 6/12/2017 | NCBI |
| CY264798 | A/Kansas/12/2017(H3N2) | 6/12/2017 | NCBI |
| CY264806 | A/Minnesota/46/2017(H3N2) | 8/12/2017 | NCBI |
| CY264814 | A/Missouri/45/2017(H3N2) | 26/11/2017 | NCBI |
| CY264822 | A/Missouri/46/2017(H3N2) | 6/12/2017 | NCBI |
| CY264830 | A/Texas/312/2017(H3N2) | 28/11/2017 | NCBI |

| | | | |
|---|---|---|---|
| CY264838 | A/Texas/313/2017(H3N2) | 7/12/2017 | NCBI |
| CY264846 | A/NewYork/50/2017(H3N2) | 7/12/2017 | NCBI |
| CY264854 | A/NewYork/51/2017(H3N2) | 8/12/2017 | NCBI |
| CY264862 | A/Montana/40/2017(H3N2) | 1/12/2017 | NCBI |
| CY264870 | A/Montana/41/2017(H3N2) | 7/12/2017 | NCBI |
| CY264878 | A/Montana/42/2017(H3N2) | 11/12/2017 | NCBI |
| CY264886 | A/Montana/43/2017(H3N2) | 14/12/2017 | NCBI |
| CY264894 | A/Ohio/51/2017(H3N2) | 11/12/2017 | NCBI |
| CY264902 | A/Ohio/52/2017(H3N2) | 12/12/2017 | NCBI |
| CY264910 | A/Ohio/53/2017(H3N2) | 14/12/2017 | NCBI |
| CY264918 | A/Florida/100/2017(H3N2) | 8/12/2017 | NCBI |
| CY264926 | A/Florida/101/2017(H3N2) | 11/12/2017 | NCBI |
| CY264934 | A/NewJersey/36/2017(H3N2) | 27/11/2017 | NCBI |
| CY264949 | A/NorthDakota/33/2017(H3N2) | 18/12/2017 | NCBI |
| CY264957 | A/NorthDakota/31/2017(H3N2) | 15/12/2017 | NCBI |
| CY264965 | A/California/140/2017(H3N2) | 12/11/2017 | NCBI |
| CY264973 | A/NorthCarolina/46/2017(H3N2) | 26/11/2017 | NCBI |
| CY264981 | A/NorthCarolina/43/2017(H3N2) | 22/11/2017 | NCBI |
| CY264989 | A/Pennsylvania/274/2017(H3N2) | 15/12/2017 | NCBI |
| CY264997 | A/Indiana/33/2017(H3N2) | 11/12/2017 | NCBI |
| CY265005 | A/Indiana/35/2017(H3N2) | 13/12/2017 | NCBI |
| CY265013 | A/Oklahoma/34/2017(H3N2) | 3/12/2017 | NCBI |
| CY265021 | A/Oklahoma/37/2017(H3N2) | 13/12/2017 | NCBI |
| CY265029 | A/Nevada/29/2017(H3N2) | 3/12/2017 | NCBI |
| CY265037 | A/Nevada/31/2017(H3N2) | 17/12/2017 | NCBI |
| CY265045 | A/Nevada/30/2017(H3N2) | 7/12/2017 | NCBI |
| CY265053 | A/Colorado/24/2017(H3N2) | 4/12/2017 | NCBI |
| CY265061 | A/Colorado/25/2017(H3N2) | 16/12/2017 | NCBI |
| CY265069 | A/Hawaii/61/2017(H3N2) | 27/11/2017 | NCBI |
| CY265077 | A/Hawaii/62/2017(H3N2) | 6/12/2017 | NCBI |
| CY265085 | A/Alaska/75/2017(H3N2) | 27/11/2017 | NCBI |
| CY265093 | A/Alaska/78/2017(H3N2) | 10/12/2017 | NCBI |
| CY265101 | A/Washington/321/2017(H3N2) | 25/11/2017 | NCBI |
| CY265109 | A/Washington/323/2017(H3N2) | 14/12/2017 | NCBI |
| CY265117 | A/Arizona/71/2017(H3N2) | 8/12/2017 | NCBI |
| CY265125 | A/Arizona/72/2017(H3N2) | 11/12/2017 | NCBI |
| CY265133 | A/NewMexico/32/2017(H3N2) | 13/12/2017 | NCBI |

| | | | |
|---|---|---|---|
| CY265141 | A/Utah/43/2017(H3N2) | 8/12/2017 | NCBI |
| CY265149 | A/Utah/44/2017(H3N2) | 9/12/2017 | NCBI |
| CY265157 | A/California/147/2017(H3N2) | 8/12/2017 | NCBI |
| CY265165 | A/California/148/2017(H3N2) | 10/12/2017 | NCBI |
| CY265173 | A/California/149/2017(H3N2) | 10/12/2017 | NCBI |
| CY265181 | A/California/152/2017(H3N2) | 12/12/2017 | NCBI |
| CY265189 | A/California/150/2017(H3N2) | 11/12/2017 | NCBI |
| CY265197 | A/California/151/2017(H3N2) | 11/12/2017 | NCBI |
| CY265205 | A/California/144/2017(H3N2) | 20/11/2017 | NCBI |
| CY265213 | A/Nevada/33/2017(H3N2) | 22/12/2017 | NCBI |
| CY266351 | A/Nicaragua/6461_14/2017(H3N2) | 15/7/2017 | NCBI |
| CY266359 | A/Nicaragua/9489_07/2017(H3N2) | 15/7/2017 | NCBI |
| CY266367 | A/Nicaragua/4769_05/2017(H3N2) | 17/7/2017 | NCBI |
| CY266375 | A/Nicaragua/5935_12/2016(H3N2) | 8/11/2016 | NCBI |
| CY266383 | A/Nicaragua/9560_01/2017(H3N2) | 15/9/2017 | NCBI |
| CY266391 | A/Nicaragua/6884_14/2017(H3N2) | 20/7/2017 | NCBI |
| CY266399 | A/Nicaragua/7577_11/2017(H3N2) | 19/8/2017 | NCBI |
| CY266407 | A/Nicaragua/9131_06/2017(H3N2) | 10/8/2017 | NCBI |
| CY266415 | A/Nicaragua/9506_04/2017(H3N2) | 28/8/2017 | NCBI |
| CY266423 | A/Nicaragua/8080_05/2017(H3N2) | 22/8/2017 | NCBI |
| CY266431 | A/Nicaragua/2010_03/2017(H3N2) | 7/8/2017 | NCBI |
| CY266439 | A/Nicaragua/8190_02/2017(H3N2) | 18/8/2017 | NCBI |
| CY266447 | A/Nicaragua/9130_04/2017(H3N2) | 18/8/2017 | NCBI |
| CY266455 | A/Nicaragua/9037_01/2017(H3N2) | 27/8/2017 | NCBI |
| CY266463 | A/Nicaragua/9076_10/2017(H3N2) | 8/8/2017 | NCBI |
| CY266471 | A/Nicaragua/8078_07/2017(H3N2) | 28/6/2017 | NCBI |
| CY266479 | A/Nicaragua/7803_01/2017(H3N2) | 15/8/2017 | NCBI |
| CY266487 | A/Nicaragua/8730_05/2017(H3N2) | 24/8/2017 | NCBI |
| CY266495 | A/Nicaragua/6186_07/2017(H3N2) | 7/9/2017 | NCBI |
| CY266503 | A/Nicaragua/6714_05/2017(H3N2) | 9/8/2017 | NCBI |
| CY266511 | A/Nicaragua/9713_01/2017(H3N2) | 11/7/2017 | NCBI |
| CY266519 | A/Nicaragua/9084_02/2017(H3N2) | 29/9/2017 | NCBI |
| CY266527 | A/Nicaragua/9526_02/2017(H3N2) | 22/7/2017 | NCBI |
| CY266535 | A/Nicaragua/6247_06/2017(H3N2) | 26/8/2017 | NCBI |
| CY266543 | A/Nicaragua/6850_13/2017(H3N2) | 2/9/2017 | NCBI |
| CY266551 | A/Nicaragua/5264_03/2017(H3N2) | 30/8/2017 | NCBI |
| CY266559 | A/Nicaragua/6802_16/2017(H3N2) | 24/8/2017 | NCBI |

| | | | |
|---|---|---|---|
| CY266567 | A/Nicaragua/9537_03/2017(H3N2) | 9/9/2017 | NCBI |
| CY266575 | A/Nicaragua/8053_08/2017(H3N2) | 2/8/2017 | NCBI |
| CY266583 | A/Nicaragua/8272_05/2017(H3N2) | 18/8/2017 | NCBI |
| CY266591 | A/Nicaragua/6267_08/2017(H3N2) | 13/7/2017 | NCBI |
| CY266599 | A/Nicaragua/9155_07/2017(H3N2) | 9/8/2017 | NCBI |
| CY266607 | A/Nicaragua/9474_05/2017(H3N2) | 11/8/2017 | NCBI |
| CY266615 | A/Nicaragua/9379_04/2017(H3N2) | 14/10/2017 | NCBI |
| CY266623 | A/Nicaragua/9522_02/2017(H3N2) | 1/8/2017 | NCBI |
| CY266631 | A/Nicaragua/8687_13/2017(H3N2) | 16/8/2017 | NCBI |
| CY266639 | A/Nicaragua/9146_04/2017(H3N2) | 21/9/2017 | NCBI |
| CY266647 | A/Nicaragua/9508_06/2017(H3N2) | 11/10/2017 | NCBI |
| CY266655 | A/Nicaragua/8727_10/2017(H3N2) | 21/7/2017 | NCBI |
| CY266663 | A/Nicaragua/6919_17/2017(H3N2) | 23/8/2017 | NCBI |
| CY266671 | A/Nicaragua/6933_28/2017(H3N2) | 28/8/2017 | NCBI |
| CY266679 | A/Nicaragua/9267_07/2017(H3N2) | 21/7/2017 | NCBI |
| CY266687 | A/Nicaragua/9116_07/2017(H3N2) | 23/8/2017 | NCBI |
| CY266695 | A/Nicaragua/9554_04/2017(H3N2) | 21/8/2017 | NCBI |
| CY266703 | A/Nicaragua/9452_07/2017(H3N2) | 21/7/2017 | NCBI |
| CY266711 | A/Nicaragua/8765_10/2017(H3N2) | 17/8/2017 | NCBI |
| CY266719 | A/Nicaragua/8689_12/2017(H3N2) | 24/8/2017 | NCBI |
| CY266727 | A/Nicaragua/9707_02/2017(H3N2) | 21/9/2017 | NCBI |
| CY266735 | A/Nicaragua/6651_11/2017(H3N2) | 31/7/2017 | NCBI |
| CY266743 | A/Nicaragua/8775_05/2017(H3N2) | 11/8/2017 | NCBI |
| CY266751 | A/Nicaragua/7859_01/2017(H3N2) | 17/8/2017 | NCBI |
| CY266759 | A/Nicaragua/7519_06/2017(H3N2) | 11/8/2017 | NCBI |
| CY266767 | A/Nicaragua/7555_08/2017(H3N2) | 26/9/2017 | NCBI |
| CY266775 | A/Nicaragua/9530_07/2017(H3N2) | 29/9/2017 | NCBI |
| CY266783 | A/Nicaragua/8540_14/2017(H3N2) | 30/7/2017 | NCBI |
| CY266791 | A/Nicaragua/9086_06/2017(H3N2) | 13/8/2017 | NCBI |
| CY266799 | A/Nicaragua/8200_02/2017(H3N2) | 15/8/2017 | NCBI |
| CY266807 | A/Nicaragua/6207_14/2017(H3N2) | 23/9/2017 | NCBI |
| CY266815 | A/Nicaragua/9828_01/2017(H3N2) | 11/8/2017 | NCBI |
| CY266823 | A/Nicaragua/8714_08/2017(H3N2) | 13/9/2017 | NCBI |
| CY266831 | A/Nicaragua/5506_06/2017(H3N2) | 11/8/2017 | NCBI |
| CY266839 | A/Nicaragua/5935_13/2017(H3N2) | 7/8/2017 | NCBI |
| CY266847 | A/Nicaragua/6363_05/2017(H3N2) | 12/8/2017 | NCBI |
| CY266855 | A/Nicaragua/8733_12/2017(H3N2) | 19/8/2017 | NCBI |

| | | | |
|---|---|---|---|
| CY266863 | A/Nicaragua/9065_11/2017(H3N2) | 28/7/2017 | NCBI |
| CY266871 | A/Nicaragua/9138_04/2017(H3N2) | 28/7/2017 | NCBI |
| CY266879 | A/Nicaragua/9872_01/2017(H3N2) | 11/9/2017 | NCBI |
| CY266887 | A/Nicaragua/8176_07/2017(H3N2) | 29/8/2017 | NCBI |
| CY266895 | A/Nicaragua/9477_07/2017(H3N2) | 13/8/2017 | NCBI |
| MG914099 | InfluenzaAvirus | 14/11/2017 | NCBI |
| CY002064 | A/NewYork/392/2004(H3N2) | 21/12/2004 | NCBI |
| AF348176 | A/HongKong/1/1968(H3N2) | 1968 | NCBI |
| AF348179 | A/HongKong/1/1968(H3N2) | 1968 | NCBI |
| CY002496 | A/Memphis/1/1971(H3N2) | 1971 | NCBI |
| CY002096 | A/Memphis/102/1972(H3N2) | 1972 | NCBI |
| DQ508833 | A/Beijing/353/1989(H3N2) | 1989 | NCBI |
| DQ508865 | A/Panama/2007/1999(H3N2) | 1999 | NCBI |
| DQ487340 | A/Panama/2007/1999(H3N2) | 1999 | NCBI |
| DQ508929 | A/Udorn/307/1972(H3N2) | 1972 | NCBI |
| DQ487341 | A/Moscow/10/1999(H3N2) | 1999 | NCBI |
| CY002424 | A/NewYork/360/2004(H3N2) | 18/11/2004 | NCBI |
| CY006707 | A/Memphis/18/1978(H3N2) | 1978 | NCBI |
| CY000001 | A/NewYork/61A/2003(H3N2) | 20/12/2003 | NCBI |
| CY000009 | A/NewYork/28/2003(H3N2) | 19/11/2003 | NCBI |
| CY000017 | A/NewYork/30/2003(H3N2) | 2/12/2003 | NCBI |
| CY000025 | A/NewYork/32/2003(H3N2) | 29/11/2003 | NCBI |
| CY000033 | A/NewYork/33/2004(H3N2) | 9/1/2004 | NCBI |
| CY000041 | A/NewYork/34/2003(H3N2) | 3/12/2003 | NCBI |
| CY000049 | A/NewYork/35/2003(H3N2) | 4/12/2003 | NCBI |
| CY000057 | A/NewYork/36/2003(H3N2) | 10/12/2003 | NCBI |
| CY000065 | A/NewYork/45/2003(H3N2) | 9/12/2003 | NCBI |
| CY000073 | A/NewYork/47/2003(H3N2) | 12/12/2003 | NCBI |
| CY000081 | A/NewYork/48/2003(H3N2) | 9/12/2003 | NCBI |
| CY000089 | A/NewYork/50/2003(H3N2) | 8/12/2003 | NCBI |
| CY000097 | A/NewYork/54/2003(H3N2) | 10/12/2003 | NCBI |
| CY000105 | A/NewYork/60A/2003(H3N2) | 15/12/2003 | NCBI |
| CY000113 | A/NewYork/110/2002(H3N2) | 29/1/2002 | NCBI |
| CY000129 | A/NewYork/16/2003(H3N2) | 11/11/2003 | NCBI |
| CY000137 | A/NewYork/26/2003(H3N2) | 2/12/2003 | NCBI |
| CY000153 | A/NewYork/41/2003(H3N2) | 8/12/2003 | NCBI |
| CY000161 | A/NewYork/42/2003(H3N2) | 8/12/2003 | NCBI |

| | | | |
|---|---|---|---|
| CY000169 | A/NewYork/43/2003(H3N2) | 10/12/2003 | NCBI |
| CY000177 | A/NewYork/44/2003(H3N2) | 8/12/2003 | NCBI |
| CY000185 | A/NewYork/83/2001(H3N2) | 15/12/2001 | NCBI |
| CY000193 | A/NewYork/23/2003(H3N2) | 16/12/2003 | NCBI |
| CY000201 | A/NewYork/84/2001(H3N2) | 29/12/2001 | NCBI |
| CY000209 | A/NewYork/91/2002(H3N2) | 17/1/2002 | NCBI |
| CY000217 | A/NewYork/95/2002(H3N2) | 26/2/2002 | NCBI |
| CY000225 | A/NewYork/100/2002(H3N2) | 22/1/2002 | NCBI |
| CY000233 | A/NewYork/113/2002(H3N2) | 31/1/2002 | NCBI |
| CY000241 | A/NewYork/121/2002(H3N2) | 26/2/2002 | NCBI |
| CY000249 | A/NewYork/19/2003(H3N2) | 18/12/2003 | NCBI |
| CY000257 | A/NewYork/52/2004(H3N2) | 5/2/2004 | NCBI |
| CY000265 | A/NewYork/53/2003(H3N2) | 11/12/2003 | NCBI |
| CY000281 | A/NewYork/86/2002(H3N2) | 1/3/2002 | NCBI |
| CY000289 | A/NewYork/92/2002(H3N2) | 18/1/2002 | NCBI |
| CY000313 | A/NewYork/129/2002(H3N2) | 12/2/2002 | NCBI |
| CY000321 | A/NewYork/131/2001(H3N2) | 23/12/2001 | NCBI |
| CY000329 | A/NewYork/133/2002(H3N2) | 6/2/2002 | NCBI |
| CY000337 | A/NewYork/135/2002(H3N2) | 14/2/2002 | NCBI |
| CY000345 | A/NewYork/15/2003(H3N2) | 12/11/2003 | NCBI |
| CY000353 | A/NewYork/21/2003(H3N2) | 26/11/2003 | NCBI |
| CY000361 | A/NewYork/22/2003(H3N2) | 2/12/2003 | NCBI |
| CY000369 | A/NewYork/31/2004(H3N2) | 5/1/2004 | NCBI |
| CY000377 | A/NewYork/49/2003(H3N2) | 18/12/2003 | NCBI |
| CY000385 | A/NewYork/85/2001(H3N2) | 31/12/2001 | NCBI |
| CY000417 | A/NewYork/103/2002(H3N2) | 25/1/2002 | NCBI |
| CY000425 | A/NewYork/128/2002(H3N2) | 25/1/2002 | NCBI |
| CY000433 | A/NewYork/132/2002(H3N2) | 27/1/2002 | NCBI |
| CY000441 | A/NewYork/134/2002(H3N2) | 4/1/2002 | NCBI |
| CY000457 | A/NewYork/149/1999(H3N2) | 29/11/1999 | NCBI |
| CY000465 | A/NewYork/150/2000(H3N2) | 28/1/2000 | NCBI |
| CY000473 | A/NewYork/2/2003(H3N2) | 27/10/2003 | NCBI |
| CY000481 | A/NewYork/71/2001(H3N2) | 22/11/2001 | NCBI |
| CY000489 | A/NewYork/106/2002(H3N2) | 20/1/2002 | NCBI |
| CY000497 | A/NewYork/108/2002(H3N2) | 31/1/2002 | NCBI |
| CY000505 | A/NewYork/4/2003(H3N2) | 8/12/2003 | NCBI |
| CY000513 | A/NewYork/11/2003(H3N2) | 31/10/2003 | NCBI |

| | | | |
|---|---|---|---|
| CY000521 | A/NewYork/29/2003(H3N2) | 12/12/2003 | NCBI |
| CY000529 | A/NewYork/109/2002(H3N2) | 30/1/2002 | NCBI |
| CY000537 | A/NewYork/114/2002(H3N2) | 5/3/2002 | NCBI |
| CY000545 | A/NewYork/115/2002(H3N2) | 7/2/2002 | NCBI |
| CY000553 | A/NewYork/116/2002(H3N2) | 30/1/2002 | NCBI |
| CY000561 | A/NewYork/69/2004(H3N2) | 6/1/2004 | NCBI |
| CY000569 | A/NewYork/80/2001(H3N2) | 26/12/2001 | NCBI |
| CY000584 | A/NewYork/127/2001(H3N2) | 31/12/2001 | NCBI |
| CY000585 | A/NewYork/130/2002(H3N2) | 11/3/2002 | NCBI |
| CY000593 | A/NewYork/147/1999(H3N2) | 23/11/1999 | NCBI |
| CY000601 | A/NewYork/151/1999(H3N2) | 4/12/1999 | NCBI |
| CY000609 | A/NewYork/154/2000(H3N2) | 24/1/2000 | NCBI |
| CY000617 | A/NewYork/161/1999(H3N2) | 16/12/1999 | NCBI |
| CY000625 | A/NewYork/120/2002(H3N2) | 19/2/2002 | NCBI |
| CY000633 | A/NewYork/143/1999(H3N2) | 26/11/1999 | NCBI |
| CY000641 | A/NewYork/163/1999(H3N2) | 22/12/1999 | NCBI |
| CY000649 | A/NewYork/167/1999(H3N2) | 30/12/1999 | NCBI |
| CY000657 | A/NewYork/169/2000(H3N2) | 3/1/2000 | NCBI |
| CY000665 | A/NewYork/170/2000(H3N2) | 5/1/2000 | NCBI |
| CY000673 | A/NewYork/171/1999(H3N2) | 30/12/1999 | NCBI |
| CY000681 | A/NewYork/172/1999(H3N2) | 26/12/1999 | NCBI |
| CY000689 | A/NewYork/173/2000(H3N2) | 2/2/2000 | NCBI |
| CY000697 | A/NewYork/174/2000(H3N2) | 4/1/2000 | NCBI |
| CY000705 | A/NewYork/175/2000(H3N2) | 6/1/2000 | NCBI |
| CY000713 | A/NewYork/176/2000(H3N2) | 1/3/2000 | NCBI |
| CY000721 | A/NewYork/177/1999(H3N2) | 25/10/1999 | NCBI |
| CY000729 | A/NewYork/179/1999(H3N2) | 24/11/1999 | NCBI |
| CY000737 | A/NewYork/180/2000(H3N2) | 9/2/2000 | NCBI |
| CY000745 | A/NewYork/185/1999(H3N2) | 25/12/1999 | NCBI |
| CY000753 | A/NewYork/192/2003(H3N2) | 4/2/2003 | NCBI |
| CY000761 | A/NewYork/10/2004(H3N2) | 30/1/2004 | NCBI |
| CY000777 | A/NewYork/38/2003(H3N2) | 21/12/2003 | NCBI |
| CY000785 | A/NewYork/46/2003(H3N2) | 9/12/2003 | NCBI |
| CY000793 | A/NewYork/117/2002(H3N2) | 1/2/2002 | NCBI |
| CY000801 | A/NewYork/141/1999(H3N2) | 9/11/1999 | NCBI |
| CY000825 | A/NewYork/157/1999(H3N2) | 15/12/1999 | NCBI |
| CY000865 | A/NewYork/193/2003(H3N2) | 19/2/2003 | NCBI |

| | | | |
|---|---|---|---|
| CY000901 | A/NewYork/13/2003(H3N2) | 12/11/2003 | NCBI |
| CY000909 | A/NewYork/14/2003(H3N2) | 3/12/2003 | NCBI |
| CY000933 | A/NewYork/111/2002(H3N2) | 4/2/2002 | NCBI |
| CY000941 | A/NewYork/119/2002(H3N2) | 29/1/2002 | NCBI |
| CY000957 | A/NewYork/59/2003(H3N2) | 16/12/2003 | NCBI |
| CY000965 | A/NewYork/64/2003(H3N2) | 26/12/2003 | NCBI |
| CY000973 | A/NewYork/67/2003(H3N2) | 29/12/2003 | NCBI |
| CY000981 | A/NewYork/160/2000(H3N2) | 3/1/2000 | NCBI |
| CY000989 | A/NewYork/138/1999(H3N2) | 3/10/1999 | NCBI |
| CY000997 | A/NewYork/165/2000(H3N2) | 18/1/2000 | NCBI |
| CY001005 | A/NewYork/189/1999(H3N2) | 20/12/1999 | NCBI |
| CY001013 | A/NewYork/198/2003(H3N2) | 8/3/2003 | NCBI |
| CY001021 | A/NewYork/1/2003(H3N2) | 28/10/2003 | NCBI |
| CY001029 | A/NewYork/6/2004(H3N2) | 6/1/2004 | NCBI |
| CY001037 | A/NewYork/7/2003(H3N2) | 31/10/2003 | NCBI |
| CY001045 | A/NewYork/8/2003(H3N2) | 29/12/2003 | NCBI |
| CY001061 | A/NewYork/18/2003(H3N2) | 1/12/2003 | NCBI |
| CY001064 | A/NewYork/51/2003(H3N2) | 10/12/2003 | NCBI |
| CY001080 | A/NewYork/99/2002(H3N2) | 23/1/2002 | NCBI |
| CY001088 | A/NewYork/24/2003(H3N2) | 1/12/2003 | NCBI |
| CY001096 | A/NewYork/39/2003(H3N2) | 4/12/2003 | NCBI |
| CY001104 | A/NewYork/101/2002(H3N2) | 4/2/2002 | NCBI |
| CY001112 | A/NewYork/27/2003(H3N2) | 28/10/2003 | NCBI |
| CY001120 | A/NewYork/137/1999(H3N2) | 4/10/1999 | NCBI |
| CY001128 | A/NewYork/105/2002(H3N2) | 5/2/2002 | NCBI |
| CY001144 | A/NewYork/107/2002(H3N2) | 16/1/2002 | NCBI |
| CY001152 | A/NewYork/74/2002(H3N2) | 19/1/2002 | NCBI |
| CY001160 | A/NewYork/202/2003(H3N2) | 5/4/2003 | NCBI |
| CY001168 | A/NewYork/94/2001(H3N2) | 25/12/2001 | NCBI |
| CY001176 | A/NewYork/181/1999(H3N2) | 5/12/1999 | NCBI |
| CY001197 | A/NewYork/122/2002(H3N2) | 21/3/2002 | NCBI |
| CY001205 | A/NewYork/56/2003(H3N2) | 16/12/2003 | NCBI |
| CY001213 | A/NewYork/63/2003(H3N2) | 29/12/2003 | NCBI |
| CY001221 | A/NewYork/65/2003(H3N2) | 29/12/2003 | NCBI |
| CY001229 | A/NewYork/70/2004(H3N2) | 6/1/2004 | NCBI |
| CY001245 | A/NewYork/162/2000(H3N2) | 9/2/2000 | NCBI |
| CY001261 | A/NewYork/81/2002(H3N2) | 19/2/2002 | NCBI |

| | | | |
|---|---|---|---|
| CY001269 | A/NewYork/164/1999(H3N2) | 27/12/1999 | NCBI |
| CY001285 | A/NewYork/9/2003(H3N2) | 31/10/2003 | NCBI |
| CY001293 | A/NewYork/37/2003(H3N2) | 4/12/2003 | NCBI |
| CY001301 | A/NewYork/75/2002(H3N2) | 12/2/2002 | NCBI |
| CY001309 | A/NewYork/97/2002(H3N2) | 13/2/2002 | NCBI |
| CY001317 | A/NewYork/102/2002(H3N2) | 23/1/2002 | NCBI |
| CY001333 | A/NewYork/118/2002(H3N2) | 30/1/2002 | NCBI |
| CY001397 | A/NewYork/156/2000(H3N2) | 5/1/2000 | NCBI |
| CY001405 | A/NewYork/213/2003(H3N2) | 22/4/2003 | NCBI |
| CY001413 | A/NewYork/263/1999(H3N2) | 10/2/1999 | NCBI |
| CY001421 | A/NewYork/20/2003(H3N2) | 1/12/2003 | NCBI |
| CY001453 | A/NewYork/184/1999(H3N2) | 20/12/1999 | NCBI |
| CY001477 | A/NewYork/240/1998(H3N2) | 28/12/1998 | NCBI |
| CY001485 | A/NewYork/249/1998(H3N2) | 12/12/1998 | NCBI |
| CY001493 | A/NewYork/250/1998(H3N2) | 12/12/1998 | NCBI |
| CY001504 | A/NewYork/247/1998(H3N2) | 2/11/1998 | NCBI |
| CY001512 | A/NewYork/58/2003(H3N2) | 16/12/2003 | NCBI |
| CY001520 | A/NewYork/142/2000(H3N2) | 6/2/2000 | NCBI |
| CY001552 | A/NewYork/215/2003(H3N2) | 22/4/2003 | NCBI |
| CY001568 | A/NewYork/224/1998(H3N2) | 28/12/1998 | NCBI |
| CY001576 | A/NewYork/253/1999(H3N2) | 26/3/1999 | NCBI |
| CY001592 | A/NewYork/259/1999(H3N2) | 2/2/1999 | NCBI |
| CY001600 | A/NewYork/262/1999(H3N2) | 25/1/1999 | NCBI |
| CY001616 | A/NewYork/265/1999(H3N2) | 29/1/1999 | NCBI |
| CY001632 | A/NewYork/268/2003(H3N2) | 27/11/2003 | NCBI |
| CY001648 | A/NewYork/270/2003(H3N2) | 18/11/2003 | NCBI |
| CY001656 | A/NewYork/284/1999(H3N2) | 8/3/1999 | NCBI |
| CY001664 | A/NewYork/286/1999(H3N2) | 19/3/1999 | NCBI |
| CY001672 | A/NewYork/287/1998(H3N2) | 29/12/1998 | NCBI |
| CY001688 | A/NewYork/248/1999(H3N2) | 2/2/1999 | NCBI |
| CY001696 | A/NewYork/255/1999(H3N2) | 10/2/1999 | NCBI |
| CY001704 | A/NewYork/257/1999(H3N2) | 3/2/1999 | NCBI |
| CY001720 | A/NewYork/273/2001(H3N2) | 29/12/2001 | NCBI |
| CY001728 | A/NewYork/274/2002(H3N2) | 26/2/2002 | NCBI |
| CY001736 | A/NewYork/276/2002(H3N2) | 7/3/2002 | NCBI |
| CY001744 | A/NewYork/277/1999(H3N2) | 8/1/1999 | NCBI |
| CY001752 | A/NewYork/279/1999(H3N2) | 23/2/1999 | NCBI |

| | | | |
|---|---|---|---|
| CY001760 | A/NewYork/280/1999(H3N2) | 12/4/1999 | NCBI |
| CY001768 | A/NewYork/282/1999(H3N2) | 11/2/1999 | NCBI |
| CY001776 | A/NewYork/288/1999(H3N2) | 26/2/1999 | NCBI |
| CY001784 | A/NewYork/289/1998(H3N2) | 4/12/1998 | NCBI |
| CY001792 | A/NewYork/290/1999(H3N2) | 17/2/1999 | NCBI |
| CY001800 | A/NewYork/304/1998(H3N2) | 30/12/1998 | NCBI |
| CY001808 | A/NewYork/311/1999(H3N2) | 1/3/1999 | NCBI |
| CY001816 | A/NewYork/313/1998(H3N2) | 28/12/1998 | NCBI |
| CY001824 | A/NewYork/314/1999(H3N2) | 3/2/1999 | NCBI |
| CY001840 | A/NewYork/317/1999(H3N2) | 4/1/1999 | NCBI |
| CY001856 | A/NewYork/320/1999(H3N2) | 26/3/1999 | NCBI |
| CY001864 | A/NewYork/321/1999(H3N2) | 17/1/1999 | NCBI |
| CY001872 | A/NewYork/323/1999(H3N2) | 25/1/1999 | NCBI |
| CY001880 | A/NewYork/327/1999(H3N2) | 24/3/1999 | NCBI |
| CY001888 | A/NewYork/329/1999(H3N2) | 1/2/1999 | NCBI |
| CY001896 | A/NewYork/333/1999(H3N2) | 16/2/1999 | NCBI |
| CY001904 | A/NewYork/335/1999(H3N2) | 20/1/1999 | NCBI |
| CY001912 | A/NewYork/336/1999(H3N2) | 24/2/1999 | NCBI |
| CY001920 | A/NewYork/337/1999(H3N2) | 8/3/1999 | NCBI |
| CY001928 | A/NewYork/338/1999(H3N2) | 8/2/1999 | NCBI |
| CY001936 | A/NewYork/340/1999(H3N2) | 23/1/1999 | NCBI |
| CY001968 | A/NewYork/278/1999(H3N2) | 4/2/1999 | NCBI |
| CY001976 | A/NewYork/324/1999(H3N2) | 26/1/1999 | NCBI |
| CY001984 | A/NewYork/331/1999(H3N2) | 28/1/1999 | NCBI |
| CY001992 | A/NewYork/332/1999(H3N2) | 17/2/1999 | NCBI |
| CY002000 | A/NewYork/359/2005(H3N2) | 17/1/2005 | NCBI |
| CY002008 | A/NewYork/361/2005(H3N2) | 2/2/2005 | NCBI |
| CY002016 | A/NewYork/378/2005(H3N2) | 3/1/2005 | NCBI |
| CY002040 | A/NewYork/386/2004(H3N2) | 22/12/2004 | NCBI |
| CY002048 | A/NewYork/387/2004(H3N2) | 27/12/2004 | NCBI |
| CY002056 | A/NewYork/391/2005(H3N2) | 7/1/2005 | NCBI |
| CY002072 | A/NewYork/396/2005(H3N2) | 4/3/2005 | NCBI |
| CY002088 | A/Memphis/66/1986(H3N2) | 1986 | NCBI |
| CY002112 | A/Memphis/59/1999(H3N2) | 1999 | NCBI |
| CY002128 | A/NewYork/275/2002(H3N2) | 4/1/2002 | NCBI |
| CY002136 | A/NewYork/283/1999(H3N2) | 12/2/1999 | NCBI |
| CY002160 | A/NewYork/316/1999(H3N2) | 26/1/1999 | NCBI |

| | | | |
|---|---|---|---|
| CY002168 | A/NewYork/322/1999(H3N2) | 18/3/1999 | NCBI |
| CY002176 | A/NewYork/356/2004(H3N2) | 1/12/2004 | NCBI |
| CY002184 | A/NewYork/357/2005(H3N2) | 15/2/2005 | NCBI |
| CY002192 | A/NewYork/366/2004(H3N2) | 20/12/2004 | NCBI |
| CY002208 | A/NewYork/370/2004(H3N2) | 20/12/2004 | NCBI |
| CY002216 | A/NewYork/371/2004(H3N2) | 9/12/2004 | NCBI |
| CY002224 | A/NewYork/372/2004(H3N2) | 22/12/2004 | NCBI |
| CY002232 | A/NewYork/374/2004(H3N2) | 27/12/2004 | NCBI |
| CY002248 | A/NewYork/381/2004(H3N2) | 28/12/2004 | NCBI |
| CY002256 | A/NewYork/383/2004(H3N2) | 29/12/2004 | NCBI |
| CY002272 | A/Memphis/24/1995(H3N2) | 1995 | NCBI |
| CY002296 | A/NewYork/347/1999(H3N2) | 19/1/1999 | NCBI |
| CY002312 | A/NewYork/153/1999(H3N2) | 9/12/1999 | NCBI |
| CY002328 | A/NewYork/77/2001(H3N2) | 26/12/2001 | NCBI |
| CY002336 | A/NewYork/266/1999(H3N2) | 6/5/1999 | NCBI |
| CY002344 | A/NewYork/272/2003(H3N2) | 28/11/2003 | NCBI |
| CY002368 | A/NewYork/325/1999(H3N2) | 26/2/1999 | NCBI |
| CY002376 | A/NewYork/328/1998(H3N2) | 28/12/1998 | NCBI |
| CY002384 | A/NewYork/330/1998(H3N2) | 30/12/1998 | NCBI |
| CY002432 | A/NewYork/362/2004(H3N2) | 8/12/2004 | NCBI |
| CY002440 | A/NewYork/365/2004(H3N2) | 16/12/2004 | NCBI |
| CY002448 | A/NewYork/367/2005(H3N2) | 16/2/2005 | NCBI |
| CY002456 | A/NewYork/373/2005(H3N2) | 22/2/2005 | NCBI |
| CY002464 | A/NewYork/384/2005(H3N2) | 6/1/2005 | NCBI |
| CY002488 | A/NewYork/393/2005(H3N2) | 23/3/2005 | NCBI |
| CY002520 | A/NewYork/204/2003(H3N2) | 16/8/2003 | NCBI |
| CY002552 | A/NewYork/252/1999(H3N2) | 12/1/1999 | NCBI |
| CY002560 | A/NewYork/261/1999(H3N2) | 9/4/1999 | NCBI |
| CY002576 | A/NewYork/339/1999(H3N2) | 9/2/1999 | NCBI |
| CY002592 | A/NewYork/377/2004(H3N2) | 20/12/2004 | NCBI |
| CY002712 | A/NewYork/319/2004(H3N2) | 8/11/2004 | NCBI |
| CY002720 | A/NewYork/352/2005(H3N2) | 4/1/2005 | NCBI |
| CY002728 | A/NewYork/363/2004(H3N2) | 15/12/2004 | NCBI |
| CY002736 | A/NewYork/394/2005(H3N2) | 4/1/2005 | NCBI |
| CY002744 | A/Memphis/109/1972(H3N2) | 1972 | NCBI |
| CY002784 | A/NewYork/232/2004(H3N2) | 9/12/2004 | NCBI |
| CY002816 | A/NewYork/301/2001(H3N2) | 21/1/2001 | NCBI |

| | | | |
|---|---|---|---|
| CY002904 | A/Christchurch/339/2004(H3N2) | 24/10/2004 | NCBI |
| CY002905 | A/Christchurch/10/2004(H3N2) | 20/7/2004 | NCBI |
| CY002906 | A/Christchurch/13/2004(H3N2) | 20/7/2004 | NCBI |
| CY002914 | A/Christchurch/15/2004(H3N2) | 12/8/2004 | NCBI |
| CY002922 | A/Christchurch/14/2004(H3N2) | 7/8/2004 | NCBI |
| CY002930 | A/Christchurch/89/2004(H3N2) | 3/9/2004 | NCBI |
| CY002938 | A/Christchurch/90/2004(H3N2) | 7/9/2004 | NCBI |
| CY002946 | A/Christchurch/184/2004(H3N2) | 22/9/2004 | NCBI |
| CY002954 | A/Ashburton/280/2004(H3N2) | 12/10/2004 | NCBI |
| CY002962 | A/Christchurch/297/2004(H3N2) | 13/10/2004 | NCBI |
| CY003032 | A/NewYork/351/2004(H3N2) | 6/11/2004 | NCBI |
| CY003056 | A/NewYork/395/2005(H3N2) | 24/1/2005 | NCBI |
| CY003072 | A/NewYork/123/2004(H3N2) | 16/12/2004 | NCBI |
| CY003080 | A/NewYork/401/2001(H3N2) | 21/11/2001 | NCBI |
| CY003088 | A/NewYork/402/2001(H3N2) | 27/12/2001 | NCBI |
| CY003096 | A/NewYork/403/2002(H3N2) | 4/1/2002 | NCBI |
| CY003104 | A/NewYork/404/2002(H3N2) | 5/1/2002 | NCBI |
| CY003120 | A/NewYork/406/2002(H3N2) | 8/1/2002 | NCBI |
| CY003123 | A/NewYork/408/2002(H3N2) | 11/1/2002 | NCBI |
| CY003136 | A/NewYork/409/2002(H3N2) | 14/1/2002 | NCBI |
| CY003144 | A/NewYork/410/2002(H3N2) | 15/1/2002 | NCBI |
| CY003152 | A/NewYork/411/2002(H3N2) | 22/1/2002 | NCBI |
| CY003160 | A/NewYork/412/2002(H3N2) | 1/2/2002 | NCBI |
| CY003168 | A/NewYork/413/2002(H3N2) | 8/2/2002 | NCBI |
| CY003176 | A/NewYork/414/2002(H3N2) | 12/2/2002 | NCBI |
| CY003184 | A/NewYork/415/2002(H3N2) | 22/2/2002 | NCBI |
| CY003192 | A/NewYork/416/2002(H3N2) | 28/2/2002 | NCBI |
| CY003200 | A/NewYork/418/2002(H3N2) | 16/1/2002 | NCBI |
| CY003208 | A/NewYork/419/2002(H3N2) | 18/1/2002 | NCBI |
| CY003240 | A/NewYork/431/2000(H3N2) | 3/1/2000 | NCBI |
| CY003248 | A/NewYork/433/2000(H3N2) | 5/1/2000 | NCBI |
| CY003256 | A/NewYork/434/2000(H3N2) | 9/1/2000 | NCBI |
| CY003264 | A/NewYork/436/2000(H3N2) | 10/1/2000 | NCBI |
| CY003272 | A/NewYork/437/2000(H3N2) | 12/1/2000 | NCBI |
| CY003280 | A/NewYork/438/2000(H3N2) | 18/1/2000 | NCBI |
| CY003408 | A/NewYork/236/2004(H3N2) | 27/12/2004 | NCBI |
| CY003416 | A/NewYork/237/2004(H3N2) | 16/12/2004 | NCBI |

| | | | |
|---|---|---|---|
| CY003424 | A/NewYork/407/2002(H3N2) | 8/1/2002 | NCBI |
| CY003432 | A/NewYork/421/1999(H3N2) | 31/12/1999 | NCBI |
| CY003440 | A/NewYork/425/1999(H3N2) | 17/12/1999 | NCBI |
| CY003448 | A/NewYork/430/2000(H3N2) | 3/1/2000 | NCBI |
| CY003456 | A/NewYork/439/2000(H3N2) | 23/1/2000 | NCBI |
| CY003488 | A/HongKong/46/1980(H3N2) | 1980 | NCBI |
| CY003496 | A/HongKong/14/1974(H3N2) | 1974 | NCBI |
| CY003504 | A/HongKong/6/1985(H3N2) | 1985 | NCBI |
| CY003512 | A/HongKong/2/1988(H3N2) | 1988 | NCBI |
| CY003520 | A/HongKong/24/1985(H3N2) | 1985 | NCBI |
| CY003528 | A/HongKong/11/1973(H3N2) | 1973 | NCBI |
| CY003536 | A/HongKong/7/1985(H3N2) | 1985 | NCBI |
| CY003544 | A/HongKong/7/1987(H3N2) | 1987 | NCBI |
| CY003552 | A/HongKong/6/1972(H3N2) | 1972 | NCBI |
| CY003560 | A/NewYork/448/1998(H3N2) | 23/11/1998 | NCBI |
| CY003568 | A/NewYork/449/1999(H3N2) | 4/1/1999 | NCBI |
| CY003576 | A/NewYork/450/1999(H3N2) | 8/1/1999 | NCBI |
| CY003584 | A/NewYork/451/1999(H3N2) | 11/1/1999 | NCBI |
| CY003592 | A/NewYork/453/1999(H3N2) | 12/1/1999 | NCBI |
| CY003600 | A/NewYork/454/1999(H3N2) | 13/1/1999 | NCBI |
| CY003608 | A/NewYork/456/1999(H3N2) | 15/1/1999 | NCBI |
| CY003616 | A/NewYork/457/1999(H3N2) | 15/1/1999 | NCBI |
| CY003624 | A/NewYork/458/1999(H3N2) | 22/1/1999 | NCBI |
| CY003632 | A/NewYork/459/1999(H3N2) | 25/1/1999 | NCBI |
| CY003640 | A/NewYork/463/2005(H3N2) | 9/2/2005 | NCBI |
| CY003648 | A/NewYork/464/2005(H3N2) | 7/3/2005 | NCBI |
| CY003656 | A/NewYork/468/2004(H3N2) | 17/9/2004 | NCBI |
| CY003664 | A/NewYork/471/2004(H3N2) | 8/11/2004 | NCBI |
| CY003680 | A/NewYork/485/2003(H3N2) | 20/1/2003 | NCBI |
| CY003712 | A/HongKong/14/1992(H3N2) | 1992 | NCBI |
| CY003720 | A/HongKong/26/1983(H3N2) | 1983 | NCBI |
| CY003728 | A/HongKong/43/1975(H3N2) | 1975 | NCBI |
| CY003736 | A/HongKong/5/1983(H3N2) | 1983 | NCBI |
| CY003744 | A/HongKong/7/1984(H3N2) | 1984 | NCBI |
| CY003777 | A/NewYork/420/2002(H3N2) | 28/1/2002 | NCBI |
| CY003817 | A/NewYork/432/2000(H3N2) | 5/1/2000 | NCBI |
| CY003825 | A/NewYork/440/2000(H3N2) | 24/1/2000 | NCBI |

| | | | |
|---|---|---|---|
| CY006043 | A/HongKong/45/1980(H3N2) | 1980 | NCBI |
| CY006044 | A/Beijing/39/1975(H3N2) | 1975 | NCBI |
| CY006052 | A/HongKong/1/1982(H3N2) | 1982 | NCBI |
| CY006060 | A/NewYork/452/1999(H3N2) | 11/1/1999 | NCBI |
| CY006068 | A/NewYork/455/1999(H3N2) | 14/1/1999 | NCBI |
| CY006076 | A/NewYork/461/2005(H3N2) | 2/2/2005 | NCBI |
| CY006084 | A/NewYork/465/2005(H3N2) | 8/3/2005 | NCBI |
| CY006092 | A/NewYork/470/2004(H3N2) | 27/9/2004 | NCBI |
| CY006227 | A/NewYork/503/1997(H3N2) | 29/11/1997 | NCBI |
| CY006235 | A/NewYork/505/1997(H3N2) | 2/12/1997 | NCBI |
| CY006243 | A/NewYork/508/1997(H3N2) | 22/12/1997 | NCBI |
| CY006251 | A/NewYork/510/1998(H3N2) | 20/2/1998 | NCBI |
| CY006259 | A/NewYork/511/1997(H3N2) | 17/12/1997 | NCBI |
| CY006267 | A/NewYork/513/1997(H3N2) | 24/12/1997 | NCBI |
| CY006275 | A/NewYork/515/1997(H3N2) | 21/12/1997 | NCBI |
| CY006283 | A/NewYork/517/1998(H3N2) | 3/1/1998 | NCBI |
| CY006291 | A/NewYork/462/2005(H3N2) | 8/2/2005 | NCBI |
| CY006299 | A/HongKong/3/1969(H3N2) | 1969 | NCBI |
| CY006315 | A/HongKong/14/1983(H3N2) | 1983 | NCBI |
| CY006323 | A/HongKong/4/1984(H3N2) | 1984 | NCBI |
| CY006339 | A/Nanchang/0058/1994(H3N2) | 1994 | NCBI |
| CY006347 | A/Nanchang/58/1993(H3N2) | 1993 | NCBI |
| CY006371 | A/NewYork/466/2004(H3N2) | 27/11/2004 | NCBI |
| CY006379 | A/NewYork/467/2004(H3N2) | 17/9/2004 | NCBI |
| CY006459 | A/NewYork/512/1998(H3N2) | 29/1/1998 | NCBI |
| CY006467 | A/NewYork/516/1997(H3N2) | 26/12/1997 | NCBI |
| CY006539 | A/NewYork/528/1998(H3N2) | 8/1/1998 | NCBI |
| CY006571 | A/NewYork/534/1998(H3N2) | 29/1/1998 | NCBI |
| CY006579 | A/NewYork/539/1998(H3N2) | 10/2/1998 | NCBI |
| CY006659 | A/NewYork/435/2000(H3N2) | 9/1/2000 | NCBI |
| CY006683 | A/HongKong/46/1971(H3N2) | 1971 | NCBI |
| CY006691 | A/Memphis/2/1978(H3N2) | 1978 | NCBI |
| CY006699 | A/Memphis/12/1978(H3N2) | 1978 | NCBI |
| CY006715 | A/Memphis/101/1974(H3N2) | 1974 | NCBI |
| CY006723 | A/Memphis/137/1976(H3N2) | 1976 | NCBI |
| CY006731 | A/Memphis/1/1977(H3N2) | 1977 | NCBI |
| CY006739 | A/Memphis/3/1977(H3N2) | 1977 | NCBI |

| | | | |
|---|---|---|---|
| CY006755 | A/Nanjing/2/1982(H3N2) | 1982 | NCBI |
| CY006787 | A/NewYork/506/1998(H3N2) | 2/2/1998 | NCBI |
| CY006811 | A/Memphis/3/1973(H3N2) | 1973 | NCBI |
| CY006827 | A/Memphis/103/1974(H3N2) | 1974 | NCBI |
| CY006835 | A/Memphis/110/1976(H3N2) | 1976 | NCBI |
| CY006851 | A/Nanjing/36/1983(H3N2) | 1983 | NCBI |
| CY006883 | A/Memphis/103/1976(H3N2) | 1976 | NCBI |
| CY006891 | A/Memphis/9/1980(H3N2) | 1980 | NCBI |
| CY006899 | A/NewYork/460/1999(H3N2) | 25/1/1999 | NCBI |
| CY006907 | A/HongKong/49/1974(H3N2) | 1974 | NCBI |
| CY006923 | A/Canterbury/382/2003(H3N2) | 14/7/2003 | NCBI |
| CY006931 | A/Canterbury/384/2003(H3N2) | 15/7/2003 | NCBI |
| CY006939 | A/Canterbury/386/2003(H3N2) | 16/7/2003 | NCBI |
| CY006955 | A/Canterbury/390/2003(H3N2) | 8/7/2003 | NCBI |
| CY006963 | A/Canterbury/391/2003(H3N2) | 19/8/2003 | NCBI |
| CY006971 | A/Canterbury/392/2003(H3N2) | 17/8/2003 | NCBI |
| CY006979 | A/Canterbury/393/2003(H3N2) | 16/6/2003 | NCBI |
| CY006987 | A/Canterbury/394/2003(H3N2) | 30/6/2003 | NCBI |
| CY006995 | A/Canterbury/395/2003(H3N2) | 26/7/2003 | NCBI |
| CY007003 | A/Canterbury/397/2003(H3N2) | 18/8/2003 | NCBI |
| CY007011 | A/Canterbury/398/2003(H3N2) | 14/8/2003 | NCBI |
| CY007035 | A/Canterbury/403/2003(H3N2) | 5/7/2003 | NCBI |
| CY007043 | A/Canterbury/404/2003(H3N2) | 5/7/2003 | NCBI |
| CY007051 | A/Canterbury/405/2003(H3N2) | 5/7/2003 | NCBI |
| CY007059 | A/Canterbury/406/2003(H3N2) | 5/7/2003 | NCBI |
| CY007075 | A/Canterbury/410/2003(H3N2) | 6/7/2003 | NCBI |
| CY007083 | A/Canterbury/411/2003(H3N2) | 15/7/2003 | NCBI |
| CY007091 | A/Canterbury/412/2003(H3N2) | 15/7/2003 | NCBI |
| CY007099 | A/Canterbury/416/2003(H3N2) | 15/7/2003 | NCBI |
| CY007115 | A/Canterbury/418/2003(H3N2) | 15/7/2003 | NCBI |
| CY007123 | A/Canterbury/420/2003(H3N2) | 16/7/2003 | NCBI |
| CY007131 | A/Canterbury/423/2003(H3N2) | 14/7/2003 | NCBI |
| CY007139 | A/Canterbury/424/2003(H3N2) | 15/7/2003 | NCBI |
| CY007187 | A/Canterbury/434/2003(H3N2) | 14/7/2003 | NCBI |
| CY007267 | A/Canterbury/444/2003(H3N2) | 17/8/2003 | NCBI |
| CY007275 | A/Whanganui/128/2004(H3N2) | 10/9/2004 | NCBI |
| CY007291 | A/BayofPlenty/279/2004(H3N2) | 11/10/2004 | NCBI |

| | | | |
|---|---|---|---|
| CY007299 | A/BayofPlenty/332/2004(H3N2) | 17/10/2004 | NCBI |
| CY007307 | A/Tairawhiti/369/2004(H3N2) | 10/9/2004 | NCBI |
| CY007331 | A/Whanganui/417/2004(H3N2) | 10/9/2004 | NCBI |
| CY007347 | A/Canterbury/12/2004(H3N2) | 5/8/2004 | NCBI |
| CY007371 | A/Canterbury/18/2004(H3N2) | 15/8/2004 | NCBI |
| CY007379 | A/Canterbury/19/2004(H3N2) | 17/8/2004 | NCBI |
| CY007387 | A/Canterbury/20/2004(H3N2) | 18/8/2004 | NCBI |
| CY007403 | A/Canterbury/23/2004(H3N2) | 19/8/2004 | NCBI |
| CY007435 | A/Canterbury/102/2004(H3N2) | 9/9/2004 | NCBI |
| CY007443 | A/Canterbury/103/2004(H3N2) | 9/9/2004 | NCBI |
| CY007451 | A/Canterbury/104/2004(H3N2) | 7/9/2004 | NCBI |
| CY007459 | A/Canterbury/105/2004(H3N2) | 7/9/2004 | NCBI |
| CY007475 | A/Canterbury/107/2004(H3N2) | 12/9/2004 | NCBI |
| CY007483 | A/Canterbury/201/2004(H3N2) | 16/9/2004 | NCBI |
| CY007491 | A/Canterbury/202/2004(H3N2) | 15/9/2004 | NCBI |
| CY007499 | A/Canterbury/205/2004(H3N2) | 15/9/2004 | NCBI |
| CY007507 | A/Canterbury/206/2004(H3N2) | 24/9/2004 | NCBI |
| CY007531 | A/Canterbury/209/2004(H3N2) | 23/9/2004 | NCBI |
| CY007547 | A/Canterbury/303/2004(H3N2) | 25/9/2004 | NCBI |
| CY007555 | A/Canterbury/304/2004(H3N2) | 4/10/2004 | NCBI |
| CY007563 | A/Canterbury/311/2004(H3N2) | 7/10/2004 | NCBI |
| CY007579 | A/Canterbury/315/2004(H3N2) | 28/9/2004 | NCBI |
| CY007587 | A/Canterbury/01/2002(H3N2) | 26/4/2002 | NCBI |
| CY007595 | A/Canterbury/02/2002(H3N2) | 29/4/2002 | NCBI |
| CY007603 | A/Canterbury/05/2002(H3N2) | 13/4/2002 | NCBI |
| CY007651 | A/Canterbury/16/2002(H3N2) | 4/5/2002 | NCBI |
| CY007659 | A/Canterbury/18/2002(H3N2) | 6/5/2002 | NCBI |
| CY007667 | A/Canterbury/22/2002(H3N2) | 10/5/2002 | NCBI |
| CY007675 | A/Canterbury/35/2002(H3N2) | 13/6/2002 | NCBI |
| CY007683 | A/Canterbury/41/2002(H3N2) | 10/6/2002 | NCBI |
| CY007691 | A/Canterbury/47/2002(H3N2) | 8/6/2002 | NCBI |
| CY007699 | A/Canterbury/48/2002(H3N2) | 9/6/2002 | NCBI |
| CY007707 | A/Canterbury/49/2002(H3N2) | 7/6/2002 | NCBI |
| CY007715 | A/Canterbury/50/2002(H3N2) | 11/6/2002 | NCBI |
| CY007723 | A/Canterbury/53/2002(H3N2) | 10/6/2002 | NCBI |
| CY007731 | A/Canterbury/56/2002(H3N2) | 25/6/2002 | NCBI |
| CY007739 | A/Canterbury/59/2002(H3N2) | 2/7/2002 | NCBI |

| | | | |
|---|---|---|---|
| CY007747 | A/Canterbury/68/2002(H3N2) | 5/7/2002 | NCBI |
| CY007755 | A/Canterbury/69/2002(H3N2) | 6/7/2002 | NCBI |
| CY007763 | A/Canterbury/75/2002(H3N2) | 11/7/2002 | NCBI |
| CY007771 | A/Canterbury/80/2002(H3N2) | 11/7/2002 | NCBI |
| CY007779 | A/Canterbury/81/2002(H3N2) | 9/7/2002 | NCBI |
| CY007787 | A/Canterbury/102/2002(H3N2) | 23/5/2002 | NCBI |
| CY007795 | A/Canterbury/01/2005(H3N2) | 4/3/2005 | NCBI |
| CY007803 | A/Canterbury/02/2005(H3N2) | 4/3/2005 | NCBI |
| CY007811 | A/Canterbury/03/2005(H3N2) | 6/3/2005 | NCBI |
| CY007819 | A/Canterbury/409/2003(H3N2) | 6/7/2003 | NCBI |
| CY007883 | A/Canterbury/31/2002(H3N2) | 20/6/2002 | NCBI |
| CY007923 | A/Canterbury/58/2002(H3N2) | 25/6/2002 | NCBI |
| CY007939 | A/Canterbury/72/2002(H3N2) | 6/7/2002 | NCBI |
| CY007947 | A/Canterbury/79/2002(H3N2) | 9/7/2002 | NCBI |
| CY007955 | A/Canterbury/16/2005(H3N2) | 20/5/2005 | NCBI |
| CY007971 | A/Guandong/243/1972(H3N2) | 1972 | NCBI |
| CY007979 | A/NewYork/526/1997(H3N2) | 29/12/1997 | NCBI |
| CY007987 | A/Canterbury/108/2004(H3N2) | 12/9/2004 | NCBI |
| CY007995 | A/Canterbury/109/2004(H3N2) | 12/9/2004 | NCBI |
| CY008003 | A/Canterbury/04/2002(H3N2) | 12/4/2002 | NCBI |
| CY008011 | A/Canterbury/13/2002(H3N2) | 19/4/2002 | NCBI |
| CY008019 | A/Canterbury/21/2002(H3N2) | 7/5/2002 | NCBI |
| CY008027 | A/Canterbury/46/2002(H3N2) | 9/6/2002 | NCBI |
| CY008035 | A/Canterbury/57/2002(H3N2) | 27/6/2002 | NCBI |
| CY008043 | A/Canterbury/24/2005(H3N2) | 4/5/2005 | NCBI |
| CY008051 | A/Canterbury/29/2005(H3N2) | 27/5/2005 | NCBI |
| CY008059 | A/Canterbury/67/2005(H3N2) | 21/6/2005 | NCBI |
| CY008067 | A/Canterbury/127/2005(H3N2) | 2/7/2005 | NCBI |
| CY008075 | A/Canterbury/129/2005(H3N2) | 3/7/2005 | NCBI |
| CY008083 | A/Canterbury/166/2005(H3N2) | 8/7/2005 | NCBI |
| CY008091 | A/Canterbury/206/2005(H3N2) | 13/7/2005 | NCBI |
| CY008099 | A/Canterbury/212/2005(H3N2) | 20/7/2005 | NCBI |
| CY008107 | A/Canterbury/259/2005(H3N2) | 1/8/2005 | NCBI |
| CY008115 | A/Memphis/2/1977(H3N2) | 1977 | NCBI |
| CY008123 | A/Memphis/4/1977(H3N2) | 1977 | NCBI |
| CY008131 | A/Canterbury/42/2000(H3N2) | 25/8/2000 | NCBI |
| CY008139 | A/Canterbury/39/2000(H3N2) | 27/8/2000 | NCBI |

| | | | |
|---|---|---|---|
| CY008156 | A/Beijing/1/1968(H3N2) | 1968 | NCBI |
| CY008164 | A/NewYork/472/2004(H3N2) | 27/9/2004 | NCBI |
| CY008180 | A/NewYork/502/1998(H3N2) | 20/3/1998 | NCBI |
| CY008212 | A/Tairawhiti/223/2004(H3N2) | 18/9/2004 | NCBI |
| CY008236 | A/Canterbury/309/2004(H3N2) | 6/10/2004 | NCBI |
| CY008276 | A/Canterbury/10/2002(H3N2) | 24/4/2002 | NCBI |
| CY008300 | A/Canterbury/61/2002(H3N2) | 30/6/2002 | NCBI |
| CY008364 | A/Canterbury/186/2005(H3N2) | 11/7/2005 | NCBI |
| CY008372 | A/Canterbury/204/2005(H3N2) | 18/7/2005 | NCBI |
| CY008404 | A/Canterbury/236/2005(H3N2) | 31/7/2005 | NCBI |
| CY008452 | A/Memphis/5/1985(H3N2) | 1985 | NCBI |
| CY008460 | A/Memphis/103/1972(H3N2) | 1972 | NCBI |
| CY008500 | A/Canterbury/88/2000(H3N2) | 2/10/2000 | NCBI |
| CY008508 | A/Canterbury/99/2000(H3N2) | 27/10/2000 | NCBI |
| CY008684 | A/Memphis/105/1972(H3N2) | 1972 | NCBI |
| CY008884 | A/NewYork/475/2003(H3N2) | 8/12/2003 | NCBI |
| CY009004 | A/HongKong/33/1973(H3N2) | 1973 | NCBI |
| CY009012 | A/Nanchang/12/1993(H3N2) | 1993 | NCBI |
| CY009028 | A/Canterbury/26/2005(H3N2) | 25/5/2005 | NCBI |
| CY009044 | A/Canterbury/124/2005(H3N2) | 1/7/2005 | NCBI |
| CY009100 | A/Canterbury/3/2000(H3N2) | 5/5/2000 | NCBI |
| CY009148 | A/Canterbury/58/2000(H3N2) | 5/9/2000 | NCBI |
| CY009260 | A/NewYork/499/2004(H3N2) | 15/1/2004 | NCBI |
| CY009460 | A/NewYork/560/1997(H3N2) | 6/1/1997 | NCBI |
| CY009668 | A/NewYork/574/1996(H3N2) | 12/12/1996 | NCBI |
| CY009740 | A/NewYork/590/1996(H3N2) | 10/12/1996 | NCBI |
| CY009852 | A/Canterbury/146/2001(H3N2) | 6/8/2001 | NCBI |
| CY009900 | A/NewYork/584/1996(H3N2) | 31/12/1996 | NCBI |
| CY009908 | A/NewYork/593/1996(H3N2) | 9/12/1996 | NCBI |
| CY010084 | A/Canterbury/250/2005(H3N2) | 16/8/2005 | NCBI |
| CY010596 | A/NewYork/578/1996(H3N2) | 27/12/1996 | NCBI |
| CY010724 | A/NewYork/634/1995(H3N2) | 8/12/1995 | NCBI |
| CY010748 | A/NewYork/648/1995(H3N2) | 14/12/1995 | NCBI |
| CY010796 | A/NewYork/555/1996(H3N2) | 23/12/1996 | NCBI |
| CY010812 | A/NewYork/623/1995(H3N2) | 29/12/1995 | NCBI |
| CY011128 | A/NewYork/599/1996(H3N2) | 16/12/1996 | NCBI |
| CY011256 | A/NewYork/557/1996(H3N2) | 21/11/1996 | NCBI |

| | | | |
|---|---|---|---|
| CY011264 | A/NewYork/559/1996(H3N2) | 25/11/1996 | NCBI |
| CY011288 | A/Memphis/5/1986(H3N2) | 1986 | NCBI |
| CY011440 | A/NewYork/632/1996(H3N2) | 1/3/1996 | NCBI |
| CY011456 | A/NewYork/636/1996(H3N2) | 22/1/1996 | NCBI |
| CY011560 | A/NewYork/679/1995(H3N2) | 3/3/1995 | NCBI |
| CY011568 | A/NewYork/693/1995(H3N2) | 21/3/1995 | NCBI |
| CY011616 | A/Wellington/22/2001(H3N2) | 9/7/2001 | NCBI |
| CY011640 | A/Wellington/66/2002(H3N2) | 3/7/2002 | NCBI |
| CY011648 | A/Waikato/21/2002(H3N2) | 1/7/2002 | NCBI |
| CY011752 | A/Waikato/61/2003(H3N2) | 30/6/2003 | NCBI |
| CY011760 | A/Waikato/75/2003(H3N2) | 2/7/2003 | NCBI |
| CY011768 | A/Waikato/120/2003(H3N2) | 8/7/2003 | NCBI |
| CY011816 | A/NewYork/652/1996(H3N2) | 1/4/1996 | NCBI |
| CY011904 | A/NewYork/720/1994(H3N2) | 24/1/1994 | NCBI |
| CY011944 | A/NewYork/752/1993(H3N2) | 30/12/1993 | NCBI |
| CY011960 | A/Waikato/5/2000(H3N2) | 30/8/2000 | NCBI |
| CY012056 | A/Waikato/51/2002(H3N2) | 12/8/2002 | NCBI |
| CY012064 | A/Wellington/34/2003(H3N2) | 30/6/2003 | NCBI |
| CY012080 | A/Waikato/91/2003(H3N2) | 28/6/2003 | NCBI |
| CY012088 | A/Waikato/102/2003(H3N2) | 8/7/2003 | NCBI |
| CY012096 | A/Waikato/150/2003(H3N2) | 7/8/2003 | NCBI |
| CY012104 | A/Wellington/01/2004(H3N2) | 26/1/2004 | NCBI |
| CY012112 | A/Wellington/6/2004(H3N2) | 18/8/2004 | NCBI |
| CY012120 | A/NewYork/756/1993(H3N2) | 2/2/1993 | NCBI |
| CY012128 | A/NewYork/758/1993(H3N2) | 3/2/1993 | NCBI |
| CY012136 | A/NewYork/759/1993(H3N2) | 8/3/1993 | NCBI |
| CY012144 | A/NewYork/760/1993(H3N2) | 9/2/1993 | NCBI |
| CY012152 | A/NewYork/761/1993(H3N2) | 3/3/1993 | NCBI |
| CY012160 | A/NewYork/762/1993(H3N2) | 4/2/1993 | NCBI |
| CY012168 | A/NewYork/763/1993(H3N2) | 26/3/1993 | NCBI |
| CY012176 | A/NewYork/764/1993(H3N2) | 23/2/1993 | NCBI |
| CY012184 | A/NewYork/765/1993(H3N2) | 3/3/1993 | NCBI |
| CY012208 | A/NewYork/598/1996(H3N2) | 17/12/1996 | NCBI |
| CY012224 | A/NewYork/657/1994(H3N2) | 2/12/1994 | NCBI |
| CY012248 | A/NewYork/677/1995(H3N2) | 18/2/1995 | NCBI |
| CY012312 | A/Wellington/34/2001(H3N2) | 28/8/2001 | NCBI |
| CY012320 | A/Waikato/105/2001(H3N2) | 13/9/2001 | NCBI |

| | | | |
|---|---|---|---|
| CY012336 | A/Wellington/63/2002(H3N2) | 2002 | NCBI |
| CY012344 | A/Waikato/155/2003(H3N2) | 22/6/2003 | NCBI |
| CY012360 | A/Waikato/29/2003(H3N2) | 23/6/2003 | NCBI |
| CY012368 | A/Waikato/115/2003(H3N2) | 6/7/2003 | NCBI |
| CY012376 | A/Wellington/53/2003(H3N2) | 16/7/2003 | NCBI |
| CY012384 | A/Waikato/122/2003(H3N2) | 25/6/2003 | NCBI |
| CY012392 | A/Waikato/129/2003(H3N2) | 11/7/2003 | NCBI |
| CY012400 | A/Waikato/133/2003(H3N2) | 8/7/2003 | NCBI |
| CY012408 | A/Waikato/139/2003(H3N2) | 24/7/2003 | NCBI |
| CY012416 | A/Waikato/147/2003(H3N2) | 28/7/2003 | NCBI |
| CY012424 | A/Waikato/148/2003(H3N2) | 25/7/2003 | NCBI |
| CY012432 | A/Canterbury/97/2000(H3N2) | 25/10/2000 | NCBI |
| CY012456 | A/NewYork/661/1994(H3N2) | 15/12/1994 | NCBI |
| CY012464 | A/NewYork/663/1994(H3N2) | 20/12/1994 | NCBI |
| CY012480 | A/NewYork/666/1995(H3N2) | 8/2/1995 | NCBI |
| CY012488 | A/NewYork/667/1995(H3N2) | 8/1/1995 | NCBI |
| CY012504 | A/NewYork/678/1995(H3N2) | 20/1/1995 | NCBI |
| CY012568 | A/NewYork/746/1994(H3N2) | 21/2/1994 | NCBI |
| CY012576 | A/NewYork/747/1994(H3N2) | 12/1/1994 | NCBI |
| CY012584 | A/NewYork/748/1994(H3N2) | 17/1/1994 | NCBI |
| CY012592 | A/NewYork/749/1994(H3N2) | 11/1/1994 | NCBI |
| CY012616 | A/Dunedin/1/2000(H3N2) | 21/1/2000 | NCBI |
| CY012624 | A/Waikato/4/2000(H3N2) | 28/8/2000 | NCBI |
| CY012632 | A/Waikato/7/2000(H3N2) | 8/9/2000 | NCBI |
| CY012656 | A/Wellington/47/2003(H3N2) | 7/7/2003 | NCBI |
| CY012664 | A/Wellington/49/2003(H3N2) | 8/7/2003 | NCBI |
| CY012672 | A/Wellington/14/2004(H3N2) | 31/8/2004 | NCBI |
| CY012680 | A/Wellington/8/2004(H3N2) | 1/9/2004 | NCBI |
| CY012688 | A/Wellington/22/2004(H3N2) | 12/9/2004 | NCBI |
| CY012696 | A/Wellington/31/2004(H3N2) | 14/9/2004 | NCBI |
| CY012704 | A/Waikato/21/2004(H3N2) | 5/9/2004 | NCBI |
| CY012712 | A/Wellington/52/2004(H3N2) | 23/9/2004 | NCBI |
| CY012720 | A/Waikato/40/2004(H3N2) | 21/9/2004 | NCBI |
| CY012728 | A/NewYork/767/1993(H3N2) | 5/4/1993 | NCBI |
| CY012736 | A/NewYork/770/1993(H3N2) | 4/3/1993 | NCBI |
| CY012744 | A/NewYork/771/1993(H3N2) | 19/4/1993 | NCBI |
| CY012752 | A/NewYork/775/1993(H3N2) | 11/3/1993 | NCBI |

| | | | |
|---|---|---|---|
| CY012760 | A/NewYork/777/1993(H3N2) | 20/3/1993 | NCBI |
| CY012768 | A/NewYork/778/1993(H3N2) | 2/2/1993 | NCBI |
| CY012776 | A/NewYork/779/1993(H3N2) | 1/2/1993 | NCBI |
| CY012784 | A/NewYork/786/1993(H3N2) | 16/3/1993 | NCBI |
| CY012792 | A/NewYork/4/2006(H3N2) | 22/3/2006 | NCBI |
| CY012848 | A/NewYork/554/1996(H3N2) | 12/11/1996 | NCBI |
| CY012920 | A/NewYork/690/1995(H3N2) | 18/1/1995 | NCBI |
| CY012944 | A/NewYork/723/1994(H3N2) | 2/1/1994 | NCBI |
| CY012952 | A/NewYork/724/1994(H3N2) | 27/1/1994 | NCBI |
| CY012960 | A/NewYork/725/1994(H3N2) | 30/1/1994 | NCBI |
| CY012968 | A/NewYork/735/1994(H3N2) | 14/1/1994 | NCBI |
| CY012976 | A/NewYork/736/1994(H3N2) | 21/1/1994 | NCBI |
| CY012984 | A/NewYork/738/1994(H3N2) | 4/2/1994 | NCBI |
| CY012992 | A/NewYork/739/1994(H3N2) | 14/2/1994 | NCBI |
| CY013048 | A/Waikato/6/2000(H3N2) | 8/9/2000 | NCBI |
| CY013080 | A/Wellington/9/2002(H3N2) | 2002 | NCBI |
| CY013088 | A/Waikato/25/2002(H3N2) | 18/7/2002 | NCBI |
| CY013096 | A/Waikato/36/2002(H3N2) | 23/7/2002 | NCBI |
| CY013112 | A/Waikato/21/2003(H3N2) | 21/6/2003 | NCBI |
| CY013120 | A/Waikato/46/2003(H3N2) | 24/6/2003 | NCBI |
| CY013128 | A/Waikato/53/2003(H3N2) | 2/7/2003 | NCBI |
| CY013136 | A/Waikato/54/2003(H3N2) | 3/7/2003 | NCBI |
| CY013144 | A/Waikato/94/2003(H3N2) | 27/6/2003 | NCBI |
| CY013152 | A/Wellington/35/2004(H3N2) | 21/9/2004 | NCBI |
| CY013160 | A/Wellington/45/2004(H3N2) | 21/9/2004 | NCBI |
| CY013168 | A/NewYork/754/1993(H3N2) | 16/1/1993 | NCBI |
| CY013176 | A/NewYork/755/1993(H3N2) | 12/3/1993 | NCBI |
| CY013184 | A/NewYork/781/1993(H3N2) | 26/3/1993 | NCBI |
| CY013192 | A/NewYork/782/1993(H3N2) | 3/4/1993 | NCBI |
| CY013200 | A/NewYork/783/1993(H3N2) | 1/4/1993 | NCBI |
| CY013208 | A/NewYork/785/1993(H3N2) | 26/3/1993 | NCBI |
| CY013216 | A/WesternAustralia/15/2001(H3N2) | 25/9/2001 | NCBI |
| CY013224 | A/WesternAustralia/23/2002(H3N2) | 25/6/2002 | NCBI |
| CY013232 | A/NewYork/5/2006(H3N2) | 5/4/2006 | NCBI |
| CY013240 | A/Canterbury/232/2005(H3N2) | 16/7/2005 | NCBI |
| CY013319 | A/NewYork/676/1995(H3N2) | 18/1/1995 | NCBI |
| CY013351 | A/NewYork/743/1994(H3N2) | 7/1/1994 | NCBI |

| | | | |
|---|---|---|---|
| CY013367 | A/NewYork/745/1994(H3N2) | 7/2/1994 | NCBI |
| CY013389 | A/Waikato/16/2000(H3N2) | 27/9/2000 | NCBI |
| CY013397 | A/Waikato/17/2000(H3N2) | 22/9/2000 | NCBI |
| CY013405 | A/Wellington/28/2000(H3N2) | 2000 | NCBI |
| CY013413 | A/Waikato/31/2002(H3N2) | 26/7/2002 | NCBI |
| CY013421 | A/Dunedin/12/2002(H3N2) | 23/7/2002 | NCBI |
| CY013429 | A/Waikato/15/2003(H3N2) | 16/6/2003 | NCBI |
| CY013437 | A/Waikato/108/2003(H3N2) | 7/7/2003 | NCBI |
| CY013445 | A/Dunedin/38/2003(H3N2) | 2/8/2003 | NCBI |
| CY013453 | A/Wellington/23/2004(H3N2) | 23/8/2004 | NCBI |
| CY013461 | A/Waikato/12/2004(H3N2) | 1/9/2004 | NCBI |
| CY013469 | A/Wellington/18/2004(H3N2) | 6/9/2004 | NCBI |
| CY013477 | A/Wellington/27/2004(H3N2) | 11/9/2004 | NCBI |
| CY013485 | A/Wellington/44/2004(H3N2) | 21/9/2004 | NCBI |
| CY013493 | A/Wellington/38/2004(H3N2) | 20/9/2004 | NCBI |
| CY013501 | A/Waikato/22/2004(H3N2) | 7/9/2004 | NCBI |
| CY013509 | A/Waikato/26/2004(H3N2) | 16/9/2004 | NCBI |
| CY013525 | A/Waikato/35/2004(H3N2) | 20/9/2004 | NCBI |
| CY013533 | A/Waikato/43/2004(H3N2) | 21/9/2004 | NCBI |
| CY013549 | A/Wellington/62/2004(H3N2) | 5/10/2004 | NCBI |
| CY013605 | A/NewYork/757/1993(H3N2) | 26/3/1993 | NCBI |
| CY013613 | A/NewYork/766/1993(H3N2) | 15/2/1993 | NCBI |
| CY013621 | A/NewYork/768/1993(H3N2) | 9/3/1993 | NCBI |
| CY013629 | A/NewYork/769/1993(H3N2) | 16/3/1993 | NCBI |
| CY013637 | A/NewYork/772/1993(H3N2) | 10/3/1993 | NCBI |
| CY013645 | A/NewYork/773/1993(H3N2) | 31/3/1993 | NCBI |
| CY013653 | A/NewYork/774/1993(H3N2) | 25/2/1993 | NCBI |
| CY013661 | A/NewYork/776/1993(H3N2) | 18/2/1993 | NCBI |
| CY013669 | A/NewYork/780/1993(H3N2) | 2/3/1993 | NCBI |
| CY013677 | A/NewYork/784/1993(H3N2) | 19/4/1993 | NCBI |
| CY013685 | A/NewYork/787/1993(H3N2) | 10/3/1993 | NCBI |
| CY013693 | A/NewYork/788/1993(H3N2) | 26/2/1993 | NCBI |
| CY013701 | A/NewYork/789/1993(H3N2) | 5/4/1993 | NCBI |
| CY013709 | A/NewYork/790/1993(H3N2) | 4/2/1993 | NCBI |
| CY013717 | A/NewYork/791/1993(H3N2) | 11/3/1993 | NCBI |
| CY013725 | A/NewYork/792/1993(H3N2) | 9/3/1993 | NCBI |
| CY013733 | A/NewYork/794/1993(H3N2) | 4/3/1993 | NCBI |

| | | | |
|---|---|---|---|
| CY013741 | A/NewYork/795/1993(H3N2) | 4/2/1993 | NCBI |
| CY013749 | A/NewYork/797/1993(H3N2) | 16/3/1993 | NCBI |
| CY013757 | A/NewYork/798/1993(H3N2) | 9/3/1993 | NCBI |
| CY013765 | A/NewYork/799/1993(H3N2) | 10/3/1993 | NCBI |
| CY013773 | A/NewYork/801/1993(H3N2) | 10/3/1993 | NCBI |
| CY013781 | A/NewYork/802/1993(H3N2) | 2/3/1993 | NCBI |
| CY013789 | A/NewYork/803/1993(H3N2) | 9/3/1993 | NCBI |
| CY013805 | A/NewYork/C4/2002(H3N2) | 1/2/2002 | NCBI |
| CY013895 | A/Wellington/26/2000(H3N2) | 2000 | NCBI |
| CY013903 | A/Wellington/30/2000(H3N2) | 25/11/2000 | NCBI |
| CY013911 | A/Wellington/4/2004(H3N2) | 18/8/2004 | NCBI |
| CY013919 | A/Waikato/3/2004(H3N2) | 3/9/2004 | NCBI |
| CY013927 | A/Wellington/59/2004(H3N2) | 28/9/2004 | NCBI |
| CY013935 | A/Waikato/51/2004(H3N2) | 28/9/2004 | NCBI |
| CY013943 | A/Waikato/56/2004(H3N2) | 29/9/2004 | NCBI |
| CY013951 | A/Waikato/64/2004(H3N2) | 5/10/2004 | NCBI |
| CY013959 | A/Waikato/68/2004(H3N2) | 2/10/2004 | NCBI |
| CY013967 | A/Waikato/69/2004(H3N2) | 8/10/2004 | NCBI |
| CY013975 | A/Waikato/71/2004(H3N2) | 14/10/2004 | NCBI |
| CY013983 | A/Wellington/64/2004(H3N2) | 19/10/2004 | NCBI |
| CY013991 | A/Waikato/72/2004(H3N2) | 19/10/2004 | NCBI |
| CY013999 | A/Waikato/73/2004(H3N2) | 15/10/2004 | NCBI |
| CY014015 | A/Waikato/1/2005(H3N2) | 25/1/2005 | NCBI |
| CY014023 | A/Wellington/1/2005(H3N2) | 9/7/2005 | NCBI |
| CY014031 | A/Wellington/2/2005(H3N2) | 11/7/2005 | NCBI |
| CY014039 | A/Otago/1/2005(H3N2) | 1/7/2005 | NCBI |
| CY014047 | A/Wellington/3/2005(H3N2) | 13/7/2005 | NCBI |
| CY014055 | A/Southland/5/2005(H3N2) | 8/8/2005 | NCBI |
| CY014063 | A/Waikato/8/2005(H3N2) | 9/8/2005 | NCBI |
| CY014071 | A/Waikato/7/2005(H3N2) | 10/8/2005 | NCBI |
| CY014079 | A/Waikato/16/2005(H3N2) | 9/8/2005 | NCBI |
| CY014087 | A/Otago/02/2005(H3N2) | 11/8/2005 | NCBI |
| CY014095 | A/Wellington/8/2005(H3N2) | 23/8/2005 | NCBI |
| CY014103 | A/Wellington/9/2005(H3N2) | 23/8/2005 | NCBI |
| CY014111 | A/Waikato/9/2005(H3N2) | 15/8/2005 | NCBI |
| CY014119 | A/Otago/4/2005(H3N2) | 31/8/2005 | NCBI |
| CY014127 | A/Otago/3/2005(H3N2) | 31/8/2005 | NCBI |

| | | | |
|---|---|---|---|
| CY014135 | A/Waikato/12/2005(H3N2) | 13/9/2005 | NCBI |
| CY014143 | A/NewYork/793/1993(H3N2) | 2/2/1993 | NCBI |
| CY014151 | A/NewYork/796/1993(H3N2) | 18/2/1993 | NCBI |
| CY014159 | A/NewYork/7/2006(H3N2) | 5/4/2006 | NCBI |
| CY015516 | A/NewYork/601/1996(H3N2) | 9/12/1996 | NCBI |
| CY015540 | A/NewYork/737/1994(H3N2) | 3/2/1994 | NCBI |
| CY015548 | A/Dunedin/1/2002(H3N2) | 7/6/2002 | NCBI |
| CY015556 | A/Waikato/23/2002(H3N2) | 3/7/2002 | NCBI |
| CY015564 | A/Waikato/16/2004(H3N2) | 9/9/2004 | NCBI |
| CY015572 | A/Wellington/34/2004(H3N2) | 19/9/2004 | NCBI |
| CY015588 | A/Waikato/2/2005(H3N2) | 26/1/2005 | NCBI |
| CY015596 | A/Waikato/3/2005(H3N2) | 13/4/2005 | NCBI |
| CY015604 | A/Wellington/4/2005(H3N2) | 12/7/2005 | NCBI |
| CY015612 | A/Wellington/5/2005(H3N2) | 26/7/2005 | NCBI |
| CY015620 | A/Wellington/6/2005(H3N2) | 14/8/2005 | NCBI |
| CY015628 | A/Waikato/15/2005(H3N2) | 11/8/2005 | NCBI |
| CY015636 | A/Waikato/10/2005(H3N2) | 26/8/2005 | NCBI |
| CY015644 | A/WesternAustralia/2/2000(H3N2) | 20/9/2000 | NCBI |
| CY015652 | A/WesternAustralia/3/2000(H3N2) | 12/9/2000 | NCBI |
| CY015660 | A/WesternAustralia/8/2000(H3N2) | 17/10/2000 | NCBI |
| CY015668 | A/WesternAustralia/11/2000(H3N2) | 14/10/2000 | NCBI |
| CY015676 | A/WesternAustralia/13/2001(H3N2) | 11/7/2001 | NCBI |
| CY015684 | A/WesternAustralia/14/2001(H3N2) | 26/9/2001 | NCBI |
| CY015692 | A/WesternAustralia/16/2001(H3N2) | 25/9/2001 | NCBI |
| CY015700 | A/WesternAustralia/17/2001(H3N2) | 2/10/2001 | NCBI |
| CY015708 | A/WesternAustralia/25/2002(H3N2) | 10/7/2002 | NCBI |
| CY015716 | A/WesternAustralia/26/2002(H3N2) | 8/7/2002 | NCBI |
| CY015724 | A/WesternAustralia/28/2002(H3N2) | 6/8/2002 | NCBI |
| CY015732 | A/WesternAustralia/29/2002(H3N2) | 21/8/2002 | NCBI |
| CY015740 | A/WesternAustralia/30/2002(H3N2) | 19/8/2002 | NCBI |
| CY015748 | A/WesternAustralia/31/2002(H3N2) | 26/8/2002 | NCBI |
| CY015756 | A/WesternAustralia/35/2002(H3N2) | 29/9/2002 | NCBI |
| CY015764 | A/WesternAustralia/36/2002(H3N2) | 15/10/2002 | NCBI |
| CY015772 | A/WesternAustralia/37/2003(H3N2) | 9/9/2003 | NCBI |
| CY015780 | A/WesternAustralia/38/2003(H3N2) | 8/9/2003 | NCBI |
| CY015788 | A/WesternAustralia/39/2003(H3N2) | 24/9/2003 | NCBI |
| CY015796 | A/WesternAustralia/40/2003(H3N2) | 17/9/2003 | NCBI |

| | | | |
|---|---|---|---|
| CY015804 | A/WesternAustralia/41/2003(H3N2) | 23/9/2003 | NCBI |
| CY015812 | A/WesternAustralia/42/2003(H3N2) | 17/9/2003 | NCBI |
| CY015820 | A/WesternAustralia/43/2003(H3N2) | 12/9/2003 | NCBI |
| CY015828 | A/WesternAustralia/44/2003(H3N2) | 22/10/2003 | NCBI |
| CY015836 | A/WesternAustralia/45/2003(H3N2) | 9/9/2003 | NCBI |
| CY015844 | A/WesternAustralia/46/2003(H3N2) | 10/9/2003 | NCBI |
| CY015852 | A/WesternAustralia/47/2003(H3N2) | 3/10/2003 | NCBI |
| CY015860 | A/WesternAustralia/48/2003(H3N2) | 6/10/2003 | NCBI |
| CY015868 | A/WesternAustralia/50/2003(H3N2) | 17/9/2003 | NCBI |
| CY015876 | A/WesternAustralia/51/2004(H3N2) | 21/9/2004 | NCBI |
| CY015884 | A/WesternAustralia/52/2004(H3N2) | 21/9/2004 | NCBI |
| CY015892 | A/WesternAustralia/53/2004(H3N2) | 29/9/2004 | NCBI |
| CY015900 | A/WesternAustralia/54/2004(H3N2) | 18/10/2004 | NCBI |
| CY015908 | A/WesternAustralia/55/2004(H3N2) | 14/9/2004 | NCBI |
| CY015916 | A/WesternAustralia/56/2004(H3N2) | 4/10/2004 | NCBI |
| CY015924 | A/WesternAustralia/57/2004(H3N2) | 7/10/2004 | NCBI |
| CY015932 | A/WesternAustralia/58/2004(H3N2) | 1/10/2004 | NCBI |
| CY015940 | A/WesternAustralia/59/2004(H3N2) | 17/10/2004 | NCBI |
| CY015948 | A/WesternAustralia/60/2004(H3N2) | 13/9/2004 | NCBI |
| CY015956 | A/WesternAustralia/61/2004(H3N2) | 14/10/2004 | NCBI |
| CY015964 | A/WesternAustralia/62/2004(H3N2) | 19/10/2004 | NCBI |
| CY015972 | A/WesternAustralia/63/2004(H3N2) | 9/10/2004 | NCBI |
| CY015980 | A/WesternAustralia/64/2004(H3N2) | 28/10/2004 | NCBI |
| CY015988 | A/WesternAustralia/65/2005(H3N2) | 4/1/2005 | NCBI |
| CY015996 | A/WesternAustralia/66/2005(H3N2) | 28/6/2005 | NCBI |
| CY016004 | A/WesternAustralia/68/2005(H3N2) | 4/7/2005 | NCBI |
| CY016012 | A/WesternAustralia/70/2005(H3N2) | 11/7/2005 | NCBI |
| CY016020 | A/WesternAustralia/73/2005(H3N2) | 2005 | NCBI |
| CY016028 | A/WesternAustralia/74/2005(H3N2) | 29/7/2005 | NCBI |
| CY016036 | A/WesternAustralia/75/2005(H3N2) | 27/7/2005 | NCBI |
| CY016044 | A/WesternAustralia/78/2005(H3N2) | 2005 | NCBI |
| CY016060 | A/Memphis/2/1986(H3N2) | 1986 | NCBI |
| CY016068 | A/NewSouthWales/13/1999(H3N2) | 1999 | NCBI |
| CY016076 | A/NewSouthWales/20/1999(H3N2) | 1999 | NCBI |
| CY016084 | A/NewSouthWales/25/2000(H3N2) | 2000 | NCBI |
| CY016092 | A/NewSouthWales/27/2000(H3N2) | 2000 | NCBI |
| CY016100 | A/NewSouthWales/28/2000(H3N2) | 2000 | NCBI |

| | | | |
|---|---|---|---|
| CY016108 | A/NewSouthWales/29/2000(H3N2) | 2000 | NCBI |
| CY016116 | A/NewSouthWales/36/2000(H3N2) | 2000 | NCBI |
| CY016212 | A/Waikato/1/2004(H3N2) | 12/8/2004 | NCBI |
| CY016268 | A/NewSouthWales/33/2000(H3N2) | 2000 | NCBI |
| CY016427 | A/NewYork/706/1993(H3N2) | 30/12/1993 | NCBI |
| CY016443 | A/Wellington/61/2004(H3N2) | 7/10/2004 | NCBI |
| CY016451 | A/Waikato/70/2004(H3N2) | 7/10/2004 | NCBI |
| CY016467 | A/Waikato/18/2005(H3N2) | 9/8/2005 | NCBI |
| CY016475 | A/Wellington/7/2005(H3N2) | 23/8/2005 | NCBI |
| CY016483 | A/NewYork/800/1993(H3N2) | 11/3/1993 | NCBI |
| CY016491 | A/WesternAustralia/10/2000(H3N2) | 19/10/2000 | NCBI |
| CY016499 | A/WesternAustralia/12/2000(H3N2) | 11/10/2000 | NCBI |
| CY016507 | A/NewSouthWales/2/1999(H3N2) | 1999 | NCBI |
| CY016515 | A/NewSouthWales/5/1999(H3N2) | 1999 | NCBI |
| CY016523 | A/NewSouthWales/6/1999(H3N2) | 1999 | NCBI |
| CY016531 | A/NewSouthWales/8/1999(H3N2) | 1999 | NCBI |
| CY016539 | A/NewSouthWales/15/1999(H3N2) | 1999 | NCBI |
| CY016547 | A/NewSouthWales/16/1999(H3N2) | 1999 | NCBI |
| CY016555 | A/NewSouthWales/21/1999(H3N2) | 1999 | NCBI |
| CY016571 | A/NewSouthWales/32/2000(H3N2) | 2000 | NCBI |
| CY016579 | A/NewSouthWales/34/2000(H3N2) | 2000 | NCBI |
| CY016587 | A/NewSouthWales/35/2000(H3N2) | 2000 | NCBI |
| CY016603 | A/SouthAustralia/20/2005(H3N2) | 2005 | NCBI |
| CY016627 | A/NewSouthWales/4/1999(H3N2) | 1999 | NCBI |
| CY016635 | A/NewSouthWales/17/1999(H3N2) | 1999 | NCBI |
| CY016651 | A/NewSouthWales/22/1999(H3N2) | 1999 | NCBI |
| CY016659 | A/SouthAustralia/23/2005(H3N2) | 2005 | NCBI |
| CY016707 | A/SouthAustralia/59/2000(H3N2) | 2000 | NCBI |
| CY016715 | A/SouthAustralia/62/2000(H3N2) | 2000 | NCBI |
| CY016739 | A/SouthAustralia/66/2000(H3N2) | 2000 | NCBI |
| CY016747 | A/SouthAustralia/67/2000(H3N2) | 2000 | NCBI |
| CY016755 | A/SouthAustralia/68/2000(H3N2) | 2000 | NCBI |
| CY016763 | A/SouthAustralia/71/2000(H3N2) | 2000 | NCBI |
| CY016771 | A/SouthAustralia/74/2000(H3N2) | 2000 | NCBI |
| CY016979 | A/WesternAustralia/69/2005(H3N2) | 5/7/2005 | NCBI |
| CY016987 | A/WesternAustralia/72/2005(H3N2) | 21/7/2005 | NCBI |
| CY016995 | A/NewYork/6/2006(H3N2) | 6/4/2006 | NCBI |

| | | | |
|---|---|---|---|
| CY017083 | A/WesternAustralia/27/2002(H3N2) | 21/7/2002 | NCBI |
| CY017091 | A/WesternAustralia/32/2002(H3N2) | 26/8/2002 | NCBI |
| CY017099 | A/WesternAustralia/76/2005(H3N2) | 1/8/2005 | NCBI |
| CY017107 | A/WesternAustralia/79/2005(H3N2) | 3/8/2005 | NCBI |
| CY017131 | A/SouthAustralia/22/2005(H3N2) | 2005 | NCBI |
| CY017163 | A/SouthAustralia/52/2000(H3N2) | 2000 | NCBI |
| CY017171 | A/SouthAustralia/76/2000(H3N2) | 2000 | NCBI |
| CY017259 | A/Queensland/1/2000(H3N2) | 15/9/2000 | NCBI |
| CY017267 | A/Queensland/4/2000(H3N2) | 5/9/2000 | NCBI |
| CY017283 | A/NewYork/680/1995(H3N2) | 8/2/1995 | NCBI |
| CY017291 | A/NewYork/681/1995(H3N2) | 20/2/1995 | NCBI |
| CY017307 | A/NewYork/704/1994(H3N2) | 3/1/1994 | NCBI |
| CY017323 | A/WesternAustralia/1/2000(H3N2) | 12/9/2000 | NCBI |
| CY017331 | A/WesternAustralia/5/2000(H3N2) | 25/9/2000 | NCBI |
| CY017339 | A/WesternAustralia/7/2000(H3N2) | 11/9/2000 | NCBI |
| CY017347 | A/WesternAustralia/9/2000(H3N2) | 8/10/2000 | NCBI |
| CY017355 | A/WesternAustralia/24/2002(H3N2) | 25/6/2002 | NCBI |
| CY017387 | A/SouthAustralia/75/2000(H3N2) | 2000 | NCBI |
| CY017395 | A/SouthAustralia/77/2000(H3N2) | 2000 | NCBI |
| CY017443 | A/Queensland/2/2000(H3N2) | 16/7/2000 | NCBI |
| CY017451 | A/Queensland/3/2000(H3N2) | 4/9/2000 | NCBI |
| CY017459 | A/Queensland/5/2000(H3N2) | 13/8/2000 | NCBI |
| CY017467 | A/Queensland/8/2000(H3N2) | 20/9/2000 | NCBI |
| CY017475 | A/Queensland/9/2000(H3N2) | 28/8/2000 | NCBI |
| CY017483 | A/Queensland/11/2001(H3N2) | 11/8/2001 | NCBI |
| CY017491 | A/Queensland/12/2001(H3N2) | 24/8/2001 | NCBI |
| CY017499 | A/Queensland/15/2001(H3N2) | 13/9/2001 | NCBI |
| CY017507 | A/Queensland/16/2001(H3N2) | 17/8/2001 | NCBI |
| CY017515 | A/Queensland/18/2001(H3N2) | 24/8/2001 | NCBI |
| CY017523 | A/Queensland/20/2001(H3N2) | 5/8/2001 | NCBI |
| CY017531 | A/Queensland/21/2002(H3N2) | 26/7/2002 | NCBI |
| CY017539 | A/Queensland/23/2002(H3N2) | 13/8/2002 | NCBI |
| CY017547 | A/Queensland/24/2002(H3N2) | 25/7/2002 | NCBI |
| CY017555 | A/Queensland/26/2002(H3N2) | 6/8/2002 | NCBI |
| CY017563 | A/Queensland/33/2003(H3N2) | 25/8/2003 | NCBI |
| CY017571 | A/Queensland/36/2003(H3N2) | 10/8/2003 | NCBI |
| CY017579 | A/Queensland/39/2003(H3N2) | 11/9/2003 | NCBI |

| | | | |
|---|---|---|---|
| CY017587 | A/Queensland/47/2004(H3N2) | 6/10/2004 | NCBI |
| CY017595 | A/Queensland/48/2004(H3N2) | 30/9/2004 | NCBI |
| CY017603 | A/Queensland/49/2004(H3N2) | 1/10/2004 | NCBI |
| CY017611 | A/Queensland/55/2005(H3N2) | 2/8/2005 | NCBI |
| CY017619 | A/Queensland/57/2005(H3N2) | 22/8/2005 | NCBI |
| CY017627 | A/Queensland/17/2001(H3N2) | 24/8/2001 | NCBI |
| CY017797 | A/WesternAustralia/49/2003(H3N2) | 15/9/2003 | NCBI |
| CY017805 | A/SouthAustralia/2/2000(H3N2) | 2000 | NCBI |
| CY017821 | A/SouthAustralia/32/2005(H3N2) | 2005 | NCBI |
| CY017885 | A/Queensland/7/2000(H3N2) | 19/9/2000 | NCBI |
| CY017893 | A/Queensland/19/2001(H3N2) | 14/7/2001 | NCBI |
| CY017901 | A/Queensland/22/2002(H3N2) | 15/7/2002 | NCBI |
| CY017909 | A/Queensland/27/2002(H3N2) | 13/8/2002 | NCBI |
| CY017917 | A/Queensland/28/2002(H3N2) | 12/8/2002 | NCBI |
| CY017925 | A/Queensland/29/2002(H3N2) | 17/8/2002 | NCBI |
| CY017933 | A/Queensland/37/2003(H3N2) | 19/8/2003 | NCBI |
| CY017941 | A/Queensland/38/2003(H3N2) | 3/9/2003 | NCBI |
| CY017949 | A/Queensland/40/2003(H3N2) | 10/9/2003 | NCBI |
| CY017965 | A/Queensland/43/2004(H3N2) | 24/10/2004 | NCBI |
| CY017973 | A/Queensland/46/2004(H3N2) | 9/10/2004 | NCBI |
| CY017981 | A/Queensland/51/2005(H3N2) | 12/7/2005 | NCBI |
| CY017989 | A/Queensland/52/2005(H3N2) | 22/7/2005 | NCBI |
| CY018925 | A/WesternAustralia/67/2005(H3N2) | 9/7/2005 | NCBI |
| CY018941 | A/SouthAustralia/42/2005(H3N2) | 2005 | NCBI |
| CY018957 | A/Queensland/14/2001(H3N2) | 3/9/2001 | NCBI |
| CY018965 | A/Queensland/30/2002(H3N2) | 23/8/2002 | NCBI |
| CY018973 | A/Queensland/31/2003(H3N2) | 12/8/2003 | NCBI |
| CY018981 | A/Queensland/32/2003(H3N2) | 20/8/2003 | NCBI |
| CY018989 | A/Queensland/34/2003(H3N2) | 25/8/2003 | NCBI |
| CY018997 | A/Queensland/35/2003(H3N2) | 27/7/2003 | NCBI |
| CY019005 | A/Queensland/41/2004(H3N2) | 7/10/2004 | NCBI |
| CY019013 | A/Queensland/53/2005(H3N2) | 26/7/2005 | NCBI |
| CY019021 | A/Queensland/54/2005(H3N2) | 11/8/2005 | NCBI |
| CY019029 | A/Queensland/56/2005(H3N2) | 16/8/2005 | NCBI |
| CY019141 | A/NewYork/900/2004(H3N2) | 22/12/2004 | NCBI |
| CY019149 | A/NewYork/900/2004(H3N2) | 22/12/2004 | NCBI |
| CY019173 | A/NewYork/905/2004(H3N2) | 27/12/2004 | NCBI |

| | | | |
|---|---|---|---|
| CY019189 | A/NewYork/918/2005(H3N2) | 10/1/2005 | NCBI |
| CY019245 | A/NewYork/908/2004(H3N2) | 27/12/2004 | NCBI |
| CY019269 | A/NewYork/913/2005(H3N2) | 10/1/2005 | NCBI |
| CY019325 | A/NewYork/928/2006(H3N2) | 28/2/2006 | NCBI |
| CY019333 | A/NewYork/933/2006(H3N2) | 1/3/2006 | NCBI |
| CY019747 | A/Queensland/60/2005(H3N2) | 18/8/2005 | NCBI |
| CY019827 | A/NewYork/923/2006(H3N2) | 23/2/2006 | NCBI |
| CY019859 | A/NewYork/938/2006(H3N2) | 2/3/2006 | NCBI |
| CY019891 | A/Albany/11/1968(H3N2) | 1968 | NCBI |
| CY019899 | A/Albany/1/1969(H3N2) | 1969 | NCBI |
| CY019907 | A/Albany/19/1968(H3N2) | 1968 | NCBI |
| CY019915 | A/Albany/3/1969(H3N2) | 1969 | NCBI |
| CY019931 | A/WesternAustralia/34/2002(H3N2) | 28/8/2002 | NCBI |
| CY019981 | A/Canterbury/50/2000(H3N2) | 2/8/2000 | NCBI |
| CY019989 | A/Wellington/21/2000(H3N2) | 19/9/2000 | NCBI |
| CY020005 | A/WesternAustralia/71/2005(H3N2) | 20/10/2005 | NCBI |
| CY020013 | A/SouthAustralia/49/2000(H3N2) | 2000 | NCBI |
| CY020021 | A/SouthAustralia/80/2000(H3N2) | 2000 | NCBI |
| CY020029 | A/Queensland/44/2004(H3N2) | 9/10/2004 | NCBI |
| CY020037 | A/Queensland/45/2004(H3N2) | 15/9/2004 | NCBI |
| CY020045 | A/Queensland/50/2004(H3N2) | 27/10/2004 | NCBI |
| CY020197 | A/NewSouthWales/7/1999(H3N2) | 1999 | NCBI |
| CY020205 | A/NewSouthWales/19/1999(H3N2) | 1999 | NCBI |
| CY020213 | A/SouthAustralia/15/2000(H3N2) | 2000 | NCBI |
| CY020221 | A/Albany/14/1978(H3N2) | 1978 | NCBI |
| CY020301 | A/SouthAustralia/61/2005(H3N2) | 2005 | NCBI |
| CY020309 | A/Queensland/10/2000(H3N2) | 2000 | NCBI |
| CY020341 | A/Wellington/55/2004(H3N2) | 27/9/2004 | NCBI |
| CY020493 | A/Queensland/6/2000(H3N2) | 2000 | NCBI |
| CY021061 | A/NewSouthWales/3/1999(H3N2) | 1999 | NCBI |
| CY021077 | A/Albany/42/1975(H3N2) | 1975 | NCBI |
| CY021093 | A/Albany/20/1974(H3N2) | 1974 | NCBI |
| CY021101 | A/Albany/4/1977(H3N2) | 1977 | NCBI |
| CY021109 | A/Albany/17/1968(H3N2) | 1968 | NCBI |
| CY021117 | A/Albany/6/1970(H3N2) | 1970 | NCBI |
| CY021597 | A/Memphis/3/1971(H3N2) | 1971 | NCBI |
| CY021741 | A/SouthAustralia/11/2000(H3N2) | 2000 | NCBI |

| | | | |
|---|---|---|---|
| CY021765 | A/SouthAustralia/53/2005(H3N2) | 2005 | NCBI |
| CY021773 | A/SouthAustralia/72/2000(H3N2) | 2000 | NCBI |
| CY021781 | A/SouthAustralia/81/2000(H3N2) | 2000 | NCBI |
| CY021829 | A/Albany/15/1976(H3N2) | 1976 | NCBI |
| CY021837 | A/Albany/4/1969(H3N2) | 1969 | NCBI |
| CY021845 | A/Albany/10/1968(H3N2) | 1968 | NCBI |
| CY021917 | A/SouthAustralia/16/2000(H3N2) | 2000 | NCBI |
| CY021925 | A/SouthAustralia/47/2000(H3N2) | 2000 | NCBI |
| CY021941 | A/Albany/1/1976(H3N2) | 1976 | NCBI |
| CY021949 | A/SouthAustralia/73/2000(H3N2) | 2000 | NCBI |
| CY021965 | A/Waikato/6/2005(H3N2) | 12/8/2005 | NCBI |
| CY022085 | A/Albany/6/1968(H3N2) | 1968 | NCBI |
| CY022181 | A/Auckland/607/2001(H3N2) | 15/8/2001 | NCBI |
| CY022189 | A/Auckland/611/2002(H3N2) | 9/9/2002 | NCBI |
| CY022197 | A/Auckland/613/2002(H3N2) | 1/8/2002 | NCBI |
| CY022205 | A/Auckland/614/2002(H3N2) | 14/1/2002 | NCBI |
| CY022253 | A/WesternAustralia/4/2000(H3N2) | 20/9/2000 | NCBI |
| CY022309 | A/Memphis/104/1976(H3N2) | 1976 | NCBI |
| CY022501 | A/Auckland/588/2000(H3N2) | 25/9/2000 | NCBI |
| CY022517 | A/Auckland/593/2000(H3N2) | 9/10/2000 | NCBI |
| CY022525 | A/Auckland/602/2001(H3N2) | 19/8/2001 | NCBI |
| CY022549 | A/Auckland/608/2002(H3N2) | 8/6/2002 | NCBI |
| CY022557 | A/Auckland/609/2002(H3N2) | 18/6/2002 | NCBI |
| CY022565 | A/Auckland/612/2002(H3N2) | 14/8/2002 | NCBI |
| CY022573 | A/Auckland/616/2005(H3N2) | 29/9/2005 | NCBI |
| CY022597 | A/Auckland/621/2005(H3N2) | 30/1/2005 | NCBI |
| CY022938 | A/Albany/1/1970(H3N2) | 1970 | NCBI |
| CY022946 | A/Albany/3/1970(H3N2) | 1970 | NCBI |
| CY023002 | A/Auckland/589/2000(H3N2) | 28/9/2000 | NCBI |
| CY023026 | A/Auckland/596/2000(H3N2) | 30/10/2000 | NCBI |
| CY023034 | A/Auckland/599/2000(H3N2) | 22/9/2000 | NCBI |
| CY023042 | A/Auckland/600/2000(H3N2) | 6/5/2000 | NCBI |
| CY023050 | A/Auckland/601/2000(H3N2) | 11/9/2000 | NCBI |
| CY023058 | A/Auckland/603/2001(H3N2) | 2/10/2001 | NCBI |
| CY023066 | A/Auckland/604/2001(H3N2) | 28/9/2001 | NCBI |
| CY023074 | A/Auckland/615/2002(H3N2) | 4/3/2002 | NCBI |
| CY023082 | A/Auckland/617/2005(H3N2) | 28/9/2005 | NCBI |

| | | | |
|---|---|---|---|
| CY019157 | A/NewYork/900/2004(H3N2) | 22/12/2004 | NCBI |
| CY019181 | A/NewYork/905/2004(H3N2) | 27/12/2004 | NCBI |
| CY019253 | A/NewYork/908/2004(H3N2) | 27/12/2004 | NCBI |
| CY019301 | A/NewYork/918/2005(H3N2) | 10/1/2005 | NCBI |
| CY019835 | A/NewYork/928/2006(H3N2) | 28/2/2006 | NCBI |
| CY019843 | A/NewYork/933/2006(H3N2) | 1/3/2006 | NCBI |
| CY020061 | A/NewYork/923/2006(H3N2) | 23/2/2006 | NCBI |
| CY020117 | A/NewYork/938/2006(H3N2) | 2/3/2006 | NCBI |
| CY019165 | A/NewYork/900/2004(H3N2) | 22/12/2004 | NCBI |
| CY019261 | A/NewYork/908/2004(H3N2) | 27/12/2004 | NCBI |
| CY019285 | A/NewYork/913/2005(H3N2) | 10/1/2005 | NCBI |
| CY019309 | A/NewYork/918/2005(H3N2) | 10/1/2005 | NCBI |
| CY019851 | A/NewYork/933/2006(H3N2) | 1/3/2006 | NCBI |
| CY019939 | A/NewYork/928/2006(H3N2) | 28/2/2006 | NCBI |
| CY020069 | A/NewYork/923/2006(H3N2) | 23/2/2006 | NCBI |
| CY020125 | A/NewYork/938/2006(H3N2) | 2/3/2006 | NCBI |
| CY019293 | A/NewYork/913/2005(H3N2) | 10/1/2005 | NCBI |
| CY019317 | A/NewYork/918/2005(H3N2) | 10/1/2005 | NCBI |
| CY019811 | A/NewYork/900/2004(H3N2) | 22/12/2004 | NCBI |
| CY019819 | A/NewYork/908/2004(H3N2) | 27/12/2004 | NCBI |
| CY020077 | A/NewYork/923/2006(H3N2) | 23/2/2006 | NCBI |
| CY020085 | A/NewYork/928/2006(H3N2) | 28/2/2006 | NCBI |
| CY020101 | A/NewYork/933/2006(H3N2) | 1/3/2006 | NCBI |
| CY020133 | A/NewYork/938/2006(H3N2) | 2/3/2006 | NCBI |
| CY020053 | A/NewYork/918/2005(H3N2) | 10/1/2005 | NCBI |
| CY020093 | A/NewYork/928/2006(H3N2) | 28/2/2006 | NCBI |
| CY020109 | A/NewYork/933/2006(H3N2) | 1/3/2006 | NCBI |
| CY020357 | A/NewYork/923/2006(H3N2) | 23/2/2006 | NCBI |
| CY020365 | A/NewYork/938/2006(H3N2) | 2/3/2006 | NCBI |
| CY020533 | A/NewYork/913/2005(H3N2) | 10/1/2005 | NCBI |
| CY021989 | A/NewYork/908/2004(H3N2) | 27/12/2004 | NCBI |
| CY081025 | A/Boston/16/2009(H3N2) | 12/2/2009 | NCBI |
| CY095709 | A/NymcX-157(PuertoRico/8/1934-NewYork/55/2004)(H3N | 2004 | NCBI |
| JN638733 | A/Indiana/08/2011(H3N2) | 27/7/2011 | NCBI |
| JQ070792 | A/Indiana/10/2011(H3N2) | 22/10/2011 | NCBI |
| JQ070784 | A/Indiana/10/2011(H3N2) | 22/10/2011 | NCBI |
| CY104605 | A/KhanhHoa/KH37/2008(mixed) | 24/1/2008 | NCBI |

| | | | |
|---|---|---|---|
| CY088030 | A/England/425/2003(H3N2) | 29/10/2003 | NCBI |
| CY088078 | A/England/754/2003(H3N2) | 2003-11 | NCBI |
| CY088142 | A/Scotland/79/2003(H3N2) | 8/11/2003 | NCBI |
| CY088158 | A/England/470/2003(H3N2) | 5/11/2003 | NCBI |
| CY088166 | A/England/786/2003(H3N2) | 2003-11 | NCBI |
| CY088198 | A/England/516/2003(H3N2) | 20/11/2003 | NCBI |
| CY088238 | A/England/788/2003(H3N2) | 2003-11 | NCBI |
| CY088262 | A/England/484/2003(H3N2) | 26/11/2003 | NCBI |
| CY088286 | A/England/492/2003(H3N2) | 21/11/2003 | NCBI |
| CY088294 | A/England/491/2003(H3N2) | 24/11/2003 | NCBI |
| CY088302 | A/England/493/2003(H3N2) | 23/11/2003 | NCBI |
| CY088310 | A/England/528/2003(H3N2) | 2003-12 | NCBI |
| CY088382 | A/England/532/2003(H3N2) | 27/11/2003 | NCBI |
| CY088438 | A/England/584/2003(H3N2) | 16/11/2003 | NCBI |
| CY107074 | A/England/425/2003(H3N2) | 29/10/2003 | NCBI |
| CY107082 | A/England/754/2003(H3N2) | 6/11/2003 | NCBI |
| CY107090 | A/Scotland/79/2003(H3N2) | 8/11/2003 | NCBI |
| CY107098 | A/England/470/2003(H3N2) | 5/11/2003 | NCBI |
| CY107106 | A/England/786/2003(H3N2) | 27/11/2003 | NCBI |
| CY107114 | A/England/516/2003(H3N2) | 20/11/2003 | NCBI |
| CY107122 | A/England/788/2003(H3N2) | 19/11/2003 | NCBI |
| CY107130 | A/England/484/2003(H3N2) | 26/11/2003 | NCBI |
| CY107138 | A/England/492/2003(H3N2) | 21/11/2003 | NCBI |
| CY107146 | A/England/491/2003(H3N2) | 24/11/2003 | NCBI |
| CY107154 | A/England/493/2003(H3N2) | 23/11/2003 | NCBI |
| CY107162 | A/England/528/2003(H3N2) | 4/12/2003 | NCBI |
| CY107170 | A/England/532/2003(H3N2) | 27/11/2003 | NCBI |
| CY107178 | A/England/584/2003(H3N2) | 16/11/2003 | NCBI |
| CY112369 | A/Auckland/4382/1982(mixed) | 1982 | NCBI |
| CY112564 | A/Madrid/G102/1993(H3N2) | 1993 | NCBI |
| CY117900 | A/Malaysia/23721/2002(mixed) | 2/12/2002 | NCBI |
| CY117960 | A/Malaysia/25804/2003(mixed) | 16/9/2003 | NCBI |
| CY117971 | A/Malaysia/25890/2003(mixed) | 25/9/2003 | NCBI |
| CY121752 | A/Finland/97/2009(H3N2) | 2009 | NCBI |
| CY121776 | A/Finland/97/2009(H3N2) | 2009 | NCBI |
| JX905404 | A/Florida/22/2009(H3N2) | 2009 | NCBI |
| KF014128 | A/Singapore/C2009.485b/2009(H3N2) | 26/6/2009 | NCBI |

| | | | |
|---|---|---|---|
| KF014126 | A/Singapore/C2009.496/2009(H3N2) | 30/6/2009 | NCBI |
| KF014127 | A/Singapore/C2009.485a/2009(H3N2) | 26/6/2009 | NCBI |
| KF014129 | A/Singapore/C2009.515/2009(H3N2) | 7/7/2009 | NCBI |
| KF014130 | A/Singapore/C2009.863/2009(H3N2) | 11/11/2009 | NCBI |
| KF014131 | A/Singapore/C2011.173/2011(H3N2) | 4/3/2011 | NCBI |
| KF014132 | A/Singapore/C2011.244/2011(H3N2) | 30/3/2011 | NCBI |
| KF014133 | A/Singapore/C2011.301/2011(H3N2) | 20/4/2011 | NCBI |
| KF014134 | A/Singapore/C2011.384/2011(H3N2) | 20/5/2011 | NCBI |
| KF014135 | A/Singapore/C2011.411/2011(H3N2) | 30/5/2011 | NCBI |
| KF014136 | A/Singapore/C2011.422/2011(H3N2) | 3/6/2011 | NCBI |
| KF014137 | A/Singapore/C2011.452/2011(H3N2) | 14/6/2011 | NCBI |
| KF014138 | A/Singapore/C2011.458/2011(H3N2) | 16/6/2011 | NCBI |
| KF014139 | A/Singapore/C2011.471/2011(H3N2) | 21/6/2011 | NCBI |
| KF014140 | A/Singapore/C2011.477/2011(H3N2) | 23/6/2011 | NCBI |
| KF014141 | A/Singapore/C2011.493/2011(H3N2) | 29/6/2011 | NCBI |
| KF014142 | A/Singapore/C2011.496/2011(H3N2) | 30/6/2011 | NCBI |
| KF014143 | A/Singapore/C2011.512/2011(H3N2) | 6/7/2011 | NCBI |
| KF014144 | A/Singapore/C2011.518/2011(H3N2) | 8/7/2011 | NCBI |
| KF014146 | A/Singapore/C2011.564/2011(H3N2) | 25/7/2011 | NCBI |
| KF014147 | A/Singapore/C2011.573/2011(H3N2) | 28/7/2011 | NCBI |
| KF014148 | A/Singapore/C2011.614/2011(H3N2) | 12/8/2011 | NCBI |
| KF014149 | A/Singapore/C2011.641/2011(H3N2) | 22/8/2011 | NCBI |
| KF014150 | A/Singapore/C2011.647/2011(H3N2) | 24/8/2011 | NCBI |
| KF014151 | A/Singapore/C2011.803/2011(H3N2) | 20/10/2011 | NCBI |
| KF014152 | A/Singapore/C2011.825/2011(H3N2) | 28/10/2011 | NCBI |
| KF014153 | A/Singapore/H2009.389b/2009(H3N2) | 22/5/2009 | NCBI |
| KF014156 | A/Singapore/H2009.485/2009(H3N2) | 26/6/2009 | NCBI |
| KF014158 | A/Singapore/H2009.510/2009(H3N2) | 5/7/2009 | NCBI |
| KF014159 | A/Singapore/H2009.518/2009(H3N2) | 8/7/2009 | NCBI |
| KF014160 | A/Singapore/H2009.679/2009(H3N2) | 5/9/2009 | NCBI |
| KF014161 | A/Singapore/H2010.211/2010(H3N2) | 18/3/2010 | NCBI |
| KF014162 | A/Singapore/H2010.301/2010(H3N2) | 20/4/2010 | NCBI |
| KF014165 | A/Singapore/H2010.384/2010(H3N2) | 20/5/2010 | NCBI |
| KF014166 | A/Singapore/H2010.389/2010(H3N2) | 22/5/2010 | NCBI |
| KF014170 | A/Singapore/H2010.619/2010(H3N2) | 14/8/2010 | NCBI |
| KF014172 | A/Singapore/H2010.797/2010(H3N2) | 18/10/2010 | NCBI |
| KF014173 | A/Singapore/H2010.822/2010(H3N2) | 27/10/2010 | NCBI |

| | | | |
|---|---|---|---|
| KF014175 | A/Singapore/H2011.447/2011(H3N2) | 12/6/2011 | NCBI |
| KF014176 | A/Singapore/H2011.460a/2011(H3N2) | 17/6/2011 | NCBI |
| KF014178 | A/Singapore/H2011.463/2011(H3N2) | 18/6/2011 | NCBI |
| KF014179 | A/Singapore/H2011.471a/2011(H3N2) | 21/6/2011 | NCBI |
| KF014180 | A/Singapore/H2011.471b/2011(H3N2) | 21/6/2011 | NCBI |
| KF014181 | A/Singapore/H2011.471c/2011(H3N2) | 21/6/2011 | NCBI |
| KF014182 | A/Singapore/H2011.474/2011(H3N2) | 22/6/2011 | NCBI |
| KF014183 | A/Singapore/H2011.479/2011(H3N2) | 24/6/2011 | NCBI |
| KF014184 | A/Singapore/H2011.482/2011(H3N2) | 25/6/2011 | NCBI |
| KF014185 | A/Singapore/H2011.496/2011(H3N2) | 30/6/2011 | NCBI |
| KF014186 | A/Singapore/H2011.499a/2011(H3N2) | 1/7/2011 | NCBI |
| KF014188 | A/Singapore/H2011.507/2011(H3N2) | 4/7/2011 | NCBI |
| KF014190 | A/Singapore/H2011.518/2011(H3N2) | 8/7/2011 | NCBI |
| KF014191 | A/Singapore/H2011.570/2011(H3N2) | 27/7/2011 | NCBI |
| KF014193 | A/Singapore/H2011.751/2011(H3N2) | 1/10/2011 | NCBI |
| KF014194 | A/Singapore/H2011.797/2011(H3N2) | 18/10/2011 | NCBI |
| KF014195 | A/Singapore/H2011.808a/2011(H3N2) | 22/10/2011 | NCBI |
| KF014155 | A/Singapore/H2009.332C/2009(H3N2) | 1/5/2009 | NCBI |
| KF014154 | A/Singapore/H2009.334C/2009(H3N2) | 2/5/2009 | NCBI |
| KF014157 | A/Singapore/H2009.471C/2009(H3N2) | 21/6/2009 | NCBI |
| KF014163 | A/Singapore/H2010.321C/2010(H3N2) | 27/4/2010 | NCBI |
| KF014164 | A/Singapore/H2010.370C/2010(H3N2) | 15/5/2010 | NCBI |
| KF014167 | A/Singapore/H2010.471C/2010(H3N2) | 21/6/2010 | NCBI |
| KF014168 | A/Singapore/H2010.559C/2010(H3N2) | 23/7/2010 | NCBI |
| KF014169 | A/Singapore/H2010.564C/2010(H3N2) | 25/7/2010 | NCBI |
| KF014171 | A/Singapore/H2010.666C/2010(H3N2) | 31/8/2010 | NCBI |
| KF014174 | A/Singapore/H2011.140C/2011(H3N2) | 20/2/2011 | NCBI |
| KF014177 | A/Singapore/H2011.460cC/2011(H3N2) | 17/6/2011 | NCBI |
| KF014192 | A/Singapore/H2011.704C/2011(H3N2) | 14/9/2011 | NCBI |
| KF014196 | A/Singapore/H2011.808bC/2011(H3N2) | 22/10/2011 | NCBI |
| KF014145 | A/Singapore/C2011.545C/2011(H3N2) | 18/7/2011 | NCBI |
| KF014198 | A/Singapore/C2009.458V/2009(H3N2) | 16/6/2009 | NCBI |
| KF014203 | A/Singapore/C2010.937V/2010(H3N2) | 8/12/2010 | NCBI |
| KF014204 | A/Singapore/C2011.027V/2011(H3N2) | 10/1/2011 | NCBI |
| KF014208 | A/Singapore/C2011.362V/2011(H3N2) | 12/5/2011 | NCBI |
| KF014218 | A/Singapore/C2011.507V/2011(H3N2) | 4/7/2011 | NCBI |
| KF014223 | A/Singapore/C2011.584V/2011(H3N2) | 1/8/2011 | NCBI |

| | | | |
|---|---|---|---|
| KF790212 | A/Alaska/02/2013(H3N2) | 20/2/2013 | NCBI |
| CY163648 | A/Wisconsin/67/2005(H3N2) | 2005 | NCBI |
| CY163656 | A/Wisconsin/67/2005(H3N2) | 2005 | NCBI |
| CY163664 | A/Wisconsin/67/2005(H3N2) | 2005 | NCBI |
| CY163680 | A/Wisconsin/67/2005(H3N2) | 2005 | NCBI |
| CY163688 | A/Wisconsin/67/2005(H3N2) | 2005 | NCBI |
| CY163696 | A/Wisconsin/67/2005(H3N2) | 2005 | NCBI |
| CY163704 | A/Wisconsin/67/2005(H3N2) | 2005 | NCBI |
| CY163712 | A/Wisconsin/67/2005(H3N2) | 2005 | NCBI |
| CY163720 | A/Wisconsin/67/2005(H3N2) | 2005 | NCBI |
| CY163728 | A/Wisconsin/67/2005(H3N2) | 2005 | NCBI |
| CY163736 | A/Wisconsin/67/2005(H3N2) | 2005 | NCBI |
| CY163744 | A/Wisconsin/67/2005(H3N2) | 2005 | NCBI |
| CY163872 | A/Wisconsin/67/2005(H3N2) | 2005 | NCBI |
| CY163880 | A/Wisconsin/67/2005(H3N2) | 2005 | NCBI |
| CY163888 | A/Wisconsin/67/2005(H3N2) | 2005 | NCBI |
| CY163896 | A/Wisconsin/67/2005(H3N2) | 2005 | NCBI |
| CY163904 | A/Wisconsin/67/2005(H3N2) | 2005 | NCBI |
| CY163912 | A/Wisconsin/67/2005(H3N2) | 2005 | NCBI |
| CY163920 | A/Wisconsin/67/2005(H3N2) | 2005 | NCBI |
| CY163928 | A/Wisconsin/67/2005(H3N2) | 2005 | NCBI |
| CY163936 | A/Wisconsin/67/2005(H3N2) | 2005 | NCBI |
| CY163944 | A/Wisconsin/67/2005(H3N2) | 2005 | NCBI |
| CY163952 | A/Wisconsin/67/2005(H3N2) | 2005 | NCBI |
| CY163960 | A/Wisconsin/67/2005(H3N2) | 2005 | NCBI |
| CY163968 | A/Wisconsin/67/2005(H3N2) | 2005 | NCBI |
| CY163976 | A/Wisconsin/67/2005(H3N2) | 2005 | NCBI |
| CY163984 | A/Wisconsin/67/2005(H3N2) | 2005 | NCBI |
| CY163992 | A/Wisconsin/67/2005(H3N2) | 2005 | NCBI |
| CY164000 | A/Wisconsin/67/2005(H3N2) | 2005 | NCBI |
| CY164008 | A/Wisconsin/67/2005(H3N2) | 2005 | NCBI |
| CY164016 | A/Wisconsin/67/2005(H3N2) | 2005 | NCBI |
| CY164024 | A/Wisconsin/67/2005(H3N2) | 2005 | NCBI |
| CY164032 | A/Wisconsin/67/2005(H3N2) | 2005 | NCBI |
| CY164040 | A/Wisconsin/67/2005(H3N2) | 2005 | NCBI |
| CY164048 | A/Wisconsin/67/2005(H3N2) | 2005 | NCBI |
| CY164056 | A/Wisconsin/67/2005(H3N2) | 2005 | NCBI |

| | | | |
|---|---|---|---|
| CY164064 | A/Wisconsin/67/2005(H3N2) | 2005 | NCBI |
| CY164072 | A/Wisconsin/67/2005(H3N2) | 2005 | NCBI |
| CY164080 | A/Wisconsin/67/2005(H3N2) | 2005 | NCBI |
| CY164088 | A/Wisconsin/67/2005(H3N2) | 2005 | NCBI |
| CY164096 | A/Wisconsin/67/2005(H3N2) | 2005 | NCBI |
| CY164104 | A/Wisconsin/67/2005(H3N2) | 2005 | NCBI |
| CY164112 | A/Wisconsin/67/2005(H3N2) | 2005 | NCBI |
| KR057600 | A/Hawaii/36/2014(H3N2) | 8/8/2014 | NCBI |
| KM438090 | A/Quebec/26-031205/2005(H3N2) | 3/12/2005 | NCBI |
| KT837039 | A/Florida/35/2014(H3N2) | 23/9/2014 | NCBI |
| KT837205 | A/Florida/35/2014(H3N2) | 23/9/2014 | NCBI |
| KT837098 | A/Minnesota/22/2014(H3N2) | 20/9/2014 | NCBI |
| KT837116 | A/Pennsylvania/40/2014(H3N2) | 13/9/2014 | NCBI |
| KT837240 | A/Minnesota/21/2014(H3N2) | 17/9/2014 | NCBI |
| KT837148 | A/NewMexico/27/2014(H3N2) | 18/9/2014 | NCBI |
| KT837701 | A/Connecticut/21/2014(H3N2) | 14/10/2014 | NCBI |
| KT837429 | A/Florida/45/2014(H3N2) | 12/10/2014 | NCBI |
| KT838115 | A/Louisiana/14/2014(H3N2) | 22/10/2014 | NCBI |
| KT837480 | A/Louisiana/08/2014(H3N2) | 11/10/2014 | NCBI |
| KT838021 | A/NewHampshire/21/2014(H3N2) | 17/10/2014 | NCBI |
| KT838085 | A/Texas/42/2014(H3N2) | 22/10/2014 | NCBI |
| KT837987 | A/Florida/38/2014(H3N2) | 7/10/2014 | NCBI |
| KT838086 | A/Florida/37/2014(H3N2) | 7/10/2014 | NCBI |
| KT837746 | A/Texas/37/2014(H3N2) | 1/10/2014 | NCBI |
| KT837526 | A/Texas/37/2014(H3N2) | 1/10/2014 | NCBI |
| KT838053 | A/Florida/39/2014(H3N2) | 7/10/2014 | NCBI |
| KT838093 | A/Mississippi/08/2014(H3N2) | 23/10/2014 | NCBI |
| KT837973 | A/Louisiana/12/2014(H3N2) | 21/10/2014 | NCBI |
| KT837965 | A/Florida/40/2014(H3N2) | 7/10/2014 | NCBI |
| KT838111 | A/Mississippi/03/2014(H3N2) | 15/10/2014 | NCBI |
| KT838266 | A/Colorado/15/2014(H3N2) | 4/10/2014 | NCBI |
| KT838178 | A/Mississippi/04/2014(H3N2) | 16/10/2014 | NCBI |
| KT838234 | A/Mississippi/05/2014(H3N2) | 14/10/2014 | NCBI |
| KT838215 | A/Wyoming/08/2014(H3N2) | 9/10/2014 | NCBI |
| KT838071 | A/Maine/04/2014(H3N2) | 9/10/2014 | NCBI |
| KU590422 | A/Missouri/42/2015(H3N2) | 4/10/2015 | NCBI |
| KU589464 | A/Tennessee/20/2015(H3N2) | 19/8/2015 | NCBI |

| | | | |
|---|---|---|---|
| KU590960 | A/NewMexico/26/2015(H3N2) | 13/9/2015 | NCBI |
| KU589572 | A/Michigan/53/2015(H3N2) | 12/9/2015 | NCBI |
| KU589944 | A/Michigan/54/2015(H3N2) | 19/9/2015 | NCBI |
| KU589588 | A/Michigan/55/2015(H3N2) | 18/9/2015 | NCBI |
| KU589596 | A/Idaho/25/2015(H3N2) | 18/9/2015 | NCBI |
| KU589604 | A/Vermont/08/2015(H3N2) | 19/8/2015 | NCBI |
| KU589612 | A/Florida/80/2015(H3N2) | 28/10/2015 | NCBI |
| KU591723 | A/Michigan/67/2015(H3N2) | 10/10/2015 | NCBI |
| KU589683 | A/Alaska/151/2015(H3N2) | 21/7/2015 | NCBI |
| KU589691 | A/Alaska/152/2015(H3N2) | 20/7/2015 | NCBI |
| KU590124 | A/Alaska/154/2015(H3N2) | 24/7/2015 | NCBI |
| KU589707 | A/Alaska/155/2015(H3N2) | 8/8/2015 | NCBI |
| KU589715 | A/Alaska/156/2015(H3N2) | 3/8/2015 | NCBI |
| KU589723 | A/Ohio/27/2015(H3N2) | 17/9/2015 | NCBI |
| KU589731 | A/Ohio/28/2015(H3N2) | 17/9/2015 | NCBI |
| KU589739 | A/Massachusetts/26/2015(H3N2) | 24/9/2015 | NCBI |
| KU589747 | A/NewMexico/23/2015(H3N2) | 12/9/2015 | NCBI |
| KU589755 | A/NewMexico/25/2015(H3N2) | 10/9/2015 | NCBI |
| KU589763 | A/Iowa/40/2015(H3N2) | 4/9/2015 | NCBI |
| KU589771 | A/Iowa/41/2015(H3N2) | 10/9/2015 | NCBI |
| KU590279 | A/Iowa/42/2015(H3N2) | 10/9/2015 | NCBI |
| KU589785 | A/Iowa/43/2015(H3N2) | 14/9/2015 | NCBI |
| KU589793 | A/Wyoming/18/2015(H3N2) | 22/9/2015 | NCBI |
| KU590433 | A/Massachusetts/25/2015(H3N2) | 10/9/2015 | NCBI |
| KU590441 | A/Indiana/17/2015(H3N2) | 31/8/2015 | NCBI |
| KU590449 | A/Indiana/19/2015(H3N2) | 1/9/2015 | NCBI |
| KU590473 | A/Indiana/22/2015(H3N2) | 1/9/2015 | NCBI |
| KU590155 | A/Indiana/25/2015(H3N2) | 3/9/2015 | NCBI |
| KU590163 | A/NewHampshire/39/2015(H3N2) | 9/9/2015 | NCBI |
| KU590171 | A/NewHampshire/40/2015(H3N2) | 14/9/2015 | NCBI |
| KU590178 | A/Minnesota/41/2015(H3N2) | 14/9/2015 | NCBI |
| KU590186 | A/Florida/69/2015(H3N2) | 15/9/2015 | NCBI |
| KU590194 | A/Florida/70/2015(H3N2) | 15/9/2015 | NCBI |
| KU590202 | A/Florida/71/2015(H3N2) | 15/9/2015 | NCBI |
| KU589904 | A/Kentucky/15/2015(H3N2) | 28/9/2015 | NCBI |
| KU590210 | A/Florida/72/2015(H3N2) | 15/9/2015 | NCBI |
| KU590226 | A/Wisconsin/81/2015(H3N2) | 18/9/2015 | NCBI |

| | | | |
|---|---|---|---|
| KU590234 | A/Wisconsin/82/2015(H3N2) | 17/9/2015 | NCBI |
| KU590242 | A/Tennessee/21/2015(H3N2) | 15/9/2015 | NCBI |
| KU590744 | A/Montana/28/2015(H3N2) | 24/9/2015 | NCBI |
| KU590752 | A/Montana/29/2015(H3N2) | 26/9/2015 | NCBI |
| KU589974 | A/Alaska/157/2015(H3N2) | 13/8/2015 | NCBI |
| KU590313 | A/Alaska/158/2015(H3N2) | 14/8/2015 | NCBI |
| KU590329 | A/Alaska/162/2015(H3N2) | 19/8/2015 | NCBI |
| KU590218 | A/Wisconsin/80/2015(H3N2) | 14/9/2015 | NCBI |
| KU590358 | A/NorthCarolina/32/2015(H3N2) | 28/9/2015 | NCBI |
| KU590465 | A/Indiana/21/2015(H3N2) | 1/9/2015 | NCBI |
| KU590762 | A/Alaska/222/2015(H3N2) | 8/9/2015 | NCBI |
| KU590778 | A/Alaska/239/2015(H3N2) | 16/9/2015 | NCBI |
| KU591884 | A/Hawaii/62/2015(H3N2) | 28/9/2015 | NCBI |
| KU591645 | A/Florida/86/2015(H3N2) | 2/11/2015 | NCBI |
| KU590828 | A/California/98/2015(H3N2) | 19/10/2015 | NCBI |
| KU591342 | A/Texas/71/2015(H3N2) | 11/11/2015 | NCBI |
| KU592096 | A/Hawaii/69/2015(H3N2) | 4/10/2015 | NCBI |
| KU592104 | A/Hawaii/70/2015(H3N2) | 5/10/2015 | NCBI |
| KU591606 | A/Hawaii/72/2015(H3N2) | 8/10/2015 | NCBI |
| KU591614 | A/Florida/82/2015(H3N2) | 26/10/2015 | NCBI |
| KU591661 | A/SouthCarolina/10/2015(H3N2) | 6/10/2015 | NCBI |
| KU591962 | A/Washington/57/2015(H3N2) | 7/11/2015 | NCBI |
| KU591975 | A/Michigan/60/2015(H3N2) | 5/10/2015 | NCBI |
| KX003377 | A/Michigan/60/2015(H3N2) | 5/10/2015 | NCBI |
| KX003437 | A/Nevada/37/2015(H3N2) | 13/11/2015 | NCBI |
| KX003771 | A/Alaska/119/2015(H3N2) | 8/7/2015 | NCBI |
| KU590298 | A/NewJersey/52/2015(H3N2) | 1/9/2015 | NCBI |
| KX416168 | A/Michigan/50/2016(H3N2) | 16/2/2016 | NCBI |
| KX417054 | A/Nevada/22/2016(H3N2) | 4/3/2016 | NCBI |
| KX411973 | A/Arizona/22/2015(H3N2) | 7/10/2015 | NCBI |
| KX412037 | A/Iowa/44/2015(H3N2) | 21/10/2015 | NCBI |
| KX412053 | A/Iowa/46/2015(H3N2) | 21/10/2015 | NCBI |
| KX412355 | A/Pennsylvania/33/2015(H3N2) | 21/9/2015 | NCBI |
| KX412494 | A/Texas/72/2015(H3N2) | 8/11/2015 | NCBI |
| KX412526 | A/Texas/74/2015(H3N2) | 21/12/2015 | NCBI |
| KX412638 | A/Pennsylvania/52/2015(H3N2) | 26/12/2015 | NCBI |
| KX412710 | A/Washington/69/2015(H3N2) | 28/12/2015 | NCBI |

| | | | |
|---|---|---|---|
| KX412758 | A/NorthCarolina/04/2016(H3N2) | 13/1/2016 | NCBI |
| KX412806 | A/Georgia/03/2016(H3N2) | 12/1/2016 | NCBI |
| KX412854 | A/Pennsylvania/03/2016(H3N2) | 23/1/2016 | NCBI |
| KX412870 | A/Texas/06/2016(H3N2) | 20/1/2016 | NCBI |
| KX413230 | A/Texas/83/2016(H3N2) | 26/2/2016 | NCBI |
| KX413254 | A/Texas/88/2016(H3N2) | 24/2/2016 | NCBI |
| KX413303 | A/Michigan/66/2015(H3N2) | 15/10/2015 | NCBI |
| KX413323 | A/Iowa/54/2015(H3N2) | 1/11/2015 | NCBI |
| KX413393 | A/Pennsylvania/45/2015(H3N2) | 12/11/2015 | NCBI |
| KX413409 | A/Florida/89/2015(H3N2) | 14/11/2015 | NCBI |
| KX413441 | A/California/111/2015(H3N2) | 20/11/2015 | NCBI |
| KX413473 | A/Arizona/29/2015(H3N2) | 27/11/2015 | NCBI |
| KX413545 | A/Minnesota/51/2015(H3N2) | 14/12/2015 | NCBI |
| KX413585 | A/Hawaii/78/2015(H3N2) | 5/12/2015 | NCBI |
| KX413593 | A/Hawaii/79/2015(H3N2) | 11/12/2015 | NCBI |
| KX413625 | A/California/117/2015(H3N2) | 5/12/2015 | NCBI |
| KX413681 | A/Minnesota/54/2015(H3N2) | 20/12/2015 | NCBI |
| KX413745 | A/Michigan/78/2015(H3N2) | 14/12/2015 | NCBI |
| KX413777 | A/Arizona/44/2015(H3N2) | 21/12/2015 | NCBI |
| KX413881 | A/California/130/2015(H3N2) | 29/12/2015 | NCBI |
| KX414009 | A/Washington/04/2016(H3N2) | 8/1/2016 | NCBI |
| KX414025 | A/Washington/05/2016(H3N2) | 12/1/2016 | NCBI |
| KX414105 | A/Indiana/10/2016(H3N2) | 19/1/2016 | NCBI |
| KX414113 | A/Indiana/11/2016(H3N2) | 17/1/2016 | NCBI |
| KX414249 | A/Washington/17/2016(H3N2) | 19/1/2016 | NCBI |
| KX414265 | A/Washington/18/2016(H3N2) | 20/1/2016 | NCBI |
| KX916204 | A/Idaho/33/2016(H3N2) | 7/6/2016 | NCBI |
| KX917164 | A/Idaho/16/2016(H3N2) | 7/3/2016 | NCBI |
| KX917324 | A/Wyoming/15/2016(H3N2) | 23/3/2016 | NCBI |
| KX916140 | A/Pennsylvania/73/2016(H3N2) | 4/5/2016 | NCBI |
| KX916172 | A/Hawaii/49/2016(H3N2) | 18/4/2016 | NCBI |
| KX916452 | A/California/118/2016(H3N2) | 30/5/2016 | NCBI |
| KX916516 | A/SouthCarolina/15/2016(H3N2) | 11/5/2016 | NCBI |
| KX916628 | A/SouthCarolina/17/2016(H3N2) | 23/5/2016 | NCBI |
| KX916764 | A/Washington/74/2016(H3N2) | 22/4/2016 | NCBI |
| KX917068 | A/Kentucky/06/2016(H3N2) | 19/2/2016 | NCBI |
| KX917404 | A/NewYork/62/2016(H3N2) | 11/4/2016 | NCBI |

| | | | |
|---|---|---|---|
| KX917428 | A/Mississippi/16/2016(H3N2) | 18/4/2016 | NCBI |
| KY116230 | A/Pennsylvania/89/2016(H3N2) | 27/9/2016 | NCBI |
| KY116254 | A/Florida/76/2016(H3N2) | 19/9/2016 | NCBI |
| KY116350 | A/Minnesota/73/2016(H3N2) | 4/10/2016 | NCBI |
| KY116454 | A/Florida/79/2016(H3N2) | 3/10/2016 | NCBI |
| KY116934 | A/Alaska/232/2015(H3N2) | 9/9/2015 | NCBI |
| KY115830 | A/NorthCarolina/54/2016(H3N2) | 20/9/2016 | NCBI |
| KY115854 | A/NorthCarolina/53/2016(H3N2) | 8/9/2016 | NCBI |
| KY115910 | A/Michigan/104/2016(H3N2) | 17/9/2016 | NCBI |
| KY115934 | A/Wisconsin/83/2016(H3N2) | 24/9/2016 | NCBI |
| KY116110 | A/Maine/22/2016(H3N2) | 17/9/2016 | NCBI |
| KY116150 | A/NewYork/99/2016(H3N2) | 23/9/2016 | NCBI |
| KY116174 | A/NewYork/98/2016(H3N2) | 21/9/2016 | NCBI |
| KY116190 | A/Wisconsin/85/2016(H3N2) | 2/10/2016 | NCBI |
| KY289632 | A/California/140/2016(H3N2) | 17/10/2016 | NCBI |
| KY289109 | A/Florida/81/2016(H3N2) | 3/10/2016 | NCBI |
| KY348531 | A/HongKong/01/1968(H3N2) | 1968 | NCBI |
| CY210385 | A/Delaware/32/2016(H3N2) | 15/11/2016 | NCBI |
| CY218937 | A/Washington/90/2016(H3N2) | 5/12/2016 | NCBI |
| CY220055 | A/Georgia/01/2017(H3N2) | 10/1/2017 | NCBI |
| CY222709 | A/California/13/2017(H3N2) | 23/1/2017 | NCBI |
| CY223822 | A/RhodeIsland/04/2017(H3N2) | 30/1/2017 | NCBI |
| CY223862 | A/Maine/11/2017(H3N2) | 15/2/2017 | NCBI |
| CY226639 | A/Tennessee/06/2017(H3N2) | 4/1/2017 | NCBI |
| CY226855 | A/Washington/14/2017(H3N2) | 9/2/2017 | NCBI |
| CY226887 | A/Wisconsin/17/2017(H3N2) | 13/2/2017 | NCBI |
| CY226950 | A/Texas/208/2016(H3N2) | 19/12/2016 | NCBI |
| CY224934 | A/Connecticut/08/2017(H3N2) | 31/1/2017 | NCBI |
| CY225006 | A/Delaware/10/2017(H3N2) | 9/2/2017 | NCBI |
| CY225022 | A/Delaware/11/2017(H3N2) | 9/2/2017 | NCBI |
| CY227270 | A/NewYork/12/2017(H3N2) | 21/2/2017 | NCBI |
| CY227310 | A/NewJersey/17/2017(H3N2) | 27/2/2017 | NCBI |
| CY230029 | A/Michigan/26/2017(H3N2) | 18/2/2017 | NCBI |
| CY230077 | A/Michigan/31/2017(H3N2) | 7/3/2017 | NCBI |
| CY230125 | A/Michigan/32/2017(H3N2) | 9/3/2017 | NCBI |
| CY230221 | A/Wisconsin/28/2017(H3N2) | 7/3/2017 | NCBI |
| CY227886 | A/Louisiana/15/2017(H3N2) | 16/3/2017 | NCBI |

| | | | |
|---|---|---|---|
| CY227974 | A/Delaware/33/2016(H3N2) | 15/11/2016 | NCBI |
| CY228421 | A/Florida/22/2017(H3N2) | 20/3/2017 | NCBI |
| CY228437 | A/Florida/23/2017(H3N2) | 23/3/2017 | NCBI |
| CY228477 | A/Massachusetts/10/2017(H3N2) | 9/3/2017 | NCBI |
| CY228501 | A/Virginia/20/2017(H3N2) | 17/3/2017 | NCBI |
| CY228525 | A/Vermont/10/2017(H3N2) | 13/3/2017 | NCBI |
| CY228557 | A/Tennessee/31/2017(H3N2) | 28/3/2017 | NCBI |
| CY228573 | A/Delaware/23/2017(H3N2) | 29/3/2017 | NCBI |
| CY228669 | A/Maine/17/2017(H3N2) | 30/3/2017 | NCBI |
| CY228789 | A/NewHampshire/17/2017(H3N2) | 28/3/2017 | NCBI |
| CY238266 | A/Texas/84/2017(H3N2) | 7/2/2017 | NCBI |
| CY238449 | A/Texas/90/2017(H3N2) | 14/2/2017 | NCBI |
| CY238513 | A/Texas/98/2017(H3N2) | 12/3/2017 | NCBI |
| CY235504 | A/Alaska/13/2017(H3N2) | 3/4/2017 | NCBI |
| CY235544 | A/Nevada/10/2017(H3N2) | 22/3/2017 | NCBI |
| CY235560 | A/Nevada/11/2017(H3N2) | 25/3/2017 | NCBI |
| CY235576 | A/Hawaii/19/2017(H3N2) | 27/3/2017 | NCBI |
| CY235600 | A/Arizona/24/2017(H3N2) | 9/4/2017 | NCBI |
| CY235624 | A/Washington/40/2017(H3N2) | 17/4/2017 | NCBI |
| CY235640 | A/Alaska/16/2017(H3N2) | 22/4/2017 | NCBI |
| CY235656 | A/Utah/26/2017(H3N2) | 24/4/2017 | NCBI |
| CY235672 | A/Utah/27/2017(H3N2) | 25/4/2017 | NCBI |
| CY235688 | A/Nevada/12/2017(H3N2) | 25/4/2017 | NCBI |
| CY235704 | A/Arizona/28/2017(H3N2) | 29/4/2017 | NCBI |
| CY235728 | A/Hawaii/21/2017(H3N2) | 6/4/2017 | NCBI |
| CY235760 | A/Idaho/25/2017(H3N2) | 28/4/2017 | NCBI |
| CY242118 | A/Wisconsin/19/2017(H3N2) | 23/1/2017 | NCBI |
| CY242134 | A/Texas/71/2017(H3N2) | 18/3/2017 | NCBI |
| CY255604 | A/SouthCarolina/16/2017(H3N2) | 4/9/2017 | NCBI |
| CY250332 | A/Maryland/47/2017(H3N2) | 11/9/2017 | NCBI |
| CY250483 | A/Wisconsin/328/2017(H3N2) | 25/9/2017 | NCBI |
| CY250491 | A/Wisconsin/327/2017(H3N2) | 22/9/2017 | NCBI |
| CY250499 | A/Pennsylvania/263/2017(H3N2) | 1/10/2017 | NCBI |
| CY250507 | A/Missouri/30/2017(H3N2) | 26/9/2017 | NCBI |
| CY266018 | A/Illinois/40/2017(H3N2) | 8/10/2017 | NCBI |
| CY266034 | A/Illinois/39/2017(H3N2) | 4/10/2017 | NCBI |
| CY266050 | A/Florida/84/2017(H3N2) | 23/10/2017 | NCBI |

| | | | |
|---|---|---|---|
| CY266066 | A/Florida/85/2017(H3N2) | 24/10/2017 | NCBI |
| CY266082 | A/Florida/87/2017(H3N2) | 25/10/2017 | NCBI |
| CY266098 | A/Florida/86/2017(H3N2) | 24/10/2017 | NCBI |
| CY266114 | A/NorthCarolina/40/2017(H3N2) | 7/10/2017 | NCBI |
| CY266130 | A/Florida/88/2017(H3N2) | 8/11/2017 | NCBI |
| CY266146 | A/Florida/89/2017(H3N2) | 9/11/2017 | NCBI |
| CY266162 | A/Florida/90/2017(H3N2) | 9/11/2017 | NCBI |
| CY266178 | A/Florida/93/2017(H3N2) | 13/11/2017 | NCBI |
| MG914107 | InfluenzaAvirus | 13/12/2017 | NCBI |
| MG914115 | InfluenzaAvirus | 19/12/2017 | NCBI |
| MG914123 | InfluenzaAvirus | 3/10/2017 | NCBI |
| MG914131 | InfluenzaAvirus | 11/10/2017 | NCBI |
| MG914139 | InfluenzaAvirus | 23/10/2017 | NCBI |
| MG914147 | InfluenzaAvirus | 6/11/2017 | NCBI |
| MG914155 | InfluenzaAvirus | 6/11/2017 | NCBI |
| MG914163 | InfluenzaAvirus | 8/11/2017 | NCBI |
| MG914171 | InfluenzaAvirus | 6/12/2017 | NCBI |
| MG928493 | InfluenzaAvirus | 2017-06 | NCBI |
| MG928485 | InfluenzaAvirus | 20/2/2017 | NCBI |
| MG928477 | InfluenzaAvirus | 20/2/2017 | NCBI |
| MG928445 | InfluenzaAvirus | 2017-02 | NCBI |
| MG928469 | InfluenzaAvirus | 20/2/2017 | NCBI |
| MG928461 | InfluenzaAvirus | 20/2/2017 | NCBI |
| MG928453 | InfluenzaAvirus | 20/2/2017 | NCBI |
| EPI347506 | A/Kentucky/05/2011(H3N2) | 4/11/2011 | GISAID |
| EPI347544 | A/Utah/12/2011(H3N2) | 6/11/2011 | GISAID |
| EPI348478 | A/ChiangRai/277/2011(H3N2) | 20/9/2011 | GISAID |
| EPI349103 | A/Victoria/361/2011(H3N2) | 24/10/2011 | GISAID |
| EPI353512 | A/HongKong/2000/2009(H3N2) | 7/4/2009 | GISAID |
| EPI353514 | A/Kentucky/05/2011(H3N2) | 4/11/2011 | GISAID |
| EPI353906 | A/Victoria/361/2011(H3N2) | 24/10/2011 | GISAID |
| EPI357622 | A/GOROKA/5/2011(H3N2) | 27/7/2011 | GISAID |
| EPI357660 | A/PERTH/219/2011(H3N2) | 17/9/2011 | GISAID |
| EPI357669 | A/BRISBANE/382/2011(H3N2) | 6/9/2011 | GISAID |
| EPI357675 | A/BRISBANE/395/2011(H3N2) | 20/10/2011 | GISAID |
| EPI363012 | A/Washington/16/2012(H3N2) | 2/3/2012 | GISAID |
| EPI363019 | A/California/17/2012(H3N2) | 25/1/2012 | GISAID |

| | | | |
|---|---|---|---|
| EPI370074 | A/VICTORIA/893/2011(H3N2) | 7/12/2011 | GISAID |
| EPI370111 | A/PHILIPPINES/3969/2011(H3N2) | 25/10/2011 | GISAID |
| EPI370132 | A/PERTH/305/2011(H3N2) | 12/12/2011 | GISAID |
| EPI370318 | A/CAMBODIA/74/2011(H3N2) | 17/11/2011 | GISAID |
| EPI370329 | A/PHILIPPINES/3948/2011(H3N2) | 21/10/2011 | GISAID |
| EPI371776 | A/Nevada/12/2012(H3N2) | 30/3/2012 | GISAID |
| EPI371792 | A/California/22/2012(H3N2) | 20/2/2012 | GISAID |
| EPI371802 | A/NewMexico/07/2012(H3N2) | 16/2/2012 | GISAID |
| EPI371819 | A/Michigan/03/2012(H3N2) | 6/3/2012 | GISAID |
| EPI372003 | A/RhodeIsland/01/2012(H3N2) | 30/1/2012 | GISAID |
| EPI376886 | A/TASMANIA/1/2012(H3N2) | 20/2/2012 | GISAID |
| EPI376894 | A/TOWNSVILLE/1/2012(H3N2) | 21/1/2012 | GISAID |
| EPI377004 | A/SOUTHAUSTRALIA/313/2011(H3N2) | 18/11/2011 | GISAID |
| EPI378515 | A/Indiana/07/2012(H3N2) | 16/7/2012 | GISAID |
| EPI378523 | A/Indiana/09/2012(H3N2) | 16/7/2012 | GISAID |
| EPI379447 | A/PERTH/41/2012(H3N2) | 8/4/2012 | GISAID |
| EPI379508 | A/GOROKA/5/2012(H3N2) | 5/3/2012 | GISAID |
| EPI379585 | A/VICTORIA/802/2012(H3N2) | 9/5/2012 | GISAID |
| EPI384818 | A/RhodeIsland/01/2012(H3N2) | 30/1/2012 | GISAID |
| EPI384820 | A/Ohio/02/2012(H3N2) | 8/3/2012 | GISAID |
| EPI388305 | A/Ohio/14/2012(H3N2) | 29/7/2012 | GISAID |
| EPI388339 | A/Ohio/16/2012(H3N2) | 28/7/2012 | GISAID |
| EPI388346 | A/Indiana/15/2012(H3N2) | 29/7/2012 | GISAID |
| EPI388352 | A/Indiana/20/2012(H3N2) | 29/7/2012 | GISAID |
| EPI393298 | A/Indiana/21/2012(H3N2) | 31/7/2012 | GISAID |
| EPI397997 | A/Ohio/17/2012(H3N2) | 28/7/2012 | GISAID |
| EPI392708 | A/Indiana/59/2012(H3N2) | 2/8/2012 | GISAID |
| EPI393306 | A/Indiana/18/2012(H3N2) | 29/7/2012 | GISAID |
| EPI393311 | A/WestVirginia/09/2012(H3N2) | 6/8/2012 | GISAID |
| EPI393317 | A/Ohio/38/2012(H3N2) | 4/8/2012 | GISAID |
| EPI393511 | A/Ohio/52/2012(H3N2) | 5/8/2012 | GISAID |
| EPI393519 | A/Indiana/65/2012(H3N2) | 2/8/2012 | GISAID |
| EPI393661 | A/Ohio/62/2012(H3N2) | 13/8/2012 | GISAID |
| EPI393667 | A/Ohio/28/2012(H3N2) | 1/8/2012 | GISAID |
| EPI393673 | A/Ohio/71/2012(H3N2) | 13/8/2012 | GISAID |
| EPI393681 | A/WestVirginia/10/2012(H3N2) | 6/8/2012 | GISAID |
| EPI393688 | A/Ohio/23/2012(H3N2) | 30/7/2012 | GISAID |

| | | | |
|---|---|---|---|
| EPI393695 | A/WestVirginia/16/2012(H3N2) | 9/8/2012 | GISAID |
| EPI393702 | A/Ohio/64/2012(H3N2) | 10/8/2012 | GISAID |
| EPI394077 | A/Wisconsin/27/2012(H3N2) | 17/8/2012 | GISAID |
| EPI394116 | A/TASMANIA/12/2012(H3N2) | 24/6/2012 | GISAID |
| EPI394124 | A/NEWCALEDONIA/1/2012(H3N2) | 26/3/2012 | GISAID |
| EPI394138 | A/TASMANIA/26/2012(H3N2) | 26/6/2012 | GISAID |
| EPI394149 | A/CHRISTCHURCH/503/2012(H3N2) | 3/7/2012 | GISAID |
| EPI394152 | A/CHRISTCHURCH/516/2012(H3N2) | 7/7/2012 | GISAID |
| EPI394913 | A/Indiana/132/2012(H3N2) | 3/8/2012 | GISAID |
| EPI394921 | A/Ohio/69/2012(H3N2) | 13/8/2012 | GISAID |
| EPI394933 | A/Wisconsin/23/2012(H3N2) | 15/8/2012 | GISAID |
| EPI394937 | A/Ohio/75/2012(H3N2) | 16/8/2012 | GISAID |
| EPI394944 | A/Maryland/22/2012(H3N2) | 16/8/2012 | GISAID |
| EPI395259 | A/Ohio/35/2012(H3N2) | 4/8/2012 | GISAID |
| EPI395267 | A/Ohio/65/2012(H3N2) | 11/8/2012 | GISAID |
| EPI397210 | A/Maryland/25/2012(H3N2) | 18/8/2012 | GISAID |
| EPI395344 | A/Wisconsin/28/2012(H3N2) | 28/8/2012 | GISAID |
| EPI395351 | A/Michigan/20/2012(H3N2) | 16/8/2012 | GISAID |
| EPI395357 | A/Pennsylvania/17/2012(H3N2) | 23/8/2012 | GISAID |
| EPI397344 | A/Michigan/19/2012(H3N2) | 16/8/2012 | GISAID |
| EPI396175 | A/Ohio/80/2012(H3N2) | 19/8/2012 | GISAID |
| EPI396183 | A/Indiana/77/2012(H3N2) | 2/8/2012 | GISAID |
| EPI396192 | A/Ohio/77/2012(H3N2) | 15/8/2012 | GISAID |
| EPI397006 | A/Ohio/78/2012(H3N2) | 9/8/2012 | GISAID |
| EPI396200 | A/Ohio/79/2012(H3N2) | 9/8/2012 | GISAID |
| EPI396208 | A/Indiana/149/2012(H3N2) | 5/8/2012 | GISAID |
| EPI397211 | A/Indiana/113/2012(H3N2) | 5/8/2012 | GISAID |
| EPI397340 | A/Indiana/150/2012(H3N2) | 8/8/2012 | GISAID |
| EPI397092 | A/Indiana/57/2012(H3N2) | 2/8/2012 | GISAID |
| EPI396230 | A/Indiana/147/2012(H3N2) | 9/8/2012 | GISAID |
| EPI397206 | A/Ohio/34/2012(H3N2) | 3/8/2012 | GISAID |
| EPI396239 | A/Indiana/103/2012(H3N2) | 5/8/2012 | GISAID |
| EPI396246 | A/Indiana/125/2012(H3N2) | 3/8/2012 | GISAID |
| EPI396254 | A/Indiana/126/2012(H3N2) | 4/8/2012 | GISAID |
| EPI396262 | A/Indiana/127/2012(H3N2) | 4/8/2012 | GISAID |
| EPI396270 | A/Indiana/133/2012(H3N2) | 3/8/2012 | GISAID |
| EPI396277 | A/Indiana/135/2012(H3N2) | 8/8/2012 | GISAID |

| | | | |
|---|---|---|---|
| EPI396280 | A/Indiana/144/2012(H3N2) | 8/8/2012 | GISAID |
| EPI398000 | A/Ohio/43/2012(H3N2) | 6/8/2012 | GISAID |
| EPI398040 | A/Ohio/57/2012(H3N2) | 5/8/2012 | GISAID |
| EPI396291 | A/Ohio/63/2012(H3N2) | 10/8/2012 | GISAID |
| EPI396299 | A/Maryland/23/2012(H3N2) | 18/8/2012 | GISAID |
| EPI397054 | A/PuertoRico/36/2012(H3N2) | 27/6/2012 | GISAID |
| EPI397106 | A/Indiana/115/2012(H3N2) | 4/8/2012 | GISAID |
| EPI397905 | A/Hawaii/03/2012(H3N2) | 12/7/2012 | GISAID |
| EPI397351 | A/Ohio/37/2012(H3N2) | 5/8/2012 | GISAID |
| EPI397913 | A/Indiana/123/2012(H3N2) | 4/8/2012 | GISAID |
| EPI397385 | A/Indiana/138/2012(H3N2) | 6/8/2012 | GISAID |
| EPI397396 | A/Indiana/93/2012(H3N2) | 6/8/2012 | GISAID |
| EPI397404 | A/Indiana/99/2012(H3N2) | 3/8/2012 | GISAID |
| EPI397412 | A/Michigan/16/2012(H3N2) | 15/8/2012 | GISAID |
| EPI397418 | A/Michigan/23/2012(H3N2) | 16/8/2012 | GISAID |
| EPI397426 | A/Minnesota/15/2012(H3N2) | 5/9/2012 | GISAID |
| EPI397432 | A/Ohio/81/2012(H3N2) | 17/8/2012 | GISAID |
| EPI397962 | A/Wisconsin/30/2012(H3N2) | 20/8/2012 | GISAID |
| EPI397446 | A/Wisconsin/31/2012(H3N2) | 22/8/2012 | GISAID |
| EPI397503 | A/Indiana/104/2012(H3N2) | 3/8/2012 | GISAID |
| EPI397654 | A/Indiana/90/2012(H3N2) | 2/8/2012 | GISAID |
| EPI397516 | A/Ohio/36/2012(H3N2) | 2/8/2012 | GISAID |
| EPI397524 | A/Ohio/44/2012(H3N2) | 8/8/2012 | GISAID |
| EPI397532 | A/Ohio/47/2012(H3N2) | 7/8/2012 | GISAID |
| EPI397540 | A/Ohio/56/2012(H3N2) | 10/8/2012 | GISAID |
| EPI397677 | A/Ohio/83/2012(H3N2) | 4/9/2012 | GISAID |
| EPI397548 | A/Ohio/84/2012(H3N2) | 13/8/2012 | GISAID |
| EPI397555 | A/Ohio/86/2012(H3N2) | 15/8/2012 | GISAID |
| EPI397562 | A/Wisconsin/29/2012(H3N2) | 20/8/2012 | GISAID |
| EPI397751 | A/Ohio/87/2012(H3N2) | 12/8/2012 | GISAID |
| EPI397759 | A/Wisconsin/35/2012(H3N2) | 22/8/2012 | GISAID |
| EPI397876 | A/Texas/50/2012(H3N2) | 15/4/2012 | GISAID |
| EPI397883 | A/Hawaii/22/2012(H3N2) | 9/7/2012 | GISAID |
| EPI397925 | A/Ohio/18/2012(H3N2) | 27/7/2012 | GISAID |
| EPI398037 | A/Ohio/19/2012(H3N2) | 29/7/2012 | GISAID |
| EPI397961 | A/Ohio/88/2012(H3N2) | 10/9/2012 | GISAID |
| EPI398417 | A/Texas/50/2012(H3N2) | 15/4/2012 | GISAID |

| | | | |
|---|---|---|---|
| EPI398424 | A/Hawaii/22/2012(H3N2) | 9/7/2012 | GISAID |
| EPI404890 | A/Iowa/16/2012(H3N2) | 17/11/2012 | GISAID |
| EPI404971 | A/Iowa/14/2012(H3N2) | 18/10/2012 | GISAID |
| EPI404989 | A/Pennsylvania/09/2012(H3N2) | 21/4/2012 | GISAID |
| EPI404996 | A/Pennsylvania/09/2012(H3N2) | 21/4/2012 | GISAID |
| EPI407279 | A/Minnesota/18/2012(H3N2) | 13/11/2012 | GISAID |
| EPI416138 | A/Michigan/32/2012(H3N2) | 18/11/2012 | GISAID |
| EPI416145 | A/Michigan/33/2012(H3N2) | 3/12/2012 | GISAID |
| EPI417231 | A/Arizona/09/2012(H3N2) | 22/10/2012 | GISAID |
| EPI418017 | A/Victoria/361/2011(H3N2) | 24/10/2011 | GISAID |
| EPI426140 | A/NewYork/39/2012(H3N2) | 20/10/2012 | GISAID |
| EPI436261 | A/NewYork/39/2012(H3N2) | 20/10/2012 | GISAID |
| EPI443697 | A/Wyoming/03/2003(H3N2) | 13/2/2003 | GISAID |
| EPI443709 | A/Washington/07/2013(H3N2) | 7/1/2013 | GISAID |
| EPI443733 | A/Alaska/02/2013(H3N2) | 20/2/2013 | GISAID |
| EPI445653 | A/Alaska/02/2013(H3N2) | 20/2/2013 | GISAID |
| EPI445712 | A/NewYork/39/2012(H3N2) | 20/10/2012 | GISAID |
| EPI450257 | A/NEWCASTLE/1/2013(H3N2) | 14/1/2013 | GISAID |
| EPI450293 | A/SOUTHAUSTRALIA/2/2013(H3N2) | 9/1/2013 | GISAID |
| EPI450524 | A/Nanjing/M1/2013(H3N2) | 26/4/2013 | GISAID |
| EPI466829 | A/Utah/07/2013(H3N2) | 5/7/2013 | GISAID |
| EPI466845 | A/Brazil/0328/2013(H3N2) | 22/5/2013 | GISAID |
| EPI466866 | A/Maryland/04/2013(H3N2) | 6/3/2013 | GISAID |
| EPI466889 | A/Ukraine/387/2013(H3N2) | 6/4/2013 | GISAID |
| EPI466905 | A/AmericanSamoa/4786/2013(H3N2) | 22/2/2013 | GISAID |
| EPI467440 | A/AmericanSamoa/4786/2013(H3N2) | 22/2/2013 | GISAID |
| EPI467448 | A/Victoria/505/2013(H3N2) | 17/6/2013 | GISAID |
| EPI477479 | A/SRILANKA/27/2013(H3N2) | 27/2/2013 | GISAID |
| EPI477495 | A/SYDNEY/506/2013(H3N2) | 4/3/2013 | GISAID |
| EPI484516 | A/Arkansas/13/2013(H3N2) | 15/8/2013 | GISAID |
| EPI484541 | A/Massachusetts/11/2013(H3N2) | 10/7/2013 | GISAID |
| EPI484551 | A/CostaRica/4009/2013(H3N2) | 23/7/2013 | GISAID |
| EPI484737 | A/CostaRica/4009/2013(H3N2) | 23/7/2013 | GISAID |
| EPI484565 | A/Georgia/03/2013(H3N2) | 14/7/2013 | GISAID |
| EPI484573 | A/Bangladesh/9334/2013(H3N2) | 7/5/2013 | GISAID |
| EPI484635 | A/Stockholm/42/2011(H3N2) | 26/12/2011 | GISAID |
| EPI484643 | A/Stockholm/19/2012(H3N2) | 14/4/2012 | GISAID |

| | | | |
|---|---|---|---|
| EPI484651 | A/Stockholm/29/2011(H3N2) | 31/10/2011 | GISAID |
| EPI484659 | A/Stockholm/12/2012(H3N2) | 12/1/2012 | GISAID |
| EPI484667 | A/Stockholm/43/2011(H3N2) | 28/12/2011 | GISAID |
| EPI484675 | A/Stockholm/14/2012(H3N2) | 11/3/2012 | GISAID |
| EPI484683 | A/Stockholm/26/2011(H3N2) | 6/10/2011 | GISAID |
| EPI485800 | A/CostaRica/4700/2013(H3N2) | 26/7/2013 | GISAID |
| EPI485808 | A/CostaRica/4700/2013(H3N2) | 26/7/2013 | GISAID |
| EPI489300 | A/Utah/07/2013(H3N2) | 5/7/2013 | GISAID |
| EPI489312 | A/Massachusetts/11/2013(H3N2) | 10/7/2013 | GISAID |
| EPI490093 | A/NewYork/39/2012(H3N2) | 20/10/2012 | GISAID |
| EPI492686 | A/Massachusetts/11/2013(H3N2) | 10/7/2013 | GISAID |
| EPI492690 | A/CostaRica/4700/2013(H3N2) | 26/7/2013 | GISAID |
| EPI492717 | A/Bangladesh/9269/2013(H3N2) | 12/5/2013 | GISAID |
| EPI497661 | A/Victoria/506/2013(H3N2) | 16/6/2013 | GISAID |
| EPI497668 | A/Utah/07/2013(H3N2) | 5/7/2013 | GISAID |
| EPI499306 | A/Pennsylvania/12/2013(H3N2) | 9/11/2013 | GISAID |
| EPI499308 | A/HongKong/5643/2013(H3N2) | 12/10/2013 | GISAID |
| EPI499365 | A/Wyoming/03/2003(H3N2) | 13/2/2003 | GISAID |
| EPI503938 | A/Washington/18/2013(H3N2) | 29/11/2013 | GISAID |
| EPI503951 | A/Washington/18/2013(H3N2) | 29/11/2013 | GISAID |
| EPI504746 | A/Stockholm/40/2013(H3N2) | 21/12/2013 | GISAID |
| EPI504749 | A/Stockholm/2/2014(H3N2) | 2/1/2014 | GISAID |
| EPI506693 | A/Hangzhou/B48/2013(H3N2) | 29/1/2013 | GISAID |
| EPI506701 | A/Hangzhou/A121/2013(H3N2) | 17/2/2013 | GISAID |
| EPI506709 | A/Hangzhou/A126/2013(H3N2) | 19/2/2013 | GISAID |
| EPI506717 | A/Hangzhou/A155/2013(H3N2) | 25/2/2013 | GISAID |
| EPI506725 | A/Hangzhou/A158/2013(H3N2) | 1/3/2013 | GISAID |
| EPI506733 | A/Hangzhou/A429/2013(H3N2) | 8/7/2013 | GISAID |
| EPI506741 | A/Hangzhou/A469/2013(H3N2) | 29/7/2013 | GISAID |
| EPI506749 | A/Hangzhou/A511/2013(H3N2) | 15/8/2013 | GISAID |
| EPI506757 | A/Hangzhou/A544/2013(H3N2) | 27/8/2013 | GISAID |
| EPI506765 | A/Hangzhou/A601/2013(H3N2) | 15/9/2013 | GISAID |
| EPI506773 | A/Hangzhou/A649/2013(H3N2) | 10/10/2013 | GISAID |
| EPI506781 | A/Hangzhou/A662/2013(H3N2) | 10/10/2013 | GISAID |
| EPI506789 | A/Hangzhou/A666/2013(H3N2) | 10/10/2013 | GISAID |
| EPI506797 | A/Hangzhou/A683/2013(H3N2) | 15/10/2013 | GISAID |
| EPI506805 | A/Hangzhou/B599/2013(H3N2) | 15/10/2013 | GISAID |

| | | | |
|---|---|---|---|
| EPI506813 | A/Hangzhou/B600/2013(H3N2) | 15/10/2013 | GISAID |
| EPI506821 | A/Hangzhou/A691/2013(H3N2) | 18/10/2013 | GISAID |
| EPI506829 | A/Hangzhou/A698/2013(H3N2) | 22/10/2013 | GISAID |
| EPI506837 | A/Hangzhou/A765/2013(H3N2) | 15/11/2013 | GISAID |
| EPI506845 | A/Hangzhou/A773/2013(H3N2) | 15/11/2013 | GISAID |
| EPI506853 | A/Hangzhou/B690/2013(H3N2) | 15/11/2013 | GISAID |
| EPI506861 | A/Hangzhou/A776/2013(H3N2) | 18/11/2013 | GISAID |
| EPI506869 | A/Hangzhou/A779/2013(H3N2) | 18/11/2013 | GISAID |
| EPI506877 | A/Hangzhou/A784/2013(H3N2) | 21/11/2013 | GISAID |
| EPI506885 | A/Hangzhou/A789/2013(H3N2) | 21/11/2013 | GISAID |
| EPI506893 | A/Hangzhou/B709/2013(H3N2) | 21/11/2013 | GISAID |
| EPI506901 | A/Hangzhou/A796/2013(H3N2) | 25/11/2013 | GISAID |
| EPI506909 | A/Hangzhou/A800/2013(H3N2) | 25/11/2013 | GISAID |
| EPI506917 | A/Hangzhou/B718/2013(H3N2) | 25/11/2013 | GISAID |
| EPI506925 | A/Hangzhou/B721/2013(H3N2) | 25/11/2013 | GISAID |
| EPI506933 | A/Hangzhou/A803/2013(H3N2) | 27/11/2013 | GISAID |
| EPI506941 | A/Hangzhou/A806/2013(H3N2) | 27/11/2013 | GISAID |
| EPI506949 | A/Hangzhou/B728/2013(H3N2) | 27/11/2013 | GISAID |
| EPI506957 | A/Hangzhou/B735/2013(H3N2) | 27/11/2013 | GISAID |
| EPI506965 | A/Hangzhou/A812/2013(H3N2) | 4/12/2013 | GISAID |
| EPI506973 | A/Hangzhou/A816/2013(H3N2) | 4/12/2013 | GISAID |
| EPI506981 | A/Hangzhou/A825/2013(H3N2) | 4/12/2013 | GISAID |
| EPI507594 | A/Almaty/2958/2013(H3N2) | 27/1/2013 | GISAID |
| EPI508248 | A/Stockholm/4/2014(H3N2) | 1/2/2014 | GISAID |
| EPI508269 | A/Gavle/1/2014(H3N2) | 6/2/2014 | GISAID |
| EPI508273 | A/Sweden/1/2014(H3N2) | 12/2/2014 | GISAID |
| EPI513291 | A/CostaRica/4700/2013(H3N2) | 26/7/2013 | GISAID |
| EPI515774 | A/Stockholm/5/2014(H3N2) | 6/2/2014 | GISAID |
| EPI515296 | A/Stockholm/6/2014(H3N2) | 6/2/2014 | GISAID |
| EPI516458 | A/Stockholm/13/2014(H3N2) | 21/2/2014 | GISAID |
| EPI516676 | A/Maryland/03/2014(H3N2) | 12/1/2014 | GISAID |
| EPI516844 | A/Iowa/04/2013(H3N2) | 1/10/2013 | GISAID |
| EPI526757 | A/PERTH/258/2013(H3N2) | 16/10/2013 | GISAID |
| EPI526801 | A/NORTHWESTAUCKLAND/6/2013(H3N2) | 18/9/2013 | GISAID |
| EPI529530 | A/BRISBANE/165/2013(H3N2) | 9/9/2013 | GISAID |
| EPI529546 | A/MALAYSIA/10/2013(H3N2) | 13/11/2013 | GISAID |
| EPI529557 | A/SYDNEY/83/2013(H3N2) | 21/11/2013 | GISAID |

| | | | |
|---|---|---|---|
| EPI529568 | A/CHIANGRAI/337/2013(H3N2) | 5/11/2013 | GISAID |
| EPI529587 | A/SONGKHLA/3076/2013(H3N2) | 18/11/2013 | GISAID |
| EPI531450 | A/Stockholm/15/2014(H3N2) | 7/4/2014 | GISAID |
| EPI537000 | A/Nebraska/04/2014(H3N2) | 11/3/2014 | GISAID |
| EPI537007 | A/Montana/27/2013(H3N2) | 27/12/2013 | GISAID |
| EPI537015 | A/Texas/50/2012(H3N2) | 15/4/2012 | GISAID |
| EPI537022 | A/California/02/2014(H3N2) | 16/1/2014 | GISAID |
| EPI537030 | A/Nebraska/04/2014(H3N2) | 11/3/2014 | GISAID |
| EPI537038 | A/Palau/6759/2014(H3N2) | 26/3/2014 | GISAID |
| EPI537046 | A/NewYork/39/2012(H3N2) | 20/10/2012 | GISAID |
| EPI537054 | A/Louisiana/39/2013(H3N2) | 29/12/2013 | GISAID |
| EPI537081 | A/NewYork/05/2014(H3N2) | 21/1/2014 | GISAID |
| EPI537985 | A/NewYork/05/2014(H3N2) | 21/1/2014 | GISAID |
| EPI540526 | A/Switzerland/9715293/2013(H3N2) | 6/12/2013 | GISAID |
| EPI541636 | A/Alaska/18/2014(H3N2) | 2/6/2014 | GISAID |
| EPI541651 | A/Washington/18/2013(H3N2) | 29/11/2013 | GISAID |
| EPI541659 | A/Switzerland/9715293/2013(H3N2) | 2013 | GISAID |
| EPI542216 | A/Stockholm/17/2014(H3N2) | 23/7/2014 | GISAID |
| EPI542224 | A/Stockholm/18/2014(H3N2) | 16/8/2014 | GISAID |
| EPI543234 | A/Stockholm/06/2014(H3N2) | 6/2/2014 | GISAID |
| EPI543523 | A/RioGrandedoSul/23/2014(H3N2) | 20/1/2014 | GISAID |
| EPI543531 | A/RioGrandedoSul/23/2014(H3N2) | 20/1/2014 | GISAID |
| EPI543551 | A/Parana/87/2014(H3N2) | 17/3/2014 | GISAID |
| EPI543666 | A/Maryland/25/2014(H3N2) | 27/8/2014 | GISAID |
| EPI544084 | A/SouthAustralia/55/2014(H3N2) | 29/6/2014 | GISAID |
| EPI544087 | A/SouthAustralia/55/2014(H3N2) | 29/6/2014 | GISAID |
| EPI544405 | A/Minnesota/21/2014(H3N2) | 17/9/2014 | GISAID |
| EPI544413 | A/Michigan/13/2014(H3N2) | 18/9/2014 | GISAID |
| EPI544421 | A/Texas/37/2014(H3N2) | 1/10/2014 | GISAID |
| EPI544429 | A/Texas/38/2014(H3N2) | 6/10/2014 | GISAID |
| EPI544437 | A/Michigan/12/2014(H3N2) | 16/9/2014 | GISAID |
| EPI544445 | A/NewMexico/27/2014(H3N2) | 18/9/2014 | GISAID |
| EPI544453 | A/Florida/35/2014(H3N2) | 23/9/2014 | GISAID |
| EPI544461 | A/Michigan/11/2014(H3N2) | 13/9/2014 | GISAID |
| EPI544476 | A/Maryland/27/2014(H3N2) | 17/9/2014 | GISAID |
| EPI544484 | A/Pennsylvania/40/2014(H3N2) | 13/9/2014 | GISAID |
| EPI544492 | A/Michigan/14/2014(H3N2) | 19/9/2014 | GISAID |

| | | | |
|---|---|---|---|
| EPI544500 | A/Michigan/15/2014(H3N2) | 24/9/2014 | GISAID |
| EPI544508 | A/Minnesota/22/2014(H3N2) | 20/9/2014 | GISAID |
| EPI544519 | A/Stockholm/06/2014(H3N2) | 6/2/2014 | GISAID |
| EPI544526 | A/NorthCarolina/13/2014(H3N2) | 15/4/2014 | GISAID |
| EPI544663 | A/Stockholm/21/2014(H3N2) | 15/10/2014 | GISAID |
| EPI545581 | A/Tak/1845/2014(H3N2) | 20/7/2014 | GISAID |
| EPI545589 | A/Kentucky/10/2014(H3N2) | 4/10/2014 | GISAID |
| EPI545597 | A/Wisconsin/26/2014(H3N2) | 25/9/2014 | GISAID |
| EPI545604 | A/Wisconsin/25/2014(H3N2) | 25/9/2014 | GISAID |
| EPI545612 | A/NorthCarolina/20/2014(H3N2) | 8/9/2014 | GISAID |
| EPI545622 | A/Wyoming/08/2014(H3N2) | 9/10/2014 | GISAID |
| EPI545630 | A/Wyoming/09/2014(H3N2) | 9/10/2014 | GISAID |
| EPI545643 | A/Mississippi/06/2014(H3N2) | 14/10/2014 | GISAID |
| EPI545648 | A/Mississippi/03/2014(H3N2) | 15/10/2014 | GISAID |
| EPI545656 | A/Mississippi/04/2014(H3N2) | 16/10/2014 | GISAID |
| EPI545664 | A/Mississippi/05/2014(H3N2) | 14/10/2014 | GISAID |
| EPI545672 | A/Colorado/15/2014(H3N2) | 4/10/2014 | GISAID |
| EPI545694 | A/Maine/04/2014(H3N2) | 9/10/2014 | GISAID |
| EPI545702 | A/Pennsylvania/45/2014(H3N2) | 25/9/2014 | GISAID |
| EPI545710 | A/Florida/34/2014(H3N2) | 30/9/2014 | GISAID |
| EPI545718 | A/Florida/39/2014(H3N2) | 7/10/2014 | GISAID |
| EPI545726 | A/Florida/40/2014(H3N2) | 7/10/2014 | GISAID |
| EPI545734 | A/Florida/36/2014(H3N2) | 3/10/2014 | GISAID |
| EPI545742 | A/Florida/38/2014(H3N2) | 7/10/2014 | GISAID |
| EPI545749 | A/Florida/37/2014(H3N2) | 7/10/2014 | GISAID |
| EPI545757 | A/Pennsylvania/44/2014(H3N2) | 27/9/2014 | GISAID |
| EPI545772 | A/NorthCarolina/21/2014(H3N2) | 6/10/2014 | GISAID |
| EPI547601 | A/Palau/6775/2014(H3N2) | 26/3/2014 | GISAID |
| EPI547711 | A/Argentina/11071/2014(H3N2) | 4/7/2014 | GISAID |
| EPI547719 | A/Argentina/11093/2014(H3N2) | 1/7/2014 | GISAID |
| EPI547727 | A/Argentina/11126/2014(H3N2) | 17/7/2014 | GISAID |
| EPI547735 | A/Bolivia/667/2014(H3N2) | 17/6/2014 | GISAID |
| EPI547743 | A/Bolivia/716/2014(H3N2) | 18/6/2014 | GISAID |
| EPI547751 | A/Bolivia/751/2014(H3N2) | 24/6/2014 | GISAID |
| EPI547766 | A/California/15/2014(H3N2) | 25/8/2014 | GISAID |
| EPI547774 | A/California/16/2014(H3N2) | 25/8/2014 | GISAID |
| EPI547782 | A/California/17/2014(H3N2) | 11/8/2014 | GISAID |

| | | | |
|---|---|---|---|
| EPI547790 | A/California/18/2014(H3N2) | 8/8/2014 | GISAID |
| EPI547798 | A/California/19/2014(H3N2) | 4/9/2014 | GISAID |
| EPI547806 | A/CostaRica/2303/2014(H3N2) | 9/7/2014 | GISAID |
| EPI547814 | A/DominicanRepublic/8483/2014(H3N2) | 28/5/2014 | GISAID |
| EPI547821 | A/DominicanRepublic/8490/2014(H3N2) | 22/5/2014 | GISAID |
| EPI547829 | A/Florida/35/2014(H3N2) | 23/9/2014 | GISAID |
| EPI547837 | A/Florida/37/2014(H3N2) | 7/10/2014 | GISAID |
| EPI547845 | A/Florida/38/2014(H3N2) | 7/10/2014 | GISAID |
| EPI547853 | A/Florida/39/2014(H3N2) | 7/10/2014 | GISAID |
| EPI547861 | A/Florida/40/2014(H3N2) | 7/10/2014 | GISAID |
| EPI547869 | A/Hawaii/35/2014(H3N2) | 10/8/2014 | GISAID |
| EPI547877 | A/Hawaii/36/2014(H3N2) | 8/8/2014 | GISAID |
| EPI547885 | A/Hawaii/37/2014(H3N2) | 13/8/2014 | GISAID |
| EPI547893 | A/Hawaii/38/2014(H3N2) | 11/8/2014 | GISAID |
| EPI547901 | A/Hawaii/39/2014(H3N2) | 13/8/2014 | GISAID |
| EPI547909 | A/Maine/04/2014(H3N2) | 9/10/2014 | GISAID |
| EPI547917 | A/Mexico/711/2014(H3N2) | 12/2/2014 | GISAID |
| EPI547925 | A/Michigan/08/2014(H3N2) | 14/8/2014 | GISAID |
| EPI547933 | A/Michigan/09/2014(H3N2) | 1/9/2014 | GISAID |
| EPI547941 | A/Michigan/10/2014(H3N2) | 10/9/2014 | GISAID |
| EPI547949 | A/Minnesota/21/2014(H3N2) | 17/9/2014 | GISAID |
| EPI547957 | A/Minnesota/22/2014(H3N2) | 20/9/2014 | GISAID |
| EPI547969 | A/NewHampshire/18/2014(H3N2) | 28/8/2014 | GISAID |
| EPI547977 | A/NewJersey/22/2014(H3N2) | 16/9/2014 | GISAID |
| EPI547985 | A/NewMexico/27/2014(H3N2) | 18/9/2014 | GISAID |
| EPI547993 | A/Newcastle/22/2014(H3N2) | 16/6/2014 | GISAID |
| EPI548001 | A/Nicaragua/7402/2014(H3N2) | 25/6/2014 | GISAID |
| EPI548009 | A/NorthCarolina/21/2014(H3N2) | 6/10/2014 | GISAID |
| EPI548017 | A/Paraguay/0049/2014(H3N2) | 3/7/2014 | GISAID |
| EPI548025 | A/Paraguay/0095/2014(H3N2) | 16/7/2014 | GISAID |
| EPI548033 | A/Pennsylvania/40/2014(H3N2) | 13/9/2014 | GISAID |
| EPI548046 | A/Texas/37/2014(H3N2) | 1/10/2014 | GISAID |
| EPI548054 | A/Texas/39/2014(H3N2) | 20/8/2014 | GISAID |
| EPI548062 | A/Virginia/16/2014(H3N2) | 14/10/2014 | GISAID |
| EPI548070 | A/Washington/18/2014(H3N2) | 6/10/2014 | GISAID |
| EPI548078 | A/Wisconsin/27/2014(H3N2) | 5/8/2014 | GISAID |
| EPI548086 | A/Finland/440/2014(H3N2) | 2014 | GISAID |

| | | | |
|---|---|---|---|
| EPI548094 | A/Stockholm/22/2014(H3N2) | 18/10/2014 | GISAID |
| EPI548102 | A/Stockholm/25/2014(H3N2) | 21/10/2014 | GISAID |
| EPI548110 | A/Stockholm/23/2014(H3N2) | 25/10/2014 | GISAID |
| EPI548118 | A/Malmoe/4/2014(H3N2) | 19/10/2014 | GISAID |
| EPI548133 | A/Stockholm/24/2014(H3N2) | 26/9/2014 | GISAID |
| EPI548860 | A/Arkansas/08/2014(H3N2) | 1/10/2014 | GISAID |
| EPI548868 | A/Montana/11/2014(H3N2) | 17/10/2014 | GISAID |
| EPI548877 | A/Colorado/21/2014(H3N2) | 21/10/2014 | GISAID |
| EPI553599 | A/Colorado/19/2014(H3N2) | 18/10/2014 | GISAID |
| EPI548892 | A/Brazil/3972/2014(H3N2) | 6/7/2014 | GISAID |
| EPI548900 | A/Bangladesh/10009/2014(H3N2) | 9/6/2014 | GISAID |
| EPI548908 | A/Hawaii/46/2014(H3N2) | 20/7/2014 | GISAID |
| EPI548916 | A/Hawaii/45/2014(H3N2) | 26/7/2014 | GISAID |
| EPI548924 | A/Mississippi/04/2014(H3N2) | 16/10/2014 | GISAID |
| EPI548932 | A/NorthDakota/06/2014(H3N2) | 1/10/2014 | GISAID |
| EPI548942 | A/Brazil/0769/2014(H3N2) | 19/2/2014 | GISAID |
| EPI548950 | A/Colorado/15/2014(H3N2) | 4/10/2014 | GISAID |
| EPI548958 | A/Brazil/9879/2014(H3N2) | 17/2/2014 | GISAID |
| EPI548966 | A/Wyoming/08/2014(H3N2) | 9/10/2014 | GISAID |
| EPI548974 | A/Mississippi/03/2014(H3N2) | 15/10/2014 | GISAID |
| EPI548982 | A/Mississippi/07/2014(H3N2) | 6/10/2014 | GISAID |
| EPI548990 | A/NewMexico/28/2014(H3N2) | 15/10/2014 | GISAID |
| EPI548998 | A/Brazil/2035/2014(H3N2) | 7/5/2014 | GISAID |
| EPI549006 | A/Brazil/623/2014(H3N2) | 7/7/2014 | GISAID |
| EPI549014 | A/Brazil/367/2014(H3N2) | 21/7/2014 | GISAID |
| EPI549022 | A/Pennsylvania/42/2014(H3N2) | 19/9/2014 | GISAID |
| EPI549030 | A/Brazil/6069/2014(H3N2) | 3/3/2014 | GISAID |
| EPI549038 | A/Mississippi/05/2014(H3N2) | 14/10/2014 | GISAID |
| EPI549046 | A/Hawaii/44/2014(H3N2) | 18/7/2014 | GISAID |
| EPI549053 | A/Hawaii/47/2014(H3N2) | 18/7/2014 | GISAID |
| EPI549061 | A/NewHampshire/19/2014(H3N2) | 28/9/2014 | GISAID |
| EPI549069 | A/Brazil/4146/2014(H3N2) | 7/6/2014 | GISAID |
| EPI549077 | A/Delaware/19/2014(H3N2) | 11/10/2014 | GISAID |
| EPI549085 | A/Connecticut/19/2014(H3N2) | 3/10/2014 | GISAID |
| EPI549116 | A/Arkansas/06/2014(H3N2) | 24/9/2014 | GISAID |
| EPI549124 | A/Arkansas/07/2014(H3N2) | 29/9/2014 | GISAID |
| EPI549132 | A/Arkansas/05/2014(H3N2) | 24/9/2014 | GISAID |

| | | | |
|---|---|---|---|
| EPI549140 | A/Pennsylvania/43/2014(H3N2) | 24/9/2014 | GISAID |
| EPI549148 | A/Minnesota/28/2014(H3N2) | 18/10/2014 | GISAID |
| EPI549156 | A/Texas/41/2014(H3N2) | 20/10/2014 | GISAID |
| EPI550842 | A/HongKong/5738/2014(H3N2) | 30/4/2014 | GISAID |
| EPI550857 | A/Alabama/05/2014(H3N2) | 20/10/2014 | GISAID |
| EPI550872 | A/Colorado/16/2014(H3N2) | 3/10/2014 | GISAID |
| EPI550895 | A/Connecticut/21/2014(H3N2) | 14/10/2014 | GISAID |
| EPI550903 | A/Florida/35/2014(H3N2) | 23/9/2014 | GISAID |
| EPI550911 | A/Florida/45/2014(H3N2) | 12/10/2014 | GISAID |
| EPI550919 | A/Indiana/10/2014(H3N2) | 25/10/2014 | GISAID |
| EPI550927 | A/Louisiana/06/2014(H3N2) | 7/10/2014 | GISAID |
| EPI550935 | A/Louisiana/07/2014(H3N2) | 10/10/2014 | GISAID |
| EPI550943 | A/Louisiana/08/2014(H3N2) | 11/10/2014 | GISAID |
| EPI550951 | A/Louisiana/09/2014(H3N2) | 11/10/2014 | GISAID |
| EPI550959 | A/Louisiana/11/2014(H3N2) | 21/10/2014 | GISAID |
| EPI550967 | A/Louisiana/14/2014(H3N2) | 22/10/2014 | GISAID |
| EPI550975 | A/Maine/05/2014(H3N2) | 14/10/2014 | GISAID |
| EPI550983 | A/Maryland/26/2014(H3N2) | 16/9/2014 | GISAID |
| EPI550996 | A/NewHampshire/21/2014(H3N2) | 17/10/2014 | GISAID |
| EPI551004 | A/NorthCarolina/21/2014(H3N2) | 6/10/2014 | GISAID |
| EPI551012 | A/Oregon/05/2014(H3N2) | 9/10/2014 | GISAID |
| EPI551020 | A/Oregon/06/2014(H3N2) | 9/10/2014 | GISAID |
| EPI551035 | A/Oregon/09/2014(H3N2) | 5/10/2014 | GISAID |
| EPI551043 | A/Texas/34/2014(H3N2) | 10/6/2014 | GISAID |
| EPI551051 | A/Texas/37/2014(H3N2) | 1/10/2014 | GISAID |
| EPI551059 | A/Texas/40/2014(H3N2) | 22/9/2014 | GISAID |
| EPI551067 | A/Texas/42/2014(H3N2) | 22/10/2014 | GISAID |
| EPI551075 | A/Texas/43/2014(H3N2) | 15/10/2014 | GISAID |
| EPI551220 | A/ENG/516/2014(H3N2) | 24/10/2014 | GISAID |
| EPI551562 | A/CANBERRA/82/2014(H3N2) | 31/7/2014 | GISAID |
| EPI551603 | A/NEWCASTLE/54/2014(H3N2) | 5/8/2014 | GISAID |
| EPI551849 | A/VICTORIA/5060/2014(H3N2) | 19/9/2014 | GISAID |
| EPI556534 | A/Indiana/17/2014(H3N2) | 11/11/2014 | GISAID |
| EPI552085 | A/Ohio/03/2014(H3N2) | 29/10/2014 | GISAID |
| EPI552093 | A/Indiana/12/2014(H3N2) | 5/11/2014 | GISAID |
| EPI552101 | A/Indiana/13/2014(H3N2) | 7/11/2014 | GISAID |
| EPI552109 | A/Louisiana/12/2014(H3N2) | 21/10/2014 | GISAID |

| | | | |
|---|---|---|---|
| EPI552117 | A/NorthDakota/09/2014(H3N2) | 24/10/2014 | GISAID |
| EPI552125 | A/Missouri/09/2014(H3N2) | 3/11/2014 | GISAID |
| EPI552133 | A/Mississippi/12/2014(H3N2) | 5/11/2014 | GISAID |
| EPI552141 | A/Alabama/07/2014(H3N2) | 23/10/2014 | GISAID |
| EPI552149 | A/Mississippi/08/2014(H3N2) | 23/10/2014 | GISAID |
| EPI552157 | A/Mississippi/10/2014(H3N2) | 3/11/2014 | GISAID |
| EPI552165 | A/Mississippi/09/2014(H3N2) | 29/10/2014 | GISAID |
| EPI552173 | A/Montana/14/2014(H3N2) | 28/10/2014 | GISAID |
| EPI552179 | A/Montana/15/2014(H3N2) | 30/10/2014 | GISAID |
| EPI552187 | A/Texas/48/2014(H3N2) | 27/10/2014 | GISAID |
| EPI552197 | A/Indiana/15/2014(H3N2) | 10/11/2014 | GISAID |
| EPI552205 | A/Texas/46/2014(H3N2) | 30/10/2014 | GISAID |
| EPI552213 | A/Iowa/16/2014(H3N2) | 21/10/2014 | GISAID |
| EPI552227 | A/Indiana/16/2014(H3N2) | 10/11/2014 | GISAID |
| EPI552235 | A/Iowa/15/2014(H3N2) | 20/10/2014 | GISAID |
| EPI552243 | A/Louisiana/13/2014(H3N2) | 21/10/2014 | GISAID |
| EPI552251 | A/Utah/21/2014(H3N2) | 6/11/2014 | GISAID |
| EPI552259 | A/Utah/19/2014(H3N2) | 7/11/2014 | GISAID |
| EPI552267 | A/Utah/20/2014(H3N2) | 5/11/2014 | GISAID |
| EPI552275 | A/Utah/17/2014(H3N2) | 6/11/2014 | GISAID |
| EPI552283 | A/Utah/18/2014(H3N2) | 2/11/2014 | GISAID |
| EPI552302 | A/Kansas/08/2014(H3N2) | 29/10/2014 | GISAID |
| EPI552309 | A/Kansas/07/2014(H3N2) | 27/10/2014 | GISAID |
| EPI552317 | A/Maryland/33/2014(H3N2) | 29/10/2014 | GISAID |
| EPI552325 | A/Ohio/04/2014(H3N2) | 29/10/2014 | GISAID |
| EPI552335 | A/Alabama/09/2014(H3N2) | 5/11/2014 | GISAID |
| EPI552343 | A/Nevada/18/2014(H3N2) | 10/11/2014 | GISAID |
| EPI552351 | A/Nevada/17/2014(H3N2) | 10/11/2014 | GISAID |
| EPI552358 | A/Utah/22/2014(H3N2) | 8/11/2014 | GISAID |
| EPI552366 | A/Mississippi/11/2014(H3N2) | 4/11/2014 | GISAID |
| EPI552374 | A/Idaho/04/2014(H3N2) | 5/11/2014 | GISAID |
| EPI552395 | A/Indiana/11/2014(H3N2) | 5/11/2014 | GISAID |
| EPI552417 | A/Karlstad/1/2014(H3N2) | 6/11/2014 | GISAID |
| EPI552425 | A/Stockholm/26/2014(H3N2) | 3/11/2014 | GISAID |
| EPI552535 | A/Michigan/15/2014(H3N2) | 24/9/2014 | GISAID |
| EPI552544 | A/Palau/6759/2014(H3N2) | 26/3/2014 | GISAID |
| EPI552560 | A/SHIMANE/96/2014(H3N2) | 9/9/2014 | GISAID |

| | | | |
|---|---|---|---|
| EPI552568 | A/SAITAMA/96/2014(H3N2) | 11/9/2014 | GISAID |
| EPI552674 | A/Gothenburg/4/2014(H3N2) | 6/11/2014 | GISAID |
| EPI552682 | A/Sweden/76/2014(H3N2) | 2/11/2014 | GISAID |
| EPI552690 | A/Stockholm/29/2014(H3N2) | 18/11/2014 | GISAID |
| EPI552698 | A/Stockholm/28/2014(H3N2) | 2/12/2014 | GISAID |
| EPI552706 | A/Malmoe/5/2014(H3N2) | 28/11/2014 | GISAID |
| EPI552722 | A/Sweden/75/2014(H3N2) | 4/12/2014 | GISAID |
| EPI552785 | A/Iowa/13/2014(H3N2) | 9/10/2014 | GISAID |
| EPI552799 | A/NewHampshire/23/2014(H3N2) | 19/11/2014 | GISAID |
| EPI552807 | A/NorthCarolina/31/2014(H3N2) | 13/11/2014 | GISAID |
| EPI552823 | A/Louisiana/20/2014(H3N2) | 19/11/2014 | GISAID |
| EPI552831 | A/Louisiana/22/2014(H3N2) | 19/11/2014 | GISAID |
| EPI552839 | A/Louisiana/17/2014(H3N2) | 18/11/2014 | GISAID |
| EPI552847 | A/Louisiana/21/2014(H3N2) | 19/11/2014 | GISAID |
| EPI552855 | A/Louisiana/18/2014(H3N2) | 18/11/2014 | GISAID |
| EPI552876 | A/Wyoming/14/2014(H3N2) | 14/11/2014 | GISAID |
| EPI552892 | A/NorthCarolina/35/2014(H3N2) | 14/11/2014 | GISAID |
| EPI552900 | A/NewJersey/27/2014(H3N2) | 19/11/2014 | GISAID |
| EPI552905 | A/Wyoming/11/2014(H3N2) | 7/11/2014 | GISAID |
| EPI552913 | A/NewJersey/26/2014(H3N2) | 17/11/2014 | GISAID |
| EPI552930 | A/Delaware/22/2014(H3N2) | 21/10/2014 | GISAID |
| EPI556759 | A/Nevada/20/2014(H3N2) | 27/10/2014 | GISAID |
| EPI552948 | A/NorthCarolina/28/2014(H3N2) | 4/11/2014 | GISAID |
| EPI552951 | A/Iowa/14/2014(H3N2) | 12/10/2014 | GISAID |
| EPI552959 | A/NorthCarolina/34/2014(H3N2) | 13/11/2014 | GISAID |
| EPI552967 | A/Louisiana/19/2014(H3N2) | 18/11/2014 | GISAID |
| EPI552975 | A/NorthCarolina/29/2014(H3N2) | 8/11/2014 | GISAID |
| EPI552983 | A/Wyoming/15/2014(H3N2) | 18/11/2014 | GISAID |
| EPI552991 | A/NorthCarolina/32/2014(H3N2) | 13/11/2014 | GISAID |
| EPI553000 | A/Alaska/47/2014(H3N2) | 5/10/2014 | GISAID |
| EPI553005 | A/Alaska/46/2014(H3N2) | 4/10/2014 | GISAID |
| EPI553016 | A/Texas/52/2014(H3N2) | 5/11/2014 | GISAID |
| EPI553026 | A/Maryland/34/2014(H3N2) | 23/10/2014 | GISAID |
| EPI553041 | A/Missouri/11/2014(H3N2) | 18/11/2014 | GISAID |
| EPI553049 | A/Massachusetts/14/2014(H3N2) | 19/11/2014 | GISAID |
| EPI553057 | A/Texas/57/2014(H3N2) | 25/11/2014 | GISAID |
| EPI553065 | A/Missouri/10/2014(H3N2) | 17/11/2014 | GISAID |

| | | | |
|---|---|---|---|
| EPI553073 | A/Delaware/24/2014(H3N2) | 11/11/2014 | GISAID |
| EPI553081 | A/Delaware/25/2014(H3N2) | 20/11/2014 | GISAID |
| EPI553092 | A/Georgia/05/2014(H3N2) | 20/11/2014 | GISAID |
| EPI553100 | A/Georgia/06/2014(H3N2) | 20/11/2014 | GISAID |
| EPI553108 | A/Georgia/04/2014(H3N2) | 20/11/2014 | GISAID |
| EPI553116 | A/Georgia/07/2014(H3N2) | 20/11/2014 | GISAID |
| EPI553119 | A/Illinois/05/2014(H3N2) | 3/11/2014 | GISAID |
| EPI553126 | A/Kansas/10/2014(H3N2) | 19/11/2014 | GISAID |
| EPI553259 | A/SENDAI/72/2014(H3N2) | 6/11/2014 | GISAID |
| EPI553374 | A/Singapore/OHC005/2013(H3N2) | 11/12/2013 | GISAID |
| EPI553382 | A/Singapore/OHC008/2013(H3N2) | 11/12/2013 | GISAID |
| EPI553390 | A/Singapore/OHC010/2013(H3N2) | 12/12/2013 | GISAID |
| EPI553398 | A/Singapore/OHC015/2013(H3N2) | 13/12/2013 | GISAID |
| EPI553406 | A/Singapore/OHC016/2013(H3N2) | 13/12/2013 | GISAID |
| EPI553414 | A/Singapore/OHC019/2013(H3N2) | 13/12/2013 | GISAID |
| EPI553422 | A/Singapore/OHC022/2013(H3N2) | 13/12/2013 | GISAID |
| EPI553430 | A/Singapore/OHC009/2013(H3N2) | 12/12/2013 | GISAID |
| EPI553438 | A/Singapore/TT938/2013(H3N2) | 12/12/2013 | GISAID |
| EPI553446 | A/Singapore/TT973/2013(H3N2) | 16/12/2013 | GISAID |
| EPI553454 | A/Singapore/CGH021/2013(H3N2) | 30/12/2013 | GISAID |
| EPI553462 | A/Singapore/TT979/2013(H3N2) | 19/12/2013 | GISAID |
| EPI553499 | A/Colorado/24/2014(H3N2) | 7/11/2014 | GISAID |
| EPI553507 | A/Wisconsin/46/2014(H3N2) | 28/10/2014 | GISAID |
| EPI553528 | A/Florida/43/2014(H3N2) | 6/10/2014 | GISAID |
| EPI553536 | A/Arkansas/01/2014(H3N2) | 17/9/2014 | GISAID |
| EPI553544 | A/Florida/45/2014(H3N2) | 12/10/2014 | GISAID |
| EPI553552 | A/Arkansas/02/2014(H3N2) | 17/9/2014 | GISAID |
| EPI553560 | A/Florida/41/2014(H3N2) | 2014 | GISAID |
| EPI553575 | A/Arkansas/04/2014(H3N2) | 22/9/2014 | GISAID |
| EPI553583 | A/NewYork/26/2014(H3N2) | 14/10/2014 | GISAID |
| EPI553609 | A/Hawaii/48/2014(H3N2) | 18/10/2014 | GISAID |
| EPI553617 | A/Hawaii/49/2014(H3N2) | 18/10/2014 | GISAID |
| EPI553625 | A/Arkansas/12/2014(H3N2) | 21/10/2014 | GISAID |
| EPI553633 | A/Iowa/17/2014(H3N2) | 25/10/2014 | GISAID |
| EPI553641 | A/Arkansas/11/2014(H3N2) | 20/10/2014 | GISAID |
| EPI553649 | A/Mississippi/08/2014(H3N2) | 23/10/2014 | GISAID |
| EPI553660 | A/Alaska/40/2014(H3N2) | 8/9/2014 | GISAID |

| | | | |
|---|---|---|---|
| EPI553672 | A/NorthCarolina/24/2014(H3N2) | 24/9/2014 | GISAID |
| EPI553680 | A/Texas/42/2014(H3N2) | 22/10/2014 | GISAID |
| EPI553695 | A/Montana/20/2014(H3N2) | 11/11/2014 | GISAID |
| EPI553704 | A/NewYork/24/2014(H3N2) | 10/10/2014 | GISAID |
| EPI553722 | A/Wisconsin/49/2014(H3N2) | 9/11/2014 | GISAID |
| EPI553730 | A/Wisconsin/34/2014(H3N2) | 30/9/2014 | GISAID |
| EPI553738 | A/Texas/62/2014(H3N2) | 3/11/2014 | GISAID |
| EPI553746 | A/Wisconsin/48/2014(H3N2) | 8/11/2014 | GISAID |
| EPI553754 | A/Indiana/19/2014(H3N2) | 12/11/2014 | GISAID |
| EPI553762 | A/Maine/06/2014(H3N2) | 18/10/2014 | GISAID |
| EPI553770 | A/Louisiana/12/2014(H3N2) | 21/10/2014 | GISAID |
| EPI553778 | A/Louisiana/14/2014(H3N2) | 22/10/2014 | GISAID |
| EPI553786 | A/NewHampshire/21/2014(H3N2) | 17/10/2014 | GISAID |
| EPI553794 | A/Ohio/05/2014(H3N2) | 30/9/2014 | GISAID |
| EPI553802 | A/Maryland/18/2014(H3N2) | 2/5/2014 | GISAID |
| EPI553810 | A/Connecticut/21/2014(H3N2) | 14/10/2014 | GISAID |
| EPI553818 | A/Illinois/04/2014(H3N2) | 27/10/2014 | GISAID |
| EPI553826 | A/Alaska/52/2014(H3N2) | 3/9/2014 | GISAID |
| EPI553834 | A/Louisiana/08/2014(H3N2) | 11/10/2014 | GISAID |
| EPI553842 | A/Wisconsin/44/2014(H3N2) | 23/10/2014 | GISAID |
| EPI553850 | A/Washington/19/2014(H3N2) | 22/10/2014 | GISAID |
| EPI553858 | A/NewYork/19/2014(H3N2) | 23/9/2014 | GISAID |
| EPI553866 | A/NewYork/20/2014(H3N2) | 25/9/2014 | GISAID |
| EPI553874 | A/Maryland/28/2014(H3N2) | 14/10/2014 | GISAID |
| EPI553882 | A/Hawaii/40/2014(H3N2) | 7/9/2014 | GISAID |
| EPI553890 | A/NewYork/18/2014(H3N2) | 18/9/2014 | GISAID |
| EPI553898 | A/NewYork/36/2014(H3N2) | 23/10/2014 | GISAID |
| EPI553906 | A/Arkansas/10/2014(H3N2) | 20/10/2014 | GISAID |
| EPI553919 | A/NorthCarolina/23/2014(H3N2) | 9/9/2014 | GISAID |
| EPI553927 | A/Maryland/32/2014(H3N2) | 15/10/2014 | GISAID |
| EPI553935 | A/NewYork/28/2014(H3N2) | 14/10/2014 | GISAID |
| EPI553943 | A/NewYork/31/2014(H3N2) | 17/10/2014 | GISAID |
| EPI553951 | A/NewYork/34/2014(H3N2) | 17/10/2014 | GISAID |
| EPI553959 | A/NewYork/35/2014(H3N2) | 21/10/2014 | GISAID |
| EPI553967 | A/Hawaii/36/2014(H3N2) | 8/8/2014 | GISAID |
| EPI553975 | A/Maryland/19/2014(H3N2) | 1/5/2014 | GISAID |
| EPI555044 | A/Astrakhan/RII23/2014(H3N2) | 1/12/2014 | GISAID |

| | | | |
|---|---|---|---|
| EPI555102 | A/Stockholm/30/2014(H3N2) | 4/12/2014 | GISAID |
| EPI555110 | A/Stockholm/31/2014(H3N2) | 26/11/2014 | GISAID |
| EPI555233 | A/RhodeIsland/21/2014(H3N2) | 30/11/2014 | GISAID |
| EPI555241 | A/Texas/55/2014(H3N2) | 18/11/2014 | GISAID |
| EPI555249 | A/NorthDakota/10/2014(H3N2) | 4/12/2014 | GISAID |
| EPI555263 | A/Massachusetts/18/2014(H3N2) | 30/11/2014 | GISAID |
| EPI555271 | A/Louisiana/28/2014(H3N2) | 24/11/2014 | GISAID |
| EPI555279 | A/Massachusetts/17/2014(H3N2) | 2/12/2014 | GISAID |
| EPI555287 | A/Wisconsin/57/2014(H3N2) | 9/12/2014 | GISAID |
| EPI555295 | A/Wisconsin/59/2014(H3N2) | 9/12/2014 | GISAID |
| EPI555303 | A/Wisconsin/58/2014(H3N2) | 9/12/2014 | GISAID |
| EPI555311 | A/Pennsylvania/47/2014(H3N2) | 28/11/2014 | GISAID |
| EPI555319 | A/Texas/54/2014(H3N2) | 17/11/2014 | GISAID |
| EPI555327 | A/Missouri/13/2014(H3N2) | 1/12/2014 | GISAID |
| EPI555335 | A/Wisconsin/60/2014(H3N2) | 10/12/2014 | GISAID |
| EPI555343 | A/Wisconsin/56/2014(H3N2) | 9/12/2014 | GISAID |
| EPI555351 | A/Wisconsin/55/2014(H3N2) | 8/12/2014 | GISAID |
| EPI555359 | A/Washington/37/2014(H3N2) | 8/12/2014 | GISAID |
| EPI555367 | A/Minnesota/35/2014(H3N2) | 29/11/2014 | GISAID |
| EPI555375 | A/Wisconsin/54/2014(H3N2) | 8/12/2014 | GISAID |
| EPI555383 | A/Colorado/28/2014(H3N2) | 20/11/2014 | GISAID |
| EPI555402 | A/Wisconsin/53/2014(H3N2) | 2/12/2014 | GISAID |
| EPI555415 | A/NewHampshire/28/2014(H3N2) | 28/11/2014 | GISAID |
| EPI555427 | A/Utah/17/2014(H3N2) | 6/11/2014 | GISAID |
| EPI555435 | A/Alabama/08/2014(H3N2) | 5/11/2014 | GISAID |
| EPI555443 | A/Michigan/19/2014(H3N2) | 14/11/2014 | GISAID |
| EPI555451 | A/Kansas/12/2014(H3N2) | 8/12/2014 | GISAID |
| EPI555459 | A/Texas/69/2014(H3N2) | 24/11/2014 | GISAID |
| EPI555467 | A/Illinois/07/2014(H3N2) | 10/11/2014 | GISAID |
| EPI555479 | A/Florida/66/2014(H3N2) | 8/12/2014 | GISAID |
| EPI555487 | A/Hawaii/60/2014(H3N2) | 3/11/2014 | GISAID |
| EPI555499 | A/Texas/56/2014(H3N2) | 25/11/2014 | GISAID |
| EPI555507 | A/NorthDakota/11/2014(H3N2) | 2/12/2014 | GISAID |
| EPI555515 | A/Oklahoma/10/2014(H3N2) | 24/11/2014 | GISAID |
| EPI555524 | A/Delaware/31/2014(H3N2) | 7/12/2014 | GISAID |
| EPI555538 | A/NewMexico/30/2014(H3N2) | 28/11/2014 | GISAID |
| EPI555546 | A/Idaho/07/2014(H3N2) | 25/11/2014 | GISAID |

| | | | |
|---|---|---|---|
| EPI555559 | A/Kentucky/14/2014(H3N2) | 26/11/2014 | GISAID |
| EPI555567 | A/Minnesota/36/2014(H3N2) | 23/11/2014 | GISAID |
| EPI555575 | A/Texas/66/2014(H3N2) | 30/10/2014 | GISAID |
| EPI556107 | A/Pennsylvania/48/2014(H3N2) | 28/11/2014 | GISAID |
| EPI556122 | A/Mississippi/16/2014(H3N2) | 8/12/2014 | GISAID |
| EPI556147 | A/Wyoming/17/2014(H3N2) | 24/11/2014 | GISAID |
| EPI556155 | A/Alabama/09/2014(H3N2) | 5/11/2014 | GISAID |
| EPI556163 | A/Texas/70/2014(H3N2) | 24/11/2014 | GISAID |
| EPI556171 | A/Mississippi/15/2014(H3N2) | 25/11/2014 | GISAID |
| EPI556190 | A/Wisconsin/52/2014(H3N2) | 2/12/2014 | GISAID |
| EPI556203 | A/NorthCarolina/31/2014(H3N2) | 13/11/2014 | GISAID |
| EPI556211 | A/Ohio/08/2014(H3N2) | 22/11/2014 | GISAID |
| EPI556219 | A/Michigan/18/2014(H3N2) | 10/11/2014 | GISAID |
| EPI556227 | A/Kentucky/15/2014(H3N2) | 2/12/2014 | GISAID |
| EPI556239 | A/Washington/35/2014(H3N2) | 1/12/2014 | GISAID |
| EPI556247 | A/NewMexico/29/2014(H3N2) | 18/11/2014 | GISAID |
| EPI556255 | A/NewJersey/29/2014(H3N2) | 20/11/2014 | GISAID |
| EPI556270 | A/RhodeIsland/22/2014(H3N2) | 8/12/2014 | GISAID |
| EPI556278 | A/Texas/68/2014(H3N2) | 24/11/2014 | GISAID |
| EPI556286 | A/Pennsylvania/49/2014(H3N2) | 28/11/2014 | GISAID |
| EPI556294 | A/Ohio/07/2014(H3N2) | 19/11/2014 | GISAID |
| EPI556302 | A/Oklahoma/09/2014(H3N2) | 24/11/2014 | GISAID |
| EPI556317 | A/Florida/65/2014(H3N2) | 5/12/2014 | GISAID |
| EPI556325 | A/Nevada/22/2014(H3N2) | 4/12/2014 | GISAID |
| EPI556333 | A/Nevada/21/2014(H3N2) | 2/12/2014 | GISAID |
| EPI556341 | A/Massachusetts/16/2014(H3N2) | 3/12/2014 | GISAID |
| EPI556349 | A/Michigan/17/2014(H3N2) | 10/11/2014 | GISAID |
| EPI556357 | A/Indiana/21/2014(H3N2) | 27/11/2014 | GISAID |
| EPI556365 | A/Maryland/40/2014(H3N2) | 25/11/2014 | GISAID |
| EPI556467 | A/FUKUSHIMA/103/2014(H3N2) | 25/11/2014 | GISAID |
| EPI556512 | A/Washington/33/2014(H3N2) | 21/11/2014 | GISAID |
| EPI556523 | A/NorthCarolina/33/2014(H3N2) | 13/11/2014 | GISAID |
| EPI556531 | A/Massachusetts/19/2014(H3N2) | 2/12/2014 | GISAID |
| EPI556548 | A/Nebraska/15/2014(H3N2) | 5/12/2014 | GISAID |
| EPI556556 | A/Vermont/09/2014(H3N2) | 14/12/2014 | GISAID |
| EPI556564 | A/California/40/2014(H3N2) | 10/12/2014 | GISAID |
| EPI556574 | A/Massachusetts/20/2014(H3N2) | 18/11/2014 | GISAID |

| | | | |
|---|---|---|---|
| EPI556582 | A/Vermont/10/2014(H3N2) | 13/12/2014 | GISAID |
| EPI556590 | A/Nebraska/14/2014(H3N2) | 11/12/2014 | GISAID |
| EPI556613 | A/Idaho/09/2014(H3N2) | 18/12/2014 | GISAID |
| EPI556622 | A/Oregon/11/2014(H3N2) | 21/11/2014 | GISAID |
| EPI556628 | A/Michigan/26/2014(H3N2) | 25/11/2014 | GISAID |
| EPI556637 | A/Oregon/13/2014(H3N2) | 18/11/2014 | GISAID |
| EPI556645 | A/Minnesota/34/2014(H3N2) | 12/11/2014 | GISAID |
| EPI556653 | A/Illinois/09/2014(H3N2) | 1/12/2014 | GISAID |
| EPI556660 | A/Minnesota/37/2014(H3N2) | 16/11/2014 | GISAID |
| EPI556668 | A/Hawaii/59/2014(H3N2) | 13/11/2014 | GISAID |
| EPI556676 | A/Virginia/30/2014(H3N2) | 11/12/2014 | GISAID |
| EPI556685 | A/Illinois/05/2014(H3N2) | 3/11/2014 | GISAID |
| EPI556695 | A/Wyoming/15/2014(H3N2) | 18/11/2014 | GISAID |
| EPI556704 | A/Louisiana/17/2014(H3N2) | 18/11/2014 | GISAID |
| EPI556712 | A/Washington/36/2014(H3N2) | 21/11/2014 | GISAID |
| EPI556725 | A/Georgia/13/2014(H3N2) | 14/11/2014 | GISAID |
| EPI556733 | A/Florida/73/2014(H3N2) | 11/12/2014 | GISAID |
| EPI556741 | A/Florida/76/2014(H3N2) | 1/12/2014 | GISAID |
| EPI556749 | A/Idaho/08/2014(H3N2) | 16/12/2014 | GISAID |
| EPI556757 | A/Virginia/31/2014(H3N2) | 11/12/2014 | GISAID |
| EPI556783 | A/Maryland/26/2014(H3N2) | 16/9/2014 | GISAID |
| EPI556791 | A/Louisiana/20/2014(H3N2) | 19/11/2014 | GISAID |
| EPI556816 | A/Texas/50/2012(H3N2) | 15/4/2012 | GISAID |
| EPI556824 | A/NewHampshire/23/2014(H3N2) | 19/11/2014 | GISAID |
| EPI557603 | A/TOYAMA/124/2014(H3N2) | 10/11/2014 | GISAID |
| EPI557631 | A/TOKYO/41724/2014(H3N2) | 4/12/2014 | GISAID |
| EPI558127 | A/Texas/89/2014(H3N2) | 7/12/2014 | GISAID |
| EPI558135 | A/Texas/72/2014(H3N2) | 8/12/2014 | GISAID |
| EPI558146 | A/Arkansas/14/2014(H3N2) | 8/12/2014 | GISAID |
| EPI558154 | A/Nevada/27/2014(H3N2) | 22/12/2014 | GISAID |
| EPI558165 | A/Texas/79/2014(H3N2) | 15/12/2014 | GISAID |
| EPI558168 | A/California/27/2014(H3N2) | 10/10/2014 | GISAID |
| EPI558176 | A/Nevada/24/2014(H3N2) | 4/12/2014 | GISAID |
| EPI558184 | A/Texas/73/2014(H3N2) | 8/12/2014 | GISAID |
| EPI558199 | A/Vermont/11/2014(H3N2) | 18/12/2014 | GISAID |
| EPI558205 | A/Florida/58/2014(H3N2) | 8/11/2014 | GISAID |
| EPI558213 | A/Texas/74/2014(H3N2) | 8/12/2014 | GISAID |

| | | | |
|---|---|---|---|
| EPI558221 | A/California/42/2014(H3N2) | 4/12/2014 | GISAID |
| EPI558241 | A/Ohio/10/2014(H3N2) | 9/12/2014 | GISAID |
| EPI558249 | A/Vermont/12/2014(H3N2) | 11/12/2014 | GISAID |
| EPI558257 | A/NewHampshire/30/2014(H3N2) | 1/12/2014 | GISAID |
| EPI558265 | A/Maine/01/2015(H3N2) | 1/1/2015 | GISAID |
| EPI558271 | A/Texas/80/2014(H3N2) | 2/12/2014 | GISAID |
| EPI558279 | A/Maine/08/2014(H3N2) | 30/12/2014 | GISAID |
| EPI558287 | A/Virginia/35/2014(H3N2) | 22/12/2014 | GISAID |
| EPI558295 | A/Utah/27/2014(H3N2) | 5/12/2014 | GISAID |
| EPI558303 | A/Nevada/26/2014(H3N2) | 12/12/2014 | GISAID |
| EPI558311 | A/Delaware/36/2014(H3N2) | 16/12/2014 | GISAID |
| EPI558326 | A/Delaware/35/2014(H3N2) | 7/12/2014 | GISAID |
| EPI558336 | A/Tennessee/18/2014(H3N2) | 9/12/2014 | GISAID |
| EPI558342 | A/Utah/34/2014(H3N2) | 5/12/2014 | GISAID |
| EPI558350 | A/NorthDakota/14/2014(H3N2) | 3/12/2014 | GISAID |
| EPI558355 | A/Colorado/25/2014(H3N2) | 9/11/2014 | GISAID |
| EPI558363 | A/Texas/82/2014(H3N2) | 11/12/2014 | GISAID |
| EPI558371 | A/Texas/81/2014(H3N2) | 10/12/2014 | GISAID |
| EPI558379 | A/Nevada/25/2014(H3N2) | 15/12/2014 | GISAID |
| EPI558385 | A/Alabama/12/2014(H3N2) | 9/12/2014 | GISAID |
| EPI558393 | A/Virginia/33/2014(H3N2) | 8/12/2014 | GISAID |
| EPI558401 | A/Missouri/16/2014(H3N2) | 10/12/2014 | GISAID |
| EPI558406 | A/Mississippi/19/2014(H3N2) | 8/12/2014 | GISAID |
| EPI558410 | A/Virginia/34/2014(H3N2) | 11/12/2014 | GISAID |
| EPI558418 | A/Tennessee/16/2014(H3N2) | 8/12/2014 | GISAID |
| EPI558426 | A/Utah/33/2014(H3N2) | 2/12/2014 | GISAID |
| EPI558434 | A/Tennessee/17/2014(H3N2) | 8/12/2014 | GISAID |
| EPI558439 | A/Utah/35/2014(H3N2) | 7/12/2014 | GISAID |
| EPI558445 | A/Alabama/10/2014(H3N2) | 4/12/2014 | GISAID |
| EPI558453 | A/Tennessee/15/2014(H3N2) | 9/12/2014 | GISAID |
| EPI558461 | A/Montana/27/2014(H3N2) | 9/12/2014 | GISAID |
| EPI558467 | A/Alabama/11/2014(H3N2) | 8/12/2014 | GISAID |
| EPI558473 | A/Ohio/09/2014(H3N2) | 4/12/2014 | GISAID |
| EPI558481 | A/Wyoming/19/2014(H3N2) | 10/12/2014 | GISAID |
| EPI558487 | A/Mississippi/20/2014(H3N2) | 9/12/2014 | GISAID |
| EPI558495 | A/Texas/75/2014(H3N2) | 2/12/2014 | GISAID |
| EPI558508 | A/Indiana/25/2014(H3N2) | 5/12/2014 | GISAID |

| | | | |
|---|---|---|---|
| EPI558516 | A/Virginia/32/2014(H3N2) | 10/12/2014 | GISAID |
| EPI558524 | A/Texas/71/2014(H3N2) | 8/12/2014 | GISAID |
| EPI558532 | A/Maine/07/2014(H3N2) | 29/12/2014 | GISAID |
| EPI558544 | A/Montana/26/2014(H3N2) | 8/12/2014 | GISAID |
| EPI558552 | A/Washington/39/2014(H3N2) | 23/12/2014 | GISAID |
| EPI558560 | A/Delaware/37/2014(H3N2) | 16/12/2014 | GISAID |
| EPI558568 | A/Pennsylvania/52/2014(H3N2) | 2/12/2014 | GISAID |
| EPI558574 | A/Louisiana/38/2014(H3N2) | 6/12/2014 | GISAID |
| EPI558580 | A/California/44/2014(H3N2) | 4/12/2014 | GISAID |
| EPI558592 | A/Virginia/29/2014(H3N2) | 9/12/2014 | GISAID |
| EPI558597 | A/Arkansas/15/2014(H3N2) | 9/12/2014 | GISAID |
| EPI558965 | A/Sweden/77/2014(H3N2) | 18/12/2014 | GISAID |
| EPI558987 | A/Stockholm/34/2014(H3N2) | 30/12/2014 | GISAID |
| EPI558995 | A/Stockholm/35/2014(H3N2) | 27/12/2014 | GISAID |
| EPI559003 | A/Skovde/1/2014(H3N2) | 31/12/2014 | GISAID |
| EPI559011 | A/Skovde/1/2015(H3N2) | 1/1/2015 | GISAID |
| EPI559438 | A/Nagasaki/13N053/2014(H3N2) | 20/2/2014 | GISAID |
| EPI560420 | A/Shaanxi-Changan/1452/2014(H3N2) | 3/11/2014 | GISAID |
| EPI560439 | A/Beijing-Xicheng/13100/2014(H3N2) | 4/8/2014 | GISAID |
| EPI560458 | A/Nevada/29/2014(H3N2) | 23/12/2014 | GISAID |
| EPI560463 | A/Ethiopia/1141/2014(H3N2) | 21/10/2014 | GISAID |
| EPI560471 | A/Iquique/62331/2014(H3N2) | 16/9/2014 | GISAID |
| EPI560479 | A/Georgia/20/2014(H3N2) | 3/12/2014 | GISAID |
| EPI560498 | A/Laos/768/2014(H3N2) | 6/8/2014 | GISAID |
| EPI560506 | A/Laos/842/2014(H3N2) | 25/8/2014 | GISAID |
| EPI560514 | A/Kenya/126/2014(H3N2) | 27/8/2014 | GISAID |
| EPI560522 | A/Laos/864/2014(H3N2) | 26/8/2014 | GISAID |
| EPI560530 | A/Georgia/19/2014(H3N2) | 14/11/2014 | GISAID |
| EPI560538 | A/Georgia/16/2014(H3N2) | 27/11/2014 | GISAID |
| EPI560553 | A/Laos/859/2014(H3N2) | 26/8/2014 | GISAID |
| EPI560561 | A/Laos/899/2014(H3N2) | 8/9/2014 | GISAID |
| EPI560569 | A/Georgia/18/2014(H3N2) | 29/11/2014 | GISAID |
| EPI560577 | A/Georgia/23/2014(H3N2) | 26/11/2014 | GISAID |
| EPI560582 | A/Mexico/2971/2014(H3N2) | 4/12/2014 | GISAID |
| EPI560590 | A/Santiago/70123/2014(H3N2) | 26/10/2014 | GISAID |
| EPI560602 | A/Ontario/RV3241/2014(H3N2) | 25/10/2014 | GISAID |
| EPI560610 | A/Busan/1996/2014(H3N2) | 15/12/2014 | GISAID |

| | | | |
|---|---|---|---|
| EPI560615 | A/Mexico/2974/2014(H3N2) | 8/12/2014 | GISAID |
| EPI560620 | A/Quebec/RV3213/2014(H3N2) | 6/10/2014 | GISAID |
| EPI560628 | A/Kenya/124/2014(H3N2) | 27/8/2014 | GISAID |
| EPI560636 | A/Kenya/125/2014(H3N2) | 21/8/2014 | GISAID |
| EPI560644 | A/Kenya/128/2014(H3N2) | 26/8/2014 | GISAID |
| EPI565851 | A/Ethiopia/546/2014(H3N2) | 23/10/2014 | GISAID |
| EPI560653 | A/Georgia/15/2014(H3N2) | 13/11/2014 | GISAID |
| EPI565563 | A/Ethiopia/1153/2014(H3N2) | 11/11/2014 | GISAID |
| EPI560664 | A/Kenya/123/2014(H3N2) | 19/8/2014 | GISAID |
| EPI560675 | A/Hawaii/63/2014(H3N2) | 15/12/2014 | GISAID |
| EPI560683 | A/Missouri/18/2014(H3N2) | 10/12/2014 | GISAID |
| EPI560691 | A/Virginia/36/2014(H3N2) | 25/11/2014 | GISAID |
| EPI560699 | A/Santiago/68839/2014(H3N2) | 20/10/2014 | GISAID |
| EPI560707 | A/Florida/80/2014(H3N2) | 18/12/2014 | GISAID |
| EPI560715 | A/Paraguay/0094/2014(H3N2) | 1/8/2014 | GISAID |
| EPI560723 | A/Pennsylvania/54/2014(H3N2) | 13/12/2014 | GISAID |
| EPI560730 | A/Manitoba/RV3623/2014(H3N2) | 11/12/2014 | GISAID |
| EPI560735 | A/Alberta/RV3246/2014(H3N2) | 12/10/2014 | GISAID |
| EPI560747 | A/Alberta/RV3215/2014(H3N2) | 4/10/2014 | GISAID |
| EPI560755 | A/Kenya/122/2014(H3N2) | 25/8/2014 | GISAID |
| EPI560760 | A/Alberta/RV3240/2014(H3N2) | 16/10/2014 | GISAID |
| EPI560768 | A/Laos/820/2014(H3N2) | 18/8/2014 | GISAID |
| EPI623487 | A/Alberta/RV3588/2014(H3N2) | 18/11/2014 | GISAID |
| EPI565661 | A/Ontario/RV3606/2014(H3N2) | 7/12/2014 | GISAID |
| EPI560783 | A/Kenya/127/2014(H3N2) | 27/8/2014 | GISAID |
| EPI560791 | A/Xinjiang-Tianshan/11190/2014(H3N2) | 4/11/2014 | GISAID |
| EPI562067 | A/Pennsylvania/56/2014(H3N2) | 1/12/2014 | GISAID |
| EPI562097 | A/YOKOHAMA/184/2014(H3N2) | 25/12/2014 | GISAID |
| EPI562104 | A/NAGANO/3652/2014(H3N2) | 1/12/2014 | GISAID |
| EPI562136 | A/NIIGATA-C/60/2014(H3N2) | 9/11/2014 | GISAID |
| EPI562142 | A/OSAKA-C/2020/2014(H3N2) | 31/10/2014 | GISAID |
| EPI562376 | A/Nagasaki/13N063/2014(H3N2) | 3/3/2014 | GISAID |
| EPI562622 | A/Myanmar/13M002/2013(H3N2) | 27/6/2013 | GISAID |
| EPI562630 | A/Myanmar/13M004/2013(H3N2) | 2/7/2013 | GISAID |
| EPI584717 | A/Louisiana/20/2014(H3N2) | 19/11/2014 | GISAID |
| EPI569276 | A/NewCaledonia/71/2014(H3N2) | 13/8/2014 | GISAID |
| EPI564072 | A/California/43/2014(H3N2) | 23/12/2014 | GISAID |

| | | | |
|---|---|---|---|
| EPI564080 | A/Maine/04/2015(H3N2) | 2/1/2015 | GISAID |
| EPI564088 | A/Wisconsin/69/2014(H3N2) | 17/12/2014 | GISAID |
| EPI564093 | A/California/51/2014(H3N2) | 19/10/2014 | GISAID |
| EPI564101 | A/Missouri/20/2014(H3N2) | 26/12/2014 | GISAID |
| EPI564106 | A/Missouri/21/2014(H3N2) | 17/12/2014 | GISAID |
| EPI564110 | A/Massachusetts/25/2014(H3N2) | 11/12/2014 | GISAID |
| EPI564118 | A/Maryland/45/2014(H3N2) | 12/12/2014 | GISAID |
| EPI564126 | A/RhodeIsland/01/2015(H3N2) | 1/1/2015 | GISAID |
| EPI564131 | A/Massachusetts/23/2014(H3N2) | 14/12/2014 | GISAID |
| EPI564139 | A/Minnesota/38/2014(H3N2) | 7/12/2014 | GISAID |
| EPI564147 | A/Massachusetts/22/2014(H3N2) | 17/12/2014 | GISAID |
| EPI564155 | A/Wisconsin/68/2014(H3N2) | 16/12/2014 | GISAID |
| EPI564163 | A/Missouri/19/2014(H3N2) | 23/12/2014 | GISAID |
| EPI564171 | A/RhodeIsland/23/2014(H3N2) | 19/12/2014 | GISAID |
| EPI564179 | A/Maryland/44/2014(H3N2) | 10/12/2014 | GISAID |
| EPI564187 | A/Washington/48/2014(H3N2) | 19/12/2014 | GISAID |
| EPI564195 | A/Texas/02/2015(H3N2) | 6/1/2015 | GISAID |
| EPI564203 | A/Pennsylvania/01/2015(H3N2) | 2/1/2015 | GISAID |
| EPI564211 | A/Pennsylvania/02/2015(H3N2) | 9/1/2015 | GISAID |
| EPI564225 | A/Minnesota/31/2014(H3N2) | 28/10/2014 | GISAID |
| EPI564241 | A/Hawaii/63/2014(H3N2) | 15/12/2014 | GISAID |
| EPI564254 | A/Colorado/38/2014(H3N2) | 3/12/2014 | GISAID |
| EPI566172 | A/Wyoming/18/2014(H3N2) | 9/12/2014 | GISAID |
| EPI564268 | A/DistrictOfColumbia/08/2014(H3N2) | 27/12/2014 | GISAID |
| EPI564282 | A/DistrictOfColumbia/10/2014(H3N2) | 28/12/2014 | GISAID |
| EPI565397 | A/Nonthaburi/01/2015(H3N2) | 26/12/2014 | GISAID |
| EPI565401 | A/Guatemala/4645/2014(H3N2) | 12/11/2014 | GISAID |
| EPI564298 | A/Paraguay/3144/2014(H3N2) | 20/11/2014 | GISAID |
| EPI565405 | A/Guatemala/4647/2014(H3N2) | 14/11/2014 | GISAID |
| EPI564307 | A/Ethiopia/155/2014(H3N2) | 27/11/2014 | GISAID |
| EPI564312 | A/Ethiopia/1146/2014(H3N2) | 23/10/2014 | GISAID |
| EPI564320 | A/Indiana/24/2014(H3N2) | 1/12/2014 | GISAID |
| EPI564325 | A/Guatemala/4638/2014(H3N2) | 12/11/2014 | GISAID |
| EPI564330 | A/Guatemala/5211/2014(H3N2) | 13/11/2014 | GISAID |
| EPI564336 | A/Tennessee/23/2014(H3N2) | 4/12/2014 | GISAID |
| EPI612335 | A/Kentucky/30/2014(H3N2) | 1/12/2014 | GISAID |
| EPI564352 | A/Iowa/20/2014(H3N2) | 14/11/2014 | GISAID |

| | | | |
|---|---|---|---|
| EPI564362 | A/Illinois/12/2014(H3N2) | 3/12/2014 | GISAID |
| EPI564365 | A/Minnesota/39/2014(H3N2) | 4/12/2014 | GISAID |
| EPI564374 | A/Delaware/38/2014(H3N2) | 2/12/2014 | GISAID |
| EPI564382 | A/Missouri/17/2014(H3N2) | 4/12/2014 | GISAID |
| EPI564395 | A/Minnesota/41/2014(H3N2) | 23/11/2014 | GISAID |
| EPI564414 | A/Tennessee/20/2014(H3N2) | 15/12/2014 | GISAID |
| EPI564422 | A/NorthCarolina/41/2014(H3N2) | 3/12/2014 | GISAID |
| EPI564430 | A/Alberta/RV3246/2014(H3N2) | 12/10/2014 | GISAID |
| EPI564441 | A/Wyoming/20/2014(H3N2) | 9/12/2014 | GISAID |
| EPI564449 | A/HongKong/7617/2014(H3N2) | 29/11/2014 | GISAID |
| EPI564457 | A/Indiana/26/2014(H3N2) | 8/12/2014 | GISAID |
| EPI564465 | A/Illinois/14/2014(H3N2) | 18/12/2014 | GISAID |
| EPI564473 | A/Colorado/35/2014(H3N2) | 12/12/2014 | GISAID |
| EPI564481 | A/Kentucky/21/2014(H3N2) | 17/12/2014 | GISAID |
| EPI564492 | A/Montana/28/2014(H3N2) | 12/12/2014 | GISAID |
| EPI564498 | A/Montana/30/2014(H3N2) | 18/12/2014 | GISAID |
| EPI564506 | A/Illinois/11/2014(H3N2) | 2/12/2014 | GISAID |
| EPI564520 | A/NewHampshire/32/2014(H3N2) | 8/12/2014 | GISAID |
| EPI564535 | A/Ontario/RV3241/2014(H3N2) | 25/10/2014 | GISAID |
| EPI564543 | A/Delaware/39/2014(H3N2) | 29/12/2014 | GISAID |
| EPI564554 | A/WestVirginia/06/2014(H3N2) | 14/12/2014 | GISAID |
| EPI564561 | A/Colorado/36/2014(H3N2) | 17/12/2014 | GISAID |
| EPI564574 | A/Delaware/42/2014(H3N2) | 22/12/2014 | GISAID |
| EPI566185 | A/Idaho/10/2014(H3N2) | 18/12/2014 | GISAID |
| EPI564585 | A/NewYork/39/2014(H3N2) | 31/12/2014 | GISAID |
| EPI564592 | A/NewYork/40/2014(H3N2) | 29/12/2014 | GISAID |
| EPI564612 | A/Arkansas/13/2014(H3N2) | 28/10/2014 | GISAID |
| EPI564618 | A/Washington/46/2014(H3N2) | 17/12/2014 | GISAID |
| EPI564630 | A/NorthCarolina/40/2014(H3N2) | 4/12/2014 | GISAID |
| EPI564634 | A/Hawaii/65/2014(H3N2) | 24/12/2014 | GISAID |
| EPI564642 | A/Nevada/30/2014(H3N2) | 17/12/2014 | GISAID |
| EPI564650 | A/Pennsylvania/65/2014(H3N2) | 9/12/2014 | GISAID |
| EPI564658 | A/NewHampshire/29/2014(H3N2) | 4/12/2014 | GISAID |
| EPI564666 | A/Pennsylvania/64/2014(H3N2) | 11/12/2014 | GISAID |
| EPI564674 | A/Pennsylvania/63/2014(H3N2) | 13/12/2014 | GISAID |
| EPI564679 | A/Ethiopia/1147/2014(H3N2) | 27/10/2014 | GISAID |
| EPI564692 | A/Virginia/37/2014(H3N2) | 5/12/2014 | GISAID |

| | | | |
|---|---|---|---|
| EPI564700 | A/Georgia/21/2014(H3N2) | 2/12/2014 | GISAID |
| EPI564716 | A/Missouri/14/2014(H3N2) | 2/12/2014 | GISAID |
| EPI564724 | A/DistrictOfColumbia/05/2014(H3N2) | 19/12/2014 | GISAID |
| EPI564732 | A/DistrictOfColumbia/04/2014(H3N2) | 26/12/2014 | GISAID |
| EPI564737 | A/DistrictOfColumbia/03/2014(H3N2) | 30/10/2014 | GISAID |
| EPI564745 | A/Florida/04/2015(H3N2) | 7/1/2015 | GISAID |
| EPI564753 | A/Florida/05/2015(H3N2) | 11/1/2015 | GISAID |
| EPI564766 | A/Ohio/14/2014(H3N2) | 9/11/2014 | GISAID |
| EPI564771 | A/California/50/2014(H3N2) | 18/10/2014 | GISAID |
| EPI564782 | A/California/52/2014(H3N2) | 24/10/2014 | GISAID |
| EPI564790 | A/Pennsylvania/51/2014(H3N2) | 2/12/2014 | GISAID |
| EPI564794 | A/California/53/2014(H3N2) | 26/10/2014 | GISAID |
| EPI564802 | A/Washington/47/2014(H3N2) | 17/12/2014 | GISAID |
| EPI564839 | A/Myanmar/13M016/2013(H3N2) | 11/7/2013 | GISAID |
| EPI564847 | A/Myanmar/13M029/2013(H3N2) | 15/7/2013 | GISAID |
| EPI564855 | A/Myanmar/13M056/2013(H3N2) | 16/7/2013 | GISAID |
| EPI564863 | A/Myanmar/13M082/2013(H3N2) | 19/7/2013 | GISAID |
| EPI565359 | A/Maine/03/2015(H3N2) | 1/1/2015 | GISAID |
| EPI565365 | A/Mexico/2789/2014(H3N2) | 26/11/2014 | GISAID |
| EPI565380 | A/Ethiopia/1140/2014(H3N2) | 20/10/2014 | GISAID |
| EPI565385 | A/Mexico/2777/2014(H3N2) | 24/11/2014 | GISAID |
| EPI565393 | A/Venezuela/07/2014(H3N2) | 4/8/2014 | GISAID |
| EPI565425 | A/Colorado/39/2014(H3N2) | 29/11/2014 | GISAID |
| EPI565433 | A/Tanzania/3580/2014(H3N2) | 26/11/2014 | GISAID |
| EPI565470 | A/Kentucky/01/2015(H3N2) | 1/1/2015 | GISAID |
| EPI565478 | A/Florida/85/2014(H3N2) | 3/12/2014 | GISAID |
| EPI565490 | A/Florida/09/2015(H3N2) | 5/1/2015 | GISAID |
| EPI565498 | A/Ohio/12/2014(H3N2) | 27/12/2014 | GISAID |
| EPI565506 | A/Florida/07/2015(H3N2) | 5/1/2015 | GISAID |
| EPI565514 | A/Utah/01/2015(H3N2) | 2/1/2015 | GISAID |
| EPI565522 | A/Montana/02/2015(H3N2) | 9/1/2015 | GISAID |
| EPI565528 | A/Florida/06/2015(H3N2) | 3/1/2015 | GISAID |
| EPI565536 | A/Montana/01/2015(H3N2) | 6/1/2015 | GISAID |
| EPI565544 | A/Tanzania/3562/2014(H3N2) | 20/11/2014 | GISAID |
| EPI565549 | A/Delaware/43/2014(H3N2) | 17/12/2014 | GISAID |
| EPI565557 | A/Tanzania/3557/2014(H3N2) | 18/11/2014 | GISAID |
| EPI565571 | A/Utah/02/2015(H3N2) | 8/1/2015 | GISAID |

| | | | |
|---|---|---|---|
| EPI565579 | A/WestVirginia/01/2015(H3N2) | 2/1/2015 | GISAID |
| EPI565587 | A/Tanzania/562/2014(H3N2) | 24/11/2014 | GISAID |
| EPI565595 | A/Washington/52/2014(H3N2) | 15/12/2014 | GISAID |
| EPI565603 | A/Wisconsin/01/2015(H3N2) | 2/1/2015 | GISAID |
| EPI565611 | A/Wisconsin/02/2015(H3N2) | 6/1/2015 | GISAID |
| EPI565619 | A/Tanzania/542/2014(H3N2) | 14/11/2014 | GISAID |
| EPI565624 | A/Mississippi/17/2014(H3N2) | 13/11/2014 | GISAID |
| EPI565636 | A/Arkansas/01/2015(H3N2) | 8/1/2015 | GISAID |
| EPI565648 | A/Oregon/01/2015(H3N2) | 1/1/2015 | GISAID |
| EPI565658 | A/Ontario/RV3531/2014(H3N2) | 3/11/2014 | GISAID |
| EPI565665 | A/Alberta/RV3591/2014(H3N2) | 21/11/2014 | GISAID |
| EPI565675 | A/Ontario/RV3609/2014(H3N2) | 18/12/2014 | GISAID |
| EPI565680 | A/HongKong/7615/2014(H3N2) | 29/11/2014 | GISAID |
| EPI565694 | A/Paraguay/4413/2014(H3N2) | 4/12/2014 | GISAID |
| EPI565707 | A/Tanzania/594/2014(H3N2) | 5/12/2014 | GISAID |
| EPI565715 | A/Tanzania/596/2014(H3N2) | 5/12/2014 | GISAID |
| EPI565723 | A/Pennsylvania/67/2014(H3N2) | 4/12/2014 | GISAID |
| EPI565731 | A/Tanzania/590/2014(H3N2) | 3/12/2014 | GISAID |
| EPI565739 | A/Tanzania/599/2014(H3N2) | 6/12/2014 | GISAID |
| EPI565747 | A/Santiago/68839/2014(H3N2) | 20/10/2014 | GISAID |
| EPI565755 | A/Ethiopia/159/2014(H3N2) | 4/9/2014 | GISAID |
| EPI565763 | A/Seoul/2007/2014(H3N2) | 22/12/2014 | GISAID |
| EPI565771 | A/Tanzania/3552/2014(H3N2) | 14/11/2014 | GISAID |
| EPI565777 | A/Delaware/02/2015(H3N2) | 2/1/2015 | GISAID |
| EPI565785 | A/Delaware/01/2015(H3N2) | 2/1/2015 | GISAID |
| EPI565792 | A/Tanzania/2665/2014(H3N2) | 25/11/2014 | GISAID |
| EPI565797 | A/Tanzania/588/2014(H3N2) | 1/12/2014 | GISAID |
| EPI565804 | A/Mexico/836/2014(H3N2) | 2/12/2014 | GISAID |
| EPI565812 | A/Tanzania/3588/2014(H3N2) | 3/12/2014 | GISAID |
| EPI565820 | A/Tanzania/609/2014(H3N2) | 12/12/2014 | GISAID |
| EPI565828 | A/Tanzania/3589/2014(H3N2) | 3/12/2014 | GISAID |
| EPI565836 | A/Mexico/2865/2014(H3N2) | 1/12/2014 | GISAID |
| EPI565844 | A/Mexico/3005/2014(H3N2) | 9/12/2014 | GISAID |
| EPI565849 | A/Mexico/3040/2014(H3N2) | 17/12/2014 | GISAID |
| EPI565859 | A/Florida/02/2015(H3N2) | 2/1/2015 | GISAID |
| EPI565867 | A/Florida/03/2015(H3N2) | 2/1/2015 | GISAID |
| EPI565875 | A/Vietnam/594/2014(H3N2) | 15/8/2014 | GISAID |

| | | | |
|---|---|---|---|
| EPI565883 | A/Tanzania/3555/2014(H3N2) | 16/11/2014 | GISAID |
| EPI565891 | A/Tanzania/3561/2014(H3N2) | 20/11/2014 | GISAID |
| EPI565903 | A/Tanzania/597/2014(H3N2) | 5/12/2014 | GISAID |
| EPI565911 | A/Tanzania/3556/2014(H3N2) | 18/11/2014 | GISAID |
| EPI565919 | A/Florida/01/2015(H3N2) | 2/1/2015 | GISAID |
| EPI565927 | A/Tanzania/3559/2014(H3N2) | 19/11/2014 | GISAID |
| EPI565935 | A/Tanzania/553/2014(H3N2) | 19/11/2014 | GISAID |
| EPI565945 | A/Ohio/15/2014(H3N2) | 11/11/2014 | GISAID |
| EPI565953 | A/Florida/12/2015(H3N2) | 7/1/2015 | GISAID |
| EPI565961 | A/Florida/11/2015(H3N2) | 6/1/2015 | GISAID |
| EPI565969 | A/Florida/13/2015(H3N2) | 11/1/2015 | GISAID |
| EPI565977 | A/Ethiopia/1159/2014(H3N2) | 12/11/2014 | GISAID |
| EPI565985 | A/Florida/10/2015(H3N2) | 2/1/2015 | GISAID |
| EPI565993 | A/Nonthaburi/2421/2014(H3N2) | 9/12/2014 | GISAID |
| EPI566001 | A/Tanzania/3594/2014(H3N2) | 11/12/2014 | GISAID |
| EPI566009 | A/Guatemala/179/2014(H3N2) | 4/11/2014 | GISAID |
| EPI566017 | A/Chanthaburi/2419/2014(H3N2) | 11/12/2014 | GISAID |
| EPI566025 | A/Tanzania/1906/2014(H3N2) | 9/12/2014 | GISAID |
| EPI566147 | A/Idaho/12/2014(H3N2) | 16/12/2014 | GISAID |
| EPI566159 | A/NewYork/38/2014(H3N2) | 31/12/2014 | GISAID |
| EPI566167 | A/Texas/85/2014(H3N2) | 11/12/2014 | GISAID |
| EPI566180 | A/Texas/84/2014(H3N2) | 11/12/2014 | GISAID |
| EPI566193 | A/Texas/88/2014(H3N2) | 5/12/2014 | GISAID |
| EPI566194 | A/Myanmar/13M006/2013(H3N2) | 4/7/2014 | GISAID |
| EPI566202 | A/Myanmar/13M011/2013(H3N2) | 9/7/2013 | GISAID |
| EPI566210 | A/Myanmar/13M020/2013(H3N2) | 11/7/2013 | GISAID |
| EPI566218 | A/Myanmar/13M035/2013(H3N2) | 15/7/2013 | GISAID |
| EPI566325 | A/Sweden/1/2015(H3N2) | 2/1/2015 | GISAID |
| EPI566340 | A/Sweden/4/2015(H3N2) | 14/1/2015 | GISAID |
| EPI566343 | A/Gavle/1/2015(H3N2) | 12/1/2015 | GISAID |
| EPI566352 | A/Lund/1/2015(H3N2) | 12/1/2015 | GISAID |
| EPI566383 | A/Stockholm/7/2015(H3N2) | 13/1/2015 | GISAID |
| EPI566386 | A/Stockholm/8/2015(H3N2) | 25/1/2015 | GISAID |
| EPI566389 | A/Stockholm/9/2015(H3N2) | 27/1/2015 | GISAID |
| EPI566439 | A/Stockholm/39/2014(H3N2) | 12/12/2014 | GISAID |
| EPI566527 | A/Myanmar/13M044/2013(H3N2) | 15/7/2013 | GISAID |
| EPI566535 | A/Myanmar/13M061/2013(H3N2) | 16/7/2013 | GISAID |

| | | | |
|---|---|---|---|
| EPI566543 | A/Myanmar/13M062/2013(H3N2) | 17/7/2013 | GISAID |
| EPI566615 | A/Myanmar/13M070/2013(H3N2) | 17/7/2013 | GISAID |
| EPI566623 | A/Myanmar/13M076/2013(H3N2) | 19/7/2013 | GISAID |
| EPI566631 | A/Myanmar/13M084/2013(H3N2) | 19/7/2013 | GISAID |
| EPI566639 | A/Myanmar/13M085/2013(H3N2) | 22/7/2013 | GISAID |
| EPI566673 | A/Myanmar/13M087/2013(H3N2) | 22/7/2013 | GISAID |
| EPI566681 | A/Myanmar/13M088/2013(H3N2) | 22/7/2013 | GISAID |
| EPI566707 | A/Myanmar/13M089/2013(H3N2) | 22/7/2013 | GISAID |
| EPI566715 | A/Myanmar/13M098/2013(H3N2) | 24/7/2013 | GISAID |
| EPI566723 | A/Myanmar/13M102/2013(H3N2) | 24/7/2013 | GISAID |
| EPI566731 | A/Myanmar/13M105/2013(H3N2) | 25/7/2013 | GISAID |
| EPI566739 | A/Myanmar/13M109/2013(H3N2) | 25/7/2013 | GISAID |
| EPI566747 | A/Myanmar/13M118/2013(H3N2) | 26/7/2013 | GISAID |
| EPI566755 | A/Myanmar/13M124/2013(H3N2) | 16/7/2013 | GISAID |
| EPI566763 | A/Myanmar/13M300/2013(H3N2) | 22/8/2013 | GISAID |
| EPI566863 | A/MIYAGI/52/2014(H3N2) | 1/12/2014 | GISAID |
| EPI566871 | A/FUKUI/61/2014(H3N2) | 7/12/2014 | GISAID |
| EPI566901 | A/NIIGATA/35/2015(H3N2) | 11/1/2015 | GISAID |
| EPI566924 | A/Nagasaki/13N100/2014(H3N2) | 29/3/2014 | GISAID |
| EPI566932 | A/Nagasaki/13N101/2014(H3N2) | 29/7/2014 | GISAID |
| EPI566940 | A/Nagasaki/14N005/2014(H3N2) | 22/9/2014 | GISAID |
| EPI567319 | A/Texas/04/2015(H3N2) | 7/1/2015 | GISAID |
| EPI567327 | A/Alabama/01/2015(H3N2) | 6/1/2015 | GISAID |
| EPI567335 | A/Michigan/31/2014(H3N2) | 22/12/2014 | GISAID |
| EPI567343 | A/Michigan/30/2014(H3N2) | 16/12/2014 | GISAID |
| EPI567351 | A/NorthCarolina/06/2015(H3N2) | 7/1/2015 | GISAID |
| EPI567359 | A/Michigan/28/2014(H3N2) | 29/12/2014 | GISAID |
| EPI567367 | A/NorthCarolina/04/2015(H3N2) | 13/1/2015 | GISAID |
| EPI567375 | A/Mississippi/02/2015(H3N2) | 5/1/2015 | GISAID |
| EPI567383 | A/Michigan/03/2015(H3N2) | 2/1/2015 | GISAID |
| EPI567391 | A/Florida/92/2014(H3N2) | 22/12/2014 | GISAID |
| EPI567402 | A/Washington/02/2015(H3N2) | 2/1/2015 | GISAID |
| EPI567410 | A/NorthCarolina/05/2015(H3N2) | 8/1/2015 | GISAID |
| EPI567424 | A/Kentucky/24/2014(H3N2) | 22/12/2014 | GISAID |
| EPI567432 | A/Texas/05/2015(H3N2) | 4/1/2015 | GISAID |
| EPI567440 | A/Texas/06/2015(H3N2) | 9/1/2015 | GISAID |
| EPI567448 | A/Kentucky/27/2014(H3N2) | 23/12/2014 | GISAID |

| | | | |
|---|---|---|---|
| EPI567456 | A/Michigan/32/2014(H3N2) | 26/12/2014 | GISAID |
| EPI567462 | A/NorthDakota/02/2015(H3N2) | 1/1/2015 | GISAID |
| EPI567468 | A/Michigan/27/2014(H3N2) | 12/12/2014 | GISAID |
| EPI567476 | A/Washington/57/2014(H3N2) | 29/12/2014 | GISAID |
| EPI567484 | A/Florida/14/2015(H3N2) | 18/1/2015 | GISAID |
| EPI567496 | A/Wisconsin/03/2015(H3N2) | 2/1/2015 | GISAID |
| EPI567504 | A/Washington/01/2015(H3N2) | 21/1/2015 | GISAID |
| EPI567512 | A/Michigan/02/2015(H3N2) | 5/1/2015 | GISAID |
| EPI567539 | A/Tennessee/01/2015(H3N2) | 11/1/2015 | GISAID |
| EPI567554 | A/NorthCarolina/02/2015(H3N2) | 28/1/2015 | GISAID |
| EPI567562 | A/Texas/107/2014(H3N2) | 15/12/2014 | GISAID |
| EPI567570 | A/Texas/106/2014(H3N2) | 20/12/2014 | GISAID |
| EPI567583 | A/Texas/105/2014(H3N2) | 26/12/2014 | GISAID |
| EPI567591 | A/Kentucky/25/2014(H3N2) | 22/12/2014 | GISAID |
| EPI567604 | A/Wisconsin/72/2014(H3N2) | 19/12/2014 | GISAID |
| EPI567612 | A/Mississippi/01/2015(H3N2) | 4/1/2015 | GISAID |
| EPI567620 | A/Washington/53/2014(H3N2) | 26/12/2014 | GISAID |
| EPI567628 | A/Wisconsin/75/2014(H3N2) | 22/12/2014 | GISAID |
| EPI567788 | A/Victoria/186/1982(H3N2) | 1982 | GISAID |
| EPI567796 | A/Sydney/2/1993(H3N2) | 1993 | GISAID |
| EPI567881 | A/Vietnam/13VH3-2/2012(H3N2) | 28/12/2012 | GISAID |
| EPI567889 | A/Vietnam/13VH3-3/2012(H3N2) | 23/7/2012 | GISAID |
| EPI567910 | A/Vietnam/13VH3-4/2013(H3N2) | 13/4/2013 | GISAID |
| EPI567927 | A/Vietnam/13VH3-8/2013(H3N2) | 25/9/2013 | GISAID |
| EPI567926 | A/Vietnam/13VH3-8/2013(H3N2) | 25/9/2013 | GISAID |
| EPI567935 | A/Vietnam/13VH3-9/2012(H3N2) | 27/7/2012 | GISAID |
| EPI567943 | A/Vietnam/13VH3-10/2012(H3N2) | 25/7/2012 | GISAID |
| EPI568050 | A/Myanmar/14M039/2014(H3N2) | 24/6/2014 | GISAID |
| EPI568058 | A/Myanmar/14M059/2014(H3N2) | 14/7/2014 | GISAID |
| EPI568066 | A/Myanmar/14M122/2014(H3N2) | 8/7/2014 | GISAID |
| EPI568074 | A/Myanmar/14M187/2014(H3N2) | 2/7/2014 | GISAID |
| EPI568082 | A/Myanmar/14M240/2014(H3N2) | 4/8/2014 | GISAID |
| EPI568090 | A/Myanmar/14M391/2014(H3N2) | 22/8/2014 | GISAID |
| EPI568098 | A/Myanmar/14M465/2014(H3N2) | 8/9/2014 | GISAID |
| EPI569302 | A/Utah/19/2014(H3N2) | 7/11/2014 | GISAID |
| EPI569312 | A/Louisiana/21/2014(H3N2) | 19/11/2014 | GISAID |
| EPI570014 | A/Sweden/8/2015(H3N2) | 1/2/2015 | GISAID |

| | | | |
|---|---|---|---|
| EPI570118 | A/Sweden/11/2015(H3N2) | 15/2/2015 | GISAID |
| EPI570137 | A/Sweden/12/2015(H3N2) | 15/2/2015 | GISAID |
| EPI571782 | A/RhodeIsland/03/2015(H3N2) | 6/1/2015 | GISAID |
| EPI571803 | A/Tanzania/3565/2014(H3N2) | 20/11/2014 | GISAID |
| EPI571811 | A/Tanzania/3576/2014(H3N2) | 24/11/2014 | GISAID |
| EPI571819 | A/Tanzania/3577/2014(H3N2) | 24/11/2014 | GISAID |
| EPI571827 | A/NewYork/09/2015(H3N2) | 8/1/2015 | GISAID |
| EPI571854 | A/RhodeIsland/02/2015(H3N2) | 5/1/2015 | GISAID |
| EPI571879 | A/Oklahoma/02/2015(H3N2) | 3/1/2015 | GISAID |
| EPI571887 | A/Tanzania/3560/2014(H3N2) | 19/11/2014 | GISAID |
| EPI571902 | A/NorthCarolina/08/2015(H3N2) | 6/1/2015 | GISAID |
| EPI571915 | A/Oregon/02/2015(H3N2) | 2/1/2015 | GISAID |
| EPI571930 | A/Utah/03/2015(H3N2) | 12/1/2015 | GISAID |
| EPI571945 | A/Nebraska/01/2015(H3N2) | 5/1/2015 | GISAID |
| EPI571960 | A/Nevada/04/2015(H3N2) | 7/1/2015 | GISAID |
| EPI571975 | A/NewHampshire/05/2015(H3N2) | 22/1/2015 | GISAID |
| EPI572009 | A/NorthDakota/03/2015(H3N2) | 1/1/2015 | GISAID |
| EPI572017 | A/Montana/04/2015(H3N2) | 18/1/2015 | GISAID |
| EPI572052 | A/NewHampshire/04/2015(H3N2) | 13/1/2015 | GISAID |
| EPI572060 | A/NewYork/04/2015(H3N2) | 22/1/2015 | GISAID |
| EPI572071 | A/Massachusetts/02/2015(H3N2) | 20/1/2015 | GISAID |
| EPI572105 | A/Missouri/04/2015(H3N2) | 15/1/2015 | GISAID |
| EPI572119 | A/Arkansas/02/2015(H3N2) | 10/1/2015 | GISAID |
| EPI572134 | A/Michigan/04/2015(H3N2) | 6/1/2015 | GISAID |
| EPI572199 | A/California/01/2015(H3N2) | 4/1/2015 | GISAID |
| EPI572227 | A/Missouri/01/2015(H3N2) | 3/1/2015 | GISAID |
| EPI572249 | A/NorthCarolina/07/2015(H3N2) | 14/1/2015 | GISAID |
| EPI577029 | A/Canberra/82/2014(H3N2) | 31/7/2014 | GISAID |
| EPI574877 | A/Colorado/02/2015(H3N2) | 9/1/2015 | GISAID |
| EPI574885 | A/Tennessee/09/2015(H3N2) | 2/2/2015 | GISAID |
| EPI574894 | A/Michigan/07/2015(H3N2) | 17/1/2015 | GISAID |
| EPI574902 | A/Tennessee/07/2015(H3N2) | 2/2/2015 | GISAID |
| EPI574910 | A/Tennessee/08/2015(H3N2) | 3/2/2015 | GISAID |
| EPI574921 | A/Michigan/05/2015(H3N2) | 8/1/2015 | GISAID |
| EPI574944 | A/Kansas/03/2015(H3N2) | 18/1/2015 | GISAID |
| EPI574963 | A/Indiana/01/2015(H3N2) | 8/1/2015 | GISAID |
| EPI574971 | A/Utah/05/2015(H3N2) | 12/1/2015 | GISAID |

| | | | |
|---|---|---|---|
| EPI574992 | A/Colorado/04/2015(H3N2) | 5/1/2015 | GISAID |
| EPI575017 | A/RhodeIsland/06/2015(H3N2) | 6/1/2015 | GISAID |
| EPI575040 | A/NewJersey/03/2015(H3N2) | 19/1/2015 | GISAID |
| EPI575060 | A/NewYork/07/2015(H3N2) | 7/1/2015 | GISAID |
| EPI575079 | A/NorthCarolina/11/2015(H3N2) | 9/2/2015 | GISAID |
| EPI575087 | A/Ohio/04/2015(H3N2) | 6/2/2015 | GISAID |
| EPI575095 | A/Florida/19/2015(H3N2) | 1/2/2015 | GISAID |
| EPI575108 | A/Iowa/31/2014(H3N2) | 25/12/2014 | GISAID |
| EPI575122 | A/Iowa/04/2015(H3N2) | 6/1/2015 | GISAID |
| EPI575158 | A/Cambodia/1384/2014(H3N2) | 17/12/2014 | GISAID |
| EPI575166 | A/NewMexico/01/2015(H3N2) | 3/1/2015 | GISAID |
| EPI575174 | A/Florida/20/2015(H3N2) | 30/1/2015 | GISAID |
| EPI575183 | A/Hawaii/02/2015(H3N2) | 3/2/2015 | GISAID |
| EPI575191 | A/Florida/18/2015(H3N2) | 2/2/2015 | GISAID |
| EPI575199 | A/Utah/06/2015(H3N2) | 29/1/2015 | GISAID |
| EPI575212 | A/Utah/07/2015(H3N2) | 10/2/2015 | GISAID |
| EPI575240 | A/Delaware/05/2015(H3N2) | 30/1/2015 | GISAID |
| EPI575248 | A/NewYork/15/2015(H3N2) | 5/2/2015 | GISAID |
| EPI575256 | A/NewHampshire/06/2015(H3N2) | 29/1/2015 | GISAID |
| EPI575264 | A/Tennessee/24/2014(H3N2) | 2/12/2014 | GISAID |
| EPI575271 | A/NewHampshire/07/2015(H3N2) | 28/1/2015 | GISAID |
| EPI575279 | A/Nevada/03/2015(H3N2) | 8/1/2015 | GISAID |
| EPI575287 | A/Nevada/02/2015(H3N2) | 14/1/2015 | GISAID |
| EPI575295 | A/Texas/93/2014(H3N2) | 16/12/2014 | GISAID |
| EPI575303 | A/Texas/96/2014(H3N2) | 26/12/2014 | GISAID |
| EPI575310 | A/Nevada/09/2015(H3N2) | 5/2/2015 | GISAID |
| EPI575318 | A/Nevada/10/2015(H3N2) | 14/2/2015 | GISAID |
| EPI575326 | A/Wisconsin/05/2015(H3N2) | 27/1/2015 | GISAID |
| EPI575334 | A/Kansas/02/2015(H3N2) | 30/1/2015 | GISAID |
| EPI575344 | A/Cambodia/1370/2014(H3N2) | 11/12/2014 | GISAID |
| EPI575352 | A/Cambodia/1372/2014(H3N2) | 11/12/2014 | GISAID |
| EPI575370 | A/Tanzania/613/2014(H3N2) | 17/12/2014 | GISAID |
| EPI575378 | A/Maine/07/2015(H3N2) | 17/1/2015 | GISAID |
| EPI575386 | A/Nebraska/03/2015(H3N2) | 9/1/2015 | GISAID |
| EPI594994 | A/Chita/262/2014(H3N2) | 16/12/2014 | GISAID |
| EPI575400 | A/Chita/256/2014(H3N2) | 11/12/2014 | GISAID |
| EPI575408 | A/Tanzania/3595/2014(H3N2) | 11/12/2014 | GISAID |

| | | | |
|---|---|---|---|
| EPI575416 | A/Chita/260/2014(H3N2) | 12/12/2014 | GISAID |
| EPI575427 | A/Chita/258/2014(H3N2) | 15/12/2014 | GISAID |
| EPI575435 | A/Tanzania/3591/2014(H3N2) | 8/12/2014 | GISAID |
| EPI575443 | A/NewYork/12/2015(H3N2) | 22/1/2015 | GISAID |
| EPI575451 | A/Massachusetts/03/2015(H3N2) | 20/1/2015 | GISAID |
| EPI575462 | A/Chita/264/2014(H3N2) | 12/12/2014 | GISAID |
| EPI575470 | A/Chita/268/2014(H3N2) | 10/12/2014 | GISAID |
| EPI575478 | A/Congo/2471/2014(H3N2) | 9/12/2014 | GISAID |
| EPI575486 | A/Congo/2458/2014(H3N2) | 4/12/2014 | GISAID |
| EPI575501 | A/Washington/04/2015(H3N2) | 30/1/2015 | GISAID |
| EPI575509 | A/Kansas/01/2015(H3N2) | 1/2/2015 | GISAID |
| EPI575517 | A/Georgia/30/2014(H3N2) | 8/12/2014 | GISAID |
| EPI575525 | A/Pennsylvania/05/2015(H3N2) | 6/2/2015 | GISAID |
| EPI575533 | A/Congo/2461/2014(H3N2) | 8/12/2014 | GISAID |
| EPI575541 | A/Pennsylvania/03/2015(H3N2) | 5/2/2015 | GISAID |
| EPI575549 | A/NewYork/16/2015(H3N2) | 9/2/2015 | GISAID |
| EPI575557 | A/NewJersey/02/2015(H3N2) | 26/1/2015 | GISAID |
| EPI575564 | A/Chita/266/2014(H3N2) | 3/12/2014 | GISAID |
| EPI575572 | A/Tomsk/254/2014(H3N2) | 12/12/2014 | GISAID |
| EPI575580 | A/Astrakhan/24/2014(H3N2) | 16/12/2014 | GISAID |
| EPI575588 | A/Texas/09/2015(H3N2) | 3/2/2015 | GISAID |
| EPI575597 | A/NewYork/14/2015(H3N2) | 5/2/2015 | GISAID |
| EPI575605 | A/Nebraska/05/2015(H3N2) | 31/1/2015 | GISAID |
| EPI575614 | A/WestVirginia/02/2015(H3N2) | 28/1/2015 | GISAID |
| EPI575621 | A/Delaware/06/2015(H3N2) | 28/1/2015 | GISAID |
| EPI575632 | A/NewYork/13/2015(H3N2) | 10/2/2015 | GISAID |
| EPI575649 | A/Georgia/24/2014(H3N2) | 1/12/2014 | GISAID |
| EPI575661 | A/Kazakhstan/15/2014(H3N2) | 29/12/2014 | GISAID |
| EPI575669 | A/Kazakhstan/22/2015(H3N2) | 2/1/2015 | GISAID |
| EPI575677 | A/Kazakhstan/29/2015(H3N2) | 6/1/2015 | GISAID |
| EPI575685 | A/Kazakhstan/27/2015(H3N2) | 5/1/2015 | GISAID |
| EPI575693 | A/Kazakhstan/30/2015(H3N2) | 6/1/2015 | GISAID |
| EPI575701 | A/Kazakhstan/33/2015(H3N2) | 6/1/2015 | GISAID |
| EPI575709 | A/Kazakhstan/20/2015(H3N2) | 5/1/2015 | GISAID |
| EPI575717 | A/Kazakhstan/16/2015(H3N2) | 1/1/2015 | GISAID |
| EPI575725 | A/Kazakhstan/35/2015(H3N2) | 7/1/2015 | GISAID |
| EPI575733 | A/Kazakhstan/31/2015(H3N2) | 6/1/2015 | GISAID |

| | | | |
|---|---|---|---|
| EPI575741 | A/Kazakhstan/25/2015(H3N2) | 5/1/2015 | GISAID |
| EPI575749 | A/Kazakhstan/32/2015(H3N2) | 13/1/2015 | GISAID |
| EPI575757 | A/Kazakhstan/42/2015(H3N2) | 13/1/2015 | GISAID |
| EPI575765 | A/Kazakhstan/36/2015(H3N2) | 13/1/2015 | GISAID |
| EPI575773 | A/Kazakhstan/34/2015(H3N2) | 6/1/2015 | GISAID |
| EPI575781 | A/Kazakhstan/13/2015(H3N2) | 13/1/2015 | GISAID |
| EPI575789 | A/Kazakhstan/05/2015(H3N2) | 13/1/2015 | GISAID |
| EPI575797 | A/Kazakhstan/26/2015(H3N2) | 5/1/2015 | GISAID |
| EPI575805 | A/Kazakhstan/03/2015(H3N2) | 13/1/2015 | GISAID |
| EPI575813 | A/Kazakhstan/49/2015(H3N2) | 13/1/2015 | GISAID |
| EPI575821 | A/Georgia/39/2014(H3N2) | 16/12/2014 | GISAID |
| EPI575840 | A/Pennsylvania/04/2015(H3N2) | 27/1/2015 | GISAID |
| EPI576096 | A/WAKAYAMA/167/2014(H3N2) | 17/12/2014 | GISAID |
| EPI576098 | A/TOKYO/34558/2014(H3N2) | 22/12/2014 | GISAID |
| EPI576116 | A/IBARAKI/1/2015(H3N2) | 9/1/2015 | GISAID |
| EPI576132 | A/TOKYO/41945/2015(H3N2) | 20/1/2015 | GISAID |
| EPI576136 | A/IWATE/7/2015(H3N2) | 21/1/2015 | GISAID |
| EPI577583 | A/Niigata/14F002/2015(H3N2) | 13/1/2015 | GISAID |
| EPI577588 | A/Niigata/14F004/2015(H3N2) | 13/1/2015 | GISAID |
| EPI577596 | A/Niigata/14F005/2015(H3N2) | 13/1/2015 | GISAID |
| EPI577604 | A/Niigata/14F007/2015(H3N2) | 13/1/2015 | GISAID |
| EPI577612 | A/Niigata/14F009/2015(H3N2) | 13/1/2015 | GISAID |
| EPI577620 | A/Kyoto/14K001/2014(H3N2) | 8/12/2014 | GISAID |
| EPI577628 | A/Kyoto/14K002/2014(H3N2) | 8/12/2014 | GISAID |
| EPI577636 | A/Kyoto/14K003/2014(H3N2) | 9/12/2014 | GISAID |
| EPI577644 | A/Kyoto/14K005/2014(H3N2) | 9/12/2014 | GISAID |
| EPI577652 | A/Kyoto/14K007/2014(H3N2) | 15/12/2014 | GISAID |
| EPI577660 | A/Gunma/14G002/2015(H3N2) | 10/1/2015 | GISAID |
| EPI577668 | A/Gunma/14G003/2015(H3N2) | 13/1/2015 | GISAID |
| EPI577676 | A/Gunma/14G004/2015(H3N2) | 13/1/2015 | GISAID |
| EPI577684 | A/Gunma/14G005/2015(H3N2) | 13/1/2015 | GISAID |
| EPI577692 | A/Gunma/14G008/2015(H3N2) | 14/1/2015 | GISAID |
| EPI577700 | A/Nagasaki/14N014/2014(H3N2) | 12/12/2014 | GISAID |
| EPI577708 | A/Nagasaki/14N015/2014(H3N2) | 15/12/2014 | GISAID |
| EPI577716 | A/Nagasaki/14N017/2014(H3N2) | 24/12/2014 | GISAID |
| EPI577724 | A/Nagasaki/14N018/2014(H3N2) | 25/12/2014 | GISAID |
| EPI577732 | A/Nagasaki/14N023/2014(H3N2) | 28/12/2014 | GISAID |

| | | | |
|---|---|---|---|
| EPI577748 | A/Nagasaki/14N010/2014(H3N2) | 4/11/2014 | GISAID |
| EPI578251 | A/Mississippi/04/2015(H3N2) | 9/1/2015 | GISAID |
| EPI578259 | A/Florida/22/2015(H3N2) | 13/1/2015 | GISAID |
| EPI578267 | A/Mississippi/05/2015(H3N2) | 15/1/2015 | GISAID |
| EPI578275 | A/Michigan/09/2015(H3N2) | 12/1/2015 | GISAID |
| EPI578283 | A/Pennsylvania/07/2015(H3N2) | 9/1/2015 | GISAID |
| EPI578298 | A/Pennsylvania/08/2015(H3N2) | 7/1/2015 | GISAID |
| EPI578306 | A/Texas/13/2015(H3N2) | 10/1/2015 | GISAID |
| EPI578314 | A/Alaska/01/2015(H3N2) | 9/1/2015 | GISAID |
| EPI578322 | A/NewMexico/02/2015(H3N2) | 9/1/2015 | GISAID |
| EPI578337 | A/Hawaii/04/2015(H3N2) | 10/1/2015 | GISAID |
| EPI578363 | A/Pennsylvania/06/2015(H3N2) | 7/1/2015 | GISAID |
| EPI578376 | A/Pennsylvania/09/2015(H3N2) | 7/1/2015 | GISAID |
| EPI578384 | A/Texas/12/2015(H3N2) | 10/1/2015 | GISAID |
| EPI578398 | A/NewMexico/04/2015(H3N2) | 11/1/2015 | GISAID |
| EPI578406 | A/Pennsylvania/10/2015(H3N2) | 8/1/2015 | GISAID |
| EPI578414 | A/NewHampshire/10/2015(H3N2) | 11/1/2015 | GISAID |
| EPI578422 | A/Delaware/09/2015(H3N2) | 24/1/2015 | GISAID |
| EPI578430 | A/HongKong/4801/2014(H3N2) | 2014 | GISAID |
| EPI578438 | A/Congo/2538/2014(H3N2) | 18/12/2014 | GISAID |
| EPI578446 | A/HongKong/7127/2014(H3N2) | 2014 | GISAID |
| EPI578454 | A/Congo/2544/2014(H3N2) | 17/12/2014 | GISAID |
| EPI578469 | A/Kazakhstan/951/2015(H3N2) | 13/1/2015 | GISAID |
| EPI578477 | A/Kazakhstan/37/2015(H3N2) | 13/1/2015 | GISAID |
| EPI578485 | A/Kazakhstan/47/2015(H3N2) | 13/1/2015 | GISAID |
| EPI578493 | A/Kazakhstan/38/2015(H3N2) | 8/1/2015 | GISAID |
| EPI578501 | A/Idaho/08/2015(H3N2) | 5/1/2015 | GISAID |
| EPI578509 | A/Iowa/06/2015(H3N2) | 15/1/2015 | GISAID |
| EPI578517 | A/California/09/2015(H3N2) | 6/1/2015 | GISAID |
| EPI578525 | A/Iowa/05/2015(H3N2) | 13/1/2015 | GISAID |
| EPI578533 | A/Laos/732/2014(H3N2) | 21/11/2014 | GISAID |
| EPI578560 | A/California/08/2015(H3N2) | 5/1/2015 | GISAID |
| EPI578571 | A/California/06/2015(H3N2) | 1/1/2015 | GISAID |
| EPI578579 | A/Maine/09/2015(H3N2) | 13/2/2015 | GISAID |
| EPI578587 | A/California/04/2015(H3N2) | 7/1/2015 | GISAID |
| EPI578595 | A/Nebraska/07/2015(H3N2) | 14/1/2015 | GISAID |
| EPI578603 | A/Maine/10/2015(H3N2) | 18/2/2015 | GISAID |

| | | | |
|---|---|---|---|
| EPI578611 | A/Missouri/07/2015(H3N2) | 15/1/2015 | GISAID |
| EPI578619 | A/Massachusetts/05/2015(H3N2) | 12/1/2015 | GISAID |
| EPI578627 | A/Minnesota/01/2015(H3N2) | 4/1/2015 | GISAID |
| EPI578635 | A/Minnesota/02/2015(H3N2) | 7/1/2015 | GISAID |
| EPI578649 | A/Georgia/02/2015(H3N2) | 2015 | GISAID |
| EPI578657 | A/Georgia/01/2015(H3N2) | 2015 | GISAID |
| EPI578665 | A/Virginia/02/2015(H3N2) | 4/1/2015 | GISAID |
| EPI578673 | A/Virginia/06/2015(H3N2) | 3/1/2015 | GISAID |
| EPI578681 | A/Virginia/03/2015(H3N2) | 4/1/2015 | GISAID |
| EPI578694 | A/Congo/2611/2014(H3N2) | 29/12/2014 | GISAID |
| EPI578702 | A/Congo/2557/2014(H3N2) | 18/12/2014 | GISAID |
| EPI578710 | A/Washington/03/2015(H3N2) | 13/1/2015 | GISAID |
| EPI578718 | A/WestVirginia/03/2015(H3N2) | 12/1/2015 | GISAID |
| EPI578726 | A/Washington/17/2015(H3N2) | 14/1/2015 | GISAID |
| EPI578734 | A/Washington/08/2015(H3N2) | 12/1/2015 | GISAID |
| EPI578742 | A/Texas/22/2015(H3N2) | 13/1/2015 | GISAID |
| EPI578750 | A/Wisconsin/10/2015(H3N2) | 10/1/2015 | GISAID |
| EPI578758 | A/Wisconsin/11/2015(H3N2) | 11/1/2015 | GISAID |
| EPI578773 | A/Delaware/14/2015(H3N2) | 15/1/2015 | GISAID |
| EPI578781 | A/Delaware/12/2015(H3N2) | 9/1/2015 | GISAID |
| EPI578794 | A/Michigan/11/2015(H3N2) | 9/1/2015 | GISAID |
| EPI578802 | A/Texas/21/2015(H3N2) | 14/1/2015 | GISAID |
| EPI578810 | A/Delaware/10/2015(H3N2) | 1/1/2015 | GISAID |
| EPI578818 | A/Kansas/05/2015(H3N2) | 14/1/2015 | GISAID |
| EPI578826 | A/Kansas/06/2015(H3N2) | 15/1/2015 | GISAID |
| EPI578834 | A/Texas/20/2015(H3N2) | 13/1/2015 | GISAID |
| EPI578842 | A/Michigan/13/2015(H3N2) | 9/1/2015 | GISAID |
| EPI578850 | A/Michigan/12/2015(H3N2) | 8/1/2015 | GISAID |
| EPI578858 | A/Cambodia/870/2014(H3N2) | 5/8/2014 | GISAID |
| EPI578866 | A/Illinois/02/2015(H3N2) | 14/1/2015 | GISAID |
| EPI578874 | A/Uganda/4530/2014(H3N2) | 2014 | GISAID |
| EPI578881 | A/Tennessee/06/2015(H3N2) | 6/1/2015 | GISAID |
| EPI578889 | A/Tennessee/05/2015(H3N2) | 6/1/2015 | GISAID |
| EPI578897 | A/Michigan/08/2015(H3N2) | 15/1/2015 | GISAID |
| EPI578910 | A/Michigan/10/2015(H3N2) | 15/1/2015 | GISAID |
| EPI579060 | A/Kazakhstan/39/2015(H3N2) | 8/1/2015 | GISAID |
| EPI579066 | A/NewMexico/06/2015(H3N2) | 27/2/2015 | GISAID |

| | | | |
|---|---|---|---|
| EPI579074 | A/NewYork/18/2015(H3N2) | 23/1/2015 | GISAID |
| EPI579086 | A/Maine/15/2015(H3N2) | 16/1/2015 | GISAID |
| EPI579094 | A/Maine/13/2015(H3N2) | 16/1/2015 | GISAID |
| EPI579102 | A/Maine/14/2015(H3N2) | 16/1/2015 | GISAID |
| EPI579110 | A/Missouri/09/2015(H3N2) | 24/1/2015 | GISAID |
| EPI579118 | A/Delaware/16/2015(H3N2) | 19/1/2015 | GISAID |
| EPI579140 | A/Texas/34/2015(H3N2) | 26/1/2015 | GISAID |
| EPI579148 | A/Texas/32/2015(H3N2) | 26/1/2015 | GISAID |
| EPI579156 | A/Texas/30/2015(H3N2) | 26/1/2015 | GISAID |
| EPI579169 | A/Texas/27/2015(H3N2) | 28/1/2015 | GISAID |
| EPI579177 | A/Texas/33/2015(H3N2) | 27/1/2015 | GISAID |
| EPI579185 | A/Texas/31/2015(H3N2) | 27/1/2015 | GISAID |
| EPI579193 | A/NewJersey/09/2015(H3N2) | 26/1/2015 | GISAID |
| EPI579201 | A/NewJersey/10/2015(H3N2) | 26/1/2015 | GISAID |
| EPI579209 | A/NewHampshire/12/2015(H3N2) | 1/2/2015 | GISAID |
| EPI579217 | A/WestVirginia/06/2015(H3N2) | 22/1/2015 | GISAID |
| EPI579225 | A/WestVirginia/07/2015(H3N2) | 25/1/2015 | GISAID |
| EPI579244 | A/Wisconsin/15/2015(H3N2) | 3/2/2015 | GISAID |
| EPI579264 | A/Texas/11/2015(H3N2) | 16/2/2015 | GISAID |
| EPI579270 | A/NewHampshire/18/2015(H3N2) | 11/2/2015 | GISAID |
| EPI579278 | A/Wisconsin/07/2015(H3N2) | 13/2/2015 | GISAID |
| EPI579286 | A/Washington/05/2015(H3N2) | 6/2/2015 | GISAID |
| EPI579298 | A/Washington/16/2015(H3N2) | 13/2/2015 | GISAID |
| EPI579303 | A/Wisconsin/08/2015(H3N2) | 6/2/2015 | GISAID |
| EPI579309 | A/Wisconsin/14/2015(H3N2) | 10/2/2015 | GISAID |
| EPI579314 | A/Wisconsin/09/2015(H3N2) | 18/2/2015 | GISAID |
| EPI579323 | A/Missouri/06/2015(H3N2) | 9/2/2015 | GISAID |
| EPI579329 | A/RhodeIsland/08/2015(H3N2) | 4/2/2015 | GISAID |
| EPI579335 | A/Virginia/04/2015(H3N2) | 4/2/2015 | GISAID |
| EPI579342 | A/NewJersey/04/2015(H3N2) | 1/2/2015 | GISAID |
| EPI579349 | A/Delaware/13/2015(H3N2) | 24/2/2015 | GISAID |
| EPI622303 | A/NewJersey/05/2015(H3N2) | 23/2/2015 | GISAID |
| EPI579359 | A/Victoria/5006/2014(H3N2) | 25/8/2014 | GISAID |
| EPI579366 | A/Cambodia/1244/2014(H3N2) | 11/11/2014 | GISAID |
| EPI579380 | A/Utah/11/2015(H3N2) | 20/2/2015 | GISAID |
| EPI579388 | A/Idaho/10/2015(H3N2) | 5/1/2015 | GISAID |
| EPI579396 | A/Nevada/15/2015(H3N2) | 23/1/2015 | GISAID |

| | | | |
|---|---|---|---|
| EPI579404 | A/NewMexico/07/2015(H3N2) | 20/1/2015 | GISAID |
| EPI579412 | A/NewMexico/08/2015(H3N2) | 19/1/2015 | GISAID |
| EPI579420 | A/Arkansas/05/2015(H3N2) | 20/1/2015 | GISAID |
| EPI579428 | A/Arkansas/04/2015(H3N2) | 21/1/2015 | GISAID |
| EPI579436 | A/Texas/26/2015(H3N2) | 19/1/2015 | GISAID |
| EPI579444 | A/Texas/24/2015(H3N2) | 21/1/2015 | GISAID |
| EPI579452 | A/Texas/29/2015(H3N2) | 24/1/2015 | GISAID |
| EPI579460 | A/Texas/23/2015(H3N2) | 21/1/2015 | GISAID |
| EPI579468 | A/Texas/28/2015(H3N2) | 21/1/2015 | GISAID |
| EPI579476 | A/Washington/06/2015(H3N2) | 27/1/2015 | GISAID |
| EPI579484 | A/Idaho/04/2015(H3N2) | 26/1/2015 | GISAID |
| EPI579492 | A/Texas/16/2015(H3N2) | 4/2/2015 | GISAID |
| EPI579500 | A/Texas/18/2015(H3N2) | 8/2/2015 | GISAID |
| EPI622345 | A/Iowa/09/2015(H3N2) | 3/2/2015 | GISAID |
| EPI579516 | A/Nevada/13/2015(H3N2) | 11/2/2015 | GISAID |
| EPI579522 | A/Idaho/05/2015(H3N2) | 28/1/2015 | GISAID |
| EPI579530 | A/Idaho/09/2015(H3N2) | 4/2/2015 | GISAID |
| EPI622096 | A/Idaho/06/2015(H3N2) | 9/2/2015 | GISAID |
| EPI579539 | A/Oklahoma/05/2015(H3N2) | 11/2/2015 | GISAID |
| EPI579546 | A/Oklahoma/06/2015(H3N2) | 11/2/2015 | GISAID |
| EPI579553 | A/California/16/2015(H3N2) | 12/2/2015 | GISAID |
| EPI579559 | A/California/12/2015(H3N2) | 15/2/2015 | GISAID |
| EPI579567 | A/Kansas/07/2015(H3N2) | 16/1/2015 | GISAID |
| EPI579603 | A/Wisconsin/18/2015(H3N2) | 19/1/2015 | GISAID |
| EPI579611 | A/Wisconsin/19/2015(H3N2) | 22/1/2015 | GISAID |
| EPI579623 | A/Utah/10/2015(H3N2) | 19/1/2015 | GISAID |
| EPI579631 | A/Florida/21/2015(H3N2) | 21/1/2015 | GISAID |
| EPI579639 | A/Florida/24/2015(H3N2) | 20/1/2015 | GISAID |
| EPI579647 | A/Ohio/01/2015(H3N2) | 16/1/2015 | GISAID |
| EPI579655 | A/Ohio/02/2015(H3N2) | 17/1/2015 | GISAID |
| EPI579663 | A/Ohio/03/2015(H3N2) | 21/1/2015 | GISAID |
| EPI579671 | A/Colorado/05/2015(H3N2) | 7/1/2015 | GISAID |
| EPI579679 | A/Colorado/06/2015(H3N2) | 8/1/2015 | GISAID |
| EPI579687 | A/Colorado/07/2015(H3N2) | 14/1/2015 | GISAID |
| EPI579695 | A/Kentucky/02/2015(H3N2) | 6/1/2015 | GISAID |
| EPI579703 | A/Kentucky/03/2015(H3N2) | 16/1/2015 | GISAID |
| EPI579711 | A/Kentucky/04/2015(H3N2) | 22/1/2015 | GISAID |

| | | | |
|---|---|---|---|
| EPI579719 | A/NorthDakota/04/2015(H3N2) | 15/1/2015 | GISAID |
| EPI579726 | A/Wyoming/02/2015(H3N2) | 15/1/2015 | GISAID |
| EPI579734 | A/NorthCarolina/10/2015(H3N2) | 13/1/2015 | GISAID |
| EPI579742 | A/Indiana/03/2015(H3N2) | 13/1/2015 | GISAID |
| EPI579748 | A/Tennessee/11/2015(H3N2) | 3/2/2015 | GISAID |
| EPI579756 | A/Wisconsin/13/2015(H3N2) | 22/2/2015 | GISAID |
| EPI579764 | A/Washington/12/2015(H3N2) | 18/2/2015 | GISAID |
| EPI579779 | A/Washington/13/2015(H3N2) | 25/2/2015 | GISAID |
| EPI622100 | A/NewMexico/09/2015(H3N2) | 18/1/2015 | GISAID |
| EPI579796 | A/Wisconsin/20/2015(H3N2) | 24/2/2015 | GISAID |
| EPI579801 | A/NewHampshire/13/2015(H3N2) | 18/2/2015 | GISAID |
| EPI579809 | A/Nevada/18/2015(H3N2) | 18/2/2015 | GISAID |
| EPI579817 | A/California/19/2015(H3N2) | 27/1/2015 | GISAID |
| EPI579825 | A/NewYork/19/2015(H3N2) | 17/2/2015 | GISAID |
| EPI579833 | A/Oregon/04/2015(H3N2) | 14/2/2015 | GISAID |
| EPI579841 | A/Kansas/08/2015(H3N2) | 19/2/2015 | GISAID |
| EPI579846 | A/NewMexico/10/2015(H3N2) | 7/2/2015 | GISAID |
| EPI579854 | A/Delaware/18/2015(H3N2) | 23/2/2015 | GISAID |
| EPI579859 | A/Nevada/17/2015(H3N2) | 18/2/2015 | GISAID |
| EPI579866 | A/Minnesota/03/2015(H3N2) | 13/1/2015 | GISAID |
| EPI579874 | A/NewJersey/12/2015(H3N2) | 29/1/2015 | GISAID |
| EPI579879 | A/Hawaii/05/2015(H3N2) | 18/1/2015 | GISAID |
| EPI579884 | A/Pennsylvania/11/2015(H3N2) | 7/1/2015 | GISAID |
| EPI579889 | A/NewYork/20/2015(H3N2) | 25/2/2015 | GISAID |
| EPI579897 | A/California/20/2015(H3N2) | 20/1/2015 | GISAID |
| EPI622142 | A/Nevada/19/2015(H3N2) | 18/2/2015 | GISAID |
| EPI579913 | A/NewJersey/11/2015(H3N2) | 21/2/2015 | GISAID |
| EPI579921 | A/Florida/25/2015(H3N2) | 15/1/2015 | GISAID |
| EPI580481 | A/Lebanon/14L41/2014(H3N2) | 9/1/2014 | GISAID |
| EPI580489 | A/Lebanon/14L45/2014(H3N2) | 22/1/2014 | GISAID |
| EPI580497 | A/Lebanon/14L56/2014(H3N2) | 31/1/2014 | GISAID |
| EPI580505 | A/Lebanon/14L78/2014(H3N2) | 10/2/2014 | GISAID |
| EPI580521 | A/Niigata/14F067/2015(H3N2) | 19/1/2015 | GISAID |
| EPI580529 | A/Niigata/14F197/2015(H3N2) | 26/1/2015 | GISAID |
| EPI580545 | A/Gunma/14G011/2015(H3N2) | 14/1/2015 | GISAID |
| EPI580561 | A/Nagasaki/14N024/2015(H3N2) | 5/1/2015 | GISAID |
| EPI580569 | A/Nagasaki/14N039/2015(H3N2) | 14/1/2015 | GISAID |

| | | | |
|---|---|---|---|
| EPI580577 | A/Nagasaki/14N072/2015(H3N2) | 5/1/2015 | GISAID |
| EPI580585 | A/Okinawa/14T003/2015(H3N2) | 6/1/2015 | GISAID |
| EPI580593 | A/Okinawa/14T004/2015(H3N2) | 13/1/2015 | GISAID |
| EPI580601 | A/Okinawa/14T006/2015(H3N2) | 20/1/2015 | GISAID |
| EPI580609 | A/Okinawa/14T007/2015(H3N2) | 22/1/2015 | GISAID |
| EPI580617 | A/Nagasaki/14N012/2014(H3N2) | 21/11/2014 | GISAID |
| EPI580625 | A/Nagasaki/14N013/2014(H3N2) | 9/12/2014 | GISAID |
| EPI582005 | A/Stockholm/12/2015(H3N2) | 10/2/2015 | GISAID |
| EPI582013 | A/Stockholm/13/2015(H3N2) | 10/2/2015 | GISAID |
| EPI582021 | A/Stockholm/14/2015(H3N2) | 18/2/2015 | GISAID |
| EPI582029 | A/Stockholm/15/2015(H3N2) | 13/2/2015 | GISAID |
| EPI582037 | A/Stockholm/16/2015(H3N2) | 18/2/2015 | GISAID |
| EPI582045 | A/Stockholm/17/2015(H3N2) | 11/2/2015 | GISAID |
| EPI582053 | A/Stockholm/18/2015(H3N2) | 11/2/2015 | GISAID |
| EPI582088 | A/Sweden/14/2015(H3N2) | 22/2/2015 | GISAID |
| EPI582096 | A/Stockholm/19/2015(H3N2) | 25/2/2015 | GISAID |
| EPI582104 | A/Stockholm/20/2015(H3N2) | 24/2/2015 | GISAID |
| EPI582138 | A/Sweden/15/2015(H3N2) | 27/2/2015 | GISAID |
| EPI582146 | A/Sweden/16/2015(H3N2) | 5/3/2015 | GISAID |
| EPI582154 | A/Sweden/17/2015(H3N2) | 5/3/2015 | GISAID |
| EPI582218 | A/Sweden/19/2015(H3N2) | 5/3/2015 | GISAID |
| EPI582274 | A/Malmoe/2/2015(H3N2) | 23/2/2015 | GISAID |
| EPI583972 | A/SAPPORO/6/2015(H3N2) | 10/1/2015 | GISAID |
| EPI583978 | A/KANAGAWA/15/2015(H3N2) | 17/1/2015 | GISAID |
| EPI583986 | A/WAKAYAMA-C/104/2014(H3N2) | 6/12/2014 | GISAID |
| EPI583994 | A/HYOGO/3213/2015(H3N2) | 17/2/2015 | GISAID |
| EPI584000 | A/AOMORI/3/2015(H3N2) | 8/1/2015 | GISAID |
| EPI584559 | A/Hawaii/07/2015(H3N2) | 5/2/2015 | GISAID |
| EPI584567 | A/Virginia/26/2014(H3N2) | 25/11/2014 | GISAID |
| EPI584575 | A/Alabama/04/2015(H3N2) | 22/1/2015 | GISAID |
| EPI584583 | A/Maryland/50/2014(H3N2) | 17/12/2014 | GISAID |
| EPI584584 | A/Pennsylvania/60/2014(H3N2) | 22/12/2014 | GISAID |
| EPI584592 | A/Massachusetts/27/2014(H3N2) | 21/12/2014 | GISAID |
| EPI584605 | A/Florida/17/2015(H3N2) | 26/1/2015 | GISAID |
| EPI584613 | A/Tennessee/25/2014(H3N2) | 8/12/2014 | GISAID |
| EPI584626 | A/Laos/1115/2014(H3N2) | 20/10/2014 | GISAID |
| EPI584634 | A/Florida/89/2014(H3N2) | 18/12/2014 | GISAID |

| | | | |
|---|---|---|---|
| EPI584642 | A/Virginia/42/2014(H3N2) | 9/12/2014 | GISAID |
| EPI584650 | A/Uganda/583/2014(H3N2) | 17/4/2014 | GISAID |
| EPI584677 | A/Uganda/1590/2014(H3N2) | 26/5/2014 | GISAID |
| EPI584685 | A/Washington/18/2015(H3N2) | 21/1/2015 | GISAID |
| EPI584699 | A/Washington/61/2014(H3N2) | 9/12/2014 | GISAID |
| EPI584714 | A/Virginia/41/2014(H3N2) | 28/12/2014 | GISAID |
| EPI585299 | A/Karlstad/3/2015(H3N2) | 3/4/2015 | GISAID |
| EPI585339 | A/Sweden/20/2015(H3N2) | 8/4/2015 | GISAID |
| EPI586564 | A/SAPPORO/38/2015(H3N2) | 9/3/2015 | GISAID |
| EPI586568 | A/KOCHI/15/2015(H3N2) | 9/2/2015 | GISAID |
| EPI587444 | A/NewZealand/316/2014(H3N2) | 21/4/2014 | GISAID |
| EPI588494 | A/Nevada/21/2015(H3N2) | 2/3/2015 | GISAID |
| EPI588514 | A/Hawaii/08/2015(H3N2) | 5/3/2015 | GISAID |
| EPI588522 | A/Texas/38/2015(H3N2) | 27/2/2015 | GISAID |
| EPI588530 | A/Texas/36/2015(H3N2) | 8/3/2015 | GISAID |
| EPI588537 | A/Oregon/05/2015(H3N2) | 8/3/2015 | GISAID |
| EPI588545 | A/NewMexico/12/2015(H3N2) | 10/3/2015 | GISAID |
| EPI588553 | A/Idaho/12/2015(H3N2) | 13/3/2015 | GISAID |
| EPI588568 | A/Georgia/03/2015(H3N2) | 6/1/2015 | GISAID |
| EPI588576 | A/NorthDakota/05/2015(H3N2) | 3/2/2015 | GISAID |
| EPI588583 | A/Kentucky/08/2015(H3N2) | 24/2/2015 | GISAID |
| EPI588591 | A/Indiana/10/2015(H3N2) | 3/3/2015 | GISAID |
| EPI588605 | A/Kentucky/07/2015(H3N2) | 24/2/2015 | GISAID |
| EPI588620 | A/Montana/07/2015(H3N2) | 7/3/2015 | GISAID |
| EPI588628 | A/Texas/39/2015(H3N2) | 18/2/2015 | GISAID |
| EPI588636 | A/Wisconsin/23/2015(H3N2) | 26/2/2015 | GISAID |
| EPI588644 | A/Wisconsin/22/2015(H3N2) | 27/2/2015 | GISAID |
| EPI588657 | A/Massachusetts/09/2015(H3N2) | 25/2/2015 | GISAID |
| EPI588665 | A/Maine/17/2015(H3N2) | 2/3/2015 | GISAID |
| EPI588673 | A/Maine/18/2015(H3N2) | 3/3/2015 | GISAID |
| EPI588681 | A/Maryland/01/2015(H3N2) | 22/1/2015 | GISAID |
| EPI588689 | A/Maryland/02/2015(H3N2) | 24/1/2015 | GISAID |
| EPI588697 | A/Maryland/03/2015(H3N2) | 16/2/2015 | GISAID |
| EPI588712 | A/Texas/37/2015(H3N2) | 3/3/2015 | GISAID |
| EPI588720 | A/NewJersey/14/2015(H3N2) | 12/3/2015 | GISAID |
| EPI588728 | A/NewJersey/15/2015(H3N2) | 2/3/2015 | GISAID |
| EPI588736 | A/Wisconsin/21/2015(H3N2) | 9/3/2015 | GISAID |

| | | | |
|---|---|---|---|
| EPI588744 | A/NewHampshire/15/2015(H3N2) | 25/2/2015 | GISAID |
| EPI588752 | A/NewHampshire/14/2015(H3N2) | 27/2/2015 | GISAID |
| EPI588760 | A/NewHampshire/16/2015(H3N2) | 3/3/2015 | GISAID |
| EPI588768 | A/Maine/20/2015(H3N2) | 11/3/2015 | GISAID |
| EPI588776 | A/Wisconsin/24/2015(H3N2) | 9/3/2015 | GISAID |
| EPI588784 | A/Washington/21/2015(H3N2) | 6/3/2015 | GISAID |
| EPI588792 | A/Vermont/05/2015(H3N2) | 13/3/2015 | GISAID |
| EPI588800 | A/RhodeIsland/12/2015(H3N2) | 30/3/2015 | GISAID |
| EPI588808 | A/Delaware/22/2015(H3N2) | 8/3/2015 | GISAID |
| EPI588816 | A/NewJersey/13/2015(H3N2) | 21/3/2015 | GISAID |
| EPI588831 | A/Michigan/17/2015(H3N2) | 17/2/2015 | GISAID |
| EPI588839 | A/Florida/27/2015(H3N2) | 5/3/2015 | GISAID |
| EPI588847 | A/NorthCarolina/16/2015(H3N2) | 15/3/2015 | GISAID |
| EPI588855 | A/NorthCarolina/15/2015(H3N2) | 17/3/2015 | GISAID |
| EPI588863 | A/NorthDakota/06/2015(H3N2) | 8/3/2015 | GISAID |
| EPI588871 | A/Ohio/10/2015(H3N2) | 21/3/2015 | GISAID |
| EPI588879 | A/Ohio/11/2015(H3N2) | 20/3/2015 | GISAID |
| EPI588894 | A/Florida/26/2015(H3N2) | 17/3/2015 | GISAID |
| EPI588902 | A/Kentucky/06/2015(H3N2) | 22/3/2015 | GISAID |
| EPI588910 | A/Colombia/4468/2015(H3N2) | 17/2/2015 | GISAID |
| EPI588918 | A/Colombia/2473/2015(H3N2) | 13/2/2015 | GISAID |
| EPI588926 | A/Colombia/4352/2015(H3N2) | 16/1/2015 | GISAID |
| EPI588930 | A/NAGANO/2202/2015(H3N2) | 12/3/2015 | GISAID |
| EPI594983 | A/NewJersey/06/2015(H3N2) | 2/3/2015 | GISAID |
| EPI594992 | A/Massachusetts/15/2015(H3N2) | 19/2/2015 | GISAID |
| EPI595002 | A/Kansas/04/2015(H3N2) | 3/3/2015 | GISAID |
| EPI595010 | A/NorthCarolina/24/2015(H3N2) | 27/1/2015 | GISAID |
| EPI595018 | A/NorthCarolina/25/2015(H3N2) | 25/1/2015 | GISAID |
| EPI595034 | A/Tennessee/10/2015(H3N2) | 27/1/2015 | GISAID |
| EPI595043 | A/Wisconsin/16/2015(H3N2) | 4/3/2015 | GISAID |
| EPI595051 | A/Texas/19/2015(H3N2) | 2/3/2015 | GISAID |
| EPI602616 | A/Pennsylvania/09/2015(H3N2) | 7/1/2015 | GISAID |
| EPI607577 | A/England/513/2014(H3N2) | 12/10/2014 | GISAID |
| EPI607585 | A/England/516/2014(H3N2) | 24/10/2014 | GISAID |
| EPI607593 | A/England/599/2014(H3N2) | 16/12/2014 | GISAID |
| EPI607601 | A/England/591/2014(H3N2) | 23/12/2014 | GISAID |
| EPI607609 | A/England/50200164/2015(H3N2) | 2/1/2015 | GISAID |

| | | | |
|---|---|---|---|
| EPI607617 | A/England/50200187/2015(H3N2) | 2/1/2015 | GISAID |
| EPI607625 | A/England/50220068/2015(H3N2) | 2/1/2015 | GISAID |
| EPI607633 | A/England/50220079/2015(H3N2) | 2/1/2015 | GISAID |
| EPI607641 | A/England/50220082/2015(H3N2) | 2/1/2015 | GISAID |
| EPI607649 | A/England/50240248/2015(H3N2) | 5/1/2015 | GISAID |
| EPI607657 | A/England/10/2015(H3N2) | 6/1/2015 | GISAID |
| EPI607665 | A/England/11/2015(H3N2) | 5/1/2015 | GISAID |
| EPI607673 | A/England/12/2015(H3N2) | 5/1/2015 | GISAID |
| EPI607681 | A/England/49/2015(H3N2) | 6/1/2015 | GISAID |
| EPI607689 | A/England/13/2015(H3N2) | 5/1/2015 | GISAID |
| EPI607697 | A/England/50/2015(H3N2) | 6/1/2015 | GISAID |
| EPI607705 | A/England/50260150/2015(H3N2) | 5/1/2015 | GISAID |
| EPI607713 | A/England/50260151/2015(H3N2) | 7/1/2015 | GISAID |
| EPI607721 | A/England/18/2015(H3N2) | 7/1/2015 | GISAID |
| EPI607729 | A/England/50280041/2015(H3N2) | 7/1/2015 | GISAID |
| EPI607737 | A/England/50280054/2015(H3N2) | 7/1/2015 | GISAID |
| EPI607745 | A/England/50280058/2015(H3N2) | 8/1/2015 | GISAID |
| EPI607753 | A/England/50300008/2015(H3N2) | 7/1/2015 | GISAID |
| EPI607761 | A/England/50300025/2015(H3N2) | 8/1/2015 | GISAID |
| EPI607769 | A/England/50300033/2015(H3N2) | 9/1/2015 | GISAID |
| EPI607777 | A/England/50300044/2015(H3N2) | 7/1/2015 | GISAID |
| EPI607785 | A/England/50320276/2015(H3N2) | 12/1/2015 | GISAID |
| EPI607793 | A/England/32/2015(H3N2) | 9/1/2015 | GISAID |
| EPI607801 | A/England/57/2015(H3N2) | 12/1/2015 | GISAID |
| EPI607809 | A/England/50360024/2015(H3N2) | 13/1/2015 | GISAID |
| EPI607817 | A/England/50360027/2015(H3N2) | 14/1/2015 | GISAID |
| EPI607825 | A/England/60/2015(H3N2) | 13/1/2015 | GISAID |
| EPI607833 | A/England/50360038/2015(H3N2) | 14/1/2015 | GISAID |
| EPI607841 | A/England/50380052/2015(H3N2) | 12/1/2015 | GISAID |
| EPI607849 | A/England/72/2015(H3N2) | 15/1/2015 | GISAID |
| EPI607857 | A/England/50400278/2015(H3N2) | 16/1/2015 | GISAID |
| EPI607865 | A/England/50400282/2015(H3N2) | 15/1/2015 | GISAID |
| EPI607873 | A/England/50400284/2015(H3N2) | 15/1/2015 | GISAID |
| EPI607881 | A/England/87/2015(H3N2) | 22/1/2015 | GISAID |
| EPI607889 | A/England/50520248/2015(H3N2) | 26/1/2015 | GISAID |
| EPI607901 | A/England/105/2015(H3N2) | 26/1/2015 | GISAID |
| EPI607899 | A/England/105/2015(H3N2) | 26/1/2015 | GISAID |

| | | | |
|---|---|---|---|
| EPI607914 | A/England/127/2015(H3N2) | 29/1/2015 | GISAID |
| EPI607915 | A/England/127/2015(H3N2) | 29/1/2015 | GISAID |
| EPI607930 | A/England/112/2015(H3N2) | 26/1/2015 | GISAID |
| EPI607932 | A/England/112/2015(H3N2) | 26/1/2015 | GISAID |
| EPI607945 | A/England/113/2015(H3N2) | 28/1/2015 | GISAID |
| EPI607947 | A/England/113/2015(H3N2) | 28/1/2015 | GISAID |
| EPI607960 | A/England/114/2015(H3N2) | 30/1/2015 | GISAID |
| EPI607962 | A/England/114/2015(H3N2) | 30/1/2015 | GISAID |
| EPI607975 | A/England/50660076/2015(H3N2) | 3/2/2015 | GISAID |
| EPI607977 | A/England/50660076/2015(H3N2) | 3/2/2015 | GISAID |
| EPI607990 | A/England/50740149/2015(H3N2) | 10/2/2015 | GISAID |
| EPI607991 | A/England/50740149/2015(H3N2) | 10/2/2015 | GISAID |
| EPI608006 | A/England/50740161/2015(H3N2) | 10/2/2015 | GISAID |
| EPI608007 | A/England/50740161/2015(H3N2) | 10/2/2015 | GISAID |
| EPI608022 | A/England/50780065/2015(H3N2) | 11/2/2015 | GISAID |
| EPI608023 | A/England/50780065/2015(H3N2) | 11/2/2015 | GISAID |
| EPI608038 | A/England/166/2015(H3N2) | 12/2/2015 | GISAID |
| EPI608041 | A/England/166/2015(H3N2) | 12/2/2015 | GISAID |
| EPI608052 | A/England/50800025/2015(H3N2) | 12/2/2015 | GISAID |
| EPI608051 | A/England/50800025/2015(H3N2) | 12/2/2015 | GISAID |
| EPI608067 | A/England/169/2015(H3N2) | 13/2/2015 | GISAID |
| EPI608068 | A/England/169/2015(H3N2) | 13/2/2015 | GISAID |
| EPI610547 | A/YAMAGATA/64/2015(H3N2) | 3/3/2015 | GISAID |
| EPI610549 | A/KANAGAWA/36/2015(H3N2) | 2/2/2015 | GISAID |
| EPI610551 | A/MIYAGI/12/2015(H3N2) | 9/1/2015 | GISAID |
| EPI610553 | A/MIYAGI/21/2015(H3N2) | 1/2/2015 | GISAID |
| EPI611225 | A/England/43960141/2014(H3N2) | 22/9/2014 | GISAID |
| EPI611249 | A/England/44020584/2014(H3N2) | 26/9/2014 | GISAID |
| EPI611263 | A/England/44080181/2014(H3N2) | 12/9/2014 | GISAID |
| EPI611271 | A/Scotland/2/2014(H3N2) | 27/8/2014 | GISAID |
| EPI611279 | A/England/44160198/2014(H3N2) | 2/10/2014 | GISAID |
| EPI611287 | A/England/44160199/2014(H3N2) | 2/10/2014 | GISAID |
| EPI611295 | A/England/511/2014(H3N2) | 2/10/2014 | GISAID |
| EPI611303 | A/England/44160201/2014(H3N2) | 2/10/2014 | GISAID |
| EPI611311 | A/England/44160202/2014(H3N2) | 2/10/2014 | GISAID |
| EPI611319 | A/England/512/2014(H3N2) | 2/10/2014 | GISAID |
| EPI611327 | A/England/44320792/2014(H3N2) | 10/10/2014 | GISAID |

| | | | |
|---|---|---|---|
| EPI611335 | A/England/44320793/2014(H3N2) | 16/10/2014 | GISAID |
| EPI611343 | A/England/44320794/2014(H3N2) | 15/10/2014 | GISAID |
| EPI611351 | A/England/44320795/2014(H3N2) | 18/10/2014 | GISAID |
| EPI611359 | A/England/44540273/2014(H3N2) | 28/10/2014 | GISAID |
| EPI611367 | A/England/44640016/2014(H3N2) | 9/11/2014 | GISAID |
| EPI611375 | A/England/525/2014(H3N2) | 20/11/2014 | GISAID |
| EPI611383 | A/England/44820080/2014(H3N2) | 20/11/2014 | GISAID |
| EPI611391 | A/England/44840003/2014(H3N2) | 24/11/2014 | GISAID |
| EPI611399 | A/England/528/2014(H3N2) | 20/11/2014 | GISAID |
| EPI611407 | A/England/44840067/2014(H3N2) | 21/11/2014 | GISAID |
| EPI611415 | A/England/529/2014(H3N2) | 21/11/2014 | GISAID |
| EPI611423 | A/England/44840069/2014(H3N2) | 21/11/2014 | GISAID |
| EPI611431 | A/England/44860850/2014(H3N2) | 21/11/2014 | GISAID |
| EPI611440 | A/England/44920021/2014(H3N2) | 28/11/2014 | GISAID |
| EPI611448 | A/England/530/2014(H3N2) | 26/11/2014 | GISAID |
| EPI611456 | A/England/44940038/2014(H3N2) | 2/12/2014 | GISAID |
| EPI611464 | A/England/45000006/2014(H3N2) | 4/12/2014 | GISAID |
| EPI611472 | A/England/45000008/2014(H3N2) | 5/12/2014 | GISAID |
| EPI611480 | A/England/45000010/2014(H3N2) | 3/12/2014 | GISAID |
| EPI611488 | A/England/45000025/2014(H3N2) | 1/12/2014 | GISAID |
| EPI611496 | A/England/533/2014(H3N2) | 1/12/2014 | GISAID |
| EPI611504 | A/England/45000028/2014(H3N2) | 27/11/2014 | GISAID |
| EPI611512 | A/England/45040065/2014(H3N2) | 9/12/2014 | GISAID |
| EPI611536 | A/England/536/2014(H3N2) | 8/12/2014 | GISAID |
| EPI611544 | A/England/45080158/2014(H3N2) | 9/12/2014 | GISAID |
| EPI611552 | A/England/45080193/2014(H3N2) | 10/12/2014 | GISAID |
| EPI611560 | A/England/45080198/2014(H3N2) | 10/12/2014 | GISAID |
| EPI611568 | A/England/45080199/2014(H3N2) | 11/12/2014 | GISAID |
| EPI611576 | A/England/45080202/2014(H3N2) | 8/12/2014 | GISAID |
| EPI611584 | A/England/45080206/2014(H3N2) | 8/12/2014 | GISAID |
| EPI611592 | A/England/45100189/2014(H3N2) | 11/12/2014 | GISAID |
| EPI611600 | A/England/45100191/2014(H3N2) | 10/12/2014 | GISAID |
| EPI611610 | A/England/45100197/2014(H3N2) | 26/11/2014 | GISAID |
| EPI611618 | A/England/45140001/2014(H3N2) | 16/12/2014 | GISAID |
| EPI611626 | A/England/45140005/2014(H3N2) | 15/12/2014 | GISAID |
| EPI611634 | A/England/45140006/2014(H3N2) | 12/12/2014 | GISAID |
| EPI611642 | A/England/45160088/2014(H3N2) | 18/12/2014 | GISAID |

| | | | |
|---|---|---|---|
| EPI611650 | A/England/45160092/2014(H3N2) | 17/12/2014 | GISAID |
| EPI611658 | A/England/45160094/2014(H3N2) | 16/12/2014 | GISAID |
| EPI611666 | A/England/601/2014(H3N2) | 15/12/2014 | GISAID |
| EPI611674 | A/England/45180212/2014(H3N2) | 16/12/2014 | GISAID |
| EPI611682 | A/England/543/2014(H3N2) | 17/12/2014 | GISAID |
| EPI611690 | A/England/45180219/2014(H3N2) | 17/12/2014 | GISAID |
| EPI611698 | A/England/45180220/2014(H3N2) | 17/12/2014 | GISAID |
| EPI611706 | A/England/544/2014(H3N2) | 18/12/2014 | GISAID |
| EPI611714 | A/England/545/2014(H3N2) | 16/12/2014 | GISAID |
| EPI611722 | A/England/45180226/2014(H3N2) | 17/12/2014 | GISAID |
| EPI611730 | A/England/45180235/2014(H3N2) | 19/12/2014 | GISAID |
| EPI611738 | A/England/602/2014(H3N2) | 17/12/2014 | GISAID |
| EPI611746 | A/England/45200044/2014(H3N2) | 18/12/2014 | GISAID |
| EPI611754 | A/England/547/2014(H3N2) | 18/12/2014 | GISAID |
| EPI611762 | A/England/45200047/2014(H3N2) | 18/12/2014 | GISAID |
| EPI611770 | A/England/45200054/2014(H3N2) | 17/12/2014 | GISAID |
| EPI611778 | A/England/575/2014(H3N2) | 22/12/2014 | GISAID |
| EPI611786 | A/England/45220034/2014(H3N2) | 19/12/2014 | GISAID |
| EPI611794 | A/England/556/2014(H3N2) | 18/12/2014 | GISAID |
| EPI611802 | A/England/45240246/2014(H3N2) | 22/12/2014 | GISAID |
| EPI611810 | A/England/558/2014(H3N2) | 23/12/2014 | GISAID |
| EPI611818 | A/England/45240252/2014(H3N2) | 18/12/2014 | GISAID |
| EPI611826 | A/England/45240260/2014(H3N2) | 22/12/2014 | GISAID |
| EPI611834 | A/England/45240273/2014(H3N2) | 22/12/2014 | GISAID |
| EPI611842 | A/England/603/2014(H3N2) | 19/12/2014 | GISAID |
| EPI611850 | A/England/50100002/2014(H3N2) | 23/12/2014 | GISAID |
| EPI611858 | A/England/50100063/2014(H3N2) | 24/12/2014 | GISAID |
| EPI611866 | A/England/50100065/2014(H3N2) | 24/12/2014 | GISAID |
| EPI611874 | A/England/563/2014(H3N2) | 24/12/2014 | GISAID |
| EPI611882 | A/England/50120064/2014(H3N2) | 29/12/2014 | GISAID |
| EPI611890 | A/England/564/2014(H3N2) | 23/12/2014 | GISAID |
| EPI611898 | A/England/606/2014(H3N2) | 29/12/2014 | GISAID |
| EPI611906 | A/England/50120073/2014(H3N2) | 24/12/2014 | GISAID |
| EPI611914 | A/England/50120074/2014(H3N2) | 23/12/2014 | GISAID |
| EPI611922 | A/England/50120128/2014(H3N2) | 30/12/2014 | GISAID |
| EPI611930 | A/England/50120129/2014(H3N2) | 23/12/2014 | GISAID |
| EPI611938 | A/England/50140078/2014(H3N2) | 30/12/2014 | GISAID |

| | | | |
|---|---|---|---|
| EPI611946 | A/England/50140086/2014(H3N2) | 29/12/2014 | GISAID |
| EPI611954 | A/England/50140112/2014(H3N2) | 29/12/2014 | GISAID |
| EPI611962 | A/England/50180052/2014(H3N2) | 29/12/2014 | GISAID |
| EPI611970 | A/England/50200102/2014(H3N2) | 21/12/2014 | GISAID |
| EPI611978 | A/England/50200115/2014(H3N2) | 31/12/2014 | GISAID |
| EPI611986 | A/England/50200119/2014(H3N2) | 30/12/2014 | GISAID |
| EPI611994 | A/England/50200132/2014(H3N2) | 29/12/2014 | GISAID |
| EPI612002 | A/England/50200133/2014(H3N2) | 29/12/2014 | GISAID |
| EPI612010 | A/England/50200134/2014(H3N2) | 30/12/2014 | GISAID |
| EPI612018 | A/England/50200158/2014(H3N2) | 31/12/2014 | GISAID |
| EPI612026 | A/England/50200190/2014(H3N2) | 31/12/2014 | GISAID |
| EPI612034 | A/England/50220066/2014(H3N2) | 2/1/2014 | GISAID |
| EPI612042 | A/England/50220076/2015(H3N2) | 6/1/2015 | GISAID |
| EPI612050 | A/England/50220894/2015(H3N2) | 6/1/2015 | GISAID |
| EPI612058 | A/England/50220895/(H3N2) | 6/1/2015 | GISAID |
| EPI612066 | A/England/50320294/2015(H3N2) | 13/1/2015 | GISAID |
| EPI612074 | A/England/50360756/2015(H3N2) | 6/1/2015 | GISAID |
| EPI612082 | A/England/50380175/2015(H3N2) | 6/1/2015 | GISAID |
| EPI612090 | A/England/50400281/2015(H3N2) | 15/1/2015 | GISAID |
| EPI612098 | A/England/77/2015(H3N2) | 12/1/2015 | GISAID |
| EPI612106 | A/England/50620849/2015(H3N2) | 30/1/2015 | GISAID |
| EPI612114 | A/England/50660120/2015(H3N2) | 3/2/2015 | GISAID |
| EPI612124 | A/England/50880445/2015(H3N2) | 7/2/2015 | GISAID |
| EPI612132 | A/England/50880448/2015(H3N2) | 9/2/2015 | GISAID |
| EPI612140 | A/England/50960080/2015(H3N2) | 11/2/2015 | GISAID |
| EPI612148 | A/England/50960083/2015(H3N2) | 27/1/2015 | GISAID |
| EPI612156 | A/England/51060001/2015(H3N2) | 3/3/2015 | GISAID |
| EPI612164 | A/England/51200124/2015(H3N2) | 7/3/2015 | GISAID |
| EPI612172 | A/England/51440624/2015(H3N2) | 1/4/2015 | GISAID |
| EPI612180 | A/England/51440626/2015(H3N2) | 1/4/2015 | GISAID |
| EPI612188 | A/England/51440629/2015(H3N2) | 1/4/2015 | GISAID |
| EPI612196 | A/Arizona/01/2015(H3N2) | 30/1/2015 | GISAID |
| EPI612204 | A/Wisconsin/27/2015(H3N2) | 26/1/2015 | GISAID |
| EPI612222 | A/Georgia/29/2014(H3N2) | 6/12/2014 | GISAID |
| EPI612230 | A/Georgia/07/2015(H3N2) | 2/1/2015 | GISAID |
| EPI612238 | A/Louisiana/39/2014(H3N2) | 10/12/2014 | GISAID |
| EPI612248 | A/Texas/94/2014(H3N2) | 23/12/2014 | GISAID |

| | | | |
|---|---|---|---|
| EPI612256 | A/Tanzania/587/2014(H3N2) | 1/12/2014 | GISAID |
| EPI612270 | A/Michigan/21/2015(H3N2) | 19/1/2015 | GISAID |
| EPI612279 | A/Michigan/22/2015(H3N2) | 16/1/2015 | GISAID |
| EPI612287 | A/Vietnam/629/2014(H3N2) | 15/12/2014 | GISAID |
| EPI612295 | A/Idaho/11/2015(H3N2) | 2/1/2015 | GISAID |
| EPI612315 | A/Washington/29/2015(H3N2) | 28/1/2015 | GISAID |
| EPI612323 | A/Maryland/53/2014(H3N2) | 17/12/2014 | GISAID |
| EPI612331 | A/Alabama/02/2015(H3N2) | 26/1/2015 | GISAID |
| EPI612343 | A/Washington/30/2015(H3N2) | 30/1/2015 | GISAID |
| EPI612358 | A/Georgia/28/2014(H3N2) | 3/12/2014 | GISAID |
| EPI612373 | A/Indiana/06/2015(H3N2) | 21/1/2015 | GISAID |
| EPI612381 | A/Georgia/38/2014(H3N2) | 17/12/2014 | GISAID |
| EPI612389 | A/NorthCarolina/23/2015(H3N2) | 9/2/2015 | GISAID |
| EPI612397 | A/Washington/60/2014(H3N2) | 23/12/2014 | GISAID |
| EPI612406 | A/Vermont/22/2014(H3N2) | 20/12/2014 | GISAID |
| EPI612414 | A/Georgia/45/2014(H3N2) | 15/12/2014 | GISAID |
| EPI612429 | A/Pennsylvania/10/2015(H3N2) | 8/1/2015 | GISAID |
| EPI612437 | A/Pennsylvania/07/2015(H3N2) | 9/1/2015 | GISAID |
| EPI612445 | A/Texas/43/2015(H3N2) | 28/3/2015 | GISAID |
| EPI612454 | A/Texas/95/2014(H3N2) | 26/12/2014 | GISAID |
| EPI612462 | A/Georgia/27/2014(H3N2) | 2014 | GISAID |
| EPI614429 | A/HongKong/7127/2014(H3N2) | 2014 | GISAID |
| EPI614437 | A/HongKong/4801/2014(H3N2) | 2014 | GISAID |
| EPI614444 | A/Switzerland/9715293/2013(H3N2) | 6/12/2013 | GISAID |
| EPI614462 | A/SouthAustralia/55/2014(H3N2) | 29/6/2014 | GISAID |
| EPI614775 | A/Kazakhstan/159/2015(H3N2) | 9/1/2015 | GISAID |
| EPI614783 | A/Congo/2495/2014(H3N2) | 11/12/2014 | GISAID |
| EPI614791 | A/Kazakhstan/184/2015(H3N2) | 26/1/2015 | GISAID |
| EPI614799 | A/Kazakhstan/266/2015(H3N2) | 13/1/2015 | GISAID |
| EPI614807 | A/Kazakhstan/166/2015(H3N2) | 29/1/2015 | GISAID |
| EPI614815 | A/Kazakhstan/291/2015(H3N2) | 6/2/2015 | GISAID |
| EPI614823 | A/Kazakhstan/161/2015(H3N2) | 14/1/2015 | GISAID |
| EPI614831 | A/Kazakhstan/158/2015(H3N2) | 23/1/2015 | GISAID |
| EPI614839 | A/Kazakhstan/274/2015(H3N2) | 21/1/2015 | GISAID |
| EPI622002 | A/Washington/31/2015(H3N2) | 15/3/2015 | GISAID |
| EPI622010 | A/Kazakhstan/194/2014(H3N2) | 29/11/2014 | GISAID |
| EPI622023 | A/SouthCarolina/07/2015(H3N2) | 5/2/2015 | GISAID |

| | | | |
|---|---|---|---|
| EPI622031 | A/California/47/2015(H3N2) | 19/2/2015 | GISAID |
| EPI622039 | A/California/24/2015(H3N2) | 3/3/2015 | GISAID |
| EPI622058 | A/Paraguay/0097/2014(H3N2) | 29/8/2014 | GISAID |
| EPI622073 | A/Iowa/11/2015(H3N2) | 5/3/2015 | GISAID |
| EPI622109 | A/Oregon/06/2015(H3N2) | 4/3/2015 | GISAID |
| EPI622119 | A/NewJersey/16/2015(H3N2) | 3/3/2015 | GISAID |
| EPI622155 | A/RhodeIsland/15/2015(H3N2) | 12/3/2015 | GISAID |
| EPI622163 | A/Maine/16/2015(H3N2) | 2/3/2015 | GISAID |
| EPI622180 | A/Chita/14/2014(H3N2) | 22/12/2014 | GISAID |
| EPI622188 | A/Georgia/05/2015(H3N2) | 2/1/2015 | GISAID |
| EPI622196 | A/Chita/12/2014(H3N2) | 25/12/2014 | GISAID |
| EPI622204 | A/Georgia/06/2015(H3N2) | 2/1/2015 | GISAID |
| EPI622218 | A/Khabarovsk/06/2015(H3N2) | 20/1/2015 | GISAID |
| EPI622226 | A/Florida/31/2015(H3N2) | 5/2/2015 | GISAID |
| EPI622234 | A/Khabarovsk/05/2015(H3N2) | 16/1/2015 | GISAID |
| EPI622242 | A/Khabarovsk/07/2015(H3N2) | 20/1/2015 | GISAID |
| EPI622250 | A/Vermont/06/2015(H3N2) | 8/3/2015 | GISAID |
| EPI622272 | A/Laos/732/2014(H3N2) | 21/11/2014 | GISAID |
| EPI622322 | A/Delaware/25/2014(H3N2) | 20/11/2014 | GISAID |
| EPI622335 | A/Florida/41/2015(H3N2) | 2015 | GISAID |
| EPI622353 | A/NewHampshire/09/2015(H3N2) | 11/1/2015 | GISAID |
| EPI622361 | A/Texas/56/2015(H3N2) | 31/1/2015 | GISAID |
| EPI622424 | A/RhodeIsland/13/2015(H3N2) | 2/3/2015 | GISAID |
| EPI623426 | A/Wisconsin/64/2014(H3N2) | 22/11/2014 | GISAID |
| EPI623465 | A/Washington/20/2014(H3N2) | 10/10/2014 | GISAID |
| EPI624003 | A/RhodeIsland/24/2015(H3N2) | 3/4/2015 | GISAID |
| EPI624011 | A/NewJersey/49/2015(H3N2) | 14/4/2015 | GISAID |
| EPI624019 | A/Wisconsin/36/2015(H3N2) | 4/4/2015 | GISAID |
| EPI624027 | A/SouthDakota/15/2015(H3N2) | 5/3/2015 | GISAID |
| EPI624035 | A/Congo/2588/2014(H3N2) | 22/12/2014 | GISAID |
| EPI624043 | A/SouthDakota/05/2015(H3N2) | 3/2/2015 | GISAID |
| EPI624059 | A/SouthDakota/34/2015(H3N2) | 11/3/2015 | GISAID |
| EPI624067 | A/SouthDakota/14/2015(H3N2) | 27/2/2015 | GISAID |
| EPI624075 | A/SouthDakota/30/2015(H3N2) | 13/2/2015 | GISAID |
| EPI624083 | A/Florida/49/2015(H3N2) | 26/4/2015 | GISAID |
| EPI624102 | A/SouthDakota/29/2015(H3N2) | 24/2/2015 | GISAID |
| EPI624110 | A/SouthDakota/25/2015(H3N2) | 29/1/2015 | GISAID |

| | | | |
|---|---|---|---|
| EPI624118 | A/SouthDakota/35/2015(H3N2) | 14/3/2015 | GISAID |
| EPI624127 | A/Washington/37/2015(H3N2) | 6/5/2015 | GISAID |
| EPI624135 | A/Idaho/17/2015(H3N2) | 14/3/2015 | GISAID |
| EPI624143 | A/NewMexico/18/2015(H3N2) | 18/3/2015 | GISAID |
| EPI624158 | A/Texas/61/2015(H3N2) | 8/3/2015 | GISAID |
| EPI624166 | A/Indiana/14/2015(H3N2) | 8/4/2015 | GISAID |
| EPI624174 | A/Alaska/09/2015(H3N2) | 18/4/2015 | GISAID |
| EPI624192 | A/Idaho/23/2015(H3N2) | 14/4/2015 | GISAID |
| EPI624200 | A/Congo/2526/2014(H3N2) | 15/12/2014 | GISAID |
| EPI624209 | A/Congo/2603/2014(H3N2) | 29/12/2014 | GISAID |
| EPI624217 | A/Nevada/36/2015(H3N2) | 6/4/2015 | GISAID |
| EPI624225 | A/Arizona/07/2015(H3N2) | 14/4/2015 | GISAID |
| EPI624233 | A/Hawaii/37/2015(H3N2) | 14/4/2015 | GISAID |
| EPI624241 | A/California/82/2015(H3N2) | 28/4/2015 | GISAID |
| EPI624249 | A/Massachusetts/19/2015(H3N2) | 13/4/2015 | GISAID |
| EPI624257 | A/Alaska/10/2015(H3N2) | 12/4/2015 | GISAID |
| EPI624265 | A/Minnesota/25/2015(H3N2) | 1/4/2015 | GISAID |
| EPI624273 | A/NewMexico/19/2015(H3N2) | 1/4/2015 | GISAID |
| EPI624281 | A/Florida/45/2015(H3N2) | 18/4/2015 | GISAID |
| EPI624296 | A/Minnesota/30/2015(H3N2) | 18/5/2015 | GISAID |
| EPI624304 | A/Hawaii/33/2015(H3N2) | 19/4/2015 | GISAID |
| EPI624312 | A/Florida/50/2015(H3N2) | 3/6/2015 | GISAID |
| EPI624320 | A/Congo/2586/2014(H3N2) | 22/12/2014 | GISAID |
| EPI624328 | A/Wisconsin/66/2015(H3N2) | 21/4/2015 | GISAID |
| EPI624336 | A/Minnesota/28/2015(H3N2) | 21/4/2015 | GISAID |
| EPI624344 | A/SouthDakota/11/2015(H3N2) | 31/3/2015 | GISAID |
| EPI624358 | A/Minnesota/27/2015(H3N2) | 12/4/2015 | GISAID |
| EPI624366 | A/SouthDakota/01/2015(H3N2) | 5/1/2015 | GISAID |
| EPI624375 | A/Minnesota/29/2015(H3N2) | 16/5/2015 | GISAID |
| EPI624949 | A/Kazakhstan/1075/2015(H3N2) | 18/2/2015 | GISAID |
| EPI624957 | A/Kazakhstan/1060/2015(H3N2) | 16/2/2015 | GISAID |
| EPI624965 | A/Kazakhstan/1046/2015(H3N2) | 9/2/2015 | GISAID |
| EPI624980 | A/Kazakhstan/123/2015(H3N2) | 16/1/2015 | GISAID |
| EPI624988 | A/Kazakhstan/183/2015(H3N2) | 27/1/2015 | GISAID |
| EPI624996 | A/Ternopil/407/2015(H3N2) | 7/4/2015 | GISAID |
| EPI625004 | A/Dnipropetrovsk/442/2015(H3N2) | 15/4/2015 | GISAID |
| EPI625012 | A/Khmelnitsky/459/2015(H3N2) | 14/4/2015 | GISAID |

| | | | |
|---|---|---|---|
| EPI625020 | A/Kazakhstan/1099/2015(H3N2) | 23/2/2015 | GISAID |
| EPI625028 | A/Khmelnitsky/464/2015(H3N2) | 14/4/2015 | GISAID |
| EPI626308 | A/Nigeria/942/2014(H3N2) | 22/4/2014 | GISAID |
| EPI626316 | A/Nigeria/120/2014(H3N2) | 10/6/2014 | GISAID |
| EPI626324 | A/Massachusetts/16/2015(H3N2) | 5/3/2015 | GISAID |
| EPI626344 | A/Kazakhstan/873/2015(H3N2) | 12/1/2015 | GISAID |
| EPI626352 | A/Kazakhstan/286/2015(H3N2) | 2/2/2015 | GISAID |
| EPI626360 | A/Kazakhstan/112/2015(H3N2) | 7/1/2015 | GISAID |
| EPI626375 | A/StPetersburg/03/2015(H3N2) | 17/1/2015 | GISAID |
| EPI626383 | A/StPetersburg/22/2015(H3N2) | 26/1/2015 | GISAID |
| EPI626391 | A/Kazakhstan/341/2015(H3N2) | 9/2/2015 | GISAID |
| EPI626406 | A/SouthDakota/20/2015(H3N2) | 6/1/2015 | GISAID |
| EPI626414 | A/Kazakhstan/122/2015(H3N2) | 13/1/2015 | GISAID |
| EPI626422 | A/Ukraine/151/2015(H3N2) | 27/2/2015 | GISAID |
| EPI626430 | A/Kazakhstan/113/2015(H3N2) | 13/1/2015 | GISAID |
| EPI626445 | A/Kazakhstan/352/2015(H3N2) | 14/2/2015 | GISAID |
| EPI626453 | A/Washington/35/2015(H3N2) | 18/3/2015 | GISAID |
| EPI626461 | A/Washington/34/2015(H3N2) | 16/3/2015 | GISAID |
| EPI626480 | A/Kharkov/241/2015(H3N2) | 25/2/2015 | GISAID |
| EPI626488 | A/California/78/2015(H3N2) | 3/3/2015 | GISAID |
| EPI626496 | A/California/81/2015(H3N2) | 7/3/2015 | GISAID |
| EPI626504 | A/Minnesota/05/2015(H3N2) | 11/1/2015 | GISAID |
| EPI626512 | A/Delaware/35/2015(H3N2) | 2/3/2015 | GISAID |
| EPI626520 | A/Hawaii/27/2015(H3N2) | 28/3/2015 | GISAID |
| EPI626533 | A/Washington/33/2015(H3N2) | 12/3/2015 | GISAID |
| EPI626541 | A/Idaho/17/2015(H3N2) | 14/3/2015 | GISAID |
| EPI626549 | A/Nigeria/71/2014(H3N2) | 2/6/2014 | GISAID |
| EPI626557 | A/Nigeria/928/2014(H3N2) | 14/5/2014 | GISAID |
| EPI626565 | A/Maryland/12/2015(H3N2) | 2/3/2015 | GISAID |
| EPI626573 | A/ENG/507/2014(H3N2) | 24/8/2014 | GISAID |
| EPI626581 | A/ENG/44060047/2014(H3N2) | 24/9/2014 | GISAID |
| EPI626589 | A/ENG/44760132/2014(H3N2) | 20/11/2014 | GISAID |
| EPI626597 | A/ENG/44780079/2014(H3N2) | 20/11/2014 | GISAID |
| EPI626638 | A/ENG/542/2014(H3N2) | 12/12/2014 | GISAID |
| EPI626651 | A/ENG/45160093/2014(H3N2) | 17/12/2014 | GISAID |
| EPI626694 | A/ENG/50100055/2014(H3N2) | 23/12/2014 | GISAID |
| EPI626702 | A/ENG/50100064/2014(H3N2) | 24/12/2014 | GISAID |

| | | | |
|---|---|---|---|
| EPI626807 | A/ENG/50200185/2014(H3N2) | 30/12/2014 | GISAID |
| EPI626860 | A/ENG/50360752/2015(H3N2) | 5/1/2015 | GISAID |
| EPI626873 | A/ENG/50400311/2015(H3N2) | 15/1/2015 | GISAID |
| EPI626881 | A/ENG/75/2015(H3N2) | 15/1/2015 | GISAID |
| EPI626889 | A/ENG/50400317/2015(H3N2) | 15/1/2015 | GISAID |
| EPI626930 | A/ENG/50800026/2015(H3N2) | 12/2/2015 | GISAID |
| EPI626943 | A/ENG/51020042/2015(H3N2) | 3/3/2015 | GISAID |
| EPI626967 | A/ENG/52840541/2015(H3N2) | 3/7/2015 | GISAID |
| EPI629536 | A/StPetersburg/101/2015(H3N2) | 16/2/2015 | GISAID |
| EPI629544 | A/Khabarovsk/19/2015(H3N2) | 4/2/2015 | GISAID |
| EPI629552 | A/Florida/51/2015(H3N2) | 9/6/2015 | GISAID |
| EPI629560 | A/Arizona/09/2015(H3N2) | 21/6/2015 | GISAID |
| EPI629582 | A/Peru/27/2015(H3N2) | 13/4/2015 | GISAID |
| EPI629590 | A/Peru/30/2015(H3N2) | 23/4/2015 | GISAID |
| EPI629605 | A/StPetersburg/103/2015(H3N2) | 20/2/2015 | GISAID |
| EPI629613 | A/Hawaii/12/2015(H3N2) | 10/2/2015 | GISAID |
| EPI629628 | A/Hawaii/15/2015(H3N2) | 15/2/2015 | GISAID |
| EPI629636 | A/Peru/28/2015(H3N2) | 9/4/2015 | GISAID |
| EPI629644 | A/Hawaii/23/2015(H3N2) | 25/2/2015 | GISAID |
| EPI629659 | A/Hawaii/24/2015(H3N2) | 4/3/2015 | GISAID |
| EPI629674 | A/Laos/122/2015(H3N2) | 2/2/2015 | GISAID |
| EPI629689 | A/Kazakhstan/1080/2015(H3N2) | 19/2/2015 | GISAID |
| EPI629697 | A/Alaska/18/2015(H3N2) | 1/6/2015 | GISAID |
| EPI629705 | A/Alaska/13/2015(H3N2) | 5/6/2015 | GISAID |
| EPI629713 | A/Alaska/21/2015(H3N2) | 8/6/2015 | GISAID |
| EPI629721 | A/Alaska/25/2015(H3N2) | 17/6/2015 | GISAID |
| EPI629729 | A/Hawaii/14/2015(H3N2) | 17/2/2015 | GISAID |
| EPI631490 | A/Minnesota/26/2015(H3N2) | 2/4/2015 | GISAID |
| EPI631498 | A/Arkansas/06/2015(H3N2) | 17/2/2015 | GISAID |
| EPI631513 | A/Florida/46/2015(H3N2) | 18/4/2015 | GISAID |
| EPI631521 | A/Indiana/13/2015(H3N2) | 17/2/2015 | GISAID |
| EPI631536 | A/Santiago/33417/2015(H3N2) | 18/5/2015 | GISAID |
| EPI631544 | A/Brazil/9056/2015(H3N2) | 15/5/2015 | GISAID |
| EPI631571 | A/Santiago/39170/2015(H3N2) | 9/6/2015 | GISAID |
| EPI631579 | A/NewJersey/48/2015(H3N2) | 14/4/2015 | GISAID |
| EPI631587 | A/NewJersey/50/2015(H3N2) | 5/7/2015 | GISAID |
| EPI631595 | A/Florida/52/2015(H3N2) | 22/6/2015 | GISAID |

| | | | |
|---|---|---|---|
| EPI631606 | A/Wisconsin/69/2015(H3N2) | 1/7/2015 | GISAID |
| EPI631614 | A/Illinois/12/2015(H3N2) | 30/6/2015 | GISAID |
| EPI631622 | A/Florida/55/2015(H3N2) | 4/7/2015 | GISAID |
| EPI631638 | A/Alaska/26/2015(H3N2) | 14/6/2015 | GISAID |
| EPI631646 | A/Alaska/17/2015(H3N2) | 1/6/2015 | GISAID |
| EPI631654 | A/Kentucky/11/2015(H3N2) | 17/2/2015 | GISAID |
| EPI631662 | A/Hawaii/28/2015(H3N2) | 4/4/2015 | GISAID |
| EPI631670 | A/Brazil/187/2015(H3N2) | 4/5/2015 | GISAID |
| EPI631678 | A/Minnesota/23/2015(H3N2) | 19/2/2015 | GISAID |
| EPI631693 | A/SouthDakota/23/2015(H3N2) | 29/1/2015 | GISAID |
| EPI631701 | A/Peru/34/2015(H3N2) | 24/4/2015 | GISAID |
| EPI631709 | A/Peru/36/2015(H3N2) | 4/5/2015 | GISAID |
| EPI631717 | A/Peru/37/2015(H3N2) | 10/5/2015 | GISAID |
| EPI631725 | A/ElSalvador/283/2015(H3N2) | 9/3/2015 | GISAID |
| EPI631733 | A/Michigan/37/2015(H3N2) | 27/6/2015 | GISAID |
| EPI631749 | A/Brazil/9492/2015(H3N2) | 15/3/2015 | GISAID |
| EPI631757 | A/NorthCarolina/26/2015(H3N2) | 30/6/2015 | GISAID |
| EPI631765 | A/Brazil/9517/2015(H3N2) | 21/3/2015 | GISAID |
| EPI631787 | A/Idaho/18/2015(H3N2) | 27/3/2015 | GISAID |
| EPI631802 | A/California/73/2015(H3N2) | 20/2/2015 | GISAID |
| EPI631810 | A/Hawaii/45/2015(H3N2) | 8/6/2015 | GISAID |
| EPI631818 | A/Massachusetts/18/2015(H3N2) | 29/3/2015 | GISAID |
| EPI631826 | A/Iowa/14/2015(H3N2) | 12/3/2015 | GISAID |
| EPI631834 | A/Alabama/08/2015(H3N2) | 14/2/2015 | GISAID |
| EPI631842 | A/Michigan/27/2015(H3N2) | 24/2/2015 | GISAID |
| EPI631855 | A/Florida/43/2015(H3N2) | 31/3/2015 | GISAID |
| EPI631863 | A/Washington/36/2015(H3N2) | 10/5/2015 | GISAID |
| EPI631871 | A/Nebraska/16/2015(H3N2) | 23/3/2015 | GISAID |
| EPI631879 | A/Maryland/13/2015(H3N2) | 4/3/2015 | GISAID |
| EPI631887 | A/NewCaledonia/71/2014NIB-91(H3N2) | 2014 | GISAID |
| EPI631895 | A/Mississippi/10/2015(H3N2) | 23/2/2015 | GISAID |
| EPI631903 | A/SouthDakota/12/2015(H3N2) | 4/4/2015 | GISAID |
| EPI631911 | A/Washington/33/2014(H3N2) | 21/11/2014 | GISAID |
| EPI633798 | A/StPetersburg/70/2015(H3N2) | 30/1/2015 | GISAID |
| EPI633806 | A/Indiana/04/2015(H3N2) | 18/1/2015 | GISAID |
| EPI633821 | A/Virginia/10/2015(H3N2) | 20/1/2015 | GISAID |
| EPI633829 | A/California/44/2015(H3N2) | 4/1/2015 | GISAID |

| | | | |
|---|---|---|---|
| EPI633837 | A/RhodeIsland/07/2015(H3N2) | 19/1/2015 | GISAID |
| EPI633852 | A/Hawaii/43/2015(H3N2) | 8/7/2015 | GISAID |
| EPI633874 | A/Iowa/13/2015(H3N2) | 30/1/2015 | GISAID |
| EPI633888 | A/Nevada/33/2015(H3N2) | 24/1/2015 | GISAID |
| EPI633896 | A/Hawaii/44/2015(H3N2) | 9/7/2015 | GISAID |
| EPI633904 | A/Brazil/70/2015(H3N2) | 5/1/2015 | GISAID |
| EPI633912 | A/Nevada/12/2015(H3N2) | 30/1/2015 | GISAID |
| EPI633920 | A/Montana/09/2015(H3N2) | 23/1/2015 | GISAID |
| EPI633928 | A/SouthDakota/22/2015(H3N2) | 29/1/2015 | GISAID |
| EPI633936 | A/Wyoming/14/2015(H3N2) | 22/1/2015 | GISAID |
| EPI633944 | A/Nevada/07/2015(H3N2) | 19/1/2015 | GISAID |
| EPI633952 | A/Nevada/32/2015(H3N2) | 23/1/2015 | GISAID |
| EPI633960 | A/Colorado/09/2015(H3N2) | 22/1/2015 | GISAID |
| EPI633985 | A/Oklahoma/15/2015(H3N2) | 30/1/2015 | GISAID |
| EPI633993 | A/NewMexico/21/2015(H3N2) | 17/1/2015 | GISAID |
| EPI634001 | A/Khabarovsk/11/2015(H3N2) | 20/1/2015 | GISAID |
| EPI634022 | A/ElSalvador/49/2015(H3N2) | 9/1/2015 | GISAID |
| EPI634030 | A/ElSalvador/63/2015(H3N2) | 20/1/2015 | GISAID |
| EPI634038 | A/Nevada/23/2015(H3N2) | 26/1/2015 | GISAID |
| EPI634053 | A/Ohio/21/2015(H3N2) | 29/7/2015 | GISAID |
| EPI634061 | A/Ohio/20/2015(H3N2) | 23/7/2015 | GISAID |
| EPI634069 | A/Victoria/503/2015(H3N2) | 31/3/2015 | GISAID |
| EPI634077 | A/Fiji/02/2015(H3N2) | 2/3/2015 | GISAID |
| EPI634085 | A/SouthAustralia/09/2015(H3N2) | 12/2/2015 | GISAID |
| EPI634100 | A/Laos/046/2015(H3N2) | 14/1/2015 | GISAID |
| EPI634108 | A/Laos/052/2015(H3N2) | 14/1/2015 | GISAID |
| EPI634123 | A/Brisbane/47/2015(H3N2) | 18/3/2015 | GISAID |
| EPI634138 | A/Brazil/237/2015(H3N2) | 14/5/2015 | GISAID |
| EPI634151 | A/Missouri/11/2015(H3N2) | 24/1/2015 | GISAID |
| EPI634159 | A/Kansas/09/2015(H3N2) | 30/1/2015 | GISAID |
| EPI634167 | A/Khabarovsk/04/2015(H3N2) | 16/1/2015 | GISAID |
| EPI634175 | A/NewJersey/40/2015(H3N2) | 19/1/2015 | GISAID |
| EPI634183 | A/Virginia/09/2015(H3N2) | 19/1/2015 | GISAID |
| EPI634191 | A/Khabarovsk/07/2015(H3N2) | 20/1/2015 | GISAID |
| EPI634199 | A/Missouri/05/2015(H3N2) | 28/1/2015 | GISAID |
| EPI634207 | A/Wisconsin/30/2015(H3N2) | 26/1/2015 | GISAID |
| EPI638978 | A/Alaska/33/2015(H3N2) | 19/6/2015 | GISAID |

| | | | |
|---|---|---|---|
| EPI638986 | A/Alaska/35/2015(H3N2) | 17/6/2015 | GISAID |
| EPI638994 | A/Alaska/29/2015(H3N2) | 18/6/2015 | GISAID |
| EPI639002 | A/Alaska/38/2015(H3N2) | 21/6/2015 | GISAID |
| EPI639010 | A/Dominica/672/2015(H3N2) | 23/3/2015 | GISAID |
| EPI639018 | A/Alaska/34/2015(H3N2) | 19/6/2015 | GISAID |
| EPI639026 | A/Dominica/642/2015(H3N2) | 14/3/2015 | GISAID |
| EPI639034 | A/Iowa/20/2015(H3N2) | 25/7/2015 | GISAID |
| EPI639042 | A/Iowa/30/2015(H3N2) | 30/7/2015 | GISAID |
| EPI639057 | A/Dominica/529/2015(H3N2) | 10/3/2015 | GISAID |
| EPI639065 | A/Mexico/1797/2015(H3N2) | 6/7/2015 | GISAID |
| EPI639073 | A/Alaska/66/2015(H3N2) | 24/6/2015 | GISAID |
| EPI639081 | A/Mexico/1783/2015(H3N2) | 10/7/2015 | GISAID |
| EPI639089 | A/Alaska/68/2015(H3N2) | 25/6/2015 | GISAID |
| EPI639104 | A/Mexico/1670/2015(H3N2) | 28/5/2015 | GISAID |
| EPI639112 | A/Ecuador/2384/2015(H3N2) | 5/6/2015 | GISAID |
| EPI639120 | A/Ecuador/1670/2015(H3N2) | 18/2/2015 | GISAID |
| EPI639128 | A/Ecuador/1668/2015(H3N2) | 6/6/2015 | GISAID |
| EPI639136 | A/Ecuador/1673/2015(H3N2) | 6/8/2015 | GISAID |
| EPI639149 | A/Alaska/65/2015(H3N2) | 26/6/2015 | GISAID |
| EPI639157 | A/Hawaii/47/2015(H3N2) | 5/7/2015 | GISAID |
| EPI639165 | A/Alaska/69/2015(H3N2) | 26/6/2015 | GISAID |
| EPI639173 | A/Hawaii/46/2015(H3N2) | 7/7/2015 | GISAID |
| EPI639181 | A/Brazil/8004/2015(H3N2) | 10/2/2015 | GISAID |
| EPI639189 | A/Florida/57/2015(H3N2) | 26/7/2015 | GISAID |
| EPI639197 | A/Santiago/37245/2015(H3N2) | 28/5/2015 | GISAID |
| EPI639205 | A/Santiago/40992/2015(H3N2) | 15/6/2015 | GISAID |
| EPI639213 | A/Santiago/42575/2015(H3N2) | 24/6/2015 | GISAID |
| EPI639221 | A/Santiago/44139/2015(H3N2) | 26/6/2015 | GISAID |
| EPI639229 | A/Santiago/35385/2015(H3N2) | 28/5/2015 | GISAID |
| EPI639237 | A/Florida/58/2015(H3N2) | 25/7/2015 | GISAID |
| EPI639252 | A/Michigan/38/2015(H3N2) | 12/6/2015 | GISAID |
| EPI639260 | A/Valparaiso/42142/2015(H3N2) | 23/6/2015 | GISAID |
| EPI639271 | A/Alaska/37/2015(H3N2) | 21/6/2015 | GISAID |
| EPI639279 | A/Alaska/36/2015(H3N2) | 16/6/2015 | GISAID |
| EPI639287 | A/Washington/39/2015(H3N2) | 14/7/2015 | GISAID |
| EPI639295 | A/Alaska/30/2015(H3N2) | 19/6/2015 | GISAID |
| EPI639303 | A/Alaska/31/2015(H3N2) | 20/6/2015 | GISAID |

| | | | |
|---|---|---|---|
| EPI642352 | A/Maryland/58/2014(H3N2) | 28/12/2014 | GISAID |
| EPI642513 | A/Michigan/39/2015(H3N2) | 12/6/2015 | GISAID |
| EPI642560 | A/Stockholm/40/2015(H3N2) | 19/8/2015 | GISAID |
| EPI643036 | A/Florida/59/2015(H3N2) | 9/8/2015 | GISAID |
| EPI643044 | A/Florida/64/2015(H3N2) | 17/8/2015 | GISAID |
| EPI643052 | A/Florida/62/2015(H3N2) | 18/8/2015 | GISAID |
| EPI643060 | A/Florida/61/2015(H3N2) | 19/8/2015 | GISAID |
| EPI643068 | A/Florida/63/2015(H3N2) | 18/8/2015 | GISAID |
| EPI643510 | A/Alaska/148/2015(H3N2) | 26/7/2015 | GISAID |
| EPI643518 | A/Alaska/150/2015(H3N2) | 19/7/2015 | GISAID |
| EPI643526 | A/Alaska/149/2015(H3N2) | 7/7/2015 | GISAID |
| EPI643534 | A/Alaska/135/2015(H3N2) | 2/8/2015 | GISAID |
| EPI643542 | A/Alaska/146/2015(H3N2) | 23/7/2015 | GISAID |
| EPI643556 | A/Alaska/142/2015(H3N2) | 10/8/2015 | GISAID |
| EPI643571 | A/Paisvasco/635/2015(H3N2) | 2/2/2015 | GISAID |
| EPI643579 | A/Alaska/144/2015(H3N2) | 3/8/2015 | GISAID |
| EPI643587 | A/Alaska/141/2015(H3N2) | 5/8/2015 | GISAID |
| EPI643595 | A/Alaska/138/2015(H3N2) | 3/8/2015 | GISAID |
| EPI643603 | A/Texas/65/2015(H3N2) | 6/8/2015 | GISAID |
| EPI643611 | A/Alaska/137/2015(H3N2) | 2/8/2015 | GISAID |
| EPI643619 | A/Panama/316597/2015(H3N2) | 24/6/2015 | GISAID |
| EPI643627 | A/Alaska/140/2015(H3N2) | 11/8/2015 | GISAID |
| EPI643635 | A/Alaska/143/2015(H3N2) | 6/8/2015 | GISAID |
| EPI643643 | A/Alaska/147/2015(H3N2) | 24/7/2015 | GISAID |
| EPI643651 | A/Alaska/139/2015(H3N2) | 4/8/2015 | GISAID |
| EPI643659 | A/Sassari/8/2015(H3N2) | 2/2/2015 | GISAID |
| EPI643674 | A/Italy/FVG-89/2015(H3N2) | 13/2/2015 | GISAID |
| EPI643682 | A/Serbia/NS1241/2015(H3N2) | 19/3/2015 | GISAID |
| EPI643690 | A/Iowa/22/2015(H3N2) | 30/7/2015 | GISAID |
| EPI643698 | A/Moscow/134/2015(H3N2) | 6/4/2015 | GISAID |
| EPI643706 | A/Guatemala/9704/2015(H3N2) | 25/5/2015 | GISAID |
| EPI643714 | A/Argentina/229/2015(H3N2) | 30/7/2015 | GISAID |
| EPI643722 | A/Iowa/23/2015(H3N2) | 30/7/2015 | GISAID |
| EPI643730 | A/Iowa/29/2015(H3N2) | 30/7/2015 | GISAID |
| EPI643738 | A/Argentina/235/2015(H3N2) | 29/7/2015 | GISAID |
| EPI643746 | A/Argentina/236/2015(H3N2) | 30/7/2015 | GISAID |
| EPI643761 | A/Argentina/254/2015(H3N2) | 27/7/2015 | GISAID |

| | | | |
|---|---|---|---|
| EPI643769 | A/Argentina/250/2015(H3N2) | 3/8/2015 | GISAID |
| EPI643780 | A/Argentina/183/2015(H3N2) | 27/7/2015 | GISAID |
| EPI643788 | A/Argentina/255/2015(H3N2) | 4/8/2015 | GISAID |
| EPI643803 | A/Iowa/31/2015(H3N2) | 30/7/2015 | GISAID |
| EPI643816 | A/Argentina/261/2015(H3N2) | 4/8/2015 | GISAID |
| EPI643824 | A/Missouri/12/2015(H3N2) | 31/1/2015 | GISAID |
| EPI643832 | A/Argentina/196/2015(H3N2) | 27/7/2015 | GISAID |
| EPI643840 | A/Serbia/NS1010/2015(H3N2) | 6/2/2015 | GISAID |
| EPI643848 | A/Argentina/218/2015(H3N2) | 27/7/2015 | GISAID |
| EPI643856 | A/Ecuador/2369/2015(H3N2) | 25/4/2015 | GISAID |
| EPI643864 | A/Argentina/234/2015(H3N2) | 29/7/2015 | GISAID |
| EPI643872 | A/Argentina/166/2015(H3N2) | 25/7/2015 | GISAID |
| EPI643880 | A/Argentina/135/2015(H3N2) | 19/7/2015 | GISAID |
| EPI643888 | A/Iowa/25/2015(H3N2) | 30/7/2015 | GISAID |
| EPI643896 | A/Jordan/30024/2015(H3N2) | 5/2/2015 | GISAID |
| EPI643904 | A/Iowa/17/2015(H3N2) | 29/1/2015 | GISAID |
| EPI643919 | A/Maine/24/2015(H3N2) | 29/1/2015 | GISAID |
| EPI643927 | A/Brazil/56/2015(H3N2) | 27/3/2015 | GISAID |
| EPI643935 | A/Iowa/19/2015(H3N2) | 22/7/2015 | GISAID |
| EPI643943 | A/Jordan/10702/2015(H3N2) | 22/3/2015 | GISAID |
| EPI643951 | A/Iowa/24/2015(H3N2) | 30/7/2015 | GISAID |
| EPI643959 | A/Jordan/30052/2015(H3N2) | 14/2/2015 | GISAID |
| EPI643967 | A/California/53/2015(H3N2) | 14/1/2015 | GISAID |
| EPI643975 | A/Massachusetts/12/2015(H3N2) | 26/1/2015 | GISAID |
| EPI643983 | A/Iowa/28/2015(H3N2) | 30/7/2015 | GISAID |
| EPI643991 | A/Oregon/10/2015(H3N2) | 11/1/2015 | GISAID |
| EPI643999 | A/Bangladesh/3400/2015(H3N2) | 25/5/2015 | GISAID |
| EPI644007 | A/Iowa/12/2015(H3N2) | 31/1/2015 | GISAID |
| EPI644015 | A/Colombia/5614/2015(H3N2) | 21/7/2015 | GISAID |
| EPI644023 | A/Guatemala/27/2015(H3N2) | 12/6/2015 | GISAID |
| EPI644031 | A/Texas/22/2015(H3N2) | 13/1/2015 | GISAID |
| EPI644039 | A/Colombia/5615/2015(H3N2) | 23/7/2015 | GISAID |
| EPI644047 | A/Guatemala/142/2015(H3N2) | 18/6/2015 | GISAID |
| EPI644055 | A/Panama/316634/2015(H3N2) | 17/7/2015 | GISAID |
| EPI644063 | A/Brazil/96/2015(H3N2) | 11/2/2015 | GISAID |
| EPI644071 | A/Moscow/99/2015(H3N2) | 10/3/2015 | GISAID |
| EPI644079 | A/Arizona/11/2015(H3N2) | 14/7/2015 | GISAID |

| | | | |
|---|---|---|---|
| EPI644087 | A/Arizona/10/2015(H3N2) | 1/8/2015 | GISAID |
| EPI644095 | A/Guatemala/9922/2015(H3N2) | 5/6/2015 | GISAID |
| EPI644103 | A/Alaska/145/2015(H3N2) | 24/7/2015 | GISAID |
| EPI644283 | A/Singapore/KK943/2014(H3N2) | 7/10/2014 | GISAID |
| EPI646110 | A/OKAYAMA/2/2015(H3N2) | 14/1/2015 | GISAID |
| EPI647812 | A/Hawaii/35/2015(H3N2) | 26/5/2015 | GISAID |
| EPI647820 | A/Hawaii/36/2015(H3N2) | 11/6/2015 | GISAID |
| EPI647828 | A/Hawaii/34/2015(H3N2) | 21/5/2015 | GISAID |
| EPI647836 | A/NewZealand/521/2015(H3N2) | 2/7/2015 | GISAID |
| EPI647844 | A/NewZealand/324/2015(H3N2) | 15/6/2015 | GISAID |
| EPI647852 | A/Brazil/2837/2015(H3N2) | 2/5/2015 | GISAID |
| EPI647860 | A/Nonthaburi/333/2015(H3N2) | 27/7/2015 | GISAID |
| EPI647868 | A/NewZealand/322/2015(H3N2) | 16/6/2015 | GISAID |
| EPI647876 | A/Rayong/335/2015(H3N2) | 21/7/2015 | GISAID |
| EPI647884 | A/Nonthaburi/340/2015(H3N2) | 22/7/2015 | GISAID |
| EPI647892 | A/Nonthaburi/345/2015(H3N2) | 2/8/2015 | GISAID |
| EPI647900 | A/NewZealand/381/2015(H3N2) | 22/6/2015 | GISAID |
| EPI647908 | A/NewZealand/519/2015(H3N2) | 3/7/2015 | GISAID |
| EPI647916 | A/NewZealand/323/2015(H3N2) | 16/6/2015 | GISAID |
| EPI647924 | A/NewZealand/385/2015(H3N2) | 21/6/2015 | GISAID |
| EPI647932 | A/NewZealand/500/2015(H3N2) | 1/7/2015 | GISAID |
| EPI647940 | A/NewZealand/282/2015(H3N2) | 12/6/2015 | GISAID |
| EPI647948 | A/NewZealand/383/2015(H3N2) | 22/6/2015 | GISAID |
| EPI647956 | A/NewZealand/382/2015(H3N2) | 22/6/2015 | GISAID |
| EPI647964 | A/NewZealand/476/2015(H3N2) | 1/7/2015 | GISAID |
| EPI647972 | A/Oregon/16/2015(H3N2) | 15/7/2015 | GISAID |
| EPI647980 | A/Oregon/18/2015(H3N2) | 18/7/2015 | GISAID |
| EPI647988 | A/Paraguay/3585/2015(H3N2) | 26/5/2015 | GISAID |
| EPI647996 | A/Paraguay/6248/2015(H3N2) | 3/8/2015 | GISAID |
| EPI648004 | A/Paraguay/3371/2015(H3N2) | 22/5/2015 | GISAID |
| EPI648012 | A/Paraguay/6991/2015(H3N2) | 4/8/2015 | GISAID |
| EPI648020 | A/Paraguay/5554/2015(H3N2) | 4/6/2015 | GISAID |
| EPI648028 | A/Guatemala/18/2015(H3N2) | 5/5/2015 | GISAID |
| EPI648036 | A/Wisconsin/66/2015(H3N2) | 21/4/2015 | GISAID |
| EPI649378 | A/Guatemala/21/2015(H3N2) | 5/5/2015 | GISAID |
| EPI649386 | A/Hokkaido/06/2015(H3N2) | 9/2/2015 | GISAID |
| EPI649394 | A/Alabama/09/2015(H3N2) | 14/2/2015 | GISAID |

| | | | |
|---|---|---|---|
| EPI649402 | A/NewZealand/588/2015(H3N2) | 15/6/2015 | GISAID |
| EPI649410 | A/Guatemala/19/2015(H3N2) | 5/5/2015 | GISAID |
| EPI649418 | A/Ohio/26/2015(H3N2) | 17/8/2015 | GISAID |
| EPI649426 | A/Iowa/26/2015(H3N2) | 30/7/2015 | GISAID |
| EPI649434 | A/Alabama/07/2015(H3N2) | 11/2/2015 | GISAID |
| EPI649442 | A/Guatemala/15/2015(H3N2) | 3/5/2015 | GISAID |
| EPI649450 | A/Guatemala/5015/2015(H3N2) | 18/5/2015 | GISAID |
| EPI649458 | A/Guatemala/5915/2015(H3N2) | 11/6/2015 | GISAID |
| EPI649466 | A/Massachusetts/23/2015(H3N2) | 5/8/2015 | GISAID |
| EPI649474 | A/Ecuador/1673/2015(H3N2) | 6/8/2015 | GISAID |
| EPI649482 | A/NewYork/64/2015(H3N2) | 10/8/2015 | GISAID |
| EPI649490 | A/Prachuapkhirikhan/289/2015(H3N2) | 2/7/2015 | GISAID |
| EPI649498 | A/Gifu/46/2014(H3N2) | 12/12/2014 | GISAID |
| EPI649506 | A/Nakhonratchaisma/35/2015(H3N2) | 16/5/2015 | GISAID |
| EPI649514 | A/NewZealand/431/2015(H3N2) | 27/5/2015 | GISAID |
| EPI649522 | A/NewYork/57/2015(H3N2) | 21/7/2015 | GISAID |
| EPI649533 | A/NewZealand/87/2015(H3N2) | 21/2/2015 | GISAID |
| EPI649541 | A/Hyogo/4001/2014(H3N2) | 22/1/2015 | GISAID |
| EPI649549 | A/Chanthaburi/46/2015(H3N2) | 12/6/2015 | GISAID |
| EPI649557 | A/Chanthaburi/40/2015(H3N2) | 2/6/2015 | GISAID |
| EPI649565 | A/NewZealand/562/2015(H3N2) | 6/7/2015 | GISAID |
| EPI649573 | A/NewYork/62/2015(H3N2) | 4/8/2015 | GISAID |
| EPI649588 | A/NewZealand/683/2015(H3N2) | 6/6/2015 | GISAID |
| EPI649596 | A/Fukui/78/2015(H3N2) | 2/3/2015 | GISAID |
| EPI649604 | A/Novosibirsk/64/2015(H3N2) | 27/2/2015 | GISAID |
| EPI649612 | A/Moscow/104/2015(H3N2) | 11/3/2015 | GISAID |
| EPI649620 | A/Bangkok/43/2015(H3N2) | 5/6/2015 | GISAID |
| EPI649628 | A/NorthCarolina/28/2015(H3N2) | 20/8/2015 | GISAID |
| EPI649636 | A/Ohio/23/2015(H3N2) | 3/8/2015 | GISAID |
| EPI649651 | A/Buryatia/19/2015(H3N2) | 26/1/2015 | GISAID |
| EPI649669 | A/Chiba/61/2014(H3N2) | 15/1/2015 | GISAID |
| EPI649677 | A/NewYork/59/2015(H3N2) | 29/7/2015 | GISAID |
| EPI649691 | A/NewZealand/536/2015(H3N2) | 7/7/2015 | GISAID |
| EPI649699 | A/NewZealand/686/2015(H3N2) | 4/6/2015 | GISAID |
| EPI649707 | A/Mexico/1384/2015(H3N2) | 24/3/2015 | GISAID |
| EPI649715 | A/NewZealand/532/2015(H3N2) | 2/7/2015 | GISAID |
| EPI649723 | A/NewYork/60/2015(H3N2) | 1/8/2015 | GISAID |

| | | | |
|---|---|---|---|
| EPI649731 | A/Chanthaburi/49/2015(H3N2) | 19/5/2015 | GISAID |
| EPI649739 | A/NewZealand/526/2015(H3N2) | 5/7/2015 | GISAID |
| EPI649747 | A/NewZealand/588/2015(H3N2) | 15/6/2015 | GISAID |
| EPI649755 | A/Mexico/1402/2015(H3N2) | 18/3/2015 | GISAID |
| EPI649763 | A/Ohio/24/2015(H3N2) | 3/8/2015 | GISAID |
| EPI649771 | A/Mexico/1401/2015(H3N2) | 1/4/2015 | GISAID |
| EPI649779 | A/NorthCarolina/29/2015(H3N2) | 17/8/2015 | GISAID |
| EPI649787 | A/Mexico/1233/2015(H3N2) | 2/3/2015 | GISAID |
| EPI649795 | A/Wisconsin/74/2015(H3N2) | 1/9/2015 | GISAID |
| EPI649803 | A/Wisconsin/75/2015(H3N2) | 1/9/2015 | GISAID |
| EPI649811 | A/NewYork/63/2015(H3N2) | 6/8/2015 | GISAID |
| EPI649819 | A/Gifu/46/2014(H3N2) | 12/12/2014 | GISAID |
| EPI649827 | A/Fukui/78/2015(H3N2) | 2/3/2015 | GISAID |
| EPI649842 | A/NewYork/58/2015(H3N2) | 3/8/2015 | GISAID |
| EPI649850 | A/Hokkaido/06/2015(H3N2) | 9/2/2015 | GISAID |
| EPI649858 | A/Chiba/61/2014(H3N2) | 15/1/2015 | GISAID |
| EPI649866 | A/Iowa/37/2015(H3N2) | 30/7/2015 | GISAID |
| EPI649874 | A/Iowa/38/2015(H3N2) | 30/7/2015 | GISAID |
| EPI649882 | A/Virginia/32/2015(H3N2) | 17/6/2015 | GISAID |
| EPI649890 | A/Mexico/1546/2015(H3N2) | 6/5/2015 | GISAID |
| EPI649905 | A/Moscow/102/2015(H3N2) | 14/3/2015 | GISAID |
| EPI649913 | A/N.Novgorod/143/2015(H3N2) | 18/3/2015 | GISAID |
| EPI649921 | A/Vladimir/137/2015(H3N2) | 12/3/2015 | GISAID |
| EPI649929 | A/Guatemala/13/2015(H3N2) | 12/3/2015 | GISAID |
| EPI649937 | A/Guatemala/15/2015(H3N2) | 3/5/2015 | GISAID |
| EPI649945 | A/Brazil/114/2015(H3N2) | 9/4/2015 | GISAID |
| EPI649953 | A/Guatemala/5015/2015(H3N2) | 18/5/2015 | GISAID |
| EPI649961 | A/Guatemala/3315/2015(H3N2) | 18/3/2015 | GISAID |
| EPI649969 | A/Massachusetts/22/2015(H3N2) | 7/8/2015 | GISAID |
| EPI649977 | A/Guatemala/13/2015(H3N2) | 12/3/2015 | GISAID |
| EPI649985 | A/NewMexico/22/2015(H3N2) | 27/7/2015 | GISAID |
| EPI649993 | A/Montana/17/2015(H3N2) | 13/8/2015 | GISAID |
| EPI650001 | A/Iowa/34/2015(H3N2) | 30/7/2015 | GISAID |
| EPI650009 | A/Montana/16/2015(H3N2) | 8/8/2015 | GISAID |
| EPI650017 | A/Hyogo/4001/2014(H3N2) | 22/1/2015 | GISAID |
| EPI651432 | A/Alaska/86/2015(H3N2) | 6/7/2015 | GISAID |
| EPI651440 | A/Arizona/13/2015(H3N2) | 5/8/2015 | GISAID |

| | | | |
|---|---|---|---|
| EPI651448 | A/Hawaii/48/2015(H3N2) | 17/7/2015 | GISAID |
| EPI651456 | A/Hawaii/49/2015(H3N2) | 23/7/2015 | GISAID |
| EPI651464 | A/Alaska/88/2015(H3N2) | 1/7/2015 | GISAID |
| EPI651480 | A/Alaska/82/2015(H3N2) | 6/7/2015 | GISAID |
| EPI651488 | A/Connecticut/01/2015(H3N2) | 12/7/2015 | GISAID |
| EPI651496 | A/Alaska/72/2015(H3N2) | 22/6/2015 | GISAID |
| EPI651504 | A/Alaska/87/2015(H3N2) | 3/7/2015 | GISAID |
| EPI651512 | A/Alaska/83/2015(H3N2) | 6/7/2015 | GISAID |
| EPI651520 | A/Alaska/27/2015(H3N2) | 14/6/2015 | GISAID |
| EPI651528 | A/Alaska/85/2015(H3N2) | 6/7/2015 | GISAID |
| EPI651536 | A/Alaska/71/2015(H3N2) | 26/6/2015 | GISAID |
| EPI651544 | A/Wisconsin/72/2015(H3N2) | 28/8/2015 | GISAID |
| EPI651552 | A/Alaska/73/2015(H3N2) | 23/6/2015 | GISAID |
| EPI651560 | A/Michigan/40/2015(H3N2) | 7/7/2015 | GISAID |
| EPI651568 | A/Alaska/79/2015(H3N2) | 24/6/2015 | GISAID |
| EPI651576 | A/Alaska/94/2015(H3N2) | 2/7/2015 | GISAID |
| EPI651584 | A/Arizona/14/2015(H3N2) | 26/8/2015 | GISAID |
| EPI651592 | A/Alaska/91/2015(H3N2) | 3/7/2015 | GISAID |
| EPI651600 | A/Alaska/77/2015(H3N2) | 25/6/2015 | GISAID |
| EPI651608 | A/Alaska/92/2015(H3N2) | 2/7/2015 | GISAID |
| EPI651616 | A/Alaska/78/2015(H3N2) | 28/6/2015 | GISAID |
| EPI651624 | A/Alaska/93/2015(H3N2) | 1/7/2015 | GISAID |
| EPI651632 | A/Alaska/75/2015(H3N2) | 23/6/2015 | GISAID |
| EPI651640 | A/Alaska/90/2015(H3N2) | 3/7/2015 | GISAID |
| EPI651648 | A/Alabama/12/2015(H3N2) | 5/9/2015 | GISAID |
| EPI651656 | A/Alabama/10/2015(H3N2) | 8/9/2015 | GISAID |
| EPI651664 | A/Minnesota/40/2015(H3N2) | 1/9/2015 | GISAID |
| EPI651672 | A/Alabama/11/2015(H3N2) | 8/9/2015 | GISAID |
| EPI651680 | A/Alaska/84/2015(H3N2) | 6/7/2015 | GISAID |
| EPI651696 | A/Alaska/89/2015(H3N2) | 1/7/2015 | GISAID |
| EPI651704 | A/Alaska/80/2015(H3N2) | 30/6/2015 | GISAID |
| EPI651712 | A/Alaska/95/2015(H3N2) | 1/7/2015 | GISAID |
| EPI651720 | A/NewHampshire/38/2015(H3N2) | 2/9/2015 | GISAID |
| EPI651727 | A/Connecticut/02/2015(H3N2) | 29/8/2015 | GISAID |
| EPI651735 | A/Texas/66/2015(H3N2) | 25/8/2015 | GISAID |
| EPI651743 | A/Hawaii/52/2015(H3N2) | 19/8/2015 | GISAID |
| EPI651751 | A/Wisconsin/77/2015(H3N2) | 1/9/2015 | GISAID |

| | | | |
|---|---|---|---|
| EPI651759 | A/Hawaii/53/2015(H3N2) | 23/8/2015 | GISAID |
| EPI651767 | A/NewYork/65/2015(H3N2) | 25/8/2015 | GISAID |
| EPI651775 | A/Florida/68/2015(H3N2) | 2/9/2015 | GISAID |
| EPI651783 | A/Hawaii/54/2015(H3N2) | 25/8/2015 | GISAID |
| EPI651791 | A/Alaska/81/2015(H3N2) | 2/7/2015 | GISAID |
| EPI653163 | A/Michigan/43/2015(H3N2) | 24/8/2015 | GISAID |
| EPI653171 | A/Michigan/44/2015(H3N2) | 21/8/2015 | GISAID |
| EPI653179 | A/Michigan/41/2015(H3N2) | 15/8/2015 | GISAID |
| EPI653187 | A/Alaska/43/2015(H3N2) | 19/6/2015 | GISAID |
| EPI653201 | A/HongKong/4801/2014(H3N2) | 26/2/2014 | GISAID |
| EPI653209 | A/Switzerland/9715293/2013NIB-88(H3N2) | 2013 | GISAID |
| EPI662361 | A/Stockholm/42/2015(H3N2) | 26/9/2015 | GISAID |
| EPI662369 | A/Uruguay/86/2015(H3N2) | 11/8/2015 | GISAID |
| EPI662377 | A/Hawaii/55/2015(H3N2) | 29/8/2015 | GISAID |
| EPI662385 | A/Uruguay/44/2015(H3N2) | 8/7/2015 | GISAID |
| EPI662394 | A/Florida/73/2015(H3N2) | 14/9/2015 | GISAID |
| EPI662402 | A/Colombia/5402/2015(H3N2) | 25/6/2015 | GISAID |
| EPI662410 | A/Colombia/235/2015(H3N2) | 18/1/2015 | GISAID |
| EPI662418 | A/Colombia/5438/2015(H3N2) | 30/6/2015 | GISAID |
| EPI662426 | A/Colombia/5364/2015(H3N2) | 30/6/2015 | GISAID |
| EPI662434 | A/Wisconsin/70/2015(H3N2) | 6/8/2015 | GISAID |
| EPI662442 | A/Colombia/241/2015(H3N2) | 7/6/2015 | GISAID |
| EPI662450 | A/Hawaii/56/2015(H3N2) | 2/9/2015 | GISAID |
| EPI662458 | A/Wisconsin/71/2015(H3N2) | 2/8/2015 | GISAID |
| EPI662466 | A/Colombia/5215/2015(H3N2) | 4/6/2015 | GISAID |
| EPI662474 | A/Uruguay/122/2015(H3N2) | 13/8/2015 | GISAID |
| EPI662482 | A/Hawaii/57/2015(H3N2) | 13/9/2015 | GISAID |
| EPI662490 | A/Uruguay/39/2015(H3N2) | 16/6/2015 | GISAID |
| EPI662498 | A/Uruguay/43/2015(H3N2) | 6/7/2015 | GISAID |
| EPI662506 | A/Uruguay/54/2015(H3N2) | 20/7/2015 | GISAID |
| EPI662514 | A/Uruguay/55/2015(H3N2) | 21/7/2015 | GISAID |
| EPI662522 | A/NewYork/64/2015(H3N2) | 10/8/2015 | GISAID |
| EPI662537 | A/Alaska/77/2015(H3N2) | 25/6/2015 | GISAID |
| EPI662545 | A/Alaska/75/2015(H3N2) | 23/6/2015 | GISAID |
| EPI662553 | A/Alaska/80/2015(H3N2) | 30/6/2015 | GISAID |
| EPI662561 | A/Alaska/79/2015(H3N2) | 24/6/2015 | GISAID |
| EPI662569 | A/Alaska/93/2015(H3N2) | 1/7/2015 | GISAID |

| | | | |
|---|---|---|---|
| EPI662577 | A/Alaska/83/2015(H3N2) | 6/7/2015 | GISAID |
| EPI662585 | A/Alaska/86/2015(H3N2) | 6/7/2015 | GISAID |
| EPI667522 | A/Massachusetts/14/2014(H3N2) | 19/11/2014 | GISAID |
| EPI667530 | A/Hawaii/34/2015(H3N2) | 21/5/2015 | GISAID |
| EPI667548 | A/Alaska/45/2015(H3N2) | 23/6/2015 | GISAID |
| EPI667556 | A/Alaska/46/2015(H3N2) | 21/6/2015 | GISAID |
| EPI667564 | A/Alaska/48/2015(H3N2) | 21/6/2015 | GISAID |
| EPI667572 | A/DominicanRepublic/8946/2015(H3N2) | 11/3/2015 | GISAID |
| EPI667580 | A/Brazil/8433/2015(H3N2) | 16/4/2015 | GISAID |
| EPI667588 | A/Brazil/9850/2015(H3N2) | 13/3/2015 | GISAID |
| EPI667596 | A/Brazil/1804/2015(H3N2) | 20/3/2015 | GISAID |
| EPI667604 | A/Brazil/440/2015(H3N2) | 18/2/2015 | GISAID |
| EPI667620 | A/Brazil/8925/2015(H3N2) | 17/3/2015 | GISAID |
| EPI667628 | A/Brazil/4996/2015(H3N2) | 16/3/2015 | GISAID |
| EPI667636 | A/Brazil/6261/2015(H3N2) | 18/2/2015 | GISAID |
| EPI667652 | A/Ecuador/2404/2015(H3N2) | 21/5/2015 | GISAID |
| EPI667660 | A/Ecuador/572/2015(H3N2) | 14/5/2015 | GISAID |
| EPI667668 | A/Ecuador/1695/2015(H3N2) | 29/6/2015 | GISAID |
| EPI667676 | A/Ecuador/821/2015(H3N2) | 5/2/2015 | GISAID |
| EPI667684 | A/Ecuador/1845/2015(H3N2) | 8/5/2015 | GISAID |
| EPI667692 | A/Alaska/67/2015(H3N2) | 1/7/2015 | GISAID |
| EPI667700 | A/NewYork/55/2015(H3N2) | 10/7/2015 | GISAID |
| EPI667708 | A/Ohio/18/2015(H3N2) | 23/7/2015 | GISAID |
| EPI667716 | A/Oregon/11/2015(H3N2) | 30/6/2015 | GISAID |
| EPI667727 | A/Nebraska/17/2015(H3N2) | 15/7/2015 | GISAID |
| EPI667735 | A/Laos/218/2015(H3N2) | 25/2/2015 | GISAID |
| EPI667749 | A/Laos/372/2015(H3N2) | 20/4/2015 | GISAID |
| EPI667757 | A/Laos/529/2015(H3N2) | 25/5/2015 | GISAID |
| EPI667765 | A/Laos/535/2015(H3N2) | 25/5/2015 | GISAID |
| EPI667773 | A/Laos/179/2015(H3N2) | 29/4/2015 | GISAID |
| EPI667781 | A/Utah/15/2015(H3N2) | 13/7/2015 | GISAID |
| EPI667786 | A/Acre/133946-IEC/2015(H3N2) | 23/1/2015 | GISAID |
| EPI667801 | A/Para/134244-IEC/2015(H3N2) | 18/3/2015 | GISAID |
| EPI667809 | A/Para/134324-IEC/2015(H3N2) | 23/3/2015 | GISAID |
| EPI667817 | A/Para/134473-IEC/2015(H3N2) | 31/3/2015 | GISAID |
| EPI667825 | A/Para/134575-IEC/2015(H3N2) | 6/4/2015 | GISAID |
| EPI667833 | A/RioGrandeDoNorte/134592-IEC/2015(H3N2) | 31/3/2015 | GISAID |

| | | | |
|---|---|---|---|
| EPI667841 | A/RioGrandeDoNorte/134696-IEC/2015(H3N2) | 8/4/2015 | GISAID |
| EPI667849 | A/Para/134701-IEC/2015(H3N2) | 4/4/2015 | GISAID |
| EPI667864 | A/PuertoRico/04/2015(H3N2) | 13/7/2015 | GISAID |
| EPI667872 | A/PuertoRico/05/2015(H3N2) | 14/7/2015 | GISAID |
| EPI667880 | A/PuertoRico/06/2015(H3N2) | 21/7/2015 | GISAID |
| EPI667888 | A/PuertoRico/07/2015(H3N2) | 7/8/2015 | GISAID |
| EPI667896 | A/PuertoRico/08/2015(H3N2) | 10/8/2015 | GISAID |
| EPI667910 | A/Dominica/645/2015(H3N2) | 16/3/2015 | GISAID |
| EPI667918 | A/St.VincentAndGrenadines/422/2015(H3N2) | 10/2/2015 | GISAID |
| EPI667926 | A/St.VincentAndGrenadines/424/2015(H3N2) | 16/2/2015 | GISAID |
| EPI667941 | A/Oregon/17/2015(H3N2) | 1/7/2015 | GISAID |
| EPI667956 | A/Victoria/511/2015(H3N2) | 16/5/2015 | GISAID |
| EPI667964 | A/Omsk/141/2015(H3N2) | 26/1/2015 | GISAID |
| EPI667972 | A/Komi/99/2015(H3N2) | 27/1/2015 | GISAID |
| EPI667980 | A/Chelyabinsk/192/2015(H3N2) | 20/2/2015 | GISAID |
| EPI667988 | A/Florida/60/2015(H3N2) | 9/8/2015 | GISAID |
| EPI667996 | A/Alaska/160/2015(H3N2) | 18/8/2015 | GISAID |
| EPI668012 | A/Tennessee/18/2015(H3N2) | 18/8/2015 | GISAID |
| EPI668020 | A/Tennessee/20/2015(H3N2) | 19/8/2015 | GISAID |
| EPI668028 | A/Tennessee/20/2015(H3N2) | 19/8/2015 | GISAID |
| EPI668036 | A/Argentina/112/2015(H3N2) | 18/7/2015 | GISAID |
| EPI668044 | A/Argentina/113/2015(H3N2) | 14/7/2015 | GISAID |
| EPI668052 | A/Argentina/114/2015(H3N2) | 16/7/2015 | GISAID |
| EPI668060 | A/Argentina/116/2015(H3N2) | 15/7/2015 | GISAID |
| EPI668068 | A/Argentina/120/2015(H3N2) | 18/7/2015 | GISAID |
| EPI668076 | A/Argentina/121/2015(H3N2) | 16/7/2015 | GISAID |
| EPI668084 | A/Argentina/132/2015(H3N2) | 22/7/2015 | GISAID |
| EPI668092 | A/Argentina/133/2015(H3N2) | 21/7/2015 | GISAID |
| EPI668113 | A/NewHampshire/38/2015(H3N2) | 2/9/2015 | GISAID |
| EPI668121 | A/NewYork/65/2015(H3N2) | 25/8/2015 | GISAID |
| EPI668129 | A/Argentina/11573/2015(H3N2) | 20/7/2015 | GISAID |
| EPI668137 | A/Minnesota/40/2015(H3N2) | 1/9/2015 | GISAID |
| EPI668145 | A/Alabama/10/2015(H3N2) | 8/9/2015 | GISAID |
| EPI668153 | A/Indiana/18/2015(H3N2) | 1/9/2015 | GISAID |
| EPI668161 | A/Indiana/26/2015(H3N2) | 3/9/2015 | GISAID |
| EPI668169 | A/Indiana/27/2015(H3N2) | 3/9/2015 | GISAID |
| EPI668184 | A/Montana/18/2015(H3N2) | 11/9/2015 | GISAID |

| | | | |
|---|---|---|---|
| EPI668192 | A/Montana/19/2015(H3N2) | 13/9/2015 | GISAID |
| EPI668200 | A/Montana/20/2015(H3N2) | 15/9/2015 | GISAID |
| EPI668208 | A/Montana/21/2015(H3N2) | 14/9/2015 | GISAID |
| EPI668216 | A/Montana/23/2015(H3N2) | 17/9/2015 | GISAID |
| EPI668231 | A/Montana/25/2015(H3N2) | 19/9/2015 | GISAID |
| EPI668239 | A/Montana/26/2015(H3N2) | 17/9/2015 | GISAID |
| EPI668247 | A/Montana/27/2015(H3N2) | 20/9/2015 | GISAID |
| EPI668255 | A/Kentucky/15/2015(H3N2) | 28/9/2015 | GISAID |
| EPI668263 | A/Wisconsin/83/2015(H3N2) | 24/9/2015 | GISAID |
| EPI668271 | A/Wisconsin/84/2015(H3N2) | 1/10/2015 | GISAID |
| EPI668279 | A/Wisconsin/76/2015(H3N2) | 1/9/2015 | GISAID |
| EPI668287 | A/Michigan/42/2015(H3N2) | 18/8/2015 | GISAID |
| EPI668311 | A/Michigan/52/2015(H3N2) | 14/9/2015 | GISAID |
| EPI668319 | A/Michigan/53/2015(H3N2) | 12/9/2015 | GISAID |
| EPI668327 | A/Michigan/54/2015(H3N2) | 19/9/2015 | GISAID |
| EPI668335 | A/Michigan/55/2015(H3N2) | 18/9/2015 | GISAID |
| EPI668343 | A/Montana/28/2015(H3N2) | 24/9/2015 | GISAID |
| EPI668351 | A/Montana/29/2015(H3N2) | 26/9/2015 | GISAID |
| EPI669734 | A/Alaska/140/2015(H3N2) | 11/8/2015 | GISAID |
| EPI669976 | A/Stockholm/43/2015(H3N2) | 7/10/2015 | GISAID |
| EPI669984 | A/Stockholm/44/2015(H3N2) | 9/10/2015 | GISAID |
| EPI669992 | A/Malmoe/5/2015(H3N2) | 10/10/2015 | GISAID |
| EPI670000 | A/Stockholm/45/2015(H3N2) | 9/10/2015 | GISAID |
| EPI670351 | A/Alaska/40/2015(H3N2) | 19/6/2015 | GISAID |
| EPI670367 | A/Alaska/45/2015(H3N2) | 23/6/2015 | GISAID |
| EPI670388 | A/Alaska/55/2015(H3N2) | 18/6/2015 | GISAID |
| EPI670396 | A/Alaska/56/2015(H3N2) | 18/6/2015 | GISAID |
| EPI670404 | A/Alaska/57/2015(H3N2) | 18/6/2015 | GISAID |
| EPI670412 | A/Alaska/59/2015(H3N2) | 17/6/2015 | GISAID |
| EPI670420 | A/Alaska/61/2015(H3N2) | 18/6/2015 | GISAID |
| EPI670428 | A/Alaska/62/2015(H3N2) | 21/6/2015 | GISAID |
| EPI670436 | A/Alaska/63/2015(H3N2) | 24/6/2015 | GISAID |
| EPI670444 | A/Alaska/64/2015(H3N2) | 24/6/2015 | GISAID |
| EPI670459 | A/Colombia/5433/2015(H3N2) | 10/7/2015 | GISAID |
| EPI670467 | A/Alaska/96/2015(H3N2) | 30/6/2015 | GISAID |
| EPI670475 | A/Alaska/97/2015(H3N2) | 2/7/2015 | GISAID |
| EPI670483 | A/Alaska/98/2015(H3N2) | 3/7/2015 | GISAID |

| | | | |
|---|---|---|---|
| EPI670491 | A/Alaska/99/2015(H3N2) | 3/7/2015 | GISAID |
| EPI670499 | A/Alaska/100/2015(H3N2) | 3/7/2015 | GISAID |
| EPI670507 | A/Alaska/101/2015(H3N2) | 4/7/2015 | GISAID |
| EPI670515 | A/Alaska/102/2015(H3N2) | 28/6/2015 | GISAID |
| EPI670523 | A/Alaska/103/2015(H3N2) | 4/7/2015 | GISAID |
| EPI670531 | A/Alaska/112/2015(H3N2) | 4/7/2015 | GISAID |
| EPI670539 | A/Alaska/113/2015(H3N2) | 10/7/2015 | GISAID |
| EPI670547 | A/Alaska/114/2015(H3N2) | 6/7/2015 | GISAID |
| EPI670555 | A/Alaska/115/2015(H3N2) | 4/7/2015 | GISAID |
| EPI670563 | A/Oregon/14/2015(H3N2) | 29/6/2015 | GISAID |
| EPI670571 | A/Acre/134806-IEC/2015(H3N2) | 16/4/2015 | GISAID |
| EPI670579 | A/Acre/135094-IEC/2015(H3N2) | 29/4/2015 | GISAID |
| EPI670587 | A/RioGrandeDoNorte/135219-IEC/2015(H3N2) | 13/5/2015 | GISAID |
| EPI670595 | A/Acre/135356-IEC/2015(H3N2) | 18/5/2015 | GISAID |
| EPI670603 | A/RioGrandeDoNorte/135497-IEC/2015(H3N2) | 29/5/2015 | GISAID |
| EPI670611 | A/PuertoRico/03/2015(H3N2) | 7/7/2015 | GISAID |
| EPI670619 | A/PuertoRico/09/2015(H3N2) | 11/8/2015 | GISAID |
| EPI670627 | A/FrenchGuiana/6101/2015(H3N2) | 5/3/2015 | GISAID |
| EPI670642 | A/Guadeloupe/9095/2015(H3N2) | 4/3/2015 | GISAID |
| EPI670650 | A/Guadeloupe/9096/2015(H3N2) | 24/2/2015 | GISAID |
| EPI670658 | A/Guadeloupe/9153/2015(H3N2) | 25/2/2015 | GISAID |
| EPI670670 | A/FrenchGuiana/8217/2015(H3N2) | 8/6/2015 | GISAID |
| EPI670676 | A/FrenchGuiana/9189/2015(H3N2) | 27/6/2015 | GISAID |
| EPI670691 | A/Guatemala/6315/2015(H3N2) | 25/6/2015 | GISAID |
| EPI670699 | A/Guatemala/6415/2015(H3N2) | 1/7/2015 | GISAID |
| EPI670707 | A/Guatemala/4153/2015(H3N2) | 7/3/2015 | GISAID |
| EPI670715 | A/Guatemala/5848/2015(H3N2) | 15/6/2015 | GISAID |
| EPI670723 | A/Guatemala/8802/2015(H3N2) | 23/3/2015 | GISAID |
| EPI670731 | A/Guatemala/9151/2015(H3N2) | 21/4/2015 | GISAID |
| EPI670745 | A/Brazil/4388/2015(H3N2) | 5/5/2015 | GISAID |
| EPI670753 | A/Brazil/0168/2015(H3N2) | 21/5/2015 | GISAID |
| EPI670761 | A/Brazil/1790/2015(H3N2) | 27/5/2015 | GISAID |
| EPI670769 | A/Brazil/2081/2015(H3N2) | 29/5/2015 | GISAID |
| EPI670777 | A/Brazil/2047/2015(H3N2) | 24/4/2015 | GISAID |
| EPI670785 | A/Brazil/7588/2015(H3N2) | 11/5/2015 | GISAID |
| EPI670797 | A/Tennessee/19/2015(H3N2) | 18/8/2015 | GISAID |
| EPI670812 | A/NewJersey/52/2015(H3N2) | 1/9/2015 | GISAID |

| | | | |
|---|---|---|---|
| EPI670820 | A/Alaska/206/2015(H3N2) | 7/9/2015 | GISAID |
| EPI670835 | A/Alaska/208/2015(H3N2) | 2/9/2015 | GISAID |
| EPI670843 | A/Ohio/27/2015(H3N2) | 17/9/2015 | GISAID |
| EPI670851 | A/Ohio/28/2015(H3N2) | 17/9/2015 | GISAID |
| EPI670859 | A/Ohio/29/2015(H3N2) | 17/9/2015 | GISAID |
| EPI670867 | A/Massachusetts/26/2015(H3N2) | 24/9/2015 | GISAID |
| EPI670875 | A/Kentucky/15/2015(H3N2) | 28/9/2015 | GISAID |
| EPI670886 | A/Bangladesh/7004/2015(H3N2) | 12/8/2015 | GISAID |
| EPI670894 | A/Bangladesh/7006/2015(H3N2) | 31/8/2015 | GISAID |
| EPI670902 | A/Bangladesh/3014/2015(H3N2) | 24/8/2015 | GISAID |
| EPI670910 | A/Bangladesh/8016/2015(H3N2) | 24/8/2015 | GISAID |
| EPI670918 | A/Bangladesh/1005/2015(H3N2) | 26/8/2015 | GISAID |
| EPI670926 | A/Bangladesh/803003/2015(H3N2) | 10/8/2015 | GISAID |
| EPI670934 | A/Bangladesh/3007/2015(H3N2) | 20/8/2015 | GISAID |
| EPI670942 | A/Bangladesh/9015/2015(H3N2) | 29/8/2015 | GISAID |
| EPI670950 | A/Florida/74/2015(H3N2) | 3/10/2015 | GISAID |
| EPI670958 | A/Florida/75/2015(H3N2) | 1/10/2015 | GISAID |
| EPI670966 | A/NorthCarolina/32/2015(H3N2) | 28/9/2015 | GISAID |
| EPI670974 | A/NorthCarolina/32/2015(H3N2) | 28/9/2015 | GISAID |
| EPI670982 | A/Maine/39/2015(H3N2) | 7/10/2015 | GISAID |
| EPI670990 | A/Maine/40/2015(H3N2) | 9/10/2015 | GISAID |
| EPI670998 | A/Maine/41/2015(H3N2) | 12/10/2015 | GISAID |
| EPI671006 | A/Missouri/41/2015(H3N2) | 4/10/2015 | GISAID |
| EPI671014 | A/Missouri/42/2015(H3N2) | 4/10/2015 | GISAID |
| EPI671022 | A/Montana/30/2015(H3N2) | 6/10/2015 | GISAID |
| EPI671030 | A/Montana/31/2015(H3N2) | 7/10/2015 | GISAID |
| EPI671038 | A/Montana/32/2015(H3N2) | 12/10/2015 | GISAID |
| EPI671046 | A/Arizona/20/2015(H3N2) | 4/10/2015 | GISAID |
| EPI671054 | A/Arizona/21/2015(H3N2) | 2/10/2015 | GISAID |
| EPI671062 | A/Arizona/22/2015(H3N2) | 7/10/2015 | GISAID |
| EPI671070 | A/Oregon/19/2015(H3N2) | 8/10/2015 | GISAID |
| EPI671078 | A/Oregon/20/2015(H3N2) | 9/10/2015 | GISAID |
| EPI671086 | A/Oregon/21/2015(H3N2) | 9/10/2015 | GISAID |
| EPI671094 | A/Oregon/22/2015(H3N2) | 9/10/2015 | GISAID |
| EPI671102 | A/Oregon/23/2015(H3N2) | 12/10/2015 | GISAID |
| EPI671110 | A/Oregon/24/2015(H3N2) | 12/10/2015 | GISAID |
| EPI671118 | A/Alaska/253/2015(H3N2) | 2/10/2015 | GISAID |

| | | | |
|---|---|---|---|
| EPI671126 | A/Alaska/254/2015(H3N2) | 5/10/2015 | GISAID |
| EPI671134 | A/Alaska/255/2015(H3N2) | 6/10/2015 | GISAID |
| EPI671142 | A/Alaska/256/2015(H3N2) | 5/10/2015 | GISAID |
| EPI671150 | A/Washington/40/2015(H3N2) | 1/9/2015 | GISAID |
| EPI671158 | A/Washington/41/2015(H3N2) | 31/8/2015 | GISAID |
| EPI671166 | A/Washington/42/2015(H3N2) | 31/8/2015 | GISAID |
| EPI671174 | A/Washington/43/2015(H3N2) | 31/8/2015 | GISAID |
| EPI671182 | A/Washington/44/2015(H3N2) | 7/9/2015 | GISAID |
| EPI671190 | A/Wyoming/18/2015(H3N2) | 22/9/2015 | GISAID |
| EPI671198 | A/Montana/28/2015(H3N2) | 24/9/2015 | GISAID |
| EPI671206 | A/Montana/29/2015(H3N2) | 26/9/2015 | GISAID |
| EPI671214 | A/Minnesota/42/2015(H3N2) | 14/9/2015 | GISAID |
| EPI671222 | A/Minnesota/43/2015(H3N2) | 17/9/2015 | GISAID |
| EPI671230 | A/Hawaii/58/2015(H3N2) | 15/9/2015 | GISAID |
| EPI671238 | A/Hawaii/60/2015(H3N2) | 26/9/2015 | GISAID |
| EPI671243 | A/Hawaii/61/2015(H3N2) | 28/9/2015 | GISAID |
| EPI671251 | A/Idaho/25/2015(H3N2) | 18/9/2015 | GISAID |
| EPI671266 | A/Texas/69/2015(H3N2) | 6/10/2015 | GISAID |
| EPI671273 | A/Texas/70/2015(H3N2) | 6/10/2015 | GISAID |
| EPI671281 | A/Vermont/08/2015(H3N2) | 19/8/2015 | GISAID |
| EPI671289 | A/Vermont/09/2015(H3N2) | 30/9/2015 | GISAID |
| EPI671297 | A/Pennsylvania/36/2015(H3N2) | 2/10/2015 | GISAID |
| EPI671305 | A/NewHampshire/41/2015(H3N2) | 10/10/2015 | GISAID |
| EPI673079 | A/KOBE/262/2015(H3N2) | 19/3/2015 | GISAID |
| EPI673591 | A/Bangladesh/5968/2015(H3N2) | 10/8/2015 | GISAID |
| EPI673599 | A/California/85/2015(H3N2) | 9/9/2015 | GISAID |
| EPI673607 | A/Bangladesh/22014/2015(H3N2) | 20/8/2015 | GISAID |
| EPI673615 | A/California/91/2015(H3N2) | 1/10/2015 | GISAID |
| EPI673623 | A/California/87/2015(H3N2) | 1/10/2015 | GISAID |
| EPI673631 | A/California/89/2015(H3N2) | 24/9/2015 | GISAID |
| EPI673639 | A/Guatemala/8147/2015(H3N2) | 19/3/2015 | GISAID |
| EPI673647 | A/California/83/2015(H3N2) | 10/9/2015 | GISAID |
| EPI673655 | A/California/86/2015(H3N2) | 5/9/2015 | GISAID |
| EPI673663 | A/Bangladesh/3937/2015(H3N2) | 17/8/2015 | GISAID |
| EPI673670 | A/Bangladesh/12004/2015(H3N2) | 11/8/2015 | GISAID |
| EPI673678 | A/SouthCarolina/09/2015(H3N2) | 2015 | GISAID |
| EPI673686 | A/Guatemala/363/2015(H3N2) | 16/4/2015 | GISAID |

| | | | |
|---|---|---|---|
| EPI673701 | A/Iowa/50/2015(H3N2) | 14/10/2015 | GISAID |
| EPI673709 | A/NewMexico/26/2015(H3N2) | 13/9/2015 | GISAID |
| EPI673724 | A/Delaware/41/2015(H3N2) | 14/10/2015 | GISAID |
| EPI673732 | A/Arkansas/09/2015(H3N2) | 12/10/2015 | GISAID |
| EPI673740 | A/Arkansas/08/2015(H3N2) | 12/10/2015 | GISAID |
| EPI673748 | A/NorthCarolina/33/2015(H3N2) | 16/10/2015 | GISAID |
| EPI673756 | A/Maine/43/2015(H3N2) | 19/10/2015 | GISAID |
| EPI673764 | A/Maine/42/2015(H3N2) | 15/10/2015 | GISAID |
| EPI673772 | A/Massachusetts/28/2015(H3N2) | 10/10/2015 | GISAID |
| EPI673784 | A/Mississippi/13/2015(H3N2) | 13/8/2015 | GISAID |
| EPI673792 | A/Alaska/116/2015(H3N2) | 6/7/2015 | GISAID |
| EPI673800 | A/Washington/49/2015(H3N2) | 4/10/2015 | GISAID |
| EPI673808 | A/Wisconsin/85/2015(H3N2) | 16/10/2015 | GISAID |
| EPI673816 | A/Washington/50/2015(H3N2) | 11/10/2015 | GISAID |
| EPI673824 | A/WestVirginia/23/2015(H3N2) | 8/10/2015 | GISAID |
| EPI673832 | A/Pennsylvania/42/2015(H3N2) | 19/10/2015 | GISAID |
| EPI673840 | A/Florida/81/2015(H3N2) | 5/10/2015 | GISAID |
| EPI673853 | A/Bangladesh/5011/2015(H3N2) | 29/8/2015 | GISAID |
| EPI673861 | A/Guatemala/9877/2015(H3N2) | 11/3/2015 | GISAID |
| EPI673869 | A/Hawaii/67/2015(H3N2) | 5/10/2015 | GISAID |
| EPI673877 | A/Hawaii/65/2015(H3N2) | 4/10/2015 | GISAID |
| EPI673885 | A/Hawaii/66/2015(H3N2) | 3/10/2015 | GISAID |
| EPI673896 | A/Hawaii/68/2015(H3N2) | 5/10/2015 | GISAID |
| EPI673904 | A/Guatemala/8136/2015(H3N2) | 9/3/2015 | GISAID |
| EPI674432 | A/Laos/I660/2015(H3N2) | 29/6/2015 | GISAID |
| EPI674440 | A/Laos/I706/2015(H3N2) | 7/7/2015 | GISAID |
| EPI674448 | A/Laos/I730/2015(H3N2) | 14/7/2015 | GISAID |
| EPI674456 | A/Laos/I751/2015(H3N2) | 22/7/2015 | GISAID |
| EPI674464 | A/Laos/I784/2015(H3N2) | 27/7/2015 | GISAID |
| EPI674472 | A/Laos/I805/2015(H3N2) | 3/8/2015 | GISAID |
| EPI674480 | A/Laos/I810/2015(H3N2) | 3/8/2015 | GISAID |
| EPI674488 | A/Laos/I847/2015(H3N2) | 10/8/2015 | GISAID |
| EPI674496 | A/Laos/I889/2015(H3N2) | 23/8/2015 | GISAID |
| EPI674504 | A/Laos/I917/2015(H3N2) | 2/9/2015 | GISAID |
| EPI674512 | A/Laos/I924/2015(H3N2) | 7/9/2015 | GISAID |
| EPI674520 | A/Laos/I929/2015(H3N2) | 9/9/2015 | GISAID |
| EPI674528 | A/Laos/I935/2015(H3N2) | 7/9/2015 | GISAID |

| | | | |
|---|---|---|---|
| EPI674536 | A/Laos/I936/2015(H3N2) | 7/9/2015 | GISAID |
| EPI674544 | A/Laos/I941/2015(H3N2) | 8/9/2015 | GISAID |
| EPI674560 | A/Laos/O587/2015(H3N2) | 21/8/2015 | GISAID |
| EPI674568 | A/Laos/SA358/2015(H3N2) | 22/8/2015 | GISAID |
| EPI674576 | A/Laos/SA373/2015(H3N2) | 7/9/2015 | GISAID |
| EPI674584 | A/Laos/SA380/2015(H3N2) | 7/9/2015 | GISAID |
| EPI674761 | A/Stockholm/54/2015(H3N2) | 30/10/2015 | GISAID |
| EPI675083 | A/Virginia/51/2014(H3N2) | 29/12/2014 | GISAID |
| EPI675090 | A/Argentina/116/2015(H3N2) | 15/7/2015 | GISAID |
| EPI675098 | A/Oklahoma/20/2015(H3N2) | 9/10/2015 | GISAID |
| EPI675118 | A/Peru/6915/2015(H3N2) | 25/8/2015 | GISAID |
| EPI675126 | A/Kyrgyzstan/82/2015(H3N2) | 12/2/2015 | GISAID |
| EPI675139 | A/Alaska/51/2015(H3N2) | 15/6/2015 | GISAID |
| EPI675153 | A/Alaska/58/2015(H3N2) | 17/6/2015 | GISAID |
| EPI675161 | A/Brazil/4390/2015(H3N2) | 5/5/2015 | GISAID |
| EPI675176 | A/Alaska/60/2015(H3N2) | 15/6/2015 | GISAID |
| EPI675184 | A/Brazil/2002/2015(H3N2) | 22/5/2015 | GISAID |
| EPI675192 | A/Michigan/62/2015(H3N2) | 29/9/2015 | GISAID |
| EPI675206 | A/Alaska/258/2015(H3N2) | 12/10/2015 | GISAID |
| EPI675214 | A/Alaska/259/2015(H3N2) | 20/10/2015 | GISAID |
| EPI675222 | A/Alaska/257/2015(H3N2) | 11/10/2015 | GISAID |
| EPI675230 | A/Alberta/RV2387/2015(H3N2) | 10/10/2015 | GISAID |
| EPI675238 | A/Indiana/28/2015(H3N2) | 28/9/2015 | GISAID |
| EPI675246 | A/Alberta/RV2382/2015(H3N2) | 1/10/2015 | GISAID |
| EPI675254 | A/Alberta/RV2386/2015(H3N2) | 10/10/2015 | GISAID |
| EPI675262 | A/Alberta/RV2385/2015(H3N2) | 10/10/2015 | GISAID |
| EPI675270 | A/BritishColumbia/RV2356/2015(H3N2) | 9/9/2015 | GISAID |
| EPI675278 | A/BritishColumbia/RV2357/2015(H3N2) | 11/9/2015 | GISAID |
| EPI675286 | A/BritishColumbia/RV2358/2015(H3N2) | 14/9/2015 | GISAID |
| EPI675294 | A/Alberta/RV2339/2015(H3N2) | 12/9/2015 | GISAID |
| EPI675302 | A/Kyrgyzstan/10/2015(H3N2) | 9/1/2015 | GISAID |
| EPI675391 | A/Alberta/RV2383/2015(H3N2) | 9/10/2015 | GISAID |
| EPI675398 | A/Colorado/16/2015(H3N2) | 21/10/2015 | GISAID |
| EPI675386 | A/Alberta/RV2338/2015(H3N2) | 9/9/2015 | GISAID |
| EPI675401 | A/Florida/80/2015(H3N2) | 28/10/2015 | GISAID |
| EPI675408 | A/Luxembourg/15008843/2015(H3N2) | 23/2/2015 | GISAID |
| EPI675417 | A/Texas/71/2015(H3N2) | 11/11/2015 | GISAID |

| | | | |
|---|---|---|---|
| EPI675425 | A/Maryland/18/2015(H3N2) | 29/10/2015 | GISAID |
| EPI675433 | A/Maryland/19/2015(H3N2) | 1/11/2015 | GISAID |
| EPI675444 | A/Louisiana/07/2015(H3N2) | 11/11/2015 | GISAID |
| EPI675452 | A/Washington/55/2015(H3N2) | 7/10/2015 | GISAID |
| EPI675460 | A/Washington/56/2015(H3N2) | 20/10/2015 | GISAID |
| EPI675468 | A/Wisconsin/86/2015(H3N2) | 19/10/2015 | GISAID |
| EPI675489 | A/Argentina/149/2015(H3N2) | 23/7/2015 | GISAID |
| EPI675497 | A/Argentina/156/2015(H3N2) | 25/7/2015 | GISAID |
| EPI675512 | A/Argentina/157/2015(H3N2) | 27/7/2015 | GISAID |
| EPI675520 | A/Argentina/158/2015(H3N2) | 27/7/2015 | GISAID |
| EPI675792 | A/NewZealand/1703/2015(H3N2) | 13/8/2015 | GISAID |
| EPI676550 | A/Alberta/RV2362/2015(H3N2) | 18/9/2015 | GISAID |
| EPI676558 | A/Alberta/RV2363/2015(H3N2) | 18/9/2015 | GISAID |
| EPI676566 | A/Ontario/RV2371/2015(H3N2) | 17/9/2015 | GISAID |
| EPI676574 | A/Alberta/RV2372/2015(H3N2) | 29/9/2015 | GISAID |
| EPI676582 | A/Alberta/RV2373/2015(H3N2) | 28/9/2015 | GISAID |
| EPI676596 | A/Ontario/RV2377/2015(H3N2) | 7/10/2015 | GISAID |
| EPI676604 | A/Ontario/RV2377/2015(H3N2) | 7/10/2015 | GISAID |
| EPI676612 | A/Ontario/RV2378/2015(H3N2) | 7/10/2015 | GISAID |
| EPI676620 | A/Ontario/RV2378/2015(H3N2) | 7/10/2015 | GISAID |
| EPI676628 | A/Ontario/RV2379/2015(H3N2) | 7/10/2015 | GISAID |
| EPI676636 | A/Ontario/RV2379/2015(H3N2) | 7/10/2015 | GISAID |
| EPI676644 | A/Ontario/RV2380/2015(H3N2) | 7/10/2015 | GISAID |
| EPI676652 | A/Ontario/RV2380/2015(H3N2) | 7/10/2015 | GISAID |
| EPI676660 | A/SouthDakota/36/2015(H3N2) | 1/8/2015 | GISAID |
| EPI676668 | A/SouthDakota/37/2015(H3N2) | 11/8/2015 | GISAID |
| EPI676676 | A/SouthDakota/39/2015(H3N2) | 13/10/2015 | GISAID |
| EPI676684 | A/NewYork/71/2015(H3N2) | 3/11/2015 | GISAID |
| EPI676692 | A/Florida/84/2015(H3N2) | 6/11/2015 | GISAID |
| EPI676700 | A/Oregon/25/2015(H3N2) | 19/10/2015 | GISAID |
| EPI676708 | A/Oregon/25/2015(H3N2) | 19/10/2015 | GISAID |
| EPI676716 | A/Oregon/27/2015(H3N2) | 23/10/2015 | GISAID |
| EPI676724 | A/PuertoRico/02/2015(H3N2) | 3/7/2015 | GISAID |
| EPI676732 | A/Argentina/163/2015(H3N2) | 24/7/2015 | GISAID |
| EPI676745 | A/Argentina/169/2015(H3N2) | 24/7/2015 | GISAID |
| EPI676758 | A/Argentina/181/2015(H3N2) | 28/7/2015 | GISAID |
| EPI676766 | A/Nebraska/18/2015(H3N2) | 4/9/2015 | GISAID |

| | | | |
|---|---|---|---|
| EPI676774 | A/Alaska/210/2015(H3N2) | 7/9/2015 | GISAID |
| EPI676788 | A/Alaska/212/2015(H3N2) | 30/8/2015 | GISAID |
| EPI676796 | A/Alaska/213/2015(H3N2) | 30/8/2015 | GISAID |
| EPI676804 | A/Alaska/214/2015(H3N2) | 31/8/2015 | GISAID |
| EPI676812 | A/Alaska/215/2015(H3N2) | 4/9/2015 | GISAID |
| EPI676820 | A/Washington/45/2015(H3N2) | 16/6/2015 | GISAID |
| EPI676828 | A/Washington/47/2015(H3N2) | 19/9/2015 | GISAID |
| EPI676836 | A/Washington/48/2015(H3N2) | 19/9/2015 | GISAID |
| EPI676844 | A/Liaoning-Baita/1293/2015(H3N2) | 6/5/2015 | GISAID |
| EPI676857 | A/Hunan-Wuling/11088/2015(H3N2) | 15/6/2015 | GISAID |
| EPI676865 | A/Jilin-Yanji/1179/2015(H3N2) | 3/3/2015 | GISAID |
| EPI676873 | A/Arizona/15/2015(H3N2) | 20/9/2015 | GISAID |
| EPI676881 | A/Arizona/16/2015(H3N2) | 28/9/2015 | GISAID |
| EPI676889 | A/Arizona/17/2015(H3N2) | 28/9/2015 | GISAID |
| EPI676897 | A/Arizona/18/2015(H3N2) | 30/9/2015 | GISAID |
| EPI676905 | A/California/84/2015(H3N2) | 30/8/2015 | GISAID |
| EPI676913 | A/Georgia/12/2015(H3N2) | 2015 | GISAID |
| EPI676921 | A/Georgia/13/2015(H3N2) | 9/10/2015 | GISAID |
| EPI676929 | A/Florida/76/2015(H3N2) | 20/10/2015 | GISAID |
| EPI676937 | A/Florida/76/2015(H3N2) | 20/10/2015 | GISAID |
| EPI676945 | A/Florida/77/2015(H3N2) | 24/10/2015 | GISAID |
| EPI676953 | A/Florida/77/2015(H3N2) | 24/10/2015 | GISAID |
| EPI676961 | A/Florida/78/2015(H3N2) | 22/10/2015 | GISAID |
| EPI676969 | A/Iowa/44/2015(H3N2) | 21/10/2015 | GISAID |
| EPI676977 | A/Iowa/44/2015(H3N2) | 21/10/2015 | GISAID |
| EPI676985 | A/Iowa/45/2015(H3N2) | 22/10/2015 | GISAID |
| EPI676993 | A/Iowa/45/2015(H3N2) | 22/10/2015 | GISAID |
| EPI677001 | A/Iowa/46/2015(H3N2) | 21/10/2015 | GISAID |
| EPI677009 | A/Iowa/46/2015(H3N2) | 21/10/2015 | GISAID |
| EPI677017 | A/Iowa/47/2015(H3N2) | 22/10/2015 | GISAID |
| EPI677025 | A/Iowa/48/2015(H3N2) | 24/10/2015 | GISAID |
| EPI677033 | A/Iowa/49/2015(H3N2) | 24/10/2015 | GISAID |
| EPI677041 | A/Alaska/216/2015(H3N2) | 6/9/2015 | GISAID |
| EPI677049 | A/Alaska/217/2015(H3N2) | 7/9/2015 | GISAID |
| EPI677057 | A/Alaska/218/2015(H3N2) | 7/9/2015 | GISAID |
| EPI677065 | A/Alaska/219/2015(H3N2) | 5/9/2015 | GISAID |
| EPI677073 | A/Alaska/220/2015(H3N2) | 4/9/2015 | GISAID |

| | | | |
|---|---|---|---|
| EPI677081 | A/Alaska/221/2015(H3N2) | 3/9/2015 | GISAID |
| EPI677089 | A/Alaska/222/2015(H3N2) | 8/9/2015 | GISAID |
| EPI677118 | A/Alaska/227/2015(H3N2) | 11/9/2015 | GISAID |
| EPI677126 | A/Alaska/228/2015(H3N2) | 12/9/2015 | GISAID |
| EPI677134 | A/Alaska/229/2015(H3N2) | 13/9/2015 | GISAID |
| EPI677142 | A/Alaska/230/2015(H3N2) | 14/9/2015 | GISAID |
| EPI677150 | A/Alaska/231/2015(H3N2) | 15/9/2015 | GISAID |
| EPI677158 | A/Alaska/232/2015(H3N2) | 9/9/2015 | GISAID |
| EPI677178 | A/Alaska/235/2015(H3N2) | 16/9/2015 | GISAID |
| EPI677186 | A/Alaska/236/2015(H3N2) | 16/9/2015 | GISAID |
| EPI677194 | A/Alaska/237/2015(H3N2) | 12/9/2015 | GISAID |
| EPI677202 | A/Alaska/238/2015(H3N2) | 16/9/2015 | GISAID |
| EPI677210 | A/Alaska/239/2015(H3N2) | 16/9/2015 | GISAID |
| EPI677218 | A/Alaska/240/2015(H3N2) | 15/9/2015 | GISAID |
| EPI677226 | A/Alaska/241/2015(H3N2) | 16/9/2015 | GISAID |
| EPI677234 | A/Alaska/243/2015(H3N2) | 14/9/2015 | GISAID |
| EPI677242 | A/Alaska/244/2015(H3N2) | 14/9/2015 | GISAID |
| EPI677250 | A/Alaska/245/2015(H3N2) | 10/9/2015 | GISAID |
| EPI677258 | A/Alaska/246/2015(H3N2) | 13/9/2015 | GISAID |
| EPI677266 | A/Alaska/247/2015(H3N2) | 17/9/2015 | GISAID |
| EPI677274 | A/Alaska/249/2015(H3N2) | 17/9/2015 | GISAID |
| EPI677282 | A/Hawaii/62/2015(H3N2) | 28/9/2015 | GISAID |
| EPI677290 | A/Hawaii/63/2015(H3N2) | 28/9/2015 | GISAID |
| EPI677305 | A/Michigan/58/2015(H3N2) | 4/10/2015 | GISAID |
| EPI677313 | A/Hawaii/69/2015(H3N2) | 4/10/2015 | GISAID |
| EPI677321 | A/Hawaii/69/2015(H3N2) | 4/10/2015 | GISAID |
| EPI677329 | A/Hawaii/70/2015(H3N2) | 5/10/2015 | GISAID |
| EPI677337 | A/Hawaii/70/2015(H3N2) | 5/10/2015 | GISAID |
| EPI677345 | A/Hawaii/71/2015(H3N2) | 9/10/2015 | GISAID |
| EPI677353 | A/Colorado/15/2015(H3N2) | 11/10/2015 | GISAID |
| EPI677361 | A/Alaska/260/2015(H3N2) | 22/10/2015 | GISAID |
| EPI677369 | A/Alaska/261/2015(H3N2) | 26/10/2015 | GISAID |
| EPI677377 | A/Alaska/262/2015(H3N2) | 25/10/2015 | GISAID |
| EPI677385 | A/Alaska/262/2015(H3N2) | 25/10/2015 | GISAID |
| EPI677393 | A/Washington/53/2015(H3N2) | 17/10/2015 | GISAID |
| EPI677401 | A/Washington/54/2015(H3N2) | 17/10/2015 | GISAID |
| EPI677414 | A/WestVirginia/24/2015(H3N2) | 19/10/2015 | GISAID |

| | | | |
|---|---|---|---|
| EPI677422 | A/Hawaii/72/2015(H3N2) | 8/10/2015 | GISAID |
| EPI677430 | A/Hawaii/72/2015(H3N2) | 8/10/2015 | GISAID |
| EPI677438 | A/Hawaii/73/2015(H3N2) | 12/10/2015 | GISAID |
| EPI677446 | A/Hawaii/74/2015(H3N2) | 9/10/2015 | GISAID |
| EPI677454 | A/Hawaii/75/2015(H3N2) | 17/10/2015 | GISAID |
| EPI677462 | A/Florida/82/2015(H3N2) | 26/10/2015 | GISAID |
| EPI677470 | A/Florida/82/2015(H3N2) | 26/10/2015 | GISAID |
| EPI677478 | A/Wisconsin/87/2015(H3N2) | 29/10/2015 | GISAID |
| EPI677486 | A/Louisiana/06/2015(H3N2) | 2/11/2015 | GISAID |
| EPI677494 | A/Florida/85/2015(H3N2) | 2/11/2015 | GISAID |
| EPI677502 | A/Florida/86/2015(H3N2) | 2/11/2015 | GISAID |
| EPI677510 | A/Florida/87/2015(H3N2) | 8/11/2015 | GISAID |
| EPI677518 | A/Minnesota/47/2015(H3N2) | 16/11/2015 | GISAID |
| EPI677526 | A/SouthCarolina/10/2015(H3N2) | 6/10/2015 | GISAID |
| EPI677534 | A/SouthCarolina/10/2015(H3N2) | 6/10/2015 | GISAID |
| EPI677542 | A/SouthCarolina/11/2015(H3N2) | 27/10/2015 | GISAID |
| EPI677550 | A/Wyoming/19/2015(H3N2) | 3/11/2015 | GISAID |
| EPI677558 | A/California/93/2015(H3N2) | 20/10/2015 | GISAID |
| EPI677566 | A/California/94/2015(H3N2) | 20/10/2015 | GISAID |
| EPI677574 | A/California/95/2015(H3N2) | 20/10/2015 | GISAID |
| EPI677582 | A/California/98/2015(H3N2) | 19/10/2015 | GISAID |
| EPI677590 | A/Washington/57/2015(H3N2) | 7/11/2015 | GISAID |
| EPI677598 | A/Washington/57/2015(H3N2) | 7/11/2015 | GISAID |
| EPI677610 | A/Washington/58/2015(H3N2) | 9/11/2015 | GISAID |
| EPI677618 | A/Michigan/66/2015(H3N2) | 15/10/2015 | GISAID |
| EPI677626 | A/Michigan/67/2015(H3N2) | 10/10/2015 | GISAID |
| EPI679143 | A/England/53340242/2015(H3N2) | 1/8/2015 | GISAID |
| EPI679164 | A/England/334/2015(H3N2) | 6/9/2015 | GISAID |
| EPI679172 | A/England/53860135/2015(H3N2) | 14/9/2015 | GISAID |
| EPI679229 | A/England/54380710/2015(H3N2) | 2/10/2015 | GISAID |
| EPI679274 | A/England/54760378/2015(H3N2) | 16/11/2015 | GISAID |
| EPI679342 | A/Alaska/104/2015(H3N2) | 4/7/2015 | GISAID |
| EPI679350 | A/Alaska/105/2015(H3N2) | 4/7/2015 | GISAID |
| EPI679358 | A/Alaska/106/2015(H3N2) | 27/6/2015 | GISAID |
| EPI679366 | A/Alaska/107/2015(H3N2) | 3/7/2015 | GISAID |
| EPI679375 | A/Alaska/108/2015(H3N2) | 6/7/2015 | GISAID |
| EPI679383 | A/Alaska/109/2015(H3N2) | 7/7/2015 | GISAID |

| | | | |
|---|---|---|---|
| EPI679398 | A/Alaska/111/2015(H3N2) | 9/7/2015 | GISAID |
| EPI679406 | A/Alaska/248/2015(H3N2) | 19/9/2015 | GISAID |
| EPI679414 | A/Alaska/251/2015(H3N2) | 18/9/2015 | GISAID |
| EPI679422 | A/Alberta/RV2372/2015(H3N2) | 29/9/2015 | GISAID |
| EPI679430 | A/California/96/2015(H3N2) | 21/10/2015 | GISAID |
| EPI679438 | A/California/97/2015(H3N2) | 24/10/2015 | GISAID |
| EPI679446 | A/Florida/79/2015(H3N2) | 26/10/2015 | GISAID |
| EPI679454 | A/HongKong/15590/2015(H3N2) | 9/10/2015 | GISAID |
| EPI679463 | A/HongKong/15607/2015(H3N2) | 14/10/2015 | GISAID |
| EPI679471 | A/HongKong/15609/2015(H3N2) | 17/10/2015 | GISAID |
| EPI679484 | A/Iowa/54/2015(H3N2) | 1/11/2015 | GISAID |
| EPI679492 | A/Laos/438/2015(H3N2) | 27/6/2015 | GISAID |
| EPI679500 | A/Laos/769/2015(H3N2) | 27/7/2015 | GISAID |
| EPI679508 | A/Laos/794/2015(H3N2) | 5/8/2015 | GISAID |
| EPI679514 | A/Montana/24/2015(H3N2) | 18/9/2015 | GISAID |
| EPI679527 | A/NewYork/67/2015(H3N2) | 6/9/2015 | GISAID |
| EPI679540 | A/NewYork/69/2015(H3N2) | 12/10/2015 | GISAID |
| EPI679549 | A/Oklahoma/21/2015(H3N2) | 18/11/2015 | GISAID |
| EPI679557 | A/Pernambuco/135596-IEC/2015(H3N2) | 28/5/2015 | GISAID |
| EPI679565 | A/Pernambuco/135597-IEC/2015(H3N2) | 1/6/2015 | GISAID |
| EPI679573 | A/Pernambuco/136136-IEC/2015(H3N2) | 22/6/2015 | GISAID |
| EPI679585 | A/Peru/69/2015(H3N2) | 21/7/2015 | GISAID |
| EPI679598 | A/PuertoRico/04/2015(H3N2) | 13/7/2015 | GISAID |
| EPI679610 | A/PuertoRico/06/2015(H3N2) | 21/7/2015 | GISAID |
| EPI679618 | A/PuertoRico/07/2015(H3N2) | 7/8/2015 | GISAID |
| EPI679626 | A/PuertoRico/08/2015(H3N2) | 10/8/2015 | GISAID |
| EPI679634 | A/PuertoRico/10/2015(H3N2) | 16/8/2015 | GISAID |
| EPI679650 | A/PuertoRico/12/2015(H3N2) | 24/8/2015 | GISAID |
| EPI679658 | A/PuertoRico/13/2015(H3N2) | 28/8/2015 | GISAID |
| EPI679672 | A/PuertoRico/15/2015(H3N2) | 30/8/2015 | GISAID |
| EPI679680 | A/PuertoRico/16/2015(H3N2) | 3/9/2015 | GISAID |
| EPI679694 | A/PuertoRico/20/2015(H3N2) | 9/9/2015 | GISAID |
| EPI679702 | A/PuertoRico/22/2015(H3N2) | 9/9/2015 | GISAID |
| EPI679710 | A/PuertoRico/23/2015(H3N2) | 12/9/2015 | GISAID |
| EPI679718 | A/PuertoRico/24/2015(H3N2) | 12/9/2015 | GISAID |
| EPI679726 | A/PuertoRico/25/2015(H3N2) | 11/9/2015 | GISAID |
| EPI679734 | A/PuertoRico/26/2015(H3N2) | 10/8/2015 | GISAID |

| | | | |
|---|---|---|---|
| EPI679749 | A/SouthDakota/38/2015(H3N2) | 9/9/2015 | GISAID |
| EPI679757 | A/Texas/64/2015(H3N2) | 1/7/2015 | GISAID |
| EPI679765 | A/Washington/59/2015(H3N2) | 21/10/2015 | GISAID |
| EPI679896 | A/Alaska/163/2015(H3N2) | 4/8/2015 | GISAID |
| EPI679904 | A/Alaska/164/2015(H3N2) | 3/8/2015 | GISAID |
| EPI679912 | A/Alaska/165/2015(H3N2) | 19/8/2015 | GISAID |
| EPI679920 | A/Alaska/166/2015(H3N2) | 15/8/2015 | GISAID |
| EPI679928 | A/Alaska/167/2015(H3N2) | 15/8/2015 | GISAID |
| EPI679936 | A/Alaska/168/2015(H3N2) | 15/8/2015 | GISAID |
| EPI679944 | A/Alaska/170/2015(H3N2) | 21/8/2015 | GISAID |
| EPI679952 | A/Alaska/171/2015(H3N2) | 25/8/2015 | GISAID |
| EPI679960 | A/Alaska/172/2015(H3N2) | 26/8/2015 | GISAID |
| EPI679968 | A/Alaska/174/2015(H3N2) | 30/8/2015 | GISAID |
| EPI679981 | A/Alaska/176/2015(H3N2) | 19/8/2015 | GISAID |
| EPI679989 | A/Alaska/177/2015(H3N2) | 21/8/2015 | GISAID |
| EPI679997 | A/Alaska/178/2015(H3N2) | 18/8/2015 | GISAID |
| EPI680005 | A/Alaska/209/2015(H3N2) | 7/9/2015 | GISAID |
| EPI680013 | A/Arizona/19/2015(H3N2) | 29/9/2015 | GISAID |
| EPI680021 | A/Arizona/23/2015(H3N2) | 26/10/2015 | GISAID |
| EPI680029 | A/Arizona/24/2015(H3N2) | 2/11/2015 | GISAID |
| EPI680037 | A/Arizona/25/2015(H3N2) | 2/11/2015 | GISAID |
| EPI680045 | A/Arizona/27/2015(H3N2) | 18/10/2015 | GISAID |
| EPI680053 | A/California/107/2015(H3N2) | 18/10/2015 | GISAID |
| EPI680061 | A/Colorado/21/2015(H3N2) | 19/11/2015 | GISAID |
| EPI680069 | A/Colorado/23/2015(H3N2) | 22/11/2015 | GISAID |
| EPI680077 | A/Colorado/24/2015(H3N2) | 17/11/2015 | GISAID |
| EPI680085 | A/Hawaii/64/2015(H3N2) | 28/9/2015 | GISAID |
| EPI680093 | A/Hawaii/77/2015(H3N2) | 2/11/2015 | GISAID |
| EPI680106 | A/Nigeria/253/2015(H3N2) | 8/9/2015 | GISAID |
| EPI680114 | A/Texas/72/2015(H3N2) | 8/11/2015 | GISAID |
| EPI680122 | A/Wisconsin/86/2015(H3N2) | 19/10/2015 | GISAID |
| EPI680131 | A/Alaska/179/2015(H3N2) | 16/8/2015 | GISAID |
| EPI680139 | A/Alaska/184/2015(H3N2) | 20/8/2015 | GISAID |
| EPI680147 | A/Alaska/187/2015(H3N2) | 19/8/2015 | GISAID |
| EPI680155 | A/Alaska/188/2015(H3N2) | 15/8/2015 | GISAID |
| EPI680163 | A/Alaska/189/2015(H3N2) | 13/8/2015 | GISAID |
| EPI680177 | A/Alaska/191/2015(H3N2) | 18/8/2015 | GISAID |

| | | | |
|---|---|---|---|
| EPI680185 | A/Alaska/192/2015(H3N2) | 16/8/2015 | GISAID |
| EPI680193 | A/Alaska/193/2015(H3N2) | 17/8/2015 | GISAID |
| EPI680201 | A/Bangladesh/3937/2015(H3N2) | 17/8/2015 | GISAID |
| EPI680209 | A/Florida/80/2015(H3N2) | 28/10/2015 | GISAID |
| EPI680217 | A/Idaho/25/2015(H3N2) | 18/9/2015 | GISAID |
| EPI680225 | A/Michigan/53/2015(H3N2) | 12/9/2015 | GISAID |
| EPI680233 | A/Michigan/54/2015(H3N2) | 19/9/2015 | GISAID |
| EPI680241 | A/Michigan/55/2015(H3N2) | 18/9/2015 | GISAID |
| EPI680249 | A/Michigan/60/2015(H3N2) | 5/10/2015 | GISAID |
| EPI680257 | A/NewMexico/26/2015(H3N2) | 13/9/2015 | GISAID |
| EPI680265 | A/Nigeria/299/2015(H3N2) | 21/9/2015 | GISAID |
| EPI680273 | A/Nigeria/300/2015(H3N2) | 22/9/2015 | GISAID |
| EPI680281 | A/Vermont/08/2015(H3N2) | 19/8/2015 | GISAID |
| EPI683254 | A/Laos/648/2015(H3N2) | 22/6/2015 | GISAID |
| EPI683262 | A/Laos/738/2015(H3N2) | 15/7/2015 | GISAID |
| EPI683278 | A/Alaska/49/2015(H3N2) | 15/6/2015 | GISAID |
| EPI683286 | A/Alaska/49/2015(H3N2) | 15/6/2015 | GISAID |
| EPI683294 | A/Alaska/50/2015(H3N2) | 16/6/2015 | GISAID |
| EPI683302 | A/Alaska/50/2015(H3N2) | 16/6/2015 | GISAID |
| EPI683310 | A/St.VincentAndGrenadines/423/2015(H3N2) | 12/2/2015 | GISAID |
| EPI683318 | A/Victoria/511/2015(H3N2) | 16/5/2015 | GISAID |
| EPI683326 | A/Victoria/511/2015(H3N2) | 16/5/2015 | GISAID |
| EPI683334 | A/SouthAustralia/21/2015(H3N2) | 13/3/2015 | GISAID |
| EPI683348 | A/Alaska/151/2015(H3N2) | 21/7/2015 | GISAID |
| EPI683356 | A/Alaska/151/2015(H3N2) | 21/7/2015 | GISAID |
| EPI683364 | A/Alaska/152/2015(H3N2) | 20/7/2015 | GISAID |
| EPI683372 | A/Alaska/152/2015(H3N2) | 20/7/2015 | GISAID |
| EPI683387 | A/Alaska/153/2015(H3N2) | 3/8/2015 | GISAID |
| EPI683395 | A/Alaska/154/2015(H3N2) | 24/7/2015 | GISAID |
| EPI683403 | A/Alaska/154/2015(H3N2) | 24/7/2015 | GISAID |
| EPI683411 | A/Alaska/155/2015(H3N2) | 8/8/2015 | GISAID |
| EPI683419 | A/Alaska/155/2015(H3N2) | 8/8/2015 | GISAID |
| EPI683427 | A/Alaska/156/2015(H3N2) | 3/8/2015 | GISAID |
| EPI683435 | A/Alaska/156/2015(H3N2) | 3/8/2015 | GISAID |
| EPI683443 | A/Blagoveshchensk/19/2015(H3N2) | 2/3/2015 | GISAID |
| EPI683451 | A/Blagoveshchensk/19/2015(H3N2) | 2/3/2015 | GISAID |
| EPI683459 | A/Irkutsk/88/2015(H3N2) | 17/2/2015 | GISAID |

| | | | |
|---|---|---|---|
| EPI683467 | A/Irkutsk/88/2015(H3N2) | 17/2/2015 | GISAID |
| EPI683475 | A/Krasnoyarsk/324/2015(H3N2) | 4/2/2015 | GISAID |
| EPI683489 | A/Yekaterinburg/239/2015(H3N2) | 5/3/2015 | GISAID |
| EPI683497 | A/Yekaterinburg/239/2015(H3N2) | 5/3/2015 | GISAID |
| EPI683505 | A/Alaska/157/2015(H3N2) | 13/8/2015 | GISAID |
| EPI683513 | A/Alaska/157/2015(H3N2) | 13/8/2015 | GISAID |
| EPI683521 | A/Alaska/158/2015(H3N2) | 14/8/2015 | GISAID |
| EPI683529 | A/Alaska/158/2015(H3N2) | 14/8/2015 | GISAID |
| EPI683537 | A/Alaska/159/2015(H3N2) | 17/8/2015 | GISAID |
| EPI683545 | A/Alaska/159/2015(H3N2) | 17/8/2015 | GISAID |
| EPI683553 | A/Alaska/162/2015(H3N2) | 19/8/2015 | GISAID |
| EPI683561 | A/Alaska/162/2015(H3N2) | 19/8/2015 | GISAID |
| EPI683569 | A/Argentina/23/2015(H3N2) | 15/6/2015 | GISAID |
| EPI683577 | A/Argentina/23/2015(H3N2) | 15/6/2015 | GISAID |
| EPI683585 | A/Argentina/24/2015(H3N2) | 15/6/2015 | GISAID |
| EPI683593 | A/Argentina/24/2015(H3N2) | 15/6/2015 | GISAID |
| EPI683601 | A/Argentina/25/2015(H3N2) | 15/6/2015 | GISAID |
| EPI683609 | A/Argentina/25/2015(H3N2) | 15/6/2015 | GISAID |
| EPI683617 | A/Argentina/103/2015(H3N2) | 30/6/2015 | GISAID |
| EPI683625 | A/Argentina/103/2015(H3N2) | 30/6/2015 | GISAID |
| EPI683633 | A/Argentina/104/2015(H3N2) | 10/7/2015 | GISAID |
| EPI683641 | A/Argentina/104/2015(H3N2) | 10/7/2015 | GISAID |
| EPI683649 | A/Argentina/11592/2015(H3N2) | 22/7/2015 | GISAID |
| EPI683657 | A/Argentina/11592/2015(H3N2) | 22/7/2015 | GISAID |
| EPI683665 | A/Argentina/11594/2015(H3N2) | 13/7/2015 | GISAID |
| EPI683673 | A/Argentina/11594/2015(H3N2) | 13/7/2015 | GISAID |
| EPI683681 | A/Argentina/11596/2015(H3N2) | 13/7/2015 | GISAID |
| EPI683689 | A/Argentina/11596/2015(H3N2) | 13/7/2015 | GISAID |
| EPI683697 | A/Argentina/11611/2015(H3N2) | 27/7/2015 | GISAID |
| EPI683705 | A/Argentina/11611/2015(H3N2) | 27/7/2015 | GISAID |
| EPI683713 | A/Argentina/11628/2015(H3N2) | 30/7/2015 | GISAID |
| EPI683721 | A/Argentina/11628/2015(H3N2) | 30/7/2015 | GISAID |
| EPI683729 | A/Argentina/11639/2015(H3N2) | 11/8/2015 | GISAID |
| EPI683742 | A/Argentina/11640/2015(H3N2) | 11/8/2015 | GISAID |
| EPI683750 | A/Argentina/11640/2015(H3N2) | 11/8/2015 | GISAID |
| EPI683758 | A/Argentina/11652/2015(H3N2) | 10/8/2015 | GISAID |
| EPI683766 | A/Argentina/11652/2015(H3N2) | 10/8/2015 | GISAID |

| | | | |
|---|---|---|---|
| EPI683774 | A/Argentina/11656/2015(H3N2) | 11/8/2015 | GISAID |
| EPI683782 | A/Argentina/11656/2015(H3N2) | 11/8/2015 | GISAID |
| EPI683790 | A/Argentina/11659/2015(H3N2) | 12/8/2015 | GISAID |
| EPI683798 | A/Argentina/11659/2015(H3N2) | 12/8/2015 | GISAID |
| EPI683806 | A/Massachusetts/25/2015(H3N2) | 10/9/2015 | GISAID |
| EPI683814 | A/Massachusetts/25/2015(H3N2) | 10/9/2015 | GISAID |
| EPI683822 | A/Indiana/17/2015(H3N2) | 31/8/2015 | GISAID |
| EPI683830 | A/Indiana/17/2015(H3N2) | 31/8/2015 | GISAID |
| EPI683838 | A/Indiana/19/2015(H3N2) | 1/9/2015 | GISAID |
| EPI683846 | A/Indiana/19/2015(H3N2) | 1/9/2015 | GISAID |
| EPI683854 | A/Indiana/20/2015(H3N2) | 1/9/2015 | GISAID |
| EPI683862 | A/Indiana/20/2015(H3N2) | 1/9/2015 | GISAID |
| EPI683870 | A/Indiana/21/2015(H3N2) | 1/9/2015 | GISAID |
| EPI683878 | A/Indiana/21/2015(H3N2) | 1/9/2015 | GISAID |
| EPI683886 | A/Indiana/22/2015(H3N2) | 1/9/2015 | GISAID |
| EPI683894 | A/Indiana/22/2015(H3N2) | 1/9/2015 | GISAID |
| EPI683909 | A/Indiana/24/2015(H3N2) | 3/9/2015 | GISAID |
| EPI683917 | A/Indiana/25/2015(H3N2) | 3/9/2015 | GISAID |
| EPI683925 | A/Indiana/25/2015(H3N2) | 3/9/2015 | GISAID |
| EPI683933 | A/NewHampshire/39/2015(H3N2) | 9/9/2015 | GISAID |
| EPI683941 | A/NewHampshire/39/2015(H3N2) | 9/9/2015 | GISAID |
| EPI683949 | A/NewHampshire/40/2015(H3N2) | 14/9/2015 | GISAID |
| EPI683957 | A/NewHampshire/40/2015(H3N2) | 14/9/2015 | GISAID |
| EPI683965 | A/Minnesota/41/2015(H3N2) | 14/9/2015 | GISAID |
| EPI683973 | A/Minnesota/41/2015(H3N2) | 14/9/2015 | GISAID |
| EPI683981 | A/Florida/69/2015(H3N2) | 15/9/2015 | GISAID |
| EPI683989 | A/Florida/69/2015(H3N2) | 15/9/2015 | GISAID |
| EPI683997 | A/Florida/70/2015(H3N2) | 15/9/2015 | GISAID |
| EPI684005 | A/Florida/70/2015(H3N2) | 15/9/2015 | GISAID |
| EPI684013 | A/Florida/71/2015(H3N2) | 15/9/2015 | GISAID |
| EPI684021 | A/Florida/71/2015(H3N2) | 15/9/2015 | GISAID |
| EPI684029 | A/Florida/72/2015(H3N2) | 15/9/2015 | GISAID |
| EPI684037 | A/Florida/72/2015(H3N2) | 15/9/2015 | GISAID |
| EPI684045 | A/Wisconsin/80/2015(H3N2) | 14/9/2015 | GISAID |
| EPI684053 | A/Wisconsin/80/2015(H3N2) | 14/9/2015 | GISAID |
| EPI684061 | A/Wisconsin/81/2015(H3N2) | 18/9/2015 | GISAID |
| EPI684069 | A/Wisconsin/81/2015(H3N2) | 18/9/2015 | GISAID |

| | | | |
|---|---|---|---|
| EPI684077 | A/Wisconsin/82/2015(H3N2) | 17/9/2015 | GISAID |
| EPI684085 | A/Wisconsin/82/2015(H3N2) | 17/9/2015 | GISAID |
| EPI684093 | A/Tennessee/21/2015(H3N2) | 15/9/2015 | GISAID |
| EPI684101 | A/Tennessee/21/2015(H3N2) | 15/9/2015 | GISAID |
| EPI684109 | A/Ohio/27/2015(H3N2) | 17/9/2015 | GISAID |
| EPI684117 | A/Ohio/28/2015(H3N2) | 17/9/2015 | GISAID |
| EPI684131 | A/Massachusetts/26/2015(H3N2) | 24/9/2015 | GISAID |
| EPI684139 | A/NewMexico/23/2015(H3N2) | 12/9/2015 | GISAID |
| EPI684147 | A/NewMexico/23/2015(H3N2) | 12/9/2015 | GISAID |
| EPI684155 | A/NewMexico/25/2015(H3N2) | 10/9/2015 | GISAID |
| EPI684163 | A/NewMexico/25/2015(H3N2) | 10/9/2015 | GISAID |
| EPI684171 | A/Iowa/40/2015(H3N2) | 4/9/2015 | GISAID |
| EPI684179 | A/Iowa/40/2015(H3N2) | 4/9/2015 | GISAID |
| EPI684187 | A/Iowa/41/2015(H3N2) | 10/9/2015 | GISAID |
| EPI684195 | A/Iowa/41/2015(H3N2) | 10/9/2015 | GISAID |
| EPI684203 | A/Iowa/42/2015(H3N2) | 10/9/2015 | GISAID |
| EPI684211 | A/Iowa/42/2015(H3N2) | 10/9/2015 | GISAID |
| EPI684219 | A/Iowa/43/2015(H3N2) | 14/9/2015 | GISAID |
| EPI684227 | A/Iowa/43/2015(H3N2) | 14/9/2015 | GISAID |
| EPI684235 | A/Wyoming/18/2015(H3N2) | 22/9/2015 | GISAID |
| EPI684243 | A/Bolivia/426/2015(H3N2) | 8/7/2015 | GISAID |
| EPI684251 | A/Bolivia/499/2015(H3N2) | 14/8/2015 | GISAID |
| EPI684259 | A/Bolivia/502/2015(H3N2) | 18/8/2015 | GISAID |
| EPI684267 | A/Michigan/67/2015(H3N2) | 10/10/2015 | GISAID |
| EPI684283 | A/SouthAfrica/6270/2015(H3N2) | 2/10/2015 | GISAID |
| EPI684291 | A/SouthAfrica/4940/2015(H3N2) | 12/8/2015 | GISAID |
| EPI684299 | A/SouthAfrica/6131/2015(H3N2) | 30/9/2015 | GISAID |
| EPI684307 | A/SouthAfrica/5390/2015(H3N2) | 28/8/2015 | GISAID |
| EPI684315 | A/Norway/2178/2014(H3N2) | 2014 | GISAID |
| EPI685587 | A/Montana/28/2015(H3N2) | 24/9/2015 | GISAID |
| EPI685783 | A/Hawaii/43/2015(H3N2) | 8/7/2015 | GISAID |
| EPI685791 | A/Alberta/RV2373/2015(H3N2) | 28/9/2015 | GISAID |
| EPI685799 | A/Alberta/RV2382/2015(H3N2) | 1/10/2015 | GISAID |
| EPI685807 | A/Alberta/RV2383/2015(H3N2) | 9/10/2015 | GISAID |
| EPI685815 | A/Alberta/RV2385/2015(H3N2) | 10/10/2015 | GISAID |
| EPI685823 | A/Alberta/RV2386/2015(H3N2) | 10/10/2015 | GISAID |
| EPI685831 | A/Alberta/RV2387/2015(H3N2) | 10/10/2015 | GISAID |

| | | | |
|---|---|---|---|
| EPI685847 | A/Laos/724/2015(H3N2) | 13/7/2015 | GISAID |
| EPI685855 | A/Laos/844/2015(H3N2) | 12/8/2015 | GISAID |
| EPI685863 | A/Laos/853/2015(H3N2) | 12/8/2015 | GISAID |
| EPI685871 | A/Laos/351/2015(H3N2) | 11/8/2015 | GISAID |
| EPI685879 | A/Laos/351/2015(H3N2) | 11/8/2015 | GISAID |
| EPI685887 | A/Laos/884/2015(H3N2) | 24/8/2015 | GISAID |
| EPI685895 | A/Laos/898/2015(H3N2) | 24/8/2015 | GISAID |
| EPI685903 | A/Laos/913/2015(H3N2) | 31/8/2015 | GISAID |
| EPI685911 | A/Laos/928/2015(H3N2) | 9/9/2015 | GISAID |
| EPI685919 | A/Laos/375/2015(H3N2) | 8/9/2015 | GISAID |
| EPI685927 | A/Laos/375/2015(H3N2) | 8/9/2015 | GISAID |
| EPI685942 | A/Michigan/60/2015(H3N2) | 5/10/2015 | GISAID |
| EPI685950 | A/Michigan/60/2015(H3N2) | 5/10/2015 | GISAID |
| EPI685963 | A/Washington/61/2015(H3N2) | 1/11/2015 | GISAID |
| EPI685971 | A/FrenchGuiana/5020/2015(H3N2) | 13/7/2015 | GISAID |
| EPI685979 | A/Guatemala/142/2015(H3N2) | 18/6/2015 | GISAID |
| EPI685987 | A/Washington/45/2015(H3N2) | 16/6/2015 | GISAID |
| EPI686000 | A/Missouri/42/2015(H3N2) | 4/10/2015 | GISAID |
| EPI686008 | A/Pennsylvania/40/2015(H3N2) | 22/10/2015 | GISAID |
| EPI686016 | A/Alaska/183/2015(H3N2) | 16/8/2015 | GISAID |
| EPI686024 | A/Alaska/185/2015(H3N2) | 20/8/2015 | GISAID |
| EPI686032 | A/Alaska/222/2015(H3N2) | 8/9/2015 | GISAID |
| EPI686040 | A/Alaska/225/2015(H3N2) | 11/9/2015 | GISAID |
| EPI686048 | A/Alaska/239/2015(H3N2) | 16/9/2015 | GISAID |
| EPI686056 | A/Alaska/242/2015(H3N2) | 13/9/2015 | GISAID |
| EPI686064 | A/Hawaii/62/2015(H3N2) | 28/9/2015 | GISAID |
| EPI686072 | A/PuertoRico/12/2015(H3N2) | 24/8/2015 | GISAID |
| EPI686080 | A/PuertoRico/17/2015(H3N2) | 9/9/2015 | GISAID |
| EPI686088 | A/PuertoRico/19/2015(H3N2) | 8/9/2015 | GISAID |
| EPI686096 | A/PuertoRico/20/2015(H3N2) | 9/9/2015 | GISAID |
| EPI686104 | A/HongKong/15590/2015(H3N2) | 9/10/2015 | GISAID |
| EPI686112 | A/HongKong/15609/2015(H3N2) | 17/10/2015 | GISAID |
| EPI686120 | A/HongKong/15611/2015(H3N2) | 19/10/2015 | GISAID |
| EPI686128 | A/Washington/51/2015(H3N2) | 9/10/2015 | GISAID |
| EPI686143 | A/Nebraska/19/2015(H3N2) | 23/10/2015 | GISAID |
| EPI686151 | A/Florida/86/2015(H3N2) | 2/11/2015 | GISAID |
| EPI686159 | A/Colorado/19/2015(H3N2) | 15/11/2015 | GISAID |

| | | | |
|---|---|---|---|
| EPI686167 | A/Colorado/21/2015(H3N2) | 19/11/2015 | GISAID |
| EPI686182 | A/Colorado/24/2015(H3N2) | 17/11/2015 | GISAID |
| EPI686190 | A/California/94/2015(H3N2) | 20/10/2015 | GISAID |
| EPI686198 | A/California/98/2015(H3N2) | 19/10/2015 | GISAID |
| EPI686206 | A/Texas/71/2015(H3N2) | 11/11/2015 | GISAID |
| EPI686214 | A/Maryland/17/2015(H3N2) | 8/10/2015 | GISAID |
| EPI686222 | A/Washington/62/2015(H3N2) | 3/12/2015 | GISAID |
| EPI686230 | A/Texas/75/2015(H3N2) | 8/12/2015 | GISAID |
| EPI686238 | A/Hawaii/76/2015(H3N2) | 18/10/2015 | GISAID |
| EPI686246 | A/California/101/2015(H3N2) | 4/11/2015 | GISAID |
| EPI686254 | A/California/106/2015(H3N2) | 3/11/2015 | GISAID |
| EPI686275 | A/Pennsylvania/50/2015(H3N2) | 23/11/2015 | GISAID |
| EPI686286 | A/Colorado/27/2015(H3N2) | 8/12/2015 | GISAID |
| EPI686294 | A/Florida/93/2015(H3N2) | 4/12/2015 | GISAID |
| EPI686302 | A/California/112/2015(H3N2) | 18/11/2015 | GISAID |
| EPI686310 | A/California/113/2015(H3N2) | 28/11/2015 | GISAID |
| EPI686318 | A/Alaska/266/2015(H3N2) | 26/11/2015 | GISAID |
| EPI686326 | A/Alaska/267/2015(H3N2) | 27/11/2015 | GISAID |
| EPI686748 | A/Halmstad/1/2015(H3N2) | 15/11/2015 | GISAID |
| EPI686756 | A/Skovde/5/2015(H3N2) | 17/11/2015 | GISAID |
| EPI686780 | A/Lund/2/2015(H3N2) | 16/11/2015 | GISAID |
| EPI686791 | A/Stockholm/58/2015(H3N2) | 17/11/2015 | GISAID |
| EPI686836 | A/Stockholm/61/2015(H3N2) | 23/11/2015 | GISAID |
| EPI686860 | A/Gothenburg/1/2015(H3N2) | 21/11/2015 | GISAID |
| EPI686868 | A/Stockholm/63/2015(H3N2) | 23/11/2015 | GISAID |
| EPI686876 | A/Stockholm/64/2015(H3N2) | 21/11/2015 | GISAID |
| EPI686884 | A/Kungalv/1/2015(H3N2) | 3/12/2015 | GISAID |
| EPI686924 | A/Uppsala/7/2015(H3N2) | 30/10/2015 | GISAID |
| EPI687165 | A/Stockholm/65/2015(H3N2) | 20/11/2015 | GISAID |
| EPI232453 | A/Victoria/208/2009(H3N2) | 2/6/2009 | GISAID |
| EPI232529 | A/Florida/22/2009(H3N2) | 2009 | GISAID |
| EPI232547 | A/HongKong/26560/2009(H3N2) | 18/9/2009 | GISAID |
| EPI232594 | A/Philippines/2191/2009(H3N2) | 26/5/2009 | GISAID |
| EPI241634 | A/Colombia/4335/2009(H3N2) | 2009 | GISAID |
| EPI254583 | A/Bangladesh/5014/2009(H3N2) | 17/5/2009 | GISAID |
| EPI270296 | A/India/2710/2009(H3N2) | 19/9/2009 | GISAID |
| EPI278760 | A/Panama/307149/2010(H3N2) | 8/6/2010 | GISAID |

| | | | |
|---|---|---|---|
| EPI278805 | A/Alabama/05/2010(H3N2) | 13/7/2010 | GISAID |
| EPI279171 | A/SRILANKA/7/2010(H3N2) | 25/5/2010 | GISAID |
| EPI279177 | A/CAMBODIA/1/2010(H3N2) | 21/6/2010 | GISAID |
| EPI280005 | A/Brisbane/11/2010(H3N2) | 26/5/2010 | GISAID |
| EPI293792 | A/Tianjin-Hexi/18/2010(H3N2) | 5/1/2010 | GISAID |
| EPI293800 | A/Beijing-Zhaoyang/141/2010(H3N2) | 6/1/2010 | GISAID |
| EPI293808 | A/Tianjin-Nankai/1842/2010(H3N2) | 1/3/2010 | GISAID |
| EPI293816 | A/Tianjin-Nankai/1912/2010(H3N2) | 24/3/2010 | GISAID |
| EPI293824 | A/Heilongjiang-Longsha/160/2010(H3N2) | 9/4/2010 | GISAID |
| EPI293832 | A/Tianjin-Hexi/1266/2010(H3N2) | 12/5/2010 | GISAID |
| EPI293864 | A/Anhui-Baohe/1456/2010(H3N2) | 17/8/2010 | GISAID |
| EPI293872 | A/Hubei-Qiaokou/1271/2010(H3N2) | 6/9/2010 | GISAID |
| EPI293880 | A/Shanghai-Luwan/1568/2010(H3N2) | 19/9/2010 | GISAID |
| EPI293888 | A/Hubei-Wuchang/1271/2010(H3N2) | 10/10/2010 | GISAID |
| EPI295112 | A/Kenya/6063/2010(H3N2) | 27/7/2010 | GISAID |
| EPI295119 | A/Kenya/6063/2010(H3N2) | 27/7/2010 | GISAID |
| EPI295127 | A/Kenya/6063/2010(H3N2) | 27/7/2010 | GISAID |
| EPI295216 | A/Mexico/4553/2010(H3N2) | 3/9/2010 | GISAID |
| EPI295224 | A/Mexico/4553/2010(H3N2) | 3/9/2010 | GISAID |
| EPI302259 | A/Taiwan/851/2010(H3N2) | 15/9/2010 | GISAID |
| EPI304954 | A/PHILIPPINES/16/2009(H3N2) | 5/3/2009 | GISAID |
| EPI334627 | A/Texas/24/2010(H3N2) | 31/12/2010 | GISAID |
| EPI331237 | A/Massachusetts/06/2010(H3N2) | 31/12/2010 | GISAID |
| EPI309555 | A/NewJersey/01/2011(H3N2) | 11/1/2011 | GISAID |
| EPI309581 | A/Georgia/01/2011(H3N2) | 10/2/2011 | GISAID |
| EPI309608 | A/Ontario/RV0064/2011(H3N2) | 2/1/2011 | GISAID |
| EPI316063 | A/Illinois/03/2011(H3N2) | 22/1/2011 | GISAID |
| EPI316200 | A/Texas/06/2011(H3N2) | 5/2/2011 | GISAID |
| EPI319642 | A/Alaska/15/2011(H3N2) | 1/3/2011 | GISAID |
| EPI319694 | A/Illinois/04/2011(H3N2) | 22/1/2011 | GISAID |
| EPI319790 | A/NorthCarolina/05/2011(H3N2) | 27/1/2011 | GISAID |
| EPI325036 | A/Wisconsin/22/2011(H3N2) | 15/4/2011 | GISAID |
| EPI326272 | A/California/16/2011(H3N2) | 16/1/2011 | GISAID |
| EPI326342 | A/Iowa/19/2010(H3N2) | 30/12/2010 | GISAID |
| EPI330100 | A/Beijing-Xicheng/1293/2011(H3N2) | 8/2/2011 | GISAID |
| EPI330108 | A/Beijing-Xicheng/1409/2011(H3N2) | 22/2/2011 | GISAID |
| EPI330513 | A/Shanghai-Luwan/188/2011(H3N2) | 14/2/2011 | GISAID |

| | | | |
|---|---|---|---|
| EPI330521 | A/Heilongjiang-Xiangyang/1125/2011(H3N2) | 9/3/2011 | GISAID |
| EPI330537 | A/Jilin-Chuanying/1156/2011(H3N2) | 3/3/2011 | GISAID |
| EPI330545 | A/Chongqing-Wanzhou/182/2011(H3N2) | 20/2/2011 | GISAID |
| EPI330553 | A/Tianjin-Hedong/195/2011(H3N2) | 15/2/2011 | GISAID |
| EPI330561 | A/Heilongjiang-Nangang/124/2011(H3N2) | 8/3/2011 | GISAID |
| EPI330753 | A/Hunan-Beihu/1143/2011(H3N2) | 11/3/2011 | GISAID |
| EPI330761 | A/Hunan-Yanfeng/165/2011(H3N2) | 14/4/2011 | GISAID |
| EPI330769 | A/Heilongjiang-Taoshan/147/2011(H3N2) | 15/4/2011 | GISAID |
| EPI330785 | A/Shanghai-Fengxian/1170/2011(H3N2) | 10/5/2011 | GISAID |
| EPI331283 | A/VICTORIA/506/2011(H3N2) | 15/1/2011 | GISAID |
| EPI331665 | A/TOWNSVILLE/117/2010(H3N2) | 16/12/2010 | GISAID |
| EPI332627 | A/NAURU/5/2011(H3N2) | 21/1/2011 | GISAID |
| EPI333152 | A/Indiana/08/2011(H3N2) | 27/7/2011 | GISAID |
| EPI335923 | A/Iowa/19/2010(H3N2) | 30/12/2010 | GISAID |
| EPI341665 | A/Guangdong-Luohu/1256/2009(H3N2) | 1/6/2009 | GISAID |
| EPI344389 | A/Indiana/10/2011(H3N2) | 22/10/2011 | GISAID |
| EPI344397 | A/Indiana/10/2011(H3N2) | 22/10/2011 | GISAID |
| EPI344405 | A/Indiana/08/2011(H3N2) | 27/7/2011 | GISAID |
| EPI346284 | A/SINGAPORE/41/2011(H3N2) | 16/5/2011 | GISAID |
| EPI346356 | A/VICTORIA/103/2011(H3N2) | 20/9/2011 | GISAID |
| EPI346424 | A/Iowa/09/2011(H3N2) | 14/11/2011 | GISAID |
| EPI346432 | A/Iowa/07/2011(H3N2) | 14/11/2011 | GISAID |
| EPI346440 | A/Iowa/08/2011(H3N2) | 14/11/2011 | GISAID |
