## Supplementary material for "Phylogenetic Tree-based Pipeline for Uncovering Mutational Patterns during Influenza Virus Evolution": Table S2

**Table S2.** GISAID acknowledgment table for sequences used in this study.

| *We acknowledge the authors, originating and submitting laboratories of the sequences from GISAID's EpiFlu* ™ Database on which this research is based. The list is detailed below. | | | | | | | |
|---|---|---|---|---|---|---|---|
| *All submitters of data may be contacted directly via the GISAID website* ***www.gisaid.org*** | | | | | | | |
| **Segment ID** | **Segment** | **Country** | **Collection date** | **Isolate name** | **Originating Lab** | **Submitting Lab** | **Authors** |
| EPI232453 | HA | Australia | 2009-Jun-02 | A/Victoria/208/2009 | WHO Collaborating Centre for Reference and Research on Influenza | Centers for Disease Control and Prevention | |
| EPI232529 | HA | United States | 2009-Jan-01 | A/Florida/22/2009 | | Centers for Disease Control and Prevention | |
| EPI232547 | HA | China | 2009-Sep-18 | A/Hong Kong/26560/2009 | Government Virus Unit | Centers for Disease Control and Prevention | |
| EPI232594 | HA | Philippines | 2009-May-26 | A/Philippines/2191/2009 | Thailand MOPH-U.S. CDC Collaboration (IEIP) | Centers for Disease Control and Prevention | |
| EPI241634 | HA | Colombia | 2009-Jan-01 | A/Colombia/4335/2009 | | Centers for Disease Control and Prevention | |
| EPI254583 | HA | Bangladesh | 2009-May-17 | A/Bangladesh/5014/2009 | Institute of Epidemiology Disease Control and Research (IEDCR) & Bangladesh National Influenza Centre (NIC) | Centers for Disease Control and Prevention | |
| EPI270296 | HA | India | 2009-Sep-19 | A/India/2710/2009 | National Institute of Virology | Centers for Disease Control and Prevention | |
| EPI278760 | HA | Panama | 2010-Jun-08 | A/Panama/307149/2010 | Instituto Conmemorativo Gorgas de Estudios de la Salud | Centers for Disease Control and Prevention | Yu Lan,Xiyan Li,Xiang Zhao,Yanhui Cheng,minju Tan,Bin Liu,Yaoyao Chen,Lei Yang,Dayan Wang,Dexin Li,Yuelong Shu |
| EPI278805 | HA | United States | 2010-Jul-13 | A/Alabama/05/2010 | U.S. Air Force School of Aerospace Medicine | Centers for Disease Control and Prevention | Yu Lan,Xiyan Li,Xiang Zhao,Yanhui Cheng,minju Tan,Bin Liu,Yaoyao Chen,Lei Yang,Dayan Wang,Dexin Li,Yuelong Shu |
| EPI279171 | HA | Sri Lanka | 2010-May-25 | A/SRI LANKA/7/2010 | Medical Research Institute | WHO Collaborating Centre for Reference and Research on Influenza | Yu Lan,Xiyan Li,Xiang Zhao,Yanhui Cheng,minju Tan,Bin Liu,Yaoyao Chen,Lei Yang,Dayan Wang,Dexin Li,Yuelong Shu |
| EPI279177 | HA | Cambodia | 2010-Jun-21 | A/CAMBODIA/1/2010 | Institute Pasteur du Cambodia | WHO Collaborating Centre for Reference and Research on Influenza | Yu Lan,Xiyan Li,Xiang Zhao,Yanhui Cheng,minju Tan,Bin Liu,Yaoyao Chen,Lei Yang,Dayan Wang,Dexin Li,Yuelong Shu |
| EPI280005 | HA | Australia | 2010-May-26 | A/Brisbane/11/2010 | WHO Collaborating Centre for Reference and Research on Influenza | Centers for Disease Control and Prevention | Yu Lan,Xiyan Li,Xiang Zhao,Yanhui Cheng,minju Tan,Bin Liu,Yaoyao Chen,Lei Yang,Dayan Wang,Dexin Li,Yuelong Shu |
| EPI293792 | HA | China | 2010-Jan-05 | A/Tianjin-Hexi/18/2010 | WHO Chinese National Influenza Center | WHO Chinese National Influenza Center | Yu Lan,Xiyan Li,Xiang Zhao,Yanhui Cheng,minju Tan,Bin Liu,Yaoyao Chen,Lei Yang,Dayan Wang,Dexin Li,Yuelong Shu |

| | | | | | | | |
|---|---|---|---|---|---|---|---|
| EPI293800 | HA | China | 2010-Jan-06 | A/Beijing-Zhaoyang/141/2010 | WHO Chinese National Influenza Center | WHO Chinese National Influenza Center | Yu Lan,Xiyan Li,Xiang Zhao,Yanhui Cheng,minju Tan,Bin Liu,Yaoyao Chen,Lei Yang,Dayan Wang,Dexin Li,Yuelong Shu |
| EPI293808 | HA | China | 2010-Mar-01 | A/Tianjin-Nankai/1842/2010 | WHO Chinese National Influenza Center | WHO Chinese National Influenza Center | Yu Lan,Xiyan Li,Xiang Zhao,Yanhui Cheng,minju Tan,Bin Liu,Yaoyao Chen,Lei Yang,Dayan Wang,Dexin Li,Yuelong Shu |
| EPI293816 | HA | China | 2010-Mar-24 | A/Tianjin-Nankai/1912/2010 | WHO Chinese National Influenza Center | WHO Chinese National Influenza Center | Yu Lan,Xiyan Li,Xiang Zhao,Yanhui Cheng,minju Tan,Bin Liu,Yaoyao Chen,Lei Yang,Dayan Wang,Dexin Li,Yuelong Shu |
| EPI293824 | HA | China | 2010-Apr-09 | A/Heilongjiang-Longsha/160/2010 | WHO Chinese National Influenza Center | WHO Chinese National Influenza Center | Yu Lan,Xiyan Li,Xiang Zhao,Yanhui Cheng,minju Tan,Bin Liu,Yaoyao Chen,Lei Yang,Dayan Wang,Dexin Li,Yuelong Shu |
| EPI293832 | HA | China | 2010-May-12 | A/Tianjin-Hexi/1266/2010 | WHO Chinese National Influenza Center | WHO Chinese National Influenza Center | |
| EPI293864 | HA | China | 2010-Aug-17 | A/Anhui-Baohe/1456/2010 | WHO Chinese National Influenza Center | WHO Chinese National Influenza Center | Deng,Y-M; Iannello,P; Caldwell,N; Leang,S-K; Komadina,N |
| EPI293872 | HA | China | 2010-Sep-06 | A/Hubei-Qiaokou/1271/2010 | WHO Chinese National Influenza Center | WHO Chinese National Influenza Center | Deng,Y-M; Iannello,P; Caldwell,N; Leang,S-K; Komadina,N |
| EPI293880 | HA | China | 2010-Sep-19 | A/Shanghai-Luwan/1568/2010 | WHO Chinese National Influenza Center | WHO Chinese National Influenza Center | |
| EPI293888 | HA | China | 2010-Oct-10 | A/Hubei-Wuchang/1271/2010 | WHO Chinese National Influenza Center | WHO Chinese National Influenza Center | |
| EPI295112 | HA | Kenya | 2010-Jul-27 | A/Kenya/6063/2010 | CDC-Kenya | Centers for Disease Control and Prevention | |
| EPI295119 | HA | Kenya | 2010-Jul-27 | A/Kenya/6063/2010 | CDC-Kenya | Centers for Disease Control and Prevention | |
| EPI295127 | HA | Kenya | 2010-Jul-27 | A/Kenya/6063/2010 | CDC-Kenya | Centers for Disease Control and Prevention | |
| EPI295216 | HA | Mexico | 2010-Sep-03 | A/Mexico/4553/2010 | Laboratorio de Virus Respiratorio | Centers for Disease Control and Prevention | |
| EPI295224 | HA | Mexico | 2010-Sep-03 | A/Mexico/4553/2010 | Laboratorio de Virus Respiratorio | Centers for Disease Control and Prevention | |
| EPI302259 | HA | Taiwan | 2010-Sep-15 | A/Taiwan/851/2010 | ADImmune Corporation | Centers for Disease Control and Prevention | |
| EPI304954 | HA | Philippines | 2009-Mar-05 | A/PHILIPPINES/16/2009 | Research Institute of Tropical Medicine | WHO Collaborating Centre for Reference and Research on Influenza | |
| EPI309555 | HA | United States | 2011-Jan-11 | A/New Jersey/01/2011 | New Jersey Department of Health & Senior Services | Centers for Disease Control and Prevention | |
| EPI309581 | HA | United States | 2011-Feb-10 | A/Georgia/01/2011 | Centers for Disease Control and Prevention | Centers for Disease Control and Prevention | |
| EPI309608 | HA | Canada | 2011-Jan-02 | A/Ontario/RV0064/2011 | National Microbiology Laboratory, Health Canada | Centers for Disease Control and Prevention | |
| EPI316063 | HA | United States | 2011-Jan-22 | A/Illinois/03/2011 | Illinois Department of Public Health-Chicago | Centers for Disease Control and Prevention | |
| EPI316200 | HA | United States | 2011-Feb-05 | A/Texas/06/2011 | Dallas County Health and Human Services | Centers for Disease Control and Prevention | |

| | | | | | | |
|---|---|---|---|---|---|---|
| EPI319642 | HA | United States | 2011-Mar-01 | A/Alaska/15/2011 | Alaska State Virology Lab | Centers for Disease Control and Prevention |
| EPI319694 | HA | United States | 2011-Jan-22 | A/Illinois/04/2011 | Illinois Department of Public Health-Chicago | Centers for Disease Control and Prevention |
| EPI319790 | HA | United States | 2011-Jan-27 | A/North Carolina/05/2011 | North Carolina State Laboratory of Public Health | Centers for Disease Control and Prevention |
| EPI325036 | HA | United States | 2011-Apr-15 | A/Wisconsin/22/2011 | Wisconsin State Laboratory of Hygiene | Centers for Disease Control and Prevention |
| EPI326272 | HA | United States | 2011-Jan-16 | A/California/16/2011 | California Department of Health Services | Centers for Disease Control and Prevention |
| EPI326342 | HA | United States | 2010-Dec-30 | A/Iowa/19/2010 | WHO Collaborating Centre for Reference and Research on Influenza | Centers for Disease Control and Prevention |
| EPI330100 | HA | China | 2011-Feb-08 | A/Beijing-Xicheng/1293/2011 | WHO Chinese National Influenza Center | WHO Chinese National Influenza Center |
| EPI330108 | HA | China | 2011-Feb-22 | A/Beijing-Xicheng/1409/2011 | WHO Chinese National Influenza Center | WHO Chinese National Influenza Center |
| EPI330513 | HA | China | 2011-Feb-14 | A/Shanghai-Luwan/188/2011 | WHO Chinese National Influenza Center | WHO Chinese National Influenza Center |
| EPI330521 | HA | China | 2011-Mar-09 | A/Heilongjiang-Xiangyang/1125/2011 | WHO Chinese National Influenza Center | WHO Chinese National Influenza Center |
| EPI330537 | HA | China | 2011-Mar-03 | A/Jilin-Chuanying/1156/2011 | WHO Chinese National Influenza Center | WHO Chinese National Influenza Center |
| EPI330545 | HA | China | 2011-Feb-20 | A/Chongqing-Wanzhou/182/2011 | WHO Chinese National Influenza Center | WHO Chinese National Influenza Center |
| EPI330553 | HA | China | 2011-Feb-15 | A/Tianjin-Hedong/195/2011 | WHO Chinese National Influenza Center | WHO Chinese National Influenza Center |
| EPI330561 | HA | China | 2011-Mar-08 | A/Heilongjiang-Nangang/124/2011 | WHO Chinese National Influenza Center | WHO Chinese National Influenza Center |
| EPI330753 | HA | China | 2011-Mar-11 | A/Hunan-Beihu/1143/2011 | WHO Chinese National Influenza Center | WHO Chinese National Influenza Center |
| EPI330761 | HA | China | 2011-Apr-14 | A/Hunan-Yanfeng/165/2011 | WHO Chinese National Influenza Center | WHO Chinese National Influenza Center |
| EPI330769 | HA | China | 2011-Apr-15 | A/Heilongjiang-Taoshan/147/2011 | WHO Chinese National Influenza Center | WHO Chinese National Influenza Center |
| EPI330785 | HA | China | 2011-May-10 | A/Shanghai-Fengxian/1170/2011 | WHO Chinese National Influenza Center | WHO Chinese National Influenza Center |
| EPI331237 | HA | United States | 2010-Dec-31 | A/Massachusetts/06/2010 | Massachusetts Department of Public Health | Centers for Disease Control and Prevention |
| EPI331283 | HA | Australia | 2011-Jan-15 | A/VICTORIA/506/2011 | Monash Medical Centre | WHO Collaborating Centre for Reference and Research on Influenza |
| EPI331665 | HA | Australia | 2010-Dec-16 | A/TOWNSVILLE/117/2010 | Queensland Health Scientific Services | WHO Collaborating Centre for Reference and Research on Influenza |
| EPI332627 | HA | Nauru | 2011-Jan-21 | A/NAURU/5/2011 | Republic of Nauru Hospital | WHO Collaborating Centre for Reference and Research on Influenza |
| EPI333152 | HA | United States | 2011-Jul-27 | A/Indiana/08/2011 | Indiana State Department of Health Laboratories | Centers for Disease Control and Prevention |

| | | | | | | |
|---|---|---|---|---|---|---|
| EPI334627 | HA | United States | 2010-Dec-31 | A/Texas/24/2010 | Dallas County Health and Human Services | Centers for Disease Control and Prevention |
| EPI335923 | HA | United States | 2010-Dec-30 | A/Iowa/19/2010 | Iowa State Hygienic Laboratory | Centers for Disease Control and Prevention |
| EPI341665 | HA | China | 2009-Jun-01 | A/Guangdong-Luohu/1256/2009 | WHO Chinese National Influenza Center | Centers for Disease Control and Prevention |
| EPI344389 | HA | United States | 2011-Oct-22 | A/Indiana/10/2011 | Indiana State Department of Health Laboratories | Centers for Disease Control and Prevention |
| EPI344397 | HA | United States | 2011-Oct-22 | A/Indiana/10/2011 | Indiana State Department of Health Laboratories | Centers for Disease Control and Prevention |
| EPI344405 | HA | United States | 2011-Jul-27 | A/Indiana/08/2011 | Indiana State Department of Health Laboratories | Centers for Disease Control and Prevention |
| EPI346284 | HA | Singapore | 2011-May-16 | A/SINGAPORE/41/2011 | Ministry of Health, Singapore | WHO Collaborating Centre for Reference and Research on Influenza |
| EPI346356 | HA | Australia | 2011-Sep-20 | A/VICTORIA/103/2011 | Victorian Infectious Diseases Reference Laboratory | WHO Collaborating Centre for Reference and Research on Influenza |
| EPI346424 | HA | United States | 2011-Nov-14 | A/Iowa/09/2011 | Iowa State Hygienic Laboratory | Centers for Disease Control and Prevention |
| EPI346432 | HA | United States | 2011-Nov-14 | A/Iowa/07/2011 | Iowa State Hygienic Laboratory | Centers for Disease Control and Prevention |
| EPI346440 | HA | United States | 2011-Nov-14 | A/Iowa/08/2011 | Iowa State Hygienic Laboratory | Centers for Disease Control and Prevention |
| EPI347506 | HA | United States | 2011-Nov-04 | A/Kentucky/05/2011 | Kentucky Division of Laboratory Services | Centers for Disease Control and Prevention |
| EPI347544 | HA | United States | 2011-Nov-06 | A/Utah/12/2011 | Utah Department of Health | Centers for Disease Control and Prevention |
| EPI348478 | HA | Thailand | 2011-Sep-20 | A/Chiang Rai/277/2011 | WHO National Influenza Centre, National Institute of Medical Research (NIMR) | Centers for Disease Control and Prevention |
| EPI349103 | HA | Australia | 2011-Oct-24 | A/Victoria/361/2011 | Melbourne Pathology | WHO Collaborating Centre for Reference and Research on Influenza |
| EPI353512 | HA | Hong Kong (SAR) | 2009-Apr-07 | A/Hong Kong/2000/2009 | Government Virus Unit | Centers for Disease Control and Prevention |
| EPI353514 | HA | United States | 2011-Nov-04 | A/Kentucky/05/2011 | Centers for Disease Control and Prevention | Centers for Disease Control and Prevention |
| EPI353906 | HA | Australia | 2011-Oct-24 | A/Victoria/361/2011 | WHO Collaborating Centre for Reference and Research on Influenza | Centers for Disease Control and Prevention |
| EPI357622 | HA | Papua New Guinea | 2011-Jul-27 | A/GOROKA/5/2011 | Papua New Guinea Institute of Medical Research | WHO Collaborating Centre for Reference and Research on Influenza |
| EPI357660 | HA | Australia | 2011-Sep-17 | A/PERTH/219/2011 | Pathwest QE II Medical Centre | WHO Collaborating Centre for Reference and Research on Influenza |
| EPI357669 | HA | Australia | 2011-Sep-06 | A/BRISBANE/382/2011 | Queensland Health Scientific Services | WHO Collaborating Centre for Reference and Research on Influenza |

| | | | | | | | |
|---|---|---|---|---|---|---|---|
| EPI357675 | HA | Australia | 2011-Oct-20 | A/BRISBANE/395/2011 | Queensland Health Scientific Services | WHO Collaborating Centre for Reference and Research on Influenza | Deng,Y-M; Iannello,P; Caldwell,N; Jelley,L; Komadina,N |
| EPI363012 | HA | United States | 2012-Mar-02 | A/Washington/16/2012 | Spokane Regional Health District | Centers for Disease Control and Prevention | Deng,Y-M; Iannello,P; Caldwell,N; Jelley,L; Komadina,N |
| EPI363019 | HA | United States | 2012-Jan-25 | A/California/17/2012 | California Department of Health Services | Centers for Disease Control and Prevention | Deng,Y-M; Iannello,P; Caldwell,N; Jelley,L; Komadina,N |
| EPI370074 | HA | Australia | 2011-Dec-07 | A/VICTORIA/893/2011 | Austin Health | WHO Collaborating Centre for Reference and Research on Influenza | Deng,Y-M; Iannello,P; Caldwell,N; Jelley,L; Komadina,N |
| EPI370111 | HA | Philippines | 2011-Oct-25 | A/PHILIPPINES/3969/2011 | Research Institute of Tropical Medicine | WHO Collaborating Centre for Reference and Research on Influenza | |
| EPI370132 | HA | Australia | 2011-Dec-12 | A/PERTH/305/2011 | Pathwest QE II Medical Centre | WHO Collaborating Centre for Reference and Research on Influenza | |
| EPI370318 | HA | Cambodia | 2011-Nov-17 | A/CAMBODIA/74/2011 | USAMC-AFRIMS Department of Virology | WHO Collaborating Centre for Reference and Research on Influenza | |
| EPI370329 | HA | Philippines | 2011-Oct-21 | A/PHILIPPINES/3948/2011 | Research Institute of Tropical Medicine | WHO Collaborating Centre for Reference and Research on Influenza | |
| EPI371776 | HA | United States | 2012-Mar-30 | A/Nevada/12/2012 | Southern Nevada Public Health Lab | Centers for Disease Control and Prevention | |
| EPI371792 | HA | United States | 2012-Feb-20 | A/California/22/2012 | California Department of Health Services | Centers for Disease Control and Prevention | |
| EPI371802 | HA | United States | 2012-Feb-16 | A/New Mexico/07/2012 | New Mexico Department of Health | Centers for Disease Control and Prevention | |
| EPI371819 | HA | United States | 2012-Mar-06 | A/Michigan/03/2012 | Michigan Department of Community Health | Centers for Disease Control and Prevention | |
| EPI372003 | HA | United States | 2012-Jan-30 | A/Rhode Island/01/2012 | Rhode Island Department of Health | Centers for Disease Control and Prevention | |
| EPI376886 | HA | Australia | 2012-Feb-20 | A/TASMANIA/1/2012 | Royal Hobart Hospital | WHO Collaborating Centre for Reference and Research on Influenza | |
| EPI376894 | HA | Australia | 2012-Jan-21 | A/TOWNSVILLE/1/2012 | Queensland Health Scientific Services | WHO Collaborating Centre for Reference and Research on Influenza | |
| EPI377004 | HA | Australia | 2011-Nov-18 | A/SOUTH AUSTRALIA/313/2011 | Institute of Medical and Veterinary Science (IMVS) | WHO Collaborating Centre for Reference and Research on Influenza | |
| EPI378515 | HA | United States | 2012-Jul-16 | A/Indiana/07/2012 | Indiana State Department of Health Laboratories | Centers for Disease Control and Prevention | |
| EPI378523 | HA | United States | 2012-Jul-16 | A/Indiana/09/2012 | Indiana State Department of Health Laboratories | Centers for Disease Control and Prevention | |
| EPI379447 | HA | Australia | 2012-Apr-08 | A/PERTH/41/2012 | Pathwest QE II Medical Centre | WHO Collaborating Centre for Reference and Research on Influenza | Deng,Y-M; Iannello,P; Caldwell,N; Jelley,L; Komadina,N |
| EPI379508 | HA | Papua New Guinea | 2012-Mar-05 | A/GOROKA/5/2012 | Papua New Guinea Institute of Medical Research | WHO Collaborating Centre for Reference and Research on Influenza | Deng,Y-M; Iannello,P; Caldwell,N; Jelley,L; Komadina,N |

| | | | | | | | |
|---|---|---|---|---|---|---|---|
| EPI379585 | HA | Australia | 2012-May-09 | A/VICTORIA/802/2012 | Austin Health | WHO Collaborating Centre for Reference and Research on Influenza | |
| EPI384818 | HA | United States | 2012-Jan-30 | A/Rhode Island/01/2012 | Rhode Island Department of Health | Centers for Disease Control and Prevention | Deng,Y-M; Iannello,P; Caldwell,N; Jelley,L; Komadina,N. |
| EPI384820 | HA | United States | 2012-Mar-08 | A/Ohio/02/2012 | WHO Collaborating Centre for Reference and Research on Influenza | Centers for Disease Control and Prevention | Deng,Y-M; Iannello,P; Caldwell,N; Komadina,N |
| EPI388305 | HA | United States | 2012-Jul-29 | A/Ohio/14/2012 | Ohio Department of Health Laboratories | Centers for Disease Control and Prevention | Deng,Y-M; Iannello,P; Caldwell,N; Komadina,N |
| EPI388339 | HA | United States | 2012-Jul-28 | A/Ohio/16/2012 | Ohio Department of Health Laboratories | Centers for Disease Control and Prevention | |
| EPI388346 | HA | United States | 2012-Jul-29 | A/Indiana/15/2012 | Indiana State Department of Health Laboratories | Centers for Disease Control and Prevention | |
| EPI388352 | HA | United States | 2012-Jul-29 | A/Indiana/20/2012 | Indiana State Department of Health Laboratories | Centers for Disease Control and Prevention | |
| EPI392708 | HA | United States | 2012-Aug-02 | A/Indiana/59/2012 | Indiana State Department of Health Laboratories | Centers for Disease Control and Prevention | |
| EPI393298 | HA | United States | 2012-Jul-31 | A/Indiana/21/2012 | Indiana State Department of Health Laboratories | Centers for Disease Control and Prevention | Deng,Y-M; Iannello,P; Caldwell,N; Komadina,N. |
| EPI393306 | HA | United States | 2012-Jul-29 | A/Indiana/18/2012 | Indiana State Department of Health Laboratories | Centers for Disease Control and Prevention | |
| EPI393311 | HA | United States | 2012-Aug-06 | A/West Virginia/09/2012 | West Virginia Office of Laboratory Services | Centers for Disease Control and Prevention | Deng,Y-M; Iannello,P; Caldwell,N; Komadina,N. |
| EPI393317 | HA | United States | 2012-Aug-04 | A/Ohio/38/2012 | Ohio Department of Health Laboratories | Centers for Disease Control and Prevention | Deng,Y-M; Iannello,P; Caldwell,N; Komadina,N. |
| EPI393511 | HA | United States | 2012-Aug-05 | A/Ohio/52/2012 | Ohio Department of Health Laboratories | Centers for Disease Control and Prevention | Deng,Y-M; Iannello,P; Caldwell,N; Komadina,N. |
| EPI393519 | HA | United States | 2012-Aug-02 | A/Indiana/65/2012 | Indiana State Department of Health Laboratories | Centers for Disease Control and Prevention | |
| EPI393661 | HA | United States | 2012-Aug-13 | A/Ohio/62/2012 | Ohio Department of Health Laboratories | Centers for Disease Control and Prevention | |
| EPI393667 | HA | United States | 2012-Aug-01 | A/Ohio/28/2012 | Ohio Department of Health Laboratories | Centers for Disease Control and Prevention | |
| EPI393673 | HA | United States | 2012-Aug-13 | A/Ohio/71/2012 | Ohio Department of Health Laboratories | Centers for Disease Control and Prevention | |
| EPI393681 | HA | United States | 2012-Aug-06 | A/West Virginia/10/2012 | West Virginia Office of Laboratory Services | Centers for Disease Control and Prevention | |
| EPI393688 | HA | United States | 2012-Jul-30 | A/Ohio/23/2012 | Ohio Department of Health Laboratories | Centers for Disease Control and Prevention | Shu, B.; Garten, R.; Emery, S.; Balish, A.; Smith, C.; Barnes, J.; Lindstrom, S.; Klimov, A.; Cox, N. |
| EPI393695 | HA | United States | 2012-Aug-09 | A/West Virginia/16/2012 | West Virginia Office of Laboratory Services | Centers for Disease Control and Prevention | Shu, B.; Garten, R.; Emery, S.; Balish, A.; Smith, C.; Barnes, J.; Lindstrom, S.; Klimov, A.; Cox, N. |
| EPI393702 | HA | United States | 2012-Aug-10 | A/Ohio/64/2012 | Ohio Department of Health Laboratories | Centers for Disease Control and Prevention | Deng,Y-M.; Iannello,P.; Caldwell,N.; Leang,S-K; |
| EPI394077 | HA | United States | 2012-Aug-17 | A/Wisconsin/27/2012 | Wisconsin State Laboratory of Hygiene | Centers for Disease Control and Prevention | Shu, B.; Garten, R.; Emery, S.; Balish, A.; Smith, C.; Barnes, J.; Lindstrom, S.; Klimov, A.; Cox, N. |

| | | | | | | | |
|---|---|---|---|---|---|---|---|
| EPI394116 | HA | Australia | 2012-Jun-24 | A/TASMANIA/12/2012 | Royal Hobart Hospital | WHO Collaborating Centre for Reference and Research on Influenza | Shu, B.; Garten, R.; Emery, S.; Balish, A.; Smith, C.; Barnes, J.; Lindstrom, S.; Klimov, A.; Cox, N. |
| EPI394124 | HA | New Caledonia | 2012-Mar-26 | A/NEW CALEDONIA/1/2012 | Institut Pasteur New Caledonia | WHO Collaborating Centre for Reference and Research on Influenza | |
| EPI394138 | HA | Australia | 2012-Jun-26 | A/TASMANIA/26/2012 | Royal Hobart Hospital | WHO Collaborating Centre for Reference and Research on Influenza | Shu,B;Emery,S;Balish,A; Garten,R; Smith,C; Barnes,J;Lindstrom,S; |
| EPI394149 | HA | New Zealand | 2012-Jul-03 | A/CHRISTCHURCH/503/2012 | Canterbury Health Services | WHO Collaborating Centre for Reference and Research on Influenza | Deng,Y-M.; Iannello,P.; Caldwell,N.; Leang,S-K; Komadina,N. |
| EPI394152 | HA | New Zealand | 2012-Jul-07 | A/CHRISTCHURCH/516/2012 | Canterbury Health Services | WHO Collaborating Centre for Reference and Research on Influenza | Deng,Y-M.; Iannello,P.; Caldwell,N.; Leang,S-K; Komadina,N. |
| EPI394913 | HA | United States | 2012-Aug-03 | A/Indiana/132/2012 | Indiana State Department of Health Laboratories | Centers for Disease Control and Prevention | Lan,Yu;Wang,Dayan;Li,Xiyan;Huang,Weijuan;Zhao,Xiang;Chen,Yaoyao;Yang,Lei;Shu,Yuelong |
| EPI394921 | HA | United States | 2012-Aug-13 | A/Ohio/69/2012 | Ohio Department of Health Laboratories | Centers for Disease Control and Prevention | Lan,Yu;Wang,Dayan;Li,Xiyan;Huang,Weijuan;Zhao,Xiang;Chen,Yaoyao;Yang,Lei;Shu,Yuelong |
| EPI394933 | HA | United States | 2012-Aug-15 | A/Wisconsin/23/2012 | Wisconsin State Laboratory of Hygiene | Centers for Disease Control and Prevention | Lan,Yu;Wang,Dayan;Li,Xiyan;Huang,Weijuan;Zhao,Xiang;Chen,Yaoyao;Yang,Lei;Shu,Yuelong |
| EPI394937 | HA | United States | 2012-Aug-16 | A/Ohio/75/2012 | Ohio Department of Health Laboratories | Centers for Disease Control and Prevention | Lan,Yu;Wang,Dayan;Li,Xiyan;Huang,Weijuan;Zhao,Xiang;Chen,Yaoyao;Yang,Lei;Shu,Yuelong |
| EPI394944 | HA | United States | 2012-Aug-16 | A/Maryland/22/2012 | Maryland Department of Health and Mental Hygiene | Centers for Disease Control and Prevention | Lan,Yu;Wang,Dayan;Li,Xiyan;Huang,Weijuan;Zhao,Xiang;Chen,Yaoyao;Yang,Lei;Shu,Yuelong |
| EPI395259 | HA | United States | 2012-Aug-04 | A/Ohio/35/2012 | Ohio Department of Health Laboratories | Centers for Disease Control and Prevention | Lan,Yu;Wang,Dayan;Li,Xiyan;Huang,Weijuan;Zhao,Xiang;Chen,Yaoyao;Yang,Lei;Shu,Yuelong |
| EPI395267 | HA | United States | 2012-Aug-11 | A/Ohio/65/2012 | Ohio Department of Health Laboratories | Centers for Disease Control and Prevention | Lan,Yu;Wang,Dayan;Li,Xiyan;Huang,Weijuan;Zhao,Xiang;Chen,Yaoyao;Yang,Lei;Shu,Yuelong |
| EPI395344 | HA | United States | 2012-Aug-28 | A/Wisconsin/28/2012 | Wisconsin State Laboratory of Hygiene | Centers for Disease Control and Prevention | Lan,Yu;Wang,Dayan;Li,Xiyan;Huang,Weijuan;Zhao,Xiang;Chen,Yaoyao;Yang,Lei;Shu,Yuelong |
| EPI395351 | HA | United States | 2012-Aug-16 | A/Michigan/20/2012 | Michigan Department of Community Health | Centers for Disease Control and Prevention | Lan,Yu;Wang,Dayan;Li,Xiyan;Huang,Weijuan;Zhao,Xiang;Chen,Yaoyao;Yang,Lei;Shu,Yuelong |
| EPI395357 | HA | United States | 2012-Aug-23 | A/Pennsylvania/17/2012 | Pennsylvania Department of Health | Centers for Disease Control and Prevention | Lan,Yu;Wang,Dayan;Li,Xiyan;Huang,Weijuan;Zhao,Xiang;Chen,Yaoyao;Yang,Lei;Shu,Yuelong |
| EPI396175 | HA | United States | 2012-Aug-19 | A/Ohio/80/2012 | Ohio Department of Health Laboratories | Centers for Disease Control and Prevention | |
| EPI396183 | HA | United States | 2012-Aug-02 | A/Indiana/77/2012 | Indiana State Department of Health Laboratories | Centers for Disease Control and Prevention | |

| | | | | | | | |
|---|---|---|---|---|---|---|---|
| EPI396192 | HA | United States | 2012-Aug-15 | A/Ohio/77/2012 | Ohio Department of Health Laboratories | Centers for Disease Control and Prevention | |
| EPI396200 | HA | United States | 2012-Aug-09 | A/Ohio/79/2012 | Ohio Department of Health Laboratories | Centers for Disease Control and Prevention | |
| EPI396208 | HA | United States | 2012-Aug-05 | A/Indiana/149/2012 | Indiana State Department of Health Laboratories | Centers for Disease Control and Prevention | |
| EPI396230 | HA | United States | 2012-Aug-09 | A/Indiana/147/2012 | Indiana State Department of Health Laboratories | Centers for Disease Control and Prevention | |
| EPI396239 | HA | United States | 2012-Aug-05 | A/Indiana/103/2012 | Indiana State Department of Health Laboratories | Centers for Disease Control and Prevention | |
| EPI396246 | HA | United States | 2012-Aug-03 | A/Indiana/125/2012 | Indiana State Department of Health Laboratories | Centers for Disease Control and Prevention | |
| EPI396254 | HA | United States | 2012-Aug-04 | A/Indiana/126/2012 | Indiana State Department of Health Laboratories | Centers for Disease Control and Prevention | |
| EPI396262 | HA | United States | 2012-Aug-04 | A/Indiana/127/2012 | Indiana State Department of Health Laboratories | Centers for Disease Control and Prevention | |
| EPI396270 | HA | United States | 2012-Aug-03 | A/Indiana/133/2012 | Indiana State Department of Health Laboratories | Centers for Disease Control and Prevention | Deng,Y-M; Iannello,P; Komadina,N |
| EPI396277 | HA | United States | 2012-Aug-08 | A/Indiana/135/2012 | Indiana State Department of Health Laboratories | Centers for Disease Control and Prevention | |
| EPI396280 | HA | United States | 2012-Aug-08 | A/Indiana/144/2012 | Indiana State Department of Health Laboratories | Centers for Disease Control and Prevention | |
| EPI396291 | HA | United States | 2012-Aug-10 | A/Ohio/63/2012 | Ohio Department of Health Laboratories | Centers for Disease Control and Prevention | |
| EPI396299 | HA | United States | 2012-Aug-18 | A/Maryland/23/2012 | Maryland Department of Health and Mental Hygiene | Centers for Disease Control and Prevention | |
| EPI397006 | HA | United States | 2012-Aug-09 | A/Ohio/78/2012 | Ohio Department of Health Laboratories | Centers for Disease Control and Prevention | |
| EPI397054 | HA | Puerto Rico | 2012-Jun-27 | A/Puerto Rico/36/2012 | Puerto Rico Department of Health | Centers for Disease Control and Prevention | |
| EPI397092 | HA | United States | 2012-Aug-02 | A/Indiana/57/2012 | Indiana State Department of Health Laboratories | Centers for Disease Control and Prevention | |
| EPI397106 | HA | United States | 2012-Aug-04 | A/Indiana/115/2012 | Indiana State Department of Health Laboratories | Centers for Disease Control and Prevention | |
| EPI397206 | HA | United States | 2012-Aug-03 | A/Ohio/34/2012 | Ohio Department of Health Laboratories | Centers for Disease Control and Prevention | |
| EPI397210 | HA | United States | 2012-Aug-18 | A/Maryland/25/2012 | Maryland Department of Health and Mental Hygiene | Centers for Disease Control and Prevention | |
| EPI397211 | HA | United States | 2012-Aug-05 | A/Indiana/113/2012 | Indiana State Department of Health Laboratories | Centers for Disease Control and Prevention | |
| EPI397340 | HA | United States | 2012-Aug-08 | A/Indiana/150/2012 | Indiana State Department of Health Laboratories | Centers for Disease Control and Prevention | |
| EPI397344 | HA | United States | 2012-Aug-16 | A/Michigan/19/2012 | Michigan Department of Community Health | Centers for Disease Control and Prevention | |
| EPI397351 | HA | United States | 2012-Aug-05 | A/Ohio/37/2012 | Ohio Department of Health Laboratories | Centers for Disease Control and Prevention | |
| EPI397385 | HA | United States | 2012-Aug-06 | A/Indiana/138/2012 | Indiana State Department of Health Laboratories | Centers for Disease Control and Prevention | |
| EPI397396 | HA | United States | 2012-Aug-06 | A/Indiana/93/2012 | Indiana State Department of Health Laboratories | Centers for Disease Control and Prevention | |

| | | | | | | | |
|---|---|---|---|---|---|---|---|
| EPI397404 | HA | United States | 2012-Aug-03 | A/Indiana/99/2012 | Indiana State Department of Health Laboratories | Centers for Disease Control and Prevention | |
| EPI397412 | HA | United States | 2012-Aug-15 | A/Michigan/16/2012 | Michigan Department of Community Health | Centers for Disease Control and Prevention | |
| EPI397418 | HA | United States | 2012-Aug-16 | A/Michigan/23/2012 | Michigan Department of Community Health | Centers for Disease Control and Prevention | |
| EPI397426 | HA | United States | 2012-Sep-05 | A/Minnesota/15/2012 | Minnesota Department of Health | Centers for Disease Control and Prevention | |
| EPI397432 | HA | United States | 2012-Aug-17 | A/Ohio/81/2012 | Ohio Department of Health Laboratories | Centers for Disease Control and Prevention | |
| EPI397446 | HA | United States | 2012-Aug-22 | A/Wisconsin/31/2012 | Wisconsin State Laboratory of Hygiene | Centers for Disease Control and Prevention | |
| EPI397503 | HA | United States | 2012-Aug-03 | A/Indiana/104/2012 | Indiana State Department of Health Laboratories | Centers for Disease Control and Prevention | Deng,Y-M; Iannello,P; Caldwell,N; Jelley,L; Komadina,N |
| EPI397516 | HA | United States | 2012-Aug-02 | A/Ohio/36/2012 | Ohio Department of Health Laboratories | Centers for Disease Control and Prevention | |
| EPI397524 | HA | United States | 2012-Aug-08 | A/Ohio/44/2012 | Ohio Department of Health Laboratories | Centers for Disease Control and Prevention | |
| EPI397532 | HA | United States | 2012-Aug-07 | A/Ohio/47/2012 | Ohio Department of Health Laboratories | Centers for Disease Control and Prevention | |
| EPI397540 | HA | United States | 2012-Aug-10 | A/Ohio/56/2012 | Ohio Department of Health Laboratories | Centers for Disease Control and Prevention | |
| EPI397548 | HA | United States | 2012-Aug-13 | A/Ohio/84/2012 | Ohio Department of Health Laboratories | Centers for Disease Control and Prevention | |
| EPI397555 | HA | United States | 2012-Aug-15 | A/Ohio/86/2012 | Ohio Department of Health Laboratories | Centers for Disease Control and Prevention | |
| EPI397562 | HA | United States | 2012-Aug-20 | A/Wisconsin/29/2012 | Wisconsin State Laboratory of Hygiene | Centers for Disease Control and Prevention | |
| EPI397654 | HA | United States | 2012-Aug-02 | A/Indiana/90/2012 | Indiana State Department of Health Laboratories | Centers for Disease Control and Prevention | |
| EPI397677 | HA | United States | 2012-Sep-04 | A/Ohio/83/2012 | Ohio Department of Health Laboratories | Centers for Disease Control and Prevention | Deng,Y-M; Iannello,P; Caldwell,N; Jelley,L; Komadina,N |
| EPI397751 | HA | United States | 2012-Aug-12 | A/Ohio/87/2012 | Ohio Department of Health Laboratories | Centers for Disease Control and Prevention | |
| EPI397759 | HA | United States | 2012-Aug-22 | A/Wisconsin/35/2012 | Wisconsin State Laboratory of Hygiene | Centers for Disease Control and Prevention | |
| EPI397876 | HA | United States | 2012-Apr-15 | A/Texas/50/2012 | Texas Department of State Health Services-Laboratory Services | Centers for Disease Control and Prevention | Deng,Y-M; Iannello,P; Caldwell,N; Komadina,N. |
| EPI397883 | HA | United States | 2012-Jul-09 | A/Hawaii/22/2012 | State of Hawaii Department of Health | Centers for Disease Control and Prevention | Deng,Y-M; Iannello,P; Caldwell,N; Komadina,N. |
| EPI397905 | HA | United States | 2012-Jul-12 | A/Hawaii/03/2012 | State of Hawaii Department of Health | Centers for Disease Control and Prevention | Deng,Y-M; Iannello,P; Caldwell,N; Komadina,N. |
| EPI397913 | HA | United States | 2012-Aug-04 | A/Indiana/123/2012 | Indiana State Department of Health Laboratories | Centers for Disease Control and Prevention | Deng,Y-M; Iannello,P; Caldwell,N; Komadina,N. |
| EPI397925 | HA | United States | 2012-Jul-27 | A/Ohio/18/2012 | Ohio Department of Health Laboratories | Centers for Disease Control and Prevention | |
| EPI397961 | HA | United States | 2012-Sep-10 | A/Ohio/88/2012 | Ohio Department of Health Laboratories | Centers for Disease Control and Prevention | Deng,Y-M; Iannello,P; Caldwell,N; Komadina,N. |
| EPI397962 | HA | United States | 2012-Aug-20 | A/Wisconsin/30/2012 | Wisconsin State Laboratory of Hygiene | Centers for Disease Control and Prevention | Deng,Y-M; Caldwell,N; Iannello,P; Komadina,N |

| | | | | | | | |
|---|---|---|---|---|---|---|---|
| EPI397997 | HA | United States | 2012-Jul-28 | A/Ohio/17/2012 | Ohio Department of Health Laboratories | Centers for Disease Control and Prevention | |
| EPI398000 | HA | United States | 2012-Aug-06 | A/Ohio/43/2012 | Ohio Department of Health Laboratories | Centers for Disease Control and Prevention | Shu, B.; Garten, R.; Emery, S.; Balish, A.; Smith, C.; Barnes, J.; Lindstrom, S.; Klimov, A.; Cox, N. |
| EPI398037 | HA | United States | 2012-Jul-29 | A/Ohio/19/2012 | Ohio Department of Health Laboratories | Centers for Disease Control and Prevention | Deng,Y-M.; Iannello,P.; Caldwell,N.; Leang,S-K; |
| EPI398040 | HA | United States | 2012-Aug-05 | A/Ohio/57/2012 | Ohio Department of Health Laboratories | Centers for Disease Control and Prevention | Shu, B.; Garten, R.; Emery, S.; Balish, A.; Smith, C.; Barnes, J.; Lindstrom, S.; Klimov, A.; Cox, N. |
| EPI398417 | HA | United States | 2012-Apr-15 | A/Texas/50/2012 | Texas Department of State Health Services-Laboratory Services | Centers for Disease Control and Prevention | |
| EPI398424 | HA | United States | 2012-Jul-09 | A/Hawaii/22/2012 | State of Hawaii Department of Health | Centers for Disease Control and Prevention | Deng,Y-M.; Iannello,P.; Caldwell,N.; Leang,S-K; |
| EPI404890 | HA | United States | 2012-Nov-17 | A/Iowa/16/2012 | Iowa State Hygienic Laboratory | Centers for Disease Control and Prevention | Lan,Yu;Wang,Dayan;Li,Xiyan;Huang,Weijuan;Zhao,Xiang;Chen,Yaoyao;Yang,Lei;Shu,Yuelong |
| EPI404971 | HA | United States | 2012-Oct-18 | A/Iowa/14/2012 | Iowa State Hygienic Laboratory | Centers for Disease Control and Prevention | Lan,Yu;Wang,Dayan;Li,Xiyan;Huang,Weijuan;Zhao,Xiang;Chen,Yaoyao;Yang,Lei;Shu,Yuelong |
| EPI404989 | HA | United States | 2012-Apr-21 | A/Pennsylvania/09/2012 | Pennsylvania Department of Health | Centers for Disease Control and Prevention | |
| EPI404996 | HA | United States | 2012-Apr-21 | A/Pennsylvania/09/2012 | Centers for Disease Control and Prevention | Centers for Disease Control and Prevention | |
| EPI407279 | HA | United States | 2012-Nov-13 | A/Minnesota/18/2012 | Minnesota Department of Health | Centers for Disease Control and Prevention | |
| EPI416138 | HA | United States | 2012-Nov-18 | A/Michigan/32/2012 | Michigan Department of Community Health | Centers for Disease Control and Prevention | |
| EPI416145 | HA | United States | 2012-Dec-03 | A/Michigan/33/2012 | Michigan Department of Community Health | Centers for Disease Control and Prevention | |
| EPI417231 | HA | United States | 2012-Oct-22 | A/Arizona/09/2012 | Arizona Department of Health Services | Centers for Disease Control and Prevention | |
| EPI418017 | HA | Australia | 2011-Oct-24 | A/Victoria/361/2011 | WHO Collaborating Centre for Reference and Research on Influenza | Centers for Disease Control and Prevention | |
| EPI426140 | HA | United States | 2012-Oct-20 | A/New York/39/2012 | New York State Department of Health | Centers for Disease Control and Prevention | |
| EPI436261 | HA | United States | 2012-Oct-20 | A/New York/39/2012 | New York State Department of Health | Centers for Disease Control and Prevention | |
| EPI443697 | HA | United States | 2003-Feb-13 | A/Wyoming/03/2003 | Iowa State Hygienic Laboratory | Centers for Disease Control and Prevention | |
| EPI443709 | HA | United States | 2013-Jan-07 | A/Washington/07/2013 | Spokane Regional Health District | Centers for Disease Control and Prevention | |
| EPI443733 | HA | United States | 2013-Feb-20 | A/Alaska/02/2013 | Alaska State Virology Lab | Centers for Disease Control and Prevention | |
| EPI445653 | HA | United States | 2013-Feb-20 | A/Alaska/02/2013 | Alaska State Virology Lab | Centers for Disease Control and Prevention | |
| EPI445712 | HA | United States | 2012-Oct-20 | A/New York/39/2012 | New York State Department of Health | Centers for Disease Control and Prevention | |

| | | | | | | | |
|---|---|---|---|---|---|---|---|
| EPI450257 | HA | Australia | 2013-Jan-14 | A/NEWCASTLE/1/2013 | John Hunter Hospital, Virology Unit, Clinical Microbiology | WHO Collaborating Centre for Reference and Research on Influenza | |
| EPI450293 | HA | Australia | 2013-Jan-09 | A/SOUTH AUSTRALIA/2/2013 | Institute of Medical and Veterinary Science (IMVS) | WHO Collaborating Centre for Reference and Research on Influenza | |
| EPI450524 | HA | China | 2013-Apr-26 | A/Nanjing/M1/2013 | Jiangsu Provincial Center for Disease Control and Prevention | Jiangsu Provincial Center for Disease Control & Prevention | |
| EPI466829 | HA | United States | 2013-Jul-05 | A/Utah/07/2013 | Utah Department of Health | Centers for Disease Control and Prevention | |
| EPI466845 | HA | Brazil | 2013-May-22 | A/Brazil/0328/2013 | Instituto Adolfo Lutz | Centers for Disease Control and Prevention | |
| EPI466866 | HA | United States | 2013-Mar-06 | A/Maryland/04/2013 | Maryland Department of Health and Mental Hygiene | Centers for Disease Control and Prevention | |
| EPI466889 | HA | Ukraine | 2013-Apr-06 | A/Ukraine/387/2013 | Institute of Epidemiology and Infectious Diseases AMS of Ukraine | Centers for Disease Control and Prevention | |
| EPI466905 | HA | American Samoa | 2013-Feb-22 | A/American Samoa/4786/2013 | State of Hawaii Department of Health | Centers for Disease Control and Prevention | |
| EPI467440 | HA | American Samoa | 2013-Feb-22 | A/American Samoa/4786/2013 | State of Hawaii Department of Health | Centers for Disease Control and Prevention | |
| EPI467448 | HA | Australia | 2013-Jun-17 | A/Victoria/505/2013 | WHO Collaborating Centre for Reference and Research on Influenza | Centers for Disease Control and Prevention | |
| EPI477479 | HA | Sri Lanka | 2013-Feb-27 | A/SRI LANKA/27/2013 | Medical Research Institute | WHO Collaborating Centre for Reference and Research on Influenza | Deng,Y-M., Iannello,P., Caldwell,N., Lau, H.,Komadina,N. |
| EPI477495 | HA | Australia | 2013-Mar-04 | A/SYDNEY/506/2013 | Prince of Wales Hospital | WHO Collaborating Centre for Reference and Research on Influenza | Deng,Y-M., Iannello,P., Caldwell,N., Lau, H.,Komadina,N. |
| EPI484516 | HA | United States | 2013-Aug-15 | A/Arkansas/13/2013 | Arkansas Department of Health | Centers for Disease Control and Prevention | |
| EPI484541 | HA | United States | 2013-Jul-10 | A/Massachusetts/11/2013 | Massachusetts Department of Public Health | Centers for Disease Control and Prevention | |
| EPI484551 | HA | Costa Rica | 2013-Jul-23 | A/Costa Rica/4009/2013 | Laboratorio Nacional de Influenza | Centers for Disease Control and Prevention | |
| EPI484565 | HA | United States | 2013-Jul-14 | A/Georgia/03/2013 | Georgia Public Health Laboratory | Centers for Disease Control and Prevention | |
| EPI484573 | HA | Bangladesh | 2013-May-07 | A/Bangladesh/9334/2013 | Institute of Epidemiology Disease Control and Research (IEDCR) & Bangladesh National Influenza Centre (NIC) | Centers for Disease Control and Prevention | |
| EPI484635 | HA | Sweden | 2011-Dec-26 | A/Stockholm/42/2011 | | Swedish National Institute for Communicable Disease Control | |
| EPI484643 | HA | Sweden | 2012-Apr-14 | A/Stockholm/19/2012 | | Swedish National Institute for Communicable Disease Control | |
| EPI484651 | HA | Sweden | 2011-Oct-31 | A/Stockholm/29/2011 | | Swedish National Institute for Communicable Disease Control | |
| EPI484659 | HA | Sweden | 2012-Jan-12 | A/Stockholm/12/2012 | | Swedish National Institute for Communicable Disease Control | Deng,Y-M., Iannello,P.,Caldwell,N., |
| EPI484667 | HA | Sweden | 2011-Dec-28 | A/Stockholm/43/2011 | | Swedish National Institute for Communicable Disease Control | Deng,Y-M., Iannello,P.,Caldwell,N., |

| | | | | | | |
|---|---|---|---|---|---|---|
| EPI484675 | HA | Sweden | 2012-Mar-11 | A/Stockholm/14/2012 | | Swedish National Institute for Communicable Disease Control |
| EPI484683 | HA | Sweden | 2011-Oct-06 | A/Stockholm/26/2011 | | Swedish National Institute for Communicable Disease Control |
| EPI484737 | HA | Costa Rica | 2013-Jul-23 | A/Costa Rica/4009/2013 | Laboratorio Nacional de Influenza | Centers for Disease Control and Prevention |
| EPI485800 | HA | Costa Rica | 2013-Jul-26 | A/Costa Rica/4700/2013 | Laboratorio Nacional de Influenza | Centers for Disease Control and Prevention |
| EPI485808 | HA | Costa Rica | 2013-Jul-26 | A/Costa Rica/4700/2013 | Laboratorio Nacional de Influenza | Centers for Disease Control and Prevention |
| EPI489300 | HA | United States | 2013-Jul-05 | A/Utah/07/2013 | Utah Department of Health | Centers for Disease Control and Prevention |
| EPI489312 | HA | United States | 2013-Jul-10 | A/Massachusetts/11/2013 | Massachusetts Department of Public Health | Centers for Disease Control and Prevention |
| EPI490093 | HA | United States | 2012-Oct-20 | A/New York/39/2012 | New York State Department of Health | Centers for Disease Control and Prevention |
| EPI492686 | HA | United States | 2013-Jul-10 | A/Massachusetts/11/2013 | Massachusetts Department of Public Health | Centers for Disease Control and Prevention |
| EPI492690 | HA | Costa Rica | 2013-Jul-26 | A/Costa Rica/4700/2013 | Laboratorio Nacional de Influenza | Centers for Disease Control and Prevention |
| EPI492717 | HA | Bangladesh | 2013-May-12 | A/Bangladesh/9269/2013 | Institute of Epidemiology Disease Control and Research (IEDCR) & Bangladesh National Influenza Centre (NIC) | Centers for Disease Control and Prevention |
| EPI497661 | HA | Australia | 2013-Jun-16 | A/Victoria/506/2013 | WHO Collaborating Centre for Reference and Research on Influenza | Centers for Disease Control and Prevention |
| EPI497668 | HA | United States | 2013-Jul-05 | A/Utah/07/2013 | Utah Department of Health | Centers for Disease Control and Prevention |
| EPI499306 | HA | United States | 2013-Nov-09 | A/Pennsylvania/12/2013 | Pennsylvania Department of Health | Centers for Disease Control and Prevention |
| EPI499308 | HA | Hong Kong (SAR) | 2013-Oct-12 | A/Hong Kong/5643/2013 | Government Virus Unit | Centers for Disease Control and Prevention |
| EPI499365 | HA | United States | 2003-Feb-13 | A/Wyoming/03/2003 | Iowa State Hygienic Laboratory | Centers for Disease Control and Prevention |
| EPI503938 | HA | United States | 2013-Nov-29 | A/Washington/18/2013 | Washington State Public Health Laboratory | Centers for Disease Control and Prevention |
| EPI503951 | HA | United States | 2013-Nov-29 | A/Washington/18/2013 | Washington State Public Health Laboratory | Centers for Disease Control and Prevention |
| EPI504746 | HA | Sweden | 2013-Dec-21 | A/Stockholm/40/2013 | | Public Health Agency of Sweden |
| EPI504749 | HA | Sweden | 2014-Jan-02 | A/Stockholm/2/2014 | | Public Health Agency of Sweden |
| EPI506693 | HA | China | 2013-Jan-29 | A/Hangzhou/B48/2013 | Hangzhou Center for Disease Control and Prevention | Hangzhou Center for Disease Control and Prevention |
| EPI506701 | HA | China | 2013-Feb-17 | A/Hangzhou/A121/2013 | Hangzhou Center for Disease Control and Prevention | Hangzhou Center for Disease Control and Prevention |
| EPI506709 | HA | China | 2013-Feb-19 | A/Hangzhou/A126/2013 | Hangzhou Center for Disease Control and Prevention | Hangzhou Center for Disease Control and Prevention |
| EPI506717 | HA | China | 2013-Feb-25 | A/Hangzhou/A155/2013 | Hangzhou Center for Disease Control and Prevention | Hangzhou Center for Disease Control and Prevention |

| | | | | | | | |
|---|---|---|---|---|---|---|---|
| EPI506725 | HA | China | 2013-Mar-01 | A/Hangzhou/A158/2013 | Hangzhou Center for Disease Control and Prevention | Hangzhou Center for Disease Control and Prevention | |
| EPI506733 | HA | China | 2013-Jul-08 | A/Hangzhou/A429/2013 | Hangzhou Center for Disease Control and Prevention | Hangzhou Center for Disease Control and Prevention | |
| EPI506741 | HA | China | 2013-Jul-29 | A/Hangzhou/A469/2013 | Hangzhou Center for Disease Control and Prevention | Hangzhou Center for Disease Control and Prevention | |
| EPI506749 | HA | China | 2013-Aug-15 | A/Hangzhou/A511/2013 | Hangzhou Center for Disease Control and Prevention | Hangzhou Center for Disease Control and Prevention | |
| EPI506757 | HA | China | 2013-Aug-27 | A/Hangzhou/A544/2013 | Hangzhou Center for Disease Control and Prevention | Hangzhou Center for Disease Control and Prevention | |
| EPI506765 | HA | China | 2013-Sep-15 | A/Hangzhou/A601/2013 | Hangzhou Center for Disease Control and Prevention | Hangzhou Center for Disease Control and Prevention | |
| EPI506773 | HA | China | 2013-Oct-10 | A/Hangzhou/A649/2013 | Hangzhou Center for Disease Control and Prevention | Hangzhou Center for Disease Control and Prevention | |
| EPI506781 | HA | China | 2013-Oct-10 | A/Hangzhou/A662/2013 | Hangzhou Center for Disease Control and Prevention | Hangzhou Center for Disease Control and Prevention | |
| EPI506789 | HA | China | 2013-Oct-10 | A/Hangzhou/A666/2013 | Hangzhou Center for Disease Control and Prevention | Hangzhou Center for Disease Control and Prevention | Galiano,M. |
| EPI506797 | HA | China | 2013-Oct-15 | A/Hangzhou/A683/2013 | Hangzhou Center for Disease Control and Prevention | Hangzhou Center for Disease Control and Prevention | Galiano,M. |
| EPI506805 | HA | China | 2013-Oct-15 | A/Hangzhou/B599/2013 | Hangzhou Center for Disease Control and Prevention | Hangzhou Center for Disease Control and Prevention | Galiano,M. |
| EPI506813 | HA | China | 2013-Oct-15 | A/Hangzhou/B600/2013 | Hangzhou Center for Disease Control and Prevention | Hangzhou Center for Disease Control and Prevention | Galiano,M. |
| EPI506821 | HA | China | 2013-Oct-18 | A/Hangzhou/A691/2013 | Hangzhou Center for Disease Control and Prevention | Hangzhou Center for Disease Control and Prevention | Galiano,M. |
| EPI506829 | HA | China | 2013-Oct-22 | A/Hangzhou/A698/2013 | Hangzhou Center for Disease Control and Prevention | Hangzhou Center for Disease Control and Prevention | Galiano,M. |
| EPI506837 | HA | China | 2013-Nov-15 | A/Hangzhou/A765/2013 | Hangzhou Center for Disease Control and Prevention | Hangzhou Center for Disease Control and Prevention | Galiano,M. |
| EPI506845 | HA | China | 2013-Nov-15 | A/Hangzhou/A773/2013 | Hangzhou Center for Disease Control and Prevention | Hangzhou Center for Disease Control and Prevention | Galiano,M. |
| EPI506853 | HA | China | 2013-Nov-15 | A/Hangzhou/B690/2013 | Hangzhou Center for Disease Control and Prevention | Hangzhou Center for Disease Control and Prevention | Galiano,M. |
| EPI506861 | HA | China | 2013-Nov-18 | A/Hangzhou/A776/2013 | Hangzhou Center for Disease Control and Prevention | Hangzhou Center for Disease Control and Prevention | Galiano,M. |
| EPI506869 | HA | China | 2013-Nov-18 | A/Hangzhou/A779/2013 | Hangzhou Center for Disease Control and Prevention | Hangzhou Center for Disease Control and Prevention | Galiano,M. |
| EPI506877 | HA | China | 2013-Nov-21 | A/Hangzhou/A784/2013 | Hangzhou Center for Disease Control and Prevention | Hangzhou Center for Disease Control and Prevention | Galiano,M. |
| EPI506885 | HA | China | 2013-Nov-21 | A/Hangzhou/A789/2013 | Hangzhou Center for Disease Control and Prevention | Hangzhou Center for Disease Control and Prevention | Galiano,M. |
| EPI506893 | HA | China | 2013-Nov-21 | A/Hangzhou/B709/2013 | Hangzhou Center for Disease Control and Prevention | Hangzhou Center for Disease Control and Prevention | Galiano,M. |
| EPI506901 | HA | China | 2013-Nov-25 | A/Hangzhou/A796/2013 | Hangzhou Center for Disease Control and Prevention | Hangzhou Center for Disease Control and Prevention | Galiano,M. |
| EPI506909 | HA | China | 2013-Nov-25 | A/Hangzhou/A800/2013 | Hangzhou Center for Disease Control and Prevention | Hangzhou Center for Disease Control and Prevention | Galiano,M. |
| EPI506917 | HA | China | 2013-Nov-25 | A/Hangzhou/B718/2013 | Hangzhou Center for Disease Control and Prevention | Hangzhou Center for Disease Control and Prevention | Galiano,M. |

| | | | | | | | |
|---|---|---|---|---|---|---|---|
| EPI506925 | HA | China | 2013-Nov-25 | A/Hangzhou/B721/2013 | Hangzhou Center for Disease Control and Prevention | Hangzhou Center for Disease Control and Prevention | Galiano,M. |
| EPI506933 | HA | China | 2013-Nov-27 | A/Hangzhou/A803/2013 | Hangzhou Center for Disease Control and Prevention | Hangzhou Center for Disease Control and Prevention | Galiano,M. |
| EPI506941 | HA | China | 2013-Nov-27 | A/Hangzhou/A806/2013 | Hangzhou Center for Disease Control and Prevention | Hangzhou Center for Disease Control and Prevention | Galiano,M. |
| EPI506949 | HA | China | 2013-Nov-27 | A/Hangzhou/B728/2013 | Hangzhou Center for Disease Control and Prevention | Hangzhou Center for Disease Control and Prevention | Galiano,M. |
| EPI506957 | HA | China | 2013-Nov-27 | A/Hangzhou/B735/2013 | Hangzhou Center for Disease Control and Prevention | Hangzhou Center for Disease Control and Prevention | Galiano,M. |
| EPI506965 | HA | China | 2013-Dec-04 | A/Hangzhou/A812/2013 | Hangzhou Center for Disease Control and Prevention | Hangzhou Center for Disease Control and Prevention | Galiano,M. |
| EPI506973 | HA | China | 2013-Dec-04 | A/Hangzhou/A816/2013 | Hangzhou Center for Disease Control and Prevention | Hangzhou Center for Disease Control and Prevention | Galiano,M. |
| EPI506981 | HA | China | 2013-Dec-04 | A/Hangzhou/A825/2013 | Hangzhou Center for Disease Control and Prevention | Hangzhou Center for Disease Control and Prevention | Galiano,M. |
| EPI507594 | HA | Kazakhstan | 2013-Jan-27 | A/Almaty/2958/2013 | National Institute for Medical Research | Centers for Disease Control and Prevention | Galiano,M. |
| EPI508248 | HA | Sweden | 2014-Feb-01 | A/Stockholm/4/2014 | | Public Health Agency of Sweden | Galiano,M. |
| EPI508269 | HA | Sweden | 2014-Feb-06 | A/Gavle/1/2014 | | Public Health Agency of Sweden | Galiano,M. |
| EPI508273 | HA | Sweden | 2014-Feb-12 | A/Sweden/1/2014 | | Public Health Agency of Sweden | Galiano,M. |
| EPI513291 | HA | Costa Rica | 2013-Jul-26 | A/Costa Rica/4700/2013 | Laboratorio Nacional de Influenza | Centers for Disease Control and Prevention | Galiano,M. |
| EPI515296 | HA | Sweden | 2014-Feb-06 | A/Stockholm/6/2014 | | Public Health Agency of Sweden | Galiano,M. |
| EPI515774 | HA | Sweden | 2014-Feb-06 | A/Stockholm/5/2014 | | Public Health Agency of Sweden | Galiano,M. |
| EPI516458 | HA | Sweden | 2014-Feb-21 | A/Stockholm/13/2014 | | Public Health Agency of Sweden | Galiano,M. |
| EPI516676 | HA | United States | 2014-Jan-12 | A/Maryland/03/2014 | Maryland Department of Health and Mental Hygiene | Centers for Disease Control and Prevention | Galiano,M. |
| EPI516844 | HA | United States | 2013-Oct-01 | A/Iowa/04/2013 | Iowa State Hygienic Laboratory | Centers for Disease Control and Prevention | Galiano,M. |
| EPI526757 | HA | Australia | 2013-Oct-16 | A/PERTH/258/2013 | Pathwest QE II Medical Centre | WHO Collaborating Centre for Reference and Research on Influenza | Galiano,M. |
| EPI526801 | HA | New Zealand | 2013-Sep-18 | A/NORTH WEST AUCKLAND/6/2013 | Institute of Environmental Science and Research | WHO Collaborating Centre for Reference and Research on Influenza | Galiano,M. |
| EPI529530 | HA | Australia | 2013-Sep-09 | A/BRISBANE/165/2013 | Queensland Health Scientific Services | WHO Collaborating Centre for Reference and Research on Influenza | Galiano,M. |
| EPI529546 | HA | Malaysia | 2013-Nov-13 | A/MALAYSIA/10/2013 | Institut Penyelidikan Perubatan | WHO Collaborating Centre for Reference and Research on Influenza | Galiano,M. |
| EPI529557 | HA | Australia | 2013-Nov-21 | A/SYDNEY/83/2013 | Clinical Virology Unit, CDIM | WHO Collaborating Centre for Reference and Research on Influenza | Galiano,M. |
| EPI529568 | HA | Thailand | 2013-Nov-05 | A/CHIANG RAI/337/2013 | WHO National Influenza Centre, National Institute of Medical Research (NIMR) | WHO Collaborating Centre for Reference and Research on Influenza | Galiano,M. |

| | | | | | | | |
|---|---|---|---|---|---|---|---|
| EPI529587 | HA | Thailand | 2013-Nov-18 | A/SONG KHLA/3076/2013 | WHO National Influenza Centre, National Institute of Medical Research (NIMR) | WHO Collaborating Centre for Reference and Research on Influenza | Galiano,M. |
| EPI531450 | HA | Sweden | 2014-Apr-07 | A/Stockholm/15/2014 | | Public Health Agency of Sweden | Galiano,M. |
| EPI537000 | HA | United States | 2014-Mar-11 | A/Nebraska/04/2014 | Nebraska Public Health Lab | Centers for Disease Control and Prevention | Galiano,M. |
| EPI537007 | HA | United States | 2013-Dec-27 | A/Montana/27/2013 | Montana Public Health Laboratory | Centers for Disease Control and Prevention | Galiano,M. |
| EPI537015 | HA | United States | 2012-Apr-15 | A/Texas/50/2012 | Texas Department of State Health Services-Laboratory Services | Centers for Disease Control and Prevention | Galiano,M. |
| EPI537022 | HA | United States | 2014-Jan-16 | A/California/02/2014 | California Department of Health Services | Centers for Disease Control and Prevention | Galiano,M. |
| EPI537030 | HA | United States | 2014-Mar-11 | A/Nebraska/04/2014 | Nebraska Public Health Lab | Centers for Disease Control and Prevention | Galiano,M. |
| EPI537038 | HA | Palau | 2014-Mar-26 | A/Palau/6759/2014 | State of Hawaii Department of Health | Centers for Disease Control and Prevention | Galiano,M. |
| EPI537046 | HA | United States | 2012-Oct-20 | A/New York/39/2012 | New York State Department of Health | Centers for Disease Control and Prevention | Galiano,M. |
| EPI537054 | HA | United States | 2013-Dec-29 | A/Louisiana/39/2013 | Louisiana Department of Health and Hospitals | Centers for Disease Control and Prevention | Galiano,M. |
| EPI537081 | HA | United States | 2014-Jan-21 | A/New York/05/2014 | New York State Department of Health | Centers for Disease Control and Prevention | Galiano,M. |
| EPI537985 | HA | United States | 2014-Jan-21 | A/New York/05/2014 | New York State Department of Health | Centers for Disease Control and Prevention | Galiano,M. |
| EPI540526 | HA | Switzerland | 2013-Dec-06 | A/Switzerland/9715293/2013 | National Institute for Medical Research | Centers for Disease Control and Prevention | Galiano,M. |
| EPI541636 | HA | United States | 2014-Jun-02 | A/Alaska/18/2014 | Alaska State Virology Lab | Centers for Disease Control and Prevention | Galiano,M. |
| EPI541651 | HA | United States | 2013-Nov-29 | A/Washington/18/2013 | Washington State Public Health Laboratory | Centers for Disease Control and Prevention | Galiano,M. |
| EPI541659 | HA | Switzerland | 2013-Jan-01 | A/Switzerland/9715293/2013 | National Institute for Medical Research | Centers for Disease Control and Prevention | Galiano,M. |
| EPI542216 | HA | Sweden | 2014-Jul-23 | A/Stockholm/17/2014 | | Public Health Agency of Sweden | Galiano,M. |
| EPI542224 | HA | Sweden | 2014-Aug-16 | A/Stockholm/18/2014 | | Public Health Agency of Sweden | Galiano,M. |
| EPI543234 | HA | Sweden | 2014-Feb-06 | A/Stockholm/06/2014 | National Institute for Medical Research | Centers for Disease Control and Prevention | Galiano,M. |
| EPI543523 | HA | Brazil | 2014-Jan-20 | A/Rio Grande do Sul/23/2014 | LACEN/RS - Laboratório Central de Saúde Pública do Rio Grande do Sul | Instituto Oswaldo Cruz FIOCRUZ - Laboratory of Respiratory Viruses and Measles (LVRS) | Galiano,M. |
| EPI543531 | HA | Brazil | 2014-Jan-20 | A/Rio Grande do Sul/23/2014 | LACEN/RS - Laboratório Central de Saúde Pública do Rio Grande do Sul | Instituto Oswaldo Cruz FIOCRUZ - Laboratory of Respiratory Viruses and Measles (LVRS) | Galiano,M. |
| EPI543551 | HA | Brazil | 2014-Mar-17 | A/Parana/87/2014 | Laboratório Central do Estado do Paraná - LACEN/PR | Instituto Oswaldo Cruz FIOCRUZ - Laboratory of Respiratory Viruses and Measles (LVRS) | Galiano,M. |
| EPI543666 | HA | United States | 2014-Aug-27 | A/Maryland/25/2014 | Maryland Department of Health and Mental Hygiene | Centers for Disease Control and Prevention | Galiano,M. |
| EPI544084 | HA | Australia | 2014-Jun-29 | A/South Australia/55/2014 | WHO Collaborating Centre for Reference and Research on Influenza | Centers for Disease Control and Prevention | Galiano,M. |

| | | | | | | | |
|---|---|---|---|---|---|---|---|
| EPI544087 | HA | Australia | 2014-Jun-29 | A/South Australia/55/2014 | WHO Collaborating Centre for Reference and Research on Influenza | Centers for Disease Control and Prevention | Galiano,M. |
| EPI544405 | HA | United States | 2014-Sep-17 | A/Minnesota/21/2014 | Minnesota Department of Health | Centers for Disease Control and Prevention | Galiano,M. |
| EPI544413 | HA | United States | 2014-Sep-18 | A/Michigan/13/2014 | Michigan Department of Community Health | Centers for Disease Control and Prevention | Galiano,M. |
| EPI544421 | HA | United States | 2014-Oct-01 | A/Texas/37/2014 | Texas Department of State Health Services-Laboratory Services | Centers for Disease Control and Prevention | Galiano,M. |
| EPI544429 | HA | United States | 2014-Oct-06 | A/Texas/38/2014 | Texas Department of State Health Services-Laboratory Services | Centers for Disease Control and Prevention | Galiano,M. |
| EPI544437 | HA | United States | 2014-Sep-16 | A/Michigan/12/2014 | Michigan Department of Community Health | Centers for Disease Control and Prevention | Galiano,M. |
| EPI544445 | HA | United States | 2014-Sep-18 | A/New Mexico/27/2014 | New Mexico Department of Health | Centers for Disease Control and Prevention | |
| EPI544453 | HA | United States | 2014-Sep-23 | A/Florida/35/2014 | Florida Department of Health-Tampa | Centers for Disease Control and Prevention | |
| EPI544461 | HA | United States | 2014-Sep-13 | A/Michigan/11/2014 | Michigan Department of Community Health | Centers for Disease Control and Prevention | |
| EPI544476 | HA | United States | 2014-Sep-17 | A/Maryland/27/2014 | Maryland Department of Health and Mental Hygiene | Centers for Disease Control and Prevention | |
| EPI544484 | HA | United States | 2014-Sep-13 | A/Pennsylvania/40/2014 | Pennsylvania Department of Health | Centers for Disease Control and Prevention | |
| EPI544492 | HA | United States | 2014-Sep-19 | A/Michigan/14/2014 | Michigan Department of Community Health | Centers for Disease Control and Prevention | |
| EPI544500 | HA | United States | 2014-Sep-24 | A/Michigan/15/2014 | Michigan Department of Community Health | Centers for Disease Control and Prevention | |
| EPI544508 | HA | United States | 2014-Sep-20 | A/Minnesota/22/2014 | Minnesota Department of Health | Centers for Disease Control and Prevention | |
| EPI544519 | HA | Sweden | 2014-Feb-06 | A/Stockholm/06/2014 | National Institute for Medical Research | Centers for Disease Control and Prevention | |
| EPI544526 | HA | United States | 2014-Apr-15 | A/North Carolina/13/2014 | North Carolina State Laboratory of Public Health | Centers for Disease Control and Prevention | |
| EPI544663 | HA | Sweden | 2014-Oct-15 | A/Stockholm/21/2014 | | Public Health Agency of Sweden | |
| EPI545581 | HA | Thailand | 2014-Jul-20 | A/Tak/1845/2014 | WHO National Influenza Centre, National Institute of Medical Research (NIMR) | Centers for Disease Control and Prevention | |
| EPI545589 | HA | United States | 2014-Oct-04 | A/Kentucky/10/2014 | Kentucky Division of Laboratory Services | Centers for Disease Control and Prevention | |
| EPI545597 | HA | United States | 2014-Sep-25 | A/Wisconsin/26/2014 | Wisconsin State Laboratory of Hygiene | Centers for Disease Control and Prevention | |
| EPI545604 | HA | United States | 2014-Sep-25 | A/Wisconsin/25/2014 | Wisconsin State Laboratory of Hygiene | Centers for Disease Control and Prevention | |
| EPI545612 | HA | United States | 2014-Sep-08 | A/North Carolina/20/2014 | North Carolina State Laboratory of Public Health | Centers for Disease Control and Prevention | |
| EPI545622 | HA | United States | 2014-Oct-09 | A/Wyoming/08/2014 | Wyoming Public Health Laboratory | Centers for Disease Control and Prevention | |
| EPI545630 | HA | United States | 2014-Oct-09 | A/Wyoming/09/2014 | Wyoming Public Health Laboratory | Centers for Disease Control and Prevention | |

| | | | | | | | |
|---|---|---|---|---|---|---|---|
| EPI545643 | HA | United States | 2014-Oct-14 | A/Mississippi/06/2014 | Mississippi Public Health Laboratory | Centers for Disease Control and Prevention | |
| EPI545648 | HA | United States | 2014-Oct-15 | A/Mississippi/03/2014 | Mississippi Public Health Laboratory | Centers for Disease Control and Prevention | |
| EPI545656 | HA | United States | 2014-Oct-16 | A/Mississippi/04/2014 | Mississippi Public Health Laboratory | Centers for Disease Control and Prevention | |
| EPI545664 | HA | United States | 2014-Oct-14 | A/Mississippi/05/2014 | Mississippi Public Health Laboratory | Centers for Disease Control and Prevention | |
| EPI545672 | HA | United States | 2014-Oct-04 | A/Colorado/15/2014 | Colorado Department of Health Lab | Centers for Disease Control and Prevention | |
| EPI545694 | HA | United States | 2014-Oct-09 | A/Maine/04/2014 | Maine Health and Environmental Testing Laboratory | Centers for Disease Control and Prevention | |
| EPI545702 | HA | United States | 2014-Sep-25 | A/Pennsylvania/45/2014 | Pennsylvania Department of Health | Centers for Disease Control and Prevention | Deng, Yi-Mo |
| EPI545710 | HA | United States | 2014-Sep-30 | A/Florida/34/2014 | Florida Department of Health-Jacksonville | Centers for Disease Control and Prevention | |
| EPI545718 | HA | United States | 2014-Oct-07 | A/Florida/39/2014 | Florida Department of Health-Jacksonville | Centers for Disease Control and Prevention | |
| EPI545726 | HA | United States | 2014-Oct-07 | A/Florida/40/2014 | Florida Department of Health-Jacksonville | Centers for Disease Control and Prevention | |
| EPI545734 | HA | United States | 2014-Oct-03 | A/Florida/36/2014 | Florida Department of Health-Jacksonville | Centers for Disease Control and Prevention | |
| EPI545742 | HA | United States | 2014-Oct-07 | A/Florida/38/2014 | Florida Department of Health-Jacksonville | Centers for Disease Control and Prevention | |
| EPI545749 | HA | United States | 2014-Oct-07 | A/Florida/37/2014 | Florida Department of Health-Jacksonville | Centers for Disease Control and Prevention | |
| EPI545757 | HA | United States | 2014-Sep-27 | A/Pennsylvania/44/2014 | Pennsylvania Department of Health | Centers for Disease Control and Prevention | |
| EPI545772 | HA | United States | 2014-Oct-06 | A/North Carolina/21/2014 | North Carolina State Laboratory of Public Health | Centers for Disease Control and Prevention | |
| EPI547601 | HA | Palau | 2014-Mar-26 | A/Palau/6775/2014 | State of Hawaii Department of Health | Centers for Disease Control and Prevention | |
| EPI547711 | HA | Argentina | 2014-Jul-04 | A/Argentina/11071/2014 | CEMIC University Hospital | Centers for Disease Control and Prevention | |
| EPI547719 | HA | Argentina | 2014-Jul-01 | A/Argentina/11093/2014 | CEMIC University Hospital | Centers for Disease Control and Prevention | |
| EPI547727 | HA | Argentina | 2014-Jul-17 | A/Argentina/11126/2014 | CEMIC University Hospital | Centers for Disease Control and Prevention | |
| EPI547735 | HA | Bolivia, Plurinationial State of | 2014-Jun-17 | A/Bolivia/667/2014 | CENETROP | Centers for Disease Control and Prevention | |
| EPI547743 | HA | Bolivia, Plurinationial State of | 2014-Jun-18 | A/Bolivia/716/2014 | CENETROP | Centers for Disease Control and Prevention | Takashita,Emi; Fujisaki,Seiichiro; Shirakura,Masayuki; Watanabe,Shinji; Odagiri,Takato |
| EPI547751 | HA | Bolivia, Plurinationial State of | 2014-Jun-24 | A/Bolivia/751/2014 | CENETROP | Centers for Disease Control and Prevention | Takashita,Emi; Fujisaki,Seiichiro; Shirakura,Masayuki; Watanabe,Shinji; Odagiri,Takato |
| EPI547766 | HA | United States | 2014-Aug-25 | A/California/15/2014 | California Department of Health Services | Centers for Disease Control and Prevention | Takashita,Emi; Fujisaki,Seiichiro; Shirakura,Masayuki; Watanabe,Shinji; Odagiri,Takato |

| | | | | | | | |
|---|---|---|---|---|---|---|---|
| EPI547774 | HA | United States | 2014-Aug-25 | A/California/16/2014 | California Department of Health Services | Centers for Disease Control and Prevention | |
| EPI547782 | HA | United States | 2014-Aug-11 | A/California/17/2014 | California Department of Health Services | Centers for Disease Control and Prevention | |
| EPI547790 | HA | United States | 2014-Aug-08 | A/California/18/2014 | California Department of Health Services | Centers for Disease Control and Prevention | |
| EPI547798 | HA | United States | 2014-Sep-04 | A/California/19/2014 | California Department of Health Services | Centers for Disease Control and Prevention | |
| EPI547806 | HA | Costa Rica | 2014-Jul-09 | A/Costa Rica/2303/2014 | Laboratorio Nacional de Influenza | Centers for Disease Control and Prevention | |
| EPI547814 | HA | Dominican Republic | 2014-May-28 | A/Dominican Republic/8483/2014 | Laboratorio de Investigacion / Centro de Educacion Medica y Amistad Dominico Japones (CEMADOJA) | Centers for Disease Control and Prevention | |
| EPI547821 | HA | Dominican Republic | 2014-May-22 | A/Dominican Republic/8490/2014 | Laboratorio de Investigacion / Centro de Educacion Medica y Amistad Dominico Japones (CEMADOJA) | Centers for Disease Control and Prevention | |
| EPI547829 | HA | United States | 2014-Sep-23 | A/Florida/35/2014 | Florida Department of Health-Tampa | Centers for Disease Control and Prevention | |
| EPI547837 | HA | United States | 2014-Oct-07 | A/Florida/37/2014 | Florida Department of Health-Jacksonville | Centers for Disease Control and Prevention | |
| EPI547845 | HA | United States | 2014-Oct-07 | A/Florida/38/2014 | Florida Department of Health-Jacksonville | Centers for Disease Control and Prevention | Saito, Reiko; Takemae, Nobuhiro; Saito, Takehiko; Shobugawa, Yugo; Kondo, Hiroki; Hibino, Akinobu; Masaki, Hironori |
| EPI547853 | HA | United States | 2014-Oct-07 | A/Florida/39/2014 | Florida Department of Health-Jacksonville | Centers for Disease Control and Prevention | Saito, Reiko; Takemae, Nobuhiro; Saito, Takehiko; Shobugawa, Yugo; Kondo, Hiroki; Hibino, Akinobu; Makiya, Tomoko |
| EPI547861 | HA | United States | 2014-Oct-07 | A/Florida/40/2014 | Florida Department of Health-Jacksonville | Centers for Disease Control and Prevention | Saito, Reiko; Takemae, Nobuhiro; Saito, Takehiko; Shobugawa, Yugo; Kondo, Hiroki; Hibino, Akinobu; Makiya, Tomoko |
| EPI547869 | HA | United States | 2014-Aug-10 | A/Hawaii/35/2014 | State of Hawaii Department of Health | Centers for Disease Control and Prevention | Saito, Reiko; Takemae, Nobuhiro; Saito, Takehiko; Shobugawa, Yugo; Kondo, Hiroki; Hibino, Akinobu; Shirahige, Yutaka |
| EPI547877 | HA | United States | 2014-Aug-08 | A/Hawaii/36/2014 | State of Hawaii Department of Health | Centers for Disease Control and Prevention | Saito, Reiko; Takemae, Nobuhiro; Saito, Takehiko; Shobugawa, Yugo; Kondo, Hiroki; Hibino, Akinobu; Kawashima, Takashi |
| EPI547885 | HA | United States | 2014-Aug-13 | A/Hawaii/37/2014 | State of Hawaii Department of Health | Centers for Disease Control and Prevention | Saito, Reiko; Takemae, Nobuhiro; Saito, Takehiko; Shobugawa, Yugo; Kondo, Hiroki; Hibino, Akinobu; Sato, Isamu |
| EPI547893 | HA | United States | 2014-Aug-11 | A/Hawaii/38/2014 | State of Hawaii Department of Health | Centers for Disease Control and Prevention | Saito, Reiko; Takemae, Nobuhiro; Saito, Takehiko; Shobugawa, Yugo; Kondo, Hiroki; Hibino, |
| EPI547901 | HA | United States | 2014-Aug-13 | A/Hawaii/39/2014 | State of Hawaii Department of Health | Centers for Disease Control and Prevention | Saito, Reiko; Takemae, Nobuhiro; Saito, Takehiko; Shobugawa, Yugo; Kondo, Hiroki; Hibino, |

| | | | | | | | |
|---|---|---|---|---|---|---|---|
| EPI547909 | HA | United States | 2014-Oct-09 | A/Maine/04/2014 | Maine Health and Environmental Testing Laboratory | Centers for Disease Control and Prevention | Saito, Reiko; Takemae, Nobuhiro; Saito, Takehiko; Shobugawa, Yugo; Kondo, Hiroki; Hibino, |
| EPI547917 | HA | Mexico | 2014-Feb-12 | A/Mexico/711/2014 | Laboratorio de Virus Respiratorio | Centers for Disease Control and Prevention | |
| EPI547925 | HA | United States | 2014-Aug-14 | A/Michigan/08/2014 | Michigan Department of Community Health | Centers for Disease Control and Prevention | |
| EPI547933 | HA | United States | 2014-Sep-01 | A/Michigan/09/2014 | Michigan Department of Community Health | Centers for Disease Control and Prevention | |
| EPI547941 | HA | United States | 2014-Sep-10 | A/Michigan/10/2014 | Michigan Department of Community Health | Centers for Disease Control and Prevention | |
| EPI547949 | HA | United States | 2014-Sep-17 | A/Minnesota/21/2014 | Minnesota Department of Health | Centers for Disease Control and Prevention | |
| EPI547957 | HA | United States | 2014-Sep-20 | A/Minnesota/22/2014 | Minnesota Department of Health | Centers for Disease Control and Prevention | |
| EPI547969 | HA | United States | 2014-Aug-28 | A/New Hampshire/18/2014 | New Hampshire Public Health Laboratories | Centers for Disease Control and Prevention | |
| EPI547977 | HA | United States | 2014-Sep-16 | A/New Jersey/22/2014 | New Jersey Department of Health & Senior Services | Centers for Disease Control and Prevention | |
| EPI547985 | HA | United States | 2014-Sep-18 | A/New Mexico/27/2014 | New Mexico Department of Health | Centers for Disease Control and Prevention | |
| EPI547993 | HA | Australia | 2014-Jun-16 | A/Newcastle/22/2014 | WHO Collaborating Centre for Reference and Research on Influenza | Centers for Disease Control and Prevention | |
| EPI548001 | HA | Nicaragua | 2014-Jun-25 | A/Nicaragua/7402/2014 | Laboratorio de Virologia, Direccion de Microbiologia | Centers for Disease Control and Prevention | |
| EPI548009 | HA | United States | 2014-Oct-06 | A/North Carolina/21/2014 | North Carolina State Laboratory of Public Health | Centers for Disease Control and Prevention | |
| EPI548017 | HA | Paraguay | 2014-Jul-03 | A/Paraguay/0049/2014 | Central Laboratory of Public Health | Centers for Disease Control and Prevention | |
| EPI548025 | HA | Paraguay | 2014-Jul-16 | A/Paraguay/0095/2014 | Central Laboratory of Public Health | Centers for Disease Control and Prevention | |
| EPI548033 | HA | United States | 2014-Sep-13 | A/Pennsylvania/40/2014 | Pennsylvania Department of Health | Centers for Disease Control and Prevention | |
| EPI548046 | HA | United States | 2014-Oct-01 | A/Texas/37/2014 | Texas Department of State Health Services-Laboratory Services | Centers for Disease Control and Prevention | |
| EPI548054 | HA | United States | 2014-Aug-20 | A/Texas/39/2014 | Texas Department of State Health Services-Laboratory Services | Centers for Disease Control and Prevention | |
| EPI548062 | HA | United States | 2014-Oct-14 | A/Virginia/16/2014 | Virginia Division of Consolidated Laboratories | Centers for Disease Control and Prevention | |
| EPI548070 | HA | United States | 2014-Oct-06 | A/Washington/18/2014 | Washington State Public Health Laboratory | Centers for Disease Control and Prevention | |
| EPI548078 | HA | United States | 2014-Aug-05 | A/Wisconsin/27/2014 | Wisconsin State Laboratory of Hygiene | Centers for Disease Control and Prevention | |
| EPI548086 | HA | Finland | 2014-Jan-01 | A/Finland/440/2014 | National Institute for Medical Research | Centers for Disease Control and Prevention | |
| EPI548094 | HA | Sweden | 2014-Oct-18 | A/Stockholm/22/2014 | | Public Health Agency of Sweden | |
| EPI548102 | HA | Sweden | 2014-Oct-21 | A/Stockholm/25/2014 | | Public Health Agency of Sweden | |
| EPI548110 | HA | Sweden | 2014-Oct-25 | A/Stockholm/23/2014 | | Public Health Agency of Sweden | |
| EPI548118 | HA | Sweden | 2014-Oct-19 | A/Malmoe/4/2014 | | Public Health Agency of Sweden | |

| | | | | | | |
|---|---|---|---|---|---|---|
| EPI548133 | HA | Sweden | 2014-Sep-26 | A/Stockholm/24/2014 | | Public Health Agency of Sweden |
| EPI548860 | HA | United States | 2014-Oct-01 | A/Arkansas/08/2014 | Arkansas Department of Health | Centers for Disease Control and Prevention |
| EPI548868 | HA | United States | 2014-Oct-17 | A/Montana/11/2014 | Montana Laboratory Services Bureau | Centers for Disease Control and Prevention |
| EPI548877 | HA | United States | 2014-Oct-21 | A/Colorado/21/2014 | Colorado Department of Health Lab | Centers for Disease Control and Prevention |
| EPI548892 | HA | Brazil | 2014-Jul-06 | A/Brazil/3972/2014 | Instituto Adolfo Lutz | Centers for Disease Control and Prevention |
| EPI548900 | HA | Bangladesh | 2014-Jun-09 | A/Bangladesh/10009/2014 | Institute of Epidemiology Disease Control and Research (IEDCR) & Bangladesh National Influenza Centre (NIC) | Centers for Disease Control and Prevention |
| EPI548908 | HA | United States | 2014-Jul-20 | A/Hawaii/46/2014 | State of Hawaii Department of Health | Centers for Disease Control and Prevention |
| EPI548916 | HA | United States | 2014-Jul-26 | A/Hawaii/45/2014 | State of Hawaii Department of Health | Centers for Disease Control and Prevention |
| EPI548924 | HA | United States | 2014-Oct-16 | A/Mississippi/04/2014 | Mississippi Public Health Laboratory | Centers for Disease Control and Prevention |
| EPI548932 | HA | United States | 2014-Oct-01 | A/North Dakota/06/2014 | North Dakota Department of Health | Centers for Disease Control and Prevention |
| EPI548942 | HA | Brazil | 2014-Feb-19 | A/Brazil/0769/2014 | Instituto Adolfo Lutz | Centers for Disease Control and Prevention |
| EPI548950 | HA | United States | 2014-Oct-04 | A/Colorado/15/2014 | Colorado Department of Health Lab | Centers for Disease Control and Prevention |
| EPI548958 | HA | Brazil | 2014-Feb-17 | A/Brazil/9879/2014 | Instituto Adolfo Lutz | Centers for Disease Control and Prevention |
| EPI548966 | HA | United States | 2014-Oct-09 | A/Wyoming/08/2014 | Wyoming Public Health Laboratory | Centers for Disease Control and Prevention |
| EPI548974 | HA | United States | 2014-Oct-15 | A/Mississippi/03/2014 | Mississippi Public Health Laboratory | Centers for Disease Control and Prevention |
| EPI548982 | HA | United States | 2014-Oct-06 | A/Mississippi/07/2014 | Mississippi Public Health Laboratory | Centers for Disease Control and Prevention |
| EPI548990 | HA | United States | 2014-Oct-15 | A/New Mexico/28/2014 | New Mexico Department of Health | Centers for Disease Control and Prevention |
| EPI548998 | HA | Brazil | 2014-May-07 | A/Brazil/2035/2014 | Instituto Adolfo Lutz | Centers for Disease Control and Prevention |
| EPI549006 | HA | Brazil | 2014-Jul-07 | A/Brazil/623/2014 | Instituto Adolfo Lutz | Centers for Disease Control and Prevention |
| EPI549014 | HA | Brazil | 2014-Jul-21 | A/Brazil/367/2014 | Instituto Adolfo Lutz | Centers for Disease Control and Prevention |
| EPI549022 | HA | United States | 2014-Sep-19 | A/Pennsylvania/42/2014 | Pennsylvania Department of Health | Centers for Disease Control and Prevention |
| EPI549030 | HA | Brazil | 2014-Mar-03 | A/Brazil/6069/2014 | Instituto Adolfo Lutz | Centers for Disease Control and Prevention |
| EPI549038 | HA | United States | 2014-Oct-14 | A/Mississippi/05/2014 | Mississippi Public Health Laboratory | Centers for Disease Control and Prevention |
| EPI549046 | HA | United States | 2014-Jul-18 | A/Hawaii/44/2014 | State of Hawaii Department of Health | Centers for Disease Control and Prevention |

| | | | | | | | |
|---|---|---|---|---|---|---|---|
| EPI549053 | HA | United States | 2014-Jul-18 | A/Hawaii/47/2014 | State of Hawaii Department of Health | Centers for Disease Control and Prevention | |
| EPI549061 | HA | United States | 2014-Sep-28 | A/New Hampshire/19/2014 | New Hampshire Public Health Laboratories | Centers for Disease Control and Prevention | |
| EPI549069 | HA | Brazil | 2014-Jun-07 | A/Brazil/4146/2014 | Instituto Adolfo Lutz | Centers for Disease Control and Prevention | |
| EPI549077 | HA | United States | 2014-Oct-11 | A/Delaware/19/2014 | Delaware Public Health Lab | Centers for Disease Control and Prevention | |
| EPI549085 | HA | United States | 2014-Oct-03 | A/Connecticut/19/2014 | Connecticut Department. of Public Health | Centers for Disease Control and Prevention | |
| EPI549116 | HA | United States | 2014-Sep-24 | A/Arkansas/06/2014 | Arkansas Department of Health | Centers for Disease Control and Prevention | |
| EPI549124 | HA | United States | 2014-Sep-29 | A/Arkansas/07/2014 | Arkansas Department of Health | Centers for Disease Control and Prevention | |
| EPI549132 | HA | United States | 2014-Sep-24 | A/Arkansas/05/2014 | Arkansas Department of Health | Centers for Disease Control and Prevention | |
| EPI549140 | HA | United States | 2014-Sep-24 | A/Pennsylvania/43/2014 | Pennsylvania Department of Health | Centers for Disease Control and Prevention | |
| EPI549148 | HA | United States | 2014-Oct-18 | A/Minnesota/28/2014 | Minnesota Department of Health | Centers for Disease Control and Prevention | |
| EPI549156 | HA | United States | 2014-Oct-20 | A/Texas/41/2014 | Texas Department of State Health Services-Laboratory Services | Centers for Disease Control and Prevention | |
| EPI550842 | HA | Hong Kong (SAR) | 2014-Apr-30 | A/Hong Kong/5738/2014 | National Institute for Medical Research | Centers for Disease Control and Prevention | |
| EPI550857 | HA | United States | 2014-Oct-20 | A/Alabama/05/2014 | ADPH Bureau of Clinical Laboratories | Centers for Disease Control and Prevention | |
| EPI550872 | HA | United States | 2014-Oct-03 | A/Colorado/16/2014 | Colorado Department of Health Lab | Centers for Disease Control and Prevention | |
| EPI550895 | HA | United States | 2014-Oct-14 | A/Connecticut/21/2014 | Connecticut Department. of Public Health | Centers for Disease Control and Prevention | |
| EPI550903 | HA | United States | 2014-Sep-23 | A/Florida/35/2014 | Florida Department of Health-Tampa | Centers for Disease Control and Prevention | |
| EPI550911 | HA | United States | 2014-Oct-12 | A/Florida/45/2014 | Florida Department of Health-Tampa | Centers for Disease Control and Prevention | |
| EPI550919 | HA | United States | 2014-Oct-25 | A/Indiana/10/2014 | Indiana State Department of Health Laboratories | Centers for Disease Control and Prevention | |
| EPI550927 | HA | United States | 2014-Oct-07 | A/Louisiana/06/2014 | Louisiana Department of Health and Hospitals | Centers for Disease Control and Prevention | |
| EPI550935 | HA | United States | 2014-Oct-10 | A/Louisiana/07/2014 | Louisiana Department of Health and Hospitals | Centers for Disease Control and Prevention | |
| EPI550943 | HA | United States | 2014-Oct-11 | A/Louisiana/08/2014 | Louisiana Department of Health and Hospitals | Centers for Disease Control and Prevention | Saito, Reiko; Takemae, Nobuhiro; Saito, Takehiko; Shobugawa, Yugo; Kondo, Hiroki; Hibino, Akinobu; Shirahige, Yutaka |
| EPI550951 | HA | United States | 2014-Oct-11 | A/Louisiana/09/2014 | Louisiana Department of Health and Hospitals | Centers for Disease Control and Prevention | Saito, Reiko; Takemae, Nobuhiro; Saito, Takehiko; Shobugawa, Yugo; Kondo, Hiroki; Hibino, Akinobu; Shirahige, Yutaka |

| | | | | | | | |
|---|---|---|---|---|---|---|---|
| EPI550959 | HA | United States | 2014-Oct-21 | A/Louisiana/11/2014 | Louisiana Department of Health and Hospitals | Centers for Disease Control and Prevention | Saito, Reiko; Takemae, Nobuhiro; Saito, Takehiko; Shobugawa, Yugo; Kondo, Hiroki; Hibino, Akinobu; Shirahige, Yutaka |
| EPI550967 | HA | United States | 2014-Oct-22 | A/Louisiana/14/2014 | Louisiana Department of Health and Hospitals | Centers for Disease Control and Prevention | |
| EPI550975 | HA | United States | 2014-Oct-14 | A/Maine/05/2014 | Maine Health and Environmental Testing Laboratory | Centers for Disease Control and Prevention | Saito, Reiko; Takemae, Nobuhiro; Saito, Takehiko; Shobugawa, Yugo; Kondo, Hiroki; Hibino, Akinobu; Kawashima, Takashi |
| EPI550983 | HA | United States | 2014-Sep-16 | A/Maryland/26/2014 | Maryland Department of Health and Mental Hygiene | Centers for Disease Control and Prevention | |
| EPI550996 | HA | United States | 2014-Oct-17 | A/New Hampshire/21/2014 | New Hampshire Public Health Laboratories | Centers for Disease Control and Prevention | Saito, Reiko; Takemae, Nobuhiro; Saito, Takehiko; Shobugawa, Yugo; Kondo, Hiroki; Hibino, Akinobu; Sato, Isamu |
| EPI551004 | HA | United States | 2014-Oct-06 | A/North Carolina/21/2014 | North Carolina State Laboratory of Public Health | Centers for Disease Control and Prevention | Saito, Reiko; Takemae, Nobuhiro; Saito, Takehiko; Shobugawa, Yugo; Kondo, Hiroki; Hibino, Akinobu; Sato, Isamu |
| EPI551012 | HA | United States | 2014-Oct-09 | A/Oregon/05/2014 | Oregon Public Health Laboratory | Centers for Disease Control and Prevention | Takashita,Emi; Fujisaki,Seiichiro; Shirakura,Masayuki; Watanabe,Shinji; Odagiri,Takato |
| EPI551020 | HA | United States | 2014-Oct-09 | A/Oregon/06/2014 | Oregon Public Health Laboratory | Centers for Disease Control and Prevention | |
| EPI551035 | HA | United States | 2014-Oct-05 | A/Oregon/09/2014 | Oregon Public Health Laboratory | Centers for Disease Control and Prevention | |
| EPI551043 | HA | United States | 2014-Jun-10 | A/Texas/34/2014 | Texas Department of State Health Services-Laboratory Services | Centers for Disease Control and Prevention | |
| EPI551051 | HA | United States | 2014-Oct-01 | A/Texas/37/2014 | Texas Department of State Health Services-Laboratory Services | Centers for Disease Control and Prevention | |
| EPI551059 | HA | United States | 2014-Sep-22 | A/Texas/40/2014 | Texas Department of State Health Services-Laboratory Services | Centers for Disease Control and Prevention | |
| EPI551067 | HA | United States | 2014-Oct-22 | A/Texas/42/2014 | Texas Department of State Health Services-Laboratory Services | Centers for Disease Control and Prevention | |
| EPI551075 | HA | United States | 2014-Oct-15 | A/Texas/43/2014 | Texas Department of State Health Services-Laboratory Services | Centers for Disease Control and Prevention | |
| EPI551220 | HA | United Kingdom | 2014-Oct-24 | A/ENG/516/2014 | Microbiology Services Colindale, Public Health England | Microbiology Services Colindale, Public Health England | |
| EPI551562 | HA | Australia | 2014-Jul-31 | A/CANBERRA/82/2014 | Canberra Hospital | WHO Collaborating Centre for Reference and Research on Influenza | |
| EPI551603 | HA | Australia | 2014-Aug-05 | A/NEWCASTLE/54/2014 | John Hunter Hospital | WHO Collaborating Centre for Reference and Research on Influenza | |
| EPI551849 | HA | Australia | 2014-Sep-19 | A/VICTORIA/5060/2014 | Victorian Infectious Diseases Reference Laboratory | WHO Collaborating Centre for Reference and Research on Influenza | |
| EPI552085 | HA | United States | 2014-Oct-29 | A/Ohio/03/2014 | Ohio Department of Health Laboratories | Centers for Disease Control and Prevention | |

| | | | | | | |
|---|---|---|---|---|---|---|
| EPI552093 | HA | United States | 2014-Nov-05 | A/Indiana/12/2014 | Indiana State Department of Health Laboratories | Centers for Disease Control and Prevention |
| EPI552101 | HA | United States | 2014-Nov-07 | A/Indiana/13/2014 | Indiana State Department of Health Laboratories | Centers for Disease Control and Prevention |
| EPI552109 | HA | United States | 2014-Oct-21 | A/Louisiana/12/2014 | Louisiana Department of Health and Hospitals | Centers for Disease Control and Prevention |
| EPI552117 | HA | United States | 2014-Oct-24 | A/North Dakota/09/2014 | North Dakota Department of Health | Centers for Disease Control and Prevention |
| EPI552125 | HA | United States | 2014-Nov-03 | A/Missouri/09/2014 | Missouri Department. of Health & Senior Services | Centers for Disease Control and Prevention |
| EPI552133 | HA | United States | 2014-Nov-05 | A/Mississippi/12/2014 | Mississippi Public Health Laboratory | Centers for Disease Control and Prevention |
| EPI552141 | HA | United States | 2014-Oct-23 | A/Alabama/07/2014 | ADPH Bureau of Clinical Laboratories | Centers for Disease Control and Prevention |
| EPI552149 | HA | United States | 2014-Oct-23 | A/Mississippi/08/2014 | Mississippi Public Health Laboratory | Centers for Disease Control and Prevention |
| EPI552157 | HA | United States | 2014-Nov-03 | A/Mississippi/10/2014 | Mississippi Public Health Laboratory | Centers for Disease Control and Prevention |
| EPI552165 | HA | United States | 2014-Oct-29 | A/Mississippi/09/2014 | Mississippi Public Health Laboratory | Centers for Disease Control and Prevention |
| EPI552173 | HA | United States | 2014-Oct-28 | A/Montana/14/2014 | Montana Laboratory Services Bureau | Centers for Disease Control and Prevention |
| EPI552179 | HA | United States | 2014-Oct-30 | A/Montana/15/2014 | Montana Laboratory Services Bureau | Centers for Disease Control and Prevention |
| EPI552187 | HA | United States | 2014-Oct-27 | A/Texas/48/2014 | Texas Department of State Health Services-Laboratory Services | Centers for Disease Control and Prevention |
| EPI552197 | HA | United States | 2014-Nov-10 | A/Indiana/15/2014 | Indiana State Department of Health Laboratories | Centers for Disease Control and Prevention |
| EPI552205 | HA | United States | 2014-Oct-30 | A/Texas/46/2014 | Houston Department of Health and Human Services | Centers for Disease Control and Prevention |
| EPI552213 | HA | United States | 2014-Oct-21 | A/Iowa/16/2014 | Iowa State Hygienic Laboratory | Centers for Disease Control and Prevention |
| EPI552227 | HA | United States | 2014-Nov-10 | A/Indiana/16/2014 | Indiana State Department of Health Laboratories | Centers for Disease Control and Prevention |
| EPI552235 | HA | United States | 2014-Oct-20 | A/Iowa/15/2014 | Iowa State Hygienic Laboratory | Centers for Disease Control and Prevention |
| EPI552243 | HA | United States | 2014-Oct-21 | A/Louisiana/13/2014 | Louisiana Department of Health and Hospitals | Centers for Disease Control and Prevention |
| EPI552251 | HA | United States | 2014-Nov-06 | A/Utah/21/2014 | Utah Department of Health | Centers for Disease Control and Prevention |
| EPI552259 | HA | United States | 2014-Nov-07 | A/Utah/19/2014 | Utah Department of Health | Centers for Disease Control and Prevention |
| EPI552267 | HA | United States | 2014-Nov-05 | A/Utah/20/2014 | Utah Department of Health | Centers for Disease Control and Prevention |
| EPI552275 | HA | United States | 2014-Nov-06 | A/Utah/17/2014 | Utah Department of Health | Centers for Disease Control and Prevention |
| EPI552283 | HA | United States | 2014-Nov-02 | A/Utah/18/2014 | Utah Department of Health | Centers for Disease Control and Prevention |
| EPI552302 | HA | United States | 2014-Oct-29 | A/Kansas/08/2014 | Kansas Department of Health and Environment | Centers for Disease Control and Prevention |

| | | | | | | | |
|---|---|---|---|---|---|---|---|
| EPI552309 | HA | United States | 2014-Oct-27 | A/Kansas/07/2014 | Kansas Department of Health and Environment | Centers for Disease Control and Prevention | |
| EPI552317 | HA | United States | 2014-Oct-29 | A/Maryland/33/2014 | Maryland Department of Health and Mental Hygiene | Centers for Disease Control and Prevention | |
| EPI552325 | HA | United States | 2014-Oct-29 | A/Ohio/04/2014 | Ohio Department of Health Laboratories | Centers for Disease Control and Prevention | |
| EPI552335 | HA | United States | 2014-Nov-05 | A/Alabama/09/2014 | ADPH Bureau of Clinical Laboratories | Centers for Disease Control and Prevention | |
| EPI552343 | HA | United States | 2014-Nov-10 | A/Nevada/18/2014 | Southern Nevada Public Health Lab | Centers for Disease Control and Prevention | |
| EPI552351 | HA | United States | 2014-Nov-10 | A/Nevada/17/2014 | Southern Nevada Public Health Lab | Centers for Disease Control and Prevention | |
| EPI552358 | HA | United States | 2014-Nov-08 | A/Utah/22/2014 | Utah Department of Health | Centers for Disease Control and Prevention | |
| EPI552366 | HA | United States | 2014-Nov-04 | A/Mississippi/11/2014 | Mississippi Public Health Laboratory | Centers for Disease Control and Prevention | |
| EPI552374 | HA | United States | 2014-Nov-05 | A/Idaho/04/2014 | State of Idaho Bureau of Laboratories | Centers for Disease Control and Prevention | Saito, Reiko; Takemae, Nobuhiro; Saito, Takehiko; Shobugawa, Yugo; Kondo, Hiroki; Hibino, Akinobu; Yadanar, Kyaw; Yi Yi, Myint; Khin Yi, Oo; Htay Htay, Tin |
| EPI552395 | HA | United States | 2014-Nov-05 | A/Indiana/11/2014 | Indiana State Department of Health Laboratories | Centers for Disease Control and Prevention | Saito, Reiko; Takemae, Nobuhiro; Saito, Takehiko; Shobugawa, Yugo; Kondo, Hiroki; Hibino, Akinobu; Yadanar, Kyaw; Yi Yi, Myint; Khin Yi, Oo; Htay Htay, Tin |
| EPI552417 | HA | Sweden | 2014-Nov-06 | A/Karlstad/1/2014 | | Public Health Agency of Sweden | Saito, Reiko; Takemae, Nobuhiro; Saito, Takehiko; Shobugawa, Yugo; Kondo, Hiroki; Hibino, Akinobu; Yadanar, Kyaw; Yi Yi, Myint; Khin Yi, Oo; Htay Htay, Tin |
| EPI552425 | HA | Sweden | 2014-Nov-03 | A/Stockholm/26/2014 | | Public Health Agency of Sweden | Saito, Reiko; Takemae, Nobuhiro; Saito, Takehiko; Shobugawa, Yugo; Kondo, Hiroki; Hibino, |
| EPI552535 | HA | United States | 2014-Sep-24 | A/Michigan/15/2014 | Michigan Department of Community Health | Centers for Disease Control and Prevention | Saito, Reiko; Takemae, Nobuhiro; Saito, Takehiko; Shobugawa, Yugo; Kondo, Hiroki; Hibino, |
| EPI552544 | HA | Palau | 2014-Mar-26 | A/Palau/6759/2014 | State of Hawaii Department of Health | Centers for Disease Control and Prevention | |
| EPI552560 | HA | Japan | 2014-Sep-09 | A/SHIMANE/96/2014 | Shimane Prefectural Institute of Public Health and Environmental Science | National Institute of Infectious Diseases (NIID) | |
| EPI552568 | HA | Japan | 2014-Sep-11 | A/SAITAMA/96/2014 | Saitama Institute of Public Health | National Institute of Infectious Diseases (NIID) | |
| EPI552674 | HA | Sweden | 2014-Nov-06 | A/Gothenburg/4/2014 | | Public Health Agency of Sweden | |
| EPI552682 | HA | Sweden | 2014-Nov-02 | A/Sweden/76/2014 | | Public Health Agency of Sweden | |
| EPI552690 | HA | Sweden | 2014-Nov-18 | A/Stockholm/29/2014 | | Public Health Agency of Sweden | |
| EPI552698 | HA | Sweden | 2014-Dec-02 | A/Stockholm/28/2014 | | Public Health Agency of Sweden | |
| EPI552706 | HA | Sweden | 2014-Nov-28 | A/Malmoe/5/2014 | | Public Health Agency of Sweden | |

| | | | | | | | |
|---|---|---|---|---|---|---|---|
| EPI552722 | HA | Sweden | 2014-Dec-04 | A/Sweden/75/2014 | | Public Health Agency of Sweden | |
| EPI552785 | HA | United States | 2014-Oct-09 | A/Iowa/13/2014 | Iowa State Hygienic Laboratory | Centers for Disease Control and Prevention | |
| EPI552799 | HA | United States | 2014-Nov-19 | A/New Hampshire/23/2014 | New Hampshire Public Health Laboratories | Centers for Disease Control and Prevention | |
| EPI552807 | HA | United States | 2014-Nov-13 | A/North Carolina/31/2014 | North Carolina State Laboratory of Public Health | Centers for Disease Control and Prevention | |
| EPI552823 | HA | United States | 2014-Nov-19 | A/Louisiana/20/2014 | Louisiana Department of Health and Hospitals | Centers for Disease Control and Prevention | |
| EPI552831 | HA | United States | 2014-Nov-19 | A/Louisiana/22/2014 | Louisiana Department of Health and Hospitals | Centers for Disease Control and Prevention | |
| EPI552839 | HA | United States | 2014-Nov-18 | A/Louisiana/17/2014 | Louisiana Department of Health and Hospitals | Centers for Disease Control and Prevention | Saito, Reiko; Takemae, Nobuhiro; Saito, Takehiko; Shobugawa, Yugo; Kondo, Hiroki; Hibino, Akinobu; Masaki, Hironori |
| EPI552847 | HA | United States | 2014-Nov-19 | A/Louisiana/21/2014 | Louisiana Department of Health and Hospitals | Centers for Disease Control and Prevention | Takashita,Emi; Fujisaki,Seiichiro; Shirakura,Masayuki; Watanabe,Shinji; Odagiri,Takato |
| EPI552855 | HA | United States | 2014-Nov-18 | A/Louisiana/18/2014 | Louisiana Department of Health and Hospitals | Centers for Disease Control and Prevention | Takashita,Emi; Fujisaki,Seiichiro; Shirakura,Masayuki; Watanabe,Shinji; Odagiri,Takato |
| EPI552876 | HA | United States | 2014-Nov-14 | A/Wyoming/14/2014 | Wyoming Public Health Laboratory | Centers for Disease Control and Prevention | Saito, Reiko; Takemae, Nobuhiro; Saito, Takehiko; Shobugawa, Yugo; Kondo, Hiroki; Hibino, Akinobu; Yadanar, Kyaw; Yi Yi, Myint; Khin Yi, Oo; Htay Htay, Tin |
| EPI552892 | HA | United States | 2014-Nov-14 | A/North Carolina/35/2014 | North Carolina State Laboratory of Public Health | Centers for Disease Control and Prevention | Saito, Reiko; Takemae, Nobuhiro; Saito, Takehiko; Shobugawa, Yugo; Kondo, Hiroki; Hibino, Akinobu; Yadanar, Kyaw; Yi Yi, Myint; Khin Yi, Oo; Htay Htay, Tin |
| EPI552900 | HA | United States | 2014-Nov-19 | A/New Jersey/27/2014 | New Jersey Department of Health & Senior Services | Centers for Disease Control and Prevention | Saito, Reiko; Takemae, Nobuhiro; Saito, Takehiko; Shobugawa, Yugo; Kondo, Hiroki; Hibino, Akinobu; Yadanar, Kyaw; Yi Yi, Myint; Khin Yi, Oo; Htay Htay, Tin |
| EPI552905 | HA | United States | 2014-Nov-07 | A/Wyoming/11/2014 | Wyoming Public Health Laboratory | Centers for Disease Control and Prevention | Saito, Reiko; Takemae, Nobuhiro; Saito, Takehiko; Shobugawa, Yugo; Kondo, Hiroki; Hibino, Akinobu; Yadanar, Kyaw; Yi Yi, Myint; Khin Yi, Oo; Htay Htay, Tin |
| EPI552913 | HA | United States | 2014-Nov-17 | A/New Jersey/26/2014 | New Jersey Department of Health & Senior Services | Centers for Disease Control and Prevention | Saito, Reiko; Takemae, Nobuhiro; Saito, Takehiko; Shobugawa, Yugo; Kondo, Hiroki; Hibino, Akinobu; Yadanar, Kyaw; Yi Yi, Myint; Khin Yi, Oo; Htay Htay, Tin |

| | | | | | | | |
|---|---|---|---|---|---|---|---|
| EPI552930 | HA | United States | 2014-Oct-21 | A/Delaware/22/2014 | Delaware Public Health Lab | Centers for Disease Control and Prevention | Saito, Reiko; Takemae, Nobuhiro; Saito, Takehiko; Shobugawa, Yugo; Kondo, Hiroki; Hibino, Akinobu; Yadanar, Kyaw; Yi Yi, Myint; Khin Yi, Oo; Htay Htay, Tin |
| EPI552948 | HA | United States | 2014-Nov-04 | A/North Carolina/28/2014 | North Carolina State Laboratory of Public Health | Centers for Disease Control and Prevention | Saito, Reiko; Takemae, Nobuhiro; Saito, Takehiko; Shobugawa, Yugo; Kondo, Hiroki; Hibino, Akinobu; Yadanar, Kyaw; Yi Yi, Myint; Khin Yi, Oo; Htay Htay, Tin |
| EPI552951 | HA | United States | 2014-Oct-12 | A/Iowa/14/2014 | Iowa State Hygienic Laboratory | Centers for Disease Control and Prevention | Saito, Reiko; Takemae, Nobuhiro; Saito, Takehiko; Shobugawa, Yugo; Kondo, Hiroki; Hibino, Akinobu; Yadanar, Kyaw; Yi Yi, Myint; Khin Yi, Oo; Htay Htay, Tin |
| EPI552959 | HA | United States | 2014-Nov-13 | A/North Carolina/34/2014 | North Carolina State Laboratory of Public Health | Centers for Disease Control and Prevention | Saito, Reiko; Takemae, Nobuhiro; Saito, Takehiko; Shobugawa, Yugo; Kondo, Hiroki; Hibino, Akinobu; Yadanar, Kyaw; Yi Yi, Myint; Khin Yi, Oo; Htay Htay, Tin |
| EPI552967 | HA | United States | 2014-Nov-18 | A/Louisiana/19/2014 | Louisiana Department of Health and Hospitals | Centers for Disease Control and Prevention | |
| EPI552975 | HA | United States | 2014-Nov-08 | A/North Carolina/29/2014 | North Carolina State Laboratory of Public Health | Centers for Disease Control and Prevention | |
| EPI552983 | HA | United States | 2014-Nov-18 | A/Wyoming/15/2014 | Wyoming Public Health Laboratory | Centers for Disease Control and Prevention | |
| EPI552991 | HA | United States | 2014-Nov-13 | A/North Carolina/32/2014 | North Carolina State Laboratory of Public Health | Centers for Disease Control and Prevention | |
| EPI553000 | HA | United States | 2014-Oct-05 | A/Alaska/47/2014 | Alaska State Virology Lab | Centers for Disease Control and Prevention | Saito, Reiko; Takemae, Nobuhiro; Saito, Takehiko; Shobugawa, Yugo; Kondo, Hiroki; Hibino, Akinobu; Yadanar, Kyaw; Yi Yi, Myint; Khin Yi, Oo; Htay Htay, Tin |
| EPI553005 | HA | United States | 2014-Oct-04 | A/Alaska/46/2014 | Alaska State Virology Lab | Centers for Disease Control and Prevention | |
| EPI553016 | HA | United States | 2014-Nov-05 | A/Texas/52/2014 | Houston Department of Health and Human Services | Centers for Disease Control and Prevention | |
| EPI553026 | HA | United States | 2014-Oct-23 | A/Maryland/34/2014 | Maryland Department of Health and Mental Hygiene | Centers for Disease Control and Prevention | |
| EPI553041 | HA | United States | 2014-Nov-18 | A/Missouri/11/2014 | Missouri Department. of Health & Senior Services | Centers for Disease Control and Prevention | |
| EPI553049 | HA | United States | 2014-Nov-19 | A/Massachusetts/14/2014 | Massachusetts Department of Public Health | Centers for Disease Control and Prevention | |
| EPI553057 | HA | United States | 2014-Nov-25 | A/Texas/57/2014 | Baylor College of Medicine | Centers for Disease Control and Prevention | |
| EPI553065 | HA | United States | 2014-Nov-17 | A/Missouri/10/2014 | Missouri Department. of Health & Senior Services | Centers for Disease Control and Prevention | |
| EPI553073 | HA | United States | 2014-Nov-11 | A/Delaware/24/2014 | Delaware Public Health Lab | Centers for Disease Control and Prevention | |
| EPI553081 | HA | United States | 2014-Nov-20 | A/Delaware/25/2014 | Delaware Public Health Lab | Centers for Disease Control and Prevention | |

| | | | | | | | |
|---|---|---|---|---|---|---|---|
| EPI553092 | HA | United States | 2014-Nov-20 | A/Georgia/05/2014 | Georgia Public Health Laboratory | Centers for Disease Control and Prevention | |
| EPI553100 | HA | United States | 2014-Nov-20 | A/Georgia/06/2014 | Georgia Public Health Laboratory | Centers for Disease Control and Prevention | |
| EPI553108 | HA | United States | 2014-Nov-20 | A/Georgia/04/2014 | Georgia Public Health Laboratory | Centers for Disease Control and Prevention | |
| EPI553116 | HA | United States | 2014-Nov-20 | A/Georgia/07/2014 | Georgia Public Health Laboratory | Centers for Disease Control and Prevention | |
| EPI553119 | HA | United States | 2014-Nov-03 | A/Illinois/05/2014 | Illinois Department of Public Health-Chicago | Centers for Disease Control and Prevention | |
| EPI553126 | HA | United States | 2014-Nov-19 | A/Kansas/10/2014 | Kansas Department of Health and Environment | Centers for Disease Control and Prevention | |
| EPI553259 | HA | Japan | 2014-Nov-06 | A/SENDAI/72/2014 | Sendai City Institute of Public Health | National Institute of Infectious Diseases (NIID) | |
| EPI553374 | HA | Singapore | 2013-Dec-11 | A/Singapore/OHC005/2013 | Ministry of Health, Singapore | Ministry of Health, Singapore | |
| EPI553382 | HA | Singapore | 2013-Dec-11 | A/Singapore/OHC008/2013 | Ministry of Health, Singapore | Ministry of Health, Singapore | |
| EPI553390 | HA | Singapore | 2013-Dec-12 | A/Singapore/OHC010/2013 | Ministry of Health, Singapore | Ministry of Health, Singapore | |
| EPI553398 | HA | Singapore | 2013-Dec-13 | A/Singapore/OHC015/2013 | Ministry of Health, Singapore | Ministry of Health, Singapore | |
| EPI553406 | HA | Singapore | 2013-Dec-13 | A/Singapore/OHC016/2013 | Ministry of Health, Singapore | Ministry of Health, Singapore | |
| EPI553414 | HA | Singapore | 2013-Dec-13 | A/Singapore/OHC019/2013 | Ministry of Health, Singapore | Ministry of Health, Singapore | |
| EPI553422 | HA | Singapore | 2013-Dec-13 | A/Singapore/OHC022/2013 | Ministry of Health, Singapore | Ministry of Health, Singapore | |
| EPI553430 | HA | Singapore | 2013-Dec-12 | A/Singapore/OHC009/2013 | Ministry of Health, Singapore | Ministry of Health, Singapore | |
| EPI553438 | HA | Singapore | 2013-Dec-12 | A/Singapore/TT938/2013 | Ministry of Health, Singapore | Ministry of Health, Singapore | |
| EPI553446 | HA | Singapore | 2013-Dec-16 | A/Singapore/TT973/2013 | Ministry of Health, Singapore | Ministry of Health, Singapore | |
| EPI553454 | HA | Singapore | 2013-Dec-30 | A/Singapore/CGH021/2013 | Ministry of Health, Singapore | Ministry of Health, Singapore | |
| EPI553462 | HA | Singapore | 2013-Dec-19 | A/Singapore/TT979/2013 | Ministry of Health, Singapore | Ministry of Health, Singapore | |
| EPI553499 | HA | United States | 2014-Nov-07 | A/Colorado/24/2014 | Colorado Department of Health Lab | Centers for Disease Control and Prevention | Saito, Reiko; Takemae, Nobuhiro; Saito, Takehiko; Shobugawa, Yugo; Kondo, Hiroki; Hibino, Akinobu; Yadanar, Kyaw; Yi Yi, Myint; Khin Yi, Oo; Htay Htay, Tin |
| EPI553507 | HA | United States | 2014-Oct-28 | A/Wisconsin/46/2014 | Wisconsin State Laboratory of Hygiene | Centers for Disease Control and Prevention | |
| EPI553528 | HA | United States | 2014-Oct-06 | A/Florida/43/2014 | Florida Department of Health-Tampa | Centers for Disease Control and Prevention | |
| EPI553536 | HA | United States | 2014-Sep-17 | A/Arkansas/01/2014 | Arkansas Department of Health | Centers for Disease Control and Prevention | |
| EPI553544 | HA | United States | 2014-Oct-12 | A/Florida/45/2014 | Florida Department of Health-Tampa | Centers for Disease Control and Prevention | |
| EPI553552 | HA | United States | 2014-Sep-17 | A/Arkansas/02/2014 | Arkansas Department of Health | Centers for Disease Control and Prevention | |
| EPI553560 | HA | United States | 2014-Jan-01 | A/Florida/41/2014 | Florida Department of Health-Tampa | Centers for Disease Control and Prevention | |
| EPI553575 | HA | United States | 2014-Sep-22 | A/Arkansas/04/2014 | Arkansas Department of Health | Centers for Disease Control and Prevention | |
| EPI553583 | HA | United States | 2014-Oct-14 | A/New York/26/2014 | New York State Department of Health | Centers for Disease Control and Prevention | |

| | | | | | | |
|---|---|---|---|---|---|---|
| EPI553599 | HA | United States | 2014-Oct-18 | A/Colorado/19/2014 | Colorado Department of Health Lab | Centers for Disease Control and Prevention |
| EPI553609 | HA | United States | 2014-Oct-18 | A/Hawaii/48/2014 | State of Hawaii Department of Health | Centers for Disease Control and Prevention |
| EPI553617 | HA | United States | 2014-Oct-18 | A/Hawaii/49/2014 | State of Hawaii Department of Health | Centers for Disease Control and Prevention |
| EPI553625 | HA | United States | 2014-Oct-21 | A/Arkansas/12/2014 | Arkansas Department of Health | Centers for Disease Control and Prevention |
| EPI553633 | HA | United States | 2014-Oct-25 | A/Iowa/17/2014 | Iowa State Hygienic Laboratory | Centers for Disease Control and Prevention |
| EPI553641 | HA | United States | 2014-Oct-20 | A/Arkansas/11/2014 | Arkansas Department of Health | Centers for Disease Control and Prevention |
| EPI553649 | HA | United States | 2014-Oct-23 | A/Mississippi/08/2014 | Mississippi Public Health Laboratory | Centers for Disease Control and Prevention |
| EPI553660 | HA | United States | 2014-Sep-08 | A/Alaska/40/2014 | Alaska State Virology Lab | Centers for Disease Control and Prevention |
| EPI553672 | HA | United States | 2014-Sep-24 | A/North Carolina/24/2014 | North Carolina State Laboratory of Public Health | Centers for Disease Control and Prevention |
| EPI553680 | HA | United States | 2014-Oct-22 | A/Texas/42/2014 | Texas Department of State Health Services-Laboratory Services | Centers for Disease Control and Prevention |
| EPI553695 | HA | United States | 2014-Nov-11 | A/Montana/20/2014 | Montana Laboratory Services Bureau | Centers for Disease Control and Prevention |
| EPI553704 | HA | United States | 2014-Oct-10 | A/New York/24/2014 | New York State Department of Health | Centers for Disease Control and Prevention |
| EPI553722 | HA | United States | 2014-Nov-09 | A/Wisconsin/49/2014 | Wisconsin State Laboratory of Hygiene | Centers for Disease Control and Prevention |
| EPI553730 | HA | United States | 2014-Sep-30 | A/Wisconsin/34/2014 | Wisconsin State Laboratory of Hygiene | Centers for Disease Control and Prevention |
| EPI553738 | HA | United States | 2014-Nov-03 | A/Texas/62/2014 | Texas Department of State Health Services-Laboratory Services | Centers for Disease Control and Prevention |
| EPI553746 | HA | United States | 2014-Nov-08 | A/Wisconsin/48/2014 | Wisconsin State Laboratory of Hygiene | Centers for Disease Control and Prevention |
| EPI553754 | HA | United States | 2014-Nov-12 | A/Indiana/19/2014 | Indiana State Department of Health Laboratories | Centers for Disease Control and Prevention |
| EPI553762 | HA | United States | 2014-Oct-18 | A/Maine/06/2014 | Maine Health and Environmental Testing Laboratory | Centers for Disease Control and Prevention |
| EPI553770 | HA | United States | 2014-Oct-21 | A/Louisiana/12/2014 | Louisiana Department of Health and Hospitals | Centers for Disease Control and Prevention |
| EPI553778 | HA | United States | 2014-Oct-22 | A/Louisiana/14/2014 | Louisiana Department of Health and Hospitals | Centers for Disease Control and Prevention |
| EPI553786 | HA | United States | 2014-Oct-17 | A/New Hampshire/21/2014 | New Hampshire Public Health Laboratories | Centers for Disease Control and Prevention |
| EPI553794 | HA | United States | 2014-Sep-30 | A/Ohio/05/2014 | Ohio Department of Health Laboratories | Centers for Disease Control and Prevention |
| EPI553802 | HA | United States | 2014-May-02 | A/Maryland/18/2014 | Maryland Department of Health and Mental Hygiene | Centers for Disease Control and Prevention |
| EPI553810 | HA | United States | 2014-Oct-14 | A/Connecticut/21/2014 | Connecticut Department. of Public Health | Centers for Disease Control and Prevention |
| EPI553818 | HA | United States | 2014-Oct-27 | A/Illinois/04/2014 | Illinois Department of Public Health-Chicago | Centers for Disease Control and Prevention |

| | | | | | | | |
|---|---|---|---|---|---|---|---|
| EPI553826 | HA | United States | 2014-Sep-03 | A/Alaska/52/2014 | Alaska State Virology Lab | Centers for Disease Control and Prevention | |
| EPI553834 | HA | United States | 2014-Oct-11 | A/Louisiana/08/2014 | Louisiana Department of Health and Hospitals | Centers for Disease Control and Prevention | |
| EPI553842 | HA | United States | 2014-Oct-23 | A/Wisconsin/44/2014 | Wisconsin State Laboratory of Hygiene | Centers for Disease Control and Prevention | |
| EPI553850 | HA | United States | 2014-Oct-22 | A/Washington/19/2014 | Washington State Public Health Laboratory | Centers for Disease Control and Prevention | |
| EPI553858 | HA | United States | 2014-Sep-23 | A/New York/19/2014 | New York State Department of Health | Centers for Disease Control and Prevention | |
| EPI553866 | HA | United States | 2014-Sep-25 | A/New York/20/2014 | New York State Department of Health | Centers for Disease Control and Prevention | |
| EPI553874 | HA | United States | 2014-Oct-14 | A/Maryland/28/2014 | Maryland Department of Health and Mental Hygiene | Centers for Disease Control and Prevention | Saito, Reiko; Takemae, Nobuhiro; Saito, Takehiko; Shobugawa, Yugo; Kondo, Hiroki; Hibino, Akinobu; Yadanar, Kyaw; Yi Yi, Myint; Khin Yi, Oo; Htay Htay, Tin |
| EPI553882 | HA | United States | 2014-Sep-07 | A/Hawaii/40/2014 | State of Hawaii Department of Health | Centers for Disease Control and Prevention | Saito, Reiko; Takemae, Nobuhiro; Saito, Takehiko; Shobugawa, Yugo; Kondo, Hiroki; Hibino, Akinobu; Yadanar, Kyaw; Yi Yi, Myint; Khin Yi, Oo; Htay Htay, Tin |
| EPI553890 | HA | United States | 2014-Sep-18 | A/New York/18/2014 | New York State Department of Health | Centers for Disease Control and Prevention | Saito, Reiko; Takemae, Nobuhiro; Saito, Takehiko; Shobugawa, Yugo; Kondo, Hiroki; Hibino, Akinobu; Masaki, Hironori |
| EPI553898 | HA | United States | 2014-Oct-23 | A/New York/36/2014 | New York State Department of Health | Centers for Disease Control and Prevention | Takashita,Emi; Fujisaki,Seiichiro; Shirakura,Masayuki; Watanabe,Shinji; Odagiri,Takato |
| EPI553906 | HA | United States | 2014-Oct-20 | A/Arkansas/10/2014 | Arkansas Department of Health | Centers for Disease Control and Prevention | Takashita,Emi; Fujisaki,Seiichiro; Shirakura,Masayuki; Watanabe,Shinji; Odagiri,Takato |
| EPI553919 | HA | United States | 2014-Sep-09 | A/North Carolina/23/2014 | North Carolina State Laboratory of Public Health | Centers for Disease Control and Prevention | |
| EPI553927 | HA | United States | 2014-Oct-15 | A/Maryland/32/2014 | Maryland Department of Health and Mental Hygiene | Centers for Disease Control and Prevention | |
| EPI553935 | HA | United States | 2014-Oct-14 | A/New York/28/2014 | New York State Department of Health | Centers for Disease Control and Prevention | |
| EPI553943 | HA | United States | 2014-Oct-17 | A/New York/31/2014 | New York State Department of Health | Centers for Disease Control and Prevention | |
| EPI553951 | HA | United States | 2014-Oct-17 | A/New York/34/2014 | New York State Department of Health | Centers for Disease Control and Prevention | |
| EPI553959 | HA | United States | 2014-Oct-21 | A/New York/35/2014 | New York State Department of Health | Centers for Disease Control and Prevention | |
| EPI553967 | HA | United States | 2014-Aug-08 | A/Hawaii/36/2014 | State of Hawaii Department of Health | Centers for Disease Control and Prevention | |
| EPI553975 | HA | United States | 2014-May-01 | A/Maryland/19/2014 | Maryland Department of Health and Mental Hygiene | Centers for Disease Control and Prevention | |
| EPI555044 | HA | Russian Federation | 2014-Dec-01 | A/Astrakhan/RII23/2014 | | WHO National Influenza Centre Russian Federation | |

| | | | | | | |
|---|---|---|---|---|---|---|
| EPI555102 | HA | Sweden | 2014-Dec-04 | A/Stockholm/30/2014 | | Public Health Agency of Sweden |
| EPI555110 | HA | Sweden | 2014-Nov-26 | A/Stockholm/31/2014 | | Public Health Agency of Sweden |
| EPI555233 | HA | United States | 2014-Nov-30 | A/Rhode Island/21/2014 | Rhode Island Department of Health | Centers for Disease Control and Prevention |
| EPI555241 | HA | United States | 2014-Nov-18 | A/Texas/55/2014 | Baylor College of Medicine | Centers for Disease Control and Prevention |
| EPI555249 | HA | United States | 2014-Dec-04 | A/North Dakota/10/2014 | North Dakota Department of Health | Centers for Disease Control and Prevention |
| EPI555263 | HA | United States | 2014-Nov-30 | A/Massachusetts/18/2014 | Massachusetts Department of Public Health | Centers for Disease Control and Prevention |
| EPI555271 | HA | United States | 2014-Nov-24 | A/Louisiana/28/2014 | Louisiana Department of Health and Hospitals | Centers for Disease Control and Prevention |
| EPI555279 | HA | United States | 2014-Dec-02 | A/Massachusetts/17/2014 | Massachusetts Department of Public Health | Centers for Disease Control and Prevention |
| EPI555287 | HA | United States | 2014-Dec-09 | A/Wisconsin/57/2014 | Marshfield Clinic Research Foundation | Centers for Disease Control and Prevention |
| EPI555295 | HA | United States | 2014-Dec-09 | A/Wisconsin/59/2014 | Marshfield Clinic Research Foundation | Centers for Disease Control and Prevention |
| EPI555303 | HA | United States | 2014-Dec-09 | A/Wisconsin/58/2014 | Marshfield Clinic Research Foundation | Centers for Disease Control and Prevention |
| EPI555311 | HA | United States | 2014-Nov-28 | A/Pennsylvania/47/2014 | UPMC-CLB Dept of Microbiology | Centers for Disease Control and Prevention |
| EPI555319 | HA | United States | 2014-Nov-17 | A/Texas/54/2014 | Baylor College of Medicine | Centers for Disease Control and Prevention |
| EPI555327 | HA | United States | 2014-Dec-01 | A/Missouri/13/2014 | Missouri Department. of Health & Senior Services | Centers for Disease Control and Prevention |
| EPI555335 | HA | United States | 2014-Dec-10 | A/Wisconsin/60/2014 | Marshfield Clinic Research Foundation | Centers for Disease Control and Prevention |
| EPI555343 | HA | United States | 2014-Dec-09 | A/Wisconsin/56/2014 | Marshfield Clinic Research Foundation | Centers for Disease Control and Prevention |
| EPI555351 | HA | United States | 2014-Dec-08 | A/Wisconsin/55/2014 | Marshfield Clinic Research Foundation | Centers for Disease Control and Prevention |
| EPI555359 | HA | United States | 2014-Dec-08 | A/Washington/37/2014 | Marshfield Clinic Research Foundation | Centers for Disease Control and Prevention |
| EPI555367 | HA | United States | 2014-Nov-29 | A/Minnesota/35/2014 | Minnesota Department of Health | Centers for Disease Control and Prevention |
| EPI555375 | HA | United States | 2014-Dec-08 | A/Wisconsin/54/2014 | Marshfield Clinic Research Foundation | Centers for Disease Control and Prevention |
| EPI555383 | HA | United States | 2014-Nov-20 | A/Colorado/28/2014 | Colorado Department of Health Lab | Centers for Disease Control and Prevention |
| EPI555402 | HA | United States | 2014-Dec-02 | A/Wisconsin/53/2014 | Marshfield Clinic Research Foundation | Centers for Disease Control and Prevention |
| EPI555415 | HA | United States | 2014-Nov-28 | A/New Hampshire/28/2014 | New Hampshire Public Health Laboratories | Centers for Disease Control and Prevention |
| EPI555427 | HA | United States | 2014-Nov-06 | A/Utah/17/2014 | Utah Department of Health | Centers for Disease Control and Prevention |
| EPI555435 | HA | United States | 2014-Nov-05 | A/Alabama/08/2014 | ADPH Bureau of Clinical Laboratories | Centers for Disease Control and Prevention |
| EPI555443 | HA | United States | 2014-Nov-14 | A/Michigan/19/2014 | University of Michigan SPH EPID | Centers for Disease Control and Prevention |

| | | | | | | |
|---|---|---|---|---|---|---|
| EPI555451 | HA | United States | 2014-Dec-08 | A/Kansas/12/2014 | Kansas Department of Health and Environment | Centers for Disease Control and Prevention |
| EPI555459 | HA | United States | 2014-Nov-24 | A/Texas/69/2014 | Baylor College of Medicine | Centers for Disease Control and Prevention |
| EPI555467 | HA | United States | 2014-Nov-10 | A/Illinois/07/2014 | Illinois Department of Public Health-Chicago | Centers for Disease Control and Prevention |
| EPI555479 | HA | United States | 2014-Dec-08 | A/Florida/66/2014 | Florida Department of Health-Jacksonville | Centers for Disease Control and Prevention |
| EPI555487 | HA | United States | 2014-Nov-03 | A/Hawaii/60/2014 | State of Hawaii Department of Health | Centers for Disease Control and Prevention |
| EPI555499 | HA | United States | 2014-Nov-25 | A/Texas/56/2014 | Baylor College of Medicine | Centers for Disease Control and Prevention |
| EPI555507 | HA | United States | 2014-Dec-02 | A/North Dakota/11/2014 | North Dakota Department of Health | Centers for Disease Control and Prevention |
| EPI555515 | HA | United States | 2014-Nov-24 | A/Oklahoma/10/2014 | Oklahoma State Department of Health | Centers for Disease Control and Prevention |
| EPI555524 | HA | United States | 2014-Dec-07 | A/Delaware/31/2014 | Delaware Public Health Lab | Centers for Disease Control and Prevention |
| EPI555538 | HA | United States | 2014-Nov-28 | A/New Mexico/30/2014 | New Mexico Department of Health | Centers for Disease Control and Prevention |
| EPI555546 | HA | United States | 2014-Nov-25 | A/Idaho/07/2014 | State of Idaho Bureau of Laboratories | Centers for Disease Control and Prevention |
| EPI555559 | HA | United States | 2014-Nov-26 | A/Kentucky/14/2014 | Kentucky Division of Laboratory Services | Centers for Disease Control and Prevention |
| EPI555567 | HA | United States | 2014-Nov-23 | A/Minnesota/36/2014 | Minnesota Department of Health | Centers for Disease Control and Prevention |
| EPI555575 | HA | United States | 2014-Oct-30 | A/Texas/66/2014 | Texas Department of State Health Services-Laboratory Services | Centers for Disease Control and Prevention |
| EPI556107 | HA | United States | 2014-Nov-28 | A/Pennsylvania/48/2014 | UPMC-CLB Dept of Microbiology | Centers for Disease Control and Prevention |
| EPI556122 | HA | United States | 2014-Dec-08 | A/Mississippi/16/2014 | Mississippi Public Health Laboratory | Centers for Disease Control and Prevention |
| EPI556147 | HA | United States | 2014-Nov-24 | A/Wyoming/17/2014 | Wyoming Public Health Laboratory | Centers for Disease Control and Prevention |
| EPI556155 | HA | United States | 2014-Nov-05 | A/Alabama/09/2014 | ADPH Bureau of Clinical Laboratories | Centers for Disease Control and Prevention |
| EPI556163 | HA | United States | 2014-Nov-24 | A/Texas/70/2014 | Baylor College of Medicine | Centers for Disease Control and Prevention |
| EPI556171 | HA | United States | 2014-Nov-25 | A/Mississippi/15/2014 | Mississippi Public Health Laboratory | Centers for Disease Control and Prevention |
| EPI556190 | HA | United States | 2014-Dec-02 | A/Wisconsin/52/2014 | Marshfield Clinic Research Foundation | Centers for Disease Control and Prevention |
| EPI556203 | HA | United States | 2014-Nov-13 | A/North Carolina/31/2014 | North Carolina State Laboratory of Public Health | Centers for Disease Control and Prevention |
| EPI556211 | HA | United States | 2014-Nov-22 | A/Ohio/08/2014 | Ohio Department of Health Laboratories | Centers for Disease Control and Prevention |
| EPI556219 | HA | United States | 2014-Nov-10 | A/Michigan/18/2014 | University of Michigan SPH EPID | Centers for Disease Control and Prevention |
| EPI556227 | HA | United States | 2014-Dec-02 | A/Kentucky/15/2014 | Kentucky Division of Laboratory Services | Centers for Disease Control and Prevention |

| | | | | | | | |
|---|---|---|---|---|---|---|---|
| EPI556239 | HA | United States | 2014-Dec-01 | A/Washington/35/2014 | Marshfield Clinic Research Foundation | Centers for Disease Control and Prevention | |
| EPI556247 | HA | United States | 2014-Nov-18 | A/New Mexico/29/2014 | New Mexico Department of Health | Centers for Disease Control and Prevention | |
| EPI556255 | HA | United States | 2014-Nov-20 | A/New Jersey/29/2014 | New Jersey Department of Health & Senior Services | Centers for Disease Control and Prevention | |
| EPI556270 | HA | United States | 2014-Dec-08 | A/Rhode Island/22/2014 | Rhode Island Department of Health | Centers for Disease Control and Prevention | |
| EPI556278 | HA | United States | 2014-Nov-24 | A/Texas/68/2014 | Baylor College of Medicine | Centers for Disease Control and Prevention | Takashita,Emi; Fujisaki,Seiichiro; Shirakura,Masayuki; Watanabe,Shinji; Odagiri,Takato |
| EPI556286 | HA | United States | 2014-Nov-28 | A/Pennsylvania/49/2014 | UPMC-CLB Dept of Microbiology | Centers for Disease Control and Prevention | |
| EPI556294 | HA | United States | 2014-Nov-19 | A/Ohio/07/2014 | Ohio Department of Health Laboratories | Centers for Disease Control and Prevention | |
| EPI556302 | HA | United States | 2014-Nov-24 | A/Oklahoma/09/2014 | Oklahoma State Department of Health | Centers for Disease Control and Prevention | |
| EPI556317 | HA | United States | 2014-Dec-05 | A/Florida/65/2014 | Florida Department of Health-Jacksonville | Centers for Disease Control and Prevention | |
| EPI556325 | HA | United States | 2014-Dec-04 | A/Nevada/22/2014 | Southern Nevada Public Health Lab | Centers for Disease Control and Prevention | |
| EPI556333 | HA | United States | 2014-Dec-02 | A/Nevada/21/2014 | Southern Nevada Public Health Lab | Centers for Disease Control and Prevention | |
| EPI556341 | HA | United States | 2014-Dec-03 | A/Massachusetts/16/2014 | Massachusetts Department of Public Health | Centers for Disease Control and Prevention | |
| EPI556349 | HA | United States | 2014-Nov-10 | A/Michigan/17/2014 | University of Michigan SPH EPID | Centers for Disease Control and Prevention | |
| EPI556357 | HA | United States | 2014-Nov-27 | A/Indiana/21/2014 | Indiana State Department of Health Laboratories | Centers for Disease Control and Prevention | |
| EPI556365 | HA | United States | 2014-Nov-25 | A/Maryland/40/2014 | Maryland Department of Health and Mental Hygiene | Centers for Disease Control and Prevention | |
| EPI556467 | HA | Japan | 2014-Nov-25 | A/FUKUSHIMA/103/2014 | Fukushima Prefectural Institute of Public Health | National Institute of Infectious Diseases (NIID) | |
| EPI556512 | HA | United States | 2014-Nov-21 | A/Washington/33/2014 | | Centers for Disease Control and Prevention | |
| EPI556523 | HA | United States | 2014-Nov-13 | A/North Carolina/33/2014 | North Carolina State Laboratory of Public Health | Centers for Disease Control and Prevention | |
| EPI556531 | HA | United States | 2014-Dec-02 | A/Massachusetts/19/2014 | Massachusetts Department of Public Health | Centers for Disease Control and Prevention | |
| EPI556534 | HA | United States | 2014-Nov-11 | A/Indiana/17/2014 | Indiana State Department of Health Laboratories | Centers for Disease Control and Prevention | |
| EPI556548 | HA | United States | 2014-Dec-05 | A/Nebraska/15/2014 | Nebraska Public Health Lab | Centers for Disease Control and Prevention | |
| EPI556556 | HA | United States | 2014-Dec-14 | A/Vermont/09/2014 | Vermont Department of Health Laboratory | Centers for Disease Control and Prevention | |
| EPI556564 | HA | United States | 2014-Dec-10 | A/California/40/2014 | California Department of Health Services | Centers for Disease Control and Prevention | |
| EPI556574 | HA | United States | 2014-Nov-18 | A/Massachusetts/20/2014 | Massachusetts Department of Public Health | Centers for Disease Control and Prevention | |
| EPI556582 | HA | United States | 2014-Dec-13 | A/Vermont/10/2014 | Vermont Department of Health Laboratory | Centers for Disease Control and Prevention | |

| | | | | | | |
|---|---|---|---|---|---|---|
| EPI556590 | HA | United States | 2014-Dec-11 | A/Nebraska/14/2014 | Nebraska Public Health Lab | Centers for Disease Control and Prevention |
| EPI556613 | HA | United States | 2014-Dec-18 | A/Idaho/09/2014 | State of Idaho Bureau of Laboratories | Centers for Disease Control and Prevention |
| EPI556622 | HA | United States | 2014-Nov-21 | A/Oregon/11/2014 | Oregon Public Health Laboratory | Centers for Disease Control and Prevention |
| EPI556628 | HA | United States | 2014-Nov-25 | A/Michigan/26/2014 | University of Michigan SPH EPID | Centers for Disease Control and Prevention |
| EPI556637 | HA | United States | 2014-Nov-18 | A/Oregon/13/2014 | Oregon Public Health Laboratory | Centers for Disease Control and Prevention |
| EPI556645 | HA | United States | 2014-Nov-12 | A/Minnesota/34/2014 | Minnesota Department of Health | Centers for Disease Control and Prevention |
| EPI556653 | HA | United States | 2014-Dec-01 | A/Illinois/09/2014 | Illinois Department of Public Health-Chicago | Centers for Disease Control and Prevention |
| EPI556660 | HA | United States | 2014-Nov-16 | A/Minnesota/37/2014 | Minnesota Department of Health | Centers for Disease Control and Prevention |
| EPI556668 | HA | United States | 2014-Nov-13 | A/Hawaii/59/2014 | State of Hawaii Department of Health | Centers for Disease Control and Prevention |
| EPI556676 | HA | United States | 2014-Dec-11 | A/Virginia/30/2014 | Virginia Division of Consolidated Laboratories | Centers for Disease Control and Prevention |
| EPI556685 | HA | United States | 2014-Nov-03 | A/Illinois/05/2014 | Illinois Department of Public Health-Chicago | Centers for Disease Control and Prevention |
| EPI556695 | HA | United States | 2014-Nov-18 | A/Wyoming/15/2014 | Wyoming Public Health Laboratory | Centers for Disease Control and Prevention |
| EPI556704 | HA | United States | 2014-Nov-18 | A/Louisiana/17/2014 | Louisiana Department of Health and Hospitals | Centers for Disease Control and Prevention |
| EPI556712 | HA | United States | 2014-Nov-21 | A/Washington/36/2014 | Washington State Public Health Laboratory | Centers for Disease Control and Prevention |
| EPI556725 | HA | United States | 2014-Nov-14 | A/Georgia/13/2014 | Georgia Public Health Laboratory | Centers for Disease Control and Prevention |
| EPI556733 | HA | United States | 2014-Dec-11 | A/Florida/73/2014 | Florida Department of Health-Jacksonville | Centers for Disease Control and Prevention |
| EPI556741 | HA | United States | 2014-Dec-01 | A/Florida/76/2014 | Florida Department of Health-Tampa | Centers for Disease Control and Prevention |
| EPI556749 | HA | United States | 2014-Dec-16 | A/Idaho/08/2014 | State of Idaho Bureau of Laboratories | Centers for Disease Control and Prevention |
| EPI556757 | HA | United States | 2014-Dec-11 | A/Virginia/31/2014 | Virginia Division of Consolidated Laboratories | Centers for Disease Control and Prevention |
| EPI556759 | HA | United States | 2014-Oct-27 | A/Nevada/20/2014 | Nevada State Health Laboratory | Centers for Disease Control and Prevention |
| EPI556783 | HA | United States | 2014-Sep-16 | A/Maryland/26/2014 | Maryland Department of Health and Mental Hygiene | Centers for Disease Control and Prevention |
| EPI556791 | HA | United States | 2014-Nov-19 | A/Louisiana/20/2014 | Louisiana Department of Health and Hospitals | Centers for Disease Control and Prevention |
| EPI556816 | HA | United States | 2012-Apr-15 | A/Texas/50/2012 | Texas Department of State Health Services-Laboratory Services | Centers for Disease Control and Prevention |
| EPI556824 | HA | United States | 2014-Nov-19 | A/New Hampshire/23/2014 | New Hampshire Public Health Laboratories | Centers for Disease Control and Prevention |
| EPI557603 | HA | Japan | 2014-Nov-10 | A/TOYAMA/124/2014 | Toyama Institute of Health | National Institute of Infectious Diseases (NIID) |

| | | | | | | | |
|---|---|---|---|---|---|---|---|
| EPI557631 | HA | Japan | 2014-Dec-04 | A/TOKYO/41724/2014 | Tokyo Metropolitan Institute of Public Health | National Institute of Infectious Diseases (NIID) | |
| EPI558127 | HA | United States | 2014-Dec-07 | A/Texas/89/2014 | Baylor College of Medicine | Centers for Disease Control and Prevention | |
| EPI558135 | HA | United States | 2014-Dec-08 | A/Texas/72/2014 | Texas Department of State Health Services, South Texas Laboratory | Centers for Disease Control and Prevention | |
| EPI558146 | HA | United States | 2014-Dec-08 | A/Arkansas/14/2014 | Arkansas Department of Health | Centers for Disease Control and Prevention | |
| EPI558154 | HA | United States | 2014-Dec-22 | A/Nevada/27/2014 | Nevada State Health Laboratory | Centers for Disease Control and Prevention | |
| EPI558165 | HA | United States | 2014-Dec-15 | A/Texas/79/2014 | Texas Department of State Health Services, South Texas Laboratory | Centers for Disease Control and Prevention | Chen,B.B.; Phuah,S.P.; Poh,M.K.; Chen,S.L.; Cui,L. |
| EPI558168 | HA | United States | 2014-Oct-10 | A/California/27/2014 | California Department of Health Services | Centers for Disease Control and Prevention | Chen,B.B.; Phuah,S.P.; Poh,M.K.; Chen,S.L.; Cui,L. |
| EPI558176 | HA | United States | 2014-Dec-04 | A/Nevada/24/2014 | Nevada State Health Laboratory | Centers for Disease Control and Prevention | Chen,B.B.; Phuah,S.P.; Poh,M.K.; Chen,S.L.; Cui,L. |
| EPI558184 | HA | United States | 2014-Dec-08 | A/Texas/73/2014 | Texas Department of State Health Services, South Texas Laboratory | Centers for Disease Control and Prevention | Chen,B.B.; Phuah,S.P.; Poh,M.K.; Chen,S.L.; Cui,L. |
| EPI558199 | HA | United States | 2014-Dec-18 | A/Vermont/11/2014 | Vermont Department of Health Laboratory | Centers for Disease Control and Prevention | Chen,B.B.; Phuah,S.P.; Poh,M.K.; Chen,S.L.; Cui,L. |
| EPI558205 | HA | United States | 2014-Nov-08 | A/Florida/58/2014 | Florida Department of Health-Jacksonville | Centers for Disease Control and Prevention | Chen,B.B.; Phuah,S.P.; Poh,M.K.; Chen,S.L.; Cui,L. |
| EPI558213 | HA | United States | 2014-Dec-08 | A/Texas/74/2014 | Texas Department of State Health Services, South Texas Laboratory | Centers for Disease Control and Prevention | Chen,B.B.; Phuah,S.P.; Poh,M.K.; Chen,S.L.; Cui,L. |
| EPI558221 | HA | United States | 2014-Dec-04 | A/California/42/2014 | California Department of Health Services | Centers for Disease Control and Prevention | Takashita,Emi; Fujisaki,Seiichiro; Shirakura,Masayuki; Watanabe,Shinji; Odagiri,Takato |
| EPI558241 | HA | United States | 2014-Dec-09 | A/Ohio/10/2014 | Ohio Department of Health Laboratories | Centers for Disease Control and Prevention | |
| EPI558249 | HA | United States | 2014-Dec-11 | A/Vermont/12/2014 | Vermont Department of Health Laboratory | Centers for Disease Control and Prevention | |
| EPI558257 | HA | United States | 2014-Dec-01 | A/New Hampshire/30/2014 | New Hampshire Public Health Laboratories | Centers for Disease Control and Prevention | |
| EPI558265 | HA | United States | 2015-Jan-01 | A/Maine/01/2015 | Maine Health and Environmental Testing Laboratory | Centers for Disease Control and Prevention | |
| EPI558271 | HA | United States | 2014-Dec-02 | A/Texas/80/2014 | Texas Department of State Health Services-Laboratory Services | Centers for Disease Control and Prevention | |
| EPI558279 | HA | United States | 2014-Dec-30 | A/Maine/08/2014 | Maine Health and Environmental Testing Laboratory | Centers for Disease Control and Prevention | |
| EPI558287 | HA | United States | 2014-Dec-22 | A/Virginia/35/2014 | Virginia Division of Consolidated Laboratories | Centers for Disease Control and Prevention | |
| EPI558295 | HA | United States | 2014-Dec-05 | A/Utah/27/2014 | Utah Department of Health | Centers for Disease Control and Prevention | |
| EPI558303 | HA | United States | 2014-Dec-12 | A/Nevada/26/2014 | Nevada State Health Laboratory | Centers for Disease Control and Prevention | |
| EPI558311 | HA | United States | 2014-Dec-16 | A/Delaware/36/2014 | Delaware Public Health Lab | Centers for Disease Control and Prevention | |
| EPI558326 | HA | United States | 2014-Dec-07 | A/Delaware/35/2014 | Delaware Public Health Lab | Centers for Disease Control and Prevention | |
| EPI558336 | HA | United States | 2014-Dec-09 | A/Tennessee/18/2014 | Tennessee Department of Health Laboratory-Nashville | Centers for Disease Control and Prevention | |

| | | | | | | | |
|---|---|---|---|---|---|---|---|
| EPI558342 | HA | United States | 2014-Dec-05 | A/Utah/34/2014 | Utah Department of Health | Centers for Disease Control and Prevention | |
| EPI558350 | HA | United States | 2014-Dec-03 | A/North Dakota/14/2014 | North Dakota Department of Health | Centers for Disease Control and Prevention | |
| EPI558355 | HA | United States | 2014-Nov-09 | A/Colorado/25/2014 | Colorado Department of Health Lab | Centers for Disease Control and Prevention | |
| EPI558363 | HA | United States | 2014-Dec-11 | A/Texas/82/2014 | Texas Department of State Health Services-Laboratory Services | Centers for Disease Control and Prevention | |
| EPI558371 | HA | United States | 2014-Dec-10 | A/Texas/81/2014 | Texas Department of State Health Services-Laboratory Services | Centers for Disease Control and Prevention | |
| EPI558379 | HA | United States | 2014-Dec-15 | A/Nevada/25/2014 | Nevada State Health Laboratory | Centers for Disease Control and Prevention | |
| EPI558385 | HA | United States | 2014-Dec-09 | A/Alabama/12/2014 | ADPH Bureau of Clinical Laboratories | Centers for Disease Control and Prevention | |
| EPI558393 | HA | United States | 2014-Dec-08 | A/Virginia/33/2014 | University of Virginia, Medical Labs/Microbiology | Centers for Disease Control and Prevention | Takashita,Emi; Fujisaki,Seiichiro; Shirakura,Masayuki; Watanabe,Shinji; Odagiri,Takato |
| EPI558401 | HA | United States | 2014-Dec-10 | A/Missouri/16/2014 | Missouri Department. of Health & Senior Services | Centers for Disease Control and Prevention | |
| EPI558406 | HA | United States | 2014-Dec-08 | A/Mississippi/19/2014 | Mississippi Public Health Laboratory | Centers for Disease Control and Prevention | |
| EPI558410 | HA | United States | 2014-Dec-11 | A/Virginia/34/2014 | Virginia Division of Consolidated Laboratories | Centers for Disease Control and Prevention | |
| EPI558418 | HA | United States | 2014-Dec-08 | A/Tennessee/16/2014 | Tennessee Department of Health Laboratory-Nashville | Centers for Disease Control and Prevention | |
| EPI558426 | HA | United States | 2014-Dec-02 | A/Utah/33/2014 | Utah Department of Health | Centers for Disease Control and Prevention | |
| EPI558434 | HA | United States | 2014-Dec-08 | A/Tennessee/17/2014 | Tennessee Department of Health Laboratory-Nashville | Centers for Disease Control and Prevention | |
| EPI558439 | HA | United States | 2014-Dec-07 | A/Utah/35/2014 | Utah Department of Health | Centers for Disease Control and Prevention | |
| EPI558445 | HA | United States | 2014-Dec-04 | A/Alabama/10/2014 | ADPH Bureau of Clinical Laboratories | Centers for Disease Control and Prevention | |
| EPI558453 | HA | United States | 2014-Dec-09 | A/Tennessee/15/2014 | Tennessee Department of Health Laboratory-Nashville | Centers for Disease Control and Prevention | |
| EPI558461 | HA | United States | 2014-Dec-09 | A/Montana/27/2014 | Montana Laboratory Services Bureau | Centers for Disease Control and Prevention | |
| EPI558467 | HA | United States | 2014-Dec-08 | A/Alabama/11/2014 | ADPH Bureau of Clinical Laboratories | Centers for Disease Control and Prevention | |
| EPI558473 | HA | United States | 2014-Dec-04 | A/Ohio/09/2014 | Ohio Department of Health Laboratories | Centers for Disease Control and Prevention | |
| EPI558481 | HA | United States | 2014-Dec-10 | A/Wyoming/19/2014 | Wyoming Public Health Laboratory | Centers for Disease Control and Prevention | |
| EPI558487 | HA | United States | 2014-Dec-09 | A/Mississippi/20/2014 | Mississippi Public Health Laboratory | Centers for Disease Control and Prevention | Galiano,M |
| EPI558495 | HA | United States | 2014-Dec-02 | A/Texas/75/2014 | Texas Department of State Health Services-Laboratory Services | Centers for Disease Control and Prevention | |
| EPI558508 | HA | United States | 2014-Dec-05 | A/Indiana/25/2014 | Indiana State Department of Health Laboratories | Centers for Disease Control and Prevention | |
| EPI558516 | HA | United States | 2014-Dec-10 | A/Virginia/32/2014 | Virginia Division of Consolidated Laboratories | Centers for Disease Control and Prevention | |

| | | | | | | |
|---|---|---|---|---|---|---|
| EPI558524 | HA | United States | 2014-Dec-08 | A/Texas/71/2014 | Texas Department of State Health Services, South Texas Laboratory | Centers for Disease Control and Prevention |
| EPI558532 | HA | United States | 2014-Dec-29 | A/Maine/07/2014 | Maine Health and Environmental Testing Laboratory | Centers for Disease Control and Prevention |
| EPI558544 | HA | United States | 2014-Dec-08 | A/Montana/26/2014 | Montana Laboratory Services Bureau | Centers for Disease Control and Prevention |
| EPI558552 | HA | United States | 2014-Dec-23 | A/Washington/39/2014 | Washington State Public Health Laboratory | Centers for Disease Control and Prevention |
| EPI558560 | HA | United States | 2014-Dec-16 | A/Delaware/37/2014 | Delaware Public Health Lab | Centers for Disease Control and Prevention |
| EPI558568 | HA | United States | 2014-Dec-02 | A/Pennsylvania/52/2014 | Pennsylvania Department of Health | Centers for Disease Control and Prevention |
| EPI558574 | HA | United States | 2014-Dec-06 | A/Louisiana/38/2014 | Louisiana Department of Health and Hospitals | Centers for Disease Control and Prevention |
| EPI558580 | HA | United States | 2014-Dec-04 | A/California/44/2014 | California Department of Health Services | Centers for Disease Control and Prevention |
| EPI558592 | HA | United States | 2014-Dec-09 | A/Virginia/29/2014 | Virginia Division of Consolidated Laboratories | Centers for Disease Control and Prevention |
| EPI558597 | HA | United States | 2014-Dec-09 | A/Arkansas/15/2014 | Arkansas Department of Health | Centers for Disease Control and Prevention |
| EPI558965 | HA | Sweden | 2014-Dec-18 | A/Sweden/77/2014 | | Public Health Agency of Sweden |
| EPI558987 | HA | Sweden | 2014-Dec-30 | A/Stockholm/34/2014 | | Public Health Agency of Sweden |
| EPI558995 | HA | Sweden | 2014-Dec-27 | A/Stockholm/35/2014 | | Public Health Agency of Sweden |
| EPI559003 | HA | Sweden | 2014-Dec-31 | A/Skovde/1/2014 | | Public Health Agency of Sweden |
| EPI559011 | HA | Sweden | 2015-Jan-01 | A/Skovde/1/2015 | | Public Health Agency of Sweden |
| EPI559438 | HA | Japan | 2014-Feb-20 | A/Nagasaki/13N053/2014 | | Niigata University (DPH) |
| EPI560420 | HA | China | 2014-Nov-03 | A/Shaanxi-Changan/1452/2014 | WHO Chinese National Influenza Center | Centers for Disease Control and Prevention |
| EPI560439 | HA | China | 2014-Aug-04 | A/Beijing-Xicheng/13100/2014 | WHO Chinese National Influenza Center | Centers for Disease Control and Prevention |
| EPI560458 | HA | United States | 2014-Dec-23 | A/Nevada/29/2014 | Nevada State Health Laboratory | Centers for Disease Control and Prevention |
| EPI560463 | HA | Ethiopia | 2014-Oct-21 | A/Ethiopia/1141/2014 | Ethiopian Health and Nutrition Research Institute (EHNRI) | Centers for Disease Control and Prevention |
| EPI560471 | HA | Chile | 2014-Sep-16 | A/Iquique/62331/2014 | Instituto de Salud Publica de Chile | Centers for Disease Control and Prevention |
| EPI560479 | HA | United States | 2014-Dec-03 | A/Georgia/20/2014 | Georgia Public Health Laboratory | Centers for Disease Control and Prevention |
| EPI560498 | HA | Lao, People's Democratic Republic | 2014-Aug-06 | A/Laos/768/2014 | National Center for Laboratory and Epidemiology | Centers for Disease Control and Prevention |
| EPI560506 | HA | Lao, People's Democratic Republic | 2014-Aug-25 | A/Laos/842/2014 | National Center for Laboratory and Epidemiology | Centers for Disease Control and Prevention |
| EPI560514 | HA | Kenya | 2014-Aug-27 | A/Kenya/126/2014 | CDC-Kenya | Centers for Disease Control and Prevention |
| EPI560522 | HA | Lao, People's Democratic Republic | 2014-Aug-26 | A/Laos/864/2014 | National Center for Laboratory and Epidemiology | Centers for Disease Control and Prevention |

| | | | | | | |
|---|---|---|---|---|---|---|
| EPI560530 | HA | United States | 2014-Nov-14 | A/Georgia/19/2014 | Georgia Public Health Laboratory | Centers for Disease Control and Prevention |
| EPI560538 | HA | United States | 2014-Nov-27 | A/Georgia/16/2014 | Georgia Public Health Laboratory | Centers for Disease Control and Prevention |
| EPI560553 | HA | Lao, People's Democratic Republic | 2014-Aug-26 | A/Laos/859/2014 | National Center for Laboratory and Epidemiology | Centers for Disease Control and Prevention |
| EPI560561 | HA | Lao, People's Democratic Republic | 2014-Sep-08 | A/Laos/899/2014 | National Center for Laboratory and Epidemiology | Centers for Disease Control and Prevention |
| EPI560569 | HA | United States | 2014-Nov-29 | A/Georgia/18/2014 | Georgia Public Health Laboratory | Centers for Disease Control and Prevention |
| EPI560577 | HA | United States | 2014-Nov-26 | A/Georgia/23/2014 | Georgia Public Health Laboratory | Centers for Disease Control and Prevention |
| EPI560582 | HA | Mexico | 2014-Dec-04 | A/Mexico/2971/2014 | Laboratorio de Virus Respiratorio | Centers for Disease Control and Prevention |
| EPI560590 | HA | Chile | 2014-Oct-26 | A/Santiago/70123/2014 | Instituto de Salud Publica de Chile | Centers for Disease Control and Prevention |
| EPI560602 | HA | Canada | 2014-Oct-25 | A/Ontario/RV3241/2014 | National Microbiology Laboratory, Health Canada | Centers for Disease Control and Prevention |
| EPI560610 | HA | Korea, Republic of | 2014-Dec-15 | A/Busan/1996/2014 | National Institute of Health | Centers for Disease Control and Prevention |
| EPI560615 | HA | Mexico | 2014-Dec-08 | A/Mexico/2974/2014 | Laboratorio de Virus Respiratorio | Centers for Disease Control and Prevention |
| EPI560620 | HA | Canada | 2014-Oct-06 | A/Quebec/RV3213/2014 | National Microbiology Laboratory, Health Canada | Centers for Disease Control and Prevention |
| EPI560628 | HA | Kenya | 2014-Aug-27 | A/Kenya/124/2014 | CDC-Kenya | Centers for Disease Control and Prevention |
| EPI560636 | HA | Kenya | 2014-Aug-21 | A/Kenya/125/2014 | CDC-Kenya | Centers for Disease Control and Prevention |
| EPI560644 | HA | Kenya | 2014-Aug-26 | A/Kenya/128/2014 | CDC-Kenya | Centers for Disease Control and Prevention |
| EPI560653 | HA | United States | 2014-Nov-13 | A/Georgia/15/2014 | Georgia Public Health Laboratory | Centers for Disease Control and Prevention |
| EPI560664 | HA | Kenya | 2014-Aug-19 | A/Kenya/123/2014 | CDC-Kenya | Centers for Disease Control and Prevention |
| EPI560675 | HA | United States | 2014-Dec-15 | A/Hawaii/63/2014 | State of Hawaii Department of Health | Centers for Disease Control and Prevention |
| EPI560683 | HA | United States | 2014-Dec-10 | A/Missouri/18/2014 | Missouri Department. of Health & Senior Services | Centers for Disease Control and Prevention |
| EPI560691 | HA | United States | 2014-Nov-25 | A/Virginia/36/2014 | University of Virginia, Medical Labs/Microbiology | Centers for Disease Control and Prevention |
| EPI560699 | HA | Chile | 2014-Oct-20 | A/Santiago/68839/2014 | Instituto de Salud Publica de Chile | Centers for Disease Control and Prevention |
| EPI560707 | HA | United States | 2014-Dec-18 | A/Florida/80/2014 | Florida Department of Health-Tampa | Centers for Disease Control and Prevention |
| EPI560715 | HA | Paraguay | 2014-Aug-01 | A/Paraguay/0094/2014 | Central Laboratory of Public Health | Centers for Disease Control and Prevention |
| EPI560723 | HA | United States | 2014-Dec-13 | A/Pennsylvania/54/2014 | Pennsylvania Department of Health | Centers for Disease Control and Prevention |

| | | | | | | | |
|---|---|---|---|---|---|---|---|
| EPI560730 | HA | Canada | 2014-Dec-11 | A/Manitoba/RV3623/2014 | National Microbiology Laboratory, Health Canada | Centers for Disease Control and Prevention | |
| EPI560735 | HA | Canada | 2014-Oct-12 | A/Alberta/RV3246/2014 | National Microbiology Laboratory, Health Canada | Centers for Disease Control and Prevention | |
| EPI560747 | HA | Canada | 2014-Oct-04 | A/Alberta/RV3215/2014 | National Microbiology Laboratory, Health Canada | Centers for Disease Control and Prevention | |
| EPI560755 | HA | Kenya | 2014-Aug-25 | A/Kenya/122/2014 | CDC-Kenya | Centers for Disease Control and Prevention | |
| EPI560760 | HA | Canada | 2014-Oct-16 | A/Alberta/RV3240/2014 | National Microbiology Laboratory, Health Canada | Centers for Disease Control and Prevention | |
| EPI560768 | HA | Lao, People's Democratic Republic | 2014-Aug-18 | A/Laos/820/2014 | National Center for Laboratory and Epidemiology | Centers for Disease Control and Prevention | |
| EPI560783 | HA | Kenya | 2014-Aug-27 | A/Kenya/127/2014 | CDC-Kenya | Centers for Disease Control and Prevention | |
| EPI560791 | HA | China | 2014-Nov-04 | A/Xinjiang-Tianshan/11190/2014 | WHO Chinese National Influenza Center | Centers for Disease Control and Prevention | |
| EPI562067 | HA | United States | 2014-Dec-01 | A/Pennsylvania/56/2014 | UPMC-CLB Dept of Microbiology | Centers for Disease Control and Prevention | |
| EPI562097 | HA | Japan | 2014-Dec-25 | A/YOKOHAMA/184/2014 | Yokohama City Institute of Public Health. | National Institute of Infectious Diseases (NIID) | Resende,Paola; Motta,Fernando; Born,Priscila; Mesquita,Milene; Siqueira,Marilda |
| EPI562104 | HA | Japan | 2014-Dec-01 | A/NAGANO/3652/2014 | Nagano Environmental Conservation Research Institute | National Institute of Infectious Diseases (NIID) | Resende,Paola; Motta,Fernando; Born,Priscila; Mesquita,Milene; Siqueira,Marilda |
| EPI562136 | HA | Japan | 2014-Nov-09 | A/NIIGATA-C/60/2014 | Niigata City Institute of Public Health and Environment | National Institute of Infectious Diseases (NIID) | |
| EPI562142 | HA | Japan | 2014-Oct-31 | A/OSAKA-C/2020/2014 | Osaka City Institute of Public Health and Environmental Sciences | National Institute of Infectious Diseases (NIID) | |
| EPI562376 | HA | Japan | 2014-Mar-03 | A/Nagasaki/13N063/2014 | | Niigata University (DPH) | |
| EPI562622 | HA | Myanmar | 2013-Jun-27 | A/Myanmar/13M002/2013 | | Niigata University (DPH) | |
| EPI562630 | HA | Myanmar | 2013-Jul-02 | A/Myanmar/13M004/2013 | | Niigata University (DPH) | |
| EPI564072 | HA | United States | 2014-Dec-23 | A/California/43/2014 | California Department of Health Services | Centers for Disease Control and Prevention | |
| EPI564080 | HA | United States | 2015-Jan-02 | A/Maine/04/2015 | Maine Health and Environmental Testing Laboratory | Centers for Disease Control and Prevention | Deng,Y-M.; Iannello,P.; Spirason,N.; Jelley,L.; Lau,H.; |
| EPI564088 | HA | United States | 2014-Dec-17 | A/Wisconsin/69/2014 | Marshfield Clinic Research Foundation | Centers for Disease Control and Prevention | Deng,Y-M.; Iannello,P.; Spirason,N.; Jelley,L.; Lau,H.; |
| EPI564093 | HA | United States | 2014-Oct-19 | A/California/51/2014 | California Department of Health Services | Centers for Disease Control and Prevention | Deng,Y-D.; Iannello,P.; Spirason,N.; Jelley,L.; Lau,H.; |
| EPI564101 | HA | United States | 2014-Dec-26 | A/Missouri/20/2014 | Missouri Department. of Health & Senior Services | Centers for Disease Control and Prevention | |
| EPI564106 | HA | United States | 2014-Dec-17 | A/Missouri/21/2014 | Missouri Department. of Health & Senior Services | Centers for Disease Control and Prevention | |
| EPI564110 | HA | United States | 2014-Dec-11 | A/Massachusetts/25/2014 | Massachusetts Department of Public Health | Centers for Disease Control and Prevention | Li, J.; Shao, T.J.; Yu, X.F.; Pu, X.Y.; Pan, J.C. |
| EPI564118 | HA | United States | 2014-Dec-12 | A/Maryland/45/2014 | Maryland Department of Health and Mental Hygiene | Centers for Disease Control and Prevention | Li, J.; Shao, T.J.; Yu, X.F.; Pu, X.Y.; Pan, J.C. |
| EPI564126 | HA | United States | 2015-Jan-01 | A/Rhode Island/01/2015 | Rhode Island Department of Health | Centers for Disease Control and Prevention | Li, J.; Shao, T.J.; Yu, X.F.; Pu, X.Y.; Pan, J.C. |

| | | | | | | | |
|---|---|---|---|---|---|---|---|
| EPI564131 | HA | United States | 2014-Dec-14 | A/Massachusetts/23/2014 | Massachusetts Department of Public Health | Centers for Disease Control and Prevention | Li, J.; Shao, T.J.; Yu, X.F.; Pu, X.Y.; Pan, J.C. |
| EPI564139 | HA | United States | 2014-Dec-07 | A/Minnesota/38/2014 | Minnesota Department of Health | Centers for Disease Control and Prevention | Li, J.; Shao, T.J.; Yu, X.F.; Pu, X.Y.; Pan, J.C. |
| EPI564147 | HA | United States | 2014-Dec-17 | A/Massachusetts/22/2014 | Massachusetts Department of Public Health | Centers for Disease Control and Prevention | Li, J.; Shao, T.J.; Yu, X.F.; Pu, X.Y.; Pan, J.C. |
| EPI564155 | HA | United States | 2014-Dec-16 | A/Wisconsin/68/2014 | Marshfield Clinic Research Foundation | Centers for Disease Control and Prevention | Li, J.; Shao, T.J.; Yu, X.F.; Pu, X.Y.; Pan, J.C. |
| EPI564163 | HA | United States | 2014-Dec-23 | A/Missouri/19/2014 | Missouri Department. of Health & Senior Services | Centers for Disease Control and Prevention | Li, J.; Shao, T.J.; Yu, X.F.; Pu, X.Y.; Pan, J.C. |
| EPI564171 | HA | United States | 2014-Dec-19 | A/Rhode Island/23/2014 | Rhode Island Department of Health | Centers for Disease Control and Prevention | Li, J.; Shao, T.J.; Yu, X.F.; Pu, X.Y.; Pan, J.C. |
| EPI564179 | HA | United States | 2014-Dec-10 | A/Maryland/44/2014 | Maryland Department of Health and Mental Hygiene | Centers for Disease Control and Prevention | Li, J.; Shao, T.J.; Yu, X.F.; Pu, X.Y.; Pan, J.C. |
| EPI564187 | HA | United States | 2014-Dec-19 | A/Washington/48/2014 | Marshfield Clinic Research Foundation | Centers for Disease Control and Prevention | Li, J.; Shao, T.J.; Yu, X.F.; Pu, X.Y.; Pan, J.C. |
| EPI564195 | HA | United States | 2015-Jan-06 | A/Texas/02/2015 | Texas Childrens Hospital | Centers for Disease Control and Prevention | Li, J.; Shao, T.J.; Yu, X.F.; Pu, X.Y.; Pan, J.C. |
| EPI564203 | HA | United States | 2015-Jan-02 | A/Pennsylvania/01/2015 | Pennsylvania Department of Health | Centers for Disease Control and Prevention | Li, J.; Shao, T.J.; Yu, X.F.; Pu, X.Y.; Pan, J.C. |
| EPI564211 | HA | United States | 2015-Jan-09 | A/Pennsylvania/02/2015 | Pennsylvania Department of Health | Centers for Disease Control and Prevention | |
| EPI564225 | HA | United States | 2014-Oct-28 | A/Minnesota/31/2014 | Minnesota Department of Health | Centers for Disease Control and Prevention | |
| EPI564241 | HA | United States | 2014-Dec-15 | A/Hawaii/63/2014 | State of Hawaii Department of Health | Centers for Disease Control and Prevention | |
| EPI564254 | HA | United States | 2014-Dec-03 | A/Colorado/38/2014 | Colorado Department of Health Lab | Centers for Disease Control and Prevention | |
| EPI564268 | HA | United States | 2014-Dec-27 | A/District Of Columbia/08/2014 | DC Public Health Lab | Centers for Disease Control and Prevention | |
| EPI564282 | HA | United States | 2014-Dec-28 | A/District Of Columbia/10/2014 | DC Public Health Lab | Centers for Disease Control and Prevention | |
| EPI564298 | HA | Paraguay | 2014-Nov-20 | A/Paraguay/3144/2014 | Central Laboratory of Public Health | Centers for Disease Control and Prevention | |
| EPI564307 | HA | Ethiopia | 2014-Nov-27 | A/Ethiopia/155/2014 | Ethiopian Health and Nutrition Research Institute (EHNRI) | Centers for Disease Control and Prevention | |
| EPI564312 | HA | Ethiopia | 2014-Oct-23 | A/Ethiopia/1146/2014 | Ethiopian Health and Nutrition Research Institute (EHNRI) | Centers for Disease Control and Prevention | |
| EPI564320 | HA | United States | 2014-Dec-01 | A/Indiana/24/2014 | Indiana State Department of Health Laboratories | Centers for Disease Control and Prevention | |
| EPI564325 | HA | Guatemala | 2014-Nov-12 | A/Guatemala/4638/2014 | Laboratorio Nacional De Salud Guatemala | Centers for Disease Control and Prevention | |
| EPI564330 | HA | Guatemala | 2014-Nov-13 | A/Guatemala/5211/2014 | Laboratorio Nacional De Salud Guatemala | Centers for Disease Control and Prevention | |
| EPI564336 | HA | United States | 2014-Dec-04 | A/Tennessee/23/2014 | Tennessee Department of Health Laboratory-Nashville | Centers for Disease Control and Prevention | Galiano,M. |
| EPI564352 | HA | United States | 2014-Nov-14 | A/Iowa/20/2014 | Iowa State Hygienic Laboratory | Centers for Disease Control and Prevention | Galiano,M. |
| EPI564362 | HA | United States | 2014-Dec-03 | A/Illinois/12/2014 | Illinois Department of Public Health-Chicago | Centers for Disease Control and Prevention | Galiano,M. |

| | | | | | | | |
|---|---|---|---|---|---|---|---|
| EPI564365 | HA | United States | 2014-Dec-04 | A/Minnesota/39/2014 | Minnesota Department of Health | Centers for Disease Control and Prevention | Galiano,M. |
| EPI564374 | HA | United States | 2014-Dec-02 | A/Delaware/38/2014 | Delaware Public Health Lab | Centers for Disease Control and Prevention | Galiano,M. |
| EPI564382 | HA | United States | 2014-Dec-04 | A/Missouri/17/2014 | Missouri Department. of Health & Senior Services | Centers for Disease Control and Prevention | Galiano,M. |
| EPI564395 | HA | United States | 2014-Nov-23 | A/Minnesota/41/2014 | Minnesota Department of Health | Centers for Disease Control and Prevention | Galiano,M. |
| EPI564414 | HA | United States | 2014-Dec-15 | A/Tennessee/20/2014 | Tennessee Department of Health Laboratory-Nashville | Centers for Disease Control and Prevention | Galiano,M. |
| EPI564422 | HA | United States | 2014-Dec-03 | A/North Carolina/41/2014 | North Carolina State Laboratory of Public Health | Centers for Disease Control and Prevention | Galiano,M. |
| EPI564430 | HA | Canada | 2014-Oct-12 | A/Alberta/RV3246/2014 | National Microbiology Laboratory, Health Canada | Centers for Disease Control and Prevention | Galiano,M. |
| EPI564441 | HA | United States | 2014-Dec-09 | A/Wyoming/20/2014 | Wyoming Public Health Laboratory | Centers for Disease Control and Prevention | Galiano,M. |
| EPI564449 | HA | Hong Kong (SAR) | 2014-Nov-29 | A/Hong Kong/7617/2014 | Government Virus Unit | Centers for Disease Control and Prevention | Galiano,M. |
| EPI564457 | HA | United States | 2014-Dec-08 | A/Indiana/26/2014 | Indiana State Department of Health Laboratories | Centers for Disease Control and Prevention | Galiano,M. |
| EPI564465 | HA | United States | 2014-Dec-18 | A/Illinois/14/2014 | Illinois Department of Public Health-Chicago | Centers for Disease Control and Prevention | Galiano,M. |
| EPI564473 | HA | United States | 2014-Dec-12 | A/Colorado/35/2014 | Colorado Department of Health Lab | Centers for Disease Control and Prevention | Galiano,M. |
| EPI564481 | HA | United States | 2014-Dec-17 | A/Kentucky/21/2014 | Kentucky Division of Laboratory Services | Centers for Disease Control and Prevention | Galiano,M. |
| EPI564492 | HA | United States | 2014-Dec-12 | A/Montana/28/2014 | Montana Laboratory Services Bureau | Centers for Disease Control and Prevention | Galiano,M. |
| EPI564498 | HA | United States | 2014-Dec-18 | A/Montana/30/2014 | Montana Laboratory Services Bureau | Centers for Disease Control and Prevention | Galiano,M. |
| EPI564506 | HA | United States | 2014-Dec-02 | A/Illinois/11/2014 | Illinois Department of Public Health-Chicago | Centers for Disease Control and Prevention | Galiano,M. |
| EPI564520 | HA | United States | 2014-Dec-08 | A/New Hampshire/32/2014 | New Hampshire Public Health Laboratories | Centers for Disease Control and Prevention | Galiano,M. |
| EPI564535 | HA | Canada | 2014-Oct-25 | A/Ontario/RV3241/2014 | National Microbiology Laboratory, Health Canada | Centers for Disease Control and Prevention | Galiano,M. |
| EPI564543 | HA | United States | 2014-Dec-29 | A/Delaware/39/2014 | Delaware Public Health Lab | Centers for Disease Control and Prevention | Galiano,M. |
| EPI564554 | HA | United States | 2014-Dec-14 | A/West Virginia/06/2014 | West Virginia Office of Laboratory Services | Centers for Disease Control and Prevention | Galiano,M. |
| EPI564561 | HA | United States | 2014-Dec-17 | A/Colorado/36/2014 | Colorado Department of Health Lab | Centers for Disease Control and Prevention | Galiano,M. |
| EPI564574 | HA | United States | 2014-Dec-22 | A/Delaware/42/2014 | Delaware Public Health Lab | Centers for Disease Control and Prevention | Galiano,M. |
| EPI564585 | HA | United States | 2014-Dec-31 | A/New York/39/2014 | New York City Department of Health | Centers for Disease Control and Prevention | Galiano,M. |
| EPI564592 | HA | United States | 2014-Dec-29 | A/New York/40/2014 | New York City Department of Health | Centers for Disease Control and Prevention | Galiano,M. |
| EPI564612 | HA | United States | 2014-Oct-28 | A/Arkansas/13/2014 | Arkansas Department of Health | Centers for Disease Control and Prevention | Galiano,M. |

| | | | | | | | |
|---|---|---|---|---|---|---|---|
| EPI564618 | HA | United States | 2014-Dec-17 | A/Washington/46/2014 | Seattle & King County Public Health Lab | Centers for Disease Control and Prevention | Galiano,M. |
| EPI564630 | HA | United States | 2014-Dec-04 | A/North Carolina/40/2014 | North Carolina State Laboratory of Public Health | Centers for Disease Control and Prevention | Galiano,M. |
| EPI564634 | HA | United States | 2014-Dec-24 | A/Hawaii/65/2014 | State of Hawaii Department of Health | Centers for Disease Control and Prevention | Galiano,M. |
| EPI564642 | HA | United States | 2014-Dec-17 | A/Nevada/30/2014 | Southern Nevada Public Health Lab | Centers for Disease Control and Prevention | Galiano,M. |
| EPI564650 | HA | United States | 2014-Dec-09 | A/Pennsylvania/65/2014 | UPMC-CLB Dept of Microbiology | Centers for Disease Control and Prevention | Galiano,M. |
| EPI564658 | HA | United States | 2014-Dec-04 | A/New Hampshire/29/2014 | New Hampshire Public Health Laboratories | Centers for Disease Control and Prevention | Galiano,M. |
| EPI564666 | HA | United States | 2014-Dec-11 | A/Pennsylvania/64/2014 | UPMC-CLB Dept of Microbiology | Centers for Disease Control and Prevention | Galiano,M. |
| EPI564674 | HA | United States | 2014-Dec-13 | A/Pennsylvania/63/2014 | UPMC-CLB Dept of Microbiology | Centers for Disease Control and Prevention | Galiano,M. |
| EPI564679 | HA | Ethiopia | 2014-Oct-27 | A/Ethiopia/1147/2014 | Ethiopian Health and Nutrition Research Institute (EHNRI) | Centers for Disease Control and Prevention | Galiano,M. |
| EPI564692 | HA | United States | 2014-Dec-05 | A/Virginia/37/2014 | University of Virginia, Medical Labs/Microbiology | Centers for Disease Control and Prevention | Galiano,M. |
| EPI564700 | HA | United States | 2014-Dec-02 | A/Georgia/21/2014 | Georgia Public Health Laboratory | Centers for Disease Control and Prevention | Galiano,M. |
| EPI564716 | HA | United States | 2014-Dec-02 | A/Missouri/14/2014 | Missouri Department. of Health & Senior Services | Centers for Disease Control and Prevention | Galiano,M. |
| EPI564724 | HA | United States | 2014-Dec-19 | A/District Of Columbia/05/2014 | DC Public Health Lab | Centers for Disease Control and Prevention | Galiano,M. |
| EPI564732 | HA | United States | 2014-Dec-26 | A/District Of Columbia/04/2014 | DC Public Health Lab | Centers for Disease Control and Prevention | Takashita,Emi; Fujisaki,Seiichiro; Shirakura,Masayuki; Watanabe,Shinji; Odagiri,Takato |
| EPI564737 | HA | United States | 2014-Oct-30 | A/District Of Columbia/03/2014 | DC Public Health Lab | Centers for Disease Control and Prevention | Takashita,Emi; Fujisaki,Seiichiro; Shirakura,Masayuki; Watanabe,Shinji; Odagiri,Takato |
| EPI564745 | HA | United States | 2015-Jan-07 | A/Florida/04/2015 | Florida Department of Health-Tampa | Centers for Disease Control and Prevention | Takashita,Emi; Fujisaki,Seiichiro; Shirakura,Masayuki; Watanabe,Shinji; Odagiri,Takato |
| EPI564753 | HA | United States | 2015-Jan-11 | A/Florida/05/2015 | Florida Department of Health-Tampa | Centers for Disease Control and Prevention | Galiano,M. |
| EPI564766 | HA | United States | 2014-Nov-09 | A/Ohio/14/2014 | Ohio Department of Health Laboratories | Centers for Disease Control and Prevention | Galiano,M. |
| EPI564771 | HA | United States | 2014-Oct-18 | A/California/50/2014 | California Department of Health Services | Centers for Disease Control and Prevention | Galiano,M. |
| EPI564782 | HA | United States | 2014-Oct-24 | A/California/52/2014 | California Department of Health Services | Centers for Disease Control and Prevention | Galiano,M. |
| EPI564790 | HA | United States | 2014-Dec-02 | A/Pennsylvania/51/2014 | Pennsylvania Department of Health | Centers for Disease Control and Prevention | Galiano,M. |
| EPI564794 | HA | United States | 2014-Oct-26 | A/California/53/2014 | California Department of Health Services | Centers for Disease Control and Prevention | Galiano,M. |
| EPI564802 | HA | United States | 2014-Dec-17 | A/Washington/47/2014 | Marshfield Clinic Research Foundation | Centers for Disease Control and Prevention | Galiano,M. |
| EPI564839 | HA | Myanmar | 2013-Jul-11 | A/Myanmar/13M016/2013 | | Niigata University (DPH) | Galiano,M. |

| | | | | | | | |
|---|---|---|---|---|---|---|---|
| EPI564847 | HA | Myanmar | 2013-Jul-15 | A/Myanmar/13M029/2013 | | Niigata University (DPH) | Galiano,M. |
| EPI564855 | HA | Myanmar | 2013-Jul-16 | A/Myanmar/13M056/2013 | | Niigata University (DPH) | Galiano,M. |
| EPI564863 | HA | Myanmar | 2013-Jul-19 | A/Myanmar/13M082/2013 | | Niigata University (DPH) | Galiano,M. |
| EPI565359 | HA | United States | 2015-Jan-01 | A/Maine/03/2015 | Maine Health and Environmental Testing Laboratory | Centers for Disease Control and Prevention | Galiano,M. |
| EPI565365 | HA | Mexico | 2014-Nov-26 | A/Mexico/2789/2014 | Laboratorio de Virus Respiratorio | Centers for Disease Control and Prevention | Galiano,M. |
| EPI565380 | HA | Ethiopia | 2014-Oct-20 | A/Ethiopia/1140/2014 | Ethiopian Health and Nutrition Research Institute (EHNRI) | Centers for Disease Control and Prevention | Galiano,M. |
| EPI565385 | HA | Mexico | 2014-Nov-24 | A/Mexico/2777/2014 | Laboratorio de Virus Respiratorio | Centers for Disease Control and Prevention | Galiano,M. |
| EPI565393 | HA | Venezuela, Bolivarian Republic of | 2014-Aug-04 | A/Venezuela/07/2014 | Instituto Nacional de Higiene "Rafael Rangel" | Centers for Disease Control and Prevention | Galiano,M. |
| EPI565397 | HA | Thailand | 2014-Dec-26 | A/Nonthaburi/01/2015 | WHO National Influenza Centre, National Institute of Medical Research (NIMR) | Centers for Disease Control and Prevention | Galiano,M. |
| EPI565401 | HA | Guatemala | 2014-Nov-12 | A/Guatemala/4645/2014 | Laboratorio Nacional De Salud Guatemala | Centers for Disease Control and Prevention | Galiano,M. |
| EPI565405 | HA | Guatemala | 2014-Nov-14 | A/Guatemala/4647/2014 | Laboratorio Nacional De Salud Guatemala | Centers for Disease Control and Prevention | Galiano,M. |
| EPI565425 | HA | United States | 2014-Nov-29 | A/Colorado/39/2014 | Colorado Department of Health Lab | Centers for Disease Control and Prevention | Galiano,M. |
| EPI565433 | HA | Tanzania, United Republic of | 2014-Nov-26 | A/Tanzania/3580/2014 | National Influenza Lab | Centers for Disease Control and Prevention | Galiano,M. |
| EPI565470 | HA | United States | 2015-Jan-01 | A/Kentucky/01/2015 | Kentucky Division of Laboratory Services | Centers for Disease Control and Prevention | Galiano,M. |
| EPI565478 | HA | United States | 2014-Dec-03 | A/Florida/85/2014 | Florida Department of Health-Tampa | Centers for Disease Control and Prevention | Galiano,M. |
| EPI565490 | HA | United States | 2015-Jan-05 | A/Florida/09/2015 | Florida Department of Health-Jacksonville | Centers for Disease Control and Prevention | Galiano,M. |
| EPI565498 | HA | United States | 2014-Dec-27 | A/Ohio/12/2014 | Ohio Department of Health Laboratories | Centers for Disease Control and Prevention | Galiano,M. |
| EPI565506 | HA | United States | 2015-Jan-05 | A/Florida/07/2015 | Florida Department of Health-Jacksonville | Centers for Disease Control and Prevention | Galiano,M. |
| EPI565514 | HA | United States | 2015-Jan-02 | A/Utah/01/2015 | Utah Department of Health | Centers for Disease Control and Prevention | |
| EPI565522 | HA | United States | 2015-Jan-09 | A/Montana/02/2015 | Montana Laboratory Services Bureau | Centers for Disease Control and Prevention | |
| EPI565528 | HA | United States | 2015-Jan-03 | A/Florida/06/2015 | Florida Department of Health-Jacksonville | Centers for Disease Control and Prevention | |
| EPI565536 | HA | United States | 2015-Jan-06 | A/Montana/01/2015 | Montana Laboratory Services Bureau | Centers for Disease Control and Prevention | |
| EPI565544 | HA | Tanzania, United Republic of | 2014-Nov-20 | A/Tanzania/3562/2014 | National Influenza Lab | Centers for Disease Control and Prevention | |
| EPI565549 | HA | United States | 2014-Dec-17 | A/Delaware/43/2014 | Delaware Public Health Lab | Centers for Disease Control and Prevention | |
| EPI565557 | HA | Tanzania, United Republic of | 2014-Nov-18 | A/Tanzania/3557/2014 | National Influenza Lab | Centers for Disease Control and Prevention | |

| | | | | | | | |
|---|---|---|---|---|---|---|---|
| EPI565563 | HA | Ethiopia | 2014-Nov-11 | A/Ethiopia/1153/2014 | Ethiopian Health and Nutrition Research Institute (EHNRI) | Centers for Disease Control and Prevention | |
| EPI565571 | HA | United States | 2015-Jan-08 | A/Utah/02/2015 | Utah Department of Health | Centers for Disease Control and Prevention | |
| EPI565579 | HA | United States | 2015-Jan-02 | A/West Virginia/01/2015 | West Virginia Office of Laboratory Services | Centers for Disease Control and Prevention | |
| EPI565587 | HA | Tanzania, United Republic of | 2014-Nov-24 | A/Tanzania/562/2014 | National Influenza Lab | Centers for Disease Control and Prevention | |
| EPI565595 | HA | United States | 2014-Dec-15 | A/Washington/52/2014 | Marshfield Clinic Research Foundation | Centers for Disease Control and Prevention | |
| EPI565603 | HA | United States | 2015-Jan-02 | A/Wisconsin/01/2015 | Marshfield Clinic Research Foundation | Centers for Disease Control and Prevention | |
| EPI565611 | HA | United States | 2015-Jan-06 | A/Wisconsin/02/2015 | Marshfield Clinic Research Foundation | Centers for Disease Control and Prevention | |
| EPI565619 | HA | Tanzania, United Republic of | 2014-Nov-14 | A/Tanzania/542/2014 | National Influenza Lab | Centers for Disease Control and Prevention | |
| EPI565624 | HA | United States | 2014-Nov-13 | A/Mississippi/17/2014 | Mississippi Public Health Laboratory | Centers for Disease Control and Prevention | |
| EPI565636 | HA | United States | 2015-Jan-08 | A/Arkansas/01/2015 | Arkansas Department of Health | Centers for Disease Control and Prevention | |
| EPI565648 | HA | United States | 2015-Jan-01 | A/Oregon/01/2015 | Oregon Public Health Laboratory | Centers for Disease Control and Prevention | |
| EPI565658 | HA | Canada | 2014-Nov-03 | A/Ontario/RV3531/2014 | National Microbiology Laboratory, Health Canada | Centers for Disease Control and Prevention | |
| EPI565661 | HA | Canada | 2014-Dec-07 | A/Ontario/RV3606/2014 | National Microbiology Laboratory, Health Canada | Centers for Disease Control and Prevention | |
| EPI565665 | HA | Canada | 2014-Nov-21 | A/Alberta/RV3591/2014 | National Microbiology Laboratory, Health Canada | Centers for Disease Control and Prevention | |
| EPI565675 | HA | Canada | 2014-Dec-18 | A/Ontario/RV3609/2014 | National Microbiology Laboratory, Health Canada | Centers for Disease Control and Prevention | Takashita,Emi; Fujisaki,Seiichiro; Shirakura,Masayuki; Watanabe,Shinji; Odagiri,Takato |
| EPI565680 | HA | Hong Kong (SAR) | 2014-Nov-29 | A/Hong Kong/7615/2014 | Government Virus Unit | Centers for Disease Control and Prevention | |
| EPI565694 | HA | Paraguay | 2014-Dec-04 | A/Paraguay/4413/2014 | Central Laboratory of Public Health | Centers for Disease Control and Prevention | |
| EPI565707 | HA | Tanzania, United Republic of | 2014-Dec-05 | A/Tanzania/594/2014 | National Influenza Lab | Centers for Disease Control and Prevention | Takashita,Emi; Fujisaki,Seiichiro; Shirakura,Masayuki; Watanabe,Shinji; Odagiri,Takato |
| EPI565715 | HA | Tanzania, United Republic of | 2014-Dec-05 | A/Tanzania/596/2014 | National Influenza Lab | Centers for Disease Control and Prevention | Takashita,Emi; Fujisaki,Seiichiro; Shirakura,Masayuki; Watanabe,Shinji; Odagiri,Takato |
| EPI565723 | HA | United States | 2014-Dec-04 | A/Pennsylvania/67/2014 | UPMC-CLB Dept of Microbiology | Centers for Disease Control and Prevention | |
| EPI565731 | HA | Tanzania, United Republic of | 2014-Dec-03 | A/Tanzania/590/2014 | National Influenza Lab | Centers for Disease Control and Prevention | |
| EPI565739 | HA | Tanzania, United Republic of | 2014-Dec-06 | A/Tanzania/599/2014 | National Influenza Lab | Centers for Disease Control and Prevention | |
| EPI565747 | HA | Chile | 2014-Oct-20 | A/Santiago/68839/2014 | Instituto de Salud Publica de Chile | Centers for Disease Control and Prevention | Saito, Reiko; Takemae, Nobuhiro; Saito, Takehiko; Shobugawa, Yugo; Kondo, Hiroki; Hibino, Akinobu; Masaki, Hironori |

| | | | | | | | |
|---|---|---|---|---|---|---|---|
| EPI565755 | HA | Ethiopia | 2014-Sep-04 | A/Ethiopia/159/2014 | Ethiopian Health and Nutrition Research Institute (EHNRI) | Centers for Disease Control and Prevention | Saito, Reiko; Takemae, Nobuhiro; Saito, Takehiko; Shobugawa, Yugo; Kondo, Hiroki; Hibino, Akinobu; Makiya, |
| EPI565763 | HA | Korea, Republic of | 2014-Dec-22 | A/Seoul/2007/2014 | National Institute of Health | Centers for Disease Control and Prevention | Saito, Reiko; Takemae, Nobuhiro; Saito, Takehiko; Shobugawa, Yugo; Kondo, Hiroki; Hibino, Akinobu; Shirahige, Yutaka |
| EPI565771 | HA | Tanzania, United Republic of | 2014-Nov-14 | A/Tanzania/3552/2014 | National Influenza Lab | Centers for Disease Control and Prevention | Saito, Reiko; Takemae, Nobuhiro; Saito, Takehiko; Shobugawa, Yugo; Kondo, Hiroki; Hibino, Akinobu; Asoh, Norichika; Kita, |
| EPI565777 | HA | United States | 2015-Jan-02 | A/Delaware/02/2015 | Delaware Public Health Lab | Centers for Disease Control and Prevention | Saito, Reiko; Takemae, Nobuhiro; Saito, Takehiko; Shobugawa, Yugo; Kondo, Hiroki; Hibino, Akinobu; Shirahige, Yutaka |
| EPI565785 | HA | United States | 2015-Jan-02 | A/Delaware/01/2015 | Delaware Public Health Lab | Centers for Disease Control and Prevention | Saito, Reiko; Takemae, Nobuhiro; Saito, Takehiko; Shobugawa, Yugo; Kondo, Hiroki; Hibino, |
| EPI565792 | HA | Tanzania, United Republic of | 2014-Nov-25 | A/Tanzania/2665/2014 | National Influenza Lab | Centers for Disease Control and Prevention | |
| EPI565797 | HA | Tanzania, United Republic of | 2014-Dec-01 | A/Tanzania/588/2014 | National Influenza Lab | Centers for Disease Control and Prevention | |
| EPI565804 | HA | Mexico | 2014-Dec-02 | A/Mexico/836/2014 | Laboratorio de Virus Respiratorio | Centers for Disease Control and Prevention | |
| EPI565812 | HA | Tanzania, United Republic of | 2014-Dec-03 | A/Tanzania/3588/2014 | National Influenza Lab | Centers for Disease Control and Prevention | |
| EPI565820 | HA | Tanzania, United Republic of | 2014-Dec-12 | A/Tanzania/609/2014 | National Influenza Lab | Centers for Disease Control and Prevention | |
| EPI565828 | HA | Tanzania, United Republic of | 2014-Dec-03 | A/Tanzania/3589/2014 | National Influenza Lab | Centers for Disease Control and Prevention | |
| EPI565836 | HA | Mexico | 2014-Dec-01 | A/Mexico/2865/2014 | Laboratorio de Virus Respiratorio | Centers for Disease Control and Prevention | |
| EPI565844 | HA | Mexico | 2014-Dec-09 | A/Mexico/3005/2014 | Laboratorio de Virus Respiratorio | Centers for Disease Control and Prevention | |
| EPI565849 | HA | Mexico | 2014-Dec-17 | A/Mexico/3040/2014 | Laboratorio de Virus Respiratorio | Centers for Disease Control and Prevention | |
| EPI565851 | HA | Ethiopia | 2014-Oct-23 | A/Ethiopia/546/2014 | Ethiopian Health and Nutrition Research Institute (EHNRI) | Centers for Disease Control and Prevention | |
| EPI565859 | HA | United States | 2015-Jan-02 | A/Florida/02/2015 | Florida Department of Health-Tampa | Centers for Disease Control and Prevention | |
| EPI565867 | HA | United States | 2015-Jan-02 | A/Florida/03/2015 | Florida Department of Health-Tampa | Centers for Disease Control and Prevention | |
| EPI565875 | HA | Vietnam | 2014-Aug-15 | A/Vietnam/594/2014 | National Institute of Hygiene and Epidemiology | Centers for Disease Control and Prevention | |
| EPI565883 | HA | Tanzania, United Republic of | 2014-Nov-16 | A/Tanzania/3555/2014 | National Influenza Lab | Centers for Disease Control and Prevention | |
| EPI565891 | HA | Tanzania, United Republic of | 2014-Nov-20 | A/Tanzania/3561/2014 | National Influenza Lab | Centers for Disease Control and Prevention | |
| EPI565903 | HA | Tanzania, United Republic of | 2014-Dec-05 | A/Tanzania/597/2014 | National Influenza Lab | Centers for Disease Control and Prevention | |

| | | | | | | |
|---|---|---|---|---|---|---|
| EPI565911 | HA | Tanzania, United Republic of | 2014-Nov-18 | A/Tanzania/3556/2014 | National Influenza Lab | Centers for Disease Control and Prevention |
| EPI565919 | HA | United States | 2015-Jan-02 | A/Florida/01/2015 | Florida Department of Health-Tampa | Centers for Disease Control and Prevention |
| EPI565927 | HA | Tanzania, United Republic of | 2014-Nov-19 | A/Tanzania/3559/2014 | National Influenza Lab | Centers for Disease Control and Prevention |
| EPI565935 | HA | Tanzania, United Republic of | 2014-Nov-19 | A/Tanzania/553/2014 | National Influenza Lab | Centers for Disease Control and Prevention |
| EPI565945 | HA | United States | 2014-Nov-11 | A/Ohio/15/2014 | Ohio Department of Health Laboratories | Centers for Disease Control and Prevention |
| EPI565953 | HA | United States | 2015-Jan-07 | A/Florida/12/2015 | Florida Department of Health-Tampa | Centers for Disease Control and Prevention |
| EPI565961 | HA | United States | 2015-Jan-06 | A/Florida/11/2015 | Florida Department of Health-Tampa | Centers for Disease Control and Prevention |
| EPI565969 | HA | United States | 2015-Jan-11 | A/Florida/13/2015 | Florida Department of Health-Tampa | Centers for Disease Control and Prevention |
| EPI565977 | HA | Ethiopia | 2014-Nov-12 | A/Ethiopia/1159/2014 | Ethiopian Health and Nutrition Research Institute (EHNRI) | Centers for Disease Control and Prevention |
| EPI565985 | HA | United States | 2015-Jan-02 | A/Florida/10/2015 | Florida Department of Health-Tampa | Centers for Disease Control and Prevention |
| EPI565993 | HA | Thailand | 2014-Dec-09 | A/Nonthaburi/2421/2014 | WHO National Influenza Centre, National Institute of Medical Research (NIMR) | Centers for Disease Control and Prevention |
| EPI566001 | HA | Tanzania, United Republic of | 2014-Dec-11 | A/Tanzania/3594/2014 | National Influenza Lab | Centers for Disease Control and Prevention |
| EPI566009 | HA | Guatemala | 2014-Nov-04 | A/Guatemala/179/2014 | Laboratorio Nacional De Salud Guatemala | Centers for Disease Control and Prevention |
| EPI566017 | HA | Thailand | 2014-Dec-11 | A/Chanthaburi/2419/2014 | WHO National Influenza Centre, National Institute of Medical Research (NIMR) | Centers for Disease Control and Prevention |
| EPI566025 | HA | Tanzania, United Republic of | 2014-Dec-09 | A/Tanzania/1906/2014 | National Influenza Lab | Centers for Disease Control and Prevention |
| EPI566147 | HA | United States | 2014-Dec-16 | A/Idaho/12/2014 | State of Idaho Bureau of Laboratories | Centers for Disease Control and Prevention |
| EPI566159 | HA | United States | 2014-Dec-31 | A/New York/38/2014 | New York City Department of Health | Centers for Disease Control and Prevention |
| EPI566167 | HA | United States | 2014-Dec-11 | A/Texas/85/2014 | Texas Department of State Health Services-Laboratory Services | Centers for Disease Control and Prevention |
| EPI566172 | HA | United States | 2014-Dec-09 | A/Wyoming/18/2014 | Wyoming Public Health Laboratory | Centers for Disease Control and Prevention |
| EPI566180 | HA | United States | 2014-Dec-11 | A/Texas/84/2014 | Texas Department of State Health Services-Laboratory Services | Centers for Disease Control and Prevention |
| EPI566185 | HA | United States | 2014-Dec-18 | A/Idaho/10/2014 | State of Idaho Bureau of Laboratories | Centers for Disease Control and Prevention |
| EPI566193 | HA | United States | 2014-Dec-05 | A/Texas/88/2014 | Texas Department of State Health Services-Laboratory Services | Centers for Disease Control and Prevention |
| EPI566194 | HA | Myanmar | 2014-Jul-04 | A/Myanmar/13M006/2013 | | Niigata University (DPH) |
| EPI566202 | HA | Myanmar | 2013-Jul-09 | A/Myanmar/13M011/2013 | | Niigata University (DPH) |
| EPI566210 | HA | Myanmar | 2013-Jul-11 | A/Myanmar/13M020/2013 | | Niigata University (DPH) |
| EPI566218 | HA | Myanmar | 2013-Jul-15 | A/Myanmar/13M035/2013 | | Niigata University (DPH) |

| | | | | | | | |
|---|---|---|---|---|---|---|---|
| EPI566325 | HA | Sweden | 2015-Jan-02 | A/Sweden/1/2015 | | Public Health Agency of Sweden | |
| EPI566340 | HA | Sweden | 2015-Jan-14 | A/Sweden/4/2015 | | Public Health Agency of Sweden | |
| EPI566343 | HA | Sweden | 2015-Jan-12 | A/Gavle/1/2015 | | Public Health Agency of Sweden | |
| EPI566352 | HA | Sweden | 2015-Jan-12 | A/Lund/1/2015 | | Public Health Agency of Sweden | |
| EPI566383 | HA | Sweden | 2015-Jan-13 | A/Stockholm/7/2015 | | Public Health Agency of Sweden | |
| EPI566386 | HA | Sweden | 2015-Jan-25 | A/Stockholm/8/2015 | | Public Health Agency of Sweden | |
| EPI566389 | HA | Sweden | 2015-Jan-27 | A/Stockholm/9/2015 | | Public Health Agency of Sweden | |
| EPI566439 | HA | Sweden | 2014-Dec-12 | A/Stockholm/39/2014 | | Public Health Agency of Sweden | |
| EPI566527 | HA | Myanmar | 2013-Jul-15 | A/Myanmar/13M044/2013 | | Niigata University (DPH) | |
| EPI566535 | HA | Myanmar | 2013-Jul-16 | A/Myanmar/13M061/2013 | | Niigata University (DPH) | |
| EPI566543 | HA | Myanmar | 2013-Jul-17 | A/Myanmar/13M062/2013 | | Niigata University (DPH) | |
| EPI566615 | HA | Myanmar | 2013-Jul-17 | A/Myanmar/13M070/2013 | | Niigata University (DPH) | |
| EPI566623 | HA | Myanmar | 2013-Jul-19 | A/Myanmar/13M076/2013 | | Niigata University (DPH) | |
| EPI566631 | HA | Myanmar | 2013-Jul-19 | A/Myanmar/13M084/2013 | | Niigata University (DPH) | |
| EPI566639 | HA | Myanmar | 2013-Jul-22 | A/Myanmar/13M085/2013 | | Niigata University (DPH) | |
| EPI566673 | HA | Myanmar | 2013-Jul-22 | A/Myanmar/13M087/2013 | | Niigata University (DPH) | |
| EPI566681 | HA | Myanmar | 2013-Jul-22 | A/Myanmar/13M088/2013 | | Niigata University (DPH) | |
| EPI566707 | HA | Myanmar | 2013-Jul-22 | A/Myanmar/13M089/2013 | | Niigata University (DPH) | |
| EPI566715 | HA | Myanmar | 2013-Jul-24 | A/Myanmar/13M098/2013 | | Niigata University (DPH) | |
| EPI566723 | HA | Myanmar | 2013-Jul-24 | A/Myanmar/13M102/2013 | | Niigata University (DPH) | |
| EPI566731 | HA | Myanmar | 2013-Jul-25 | A/Myanmar/13M105/2013 | | Niigata University (DPH) | |
| EPI566739 | HA | Myanmar | 2013-Jul-25 | A/Myanmar/13M109/2013 | | Niigata University (DPH) | |
| EPI566747 | HA | Myanmar | 2013-Jul-26 | A/Myanmar/13M118/2013 | | Niigata University (DPH) | |
| EPI566755 | HA | Myanmar | 2013-Jul-16 | A/Myanmar/13M124/2013 | | Niigata University (DPH) | |
| EPI566763 | HA | Myanmar | 2013-Aug-22 | A/Myanmar/13M300/2013 | | Niigata University (DPH) | |
| EPI566863 | HA | Japan | 2014-Dec-01 | A/MIYAGI/52/2014 | Miyagi Prefectural Institute of Public Health and Environment | National Institute of Infectious Diseases (NIID) | |
| EPI566871 | HA | Japan | 2014-Dec-07 | A/FUKUI/61/2014 | Fukui Prefectural Institute of Public Health | National Institute of Infectious Diseases (NIID) | |
| EPI566901 | HA | Japan | 2015-Jan-11 | A/NIIGATA/35/2015 | Niigata Prefectural Institute of Public Health and Environmental Sciences | National Institute of Infectious Diseases (NIID) | |
| EPI566924 | HA | Japan | 2014-Mar-29 | A/Nagasaki/13N100/2014 | | Niigata University (DPH) | |
| EPI566932 | HA | Japan | 2014-Jul-29 | A/Nagasaki/13N101/2014 | | Niigata University (DPH) | |
| EPI566940 | HA | Japan | 2014-Sep-22 | A/Nagasaki/14N005/2014 | | Niigata University (DPH) | |
| EPI567319 | HA | United States | 2015-Jan-07 | A/Texas/04/2015 | Baylor College of Medicine | Centers for Disease Control and Prevention | Saito, Reiko; Takemae, Nobuhiro; Saito, Takehiko; Shobugawa, Yugo; Kondo, Hiroki; Hibino, Akinobu; Masaki, Hironori |
| EPI567327 | HA | United States | 2015-Jan-06 | A/Alabama/01/2015 | ADPH Bureau of Clinical Laboratories | Centers for Disease Control and Prevention | Saito, Reiko; Takemae, Nobuhiro; Saito, Takehiko; Shobugawa, Yugo; Kondo, Hiroki; Hibino, Akinobu; Kawashima, Takashi |

| | | | | | | | |
|---|---|---|---|---|---|---|---|
| EPI567335 | HA | United States | 2014-Dec-22 | A/Michigan/31/2014 | University of Michigan SPH EPID | Centers for Disease Control and Prevention | Saito, Reiko; Takemae, Nobuhiro; Saito, Takehiko; Shobugawa, Yugo; Kondo, Hiroki; Hibino, Akinobu; Kawashima, Takashi |
| EPI567343 | HA | United States | 2014-Dec-16 | A/Michigan/30/2014 | University of Michigan SPH EPID | Centers for Disease Control and Prevention | Saito, Reiko; Takemae, Nobuhiro; Saito, Takehiko; Shobugawa, Yugo; Kondo, Hiroki; Hibino, Akinobu; Kawashima, Takashi |
| EPI567351 | HA | United States | 2015-Jan-07 | A/North Carolina/06/2015 | North Carolina State Laboratory of Public Health | Centers for Disease Control and Prevention | Saito, Reiko; Takemae, Nobuhiro; Saito, Takehiko; Shobugawa, Yugo; Kondo, Hiroki; Hibino, Akinobu; Kawashima, Takashi |
| EPI567359 | HA | United States | 2014-Dec-29 | A/Michigan/28/2014 | University of Michigan SPH EPID | Centers for Disease Control and Prevention | Saito, Reiko; Takemae, Nobuhiro; Saito, Takehiko; Shobugawa, Yugo; Kondo, Hiroki; Hibino, Akinobu; Kodo, Naoki |
| EPI567367 | HA | United States | 2015-Jan-13 | A/North Carolina/04/2015 | North Carolina State Laboratory of Public Health | Centers for Disease Control and Prevention | Saito, Reiko; Takemae, Nobuhiro; Saito, Takehiko; Shobugawa, Yugo; Kondo, Hiroki; Hibino, Akinobu; Kodo, Naoki |
| EPI567375 | HA | United States | 2015-Jan-05 | A/Mississippi/02/2015 | Mississippi Public Health Laboratory | Centers for Disease Control and Prevention | Saito, Reiko; Takemae, Nobuhiro; Saito, Takehiko; Shobugawa, Yugo; Kondo, Hiroki; Hibino, Akinobu; Kodo, Naoki |
| EPI567383 | HA | United States | 2015-Jan-02 | A/Michigan/03/2015 | University of Michigan SPH EPID | Centers for Disease Control and Prevention | Saito, Reiko; Takemae, Nobuhiro; Saito, Takehiko; Shobugawa, Yugo; Kondo, Hiroki; Hibino, Akinobu; Sato, Isamu |
| EPI567391 | HA | United States | 2014-Dec-22 | A/Florida/92/2014 | Florida Department of Health-Tampa | Centers for Disease Control and Prevention | Saito, Reiko; Takemae, Nobuhiro; Saito, Takehiko; Shobugawa, Yugo; Kondo, Hiroki; Hibino, Akinobu; Sato, Isamu |
| EPI567402 | HA | United States | 2015-Jan-02 | A/Washington/02/2015 | Marshfield Clinic Research Foundation | Centers for Disease Control and Prevention | Saito, Reiko; Takemae, Nobuhiro; Saito, Takehiko; Shobugawa, Yugo; Kondo, Hiroki; Hibino, Akinobu; Sato, Isamu |
| EPI567410 | HA | United States | 2015-Jan-08 | A/North Carolina/05/2015 | North Carolina State Laboratory of Public Health | Centers for Disease Control and Prevention | Takashita,Emi; Fujisaki,Seiichiro; Shirakura,Masayuki; Watanabe,Shinji; Odagiri,Takato |
| EPI567424 | HA | United States | 2014-Dec-22 | A/Kentucky/24/2014 | Kentucky Division of Laboratory Services | Centers for Disease Control and Prevention | Takashita,Emi; Fujisaki,Seiichiro; Shirakura,Masayuki; Watanabe,Shinji; Odagiri,Takato |
| EPI567432 | HA | United States | 2015-Jan-04 | A/Texas/05/2015 | Baylor College of Medicine | Centers for Disease Control and Prevention | |
| EPI567440 | HA | United States | 2015-Jan-09 | A/Texas/06/2015 | Baylor College of Medicine | Centers for Disease Control and Prevention | |
| EPI567448 | HA | United States | 2014-Dec-23 | A/Kentucky/27/2014 | Kentucky Division of Laboratory Services | Centers for Disease Control and Prevention | |
| EPI567456 | HA | United States | 2014-Dec-26 | A/Michigan/32/2014 | University of Michigan SPH EPID | Centers for Disease Control and Prevention | |

| | | | | | | |
|---|---|---|---|---|---|---|
| EPI567462 | HA | United States | 2015-Jan-01 | A/North Dakota/02/2015 | North Dakota Department of Health | Centers for Disease Control and Prevention |
| EPI567468 | HA | United States | 2014-Dec-12 | A/Michigan/27/2014 | University of Michigan SPH EPID | Centers for Disease Control and Prevention |
| EPI567476 | HA | United States | 2014-Dec-29 | A/Washington/57/2014 | Marshfield Clinic Research Foundation | Centers for Disease Control and Prevention |
| EPI567484 | HA | United States | 2015-Jan-18 | A/Florida/14/2015 | Florida Department of Health-Jacksonville | Centers for Disease Control and Prevention |
| EPI567496 | HA | United States | 2015-Jan-02 | A/Wisconsin/03/2015 | Marshfield Clinic Research Foundation | Centers for Disease Control and Prevention |
| EPI567504 | HA | United States | 2015-Jan-21 | A/Washington/01/2015 | Marshfield Clinic Research Foundation | Centers for Disease Control and Prevention |
| EPI567512 | HA | United States | 2015-Jan-05 | A/Michigan/02/2015 | University of Michigan SPH EPID | Centers for Disease Control and Prevention |
| EPI567539 | HA | United States | 2015-Jan-11 | A/Tennessee/01/2015 | Tennessee Department of Health Laboratory-Nashville | Centers for Disease Control and Prevention |
| EPI567554 | HA | United States | 2015-Jan-28 | A/North Carolina/02/2015 | North Carolina State Laboratory of Public Health | Centers for Disease Control and Prevention |
| EPI567562 | HA | United States | 2014-Dec-15 | A/Texas/107/2014 | Baylor College of Medicine | Centers for Disease Control and Prevention |
| EPI567570 | HA | United States | 2014-Dec-20 | A/Texas/106/2014 | Baylor College of Medicine | Centers for Disease Control and Prevention |
| EPI567583 | HA | United States | 2014-Dec-26 | A/Texas/105/2014 | Baylor College of Medicine | Centers for Disease Control and Prevention |
| EPI567591 | HA | United States | 2014-Dec-22 | A/Kentucky/25/2014 | Kentucky Division of Laboratory Services | Centers for Disease Control and Prevention |
| EPI567604 | HA | United States | 2014-Dec-19 | A/Wisconsin/72/2014 | Marshfield Clinic Research Foundation | Centers for Disease Control and Prevention |
| EPI567612 | HA | United States | 2015-Jan-04 | A/Mississippi/01/2015 | Mississippi Public Health Laboratory | Centers for Disease Control and Prevention |
| EPI567620 | HA | United States | 2014-Dec-26 | A/Washington/53/2014 | Marshfield Clinic Research Foundation | Centers for Disease Control and Prevention |
| EPI567628 | HA | United States | 2014-Dec-22 | A/Wisconsin/75/2014 | Marshfield Clinic Research Foundation | Centers for Disease Control and Prevention |
| EPI567788 | HA | Australia | 1982-Jan-01 | A/Victoria/186/1982 | | WHO Collaborating Centre for Reference and Research on Influenza |
| EPI567796 | HA | Australia | 1993-Jan-01 | A/Sydney/2/1993 | Clinical Virology Unit, CDIM | WHO Collaborating Centre for Reference and Research on Influenza |
| EPI567881 | HA | Vietnam | 2012-Dec-28 | A/Vietnam/13V H3-2/2012 | | Niigata University (DPH) |
| EPI567889 | HA | Vietnam | 2012-Jul-23 | A/Vietnam/13V H3-3/2012 | | Niigata University (DPH) |
| EPI567910 | HA | Vietnam | 2013-Apr-13 | A/Vietnam/13V H3-4/2013 | | Niigata University (DPH) |
| EPI567926 | HA | Vietnam | 2013-Sep-25 | A/Vietnam/13V H3-8/2013 | | Niigata University (DPH) |
| EPI567927 | HA | Vietnam | 2013-Sep-25 | A/Vietnam/13V H3-8/2013 | | Niigata University (DPH) |
| EPI567935 | HA | Vietnam | 2012-Jul-27 | A/Vietnam/13V H3-9/2012 | | Niigata University (DPH) |
| EPI567943 | HA | Vietnam | 2012-Jul-25 | A/Vietnam/13V H3-10/2012 | | Niigata University (DPH) |
| EPI568050 | HA | Myanmar | 2014-Jun-24 | A/Myanmar/14M039/2014 | | Niigata University (DPH) |

| | | | | | | | |
|---|---|---|---|---|---|---|---|
| EPI568058 | HA | Myanmar | 2014-Jul-14 | A/Myanmar/14M059/2014 | | Niigata University (DPH) | |
| EPI568066 | HA | Myanmar | 2014-Jul-08 | A/Myanmar/14M122/2014 | | Niigata University (DPH) | |
| EPI568074 | HA | Myanmar | 2014-Jul-02 | A/Myanmar/14M187/2014 | | Niigata University (DPH) | |
| EPI568082 | HA | Myanmar | 2014-Aug-04 | A/Myanmar/14M240/2014 | | Niigata University (DPH) | |
| EPI568090 | HA | Myanmar | 2014-Aug-22 | A/Myanmar/14M391/2014 | | Niigata University (DPH) | |
| EPI568098 | HA | Myanmar | 2014-Sep-08 | A/Myanmar/14M465/2014 | | Niigata University (DPH) | |
| EPI569276 | HA | New Caledonia | 2014-Aug-13 | A/New Caledonia/71/2014 | WHO Collaborating Centre for Reference and Research on Influenza | Centers for Disease Control and Prevention | |
| EPI569302 | HA | United States | 2014-Nov-07 | A/Utah/19/2014 | Utah Department of Health | Centers for Disease Control and Prevention | |
| EPI569312 | HA | United States | 2014-Nov-19 | A/Louisiana/21/2014 | Louisiana Department of Health and Hospitals | Centers for Disease Control and Prevention | |
| EPI570014 | HA | Sweden | 2015-Feb-01 | A/Sweden/8/2015 | | Public Health Agency of Sweden | |
| EPI570118 | HA | Sweden | 2015-Feb-15 | A/Sweden/11/2015 | | Public Health Agency of Sweden | |
| EPI570137 | HA | Sweden | 2015-Feb-15 | A/Sweden/12/2015 | | Public Health Agency of Sweden | |
| EPI571782 | HA | United States | 2015-Jan-06 | A/Rhode Island/03/2015 | Rhode Island Department of Health | Centers for Disease Control and Prevention | |
| EPI571803 | HA | Tanzania, United Republic of | 2014-Nov-20 | A/Tanzania/3565/2014 | National Influenza Lab | Centers for Disease Control and Prevention | |
| EPI571811 | HA | Tanzania, United Republic of | 2014-Nov-24 | A/Tanzania/3576/2014 | National Influenza Lab | Centers for Disease Control and Prevention | |
| EPI571819 | HA | Tanzania, United Republic of | 2014-Nov-24 | A/Tanzania/3577/2014 | National Influenza Lab | Centers for Disease Control and Prevention | |
| EPI571827 | HA | United States | 2015-Jan-08 | A/New York/09/2015 | New York City Department of Health | Centers for Disease Control and Prevention | |
| EPI571854 | HA | United States | 2015-Jan-05 | A/Rhode Island/02/2015 | Rhode Island Department of Health | Centers for Disease Control and Prevention | |
| EPI571879 | HA | United States | 2015-Jan-03 | A/Oklahoma/02/2015 | Oklahoma State Department of Health | Centers for Disease Control and Prevention | Saito, Reiko; Takemae, Nobuhiro; Saito, Takehiko; Shobugawa, Yugo; Kondo, Hiroki; Hibino, Akinobu; Yadanar, Kyaw; Yi Yi, Myint; Khin Yi, Oo; Htay Htay, Tin |
| EPI571887 | HA | Tanzania, United Republic of | 2014-Nov-19 | A/Tanzania/3560/2014 | National Influenza Lab | Centers for Disease Control and Prevention | Saito, Reiko; Takemae, Nobuhiro; Saito, Takehiko; Shobugawa, Yugo; Kondo, Hiroki; Hibino, Akinobu; Yadanar, Kyaw; Yi Yi, Myint; Khin Yi, Oo; Htay Htay, Tin |
| EPI571902 | HA | United States | 2015-Jan-06 | A/North Carolina/08/2015 | North Carolina State Laboratory of Public Health | Centers for Disease Control and Prevention | Saito, Reiko; Takemae, Nobuhiro; Saito, Takehiko; Shobugawa, Yugo; Kondo, Hiroki; Hibino, |
| EPI571915 | HA | United States | 2015-Jan-02 | A/Oregon/02/2015 | Oregon Public Health Laboratory | Centers for Disease Control and Prevention | Saito, Reiko; Takemae, Nobuhiro; Saito, Takehiko; Shobugawa, Yugo; Kondo, Hiroki; Hibino, |
| EPI571930 | HA | United States | 2015-Jan-12 | A/Utah/03/2015 | Utah Department of Health | Centers for Disease Control and Prevention | Saito, Reiko; Takemae, Nobuhiro; Saito, Takehiko; Shobugawa, Yugo; Kondo, Hiroki; Hibino, |

| | | | | | | | |
|---|---|---|---|---|---|---|---|
| EPI571945 | HA | United States | 2015-Jan-05 | A/Nebraska/01/2015 | Nebraska Public Health Lab | Centers for Disease Control and Prevention | Deng,Y-M.; Iannello,P.; Spirason,N; Jelley,L.;Lau,H.; |
| EPI571960 | HA | United States | 2015-Jan-07 | A/Nevada/04/2015 | Southern Nevada Public Health Lab | Centers for Disease Control and Prevention | Deng,Y-M.; Iannello,P.; Spirason,N; Jelley,L.;Lau,H. |
| EPI571975 | HA | United States | 2015-Jan-22 | A/New Hampshire/05/2015 | New Hampshire Public Health Laboratories | Centers for Disease Control and Prevention | |
| EPI572009 | HA | United States | 2015-Jan-01 | A/North Dakota/03/2015 | North Dakota Department of Health | Centers for Disease Control and Prevention | |
| EPI572017 | HA | United States | 2015-Jan-18 | A/Montana/04/2015 | Montana Laboratory Services Bureau | Centers for Disease Control and Prevention | |
| EPI572052 | HA | United States | 2015-Jan-13 | A/New Hampshire/04/2015 | New Hampshire Public Health Laboratories | Centers for Disease Control and Prevention | |
| EPI572060 | HA | United States | 2015-Jan-22 | A/New York/04/2015 | New York City Department of Health | Centers for Disease Control and Prevention | |
| EPI572071 | HA | United States | 2015-Jan-20 | A/Massachusetts/02/2015 | Massachusetts Department of Public Health | Centers for Disease Control and Prevention | |
| EPI572105 | HA | United States | 2015-Jan-15 | A/Missouri/04/2015 | Missouri Department. of Health & Senior Services | Centers for Disease Control and Prevention | |
| EPI572119 | HA | United States | 2015-Jan-10 | A/Arkansas/02/2015 | Arkansas Department of Health | Centers for Disease Control and Prevention | |
| EPI572134 | HA | United States | 2015-Jan-06 | A/Michigan/04/2015 | Michigan Department of Community Health | Centers for Disease Control and Prevention | |
| EPI572199 | HA | United States | 2015-Jan-04 | A/California/01/2015 | California Department of Health Services | Centers for Disease Control and Prevention | |
| EPI572227 | HA | United States | 2015-Jan-03 | A/Missouri/01/2015 | Missouri Department. of Health & Senior Services | Centers for Disease Control and Prevention | |
| EPI572249 | HA | United States | 2015-Jan-14 | A/North Carolina/07/2015 | North Carolina State Laboratory of Public Health | Centers for Disease Control and Prevention | |
| EPI574877 | HA | United States | 2015-Jan-09 | A/Colorado/02/2015 | Colorado Department of Health Lab | Centers for Disease Control and Prevention | |
| EPI574885 | HA | United States | 2015-Feb-02 | A/Tennessee/09/2015 | Tennessee Department of Health Laboratory-Nashville | Centers for Disease Control and Prevention | Takashita,Emi; Fujisaki,Seiichiro; Shirakura,Masayuki; Watanabe,Shinji; Odagiri,Takato |
| EPI574894 | HA | United States | 2015-Jan-17 | A/Michigan/07/2015 | Michigan Department of Community Health | Centers for Disease Control and Prevention | Saito, Reiko; Takemae, Nobuhiro; Saito, Takehiko; Shobugawa, Yugo; Kondo, Hiroki; Hibino, Akinobu; Yadanar, Kyaw; Yi Yi, Myint; Khin Yi, Oo; Htay Htay, Tin |
| EPI574902 | HA | United States | 2015-Feb-02 | A/Tennessee/07/2015 | Tennessee Department of Health Laboratory-Nashville | Centers for Disease Control and Prevention | Saito, Reiko; Takemae, Nobuhiro; Saito, Takehiko; Shobugawa, Yugo; Kondo, Hiroki; Hibino, Akinobu; Yadanar, Kyaw; Yi Yi, Myint; Khin Yi, Oo; Htay Htay, Tin |
| EPI574910 | HA | United States | 2015-Feb-03 | A/Tennessee/08/2015 | Tennessee Department of Health Laboratory-Nashville | Centers for Disease Control and Prevention | Saito, Reiko; Takemae, Nobuhiro; Saito, Takehiko; Shobugawa, Yugo; Kondo, Hiroki; Hibino, Akinobu; Yadanar, Kyaw; Yi Yi, Myint; Khin Yi, Oo; Htay Htay, Tin |

| | | | | | | | |
|---|---|---|---|---|---|---|---|
| EPI574921 | HA | United States | 2015-Jan-08 | A/Michigan/05/2015 | Michigan Department of Community Health | Centers for Disease Control and Prevention | Saito, Reiko; Takemae, Nobuhiro; Saito, Takehiko; Shobugawa, Yugo; Kondo, Hiroki; Hibino, Akinobu; Yadanar, Kyaw; Yi Yi, Myint; Khin Yi, Oo; Htay Htay, Tin |
| EPI574944 | HA | United States | 2015-Jan-18 | A/Kansas/03/2015 | Kansas Department of Health and Environment | Centers for Disease Control and Prevention | Saito, Reiko; Takemae, Nobuhiro; Saito, Takehiko; Shobugawa, Yugo; Kondo, Hiroki; Hibino, Akinobu; Yadanar, Kyaw; Yi Yi, Myint; Khin Yi, Oo; Htay Htay, Tin |
| EPI574963 | HA | United States | 2015-Jan-08 | A/Indiana/01/2015 | Indiana State Department of Health Laboratories | Centers for Disease Control and Prevention | Saito, Reiko; Takemae, Nobuhiro; Saito, Takehiko; Shobugawa, Yugo; Kondo, Hiroki; Hibino, Akinobu; Yadanar, Kyaw; Yi Yi, Myint; Khin Yi, Oo; Htay Htay, Tin |
| EPI574971 | HA | United States | 2015-Jan-12 | A/Utah/05/2015 | Utah Department of Health | Centers for Disease Control and Prevention | |
| EPI574992 | HA | United States | 2015-Jan-05 | A/Colorado/04/2015 | Colorado Department of Health Lab | Centers for Disease Control and Prevention | |
| EPI575017 | HA | United States | 2015-Jan-06 | A/Rhode Island/06/2015 | Rhode Island Department of Health | Centers for Disease Control and Prevention | Saito, Reiko; Takemae, Nobuhiro; Saito, Takehiko; Shobugawa, Yugo; Kondo, Hiroki; Hibino, Akinobu; Yadanar, Kyaw; Yi Yi, Myint; Khin Yi, Oo; Htay Htay, Tin |
| EPI575040 | HA | United States | 2015-Jan-19 | A/New Jersey/03/2015 | New Jersey Department of Health & Senior Services | Centers for Disease Control and Prevention | |
| EPI575060 | HA | United States | 2015-Jan-07 | A/New York/07/2015 | New York City Department of Health | Centers for Disease Control and Prevention | |
| EPI575079 | HA | United States | 2015-Feb-09 | A/North Carolina/11/2015 | North Carolina State Laboratory of Public Health | Centers for Disease Control and Prevention | |
| EPI575087 | HA | United States | 2015-Feb-06 | A/Ohio/04/2015 | Ohio Department of Health Laboratories | Centers for Disease Control and Prevention | |
| EPI575095 | HA | United States | 2015-Feb-01 | A/Florida/19/2015 | Florida Department of Health-Jacksonville | Centers for Disease Control and Prevention | |
| EPI575108 | HA | United States | 2014-Dec-25 | A/Iowa/31/2014 | Iowa State Hygienic Laboratory | Centers for Disease Control and Prevention | |
| EPI575122 | HA | United States | 2015-Jan-06 | A/Iowa/04/2015 | Iowa State Hygienic Laboratory | Centers for Disease Control and Prevention | |
| EPI575158 | HA | Cambodia | 2014-Dec-17 | A/Cambodia/1384/2014 | National Public Health Laboratory | Centers for Disease Control and Prevention | |
| EPI575166 | HA | United States | 2015-Jan-03 | A/New Mexico/01/2015 | New Mexico Department of Health | Centers for Disease Control and Prevention | |
| EPI575174 | HA | United States | 2015-Jan-30 | A/Florida/20/2015 | Florida Department of Health-Jacksonville | Centers for Disease Control and Prevention | |
| EPI575183 | HA | United States | 2015-Feb-03 | A/Hawaii/02/2015 | State of Hawaii Department of Health | Centers for Disease Control and Prevention | |
| EPI575191 | HA | United States | 2015-Feb-02 | A/Florida/18/2015 | Florida Department of Health-Jacksonville | Centers for Disease Control and Prevention | |
| EPI575199 | HA | United States | 2015-Jan-29 | A/Utah/06/2015 | Utah Department of Health | Centers for Disease Control and Prevention | |

| | | | | | | | |
|---|---|---|---|---|---|---|---|
| EPI575212 | HA | United States | 2015-Feb-10 | A/Utah/07/2015 | Utah Department of Health | Centers for Disease Control and Prevention | |
| EPI575240 | HA | United States | 2015-Jan-30 | A/Delaware/05/2015 | Delaware Public Health Lab | Centers for Disease Control and Prevention | |
| EPI575248 | HA | United States | 2015-Feb-05 | A/New York/15/2015 | New York City Department of Health | Centers for Disease Control and Prevention | |
| EPI575256 | HA | United States | 2015-Jan-29 | A/New Hampshire/06/2015 | New Hampshire Public Health Laboratories | Centers for Disease Control and Prevention | |
| EPI575264 | HA | United States | 2014-Dec-02 | A/Tennessee/24/2014 | St. Judes Childrens Research Hospital | Centers for Disease Control and Prevention | |
| EPI575271 | HA | United States | 2015-Jan-28 | A/New Hampshire/07/2015 | New Hampshire Public Health Laboratories | Centers for Disease Control and Prevention | |
| EPI575279 | HA | United States | 2015-Jan-08 | A/Nevada/03/2015 | Nevada State Health Laboratory | Centers for Disease Control and Prevention | |
| EPI575287 | HA | United States | 2015-Jan-14 | A/Nevada/02/2015 | Nevada State Health Laboratory | Centers for Disease Control and Prevention | |
| EPI575295 | HA | United States | 2014-Dec-16 | A/Texas/93/2014 | Texas Childrens Hospital | Centers for Disease Control and Prevention | |
| EPI575303 | HA | United States | 2014-Dec-26 | A/Texas/96/2014 | Texas Childrens Hospital | Centers for Disease Control and Prevention | |
| EPI575310 | HA | United States | 2015-Feb-05 | A/Nevada/09/2015 | Nevada State Health Laboratory | Centers for Disease Control and Prevention | |
| EPI575318 | HA | United States | 2015-Feb-14 | A/Nevada/10/2015 | Nevada State Health Laboratory | Centers for Disease Control and Prevention | |
| EPI575326 | HA | United States | 2015-Jan-27 | A/Wisconsin/05/2015 | Wisconsin State Laboratory of Hygiene | Centers for Disease Control and Prevention | |
| EPI575334 | HA | United States | 2015-Jan-30 | A/Kansas/02/2015 | Kansas Department of Health and Environment | Centers for Disease Control and Prevention | |
| EPI575344 | HA | Cambodia | 2014-Dec-11 | A/Cambodia/1370/2014 | National Public Health Laboratory | Centers for Disease Control and Prevention | |
| EPI575352 | HA | Cambodia | 2014-Dec-11 | A/Cambodia/1372/2014 | National Public Health Laboratory | Centers for Disease Control and Prevention | |
| EPI575370 | HA | Tanzania, United Republic of | 2014-Dec-17 | A/Tanzania/613/2014 | National Influenza Lab | Centers for Disease Control and Prevention | |
| EPI575378 | HA | United States | 2015-Jan-17 | A/Maine/07/2015 | Maine Health and Environmental Testing Laboratory | Centers for Disease Control and Prevention | |
| EPI575386 | HA | United States | 2015-Jan-09 | A/Nebraska/03/2015 | Nebraska Public Health Lab | Centers for Disease Control and Prevention | |
| EPI575400 | HA | Russian Federation | 2014-Dec-11 | A/Chita/256/2014 | Russian Academy of Medical Sciences | Centers for Disease Control and Prevention | |
| EPI575408 | HA | Tanzania, United Republic of | 2014-Dec-11 | A/Tanzania/3595/2014 | National Influenza Lab | Centers for Disease Control and Prevention | |
| EPI575416 | HA | Russian Federation | 2014-Dec-12 | A/Chita/260/2014 | Russian Academy of Medical Sciences | Centers for Disease Control and Prevention | |
| EPI575427 | HA | Russian Federation | 2014-Dec-15 | A/Chita/258/2014 | Russian Academy of Medical Sciences | Centers for Disease Control and Prevention | Saito, Reiko; Takemae, Nobuhiro; Saito, Takehiko; Shobugawa, Yugo; Kondo, Hiroki; Hibino, Akinobu; Yadanar, Kyaw; Yi Yi, Myint; Khin Yi, Oo; Htay Htay, Tin |

| | | | | | | | |
|---|---|---|---|---|---|---|---|
| EPI575435 | HA | Tanzania, United Republic of | 2014-Dec-08 | A/Tanzania/3591/2014 | National Influenza Lab | Centers for Disease Control and Prevention | Saito, Reiko; Takemae, Nobuhiro; Saito, Takehiko; Shobugawa, Yugo; Kondo, Hiroki; Hibino, Akinobu; Yadanar, Kyaw; Yi Yi, Myint; Khin Yi, Oo; Htay Htay, Tin |
| EPI575443 | HA | United States | 2015-Jan-22 | A/New York/12/2015 | New York City Department of Health | Centers for Disease Control and Prevention | Saito, Reiko; Takemae, Nobuhiro; Saito, Takehiko; Shobugawa, Yugo; Kondo, Hiroki; Hibino, Akinobu; Yadanar, Kyaw; Yi Yi, Myint; Khin Yi, Oo; Htay Htay, Tin |
| EPI575451 | HA | United States | 2015-Jan-20 | A/Massachusetts/03/2015 | Massachusetts Department of Public Health | Centers for Disease Control and Prevention | |
| EPI575462 | HA | Russian Federation | 2014-Dec-12 | A/Chita/264/2014 | Russian Academy of Medical Sciences | Centers for Disease Control and Prevention | |
| EPI575470 | HA | Russian Federation | 2014-Dec-10 | A/Chita/268/2014 | Russian Academy of Medical Sciences | Centers for Disease Control and Prevention | |
| EPI575478 | HA | Congo, the Democatic Republic of | 2014-Dec-09 | A/Congo/2471/2014 | INRB Service de Virologie | Centers for Disease Control and Prevention | |
| EPI575486 | HA | Congo, the Democatic Republic of | 2014-Dec-04 | A/Congo/2458/2014 | INRB Service de Virologie | Centers for Disease Control and Prevention | |
| EPI575501 | HA | United States | 2015-Jan-30 | A/Washington/04/2015 | Marshfield Clinic Research Foundation | Centers for Disease Control and Prevention | |
| EPI575509 | HA | United States | 2015-Feb-01 | A/Kansas/01/2015 | Kansas Department of Health and Environment | Centers for Disease Control and Prevention | |
| EPI575517 | HA | United States | 2014-Dec-08 | A/Georgia/30/2014 | Georgia Public Health Laboratory | Centers for Disease Control and Prevention | |
| EPI575525 | HA | United States | 2015-Feb-06 | A/Pennsylvania/05/2015 | Pennsylvania Department of Health | Centers for Disease Control and Prevention | |
| EPI575533 | HA | Congo, the Democatic Republic of | 2014-Dec-08 | A/Congo/2461/2014 | INRB Service de Virologie | Centers for Disease Control and Prevention | |
| EPI575541 | HA | United States | 2015-Feb-05 | A/Pennsylvania/03/2015 | Pennsylvania Department of Health | Centers for Disease Control and Prevention | |
| EPI575549 | HA | United States | 2015-Feb-09 | A/New York/16/2015 | New York City Department of Health | Centers for Disease Control and Prevention | |
| EPI575557 | HA | United States | 2015-Jan-26 | A/New Jersey/02/2015 | New Jersey Department of Health & Senior Services | Centers for Disease Control and Prevention | |
| EPI575564 | HA | Russian Federation | 2014-Dec-03 | A/Chita/266/2014 | Russian Academy of Medical Sciences | Centers for Disease Control and Prevention | |
| EPI575572 | HA | Russian Federation | 2014-Dec-12 | A/Tomsk/254/2014 | Russian Academy of Medical Sciences | Centers for Disease Control and Prevention | |
| EPI575580 | HA | Russian Federation | 2014-Dec-16 | A/Astrakhan/24/2014 | Russian Academy of Medical Sciences | Centers for Disease Control and Prevention | |
| EPI575588 | HA | United States | 2015-Feb-03 | A/Texas/09/2015 | Texas Department of State Health Services-Laboratory Services | Centers for Disease Control and Prevention | |
| EPI575597 | HA | United States | 2015-Feb-05 | A/New York/14/2015 | New York City Department of Health | Centers for Disease Control and Prevention | |
| EPI575605 | HA | United States | 2015-Jan-31 | A/Nebraska/05/2015 | Nebraska Public Health Lab | Centers for Disease Control and Prevention | |

| | | | | | | | |
|---|---|---|---|---|---|---|---|
| EPI575614 | HA | United States | 2015-Jan-28 | A/West Virginia/02/2015 | West Virginia Office of Laboratory Services | Centers for Disease Control and Prevention | |
| EPI575621 | HA | United States | 2015-Jan-28 | A/Delaware/06/2015 | Delaware Public Health Lab | Centers for Disease Control and Prevention | |
| EPI575632 | HA | United States | 2015-Feb-10 | A/New York/13/2015 | New York City Department of Health | Centers for Disease Control and Prevention | |
| EPI575649 | HA | United States | 2014-Dec-01 | A/Georgia/24/2014 | U.S. Air Force School of Aerospace Medicine | Centers for Disease Control and Prevention | |
| EPI575661 | HA | Kazakhstan | 2014-Dec-29 | A/Kazakhstan/15/2014 | CSEE | Centers for Disease Control and Prevention | |
| EPI575669 | HA | Kazakhstan | 2015-Jan-02 | A/Kazakhstan/22/2015 | CSEE | Centers for Disease Control and Prevention | |
| EPI575677 | HA | Kazakhstan | 2015-Jan-06 | A/Kazakhstan/29/2015 | CSEE | Centers for Disease Control and Prevention | |
| EPI575685 | HA | Kazakhstan | 2015-Jan-05 | A/Kazakhstan/27/2015 | CSEE | Centers for Disease Control and Prevention | |
| EPI575693 | HA | Kazakhstan | 2015-Jan-06 | A/Kazakhstan/30/2015 | CSEE | Centers for Disease Control and Prevention | |
| EPI575701 | HA | Kazakhstan | 2015-Jan-06 | A/Kazakhstan/33/2015 | CSEE | Centers for Disease Control and Prevention | |
| EPI575709 | HA | Kazakhstan | 2015-Jan-05 | A/Kazakhstan/20/2015 | CSEE | Centers for Disease Control and Prevention | |
| EPI575717 | HA | Kazakhstan | 2015-Jan-01 | A/Kazakhstan/16/2015 | CSEE | Centers for Disease Control and Prevention | |
| EPI575725 | HA | Kazakhstan | 2015-Jan-07 | A/Kazakhstan/35/2015 | CSEE | Centers for Disease Control and Prevention | Takashita,Emi; Fujisaki,Seiichiro; Shirakura,Masayuki; Watanabe,Shinji; Odagiri,Takato |
| EPI575733 | HA | Kazakhstan | 2015-Jan-06 | A/Kazakhstan/31/2015 | CSEE | Centers for Disease Control and Prevention | Takashita,Emi; Fujisaki,Seiichiro; Shirakura,Masayuki; Watanabe,Shinji; Odagiri,Takato |
| EPI575741 | HA | Kazakhstan | 2015-Jan-05 | A/Kazakhstan/25/2015 | CSEE | Centers for Disease Control and Prevention | |
| EPI575749 | HA | Kazakhstan | 2015-Jan-13 | A/Kazakhstan/32/2015 | CSEE | Centers for Disease Control and Prevention | |
| EPI575757 | HA | Kazakhstan | 2015-Jan-13 | A/Kazakhstan/42/2015 | CSEE | Centers for Disease Control and Prevention | |
| EPI575765 | HA | Kazakhstan | 2015-Jan-13 | A/Kazakhstan/36/2015 | CSEE | Centers for Disease Control and Prevention | |
| EPI575773 | HA | Kazakhstan | 2015-Jan-06 | A/Kazakhstan/34/2015 | CSEE | Centers for Disease Control and Prevention | |
| EPI575781 | HA | Kazakhstan | 2015-Jan-13 | A/Kazakhstan/13/2015 | CSEE | Centers for Disease Control and Prevention | |
| EPI575789 | HA | Kazakhstan | 2015-Jan-13 | A/Kazakhstan/05/2015 | CSEE | Centers for Disease Control and Prevention | |
| EPI575797 | HA | Kazakhstan | 2015-Jan-05 | A/Kazakhstan/26/2015 | CSEE | Centers for Disease Control and Prevention | |
| EPI575805 | HA | Kazakhstan | 2015-Jan-13 | A/Kazakhstan/03/2015 | CSEE | Centers for Disease Control and Prevention | |
| EPI575813 | HA | Kazakhstan | 2015-Jan-13 | A/Kazakhstan/49/2015 | CSEE | Centers for Disease Control and Prevention | |

| | | | | | | | |
|---|---|---|---|---|---|---|---|
| EPI575821 | HA | United States | 2014-Dec-16 | A/Georgia/39/2014 | Georgia Public Health Laboratory | Centers for Disease Control and Prevention | |
| EPI575840 | HA | United States | 2015-Jan-27 | A/Pennsylvania/04/2015 | University of Pittsburgh Medical Center Microbiology Lab | Centers for Disease Control and Prevention | |
| EPI576096 | HA | Japan | 2014-Dec-17 | A/WAKAYAMA/167/2014 | Wakayama Prefectural Research Center of Environment and Public Health | National Institute of Infectious Diseases (NIID) | |
| EPI576098 | HA | Japan | 2014-Dec-22 | A/TOKYO/34558/2014 | Tokyo Metropolitan Institute of Public Health | National Institute of Infectious Diseases (NIID) | |
| EPI576116 | HA | Japan | 2015-Jan-09 | A/IBARAKI/1/2015 | Ibaraki Prefectural Institute of Public Health | National Institute of Infectious Diseases (NIID) | |
| EPI576132 | HA | Japan | 2015-Jan-20 | A/TOKYO/41945/2015 | Tokyo Metropolitan Institute of Public Health | National Institute of Infectious Diseases (NIID) | Saito, Reiko; Takemae, Nobuhiro; Saito, Takehiko; Shobugawa, Yugo; Kondo, Hiroki; Hibino, Akinobu; Shirahige, Yutaka |
| EPI576136 | HA | Japan | 2015-Jan-21 | A/IWATE/7/2015 | Research Institute for Environmental Sciences and Public Health of Iwate Prefecture | National Institute of Infectious Diseases (NIID) | |
| EPI577029 | HA | Australia | 2014-Jul-31 | A/Canberra/82/2014 | WHO Collaborating Centre for Reference and Research on Influenza | Centers for Disease Control and Prevention | |
| EPI577583 | HA | Japan | 2015-Jan-13 | A/Niigata/14F002/2015 | | Niigata University (DPH) | |
| EPI577588 | HA | Japan | 2015-Jan-13 | A/Niigata/14F004/2015 | | Niigata University (DPH) | |
| EPI577596 | HA | Japan | 2015-Jan-13 | A/Niigata/14F005/2015 | | Niigata University (DPH) | |
| EPI577604 | HA | Japan | 2015-Jan-13 | A/Niigata/14F007/2015 | | Niigata University (DPH) | |
| EPI577612 | HA | Japan | 2015-Jan-13 | A/Niigata/14F009/2015 | | Niigata University (DPH) | |
| EPI577620 | HA | Japan | 2014-Dec-08 | A/Kyoto/14K001/2014 | | Niigata University (DPH) | |
| EPI577628 | HA | Japan | 2014-Dec-08 | A/Kyoto/14K002/2014 | | Niigata University (DPH) | |
| EPI577636 | HA | Japan | 2014-Dec-09 | A/Kyoto/14K003/2014 | | Niigata University (DPH) | |
| EPI577644 | HA | Japan | 2014-Dec-09 | A/Kyoto/14K005/2014 | | Niigata University (DPH) | |
| EPI577652 | HA | Japan | 2014-Dec-15 | A/Kyoto/14K007/2014 | | Niigata University (DPH) | |
| EPI577660 | HA | Japan | 2015-Jan-10 | A/Gunma/14G002/2015 | | Niigata University (DPH) | |
| EPI577668 | HA | Japan | 2015-Jan-13 | A/Gunma/14G003/2015 | | Niigata University (DPH) | |
| EPI577676 | HA | Japan | 2015-Jan-13 | A/Gunma/14G004/2015 | | Niigata University (DPH) | |
| EPI577684 | HA | Japan | 2015-Jan-13 | A/Gunma/14G005/2015 | | Niigata University (DPH) | |
| EPI577692 | HA | Japan | 2015-Jan-14 | A/Gunma/14G008/2015 | | Niigata University (DPH) | |
| EPI577700 | HA | Japan | 2014-Dec-12 | A/Nagasaki/14N014/2014 | | Niigata University (DPH) | |
| EPI577708 | HA | Japan | 2014-Dec-15 | A/Nagasaki/14N015/2014 | | Niigata University (DPH) | |
| EPI577716 | HA | Japan | 2014-Dec-24 | A/Nagasaki/14N017/2014 | | Niigata University (DPH) | |
| EPI577724 | HA | Japan | 2014-Dec-25 | A/Nagasaki/14N018/2014 | | Niigata University (DPH) | |
| EPI577732 | HA | Japan | 2014-Dec-28 | A/Nagasaki/14N023/2014 | | Niigata University (DPH) | |
| EPI577748 | HA | Japan | 2014-Nov-04 | A/Nagasaki/14N010/2014 | | Niigata University (DPH) | |
| EPI578251 | HA | United States | 2015-Jan-09 | A/Mississippi/04/2015 | Mississippi Public Health Laboratory | Centers for Disease Control and Prevention | |

| | | | | | | |
|---|---|---|---|---|---|---|
| EPI578259 | HA | United States | 2015-Jan-13 | A/Florida/22/2015 | Florida Department of Health-Tampa | Centers for Disease Control and Prevention |
| EPI578267 | HA | United States | 2015-Jan-15 | A/Mississippi/05/2015 | Mississippi Public Health Laboratory | Centers for Disease Control and Prevention |
| EPI578275 | HA | United States | 2015-Jan-12 | A/Michigan/09/2015 | Michigan Department of Community Health | Centers for Disease Control and Prevention |
| EPI578283 | HA | United States | 2015-Jan-09 | A/Pennsylvania/07/2015 | UPMC-CLB Dept of Microbiology | Centers for Disease Control and Prevention |
| EPI578298 | HA | United States | 2015-Jan-07 | A/Pennsylvania/08/2015 | UPMC-CLB Dept of Microbiology | Centers for Disease Control and Prevention |
| EPI578306 | HA | United States | 2015-Jan-10 | A/Texas/13/2015 | Baylor College of Medicine | Centers for Disease Control and Prevention |
| EPI578314 | HA | United States | 2015-Jan-09 | A/Alaska/01/2015 | Alaska State Virology Lab | Centers for Disease Control and Prevention |
| EPI578322 | HA | United States | 2015-Jan-09 | A/New Mexico/02/2015 | New Mexico Department of Health | Centers for Disease Control and Prevention |
| EPI578337 | HA | United States | 2015-Jan-10 | A/Hawaii/04/2015 | State of Hawaii Department of Health | Centers for Disease Control and Prevention |
| EPI578363 | HA | United States | 2015-Jan-07 | A/Pennsylvania/06/2015 | UPMC-CLB Dept of Microbiology | Centers for Disease Control and Prevention |
| EPI578376 | HA | United States | 2015-Jan-07 | A/Pennsylvania/09/2015 | UPMC-CLB Dept of Microbiology | Centers for Disease Control and Prevention |
| EPI578384 | HA | United States | 2015-Jan-10 | A/Texas/12/2015 | Baylor College of Medicine | Centers for Disease Control and Prevention |
| EPI578398 | HA | United States | 2015-Jan-11 | A/New Mexico/04/2015 | New Mexico Department of Health | Centers for Disease Control and Prevention |
| EPI578406 | HA | United States | 2015-Jan-08 | A/Pennsylvania/10/2015 | UPMC-CLB Dept of Microbiology | Centers for Disease Control and Prevention |
| EPI578414 | HA | United States | 2015-Jan-11 | A/New Hampshire/10/2015 | New Hampshire Public Health Laboratories | Centers for Disease Control and Prevention |
| EPI578422 | HA | United States | 2015-Jan-24 | A/Delaware/09/2015 | Delaware Public Health Lab | Centers for Disease Control and Prevention |
| EPI578430 | HA | Hong Kong (SAR) | 2014-Jan-01 | A/Hong Kong/4801/2014 | Crick Worldwide Influenza Centre | Centers for Disease Control and Prevention |
| EPI578438 | HA | Congo, the Democatic Republic of | 2014-Dec-18 | A/Congo/2538/2014 | INRB Service de Virologie | Centers for Disease Control and Prevention |
| EPI578446 | HA | Hong Kong (SAR) | 2014-Jan-01 | A/Hong Kong/7127/2014 | Crick Worldwide Influenza Centre | Centers for Disease Control and Prevention |
| EPI578454 | HA | Congo, the Democatic Republic of | 2014-Dec-17 | A/Congo/2544/2014 | INRB Service de Virologie | Centers for Disease Control and Prevention |
| EPI578469 | HA | Kazakhstan | 2015-Jan-13 | A/Kazakhstan/951/2015 | CSEE | Centers for Disease Control and Prevention |
| EPI578477 | HA | Kazakhstan | 2015-Jan-13 | A/Kazakhstan/37/2015 | CSEE | Centers for Disease Control and Prevention |
| EPI578485 | HA | Kazakhstan | 2015-Jan-13 | A/Kazakhstan/47/2015 | CSEE | Centers for Disease Control and Prevention |
| EPI578493 | HA | Kazakhstan | 2015-Jan-08 | A/Kazakhstan/38/2015 | CSEE | Centers for Disease Control and Prevention |

| | | | | | | |
|---|---|---|---|---|---|---|
| EPI578501 | HA | United States | 2015-Jan-05 | A/Idaho/08/2015 | State of Idaho Bureau of Laboratories | Centers for Disease Control and Prevention |
| EPI578509 | HA | United States | 2015-Jan-15 | A/Iowa/06/2015 | Iowa State Hygienic Laboratory | Centers for Disease Control and Prevention |
| EPI578517 | HA | United States | 2015-Jan-06 | A/California/09/2015 | California Department of Health Services | Centers for Disease Control and Prevention |
| EPI578525 | HA | United States | 2015-Jan-13 | A/Iowa/05/2015 | Iowa State Hygienic Laboratory | Centers for Disease Control and Prevention |
| EPI578533 | HA | Lao, People's Democratic Republic | 2014-Nov-21 | A/Laos/732/2014 | National Center for Laboratory and Epidemiology | Centers for Disease Control and Prevention |
| EPI578560 | HA | United States | 2015-Jan-05 | A/California/08/2015 | California Department of Health Services | Centers for Disease Control and Prevention |
| EPI578571 | HA | United States | 2015-Jan-01 | A/California/06/2015 | California Department of Health Services | Centers for Disease Control and Prevention |
| EPI578579 | HA | United States | 2015-Feb-13 | A/Maine/09/2015 | Maine Health and Environmental Testing Laboratory | Centers for Disease Control and Prevention |
| EPI578587 | HA | United States | 2015-Jan-07 | A/California/04/2015 | California Department of Health Services | Centers for Disease Control and Prevention |
| EPI578595 | HA | United States | 2015-Jan-14 | A/Nebraska/07/2015 | Nebraska Public Health Lab | Centers for Disease Control and Prevention |
| EPI578603 | HA | United States | 2015-Feb-18 | A/Maine/10/2015 | Maine Health and Environmental Testing Laboratory | Centers for Disease Control and Prevention |
| EPI578611 | HA | United States | 2015-Jan-15 | A/Missouri/07/2015 | Missouri Department. of Health & Senior Services | Centers for Disease Control and Prevention |
| EPI578619 | HA | United States | 2015-Jan-12 | A/Massachusetts/05/2015 | Massachusetts Department of Public Health | Centers for Disease Control and Prevention |
| EPI578627 | HA | United States | 2015-Jan-04 | A/Minnesota/01/2015 | Minnesota Department of Health | Centers for Disease Control and Prevention |
| EPI578635 | HA | United States | 2015-Jan-07 | A/Minnesota/02/2015 | Minnesota Department of Health | Centers for Disease Control and Prevention |
| EPI578649 | HA | United States | 2015-Jan-01 | A/Georgia/02/2015 | Georgia Public Health Laboratory | Centers for Disease Control and Prevention |
| EPI578657 | HA | United States | 2015-Jan-01 | A/Georgia/01/2015 | Georgia Public Health Laboratory | Centers for Disease Control and Prevention |
| EPI578665 | HA | United States | 2015-Jan-04 | A/Virginia/02/2015 | University of Virginia, Medical Labs/Microbiology | Centers for Disease Control and Prevention |
| EPI578673 | HA | United States | 2015-Jan-03 | A/Virginia/06/2015 | University of Virginia, Medical Labs/Microbiology | Centers for Disease Control and Prevention |
| EPI578681 | HA | United States | 2015-Jan-04 | A/Virginia/03/2015 | University of Virginia, Medical Labs/Microbiology | Centers for Disease Control and Prevention |
| EPI578694 | HA | Congo, the Democatic Republic of | 2014-Dec-29 | A/Congo/2611/2014 | INRB Service de Virologie | Centers for Disease Control and Prevention |
| EPI578702 | HA | Congo, the Democatic Republic of | 2014-Dec-18 | A/Congo/2557/2014 | INRB Service de Virologie | Centers for Disease Control and Prevention |
| EPI578710 | HA | United States | 2015-Jan-13 | A/Washington/03/2015 | Marshfield Clinic Research Foundation | Centers for Disease Control and Prevention |
| EPI578718 | HA | United States | 2015-Jan-12 | A/West Virginia/03/2015 | West Virginia Office of Laboratory Services | Centers for Disease Control and Prevention |

| | | | | | | | |
|---|---|---|---|---|---|---|---|
| EPI578726 | HA | United States | 2015-Jan-14 | A/Washington/17/2015 | Marshfield Clinic Research Foundation | Centers for Disease Control and Prevention | |
| EPI578734 | HA | United States | 2015-Jan-12 | A/Washington/08/2015 | Marshfield Clinic Research Foundation | Centers for Disease Control and Prevention | |
| EPI578742 | HA | United States | 2015-Jan-13 | A/Texas/22/2015 | Texas Department of State Health Services-Laboratory Services | Centers for Disease Control and Prevention | |
| EPI578750 | HA | United States | 2015-Jan-10 | A/Wisconsin/10/2015 | Wisconsin State Laboratory of Hygiene | Centers for Disease Control and Prevention | |
| EPI578758 | HA | United States | 2015-Jan-11 | A/Wisconsin/11/2015 | Wisconsin State Laboratory of Hygiene | Centers for Disease Control and Prevention | |
| EPI578773 | HA | United States | 2015-Jan-15 | A/Delaware/14/2015 | Delaware Public Health Lab | Centers for Disease Control and Prevention | |
| EPI578781 | HA | United States | 2015-Jan-09 | A/Delaware/12/2015 | Delaware Public Health Lab | Centers for Disease Control and Prevention | |
| EPI578794 | HA | United States | 2015-Jan-09 | A/Michigan/11/2015 | University of Michigan SPH EPID | Centers for Disease Control and Prevention | |
| EPI578802 | HA | United States | 2015-Jan-14 | A/Texas/21/2015 | Texas Department of State Health Services-Laboratory Services | Centers for Disease Control and Prevention | |
| EPI578810 | HA | United States | 2015-Jan-01 | A/Delaware/10/2015 | Delaware Public Health Lab | Centers for Disease Control and Prevention | |
| EPI578818 | HA | United States | 2015-Jan-14 | A/Kansas/05/2015 | Kansas Department of Health and Environment | Centers for Disease Control and Prevention | |
| EPI578826 | HA | United States | 2015-Jan-15 | A/Kansas/06/2015 | Kansas Department of Health and Environment | Centers for Disease Control and Prevention | |
| EPI578834 | HA | United States | 2015-Jan-13 | A/Texas/20/2015 | Texas Department of State Health Services-Laboratory Services | Centers for Disease Control and Prevention | |
| EPI578842 | HA | United States | 2015-Jan-09 | A/Michigan/13/2015 | University of Michigan SPH EPID | Centers for Disease Control and Prevention | |
| EPI578850 | HA | United States | 2015-Jan-08 | A/Michigan/12/2015 | University of Michigan SPH EPID | Centers for Disease Control and Prevention | |
| EPI578858 | HA | Cambodia | 2014-Aug-05 | A/Cambodia/870/2014 | National Public Health Laboratory | Centers for Disease Control and Prevention | Chen,B.B.; Phuah,S.P.; Poh,M.K.; Chen,S.L.; Cui,L. |
| EPI578866 | HA | United States | 2015-Jan-14 | A/Illinois/02/2015 | Illinois Department of Public Health-Chicago | Centers for Disease Control and Prevention | Chen,B.B.; Phuah,S.P.; Poh,M.K.; Chen,S.L.; Cui,L. |
| EPI578874 | HA | Uganda | 2014-Jan-01 | A/Uganda/4530/2014 | Uganda Virus Research Institute (UVRI), National Influenza Center | Centers for Disease Control and Prevention | Chen,B.B.; Phuah,S.P.; Poh,M.K.; Chen,S.L.; Cui,L. |
| EPI578881 | HA | United States | 2015-Jan-06 | A/Tennessee/06/2015 | Tennessee Department of Health Laboratory-Nashville | Centers for Disease Control and Prevention | Chen,B.B.; Phuah,S.P.; Poh,M.K.; Chen,S.L.; Cui,L. |
| EPI578889 | HA | United States | 2015-Jan-06 | A/Tennessee/05/2015 | Tennessee Department of Health Laboratory-Nashville | Centers for Disease Control and Prevention | |
| EPI578897 | HA | United States | 2015-Jan-15 | A/Michigan/08/2015 | Michigan Department of Community Health | Centers for Disease Control and Prevention | |
| EPI578910 | HA | United States | 2015-Jan-15 | A/Michigan/10/2015 | Michigan Department of Community Health | Centers for Disease Control and Prevention | |
| EPI579060 | HA | Kazakhstan | 2015-Jan-08 | A/Kazakhstan/39/2015 | CSEE | Centers for Disease Control and Prevention | |
| EPI579066 | HA | United States | 2015-Feb-27 | A/New Mexico/06/2015 | New Mexico Department of Health | Centers for Disease Control and Prevention | |
| EPI579074 | HA | United States | 2015-Jan-23 | A/New York/18/2015 | New York City Department of Health | Centers for Disease Control and Prevention | |

| | | | | | | |
|---|---|---|---|---|---|---|
| EPI579086 | HA | United States | 2015-Jan-16 | A/Maine/15/2015 | Maine Health and Environmental Testing Laboratory | Centers for Disease Control and Prevention |
| EPI579094 | HA | United States | 2015-Jan-16 | A/Maine/13/2015 | Maine Health and Environmental Testing Laboratory | Centers for Disease Control and Prevention |
| EPI579102 | HA | United States | 2015-Jan-16 | A/Maine/14/2015 | Maine Health and Environmental Testing Laboratory | Centers for Disease Control and Prevention |
| EPI579110 | HA | United States | 2015-Jan-24 | A/Missouri/09/2015 | Missouri Department. of Health & Senior Services | Centers for Disease Control and Prevention |
| EPI579118 | HA | United States | 2015-Jan-19 | A/Delaware/16/2015 | Delaware Public Health Lab | Centers for Disease Control and Prevention |
| EPI579140 | HA | United States | 2015-Jan-26 | A/Texas/34/2015 | Texas Department of State Health Services-Laboratory Services | Centers for Disease Control and Prevention |
| EPI579148 | HA | United States | 2015-Jan-26 | A/Texas/32/2015 | Texas Department of State Health Services-Laboratory Services | Centers for Disease Control and Prevention |
| EPI579156 | HA | United States | 2015-Jan-26 | A/Texas/30/2015 | Texas Department of State Health Services-Laboratory Services | Centers for Disease Control and Prevention |
| EPI579169 | HA | United States | 2015-Jan-28 | A/Texas/27/2015 | Texas Department of State Health Services-Laboratory Services | Centers for Disease Control and Prevention |
| EPI579177 | HA | United States | 2015-Jan-27 | A/Texas/33/2015 | Texas Department of State Health Services-Laboratory Services | Centers for Disease Control and Prevention |
| EPI579185 | HA | United States | 2015-Jan-27 | A/Texas/31/2015 | Texas Department of State Health Services-Laboratory Services | Centers for Disease Control and Prevention |
| EPI579193 | HA | United States | 2015-Jan-26 | A/New Jersey/09/2015 | New Jersey Department of Health & Senior Services | Centers for Disease Control and Prevention |
| EPI579201 | HA | United States | 2015-Jan-26 | A/New Jersey/10/2015 | New Jersey Department of Health & Senior Services | Centers for Disease Control and Prevention |
| EPI579209 | HA | United States | 2015-Feb-01 | A/New Hampshire/12/2015 | New Hampshire Public Health Laboratories | Centers for Disease Control and Prevention |
| EPI579217 | HA | United States | 2015-Jan-22 | A/West Virginia/06/2015 | West Virginia Office of Laboratory Services | Centers for Disease Control and Prevention |
| EPI579225 | HA | United States | 2015-Jan-25 | A/West Virginia/07/2015 | West Virginia Office of Laboratory Services | Centers for Disease Control and Prevention |
| EPI579244 | HA | United States | 2015-Feb-03 | A/Wisconsin/15/2015 | Marshfield Clinic Research Foundation | Centers for Disease Control and Prevention |
| EPI579264 | HA | United States | 2015-Feb-16 | A/Texas/11/2015 | Texas Department of State Health Services-Laboratory Services | Centers for Disease Control and Prevention |
| EPI579270 | HA | United States | 2015-Feb-11 | A/New Hampshire/18/2015 | New Hampshire Public Health Laboratories | Centers for Disease Control and Prevention |
| EPI579278 | HA | United States | 2015-Feb-13 | A/Wisconsin/07/2015 | Wisconsin State Laboratory of Hygiene | Centers for Disease Control and Prevention |
| EPI579286 | HA | United States | 2015-Feb-06 | A/Washington/05/2015 | Marshfield Clinic Research Foundation | Centers for Disease Control and Prevention |
| EPI579298 | HA | United States | 2015-Feb-13 | A/Washington/16/2015 | Marshfield Clinic Research Foundation | Centers for Disease Control and Prevention |
| EPI579303 | HA | United States | 2015-Feb-06 | A/Wisconsin/08/2015 | Marshfield Clinic Research Foundation | Centers for Disease Control and Prevention |
| EPI579309 | HA | United States | 2015-Feb-10 | A/Wisconsin/14/2015 | Marshfield Clinic Research Foundation | Centers for Disease Control and Prevention |
| EPI579314 | HA | United States | 2015-Feb-18 | A/Wisconsin/09/2015 | Marshfield Clinic Research Foundation | Centers for Disease Control and Prevention |

| | | | | | | | |
|---|---|---|---|---|---|---|---|
| EPI579323 | HA | United States | 2015-Feb-09 | A/Missouri/06/2015 | Missouri Department. of Health & Senior Services | Centers for Disease Control and Prevention | Deng,Y-M.; Iannello,P.; Spirason,N.; Jelley,L.; Lau,H.; |
| EPI579329 | HA | United States | 2015-Feb-04 | A/Rhode Island/08/2015 | Rhode Island Department of Health | Centers for Disease Control and Prevention | |
| EPI579335 | HA | United States | 2015-Feb-04 | A/Virginia/04/2015 | Virginia Division of Consolidated Laboratories | Centers for Disease Control and Prevention | |
| EPI579342 | HA | United States | 2015-Feb-01 | A/New Jersey/04/2015 | New Jersey Department of Health & Senior Services | Centers for Disease Control and Prevention | |
| EPI579349 | HA | United States | 2015-Feb-24 | A/Delaware/13/2015 | Delaware Public Health Lab | Centers for Disease Control and Prevention | |
| EPI579359 | HA | Australia | 2014-Aug-25 | A/Victoria/5006/2014 | WHO Collaborating Centre for Reference and Research on Influenza | Centers for Disease Control and Prevention | |
| EPI579366 | HA | Cambodia | 2014-Nov-11 | A/Cambodia/1244/2014 | National Public Health Laboratory | Centers for Disease Control and Prevention | |
| EPI579380 | HA | United States | 2015-Feb-20 | A/Utah/11/2015 | Utah Department of Health | Centers for Disease Control and Prevention | |
| EPI579388 | HA | United States | 2015-Jan-05 | A/Idaho/10/2015 | State of Idaho Bureau of Laboratories | Centers for Disease Control and Prevention | |
| EPI579396 | HA | United States | 2015-Jan-23 | A/Nevada/15/2015 | Southern Nevada Public Health Lab | Centers for Disease Control and Prevention | |
| EPI579404 | HA | United States | 2015-Jan-20 | A/New Mexico/07/2015 | New Mexico Department of Health | Centers for Disease Control and Prevention | |
| EPI579412 | HA | United States | 2015-Jan-19 | A/New Mexico/08/2015 | New Mexico Department of Health | Centers for Disease Control and Prevention | |
| EPI579420 | HA | United States | 2015-Jan-20 | A/Arkansas/05/2015 | Arkansas Department of Health | Centers for Disease Control and Prevention | |
| EPI579428 | HA | United States | 2015-Jan-21 | A/Arkansas/04/2015 | Arkansas Department of Health | Centers for Disease Control and Prevention | |
| EPI579436 | HA | United States | 2015-Jan-19 | A/Texas/26/2015 | Baylor College of Medicine | Centers for Disease Control and Prevention | |
| EPI579444 | HA | United States | 2015-Jan-21 | A/Texas/24/2015 | Baylor College of Medicine | Centers for Disease Control and Prevention | |
| EPI579452 | HA | United States | 2015-Jan-24 | A/Texas/29/2015 | Baylor College of Medicine | Centers for Disease Control and Prevention | |
| EPI579460 | HA | United States | 2015-Jan-21 | A/Texas/23/2015 | Baylor College of Medicine | Centers for Disease Control and Prevention | |
| EPI579468 | HA | United States | 2015-Jan-21 | A/Texas/28/2015 | Baylor College of Medicine | Centers for Disease Control and Prevention | |
| EPI579476 | HA | United States | 2015-Jan-27 | A/Washington/06/2015 | Washington State Public Health Laboratory | Centers for Disease Control and Prevention | |
| EPI579484 | HA | United States | 2015-Jan-26 | A/Idaho/04/2015 | State of Idaho Bureau of Laboratories | Centers for Disease Control and Prevention | |
| EPI579492 | HA | United States | 2015-Feb-04 | A/Texas/16/2015 | Baylor College of Medicine | Centers for Disease Control and Prevention | |
| EPI579500 | HA | United States | 2015-Feb-08 | A/Texas/18/2015 | Baylor College of Medicine | Centers for Disease Control and Prevention | |
| EPI579516 | HA | United States | 2015-Feb-11 | A/Nevada/13/2015 | Southern Nevada Public Health Lab | Centers for Disease Control and Prevention | |
| EPI579522 | HA | United States | 2015-Jan-28 | A/Idaho/05/2015 | State of Idaho Bureau of Laboratories | Centers for Disease Control and Prevention | |

| | | | | | | |
|---|---|---|---|---|---|---|
| EPI579530 | HA | United States | 2015-Feb-04 | A/Idaho/09/2015 | State of Idaho Bureau of Laboratories | Centers for Disease Control and Prevention |
| EPI579539 | HA | United States | 2015-Feb-11 | A/Oklahoma/05/2015 | Oklahoma State Department of Health | Centers for Disease Control and Prevention |
| EPI579546 | HA | United States | 2015-Feb-11 | A/Oklahoma/06/2015 | Oklahoma State Department of Health | Centers for Disease Control and Prevention |
| EPI579553 | HA | United States | 2015-Feb-12 | A/California/16/2015 | California Department of Health Services | Centers for Disease Control and Prevention |
| EPI579559 | HA | United States | 2015-Feb-15 | A/California/12/2015 | California Department of Health Services | Centers for Disease Control and Prevention |
| EPI579567 | HA | United States | 2015-Jan-16 | A/Kansas/07/2015 | Kansas Department of Health and Environment | Centers for Disease Control and Prevention |
| EPI579603 | HA | United States | 2015-Jan-19 | A/Wisconsin/18/2015 | Marshfield Clinic Research Foundation | Centers for Disease Control and Prevention |
| EPI579611 | HA | United States | 2015-Jan-22 | A/Wisconsin/19/2015 | Marshfield Clinic Research Foundation | Centers for Disease Control and Prevention |
| EPI579623 | HA | United States | 2015-Jan-19 | A/Utah/10/2015 | Utah Department of Health | Centers for Disease Control and Prevention |
| EPI579631 | HA | United States | 2015-Jan-21 | A/Florida/21/2015 | Florida Department of Health-Jacksonville | Centers for Disease Control and Prevention |
| EPI579639 | HA | United States | 2015-Jan-20 | A/Florida/24/2015 | Florida Department of Health-Tampa | Centers for Disease Control and Prevention |
| EPI579647 | HA | United States | 2015-Jan-16 | A/Ohio/01/2015 | Ohio Department of Health Laboratories | Centers for Disease Control and Prevention |
| EPI579655 | HA | United States | 2015-Jan-17 | A/Ohio/02/2015 | Ohio Department of Health Laboratories | Centers for Disease Control and Prevention |
| EPI579663 | HA | United States | 2015-Jan-21 | A/Ohio/03/2015 | Ohio Department of Health Laboratories | Centers for Disease Control and Prevention |
| EPI579671 | HA | United States | 2015-Jan-07 | A/Colorado/05/2015 | Colorado Department of Health Lab | Centers for Disease Control and Prevention |
| EPI579679 | HA | United States | 2015-Jan-08 | A/Colorado/06/2015 | Colorado Department of Health Lab | Centers for Disease Control and Prevention |
| EPI579687 | HA | United States | 2015-Jan-14 | A/Colorado/07/2015 | Colorado Department of Health Lab | Centers for Disease Control and Prevention |
| EPI579695 | HA | United States | 2015-Jan-06 | A/Kentucky/02/2015 | Kentucky Division of Laboratory Services | Centers for Disease Control and Prevention |
| EPI579703 | HA | United States | 2015-Jan-16 | A/Kentucky/03/2015 | Kentucky Division of Laboratory Services | Centers for Disease Control and Prevention |
| EPI579711 | HA | United States | 2015-Jan-22 | A/Kentucky/04/2015 | Kentucky Division of Laboratory Services | Centers for Disease Control and Prevention |
| EPI579719 | HA | United States | 2015-Jan-15 | A/North Dakota/04/2015 | North Dakota Department of Health | Centers for Disease Control and Prevention |
| EPI579726 | HA | United States | 2015-Jan-15 | A/Wyoming/02/2015 | Wyoming Public Health Laboratory | Centers for Disease Control and Prevention |
| EPI579734 | HA | United States | 2015-Jan-13 | A/North Carolina/10/2015 | North Carolina State Laboratory of Public Health | Centers for Disease Control and Prevention |
| EPI579742 | HA | United States | 2015-Jan-13 | A/Indiana/03/2015 | Indiana State Department of Health Laboratories | Centers for Disease Control and Prevention |
| EPI579748 | HA | United States | 2015-Feb-03 | A/Tennessee/11/2015 | Tennessee Department of Health Laboratory-Nashville | Centers for Disease Control and Prevention |

| | | | | | | | |
|---|---|---|---|---|---|---|---|
| EPI579756 | HA | United States | 2015-Feb-22 | A/Wisconsin/13/2015 | Marshfield Clinic Research Foundation | Centers for Disease Control and Prevention | |
| EPI579764 | HA | United States | 2015-Feb-18 | A/Washington/12/2015 | Marshfield Clinic Research Foundation | Centers for Disease Control and Prevention | |
| EPI579779 | HA | United States | 2015-Feb-25 | A/Washington/13/2015 | Marshfield Clinic Research Foundation | Centers for Disease Control and Prevention | |
| EPI579796 | HA | United States | 2015-Feb-24 | A/Wisconsin/20/2015 | Wisconsin State Laboratory of Hygiene | Centers for Disease Control and Prevention | |
| EPI579801 | HA | United States | 2015-Feb-18 | A/New Hampshire/13/2015 | New Hampshire Public Health Laboratories | Centers for Disease Control and Prevention | |
| EPI579809 | HA | United States | 2015-Feb-18 | A/Nevada/18/2015 | Southern Nevada Public Health Lab | Centers for Disease Control and Prevention | |
| EPI579817 | HA | United States | 2015-Jan-27 | A/California/19/2015 | California Department of Health Services | Centers for Disease Control and Prevention | |
| EPI579825 | HA | United States | 2015-Feb-17 | A/New York/19/2015 | New York City Department of Health | Centers for Disease Control and Prevention | |
| EPI579833 | HA | United States | 2015-Feb-14 | A/Oregon/04/2015 | Oregon Public Health Laboratory | Centers for Disease Control and Prevention | |
| EPI579841 | HA | United States | 2015-Feb-19 | A/Kansas/08/2015 | Kansas Department of Health and Environment | Centers for Disease Control and Prevention | |
| EPI579846 | HA | United States | 2015-Feb-07 | A/New Mexico/10/2015 | New Mexico Department of Health | Centers for Disease Control and Prevention | |
| EPI579854 | HA | United States | 2015-Feb-23 | A/Delaware/18/2015 | Delaware Public Health Lab | Centers for Disease Control and Prevention | |
| EPI579859 | HA | United States | 2015-Feb-18 | A/Nevada/17/2015 | Southern Nevada Public Health Lab | Centers for Disease Control and Prevention | |
| EPI579866 | HA | United States | 2015-Jan-13 | A/Minnesota/03/2015 | Minnesota Department of Health | Centers for Disease Control and Prevention | |
| EPI579874 | HA | United States | 2015-Jan-29 | A/New Jersey/12/2015 | New Jersey Department of Health & Senior Services | Centers for Disease Control and Prevention | |
| EPI579879 | HA | United States | 2015-Jan-18 | A/Hawaii/05/2015 | State of Hawaii Department of Health | Centers for Disease Control and Prevention | Resende,Paola; Motta,Fernando; Born,Priscila; Mesquita,Milene; Siqueira,Marilda |
| EPI579884 | HA | United States | 2015-Jan-07 | A/Pennsylvania/11/2015 | UPMC-CLB Dept of Microbiology | Centers for Disease Control and Prevention | |
| EPI579889 | HA | United States | 2015-Feb-25 | A/New York/20/2015 | New York City Department of Health | Centers for Disease Control and Prevention | |
| EPI579897 | HA | United States | 2015-Jan-20 | A/California/20/2015 | California Department of Health Services | Centers for Disease Control and Prevention | |
| EPI579913 | HA | United States | 2015-Feb-21 | A/New Jersey/11/2015 | New Jersey Department of Health & Senior Services | Centers for Disease Control and Prevention | |
| EPI579921 | HA | United States | 2015-Jan-15 | A/Florida/25/2015 | Florida Department of Health-Tampa | Centers for Disease Control and Prevention | |
| EPI580481 | HA | Lebanon | 2014-Jan-09 | A/Lebanon/14L41/2014 | | Niigata University (DPH) | |
| EPI580489 | HA | Lebanon | 2014-Jan-22 | A/Lebanon/14L45/2014 | | Niigata University (DPH) | |
| EPI580497 | HA | Lebanon | 2014-Jan-31 | A/Lebanon/14L56/2014 | | Niigata University (DPH) | |
| EPI580505 | HA | Lebanon | 2014-Feb-10 | A/Lebanon/14L78/2014 | | Niigata University (DPH) | Deng,Y-D.; Iannello,P.; Spirason,N.; Jelley,L.; Lau,H.; |
| EPI580521 | HA | Japan | 2015-Jan-19 | A/Niigata/14F067/2015 | | Niigata University (DPH) | |

| | | | | | | | |
|---|---|---|---|---|---|---|---|
| EPI580529 | HA | Japan | 2015-Jan-26 | A/Niigata/14F197/2015 | | Niigata University (DPH) | |
| EPI580545 | HA | Japan | 2015-Jan-14 | A/Gunma/14G011/2015 | | Niigata University (DPH) | |
| EPI580561 | HA | Japan | 2015-Jan-05 | A/Nagasaki/14N024/2015 | | Niigata University (DPH) | Li, J.; Shao, T.J.; Yu, X.F.; Pu, X.Y.; Pan, J.C. |
| EPI580569 | HA | Japan | 2015-Jan-14 | A/Nagasaki/14N039/2015 | | Niigata University (DPH) | Li, J.; Shao, T.J.; Yu, X.F.; Pu, X.Y.; Pan, J.C. |
| EPI580577 | HA | Japan | 2015-Jan-05 | A/Nagasaki/14N072/2015 | | Niigata University (DPH) | Li, J.; Shao, T.J.; Yu, X.F.; Pu, X.Y.; Pan, J.C. |
| EPI580585 | HA | Japan | 2015-Jan-06 | A/Okinawa/14T003/2015 | | Niigata University (DPH) | Li, J.; Shao, T.J.; Yu, X.F.; Pu, X.Y.; Pan, J.C. |
| EPI580593 | HA | Japan | 2015-Jan-13 | A/Okinawa/14T004/2015 | | Niigata University (DPH) | Li, J.; Shao, T.J.; Yu, X.F.; Pu, X.Y.; Pan, J.C. |
| EPI580601 | HA | Japan | 2015-Jan-20 | A/Okinawa/14T006/2015 | | Niigata University (DPH) | Li, J.; Shao, T.J.; Yu, X.F.; Pu, X.Y.; Pan, J.C. |
| EPI580609 | HA | Japan | 2015-Jan-22 | A/Okinawa/14T007/2015 | | Niigata University (DPH) | Li, J.; Shao, T.J.; Yu, X.F.; Pu, X.Y.; Pan, J.C. |
| EPI580617 | HA | Japan | 2014-Nov-21 | A/Nagasaki/14N012/2014 | | Niigata University (DPH) | Li, J.; Shao, T.J.; Yu, X.F.; Pu, X.Y.; Pan, J.C. |
| EPI580625 | HA | Japan | 2014-Dec-09 | A/Nagasaki/14N013/2014 | | Niigata University (DPH) | Li, J.; Shao, T.J.; Yu, X.F.; Pu, X.Y.; Pan, J.C. |
| EPI582005 | HA | Sweden | 2015-Feb-10 | A/Stockholm/12/2015 | | Public Health Agency of Sweden | Li, J.; Shao, T.J.; Yu, X.F.; Pu, X.Y.; Pan, J.C. |
| EPI582013 | HA | Sweden | 2015-Feb-10 | A/Stockholm/13/2015 | | Public Health Agency of Sweden | Li, J.; Shao, T.J.; Yu, X.F.; Pu, X.Y.; Pan, J.C. |
| EPI582021 | HA | Sweden | 2015-Feb-18 | A/Stockholm/14/2015 | | Public Health Agency of Sweden | Li, J.; Shao, T.J.; Yu, X.F.; Pu, X.Y.; Pan, J.C. |
| EPI582029 | HA | Sweden | 2015-Feb-13 | A/Stockholm/15/2015 | | Public Health Agency of Sweden | Li, J.; Shao, T.J.; Yu, X.F.; Pu, X.Y.; Pan, J.C. |
| EPI582037 | HA | Sweden | 2015-Feb-18 | A/Stockholm/16/2015 | | Public Health Agency of Sweden | Li, J.; Shao, T.J.; Yu, X.F.; Pu, X.Y.; Pan, J.C. |
| EPI582045 | HA | Sweden | 2015-Feb-11 | A/Stockholm/17/2015 | | Public Health Agency of Sweden | Li, J.; Shao, T.J.; Yu, X.F.; Pu, X.Y.; Pan, J.C. |
| EPI582053 | HA | Sweden | 2015-Feb-11 | A/Stockholm/18/2015 | | Public Health Agency of Sweden | Li, J.; Shao, T.J.; Yu, X.F.; Pu, X.Y.; Pan, J.C. |
| EPI582088 | HA | Sweden | 2015-Feb-22 | A/Sweden/14/2015 | | Public Health Agency of Sweden | |
| EPI582096 | HA | Sweden | 2015-Feb-25 | A/Stockholm/19/2015 | | Public Health Agency of Sweden | |
| EPI582104 | HA | Sweden | 2015-Feb-24 | A/Stockholm/20/2015 | | Public Health Agency of Sweden | |
| EPI582138 | HA | Sweden | 2015-Feb-27 | A/Sweden/15/2015 | | Public Health Agency of Sweden | |
| EPI582146 | HA | Sweden | 2015-Mar-05 | A/Sweden/16/2015 | | Public Health Agency of Sweden | |
| EPI582154 | HA | Sweden | 2015-Mar-05 | A/Sweden/17/2015 | | Public Health Agency of Sweden | |
| EPI582218 | HA | Sweden | 2015-Mar-05 | A/Sweden/19/2015 | | Public Health Agency of Sweden | |
| EPI582274 | HA | Sweden | 2015-Feb-23 | A/Malmoe/2/2015 | | Public Health Agency of Sweden | |
| EPI583972 | HA | Japan | 2015-Jan-10 | A/SAPPORO/6/2015 | Sapporo City Institute of Public Health | National Institute of Infectious Diseases (NIID) | |
| EPI583978 | HA | Japan | 2015-Jan-17 | A/KANAGAWA/15/2015 | Kanagawa Prefectural Institute of Public Health | National Institute of Infectious Diseases (NIID) | |
| EPI583986 | HA | Japan | 2014-Dec-06 | A/WAKAYAMA-C/104/2014 | Wakayama City Institute of Public Health | National Institute of Infectious Diseases (NIID) | |

| | | | | | | | |
|---|---|---|---|---|---|---|---|
| EPI583994 | HA | Japan | 2015-Feb-17 | A/HYOGO/3213/2015 | Hyogo Prefectural Institute of Public Health and Consumer Sciences | National Institute of Infectious Diseases (NIID) | |
| EPI584000 | HA | Japan | 2015-Jan-08 | A/AOMORI/3/2015 | Aomori Prefectural Institute of Public Health and Environment | National Institute of Infectious Diseases (NIID) | |
| EPI584559 | HA | United States | 2015-Feb-05 | A/Hawaii/07/2015 | State of Hawaii Department of Health | Centers for Disease Control and Prevention | |
| EPI584567 | HA | United States | 2014-Nov-25 | A/Virginia/26/2014 | University of Virginia, Medical Labs/Microbiology | Centers for Disease Control and Prevention | |
| EPI584575 | HA | United States | 2015-Jan-22 | A/Alabama/04/2015 | ADPH Bureau of Clinical Laboratories | Centers for Disease Control and Prevention | |
| EPI584583 | HA | United States | 2014-Dec-17 | A/Maryland/50/2014 | Maryland Department of Health and Mental Hygiene | Centers for Disease Control and Prevention | |
| EPI584584 | HA | United States | 2014-Dec-22 | A/Pennsylvania/60/2014 | Pennsylvania Department of Health | Centers for Disease Control and Prevention | Galiano,M. |
| EPI584592 | HA | United States | 2014-Dec-21 | A/Massachusetts/27/2014 | Massachusetts Department of Public Health | Centers for Disease Control and Prevention | Galiano,M. |
| EPI584605 | HA | United States | 2015-Jan-26 | A/Florida/17/2015 | Florida Department of Health-Jacksonville | Centers for Disease Control and Prevention | Galiano,M. |
| EPI584613 | HA | United States | 2014-Dec-08 | A/Tennessee/25/2014 | St. Judes Childrens Research Hospital | Centers for Disease Control and Prevention | Galiano,M. |
| EPI584626 | HA | Lao, People's Democratic Republic | 2014-Oct-20 | A/Laos/1115/2014 | National Center for Laboratory and Epidemiology | Centers for Disease Control and Prevention | Galiano,M. |
| EPI584634 | HA | United States | 2014-Dec-18 | A/Florida/89/2014 | Florida Department of Health-Jacksonville | Centers for Disease Control and Prevention | Galiano,M. |
| EPI584642 | HA | United States | 2014-Dec-09 | A/Virginia/42/2014 | University of Virginia, Medical Labs/Microbiology | Centers for Disease Control and Prevention | Galiano,M. |
| EPI584650 | HA | Uganda | 2014-Apr-17 | A/Uganda/583/2014 | Uganda Virus Research Institute (UVRI), National Influenza Center | Centers for Disease Control and Prevention | Galiano,M. |
| EPI584677 | HA | Uganda | 2014-May-26 | A/Uganda/1590/2014 | Uganda Virus Research Institute (UVRI), National Influenza Center | Centers for Disease Control and Prevention | Galiano,M. |
| EPI584685 | HA | United States | 2015-Jan-21 | A/Washington/18/2015 | Marshfield Clinic Research Foundation | Centers for Disease Control and Prevention | Galiano,M. |
| EPI584699 | HA | United States | 2014-Dec-09 | A/Washington/61/2014 | | Centers for Disease Control and Prevention | Galiano,M. |
| EPI584714 | HA | United States | 2014-Dec-28 | A/Virginia/41/2014 | University of Virginia, Medical Labs/Microbiology | Centers for Disease Control and Prevention | Galiano,M. |
| EPI584717 | HA | United States | 2014-Nov-19 | A/Louisiana/20/2014 | Louisiana Department of Health and Hospitals | Centers for Disease Control and Prevention | Galiano,M. |
| EPI585299 | HA | Sweden | 2015-Apr-03 | A/Karlstad/3/2015 | | Public Health Agency of Sweden | Galiano,M. |
| EPI585339 | HA | Sweden | 2015-Apr-08 | A/Sweden/20/2015 | | Public Health Agency of Sweden | Galiano,M. |
| EPI586564 | HA | Japan | 2015-Mar-09 | A/SAPPORO/38/2015 | Sapporo City Institute of Public Health | National Institute of Infectious Diseases (NIID) | Galiano,M. |
| EPI586568 | HA | Japan | 2015-Feb-09 | A/KOCHI/15/2015 | Kochi Public Health and Sanitation Institute | National Institute of Infectious Diseases (NIID) | Galiano,M. |
| EPI587444 | HA | New Zealand | 2014-Apr-21 | A/New Zealand/316/2014 | Institute of Environmental Science and Research | WHO Collaborating Centre for Reference and Research on Influenza | Galiano,M. |
| EPI588494 | HA | United States | 2015-Mar-02 | A/Nevada/21/2015 | Southern Nevada Public Health Lab | Centers for Disease Control and Prevention | Galiano,M. |

| | | | | | | | |
|---|---|---|---|---|---|---|---|
| EPI588514 | HA | United States | 2015-Mar-05 | A/Hawaii/08/2015 | State of Hawaii Department of Health | Centers for Disease Control and Prevention | Galiano,M. |
| EPI588522 | HA | United States | 2015-Feb-27 | A/Texas/38/2015 | Baylor College of Medicine | Centers for Disease Control and Prevention | Galiano,M. |
| EPI588530 | HA | United States | 2015-Mar-08 | A/Texas/36/2015 | Baylor College of Medicine | Centers for Disease Control and Prevention | Galiano,M. |
| EPI588537 | HA | United States | 2015-Mar-08 | A/Oregon/05/2015 | Oregon Public Health Laboratory | Centers for Disease Control and Prevention | Galiano,M. |
| EPI588545 | HA | United States | 2015-Mar-10 | A/New Mexico/12/2015 | New Mexico Department of Health | Centers for Disease Control and Prevention | Galiano,M. |
| EPI588553 | HA | United States | 2015-Mar-13 | A/Idaho/12/2015 | State of Idaho Bureau of Laboratories | Centers for Disease Control and Prevention | Galiano,M. |
| EPI588568 | HA | United States | 2015-Jan-06 | A/Georgia/03/2015 | Georgia Public Health Laboratory | Centers for Disease Control and Prevention | Galiano,M. |
| EPI588576 | HA | United States | 2015-Feb-03 | A/North Dakota/05/2015 | North Dakota Department of Health | Centers for Disease Control and Prevention | Galiano,M. |
| EPI588583 | HA | United States | 2015-Feb-24 | A/Kentucky/08/2015 | Kentucky Division of Laboratory Services | Centers for Disease Control and Prevention | Galiano,M. |
| EPI588591 | HA | United States | 2015-Mar-03 | A/Indiana/10/2015 | Indiana State Department of Health Laboratories | Centers for Disease Control and Prevention | Galiano,M. |
| EPI588605 | HA | United States | 2015-Feb-24 | A/Kentucky/07/2015 | Kentucky Division of Laboratory Services | Centers for Disease Control and Prevention | Takashita,Emi; Fujisaki,Seiichiro; Shirakura,Masayuki; Watanabe,Shinji; Odagiri,Takato |
| EPI588620 | HA | United States | 2015-Mar-07 | A/Montana/07/2015 | Montana Laboratory Services Bureau | Centers for Disease Control and Prevention | Galiano,M. |
| EPI588628 | HA | United States | 2015-Feb-18 | A/Texas/39/2015 | Baylor College of Medicine | Centers for Disease Control and Prevention | Galiano,M. |
| EPI588636 | HA | United States | 2015-Feb-26 | A/Wisconsin/23/2015 | Marshfield Clinic Research Foundation | Centers for Disease Control and Prevention | Galiano,M. |
| EPI588644 | HA | United States | 2015-Feb-27 | A/Wisconsin/22/2015 | Marshfield Clinic Research Foundation | Centers for Disease Control and Prevention | Galiano,M. |
| EPI588657 | HA | United States | 2015-Feb-25 | A/Massachusetts/09/2015 | Massachusetts Department of Public Health | Centers for Disease Control and Prevention | Galiano,M. |
| EPI588665 | HA | United States | 2015-Mar-02 | A/Maine/17/2015 | Maine Health and Environmental Testing Laboratory | Centers for Disease Control and Prevention | Galiano,M. |
| EPI588673 | HA | United States | 2015-Mar-03 | A/Maine/18/2015 | Maine Health and Environmental Testing Laboratory | Centers for Disease Control and Prevention | Galiano,M. |
| EPI588681 | HA | United States | 2015-Jan-22 | A/Maryland/01/2015 | Maryland Department of Health and Mental Hygiene | Centers for Disease Control and Prevention | Galiano,M. |
| EPI588689 | HA | United States | 2015-Jan-24 | A/Maryland/02/2015 | Maryland Department of Health and Mental Hygiene | Centers for Disease Control and Prevention | Galiano,M. |
| EPI588697 | HA | United States | 2015-Feb-16 | A/Maryland/03/2015 | Maryland Department of Health and Mental Hygiene | Centers for Disease Control and Prevention | Galiano,M. |
| EPI588712 | HA | United States | 2015-Mar-03 | A/Texas/37/2015 | Texas Department of State Health Services-Laboratory Services | Centers for Disease Control and Prevention | Galiano,M. |
| EPI588720 | HA | United States | 2015-Mar-12 | A/New Jersey/14/2015 | New Jersey Department of Health & Senior Services | Centers for Disease Control and Prevention | Galiano,M. |
| EPI588728 | HA | United States | 2015-Mar-02 | A/New Jersey/15/2015 | New Jersey Department of Health & Senior Services | Centers for Disease Control and Prevention | Galiano,M. |
| EPI588736 | HA | United States | 2015-Mar-09 | A/Wisconsin/21/2015 | Wisconsin State Laboratory of Hygiene | Centers for Disease Control and Prevention | |

| | | | | | | | |
|---|---|---|---|---|---|---|---|
| EPI588744 | HA | United States | 2015-Feb-25 | A/New Hampshire/15/2015 | New Hampshire Public Health Laboratories | Centers for Disease Control and Prevention | |
| EPI588752 | HA | United States | 2015-Feb-27 | A/New Hampshire/14/2015 | New Hampshire Public Health Laboratories | Centers for Disease Control and Prevention | Takashita,Emi; Fujisaki,Seiichiro; Shirakura,Masayuki; Watanabe,Shinji; Odagiri,Takato |
| EPI588760 | HA | United States | 2015-Mar-03 | A/New Hampshire/16/2015 | New Hampshire Public Health Laboratories | Centers for Disease Control and Prevention | |
| EPI588768 | HA | United States | 2015-Mar-11 | A/Maine/20/2015 | Maine Health and Environmental Testing Laboratory | Centers for Disease Control and Prevention | |
| EPI588776 | HA | United States | 2015-Mar-09 | A/Wisconsin/24/2015 | Marshfield Clinic Research Foundation | Centers for Disease Control and Prevention | |
| EPI588784 | HA | United States | 2015-Mar-06 | A/Washington/21/2015 | Marshfield Clinic Research Foundation | Centers for Disease Control and Prevention | |
| EPI588792 | HA | United States | 2015-Mar-13 | A/Vermont/05/2015 | Vermont Department of Health Laboratory | Centers for Disease Control and Prevention | |
| EPI588800 | HA | United States | 2015-Mar-30 | A/Rhode Island/12/2015 | Rhode Island Department of Health | Centers for Disease Control and Prevention | |
| EPI588808 | HA | United States | 2015-Mar-08 | A/Delaware/22/2015 | Delaware Public Health Lab | Centers for Disease Control and Prevention | |
| EPI588816 | HA | United States | 2015-Mar-21 | A/New Jersey/13/2015 | New Jersey Department of Health & Senior Services | Centers for Disease Control and Prevention | |
| EPI588831 | HA | United States | 2015-Feb-17 | A/Michigan/17/2015 | University of Michigan SPH EPID | Centers for Disease Control and Prevention | |
| EPI588839 | HA | United States | 2015-Mar-05 | A/Florida/27/2015 | Florida Department of Health-Tampa | Centers for Disease Control and Prevention | |
| EPI588847 | HA | United States | 2015-Mar-15 | A/North Carolina/16/2015 | North Carolina State Laboratory of Public Health | Centers for Disease Control and Prevention | Takashita,Emi; Fujisaki,Seiichiro; Shirakura,Masayuki; Watanabe,Shinji; Odagiri,Takato |
| EPI588855 | HA | United States | 2015-Mar-17 | A/North Carolina/15/2015 | North Carolina State Laboratory of Public Health | Centers for Disease Control and Prevention | |
| EPI588863 | HA | United States | 2015-Mar-08 | A/North Dakota/06/2015 | North Dakota Department of Health | Centers for Disease Control and Prevention | |
| EPI588871 | HA | United States | 2015-Mar-21 | A/Ohio/10/2015 | Ohio Department of Health Laboratories | Centers for Disease Control and Prevention | |
| EPI588879 | HA | United States | 2015-Mar-20 | A/Ohio/11/2015 | Ohio Department of Health Laboratories | Centers for Disease Control and Prevention | |
| EPI588894 | HA | United States | 2015-Mar-17 | A/Florida/26/2015 | Florida Department of Health-Jacksonville | Centers for Disease Control and Prevention | |
| EPI588902 | HA | United States | 2015-Mar-22 | A/Kentucky/06/2015 | Kentucky Division of Laboratory Services | Centers for Disease Control and Prevention | |
| EPI588910 | HA | Colombia | 2015-Feb-17 | A/Colombia/4468/2015 | Instituto Nacional de Salud de Columbia | Centers for Disease Control and Prevention | Saito, Reiko; Takemae, Nobuhiro; Saito, Takehiko; Shobugawa, Yugo; Kondo, Hiroki; Hibino, Akinobu; Sato, Isamu |
| EPI588918 | HA | Colombia | 2015-Feb-13 | A/Colombia/2473/2015 | Instituto Nacional de Salud de Columbia | Centers for Disease Control and Prevention | |
| EPI588926 | HA | Colombia | 2015-Jan-16 | A/Colombia/4352/2015 | Instituto Nacional de Salud de Columbia | Centers for Disease Control and Prevention | |
| EPI588930 | HA | Japan | 2015-Mar-12 | A/NAGANO/2202/2015 | Nagano Environmental Conservation Research Institute | National Institute of Infectious Diseases (NIID) | |

| | | | | | | |
|---|---|---|---|---|---|---|
| EPI594983 | HA | United States | 2015-Mar-02 | A/New Jersey/06/2015 | New Jersey Department of Health & Senior Services | Centers for Disease Control and Prevention |
| EPI594992 | HA | United States | 2015-Feb-19 | A/Massachusetts/15/2015 | Massachusetts Department of Public Health | Centers for Disease Control and Prevention |
| EPI594994 | HA | Russian Federation | 2014-Dec-16 | A/Chita/262/2014 | Russian Academy of Medical Sciences | Centers for Disease Control and Prevention |
| EPI595002 | HA | United States | 2015-Mar-03 | A/Kansas/04/2015 | Kansas Department of Health and Environment | Centers for Disease Control and Prevention |
| EPI595010 | HA | United States | 2015-Jan-27 | A/North Carolina/24/2015 | North Carolina State Laboratory of Public Health | Centers for Disease Control and Prevention |
| EPI595018 | HA | United States | 2015-Jan-25 | A/North Carolina/25/2015 | North Carolina State Laboratory of Public Health | Centers for Disease Control and Prevention |
| EPI595034 | HA | United States | 2015-Jan-27 | A/Tennessee/10/2015 | Tennessee Department of Health Laboratory-Nashville | Centers for Disease Control and Prevention |
| EPI595043 | HA | United States | 2015-Mar-04 | A/Wisconsin/16/2015 | Wisconsin State Laboratory of Hygiene | Centers for Disease Control and Prevention |
| EPI595051 | HA | United States | 2015-Mar-02 | A/Texas/19/2015 | Texas Department of State Health Services-Laboratory Services | Centers for Disease Control and Prevention |
| EPI602616 | HA | United States | 2015-Jan-07 | A/Pennsylvania/09/2015 | UPMC-CLB Dept of Microbiology | Centers for Disease Control and Prevention |
| EPI607577 | HA | United Kingdom | 2014-Oct-12 | A/England/513/2014 | Microbiology Services Colindale, Public Health England | Microbiology Services Colindale, Public Health England |
| EPI607585 | HA | United Kingdom | 2014-Oct-24 | A/England/516/2014 | Microbiology Services Colindale, Public Health England | Microbiology Services Colindale, Public Health England |
| EPI607593 | HA | United Kingdom | 2014-Dec-16 | A/England/599/2014 | Microbiology Services Colindale, Public Health England | Microbiology Services Colindale, Public Health England |
| EPI607601 | HA | United Kingdom | 2014-Dec-23 | A/England/591/2014 | Microbiology Services Colindale, Public Health England | Microbiology Services Colindale, Public Health England |
| EPI607609 | HA | United Kingdom | 2015-Jan-02 | A/England/50200164/2015 | Microbiology Services Colindale, Public Health England | Microbiology Services Colindale, Public Health England |
| EPI607617 | HA | United Kingdom | 2015-Jan-02 | A/England/50200187/2015 | Microbiology Services Colindale, Public Health England | Microbiology Services Colindale, Public Health England |
| EPI607625 | HA | United Kingdom | 2015-Jan-02 | A/England/50220068/2015 | Microbiology Services Colindale, Public Health England | Microbiology Services Colindale, Public Health England |
| EPI607633 | HA | United Kingdom | 2015-Jan-02 | A/England/50220079/2015 | Microbiology Services Colindale, Public Health England | Microbiology Services Colindale, Public Health England |
| EPI607641 | HA | United Kingdom | 2015-Jan-02 | A/England/50220082/2015 | Microbiology Services Colindale, Public Health England | Microbiology Services Colindale, Public Health England |
| EPI607649 | HA | United Kingdom | 2015-Jan-05 | A/England/50240248/2015 | Microbiology Services Colindale, Public Health England | Microbiology Services Colindale, Public Health England |
| EPI607657 | HA | United Kingdom | 2015-Jan-06 | A/England/10/2015 | Microbiology Services Colindale, Public Health England | Microbiology Services Colindale, Public Health England |
| EPI607665 | HA | United Kingdom | 2015-Jan-05 | A/England/11/2015 | Microbiology Services Colindale, Public Health England | Microbiology Services Colindale, Public Health England |
| EPI607673 | HA | United Kingdom | 2015-Jan-05 | A/England/12/2015 | Microbiology Services Colindale, Public Health England | Microbiology Services Colindale, Public Health England |
| EPI607681 | HA | United Kingdom | 2015-Jan-06 | A/England/49/2015 | Microbiology Services Colindale, Public Health England | Microbiology Services Colindale, Public Health England |
| EPI607689 | HA | United Kingdom | 2015-Jan-05 | A/England/13/2015 | Microbiology Services Colindale, Public Health England | Microbiology Services Colindale, Public Health England |

| | | | | | | | |
|---|---|---|---|---|---|---|---|
| EPI607697 | HA | United Kingdom | 2015-Jan-06 | A/England/50/2015 | Microbiology Services Colindale, Public Health England | Microbiology Services Colindale, Public Health England | |
| EPI607705 | HA | United Kingdom | 2015-Jan-05 | A/England/50260150/2015 | Microbiology Services Colindale, Public Health England | Microbiology Services Colindale, Public Health England | |
| EPI607713 | HA | United Kingdom | 2015-Jan-07 | A/England/50260151/2015 | Microbiology Services Colindale, Public Health England | Microbiology Services Colindale, Public Health England | |
| EPI607721 | HA | United Kingdom | 2015-Jan-07 | A/England/18/2015 | Microbiology Services Colindale, Public Health England | Microbiology Services Colindale, Public Health England | |
| EPI607729 | HA | United Kingdom | 2015-Jan-07 | A/England/50280041/2015 | Microbiology Services Colindale, Public Health England | Microbiology Services Colindale, Public Health England | |
| EPI607737 | HA | United Kingdom | 2015-Jan-07 | A/England/50280054/2015 | Microbiology Services Colindale, Public Health England | Microbiology Services Colindale, Public Health England | Saito, Reiko; Takemae, Nobuhiro; Saito, Takehiko; Shobugawa, Yugo; Kondo, Hiroki; Hibino, Akinobu; Masaki, Hironori |
| EPI607745 | HA | United Kingdom | 2015-Jan-08 | A/England/50280058/2015 | Microbiology Services Colindale, Public Health England | Microbiology Services Colindale, Public Health England | Saito, Reiko; Takemae, Nobuhiro; Saito, Takehiko; Shobugawa, Yugo; Kondo, Hiroki; Hibino, Akinobu; Kodo, Naoki |
| EPI607753 | HA | United Kingdom | 2015-Jan-07 | A/England/50300008/2015 | Microbiology Services Colindale, Public Health England | Microbiology Services Colindale, Public Health England | Takashita,Emi; Fujisaki,Seiichiro; Shirakura,Masayuki; Watanabe,Shinji; Odagiri,Takato |
| EPI607761 | HA | United Kingdom | 2015-Jan-08 | A/England/50300025/2015 | Microbiology Services Colindale, Public Health England | Microbiology Services Colindale, Public Health England | Takashita,Emi; Fujisaki,Seiichiro; Shirakura,Masayuki; Watanabe,Shinji; Odagiri,Takato |
| EPI607769 | HA | United Kingdom | 2015-Jan-09 | A/England/50300033/2015 | Microbiology Services Colindale, Public Health England | Microbiology Services Colindale, Public Health England | |
| EPI607777 | HA | United Kingdom | 2015-Jan-07 | A/England/50300044/2015 | Microbiology Services Colindale, Public Health England | Microbiology Services Colindale, Public Health England | |
| EPI607785 | HA | United Kingdom | 2015-Jan-12 | A/England/50320276/2015 | Microbiology Services Colindale, Public Health England | Microbiology Services Colindale, Public Health England | |
| EPI607793 | HA | United Kingdom | 2015-Jan-09 | A/England/32/2015 | Microbiology Services Colindale, Public Health England | Microbiology Services Colindale, Public Health England | |
| EPI607801 | HA | United Kingdom | 2015-Jan-12 | A/England/57/2015 | Microbiology Services Colindale, Public Health England | Microbiology Services Colindale, Public Health England | |
| EPI607809 | HA | United Kingdom | 2015-Jan-13 | A/England/50360024/2015 | Microbiology Services Colindale, Public Health England | Microbiology Services Colindale, Public Health England | |
| EPI607817 | HA | United Kingdom | 2015-Jan-14 | A/England/50360027/2015 | Microbiology Services Colindale, Public Health England | Microbiology Services Colindale, Public Health England | |
| EPI607825 | HA | United Kingdom | 2015-Jan-13 | A/England/60/2015 | Microbiology Services Colindale, Public Health England | Microbiology Services Colindale, Public Health England | |
| EPI607833 | HA | United Kingdom | 2015-Jan-14 | A/England/50360038/2015 | Microbiology Services Colindale, Public Health England | Microbiology Services Colindale, Public Health England | |
| EPI607841 | HA | United Kingdom | 2015-Jan-12 | A/England/50380052/2015 | Microbiology Services Colindale, Public Health England | Microbiology Services Colindale, Public Health England | |
| EPI607849 | HA | United Kingdom | 2015-Jan-15 | A/England/72/2015 | Microbiology Services Colindale, Public Health England | Microbiology Services Colindale, Public Health England | |
| EPI607857 | HA | United Kingdom | 2015-Jan-16 | A/England/50400278/2015 | Microbiology Services Colindale, Public Health England | Microbiology Services Colindale, Public Health England | |
| EPI607865 | HA | United Kingdom | 2015-Jan-15 | A/England/50400282/2015 | Microbiology Services Colindale, Public Health England | Microbiology Services Colindale, Public Health England | |

| | | | | | | | |
|---|---|---|---|---|---|---|---|
| EPI607873 | HA | United Kingdom | 2015-Jan-15 | A/England/50400284/2015 | Microbiology Services Colindale, Public Health England | Microbiology Services Colindale, Public Health England | |
| EPI607881 | HA | United Kingdom | 2015-Jan-22 | A/England/87/2015 | Microbiology Services Colindale, Public Health England | Microbiology Services Colindale, Public Health England | |
| EPI607889 | HA | United Kingdom | 2015-Jan-26 | A/England/50520248/2015 | Microbiology Services Colindale, Public Health England | Microbiology Services Colindale, Public Health England | |
| EPI607899 | HA | United Kingdom | 2015-Jan-26 | A/England/105/2015 | Microbiology Services Colindale, Public Health England | Microbiology Services Colindale, Public Health England | |
| EPI607901 | HA | United Kingdom | 2015-Jan-26 | A/England/105/2015 | Microbiology Services Colindale, Public Health England | Microbiology Services Colindale, Public Health England | |
| EPI607914 | HA | United Kingdom | 2015-Jan-29 | A/England/127/2015 | Microbiology Services Colindale, Public Health England | Microbiology Services Colindale, Public Health England | |
| EPI607915 | HA | United Kingdom | 2015-Jan-29 | A/England/127/2015 | Microbiology Services Colindale, Public Health England | Microbiology Services Colindale, Public Health England | |
| EPI607930 | HA | United Kingdom | 2015-Jan-26 | A/England/112/2015 | Microbiology Services Colindale, Public Health England | Microbiology Services Colindale, Public Health England | |
| EPI607932 | HA | United Kingdom | 2015-Jan-26 | A/England/112/2015 | Microbiology Services Colindale, Public Health England | Microbiology Services Colindale, Public Health England | |
| EPI607945 | HA | United Kingdom | 2015-Jan-28 | A/England/113/2015 | Microbiology Services Colindale, Public Health England | Microbiology Services Colindale, Public Health England | |
| EPI607947 | HA | United Kingdom | 2015-Jan-28 | A/England/113/2015 | Microbiology Services Colindale, Public Health England | Microbiology Services Colindale, Public Health England | |
| EPI607960 | HA | United Kingdom | 2015-Jan-30 | A/England/114/2015 | Microbiology Services Colindale, Public Health England | Microbiology Services Colindale, Public Health England | |
| EPI607962 | HA | United Kingdom | 2015-Jan-30 | A/England/114/2015 | Microbiology Services Colindale, Public Health England | Microbiology Services Colindale, Public Health England | |
| EPI607975 | HA | United Kingdom | 2015-Feb-03 | A/England/50660076/2015 | Microbiology Services Colindale, Public Health England | Microbiology Services Colindale, Public Health England | |
| EPI607977 | HA | United Kingdom | 2015-Feb-03 | A/England/50660076/2015 | Microbiology Services Colindale, Public Health England | Microbiology Services Colindale, Public Health England | |
| EPI607990 | HA | United Kingdom | 2015-Feb-10 | A/England/50740149/2015 | Microbiology Services Colindale, Public Health England | Microbiology Services Colindale, Public Health England | |
| EPI607991 | HA | United Kingdom | 2015-Feb-10 | A/England/50740149/2015 | Microbiology Services Colindale, Public Health England | Microbiology Services Colindale, Public Health England | |
| EPI608006 | HA | United Kingdom | 2015-Feb-10 | A/England/50740161/2015 | Microbiology Services Colindale, Public Health England | Microbiology Services Colindale, Public Health England | |
| EPI608007 | HA | United Kingdom | 2015-Feb-10 | A/England/50740161/2015 | Microbiology Services Colindale, Public Health England | Microbiology Services Colindale, Public Health England | |
| EPI608022 | HA | United Kingdom | 2015-Feb-11 | A/England/50780065/2015 | Microbiology Services Colindale, Public Health England | Microbiology Services Colindale, Public Health England | Saito, Reiko; Takemae, Nobuhiro; Saito, Takehiko; Shobugawa, Yugo; Kondo, Hiroki; Hibino, Akinobu; Yadanar, Kyaw; Yi Yi, Myint; Khin Yi, Oo; Htay Htay, Tin |
| EPI608023 | HA | United Kingdom | 2015-Feb-11 | A/England/50780065/2015 | Microbiology Services Colindale, Public Health England | Microbiology Services Colindale, Public Health England | Saito, Reiko; Takemae, Nobuhiro; Saito, Takehiko; Shobugawa, Yugo; Kondo, Hiroki; Hibino, Akinobu; Yadanar, Kyaw; Yi Yi, Myint; Khin Yi, Oo; Htay Htay, Tin |
| EPI608038 | HA | United Kingdom | 2015-Feb-12 | A/England/166/2015 | Microbiology Services Colindale, Public Health England | Microbiology Services Colindale, Public Health England | Saito, Reiko; Takemae, Nobuhiro; Saito, Takehiko; Shobugawa, Yugo; Kondo, Hiroki; Hibino, |

| | | | | | | | |
|---|---|---|---|---|---|---|---|
| EPI608041 | HA | United Kingdom | 2015-Feb-12 | A/England/166/2015 | Microbiology Services Colindale, Public Health England | Microbiology Services Colindale, Public Health England | Saito, Reiko; Takemae, Nobuhiro; Saito, Takehiko; Shobugawa, Yugo; Kondo, Hiroki; Hibino, |
| EPI608051 | HA | United Kingdom | 2015-Feb-12 | A/England/50800025/2015 | Microbiology Services Colindale, Public Health England | Microbiology Services Colindale, Public Health England | |
| EPI608052 | HA | United Kingdom | 2015-Feb-12 | A/England/50800025/2015 | Microbiology Services Colindale, Public Health England | Microbiology Services Colindale, Public Health England | |
| EPI608067 | HA | United Kingdom | 2015-Feb-13 | A/England/169/2015 | Microbiology Services Colindale, Public Health England | Microbiology Services Colindale, Public Health England | |
| EPI608068 | HA | United Kingdom | 2015-Feb-13 | A/England/169/2015 | Microbiology Services Colindale, Public Health England | Microbiology Services Colindale, Public Health England | |
| EPI610547 | HA | Japan | 2015-Mar-03 | A/YAMAGATA/64/2015 | Yamagata Prefectural Institute of Public Health | National Institute of Infectious Diseases (NIID) | |
| EPI610549 | HA | Japan | 2015-Feb-02 | A/KANAGAWA/36/2015 | Kanagawa Prefectural Institute of Public Health | National Institute of Infectious Diseases (NIID) | |
| EPI610551 | HA | Japan | 2015-Jan-09 | A/MIYAGI/12/2015 | Miyagi Prefectural Institute of Public Health and Environment | National Institute of Infectious Diseases (NIID) | |
| EPI610553 | HA | Japan | 2015-Feb-01 | A/MIYAGI/21/2015 | Miyagi Prefectural Institute of Public Health and Environment | National Institute of Infectious Diseases (NIID) | Saito, Reiko; Takemae, Nobuhiro; Saito, Takehiko; Shobugawa, Yugo; Kondo, Hiroki; Hibino, Akinobu; Asou, Norichika; Kita, |
| EPI611225 | HA | United Kingdom | 2014-Sep-22 | A/England/43960141/2014 | Microbiology Services Colindale, Public Health England | Microbiology Services Colindale, Public Health England | Saito, Reiko; Takemae, Nobuhiro; Saito, Takehiko; Shobugawa, Yugo; Kondo, Hiroki; Hibino, Akinobu; Masaki, Hironori |
| EPI611249 | HA | United Kingdom | 2014-Sep-26 | A/England/44020584/2014 | Microbiology Services Colindale, Public Health England | Microbiology Services Colindale, Public Health England | Saito, Reiko; Takemae, Nobuhiro; Saito, Takehiko; Shobugawa, Yugo; Kondo, Hiroki; Hibino, Akinobu; Yadanar, Kyaw; Yi Yi, Myint; Khin Yi, Oo; Htay Htay, Tin |
| EPI611263 | HA | United Kingdom | 2014-Sep-12 | A/England/44080181/2014 | Microbiology Services Colindale, Public Health England | Microbiology Services Colindale, Public Health England | |
| EPI611271 | HA | United Kingdom | 2014-Aug-27 | A/Scotland/2/2014 | Microbiology Services Colindale, Public Health England | Microbiology Services Colindale, Public Health England | |
| EPI611279 | HA | United Kingdom | 2014-Oct-02 | A/England/44160198/2014 | Microbiology Services Colindale, Public Health England | Microbiology Services Colindale, Public Health England | |
| EPI611287 | HA | United Kingdom | 2014-Oct-02 | A/England/44160199/2014 | Microbiology Services Colindale, Public Health England | Microbiology Services Colindale, Public Health England | Saito, Reiko; Takemae, Nobuhiro; Saito, Takehiko; Shobugawa, Yugo; Kondo, Hiroki; Hibino, Akinobu; Yadanar, Kyaw; Yi Yi, Myint; Khin Yi, Oo; Htay Htay, Tin |
| EPI611295 | HA | United Kingdom | 2014-Oct-02 | A/England/511/2014 | Microbiology Services Colindale, Public Health England | Microbiology Services Colindale, Public Health England | Saito, Reiko; Takemae, Nobuhiro; Saito, Takehiko; Shobugawa, Yugo; Kondo, Hiroki; Hibino, Akinobu; Yadanar, Kyaw; Yi Yi, Myint; Khin Yi, Oo; Htay Htay, Tin |
| EPI611303 | HA | United Kingdom | 2014-Oct-02 | A/England/44160201/2014 | Microbiology Services Colindale, Public Health England | Microbiology Services Colindale, Public Health England | |
| EPI611311 | HA | United Kingdom | 2014-Oct-02 | A/England/44160202/2014 | Microbiology Services Colindale, Public Health England | Microbiology Services Colindale, Public Health England | |

| | | | | | | |
|---|---|---|---|---|---|---|
| EPI611319 | HA | United Kingdom | 2014-Oct-02 | A/England/512/2014 | Microbiology Services Colindale, Public Health England | Microbiology Services Colindale, Public Health England |
| EPI611327 | HA | United Kingdom | 2014-Oct-10 | A/England/44320792/2014 | Microbiology Services Colindale, Public Health England | Microbiology Services Colindale, Public Health England |
| EPI611335 | HA | United Kingdom | 2014-Oct-16 | A/England/44320793/2014 | Microbiology Services Colindale, Public Health England | Microbiology Services Colindale, Public Health England |
| EPI611343 | HA | United Kingdom | 2014-Oct-15 | A/England/44320794/2014 | Microbiology Services Colindale, Public Health England | Microbiology Services Colindale, Public Health England |
| EPI611351 | HA | United Kingdom | 2014-Oct-18 | A/England/44320795/2014 | Microbiology Services Colindale, Public Health England | Microbiology Services Colindale, Public Health England |
| EPI611359 | HA | United Kingdom | 2014-Oct-28 | A/England/44540273/2014 | Microbiology Services Colindale, Public Health England | Microbiology Services Colindale, Public Health England |
| EPI611367 | HA | United Kingdom | 2014-Nov-09 | A/England/44640016/2014 | Microbiology Services Colindale, Public Health England | Microbiology Services Colindale, Public Health England |
| EPI611375 | HA | United Kingdom | 2014-Nov-20 | A/England/525/2014 | Microbiology Services Colindale, Public Health England | Microbiology Services Colindale, Public Health England |
| EPI611383 | HA | United Kingdom | 2014-Nov-20 | A/England/44820080/2014 | Microbiology Services Colindale, Public Health England | Microbiology Services Colindale, Public Health England |
| EPI611391 | HA | United Kingdom | 2014-Nov-24 | A/England/44840003/2014 | Microbiology Services Colindale, Public Health England | Microbiology Services Colindale, Public Health England |
| EPI611399 | HA | United Kingdom | 2014-Nov-20 | A/England/528/2014 | Microbiology Services Colindale, Public Health England | Microbiology Services Colindale, Public Health England |
| EPI611407 | HA | United Kingdom | 2014-Nov-21 | A/England/44840067/2014 | Microbiology Services Colindale, Public Health England | Microbiology Services Colindale, Public Health England |
| EPI611415 | HA | United Kingdom | 2014-Nov-21 | A/England/529/2014 | Microbiology Services Colindale, Public Health England | Microbiology Services Colindale, Public Health England |
| EPI611423 | HA | United Kingdom | 2014-Nov-21 | A/England/44840069/2014 | Microbiology Services Colindale, Public Health England | Microbiology Services Colindale, Public Health England |
| EPI611431 | HA | United Kingdom | 2014-Nov-21 | A/England/44860850/2014 | Microbiology Services Colindale, Public Health England | Microbiology Services Colindale, Public Health England |
| EPI611440 | HA | United Kingdom | 2014-Nov-28 | A/England/44920021/2014 | Microbiology Services Colindale, Public Health England | Microbiology Services Colindale, Public Health England |
| EPI611448 | HA | United Kingdom | 2014-Nov-26 | A/England/530/2014 | Microbiology Services Colindale, Public Health England | Microbiology Services Colindale, Public Health England |
| EPI611456 | HA | United Kingdom | 2014-Dec-02 | A/England/44940038/2014 | Microbiology Services Colindale, Public Health England | Microbiology Services Colindale, Public Health England |
| EPI611464 | HA | United Kingdom | 2014-Dec-04 | A/England/45000006/2014 | Microbiology Services Colindale, Public Health England | Microbiology Services Colindale, Public Health England |
| EPI611472 | HA | United Kingdom | 2014-Dec-05 | A/England/45000008/2014 | Microbiology Services Colindale, Public Health England | Microbiology Services Colindale, Public Health England |
| EPI611480 | HA | United Kingdom | 2014-Dec-03 | A/England/45000010/2014 | Microbiology Services Colindale, Public Health England | Microbiology Services Colindale, Public Health England |
| EPI611488 | HA | United Kingdom | 2014-Dec-01 | A/England/45000025/2014 | Microbiology Services Colindale, Public Health England | Microbiology Services Colindale, Public Health England |
| EPI611496 | HA | United Kingdom | 2014-Dec-01 | A/England/533/2014 | Microbiology Services Colindale, Public Health England | Microbiology Services Colindale, Public Health England |
| EPI611504 | HA | United Kingdom | 2014-Nov-27 | A/England/45000028/2014 | Microbiology Services Colindale, Public Health England | Microbiology Services Colindale, Public Health England |
| EPI611512 | HA | United Kingdom | 2014-Dec-09 | A/England/45040065/2014 | Microbiology Services Colindale, Public Health England | Microbiology Services Colindale, Public Health England |

| | | | | | | |
|---|---|---|---|---|---|---|
| EPI611536 | HA | United Kingdom | 2014-Dec-08 | A/England/536/2014 | Microbiology Services Colindale, Public Health England | Microbiology Services Colindale, Public Health England |
| EPI611544 | HA | United Kingdom | 2014-Dec-09 | A/England/45080158/2014 | Microbiology Services Colindale, Public Health England | Microbiology Services Colindale, Public Health England |
| EPI611552 | HA | United Kingdom | 2014-Dec-10 | A/England/45080193/2014 | Microbiology Services Colindale, Public Health England | Microbiology Services Colindale, Public Health England |
| EPI611560 | HA | United Kingdom | 2014-Dec-10 | A/England/45080198/2014 | Microbiology Services Colindale, Public Health England | Microbiology Services Colindale, Public Health England |
| EPI611568 | HA | United Kingdom | 2014-Dec-11 | A/England/45080199/2014 | Microbiology Services Colindale, Public Health England | Microbiology Services Colindale, Public Health England |
| EPI611576 | HA | United Kingdom | 2014-Dec-08 | A/England/45080202/2014 | Microbiology Services Colindale, Public Health England | Microbiology Services Colindale, Public Health England |
| EPI611584 | HA | United Kingdom | 2014-Dec-08 | A/England/45080206/2014 | Microbiology Services Colindale, Public Health England | Microbiology Services Colindale, Public Health England |
| EPI611592 | HA | United Kingdom | 2014-Dec-11 | A/England/45100189/2014 | Microbiology Services Colindale, Public Health England | Microbiology Services Colindale, Public Health England |
| EPI611600 | HA | United Kingdom | 2014-Dec-10 | A/England/45100191/2014 | Microbiology Services Colindale, Public Health England | Microbiology Services Colindale, Public Health England |
| EPI611610 | HA | United Kingdom | 2014-Nov-26 | A/England/45100197/2014 | Microbiology Services Colindale, Public Health England | Microbiology Services Colindale, Public Health England |
| EPI611618 | HA | United Kingdom | 2014-Dec-16 | A/England/45140001/2014 | Microbiology Services Colindale, Public Health England | Microbiology Services Colindale, Public Health England |
| EPI611626 | HA | United Kingdom | 2014-Dec-15 | A/England/45140005/2014 | Microbiology Services Colindale, Public Health England | Microbiology Services Colindale, Public Health England |
| EPI611634 | HA | United Kingdom | 2014-Dec-12 | A/England/45140006/2014 | Microbiology Services Colindale, Public Health England | Microbiology Services Colindale, Public Health England |
| EPI611642 | HA | United Kingdom | 2014-Dec-18 | A/England/45160088/2014 | Microbiology Services Colindale, Public Health England | Microbiology Services Colindale, Public Health England |
| EPI611650 | HA | United Kingdom | 2014-Dec-17 | A/England/45160092/2014 | Microbiology Services Colindale, Public Health England | Microbiology Services Colindale, Public Health England |
| EPI611658 | HA | United Kingdom | 2014-Dec-16 | A/England/45160094/2014 | Microbiology Services Colindale, Public Health England | Microbiology Services Colindale, Public Health England |
| EPI611666 | HA | United Kingdom | 2014-Dec-15 | A/England/601/2014 | Microbiology Services Colindale, Public Health England | Microbiology Services Colindale, Public Health England |
| EPI611674 | HA | United Kingdom | 2014-Dec-16 | A/England/45180212/2014 | Microbiology Services Colindale, Public Health England | Microbiology Services Colindale, Public Health England |
| EPI611682 | HA | United Kingdom | 2014-Dec-17 | A/England/543/2014 | Microbiology Services Colindale, Public Health England | Microbiology Services Colindale, Public Health England |
| EPI611690 | HA | United Kingdom | 2014-Dec-17 | A/England/45180219/2014 | Microbiology Services Colindale, Public Health England | Microbiology Services Colindale, Public Health England |
| EPI611698 | HA | United Kingdom | 2014-Dec-17 | A/England/45180220/2014 | Microbiology Services Colindale, Public Health England | Microbiology Services Colindale, Public Health England |
| EPI611706 | HA | United Kingdom | 2014-Dec-18 | A/England/544/2014 | Microbiology Services Colindale, Public Health England | Microbiology Services Colindale, Public Health England |
| EPI611714 | HA | United Kingdom | 2014-Dec-16 | A/England/545/2014 | Microbiology Services Colindale, Public Health England | Microbiology Services Colindale, Public Health England |
| EPI611722 | HA | United Kingdom | 2014-Dec-17 | A/England/45180226/2014 | Microbiology Services Colindale, Public Health England | Microbiology Services Colindale, Public Health England |
| EPI611730 | HA | United Kingdom | 2014-Dec-19 | A/England/45180235/2014 | Microbiology Services Colindale, Public Health England | Microbiology Services Colindale, Public Health England |

| | | | | | | | |
|---|---|---|---|---|---|---|---|
| EPI611738 | HA | United Kingdom | 2014-Dec-17 | A/England/602/2014 | Microbiology Services Colindale, Public Health England | Microbiology Services Colindale, Public Health England | |
| EPI611746 | HA | United Kingdom | 2014-Dec-18 | A/England/45200044/2014 | Microbiology Services Colindale, Public Health England | Microbiology Services Colindale, Public Health England | |
| EPI611754 | HA | United Kingdom | 2014-Dec-18 | A/England/547/2014 | Microbiology Services Colindale, Public Health England | Microbiology Services Colindale, Public Health England | |
| EPI611762 | HA | United Kingdom | 2014-Dec-18 | A/England/45200047/2014 | Microbiology Services Colindale, Public Health England | Microbiology Services Colindale, Public Health England | |
| EPI611770 | HA | United Kingdom | 2014-Dec-17 | A/England/45200054/2014 | Microbiology Services Colindale, Public Health England | Microbiology Services Colindale, Public Health England | |
| EPI611778 | HA | United Kingdom | 2014-Dec-22 | A/England/575/2014 | Microbiology Services Colindale, Public Health England | Microbiology Services Colindale, Public Health England | |
| EPI611786 | HA | United Kingdom | 2014-Dec-19 | A/England/45220034/2014 | Microbiology Services Colindale, Public Health England | Microbiology Services Colindale, Public Health England | |
| EPI611794 | HA | United Kingdom | 2014-Dec-18 | A/England/556/2014 | Microbiology Services Colindale, Public Health England | Microbiology Services Colindale, Public Health England | Takashita,Emi; Fujisaki,Seiichiro; Shirakura,Masayuki; Watanabe,Shinji; Odagiri,Takato |
| EPI611802 | HA | United Kingdom | 2014-Dec-22 | A/England/45240246/2014 | Microbiology Services Colindale, Public Health England | Microbiology Services Colindale, Public Health England | Takashita,Emi; Fujisaki,Seiichiro; Shirakura,Masayuki; Watanabe,Shinji; Odagiri,Takato |
| EPI611810 | HA | United Kingdom | 2014-Dec-23 | A/England/558/2014 | Microbiology Services Colindale, Public Health England | Microbiology Services Colindale, Public Health England | |
| EPI611818 | HA | United Kingdom | 2014-Dec-18 | A/England/45240252/2014 | Microbiology Services Colindale, Public Health England | Microbiology Services Colindale, Public Health England | |
| EPI611826 | HA | United Kingdom | 2014-Dec-22 | A/England/45240260/2014 | Microbiology Services Colindale, Public Health England | Microbiology Services Colindale, Public Health England | |
| EPI611834 | HA | United Kingdom | 2014-Dec-22 | A/England/45240273/2014 | Microbiology Services Colindale, Public Health England | Microbiology Services Colindale, Public Health England | |
| EPI611842 | HA | United Kingdom | 2014-Dec-19 | A/England/603/2014 | Microbiology Services Colindale, Public Health England | Microbiology Services Colindale, Public Health England | |
| EPI611850 | HA | United Kingdom | 2014-Dec-23 | A/England/50100002/2014 | Microbiology Services Colindale, Public Health England | Microbiology Services Colindale, Public Health England | |
| EPI611858 | HA | United Kingdom | 2014-Dec-24 | A/England/50100063/2014 | Microbiology Services Colindale, Public Health England | Microbiology Services Colindale, Public Health England | |
| EPI611866 | HA | United Kingdom | 2014-Dec-24 | A/England/50100065/2014 | Microbiology Services Colindale, Public Health England | Microbiology Services Colindale, Public Health England | |
| EPI611874 | HA | United Kingdom | 2014-Dec-24 | A/England/563/2014 | Microbiology Services Colindale, Public Health England | Microbiology Services Colindale, Public Health England | |
| EPI611882 | HA | United Kingdom | 2014-Dec-29 | A/England/50120064/2014 | Microbiology Services Colindale, Public Health England | Microbiology Services Colindale, Public Health England | |
| EPI611890 | HA | United Kingdom | 2014-Dec-23 | A/England/564/2014 | Microbiology Services Colindale, Public Health England | Microbiology Services Colindale, Public Health England | |
| EPI611898 | HA | United Kingdom | 2014-Dec-29 | A/England/606/2014 | Microbiology Services Colindale, Public Health England | Microbiology Services Colindale, Public Health England | |
| EPI611906 | HA | United Kingdom | 2014-Dec-24 | A/England/50120073/2014 | Microbiology Services Colindale, Public Health England | Microbiology Services Colindale, Public Health England | |
| EPI611914 | HA | United Kingdom | 2014-Dec-23 | A/England/50120074/2014 | Microbiology Services Colindale, Public Health England | Microbiology Services Colindale, Public Health England | |
| EPI611922 | HA | United Kingdom | 2014-Dec-30 | A/England/50120128/2014 | Microbiology Services Colindale, Public Health England | Microbiology Services Colindale, Public Health England | |

| | | | | | | | |
|---|---|---|---|---|---|---|---|
| EPI611930 | HA | United Kingdom | 2014-Dec-23 | A/England/50120129/2014 | Microbiology Services Colindale, Public Health England | Microbiology Services Colindale, Public Health England | |
| EPI611938 | HA | United Kingdom | 2014-Dec-30 | A/England/50140078/2014 | Microbiology Services Colindale, Public Health England | Microbiology Services Colindale, Public Health England | |
| EPI611946 | HA | United Kingdom | 2014-Dec-29 | A/England/50140086/2014 | Microbiology Services Colindale, Public Health England | Microbiology Services Colindale, Public Health England | |
| EPI611954 | HA | United Kingdom | 2014-Dec-29 | A/England/50140112/2014 | Microbiology Services Colindale, Public Health England | Microbiology Services Colindale, Public Health England | |
| EPI611962 | HA | United Kingdom | 2014-Dec-29 | A/England/50180052/2014 | Microbiology Services Colindale, Public Health England | Microbiology Services Colindale, Public Health England | |
| EPI611970 | HA | United Kingdom | 2014-Dec-21 | A/England/50200102/2014 | Microbiology Services Colindale, Public Health England | Microbiology Services Colindale, Public Health England | |
| EPI611978 | HA | United Kingdom | 2014-Dec-31 | A/England/50200115/2014 | Microbiology Services Colindale, Public Health England | Microbiology Services Colindale, Public Health England | |
| EPI611986 | HA | United Kingdom | 2014-Dec-30 | A/England/50200119/2014 | Microbiology Services Colindale, Public Health England | Microbiology Services Colindale, Public Health England | |
| EPI611994 | HA | United Kingdom | 2014-Dec-29 | A/England/50200132/2014 | Microbiology Services Colindale, Public Health England | Microbiology Services Colindale, Public Health England | Chen,B.B.; Phuah,S.P.; Poh,M.K.; Chen,S.L.; Cui,L. |
| EPI612002 | HA | United Kingdom | 2014-Dec-29 | A/England/50200133/2014 | Microbiology Services Colindale, Public Health England | Microbiology Services Colindale, Public Health England | |
| EPI612010 | HA | United Kingdom | 2014-Dec-30 | A/England/50200134/2014 | Microbiology Services Colindale, Public Health England | Microbiology Services Colindale, Public Health England | |
| EPI612018 | HA | United Kingdom | 2014-Dec-31 | A/England/50200158/2014 | Microbiology Services Colindale, Public Health England | Microbiology Services Colindale, Public Health England | |
| EPI612026 | HA | United Kingdom | 2014-Dec-31 | A/England/50200190/2014 | Microbiology Services Colindale, Public Health England | Microbiology Services Colindale, Public Health England | |
| EPI612034 | HA | United Kingdom | 2014-Jan-02 | A/England/50220066/2014 | Microbiology Services Colindale, Public Health England | Microbiology Services Colindale, Public Health England | |
| EPI612042 | HA | United Kingdom | 2015-Jan-06 | A/England/50220076/2015 | Microbiology Services Colindale, Public Health England | Microbiology Services Colindale, Public Health England | |
| EPI612050 | HA | United Kingdom | 2015-Jan-06 | A/England/50220894/2015 | Microbiology Services Colindale, Public Health England | Microbiology Services Colindale, Public Health England | |
| EPI612058 | HA | United Kingdom | 2015-Jan-06 | A/England/50220895/ | Microbiology Services Colindale, Public Health England | Microbiology Services Colindale, Public Health England | |
| EPI612066 | HA | United Kingdom | 2015-Jan-13 | A/England/50320294/2015 | Microbiology Services Colindale, Public Health England | Microbiology Services Colindale, Public Health England | |
| EPI612074 | HA | United Kingdom | 2015-Jan-06 | A/England/50360756/2015 | Microbiology Services Colindale, Public Health England | Microbiology Services Colindale, Public Health England | |
| EPI612082 | HA | United Kingdom | 2015-Jan-06 | A/England/50380175/2015 | Microbiology Services Colindale, Public Health England | Microbiology Services Colindale, Public Health England | |
| EPI612090 | HA | United Kingdom | 2015-Jan-15 | A/England/50400281/2015 | Microbiology Services Colindale, Public Health England | Microbiology Services Colindale, Public Health England | |
| EPI612098 | HA | United Kingdom | 2015-Jan-12 | A/England/77/2015 | Microbiology Services Colindale, Public Health England | Microbiology Services Colindale, Public Health England | Takashita,Emi; Fujisaki,Seiichiro; Shirakura,Masayuki; Watanabe,Shinji; Odagiri,Takato |
| EPI612106 | HA | United Kingdom | 2015-Jan-30 | A/England/50620849/2015 | Microbiology Services Colindale, Public Health England | Microbiology Services Colindale, Public Health England | |
| EPI612114 | HA | United Kingdom | 2015-Feb-03 | A/England/50660120/2015 | Microbiology Services Colindale, Public Health England | Microbiology Services Colindale, Public Health England | |
| EPI612124 | HA | United Kingdom | 2015-Feb-07 | A/England/50880445/2015 | Microbiology Services Colindale, Public Health England | Microbiology Services Colindale, Public Health England | |

| | | | | | | | |
|---|---|---|---|---|---|---|---|
| EPI612132 | HA | United Kingdom | 2015-Feb-09 | A/England/50880448/2015 | Microbiology Services Colindale, Public Health England | Microbiology Services Colindale, Public Health England | |
| EPI612140 | HA | United Kingdom | 2015-Feb-11 | A/England/50960080/2015 | Microbiology Services Colindale, Public Health England | Microbiology Services Colindale, Public Health England | |
| EPI612148 | HA | United Kingdom | 2015-Jan-27 | A/England/50960083/2015 | Microbiology Services Colindale, Public Health England | Microbiology Services Colindale, Public Health England | |
| EPI612156 | HA | United Kingdom | 2015-Mar-03 | A/England/51060001/2015 | Microbiology Services Colindale, Public Health England | Microbiology Services Colindale, Public Health England | |
| EPI612164 | HA | United Kingdom | 2015-Mar-07 | A/England/51200124/2015 | Microbiology Services Colindale, Public Health England | Microbiology Services Colindale, Public Health England | |
| EPI612172 | HA | United Kingdom | 2015-Apr-01 | A/England/51440624/2015 | Microbiology Services Colindale, Public Health England | Microbiology Services Colindale, Public Health England | |
| EPI612180 | HA | United Kingdom | 2015-Apr-01 | A/England/51440626/2015 | Microbiology Services Colindale, Public Health England | Microbiology Services Colindale, Public Health England | |
| EPI612188 | HA | United Kingdom | 2015-Apr-01 | A/England/51440629/2015 | Microbiology Services Colindale, Public Health England | Microbiology Services Colindale, Public Health England | Deng,Y-M.; Iannello,P.; Spirason,N.; Jelley,L.; Lau,H.; |
| EPI612196 | HA | United States | 2015-Jan-30 | A/Arizona/01/2015 | Arizona Department of Health Services | Centers for Disease Control and Prevention | Deng,Y-M.; Iannello,P.; Spirason,N.; Jelley,L.; Lau,H.; |
| EPI612204 | HA | United States | 2015-Jan-26 | A/Wisconsin/27/2015 | Marshfield Clinic Research Foundation | Centers for Disease Control and Prevention | |
| EPI612222 | HA | United States | 2014-Dec-06 | A/Georgia/29/2014 | Georgia Public Health Laboratory | Centers for Disease Control and Prevention | |
| EPI612230 | HA | United States | 2015-Jan-02 | A/Georgia/07/2015 | Georgia Public Health Laboratory | Centers for Disease Control and Prevention | |
| EPI612238 | HA | United States | 2014-Dec-10 | A/Louisiana/39/2014 | Louisiana Department of Health and Hospitals | Centers for Disease Control and Prevention | |
| EPI612248 | HA | United States | 2014-Dec-23 | A/Texas/94/2014 | Texas Childrens Hospital | Centers for Disease Control and Prevention | |
| EPI612256 | HA | Tanzania, United Republic of | 2014-Dec-01 | A/Tanzania/587/2014 | National Influenza Lab | Centers for Disease Control and Prevention | |
| EPI612270 | HA | United States | 2015-Jan-19 | A/Michigan/21/2015 | University of Michigan SPH EPID | Centers for Disease Control and Prevention | |
| EPI612279 | HA | United States | 2015-Jan-16 | A/Michigan/22/2015 | University of Michigan SPH EPID | Centers for Disease Control and Prevention | |
| EPI612287 | HA | Vietnam | 2014-Dec-15 | A/Vietnam/629/2014 | National Institute of Hygiene and Epidemiology | Centers for Disease Control and Prevention | |
| EPI612295 | HA | United States | 2015-Jan-02 | A/Idaho/11/2015 | State of Idaho Bureau of Laboratories | Centers for Disease Control and Prevention | |
| EPI612315 | HA | United States | 2015-Jan-28 | A/Washington/29/2015 | Marshfield Clinic Research Foundation | Centers for Disease Control and Prevention | |
| EPI612323 | HA | United States | 2014-Dec-17 | A/Maryland/53/2014 | Maryland Department of Health and Mental Hygiene | Centers for Disease Control and Prevention | |
| EPI612331 | HA | United States | 2015-Jan-26 | A/Alabama/02/2015 | ADPH Bureau of Clinical Laboratories | Centers for Disease Control and Prevention | |
| EPI612335 | HA | United States | 2014-Dec-01 | A/Kentucky/30/2014 | Kentucky Division of Laboratory Services | Centers for Disease Control and Prevention | |
| EPI612343 | HA | United States | 2015-Jan-30 | A/Washington/30/2015 | Marshfield Clinic Research Foundation | Centers for Disease Control and Prevention | |
| EPI612358 | HA | United States | 2014-Dec-03 | A/Georgia/28/2014 | Georgia Public Health Laboratory | Centers for Disease Control and Prevention | |

| | | | | | | | |
|---|---|---|---|---|---|---|---|
| EPI612373 | HA | United States | 2015-Jan-21 | A/Indiana/06/2015 | Indiana State Department of Health Laboratories | Centers for Disease Control and Prevention | |
| EPI612381 | HA | United States | 2014-Dec-17 | A/Georgia/38/2014 | Georgia Public Health Laboratory | Centers for Disease Control and Prevention | |
| EPI612389 | HA | United States | 2015-Feb-09 | A/North Carolina/23/2015 | North Carolina State Laboratory of Public Health | Centers for Disease Control and Prevention | |
| EPI612397 | HA | United States | 2014-Dec-23 | A/Washington/60/2014 | Washington State Public Health Laboratory | Centers for Disease Control and Prevention | |
| EPI612406 | HA | United States | 2014-Dec-20 | A/Vermont/22/2014 | Vermont Department of Health Laboratory | Centers for Disease Control and Prevention | |
| EPI612414 | HA | United States | 2014-Dec-15 | A/Georgia/45/2014 | Georgia Public Health Laboratory | Centers for Disease Control and Prevention | |
| EPI612429 | HA | United States | 2015-Jan-08 | A/Pennsylvania/10/2015 | UPMC-CLB Dept of Microbiology | Centers for Disease Control and Prevention | |
| EPI612437 | HA | United States | 2015-Jan-09 | A/Pennsylvania/07/2015 | UPMC-CLB Dept of Microbiology | Centers for Disease Control and Prevention | |
| EPI612445 | HA | United States | 2015-Mar-28 | A/Texas/43/2015 | Baylor College of Medicine | Centers for Disease Control and Prevention | |
| EPI612454 | HA | United States | 2014-Dec-26 | A/Texas/95/2014 | Texas Childrens Hospital | Centers for Disease Control and Prevention | |
| EPI612462 | HA | United States | 2014-Jan-01 | A/Georgia/27/2014 | Georgia Public Health Laboratory | Centers for Disease Control and Prevention | |
| EPI614429 | HA | Hong Kong (SAR) | 2014-Jan-01 | A/Hong Kong/7127/2014 | Crick Worldwide Influenza Centre | Centers for Disease Control and Prevention | |
| EPI614437 | HA | Hong Kong (SAR) | 2014-Jan-01 | A/Hong Kong/4801/2014 | Crick Worldwide Influenza Centre | Centers for Disease Control and Prevention | |
| EPI614444 | HA | Switzerland | 2013-Dec-06 | A/Switzerland/9715293/2013 | Crick Worldwide Influenza Centre | Centers for Disease Control and Prevention | |
| EPI614462 | HA | Australia | 2014-Jun-29 | A/South Australia/55/2014 | WHO Collaborating Centre for Reference and Research on Influenza | Centers for Disease Control and Prevention | |
| EPI614775 | HA | Kazakhstan | 2015-Jan-09 | A/Kazakhstan/159/2015 | CSEE | Centers for Disease Control and Prevention | Deng,Y-M.; Iannello,P.; Spirason,N.; Jelley,L.; Lau,H.; |
| EPI614783 | HA | Congo, the Democatic Republic of | 2014-Dec-11 | A/Congo/2495/2014 | INRB Service de Virologie | Centers for Disease Control and Prevention | Deng,Y-M.; Iannello,P.; Spirason,N.; Jelley,L.; Lau,H.; Komadina,N. |
| EPI614791 | HA | Kazakhstan | 2015-Jan-26 | A/Kazakhstan/184/2015 | CSEE | Centers for Disease Control and Prevention | Deng,Y-M.; Iannello,P.; Spirason,N.; Jelley,L.; Lau,H.; |
| EPI614799 | HA | Kazakhstan | 2015-Jan-13 | A/Kazakhstan/266/2015 | CSEE | Centers for Disease Control and Prevention | |
| EPI614807 | HA | Kazakhstan | 2015-Jan-29 | A/Kazakhstan/166/2015 | CSEE | Centers for Disease Control and Prevention | |
| EPI614815 | HA | Kazakhstan | 2015-Feb-06 | A/Kazakhstan/291/2015 | CSEE | Centers for Disease Control and Prevention | |
| EPI614823 | HA | Kazakhstan | 2015-Jan-14 | A/Kazakhstan/161/2015 | CSEE | Centers for Disease Control and Prevention | |
| EPI614831 | HA | Kazakhstan | 2015-Jan-23 | A/Kazakhstan/158/2015 | CSEE | Centers for Disease Control and Prevention | |
| EPI614839 | HA | Kazakhstan | 2015-Jan-21 | A/Kazakhstan/274/2015 | CSEE | Centers for Disease Control and Prevention | Li, J.; Shao, T.J.; Yu, X.F.; Pu, X.Y.; Pan, J.C. |

| | | | | | | | |
|---|---|---|---|---|---|---|---|
| EPI622002 | HA | United States | 2015-Mar-15 | A/Washington/31/2015 | Washington State Public Health Laboratory | Centers for Disease Control and Prevention | Li, J.; Shao, T.J.; Yu, X.F.; Pu, X.Y.; Pan, J.C. |
| EPI622010 | HA | Kazakhstan | 2014-Nov-29 | A/Kazakhstan/194/2014 | CSEE | Centers for Disease Control and Prevention | Li, J.; Shao, T.J.; Yu, X.F.; Pu, X.Y.; Pan, J.C. |
| EPI622023 | HA | United States | 2015-Feb-05 | A/South Carolina/07/2015 | South Carolina Department of Health | Centers for Disease Control and Prevention | Li, J.; Shao, T.J.; Yu, X.F.; Pu, X.Y.; Pan, J.C. |
| EPI622031 | HA | United States | 2015-Feb-19 | A/California/47/2015 | California Department of Health Services | Centers for Disease Control and Prevention | Li, J.; Shao, T.J.; Yu, X.F.; Pu, X.Y.; Pan, J.C. |
| EPI622039 | HA | United States | 2015-Mar-03 | A/California/24/2015 | California Department of Health Services | Centers for Disease Control and Prevention | Li, J.; Shao, T.J.; Yu, X.F.; Pu, X.Y.; Pan, J.C. |
| EPI622058 | HA | Paraguay | 2014-Aug-29 | A/Paraguay/0097/2014 | Central Laboratory of Public Health | Centers for Disease Control and Prevention | Li, J.; Shao, T.J.; Yu, X.F.; Pu, X.Y.; Pan, J.C. |
| EPI622073 | HA | United States | 2015-Mar-05 | A/Iowa/11/2015 | Iowa State Hygienic Laboratory | Centers for Disease Control and Prevention | Li, J.; Shao, T.J.; Yu, X.F.; Pu, X.Y.; Pan, J.C. |
| EPI622096 | HA | United States | 2015-Feb-09 | A/Idaho/06/2015 | State of Idaho Bureau of Laboratories | Centers for Disease Control and Prevention | |
| EPI622100 | HA | United States | 2015-Jan-18 | A/New Mexico/09/2015 | New Mexico Department of Health | Centers for Disease Control and Prevention | |
| EPI622109 | HA | United States | 2015-Mar-04 | A/Oregon/06/2015 | Oregon Public Health Laboratory | Centers for Disease Control and Prevention | |
| EPI622119 | HA | United States | 2015-Mar-03 | A/New Jersey/16/2015 | New Jersey Department of Health & Senior Services | Centers for Disease Control and Prevention | |
| EPI622142 | HA | United States | 2015-Feb-18 | A/Nevada/19/2015 | Southern Nevada Public Health Lab | Centers for Disease Control and Prevention | |
| EPI622155 | HA | United States | 2015-Mar-12 | A/Rhode Island/15/2015 | Rhode Island Department of Health | Centers for Disease Control and Prevention | |
| EPI622163 | HA | United States | 2015-Mar-02 | A/Maine/16/2015 | Maine Health and Environmental Testing Laboratory | Centers for Disease Control and Prevention | |
| EPI622180 | HA | Russian Federation | 2014-Dec-22 | A/Chita/14/2014 | State Research Center of Virology and Biotechnology (VECTOR) | Centers for Disease Control and Prevention | |
| EPI622188 | HA | United States | 2015-Jan-02 | A/Georgia/05/2015 | Georgia Public Health Laboratory | Centers for Disease Control and Prevention | |
| EPI622196 | HA | Russian Federation | 2014-Dec-25 | A/Chita/12/2014 | State Research Center of Virology and Biotechnology (VECTOR) | Centers for Disease Control and Prevention | |
| EPI622204 | HA | United States | 2015-Jan-02 | A/Georgia/06/2015 | Georgia Public Health Laboratory | Centers for Disease Control and Prevention | |
| EPI622218 | HA | Russian Federation | 2015-Jan-20 | A/Khabarovsk/06/2015 | State Research Center of Virology and Biotechnology (VECTOR) | Centers for Disease Control and Prevention | |
| EPI622226 | HA | United States | 2015-Feb-05 | A/Florida/31/2015 | Florida Department of Health-Tampa | Centers for Disease Control and Prevention | |
| EPI622234 | HA | Russian Federation | 2015-Jan-16 | A/Khabarovsk/05/2015 | State Research Center of Virology and Biotechnology (VECTOR) | Centers for Disease Control and Prevention | |
| EPI622242 | HA | Russian Federation | 2015-Jan-20 | A/Khabarovsk/07/2015 | State Research Center of Virology and Biotechnology (VECTOR) | Centers for Disease Control and Prevention | |
| EPI622250 | HA | United States | 2015-Mar-08 | A/Vermont/06/2015 | Vermont Department of Health Laboratory | Centers for Disease Control and Prevention | |
| EPI622272 | HA | Lao, People's Democratic Republic | 2014-Nov-21 | A/Laos/732/2014 | National Center for Laboratory and Epidemiology | Centers for Disease Control and Prevention | |
| EPI622303 | HA | United States | 2015-Feb-23 | A/New Jersey/05/2015 | New Jersey Department of Health & Senior Services | Centers for Disease Control and Prevention | |

| | | | | | | | |
|---|---|---|---|---|---|---|---|
| EPI622322 | HA | United States | 2014-Nov-20 | A/Delaware/25/2014 | Delaware Public Health Lab | Centers for Disease Control and Prevention | |
| EPI622335 | HA | United States | 2015-Jan-01 | A/Florida/41/2015 | Florida Department of Health-Jacksonville | Centers for Disease Control and Prevention | |
| EPI622345 | HA | United States | 2015-Feb-03 | A/Iowa/09/2015 | Iowa State Hygienic Laboratory | Centers for Disease Control and Prevention | |
| EPI622353 | HA | United States | 2015-Jan-11 | A/New Hampshire/09/2015 | New Hampshire Public Health Laboratories | Centers for Disease Control and Prevention | |
| EPI622361 | HA | United States | 2015-Jan-31 | A/Texas/56/2015 | Baylor College of Medicine | Centers for Disease Control and Prevention | |
| EPI622424 | HA | United States | 2015-Mar-02 | A/Rhode Island/13/2015 | Rhode Island Department of Health | Centers for Disease Control and Prevention | |
| EPI623426 | HA | United States | 2014-Nov-22 | A/Wisconsin/64/2014 | Wisconsin State Laboratory of Hygiene | Centers for Disease Control and Prevention | |
| EPI623465 | HA | United States | 2014-Oct-10 | A/Washington/20/2014 | Washington State Public Health Laboratory | Centers for Disease Control and Prevention | |
| EPI623487 | HA | Canada | 2014-Nov-18 | A/Alberta/RV3588/2014 | National Microbiology Laboratory, Health Canada | Centers for Disease Control and Prevention | |
| EPI624003 | HA | United States | 2015-Apr-03 | A/Rhode Island/24/2015 | Rhode Island Department of Health | Centers for Disease Control and Prevention | |
| EPI624011 | HA | United States | 2015-Apr-14 | A/New Jersey/49/2015 | New Jersey Department of Health & Senior Services | Centers for Disease Control and Prevention | |
| EPI624019 | HA | United States | 2015-Apr-04 | A/Wisconsin/36/2015 | Wisconsin State Laboratory of Hygiene | Centers for Disease Control and Prevention | |
| EPI624027 | HA | United States | 2015-Mar-05 | A/South Dakota/15/2015 | South Dakota Public Health Lab | Centers for Disease Control and Prevention | |
| EPI624035 | HA | Congo, the Democatic Republic of | 2014-Dec-22 | A/Congo/2588/2014 | INRB Service de Virologie | Centers for Disease Control and Prevention | |
| EPI624043 | HA | United States | 2015-Feb-03 | A/South Dakota/05/2015 | South Dakota Public Health Lab | Centers for Disease Control and Prevention | |
| EPI624059 | HA | United States | 2015-Mar-11 | A/South Dakota/34/2015 | South Dakota Public Health Lab | Centers for Disease Control and Prevention | |
| EPI624067 | HA | United States | 2015-Feb-27 | A/South Dakota/14/2015 | South Dakota Public Health Lab | Centers for Disease Control and Prevention | |
| EPI624075 | HA | United States | 2015-Feb-13 | A/South Dakota/30/2015 | South Dakota Public Health Lab | Centers for Disease Control and Prevention | |
| EPI624083 | HA | United States | 2015-Apr-26 | A/Florida/49/2015 | Florida Department of Health-Jacksonville | Centers for Disease Control and Prevention | |
| EPI624102 | HA | United States | 2015-Feb-24 | A/South Dakota/29/2015 | South Dakota Public Health Lab | Centers for Disease Control and Prevention | |
| EPI624110 | HA | United States | 2015-Jan-29 | A/South Dakota/25/2015 | South Dakota Public Health Lab | Centers for Disease Control and Prevention | |
| EPI624118 | HA | United States | 2015-Mar-14 | A/South Dakota/35/2015 | South Dakota Public Health Lab | Centers for Disease Control and Prevention | |
| EPI624127 | HA | United States | 2015-May-06 | A/Washington/37/2015 | Washington State Public Health Laboratory | Centers for Disease Control and Prevention | |
| EPI624135 | HA | United States | 2015-Mar-14 | A/Idaho/17/2015 | State of Idaho Bureau of Laboratories | Centers for Disease Control and Prevention | Galiano,M. |
| EPI624143 | HA | United States | 2015-Mar-18 | A/New Mexico/18/2015 | New Mexico Department of Health | Centers for Disease Control and Prevention | Galiano,M. |

| | | | | | | |
|---|---|---|---|---|---|---|
| EPI624158 | HA | United States | 2015-Mar-08 | A/Texas/61/2015 | Texas Department of State Health Services-Laboratory Services | Centers for Disease Control and Prevention |
| EPI624166 | HA | United States | 2015-Apr-08 | A/Indiana/14/2015 | Indiana State Department of Health Laboratories | Centers for Disease Control and Prevention |
| EPI624174 | HA | United States | 2015-Apr-18 | A/Alaska/09/2015 | Alaska State Virology Lab | Centers for Disease Control and Prevention |
| EPI624192 | HA | United States | 2015-Apr-14 | A/Idaho/23/2015 | State of Idaho Bureau of Laboratories | Centers for Disease Control and Prevention |
| EPI624200 | HA | Congo, the Democatic Republic of | 2014-Dec-15 | A/Congo/2526/2014 | INRB Service de Virologie | Centers for Disease Control and Prevention |
| EPI624209 | HA | Congo, the Democatic Republic of | 2014-Dec-29 | A/Congo/2603/2014 | INRB Service de Virologie | Centers for Disease Control and Prevention |
| EPI624217 | HA | United States | 2015-Apr-06 | A/Nevada/36/2015 | Southern Nevada Public Health Lab | Centers for Disease Control and Prevention |
| EPI624225 | HA | United States | 2015-Apr-14 | A/Arizona/07/2015 | Arizona Department of Health Services | Centers for Disease Control and Prevention |
| EPI624233 | HA | United States | 2015-Apr-14 | A/Hawaii/37/2015 | State of Hawaii Department of Health | Centers for Disease Control and Prevention |
| EPI624241 | HA | United States | 2015-Apr-28 | A/California/82/2015 | California Department of Health Services | Centers for Disease Control and Prevention |
| EPI624249 | HA | United States | 2015-Apr-13 | A/Massachusetts/19/2015 | Massachusetts Department of Public Health | Centers for Disease Control and Prevention |
| EPI624257 | HA | United States | 2015-Apr-12 | A/Alaska/10/2015 | Alaska State Virology Lab | Centers for Disease Control and Prevention |
| EPI624265 | HA | United States | 2015-Apr-01 | A/Minnesota/25/2015 | Minnesota Department of Health | Centers for Disease Control and Prevention |
| EPI624273 | HA | United States | 2015-Apr-01 | A/New Mexico/19/2015 | New Mexico Department of Health | Centers for Disease Control and Prevention |
| EPI624281 | HA | United States | 2015-Apr-18 | A/Florida/45/2015 | Florida Department of Health-Jacksonville | Centers for Disease Control and Prevention |
| EPI624296 | HA | United States | 2015-May-18 | A/Minnesota/30/2015 | Minnesota Department of Health | Centers for Disease Control and Prevention |
| EPI624304 | HA | United States | 2015-Apr-19 | A/Hawaii/33/2015 | State of Hawaii Department of Health | Centers for Disease Control and Prevention |
| EPI624312 | HA | United States | 2015-Jun-03 | A/Florida/50/2015 | Florida Department of Health-Tampa | Centers for Disease Control and Prevention |
| EPI624320 | HA | Congo, the Democatic Republic of | 2014-Dec-22 | A/Congo/2586/2014 | INRB Service de Virologie | Centers for Disease Control and Prevention |
| EPI624328 | HA | United States | 2015-Apr-21 | A/Wisconsin/66/2015 | Wisconsin State Laboratory of Hygiene | Centers for Disease Control and Prevention |
| EPI624336 | HA | United States | 2015-Apr-21 | A/Minnesota/28/2015 | Minnesota Department of Health | Centers for Disease Control and Prevention |
| EPI624344 | HA | United States | 2015-Mar-31 | A/South Dakota/11/2015 | South Dakota Public Health Lab | Centers for Disease Control and Prevention |
| EPI624358 | HA | United States | 2015-Apr-12 | A/Minnesota/27/2015 | Minnesota Department of Health | Centers for Disease Control and Prevention |
| EPI624366 | HA | United States | 2015-Jan-05 | A/South Dakota/01/2015 | South Dakota Public Health Lab | Centers for Disease Control and Prevention |

| | | | | | | |
|---|---|---|---|---|---|---|
| EPI624375 | HA | United States | 2015-May-16 | A/Minnesota/29/2015 | Minnesota Department of Health | Centers for Disease Control and Prevention |
| EPI624949 | HA | Kazakhstan | 2015-Feb-18 | A/Kazakhstan/1075/2015 | CSEE | Centers for Disease Control and Prevention |
| EPI624957 | HA | Kazakhstan | 2015-Feb-16 | A/Kazakhstan/1060/2015 | CSEE | Centers for Disease Control and Prevention |
| EPI624965 | HA | Kazakhstan | 2015-Feb-09 | A/Kazakhstan/1046/2015 | CSEE | Centers for Disease Control and Prevention |
| EPI624980 | HA | Kazakhstan | 2015-Jan-16 | A/Kazakhstan/123/2015 | CSEE | Centers for Disease Control and Prevention |
| EPI624988 | HA | Kazakhstan | 2015-Jan-27 | A/Kazakhstan/183/2015 | CSEE | Centers for Disease Control and Prevention |
| EPI624996 | HA | Ukraine | 2015-Apr-07 | A/Ternopil/407/2015 | Institute of Epidemiology and Infectious Diseases AMS of Ukraine | Centers for Disease Control and Prevention |
| EPI625004 | HA | Ukraine | 2015-Apr-15 | A/Dnipropetrovsk/442/2015 | Institute of Epidemiology and Infectious Diseases AMS of Ukraine | Centers for Disease Control and Prevention |
| EPI625012 | HA | Ukraine | 2015-Apr-14 | A/Khmelnitsky/459/2015 | Institute of Epidemiology and Infectious Diseases AMS of Ukraine | Centers for Disease Control and Prevention |
| EPI625020 | HA | Kazakhstan | 2015-Feb-23 | A/Kazakhstan/1099/2015 | CSEE | Centers for Disease Control and Prevention |
| EPI625028 | HA | Ukraine | 2015-Apr-14 | A/Khmelnitsky/464/2015 | Institute of Epidemiology and Infectious Diseases AMS of Ukraine | Centers for Disease Control and Prevention |
| EPI626308 | HA | Nigeria | 2014-Apr-22 | A/Nigeria/942/2014 | National Influenza Reference Laboratory | Centers for Disease Control and Prevention |
| EPI626316 | HA | Nigeria | 2014-Jun-10 | A/Nigeria/120/2014 | National Influenza Reference Laboratory | Centers for Disease Control and Prevention |
| EPI626324 | HA | United States | 2015-Mar-05 | A/Massachusetts/16/2015 | Massachusetts Department of Public Health | Centers for Disease Control and Prevention |
| EPI626344 | HA | Kazakhstan | 2015-Jan-12 | A/Kazakhstan/873/2015 | CSEE | Centers for Disease Control and Prevention |
| EPI626352 | HA | Kazakhstan | 2015-Feb-02 | A/Kazakhstan/286/2015 | CSEE | Centers for Disease Control and Prevention |
| EPI626360 | HA | Kazakhstan | 2015-Jan-07 | A/Kazakhstan/112/2015 | CSEE | Centers for Disease Control and Prevention |
| EPI626375 | HA | Russian Federation | 2015-Jan-17 | A/St Petersburg/03/2015 | Russian Academy of Medical Sciences | Centers for Disease Control and Prevention |
| EPI626383 | HA | Russian Federation | 2015-Jan-26 | A/St Petersburg/22/2015 | Russian Academy of Medical Sciences | Centers for Disease Control and Prevention |
| EPI626391 | HA | Kazakhstan | 2015-Feb-09 | A/Kazakhstan/341/2015 | CSEE | Centers for Disease Control and Prevention |
| EPI626406 | HA | United States | 2015-Jan-06 | A/South Dakota/20/2015 | South Dakota Public Health Lab | Centers for Disease Control and Prevention |
| EPI626414 | HA | Kazakhstan | 2015-Jan-13 | A/Kazakhstan/122/2015 | CSEE | Centers for Disease Control and Prevention |
| EPI626422 | HA | Ukraine | 2015-Feb-27 | A/Ukraine/151/2015 | Institute of Epidemiology and Infectious Diseases AMS of Ukraine | Centers for Disease Control and Prevention |
| EPI626430 | HA | Kazakhstan | 2015-Jan-13 | A/Kazakhstan/113/2015 | CSEE | Centers for Disease Control and Prevention |
| EPI626445 | HA | Kazakhstan | 2015-Feb-14 | A/Kazakhstan/352/2015 | CSEE | Centers for Disease Control and Prevention |

| | | | | | | | |
|---|---|---|---|---|---|---|---|
| EPI626453 | HA | United States | 2015-Mar-18 | A/Washington/35/2015 | Washington State Public Health Laboratory | Centers for Disease Control and Prevention | |
| EPI626461 | HA | United States | 2015-Mar-16 | A/Washington/34/2015 | Washington State Public Health Laboratory | Centers for Disease Control and Prevention | |
| EPI626480 | HA | Ukraine | 2015-Feb-25 | A/Kharkov/241/2015 | Institute of Epidemiology and Infectious Diseases AMS of Ukraine | Centers for Disease Control and Prevention | |
| EPI626488 | HA | United States | 2015-Mar-03 | A/California/78/2015 | California Department of Health Services | Centers for Disease Control and Prevention | |
| EPI626496 | HA | United States | 2015-Mar-07 | A/California/81/2015 | California Department of Health Services | Centers for Disease Control and Prevention | |
| EPI626504 | HA | United States | 2015-Jan-11 | A/Minnesota/05/2015 | Minnesota Department of Health | Centers for Disease Control and Prevention | |
| EPI626512 | HA | United States | 2015-Mar-02 | A/Delaware/35/2015 | Delaware Public Health Lab | Centers for Disease Control and Prevention | |
| EPI626520 | HA | United States | 2015-Mar-28 | A/Hawaii/27/2015 | State of Hawaii Department of Health | Centers for Disease Control and Prevention | |
| EPI626533 | HA | United States | 2015-Mar-12 | A/Washington/33/2015 | Washington State Public Health Laboratory | Centers for Disease Control and Prevention | |
| EPI626541 | HA | United States | 2015-Mar-14 | A/Idaho/17/2015 | State of Idaho Bureau of Laboratories | Centers for Disease Control and Prevention | |
| EPI626549 | HA | Nigeria | 2014-Jun-02 | A/Nigeria/71/2014 | National Influenza Reference Laboratory | Centers for Disease Control and Prevention | |
| EPI626557 | HA | Nigeria | 2014-May-14 | A/Nigeria/928/2014 | National Influenza Reference Laboratory | Centers for Disease Control and Prevention | |
| EPI626565 | HA | United States | 2015-Mar-02 | A/Maryland/12/2015 | Maryland Department of Health and Mental Hygiene | Centers for Disease Control and Prevention | |
| EPI626573 | HA | United Kingdom | 2014-Aug-24 | A/ENG/507/2014 | Microbiology Services Colindale, Public Health England | Microbiology Services Colindale, Public Health England | |
| EPI626581 | HA | United Kingdom | 2014-Sep-24 | A/ENG/44060047/2014 | Microbiology Services Colindale, Public Health England | Microbiology Services Colindale, Public Health England | Takashita,Emi; Fujisaki,Seiichiro; Shirakura,Masayuki; Watanabe,Shinji; Odagiri,Takato |
| EPI626589 | HA | United Kingdom | 2014-Nov-20 | A/ENG/44760132/2014 | Microbiology Services Colindale, Public Health England | Microbiology Services Colindale, Public Health England | Takashita,Emi; Fujisaki,Seiichiro; Shirakura,Masayuki; Watanabe,Shinji; Odagiri,Takato |
| EPI626597 | HA | United Kingdom | 2014-Nov-20 | A/ENG/44780079/2014 | Microbiology Services Colindale, Public Health England | Microbiology Services Colindale, Public Health England | Takashita,Emi; Fujisaki,Seiichiro; Shirakura,Masayuki; Watanabe,Shinji; Odagiri,Takato |
| EPI626638 | HA | United Kingdom | 2014-Dec-12 | A/ENG/542/2014 | Microbiology Services Colindale, Public Health England | Microbiology Services Colindale, Public Health England | Takashita,Emi; Fujisaki,Seiichiro; Shirakura,Masayuki; Watanabe,Shinji; Odagiri,Takato |
| EPI626651 | HA | United Kingdom | 2014-Dec-17 | A/ENG/45160093/2014 | Microbiology Services Colindale, Public Health England | Microbiology Services Colindale, Public Health England | Takashita,Emi; Fujisaki,Seiichiro; Shirakura,Masayuki; Watanabe,Shinji; Odagiri,Takato |
| EPI626694 | HA | United Kingdom | 2014-Dec-23 | A/ENG/50100055/2014 | Microbiology Services Colindale, Public Health England | Microbiology Services Colindale, Public Health England | Takashita,Emi; Fujisaki,Seiichiro; Shirakura,Masayuki; Watanabe,Shinji; Odagiri,Takato |
| EPI626702 | HA | United Kingdom | 2014-Dec-24 | A/ENG/50100064/2014 | Microbiology Services Colindale, Public Health England | Microbiology Services Colindale, Public Health England | Takashita,Emi; Fujisaki,Seiichiro; Shirakura,Masayuki; Watanabe,Shinji; Odagiri,Takato |
| EPI626807 | HA | United Kingdom | 2014-Dec-30 | A/ENG/50200185/2014 | Microbiology Services Colindale, Public Health England | Microbiology Services Colindale, Public Health England | |

| | | | | | | |
|---|---|---|---|---|---|---|
| EPI626860 | HA | United Kingdom | 2015-Jan-05 | A/ENG/50360752/2015 | Microbiology Services Colindale, Public Health England | Microbiology Services Colindale, Public Health England |
| EPI626873 | HA | United Kingdom | 2015-Jan-15 | A/ENG/50400311/2015 | Microbiology Services Colindale, Public Health England | Microbiology Services Colindale, Public Health England |
| EPI626881 | HA | United Kingdom | 2015-Jan-15 | A/ENG/75/2015 | Microbiology Services Colindale, Public Health England | Microbiology Services Colindale, Public Health England |
| EPI626889 | HA | United Kingdom | 2015-Jan-15 | A/ENG/50400317/2015 | Microbiology Services Colindale, Public Health England | Microbiology Services Colindale, Public Health England |
| EPI626930 | HA | United Kingdom | 2015-Feb-12 | A/ENG/50800026/2015 | Microbiology Services Colindale, Public Health England | Microbiology Services Colindale, Public Health England |
| EPI626943 | HA | United Kingdom | 2015-Mar-03 | A/ENG/51020042/2015 | Microbiology Services Colindale, Public Health England | Microbiology Services Colindale, Public Health England |
| EPI626967 | HA | United Kingdom | 2015-Jul-03 | A/ENG/52840541/2015 | Microbiology Services Colindale, Public Health England | Microbiology Services Colindale, Public Health England |
| EPI629536 | HA | Russian Federation | 2015-Feb-16 | A/St Petersburg/101/2015 | Russian Academy of Medical Sciences | Centers for Disease Control and Prevention |
| EPI629544 | HA | Russian Federation | 2015-Feb-04 | A/Khabarovsk/19/2015 | Russian Academy of Medical Sciences | Centers for Disease Control and Prevention |
| EPI629552 | HA | United States | 2015-Jun-09 | A/Florida/51/2015 | Florida Department of Health-Jacksonville | Centers for Disease Control and Prevention |
| EPI629560 | HA | United States | 2015-Jun-21 | A/Arizona/09/2015 | Arizona Department of Health Services | Centers for Disease Control and Prevention |
| EPI629582 | HA | Peru | 2015-Apr-13 | A/Peru/27/2015 | Laboratorio de Referencia Nacional Virus Respiratorios, Instituto Nacional de Salud | Centers for Disease Control and Prevention |
| EPI629590 | HA | Peru | 2015-Apr-23 | A/Peru/30/2015 | Laboratorio de Referencia Nacional Virus Respiratorios, Instituto Nacional de Salud | Centers for Disease Control and Prevention |
| EPI629605 | HA | Russian Federation | 2015-Feb-20 | A/St Petersburg/103/2015 | Russian Academy of Medical Sciences | Centers for Disease Control and Prevention |
| EPI629613 | HA | United States | 2015-Feb-10 | A/Hawaii/12/2015 | State of Hawaii Department of Health | Centers for Disease Control and Prevention |
| EPI629628 | HA | United States | 2015-Feb-15 | A/Hawaii/15/2015 | State of Hawaii Department of Health | Centers for Disease Control and Prevention |
| EPI629636 | HA | Peru | 2015-Apr-09 | A/Peru/28/2015 | Laboratorio de Referencia Nacional Virus Respiratorios, Instituto Nacional de Salud | Centers for Disease Control and Prevention |
| EPI629644 | HA | United States | 2015-Feb-25 | A/Hawaii/23/2015 | State of Hawaii Department of Health | Centers for Disease Control and Prevention |
| EPI629659 | HA | United States | 2015-Mar-04 | A/Hawaii/24/2015 | State of Hawaii Department of Health | Centers for Disease Control and Prevention |
| EPI629674 | HA | Lao, People's Democratic Republic | 2015-Feb-02 | A/Laos/122/2015 | National Center for Laboratory and Epidemiology | Centers for Disease Control and Prevention |
| EPI629689 | HA | Kazakhstan | 2015-Feb-19 | A/Kazakhstan/1080/2015 | CSEE | Centers for Disease Control and Prevention |
| EPI629697 | HA | United States | 2015-Jun-01 | A/Alaska/18/2015 | Alaska State Virology Lab | Centers for Disease Control and Prevention |
| EPI629705 | HA | United States | 2015-Jun-05 | A/Alaska/13/2015 | Alaska State Virology Lab | Centers for Disease Control and Prevention |

| | | | | | | |
|---|---|---|---|---|---|---|
| EPI629713 | HA | United States | 2015-Jun-08 | A/Alaska/21/2015 | Alaska State Virology Lab | Centers for Disease Control and Prevention |
| EPI629721 | HA | United States | 2015-Jun-17 | A/Alaska/25/2015 | Alaska State Virology Lab | Centers for Disease Control and Prevention |
| EPI629729 | HA | United States | 2015-Feb-17 | A/Hawaii/14/2015 | State of Hawaii Department of Health | Centers for Disease Control and Prevention |
| EPI631490 | HA | United States | 2015-Apr-02 | A/Minnesota/26/2015 | Minnesota Department of Health | Centers for Disease Control and Prevention |
| EPI631498 | HA | United States | 2015-Feb-17 | A/Arkansas/06/2015 | Arkansas Department of Health | Centers for Disease Control and Prevention |
| EPI631513 | HA | United States | 2015-Apr-18 | A/Florida/46/2015 | Florida Department of Health-Jacksonville | Centers for Disease Control and Prevention |
| EPI631521 | HA | United States | 2015-Feb-17 | A/Indiana/13/2015 | Indiana State Department of Health Laboratories | Centers for Disease Control and Prevention |
| EPI631536 | HA | Chile | 2015-May-18 | A/Santiago/33417/2015 | Instituto de Salud Publica de Chile | Centers for Disease Control and Prevention |
| EPI631544 | HA | Brazil | 2015-May-15 | A/Brazil/9056/2015 | Instituto Adolfo Lutz | Centers for Disease Control and Prevention |
| EPI631571 | HA | Chile | 2015-Jun-09 | A/Santiago/39170/2015 | Instituto de Salud Publica de Chile | Centers for Disease Control and Prevention |
| EPI631579 | HA | United States | 2015-Apr-14 | A/New Jersey/48/2015 | New Jersey Department of Health & Senior Services | Centers for Disease Control and Prevention |
| EPI631587 | HA | United States | 2015-Jul-05 | A/New Jersey/50/2015 | New Jersey Department of Health & Senior Services | Centers for Disease Control and Prevention |
| EPI631595 | HA | United States | 2015-Jun-22 | A/Florida/52/2015 | Florida Department of Health-Jacksonville | Centers for Disease Control and Prevention |
| EPI631606 | HA | United States | 2015-Jul-01 | A/Wisconsin/69/2015 | Wisconsin State Laboratory of Hygiene | Centers for Disease Control and Prevention |
| EPI631614 | HA | United States | 2015-Jun-30 | A/Illinois/12/2015 | Illinois Department of Public Health-Chicago | Centers for Disease Control and Prevention |
| EPI631622 | HA | United States | 2015-Jul-04 | A/Florida/55/2015 | Florida Department of Health-Tampa | Centers for Disease Control and Prevention |
| EPI631638 | HA | United States | 2015-Jun-14 | A/Alaska/26/2015 | Alaska State Virology Lab | Centers for Disease Control and Prevention |
| EPI631646 | HA | United States | 2015-Jun-01 | A/Alaska/17/2015 | Alaska State Virology Lab | Centers for Disease Control and Prevention |
| EPI631654 | HA | United States | 2015-Feb-17 | A/Kentucky/11/2015 | Kentucky Division of Laboratory Services | Centers for Disease Control and Prevention |
| EPI631662 | HA | United States | 2015-Apr-04 | A/Hawaii/28/2015 | State of Hawaii Department of Health | Centers for Disease Control and Prevention |
| EPI631670 | HA | Brazil | 2015-May-04 | A/Brazil/187/2015 | Instituto Oswaldo Cruz FIOCRUZ - Laboratory of Respiratory Viruses and Measles (LVRS) | Centers for Disease Control and Prevention |
| EPI631678 | HA | United States | 2015-Feb-19 | A/Minnesota/23/2015 | Minnesota Department of Health | Centers for Disease Control and Prevention |
| EPI631693 | HA | United States | 2015-Jan-29 | A/South Dakota/23/2015 | South Dakota Public Health Lab | Centers for Disease Control and Prevention |
| EPI631701 | HA | Peru | 2015-Apr-24 | A/Peru/34/2015 | Laboratorio de Referencia Nacional Virus Respiratorios, Instituto Nacional de Salud | Centers for Disease Control and Prevention |

| | | | | | | |
|---|---|---|---|---|---|---|
| EPI631709 | HA | Peru | 2015-May-04 | A/Peru/36/2015 | Laboratorio de Referencia Nacional Virus Respiratorios, Instituto Nacional de Salud | Centers for Disease Control and Prevention |
| EPI631717 | HA | Peru | 2015-May-10 | A/Peru/37/2015 | Laboratorio de Referencia Nacional Virus Respiratorios, Instituto Nacional de Salud | Centers for Disease Control and Prevention |
| EPI631725 | HA | El Salvador | 2015-Mar-09 | A/El Salvador/283/2015 | Contiguo a Hospital Rosales | Centers for Disease Control and Prevention |
| EPI631733 | HA | United States | 2015-Jun-27 | A/Michigan/37/2015 | Michigan Department of Community Health | Centers for Disease Control and Prevention |
| EPI631749 | HA | Brazil | 2015-Mar-15 | A/Brazil/9492/2015 | Instituto Adolfo Lutz | Centers for Disease Control and Prevention |
| EPI631757 | HA | United States | 2015-Jun-30 | A/North Carolina/26/2015 | North Carolina State Laboratory of Public Health | Centers for Disease Control and Prevention |
| EPI631765 | HA | Brazil | 2015-Mar-21 | A/Brazil/9517/2015 | Instituto Adolfo Lutz | Centers for Disease Control and Prevention |
| EPI631787 | HA | United States | 2015-Mar-27 | A/Idaho/18/2015 | State of Idaho Bureau of Laboratories | Centers for Disease Control and Prevention |
| EPI631802 | HA | United States | 2015-Feb-20 | A/California/73/2015 | California Department of Health Services | Centers for Disease Control and Prevention |
| EPI631810 | HA | United States | 2015-Jun-08 | A/Hawaii/45/2015 | State of Hawaii Department of Health | Centers for Disease Control and Prevention |
| EPI631818 | HA | United States | 2015-Mar-29 | A/Massachusetts/18/2015 | Massachusetts Department of Public Health | Centers for Disease Control and Prevention |
| EPI631826 | HA | United States | 2015-Mar-12 | A/Iowa/14/2015 | Iowa State Hygienic Laboratory | Centers for Disease Control and Prevention |
| EPI631834 | HA | United States | 2015-Feb-14 | A/Alabama/08/2015 | ADPH Bureau of Clinical Laboratories | Centers for Disease Control and Prevention |
| EPI631842 | HA | United States | 2015-Feb-24 | A/Michigan/27/2015 | Michigan Department of Community Health | Centers for Disease Control and Prevention |
| EPI631855 | HA | United States | 2015-Mar-31 | A/Florida/43/2015 | Florida Department of Health-Jacksonville | Centers for Disease Control and Prevention |
| EPI631863 | HA | United States | 2015-May-10 | A/Washington/36/2015 | Washington State Public Health Laboratory | Centers for Disease Control and Prevention |
| EPI631871 | HA | United States | 2015-Mar-23 | A/Nebraska/16/2015 | Nebraska Public Health Lab | Centers for Disease Control and Prevention |
| EPI631879 | HA | United States | 2015-Mar-04 | A/Maryland/13/2015 | Maryland Department of Health and Mental Hygiene | Centers for Disease Control and Prevention |
| EPI631887 | HA | United Kingdom | 2014-Jan-01 | A/New Caledonia/71/2014 NIB-91 | Crick Worldwide Influenza Centre | Centers for Disease Control and Prevention |
| EPI631895 | HA | United States | 2015-Feb-23 | A/Mississippi/10/2015 | Mississippi Public Health Laboratory | Centers for Disease Control and Prevention |
| EPI631903 | HA | United States | 2015-Apr-04 | A/South Dakota/12/2015 | South Dakota Public Health Lab | Centers for Disease Control and Prevention |
| EPI631911 | HA | United States | 2014-Nov-21 | A/Washington/33/2014 | Washington State Public Health Laboratory | Centers for Disease Control and Prevention |
| EPI633798 | HA | Russian Federation | 2015-Jan-30 | A/St Petersburg/70/2015 | Russian Academy of Medical Sciences | Centers for Disease Control and Prevention |
| EPI633806 | HA | United States | 2015-Jan-18 | A/Indiana/04/2015 | Indiana State Department of Health Laboratories | Centers for Disease Control and Prevention |

| | | | | | | |
|---|---|---|---|---|---|---|
| EPI633821 | HA | United States | 2015-Jan-20 | A/Virginia/10/2015 | Virginia Division of Consolidated Laboratories | Centers for Disease Control and Prevention |
| EPI633829 | HA | United States | 2015-Jan-04 | A/California/44/2015 | California Department of Health Services | Centers for Disease Control and Prevention |
| EPI633837 | HA | United States | 2015-Jan-19 | A/Rhode Island/07/2015 | Rhode Island Department of Health | Centers for Disease Control and Prevention |
| EPI633852 | HA | United States | 2015-Jul-08 | A/Hawaii/43/2015 | State of Hawaii Department of Health | Centers for Disease Control and Prevention |
| EPI633874 | HA | United States | 2015-Jan-30 | A/Iowa/13/2015 | Iowa State Hygienic Laboratory | Centers for Disease Control and Prevention |
| EPI633888 | HA | United States | 2015-Jan-24 | A/Nevada/33/2015 | Southern Nevada Public Health Lab | Centers for Disease Control and Prevention |
| EPI633896 | HA | United States | 2015-Jul-09 | A/Hawaii/44/2015 | State of Hawaii Department of Health | Centers for Disease Control and Prevention |
| EPI633904 | HA | Brazil | 2015-Jan-05 | A/Brazil/70/2015 | Instituto Oswaldo Cruz FIOCRUZ - Laboratory of Respiratory Viruses and Measles (LVRS) | Centers for Disease Control and Prevention |
| EPI633912 | HA | United States | 2015-Jan-30 | A/Nevada/12/2015 | Southern Nevada Public Health Lab | Centers for Disease Control and Prevention |
| EPI633920 | HA | United States | 2015-Jan-23 | A/Montana/09/2015 | Montana Laboratory Services Bureau | Centers for Disease Control and Prevention |
| EPI633928 | HA | United States | 2015-Jan-29 | A/South Dakota/22/2015 | South Dakota Public Health Lab | Centers for Disease Control and Prevention |
| EPI633936 | HA | United States | 2015-Jan-22 | A/Wyoming/14/2015 | Wyoming Public Health Laboratory | Centers for Disease Control and Prevention |
| EPI633944 | HA | United States | 2015-Jan-19 | A/Nevada/07/2015 | Nevada State Health Laboratory | Centers for Disease Control and Prevention |
| EPI633952 | HA | United States | 2015-Jan-23 | A/Nevada/32/2015 | Southern Nevada Public Health Lab | Centers for Disease Control and Prevention |
| EPI633960 | HA | United States | 2015-Jan-22 | A/Colorado/09/2015 | Colorado Department of Health Lab | Centers for Disease Control and Prevention |
| EPI633985 | HA | United States | 2015-Jan-30 | A/Oklahoma/15/2015 | Oklahoma State Department of Health | Centers for Disease Control and Prevention |
| EPI633993 | HA | United States | 2015-Jan-17 | A/New Mexico/21/2015 | New Mexico Department of Health | Centers for Disease Control and Prevention |
| EPI634001 | HA | Russian Federation | 2015-Jan-20 | A/Khabarovsk/11/2015 | Russian Academy of Medical Sciences | Centers for Disease Control and Prevention |
| EPI634022 | HA | El Salvador | 2015-Jan-09 | A/El Salvador/49/2015 | Contiguo a Hospital Rosales | Centers for Disease Control and Prevention |
| EPI634030 | HA | El Salvador | 2015-Jan-20 | A/El Salvador/63/2015 | Contiguo a Hospital Rosales | Centers for Disease Control and Prevention |
| EPI634038 | HA | United States | 2015-Jan-26 | A/Nevada/23/2015 | Southern Nevada Public Health Lab | Centers for Disease Control and Prevention |
| EPI634053 | HA | United States | 2015-Jul-29 | A/Ohio/21/2015 | Ohio Department of Health Laboratories | Centers for Disease Control and Prevention |
| EPI634061 | HA | United States | 2015-Jul-23 | A/Ohio/20/2015 | Ohio Department of Health Laboratories | Centers for Disease Control and Prevention |
| EPI634069 | HA | Australia | 2015-Mar-31 | A/Victoria/503/2015 | WHO Collaborating Centre for Reference and Research on Influenza | Centers for Disease Control and Prevention |

| | | | | | | |
|---|---|---|---|---|---|---|
| EPI634077 | HA | Fiji | 2015-Mar-02 | A/Fiji/02/2015 | WHO Collaborating Centre for Reference and Research on Influenza | Centers for Disease Control and Prevention |
| EPI634085 | HA | Australia | 2015-Feb-12 | A/South Australia/09/2015 | WHO Collaborating Centre for Reference and Research on Influenza | Centers for Disease Control and Prevention |
| EPI634100 | HA | Lao, People's Democratic Republic | 2015-Jan-14 | A/Laos/046/2015 | National Center for Laboratory and Epidemiology | Centers for Disease Control and Prevention |
| EPI634108 | HA | Lao, People's Democratic Republic | 2015-Jan-14 | A/Laos/052/2015 | National Center for Laboratory and Epidemiology | Centers for Disease Control and Prevention |
| EPI634123 | HA | Australia | 2015-Mar-18 | A/Brisbane/47/2015 | WHO Collaborating Centre for Reference and Research on Influenza | Centers for Disease Control and Prevention |
| EPI634138 | HA | Brazil | 2015-May-14 | A/Brazil/237/2015 | Instituto Oswaldo Cruz FIOCRUZ - Laboratory of Respiratory Viruses and Measles (LVRS) | Centers for Disease Control and Prevention |
| EPI634151 | HA | United States | 2015-Jan-24 | A/Missouri/11/2015 | Missouri Department. of Health & Senior Services | Centers for Disease Control and Prevention |
| EPI634159 | HA | United States | 2015-Jan-30 | A/Kansas/09/2015 | Kansas Department of Health and Environment | Centers for Disease Control and Prevention |
| EPI634167 | HA | Russian Federation | 2015-Jan-16 | A/Khabarovsk/04/2015 | Russian Academy of Medical Sciences | Centers for Disease Control and Prevention |
| EPI634175 | HA | United States | 2015-Jan-19 | A/New Jersey/40/2015 | New Jersey Department of Health & Senior Services | Centers for Disease Control and Prevention |
| EPI634183 | HA | United States | 2015-Jan-19 | A/Virginia/09/2015 | Virginia Division of Consolidated Laboratories | Centers for Disease Control and Prevention |
| EPI634191 | HA | Russian Federation | 2015-Jan-20 | A/Khabarovsk/07/2015 | Russian Academy of Medical Sciences | Centers for Disease Control and Prevention |
| EPI634199 | HA | United States | 2015-Jan-28 | A/Missouri/05/2015 | Missouri Department. of Health & Senior Services | Centers for Disease Control and Prevention |
| EPI634207 | HA | United States | 2015-Jan-26 | A/Wisconsin/30/2015 | Wisconsin State Laboratory of Hygiene | Centers for Disease Control and Prevention |
| EPI638978 | HA | United States | 2015-Jun-19 | A/Alaska/33/2015 | Alaska State Virology Lab | Centers for Disease Control and Prevention |
| EPI638986 | HA | United States | 2015-Jun-17 | A/Alaska/35/2015 | Alaska State Virology Lab | Centers for Disease Control and Prevention |
| EPI638994 | HA | United States | 2015-Jun-18 | A/Alaska/29/2015 | Alaska State Virology Lab | Centers for Disease Control and Prevention |
| EPI639002 | HA | United States | 2015-Jun-21 | A/Alaska/38/2015 | Alaska State Virology Lab | Centers for Disease Control and Prevention |
| EPI639010 | HA | Dominica | 2015-Mar-23 | A/Dominica/672/2015 | Caribbean Epidemiology Center | Centers for Disease Control and Prevention |
| EPI639018 | HA | United States | 2015-Jun-19 | A/Alaska/34/2015 | Alaska State Virology Lab | Centers for Disease Control and Prevention |
| EPI639026 | HA | Dominica | 2015-Mar-14 | A/Dominica/642/2015 | Caribbean Epidemiology Center | Centers for Disease Control and Prevention |
| EPI639034 | HA | United States | 2015-Jul-25 | A/Iowa/20/2015 | Iowa State Hygienic Laboratory | Centers for Disease Control and Prevention |

| | | | | | | |
|---|---|---|---|---|---|---|
| EPI639042 | HA | United States | 2015-Jul-30 | A/Iowa/30/2015 | Iowa State Hygienic Laboratory | Centers for Disease Control and Prevention |
| EPI639057 | HA | Dominica | 2015-Mar-10 | A/Dominica/529/2015 | Caribbean Epidemiology Center | Centers for Disease Control and Prevention |
| EPI639065 | HA | Mexico | 2015-Jul-06 | A/Mexico/1797/2015 | Laboratorio de Virus Respiratorio | Centers for Disease Control and Prevention |
| EPI639073 | HA | United States | 2015-Jun-24 | A/Alaska/66/2015 | Alaska State Virology Lab | Centers for Disease Control and Prevention |
| EPI639081 | HA | Mexico | 2015-Jul-10 | A/Mexico/1783/2015 | Laboratorio de Virus Respiratorio | Centers for Disease Control and Prevention |
| EPI639089 | HA | United States | 2015-Jun-25 | A/Alaska/68/2015 | Alaska State Virology Lab | Centers for Disease Control and Prevention |
| EPI639104 | HA | Mexico | 2015-May-28 | A/Mexico/1670/2015 | Laboratorio de Virus Respiratorio | Centers for Disease Control and Prevention |
| EPI639112 | HA | Ecuador | 2015-Jun-05 | A/Ecuador/2384/2015 | INSPI | Centers for Disease Control and Prevention |
| EPI639120 | HA | Ecuador | 2015-Feb-18 | A/Ecuador/1670/2015 | INSPI | Centers for Disease Control and Prevention |
| EPI639128 | HA | Ecuador | 2015-Jun-06 | A/Ecuador/1668/2015 | INSPI | Centers for Disease Control and Prevention |
| EPI639136 | HA | Ecuador | 2015-Aug-06 | A/Ecuador/1673/2015 | INSPI | Centers for Disease Control and Prevention |
| EPI639149 | HA | United States | 2015-Jun-26 | A/Alaska/65/2015 | Alaska State Virology Lab | Centers for Disease Control and Prevention |
| EPI639157 | HA | United States | 2015-Jul-05 | A/Hawaii/47/2015 | State of Hawaii Department of Health | Centers for Disease Control and Prevention |
| EPI639165 | HA | United States | 2015-Jun-26 | A/Alaska/69/2015 | Alaska State Virology Lab | Centers for Disease Control and Prevention |
| EPI639173 | HA | United States | 2015-Jul-07 | A/Hawaii/46/2015 | State of Hawaii Department of Health | Centers for Disease Control and Prevention |
| EPI639181 | HA | Brazil | 2015-Feb-10 | A/Brazil/8004/2015 | Instituto Adolfo Lutz | Centers for Disease Control and Prevention |
| EPI639189 | HA | United States | 2015-Jul-26 | A/Florida/57/2015 | Florida Department of Health-Tampa | Centers for Disease Control and Prevention |
| EPI639197 | HA | Chile | 2015-May-28 | A/Santiago/37245/2015 | Instituto de Salud Publica de Chile | Centers for Disease Control and Prevention |
| EPI639205 | HA | Chile | 2015-Jun-15 | A/Santiago/40992/2015 | Instituto de Salud Publica de Chile | Centers for Disease Control and Prevention |
| EPI639213 | HA | Chile | 2015-Jun-24 | A/Santiago/42575/2015 | Instituto de Salud Publica de Chile | Centers for Disease Control and Prevention |
| EPI639221 | HA | Chile | 2015-Jun-26 | A/Santiago/44139/2015 | Instituto de Salud Publica de Chile | Centers for Disease Control and Prevention |
| EPI639229 | HA | Chile | 2015-May-28 | A/Santiago/35385/2015 | Instituto de Salud Publica de Chile | Centers for Disease Control and Prevention |
| EPI639237 | HA | United States | 2015-Jul-25 | A/Florida/58/2015 | Florida Department of Health-Tampa | Centers for Disease Control and Prevention |
| EPI639252 | HA | United States | 2015-Jun-12 | A/Michigan/38/2015 | Michigan Department of Community Health | Centers for Disease Control and Prevention |
| EPI639260 | HA | Chile | 2015-Jun-23 | A/Valparaiso/42142/2015 | Instituto de Salud Publica de Chile | Centers for Disease Control and Prevention |

| | | | | | | |
|---|---|---|---|---|---|---|
| EPI639271 | HA | United States | 2015-Jun-21 | A/Alaska/37/2015 | Alaska State Virology Lab | Centers for Disease Control and Prevention |
| EPI639279 | HA | United States | 2015-Jun-16 | A/Alaska/36/2015 | Alaska State Virology Lab | Centers for Disease Control and Prevention |
| EPI639287 | HA | United States | 2015-Jul-14 | A/Washington/39/2015 | Washington State Public Health Laboratory | Centers for Disease Control and Prevention |
| EPI639295 | HA | United States | 2015-Jun-19 | A/Alaska/30/2015 | Alaska State Virology Lab | Centers for Disease Control and Prevention |
| EPI639303 | HA | United States | 2015-Jun-20 | A/Alaska/31/2015 | Alaska State Virology Lab | Centers for Disease Control and Prevention |
| EPI642352 | HA | United States | 2014-Dec-28 | A/Maryland/58/2014 | Maryland Department of Health and Mental Hygiene | Centers for Disease Control and Prevention |
| EPI642513 | HA | United States | 2015-Jun-12 | A/Michigan/39/2015 | Michigan Department of Community Health | Centers for Disease Control and Prevention |
| EPI642560 | HA | Sweden | 2015-Aug-19 | A/Stockholm/40/2015 | | Public Health Agency of Sweden |
| EPI643036 | HA | United States | 2015-Aug-09 | A/Florida/59/2015 | Florida Department of Health-Tampa | Centers for Disease Control and Prevention |
| EPI643044 | HA | United States | 2015-Aug-17 | A/Florida/64/2015 | Florida Department of Health-Tampa | Centers for Disease Control and Prevention |
| EPI643052 | HA | United States | 2015-Aug-18 | A/Florida/62/2015 | Florida Department of Health-Tampa | Centers for Disease Control and Prevention |
| EPI643060 | HA | United States | 2015-Aug-19 | A/Florida/61/2015 | Florida Department of Health-Tampa | Centers for Disease Control and Prevention |
| EPI643068 | HA | United States | 2015-Aug-18 | A/Florida/63/2015 | Florida Department of Health-Tampa | Centers for Disease Control and Prevention |
| EPI643510 | HA | United States | 2015-Jul-26 | A/Alaska/148/2015 | Alaska State Virology Lab | Centers for Disease Control and Prevention |
| EPI643518 | HA | United States | 2015-Jul-19 | A/Alaska/150/2015 | Alaska State Virology Lab | Centers for Disease Control and Prevention |
| EPI643526 | HA | United States | 2015-Jul-07 | A/Alaska/149/2015 | Alaska State Virology Lab | Centers for Disease Control and Prevention |
| EPI643534 | HA | United States | 2015-Aug-02 | A/Alaska/135/2015 | Alaska State Virology Lab | Centers for Disease Control and Prevention |
| EPI643542 | HA | United States | 2015-Jul-23 | A/Alaska/146/2015 | Alaska State Virology Lab | Centers for Disease Control and Prevention |
| EPI643556 | HA | United States | 2015-Aug-10 | A/Alaska/142/2015 | Alaska State Virology Lab | Centers for Disease Control and Prevention |
| EPI643571 | HA | Spain | 2015-Feb-02 | A/Paisvasco/635/2015 | Crick Worldwide Influenza Centre | Centers for Disease Control and Prevention |
| EPI643579 | HA | United States | 2015-Aug-03 | A/Alaska/144/2015 | Alaska State Virology Lab | Centers for Disease Control and Prevention |
| EPI643587 | HA | United States | 2015-Aug-05 | A/Alaska/141/2015 | Alaska State Virology Lab | Centers for Disease Control and Prevention |
| EPI643595 | HA | United States | 2015-Aug-03 | A/Alaska/138/2015 | Alaska State Virology Lab | Centers for Disease Control and Prevention |
| EPI643603 | HA | United States | 2015-Aug-06 | A/Texas/65/2015 | Texas Department of State Health Services-Laboratory Services | Centers for Disease Control and Prevention |
| EPI643611 | HA | United States | 2015-Aug-02 | A/Alaska/137/2015 | Alaska State Virology Lab | Centers for Disease Control and Prevention |

| | | | | | | |
|---|---|---|---|---|---|---|
| EPI643619 | HA | Panama | 2015-Jun-24 | A/Panama/316597/2015 | Instituto Conmemorativo Gorgas de Estudios de la Salud | Centers for Disease Control and Prevention |
| EPI643627 | HA | United States | 2015-Aug-11 | A/Alaska/140/2015 | Alaska State Virology Lab | Centers for Disease Control and Prevention |
| EPI643635 | HA | United States | 2015-Aug-06 | A/Alaska/143/2015 | Alaska State Virology Lab | Centers for Disease Control and Prevention |
| EPI643643 | HA | United States | 2015-Jul-24 | A/Alaska/147/2015 | Alaska State Virology Lab | Centers for Disease Control and Prevention |
| EPI643651 | HA | United States | 2015-Aug-04 | A/Alaska/139/2015 | Alaska State Virology Lab | Centers for Disease Control and Prevention |
| EPI643659 | HA | Italy | 2015-Feb-02 | A/Sassari/8/2015 | Crick Worldwide Influenza Centre | Centers for Disease Control and Prevention |
| EPI643674 | HA | Italy | 2015-Feb-13 | A/Italy/FVG-89/2015 | Crick Worldwide Influenza Centre | Centers for Disease Control and Prevention |
| EPI643682 | HA | Serbia | 2015-Mar-19 | A/Serbia/NS1241/2015 | Crick Worldwide Influenza Centre | Centers for Disease Control and Prevention |
| EPI643690 | HA | United States | 2015-Jul-30 | A/Iowa/22/2015 | Iowa State Hygienic Laboratory | Centers for Disease Control and Prevention |
| EPI643698 | HA | Russian Federation | 2015-Apr-06 | A/Moscow/134/2015 | Ivanovsky Research Institute of Virology RAMS | Centers for Disease Control and Prevention |
| EPI643706 | HA | Guatemala | 2015-May-25 | A/Guatemala/9704/2015 | Laboratorio Nacional De Salud Guatemala | Centers for Disease Control and Prevention |
| EPI643714 | HA | Argentina | 2015-Jul-30 | A/Argentina/229/2015 | Instituto Nacional de Enfermedades Infecciosas | Centers for Disease Control and Prevention |
| EPI643722 | HA | United States | 2015-Jul-30 | A/Iowa/23/2015 | Iowa State Hygienic Laboratory | Centers for Disease Control and Prevention |
| EPI643730 | HA | United States | 2015-Jul-30 | A/Iowa/29/2015 | Iowa State Hygienic Laboratory | Centers for Disease Control and Prevention |
| EPI643738 | HA | Argentina | 2015-Jul-29 | A/Argentina/235/2015 | Instituto Nacional de Enfermedades Infecciosas | Centers for Disease Control and Prevention |
| EPI643746 | HA | Argentina | 2015-Jul-30 | A/Argentina/236/2015 | Instituto Nacional de Enfermedades Infecciosas | Centers for Disease Control and Prevention |
| EPI643761 | HA | Argentina | 2015-Jul-27 | A/Argentina/254/2015 | Instituto Nacional de Enfermedades Infecciosas | Centers for Disease Control and Prevention |
| EPI643769 | HA | Argentina | 2015-Aug-03 | A/Argentina/250/2015 | Instituto Nacional de Enfermedades Infecciosas | Centers for Disease Control and Prevention |
| EPI643780 | HA | Argentina | 2015-Jul-27 | A/Argentina/183/2015 | Instituto Nacional de Enfermedades Infecciosas | Centers for Disease Control and Prevention |
| EPI643788 | HA | Argentina | 2015-Aug-04 | A/Argentina/255/2015 | Instituto Nacional de Enfermedades Infecciosas | Centers for Disease Control and Prevention |
| EPI643803 | HA | United States | 2015-Jul-30 | A/Iowa/31/2015 | Iowa State Hygienic Laboratory | Centers for Disease Control and Prevention |
| EPI643816 | HA | Argentina | 2015-Aug-04 | A/Argentina/261/2015 | Instituto Nacional de Enfermedades Infecciosas | Centers for Disease Control and Prevention |
| EPI643824 | HA | United States | 2015-Jan-31 | A/Missouri/12/2015 | Missouri Department. of Health & Senior Services | Centers for Disease Control and Prevention |
| EPI643832 | HA | Argentina | 2015-Jul-27 | A/Argentina/196/2015 | Instituto Nacional de Enfermedades Infecciosas | Centers for Disease Control and Prevention |
| EPI643840 | HA | Serbia | 2015-Feb-06 | A/Serbia/NS1010/2015 | Crick Worldwide Influenza Centre | Centers for Disease Control and Prevention |

| | | | | | | |
|---|---|---|---|---|---|---|
| EPI643848 | HA | Argentina | 2015-Jul-27 | A/Argentina/218/2015 | Instituto Nacional de Enfermedades Infecciosas | Centers for Disease Control and Prevention |
| EPI643856 | HA | Ecuador | 2015-Apr-25 | A/Ecuador/2369/2015 | INSPI | Centers for Disease Control and Prevention |
| EPI643864 | HA | Argentina | 2015-Jul-29 | A/Argentina/234/2015 | Instituto Nacional de Enfermedades Infecciosas | Centers for Disease Control and Prevention |
| EPI643872 | HA | Argentina | 2015-Jul-25 | A/Argentina/166/2015 | Instituto Nacional de Enfermedades Infecciosas | Centers for Disease Control and Prevention |
| EPI643880 | HA | Argentina | 2015-Jul-19 | A/Argentina/135/2015 | Instituto Nacional de Enfermedades Infecciosas | Centers for Disease Control and Prevention |
| EPI643888 | HA | United States | 2015-Jul-30 | A/Iowa/25/2015 | Iowa State Hygienic Laboratory | Centers for Disease Control and Prevention |
| EPI643896 | HA | Jordan | 2015-Feb-05 | A/Jordan/30024/2015 | Crick Worldwide Influenza Centre | Centers for Disease Control and Prevention |
| EPI643904 | HA | United States | 2015-Jan-29 | A/Iowa/17/2015 | Iowa State Hygienic Laboratory | Centers for Disease Control and Prevention |
| EPI643919 | HA | United States | 2015-Jan-29 | A/Maine/24/2015 | Maine Health and Environmental Testing Laboratory | Centers for Disease Control and Prevention |
| EPI643927 | HA | Brazil | 2015-Mar-27 | A/Brazil/56/2015 | Instituto Oswaldo Cruz FIOCRUZ - Laboratory of Respiratory Viruses and Measles (LVRS) | Centers for Disease Control and Prevention |
| EPI643935 | HA | United States | 2015-Jul-22 | A/Iowa/19/2015 | Iowa State Hygienic Laboratory | Centers for Disease Control and Prevention |
| EPI643943 | HA | Jordan | 2015-Mar-22 | A/Jordan/10702/2015 | Crick Worldwide Influenza Centre | Centers for Disease Control and Prevention |
| EPI643951 | HA | United States | 2015-Jul-30 | A/Iowa/24/2015 | Iowa State Hygienic Laboratory | Centers for Disease Control and Prevention |
| EPI643959 | HA | Jordan | 2015-Feb-14 | A/Jordan/30052/2015 | Crick Worldwide Influenza Centre | Centers for Disease Control and Prevention |
| EPI643967 | HA | United States | 2015-Jan-14 | A/California/53/2015 | California Department of Health Services | Centers for Disease Control and Prevention |
| EPI643975 | HA | United States | 2015-Jan-26 | A/Massachusetts/12/2015 | Massachusetts Department of Public Health | Centers for Disease Control and Prevention |
| EPI643983 | HA | United States | 2015-Jul-30 | A/Iowa/28/2015 | Iowa State Hygienic Laboratory | Centers for Disease Control and Prevention |
| EPI643991 | HA | United States | 2015-Jan-11 | A/Oregon/10/2015 | Oregon Public Health Laboratory | Centers for Disease Control and Prevention |
| EPI643999 | HA | Bangladesh | 2015-May-25 | A/Bangladesh/3400/2015 | Institute of Epidemiology Disease Control and Research (IEDCR) & Bangladesh National Influenza Centre (NIC) | Centers for Disease Control and Prevention |
| EPI644007 | HA | United States | 2015-Jan-31 | A/Iowa/12/2015 | Iowa State Hygienic Laboratory | Centers for Disease Control and Prevention |
| EPI644015 | HA | Colombia | 2015-Jul-21 | A/Colombia/5614/2015 | Instituto Nacional de Salud de Columbia | Centers for Disease Control and Prevention |
| EPI644023 | HA | Guatemala | 2015-Jun-12 | A/Guatemala/27/2015 | Laboratorio Nacional De Salud Guatemala | Centers for Disease Control and Prevention |
| EPI644031 | HA | United States | 2015-Jan-13 | A/Texas/22/2015 | Texas Department of State Health Services-Laboratory Services | Centers for Disease Control and Prevention |
| EPI644039 | HA | Colombia | 2015-Jul-23 | A/Colombia/5615/2015 | Instituto Nacional de Salud de Columbia | Centers for Disease Control and Prevention |

| | | | | | | |
|---|---|---|---|---|---|---|
| EPI644047 | HA | Guatemala | 2015-Jun-18 | A/Guatemala/142/2015 | Laboratorio Nacional De Salud Guatemala | Centers for Disease Control and Prevention |
| EPI644055 | HA | Panama | 2015-Jul-17 | A/Panama/316634/2015 | Instituto Conmemorativo Gorgas de Estudios de la Salud | Centers for Disease Control and Prevention |
| EPI644063 | HA | Brazil | 2015-Feb-11 | A/Brazil/96/2015 | Instituto Oswaldo Cruz FIOCRUZ - Laboratory of Respiratory Viruses and Measles (LVRS) | Centers for Disease Control and Prevention |
| EPI644071 | HA | Russian Federation | 2015-Mar-10 | A/Moscow/99/2015 | Ivanovsky Research Institute of Virology RAMS | Centers for Disease Control and Prevention |
| EPI644079 | HA | United States | 2015-Jul-14 | A/Arizona/11/2015 | Arizona Department of Health Services | Centers for Disease Control and Prevention |
| EPI644087 | HA | United States | 2015-Aug-01 | A/Arizona/10/2015 | Arizona Department of Health Services | Centers for Disease Control and Prevention |
| EPI644095 | HA | Guatemala | 2015-Jun-05 | A/Guatemala/9922/2015 | Laboratorio Nacional De Salud Guatemala | Centers for Disease Control and Prevention |
| EPI644103 | HA | United States | 2015-Jul-24 | A/Alaska/145/2015 | Alaska State Virology Lab | Centers for Disease Control and Prevention |
| EPI644283 | HA | Singapore | 2014-Oct-07 | A/Singapore/KK943/2014 | WHO Collaborating Centre for Reference and Research on Influenza | Centers for Disease Control and Prevention |
| EPI646110 | HA | Japan | 2015-Jan-14 | A/OKAYAMA/2/2015 | Okayama Prefectural Institute for Environmental Science and Public Health | National Institute of Infectious Diseases (NIID) |
| EPI647812 | HA | United States | 2015-May-26 | A/Hawaii/35/2015 | State of Hawaii Department of Health | Centers for Disease Control and Prevention |
| EPI647820 | HA | United States | 2015-Jun-11 | A/Hawaii/36/2015 | State of Hawaii Department of Health | Centers for Disease Control and Prevention |
| EPI647828 | HA | United States | 2015-May-21 | A/Hawaii/34/2015 | State of Hawaii Department of Health | Centers for Disease Control and Prevention |
| EPI647836 | HA | New Zealand | 2015-Jul-02 | A/New Zealand/521/2015 | Institute of Environmental Science & Research | Centers for Disease Control and Prevention |
| EPI647844 | HA | New Zealand | 2015-Jun-15 | A/New Zealand/324/2015 | Institute of Environmental Science & Research | Centers for Disease Control and Prevention |
| EPI647852 | HA | Brazil | 2015-May-02 | A/Brazil/2837/2015 | Instituto Adolfo Lutz | Centers for Disease Control and Prevention |
| EPI647860 | HA | Thailand | 2015-Jul-27 | A/Nonthaburi/333/2015 | WHO National Influenza Centre, National Institute of Medical Research (NIMR) | Centers for Disease Control and Prevention |
| EPI647868 | HA | New Zealand | 2015-Jun-16 | A/New Zealand/322/2015 | Institute of Environmental Science & Research | Centers for Disease Control and Prevention |
| EPI647876 | HA | Thailand | 2015-Jul-21 | A/Rayong/335/2015 | WHO National Influenza Centre, National Institute of Medical Research (NIMR) | Centers for Disease Control and Prevention |
| EPI647884 | HA | Thailand | 2015-Jul-22 | A/Nonthaburi/340/2015 | WHO National Influenza Centre, National Institute of Medical Research (NIMR) | Centers for Disease Control and Prevention |
| EPI647892 | HA | Thailand | 2015-Aug-02 | A/Nonthaburi/345/2015 | WHO National Influenza Centre, National Institute of Medical Research (NIMR) | Centers for Disease Control and Prevention |
| EPI647900 | HA | New Zealand | 2015-Jun-22 | A/New Zealand/381/2015 | Institute of Environmental Science & Research | Centers for Disease Control and Prevention |

| | | | | | | | |
|---|---|---|---|---|---|---|---|
| EPI647908 | HA | New Zealand | 2015-Jul-03 | A/New Zealand/519/2015 | Institute of Environmental Science & Research | Centers for Disease Control and Prevention | |
| EPI647916 | HA | New Zealand | 2015-Jun-16 | A/New Zealand/323/2015 | Institute of Environmental Science & Research | Centers for Disease Control and Prevention | |
| EPI647924 | HA | New Zealand | 2015-Jun-21 | A/New Zealand/385/2015 | Institute of Environmental Science & Research | Centers for Disease Control and Prevention | |
| EPI647932 | HA | New Zealand | 2015-Jul-01 | A/New Zealand/500/2015 | Institute of Environmental Science & Research | Centers for Disease Control and Prevention | |
| EPI647940 | HA | New Zealand | 2015-Jun-12 | A/New Zealand/282/2015 | Institute of Environmental Science & Research | Centers for Disease Control and Prevention | Galiano,M. |
| EPI647948 | HA | New Zealand | 2015-Jun-22 | A/New Zealand/383/2015 | Institute of Environmental Science & Research | Centers for Disease Control and Prevention | Galiano,M. |
| EPI647956 | HA | New Zealand | 2015-Jun-22 | A/New Zealand/382/2015 | Institute of Environmental Science & Research | Centers for Disease Control and Prevention | Galiano,M. |
| EPI647964 | HA | New Zealand | 2015-Jul-01 | A/New Zealand/476/2015 | Institute of Environmental Science & Research | Centers for Disease Control and Prevention | Galiano,M. |
| EPI647972 | HA | United States | 2015-Jul-15 | A/Oregon/16/2015 | Oregon Public Health Laboratory | Centers for Disease Control and Prevention | Galiano,M. |
| EPI647980 | HA | United States | 2015-Jul-18 | A/Oregon/18/2015 | Oregon Public Health Laboratory | Centers for Disease Control and Prevention | |
| EPI647988 | HA | Paraguay | 2015-May-26 | A/Paraguay/3585/2015 | Central Laboratory of Public Health | Centers for Disease Control and Prevention | |
| EPI647996 | HA | Paraguay | 2015-Aug-03 | A/Paraguay/6248/2015 | Central Laboratory of Public Health | Centers for Disease Control and Prevention | |
| EPI648004 | HA | Paraguay | 2015-May-22 | A/Paraguay/3371/2015 | Central Laboratory of Public Health | Centers for Disease Control and Prevention | |
| EPI648012 | HA | Paraguay | 2015-Aug-04 | A/Paraguay/6991/2015 | Central Laboratory of Public Health | Centers for Disease Control and Prevention | |
| EPI648020 | HA | Paraguay | 2015-Jun-04 | A/Paraguay/5554/2015 | Central Laboratory of Public Health | Centers for Disease Control and Prevention | |
| EPI648028 | HA | Guatemala | 2015-May-05 | A/Guatemala/18/2015 | Laboratorio Nacional De Salud Guatemala | Centers for Disease Control and Prevention | |
| EPI648036 | HA | United States | 2015-Apr-21 | A/Wisconsin/66/2015 | Wisconsin State Laboratory of Hygiene | Centers for Disease Control and Prevention | |
| EPI649378 | HA | Guatemala | 2015-May-05 | A/Guatemala/21/2015 | Laboratorio Nacional De Salud Guatemala | Centers for Disease Control and Prevention | |
| EPI649386 | HA | Japan | 2015-Feb-09 | A/Hokkaido/06/2015 | National Institute of Infectious Diseases (NIID) | Centers for Disease Control and Prevention | |
| EPI649394 | HA | United States | 2015-Feb-14 | A/Alabama/09/2015 | ADPH Bureau of Clinical Laboratories | Centers for Disease Control and Prevention | |
| EPI649402 | HA | New Zealand | 2015-Jun-15 | A/New Zealand/588/2015 | Institute of Environmental Science & Research | Centers for Disease Control and Prevention | |
| EPI649410 | HA | Guatemala | 2015-May-05 | A/Guatemala/19/2015 | Laboratorio Nacional De Salud Guatemala | Centers for Disease Control and Prevention | |
| EPI649418 | HA | United States | 2015-Aug-17 | A/Ohio/26/2015 | Ohio Department of Health Laboratories | Centers for Disease Control and Prevention | |
| EPI649426 | HA | United States | 2015-Jul-30 | A/Iowa/26/2015 | Iowa State Hygienic Laboratory | Centers for Disease Control and Prevention | |
| EPI649434 | HA | United States | 2015-Feb-11 | A/Alabama/07/2015 | ADPH Bureau of Clinical Laboratories | Centers for Disease Control and Prevention | |

| | | | | | | |
|---|---|---|---|---|---|---|
| EPI649442 | HA | Guatemala | 2015-May-03 | A/Guatemala/15/2015 | Laboratorio Nacional De Salud Guatemala | Centers for Disease Control and Prevention |
| EPI649450 | HA | Guatemala | 2015-May-18 | A/Guatemala/5015/2015 | Laboratorio Nacional De Salud Guatemala | Centers for Disease Control and Prevention |
| EPI649458 | HA | Guatemala | 2015-Jun-11 | A/Guatemala/5915/2015 | Laboratorio Nacional De Salud Guatemala | Centers for Disease Control and Prevention |
| EPI649466 | HA | United States | 2015-Aug-05 | A/Massachusetts/23/2015 | Massachusetts Department of Public Health | Centers for Disease Control and Prevention |
| EPI649474 | HA | Ecuador | 2015-Aug-06 | A/Ecuador/1673/2015 | INSPI | Centers for Disease Control and Prevention |
| EPI649482 | HA | United States | 2015-Aug-10 | A/New York/64/2015 | New York State Department of Health | Centers for Disease Control and Prevention |
| EPI649490 | HA | Thailand | 2015-Jul-02 | A/Prachuapkhirikhan/289/2015 | WHO National Influenza Centre, National Institute of Medical Research (NIMR) | Centers for Disease Control and Prevention |
| EPI649498 | HA | Japan | 2014-Dec-12 | A/Gifu/46/2014 | National Institute of Infectious Diseases (NIID) | Centers for Disease Control and Prevention |
| EPI649506 | HA | Thailand | 2015-May-16 | A/Nakhonratchaisma/35/2015 | WHO National Influenza Centre, National Institute of Medical Research (NIMR) | Centers for Disease Control and Prevention |
| EPI649514 | HA | New Zealand | 2015-May-27 | A/New Zealand/431/2015 | Institute of Environmental Science & Research | Centers for Disease Control and Prevention |
| EPI649522 | HA | United States | 2015-Jul-21 | A/New York/57/2015 | New York State Department of Health | Centers for Disease Control and Prevention |
| EPI649533 | HA | New Zealand | 2015-Feb-21 | A/New Zealand/87/2015 | Institute of Environmental Science & Research | Centers for Disease Control and Prevention |
| EPI649541 | HA | Japan | 2015-Jan-22 | A/Hyogo/4001/2014 | National Institute of Infectious Diseases (NIID) | Centers for Disease Control and Prevention |
| EPI649549 | HA | Thailand | 2015-Jun-12 | A/Chanthaburi/46/2015 | WHO National Influenza Centre, National Institute of Medical Research (NIMR) | Centers for Disease Control and Prevention |
| EPI649557 | HA | Thailand | 2015-Jun-02 | A/Chanthaburi/40/2015 | WHO National Influenza Centre, National Institute of Medical Research (NIMR) | Centers for Disease Control and Prevention |
| EPI649565 | HA | New Zealand | 2015-Jul-06 | A/New Zealand/562/2015 | Institute of Environmental Science & Research | Centers for Disease Control and Prevention |
| EPI649573 | HA | United States | 2015-Aug-04 | A/New York/62/2015 | New York State Department of Health | Centers for Disease Control and Prevention |
| EPI649588 | HA | New Zealand | 2015-Jun-06 | A/New Zealand/683/2015 | Institute of Environmental Science & Research | Centers for Disease Control and Prevention |
| EPI649596 | HA | Japan | 2015-Mar-02 | A/Fukui/78/2015 | National Institute of Infectious Diseases (NIID) | Centers for Disease Control and Prevention |
| EPI649604 | HA | Russian Federation | 2015-Feb-27 | A/Novosibirsk/64/2015 | State Research Center of Virology and Biotechnology (VECTOR) | Centers for Disease Control and Prevention |
| EPI649612 | HA | Russian Federation | 2015-Mar-11 | A/Moscow/104/2015 | Ivanovsky Research Institute of Virology RAMS | Centers for Disease Control and Prevention |
| EPI649620 | HA | Thailand | 2015-Jun-05 | A/Bangkok/43/2015 | WHO National Influenza Centre, National Institute of Medical Research (NIMR) | Centers for Disease Control and Prevention |
| EPI649628 | HA | United States | 2015-Aug-20 | A/North Carolina/28/2015 | North Carolina State Laboratory of Public Health | Centers for Disease Control and Prevention |

| | | | | | | |
|---|---|---|---|---|---|---|
| EPI649636 | HA | United States | 2015-Aug-03 | A/Ohio/23/2015 | Ohio Department of Health Laboratories | Centers for Disease Control and Prevention |
| EPI649651 | HA | Russian Federation | 2015-Jan-26 | A/Buryatia/19/2015 | State Research Center of Virology and Biotechnology (VECTOR) | Centers for Disease Control and Prevention |
| EPI649669 | HA | Japan | 2015-Jan-15 | A/Chiba/61/2014 | National Institute of Infectious Diseases (NIID) | Centers for Disease Control and Prevention |
| EPI649677 | HA | United States | 2015-Jul-29 | A/New York/59/2015 | New York State Department of Health | Centers for Disease Control and Prevention |
| EPI649691 | HA | New Zealand | 2015-Jul-07 | A/New Zealand/536/2015 | Institute of Environmental Science & Research | Centers for Disease Control and Prevention |
| EPI649699 | HA | New Zealand | 2015-Jun-04 | A/New Zealand/686/2015 | Institute of Environmental Science & Research | Centers for Disease Control and Prevention |
| EPI649707 | HA | Mexico | 2015-Mar-24 | A/Mexico/1384/2015 | Laboratorio de Virus Respiratorio | Centers for Disease Control and Prevention |
| EPI649715 | HA | New Zealand | 2015-Jul-02 | A/New Zealand/532/2015 | Institute of Environmental Science & Research | Centers for Disease Control and Prevention |
| EPI649723 | HA | United States | 2015-Aug-01 | A/New York/60/2015 | New York State Department of Health | Centers for Disease Control and Prevention |
| EPI649731 | HA | Thailand | 2015-May-19 | A/Chanthaburi/49/2015 | WHO National Influenza Centre, National Institute of Medical Research (NIMR) | Centers for Disease Control and Prevention |
| EPI649739 | HA | New Zealand | 2015-Jul-05 | A/New Zealand/526/2015 | Institute of Environmental Science & Research | Centers for Disease Control and Prevention |
| EPI649747 | HA | New Zealand | 2015-Jun-15 | A/New Zealand/588/2015 | Institute of Environmental Science & Research | Centers for Disease Control and Prevention |
| EPI649755 | HA | Mexico | 2015-Mar-18 | A/Mexico/1402/2015 | Laboratorio de Virus Respiratorio | Centers for Disease Control and Prevention |
| EPI649763 | HA | United States | 2015-Aug-03 | A/Ohio/24/2015 | Ohio Department of Health Laboratories | Centers for Disease Control and Prevention |
| EPI649771 | HA | Mexico | 2015-Apr-01 | A/Mexico/1401/2015 | Laboratorio de Virus Respiratorio | Centers for Disease Control and Prevention |
| EPI649779 | HA | United States | 2015-Aug-17 | A/North Carolina/29/2015 | North Carolina State Laboratory of Public Health | Centers for Disease Control and Prevention |
| EPI649787 | HA | Mexico | 2015-Mar-02 | A/Mexico/1233/2015 | Laboratorio de Virus Respiratorio | Centers for Disease Control and Prevention |
| EPI649795 | HA | United States | 2015-Sep-01 | A/Wisconsin/74/2015 | Wisconsin State Laboratory of Hygiene | Centers for Disease Control and Prevention |
| EPI649803 | HA | United States | 2015-Sep-01 | A/Wisconsin/75/2015 | Wisconsin State Laboratory of Hygiene | Centers for Disease Control and Prevention |
| EPI649811 | HA | United States | 2015-Aug-06 | A/New York/63/2015 | New York State Department of Health | Centers for Disease Control and Prevention |
| EPI649819 | HA | Japan | 2014-Dec-12 | A/Gifu/46/2014 | National Institute of Infectious Diseases (NIID) | Centers for Disease Control and Prevention |
| EPI649827 | HA | Japan | 2015-Mar-02 | A/Fukui/78/2015 | National Institute of Infectious Diseases (NIID) | Centers for Disease Control and Prevention |
| EPI649842 | HA | United States | 2015-Aug-03 | A/New York/58/2015 | New York State Department of Health | Centers for Disease Control and Prevention |
| EPI649850 | HA | Japan | 2015-Feb-09 | A/Hokkaido/06/2015 | National Institute of Infectious Diseases (NIID) | Centers for Disease Control and Prevention |
| EPI649858 | HA | Japan | 2015-Jan-15 | A/Chiba/61/2014 | National Institute of Infectious Diseases (NIID) | Centers for Disease Control and Prevention |

| | | | | | | | |
|---|---|---|---|---|---|---|---|
| EPI649866 | HA | United States | 2015-Jul-30 | A/Iowa/37/2015 | Iowa State Hygienic Laboratory | Centers for Disease Control and Prevention | |
| EPI649874 | HA | United States | 2015-Jul-30 | A/Iowa/38/2015 | Iowa State Hygienic Laboratory | Centers for Disease Control and Prevention | |
| EPI649882 | HA | United States | 2015-Jun-17 | A/Virginia/32/2015 | Virginia Division of Consolidated Laboratories | Centers for Disease Control and Prevention | |
| EPI649890 | HA | Mexico | 2015-May-06 | A/Mexico/1546/2015 | Laboratorio de Virus Respiratorio | Centers for Disease Control and Prevention | |
| EPI649905 | HA | Russian Federation | 2015-Mar-14 | A/Moscow/102/2015 | Ivanovsky Research Institute of Virology RAMS | Centers for Disease Control and Prevention | Galiano,M. |
| EPI649913 | HA | Russian Federation | 2015-Mar-18 | A/N.Novgorod/143/2015 | Ivanovsky Research Institute of Virology RAMS | Centers for Disease Control and Prevention | Galiano,M. |
| EPI649921 | HA | Russian Federation | 2015-Mar-12 | A/Vladimir/137/2015 | Ivanovsky Research Institute of Virology RAMS | Centers for Disease Control and Prevention | |
| EPI649929 | HA | Guatemala | 2015-Mar-12 | A/Guatemala/13/2015 | Laboratorio Nacional De Salud Guatemala | Centers for Disease Control and Prevention | |
| EPI649937 | HA | Guatemala | 2015-May-03 | A/Guatemala/15/2015 | Laboratorio Nacional De Salud Guatemala | Centers for Disease Control and Prevention | |
| EPI649945 | HA | Brazil | 2015-Apr-09 | A/Brazil/114/2015 | Instituto Oswaldo Cruz FIOCRUZ - Laboratory of Respiratory Viruses and Measles (LVRS) | Centers for Disease Control and Prevention | |
| EPI649953 | HA | Guatemala | 2015-May-18 | A/Guatemala/5015/2015 | Laboratorio Nacional De Salud Guatemala | Centers for Disease Control and Prevention | |
| EPI649961 | HA | Guatemala | 2015-Mar-18 | A/Guatemala/3315/2015 | Laboratorio Nacional De Salud Guatemala | Centers for Disease Control and Prevention | |
| EPI649969 | HA | United States | 2015-Aug-07 | A/Massachusetts/22/2015 | Massachusetts Department of Public Health | Centers for Disease Control and Prevention | |
| EPI649977 | HA | Guatemala | 2015-Mar-12 | A/Guatemala/13/2015 | Laboratorio Nacional De Salud Guatemala | Centers for Disease Control and Prevention | |
| EPI649985 | HA | United States | 2015-Jul-27 | A/New Mexico/22/2015 | New Mexico Department of Health | Centers for Disease Control and Prevention | |
| EPI649993 | HA | United States | 2015-Aug-13 | A/Montana/17/2015 | Montana Laboratory Services Bureau | Centers for Disease Control and Prevention | |
| EPI650001 | HA | United States | 2015-Jul-30 | A/Iowa/34/2015 | Iowa State Hygienic Laboratory | Centers for Disease Control and Prevention | |
| EPI650009 | HA | United States | 2015-Aug-08 | A/Montana/16/2015 | Montana Laboratory Services Bureau | Centers for Disease Control and Prevention | |
| EPI650017 | HA | Japan | 2015-Jan-22 | A/Hyogo/4001/2014 | National Institute of Infectious Diseases (NIID) | Centers for Disease Control and Prevention | |
| EPI651432 | HA | United States | 2015-Jul-06 | A/Alaska/86/2015 | Alaska State Virology Lab | Centers for Disease Control and Prevention | |
| EPI651440 | HA | United States | 2015-Aug-05 | A/Arizona/13/2015 | Arizona Department of Health Services | Centers for Disease Control and Prevention | |
| EPI651448 | HA | United States | 2015-Jul-17 | A/Hawaii/48/2015 | State of Hawaii Department of Health | Centers for Disease Control and Prevention | |
| EPI651456 | HA | United States | 2015-Jul-23 | A/Hawaii/49/2015 | State of Hawaii Department of Health | Centers for Disease Control and Prevention | |
| EPI651464 | HA | United States | 2015-Jul-01 | A/Alaska/88/2015 | Alaska State Virology Lab | Centers for Disease Control and Prevention | |
| EPI651480 | HA | United States | 2015-Jul-06 | A/Alaska/82/2015 | Alaska State Virology Lab | Centers for Disease Control and Prevention | |

| | | | | | | |
|---|---|---|---|---|---|---|
| EPI651488 | HA | United States | 2015-Jul-12 | A/Connecticut/01/2015 | Connecticut Department. of Public Health | Centers for Disease Control and Prevention |
| EPI651496 | HA | United States | 2015-Jun-22 | A/Alaska/72/2015 | Alaska State Virology Lab | Centers for Disease Control and Prevention |
| EPI651504 | HA | United States | 2015-Jul-03 | A/Alaska/87/2015 | Alaska State Virology Lab | Centers for Disease Control and Prevention |
| EPI651512 | HA | United States | 2015-Jul-06 | A/Alaska/83/2015 | Alaska State Virology Lab | Centers for Disease Control and Prevention |
| EPI651520 | HA | United States | 2015-Jun-14 | A/Alaska/27/2015 | Alaska State Virology Lab | Centers for Disease Control and Prevention |
| EPI651528 | HA | United States | 2015-Jul-06 | A/Alaska/85/2015 | Alaska State Virology Lab | Centers for Disease Control and Prevention |
| EPI651536 | HA | United States | 2015-Jun-26 | A/Alaska/71/2015 | Alaska State Virology Lab | Centers for Disease Control and Prevention |
| EPI651544 | HA | United States | 2015-Aug-28 | A/Wisconsin/72/2015 | Wisconsin State Laboratory of Hygiene | Centers for Disease Control and Prevention |
| EPI651552 | HA | United States | 2015-Jun-23 | A/Alaska/73/2015 | Alaska State Virology Lab | Centers for Disease Control and Prevention |
| EPI651560 | HA | United States | 2015-Jul-07 | A/Michigan/40/2015 | Michigan Department of Community Health | Centers for Disease Control and Prevention |
| EPI651568 | HA | United States | 2015-Jun-24 | A/Alaska/79/2015 | Alaska State Virology Lab | Centers for Disease Control and Prevention |
| EPI651576 | HA | United States | 2015-Jul-02 | A/Alaska/94/2015 | Alaska State Virology Lab | Centers for Disease Control and Prevention |
| EPI651584 | HA | United States | 2015-Aug-26 | A/Arizona/14/2015 | Arizona Department of Health Services | Centers for Disease Control and Prevention |
| EPI651592 | HA | United States | 2015-Jul-03 | A/Alaska/91/2015 | Alaska State Virology Lab | Centers for Disease Control and Prevention |
| EPI651600 | HA | United States | 2015-Jun-25 | A/Alaska/77/2015 | Alaska State Virology Lab | Centers for Disease Control and Prevention |
| EPI651608 | HA | United States | 2015-Jul-02 | A/Alaska/92/2015 | Alaska State Virology Lab | Centers for Disease Control and Prevention |
| EPI651616 | HA | United States | 2015-Jun-28 | A/Alaska/78/2015 | Alaska State Virology Lab | Centers for Disease Control and Prevention |
| EPI651624 | HA | United States | 2015-Jul-01 | A/Alaska/93/2015 | Alaska State Virology Lab | Centers for Disease Control and Prevention |
| EPI651632 | HA | United States | 2015-Jun-23 | A/Alaska/75/2015 | Alaska State Virology Lab | Centers for Disease Control and Prevention |
| EPI651640 | HA | United States | 2015-Jul-03 | A/Alaska/90/2015 | Alaska State Virology Lab | Centers for Disease Control and Prevention |
| EPI651648 | HA | United States | 2015-Sep-05 | A/Alabama/12/2015 | ADPH Bureau of Clinical Laboratories | Centers for Disease Control and Prevention |
| EPI651656 | HA | United States | 2015-Sep-08 | A/Alabama/10/2015 | ADPH Bureau of Clinical Laboratories | Centers for Disease Control and Prevention |
| EPI651664 | HA | United States | 2015-Sep-01 | A/Minnesota/40/2015 | Minnesota Department of Health | Centers for Disease Control and Prevention |
| EPI651672 | HA | United States | 2015-Sep-08 | A/Alabama/11/2015 | ADPH Bureau of Clinical Laboratories | Centers for Disease Control and Prevention |
| EPI651680 | HA | United States | 2015-Jul-06 | A/Alaska/84/2015 | Alaska State Virology Lab | Centers for Disease Control and Prevention |

| | | | | | | | |
|---|---|---|---|---|---|---|---|
| EPI651696 | HA | United States | 2015-Jul-01 | A/Alaska/89/2015 | Alaska State Virology Lab | Centers for Disease Control and Prevention | |
| EPI651704 | HA | United States | 2015-Jun-30 | A/Alaska/80/2015 | Alaska State Virology Lab | Centers for Disease Control and Prevention | |
| EPI651712 | HA | United States | 2015-Jul-01 | A/Alaska/95/2015 | Alaska State Virology Lab | Centers for Disease Control and Prevention | |
| EPI651720 | HA | United States | 2015-Sep-02 | A/New Hampshire/38/2015 | New Hampshire Public Health Laboratories | Centers for Disease Control and Prevention | |
| EPI651727 | HA | United States | 2015-Aug-29 | A/Connecticut/02/2015 | Connecticut Department. of Public Health | Centers for Disease Control and Prevention | |
| EPI651735 | HA | United States | 2015-Aug-25 | A/Texas/66/2015 | Texas Department of State Health Services-Laboratory Services | Centers for Disease Control and Prevention | |
| EPI651743 | HA | United States | 2015-Aug-19 | A/Hawaii/52/2015 | State of Hawaii Department of Health | Centers for Disease Control and Prevention | |
| EPI651751 | HA | United States | 2015-Sep-01 | A/Wisconsin/77/2015 | Wisconsin State Laboratory of Hygiene | Centers for Disease Control and Prevention | |
| EPI651759 | HA | United States | 2015-Aug-23 | A/Hawaii/53/2015 | State of Hawaii Department of Health | Centers for Disease Control and Prevention | |
| EPI651767 | HA | United States | 2015-Aug-25 | A/New York/65/2015 | New York State Department of Health | Centers for Disease Control and Prevention | |
| EPI651775 | HA | United States | 2015-Sep-02 | A/Florida/68/2015 | Florida Department of Health-Tampa | Centers for Disease Control and Prevention | |
| EPI651783 | HA | United States | 2015-Aug-25 | A/Hawaii/54/2015 | State of Hawaii Department of Health | Centers for Disease Control and Prevention | |
| EPI651791 | HA | United States | 2015-Jul-02 | A/Alaska/81/2015 | Alaska State Virology Lab | Centers for Disease Control and Prevention | |
| EPI653163 | HA | United States | 2015-Aug-24 | A/Michigan/43/2015 | Michigan Department of Community Health | Centers for Disease Control and Prevention | |
| EPI653171 | HA | United States | 2015-Aug-21 | A/Michigan/44/2015 | Michigan Department of Community Health | Centers for Disease Control and Prevention | |
| EPI653179 | HA | United States | 2015-Aug-15 | A/Michigan/41/2015 | Michigan Department of Community Health | Centers for Disease Control and Prevention | |
| EPI653187 | HA | United States | 2015-Jun-19 | A/Alaska/43/2015 | Alaska State Virology Lab | Centers for Disease Control and Prevention | |
| EPI653201 | HA | Hong Kong (SAR) | 2014-Feb-26 | A/Hong Kong/4801/2014 | Crick Worldwide Influenza Centre | Centers for Disease Control and Prevention | |
| EPI653209 | HA | Switzerland | 2013-Jan-01 | A/Switzerland/9715293/2013 NIB-88 | National Institute for Biological Standards and Control (NIBSC) | Centers for Disease Control and Prevention | |
| EPI662361 | HA | Sweden | 2015-Sep-26 | A/Stockholm/42/2015 | | Public Health Agency of Sweden | Takashita,Emi; Fujisaki,Seiichiro; Shirakura,Masayuki; Watanabe,Shinji; Odagiri,Takato |
| EPI662369 | HA | Uruguay | 2015-Aug-11 | A/Uruguay/86/2015 | Departamento de Laboratorio de Salud Pública (DLSP) | Centers for Disease Control and Prevention | Takashita,Emi; Fujisaki,Seiichiro; Shirakura,Masayuki; Watanabe,Shinji; Odagiri,Takato |
| EPI662377 | HA | United States | 2015-Aug-29 | A/Hawaii/55/2015 | State of Hawaii Department of Health | Centers for Disease Control and Prevention | Takashita,Emi; Fujisaki,Seiichiro; Shirakura,Masayuki; Watanabe,Shinji; Odagiri,Takato |
| EPI662385 | HA | Uruguay | 2015-Jul-08 | A/Uruguay/44/2015 | Departamento de Laboratorio de Salud Pública (DLSP) | Centers for Disease Control and Prevention | Takashita,Emi; Fujisaki,Seiichiro; Shirakura,Masayuki; Watanabe,Shinji; Odagiri,Takato |

| | | | | | | | |
|---|---|---|---|---|---|---|---|
| EPI662394 | HA | United States | 2015-Sep-14 | A/Florida/73/2015 | Florida Department of Health-Tampa | Centers for Disease Control and Prevention | Takashita,Emi; Fujisaki,Seiichiro; Shirakura,Masayuki; Watanabe,Shinji; Odagiri,Takato |
| EPI662402 | HA | Colombia | 2015-Jun-25 | A/Colombia/5402/2015 | Instituto Nacional de Salud de Columbia | Centers for Disease Control and Prevention | Takashita,Emi; Fujisaki,Seiichiro; Shirakura,Masayuki; Watanabe,Shinji; Odagiri,Takato |
| EPI662410 | HA | Colombia | 2015-Jan-18 | A/Colombia/235/2015 | Instituto Nacional de Salud de Columbia | Centers for Disease Control and Prevention | Takashita,Emi; Fujisaki,Seiichiro; Shirakura,Masayuki; Watanabe,Shinji; Odagiri,Takato |
| EPI662418 | HA | Colombia | 2015-Jun-30 | A/Colombia/5438/2015 | Instituto Nacional de Salud de Columbia | Centers for Disease Control and Prevention | |
| EPI662426 | HA | Colombia | 2015-Jun-30 | A/Colombia/5364/2015 | Instituto Nacional de Salud de Columbia | Centers for Disease Control and Prevention | |
| EPI662434 | HA | United States | 2015-Aug-06 | A/Wisconsin/70/2015 | Wisconsin State Laboratory of Hygiene | Centers for Disease Control and Prevention | |
| EPI662442 | HA | Colombia | 2015-Jun-07 | A/Colombia/241/2015 | Instituto Nacional de Salud de Columbia | Centers for Disease Control and Prevention | |
| EPI662450 | HA | United States | 2015-Sep-02 | A/Hawaii/56/2015 | State of Hawaii Department of Health | Centers for Disease Control and Prevention | |
| EPI662458 | HA | United States | 2015-Aug-02 | A/Wisconsin/71/2015 | Wisconsin State Laboratory of Hygiene | Centers for Disease Control and Prevention | |
| EPI662466 | HA | Colombia | 2015-Jun-04 | A/Colombia/5215/2015 | Instituto Nacional de Salud de Columbia | Centers for Disease Control and Prevention | |
| EPI662474 | HA | Uruguay | 2015-Aug-13 | A/Uruguay/122/2015 | Departamento de Laboratorio de Salud Pública (DLSP) | Centers for Disease Control and Prevention | |
| EPI662482 | HA | United States | 2015-Sep-13 | A/Hawaii/57/2015 | State of Hawaii Department of Health | Centers for Disease Control and Prevention | |
| EPI662490 | HA | Uruguay | 2015-Jun-16 | A/Uruguay/39/2015 | Departamento de Laboratorio de Salud Pública (DLSP) | Centers for Disease Control and Prevention | |
| EPI662498 | HA | Uruguay | 2015-Jul-06 | A/Uruguay/43/2015 | Departamento de Laboratorio de Salud Pública (DLSP) | Centers for Disease Control and Prevention | |
| EPI662506 | HA | Uruguay | 2015-Jul-20 | A/Uruguay/54/2015 | Departamento de Laboratorio de Salud Pública (DLSP) | Centers for Disease Control and Prevention | |
| EPI662514 | HA | Uruguay | 2015-Jul-21 | A/Uruguay/55/2015 | Departamento de Laboratorio de Salud Pública (DLSP) | Centers for Disease Control and Prevention | |
| EPI662522 | HA | United States | 2015-Aug-10 | A/New York/64/2015 | New York State Department of Health | Centers for Disease Control and Prevention | |
| EPI662537 | HA | United States | 2015-Jun-25 | A/Alaska/77/2015 | Alaska State Virology Lab | Centers for Disease Control and Prevention | Takashita,Emi; Fujisaki,Seiichiro; Shirakura,Masayuki; Watanabe,Shinji; Odagiri,Takato |
| EPI662545 | HA | United States | 2015-Jun-23 | A/Alaska/75/2015 | Alaska State Virology Lab | Centers for Disease Control and Prevention | |
| EPI662553 | HA | United States | 2015-Jun-30 | A/Alaska/80/2015 | Alaska State Virology Lab | Centers for Disease Control and Prevention | |
| EPI662561 | HA | United States | 2015-Jun-24 | A/Alaska/79/2015 | Alaska State Virology Lab | Centers for Disease Control and Prevention | |
| EPI662569 | HA | United States | 2015-Jul-01 | A/Alaska/93/2015 | Alaska State Virology Lab | Centers for Disease Control and Prevention | |
| EPI662577 | HA | United States | 2015-Jul-06 | A/Alaska/83/2015 | Alaska State Virology Lab | Centers for Disease Control and Prevention | |

| | | | | | | |
|---|---|---|---|---|---|---|
| EPI662585 | HA | United States | 2015-Jul-06 | A/Alaska/86/2015 | Alaska State Virology Lab | Centers for Disease Control and Prevention |
| EPI667522 | HA | United States | 2014-Nov-19 | A/Massachusetts/14/2014 | Massachusetts Department of Public Health | Centers for Disease Control and Prevention |
| EPI667530 | HA | United States | 2015-May-21 | A/Hawaii/34/2015 | State of Hawaii Department of Health | Centers for Disease Control and Prevention |
| EPI667548 | HA | United States | 2015-Jun-23 | A/Alaska/45/2015 | Alaska State Virology Lab | Centers for Disease Control and Prevention |
| EPI667556 | HA | United States | 2015-Jun-21 | A/Alaska/46/2015 | Alaska State Virology Lab | Centers for Disease Control and Prevention |
| EPI667564 | HA | United States | 2015-Jun-21 | A/Alaska/48/2015 | Alaska State Virology Lab | Centers for Disease Control and Prevention |
| EPI667572 | HA | Dominican Republic | 2015-Mar-11 | A/Dominican Republic/8946/2015 | Laboratorio de Investigacion / Centro de Educacion Medica y Amistad Dominico Japones (CEMADOJA) | Centers for Disease Control and Prevention |
| EPI667580 | HA | Brazil | 2015-Apr-16 | A/Brazil/8433/2015 | Instituto Adolfo Lutz | Centers for Disease Control and Prevention |
| EPI667588 | HA | Brazil | 2015-Mar-13 | A/Brazil/9850/2015 | Instituto Adolfo Lutz | Centers for Disease Control and Prevention |
| EPI667596 | HA | Brazil | 2015-Mar-20 | A/Brazil/1804/2015 | Instituto Adolfo Lutz | Centers for Disease Control and Prevention |
| EPI667604 | HA | Brazil | 2015-Feb-18 | A/Brazil/440/2015 | Instituto Adolfo Lutz | Centers for Disease Control and Prevention |
| EPI667620 | HA | Brazil | 2015-Mar-17 | A/Brazil/8925/2015 | Instituto Adolfo Lutz | Centers for Disease Control and Prevention |
| EPI667628 | HA | Brazil | 2015-Mar-16 | A/Brazil/4996/2015 | Instituto Adolfo Lutz | Centers for Disease Control and Prevention |
| EPI667636 | HA | Brazil | 2015-Feb-18 | A/Brazil/6261/2015 | Instituto Adolfo Lutz | Centers for Disease Control and Prevention |
| EPI667652 | HA | Ecuador | 2015-May-21 | A/Ecuador/2404/2015 | INSPI | Centers for Disease Control and Prevention |
| EPI667660 | HA | Ecuador | 2015-May-14 | A/Ecuador/572/2015 | INSPI | Centers for Disease Control and Prevention |
| EPI667668 | HA | Ecuador | 2015-Jun-29 | A/Ecuador/1695/2015 | INSPI | Centers for Disease Control and Prevention |
| EPI667676 | HA | Ecuador | 2015-Feb-05 | A/Ecuador/821/2015 | INSPI | Centers for Disease Control and Prevention |
| EPI667684 | HA | Ecuador | 2015-May-08 | A/Ecuador/1845/2015 | INSPI | Centers for Disease Control and Prevention |
| EPI667692 | HA | United States | 2015-Jul-01 | A/Alaska/67/2015 | Alaska State Virology Lab | Centers for Disease Control and Prevention |
| EPI667700 | HA | United States | 2015-Jul-10 | A/New York/55/2015 | New York State Department of Health | Centers for Disease Control and Prevention |
| EPI667708 | HA | United States | 2015-Jul-23 | A/Ohio/18/2015 | Ohio Department of Health Laboratories | Centers for Disease Control and Prevention |
| EPI667716 | HA | United States | 2015-Jun-30 | A/Oregon/11/2015 | Oregon Public Health Laboratory | Centers for Disease Control and Prevention |
| EPI667727 | HA | United States | 2015-Jul-15 | A/Nebraska/17/2015 | Nebraska Public Health Lab | Centers for Disease Control and Prevention |

| | | | | | | |
|---|---|---|---|---|---|---|
| EPI667735 | HA | Lao, People's Democratic Republic | 2015-Feb-25 | A/Laos/218/2015 | National Center for Laboratory and Epidemiology | Centers for Disease Control and Prevention |
| EPI667749 | HA | Lao, People's Democratic Republic | 2015-Apr-20 | A/Laos/372/2015 | National Center for Laboratory and Epidemiology | Centers for Disease Control and Prevention |
| EPI667757 | HA | Lao, People's Democratic Republic | 2015-May-25 | A/Laos/529/2015 | National Center for Laboratory and Epidemiology | Centers for Disease Control and Prevention |
| EPI667765 | HA | Lao, People's Democratic Republic | 2015-May-25 | A/Laos/535/2015 | National Center for Laboratory and Epidemiology | Centers for Disease Control and Prevention |
| EPI667773 | HA | Lao, People's Democratic Republic | 2015-Apr-29 | A/Laos/179/2015 | National Center for Laboratory and Epidemiology | Centers for Disease Control and Prevention |
| EPI667781 | HA | United States | 2015-Jul-13 | A/Utah/15/2015 | Utah Department of Health | Centers for Disease Control and Prevention |
| EPI667786 | HA | Brazil | 2015-Jan-23 | A/Acre/133946-IEC/2015 | National Influenza Center | Centers for Disease Control and Prevention |
| EPI667801 | HA | Brazil | 2015-Mar-18 | A/Para/134244-IEC/2015 | National Influenza Center | Centers for Disease Control and Prevention |
| EPI667809 | HA | Brazil | 2015-Mar-23 | A/Para/134324-IEC/2015 | National Influenza Center | Centers for Disease Control and Prevention |
| EPI667817 | HA | Brazil | 2015-Mar-31 | A/Para/134473-IEC/2015 | National Influenza Center | Centers for Disease Control and Prevention |
| EPI667825 | HA | Brazil | 2015-Apr-06 | A/Para/134575-IEC/2015 | National Influenza Center | Centers for Disease Control and Prevention |
| EPI667833 | HA | Brazil | 2015-Mar-31 | A/Rio Grande Do Norte/134592-IEC/2015 | National Influenza Center | Centers for Disease Control and Prevention |
| EPI667841 | HA | Brazil | 2015-Apr-08 | A/Rio Grande Do Norte/134696-IEC/2015 | National Influenza Center | Centers for Disease Control and Prevention |
| EPI667849 | HA | Brazil | 2015-Apr-04 | A/Para/134701-IEC/2015 | National Influenza Center | Centers for Disease Control and Prevention |
| EPI667864 | HA | Puerto Rico | 2015-Jul-13 | A/Puerto Rico/04/2015 | Puerto Rico Department of Health | Centers for Disease Control and Prevention |
| EPI667872 | HA | Puerto Rico | 2015-Jul-14 | A/Puerto Rico/05/2015 | Puerto Rico Department of Health | Centers for Disease Control and Prevention |
| EPI667880 | HA | Puerto Rico | 2015-Jul-21 | A/Puerto Rico/06/2015 | Puerto Rico Department of Health | Centers for Disease Control and Prevention |
| EPI667888 | HA | Puerto Rico | 2015-Aug-07 | A/Puerto Rico/07/2015 | Puerto Rico Department of Health | Centers for Disease Control and Prevention |
| EPI667896 | HA | Puerto Rico | 2015-Aug-10 | A/Puerto Rico/08/2015 | Puerto Rico Department of Health | Centers for Disease Control and Prevention |
| EPI667910 | HA | Dominica | 2015-Mar-16 | A/Dominica/645/2015 | Caribbean Epidemiology Center | Centers for Disease Control and Prevention |
| EPI667918 | HA | Saint Vincent and the Grenadines | 2015-Feb-10 | A/St. Vincent And Grenadines/422/2015 | Caribbean Epidemiology Center | Centers for Disease Control and Prevention |
| EPI667926 | HA | Saint Vincent and the Grenadines | 2015-Feb-16 | A/St. Vincent And Grenadines/424/2015 | Caribbean Epidemiology Center | Centers for Disease Control and Prevention |

| | | | | | | |
|---|---|---|---|---|---|---|
| EPI667941 | HA | United States | 2015-Jul-01 | A/Oregon/17/2015 | Oregon Public Health Laboratory | Centers for Disease Control and Prevention |
| EPI667956 | HA | Australia | 2015-May-16 | A/Victoria/511/2015 | WHO Collaborating Centre for Reference and Research on Influenza | Centers for Disease Control and Prevention |
| EPI667964 | HA | Russian Federation | 2015-Jan-26 | A/Omsk/141/2015 | State Research Center of Virology and Biotechnology (VECTOR) | Centers for Disease Control and Prevention |
| EPI667972 | HA | Russian Federation | 2015-Jan-27 | A/Komi/99/2015 | State Research Center of Virology and Biotechnology (VECTOR) | Centers for Disease Control and Prevention |
| EPI667980 | HA | Russian Federation | 2015-Feb-20 | A/Chelyabinsk/192/2015 | State Research Center of Virology and Biotechnology (VECTOR) | Centers for Disease Control and Prevention |
| EPI667988 | HA | United States | 2015-Aug-09 | A/Florida/60/2015 | Florida Department of Health-Tampa | Centers for Disease Control and Prevention |
| EPI667996 | HA | United States | 2015-Aug-18 | A/Alaska/160/2015 | Alaska State Virology Lab | Centers for Disease Control and Prevention |
| EPI668012 | HA | United States | 2015-Aug-18 | A/Tennessee/18/2015 | Tennessee Department of Health Laboratory-Nashville | Centers for Disease Control and Prevention |
| EPI668020 | HA | United States | 2015-Aug-19 | A/Tennessee/20/2015 | Tennessee Department of Health Laboratory-Nashville | Centers for Disease Control and Prevention |
| EPI668028 | HA | United States | 2015-Aug-19 | A/Tennessee/20/2015 | Tennessee Department of Health Laboratory-Nashville | Centers for Disease Control and Prevention |
| EPI668036 | HA | Argentina | 2015-Jul-18 | A/Argentina/112/2015 | Instituto Nacional de Enfermedades Infecciosas | Centers for Disease Control and Prevention |
| EPI668044 | HA | Argentina | 2015-Jul-14 | A/Argentina/113/2015 | Instituto Nacional de Enfermedades Infecciosas | Centers for Disease Control and Prevention |
| EPI668052 | HA | Argentina | 2015-Jul-16 | A/Argentina/114/2015 | Instituto Nacional de Enfermedades Infecciosas | Centers for Disease Control and Prevention |
| EPI668060 | HA | Argentina | 2015-Jul-15 | A/Argentina/116/2015 | Instituto Nacional de Enfermedades Infecciosas | Centers for Disease Control and Prevention |
| EPI668068 | HA | Argentina | 2015-Jul-18 | A/Argentina/120/2015 | Instituto Nacional de Enfermedades Infecciosas | Centers for Disease Control and Prevention |
| EPI668076 | HA | Argentina | 2015-Jul-16 | A/Argentina/121/2015 | Instituto Nacional de Enfermedades Infecciosas | Centers for Disease Control and Prevention |
| EPI668084 | HA | Argentina | 2015-Jul-22 | A/Argentina/132/2015 | Instituto Nacional de Enfermedades Infecciosas | Centers for Disease Control and Prevention |
| EPI668092 | HA | Argentina | 2015-Jul-21 | A/Argentina/133/2015 | Instituto Nacional de Enfermedades Infecciosas | Centers for Disease Control and Prevention |
| EPI668113 | HA | United States | 2015-Sep-02 | A/New Hampshire/38/2015 | New Hampshire Public Health Laboratories | Centers for Disease Control and Prevention |
| EPI668121 | HA | United States | 2015-Aug-25 | A/New York/65/2015 | New York State Department of Health | Centers for Disease Control and Prevention |
| EPI668129 | HA | Argentina | 2015-Jul-20 | A/Argentina/11573/2015 | CEMIC University Hospital | Centers for Disease Control and Prevention |
| EPI668137 | HA | United States | 2015-Sep-01 | A/Minnesota/40/2015 | Minnesota Department of Health | Centers for Disease Control and Prevention |
| EPI668145 | HA | United States | 2015-Sep-08 | A/Alabama/10/2015 | ADPH Bureau of Clinical Laboratories | Centers for Disease Control and Prevention |
| EPI668153 | HA | United States | 2015-Sep-01 | A/Indiana/18/2015 | Indiana State Department of Health Laboratories | Centers for Disease Control and Prevention |
| EPI668161 | HA | United States | 2015-Sep-03 | A/Indiana/26/2015 | Indiana State Department of Health Laboratories | Centers for Disease Control and Prevention |

| | | | | | | |
|---|---|---|---|---|---|---|
| EPI668169 | HA | United States | 2015-Sep-03 | A/Indiana/27/2015 | Indiana State Department of Health Laboratories | Centers for Disease Control and Prevention |
| EPI668184 | HA | United States | 2015-Sep-11 | A/Montana/18/2015 | Montana Public Health Laboratory | Centers for Disease Control and Prevention |
| EPI668192 | HA | United States | 2015-Sep-13 | A/Montana/19/2015 | Montana Public Health Laboratory | Centers for Disease Control and Prevention |
| EPI668200 | HA | United States | 2015-Sep-15 | A/Montana/20/2015 | Montana Public Health Laboratory | Centers for Disease Control and Prevention |
| EPI668208 | HA | United States | 2015-Sep-14 | A/Montana/21/2015 | Montana Public Health Laboratory | Centers for Disease Control and Prevention |
| EPI668216 | HA | United States | 2015-Sep-17 | A/Montana/23/2015 | Montana Public Health Laboratory | Centers for Disease Control and Prevention |
| EPI668231 | HA | United States | 2015-Sep-19 | A/Montana/25/2015 | Montana Public Health Laboratory | Centers for Disease Control and Prevention |
| EPI668239 | HA | United States | 2015-Sep-17 | A/Montana/26/2015 | Montana Public Health Laboratory | Centers for Disease Control and Prevention |
| EPI668247 | HA | United States | 2015-Sep-20 | A/Montana/27/2015 | Montana Public Health Laboratory | Centers for Disease Control and Prevention |
| EPI668255 | HA | United States | 2015-Sep-28 | A/Kentucky/15/2015 | Kentucky Division of Laboratory Services | Centers for Disease Control and Prevention |
| EPI668263 | HA | United States | 2015-Sep-24 | A/Wisconsin/83/2015 | Wisconsin State Laboratory of Hygiene | Centers for Disease Control and Prevention |
| EPI668271 | HA | United States | 2015-Oct-01 | A/Wisconsin/84/2015 | Wisconsin State Laboratory of Hygiene | Centers for Disease Control and Prevention |
| EPI668279 | HA | United States | 2015-Sep-01 | A/Wisconsin/76/2015 | Wisconsin State Laboratory of Hygiene | Centers for Disease Control and Prevention |
| EPI668287 | HA | United States | 2015-Aug-18 | A/Michigan/42/2015 | Michigan Department of Community Health | Centers for Disease Control and Prevention |
| EPI668311 | HA | United States | 2015-Sep-14 | A/Michigan/52/2015 | Michigan Department of Community Health | Centers for Disease Control and Prevention |
| EPI668319 | HA | United States | 2015-Sep-12 | A/Michigan/53/2015 | Michigan Department of Community Health | Centers for Disease Control and Prevention |
| EPI668327 | HA | United States | 2015-Sep-19 | A/Michigan/54/2015 | Michigan Department of Community Health | Centers for Disease Control and Prevention |
| EPI668335 | HA | United States | 2015-Sep-18 | A/Michigan/55/2015 | Michigan Department of Community Health | Centers for Disease Control and Prevention |
| EPI668343 | HA | United States | 2015-Sep-24 | A/Montana/28/2015 | Montana Laboratory Services Bureau | Centers for Disease Control and Prevention |
| EPI668351 | HA | United States | 2015-Sep-26 | A/Montana/29/2015 | Montana Laboratory Services Bureau | Centers for Disease Control and Prevention |
| EPI669734 | HA | United States | 2015-Aug-11 | A/Alaska/140/2015 | Alaska State Virology Lab | Centers for Disease Control and Prevention |
| EPI669976 | HA | Sweden | 2015-Oct-07 | A/Stockholm/43/2015 | | Public Health Agency of Sweden |
| EPI669984 | HA | Sweden | 2015-Oct-09 | A/Stockholm/44/2015 | | Public Health Agency of Sweden |
| EPI669992 | HA | Sweden | 2015-Oct-10 | A/Malmoe/5/2015 | | Public Health Agency of Sweden |
| EPI670000 | HA | Sweden | 2015-Oct-09 | A/Stockholm/45/2015 | | Public Health Agency of Sweden |
| EPI670351 | HA | United States | 2015-Jun-19 | A/Alaska/40/2015 | Alaska State Virology Lab | Centers for Disease Control and Prevention |
| EPI670367 | HA | United States | 2015-Jun-23 | A/Alaska/45/2015 | Alaska State Virology Lab | Centers for Disease Control and Prevention |

| | | | | | | |
|---|---|---|---|---|---|---|
| EPI670388 | HA | United States | 2015-Jun-18 | A/Alaska/55/2015 | Alaska State Virology Lab | Centers for Disease Control and Prevention |
| EPI670396 | HA | United States | 2015-Jun-18 | A/Alaska/56/2015 | Alaska State Virology Lab | Centers for Disease Control and Prevention |
| EPI670404 | HA | United States | 2015-Jun-18 | A/Alaska/57/2015 | Alaska State Virology Lab | Centers for Disease Control and Prevention |
| EPI670412 | HA | United States | 2015-Jun-17 | A/Alaska/59/2015 | Alaska State Virology Lab | Centers for Disease Control and Prevention |
| EPI670420 | HA | United States | 2015-Jun-18 | A/Alaska/61/2015 | Alaska State Virology Lab | Centers for Disease Control and Prevention |
| EPI670428 | HA | United States | 2015-Jun-21 | A/Alaska/62/2015 | Alaska State Virology Lab | Centers for Disease Control and Prevention |
| EPI670436 | HA | United States | 2015-Jun-24 | A/Alaska/63/2015 | Alaska State Virology Lab | Centers for Disease Control and Prevention |
| EPI670444 | HA | United States | 2015-Jun-24 | A/Alaska/64/2015 | Alaska State Virology Lab | Centers for Disease Control and Prevention |
| EPI670459 | HA | Colombia | 2015-Jul-10 | A/Colombia/5433/2015 | Instituto Nacional de Salud de Columbia | Centers for Disease Control and Prevention |
| EPI670467 | HA | United States | 2015-Jun-30 | A/Alaska/96/2015 | Alaska State Virology Lab | Centers for Disease Control and Prevention |
| EPI670475 | HA | United States | 2015-Jul-02 | A/Alaska/97/2015 | Alaska State Virology Lab | Centers for Disease Control and Prevention |
| EPI670483 | HA | United States | 2015-Jul-03 | A/Alaska/98/2015 | Alaska State Virology Lab | Centers for Disease Control and Prevention |
| EPI670491 | HA | United States | 2015-Jul-03 | A/Alaska/99/2015 | Alaska State Virology Lab | Centers for Disease Control and Prevention |
| EPI670499 | HA | United States | 2015-Jul-03 | A/Alaska/100/2015 | Alaska State Virology Lab | Centers for Disease Control and Prevention |
| EPI670507 | HA | United States | 2015-Jul-04 | A/Alaska/101/2015 | Alaska State Virology Lab | Centers for Disease Control and Prevention |
| EPI670515 | HA | United States | 2015-Jun-28 | A/Alaska/102/2015 | Alaska State Virology Lab | Centers for Disease Control and Prevention |
| EPI670523 | HA | United States | 2015-Jul-04 | A/Alaska/103/2015 | Alaska State Virology Lab | Centers for Disease Control and Prevention |
| EPI670531 | HA | United States | 2015-Jul-04 | A/Alaska/112/2015 | Alaska State Virology Lab | Centers for Disease Control and Prevention |
| EPI670539 | HA | United States | 2015-Jul-10 | A/Alaska/113/2015 | Alaska State Virology Lab | Centers for Disease Control and Prevention |
| EPI670547 | HA | United States | 2015-Jul-06 | A/Alaska/114/2015 | Alaska State Virology Lab | Centers for Disease Control and Prevention |
| EPI670555 | HA | United States | 2015-Jul-04 | A/Alaska/115/2015 | Alaska State Virology Lab | Centers for Disease Control and Prevention |
| EPI670563 | HA | United States | 2015-Jun-29 | A/Oregon/14/2015 | Oregon Public Health Laboratory | Centers for Disease Control and Prevention |
| EPI670571 | HA | Brazil | 2015-Apr-16 | A/Acre/134806-IEC/2015 | National Influenza Center | Centers for Disease Control and Prevention |
| EPI670579 | HA | Brazil | 2015-Apr-29 | A/Acre/135094-IEC/2015 | National Influenza Center | Centers for Disease Control and Prevention |
| EPI670587 | HA | Brazil | 2015-May-13 | A/Rio Grande Do Norte/135219-IEC/2015 | National Influenza Center | Centers for Disease Control and Prevention |

| | | | | | | | |
|---|---|---|---|---|---|---|---|
| EPI670595 | HA | Brazil | 2015-May-18 | A/Acre/135356-IEC/2015 | National Influenza Center | Centers for Disease Control and Prevention | |
| EPI670603 | HA | Brazil | 2015-May-29 | A/Rio Grande Do Norte/135497-IEC/2015 | National Influenza Center | Centers for Disease Control and Prevention | |
| EPI670611 | HA | Puerto Rico | 2015-Jul-07 | A/Puerto Rico/03/2015 | Puerto Rico Department of Health | Centers for Disease Control and Prevention | |
| EPI670619 | HA | Puerto Rico | 2015-Aug-11 | A/Puerto Rico/09/2015 | Puerto Rico Department of Health | Centers for Disease Control and Prevention | |
| EPI670627 | HA | French Guiana | 2015-Mar-05 | A/French Guiana/6101/2015 | National Influenza Center French Guiana and French Indies | Centers for Disease Control and Prevention | |
| EPI670642 | HA | Guadeloupe | 2015-Mar-04 | A/Guadeloupe/9095/2015 | National Influenza Center French Guiana and French Indies | Centers for Disease Control and Prevention | |
| EPI670650 | HA | Guadeloupe | 2015-Feb-24 | A/Guadeloupe/9096/2015 | National Influenza Center French Guiana and French Indies | Centers for Disease Control and Prevention | |
| EPI670658 | HA | Guadeloupe | 2015-Feb-25 | A/Guadeloupe/9153/2015 | National Influenza Center French Guiana and French Indies | Centers for Disease Control and Prevention | |
| EPI670670 | HA | French Guiana | 2015-Jun-08 | A/French Guiana/8217/2015 | National Influenza Center French Guiana and French Indies | Centers for Disease Control and Prevention | |
| EPI670676 | HA | French Guiana | 2015-Jun-27 | A/French Guiana/9189/2015 | National Influenza Center French Guiana and French Indies | Centers for Disease Control and Prevention | |
| EPI670691 | HA | Guatemala | 2015-Jun-25 | A/Guatemala/6315/2015 | Laboratorio Nacional De Salud Guatemala | Centers for Disease Control and Prevention | |
| EPI670699 | HA | Guatemala | 2015-Jul-01 | A/Guatemala/6415/2015 | Laboratorio Nacional De Salud Guatemala | Centers for Disease Control and Prevention | |
| EPI670707 | HA | Guatemala | 2015-Mar-07 | A/Guatemala/4153/2015 | Laboratorio Nacional De Salud Guatemala | Centers for Disease Control and Prevention | |
| EPI670715 | HA | Guatemala | 2015-Jun-15 | A/Guatemala/5848/2015 | Laboratorio Nacional De Salud Guatemala | Centers for Disease Control and Prevention | |
| EPI670723 | HA | Guatemala | 2015-Mar-23 | A/Guatemala/8802/2015 | Laboratorio Nacional De Salud Guatemala | Centers for Disease Control and Prevention | |
| EPI670731 | HA | Guatemala | 2015-Apr-21 | A/Guatemala/9151/2015 | Laboratorio Nacional De Salud Guatemala | Centers for Disease Control and Prevention | |
| EPI670745 | HA | Brazil | 2015-May-05 | A/Brazil/4388/2015 | Instituto Adolfo Lutz | Centers for Disease Control and Prevention | |
| EPI670753 | HA | Brazil | 2015-May-21 | A/Brazil/0168/2015 | Instituto Adolfo Lutz | Centers for Disease Control and Prevention | |
| EPI670761 | HA | Brazil | 2015-May-27 | A/Brazil/1790/2015 | Instituto Adolfo Lutz | Centers for Disease Control and Prevention | |
| EPI670769 | HA | Brazil | 2015-May-29 | A/Brazil/2081/2015 | Instituto Adolfo Lutz | Centers for Disease Control and Prevention | Takashita,Emi; Fujisaki,Seiichiro; Shirakura,Masayuki; Watanabe,Shinji; Odagiri,Takato |
| EPI670777 | HA | Brazil | 2015-Apr-24 | A/Brazil/2047/2015 | Instituto Adolfo Lutz | Centers for Disease Control and Prevention | |
| EPI670785 | HA | Brazil | 2015-May-11 | A/Brazil/7588/2015 | Instituto Adolfo Lutz | Centers for Disease Control and Prevention | |
| EPI670797 | HA | United States | 2015-Aug-18 | A/Tennessee/19/2015 | Tennessee Department of Health Laboratory-Nashville | Centers for Disease Control and Prevention | |
| EPI670812 | HA | United States | 2015-Sep-01 | A/New Jersey/52/2015 | New Jersey Department of Health & Senior Services | Centers for Disease Control and Prevention | |
| EPI670820 | HA | United States | 2015-Sep-07 | A/Alaska/206/2015 | Alaska State Virology Lab | Centers for Disease Control and Prevention | |

| | | | | | | |
|---|---|---|---|---|---|---|
| EPI670835 | HA | United States | 2015-Sep-02 | A/Alaska/208/2015 | Alaska State Virology Lab | Centers for Disease Control and Prevention |
| EPI670843 | HA | United States | 2015-Sep-17 | A/Ohio/27/2015 | Ohio Department of Health Laboratories | Centers for Disease Control and Prevention |
| EPI670851 | HA | United States | 2015-Sep-17 | A/Ohio/28/2015 | Ohio Department of Health Laboratories | Centers for Disease Control and Prevention |
| EPI670859 | HA | United States | 2015-Sep-17 | A/Ohio/29/2015 | Ohio Department of Health Laboratories | Centers for Disease Control and Prevention |
| EPI670867 | HA | United States | 2015-Sep-24 | A/Massachusetts/26/2015 | Massachusetts Department of Public Health | Centers for Disease Control and Prevention |
| EPI670875 | HA | United States | 2015-Sep-28 | A/Kentucky/15/2015 | Kentucky Division of Laboratory Services | Centers for Disease Control and Prevention |
| EPI670886 | HA | Bangladesh | 2015-Aug-12 | A/Bangladesh/7004/2015 | Institute of Epidemiology Disease Control and Research (IEDCR) & Bangladesh National Influenza Centre (NIC) | Centers for Disease Control and Prevention |
| EPI670894 | HA | Bangladesh | 2015-Aug-31 | A/Bangladesh/7006/2015 | Institute of Epidemiology Disease Control and Research (IEDCR) & Bangladesh National Influenza Centre (NIC) | Centers for Disease Control and Prevention |
| EPI670902 | HA | Bangladesh | 2015-Aug-24 | A/Bangladesh/3014/2015 | Institute of Epidemiology Disease Control and Research (IEDCR) & Bangladesh National Influenza Centre (NIC) | Centers for Disease Control and Prevention |
| EPI670910 | HA | Bangladesh | 2015-Aug-24 | A/Bangladesh/8016/2015 | Institute of Epidemiology Disease Control and Research (IEDCR) & Bangladesh National Influenza Centre (NIC) | Centers for Disease Control and Prevention |
| EPI670918 | HA | Bangladesh | 2015-Aug-26 | A/Bangladesh/1005/2015 | Institute of Epidemiology Disease Control and Research (IEDCR) & Bangladesh National Influenza Centre (NIC) | Centers for Disease Control and Prevention |
| EPI670926 | HA | Bangladesh | 2015-Aug-10 | A/Bangladesh/803003/2015 | Institute of Epidemiology Disease Control and Research (IEDCR) & Bangladesh National Influenza Centre (NIC) | Centers for Disease Control and Prevention |
| EPI670934 | HA | Bangladesh | 2015-Aug-20 | A/Bangladesh/3007/2015 | Institute of Epidemiology Disease Control and Research (IEDCR) & Bangladesh National Influenza Centre (NIC) | Centers for Disease Control and Prevention |
| EPI670942 | HA | Bangladesh | 2015-Aug-29 | A/Bangladesh/9015/2015 | Institute of Epidemiology Disease Control and Research (IEDCR) & Bangladesh National Influenza Centre (NIC) | Centers for Disease Control and Prevention |
| EPI670950 | HA | United States | 2015-Oct-03 | A/Florida/74/2015 | Florida Department of Health-Jacksonville | Centers for Disease Control and Prevention |
| EPI670958 | HA | United States | 2015-Oct-01 | A/Florida/75/2015 | Florida Department of Health-Jacksonville | Centers for Disease Control and Prevention |
| EPI670966 | HA | United States | 2015-Sep-28 | A/North Carolina/32/2015 | North Carolina State Laboratory of Public Health | Centers for Disease Control and Prevention |

| | | | | | | |
|---|---|---|---|---|---|---|
| EPI670974 | HA | United States | 2015-Sep-28 | A/North Carolina/32/2015 | North Carolina State Laboratory of Public Health | Centers for Disease Control and Prevention |
| EPI670982 | HA | United States | 2015-Oct-07 | A/Maine/39/2015 | Maine Health and Environmental Testing Laboratory | Centers for Disease Control and Prevention |
| EPI670990 | HA | United States | 2015-Oct-09 | A/Maine/40/2015 | Maine Health and Environmental Testing Laboratory | Centers for Disease Control and Prevention |
| EPI670998 | HA | United States | 2015-Oct-12 | A/Maine/41/2015 | Maine Health and Environmental Testing Laboratory | Centers for Disease Control and Prevention |
| EPI671006 | HA | United States | 2015-Oct-04 | A/Missouri/41/2015 | Missouri Department. of Health & Senior Services | Centers for Disease Control and Prevention |
| EPI671014 | HA | United States | 2015-Oct-04 | A/Missouri/42/2015 | Missouri Department. of Health & Senior Services | Centers for Disease Control and Prevention |
| EPI671022 | HA | United States | 2015-Oct-06 | A/Montana/30/2015 | Montana Laboratory Services Bureau | Centers for Disease Control and Prevention |
| EPI671030 | HA | United States | 2015-Oct-07 | A/Montana/31/2015 | Montana Laboratory Services Bureau | Centers for Disease Control and Prevention |
| EPI671038 | HA | United States | 2015-Oct-12 | A/Montana/32/2015 | Montana Laboratory Services Bureau | Centers for Disease Control and Prevention |
| EPI671046 | HA | United States | 2015-Oct-04 | A/Arizona/20/2015 | South Carolina Department of Health | Centers for Disease Control and Prevention |
| EPI671054 | HA | United States | 2015-Oct-02 | A/Arizona/21/2015 | Arizona Department of Health Services | Centers for Disease Control and Prevention |
| EPI671062 | HA | United States | 2015-Oct-07 | A/Arizona/22/2015 | South Carolina Department of Health | Centers for Disease Control and Prevention |
| EPI671070 | HA | United States | 2015-Oct-08 | A/Oregon/19/2015 | Oregon Public Health Laboratory | Centers for Disease Control and Prevention |
| EPI671078 | HA | United States | 2015-Oct-09 | A/Oregon/20/2015 | Oregon Public Health Laboratory | Centers for Disease Control and Prevention |
| EPI671086 | HA | United States | 2015-Oct-09 | A/Oregon/21/2015 | Oregon Public Health Laboratory | Centers for Disease Control and Prevention |
| EPI671094 | HA | United States | 2015-Oct-09 | A/Oregon/22/2015 | Oregon Public Health Laboratory | Centers for Disease Control and Prevention |
| EPI671102 | HA | United States | 2015-Oct-12 | A/Oregon/23/2015 | Oregon Public Health Laboratory | Centers for Disease Control and Prevention |
| EPI671110 | HA | United States | 2015-Oct-12 | A/Oregon/24/2015 | Oregon Public Health Laboratory | Centers for Disease Control and Prevention |
| EPI671118 | HA | United States | 2015-Oct-02 | A/Alaska/253/2015 | Alaska State Virology Lab | Centers for Disease Control and Prevention |
| EPI671126 | HA | United States | 2015-Oct-05 | A/Alaska/254/2015 | Alaska State Virology Lab | Centers for Disease Control and Prevention |
| EPI671134 | HA | United States | 2015-Oct-06 | A/Alaska/255/2015 | Alaska State Virology Lab | Centers for Disease Control and Prevention |
| EPI671142 | HA | United States | 2015-Oct-05 | A/Alaska/256/2015 | Alaska State Virology Lab | Centers for Disease Control and Prevention |
| EPI671150 | HA | United States | 2015-Sep-01 | A/Washington/40/2015 | Washington State Public Health Laboratory | Centers for Disease Control and Prevention |
| EPI671158 | HA | United States | 2015-Aug-31 | A/Washington/41/2015 | Washington State Public Health Laboratory | Centers for Disease Control and Prevention |
| EPI671166 | HA | United States | 2015-Aug-31 | A/Washington/42/2015 | Washington State Public Health Laboratory | Centers for Disease Control and Prevention |

| | | | | | | | |
|---|---|---|---|---|---|---|---|
| EPI671174 | HA | United States | 2015-Aug-31 | A/Washington/43/2015 | Washington State Public Health Laboratory | Centers for Disease Control and Prevention | |
| EPI671182 | HA | United States | 2015-Sep-07 | A/Washington/44/2015 | Washington State Public Health Laboratory | Centers for Disease Control and Prevention | |
| EPI671190 | HA | United States | 2015-Sep-22 | A/Wyoming/18/2015 | Wyoming Public Health Laboratory | Centers for Disease Control and Prevention | |
| EPI671198 | HA | United States | 2015-Sep-24 | A/Montana/28/2015 | Montana Laboratory Services Bureau | Centers for Disease Control and Prevention | |
| EPI671206 | HA | United States | 2015-Sep-26 | A/Montana/29/2015 | Montana Laboratory Services Bureau | Centers for Disease Control and Prevention | |
| EPI671214 | HA | United States | 2015-Sep-14 | A/Minnesota/42/2015 | Minnesota Department of Health | Centers for Disease Control and Prevention | |
| EPI671222 | HA | United States | 2015-Sep-17 | A/Minnesota/43/2015 | Minnesota Department of Health | Centers for Disease Control and Prevention | |
| EPI671230 | HA | United States | 2015-Sep-15 | A/Hawaii/58/2015 | State of Hawaii Department of Health | Centers for Disease Control and Prevention | |
| EPI671238 | HA | United States | 2015-Sep-26 | A/Hawaii/60/2015 | State of Hawaii Department of Health | Centers for Disease Control and Prevention | |
| EPI671243 | HA | United States | 2015-Sep-28 | A/Hawaii/61/2015 | State of Hawaii Department of Health | Centers for Disease Control and Prevention | |
| EPI671251 | HA | United States | 2015-Sep-18 | A/Idaho/25/2015 | State of Idaho Bureau of Laboratories | Centers for Disease Control and Prevention | |
| EPI671266 | HA | United States | 2015-Oct-06 | A/Texas/69/2015 | Texas Department of State Health Services-Laboratory Services | Centers for Disease Control and Prevention | |
| EPI671273 | HA | United States | 2015-Oct-06 | A/Texas/70/2015 | Texas Department of State Health Services-Laboratory Services | Centers for Disease Control and Prevention | |
| EPI671281 | HA | United States | 2015-Aug-19 | A/Vermont/08/2015 | Vermont Department of Health Laboratory | Centers for Disease Control and Prevention | |
| EPI671289 | HA | United States | 2015-Sep-30 | A/Vermont/09/2015 | Vermont Department of Health Laboratory | Centers for Disease Control and Prevention | |
| EPI671297 | HA | United States | 2015-Oct-02 | A/Pennsylvania/36/2015 | Pennsylvania Department of Health | Centers for Disease Control and Prevention | |
| EPI671305 | HA | United States | 2015-Oct-10 | A/New Hampshire/41/2015 | New Hampshire Public Health Laboratories | Centers for Disease Control and Prevention | Galiano,M. |
| EPI673079 | HA | Japan | 2015-Mar-19 | A/KOBE/262/2015 | Kobe Institute of Health | National Institute of Infectious Diseases (NIID) | Galiano,M. |
| EPI673591 | HA | Bangladesh | 2015-Aug-10 | A/Bangladesh/5968/2015 | Institute of Epidemiology Disease Control and Research (IEDCR) & Bangladesh National Influenza Centre (NIC) | Centers for Disease Control and Prevention | Galiano,M. |
| EPI673599 | HA | United States | 2015-Sep-09 | A/California/85/2015 | California Department of Health Services | Centers for Disease Control and Prevention | Galiano,M. |
| EPI673607 | HA | Bangladesh | 2015-Aug-20 | A/Bangladesh/22014/2015 | Institute of Epidemiology Disease Control and Research (IEDCR) & Bangladesh National Influenza Centre (NIC) | Centers for Disease Control and Prevention | Galiano,M. |
| EPI673615 | HA | United States | 2015-Oct-01 | A/California/91/2015 | California Department of Health Services | Centers for Disease Control and Prevention | Galiano,M. |
| EPI673623 | HA | United States | 2015-Oct-01 | A/California/87/2015 | California Department of Health Services | Centers for Disease Control and Prevention | Galiano,M. |

| | | | | | | | |
|---|---|---|---|---|---|---|---|
| EPI673631 | HA | United States | 2015-Sep-24 | A/California/89/2015 | California Department of Health Services | Centers for Disease Control and Prevention | Galiano,M. |
| EPI673639 | HA | Guatemala | 2015-Mar-19 | A/Guatemala/8147/2015 | Laboratorio Nacional De Salud Guatemala | Centers for Disease Control and Prevention | |
| EPI673647 | HA | United States | 2015-Sep-10 | A/California/83/2015 | California Department of Health Services | Centers for Disease Control and Prevention | |
| EPI673655 | HA | United States | 2015-Sep-05 | A/California/86/2015 | California Department of Health Services | Centers for Disease Control and Prevention | |
| EPI673663 | HA | Bangladesh | 2015-Aug-17 | A/Bangladesh/3937/2015 | Institute of Epidemiology Disease Control and Research (IEDCR) & Bangladesh National Influenza Centre (NIC) | Centers for Disease Control and Prevention | |
| EPI673670 | HA | Bangladesh | 2015-Aug-11 | A/Bangladesh/12004/2015 | Institute of Epidemiology Disease Control and Research (IEDCR) & Bangladesh National Influenza Centre (NIC) | Centers for Disease Control and Prevention | |
| EPI673678 | HA | United States | 2015-Jan-01 | A/South Carolina/09/2015 | South Carolina Department of Health | Centers for Disease Control and Prevention | |
| EPI673686 | HA | Guatemala | 2015-Apr-16 | A/Guatemala/363/2015 | Laboratorio Nacional De Salud Guatemala | Centers for Disease Control and Prevention | |
| EPI673701 | HA | United States | 2015-Oct-14 | A/Iowa/50/2015 | Iowa State Hygienic Laboratory | Centers for Disease Control and Prevention | |
| EPI673709 | HA | United States | 2015-Sep-13 | A/New Mexico/26/2015 | New Mexico Department of Health | Centers for Disease Control and Prevention | |
| EPI673724 | HA | United States | 2015-Oct-14 | A/Delaware/41/2015 | Delaware Public Health Lab | Centers for Disease Control and Prevention | |
| EPI673732 | HA | United States | 2015-Oct-12 | A/Arkansas/09/2015 | Arkansas Department of Health | Centers for Disease Control and Prevention | |
| EPI673740 | HA | United States | 2015-Oct-12 | A/Arkansas/08/2015 | Arkansas Department of Health | Centers for Disease Control and Prevention | |
| EPI673748 | HA | United States | 2015-Oct-16 | A/North Carolina/33/2015 | North Carolina State Laboratory of Public Health | Centers for Disease Control and Prevention | |
| EPI673756 | HA | United States | 2015-Oct-19 | A/Maine/43/2015 | Maine Health and Environmental Testing Laboratory | Centers for Disease Control and Prevention | |
| EPI673764 | HA | United States | 2015-Oct-15 | A/Maine/42/2015 | Maine Health and Environmental Testing Laboratory | Centers for Disease Control and Prevention | |
| EPI673772 | HA | United States | 2015-Oct-10 | A/Massachusetts/28/2015 | Massachusetts Department of Public Health | Centers for Disease Control and Prevention | |
| EPI673784 | HA | United States | 2015-Aug-13 | A/Mississippi/13/2015 | Mississippi Public Health Laboratory | Centers for Disease Control and Prevention | |
| EPI673792 | HA | United States | 2015-Jul-06 | A/Alaska/116/2015 | Alaska State Virology Lab | Centers for Disease Control and Prevention | |
| EPI673800 | HA | United States | 2015-Oct-04 | A/Washington/49/2015 | Washington State Public Health Laboratory | Centers for Disease Control and Prevention | |
| EPI673808 | HA | United States | 2015-Oct-16 | A/Wisconsin/85/2015 | Wisconsin State Laboratory of Hygiene | Centers for Disease Control and Prevention | |
| EPI673816 | HA | United States | 2015-Oct-11 | A/Washington/50/2015 | Washington State Public Health Laboratory | Centers for Disease Control and Prevention | |
| EPI673824 | HA | United States | 2015-Oct-08 | A/West Virginia/23/2015 | West Virginia Office of Laboratory Services | Centers for Disease Control and Prevention | |

| | | | | | | |
|---|---|---|---|---|---|---|
| EPI673832 | HA | United States | 2015-Oct-19 | A/Pennsylvania/42/2015 | Pennsylvania Department of Health | Centers for Disease Control and Prevention |
| EPI673840 | HA | United States | 2015-Oct-05 | A/Florida/81/2015 | Florida Department of Health-Tampa | Centers for Disease Control and Prevention |
| EPI673853 | HA | Bangladesh | 2015-Aug-29 | A/Bangladesh/5011/2015 | Institute of Epidemiology Disease Control and Research (IEDCR) & Bangladesh National Influenza Centre (NIC) | Centers for Disease Control and Prevention |
| EPI673861 | HA | Guatemala | 2015-Mar-11 | A/Guatemala/9877/2015 | Laboratorio Nacional De Salud Guatemala | Centers for Disease Control and Prevention |
| EPI673869 | HA | United States | 2015-Oct-05 | A/Hawaii/67/2015 | State of Hawaii Department of Health | Centers for Disease Control and Prevention |
| EPI673877 | HA | United States | 2015-Oct-04 | A/Hawaii/65/2015 | State of Hawaii Department of Health | Centers for Disease Control and Prevention |
| EPI673885 | HA | United States | 2015-Oct-03 | A/Hawaii/66/2015 | State of Hawaii Department of Health | Centers for Disease Control and Prevention |
| EPI673896 | HA | United States | 2015-Oct-05 | A/Hawaii/68/2015 | State of Hawaii Department of Health | Centers for Disease Control and Prevention |
| EPI673904 | HA | Guatemala | 2015-Mar-09 | A/Guatemala/8136/2015 | Laboratorio Nacional De Salud Guatemala | Centers for Disease Control and Prevention |
| EPI674432 | HA | Lao, People's Democratic Republic | 2015-Jun-29 | A/Laos/I660/2015 | Ministry of Health Laos | National Institute of Infectious Diseases (NIID) |
| EPI674440 | HA | Lao, People's Democratic Republic | 2015-Jul-07 | A/Laos/I706/2015 | Ministry of Health Laos | National Institute of Infectious Diseases (NIID) |
| EPI674448 | HA | Lao, People's Democratic Republic | 2015-Jul-14 | A/Laos/I730/2015 | Ministry of Health Laos | National Institute of Infectious Diseases (NIID) |
| EPI674456 | HA | Lao, People's Democratic Republic | 2015-Jul-22 | A/Laos/I751/2015 | Ministry of Health Laos | National Institute of Infectious Diseases (NIID) |
| EPI674464 | HA | Lao, People's Democratic Republic | 2015-Jul-27 | A/Laos/I784/2015 | Ministry of Health Laos | National Institute of Infectious Diseases (NIID) |
| EPI674472 | HA | Lao, People's Democratic Republic | 2015-Aug-03 | A/Laos/I805/2015 | Ministry of Health Laos | National Institute of Infectious Diseases (NIID) |
| EPI674480 | HA | Lao, People's Democratic Republic | 2015-Aug-03 | A/Laos/I810/2015 | Ministry of Health Laos | National Institute of Infectious Diseases (NIID) |
| EPI674488 | HA | Lao, People's Democratic Republic | 2015-Aug-10 | A/Laos/I847/2015 | Ministry of Health Laos | National Institute of Infectious Diseases (NIID) |
| EPI674496 | HA | Lao, People's Democratic Republic | 2015-Aug-23 | A/Laos/I889/2015 | Ministry of Health Laos | National Institute of Infectious Diseases (NIID) |
| EPI674504 | HA | Lao, People's Democratic Republic | 2015-Sep-02 | A/Laos/I917/2015 | Ministry of Health Laos | National Institute of Infectious Diseases (NIID) |

| | | | | | | |
|---|---|---|---|---|---|---|
| EPI674512 | HA | Lao, People's Democratic Republic | 2015-Sep-07 | A/Laos/I924/2015 | Ministry of Health Laos | National Institute of Infectious Diseases (NIID) |
| EPI674520 | HA | Lao, People's Democratic Republic | 2015-Sep-09 | A/Laos/I929/2015 | Ministry of Health Laos | National Institute of Infectious Diseases (NIID) |
| EPI674528 | HA | Lao, People's Democratic Republic | 2015-Sep-07 | A/Laos/I935/2015 | Ministry of Health Laos | National Institute of Infectious Diseases (NIID) |
| EPI674536 | HA | Lao, People's Democratic Republic | 2015-Sep-07 | A/Laos/I936/2015 | Ministry of Health Laos | National Institute of Infectious Diseases (NIID) |
| EPI674544 | HA | Lao, People's Democratic Republic | 2015-Sep-08 | A/Laos/I941/2015 | Ministry of Health Laos | National Institute of Infectious Diseases (NIID) |
| EPI674560 | HA | Lao, People's Democratic Republic | 2015-Aug-21 | A/Laos/O587/2015 | Ministry of Health Laos | National Institute of Infectious Diseases (NIID) |
| EPI674568 | HA | Lao, People's Democratic Republic | 2015-Aug-22 | A/Laos/SA358/2015 | Ministry of Health Laos | National Institute of Infectious Diseases (NIID) |
| EPI674576 | HA | Lao, People's Democratic Republic | 2015-Sep-07 | A/Laos/SA373/2015 | Ministry of Health Laos | National Institute of Infectious Diseases (NIID) |
| EPI674584 | HA | Lao, People's Democratic Republic | 2015-Sep-07 | A/Laos/SA380/2015 | Ministry of Health Laos | National Institute of Infectious Diseases (NIID) |
| EPI674761 | HA | Sweden | 2015-Oct-30 | A/Stockholm/54/2015 | | Public Health Agency of Sweden |
| EPI675083 | HA | United States | 2014-Dec-29 | A/Virginia/51/2014 | Virginia Division of Consolidated Laboratories | Centers for Disease Control and Prevention |
| EPI675090 | HA | Argentina | 2015-Jul-15 | A/Argentina/116/2015 | Instituto Nacional de Enfermedades Infecciosas | Centers for Disease Control and Prevention |
| EPI675098 | HA | United States | 2015-Oct-09 | A/Oklahoma/20/2015 | Oklahoma State Department of Health | Centers for Disease Control and Prevention |
| EPI675118 | HA | Peru | 2015-Aug-25 | A/Peru/6915/2015 | Laboratorio de Referencia Nacional Virus Respiratorios, Instituto Nacional de Salud | Centers for Disease Control and Prevention |
| EPI675126 | HA | Kyrgyzstan | 2015-Feb-12 | A/Kyrgyzstan/82/2015 | National Virology Laboratory, Center Microbiological Investigations | Centers for Disease Control and Prevention |
| EPI675139 | HA | United States | 2015-Jun-15 | A/Alaska/51/2015 | Alaska State Virology Lab | Centers for Disease Control and Prevention |
| EPI675153 | HA | United States | 2015-Jun-17 | A/Alaska/58/2015 | Alaska State Virology Lab | Centers for Disease Control and Prevention |
| EPI675161 | HA | Brazil | 2015-May-05 | A/Brazil/4390/2015 | Instituto Adolfo Lutz | Centers for Disease Control and Prevention |
| EPI675176 | HA | United States | 2015-Jun-15 | A/Alaska/60/2015 | Alaska State Virology Lab | Centers for Disease Control and Prevention |
| EPI675184 | HA | Brazil | 2015-May-22 | A/Brazil/2002/2015 | Instituto Adolfo Lutz | Centers for Disease Control and Prevention |

| | | | | | | | |
|---|---|---|---|---|---|---|---|
| EPI675192 | HA | United States | 2015-Sep-29 | A/Michigan/62/2015 | Michigan Department of Community Health | Centers for Disease Control and Prevention | |
| EPI675206 | HA | United States | 2015-Oct-12 | A/Alaska/258/2015 | Alaska State Virology Lab | Centers for Disease Control and Prevention | |
| EPI675214 | HA | United States | 2015-Oct-20 | A/Alaska/259/2015 | Alaska State Virology Lab | Centers for Disease Control and Prevention | |
| EPI675222 | HA | United States | 2015-Oct-11 | A/Alaska/257/2015 | Alaska State Virology Lab | Centers for Disease Control and Prevention | Galiano,M. |
| EPI675230 | HA | Canada | 2015-Oct-10 | A/Alberta/RV2387/2015 | National Microbiology Laboratory, Health Canada | Centers for Disease Control and Prevention | |
| EPI675238 | HA | United States | 2015-Sep-28 | A/Indiana/28/2015 | Indiana State Department of Health Laboratories | Centers for Disease Control and Prevention | |
| EPI675246 | HA | Canada | 2015-Oct-01 | A/Alberta/RV2382/2015 | National Microbiology Laboratory, Health Canada | Centers for Disease Control and Prevention | |
| EPI675254 | HA | Canada | 2015-Oct-10 | A/Alberta/RV2386/2015 | National Microbiology Laboratory, Health Canada | Centers for Disease Control and Prevention | |
| EPI675262 | HA | Canada | 2015-Oct-10 | A/Alberta/RV2385/2015 | National Microbiology Laboratory, Health Canada | Centers for Disease Control and Prevention | |
| EPI675270 | HA | Canada | 2015-Sep-09 | A/British Columbia/RV2356/2015 | National Microbiology Laboratory, Health Canada | Centers for Disease Control and Prevention | |
| EPI675278 | HA | Canada | 2015-Sep-11 | A/British Columbia/RV2357/2015 | National Microbiology Laboratory, Health Canada | Centers for Disease Control and Prevention | |
| EPI675286 | HA | Canada | 2015-Sep-14 | A/British Columbia/RV2358/2015 | National Microbiology Laboratory, Health Canada | Centers for Disease Control and Prevention | |
| EPI675294 | HA | Canada | 2015-Sep-12 | A/Alberta/RV2339/2015 | National Microbiology Laboratory, Health Canada | Centers for Disease Control and Prevention | |
| EPI675302 | HA | Kyrgyzstan | 2015-Jan-09 | A/Kyrgyzstan/10/2015 | National Virology Laboratory, Center Microbiological Investigations | Centers for Disease Control and Prevention | |
| EPI675386 | HA | Canada | 2015-Sep-09 | A/Alberta/RV2338/2015 | National Microbiology Laboratory, Health Canada | Centers for Disease Control and Prevention | |
| EPI675391 | HA | Canada | 2015-Oct-09 | A/Alberta/RV2383/2015 | National Microbiology Laboratory, Health Canada | Centers for Disease Control and Prevention | |
| EPI675398 | HA | United States | 2015-Oct-21 | A/Colorado/16/2015 | Colorado Department of Health Lab | Centers for Disease Control and Prevention | |
| EPI675401 | HA | United States | 2015-Oct-28 | A/Florida/80/2015 | Florida Department of Health-Jacksonville | Centers for Disease Control and Prevention | |
| EPI675408 | HA | Luxembourg | 2015-Feb-23 | A/Luxembourg/15008843/2015 | Laboratoire National de Santé | Laboratoire National de Santé | |
| EPI675417 | HA | United States | 2015-Nov-11 | A/Texas/71/2015 | Texas Department of State Health Services-Laboratory Services | Centers for Disease Control and Prevention | |
| EPI675425 | HA | United States | 2015-Oct-29 | A/Maryland/18/2015 | Maryland Department of Health and Mental Hygiene | Centers for Disease Control and Prevention | |
| EPI675433 | HA | United States | 2015-Nov-01 | A/Maryland/19/2015 | Maryland Department of Health and Mental Hygiene | Centers for Disease Control and Prevention | |
| EPI675444 | HA | United States | 2015-Nov-11 | A/Louisiana/07/2015 | Louisiana Department of Health and Hospitals | Centers for Disease Control and Prevention | |
| EPI675452 | HA | United States | 2015-Oct-07 | A/Washington/55/2015 | Washington State Public Health Laboratory | Centers for Disease Control and Prevention | |
| EPI675460 | HA | United States | 2015-Oct-20 | A/Washington/56/2015 | Washington State Public Health Laboratory | Centers for Disease Control and Prevention | |

| | | | | | | | |
|---|---|---|---|---|---|---|---|
| EPI675468 | HA | United States | 2015-Oct-19 | A/Wisconsin/86/2015 | Wisconsin State Laboratory of Hygiene | Centers for Disease Control and Prevention | |
| EPI675489 | HA | Argentina | 2015-Jul-23 | A/Argentina/149/2015 | Instituto Nacional de Enfermedades Infecciosas | Centers for Disease Control and Prevention | |
| EPI675497 | HA | Argentina | 2015-Jul-25 | A/Argentina/156/2015 | Instituto Nacional de Enfermedades Infecciosas | Centers for Disease Control and Prevention | |
| EPI675512 | HA | Argentina | 2015-Jul-27 | A/Argentina/157/2015 | Instituto Nacional de Enfermedades Infecciosas | Centers for Disease Control and Prevention | |
| EPI675520 | HA | Argentina | 2015-Jul-27 | A/Argentina/158/2015 | Instituto Nacional de Enfermedades Infecciosas | Centers for Disease Control and Prevention | |
| EPI675792 | HA | New Zealand | 2015-Aug-13 | A/New Zealand/1703/2015 | Institute of Environmental Science & Research | Centers for Disease Control and Prevention | |
| EPI676550 | HA | Canada | 2015-Sep-18 | A/Alberta/RV2362/2015 | National Microbiology Laboratory, Health Canada | Centers for Disease Control and Prevention | |
| EPI676558 | HA | Canada | 2015-Sep-18 | A/Alberta/RV2363/2015 | National Microbiology Laboratory, Health Canada | Centers for Disease Control and Prevention | |
| EPI676566 | HA | Canada | 2015-Sep-17 | A/Ontario/RV2371/2015 | National Microbiology Laboratory, Health Canada | Centers for Disease Control and Prevention | |
| EPI676574 | HA | Canada | 2015-Sep-29 | A/Alberta/RV2372/2015 | National Microbiology Laboratory, Health Canada | Centers for Disease Control and Prevention | |
| EPI676582 | HA | Canada | 2015-Sep-28 | A/Alberta/RV2373/2015 | National Microbiology Laboratory, Health Canada | Centers for Disease Control and Prevention | |
| EPI676596 | HA | Canada | 2015-Oct-07 | A/Ontario/RV2377/2015 | National Microbiology Laboratory, Health Canada | Centers for Disease Control and Prevention | |
| EPI676604 | HA | Canada | 2015-Oct-07 | A/Ontario/RV2377/2015 | National Microbiology Laboratory, Health Canada | Centers for Disease Control and Prevention | |
| EPI676612 | HA | Canada | 2015-Oct-07 | A/Ontario/RV2378/2015 | National Microbiology Laboratory, Health Canada | Centers for Disease Control and Prevention | |
| EPI676620 | HA | Canada | 2015-Oct-07 | A/Ontario/RV2378/2015 | National Microbiology Laboratory, Health Canada | Centers for Disease Control and Prevention | |
| EPI676628 | HA | Canada | 2015-Oct-07 | A/Ontario/RV2379/2015 | National Microbiology Laboratory, Health Canada | Centers for Disease Control and Prevention | |
| EPI676636 | HA | Canada | 2015-Oct-07 | A/Ontario/RV2379/2015 | National Microbiology Laboratory, Health Canada | Centers for Disease Control and Prevention | |
| EPI676644 | HA | Canada | 2015-Oct-07 | A/Ontario/RV2380/2015 | National Microbiology Laboratory, Health Canada | Centers for Disease Control and Prevention | |
| EPI676652 | HA | Canada | 2015-Oct-07 | A/Ontario/RV2380/2015 | National Microbiology Laboratory, Health Canada | Centers for Disease Control and Prevention | |
| EPI676660 | HA | United States | 2015-Aug-01 | A/South Dakota/36/2015 | South Dakota Public Health Lab | Centers for Disease Control and Prevention | |
| EPI676668 | HA | United States | 2015-Aug-11 | A/South Dakota/37/2015 | South Dakota Public Health Lab | Centers for Disease Control and Prevention | |
| EPI676676 | HA | United States | 2015-Oct-13 | A/South Dakota/39/2015 | South Dakota Public Health Lab | Centers for Disease Control and Prevention | Takashita,Emi; Fujisaki,Seiichiro; Shirakura,Masayuki; Watanabe,Shinji; Odagiri,Takato |
| EPI676684 | HA | United States | 2015-Nov-03 | A/New York/71/2015 | New York State Department of Health | Centers for Disease Control and Prevention | Takashita,Emi; Fujisaki,Seiichiro; Shirakura,Masayuki; Watanabe,Shinji; Odagiri,Takato |
| EPI676692 | HA | United States | 2015-Nov-06 | A/Florida/84/2015 | Florida Department of Health-Tampa | Centers for Disease Control and Prevention | Takashita,Emi; Fujisaki,Seiichiro; Shirakura,Masayuki; Watanabe,Shinji; Odagiri,Takato |

| | | | | | | | |
|---|---|---|---|---|---|---|---|
| EPI676700 | HA | United States | 2015-Oct-19 | A/Oregon/25/2015 | Oregon Public Health Laboratory | Centers for Disease Control and Prevention | Takashita,Emi; Fujisaki,Seiichiro; Shirakura,Masayuki; Watanabe,Shinji; Odagiri,Takato |
| EPI676708 | HA | United States | 2015-Oct-19 | A/Oregon/25/2015 | Oregon Public Health Laboratory | Centers for Disease Control and Prevention | Takashita,Emi; Fujisaki,Seiichiro; Shirakura,Masayuki; Watanabe,Shinji; Odagiri,Takato |
| EPI676716 | HA | United States | 2015-Oct-23 | A/Oregon/27/2015 | Oregon Public Health Laboratory | Centers for Disease Control and Prevention | |
| EPI676724 | HA | Puerto Rico | 2015-Jul-03 | A/Puerto Rico/02/2015 | Puerto Rico Department of Health | Centers for Disease Control and Prevention | |
| EPI676732 | HA | Argentina | 2015-Jul-24 | A/Argentina/163/2015 | Instituto Nacional de Enfermedades Infecciosas | Centers for Disease Control and Prevention | |
| EPI676745 | HA | Argentina | 2015-Jul-24 | A/Argentina/169/2015 | Instituto Nacional de Enfermedades Infecciosas | Centers for Disease Control and Prevention | |
| EPI676758 | HA | Argentina | 2015-Jul-28 | A/Argentina/181/2015 | Instituto Nacional de Enfermedades Infecciosas | Centers for Disease Control and Prevention | |
| EPI676766 | HA | United States | 2015-Sep-04 | A/Nebraska/18/2015 | Nebraska Public Health Lab | Centers for Disease Control and Prevention | |
| EPI676774 | HA | United States | 2015-Sep-07 | A/Alaska/210/2015 | Alaska State Virology Lab | Centers for Disease Control and Prevention | |
| EPI676788 | HA | United States | 2015-Aug-30 | A/Alaska/212/2015 | Alaska State Virology Lab | Centers for Disease Control and Prevention | |
| EPI676796 | HA | United States | 2015-Aug-30 | A/Alaska/213/2015 | Alaska State Virology Lab | Centers for Disease Control and Prevention | |
| EPI676804 | HA | United States | 2015-Aug-31 | A/Alaska/214/2015 | Alaska State Virology Lab | Centers for Disease Control and Prevention | |
| EPI676812 | HA | United States | 2015-Sep-04 | A/Alaska/215/2015 | Alaska State Virology Lab | Centers for Disease Control and Prevention | |
| EPI676820 | HA | United States | 2015-Jun-16 | A/Washington/45/2015 | Washington State Public Health Laboratory | Centers for Disease Control and Prevention | |
| EPI676828 | HA | United States | 2015-Sep-19 | A/Washington/47/2015 | Washington State Public Health Laboratory | Centers for Disease Control and Prevention | |
| EPI676836 | HA | United States | 2015-Sep-19 | A/Washington/48/2015 | Washington State Public Health Laboratory | Centers for Disease Control and Prevention | |
| EPI676844 | HA | China | 2015-May-06 | A/Liaoning-Baita/1293/2015 | WHO Chinese National Influenza Center | Centers for Disease Control and Prevention | |
| EPI676857 | HA | China | 2015-Jun-15 | A/Hunan-Wuling/11088/2015 | WHO Chinese National Influenza Center | Centers for Disease Control and Prevention | |
| EPI676865 | HA | China | 2015-Mar-03 | A/Jilin-Yanji/1179/2015 | WHO Chinese National Influenza Center | Centers for Disease Control and Prevention | |
| EPI676873 | HA | United States | 2015-Sep-20 | A/Arizona/15/2015 | South Carolina Department of Health | Centers for Disease Control and Prevention | |
| EPI676881 | HA | United States | 2015-Sep-28 | A/Arizona/16/2015 | Arizona Department of Health Services | Centers for Disease Control and Prevention | |
| EPI676889 | HA | United States | 2015-Sep-28 | A/Arizona/17/2015 | Arizona Department of Health Services | Centers for Disease Control and Prevention | |
| EPI676897 | HA | United States | 2015-Sep-30 | A/Arizona/18/2015 | Arizona Department of Health Services | Centers for Disease Control and Prevention | |
| EPI676905 | HA | United States | 2015-Aug-30 | A/California/84/2015 | California Department of Health Services | Centers for Disease Control and Prevention | |

| | | | | | | |
|---|---|---|---|---|---|---|
| EPI676913 | HA | United States | 2015-Jan-01 | A/Georgia/12/2015 | Georgia Public Health Laboratory | Centers for Disease Control and Prevention |
| EPI676921 | HA | United States | 2015-Oct-09 | A/Georgia/13/2015 | Georgia Public Health Laboratory | Centers for Disease Control and Prevention |
| EPI676929 | HA | United States | 2015-Oct-20 | A/Florida/76/2015 | Florida Department of Health-Tampa | Centers for Disease Control and Prevention |
| EPI676937 | HA | United States | 2015-Oct-20 | A/Florida/76/2015 | Florida Department of Health-Tampa | Centers for Disease Control and Prevention |
| EPI676945 | HA | United States | 2015-Oct-24 | A/Florida/77/2015 | Florida Department of Health-Tampa | Centers for Disease Control and Prevention |
| EPI676953 | HA | United States | 2015-Oct-24 | A/Florida/77/2015 | Florida Department of Health-Tampa | Centers for Disease Control and Prevention |
| EPI676961 | HA | United States | 2015-Oct-22 | A/Florida/78/2015 | Florida Department of Health-Jacksonville | Centers for Disease Control and Prevention |
| EPI676969 | HA | United States | 2015-Oct-21 | A/Iowa/44/2015 | Iowa State Hygienic Laboratory | Centers for Disease Control and Prevention |
| EPI676977 | HA | United States | 2015-Oct-21 | A/Iowa/44/2015 | Iowa State Hygienic Laboratory | Centers for Disease Control and Prevention |
| EPI676985 | HA | United States | 2015-Oct-22 | A/Iowa/45/2015 | Iowa State Hygienic Laboratory | Centers for Disease Control and Prevention |
| EPI676993 | HA | United States | 2015-Oct-22 | A/Iowa/45/2015 | Iowa State Hygienic Laboratory | Centers for Disease Control and Prevention |
| EPI677001 | HA | United States | 2015-Oct-21 | A/Iowa/46/2015 | Iowa State Hygienic Laboratory | Centers for Disease Control and Prevention |
| EPI677009 | HA | United States | 2015-Oct-21 | A/Iowa/46/2015 | Iowa State Hygienic Laboratory | Centers for Disease Control and Prevention |
| EPI677017 | HA | United States | 2015-Oct-22 | A/Iowa/47/2015 | Iowa State Hygienic Laboratory | Centers for Disease Control and Prevention |
| EPI677025 | HA | United States | 2015-Oct-24 | A/Iowa/48/2015 | Iowa State Hygienic Laboratory | Centers for Disease Control and Prevention |
| EPI677033 | HA | United States | 2015-Oct-24 | A/Iowa/49/2015 | Iowa State Hygienic Laboratory | Centers for Disease Control and Prevention |
| EPI677041 | HA | United States | 2015-Sep-06 | A/Alaska/216/2015 | Alaska State Virology Lab | Centers for Disease Control and Prevention |
| EPI677049 | HA | United States | 2015-Sep-07 | A/Alaska/217/2015 | Alaska State Virology Lab | Centers for Disease Control and Prevention |
| EPI677057 | HA | United States | 2015-Sep-07 | A/Alaska/218/2015 | Alaska State Virology Lab | Centers for Disease Control and Prevention |
| EPI677065 | HA | United States | 2015-Sep-05 | A/Alaska/219/2015 | Alaska State Virology Lab | Centers for Disease Control and Prevention |
| EPI677073 | HA | United States | 2015-Sep-04 | A/Alaska/220/2015 | Alaska State Virology Lab | Centers for Disease Control and Prevention |
| EPI677081 | HA | United States | 2015-Sep-03 | A/Alaska/221/2015 | Alaska State Virology Lab | Centers for Disease Control and Prevention |
| EPI677089 | HA | United States | 2015-Sep-08 | A/Alaska/222/2015 | Alaska State Virology Lab | Centers for Disease Control and Prevention |
| EPI677118 | HA | United States | 2015-Sep-11 | A/Alaska/227/2015 | Alaska State Virology Lab | Centers for Disease Control and Prevention |
| EPI677126 | HA | United States | 2015-Sep-12 | A/Alaska/228/2015 | Alaska State Virology Lab | Centers for Disease Control and Prevention |

| | | | | | | |
|---|---|---|---|---|---|---|
| EPI677134 | HA | United States | 2015-Sep-13 | A/Alaska/229/2015 | Alaska State Virology Lab | Centers for Disease Control and Prevention |
| EPI677142 | HA | United States | 2015-Sep-14 | A/Alaska/230/2015 | Alaska State Virology Lab | Centers for Disease Control and Prevention |
| EPI677150 | HA | United States | 2015-Sep-15 | A/Alaska/231/2015 | Alaska State Virology Lab | Centers for Disease Control and Prevention |
| EPI677158 | HA | United States | 2015-Sep-09 | A/Alaska/232/2015 | Alaska State Virology Lab | Centers for Disease Control and Prevention |
| EPI677178 | HA | United States | 2015-Sep-16 | A/Alaska/235/2015 | Alaska State Virology Lab | Centers for Disease Control and Prevention |
| EPI677186 | HA | United States | 2015-Sep-16 | A/Alaska/236/2015 | Alaska State Virology Lab | Centers for Disease Control and Prevention |
| EPI677194 | HA | United States | 2015-Sep-12 | A/Alaska/237/2015 | Alaska State Virology Lab | Centers for Disease Control and Prevention |
| EPI677202 | HA | United States | 2015-Sep-16 | A/Alaska/238/2015 | Alaska State Virology Lab | Centers for Disease Control and Prevention |
| EPI677210 | HA | United States | 2015-Sep-16 | A/Alaska/239/2015 | Alaska State Virology Lab | Centers for Disease Control and Prevention |
| EPI677218 | HA | United States | 2015-Sep-15 | A/Alaska/240/2015 | Alaska State Virology Lab | Centers for Disease Control and Prevention |
| EPI677226 | HA | United States | 2015-Sep-16 | A/Alaska/241/2015 | Alaska State Virology Lab | Centers for Disease Control and Prevention |
| EPI677234 | HA | United States | 2015-Sep-14 | A/Alaska/243/2015 | Alaska State Virology Lab | Centers for Disease Control and Prevention |
| EPI677242 | HA | United States | 2015-Sep-14 | A/Alaska/244/2015 | Alaska State Virology Lab | Centers for Disease Control and Prevention |
| EPI677250 | HA | United States | 2015-Sep-10 | A/Alaska/245/2015 | Alaska State Virology Lab | Centers for Disease Control and Prevention |
| EPI677258 | HA | United States | 2015-Sep-13 | A/Alaska/246/2015 | Alaska State Virology Lab | Centers for Disease Control and Prevention |
| EPI677266 | HA | United States | 2015-Sep-17 | A/Alaska/247/2015 | Alaska State Virology Lab | Centers for Disease Control and Prevention |
| EPI677274 | HA | United States | 2015-Sep-17 | A/Alaska/249/2015 | Alaska State Virology Lab | Centers for Disease Control and Prevention |
| EPI677282 | HA | United States | 2015-Sep-28 | A/Hawaii/62/2015 | State of Hawaii Department of Health | Centers for Disease Control and Prevention |
| EPI677290 | HA | United States | 2015-Sep-28 | A/Hawaii/63/2015 | State of Hawaii Department of Health | Centers for Disease Control and Prevention |
| EPI677305 | HA | United States | 2015-Oct-04 | A/Michigan/58/2015 | Michigan Department of Community Health | Centers for Disease Control and Prevention |
| EPI677313 | HA | United States | 2015-Oct-04 | A/Hawaii/69/2015 | State of Hawaii Department of Health | Centers for Disease Control and Prevention |
| EPI677321 | HA | United States | 2015-Oct-04 | A/Hawaii/69/2015 | State of Hawaii Department of Health | Centers for Disease Control and Prevention |
| EPI677329 | HA | United States | 2015-Oct-05 | A/Hawaii/70/2015 | State of Hawaii Department of Health | Centers for Disease Control and Prevention |
| EPI677337 | HA | United States | 2015-Oct-05 | A/Hawaii/70/2015 | State of Hawaii Department of Health | Centers for Disease Control and Prevention |
| EPI677345 | HA | United States | 2015-Oct-09 | A/Hawaii/71/2015 | State of Hawaii Department of Health | Centers for Disease Control and Prevention |

| | | | | | | |
|---|---|---|---|---|---|---|
| EPI677353 | HA | United States | 2015-Oct-11 | A/Colorado/15/2015 | Colorado Department of Health Lab | Centers for Disease Control and Prevention |
| EPI677361 | HA | United States | 2015-Oct-22 | A/Alaska/260/2015 | Alaska State Virology Lab | Centers for Disease Control and Prevention |
| EPI677369 | HA | United States | 2015-Oct-26 | A/Alaska/261/2015 | Alaska State Virology Lab | Centers for Disease Control and Prevention |
| EPI677377 | HA | United States | 2015-Oct-25 | A/Alaska/262/2015 | Alaska State Virology Lab | Centers for Disease Control and Prevention |
| EPI677385 | HA | United States | 2015-Oct-25 | A/Alaska/262/2015 | Alaska State Virology Lab | Centers for Disease Control and Prevention |
| EPI677393 | HA | United States | 2015-Oct-17 | A/Washington/53/2015 | Washington State Public Health Laboratory | Centers for Disease Control and Prevention |
| EPI677401 | HA | United States | 2015-Oct-17 | A/Washington/54/2015 | Washington State Public Health Laboratory | Centers for Disease Control and Prevention |
| EPI677414 | HA | United States | 2015-Oct-19 | A/West Virginia/24/2015 | West Virginia Office of Laboratory Services | Centers for Disease Control and Prevention |
| EPI677422 | HA | United States | 2015-Oct-08 | A/Hawaii/72/2015 | State of Hawaii Department of Health | Centers for Disease Control and Prevention |
| EPI677430 | HA | United States | 2015-Oct-08 | A/Hawaii/72/2015 | State of Hawaii Department of Health | Centers for Disease Control and Prevention |
| EPI677438 | HA | United States | 2015-Oct-12 | A/Hawaii/73/2015 | State of Hawaii Department of Health | Centers for Disease Control and Prevention |
| EPI677446 | HA | United States | 2015-Oct-09 | A/Hawaii/74/2015 | State of Hawaii Department of Health | Centers for Disease Control and Prevention |
| EPI677454 | HA | United States | 2015-Oct-17 | A/Hawaii/75/2015 | State of Hawaii Department of Health | Centers for Disease Control and Prevention |
| EPI677462 | HA | United States | 2015-Oct-26 | A/Florida/82/2015 | Florida Department of Health-Tampa | Centers for Disease Control and Prevention |
| EPI677470 | HA | United States | 2015-Oct-26 | A/Florida/82/2015 | Florida Department of Health-Tampa | Centers for Disease Control and Prevention |
| EPI677478 | HA | United States | 2015-Oct-29 | A/Wisconsin/87/2015 | Wisconsin State Laboratory of Hygiene | Centers for Disease Control and Prevention |
| EPI677486 | HA | United States | 2015-Nov-02 | A/Louisiana/06/2015 | Louisiana Department of Health and Hospitals | Centers for Disease Control and Prevention |
| EPI677494 | HA | United States | 2015-Nov-02 | A/Florida/85/2015 | Florida Department of Health-Tampa | Centers for Disease Control and Prevention |
| EPI677502 | HA | United States | 2015-Nov-02 | A/Florida/86/2015 | Florida Department of Health-Tampa | Centers for Disease Control and Prevention |
| EPI677510 | HA | United States | 2015-Nov-08 | A/Florida/87/2015 | Florida Department of Health-Tampa | Centers for Disease Control and Prevention |
| EPI677518 | HA | United States | 2015-Nov-16 | A/Minnesota/47/2015 | Minnesota Department of Health | Centers for Disease Control and Prevention |
| EPI677526 | HA | United States | 2015-Oct-06 | A/South Carolina/10/2015 | South Carolina Department of Health | Centers for Disease Control and Prevention |
| EPI677534 | HA | United States | 2015-Oct-06 | A/South Carolina/10/2015 | South Carolina Department of Health | Centers for Disease Control and Prevention |
| EPI677542 | HA | United States | 2015-Oct-27 | A/South Carolina/11/2015 | South Carolina Department of Health | Centers for Disease Control and Prevention |
| EPI677550 | HA | United States | 2015-Nov-03 | A/Wyoming/19/2015 | Wyoming Public Health Laboratory | Centers for Disease Control and Prevention |

| | | | | | | |
|---|---|---|---|---|---|---|
| EPI677558 | HA | United States | 2015-Oct-20 | A/California/93/2015 | California Department of Health Services | Centers for Disease Control and Prevention |
| EPI677566 | HA | United States | 2015-Oct-20 | A/California/94/2015 | California Department of Health Services | Centers for Disease Control and Prevention |
| EPI677574 | HA | United States | 2015-Oct-20 | A/California/95/2015 | California Department of Health Services | Centers for Disease Control and Prevention |
| EPI677582 | HA | United States | 2015-Oct-19 | A/California/98/2015 | California Department of Health Services | Centers for Disease Control and Prevention |
| EPI677590 | HA | United States | 2015-Nov-07 | A/Washington/57/2015 | Washington State Public Health Laboratory | Centers for Disease Control and Prevention |
| EPI677598 | HA | United States | 2015-Nov-07 | A/Washington/57/2015 | Washington State Public Health Laboratory | Centers for Disease Control and Prevention |
| EPI677610 | HA | United States | 2015-Nov-09 | A/Washington/58/2015 | Washington State Public Health Laboratory | Centers for Disease Control and Prevention |
| EPI677618 | HA | United States | 2015-Oct-15 | A/Michigan/66/2015 | Michigan Department of Community Health | Centers for Disease Control and Prevention |
| EPI677626 | HA | United States | 2015-Oct-10 | A/Michigan/67/2015 | Michigan Department of Community Health | Centers for Disease Control and Prevention |
| EPI679143 | HA | United Kingdom | 2015-Aug-01 | A/England/53340242/2015 | Microbiology Services Colindale, Public Health England | Microbiology Services Colindale, Public Health England |
| EPI679164 | HA | United Kingdom | 2015-Sep-06 | A/England/334/2015 | Microbiology Services Colindale, Public Health England | Microbiology Services Colindale, Public Health England |
| EPI679172 | HA | United Kingdom | 2015-Sep-14 | A/England/53860135/2015 | Microbiology Services Colindale, Public Health England | Microbiology Services Colindale, Public Health England |
| EPI679229 | HA | United Kingdom | 2015-Oct-02 | A/England/54380710/2015 | Microbiology Services Colindale, Public Health England | Microbiology Services Colindale, Public Health England |
| EPI679274 | HA | United Kingdom | 2015-Nov-16 | A/England/54760378/2015 | Microbiology Services Colindale, Public Health England | Microbiology Services Colindale, Public Health England |
| EPI679342 | HA | United States | 2015-Jul-04 | A/Alaska/104/2015 | Alaska State Virology Lab | Centers for Disease Control and Prevention |
| EPI679350 | HA | United States | 2015-Jul-04 | A/Alaska/105/2015 | Alaska State Virology Lab | Centers for Disease Control and Prevention |
| EPI679358 | HA | United States | 2015-Jun-27 | A/Alaska/106/2015 | Alaska State Virology Lab | Centers for Disease Control and Prevention |
| EPI679366 | HA | United States | 2015-Jul-03 | A/Alaska/107/2015 | Alaska State Virology Lab | Centers for Disease Control and Prevention |
| EPI679375 | HA | United States | 2015-Jul-06 | A/Alaska/108/2015 | Alaska State Virology Lab | Centers for Disease Control and Prevention |
| EPI679383 | HA | United States | 2015-Jul-07 | A/Alaska/109/2015 | Alaska State Virology Lab | Centers for Disease Control and Prevention |
| EPI679398 | HA | United States | 2015-Jul-09 | A/Alaska/111/2015 | Alaska State Virology Lab | Centers for Disease Control and Prevention |
| EPI679406 | HA | United States | 2015-Sep-19 | A/Alaska/248/2015 | Alaska State Virology Lab | Centers for Disease Control and Prevention |
| EPI679414 | HA | United States | 2015-Sep-18 | A/Alaska/251/2015 | Alaska State Virology Lab | Centers for Disease Control and Prevention |
| EPI679422 | HA | Canada | 2015-Sep-29 | A/Alberta/RV2372/2015 | National Microbiology Laboratory, Health Canada | Centers for Disease Control and Prevention |
| EPI679430 | HA | United States | 2015-Oct-21 | A/California/96/2015 | California Department of Health Services | Centers for Disease Control and Prevention |

| | | | | | | |
|---|---|---|---|---|---|---|
| EPI679438 | HA | United States | 2015-Oct-24 | A/California/97/2015 | California Department of Health Services | Centers for Disease Control and Prevention |
| EPI679446 | HA | United States | 2015-Oct-26 | A/Florida/79/2015 | Florida Department of Health-Jacksonville | Centers for Disease Control and Prevention |
| EPI679454 | HA | Hong Kong (SAR) | 2015-Oct-09 | A/Hong Kong/15590/2015 | Government Virus Unit | Centers for Disease Control and Prevention |
| EPI679463 | HA | Hong Kong (SAR) | 2015-Oct-14 | A/Hong Kong/15607/2015 | Government Virus Unit | Centers for Disease Control and Prevention |
| EPI679471 | HA | Hong Kong (SAR) | 2015-Oct-17 | A/Hong Kong/15609/2015 | Government Virus Unit | Centers for Disease Control and Prevention |
| EPI679484 | HA | United States | 2015-Nov-01 | A/Iowa/54/2015 | Iowa State Hygienic Laboratory | Centers for Disease Control and Prevention |
| EPI679492 | HA | Lao, People's Democratic Republic | 2015-Jun-27 | A/Laos/438/2015 | National Center for Laboratory and Epidemiology | Centers for Disease Control and Prevention |
| EPI679500 | HA | Lao, People's Democratic Republic | 2015-Jul-27 | A/Laos/769/2015 | National Center for Laboratory and Epidemiology | Centers for Disease Control and Prevention |
| EPI679508 | HA | Lao, People's Democratic Republic | 2015-Aug-05 | A/Laos/794/2015 | National Center for Laboratory and Epidemiology | Centers for Disease Control and Prevention |
| EPI679514 | HA | United States | 2015-Sep-18 | A/Montana/24/2015 | Montana Public Health Laboratory | Centers for Disease Control and Prevention |
| EPI679527 | HA | United States | 2015-Sep-06 | A/New York/67/2015 | New York State Department of Health | Centers for Disease Control and Prevention |
| EPI679540 | HA | United States | 2015-Oct-12 | A/New York/69/2015 | New York State Department of Health | Centers for Disease Control and Prevention |
| EPI679549 | HA | United States | 2015-Nov-18 | A/Oklahoma/21/2015 | Oklahoma State Department of Health | Centers for Disease Control and Prevention |
| EPI679557 | HA | Brazil | 2015-May-28 | A/Pernambuco/135596-IEC/2015 | National Influenza Center | Centers for Disease Control and Prevention |
| EPI679565 | HA | Brazil | 2015-Jun-01 | A/Pernambuco/135597-IEC/2015 | National Influenza Center | Centers for Disease Control and Prevention |
| EPI679573 | HA | Brazil | 2015-Jun-22 | A/Pernambuco/136136-IEC/2015 | National Influenza Center | Centers for Disease Control and Prevention |
| EPI679585 | HA | Peru | 2015-Jul-21 | A/Peru/69/2015 | Laboratorio de Referencia Nacional Virus Respiratorios, Instituto Nacional de Salud | Centers for Disease Control and Prevention |
| EPI679598 | HA | Puerto Rico | 2015-Jul-13 | A/Puerto Rico/04/2015 | Puerto Rico Department of Health | Centers for Disease Control and Prevention |
| EPI679610 | HA | Puerto Rico | 2015-Jul-21 | A/Puerto Rico/06/2015 | Puerto Rico Department of Health | Centers for Disease Control and Prevention |
| EPI679618 | HA | Puerto Rico | 2015-Aug-07 | A/Puerto Rico/07/2015 | Puerto Rico Department of Health | Centers for Disease Control and Prevention |
| EPI679626 | HA | Puerto Rico | 2015-Aug-10 | A/Puerto Rico/08/2015 | Puerto Rico Department of Health | Centers for Disease Control and Prevention |
| EPI679634 | HA | Puerto Rico | 2015-Aug-16 | A/Puerto Rico/10/2015 | Puerto Rico Department of Health | Centers for Disease Control and Prevention |
| EPI679650 | HA | Puerto Rico | 2015-Aug-24 | A/Puerto Rico/12/2015 | Puerto Rico Department of Health | Centers for Disease Control and Prevention |

| | | | | | | |
|---|---|---|---|---|---|---|
| EPI679658 | HA | Puerto Rico | 2015-Aug-28 | A/Puerto Rico/13/2015 | Puerto Rico Department of Health | Centers for Disease Control and Prevention |
| EPI679672 | HA | Puerto Rico | 2015-Aug-30 | A/Puerto Rico/15/2015 | Puerto Rico Department of Health | Centers for Disease Control and Prevention |
| EPI679680 | HA | Puerto Rico | 2015-Sep-03 | A/Puerto Rico/16/2015 | Puerto Rico Department of Health | Centers for Disease Control and Prevention |
| EPI679694 | HA | Puerto Rico | 2015-Sep-09 | A/Puerto Rico/20/2015 | Puerto Rico Department of Health | Centers for Disease Control and Prevention |
| EPI679702 | HA | Puerto Rico | 2015-Sep-09 | A/Puerto Rico/22/2015 | Puerto Rico Department of Health | Centers for Disease Control and Prevention |
| EPI679710 | HA | Puerto Rico | 2015-Sep-12 | A/Puerto Rico/23/2015 | Puerto Rico Department of Health | Centers for Disease Control and Prevention |
| EPI679718 | HA | Puerto Rico | 2015-Sep-12 | A/Puerto Rico/24/2015 | Puerto Rico Department of Health | Centers for Disease Control and Prevention |
| EPI679726 | HA | Puerto Rico | 2015-Sep-11 | A/Puerto Rico/25/2015 | Puerto Rico Department of Health | Centers for Disease Control and Prevention |
| EPI679734 | HA | Puerto Rico | 2015-Aug-10 | A/Puerto Rico/26/2015 | Puerto Rico Department of Health | Centers for Disease Control and Prevention |
| EPI679749 | HA | United States | 2015-Sep-09 | A/South Dakota/38/2015 | South Dakota Public Health Lab | Centers for Disease Control and Prevention |
| EPI679757 | HA | United States | 2015-Jul-01 | A/Texas/64/2015 | Dallas County Health and Human Services | Centers for Disease Control and Prevention |
| EPI679765 | HA | United States | 2015-Oct-21 | A/Washington/59/2015 | Seattle & King County Public Health Lab | Centers for Disease Control and Prevention |
| EPI679896 | HA | United States | 2015-Aug-04 | A/Alaska/163/2015 | Alaska State Virology Lab | Centers for Disease Control and Prevention |
| EPI679904 | HA | United States | 2015-Aug-03 | A/Alaska/164/2015 | Alaska State Virology Lab | Centers for Disease Control and Prevention |
| EPI679912 | HA | United States | 2015-Aug-19 | A/Alaska/165/2015 | Alaska State Virology Lab | Centers for Disease Control and Prevention |
| EPI679920 | HA | United States | 2015-Aug-15 | A/Alaska/166/2015 | Alaska State Virology Lab | Centers for Disease Control and Prevention |
| EPI679928 | HA | United States | 2015-Aug-15 | A/Alaska/167/2015 | Alaska State Virology Lab | Centers for Disease Control and Prevention |
| EPI679936 | HA | United States | 2015-Aug-15 | A/Alaska/168/2015 | Alaska State Virology Lab | Centers for Disease Control and Prevention |
| EPI679944 | HA | United States | 2015-Aug-21 | A/Alaska/170/2015 | Alaska State Virology Lab | Centers for Disease Control and Prevention |
| EPI679952 | HA | United States | 2015-Aug-25 | A/Alaska/171/2015 | Alaska State Virology Lab | Centers for Disease Control and Prevention |
| EPI679960 | HA | United States | 2015-Aug-26 | A/Alaska/172/2015 | Alaska State Virology Lab | Centers for Disease Control and Prevention |
| EPI679968 | HA | United States | 2015-Aug-30 | A/Alaska/174/2015 | Alaska State Virology Lab | Centers for Disease Control and Prevention |
| EPI679981 | HA | United States | 2015-Aug-19 | A/Alaska/176/2015 | Alaska State Virology Lab | Centers for Disease Control and Prevention |
| EPI679989 | HA | United States | 2015-Aug-21 | A/Alaska/177/2015 | Alaska State Virology Lab | Centers for Disease Control and Prevention |
| EPI679997 | HA | United States | 2015-Aug-18 | A/Alaska/178/2015 | Alaska State Virology Lab | Centers for Disease Control and Prevention |

| | | | | | | | |
|---|---|---|---|---|---|---|---|
| EPI680005 | HA | United States | 2015-Sep-07 | A/Alaska/209/2015 | Alaska State Virology Lab | Centers for Disease Control and Prevention | |
| EPI680013 | HA | United States | 2015-Sep-29 | A/Arizona/19/2015 | Arizona Department of Health Services | Centers for Disease Control and Prevention | |
| EPI680021 | HA | United States | 2015-Oct-26 | A/Arizona/23/2015 | Arizona Department of Health Services | Centers for Disease Control and Prevention | |
| EPI680029 | HA | United States | 2015-Nov-02 | A/Arizona/24/2015 | Arizona Department of Health Services | Centers for Disease Control and Prevention | |
| EPI680037 | HA | United States | 2015-Nov-02 | A/Arizona/25/2015 | Arizona Department of Health Services | Centers for Disease Control and Prevention | Galiano,M. |
| EPI680045 | HA | United States | 2015-Oct-18 | A/Arizona/27/2015 | Arizona Department of Health Services | Centers for Disease Control and Prevention | Galiano,M. |
| EPI680053 | HA | United States | 2015-Oct-18 | A/California/107/2015 | California Department of Health Services | Centers for Disease Control and Prevention | Galiano,M. |
| EPI680061 | HA | United States | 2015-Nov-19 | A/Colorado/21/2015 | Colorado Department of Health Lab | Centers for Disease Control and Prevention | |
| EPI680069 | HA | United States | 2015-Nov-22 | A/Colorado/23/2015 | Colorado Department of Health Lab | Centers for Disease Control and Prevention | |
| EPI680077 | HA | United States | 2015-Nov-17 | A/Colorado/24/2015 | Colorado Department of Health Lab | Centers for Disease Control and Prevention | |
| EPI680085 | HA | United States | 2015-Sep-28 | A/Hawaii/64/2015 | State of Hawaii Department of Health | Centers for Disease Control and Prevention | |
| EPI680093 | HA | United States | 2015-Nov-02 | A/Hawaii/77/2015 | State of Hawaii Department of Health | Centers for Disease Control and Prevention | |
| EPI680106 | HA | Nigeria | 2015-Sep-08 | A/Nigeria/253/2015 | National Influenza Reference Laboratory | Centers for Disease Control and Prevention | |
| EPI680114 | HA | United States | 2015-Nov-08 | A/Texas/72/2015 | Texas Department of State Health Services-Laboratory Services | Centers for Disease Control and Prevention | |
| EPI680122 | HA | United States | 2015-Oct-19 | A/Wisconsin/86/2015 | Wisconsin State Laboratory of Hygiene | Centers for Disease Control and Prevention | |
| EPI680131 | HA | United States | 2015-Aug-16 | A/Alaska/179/2015 | Alaska State Virology Lab | Centers for Disease Control and Prevention | |
| EPI680139 | HA | United States | 2015-Aug-20 | A/Alaska/184/2015 | Alaska State Virology Lab | Centers for Disease Control and Prevention | |
| EPI680147 | HA | United States | 2015-Aug-19 | A/Alaska/187/2015 | Alaska State Virology Lab | Centers for Disease Control and Prevention | |
| EPI680155 | HA | United States | 2015-Aug-15 | A/Alaska/188/2015 | Alaska State Virology Lab | Centers for Disease Control and Prevention | |
| EPI680163 | HA | United States | 2015-Aug-13 | A/Alaska/189/2015 | Alaska State Virology Lab | Centers for Disease Control and Prevention | |
| EPI680177 | HA | United States | 2015-Aug-18 | A/Alaska/191/2015 | Alaska State Virology Lab | Centers for Disease Control and Prevention | |
| EPI680185 | HA | United States | 2015-Aug-16 | A/Alaska/192/2015 | Alaska State Virology Lab | Centers for Disease Control and Prevention | |
| EPI680193 | HA | United States | 2015-Aug-17 | A/Alaska/193/2015 | Alaska State Virology Lab | Centers for Disease Control and Prevention | |
| EPI680201 | HA | Bangladesh | 2015-Aug-17 | A/Bangladesh/3937/2015 | Institute of Epidemiology Disease Control and Research (IEDCR) & Bangladesh National Influenza Centre (NIC) | Centers for Disease Control and Prevention | |

| | | | | | | |
|---|---|---|---|---|---|---|
| EPI680209 | HA | United States | 2015-Oct-28 | A/Florida/80/2015 | Florida Department of Health-Jacksonville | Centers for Disease Control and Prevention |
| EPI680217 | HA | United States | 2015-Sep-18 | A/Idaho/25/2015 | State of Idaho Bureau of Laboratories | Centers for Disease Control and Prevention |
| EPI680225 | HA | United States | 2015-Sep-12 | A/Michigan/53/2015 | Michigan Department of Community Health | Centers for Disease Control and Prevention |
| EPI680233 | HA | United States | 2015-Sep-19 | A/Michigan/54/2015 | Michigan Department of Community Health | Centers for Disease Control and Prevention |
| EPI680241 | HA | United States | 2015-Sep-18 | A/Michigan/55/2015 | Michigan Department of Community Health | Centers for Disease Control and Prevention |
| EPI680249 | HA | United States | 2015-Oct-05 | A/Michigan/60/2015 | Michigan Department of Community Health | Centers for Disease Control and Prevention |
| EPI680257 | HA | United States | 2015-Sep-13 | A/New Mexico/26/2015 | New Mexico Department of Health | Centers for Disease Control and Prevention |
| EPI680265 | HA | Nigeria | 2015-Sep-21 | A/Nigeria/299/2015 | National Influenza Reference Laboratory | Centers for Disease Control and Prevention |
| EPI680273 | HA | Nigeria | 2015-Sep-22 | A/Nigeria/300/2015 | National Influenza Reference Laboratory | Centers for Disease Control and Prevention |
| EPI680281 | HA | United States | 2015-Aug-19 | A/Vermont/08/2015 | Vermont Department of Health Laboratory | Centers for Disease Control and Prevention |
| EPI683254 | HA | Lao, People's Democratic Republic | 2015-Jun-22 | A/Laos/648/2015 | National Center for Laboratory and Epidemiology | Centers for Disease Control and Prevention |
| EPI683262 | HA | Lao, People's Democratic Republic | 2015-Jul-15 | A/Laos/738/2015 | National Center for Laboratory and Epidemiology | Centers for Disease Control and Prevention |
| EPI683278 | HA | United States | 2015-Jun-15 | A/Alaska/49/2015 | Alaska State Virology Lab | Centers for Disease Control and Prevention |
| EPI683286 | HA | United States | 2015-Jun-15 | A/Alaska/49/2015 | Alaska State Virology Lab | Centers for Disease Control and Prevention |
| EPI683294 | HA | United States | 2015-Jun-16 | A/Alaska/50/2015 | Alaska State Virology Lab | Centers for Disease Control and Prevention |
| EPI683302 | HA | United States | 2015-Jun-16 | A/Alaska/50/2015 | Alaska State Virology Lab | Centers for Disease Control and Prevention |
| EPI683310 | HA | Saint Vincent and the Grenadines | 2015-Feb-12 | A/St. Vincent And Grenadines/423/2015 | Caribbean Epidemiology Center | Centers for Disease Control and Prevention |
| EPI683318 | HA | Australia | 2015-May-16 | A/Victoria/511/2015 | WHO Collaborating Centre for Reference and Research on Influenza | Centers for Disease Control and Prevention |
| EPI683326 | HA | Australia | 2015-May-16 | A/Victoria/511/2015 | WHO Collaborating Centre for Reference and Research on Influenza | Centers for Disease Control and Prevention |
| EPI683334 | HA | Australia | 2015-Mar-13 | A/South Australia/21/2015 | WHO Collaborating Centre for Reference and Research on Influenza | Centers for Disease Control and Prevention |
| EPI683348 | HA | United States | 2015-Jul-21 | A/Alaska/151/2015 | Alaska State Virology Lab | Centers for Disease Control and Prevention |
| EPI683356 | HA | United States | 2015-Jul-21 | A/Alaska/151/2015 | Alaska State Virology Lab | Centers for Disease Control and Prevention |

| | | | | | | |
|---|---|---|---|---|---|---|
| EPI683364 | HA | United States | 2015-Jul-20 | A/Alaska/152/2015 | Alaska State Virology Lab | Centers for Disease Control and Prevention |
| EPI683372 | HA | United States | 2015-Jul-20 | A/Alaska/152/2015 | Alaska State Virology Lab | Centers for Disease Control and Prevention |
| EPI683387 | HA | United States | 2015-Aug-03 | A/Alaska/153/2015 | Alaska State Virology Lab | Centers for Disease Control and Prevention |
| EPI683395 | HA | United States | 2015-Jul-24 | A/Alaska/154/2015 | Alaska State Virology Lab | Centers for Disease Control and Prevention |
| EPI683403 | HA | United States | 2015-Jul-24 | A/Alaska/154/2015 | Alaska State Virology Lab | Centers for Disease Control and Prevention |
| EPI683411 | HA | United States | 2015-Aug-08 | A/Alaska/155/2015 | Alaska State Virology Lab | Centers for Disease Control and Prevention |
| EPI683419 | HA | United States | 2015-Aug-08 | A/Alaska/155/2015 | Alaska State Virology Lab | Centers for Disease Control and Prevention |
| EPI683427 | HA | United States | 2015-Aug-03 | A/Alaska/156/2015 | Alaska State Virology Lab | Centers for Disease Control and Prevention |
| EPI683435 | HA | United States | 2015-Aug-03 | A/Alaska/156/2015 | Alaska State Virology Lab | Centers for Disease Control and Prevention |
| EPI683443 | HA | Russian Federation | 2015-Mar-02 | A/Blagoveshchensk/19/2015 | State Research Center of Virology and Biotechnology (VECTOR) | Centers for Disease Control and Prevention |
| EPI683451 | HA | Russian Federation | 2015-Mar-02 | A/Blagoveshchensk/19/2015 | State Research Center of Virology and Biotechnology (VECTOR) | Centers for Disease Control and Prevention |
| EPI683459 | HA | Russian Federation | 2015-Feb-17 | A/Irkutsk/88/2015 | State Research Center of Virology and Biotechnology (VECTOR) | Centers for Disease Control and Prevention |
| EPI683467 | HA | Russian Federation | 2015-Feb-17 | A/Irkutsk/88/2015 | State Research Center of Virology and Biotechnology (VECTOR) | Centers for Disease Control and Prevention |
| EPI683475 | HA | Russian Federation | 2015-Feb-04 | A/Krasnoyarsk/324/2015 | State Research Center of Virology and Biotechnology (VECTOR) | Centers for Disease Control and Prevention |
| EPI683489 | HA | Russian Federation | 2015-Mar-05 | A/Yekaterinburg/239/2015 | State Research Center of Virology and Biotechnology (VECTOR) | Centers for Disease Control and Prevention |
| EPI683497 | HA | Russian Federation | 2015-Mar-05 | A/Yekaterinburg/239/2015 | State Research Center of Virology and Biotechnology (VECTOR) | Centers for Disease Control and Prevention |
| EPI683505 | HA | United States | 2015-Aug-13 | A/Alaska/157/2015 | Alaska State Virology Lab | Centers for Disease Control and Prevention |
| EPI683513 | HA | United States | 2015-Aug-13 | A/Alaska/157/2015 | Alaska State Virology Lab | Centers for Disease Control and Prevention |
| EPI683521 | HA | United States | 2015-Aug-14 | A/Alaska/158/2015 | Alaska State Virology Lab | Centers for Disease Control and Prevention |
| EPI683529 | HA | United States | 2015-Aug-14 | A/Alaska/158/2015 | Alaska State Virology Lab | Centers for Disease Control and Prevention |
| EPI683537 | HA | United States | 2015-Aug-17 | A/Alaska/159/2015 | Alaska State Virology Lab | Centers for Disease Control and Prevention |
| EPI683545 | HA | United States | 2015-Aug-17 | A/Alaska/159/2015 | Alaska State Virology Lab | Centers for Disease Control and Prevention |
| EPI683553 | HA | United States | 2015-Aug-19 | A/Alaska/162/2015 | Alaska State Virology Lab | Centers for Disease Control and Prevention |
| EPI683561 | HA | United States | 2015-Aug-19 | A/Alaska/162/2015 | Alaska State Virology Lab | Centers for Disease Control and Prevention |
| EPI683569 | HA | Argentina | 2015-Jun-15 | A/Argentina/23/2015 | Instituto Nacional de Enfermedades Infecciosas | Centers for Disease Control and Prevention |

| | | | | | | |
|---|---|---|---|---|---|---|
| EPI683577 | HA | Argentina | 2015-Jun-15 | A/Argentina/23/2015 | Instituto Nacional de Enfermedades Infecciosas | Centers for Disease Control and Prevention |
| EPI683585 | HA | Argentina | 2015-Jun-15 | A/Argentina/24/2015 | Instituto Nacional de Enfermedades Infecciosas | Centers for Disease Control and Prevention |
| EPI683593 | HA | Argentina | 2015-Jun-15 | A/Argentina/24/2015 | Instituto Nacional de Enfermedades Infecciosas | Centers for Disease Control and Prevention |
| EPI683601 | HA | Argentina | 2015-Jun-15 | A/Argentina/25/2015 | Instituto Nacional de Enfermedades Infecciosas | Centers for Disease Control and Prevention |
| EPI683609 | HA | Argentina | 2015-Jun-15 | A/Argentina/25/2015 | Instituto Nacional de Enfermedades Infecciosas | Centers for Disease Control and Prevention |
| EPI683617 | HA | Argentina | 2015-Jun-30 | A/Argentina/103/2015 | Instituto Nacional de Enfermedades Infecciosas | Centers for Disease Control and Prevention |
| EPI683625 | HA | Argentina | 2015-Jun-30 | A/Argentina/103/2015 | Instituto Nacional de Enfermedades Infecciosas | Centers for Disease Control and Prevention |
| EPI683633 | HA | Argentina | 2015-Jul-10 | A/Argentina/104/2015 | Instituto Nacional de Enfermedades Infecciosas | Centers for Disease Control and Prevention |
| EPI683641 | HA | Argentina | 2015-Jul-10 | A/Argentina/104/2015 | Instituto Nacional de Enfermedades Infecciosas | Centers for Disease Control and Prevention |
| EPI683649 | HA | Argentina | 2015-Jul-22 | A/Argentina/11592/2015 | CEMIC University Hospital | Centers for Disease Control and Prevention |
| EPI683657 | HA | Argentina | 2015-Jul-22 | A/Argentina/11592/2015 | CEMIC University Hospital | Centers for Disease Control and Prevention |
| EPI683665 | HA | Argentina | 2015-Jul-13 | A/Argentina/11594/2015 | CEMIC University Hospital | Centers for Disease Control and Prevention |
| EPI683673 | HA | Argentina | 2015-Jul-13 | A/Argentina/11594/2015 | CEMIC University Hospital | Centers for Disease Control and Prevention |
| EPI683681 | HA | Argentina | 2015-Jul-13 | A/Argentina/11596/2015 | CEMIC University Hospital | Centers for Disease Control and Prevention |
| EPI683689 | HA | Argentina | 2015-Jul-13 | A/Argentina/11596/2015 | CEMIC University Hospital | Centers for Disease Control and Prevention |
| EPI683697 | HA | Argentina | 2015-Jul-27 | A/Argentina/11611/2015 | CEMIC University Hospital | Centers for Disease Control and Prevention |
| EPI683705 | HA | Argentina | 2015-Jul-27 | A/Argentina/11611/2015 | CEMIC University Hospital | Centers for Disease Control and Prevention |
| EPI683713 | HA | Argentina | 2015-Jul-30 | A/Argentina/11628/2015 | CEMIC University Hospital | Centers for Disease Control and Prevention |
| EPI683721 | HA | Argentina | 2015-Jul-30 | A/Argentina/11628/2015 | CEMIC University Hospital | Centers for Disease Control and Prevention |
| EPI683729 | HA | Argentina | 2015-Aug-11 | A/Argentina/11639/2015 | CEMIC University Hospital | Centers for Disease Control and Prevention |
| EPI683742 | HA | Argentina | 2015-Aug-11 | A/Argentina/11640/2015 | CEMIC University Hospital | Centers for Disease Control and Prevention |
| EPI683750 | HA | Argentina | 2015-Aug-11 | A/Argentina/11640/2015 | CEMIC University Hospital | Centers for Disease Control and Prevention |
| EPI683758 | HA | Argentina | 2015-Aug-10 | A/Argentina/11652/2015 | CEMIC University Hospital | Centers for Disease Control and Prevention |
| EPI683766 | HA | Argentina | 2015-Aug-10 | A/Argentina/11652/2015 | CEMIC University Hospital | Centers for Disease Control and Prevention |
| EPI683774 | HA | Argentina | 2015-Aug-11 | A/Argentina/11656/2015 | CEMIC University Hospital | Centers for Disease Control and Prevention |

| | | | | | | |
|---|---|---|---|---|---|---|
| EPI683782 | HA | Argentina | 2015-Aug-11 | A/Argentina/11656/2015 | CEMIC University Hospital | Centers for Disease Control and Prevention |
| EPI683790 | HA | Argentina | 2015-Aug-12 | A/Argentina/11659/2015 | CEMIC University Hospital | Centers for Disease Control and Prevention |
| EPI683798 | HA | Argentina | 2015-Aug-12 | A/Argentina/11659/2015 | CEMIC University Hospital | Centers for Disease Control and Prevention |
| EPI683806 | HA | United States | 2015-Sep-10 | A/Massachusetts/25/2015 | Massachusetts Department of Public Health | Centers for Disease Control and Prevention |
| EPI683814 | HA | United States | 2015-Sep-10 | A/Massachusetts/25/2015 | Massachusetts Department of Public Health | Centers for Disease Control and Prevention |
| EPI683822 | HA | United States | 2015-Aug-31 | A/Indiana/17/2015 | Indiana State Department of Health Laboratories | Centers for Disease Control and Prevention |
| EPI683830 | HA | United States | 2015-Aug-31 | A/Indiana/17/2015 | Indiana State Department of Health Laboratories | Centers for Disease Control and Prevention |
| EPI683838 | HA | United States | 2015-Sep-01 | A/Indiana/19/2015 | Indiana State Department of Health Laboratories | Centers for Disease Control and Prevention |
| EPI683846 | HA | United States | 2015-Sep-01 | A/Indiana/19/2015 | Indiana State Department of Health Laboratories | Centers for Disease Control and Prevention |
| EPI683854 | HA | United States | 2015-Sep-01 | A/Indiana/20/2015 | Indiana State Department of Health Laboratories | Centers for Disease Control and Prevention |
| EPI683862 | HA | United States | 2015-Sep-01 | A/Indiana/20/2015 | Indiana State Department of Health Laboratories | Centers for Disease Control and Prevention |
| EPI683870 | HA | United States | 2015-Sep-01 | A/Indiana/21/2015 | Indiana State Department of Health Laboratories | Centers for Disease Control and Prevention |
| EPI683878 | HA | United States | 2015-Sep-01 | A/Indiana/21/2015 | Indiana State Department of Health Laboratories | Centers for Disease Control and Prevention |
| EPI683886 | HA | United States | 2015-Sep-01 | A/Indiana/22/2015 | Indiana State Department of Health Laboratories | Centers for Disease Control and Prevention |
| EPI683894 | HA | United States | 2015-Sep-01 | A/Indiana/22/2015 | Indiana State Department of Health Laboratories | Centers for Disease Control and Prevention |
| EPI683909 | HA | United States | 2015-Sep-03 | A/Indiana/24/2015 | Indiana State Department of Health Laboratories | Centers for Disease Control and Prevention |
| EPI683917 | HA | United States | 2015-Sep-03 | A/Indiana/25/2015 | Indiana State Department of Health Laboratories | Centers for Disease Control and Prevention |
| EPI683925 | HA | United States | 2015-Sep-03 | A/Indiana/25/2015 | Indiana State Department of Health Laboratories | Centers for Disease Control and Prevention |
| EPI683933 | HA | United States | 2015-Sep-09 | A/New Hampshire/39/2015 | New Hampshire Public Health Laboratories | Centers for Disease Control and Prevention |
| EPI683941 | HA | United States | 2015-Sep-09 | A/New Hampshire/39/2015 | New Hampshire Public Health Laboratories | Centers for Disease Control and Prevention |
| EPI683949 | HA | United States | 2015-Sep-14 | A/New Hampshire/40/2015 | New Hampshire Public Health Laboratories | Centers for Disease Control and Prevention |
| EPI683957 | HA | United States | 2015-Sep-14 | A/New Hampshire/40/2015 | New Hampshire Public Health Laboratories | Centers for Disease Control and Prevention |
| EPI683965 | HA | United States | 2015-Sep-14 | A/Minnesota/41/2015 | Minnesota Department of Health | Centers for Disease Control and Prevention |
| EPI683973 | HA | United States | 2015-Sep-14 | A/Minnesota/41/2015 | Minnesota Department of Health | Centers for Disease Control and Prevention |
| EPI683981 | HA | United States | 2015-Sep-15 | A/Florida/69/2015 | Florida Department of Health-Jacksonville | Centers for Disease Control and Prevention |

| | | | | | | |
|---|---|---|---|---|---|---|
| EPI683989 | HA | United States | 2015-Sep-15 | A/Florida/69/2015 | Florida Department of Health-Jacksonville | Centers for Disease Control and Prevention |
| EPI683997 | HA | United States | 2015-Sep-15 | A/Florida/70/2015 | Florida Department of Health-Jacksonville | Centers for Disease Control and Prevention |
| EPI684005 | HA | United States | 2015-Sep-15 | A/Florida/70/2015 | Florida Department of Health-Jacksonville | Centers for Disease Control and Prevention |
| EPI684013 | HA | United States | 2015-Sep-15 | A/Florida/71/2015 | Florida Department of Health-Jacksonville | Centers for Disease Control and Prevention |
| EPI684021 | HA | United States | 2015-Sep-15 | A/Florida/71/2015 | Florida Department of Health-Jacksonville | Centers for Disease Control and Prevention |
| EPI684029 | HA | United States | 2015-Sep-15 | A/Florida/72/2015 | Florida Department of Health-Jacksonville | Centers for Disease Control and Prevention |
| EPI684037 | HA | United States | 2015-Sep-15 | A/Florida/72/2015 | Florida Department of Health-Jacksonville | Centers for Disease Control and Prevention |
| EPI684045 | HA | United States | 2015-Sep-14 | A/Wisconsin/80/2015 | Wisconsin State Laboratory of Hygiene | Centers for Disease Control and Prevention |
| EPI684053 | HA | United States | 2015-Sep-14 | A/Wisconsin/80/2015 | Wisconsin State Laboratory of Hygiene | Centers for Disease Control and Prevention |
| EPI684061 | HA | United States | 2015-Sep-18 | A/Wisconsin/81/2015 | Wisconsin State Laboratory of Hygiene | Centers for Disease Control and Prevention |
| EPI684069 | HA | United States | 2015-Sep-18 | A/Wisconsin/81/2015 | Wisconsin State Laboratory of Hygiene | Centers for Disease Control and Prevention |
| EPI684077 | HA | United States | 2015-Sep-17 | A/Wisconsin/82/2015 | Wisconsin State Laboratory of Hygiene | Centers for Disease Control and Prevention |
| EPI684085 | HA | United States | 2015-Sep-17 | A/Wisconsin/82/2015 | Wisconsin State Laboratory of Hygiene | Centers for Disease Control and Prevention |
| EPI684093 | HA | United States | 2015-Sep-15 | A/Tennessee/21/2015 | Tennessee Department of Health Laboratory-Nashville | Centers for Disease Control and Prevention |
| EPI684101 | HA | United States | 2015-Sep-15 | A/Tennessee/21/2015 | Tennessee Department of Health Laboratory-Nashville | Centers for Disease Control and Prevention |
| EPI684109 | HA | United States | 2015-Sep-17 | A/Ohio/27/2015 | Ohio Department of Health Laboratories | Centers for Disease Control and Prevention |
| EPI684117 | HA | United States | 2015-Sep-17 | A/Ohio/28/2015 | Ohio Department of Health Laboratories | Centers for Disease Control and Prevention |
| EPI684131 | HA | United States | 2015-Sep-24 | A/Massachusetts/26/2015 | Massachusetts Department of Public Health | Centers for Disease Control and Prevention |
| EPI684139 | HA | United States | 2015-Sep-12 | A/New Mexico/23/2015 | New Mexico Department of Health | Centers for Disease Control and Prevention |
| EPI684147 | HA | United States | 2015-Sep-12 | A/New Mexico/23/2015 | New Mexico Department of Health | Centers for Disease Control and Prevention |
| EPI684155 | HA | United States | 2015-Sep-10 | A/New Mexico/25/2015 | New Mexico Department of Health | Centers for Disease Control and Prevention |
| EPI684163 | HA | United States | 2015-Sep-10 | A/New Mexico/25/2015 | New Mexico Department of Health | Centers for Disease Control and Prevention |
| EPI684171 | HA | United States | 2015-Sep-04 | A/Iowa/40/2015 | Iowa State Hygienic Laboratory | Centers for Disease Control and Prevention |
| EPI684179 | HA | United States | 2015-Sep-04 | A/Iowa/40/2015 | Iowa State Hygienic Laboratory | Centers for Disease Control and Prevention |
| EPI684187 | HA | United States | 2015-Sep-10 | A/Iowa/41/2015 | Iowa State Hygienic Laboratory | Centers for Disease Control and Prevention |

| | | | | | | |
|---|---|---|---|---|---|---|
| EPI684195 | HA | United States | 2015-Sep-10 | A/Iowa/41/2015 | Iowa State Hygienic Laboratory | Centers for Disease Control and Prevention |
| EPI684203 | HA | United States | 2015-Sep-10 | A/Iowa/42/2015 | Iowa State Hygienic Laboratory | Centers for Disease Control and Prevention |
| EPI684211 | HA | United States | 2015-Sep-10 | A/Iowa/42/2015 | Iowa State Hygienic Laboratory | Centers for Disease Control and Prevention |
| EPI684219 | HA | United States | 2015-Sep-14 | A/Iowa/43/2015 | Iowa State Hygienic Laboratory | Centers for Disease Control and Prevention |
| EPI684227 | HA | United States | 2015-Sep-14 | A/Iowa/43/2015 | Iowa State Hygienic Laboratory | Centers for Disease Control and Prevention |
| EPI684235 | HA | United States | 2015-Sep-22 | A/Wyoming/18/2015 | Wyoming Public Health Laboratory | Centers for Disease Control and Prevention |
| EPI684243 | HA | Bolivia, Plurinationial State of | 2015-Jul-08 | A/Bolivia/426/2015 | Instituto Nacional de Laboratoriosde Salud (INLASA) | Centers for Disease Control and Prevention |
| EPI684251 | HA | Bolivia, Plurinationial State of | 2015-Aug-14 | A/Bolivia/499/2015 | Instituto Nacional de Laboratoriosde Salud (INLASA) | Centers for Disease Control and Prevention |
| EPI684259 | HA | Bolivia, Plurinationial State of | 2015-Aug-18 | A/Bolivia/502/2015 | Instituto Nacional de Laboratoriosde Salud (INLASA) | Centers for Disease Control and Prevention |
| EPI684267 | HA | United States | 2015-Oct-10 | A/Michigan/67/2015 | Michigan Department of Community Health | Centers for Disease Control and Prevention |
| EPI684283 | HA | South Africa | 2015-Oct-02 | A/South Africa/6270/2015 | National Institute for Communicable Disease | Centers for Disease Control and Prevention |
| EPI684291 | HA | South Africa | 2015-Aug-12 | A/South Africa/4940/2015 | National Institute for Communicable Disease | Centers for Disease Control and Prevention |
| EPI684299 | HA | South Africa | 2015-Sep-30 | A/South Africa/6131/2015 | National Institute for Communicable Disease | Centers for Disease Control and Prevention |
| EPI684307 | HA | South Africa | 2015-Aug-28 | A/South Africa/5390/2015 | National Institute for Communicable Disease | Centers for Disease Control and Prevention |
| EPI684315 | HA | Norway | 2014-Jan-01 | A/Norway/2178/2014 | National Institute for Medical Research | Centers for Disease Control and Prevention |
| EPI685587 | HA | United States | 2015-Sep-24 | A/Montana/28/2015 | Montana Laboratory Services Bureau | Centers for Disease Control and Prevention |
| EPI685783 | HA | United States | 2015-Jul-08 | A/Hawaii/43/2015 | State of Hawaii Department of Health | Centers for Disease Control and Prevention |
| EPI685791 | HA | Canada | 2015-Sep-28 | A/Alberta/RV2373/2015 | National Microbiology Laboratory, Health Canada | Centers for Disease Control and Prevention |
| EPI685799 | HA | Canada | 2015-Oct-01 | A/Alberta/RV2382/2015 | National Microbiology Laboratory, Health Canada | Centers for Disease Control and Prevention |
| EPI685807 | HA | Canada | 2015-Oct-09 | A/Alberta/RV2383/2015 | National Microbiology Laboratory, Health Canada | Centers for Disease Control and Prevention |
| EPI685815 | HA | Canada | 2015-Oct-10 | A/Alberta/RV2385/2015 | National Microbiology Laboratory, Health Canada | Centers for Disease Control and Prevention |
| EPI685823 | HA | Canada | 2015-Oct-10 | A/Alberta/RV2386/2015 | National Microbiology Laboratory, Health Canada | Centers for Disease Control and Prevention |
| EPI685831 | HA | Canada | 2015-Oct-10 | A/Alberta/RV2387/2015 | National Microbiology Laboratory, Health Canada | Centers for Disease Control and Prevention |

| | | | | | | |
|---|---|---|---|---|---|---|
| EPI685847 | HA | Lao, People's Democratic Republic | 2015-Jul-13 | A/Laos/724/2015 | National Center for Laboratory and Epidemiology | Centers for Disease Control and Prevention |
| EPI685855 | HA | Lao, People's Democratic Republic | 2015-Aug-12 | A/Laos/844/2015 | National Center for Laboratory and Epidemiology | Centers for Disease Control and Prevention |
| EPI685863 | HA | Lao, People's Democratic Republic | 2015-Aug-12 | A/Laos/853/2015 | National Center for Laboratory and Epidemiology | Centers for Disease Control and Prevention |
| EPI685871 | HA | Lao, People's Democratic Republic | 2015-Aug-11 | A/Laos/351/2015 | National Center for Laboratory and Epidemiology | Centers for Disease Control and Prevention |
| EPI685879 | HA | Lao, People's Democratic Republic | 2015-Aug-11 | A/Laos/351/2015 | National Center for Laboratory and Epidemiology | Centers for Disease Control and Prevention |
| EPI685887 | HA | Lao, People's Democratic Republic | 2015-Aug-24 | A/Laos/884/2015 | National Center for Laboratory and Epidemiology | Centers for Disease Control and Prevention |
| EPI685895 | HA | Lao, People's Democratic Republic | 2015-Aug-24 | A/Laos/898/2015 | National Center for Laboratory and Epidemiology | Centers for Disease Control and Prevention |
| EPI685903 | HA | Lao, People's Democratic Republic | 2015-Aug-31 | A/Laos/913/2015 | National Center for Laboratory and Epidemiology | Centers for Disease Control and Prevention |
| EPI685911 | HA | Lao, People's Democratic Republic | 2015-Sep-09 | A/Laos/928/2015 | National Center for Laboratory and Epidemiology | Centers for Disease Control and Prevention |
| EPI685919 | HA | Lao, People's Democratic Republic | 2015-Sep-08 | A/Laos/375/2015 | National Center for Laboratory and Epidemiology | Centers for Disease Control and Prevention |
| EPI685927 | HA | Lao, People's Democratic Republic | 2015-Sep-08 | A/Laos/375/2015 | National Center for Laboratory and Epidemiology | Centers for Disease Control and Prevention |
| EPI685942 | HA | United States | 2015-Oct-05 | A/Michigan/60/2015 | Michigan Department of Community Health | Centers for Disease Control and Prevention |
| EPI685950 | HA | United States | 2015-Oct-05 | A/Michigan/60/2015 | Michigan Department of Community Health | Centers for Disease Control and Prevention |
| EPI685963 | HA | United States | 2015-Nov-01 | A/Washington/61/2015 | Washington State Public Health Laboratory | Centers for Disease Control and Prevention |
| EPI685971 | HA | French Guiana | 2015-Jul-13 | A/French Guiana/5020/2015 | National Influenza Center French Guiana and French Indies | Centers for Disease Control and Prevention |
| EPI685979 | HA | Guatemala | 2015-Jun-18 | A/Guatemala/142/2015 | Laboratorio Nacional De Salud Guatemala | Centers for Disease Control and Prevention |
| EPI685987 | HA | United States | 2015-Jun-16 | A/Washington/45/2015 | Washington State Public Health Laboratory | Centers for Disease Control and Prevention |
| EPI686000 | HA | United States | 2015-Oct-04 | A/Missouri/42/2015 | Missouri Department. of Health & Senior Services | Centers for Disease Control and Prevention |
| EPI686008 | HA | United States | 2015-Oct-22 | A/Pennsylvania/40/2015 | Pennsylvania Department of Health | Centers for Disease Control and Prevention |
| EPI686016 | HA | United States | 2015-Aug-16 | A/Alaska/183/2015 | Alaska State Virology Lab | Centers for Disease Control and Prevention |

| | | | | | | |
|---|---|---|---|---|---|---|
| EPI686024 | HA | United States | 2015-Aug-20 | A/Alaska/185/2015 | Alaska State Virology Lab | Centers for Disease Control and Prevention |
| EPI686032 | HA | United States | 2015-Sep-08 | A/Alaska/222/2015 | Alaska State Virology Lab | Centers for Disease Control and Prevention |
| EPI686040 | HA | United States | 2015-Sep-11 | A/Alaska/225/2015 | Alaska State Virology Lab | Centers for Disease Control and Prevention |
| EPI686048 | HA | United States | 2015-Sep-16 | A/Alaska/239/2015 | Alaska State Virology Lab | Centers for Disease Control and Prevention |
| EPI686056 | HA | United States | 2015-Sep-13 | A/Alaska/242/2015 | Alaska State Virology Lab | Centers for Disease Control and Prevention |
| EPI686064 | HA | United States | 2015-Sep-28 | A/Hawaii/62/2015 | State of Hawaii Department of Health | Centers for Disease Control and Prevention |
| EPI686072 | HA | Puerto Rico | 2015-Aug-24 | A/Puerto Rico/12/2015 | Puerto Rico Department of Health | Centers for Disease Control and Prevention |
| EPI686080 | HA | Puerto Rico | 2015-Sep-09 | A/Puerto Rico/17/2015 | Puerto Rico Department of Health | Centers for Disease Control and Prevention |
| EPI686088 | HA | Puerto Rico | 2015-Sep-08 | A/Puerto Rico/19/2015 | Puerto Rico Department of Health | Centers for Disease Control and Prevention |
| EPI686096 | HA | Puerto Rico | 2015-Sep-09 | A/Puerto Rico/20/2015 | Puerto Rico Department of Health | Centers for Disease Control and Prevention |
| EPI686104 | HA | Hong Kong (SAR) | 2015-Oct-09 | A/Hong Kong/15590/2015 | Government Virus Unit | Centers for Disease Control and Prevention |
| EPI686112 | HA | Hong Kong (SAR) | 2015-Oct-17 | A/Hong Kong/15609/2015 | Government Virus Unit | Centers for Disease Control and Prevention |
| EPI686120 | HA | Hong Kong (SAR) | 2015-Oct-19 | A/Hong Kong/15611/2015 | Government Virus Unit | Centers for Disease Control and Prevention |
| EPI686128 | HA | United States | 2015-Oct-09 | A/Washington/51/2015 | Washington State Public Health Laboratory | Centers for Disease Control and Prevention |
| EPI686143 | HA | United States | 2015-Oct-23 | A/Nebraska/19/2015 | Nebraska Public Health Lab | Centers for Disease Control and Prevention |
| EPI686151 | HA | United States | 2015-Nov-02 | A/Florida/86/2015 | Florida Department of Health-Tampa | Centers for Disease Control and Prevention |
| EPI686159 | HA | United States | 2015-Nov-15 | A/Colorado/19/2015 | Colorado Department of Health Lab | Centers for Disease Control and Prevention |
| EPI686167 | HA | United States | 2015-Nov-19 | A/Colorado/21/2015 | Colorado Department of Health Lab | Centers for Disease Control and Prevention |
| EPI686182 | HA | United States | 2015-Nov-17 | A/Colorado/24/2015 | Colorado Department of Health Lab | Centers for Disease Control and Prevention |
| EPI686190 | HA | United States | 2015-Oct-20 | A/California/94/2015 | California Department of Health Services | Centers for Disease Control and Prevention |
| EPI686198 | HA | United States | 2015-Oct-19 | A/California/98/2015 | California Department of Health Services | Centers for Disease Control and Prevention |
| EPI686206 | HA | United States | 2015-Nov-11 | A/Texas/71/2015 | Texas Department of State Health Services-Laboratory Services | Centers for Disease Control and Prevention |
| EPI686214 | HA | United States | 2015-Oct-08 | A/Maryland/17/2015 | Maryland Department of Health and Mental Hygiene | Centers for Disease Control and Prevention |
| EPI686222 | HA | United States | 2015-Dec-03 | A/Washington/62/2015 | Marshfield Clinic Research Foundation | Centers for Disease Control and Prevention |
| EPI686230 | HA | United States | 2015-Dec-08 | A/Texas/75/2015 | Texas Department of State Health Services-Laboratory Services | Centers for Disease Control and Prevention |

| | | | | | | |
|---|---|---|---|---|---|---|
| EPI686238 | HA | United States | 2015-Oct-18 | A/Hawaii/76/2015 | State of Hawaii Department of Health | Centers for Disease Control and Prevention |
| EPI686246 | HA | United States | 2015-Nov-04 | A/California/101/2015 | California Department of Health Services | Centers for Disease Control and Prevention |
| EPI686254 | HA | United States | 2015-Nov-03 | A/California/106/2015 | California Department of Health Services | Centers for Disease Control and Prevention |
| EPI686275 | HA | United States | 2015-Nov-23 | A/Pennsylvania/50/2015 | Pennsylvania Department of Health | Centers for Disease Control and Prevention |
| EPI686286 | HA | United States | 2015-Dec-08 | A/Colorado/27/2015 | Colorado Department of Health Lab | Centers for Disease Control and Prevention |
| EPI686294 | HA | United States | 2015-Dec-04 | A/Florida/93/2015 | Florida Department of Health-Tampa | Centers for Disease Control and Prevention |
| EPI686302 | HA | United States | 2015-Nov-18 | A/California/112/2015 | California Department of Health Services | Centers for Disease Control and Prevention |
| EPI686310 | HA | United States | 2015-Nov-28 | A/California/113/2015 | California Department of Health Services | Centers for Disease Control and Prevention |
| EPI686318 | HA | United States | 2015-Nov-26 | A/Alaska/266/2015 | Alaska State Virology Lab | Centers for Disease Control and Prevention |
| EPI686326 | HA | United States | 2015-Nov-27 | A/Alaska/267/2015 | Alaska State Virology Lab | Centers for Disease Control and Prevention |
| EPI686748 | HA | Sweden | 2015-Nov-15 | A/Halmstad/1/2015 | | Public Health Agency of Sweden |
| EPI686756 | HA | Sweden | 2015-Nov-17 | A/Skovde/5/2015 | | Public Health Agency of Sweden |
| EPI686780 | HA | Sweden | 2015-Nov-16 | A/Lund/2/2015 | | Public Health Agency of Sweden |
| EPI686791 | HA | Sweden | 2015-Nov-17 | A/Stockholm/58/2015 | | Public Health Agency of Sweden |
| EPI686836 | HA | Sweden | 2015-Nov-23 | A/Stockholm/61/2015 | | Public Health Agency of Sweden |
| EPI686860 | HA | Sweden | 2015-Nov-21 | A/Gothenburg/1/2015 | | Public Health Agency of Sweden |
| EPI686868 | HA | Sweden | 2015-Nov-23 | A/Stockholm/63/2015 | | Public Health Agency of Sweden |
| EPI686876 | HA | Sweden | 2015-Nov-21 | A/Stockholm/64/2015 | | Public Health Agency of Sweden |
| EPI686884 | HA | Sweden | 2015-Dec-03 | A/Kungalv/1/2015 | | Public Health Agency of Sweden |
| EPI686924 | HA | Sweden | 2015-Oct-30 | A/Uppsala/7/2015 | | Public Health Agency of Sweden |
| EPI687165 | HA | Sweden | 2015-Nov-20 | A/Stockholm/65/2015 | | Public Health Agency of Sweden |
